# SCANBIT facilitates identification of tumor cell populations in scRNAseq data using pseudobulked SNV calls

**DOI:** 10.64898/2026.01.27.701834

**Authors:** MV Cannon, MJ Gust, AC Gross, M Cam, JB Reinecke, L Jimenez Garcia, CH Strawser, L Ryan, M Sammons, CZ Zhang, RD Roberts

## Abstract

**Motivation:** Single cell RNAseq (scRNAseq) is an ideal tool to characterize the heterogeneity within the tumor microenvironment, however, accurate identification of tumor cells can be a challenge. Reference-based methods can be inaccurate, if reference datasets are even available. Current purpose-built methods can be inaccurate, particularly with highly heterogeneous tumor types. Improved methods are needed. We explored the use of genetic variants to distinguish tumor from normal cells within scRNAseq data.

**Results:** We characterized the limitations inherent to calling variants from scRNAseq data, quantifying how data sparsity precludes genetic distance calculation between single cells. As a novel workaround, we pooled data from transcriptionally similar cell clusters to call high quality variants and then calculated pairwise differences between cell populations and performed hierarchical clustering. We quantified confidence in genetic divergence between tumor and normal cell populations using bootstrapping. We performed extensive validation to assess accurate identification of tumor cells using ground-truth datasets. Application of our method to human scRNAseq samples highlighted the utility of our approach and revealed how mutational burden influences successful tumor cell identification.

Improved cell type assignment in scRNAseq data will facilitate analysis of tumor samples and, in turn, accelerate our understanding of the mechanisms underlying tumor progression and reveal potential biological vulnerabilities that can be exploited to develop improved treatment options.

**Availability and implementation:** Our method is publicly available as an R package: SCANBIT (Single Cell Altered Nucleotide Based Inference of Tumor) https://github.com/kidcancerlab/scanBit.

## 1) Introduction

Tumor heterogeneity and microenvironmental interactions have emerged as key aspects of cancer biology (Aynaud, et al., 2020; Rajan, et al., 2023; Tirosh and Suva, 2024). A common tool to understand the complex tumor microenvironment and interactions within a tumor is single-cell RNA sequencing (scRNAseq). This technique allows exploration of the phenotypic state of each cell from a sample and provides insights that are not possible when using bulk methods such as standard RNAseq (Wang, et al., 2023). This increased data granularity comes with a myriad of additional challenges, however, and statistical methods to properly analyze scRNAseq data is a rapidly evolving field (Andrews, et al., 2021; Jiang, et al., 2022). One of these challenges is the assignment of labels for each cell within these datasets. For cancer samples, tumor cells are often not present in reference data commonly used to identify cell types. Further, heterogeneity between patients, tumor subtypes, or even within a single patient can lead to inaccurate identification of tumor cells in scRNAseq data when using reference-based methods. Gene markers to identify tumor cells with certainty are also often not well-established. Several methods have been developed to overcome this limitation using various approaches such as copy number variation or machine learning, each with individual drawbacks and limitations (De Falco, et al., 2023; Nofech-Mozes, et al., 2023).

Osteosarcoma exhibits extraordinary genome complexity and heterogeneity, and the identification of tumor cells in scRNAseq data has been challenging (Beird, et al., 2022; Espejo Valle-Inclan, et al., 2025). To overcome these difficulties, we chose to investigate whether we could use the sequences generated during scRNAseq to identify tumor cells based on DNA substitutions accumulated in the tumor cells. We first assessed how many variants could be called on individual cells to determine if this would be adequate to calculate genetic distance. We then evaluated the potential and accuracy of calling variants on individual cells and quantified the associated limitations. We then tested if applying this method to transcriptionally clustered pseudobulked data improved results. We developed an R package called SCANBIT (Single Cell Altered Nucleotide Based Inference of Tumor) to do identify tumor cells using variant calling on pseudobulked populations.

## 2) Methods

### Mouse sample collection

All mouse experiments were performed under IACUC-approved protocols. We generated metastatic osteosarcoma samples from both F420 and K7M2 osteosarcoma cell lines injected into C57BL/6 and BALB/c mice, respectively.

We injected F420 tumor cells into the tail vein of C57BL/6 mice. For one sample, the F420 cells were transfected with an sLPmCherry construct to fluorescently mark adjacent cells (Ombrato, et al., 2019). We sacrificed the mice after 44 or 46 days, collected lung tissue and performed dissociation. For the sLPmCherry sample, we isolated tumor cells and adjacent normal cells by flow cytometry. We did not sort the non-transfected sample.

For the K7M2 model, we injected K7M2 cells transfected with sLPmCherry into the tail vein of BALB/c mice. We sacrificed mice after ten days and collected lungs for dissociation. We split the sample into CD45+, Epcam+, and double-negative fractions to ensure representation of key cell types in the samples. We performed CD45 selection using the Miltenyi CD45 depletion kit (cat# 130-052-301). Epcam selection was performed by flow cytometry after staining with R&D-EpCAM antibodies (cat #FAB9601R-100UG). CD45+, Epcam+, and double-negative cells were pooled in a 1:1:1 ratio for further processing.

We processed isolated cells for 10x 3’ scRNAseq using a 10x Chromium controller. We targeted between 1800 and 8000 cells, depending on the dissociated cell count. After standard library preparation, all samples were sequenced on an Illumina NovaSeq 6000 sequencer.

### Publicly available data

We retrieved additional data from several sources to perform downstream analyses. For experiments using 10x Flex data, we downloaded mouse eye, ovary, stomach, and intestine data from the 10x website (https://www.10xgenomics.com/datasets/40k-mixture-of-cells-dissociated-from-4-fixed-mouse-tissues-using-manual-dissociation-multiplexed-samples-4-probe-barcodes-1-standard). This data is aligned to the mm10 mouse genome.

We downloaded data from the NCBI SRA database linked to NCBI GEO IDs GSE210358, GSE222703, GSE247228, GSE201615, GSE162454, GSE169396, GSE237579, GSE184880, GSE306201, and GSE261693. This included prostate cancer, clear cell carcinoma of the kidney and adjacent normal kidney tissue, melanoma, undifferentiated pleiomorphic sarcoma, osteosarcoma, healthy bone, healthy PBMCs, ovarian cancer, breast cancer, and Ewing sarcoma samples. We downloaded raw fastq files and associated metadata for processing.

We also used data previously published by our lab. This included data available online through GEO accessions GSE252703, GSE179681 and GSE316454. Metadata for all samples is available in Supplemental Table 1.

### Data processing

We used either the hg38 human genome, mm10 mouse genome, or a custom reference made up of hg38 plus RFP and eGFP sequences (Supplemental Table 1). Each sample was aligned to the appropriate reference using Cell Ranger version 7.2.0 (10x), allowing for inclusion of intronic reads. Following alignment, we processed the data using Seurat version 5.1.0 in R version 4.3.0 (Stuart, et al., 2019). Samples were filtered to remove low-quality cells and doublets using sample-specific cutoffs (Supplemental Table 1) and processed using standard methods to generate Seurat objects with clusters calculated using Louvain clustering. We used silhouette scoring to choose clustering resolution for each sample independently. We used SingleR to perform cell type annotation using several references (Aran, et al., 2019). For mice, we used MouseRNAseqData and ImmGenData from the celldex R package (Aran, et al., 2019) as well as F420 and K7M2 osteosarcoma cell culture scRNAseq data. For human data, we used MonacoImmuneData, HumanPrimaryCellAtlasData, and BlueprintEncodeData references from the celldex R package (Aran, et al., 2019). All code for validation studies are available through GitHub (https://github.com/kidcancerlab/24_validate_snvs).

### Variant calling

Variant calling methods are incorporated into the scanBit R package (https://github.com/kidcancerlab/scanBit) as a single function “get_snp_tree()” that takes a table of cell barcodes, cluster groups, and corresponding bam file locations. The function outputs a hierarchical clustering tree and a file identifying genetically distinct clusters. Briefly, for each cluster, cell barcodes are used to subset the sample bam file down to the matching reads for that cluster using samtools (Danecek, et al., 2021; Li, et al., 2009). PCR duplicates are removed using samtools. Variants are called and filtered for each cluster independently using bcftools “mpileup” and “call” (Danecek, et al., 2011; Danecek, et al., 2021; Li, 2011). Variants from each cluster are merged into a single bcf file and indexed using bcftools. The merged bcf file is the input for a custom python script (vcfToMatrix.py) that compares the variants from each cluster to calculate a distance matrix composed of the pairwise proportion of alleles that differ between each cluster. We use this to generate a hierarchical clustering tree. To assess the confidence of each node in the hierarchical clustering tree, we utilize bootstrapping (Hillis and Bull, 1993). We sample the variants with replacement to create 10,000 simulated datasets. For each node in the tree, we calculate the bootstrap support value – the percentage of replicates that maintained the exact division of clusters defined by that node. The final output for the function is a pdf of the tree with bootstrap values and a table outlining the genetically distinct cluster groupings based on user-provided bootstrap threshold. We used a bootstrap threshold of 85% to define genetically distinct cell clusters.

### Variant calling accuracy assessment

We used 10x Flex scRNAseq data to estimate the false positive variant call rate. The 10x Flex kit uses probes to detect transcripts, with the sequence data reflecting the probe sequences and not transcript sequences. The Flex data analyzed here used probes designed against the mm10 genome. Any variant that differs from the mm10 genome is an error that occurred during probe synthesis, library PCR amplification, or sequencing. We called variants on the four Flex datasets outlined above and counted, for each read depth covered, the number of total covered sites and sites with a variant call to generate a table of the error rate at a given sequencing depth. To calculate how many total sites are commonly covered by scRNAseq data, we used human patient scRNAseq samples to count the number of sites covered by individual cells at varying read depths. We then multiplied the error rate for each read depth calculated from the 10x Flex data by the number of sites covered at corresponding read depths in the patient data for single cells to calculate the number of variant sites expected to be erroneously called. We compared the actual number of variants called in the human data to the expected number of errors to calculate the proportion of observed variants that are likely to be incorrectly called at each read depth.

### Validation using healthy mouse data from two strains

To test if pseudobulked samples can help identify genetically distinct cell populations, we used scRNAseq data from the lungs of healthy, genetically distinct C57BL/6 and BALB/c mice. We merged data from these samples and clustered the cells by transcriptional profile and sample, and then analyzed the data with scanBit. The resulting hierarchical clustering tree was inspected to determine if clusters from each strain grouped correctly.

### Validation using transgene-labelled tumor cells

To quantify how well scanBit successfully identifies genetically distinct tumor populations, we used data derived from syngeneic mouse models of cancer, where the tumor cells were transfected with either GFP, RFP, or lineage tracing barcodes. After alignment to a reference containing GFP and RFP we counted, for each cell, how many reads aligned to a transgene or lineage tracing barcode. Then, for each cluster, we then quantified the proportion of cells with expression of either. We compared this with the clusters defined as tumor by scanBit to determine if those clusters had high proportion of transgene-expressing cells. We included samples from mice injected with K7M2 cell lines as a negative control because this cell line does not have a high enough mutational burden to be genetically identified using scanBit. We excluded one sample that had no detectable transgene.

### Testing scanBit using syngeneic mouse osteosarcoma models

To mimic human patient data where the tumor cells are genetically related to the host, we used two syngeneic mouse models of osteosarcoma. The F420 cell line is derived from C57BL/6 mice, while the K7M2 cell line is derived from BALB/c mice (Khanna, et al., 2000; Zhao, et al., 2015). We analyzed two datasets comprised of 1) one K7M2 control cell culture sample, one BALB/c control lung sample, and three lung metastasis samples from BALB/c mice injected with K7M2 cells, and 2) one F420 cell culture control sample, one C57BL/6 control lung sample, and five lung metastasis samples from C57BL/6 mice injected with F420. We merged and clustered each dataset by transcriptional profile and sample and then processed them through scanBit to define genetically distinct cell populations. After the F420/C57BL/6 data was run, we noted that one cluster from one sample consisted of a roughly even mixture of tumor and healthy cells. This resulted in a tree with low bootstrap values. We re-clustered this sample with a resolution of 0.25 to produce more granular clusters and re-merged it with the rest of the samples. Re-running scanBit with the more granular clusters resulted in a tree that accurately identified tumor clusters.

### Testing scanBit on human patient tumor samples

To assess how well scanBit works on human patient tumor samples, we tested a range of cancer types, including prostate cancer, clear cell carcinoma of the kidney, melanoma, undifferentiated pleiomorphic sarcoma, osteosarcoma, ovarian cancer, breast cancer, and Ewing sarcoma samples. To ensure that genetically distinct populations were not incorrectly identified in normal tissue, we also analyzed healthy kidney tissue adjacent to the clear cell carcinoma of the kidney, healthy bone, and healthy PBMCs. We ran each sample through scanBit and recorded the bootstrap value of the highest node on the resulting tree.

### Determination of required data quantity to accurately distinguish genetically distinct clusters

To assess how many reads or cells are required to distinguish genetically distinct clusters, we downsampled human patient scRNAseq data. We omitted any cluster with fewer than 500 cells so that all clusters had the same number of cells when downsampled to a given number. We used either all, 500, 100, 50, 20, 10, 5, 4, 3, or 2 cells per cluster and ran scanBit. We recorded the bootstrap values at the highest node of the resulting trees. We also recorded the total number of UMIs for each cluster from each sample when downsampled to a given number of cells.

The scanBit R package is available on GitHub (https://github.com/kidcancerlab/scanBit), and all code used for bioinformatic validation of scanBit is also available on GitHub (https://github.com/kidcancerlab/24_validate_snvs).

## 3) Experiments and Results

### Evaluation of variant calling on single cells

To assess the potential to call variants in data from individual cells, a key consideration is the number of variants that can be accurately called given the data sparsity inherent to scRNAseq. We began by determining how many positions are covered across the genome by scRNAseq data at varying depths in human patient scRNAseq data. The cells used for this analysis had between 1,077 and 147,868 UMIs per cell with a median of 7,648. We observed an average of 137,353 sites covered using a cutoff of 2X, 15,330 sites covered using a cutoff of 10X, and 6,731 sites covered using a cutoff of 20X (Figure 1A and Supplemental Table 2). However, only non-reference (variant) sites will be informative for distinguishing genetically distinct populations. When considering only variant sites, each cell had on average, only 82.99 sites covered using a minimum read depth of 2X, 12.95 sites using a cutoff of 10X, and 5.66 sites using 20X per cell (Figure 1A and Supplemental Table 3). Variant calls represented 0.06% of all sites at 2X coverage through 0.07% at 20X (Supplemental Figure 1A). We also found that while less than 1% of cells had no variant calls at 2X, this increased to 53.3% and 79.3% of cells at 10X and 20X coverage, respectively (Supplemental Figure 1B).

**Figure 1:**
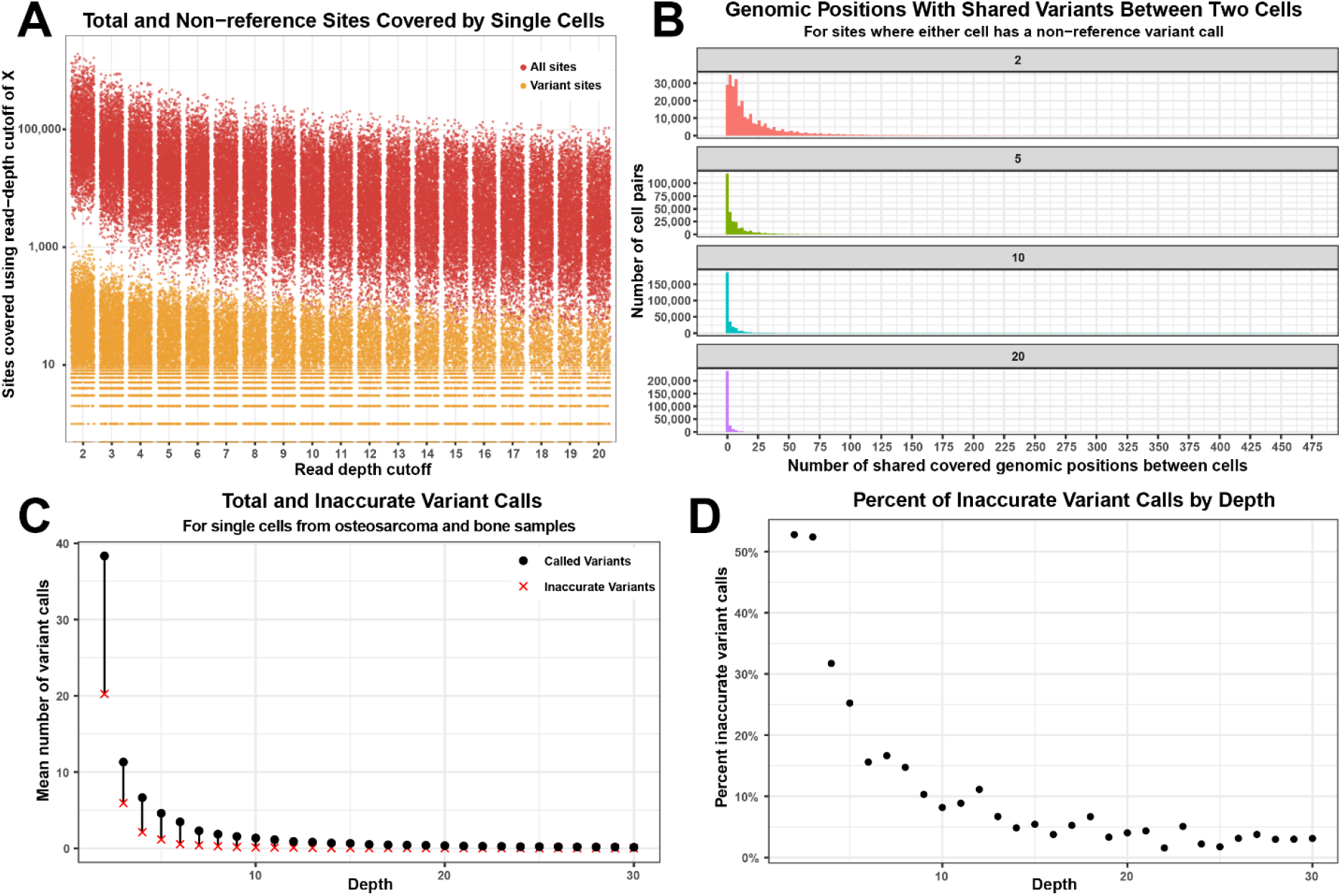
Single cells lack adequate data quantity to calculate genetic distance accurately. Genetic distance calculation between cells requires accurate variant info in adequate numbers. This quantity of calls is lacking when using individual cells. We calculated the total number of genomic sites covered to a given sequence depth in individual cells (red points in A) as well as the number of genomic sites that differ from the reference sequence and covered to a given sequence depth (orange points in A) showing that very few non-reference sites are covered. We also calculated how many sites have variant calls at shared sites in single cell data when using 2, 5, 10 or 20 as a minimum read depth threshold (B). For normal scRNAseq data from osteosarcoma and healthy bone samples, we calculated the average number of sites with variant calls at given sequence depths in individual cells and the expected number of incorrectly called variants as determined from analysis of Flex data (C). Given these values, (D) shows the expected percentage of variant calls expected to be inaccurate at given sequence depths.

Further restraining the utility of variant calls from single cell data is the requirement for variants to be shared between cells to inform the genetic relatedness of those cells. We calculated, for each sample used in the previous analysis, how many genomic positions had data between each pair of cells analyzed per sample where one of the two cells had a non-reference call (informative sites) when applying minimum read depth cutoffs of 2, 5, 10, and 20 for variant calling. We found that on average, only 21.90 sites had variant information between any two cells per sample (median of 11), even when allowing for variant calls at positions with only two reads (Figure 1B and Supplemental Figure 2). When applying more stringent cutoffs, such as 10, the average number of shared sites between any two cells in a sample was only 3.06 (median of 1), and with a cutoff of 20, we see only 1.03 sites on average shared between two individual cells (median of 0) (Figure 1B). There was considerable variance in the number of positions with shared information between cells, with some cells with similar transcriptional profiles having many shared sites while others had very few (Supplemental Figure 2).

**Figure 2:**
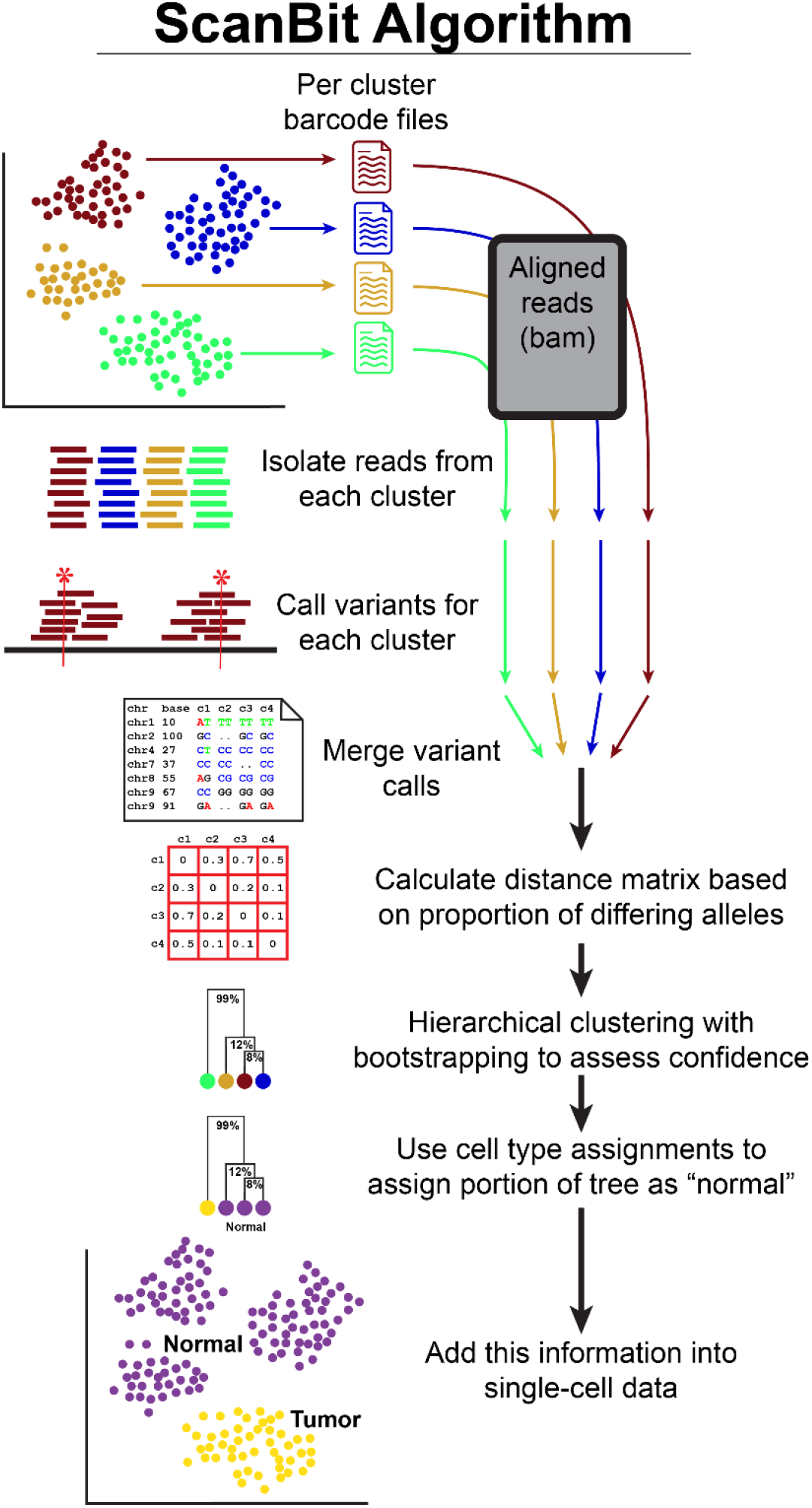
Overview of the scanBit algorithm. For each transcriptionally clustered cell population, a file is written containing all cell barcodes. For each file of barcodes, all corresponding sequence reads are retrieved from sample bam files, PCR duplicates are filtered and passed to bcftools for variant calling. Variant calls from each cluster are then merged and the pairwise proportions of alleles that differ are calculated and this is used as input for hierarchical clustering with bootstrapping to assess confidence in the division of genetically distinct populations. Further analyses are possible which use cell type annotations to assign a portion of the tree as normal based on presence of cell types unlikely to be inappropriately assigned to tumor cells. Information on genetically distinct populations can then be added back into the metadata of tumor samples.

### Evaluation of inaccurate variant calling across varying read depths

To assess the rate of incorrect variant calls made at various read depth cutoffs, we utilized 10x Flex scRNAseq data, where any variant call represents an error. We identified between six and sixteen clusters of cells per sample in the data from mouse eye, ovary, stomach, and intestine samples. We found that the number of sites covered at each read depth varied from an average of 94,938 for 2x coverage down to 17,092 sites for a coverage of 20 (Figure 1A, Supplemental Table 4). We then counted the number of sites called as non-reference (and therefore represent inaccurate variant calls) and found that these sites represented 0.031% of all sites covered at 2x through 0.003% for sites covered at 20x (Supplemental Figure 3 and Supplemental Table 4). This percentage is the rate at which non-variant sites are incorrectly called as variant (false positives).

**Figure 3:**
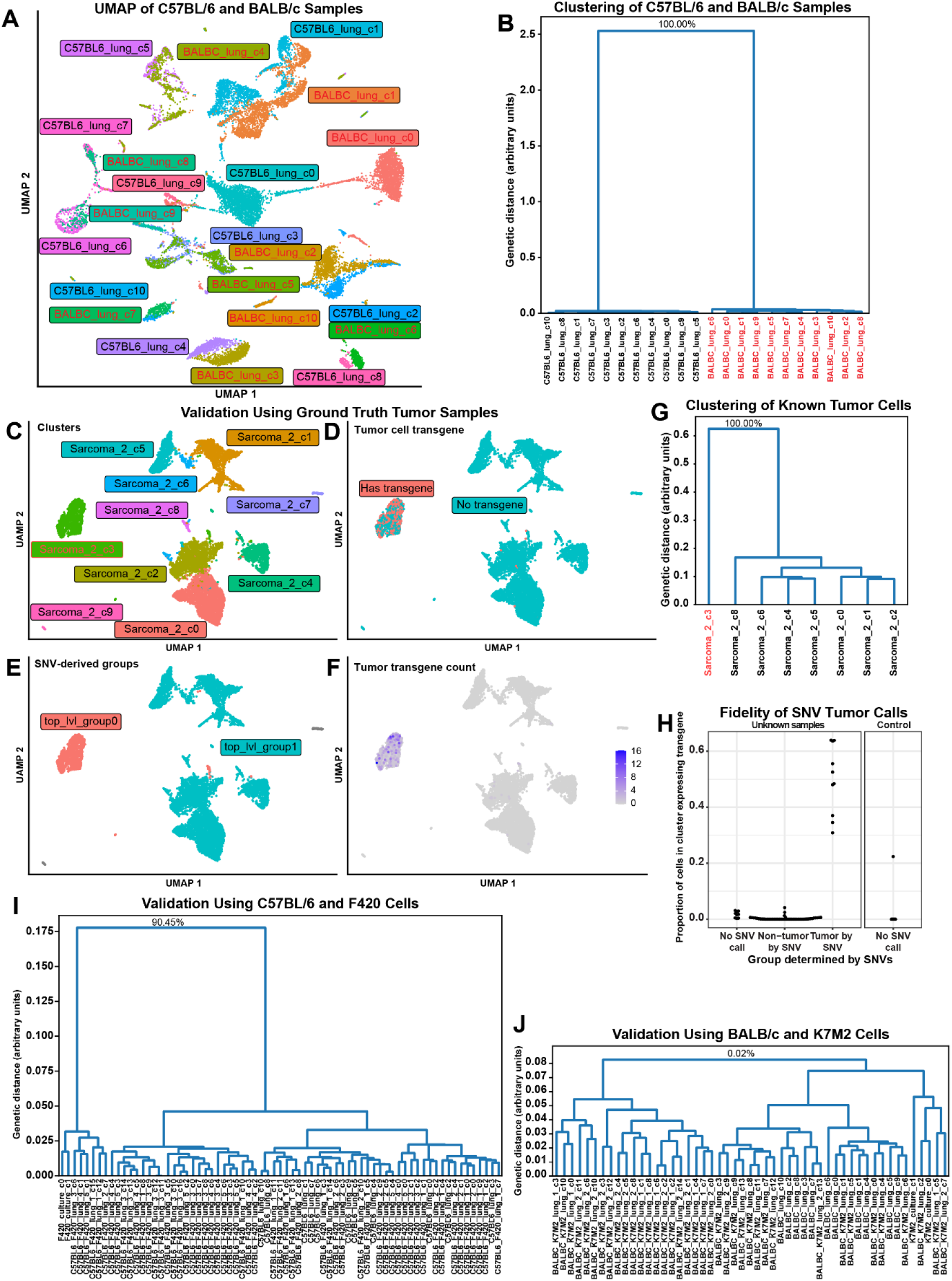
Validation of scanBit using mouse strains and genetically labeled tumor cells. To validate our approach to identifying genetically distinct cell populations we began by using scRNAseq data from genetically distinct C57BL/6 and BALB/c mouse strains which was clustered by transcriptional similarity and sample (A). We processed this data with scanBit to generate a hierarchical clustering tree (B). Bootstrap values for lower nodes omitted to save space but full figure is available in Supplemental Figure 4. The bootstrap value of the top node indicated that 100% of 10,000 resamplings correctly divided C57BL/6 from BALB/c clusters. We next validated our approach using syngeneic mouse cancer models that were genetically labeled with GFP, RFP or lineage tracing barcodes. Representative results from one sample are shown: transcriptionally defined clusters (C), cells containing tumor transgene (D), genetically distinct populations defined by scanBit (E), tumor transgene read count (F) and hierarchical clustering tree with bootstrap values (G). Bootstrap values for lower nodes omitted to save space but full figure is available in Supplemental Figure 5. (H) summarizes results from all samples, showing the proportion of cells in the cluster with tumor transgenes for clusters with no scanBit tumor call (too few cells or reads), those predicted to be non-tumor predicted to be tumor by SNV calling. Included is a control sample where the tumor cells lack adequate genetic distance for accurate scanBit results. We next validated our approach using data from control healthy mice and mice injected with F420 (I) and K7M2 (J) osteosarcoma cells as well as control cell culture cells. F420 cell clusters were easily distinguished from C57BL/6 host cells, while K7M2 cells lacked adequate genetic distance to be distinguished from host BALB/c cells. Bootstrap values for lower nodes omitted to save space but full figure is available in Supplemental Figure 6.

We used this calculated false positive variant calling rate to estimate the proportion of called variants in standard scRNAseq that likely result from sequencing errors. Above, we calculated the number of variants called for several read depth thresholds for individual cells in the human patient data (Figure 1C). When considering the total number of sites covered in these samples at each read depth and our estimates of the false positive variant call rates from the Flex data at each read depth, we calculated that we would expect 52.7%, 8.2% and 4.0% of variant calls to be false positives at 2x, 10x and 20x read depth, respectively (Figure 1D).

### Overview of variant calling approach using pseudobulked cell clusters

Because variant calling on single cells dramatically limits coverage and potentially introduces high levels of false positives, we decided to pseudobulk cells by cluster so that the combined sequence information would enable more accurate variant calling. We used as input a table of cell barcodes and corresponding transcriptional cluster identifiers. We then pooled all reads from the associated bam file for each sample and called variants for each pseudobulked cluster. The variants across all clusters are compiled, and we calculate the proportion of differing alleles between each pair of clusters as input into hierarchical clustering. We use bootstrapping of the data to assess the confidence of the genetic division (Figure 2). We then use cell type annotations to identify which of the genetically distinct populations are least likely to be inaccurately annotated tumor cell populations.

### Validation of variant calling on pseudobulked scRNAseq data from distinct mouse strains

To test this approach, we chose two genetically distinct lung single-cell datasets, one from C57BL/6 and the other from BALB/c. We had eleven clusters in both the C57BL/6 sample and the BALB/c sample (Figure 3A). The phylogenetic tree results are presented in Figure 3B and Supplemental Figure 4. We tested minimum read depth cutoffs of 5, 10, 20, and 30 to assess how this affected the results. For all read depth cutoffs used, the trees generated by our method clearly split the two mouse strains with bootstrap values of 100% (Figure 3B and Supplemental Figure 4).

**Figure 4:**
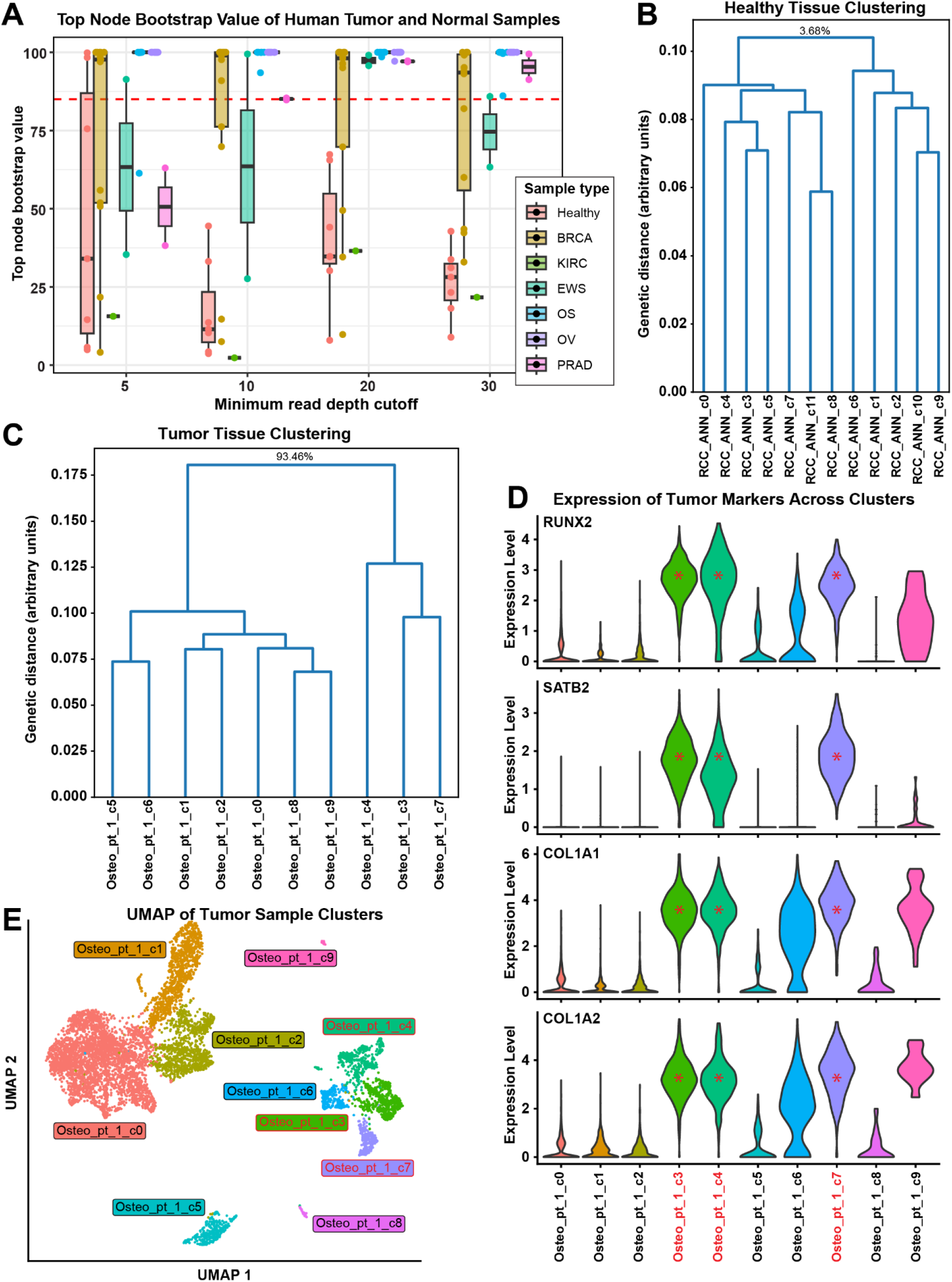
Testing on human patient samples shows dependency on tumor mutational burden. We tested scanBit on various human patient tumor and healthy samples (A). When considering the top node bootstrap values of output trees, healthy samples had low bootstrap values except when using a very low minimum read depth threshold for variant calls. Across the cancer types tested, each had varying levels of success in distinguishing tumor and normal subpopulations, presumably due to differences in mutational burden. (B) shows a representative example of results from running scanBit on a healthy sample (normal kidney tissue adjacent to clear cell carcinoma of the kidney). Bootstrap values for lower nodes omitted to save space but full figure is available in Supplemental Figure 7. (C) is representative results from running scanBit on an osteosarcoma sample. Tumor clusters are highlighted in red. Note that clusters 6 and 9 are not identified as tumor by scanBit and may have been confused for tumor cells as these clusters express a subset of tumor markers (D). Clusters identified as tumor by scanBit have red asterisks. (E) is a UMAP projection of the osteosarcoma sample from (C) and (D). Clusters identified as tumor by scanBit have red text.

### Validation using ground truth cancer scRNAseq datasets

To test samples with less extreme genetic distance, we took advantage of existing mouse data where syngeneic cancer models included transfected genes that made the identification of tumor cells possible. We included scRNAseq tumor samples from mice, in which the tumor cells expressed GFP, RFP, or lineage tracing barcodes. Only tumor cells will have reads from these genes, and the clusters containing tumor cells can therefore be identified. We analyzed six samples with GFP, two with RFP, and two with lineage tracing barcodes. One sample functioned as a negative control because K7M2 had insufficient genetic distance from the BALB/c mouse host to be distinguished using variants. After clustering (representative example in Figure 3C), we identified which cells had transgene expression (Figure 3D and F, for example). Across these samples, between 0 and 64% percent of the cells per cluster expressed a transgene used as a tumor marker. Within the clusters designated as tumor by transgene presence, between 22% and 64% of the cells had the transgene sequence detected, while non-tumor clusters had at most 4% of cells expressing the transgene. The negative control K7M2 osteosarcoma sample had one cluster with 22% of cells expressing the transgene. While the K7M2 tumor cell cluster was divided from normal cells at the root node of the hierarchical tree, the bootstrap value of this split was only 50.99% (Supplemental Figure 5).

**Figure 5:**
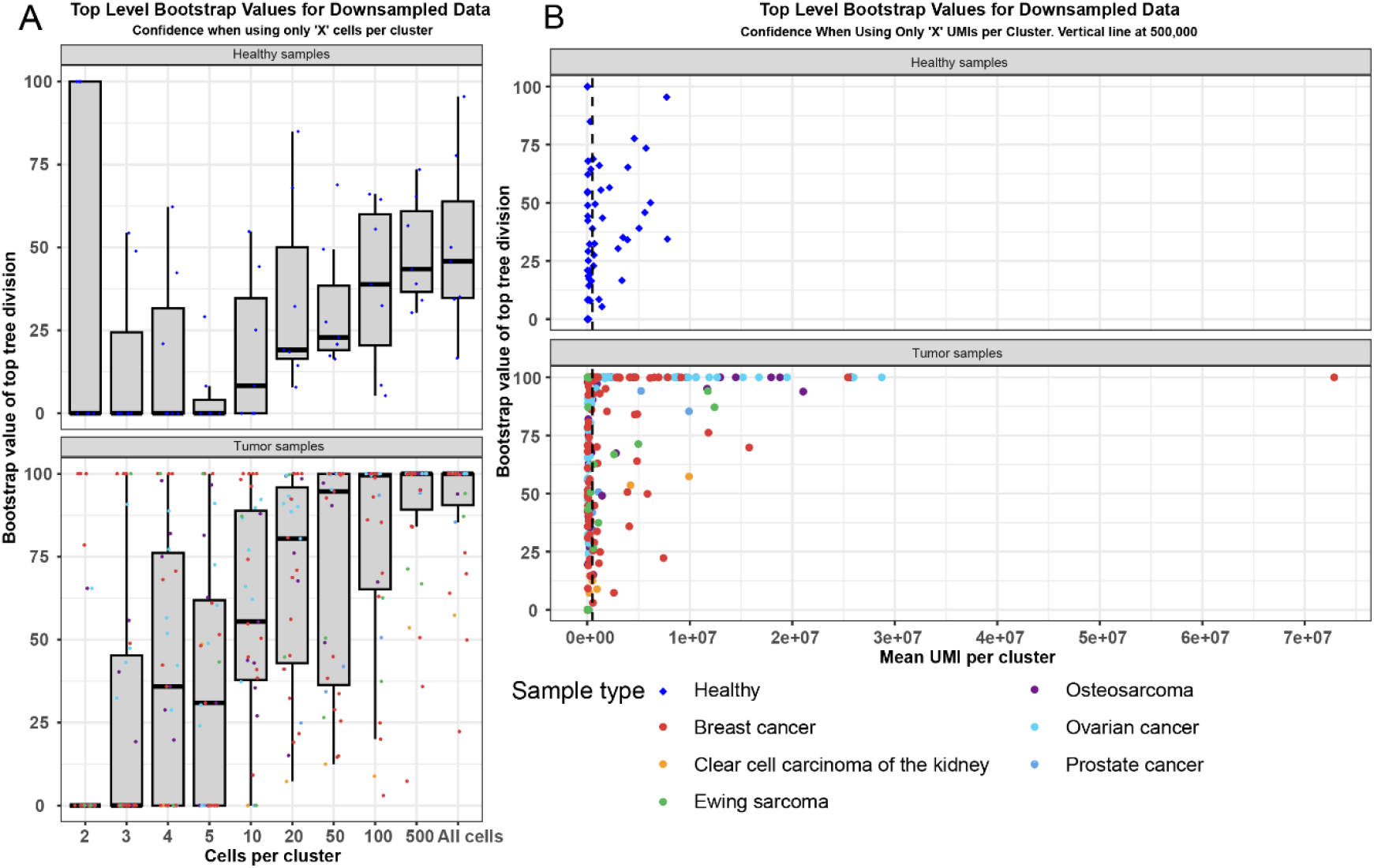
Analysis of downsampled data reveals minimum requirements for data quantity We downsampled the number of cells per cluster in both healthy and tumor samples and analyzed them using scanBit. We included all, 500, 100, 50, 20, 10, 5, 4, 3 or 2 cells per cluster for each sample and recorded the bootstrap value of the top level node in the resulting hierarchical clustering trees (A). Healthy samples had mostly low bootstrap values regardless of downsampling while bootstrap values of tumor samples dropped when fewer than 50 cells were included in each cluster. We used these data and calculated the average number of UMIs in each cluster to determine how many were required to distinguish genetically different populations (B). When a sample had fewer than around 500,000 UMIs top level bootstrap values quickly degraded.

After running scanBit on these samples, we observed that all clusters with high transgene expression (excluding the negative control) were identified as genetically distinct populations (representative example in Figure 3D, E, F, and G and summary of all samples in Figure 3H, all results in Supplemental Figure 5). The overall distance observed between the tumor cells and the syngeneic host cells (as shown by smaller values in the y-axis of the resulting genetic trees) (Figure 3G) was much lower than observed between C57BL/6 and BALB/c (Figure 3B), showing that these populations were less genetically diverged.

### Evaluation using syngeneic mouse models

As a further test, we analyzed samples from mice with two implanted osteosarcoma cell lines. The F420 cell line is derived from C57BL/6 mice, and so the majority of SNVs distinguishing one group from another should be those that have accumulated within the tumor cells (Zhao, et al., 2015). Likewise, the K7M2 cell line arose as a spontaneous osteosarcoma from BALB/c mice (Khanna, et al., 2000). These samples mimic the patient condition where the tumor cells share genetic heritage with the patient and contain accumulated genetic differences. We analyzed the combined host mouse strain, control cell culture samples, and tumor samples from mice harboring each cell line. We found that for the F420 cell line, we were able to distinguish the tumor cell populations from the host cells. We observed bootstrap values of 85.47%, 90.45%, 47.66%, and 99.50, when including minimum read depth cutoffs for variant calling of 5, 10, 20, and 30 (Figure 3I and J and Supplemental Figure 6). For the K7M2 samples, there was insufficient genetic distance to distinguish the tumor cells, either in culture or *in vivo*, from the host cells with bootstrap values no greater than 3.16% across all read depth cutoffs tested (Figure 3J and Supplemental Figure 6).

### Evaluation using human patient scRNAseq samples

We next chose to test our approach on human patient scRNAseq samples with varying level of mutational burden. We downloaded publicly available data from the SRA and GEO databases. We included healthy samples from bone, adjacent normal kidney, and PBMCs as controls, as well as osteosarcoma, prostate cancer, clear cell carcinoma of the kidney, ovarian cancer, breast cancer, and Ewing sarcoma samples. We analyzed the samples using scanBit to determine if healthy samples had uniform genetic structure and tumor samples showed genetically distinct tumor cell populations.

The healthy kidney and PBMC samples were all genetically homogeneous with bootstrap values below 67% (mean 26.7%) (Figure 4A and B and Supplemental Figure 7). For the healthy bone tissue, when using a minimum read depth threshold of five, two samples had a single cluster of cells that were genetically distinct (bootstrap values of 99.8% and 98.3%), however higher read depth cutoffs did not show this group to be genetically distinct with bootstrap values no greater than 44.5% and 34.7%, respectively (Figure 4A) suggesting that these were false positive results due to relaxed read depth thresholds. The other two healthy bone samples had bootstrap values below 34.7% (mean 23.3%) for minimum read depths of 10, 20, and 30.

For the 31 tumor samples tested, all samples had genetically distinct subpopulations of cells except the clear cell carcinoma of the kidney and four of the 13 breast cancer samples. The clear cell carcinoma of the kidney sample had a maximum bootstrap value of 36.5% across all read depths tested (Figure 4A and Supplemental Figure 7). The four breast cancer samples that did not meet the minimum bootstrap value cutoff had a maximum bootstrap value of 76.2%. The other 26 tumor samples had genetically distinct populations for at least one minimum read depth. Example validation of subpopulations in an osteosarcoma sample (Figure 4C, D, and E) shows division into two populations at the root node as well as child nodes within the tumor clusters, suggesting genetic sub-structure may exist within the tumor cells. Tumor marker genes were highly enriched within the three genetically distinct clusters (Figure 4D). Two clusters that were genetically indistinguishable from normal cells expressed some, but not all, tumor markers.

To test the minimum amount of data required to generate accurate results, we performed a downsampling analysis on the data. Using the osteosarcoma patient data, we randomly downsampled each cluster from each sample to 1, 2, 3, 4, 5, 10, 20, 50, 100, 500, or all cells. We then analyzed each sample with scanBit and recorded the bootstrap value of the top division in the resulting tree to assess when the value degraded due to insufficient input data. Because each sample was sequenced to a different read depth per cell, we also calculated the average total number of UMIs represented by each sample per cluster. The results show that for samples with a previously good top node bootstrap value, downsampling below about 50 cells (Figure 5A) or 500,000 total UMIs (Figure 5B) per cluster reduced the capacity to detect genetically distinct cell populations (Supplemental Figure 8). Note that some of the healthy samples had higher bootstrap support values here compared to Figure 4A due to filtering of clusters with fewer than 500 cells in the downsampling experiment.

## 4) Discussion

A major limitation for all analyses using single-cell technology is data sparsity. When considering if variant calling from scRNAseq reads will provide enough information to distinguish tumor from normal cells, an important factor is how many genomic positions have adequate coverage to accurately call variants. Not only are sequencing reads limited to expressed genes, but for most 10x scRNAseq data, only the 3’ UTR is covered. This dramatically limits the number of sites comparable between cells, especially between transcriptionally distinct cell types. Our results showed that while many sites were covered in each cell, a small proportion of those were called as variant, even when using as low as 2X read depth cutoff (Figure 1A). The number of useful variants is restricted further when comparing cells, where both cells need to have variants at shared genomic positions. This limits informative sites to around 22 positions on average, even when using a 2X read depth threshold, with many cells having no shared variant sites (Figure 1B). Using a more reasonable read depth requirement to more accurately call variants means that most cells lack any shared variants (Figure 1B). We also demonstrated that relaxing conventional read depth thresholds, generally 15x or more (Ajay, et al., 2011; Bentley, et al., 2008; Kishikawa, et al., 2019; Koboldt, 2020; Koboldt, et al., 2010) for variant calling results in unacceptably high rates of inaccurate variant calls (Figure 1D).

Given this, it is apparent that measuring genetic distance between individual cells using variants detected from scRNAseq data is not practical. To distinguish genetically distinct cell populations, we need to increase the number of detected variants in our data. If these populations are transcriptionally distinct, we could combine data from transcriptionally similar cells to increase sequencing depth. We therefore used a pseudobulk methodology, combining sequencing reads from transcriptionally clustered cells as a single “sample” for the purpose of variant calling. We used Louvain-based transcriptional clustering as implemented in Seurat for this study. However other clustering methods such as meta-cell analysis would also work (Baran, et al., 2019). Even using this approach, however, we considered that we may have few useful sites if we used the recommended read depth thresholds for variant calling (Ajay, et al., 2011). We therefore chose to quantify how relaxing the read depth thresholds would influence data quality.

A high read depth threshold for variant calling is used to overcome incorrect variant calls caused by inaccuracies in the sequencing data (Ajay, et al., 2011). Reducing the minimum read depth allows identification of more variants across the genome, increasing data *quantity* while reducing data *quality* and increasing false positive variant calls. Variant calling in the context in which we are working, then, is an exercise in finding the optimal tradeoff between detection sensitivity and specificity to enable reliable identification of tumor cells or subclones. Data from scRNAseq is distinct from whole genome or exome data, so we quantified false positive variant calls at varying read depth cutoffs using 10x Flex datasets. We found that while the error rate seems low, given the high number of positions covered in a typical dataset and the fact that most sites match the reference, many variant sites will be incorrectly called. Given the number of sites covered in typical scRNAseq data and the calculated error rate per genomic position covered at each read depth, at a read depth of 2, more than half of the called variants are inaccurate. This dramatically improves when using more commonly accepted thresholds. Our data, therefore, show how excessive relaxing of read depth thresholds for variant calling introduces extremely high levels of noise to the analysis.

We validated our approach using several methods. We showed that pseudobulked cell populations from two genetically distinct strains of mouse were easily distinguished. The large genetic distance between the strains made this an easy validation and is unlikely to exist when comparing normal and tumor cells from a patient. To test genetically related samples, we turned to ground truth datasets from syngeneic mouse models of cancer containing transgenic marker genes. These cell lines also derive from the same mouse strain they were implanted into and so should share a high level of genetic similarity to the host, akin to tumor cells and the patients’ normal cells. Our success with these samples validates our approach. Additionally, the inability to distinguish K7M2 osteosarcoma cells from host BALB/c demonstrates the reliance of this approach on adequate genetic distance between populations.

Given successful validation, we chose to test samples where we have no ground truth. We used two datasets: 1) one anticipated to be distinguishable from normal cells, and 2) the K7M2 dataset as a negative control. While the negative control dataset had no genetically distinct subpopulations, the F420 cells could be distinguished from the C57BL/6 cells. The tumor cells also seemed to show intra-tumoral genetic heterogeneity. One result of note from the analysis of the C57BL/6 and F420 samples is that initially, the tumor cells did not separate from the normal cells. This was due to a cluster that had a roughly even mixture of normal and tumor cells. Clustering at a higher resolution split this cluster, and afterwards, the analysis correctly identified the tumor and normal clusters. This point highlights how the pseudobulking approach is reliant on accurate clustering of cells and that mixed clusters can reduce accuracy.

We next decided to apply our approach to human tumor and healthy samples. Healthy samples in this context were negative controls because they should have no genetic substructure. We analyzed various cancer types to investigate performance in samples with different mutation accumulation rates (Lawrence, et al., 2013). Given the nature of our approach and observations so far, tumor cells with few mutations may lack adequate genetic distance to be confidently identified. The chosen tumor subtypes ranged from mutationally “cold” to “hot”. The data also had highly variable cell counts and read depths to help evaluate the limitations of our approach. For tumor samples, we saw interesting patterns in our ability to distinguish normal from tumor cells in different cancers. Osteosarcoma and ovarian cancers have always had genetically distinct populations. Distinct populations were found in roughly half of the breast cancer and Ewing sarcoma samples. For prostate cancer, distinct populations were found only when using a read depth threshold of 20 or 30. The clear cell carcinoma of the kidney sample never had distinct populations.

Differences in success rates likely arise from varying genetic distances between cancer types, but some samples could have few tumor cells or low sequencing depth. For the analysis of the osteosarcoma sample from Figure 4, it is worth noting that the genetically distinct clusters do not include clusters that could be easily mistaken for tumor cells (clusters 6 and 9 in Figure 4E). Our approach shows these populations grouping with healthy cells, demonstrating how the results provide clarity where there had been ambiguity. Nodes within the tumor portion of the tree with very high bootstrap values highlight the potential of our method to reveal intra-tumor genetic substructure. Further experiments would be needed to validate this, however.

A critical question for our approach is the amount of data required to successfully identify genetically distinct populations. This is especially important where tumor cell clusters have few cells or limited sequencing depth to ensure adequate variant count. Our downsampling analysis suggests that when the clusters contained fewer than around 50 cells, or 500,000 UMIs per cluster, the discriminatory power of our approach began to degrade (Figure 5A and B). We also note that when only two cells were included per cluster, some healthy samples incorrectly had high bootstrap values, suggesting that noise and false positive rates are higher with few cells per cluster. This analysis also demonstrated the importance of genetic distance for our approach again. We could sometimes distinguish tumor from normal cells even with very few cells per cluster in samples with high mutational load. This also suggests that mutationally “cold” samples would require high cell counts and sequencing depth.

The bootstrapping values in our results give a metric for confidence in the divisions between populations of genetically distinct cells. Values as low as 70% are sometimes used in traditional phylogenetics (Hillis and Bull, 1993; Holmes, 2003; Lemoine, et al., 2018; Simon, 2022). However, based on validation analyses, we chose to employ a more stringent bootstrap value of 85% to ensure high confidence in our results.

Our approach is implemented as a computationally efficient R package. We take advantage of computational clusters where available and can submit batch jobs using slurm or sun grid engine job schedulers to parallelize many of the operations. Run time is dependent on computational resources, but pulling sequencing reads from the bam file and calling variants on each cluster is parallelized, with each taking around eleven minutes and using roughly half a gigabyte of RAM on average. Merging data from each cluster into a single file takes less than two minutes on average, while using only 43Mb of RAM, and calculating the hierarchical clustering tree and bootstrapping takes a bit less than two minutes on average and uses an average of around one GB of RAM. Processing one sample on a computational cluster averaged around 15 minutes. If parallelization is not possible, the processing time will scale with the number of clusters and data quantity.

There are several limitations to the approach outlined here. We have traded the ability to interrogate single cells for increased variant calling accuracy and quantity by pseudobulking cells. This means that accurate cell clustering is critical to the success of this approach. For most samples, this should be a trivial limitation; however, careful clustering is critical where tumor cells are transcriptionally similar to normal cell types. Another limitation of this method is the dependence on adequate mutational burden to distinguish cell populations. This method also relies on adequate cell numbers and sequencing depth in both tumor and healthy samples, requiring a minimum of 50 cells or 500,000 UMIs per cluster. Lastly, this approach also requires a bam file, which may not always be readily available.

## 5) Conclusions

We have outlined here an approach to distinguish genetically distinct populations of cells within scRNAseq data. We quantified how the calculation of genetic distance in scRNAseq is limited by data sparsity and that the temptation to gain additional covered genomic positions through relaxation of minimum read depth standards leads to inaccurately called variants. There is a natural inclination to relax traditionally stringent data standards and processing to try to get as much information out of this data as possible. Other approaches have called variants in scRNAseq data, sometimes calling variants from as few as one read (Lareau, et al., 2024; Ludwig, et al., 2019; Marot-Lassauzaie, et al., 2024; Weng, et al., 2024; Xu, et al., 2019). The data presented here demonstrates why compromising normal standards must be resisted. While modern sequencing technologies are remarkably accurate, the large number of data points collected in genomic analysis means that even small inaccuracies can generate appreciable proportions of false positive results. Given the current technological limitations of scRNAseq, variant calling on individual cells is going to be severely limited in scope and may make certain types of analyses completely untenable.

Thoughtful use of pseudobulking of the data can expand the analyses that can be performed. Extensive validation revealed the power of this approach and key limitations related to data quality and quantity.

We would like to highlight that this approach, while effective, is best used in coordination of orthogonal methods to correctly identify tumor cells. Tumor cell identification is a challenging problem requiring multiple lines of evidence to produce accurate results.

The primary output of our method is confidently identified tumor cells in complex samples, leading to improved biological interpretations and insights from the data. Single-cell technologies are powerful tools to inform tumor biology but require careful analytical approaches. Improved methods to produce accurate interpretations using this technology will assist in our common goal of finding safe and effective treatments for cancer.

## Acknowledgements

We gratefully acknowledge the members of the Nationwide Children’s Hospital weekly bioinformatics meeting for feedback during development.

## Author contributions

Matthew V. Cannon (Conceptualization, Data Curation, Formal Analysis, Investigation, Methodology, Project Administration, Resources, Software, Supervision, Validation, Visualization, Writing – Original Draft, Writing – Review & Editing), Matthew J. Gust (Conceptualization, Formal Analysis, Methodology, Software, Validation, Visualization, Writing – Review & Editing), Amy C. Gross (Investigation, Methodology, Project Administration, Resources, Supervision, Writing – Review & Editing), Maren Cam (Investigation, Methodology, Writing – Review & Editing), James B. Reinecke (Investigation, Methodology, Resources, Supervision, Writing – Review & Editing), Leyre Jimenez Garcia (Investigation, Methodology, Writing – Review & Editing), Corinne Strawser (Validation, Writing – Review & Editing), Lindsay Ryan (Investigation, Methodology, Writing – Review & Editing), Melissa Sammons (Investigation, Methodology, Writing – Review & Editing), Cheng-Zhong Zhang (Methodology, Supervision, Writing – Review & Editing), Ryan D. Roberts (Conceptualization, Funding Acquisition, Methodology, Project Administration, Resources, Supervision, Validation, Writing – Review & Editing)

## Funding

This work was funded through support to Dr. Roberts from the Alex’s Lemonade Stand Crazy 8 initiative (RDR, CZZ), a NIH NCI R01 grant (R01CA260178, RDR) and a NIH NCI U54 grant (1U54CA232561 RDR).

## Data availability

We have implemented scanBit as an R package which is publicly available here: https://github.com/kidcancerlab/scanBit. Code to reproduce the validation experiments outlined in this manuscript is available here: https://github.com/kidcancerlab/24_validate_snvs. Data generated to support these analyses is available through the NCBI GEO database under accession GSE316454. All other data are publicly available as outlined in the methods.

## Notes

### Competing Interest Statement

The authors have declared no competing interest.

