## Supplemental figures and tables for "SCANBIT facilitates identification of tumor cell populations in scRNAseq data using pseudobulked SNV calls": combined.pdf

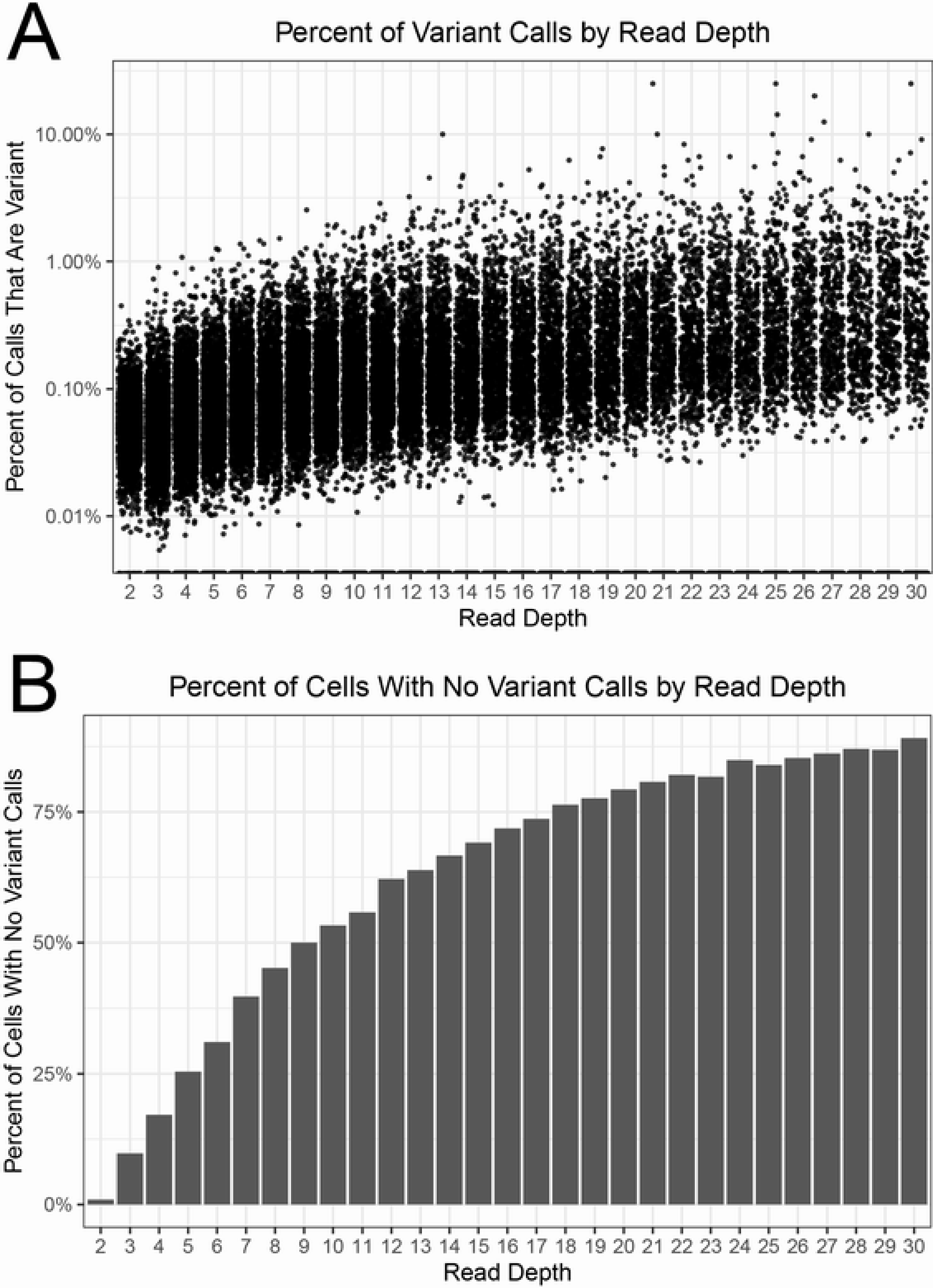
A
Percent of Variant Calls by Read Depth
Percent of Calls That Are Variant
10.00%
1.00%
0.10%
0.01%
2 3 4 5 6 7 8 9 10 11 12 13 14 15 16 17 18 19 20 21 22 23 24 25 26 27 28 29 30
Read Depth
B
Percent of Cells With No Variant Calls by Read Depth
Percent of Cells With No Variant Calls
75%
50%
25%
0%
2 3 4 5 6 7 8 9 10 11 12 13 14 15 16 17 18 19 20 21 22 23 24 25 26 27 28 29 30
Read Depth

**Supplemental Figure 1: Percent of variant calls in individual cell variant calls**
We calculated the percent of percent of all sites called as variant (A) and the proportion of cells with no variant calls (B) in individual cells from osteosarcoma and healthy bone scRNAseq data.

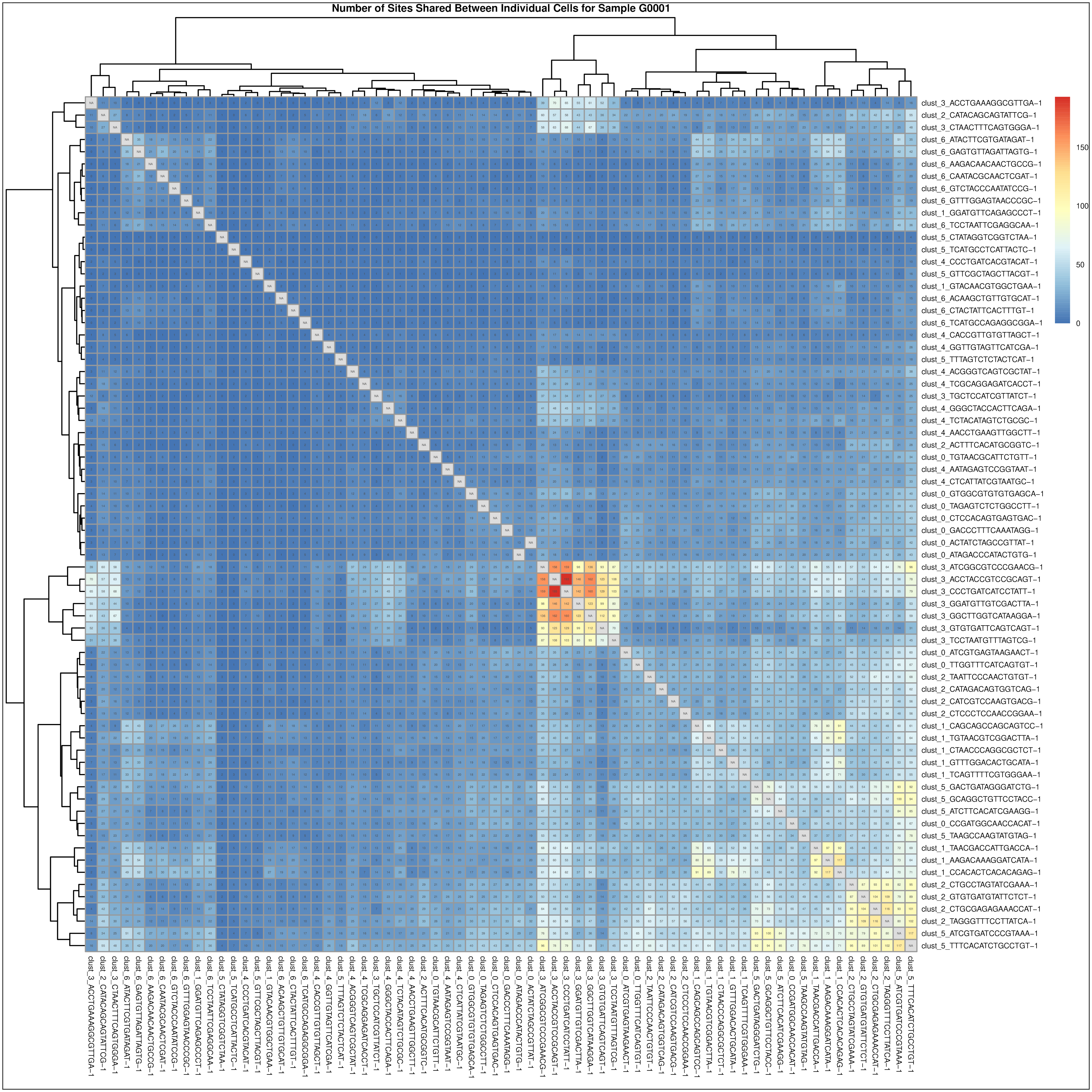
Number of Sites Shared Between Individual Cells for Sample G0001
150
100
50
0
clust_3_ACCTGAAAGGCGTTGA-1
clust_2_CATACAGCAGTATTCG-1
clust_3_CTAACTTTCAGTGGGA-1
clust_6_ATACTTCGTGATAGAT-1
clust_6_GAGTGTTAGATTAGTG-1
clust_6_AAGACAACAACTGCCG-1
clust_6_CAATACGCAACTCGAT-1
clust_6_GTCTACCCAATATCCG-1
clust_6_GTTTGGAGTAACCCGC-1
clust_1_GGATGTTCAGAGCCCT-1
clust_6_TCCTAATTCGAGGCAA-1
clust_5_CTATAGGTCGGTCTAA-1
clust_5_TCATGCCTCATTACTC-1
clust_4_CCCTGATCACGTACAT-1
clust_5_GTTCGCTAGCTTACGT-1
clust_1_GTACAACGTGGCTGAA-1
clust_6_ACAAGCTGTTGTGCAT-1
clust_6_CTACTATTCACTTTGT-1
clust_6_TCATGCCAGAGGCGGA-1
clust_4_CACCGTTGTGTTAGCT-1
clust_4_GGTTGTAGTTCATCGA-1
clust_5_TTTAGTCTCTACTCAT-1
clust_4_ACGGGTCAGTCGCTAT-1
clust_4_TCGCAGGAGATCACCT-1
clust_3_TGCTCCATCGTTATCT-1
clust_4_GGGCTACCACTTCAGA-1
clust_4_TCTACATAGTCTGCGC-1
clust_4_AACCTGAAGTTGGCTT-1
clust_2_ACTTTCACATGCGGTC-1
clust_0_TGTAACGCATTCTGTT-1
clust_4_AATAGAGTCCGGTAAT-1
clust_4_CTCATTATCGTAATGC-1
clust_0_GTGGCGTGTGTGAGCA-1
clust_0_TAGAGTCTCTGGCCTT-1
clust_0_CTCCACAGTGAGTGAC-1
clust_0_GACCCTTTCAAATAGG-1
clust_0_ACTATCTAGCCGTTAT-1
clust_0_ATAGACCCATACTGTG-1
clust_3_ATCGGCGTCCCGAACG-1
clust_3_ACCTACCGTCCGCAGT-1
clust_3_CCCTGATCATCCTATT-1
clust_3_GGATGTTGTCGACTTA-1
clust_3_GGCTTGGTCATAAGGA-1
clust_3_GTGTGATTCAGTCAGT-1
clust_3_TCCTAATGTTTAGTCG-1
clust_0_ATCGTGAGTAAGAACT-1
clust_0_TTGGTTTCATCAGTGT-1
clust_2_TAATTCCCAACTGTGT-1
clust_2_CATAGACAGTGGTCAG-1
clust_2_CATCGTCCAAGTGACG-1
clust_2_CTCCCTCCAACCGGAA-1
clust_1_CAGCAGCCAGCAGTCC-1
clust_1_TGTAACGTCGGACTTA-1
clust_1_CTAACCCAGGCGCTCT-1
clust_1_GTTTGGACACTGCATA-1
clust_1_TCAGTTTTCGTGGGAA-1
clust_5_GACTGATAGGGATCTG-1
clust_5_GCAGGCTGTTCCTACC-1
clust_5_ATCTTCACATCGAAGG-1
clust_0_CCGATGGCAACCACAT-1
clust_5_TAAGCCAAGTATGTAG-1
clust_1_TAACGACCATTGACCA-1
clust_1_AAGACAAAGGATCATA-1
clust_1_CCACACTCACACAGAG-1
clust_2_CTGCCTAGTATCGAAA-1
clust_2_GTGTGATGTATTCTCT-1
clust_2_CTGCGAGAGAAACCAT-1
clust_2_TAGGGTTTCCTTATCA-1
clust_5_ATCGTGATCCCGTAAA-1
clust_5_TTTCACATCTGCCTGT-1

**Number of Sites Shared Between Individual Cells for Sample G0002**

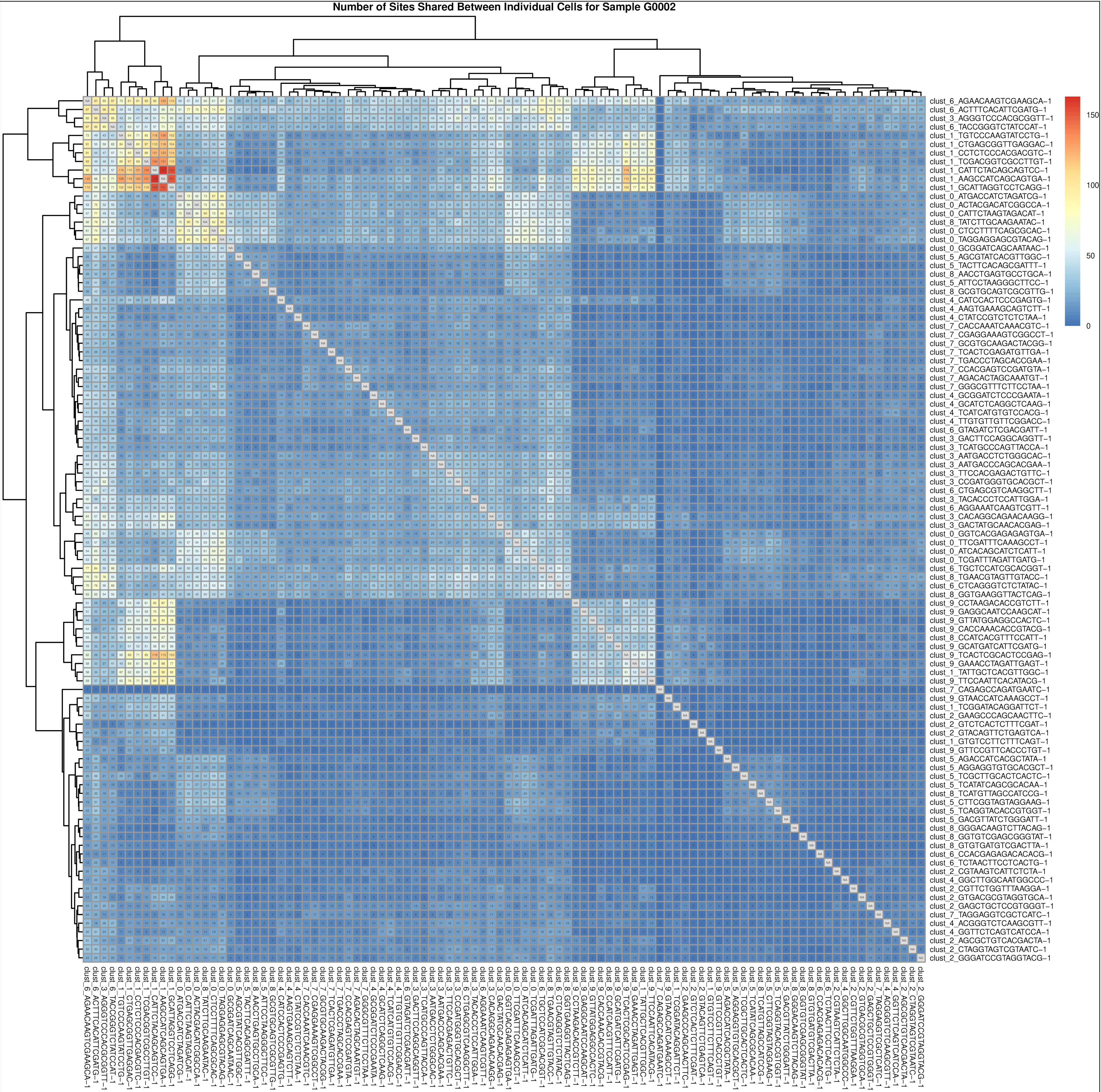

Number of Sites Shared Between Individual Cells for Sample G0003

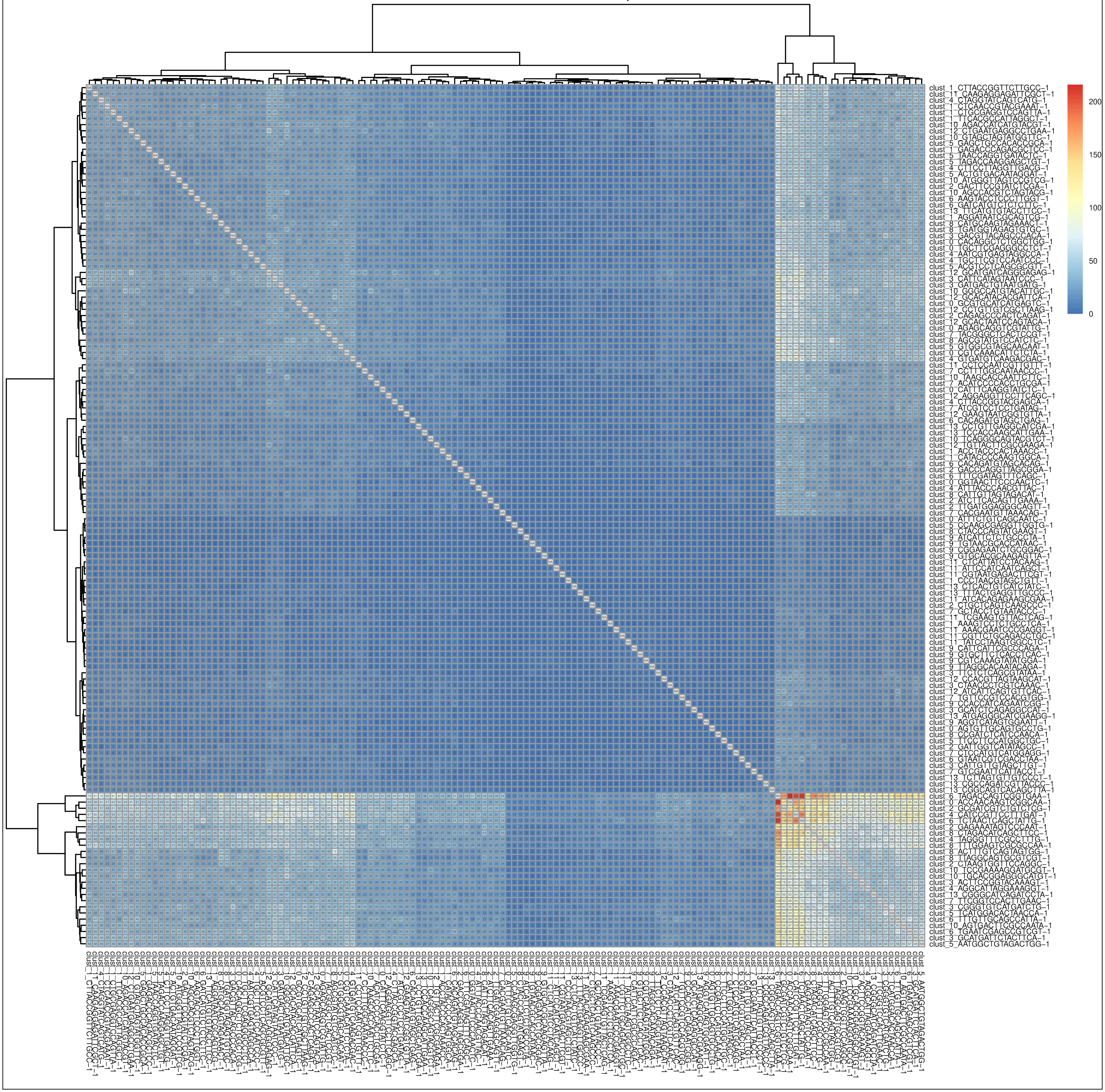

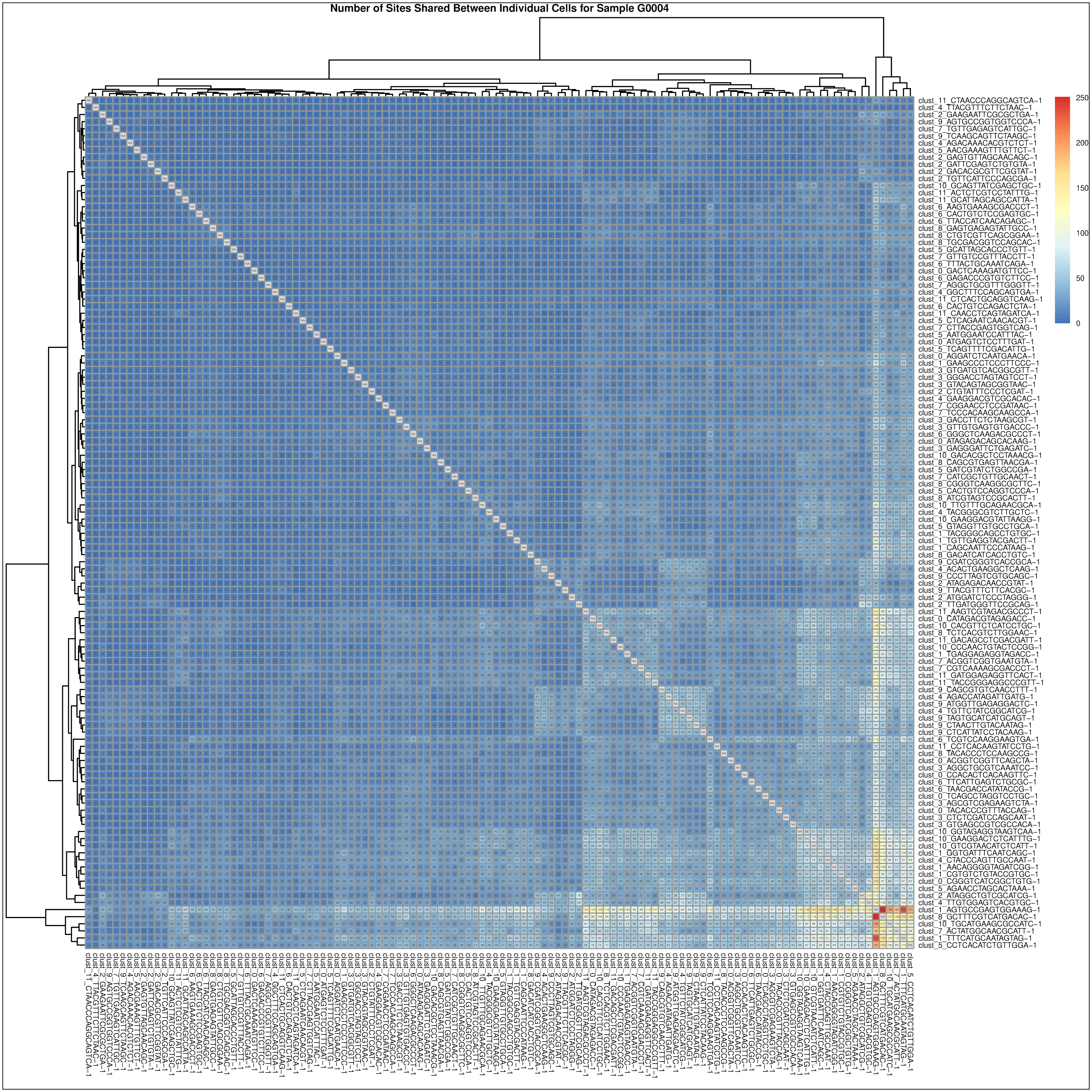
Number of Sites Shared Between Individual Cells for Sample G0004
250
200
150
100
50
0

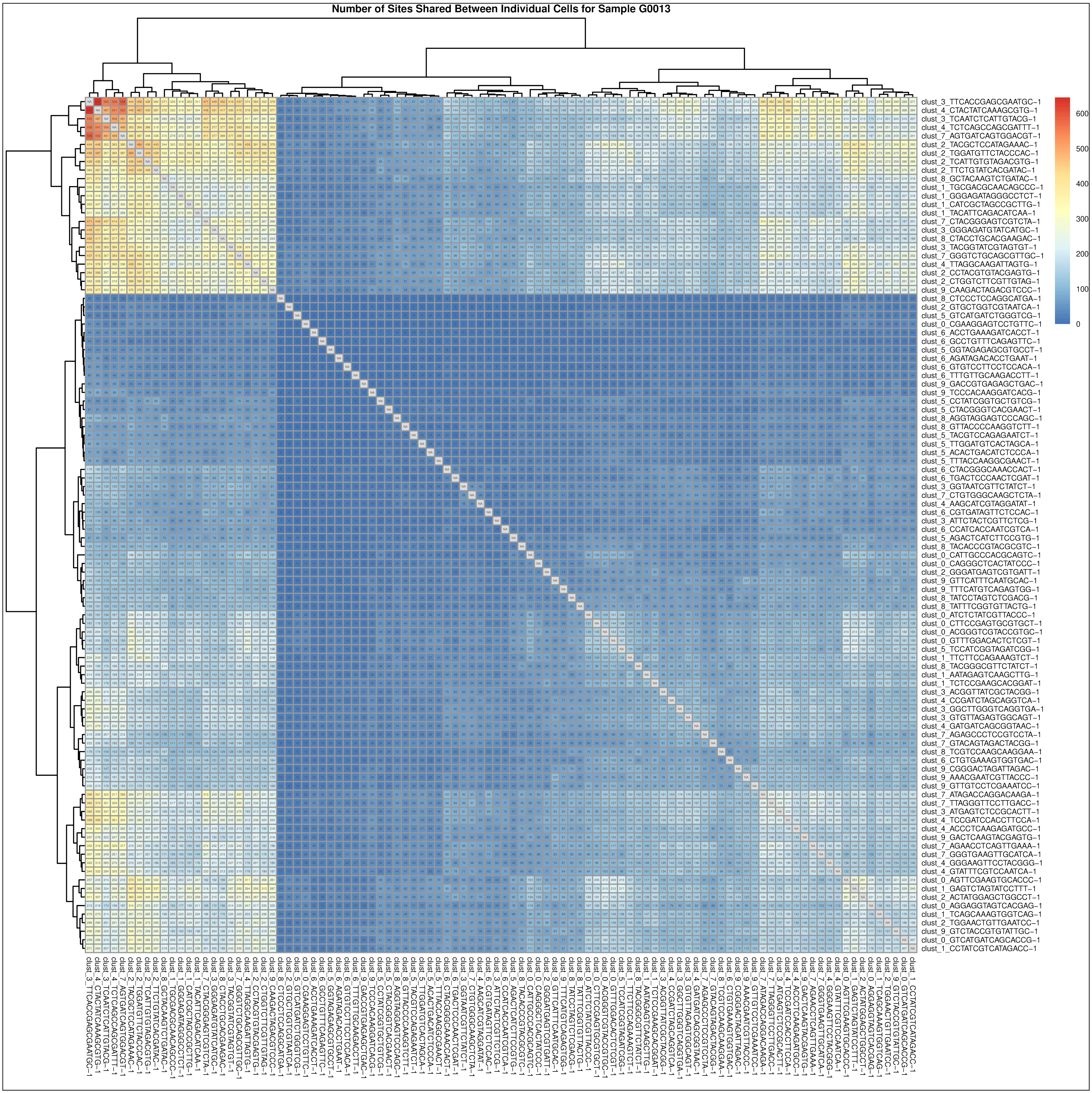
Number of Sites Shared Between Individual Cells for Sample G0013
600
500
400
300
200
100
0
clust_3_TTCACCGAGCGAATGC-1
clust_4_CTACTATCAAAGCGTG-1
clust_3_TCAATCTCATTGTACG-1
clust_4_TCTCAGCCAGCGATTT-1
clust_7_AGTGATCAGTGGACGT-1
clust_2_TACGCTCCATAGAAAC-1
clust_2_TGGATGTTCTACCCAC-1
clust_2_TCATTGTGTAGACGTG-1
clust_2_TTCTGTATCACGATAC-1
clust_8_GCTACAAGTCTGATAC-1
clust_1_TGCGACGCAACAGCCC-1
clust_1_GGGAGATAGGGCCTCT-1
clust_1_CATCGCTAGCCGCTTG-1
clust_1_TACATTCAGACATCAA-1
clust_7_CTACGGGAGTCGTCTA-1
clust_3_GGGAGATGTATCATGC-1
clust_8_CTACCTGCACGAAGAC-1
clust_3_TACGGTATCGTAGTGT-1
clust_7_GGGTCTGCAGCGTTGC-1
clust_4_TTAGGCAAGATTAGTG-1
clust_2_CCTACGTGTACGAGTG-1
clust_2_CTGGTCTTCGTTGTAG-1
clust_9_CAAGACTAGACGTCCC-1
clust_8_CTCCCTCCAGGCATGA-1
clust_2_GTGCTGGTCGTAATCA-1
clust_5_GTCATGATCTGGGTCG-1
clust_0_CGAAGGAGTCCTGTTC-1
clust_6_ACCTGAAAGATCACCT-1
clust_6_GCCTGTTTCAGAGTTC-1
clust_5_GGTAGAGAGCGTGCCT-1
clust_6_AGATAGACACCTGAAT-1
clust_6_GTGTCCTTCCTCCACA-1
clust_6_TTTGTTGCAAGACCTT-1
clust_9_GACCGTGAGAGCTGAC-1
clust_9_TCCCACAAGGATCACG-1
clust_5_CCTATCGGTGCTGTCG-1
clust_5_CTACGGGTCACGAACT-1
clust_8_AGGTAGGAGTCCCAGC-1
clust_8_GTTACCCCAAGGTCTT-1
clust_5_TACGTCCAGAGAATCT-1
clust_5_TTGGATGTCACTAGCA-1
clust_5_ACACTGACATCTCCCA-1
clust_5_TTTACCAAGGCGAACT-1
clust_6_CTACGGGCAAACCACT-1
clust_6_TGACTCCCAACTCGAT-1
clust_3_GGTAATCGTTCTATCT-1
clust_7_CTGTGGGCAAGCTCTA-1
clust_4_AAGCATCGTAGGATAT-1
clust_6_CGTGATAGTTCTCCAC-1
clust_3_ATTCTACTCGTTCTCG-1
clust_6_CCATCACCAATCGTCA-1
clust_5_AGACTCATCTTCCGTG-1
clust_8_TACACCCGTACGCGTC-1
clust_0_CATTGCCCACGCAGTC-1
clust_0_CAGGGCTCACTATCCC-1
clust_2_GGGATGAGTCGTGATT-1
clust_9_GTTCATTTCAATGCAC-1
clust_9_TTTCATGTCAGAGTGG-1
clust_8_TATCCTAGTCTCGACG-1
clust_8_TATTTCGGTGTTACTG-1
clust_0_ATCTCTATCGTTACCC-1
clust_0_CTTCCGAGTGCGTGCT-1
clust_0_ACGGGTCGTACCGTGC-1
clust_0_GTTTGGACACTCTCGT-1
clust_5_TCCATCGGTAGATCGG-1
clust_1_TTCTTCCAGAAAGTCT-1
clust_8_TACGGGCGTTCTATCT-1
clust_1_AATAGAGTCAAGCTTG-1
clust_1_TCTCCGAAGCACGGAT-1
clust_3_ACGGTTATCGCTACGG-1
clust_4_CCGATCTAGCAGGTCA-1
clust_3_GGCTTGGGTCAGGTGA-1
clust_3_GTGTTAGAGTGGCAGT-1
clust_4_GATGATCAGCGGTAAC-1
clust_7_AGAGCCCTCCGTCCTA-1
clust_7_GTACAGTAGACTACGG-1
clust_8_TCGTCCAAGCAAGGAA-1
clust_6_CTGTGAAAGTGGTGAC-1
clust_9_CGGGACTAGATTAGAC-1
clust_9_AAACGAATCGTTACCC-1
clust_9_GTTGTCCTCGAAATCC-1
clust_7_ATAGACCAGGACAAGA-1
clust_7_TTAGGGTTCCTTGACC-1
clust_3_ATGAGTCTCCGCACTT-1
clust_4_TCCGATCCACCTTCCA-1
clust_4_ACCCTCAAGAGATGCC-1
clust_9_GACTCAAGTACGAGTG-1
clust_7_AGAACCTCAGTTGAAA-1
clust_7_GGGTGAAGTTGCATCA-1
clust_4_GGGAAGTTCCTACGGG-1
clust_4_GTATTTCGTCCAATCA-1
clust_0_AGTTCGAAGTGCACCC-1
clust_1_GAGTCTAGTATCCTTT-1
clust_2_ACTATGGAGCTGGCCT-1
clust_0_AGGAGGTAGTCACGAG-1
clust_1_TCAGCAAAGTGGTCAG-1
clust_2_TGGAACTGTTGAATCC-1
clust_9_GTCTACCGTGTATTGC-1
clust_0_GTCATGATCAGCACCG-1
clust_1_CCTATCGTCATAGACC-1
NA

#### Number of Sites Shared Between Individual Cells for Sample G0014

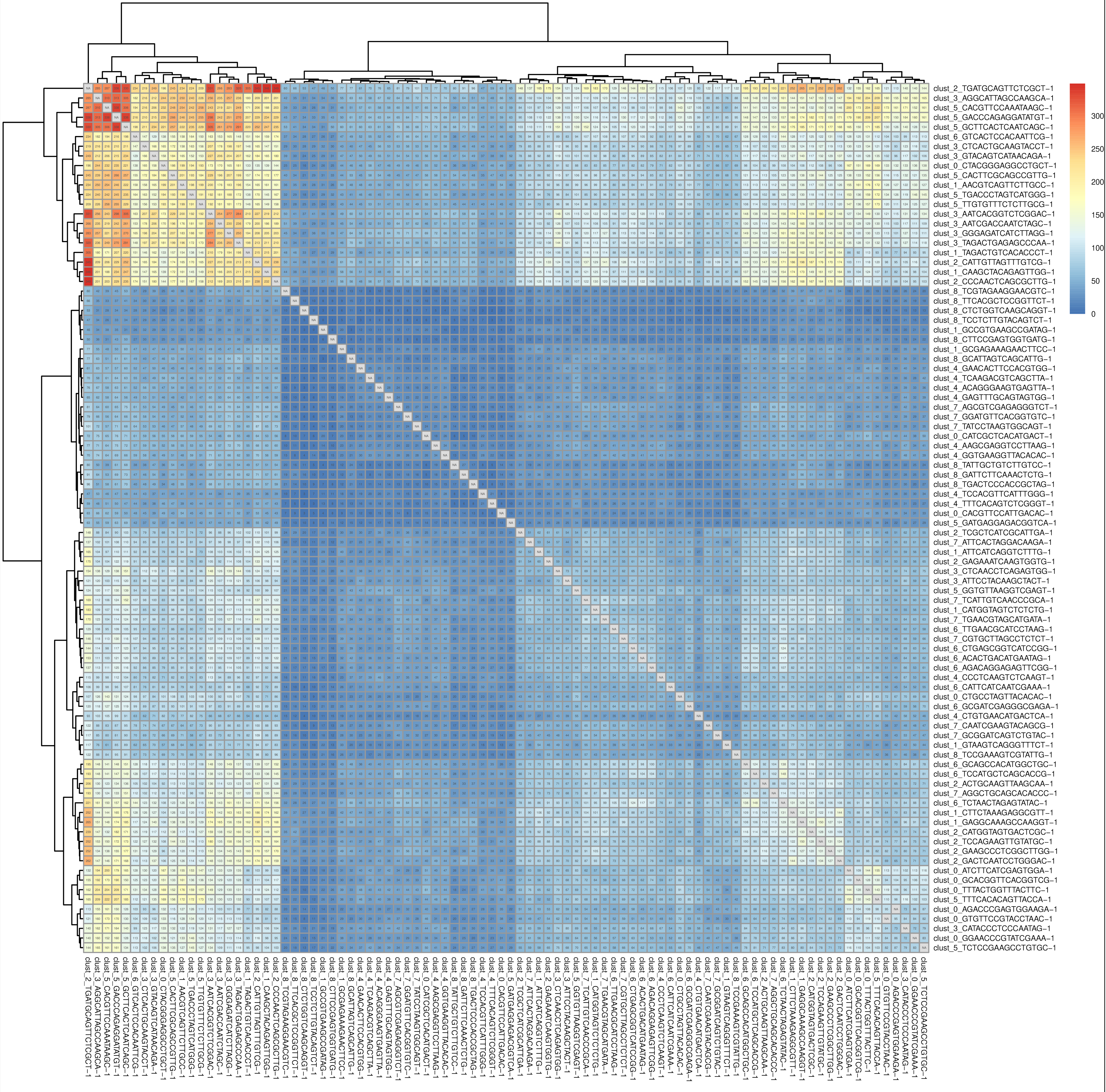

#### Number of Sites Shared Between Individual Cells for Sample G0015

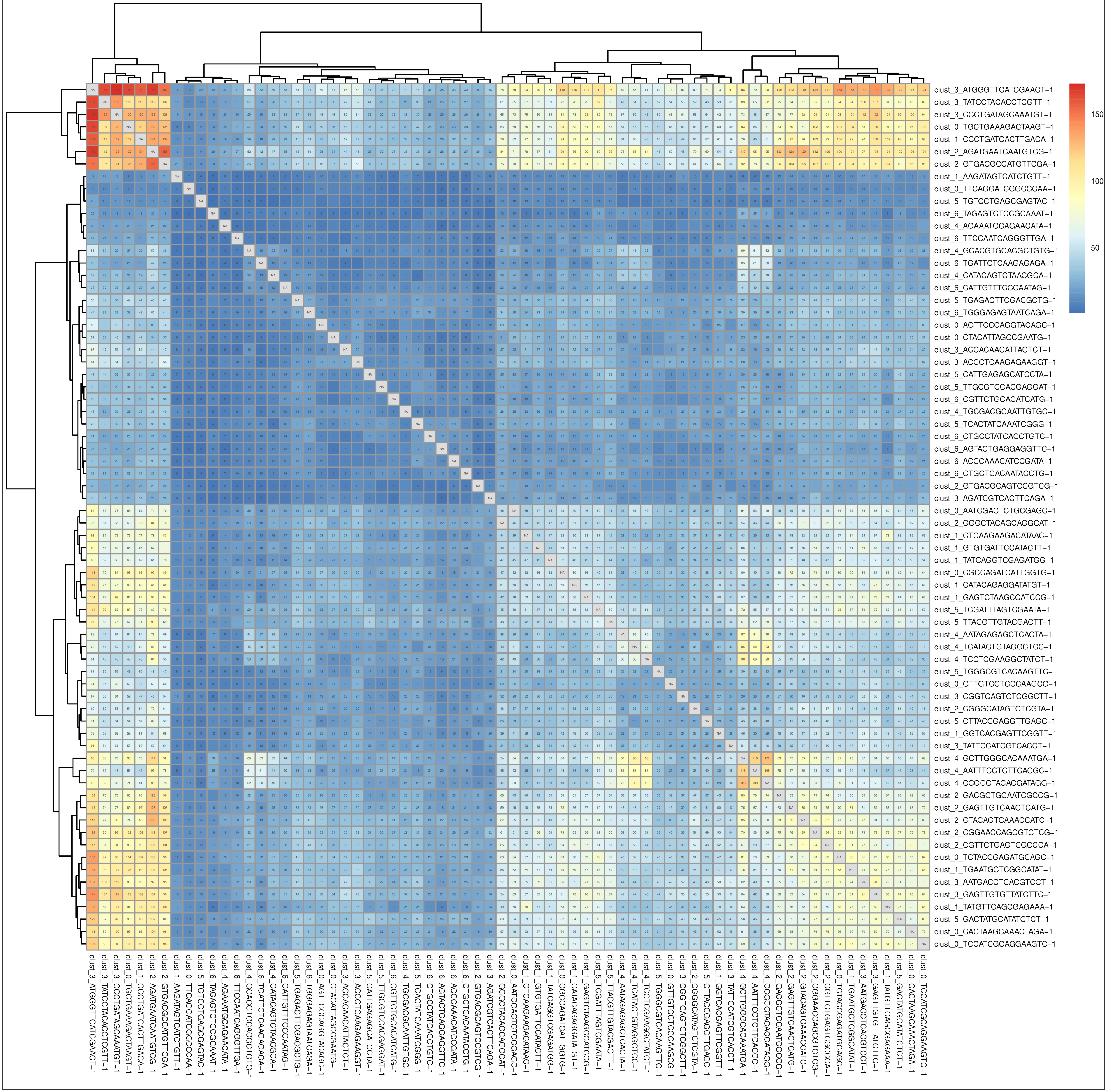

Number of Sites Shared Between Individual Cells for Sample G0016

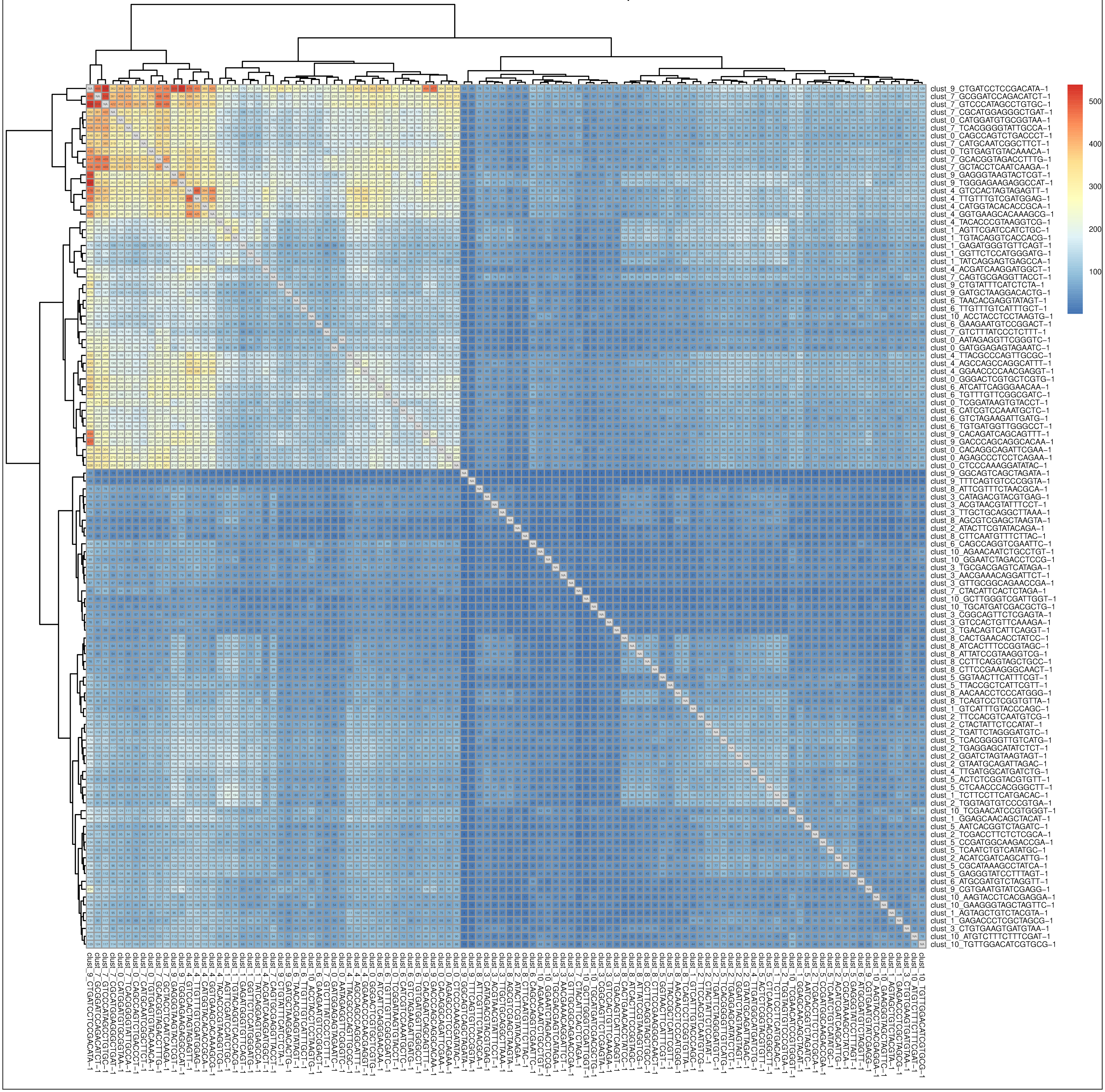

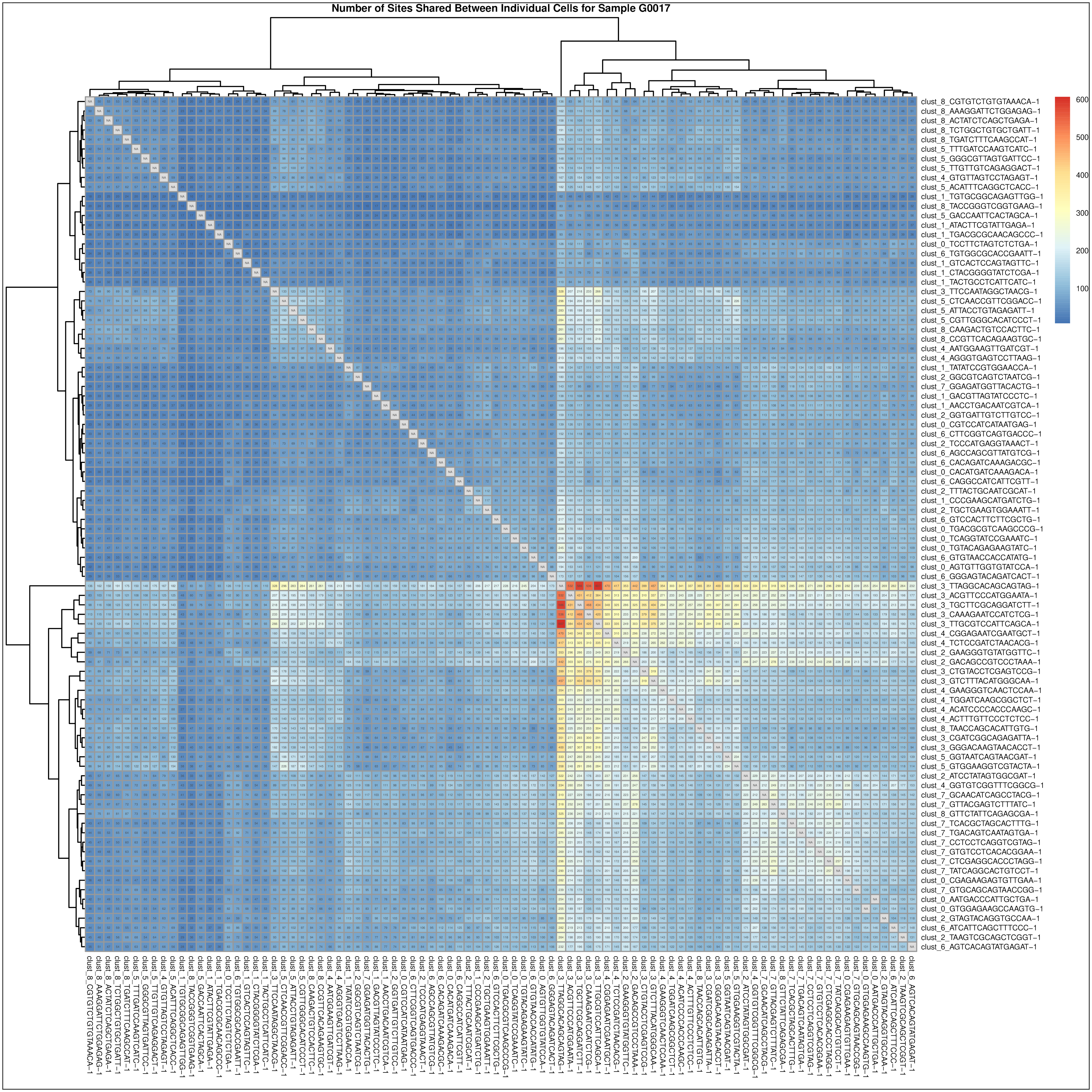
Number of Sites Shared Between Individual Cells for Sample G0017
600
500
400
300
200
100
clust_8_CGTGTCTGTGTAAACA−1
clust_8_AAAGGATTCTGGAGAG−1
clust_8_ACTATCTCAGCTGAGA−1
clust_8_TCTGGCTGTGCTGATT−1
clust_8_TGATCTTTCAAGCCAT−1
clust_5_TTTGATCCAAGTCATC−1
clust_5_GGGCGTTAGTGATTCC−1
clust_5_TTGTTGTCAGAGGACT−1
clust_4_GTGTTAGTCCTAGAGT−1
clust_5_ACATTTCAGGCTCACC−1
clust_1_TGTGCGGCAGAGTTGG−1
clust_8_TACCGGGTCGGTGAAG−1
clust_5_GACCAATTCACTAGCA−1
clust_1_ATACTTCGTATTGAGA−1
clust_1_TGACGCGCAACAGCCC−1
clust_0_TCCTTCTAGTCTCTGA−1
clust_6_TGTGGCGCACCGAATT−1
clust_1_GTCACTCCAGTAGTTC−1
clust_1_CTACGGGGTATCTCGA−1
clust_1_TACTGCCTCATTCATC−1
clust_3_TTCCAATAGGCTAACG−1
clust_5_CTCAACCGTTCGGACC−1
clust_5_ATTACCTGTAGAGATT−1
clust_5_CGTTGGGCACATCCCT−1
clust_8_CAAGACTGTCCACTTC−1
clust_8_CCGTTCACAGAAGTGC−1
clust_4_AATGGAAGTTGATCGT−1
clust_4_AGGGTGAGTCCTTAAG−1
clust_1_TATATCCGTGGAACCA−1
clust_2_GGCGTCAGTCTAATCG−1
clust_7_GGAGATGGTTACACTG−1
clust_1_GACGTTAGTATCCCTC−1
clust_1_AACCTGACAATCGTCA−1
clust_2_GGTGATTGTCTTGTCC−1
clust_0_CGTCCATCATAATGAG−1
clust_6_CTTCGGTCAGTGACCC−1
clust_2_TCCCATGAGGTAAACT−1
clust_6_AGCCAGCGTTATGTCG−1
clust_6_CACAGATCAAAGACGC−1
clust_0_CACATGATCAAAGACA−1
clust_6_CAGGCCATCATTCGTT−1
clust_2_TTTACTGCAATCGCAT−1
clust_1_CCCGAAGCATGATCTG−1
clust_2_TGCTGAAGTGGAAATT−1
clust_6_GTCCACTTCTTCGCTG−1
clust_0_TGACGCGTCAAGCCCG−1
clust_0_TCAGGTATCCGAAATC−1
clust_0_TGTACAGAGAAGTATC−1
clust_6_GTGTAACCACCATATG−1
clust_0_AGTGTTGGTGTATCCA−1
clust_6_GGGAGTACAGATCACT−1
clust_3_TTAGGCACAGCAGTAG−1
clust_3_ACGTTCCCATGGAATA−1
clust_3_TGCTTCGCAGGATCTT−1
clust_3_CAAAGAATCCATCTCG−1
clust_3_TTGCGTCCATTCAGCA−1
clust_4_CGGAGAATCGAATGCT−1
clust_4_TCTCCGATCTAACACG−1
clust_2_GAAGGGTGTATGGTTC−1
clust_2_GACAGCCGTCCCTAAA−1
clust_3_CTGTACCTCGAGTCCG−1
clust_3_GTCTTTACATGGGCAA−1
clust_4_GAAGGGTCAACTCCAA−1
clust_4_TGGATCAAGCGGCTCT−1
clust_4_ACATCCCCACCCAAGC−1
clust_4_ACTTTGTTCCCTCTCC−1
clust_8_TAACCAGCACATTGTG−1
clust_3_CGATCGGCAGAGATTA−1
clust_3_GGGACAAGTAACACCT−1
clust_5_GGTAATCAGTAACGAT−1
clust_5_GTGGAAGGTCGTACTA−1
clust_2_ATCCTATAGTGGCGAT−1
clust_4_GGTGTCGGTTTCGGCG−1
clust_7_GCAACATCAGCCTACG−1
clust_7_GTTACGAGTCTTTATC−1
clust_8_GTTCTATTCAGAGCGA−1
clust_7_TCACGCTAGCACTTTG−1
clust_7_TGACAGTCAATAGTGA−1
clust_7_CCTCCTCAGGTCGTAG−1
clust_7_GTGTCCTCACACGGAA−1
clust_7_CTCGAGGCACCCTAGG−1
clust_7_TATCAGGCACTGTCCT−1
clust_0_CGAGAAGAGTGTTGAA−1
clust_7_GTGCAGCAGTAACCGG−1
clust_0_AATGACCCATTGCTGA−1
clust_0_GTGGAGAAGCCAAGTG−1
clust_2_GTAGTACAGGTGCCAA−1
clust_6_ATCATTCAGCTTTCCC−1
clust_2_TAAGTCGCAGCTCGGT−1
clust_6_AGTCACAGTATGAGAT−1

**Number of Sites Shared Between Individual Cells for Sample G0018**

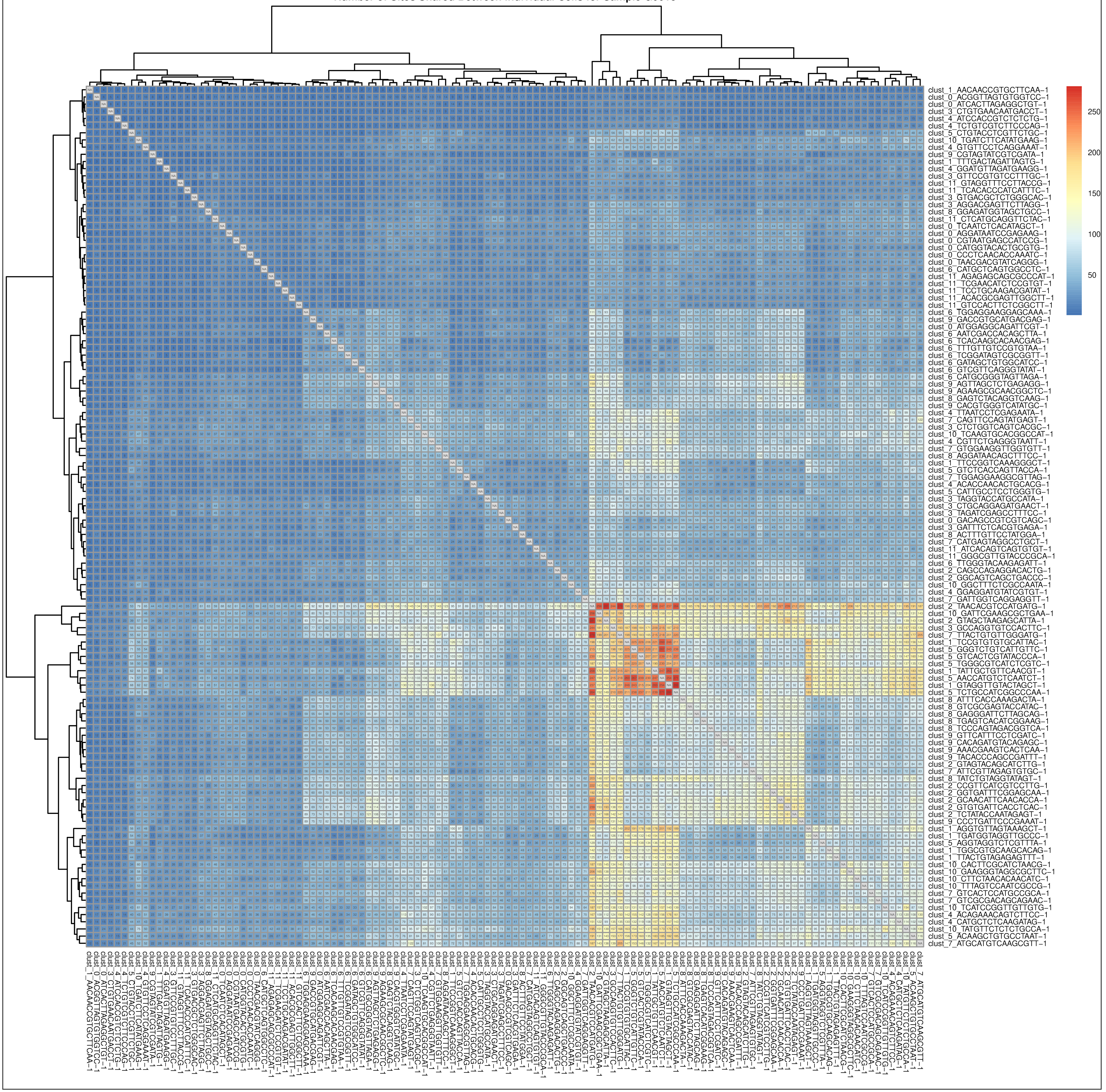

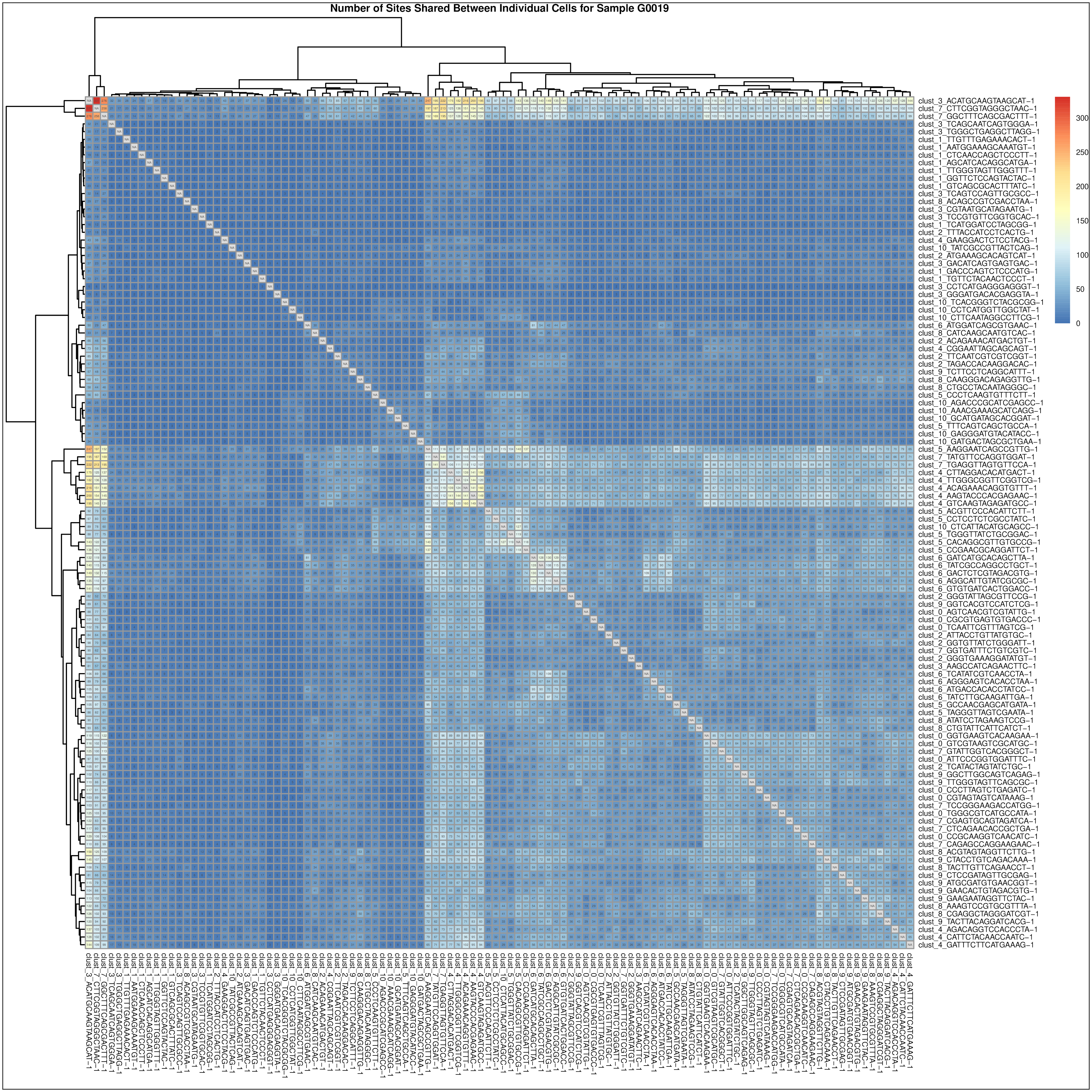
Number of Sites Shared Between Individual Cells for Sample G0019
clust_3_ACATGCAAGTAAGCAT-1
clust_7_CTTCGGTAGGGCTAAC-1
clust_7_GGCTTTCAGCGACTTT-1
clust_3_TCAGCAATCAGTGGGA-1
clust_3_TGGGCTGAGGCTTAGG-1
clust_1_TTGTTTGAGAAACACT-1
clust_1_AATGGAAAGCAAATGT-1
clust_1_CTCAACCAGCTCCCTT-1
clust_1_AGCATCACAGGCATGA-1
clust_1_TTGGGTAGTTGGGTTT-1
clust_1_GGTTCTCCAGTACTAC-1
clust_1_GTCAGCGCACTTTATC-1
clust_3_TCAGTCCAGTTGCGCC-1
clust_8_ACAGCCGTCGACCTAA-1
clust_3_CGTAATGCATAGAATG-1
clust_3_TCCGTGTTCGGTGCAC-1
clust_1_TCATGGATCCTAGCGG-1
clust_2_TTTACCATCCTCACTG-1
clust_4_GAAGGACTCTCCTACG-1
clust_10_TATCGCCGTTACTCAG-1
clust_2_ATGAAAGCACAGTCAT-1
clust_3_GACATCAGTGAGTGAC-1
clust_1_GACCCAGTCTCCCATG-1
clust_1_TGTTCTACAACTCCCT-1
clust_3_CCTCATGAGGGAGGGT-1
clust_3_GGGATGACACGAGGTA-1
clust_10_TCACGGGTCTACGCGG-1
clust_10_CCTCATGGTTGGCTAT-1
clust_10_CTTCAATAGGCCTTCG-1
clust_6_ATGGATCAGCGTGAAC-1
clust_8_CATCAAGCAATGTCAC-1
clust_2_ACAGAAACATGACTGT-1
clust_4_CGGAATTAGCAGCAGT-1
clust_2_TTCAATCGTCGTCGGT-1
clust_2_TAGACCACAAGGACAC-1
clust_9_TCTTCCTCAGGCATTT-1
clust_8_CAAGGGACAGAGGTTG-1
clust_8_CTGCCTACAATAGGGC-1
clust_5_CCCTCAAGTGTTTCTT-1
clust_10_AGACCCGCATCGAGCC-1
clust_10_AAACGAAAGCATCAGG-1
clust_10_GCATGATAGCACGGAT-1
clust_5_TTTCAGTCAGCTGCCA-1
clust_10_GAGGGATGTACATACC-1
clust_10_GATGACTAGCGCTGAA-1
clust_5_AAGGAATCAGCCGTTG-1
clust_7_TATGTTCCAGGTGGAT-1
clust_7_TGAGGTTAGTGTTCCA-1
clust_4_CTTAGGACACATGACT-1
clust_4_TTGGGCGGTTCGGTCG-1
clust_4_ACAGAAACAGGTGTTT-1
clust_4_AAGTACCCACGAGAAC-1
clust_4_GTCAAGTAGAGATGCC-1
clust_5_ACGTTCCCACATTCTT-1
clust_5_CCTCCTCTCGCCTATC-1
clust_10_CTCATTACATGCAGCC-1
clust_5_TGGGTTATCTGCGGAC-1
clust_5_CACAGGCGTTGTGCCG-1
clust_5_CCGAACGCAGGATTCT-1
clust_6_GATCATGCACAGCTTA-1
clust_6_TATCGCCAGGCCTGCT-1
clust_6_GACTCTCGTAGACGTG-1
clust_6_AGGCATTGTATCGCGC-1
clust_6_GTGTGATCACTGGACC-1
clust_2_GGGTATTAGCGTTCCG-1
clust_9_GGTCACGTCCATCTCG-1
clust_0_AGTCAACGTCGTATTG-1
clust_0_CGCGTGAGTGTGACCC-1
clust_0_TCAATTCGTTTAGTCG-1
clust_2_ATTACCTGTTATGTGC-1
clust_2_GGTGTTATCTGGGATT-1
clust_7_GGTGATTTCTGTCGTC-1
clust_2_GGGTGAAAGGATATGT-1
clust_3_AAGCCATCAGAACTTC-1
clust_6_TCATATCGTCAACCTA-1
clust_6_AGGGAGTCACACCTAA-1
clust_6_ATGACCACACCTATCC-1
clust_6_TATCTTGCAAGATTGA-1
clust_5_GCCAACGAGCATGATA-1
clust_5_TAGGGTTAGTCGAATA-1
clust_8_ATATCCTAGAAGTCCG-1
clust_8_CTGTATTCATTCATCT-1
clust_0_GGTGAAGTCACAAGAA-1
clust_0_GTCGTAAGTCGCATGC-1
clust_7_GTATTGGTCACGGGCT-1
clust_0_ATTCCCGGTGGATTTC-1
clust_2_TCATACTAGTATCTGC-1
clust_9_GGCTTGGCAGTCAGAG-1
clust_9_TTGGGTAGTTCAGCGC-1
clust_0_CCCTTAGTCTGAGATC-1
clust_0_CGTAGTAGTCATAAAG-1
clust_7_TCCGGGAAGACCATGG-1
clust_0_TGGGCGTCATGCCATA-1
clust_7_CGAGTGCAGTAGATCA-1
clust_7_CTCAGAACACCGCTGA-1
clust_0_CCGCAAGGTCAACATC-1
clust_7_CAGAGCCAGGAAGAAC-1
clust_8_ACGTAGTAGGTTCTTG-1
clust_9_CTACCTGTCAGACAAA-1
clust_8_TACTTGTTCAGAACCT-1
clust_9_CTCCGATAGTTGCGAG-1
clust_9_ATGCGATGTGAACGGT-1
clust_9_GAACACTGTAGACGTG-1
clust_9_GAAGAATAGGTTCTAC-1
clust_8_AAAGTCCGTGCGTTTA-1
clust_8_CGAGGCTAGGGATCGT-1
clust_9_TACTTACAGGATCACG-1
clust_4_AGACAGGTCCACCCTA-1
clust_4_CATTCTACAACCAATC-1
clust_4_GATTTCTTCATGAAAG-1
300
250
200
150
100
50
0

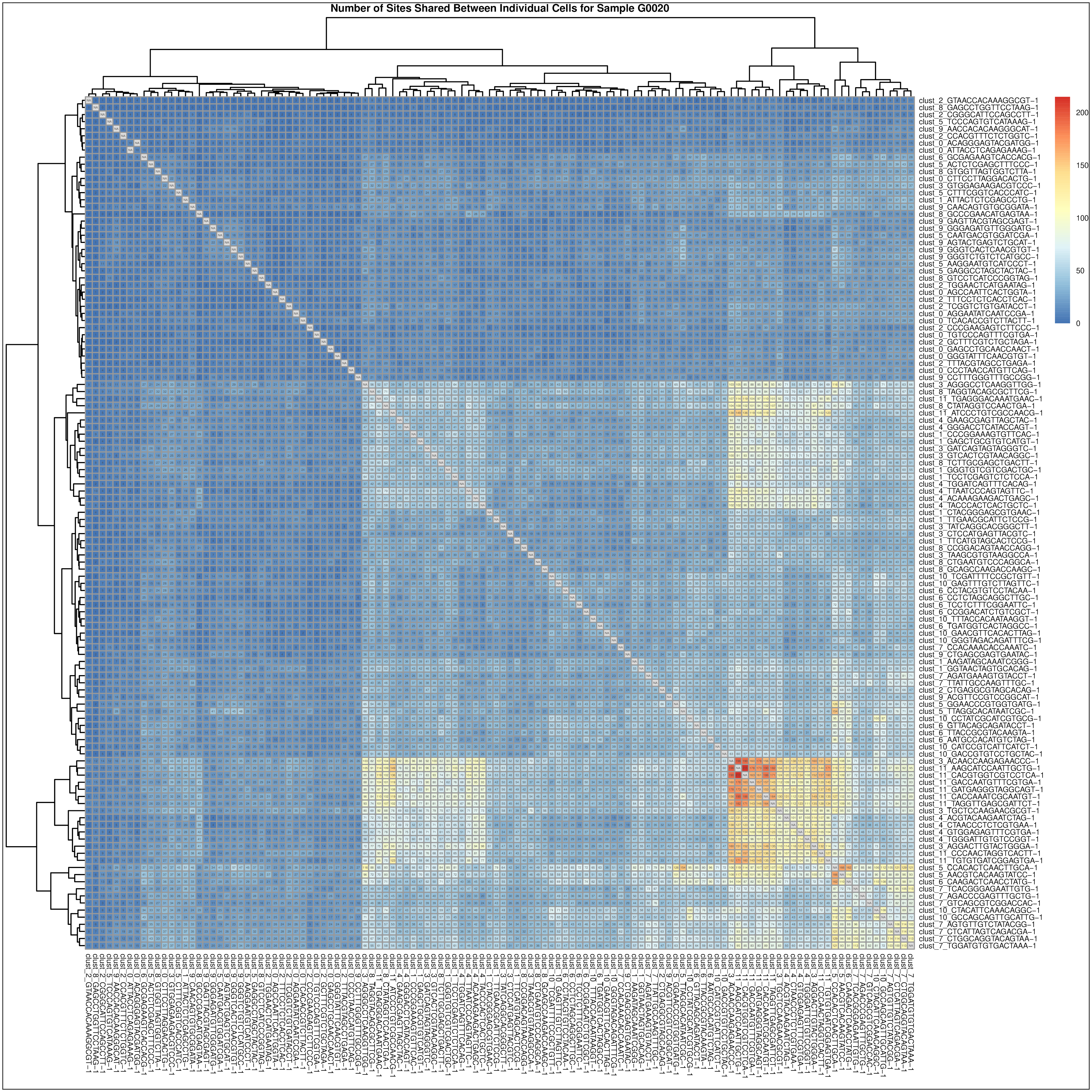
Number of Sites Shared Between Individual Cells for Sample G0020
200
150
100
50
0
clust_2_GTAACCACAAAGGCGT−1
clust_8_GAGCCTGGTTCCTAAG−1
clust_2_CGGGCATTCCAGCCTT−1
clust_5_TCCCAGTGTCATAAAG−1
clust_9_AACCACACAAGGGCAT−1
clust_2_CCACGTTTCTCTGGTC−1
clust_0_ACAGGGAGTACGATGG−1
clust_0_ATTACCTCAGAGAAAG−1
clust_6_GCGAGAAGTCACCACG−1
clust_5_ACTCTCGAGCTTTCCC−1
clust_8_GTGGTTAGTGGTCTTA−1
clust_0_CTTCCTTAGGACACTG−1
clust_3_GTGGAGAAGACGTCCC−1
clust_5_CTTTCGGTCACCCATC−1
clust_1_ATTACTCTCGAGCCTG−1
clust_9_CAACAGTGTGCGGATA−1
clust_8_GCCCGAACATGAGTAA−1
clust_9_GAGTTACGTAGCGAGT−1
clust_9_GGGAGATGTTGGGATG−1
clust_5_CAATGACGTGGATCGA−1
clust_9_AGTACTGAGTCTGCAT−1
clust_9_GGGTCACTCAAGCGTT−1
clust_9_GGGTGTCGTCTCATGC−1
clust_5_AAGGAATGTCATCCCT−1
clust_5_GAGGCCTAGCTACTAC−1
clust_8_GTCCTCATCCCGGTAG−1
clust_2_TGGAACTCATGAATAG−1
clust_0_AGCCAATTCACTGGTA−1
clust_2_TTTCCTCTCACCTCAC−1
clust_2_TCGGTCTGTGATACCT−1
clust_0_AGGAATATCAATCCGA−1
clust_0_TCACACCGTCTTACTT−1
clust_2_CCCGAAGAGTCTTCCC−1
clust_0_TGTCCCAGTTTCGTGA−1
clust_2_GCTTTCGTCTGCTAGA−1
clust_0_GAGCCTGCAACCAACT−1
clust_0_GGGTATTTCAACGTGT−1
clust_2_TTTACGTAGCCTGAGA−1
clust_0_CCCTAACCATGTTCAG−1
clust_9_CCTTTGGGTTTGCCGG−1
clust_3_AGGGCCTCAAGGTTGG−1
clust_8_TAGGTACAGCGCTTCG−1
clust_11_TGAGGGACAAATGAAC−1
clust_8_CTATAGGTCCAACTGA−1
clust_11_ATCCCTGTCGCCAACG−1
clust_4_GAAGCGAGTTAGCTAC−1
clust_4_GGGACCTCATACCAGT−1
clust_1_CCCGGAAAGTGTTCAC−1
clust_1_GAGCTGCGTGTCGATT−1
clust_3_GATCAGTAGTAGGGTC−1
clust_3_GTCACTCGTAACAGGC−1
clust_8_TCTTGCGAGCTGACTT−1
clust_1_GGGTGTCGTCGACTGC−1
clust_1_TCCTCGAGTCTCTCCA−1
clust_4_TGGATCAGTTTCACAG−1
clust_4_TTAATCCCAGTAGTTC−1
clust_4_ACAAAGAAGACTGAGC−1
clust_4_TACCCACTCACTGCTC−1
clust_1_CTACGGGAGCGTGAAC−1
clust_1_TTGAACGCATTCTCCG−1
clust_3_TATCAGGCACGGGCTT−1
clust_3_CTCCATGAGTTACGTC−1
clust_1_TTCATGTAGCACTCCG−1
clust_8_CCGGACAGTAACCAGG−1
clust_3_TAAGCGTGTAAGGCCA−1
clust_8_CTGAATGTCCCAGGCA−1
clust_8_GCAGCCAAGAACCAAC−1
clust_10_TCGATTTCCGCTGTT−1
clust_10_GAGTTTGTCTTAGTTC−1
clust_6_CCTACGTGTCCTACAA−1
clust_6_CCTCTAGCAGGCTTGC−1
clust_6_TCCTCTTTCGGAATTC−1
clust_6_CCGGACATCTGTCGCT−1
clust_10_TTTACCACAATAAGGT−1
clust_6_TGATGGTCACTAGGCC−1
clust_10_GAACTTTCACACTTAG−1
clust_10_GGGTAGACAGATTTCG−1
clust_7_CCACAAACACCAAATC−1
clust_9_CTGAGCGAGTGAATAC−1
clust_1_AAGATAGCAAATCGGG−1
clust_1_GGTAACTAGTGCACAG−1
clust_7_AGATGAAAGTGTACCT−1
clust_7_TTATTGCCAAGTTTGC−1
clust_2_CTGAGGCGTAGCACAG−1
clust_9_ACGTTCCGTCGGGCAT−1
clust_5_GGAACCCGTGGTGATG−1
clust_5_TTAGGCACATAATCGC−1
clust_10_CCTATCGCATCGTGCG−1
clust_6_GTTACAGCAGATACCT−1
clust_6_TTACCGCGTACAAGTA−1
clust_6_AATGCCACATGTCTAG−1
clust_10_CATCCGTCATTCATCT−1
clust_10_GACCGTGTCCTGCTAC−1
clust_3_ACAACCAAGAGAACCC−1
clust_11_AAGCATCCAATTGCTG−1
clust_11_CACGTGGTCGTCCTCA−1
clust_11_GACCAATGTTTCGTGA−1
clust_11_GATGAGGGTAGGCAGT−1
clust_11_CACCAAATCGCAATGT−1
clust_11_TAGGTTGAGCGATTCT−1
clust_3_TGCTCCAAGAACGCGT−1
clust_4_ACGTACAAGAATCTAG−1
clust_4_CTAACCCTCTCGTGAA−1
clust_4_GTGGAGAGTTTCGTGA−1
clust_4_TGGGATTGTGTCCGGT−1
clust_3_AGGACTTGTACTGGGA−1
clust_11_CCCAACTAGGTCACTT−1
clust_11_TGTGTGATCGGAGTGA−1
clust_5_CCACACTCAACTTGCA−1
clust_5_AACGTCACAAGTATCC−1
clust_6_CAAGACTCAACCTATG−1
clust_7_TCACGGGAGAATTGTG−1
clust_7_AGACCCGAGTTTGCTG−1
clust_7_GTCAGCGTCGGACCAC−1
clust_10_CTACATTCAAACAGGC−1
clust_10_GCCAGCAGTTGCATTG−1
clust_7_AGTGTTGTCTATACGG−1
clust_7_CTCATTAGTCAGACGA−1
clust_7_CTGGCAGGTACAGTAA−1
clust_7_TGGATGTGTGACTAAA−1

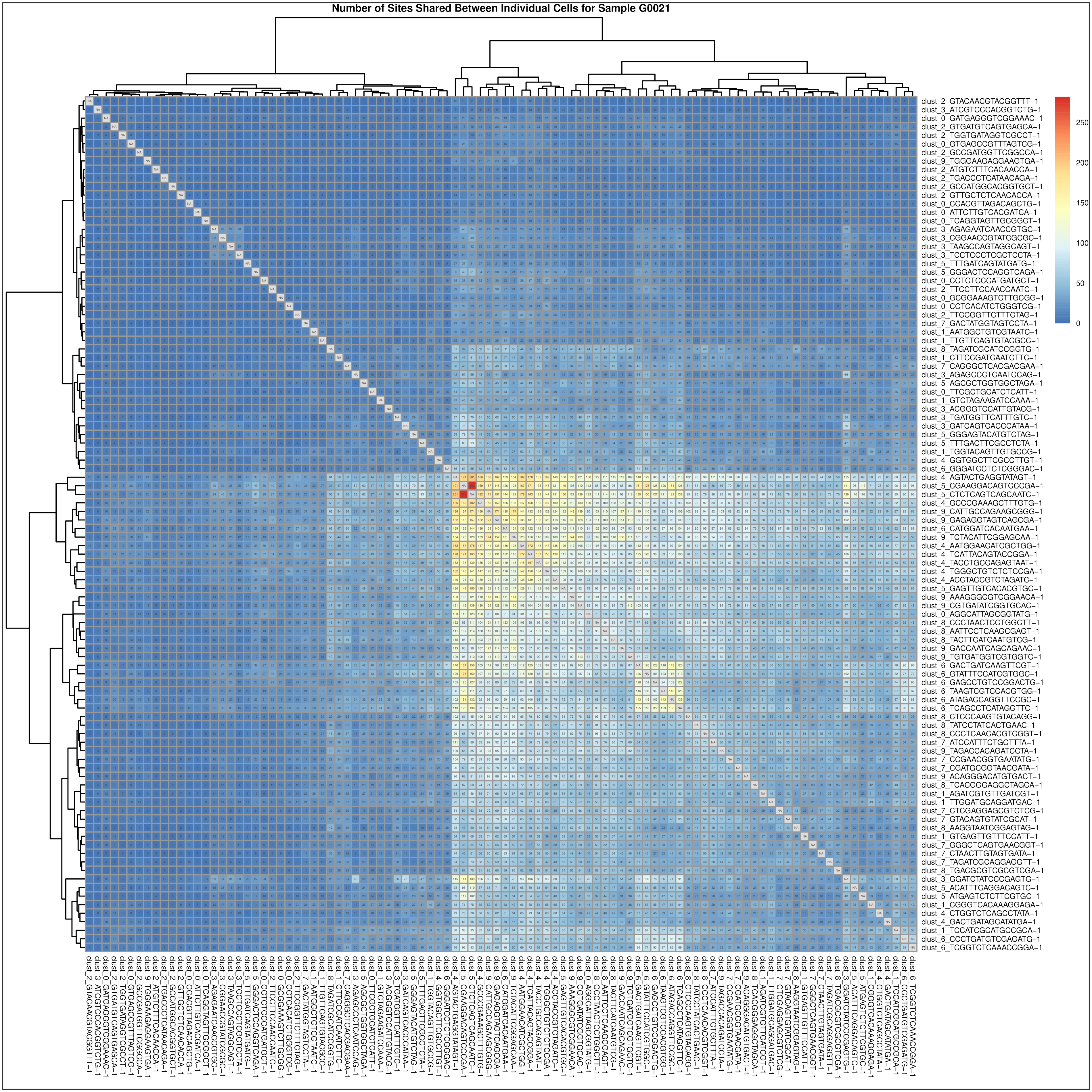
Number of Sites Shared Between Individual Cells for Sample G0021
250
200
150
100
50
0
clust_2_GTACAACGTACGGTTT−1
clust_3_ATCGTCCCACGGTCTG−1
clust_0_GATGAGGGTCGGAAAC−1
clust_2_GTGATGTCAGTGAGCA−1
clust_2_TGGTGATAGGTCGCCT−1
clust_0_GTGAGCCGTTTAGTCG−1
clust_2_GCCGATGGTTCGGCCA−1
clust_9_TGGGAAGAGGAAGTGA−1
clust_2_ATGTCTTTCACAACCA−1
clust_2_TGACCCTCATAACAGA−1
clust_2_GCCATGGCACGGTGCT−1
clust_2_GTTGCTCTCAACACCA−1
clust_0_CCACGTTAGACAGCTG−1
clust_0_ATTCTTGTCACGATCA−1
clust_0_TCAGGTAGTTGCGGCT−1
clust_3_AGAGAATCAACCGTGC−1
clust_3_CGGAACCGTATCGCGC−1
clust_3_TAAGCCAGTAGGCAGT−1
clust_3_TCCTCCCTCGCTCCTA−1
clust_5_TTTGATCAGTATGATG−1
clust_5_GGGACTCCAGGTCAGA−1
clust_0_CCTCTCCCATGATGCT−1
clust_2_TTCCTTCCAACCAATC−1
clust_0_GCGGAAAGTCTTGCGG−1
clust_0_CCTCACATCTGGGTCG−1
clust_2_TTCCGGTTCTTTCTAG−1
clust_7_GACTATGGTAGTCCTA−1
clust_1_AATGGCTGTCGTAATC−1
clust_1_TTGTTCAGTGTACGCC−1
clust_8_TAGATCGCATCCGGTG−1
clust_1_CTTCCGATCAATCTTC−1
clust_7_CAGGGCTCACGACGAA−1
clust_3_AGAGCCCTCAATCCAG−1
clust_5_AGCGCTGGTGGCTAGA−1
clust_0_TTCGCTGCATCTCATT−1
clust_1_GTCTAGAAGATCCAAA−1
clust_3_ACGGGTCCATTGTACG−1
clust_3_TGATGGTTCATTTGTC−1
clust_3_GATCAGTCACCCATAA−1
clust_5_GGGAGTACATGTCTAG−1
clust_5_TTTGACTTCGCCTCTA−1
clust_1_TGGTACAGTTGTGCCG−1
clust_4_GGTGGCTTCGCCTTGT−1
clust_6_GGGATCCTCTCGGGAC−1
clust_4_AGTACTGAGGTATAGT−1
clust_5_CGAAGGACAGTCCCGA−1
clust_5_CTCTCAGTCAGCAATC−1
clust_4_GCCCGAAAGCTTTGTG−1
clust_9_CATTGCCAGAAGCGGG−1
clust_9_GAGAGGTAGTCAGCGA−1
clust_6_CATGGATCACAATGAA−1
clust_9_TCTACATTCGGAGCAA−1
clust_4_AATGGAACATCGCTGG−1
clust_4_TCATTACAGTACCGGA−1
clust_4_TACCTGCCAGAGTAAT−1
clust_4_TGGGCTGTCTCTCCGA−1
clust_4_ACCTACCGTCTAGATC−1
clust_5_GAGTTGTCACACGTGC−1
clust_9_AAAGGGCGTCGGAACA−1
clust_9_CGTGATATCGGTGCAC−1
clust_0_AGGCATTAGCGGTATG−1
clust_8_CCCTAACTCCTGGCTT−1
clust_8_AATTCCTCAAGCGAGT−1
clust_8_TACTTCATCAATGTCG−1
clust_9_GACCAATCAGCAGAAC−1
clust_9_TGTGATGGTCGTGGTC−1
clust_6_GACTGATCAAGTTCGT−1
clust_6_GTATTTCCATCGTGGC−1
clust_6_GAGCCTGTCCGGACTG−1
clust_6_TAAGTCGTCCACGTGG−1
clust_6_ATAGACCAGGTTCCGC−1
clust_6_TCAGCCTCATAGGTTC−1
clust_8_CTCCCAAGTGTACAGG−1
clust_8_TATCCTATCACTGAAC−1
clust_8_CCCTCAACACGTCGGT−1
clust_7_ATCCATTTCTGCTTTA−1
clust_9_TAGACCACAGATCCTA−1
clust_7_CCGAACGGTGAATATG−1
clust_7_CGATGCGGTAACGATA−1
clust_9_ACAGGGACATGTGACT−1
clust_8_TCACGGGAGGCTAGCA−1
clust_1_AGATCGTGTTGATCGT−1
clust_1_TTGGATGCAGGATGAC−1
clust_7_CTCGAGGAGCGTCTCG−1
clust_7_GTACAGTGTATCGCAT−1
clust_8_AAGGTAATCGGAGTAG−1
clust_1_GTGAGTTGTTTCCATT−1
clust_7_GGGCTCAGTGAACGGT−1
clust_7_CTAACTTGTAGTGATA−1
clust_7_TAGATCGCAGGAGGTT−1
clust_8_TGACGCGTCGCGTCGA−1
clust_3_GGATCTATCCCGAGTG−1
clust_5_ACATTTCAGGACAGTC−1
clust_5_ATGAGTCTCTTCGTGC−1
clust_1_CGGGTCACAAAGGAGA−1
clust_4_CTGGTCTCAGCCTATA−1
clust_4_GACTGATAGCATATGA−1
clust_1_TCCATCGCATGCCGCA−1
clust_6_CCCTGATGTCGAGATG−1
clust_6_TCGGTCTCAAACCGGA−1

#### Number of Sites Shared Between Individual Cells for Sample G0022

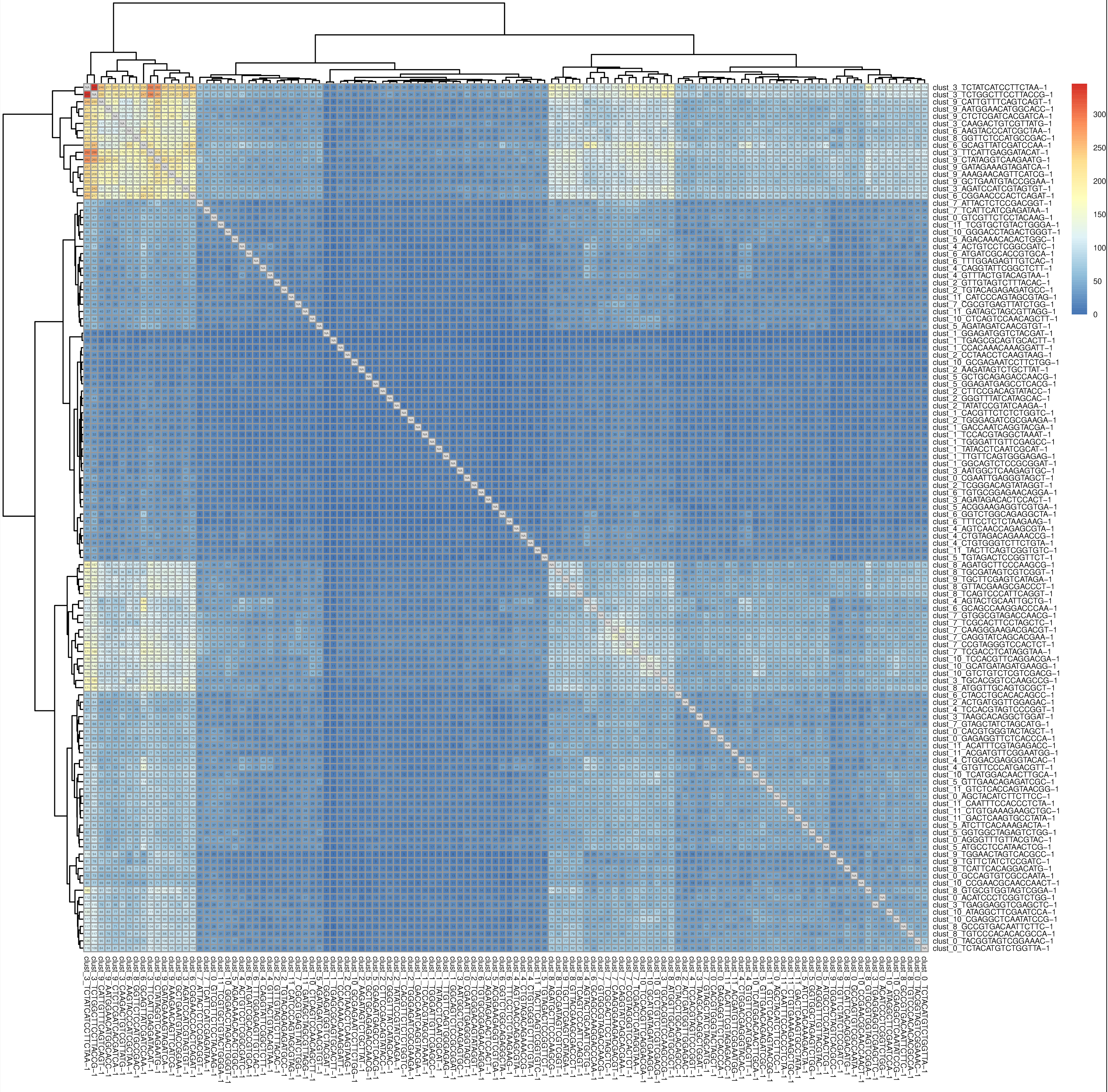

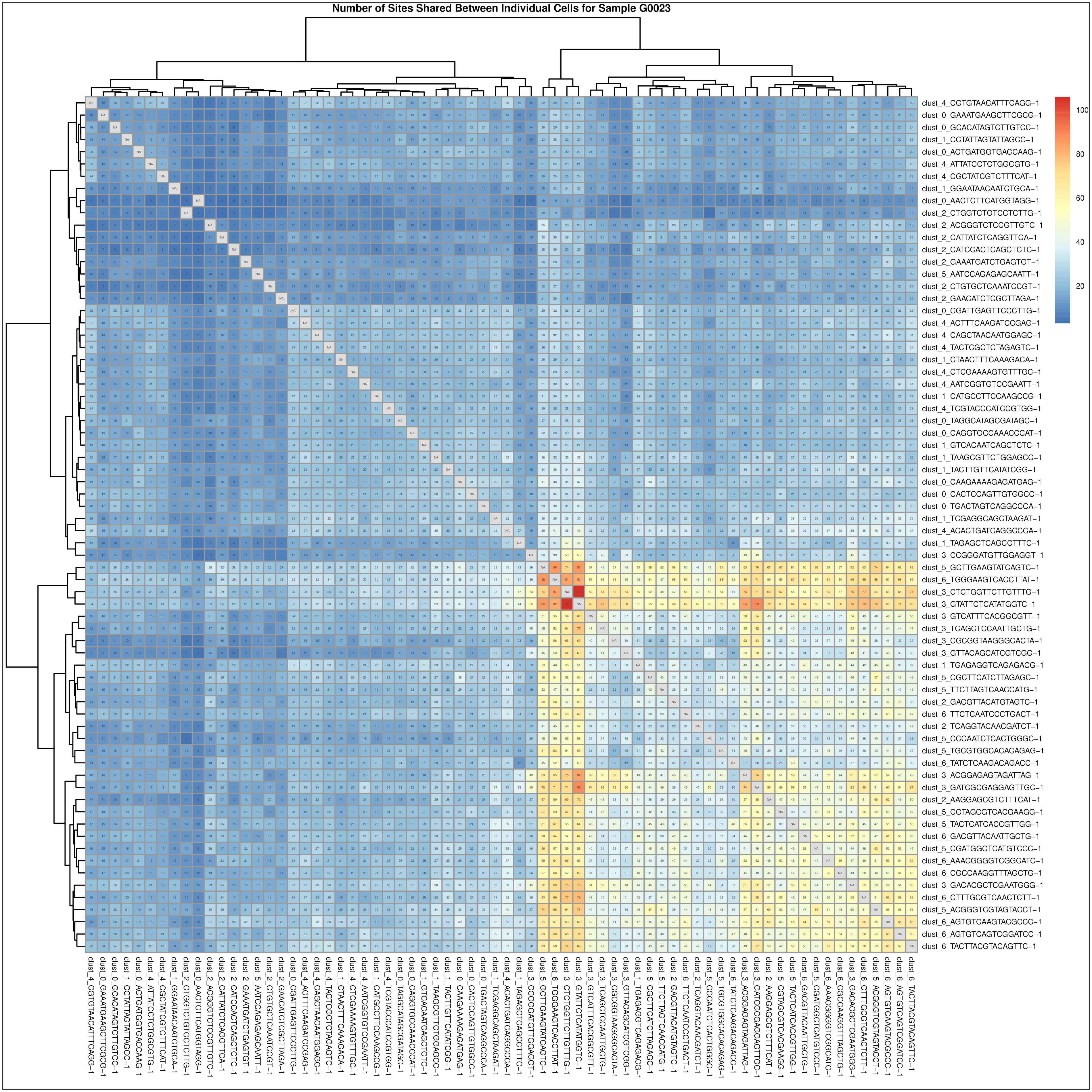
Number of Sites Shared Between Individual Cells for Sample G0023
100
80
60
40
20
clust_4_CGTGTAACATTTCAGG-1
clust_0_GAAATGAAGCTTCGCG-1
clust_0_GCACATAGTCTTGTCC-1
clust_1_CCTATTAGTATTAGCC-1
clust_0_ACTGATGGTGACCAAG-1
clust_4_ATTATCCTCTGGCGTG-1
clust_4_CGCTATCGTCTTTCAT-1
clust_1_GGAATAACAATCTGCA-1
clust_0_AACTCTTCATGGTAGG-1
clust_2_CTGGTCTGTCCTCTTG-1
clust_2_ACGGGTCTCCGTTGTC-1
clust_2_CATTATCTCAGGTTCA-1
clust_2_CATCCACTCAGCTCTC-1
clust_2_GAAATGATCTGAGTGT-1
clust_5_AATCCAGAGAGCAATT-1
clust_2_CTGTGCTCAAATCCGT-1
clust_2_GAACATCTCGCTTAGA-1
clust_0_CGATTGAGTTCCCTTG-1
clust_4_ACTTTCAAGATCCGAG-1
clust_4_CAGCTAACAATGGAGC-1
clust_4_TACTCGCTCTAGAGTC-1
clust_1_CTAACTTTCAAAGACA-1
clust_4_CTCGAAAAGTGTTTGC-1
clust_4_AATCGGTGTCCGAATT-1
clust_1_CATGCCTTCCAAGCCG-1
clust_4_TCGTACCCATCCGTGG-1
clust_0_TAGGCATAGCGATAGC-1
clust_0_CAGGTGCCAAACCCAT-1
clust_1_GTCACAATCAGCTCTC-1
clust_1_TAAGCGTTCTGGAGCC-1
clust_1_TACTTGTTCATATCGG-1
clust_0_CAAGAAAAGAGATGAG-1
clust_0_CACTCCAGTTGTGGCC-1
clust_0_TGACTAGTCAGGCCCA-1
clust_1_TCGAGGCAGCTAAGAT-1
clust_4_ACACTGATCAGGCCCA-1
clust_1_TAGAGCTCAGCCTTTC-1
clust_3_CCGGGATGTTGGAGGT-1
clust_5_GCTTGAAGTATCAGTC-1
clust_6_TGGGAAGTCACCTTAT-1
clust_3_CTCTGGTTCTTGTTTG-1
clust_3_GTATTCTCATATGGTC-1
clust_3_GTCATTTCACGGCGTT-1
clust_3_TCAGCTCCAATTGCTG-1
clust_3_CGCGGTAAGGGCACTA-1
clust_3_GTTACAGCATCGTCGG-1
clust_1_TGAGAGGTCAGAGACG-1
clust_5_CGCTTCATCTTAGAGC-1
clust_5_TTCTTAGTCAACCATG-1
clust_2_GACGTTACATGTAGTC-1
clust_6_TTCTCAATCCCTGACT-1
clust_2_TCAGGTACAACGATCT-1
clust_5_CCCAATCTCACTGGGC-1
clust_5_TGCGTGGCACACAGAG-1
clust_6_TATCTCAAGACAGACC-1
clust_3_ACGGAGAGTAGATTAG-1
clust_3_GATCGCGAGGAGTTGC-1
clust_2_AAGGAGCGTCTTTCAT-1
clust_5_CGTAGCGTCACGAAGG-1
clust_5_TACTCATCACCGTTGG-1
clust_6_GACGTTACAATTGCTG-1
clust_5_CGATGGCTCATGTCCC-1
clust_6_AAACGGGGTCGGCATC-1
clust_6_CGCCAAGGTTTAGCTG-1
clust_3_GACACGCTCGAATGGG-1
clust_6_CTTTGCGTCAACTCTT-1
clust_5_ACGGGTCGTAGTACCT-1
clust_6_AGTGTCAAGTACGCCC-1
clust_6_AGTGTCAGTCGGATCC-1
clust_6_TACTTACGTACAGTTC-1

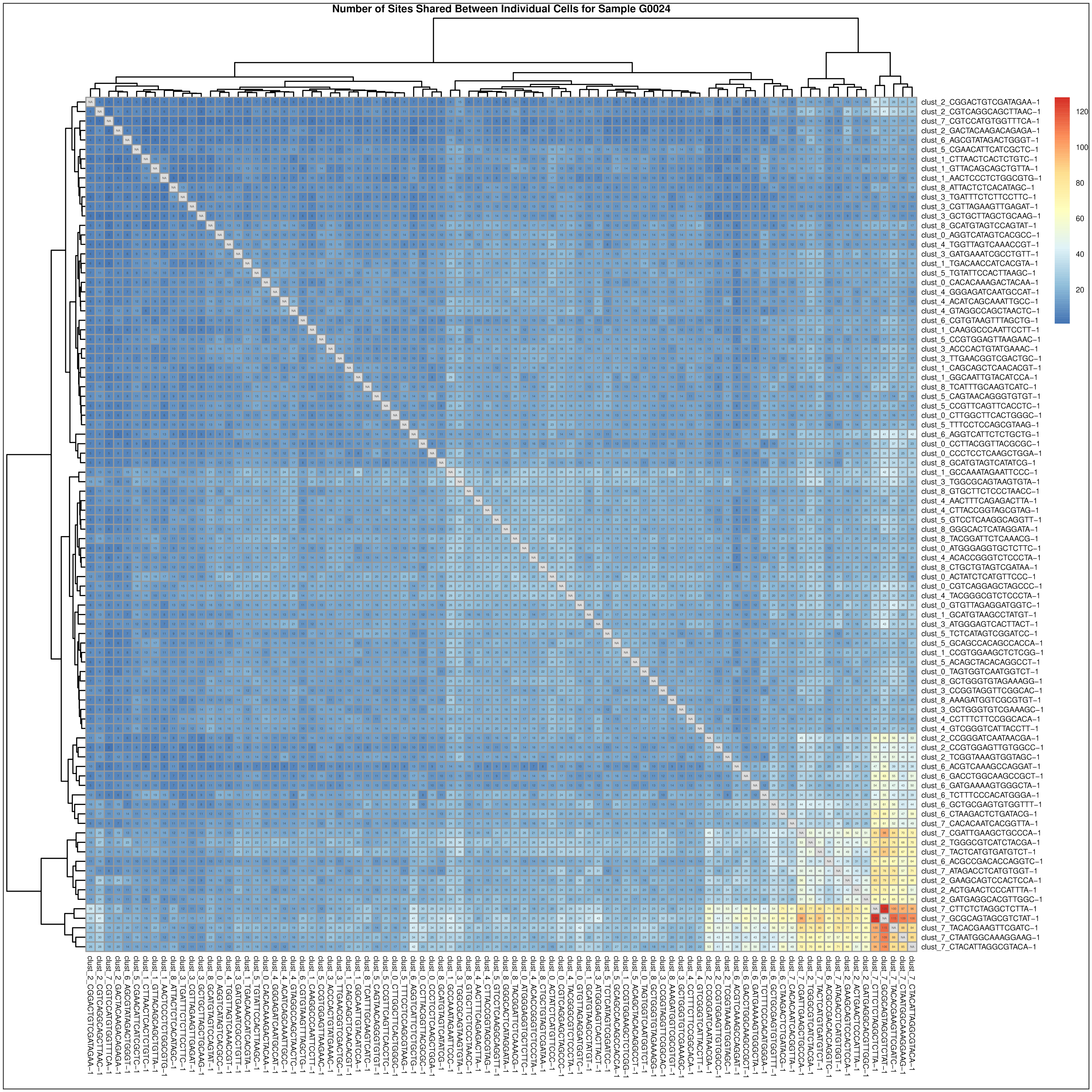

Number of Sites Shared Between Individual Cells for Sample G0024
clust_2_CGGACTGTCGATAGAA−1
clust_2_CGTCAGGCAGCTTAAC−1
clust_7_CGTCCATGTGGTTTCA−1
clust_2_GACTACAAGACAGAGA−1
clust_6_AGCGTATAGACTGGGT−1
clust_5_CGAACATTCATCGCTC−1
clust_1_CTTAACTCACTCTGTC−1
clust_1_GTTACAGCAGCTGTTA−1
clust_1_AACTCCCTCTGGCGTG−1
clust_8_ATTACTCTCACATAGC−1
clust_3_TGATTTCTCTTCCTTC−1
clust_3_CGTTAGAAGTTGAGAT−1
clust_3_GCTGCTTAGCTGCAAG−1
clust_8_GCATGTAGTCCAGTAT−1
clust_0_AGGTCATAGTCACGCC−1
clust_4_TGGTTAGTCAAACCGT−1
clust_3_GATGAAATCGCCTGTT−1
clust_1_TGACAACCATCACGTA−1
clust_5_TGTATTCCACTTAAGC−1
clust_0_CACACAAAGACTACAA−1
clust_4_GGGAGATCAATGCCAT−1
clust_4_ACATCAGCAAATTGCC−1
clust_4_GTAGGCCAGCTAACTC−1
clust_6_CGTGTAAGTTTAGCTG−1
clust_1_CAAGGCCCAATTCCTT−1
clust_5_CCGTGGAGTTAAGAAC−1
clust_3_ACCCACTGTATGAAAC−1
clust_3_TTGAACGGTCGACTGC−1
clust_1_CAGCAGCTCAACACGT−1
clust_1_GGCAATTGTACATCCA−1
clust_8_TCATTTGCAAGTCATC−1
clust_5_CAGTAACAGGGTGTGT−1
clust_5_CCGTTCAGTTCACCTC−1
clust_0_CTTGGCTTCACTGGGC−1
clust_5_TTTCCTCCAGCGTAAG−1
clust_6_AGGTCATTCTCTGCTG−1
clust_0_CCTTACGGTTACGCGC−1
clust_0_CCCTCCTCAAGCTGGA−1
clust_8_GCATGTAGTCATATCG−1
clust_1_GCCAAATAGAATTCCC−1
clust_3_TGGCGCAGTAAGTGTA−1
clust_8_GTGCTTCTCCCTAACC−1
clust_4_AACTTTCAGAGACTTA−1
clust_4_CTTACCGGTAGCGTAG−1
clust_5_GTCCTCAAGGCAGGTT−1
clust_8_GGGCACTCATAGGATA−1
clust_8_TACGGATTCTCAAACG−1
clust_0_ATGGGAGGTGCTCTTC−1
clust_4_ACACCGGGTCTCCCTA−1
clust_8_CTGCTGTAGTCGATAA−1
clust_0_ACTATCTCATGTTCCC−1
clust_0_CGTCAGGAGCTAGCCC−1
clust_4_TACGGGCGTCTCCCTA−1
clust_0_GTGTTAGAGGATGGTC−1
clust_1_GCATGTAAGCCTATGT−1
clust_3_ATGGGAGTCACTTACT−1
clust_5_TCTCATAGTCGGATCC−1
clust_5_GCAGCCACAGCCACCA−1
clust_1_CCGTGGAAGCTCTCGG−1
clust_5_ACAGCTACACAGGCCT−1
clust_0_TAGTGGTCAATGGTCT−1
clust_8_GCTGGGTGTAGAAAGG−1
clust_3_CCGGTAGGTTCGGCAC−1
clust_8_AAAGATGGTCGCGTGT−1
clust_3_GCTGGGTGTCGAAAGC−1
clust_4_CCTTTCTTCCGGCACA−1
clust_4_GTCGGGTCATTACCTT−1
clust_2_CCGGGATCAATAACGA−1
clust_2_CCGTGGAGTTGTGGCC−1
clust_2_TCGGTAAAGTGGTAGC−1
clust_6_ACGTCAAAGCCAGGAT−1
clust_6_GACCTGGCAAGCCGCT−1
clust_6_GATGAAAAGTGGGCTA−1
clust_6_TCTTTCCCACATGGGA−1
clust_6_GCTGCGAGTGTGGTTT−1
clust_6_CTAAGACTCTGATACG−1
clust_7_CACACAATCACGGTTA−1
clust_7_CGATTGAAGCTGCCCA−1
clust_2_TGGGCGTCATCTACGA−1
clust_7_TACTCATGTGATGTCT−1
clust_6_ACGCCGACACCAGGTC−1
clust_7_ATAGACCTCATGTGGT−1
clust_2_GAAGCAGTCCACTCCA−1
clust_2_ACTGAACTCCCATTTA−1
clust_2_GATGAGGCACGTTGGC−1
clust_7_CTTCTCTAGGCTCTTA−1
clust_7_GCGCAGTAGCGTCTAT−1
clust_7_TACACGAAGTTCGATC−1
clust_7_CTAATGGCAAAGGAAG−1
clust_7_CTACATTAGGCGTACA−1
120
100
80
60
40
20

**Number of Sites Shared Between Individual Cells for Sample G0025**

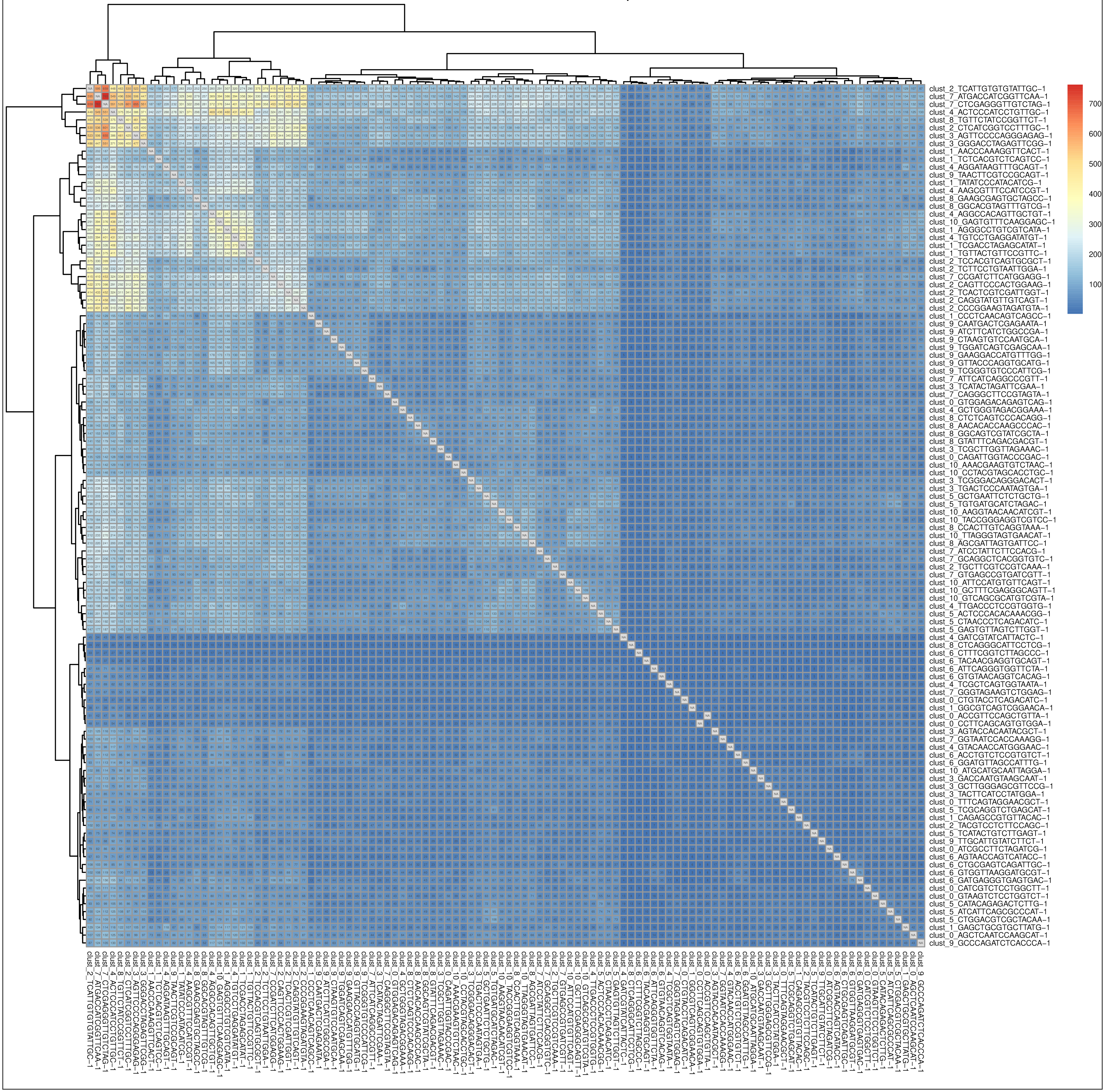

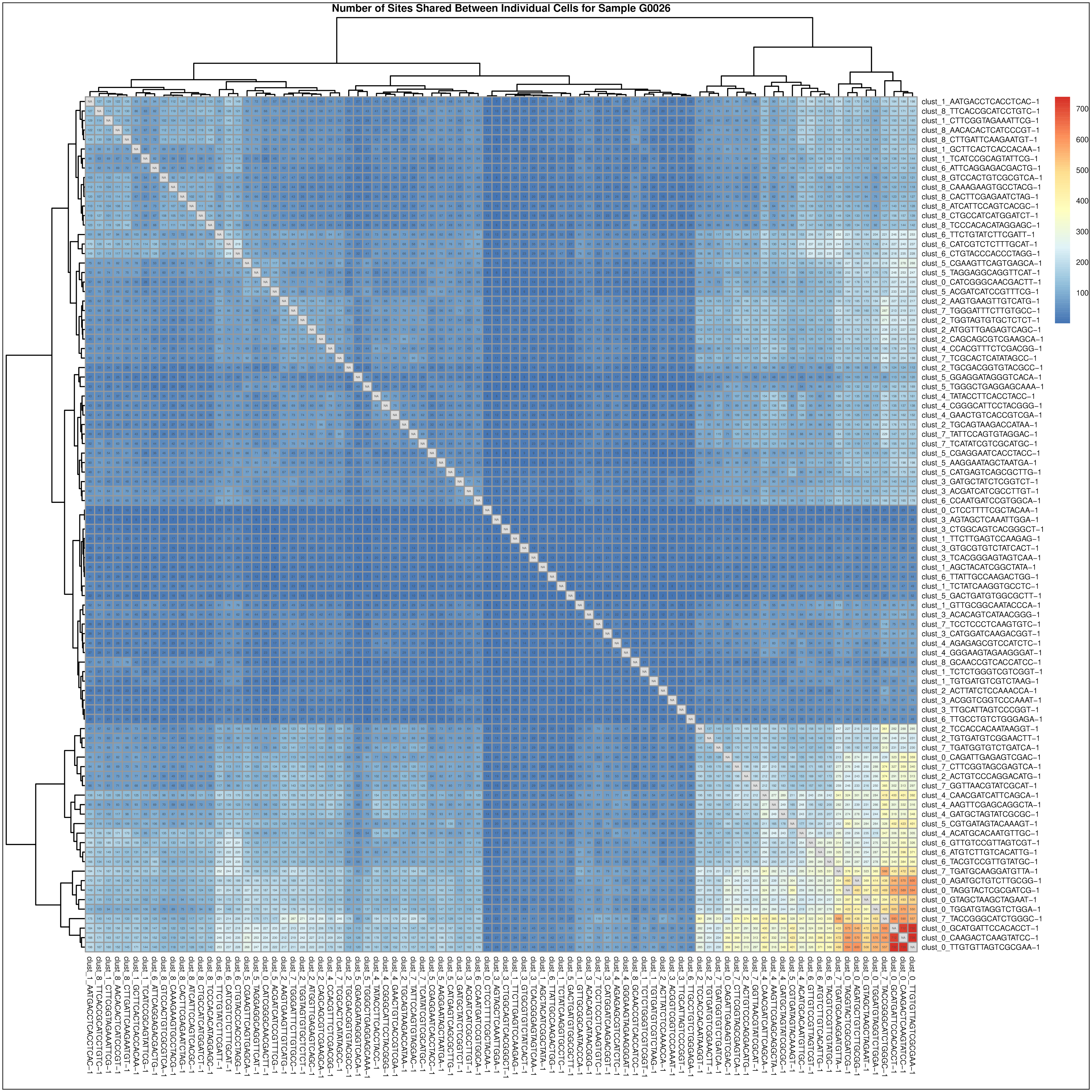

Number of Sites Shared Between Individual Cells for Sample G0026
700
600
500
400
300
200
100
clust_1_AATGACCTCACCTCAC-1
clust_8_TTCACCGCATCCTGTC-1
clust_1_CTTCGGTAGAAATTCG-1
clust_8_AACACACTCATCCCGT-1
clust_8_CTTGATTCAAGAATGT-1
clust_1_GCTTCACTCACCACAA-1
clust_1_TCATCCGCAGTATTCG-1
clust_6_ATTCAGGAGACGACTG-1
clust_8_GTCCACTGTCGCGTCA-1
clust_8_CAAAGAAGTGCCTACG-1
clust_8_CACTTCGAGAATCTAG-1
clust_8_ATCATTCCAGTCACGC-1
clust_8_CTGCCATCATGGATCT-1
clust_8_TCCCACACATAGGAGC-1
clust_6_TTCTGTATCTTCGATT-1
clust_6_CATCGTCTCTTTGCAT-1
clust_6_CTGTACCCACCCTAGG-1
clust_5_CGAAGTTCAGTGAGCA-1
clust_5_TAGGAGGCAGGTTCAT-1
clust_0_CATCGGGCAACGACTT-1
clust_5_ACGATCATCCGTTTCG-1
clust_2_AAGTGAAGTTGTCATG-1
clust_7_TGGGATTTCTTGTGCC-1
clust_2_TGGTAGTGTGCTCTCT-1
clust_2_ATGGTTGAGAGTCAGC-1
clust_2_CAGCAGCGTCGAAGCA-1
clust_4_CCACGTTTCTCGACGG-1
clust_7_TCGCACTCATATAGCC-1
clust_2_TGCGACGGTGTACGCC-1
clust_5_GGAGGATAGGGTCACA-1
clust_5_TGGGCTGAGGAGCAAA-1
clust_4_TATACCTTCACCTACC-1
clust_4_CGGGCATTCCTACGGG-1
clust_4_GAACTGTCACCGTCGA-1
clust_2_TGCAGTAAGACCATAA-1
clust_7_TATTCCAGTGTAGGAC-1
clust_7_TCATATCGTCGCATGC-1
clust_5_CGAGGAATCACCTACC-1
clust_5_AAGGAATAGCTAATGA-1
clust_5_CATGAGTCAGCGCTTG-1
clust_3_GATGCTATCTCGGTCT-1
clust_3_ACGATCATCGCCTTGT-1
clust_6_CCAATGATCCGTGGCA-1
clust_0_CTCCTTTTCGCTACAA-1
clust_3_AGTAGCTCAAATTGGA-1
clust_3_CTGGCAGTCACGGGCT-1
clust_1_TTCTTGAGTCCAAGAG-1
clust_3_GTGCGTGTCTATCACT-1
clust_3_TCACGGGAGTAGTCAA-1
clust_1_AGCTACATCGGCTATA-1
clust_6_TTATTGCCAAGACTGG-1
clust_1_TCTATCAAGGTGCCTC-1
clust_5_GACTGATGTGGCGCTT-1
clust_1_GTTGCGGCAATACCCA-1
clust_3_ACACAGTCATAACGGG-1
clust_7_TCCTCCCTCAAGTGTC-1
clust_3_CATGGATCAAGACGGT-1
clust_4_AGAGAGCGTCCATCTC-1
clust_4_GGGAAGTAGAAGGGAT-1
clust_8_GCAACCGTCACCATCC-1
clust_1_TCTCTGGGTCGTCGGT-1
clust_1_TGTGATGTCGTCTAAG-1
clust_2_ACTTATCTCCAAACCA-1
clust_3_ACGGTCGGTCCCAAAT-1
clust_3_TTGCATTAGTCCCGGT-1
clust_6_TTGCCTGTCTGGGAGA-1
clust_2_TCCACCACAATAAGGT-1
clust_2_TGTGATGTCGGAACTT-1
clust_7_TGATGGTGTCTGATCA-1
clust_0_CAGATTGAGAGTCGAC-1
clust_7_CTTCGGTAGCGAGTCA-1
clust_2_ACTGTCCCAGGACATG-1
clust_7_GGTTAACGTATCGCAT-1
clust_4_CAACGATCATTCAGCA-1
clust_4_AAGTTCGAGCAGGCTA-1
clust_4_GATGCTAGTATCGCGC-1
clust_5_CGTGATAGTACAAAGT-1
clust_4_ACATGCACAATGTTGC-1
clust_6_GTTGTCCGTTAGTCGT-1
clust_6_ATGTCTTGTCACATTG-1
clust_6_TACGTCCGTTGTATGC-1
clust_7_TGATGCAAGGATGTTA-1
clust_0_AGATGCTGTCTTGCGG-1
clust_0_TAGGTACTCGCGATCG-1
clust_0_GTAGCTAAGCTAGAAT-1
clust_0_TGGATGTAGGTCTGGA-1
clust_7_TACCGGGCATCTGGGC-1
clust_0_GCATGATTCCACACCT-1
clust_0_CAAGACTCAAGTATCC-1
clust_0_TTGTGTTAGTCGCGAA-1

Number of Sites Shared Between Individual Cells for Sample G0027

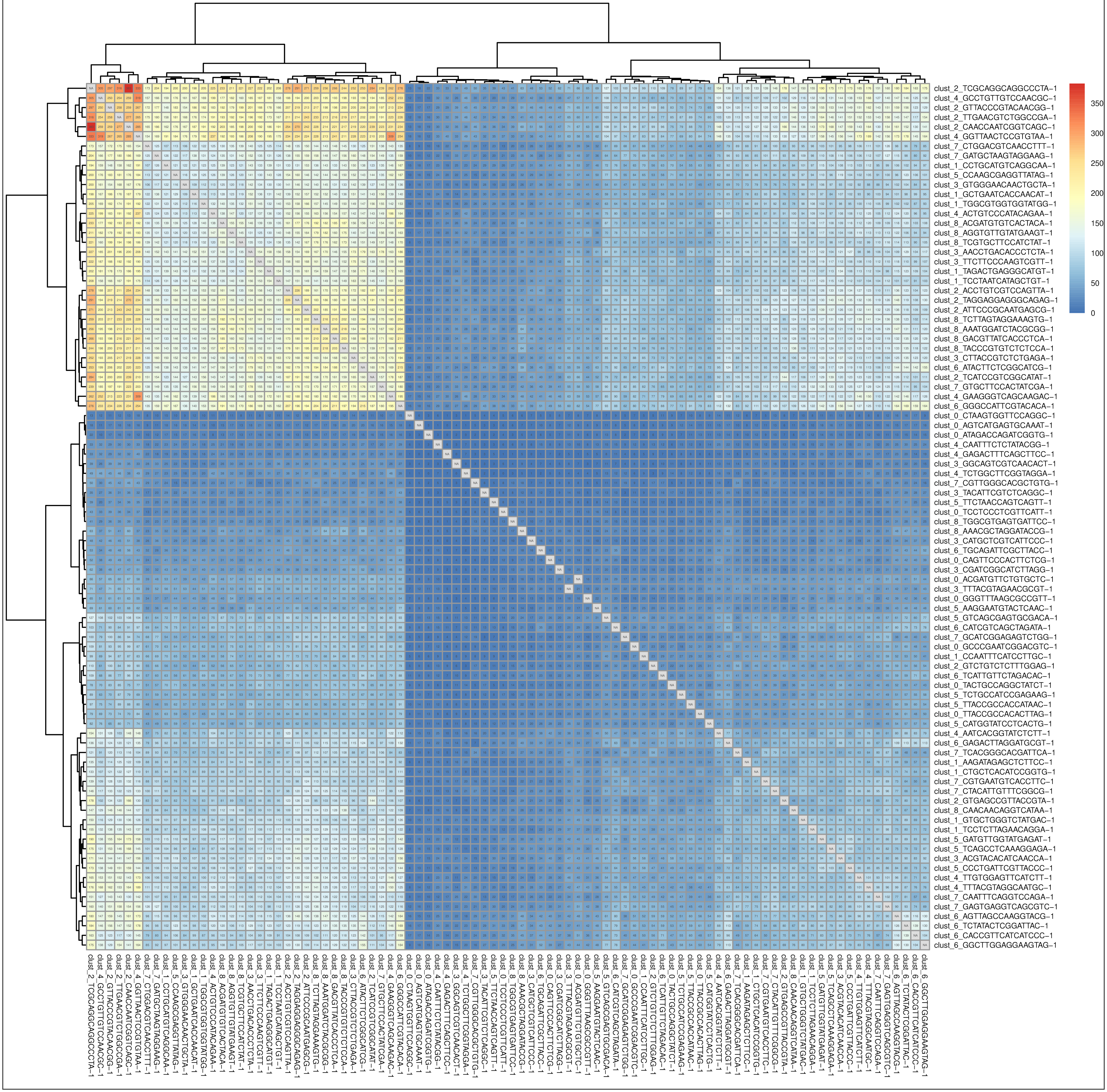

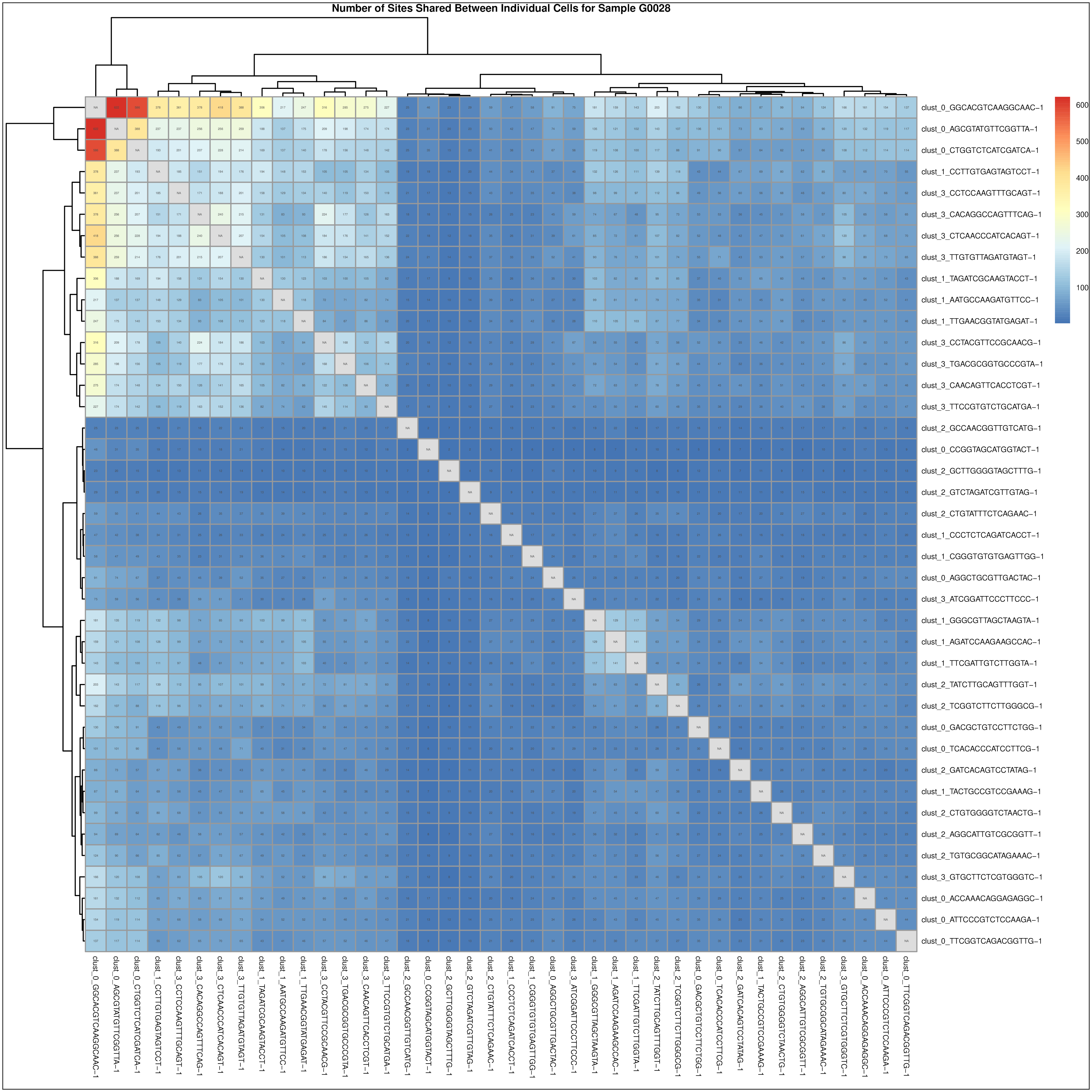
Number of Sites Shared Between Individual Cells for Sample G0028
600
500
400
300
200
100
clust_0_GGCACGTCAAGGCAAC-1
clust_0_AGCGTATGTTCGGTTA-1
clust_0_CTGGTCTCATCGATCA-1
clust_1_CCTTGTGAGTAGTCCT-1
clust_3_CCTCCAAGTTTGCAGT-1
clust_3_CACAGGCCAGTTTCAG-1
clust_3_CTCAACCCATCACAGT-1
clust_3_TTGTGTTAGATGTAGT-1
clust_1_TAGATCGCAAGTACCT-1
clust_1_AATGCCAAGATGTTCC-1
clust_1_TTGAACGGTATGAGAT-1
clust_3_CCTACGTTCCGCAACG-1
clust_3_TGACGCGGTGCCCGTA-1
clust_3_CAACAGTTCACCTCGT-1
clust_3_TTCCGTGTCTGCATGA-1
clust_2_GCCAACGGTTGTCATG-1
clust_0_CCGGTAGCATGGTACT-1
clust_2_GCTTGGGGTAGCTTTG-1
clust_2_GTCTAGATCGTTGTAG-1
clust_2_CTGTATTTCTCAGAAC-1
clust_1_CCCTCTCAGATCACCT-1
clust_1_CGGGTGTGTGAGTTGG-1
clust_0_AGGCTGCGTTGACTAC-1
clust_3_ATCGGATTCCCTTCCC-1
clust_1_GGGCGTTAGCTAAGTA-1
clust_1_AGATCCAAGAAGCCAC-1
clust_1_TTCGATTGTCTTGGTA-1
clust_2_TATCTTGCAGTTTGGT-1
clust_2_TCGGTCTTCTTGGGCG-1
clust_0_GACGCTGTCCTTCTGG-1
clust_0_TCACACCCATCCTTCG-1
clust_2_GATCACAGTCCTATAG-1
clust_1_TACTGCCGTCCGAAAG-1
clust_2_CTGTGGGGTCTAACTG-1
clust_2_AGGCATTGTCGCGGTT-1
clust_2_TGTGCGGCATAGAAAC-1
clust_3_GTGCTTCTCGTGGGTC-1
clust_0_ACCAAACAGGAGAGGC-1
clust_0_ATTCCCGTCTCCAAGA-1
clust_0_TTCGGTCAGACGGTTG-1

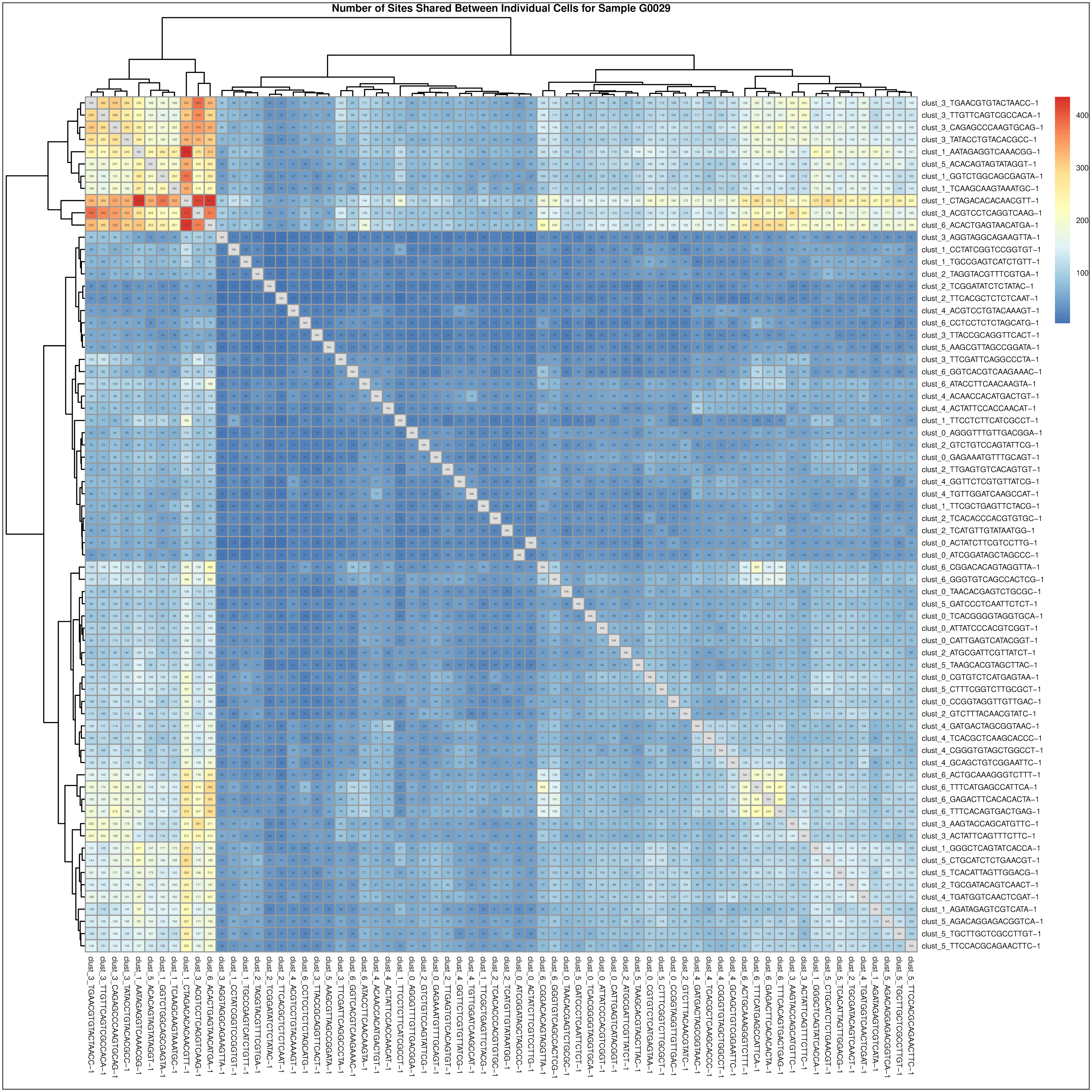
Number of Sites Shared Between Individual Cells for Sample G0029
400
300
200
100
clust_3_TGAACGTGTACTAACC−1
clust_3_TTGTTCAGTCGCCACA−1
clust_3_CAGAGCCCAAGTGCAG−1
clust_3_TATACCTGTACACGCC−1
clust_1_AATAGAGGTCAAACGG−1
clust_5_ACACAGTAGTATAGGT−1
clust_1_GGTCTGGCAGCGAGTA−1
clust_1_TCAAGCAAGTAAATGC−1
clust_1_CTAGACACACAACGTT−1
clust_3_ACGTCCTCAGGTCAAG−1
clust_6_ACACTGAGTAACATGA−1
clust_3_AGGTAGGCAGAAGTTA−1
clust_1_CCTATCGGTCCGGTGT−1
clust_1_TGCCGAGTCATCTGTT−1
clust_2_TAGGTACGTTTCGTGA−1
clust_2_TCGGATATCTCTATAC−1
clust_2_TTCACGCTCTCTCAAT−1
clust_4_ACGTCCTGTACAAAGT−1
clust_6_CCTCCTCTCTAGCATG−1
clust_3_TTACCGCAGGTTCACT−1
clust_5_AAGCGTTAGCCGGATA−1
clust_3_TTCGATTCAGGCCCTA−1
clust_6_GGTCACGTCAAGAAAC−1
clust_6_ATACCTTCAACAAGTA−1
clust_4_ACAACCACATGACTGT−1
clust_4_ACTATTCCACCAACAT−1
clust_1_TTCCTCTTCATCGCCT−1
clust_0_AGGGTTTGTTGACGGA−1
clust_2_GTCTGTCCAGTATTCG−1
clust_0_GAGAAATGTTTGCAGT−1
clust_2_TTGAGTGTCACAGTGT−1
clust_4_GGTTCTCGTGTTATCG−1
clust_4_TGTTGGATCAAGCCAT−1
clust_1_TTCGCTGAGTTCTACG−1
clust_2_TCACACCCACGTGTGC−1
clust_2_TCATGTTGTATAATGG−1
clust_0_ACTATCTTCGTCCTTG−1
clust_0_ATCGGATAGCTAGCCC−1
clust_6_CGGACACAGTAGGTTA−1
clust_6_GGGTGTCAGCCACTCG−1
clust_0_TAACACGAGTCTGCGC−1
clust_5_GATCCCTCAATTCTCT−1
clust_0_TCACGGGGTAGGTGCA−1
clust_0_ATTATCCCACGTCGGT−1
clust_0_CATTGAGTCATACGGT−1
clust_2_ATGCGATTCGTTATCT−1
clust_5_TAAGCACGTAGCTTAC−1
clust_0_CGTGTCTCATGAGTAA−1
clust_5_CTTTCGGTCTTGCGCT−1
clust_0_CCGGTAGGTTGTTGAC−1
clust_2_GTCTTTACAACGTATC−1
clust_4_GATGACTAGCGGTAAC−1
clust_4_TCACGCTCAAGCACCC−1
clust_4_CGGGTGTAGCTGGCCT−1
clust_4_GCAGCTGTCGGAATTC−1
clust_6_ACTGCAAAGGGTCTTT−1
clust_6_TTTCATGAGCCATTCA−1
clust_6_GAGACTTCACACACTA−1
clust_6_TTTCACAGTGACTGAG−1
clust_3_AAGTACCAGCATGTTC−1
clust_3_ACTATTCAGTTTCTTC−1
clust_1_GGGCTCAGTATCACCA−1
clust_5_CTGCATCTCTGAACGT−1
clust_5_TCACATTAGTTGGACG−1
clust_2_TGCGATACAGTCAACT−1
clust_4_TGATGGTCAACTCGAT−1
clust_1_AGATAGAGTCGTCATA−1
clust_5_AGACAGGAGACGGTCA−1
clust_5_TGCTTGCTCGCCTTGT−1
clust_5_TTCCACGCAGAACTTC−1

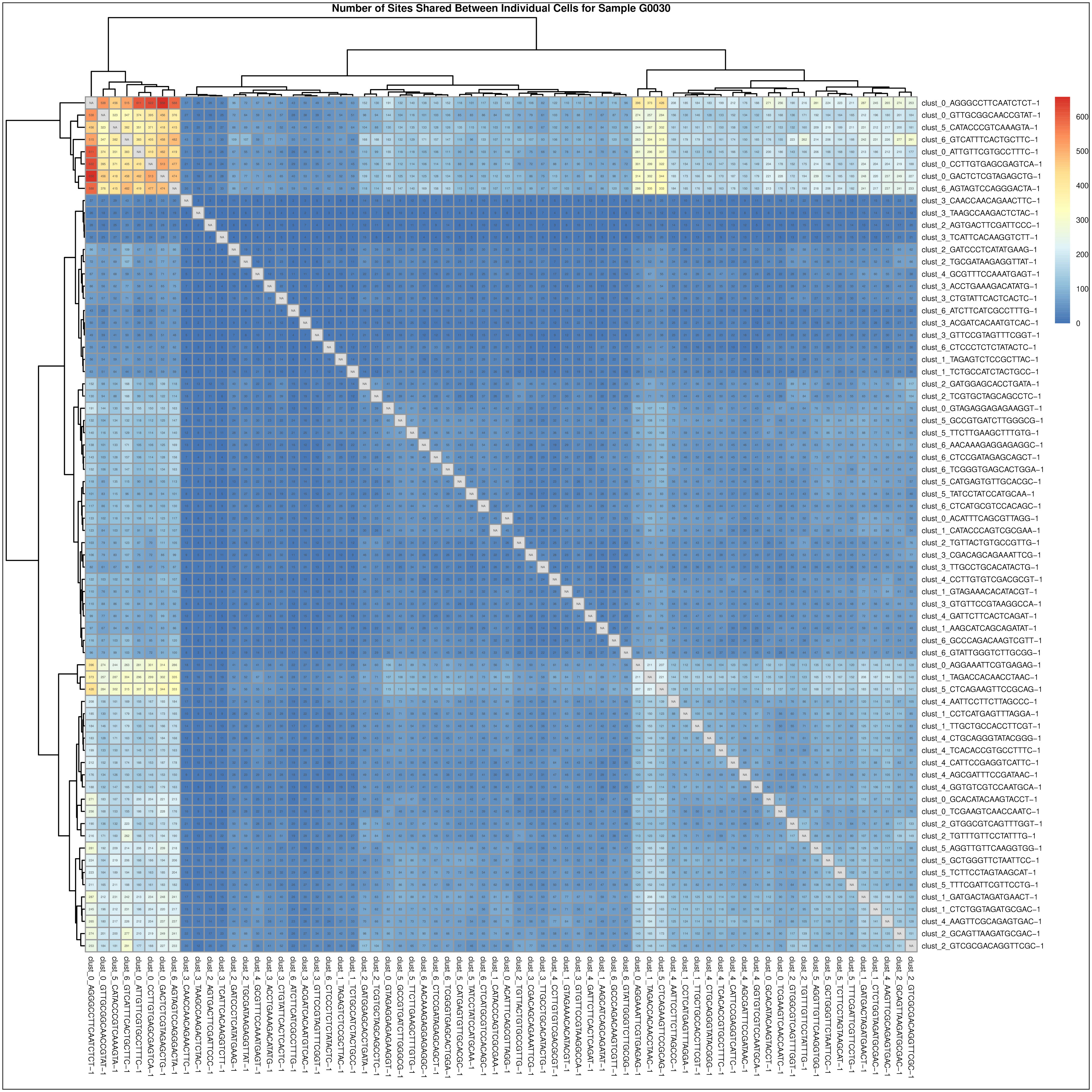
Number of Sites Shared Between Individual Cells for Sample G0030
600
500
400
300
200
100
0
clust_0_AGGGCCTTCAATCTCT-1
clust_0_GTTGCGGCAACCGTAT-1
clust_5_CATACCCGTCAAAGTA-1
clust_6_GTCATTTCACTGCTTC-1
clust_0_ATTGTTCGTGCCTTTC-1
clust_0_CCTTGTGAGCGAGTCA-1
clust_0_GACTCTCGTAGAGCTG-1
clust_6_AGTAGTCCAGGGACTA-1
clust_3_CAACCAACAGAACTTC-1
clust_3_TAAGCCAAGACTCTAC-1
clust_2_AGTGACTTCGATTCCC-1
clust_3_TCATTCACAAGGTCTT-1
clust_2_GATCCCTCATATGAAG-1
clust_2_TGCGATAAGAGGTTAT-1
clust_4_GCGTTTCCAAATGAGT-1
clust_3_ACCTGAAAGACATATG-1
clust_3_CTGTATTCACTCACTC-1
clust_6_ATCTTCATCGCCTTTG-1
clust_3_ACGATCACAATGTCAC-1
clust_3_GTTCCGTAGTTTCGGT-1
clust_6_CTCCCTCTCTATACTC-1
clust_1_TAGAGTCTCCGCTTAC-1
clust_1_TCTGCCATCTACTGCC-1
clust_2_GATGGAGCACCTGATA-1
clust_2_TCGTGCTAGCAGCCTC-1
clust_0_GTAGAGGAGAGAAGGT-1
clust_5_GCCGTGATCTTGGGCG-1
clust_5_TTCTTGAAGCTTTGTG-1
clust_6_AACAAAGAGGAGAGGC-1
clust_6_CTCCGATAGAGCAGCT-1
clust_6_TCGGGTGAGCACTGGA-1
clust_5_CATGAGTGTTGCACGC-1
clust_5_TATCCTATCCATGCAA-1
clust_6_CTCATGCGTCCACAGC-1
clust_0_ACATTTCAGCGTTAGG-1
clust_1_CATACCCAGTCGCGAA-1
clust_2_TGTTACTGTGCCGTTG-1
clust_3_CGACAGCAGAAATTCG-1
clust_3_TTGCCTGCACATACTG-1
clust_4_CCTTGTGTCGACGCGT-1
clust_1_GTAGAAACACATACGT-1
clust_3_GTGTTCCGTAAGGCCA-1
clust_4_GATTCTTCACTCAGAT-1
clust_1_AAGCATCAGCAGATAT-1
clust_6_GCCCAGACAAGTCGTT-1
clust_6_GTATTGGGTCTTGCGG-1
clust_0_AGGAAATTCGTGAGAG-1
clust_1_TAGACCACAACCTAAC-1
clust_5_CTCAGAAGTTCCGCAG-1
clust_4_AATTCCTTCTTAGCCC-1
clust_1_CCTCATGAGTTTAGGA-1
clust_1_TTGCTGCCACCTTCGT-1
clust_4_CTGCAGGGTATACGGG-1
clust_4_TCACACCGTGCCTTTC-1
clust_4_CATTCCGAGGTCATTC-1
clust_4_AGCGATTTCCGATAAC-1
clust_4_GGTGTCGTCCAATGCA-1
clust_0_GCACATACAAGTACCT-1
clust_0_TCGAAGTCAACCAATC-1
clust_2_GTGGCGTCAGTTTGGT-1
clust_2_TGTTTGTTCCTATTTG-1
clust_5_AGGTTGTTCAAGGTGG-1
clust_5_GCTGGGTTCTAATTCC-1
clust_5_TCTTCCTAGTAAGCAT-1
clust_5_TTTCGATTCGTTCCTG-1
clust_1_GATGACTAGATGAACT-1
clust_1_CTCTGGTAGATGCGAC-1
clust_4_AAGTTCGCAGAGTGAC-1
clust_2_GCAGTTAAGATGCGAC-1
clust_2_GTCGCGACAGGTTCGC-1

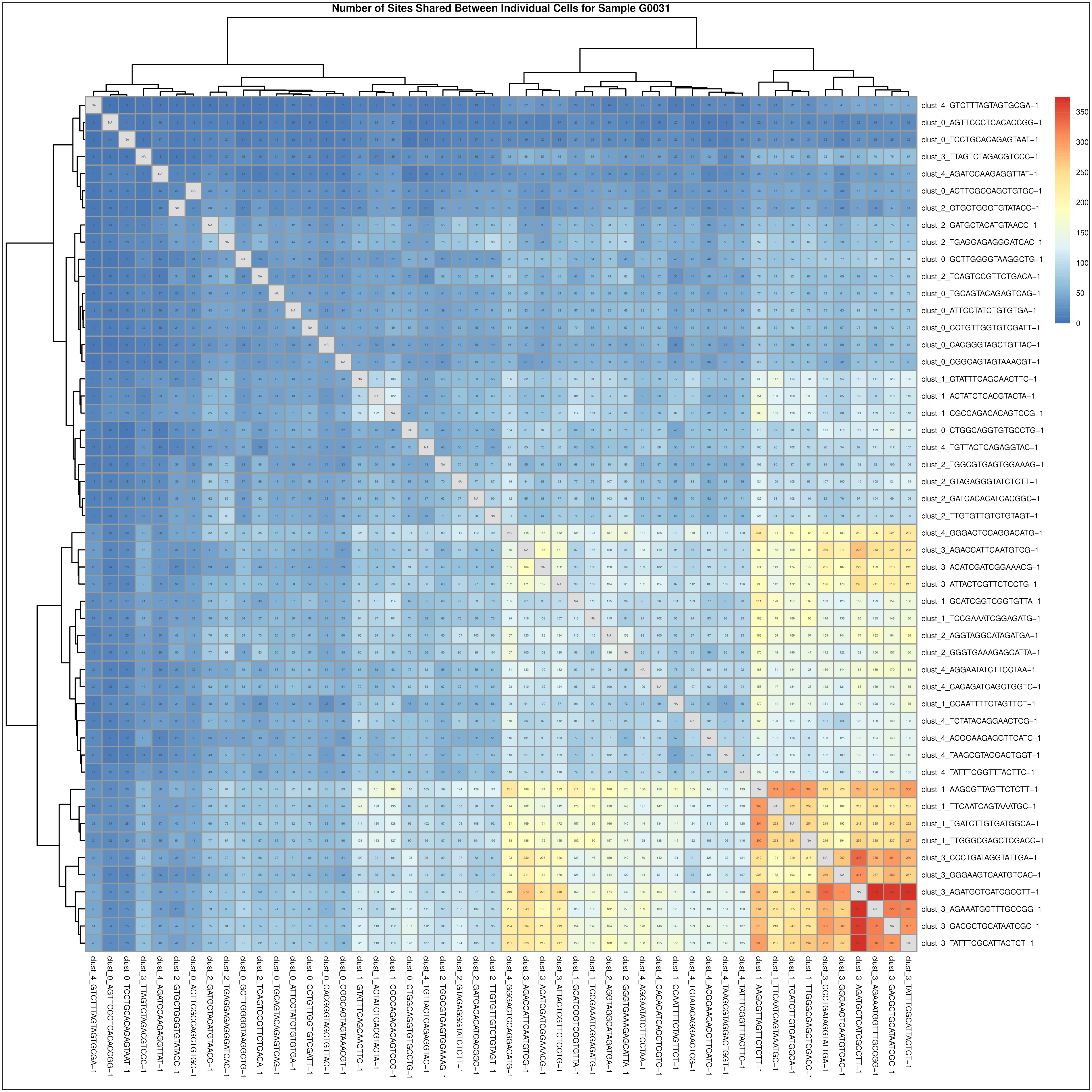
Number of Sites Shared Between Individual Cells for Sample G0031
clust_4_GTCTTTAGTAGTGCGA−1
clust_0_AGTTCCCTCACACCGG−1
clust_0_TCCTGCACAGAGTAAT−1
clust_3_TTAGTCTAGACGTCCC−1
clust_4_AGATCCAAGAGGTTAT−1
clust_0_ACTTCGCCAGCTGTGC−1
clust_2_GTGCTGGGTGTATACC−1
clust_2_GATGCTACATGTAACC−1
clust_2_TGAGGAGAGGGATCAC−1
clust_0_GCTTGGGGTAAGGCTG−1
clust_2_TCAGTCCGTTCTGACA−1
clust_0_TGCAGTACAGAGTCAG−1
clust_0_ATTCCTATCTGTGTGA−1
clust_0_CCTGTTGGTGTCGATT−1
clust_0_CACGGGTAGCTGTTAC−1
clust_0_CGGCAGTAGTAAACGT−1
clust_1_GTATTTCAGCAACTTC−1
clust_1_ACTATCTCACGTACTA−1
clust_1_CGCCAGACACAGTCCG−1
clust_0_CTGGCAGGTGTGCCTG−1
clust_4_TGTTACTCAGAGGTAC−1
clust_2_TGGCGTGAGTGGAAAG−1
clust_2_GTAGAGGGTATCTCTT−1
clust_2_GATCACACATCACGGC−1
clust_2_TTGTGTTGTCTGTAGT−1
clust_4_GGGACTCCAGGACATG−1
clust_3_AGACCATTCAATGTCG−1
clust_3_ACATCGATCGGAAACG−1
clust_3_ATTACTCGTTCTCCTG−1
clust_1_GCATCGGTCGGTGTTA−1
clust_1_TCCGAAATCGGAGATG−1
clust_2_AGGTAGGCATAGATGA−1
clust_2_GGGTGAAAGAGCATTA−1
clust_4_AGGAATATCTTCCTAA−1
clust_4_CACAGATCAGCTGGTC−1
clust_1_CCAATTTTCTAGTTCT−1
clust_4_TCTATACAGGAACTCG−1
clust_4_ACGGAAGAGGTTCATC−1
clust_4_TAAGCGTAGGACTGGT−1
clust_4_TATTTCGGTTTACTTC−1
clust_1_AAGCGTTAGTTCTCTT−1
clust_1_TTCAATCAGTAAATGC−1
clust_1_TGATCTTGTGATGGCA−1
clust_1_TTGGGCGAGCTCGACC−1
clust_3_CCCTGATAGGTATTGA−1
clust_3_GGGAAGTCAATGTCAC−1
clust_3_AGATGCTCATCGCCTT−1
clust_3_AGAAATGGTTTGCCGG−1
clust_3_GACGCTGCATAATCGC−1
clust_3_TATTTCGCATTACTCT−1
350
300
250
200
150
100
50
0

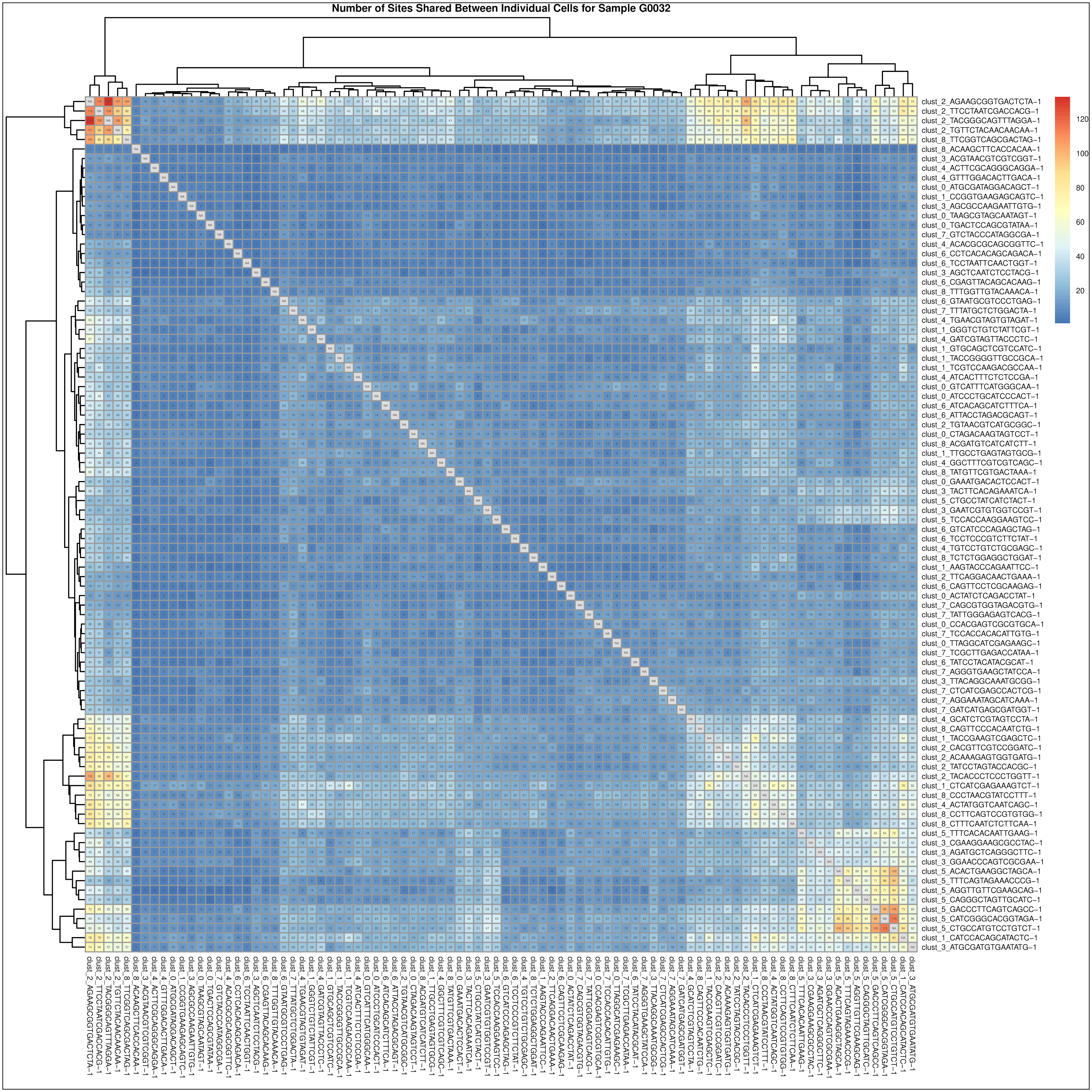
Number of Sites Shared Between Individual Cells for Sample G0032
120
100
80
60
40
20
clust_2_AGAAGCGGTGACTCTA-1
clust_2_TTCCTAATCGACCACG-1
clust_2_TACGGGCAGTTTAGGA-1
clust_2_TGTTCTACAACAACAA-1
clust_8_TTCGGTCAGCGACTAG-1
clust_8_ACAAGCTTCACCACAA-1
clust_3_ACGTAACGTCGTCGGT-1
clust_4_ACTTCGCAGGGCAGGA-1
clust_4_GTTTGGACACTTGACA-1
clust_0_ATGCGATAGGACAGCT-1
clust_1_CCGGTGAAGAGCAGTC-1
clust_3_AGCGCCAAGAATTGTG-1
clust_0_TAAGCGTAGCAATAGT-1
clust_0_TGACTCCAGCGTATAA-1
clust_7_GTCTACCCATAGGCGA-1
clust_4_ACACGCGCAGCGGTTC-1
clust_6_CCTCACACAGCAGACA-1
clust_6_TCCTAATTCAACTGGT-1
clust_3_AGCTCAATCTCCTACG-1
clust_6_CGAGTTACAGCACAAG-1
clust_8_TTTGGTTGTACAAACA-1
clust_6_GTAATGCGTCCCTGAG-1
clust_7_TTTATGCTCTGGACTA-1
clust_4_TGAACGTAGTGTAGAT-1
clust_1_GGGTCTGTCTATTCGT-1
clust_4_GATCGTAGTTACCCTC-1
clust_1_GTGCAGCTCGTCCATC-1
clust_1_TACCGGGGTTGCCGCA-1
clust_1_TCGTCCAAGACGCCAA-1
clust_4_ATCACTTTCTCTCCGA-1
clust_0_GTCATTTCATGGGCAA-1
clust_0_ATCCCTGCATCCCACT-1
clust_6_ATCACAGCATCTTTCA-1
clust_6_ATTACCTAGACGCAGT-1
clust_2_TGTAACGTCATGCGGC-1
clust_0_CTAGACAAGTAGTCCT-1
clust_8_ACGATGTCATCATCTT-1
clust_1_TTGCCTGAGTAGTGCG-1
clust_4_GGCTTTCGTCGTCAGC-1
clust_8_TATGTTCGTGACTAAA-1
clust_0_GAAATGACACTCCACT-1
clust_3_TACTTCACAGAAATCA-1
clust_5_CTGCCTATCATCTACT-1
clust_3_GAATCGTGTGGTCCGT-1
clust_5_TCCACCAAGGAAGTCC-1
clust_6_GTCATCCCAGAGCTAG-1
clust_6_TCCTCCCGTCTTCTAT-1
clust_4_TGTCCTGTCTGCGAGC-1
clust_8_TCTCTGGAGGCTGGAT-1
clust_1_AAGTACCCAGAATTCC-1
clust_2_TTCAGGACAACTGAAA-1
clust_6_CAGTTCCTCGCAAGAG-1
clust_0_ACTATCTCAGACCTAT-1
clust_7_CAGCGTGGTAGACGTG-1
clust_7_TATTGGGAGAGTCACG-1
clust_0_CCACGAGTCGCGTGCA-1
clust_7_TCCACCACACATTGTG-1
clust_0_TTAGGCATCGAGAAGC-1
clust_7_TCGCTTGAGACCATAA-1
clust_6_TATCCTACATACGCAT-1
clust_7_AGGGTGAAGCTATCCA-1
clust_3_TTACAGGCAAATGCGG-1
clust_7_CTCATCGAGCCACTCG-1
clust_7_AGGAAATAGCATCAAA-1
clust_7_GATCATGAGCGATGGT-1
clust_4_GCATCTCGTAGTCCTA-1
clust_8_CAGTTCCCACAATCTG-1
clust_1_TACCGAAGTCGAGCTC-1
clust_2_CACGTTCGTCCGGATC-1
clust_2_ACAAAGAGTGGTGATG-1
clust_2_TATCCTAGTACCACGC-1
clust_2_TACACCCTCCCTGGTT-1
clust_1_CTCATCGAGAAAGTCT-1
clust_8_CCCTAACGTATCCTTT-1
clust_4_ACTATGGTCAATCAGC-1
clust_8_CCTTCAGTCCGTGTGG-1
clust_8_CTTTCAATCTCTTCAA-1
clust_5_TTTCACACAATTGAAG-1
clust_3_CGAAGGAAGCGCCTAC-1
clust_3_AGATGCTCAGGGCTTC-1
clust_3_GGAACCCAGTCGCGAA-1
clust_5_ACACTGAAGGCTAGCA-1
clust_5_TTTCAGTAGAAACCCG-1
clust_5_AGGTTGTTCGAAGCAG-1
clust_5_CAGGGCTAGTTGCATC-1
clust_5_GACCCTTCAGTCAGCC-1
clust_5_CATCGGGCACGGTAGA-1
clust_5_CTGCCATGTCCTGTCT-1
clust_1_CATCCACAGCATACTC-1
clust_3_ATGCGATGTGAATATG-1

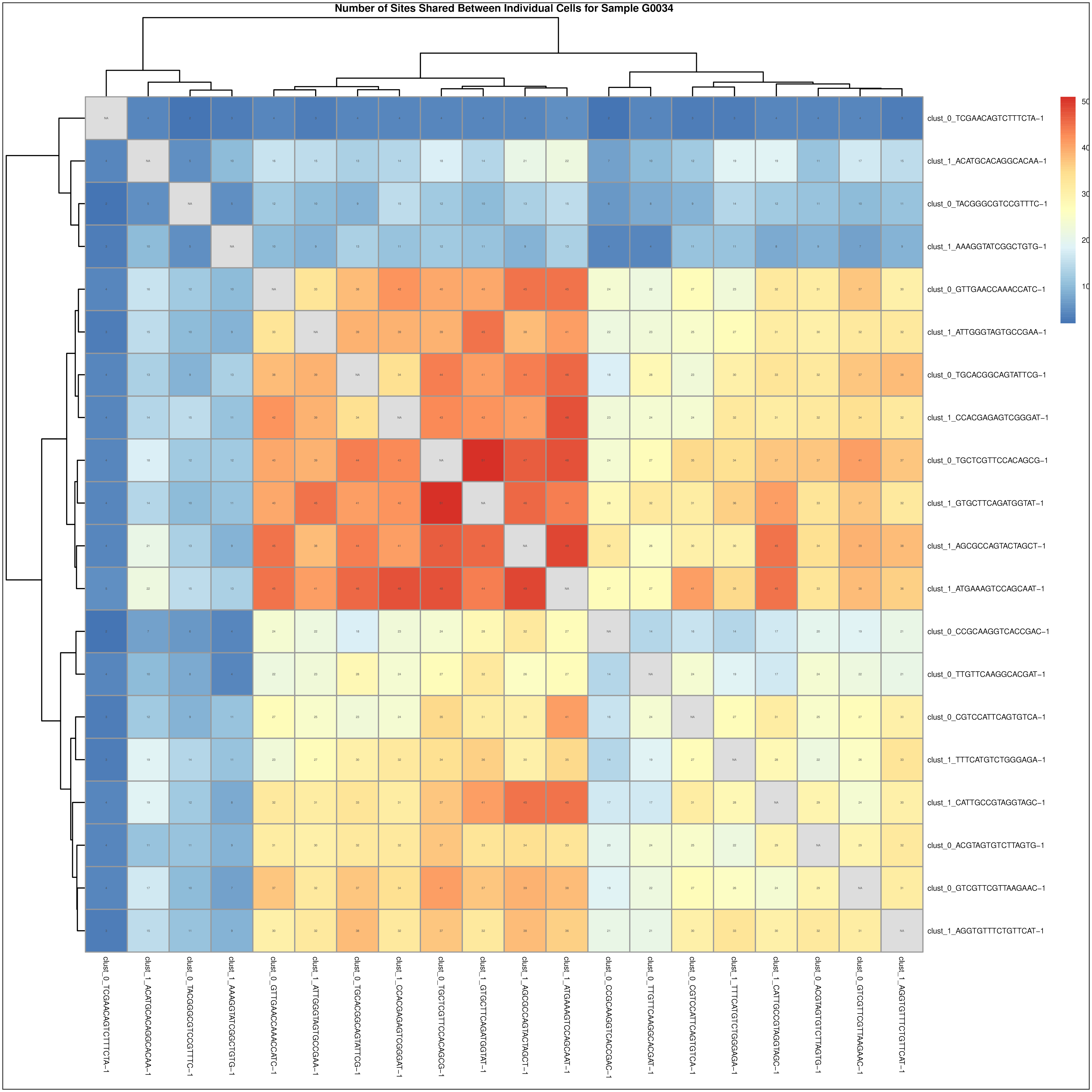
Number of Sites Shared Between Individual Cells for Sample G0034
clust_0_TCGAACAGTCTTTCTA−1
clust_1_ACATGCACAGGCACAA−1
clust_0_TACGGGCGTCCGTTTC−1
clust_1_AAAGGTATCGGCTGTG−1
clust_0_GTTGAACCAAACCATC−1
clust_1_ATTGGGTAGTGCCGAA−1
clust_0_TGCACGGCAGTATTCG−1
clust_1_CCACGAGAGTCGGGAT−1
clust_0_TGCTCGTTCCACAGCG−1
clust_1_GTGCTTCAGATGGTAT−1
clust_1_AGCGCCAGTACTAGCT−1
clust_1_ATGAAAGTCCAGCAAT−1
clust_0_CCGCAAGGTCACCGAC−1
clust_0_TTGTTCAAGGCACGAT−1
clust_0_CGTCCATTCAGTGTCA−1
clust_1_TTTCATGTCTGGGAGA−1
clust_1_CATTGCCGTAGGTAGC−1
clust_0_ACGTAGTGTCTTAGTG−1
clust_0_GTCGTTCGTTAAGAAC−1
clust_1_AGGTGTTTCTGTTCAT−1

**Number of Sites Shared Between Individual Cells for Sample G0035**

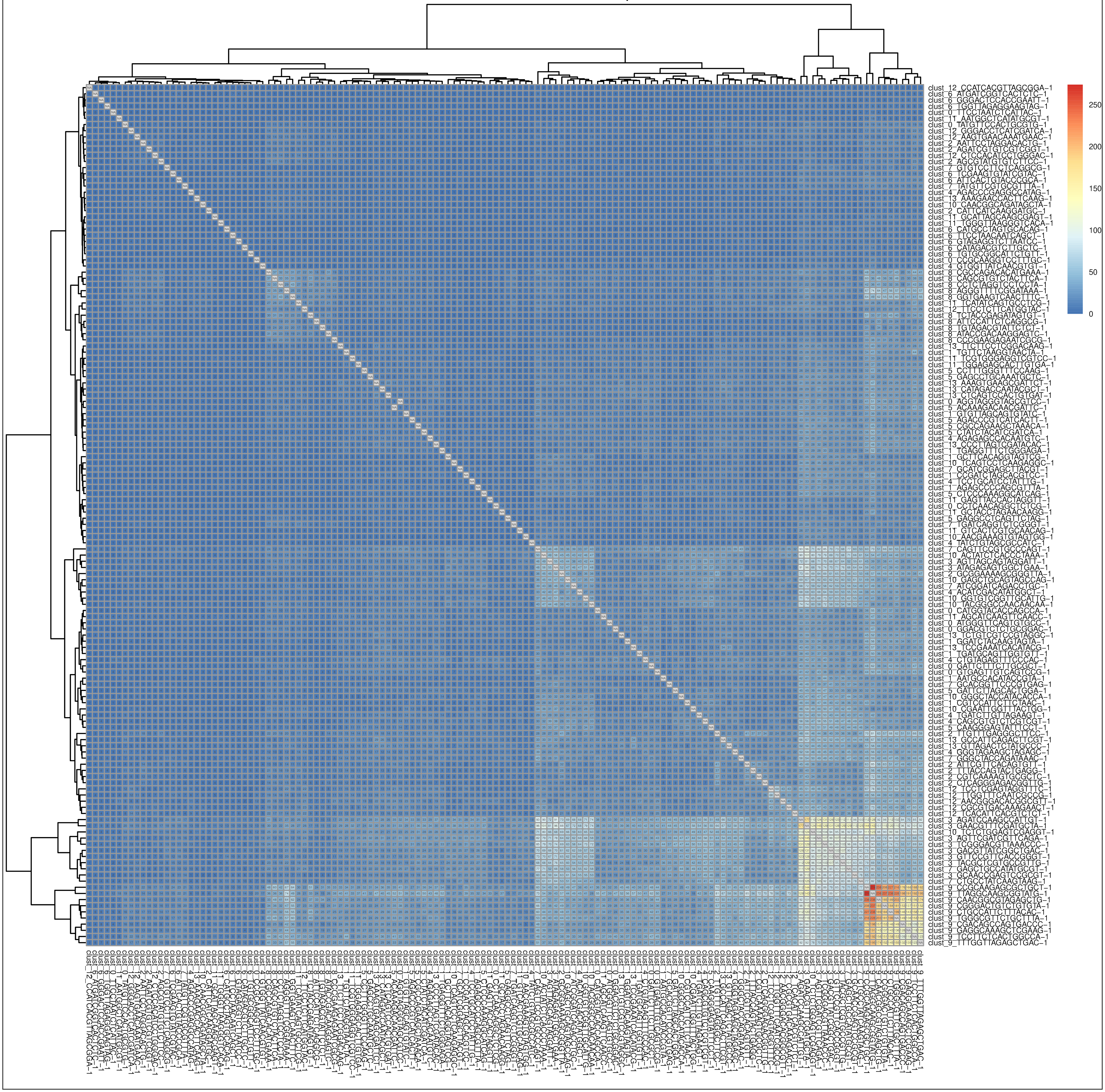

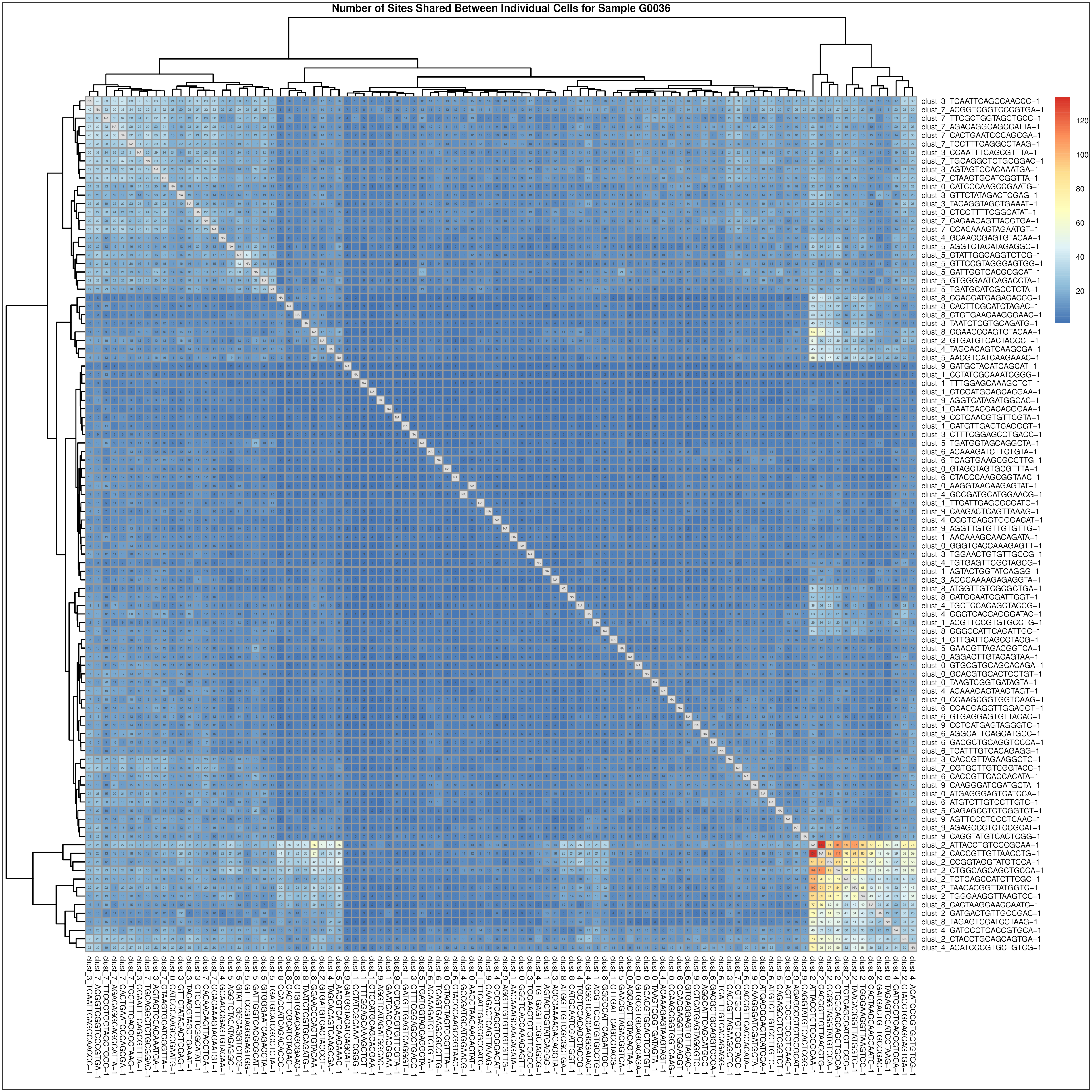
Number of Sites Shared Between Individual Cells for Sample G0036
120
100
80
60
40
20
clust_3_TCAATTCAGCCAACCC-1
clust_7_ACGGTCGGTCCCGTGA-1
clust_7_TTCGCTGGTAGCTGCC-1
clust_7_AGACAGGCAGCCATTA-1
clust_7_CACTGAATCCCAGCGA-1
clust_7_TCCTTTCAGGCCTAAG-1
clust_3_CCAATTTCAGCGTTTA-1
clust_7_TGCAGGCTCTGCGGAC-1
clust_3_AGTAGTCCACAAATGA-1
clust_7_CTAAGTGCATCGGTTA-1
clust_0_CATCCCAAGCCGAATG-1
clust_3_GTTCTATAGACTCGAG-1
clust_3_TACAGGTAGCTGAAAT-1
clust_3_CTCCTTTTCGGCATAT-1
clust_7_CACAACAGTTACCTGA-1
clust_7_CCACAAAGTAGAATGT-1
clust_4_GCAACCGAGTGTACAA-1
clust_5_AGGTCTACATAGAGGC-1
clust_5_GTATTGGCAGGTCTCG-1
clust_5_GTTCCGTAGGGAGTGG-1
clust_5_GATTGGTCACGCGCAT-1
clust_5_GTGGGAATCAGACCTA-1
clust_5_TGATGCATCGCCTCTA-1
clust_8_CCACCATCAGACACCC-1
clust_8_CACTTCGCATCTAGAC-1
clust_8_CTGTGAACAAGCGAAC-1
clust_8_TAATCTCGTGCAGATG-1
clust_8_GGAACCCAGTGTACAA-1
clust_2_GTGATGTCACTACCCT-1
clust_4_TAGCACAGTCAAGCGA-1
clust_5_AACGTCATCAAGAAAC-1
clust_9_GATGCTACATCAGCAT-1
clust_1_CCTATCGCAAATCGGG-1
clust_1_TTTGGAGCAAAGCTCT-1
clust_1_CTCCATGCAGCACGAA-1
clust_9_AGGTCATAGATGGCAC-1
clust_1_GAATCACCACACGGAA-1
clust_9_CCTCAACGTGTTCGTA-1
clust_1_GATGTTGAGTCAGGGT-1
clust_3_CTTTCGGAGCCTGACC-1
clust_5_TGATGGTAGCAGGCTA-1
clust_6_ACAAAGATCTTCTGTA-1
clust_6_TCAGTGAAGCGCCTTG-1
clust_0_GTAGCTAGTGCGTTTA-1
clust_6_CTACCCAAGCGGTAAC-1
clust_0_AAGGTAACAAGAGTAT-1
clust_4_GCCGATGCATGGAACG-1
clust_1_TTCATTGAGCGCCATC-1
clust_9_CAAGACTCAGTTAAAG-1
clust_4_CGGTCAGGTGGGACAT-1
clust_9_AGGTTGTGTTGTGTTG-1
clust_1_AACAAAGCAACAGATA-1
clust_0_GGGTCACCAAAGAGTT-1
clust_3_TGGAACTGTGTTGCCG-1
clust_4_TGTGAGTTCGCTAGCG-1
clust_1_AGTACTGGTATCAGGG-1
clust_3_ACCCAAAAGAGAGGTA-1
clust_8_ATGGTTGTCGCGCTGA-1
clust_8_CATGCAATCGATTGGT-1
clust_4_TGCTCCACAGCTACCG-1
clust_4_GGGTCACCAGGGATAC-1
clust_1_ACGTTCCGTGTGCCTG-1
clust_8_GGGCCATTCAGATTGC-1
clust_1_CTTGATTCAGCCTACG-1
clust_5_GAACGTTAGACGGTCA-1
clust_0_AGGACTTGTACAGTAA-1
clust_0_GTGCGTGCAGCACAGA-1
clust_0_GCACGTGCACTCCTGT-1
clust_0_TAAGTCGGTGATAGTA-1
clust_4_ACAAAGAGTAAGTAGT-1
clust_0_CCAAGCGGTGGTCAAG-1
clust_6_CCACGAGGTTGGAGGT-1
clust_6_GTGAGGAGTGTTACAC-1
clust_9_CCTCATGAGTAGGGTC-1
clust_6_AGGCATTCAGCATGCC-1
clust_6_GACGCTGCAGGTCCCA-1
clust_6_TCATTTGTCACAGAGG-1
clust_3_CACCGTTAGAAGGCTC-1
clust_7_CGTGCTTGTCGGTACC-1
clust_6_CACCGTTCACCACATA-1
clust_9_CAAGGGATCGATGCTA-1
clust_0_ATGAGGGAGTCATCCA-1
clust_6_ATGTCTTGTCCTTGTC-1
clust_5_CAGAGCCTCTCGGTCT-1
clust_9_AGTTCCCTCCCTCAAC-1
clust_9_AGAGCCCTCTCCGCAT-1
clust_9_CAGGTATGTCACTCGG-1
clust_2_ATTACCTGTCCGCAAA-1
clust_2_CACCGTTGTTAACCTG-1
clust_2_CCGGTAGGTATGTCCA-1
clust_2_CTGGCAGCAGCTGCCA-1
clust_2_TCTCAGCCATCTTCGC-1
clust_2_TAACACGGTTATGGTC-1
clust_2_TGGGAAGGTTAAGTCC-1
clust_8_CACTAAGCAACCAATC-1
clust_2_GATGACTGTTGCCGAC-1
clust_8_TAGAGTCCATCCTAAG-1
clust_4_GATCCCTCACCGTGCA-1
clust_2_CTACCTGCAGCAGTGA-1
clust_4_ACATCCCGTGCTGTCG-1

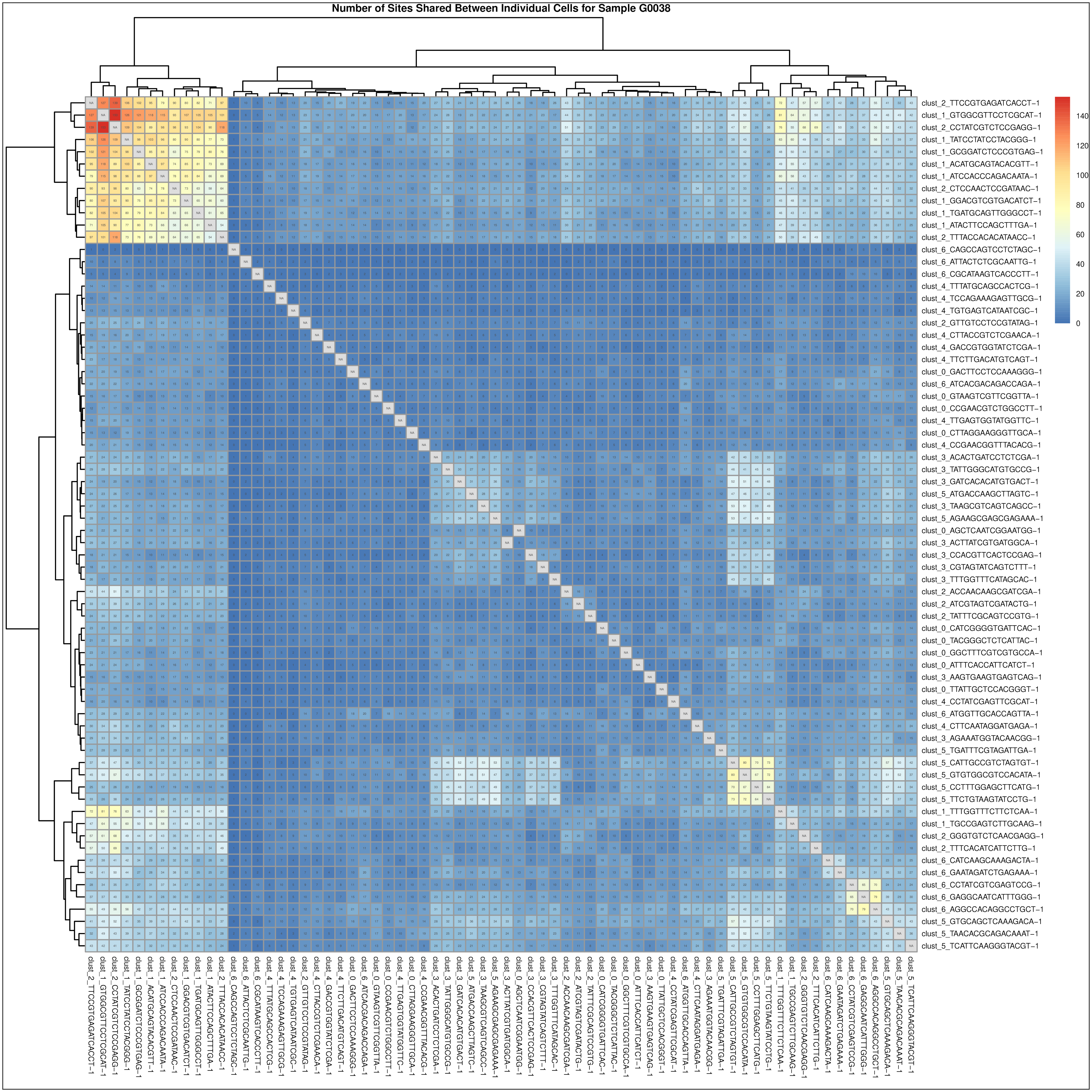
Number of Sites Shared Between Individual Cells for Sample G0038
140
120
100
80
60
40
20
0
clust_2_TTCCGTGAGATCACCT-1
clust_1_GTGGCGTTCCTCGCAT-1
clust_2_CCTATCGTCTCCGAGG-1
clust_1_TATCCTATCCTACGGG-1
clust_1_GCGGATCTCCCGTGAG-1
clust_1_ACATGCAGTACACGTT-1
clust_1_ATCCACCCAGACAATA-1
clust_2_CTCCAACTCCGATAAC-1
clust_1_GGACGTCGTGACATCT-1
clust_1_TGATGCAGTTGGGCCT-1
clust_1_ATACTTCCAGCTTTGA-1
clust_2_TTTACCACACATAACC-1
clust_6_CAGCCAGTCCTCTAGC-1
clust_6_ATTACTCTCGCAATTG-1
clust_6_CGCATAAGTCACCCTT-1
clust_4_TTTATGCAGCCACTCG-1
clust_4_TCCAGAAAGAGTTGCG-1
clust_4_TGTGAGTCATAATCGC-1
clust_2_GTTGTCCTCCGTATAG-1
clust_4_CTTACCGTCTCGAACA-1
clust_4_GACCGTGGTATCTCGA-1
clust_4_TTCTTGACATGTCAGT-1
clust_0_GACTTCCTCCAAAGGG-1
clust_6_ATCACGACAGACCAGA-1
clust_0_GTAAGTCGTTCGGTTA-1
clust_0_CCGAACGTCTGGCCTT-1
clust_4_TTGAGTGGTATGGTTC-1
clust_0_CTTAGGAAGGGTTGCA-1
clust_4_CCGAACGGTTTACACG-1
clust_3_ACACTGATCCTCTCGA-1
clust_3_TATTGGGCATGTGCCG-1
clust_3_GATCACACATGTGACT-1
clust_5_ATGACCAAGCTTAGTC-1
clust_3_TAAGCGTCAGTCAGCC-1
clust_5_AGAAGCGAGCGAGAAA-1
clust_0_AGCTCAATCGGAATGG-1
clust_3_ACTTATCGTGATGGCA-1
clust_3_CCACGTTCACTCCGAG-1
clust_3_CGTAGTATCAGTCTTT-1
clust_3_TTTGGTTTCATAGCAC-1
clust_2_ACCAACAAGCGATCGA-1
clust_2_ATCGTAGTCGATACTG-1
clust_2_TATTTCGCAGTCCGTG-1
clust_0_CATCGGGGTGATTCAC-1
clust_0_TACGGGCTCTCATTAC-1
clust_0_GGCTTTCGTCGTGCCA-1
clust_0_ATTTCACCATTCATCT-1
clust_3_AAGTGAAGTGAGTCAG-1
clust_0_TTATTGCTCCACGGGT-1
clust_4_CCTATCGAGTTCGCAT-1
clust_6_ATGGTTGCACCAGTTA-1
clust_4_CTTCAATAGGATGAGA-1
clust_3_AGAAATGGTACAACGG-1
clust_5_TGATTTCGTAGATTGA-1
clust_5_CATTGCCGTCTAGTGT-1
clust_5_GTGTGGCGTCCACATA-1
clust_5_CCTTTGGAGCTTCATG-1
clust_5_TTCTGTAAGTATCCTG-1
clust_1_TTTGGTTTCTTCTCAA-1
clust_1_TGCCGAGTCTTGCAAG-1
clust_2_GGGTGTCTCAACGAGG-1
clust_2_TTTCACATCATTCTTG-1
clust_6_CATCAAGCAAAGACTA-1
clust_6_GAATAGATCTGAGAAA-1
clust_6_CCTATCGTCGAGTCCG-1
clust_6_GAGGCAATCATTTGGG-1
clust_6_AGGCCACAGGCCTGCT-1
clust_5_GTGCAGCTCAAAGACA-1
clust_5_TAACACGCAGACAAAT-1
clust_5_TCATTCAAGGGTACGT-1

**Number of Sites Shared Between Individual Cells for Sample G0039**

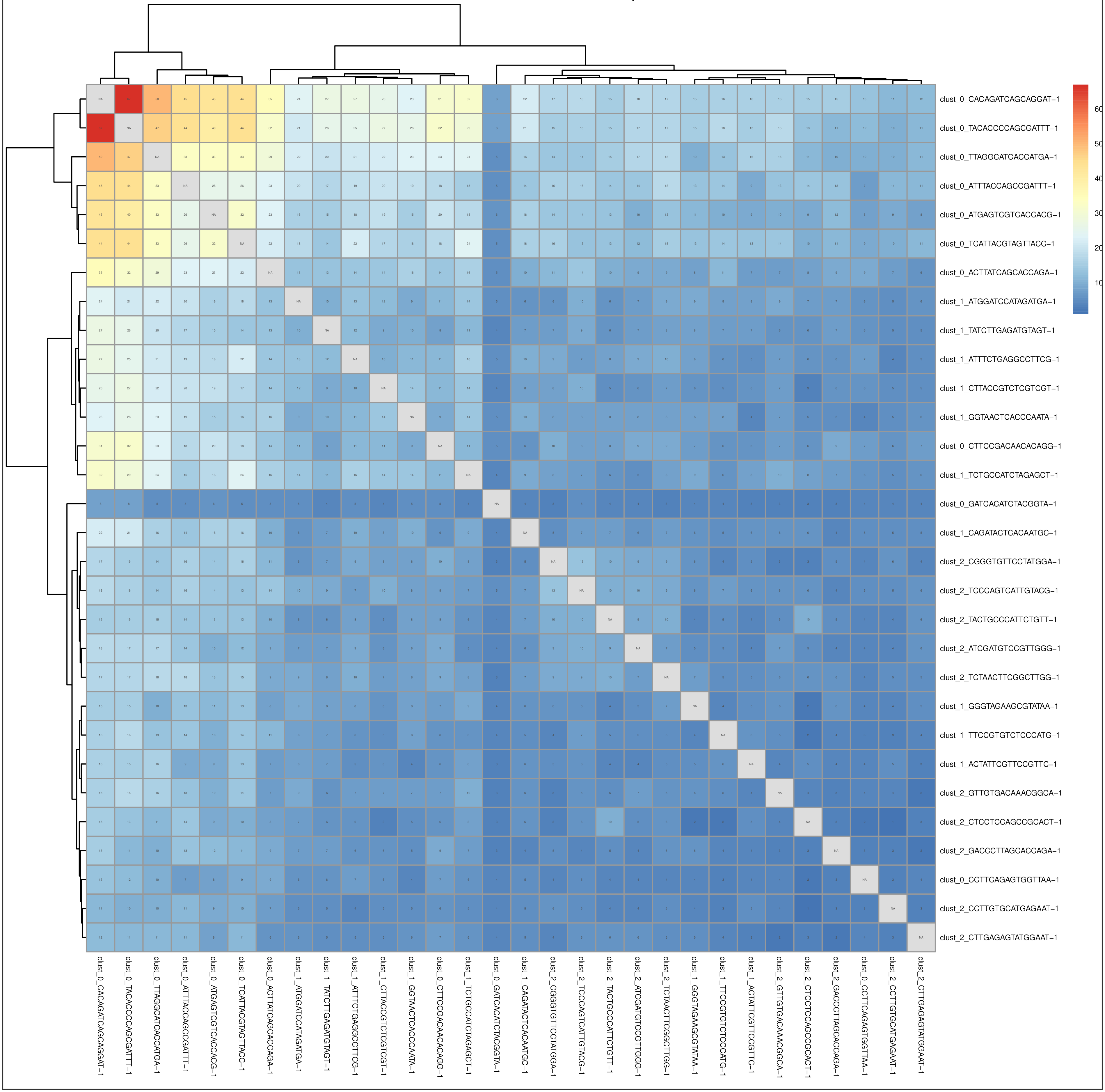

**Number of Sites Shared Between Individual Cells for Sample G0040**

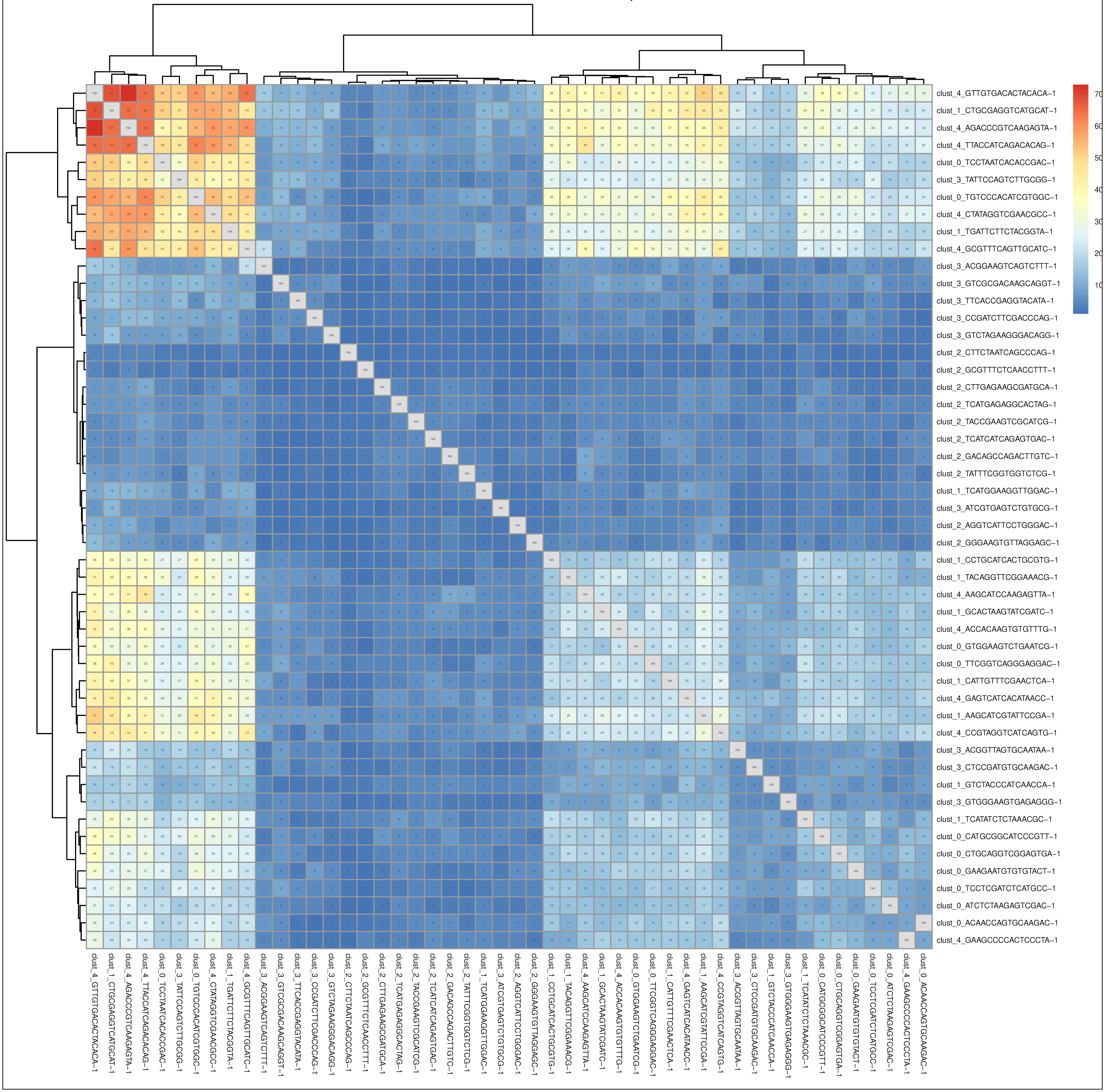

Number of Sites Shared Between Individual Cells for Sample G0041

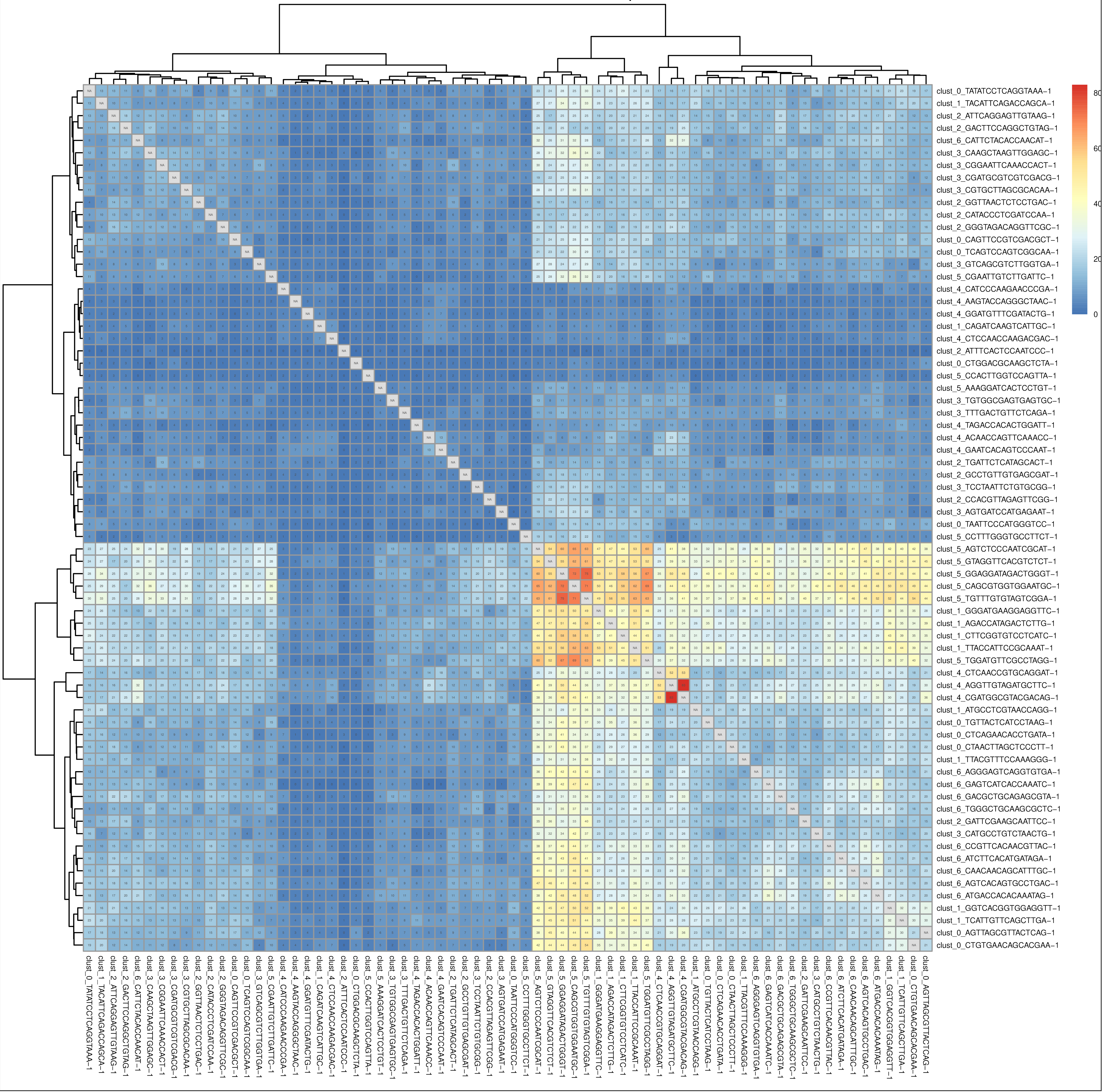

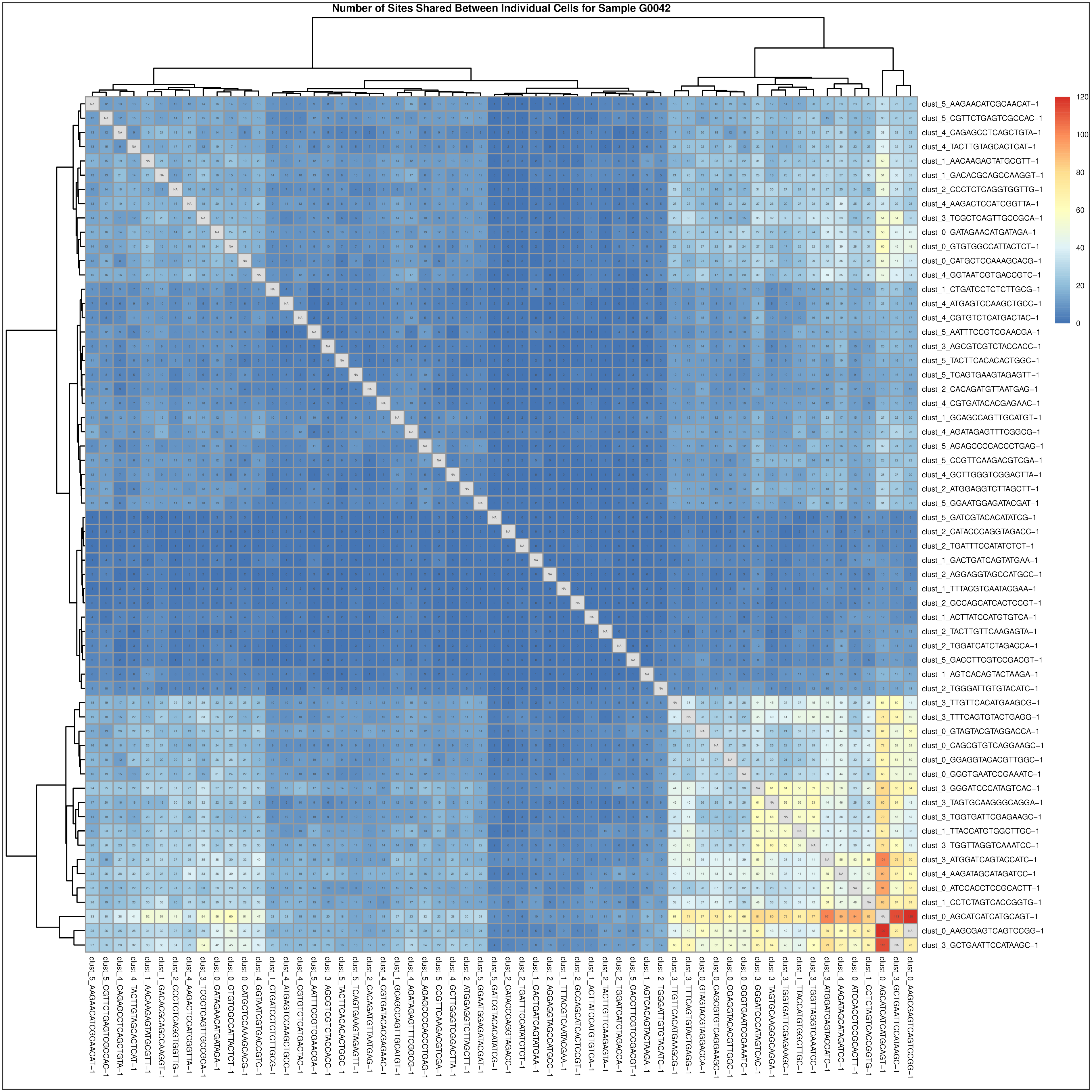
Number of Sites Shared Between Individual Cells for Sample G0042
120
100
80
60
40
20
0
clust_5_AAGAACATCGCAACAT-1
clust_5_CGTTCTGAGTCGCCAC-1
clust_4_CAGAGCCTCAGCTGTA-1
clust_4_TACTTGTAGCACTCAT-1
clust_1_AACAAGAGTATGCGTT-1
clust_1_GACACGCAGCCAAGGT-1
clust_2_CCCTCTCAGGTGGTTG-1
clust_4_AAGACTCCATCGGTTA-1
clust_3_TCGCTCAGTTGCCGCA-1
clust_0_GATAGAACATGATAGA-1
clust_0_GTGTGGCCATTACTCT-1
clust_0_CATGCTCCAAAGCACG-1
clust_4_GGTAATCGTGACCGTC-1
clust_1_CTGATCCTCTCTTGCG-1
clust_4_ATGAGTCCAAGCTGCC-1
clust_4_CGTGTCTCATGACTAC-1
clust_5_AATTTCCGTCGAACGA-1
clust_3_AGCGTCGTCTACCACC-1
clust_5_TACTTCACACACTGGC-1
clust_5_TCAGTGAAGTAGAGTT-1
clust_2_CACAGATGTTAATGAG-1
clust_4_CGTGATACACGAGAAC-1
clust_1_GCAGCCAGTTGCATGT-1
clust_4_AGATAGAGTTTCGGCG-1
clust_5_AGAGCCCCACCCTGAG-1
clust_5_CCGTTCAAGACGTCGA-1
clust_4_GCTTGGGTCGGACTTA-1
clust_2_ATGGAGGTCTTAGCTT-1
clust_5_GGAATGGAGATACGAT-1
clust_5_GATCGTACACATATCG-1
clust_2_CATACCCAGGTAGACC-1
clust_2_TGATTTCCATATCTCT-1
clust_1_GACTGATCAGTATGAA-1
clust_2_AGGAGGTAGCCATGCC-1
clust_1_TTTACGTCAATACGAA-1
clust_2_GCCAGCATCACTCCGT-1
clust_1_ACTTATCCATGTGTCA-1
clust_2_TACTTGTTCAAGAGTA-1
clust_2_TGGATCATCTAGACCA-1
clust_5_GACCTTCGTCCGACGT-1
clust_1_AGTCACAGTACTAAGA-1
clust_2_TGGGATTGTGTACATC-1
clust_3_TTGTTCACATGAAGCG-1
clust_3_TTTCAGTGTACTGAGG-1
clust_0_GTAGTACGTAGGACCA-1
clust_0_CAGCGTGTCAGGAAGC-1
clust_0_GGAGGTACACGTTGGC-1
clust_0_GGGTGAATCCGAAATC-1
clust_3_GGGATCCCATAGTCAC-1
clust_3_TAGTGCAAGGGCAGGA-1
clust_3_TGGTGATTCGAGAAGC-1
clust_1_TTACCATGTGGCTTGC-1
clust_3_TGGTTAGGTCAAATCC-1
clust_3_ATGGATCAGTACCATC-1
clust_4_AAGATAGCATAGATCC-1
clust_0_ATCCACCTCCGCACTT-1
clust_1_CCTCTAGTCACCGGTG-1
clust_0_AGCATCATCATGCAGT-1
clust_0_AAGCGAGTCAGTCCGG-1
clust_3_GCTGAATTCCATAAGC-1

Number of Sites Shared Between Individual Cells for Sample G0043

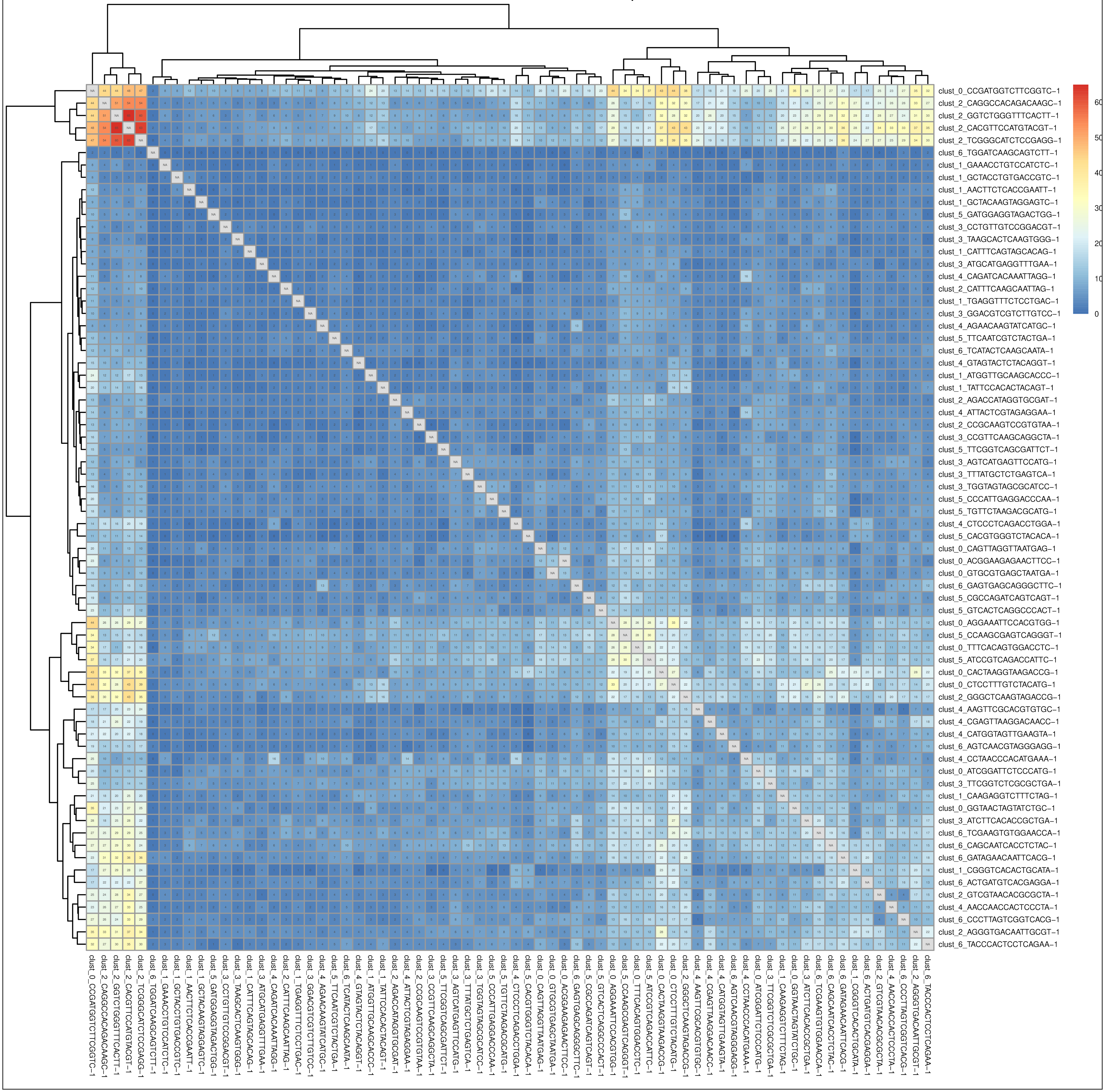

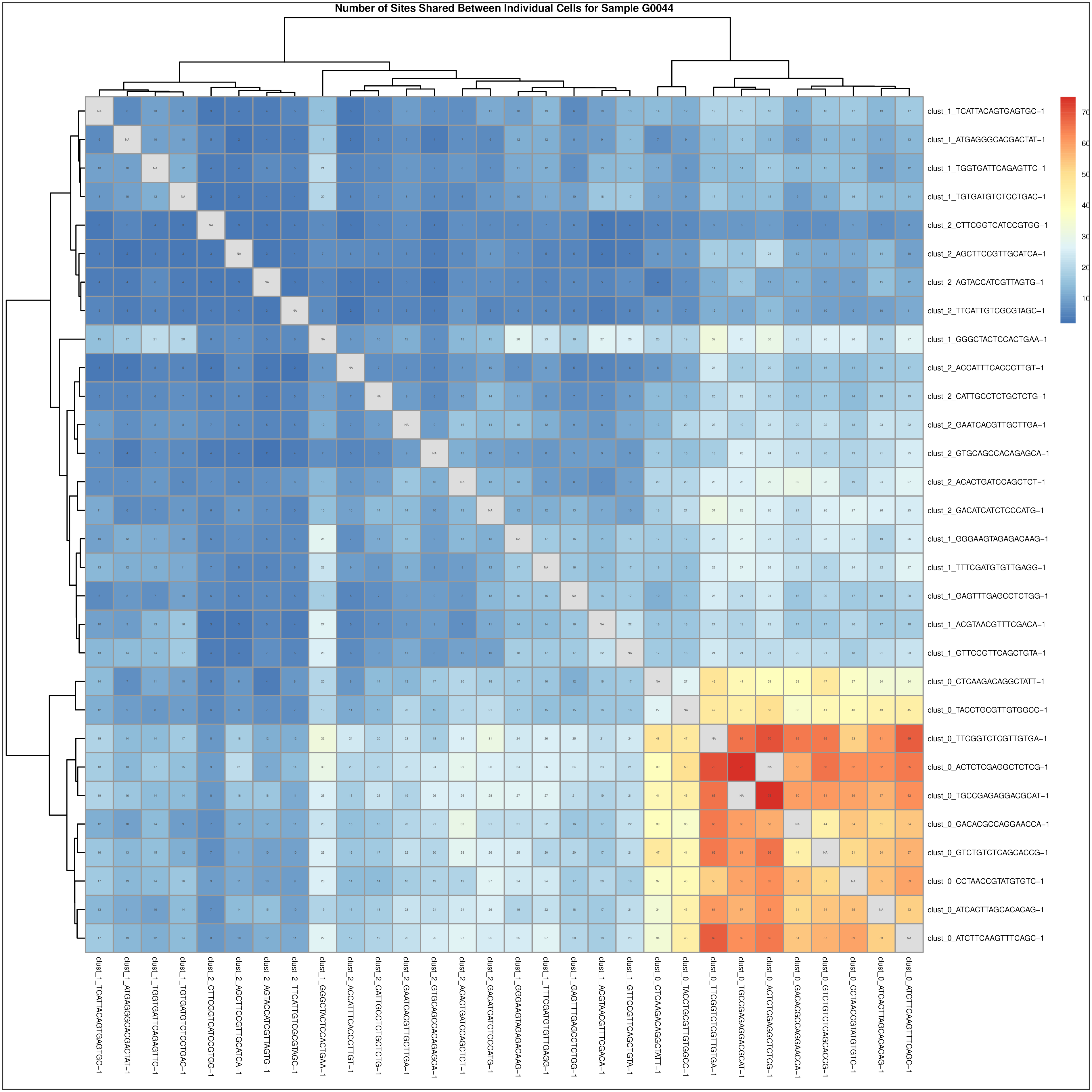
Number of Sites Shared Between Individual Cells for Sample G0044
clust_1_TCATTACAGTGAGTGC−1
clust_1_ATGAGGGCACGACTAT−1
clust_1_TGGTGATTCAGAGTTC−1
clust_1_TGTGATGTCTCCTGAC−1
clust_2_CTTCGGTCATCCGTGG−1
clust_2_AGCTTCCGTTGCATCA−1
clust_2_AGTACCATCGTTAGTG−1
clust_2_TTCATTGTCGCGTAGC−1
clust_1_GGGCTACTCCACTGAA−1
clust_2_ACCATTTCACCCTTGT−1
clust_2_CATTGCCTCTGCTCTG−1
clust_2_GAATCACGTTGCTTGA−1
clust_2_GTGCAGCCACAGAGCA−1
clust_2_ACACTGATCCAGCTCT−1
clust_2_GACATCATCTCCCATG−1
clust_1_GGGAAGTAGAGACAAG−1
clust_1_TTTCGATGTGTTGAGG−1
clust_1_GAGTTTGAGCCTCTGG−1
clust_1_ACGTAACGTTTCGACA−1
clust_1_GTTCCGTTCAGCTGTA−1
clust_0_CTCAAGACAGGCTATT−1
clust_0_TACCTGCGTTGTGGCC−1
clust_0_TTCGGTCTCGTTGTGA−1
clust_0_ACTCTCGAGGCTCTCG−1
clust_0_TGCCGAGAGGACGCAT−1
clust_0_GACACGCCAGGAACCA−1
clust_0_GTCTGTCTCAGCACCG−1
clust_0_CCTAACCGTATGTGTC−1
clust_0_ATCACTTAGCACACAG−1
clust_0_ATCTTCAAGTTTCAGC−1

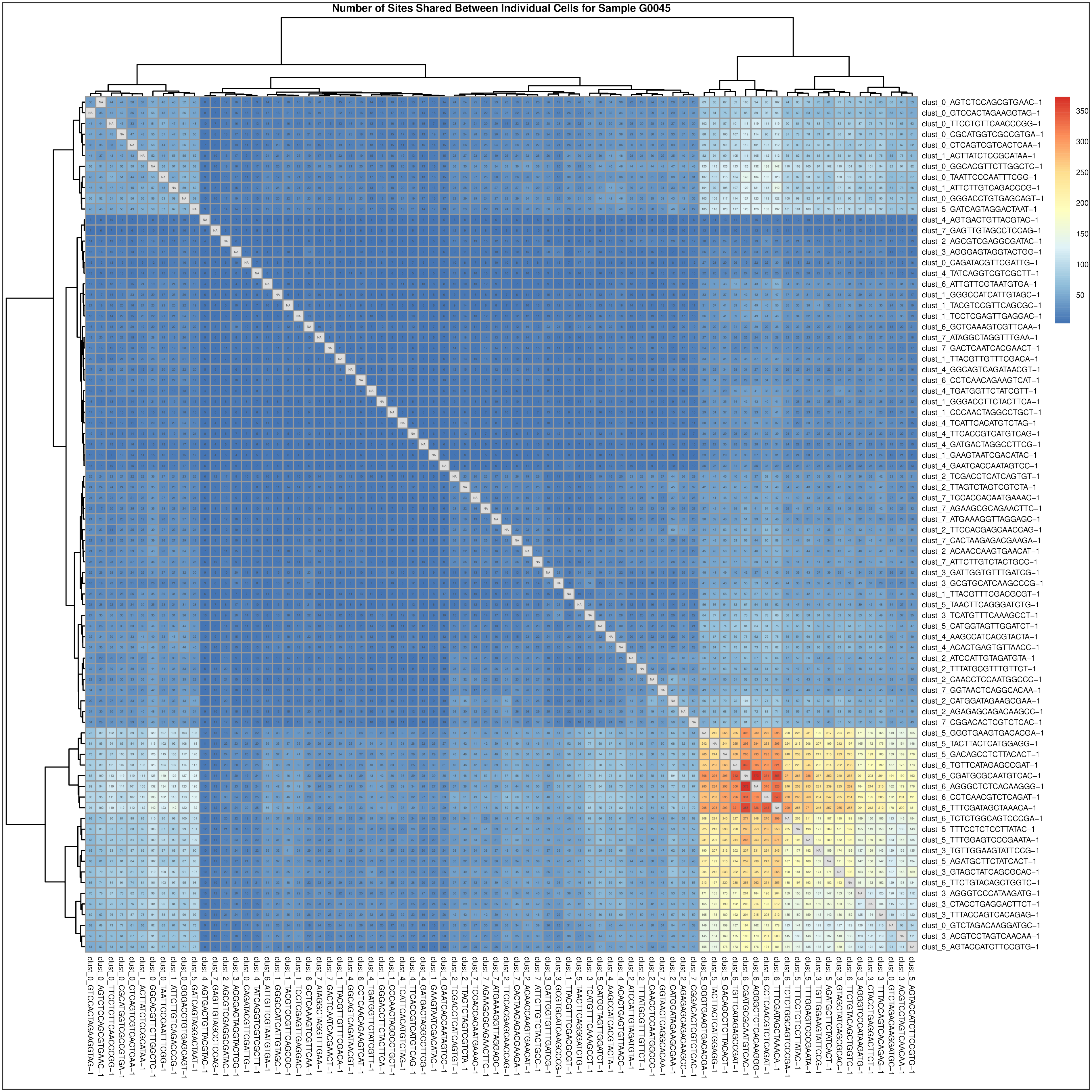
Number of Sites Shared Between Individual Cells for Sample G0045
350
300
250
200
150
100
50
clust_0_AGTCTCCAGCGTGAAC-1
clust_0_GTCCACTAGAAGGTAG-1
clust_0_TTCCTCTTCAACCCGG-1
clust_0_CGCATGGTCGCCGTGA-1
clust_0_CTCAGTCGTCACTCAA-1
clust_1_ACTTATCTCCGCATAA-1
clust_0_GGCACGTTCTTGGCTC-1
clust_0_TAATTCCCAATTTCGG-1
clust_1_ATTCTTGTCAGACCCG-1
clust_0_GGGACCTGTGAGCAGT-1
clust_5_GATCAGTAGGACTAAT-1
clust_4_AGTGACTGTTACGTAC-1
clust_7_GAGTTGTAGCCTCCAG-1
clust_2_AGCGTCGAGGCGATAC-1
clust_3_AGGGAGTAGGTACTGG-1
clust_0_CAGATACGTTCGATTG-1
clust_4_TATCAGGTCGTCGCTT-1
clust_6_ATTGTTCGTAATGTGA-1
clust_1_GGGCCATCATTGTAGC-1
clust_1_TACGTCCGTTCAGCGC-1
clust_1_TCCTCGAGTTGAGGAC-1
clust_6_GCTCAAAGTCGTTCAA-1
clust_7_ATAGGCTAGGTTTGAA-1
clust_7_GACTCAATCACGAACT-1
clust_1_TTACGTTGTTTCGACA-1
clust_4_GGCAGTCAGATAACGT-1
clust_6_CCTCAACAGAAGTCAT-1
clust_4_TGATGGTTCTATCGTT-1
clust_1_GGGACCTTCTACTTCA-1
clust_1_CCCAACTAGGCCTGCT-1
clust_4_TCATTCACATGTCTAG-1
clust_4_TTCACCGTCATGTCAG-1
clust_4_GATGACTAGGCCTTCG-1
clust_1_GAAGTAATCGACATAC-1
clust_4_GAATCACCAATAGTCC-1
clust_2_TCGACCTCATCAGTGT-1
clust_2_TTAGTCTAGTCGTCTA-1
clust_7_TCCACCACAATGAAAC-1
clust_7_AGAAGCGCAGAACTTC-1
clust_7_ATGAAAGGTTAGGAGC-1
clust_2_TTCCACGAGCAACCAG-1
clust_7_CACTAAGAGACGAAGA-1
clust_2_ACAACCAAGTGAACAT-1
clust_7_ATTCTTGTCTACTGCC-1
clust_3_GATTGGTGTTTGATCG-1
clust_3_GCGTGCATCAAGCCCG-1
clust_1_TTACGTTTCGACGCGT-1
clust_5_TAACTTCAGGGATCTG-1
clust_3_TCATGTTTCAAAGCCT-1
clust_5_CATGGTAGTTGGATCT-1
clust_4_AAGCCATCACGTACTA-1
clust_4_ACACTGAGTGTTAACC-1
clust_2_ATCCATTGTAGATGTA-1
clust_2_TTTATGCGTTTGTTCT-1
clust_2_CAACCTCCAATGGCCC-1
clust_7_GGTAACTCAGGCACAA-1
clust_2_CATGGATAGAAGCGAA-1
clust_2_AGAGAGCAGACAAGCC-1
clust_7_CGGACACTCGTCTCAC-1
clust_5_GGGTGAAGTGACACGA-1
clust_5_TACTTACTCATGGAGG-1
clust_5_GACAGCCTCTTACACT-1
clust_6_TGTTCATAGAGCCGAT-1
clust_6_CGATGCGCAATGTCAC-1
clust_6_AGGGCTCTCACAAGGG-1
clust_6_CCTCAACGTCTCAGAT-1
clust_6_TTTCGATAGCTAAACA-1
clust_6_TCTCTGGCAGTCCCGA-1
clust_5_TTTCCTCTCCTTATAC-1
clust_5_TTTGGAGTCCCGAATA-1
clust_3_TGTTGGAAGTATTCCG-1
clust_5_AGATGCTTCTATCACT-1
clust_3_GTAGCTATCAGCGCAC-1
clust_6_TTCTGTACAGCTGGTC-1
clust_3_AGGGTCCCATAAGATG-1
clust_3_CTACCTGAGGACTTCT-1
clust_3_TTTACCAGTCACAGAG-1
clust_0_GTCTAGACAAGGATGC-1
clust_3_ACGTCCTAGTCAACAA-1
clust_5_AGTACCATCTTCCGTG-1

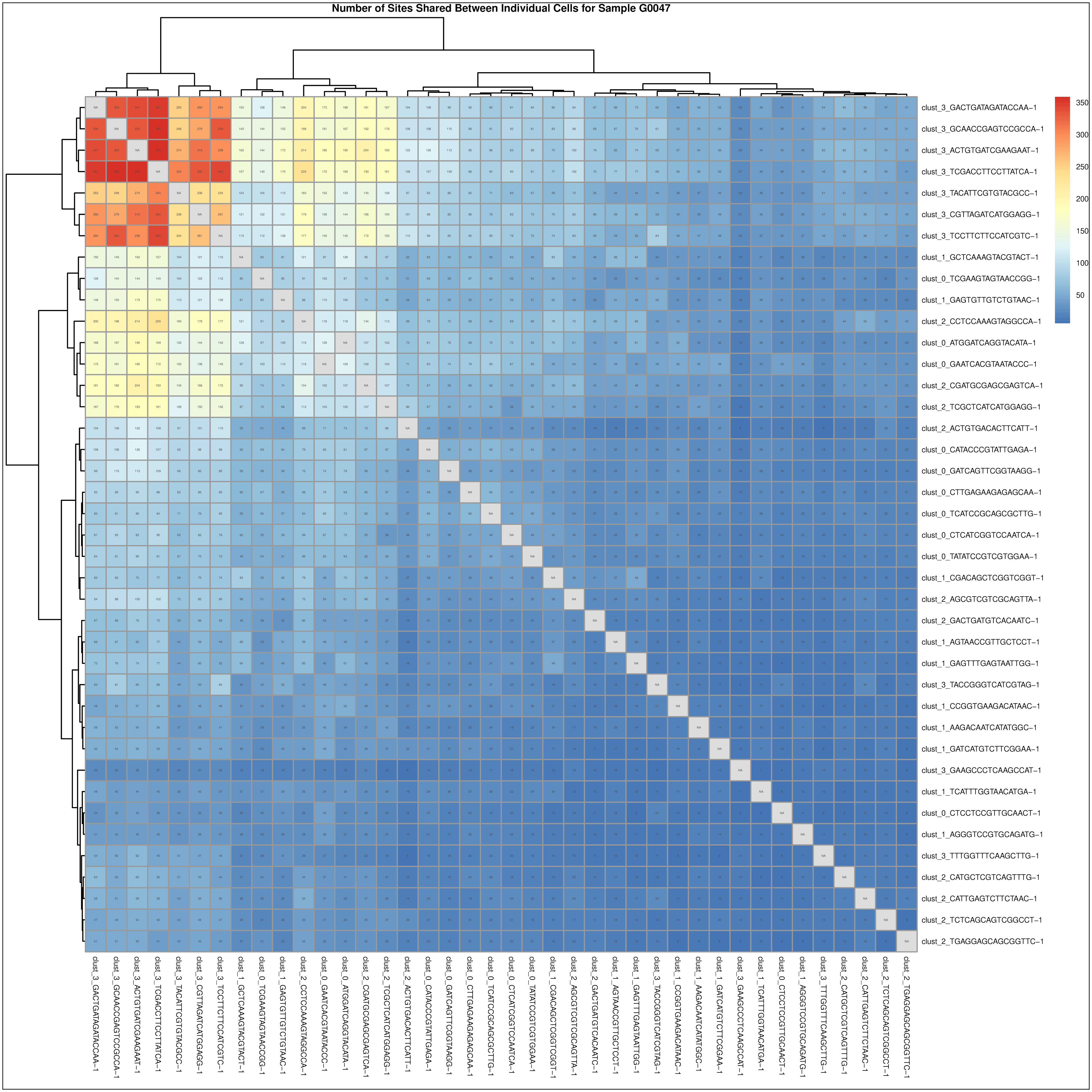
Number of Sites Shared Between Individual Cells for Sample G0047
clust_3_GACTGATAGATACCAA−1
clust_3_GCAACCGAGTCCGCCA−1
clust_3_ACTGTGATCGAAGAAT−1
clust_3_TCGACCTTCCTTATCA−1
clust_3_TACATTCGTGTACGCC−1
clust_3_CGTTAGATCATGGAGG−1
clust_3_TCCTTCTTCCATCGTC−1
clust_1_GCTCAAAGTACGTACT−1
clust_0_TCGAAGTAGTAACCGG−1
clust_1_GAGTGTTGTCTGTAAC−1
clust_2_CCTCCAAAGTAGGCCA−1
clust_0_ATGGATCAGGTACATA−1
clust_0_GAATCACGTAATACCC−1
clust_2_CGATGCGAGCGAGTCA−1
clust_2_TCGCTCATCATGGAGG−1
clust_2_ACTGTGACACTTCATT−1
clust_0_CATACCCGTATTGAGA−1
clust_0_GATCAGTTCGGTAAGG−1
clust_0_CTTGAGAAGAGAGCAA−1
clust_0_TCATCCGCAGCGCTTG−1
clust_0_CTCATCGGTCCAATCA−1
clust_0_TATATCCGTCGTGGAA−1
clust_1_CGACAGCTCGGTCGGT−1
clust_2_AGCGTCGTCGCAGTTA−1
clust_2_GACTGATGTCACAATC−1
clust_1_AGTAACCGTTGCTCCT−1
clust_1_GAGTTTGAGTAATTGG−1
clust_3_TACCGGGTCATCGTAG−1
clust_1_CCGGTGAAGACATAAC−1
clust_1_AAGACAATCATATGGC−1
clust_1_GATCATGTCTTCGGAA−1
clust_3_GAAGCCCTCAAGCCAT−1
clust_1_TCATTTGGTAACATGA−1
clust_0_CTCCTCCGTTGCAACT−1
clust_1_AGGGTCCGTGCAGATG−1
clust_3_TTTGGTTTCAAGCTTG−1
clust_2_CATGCTCGTCAGTTTG−1
clust_2_CATTGAGTCTTCTAAC−1
clust_2_TCTCAGCAGTCGGCCT−1
clust_2_TGAGGAGCAGCGGTTC−1
350
300
250
200
150
100
50

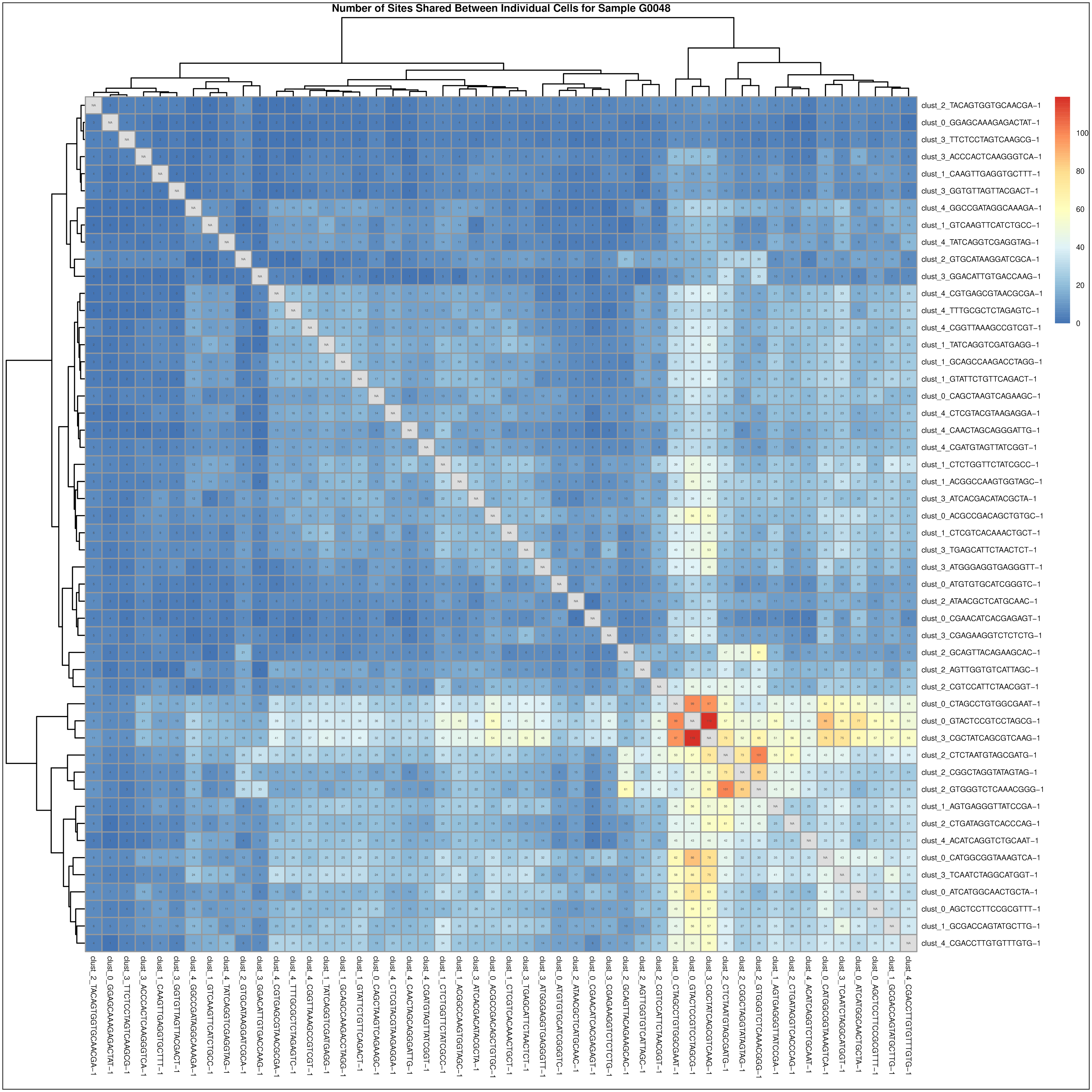
Number of Sites Shared Between Individual Cells for Sample G0048
clust_2_TACAGTGGTGCAACGA−1
clust_0_GGAGCAAAGAGACTAT−1
clust_3_TTCTCCTAGTCAAGCG−1
clust_3_ACCCACTCAAGGGTCA−1
clust_1_CAAGTTGAGGTGCTTT−1
clust_3_GGTGTTAGTTACGACT−1
clust_4_GGCCGATAGGCAAAGA−1
clust_1_GTCAAGTTCATCTGCC−1
clust_4_TATCAGGTCGAGGTAG−1
clust_2_GTGCATAAGGATCGCA−1
clust_3_GGACATTGTGACCAAG−1
clust_4_CGTGAGCGTAACGCGA−1
clust_4_TTTGCGCTCTAGAGTC−1
clust_4_CGGTTAAAGCCGTCGT−1
clust_1_TATCAGGTCGATGAGG−1
clust_1_GCAGCCAAGACCTAGG−1
clust_1_GTATTCTGTTCAGACT−1
clust_0_CAGCTAAGTCAGAAGC−1
clust_4_CTCGTACGTAAGAGGA−1
clust_4_CAACTAGCAGGGATTG−1
clust_4_CGATGTAGTTATCGGT−1
clust_1_CTCTGGTTCTATCGCC−1
clust_1_ACGGCCAAGTGGTAGC−1
clust_3_ATCACGACATACGCTA−1
clust_0_ACGCCGACAGCTGTGC−1
clust_1_CTCGTCACAAACTGCT−1
clust_3_TGAGCATTCTAACTCT−1
clust_3_ATGGGAGGTGAGGGTT−1
clust_0_ATGTGTGCATCGGGTC−1
clust_2_ATAACGCTCATGCAAC−1
clust_0_CGAACATCACGAGAGT−1
clust_3_CGAGAAGGTCTCTCTG−1
clust_2_GCAGTTACAGAAGCAC−1
clust_2_AGTTGGTGTCATTAGC−1
clust_2_CGTCCATTCTAACGGT−1
clust_0_CTAGCCTGTGGCGAAT−1
clust_0_GTACTCCGTCCTAGCG−1
clust_3_CGCTATCAGCGTCAAG−1
clust_2_CTCTAATGTAGCGATG−1
clust_2_CGGCTAGGTATAGTAG−1
clust_2_GTGGGTCTCAAACGGG−1
clust_1_AGTGAGGGTTATCCGA−1
clust_2_CTGATAGGTCACCCAG−1
clust_4_ACATCAGGTCTGCAAT−1
clust_0_CATGGCGGTAAAGTCA−1
clust_3_TCAATCTAGGCATGGT−1
clust_0_ATCATGGCAACTGCTA−1
clust_0_AGCTCCTTCCGCGTTT−1
clust_1_GCGACCAGTATGCTTG−1
clust_4_CGACCTTGTGTTTGTG−1
100
80
60
40
20
0

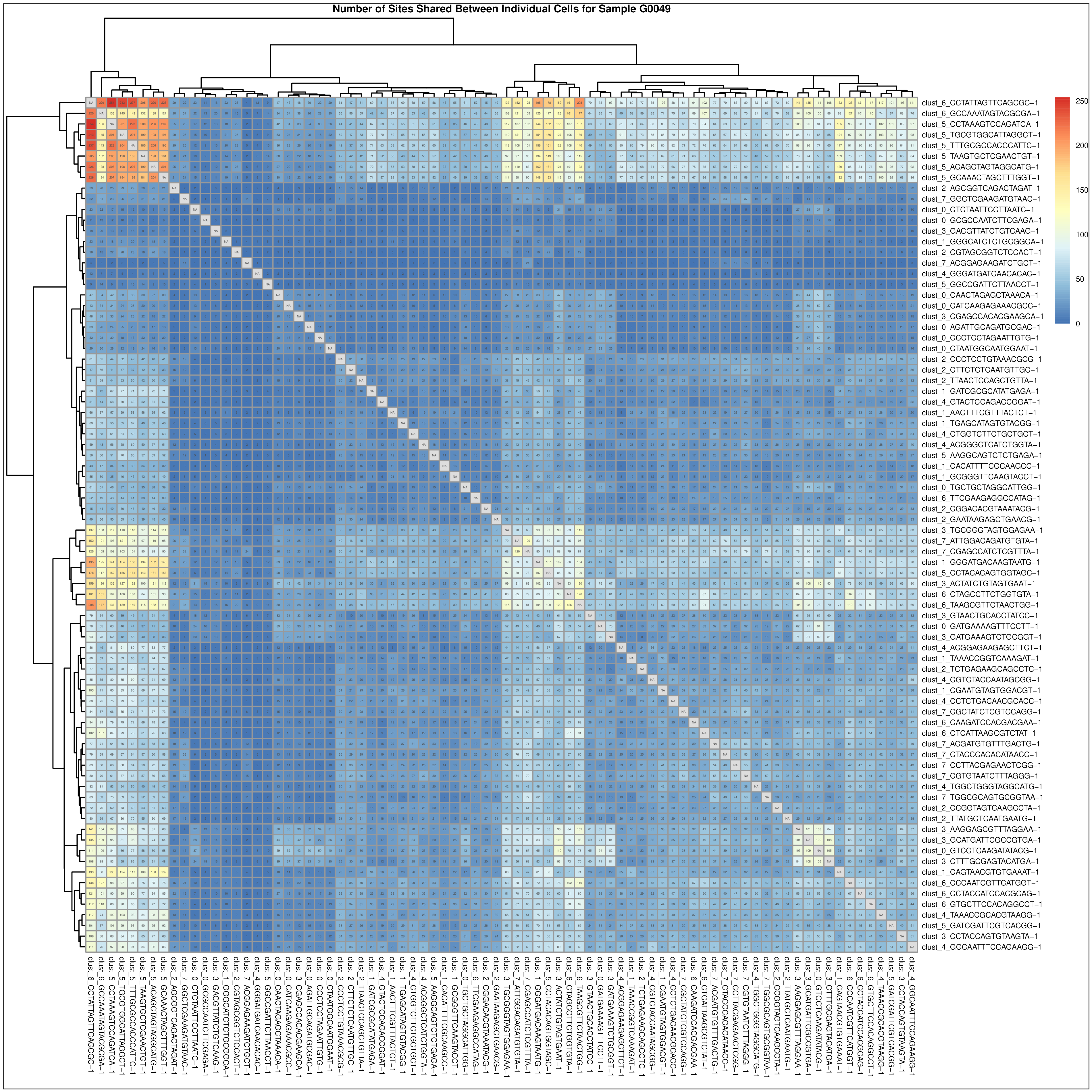

Number of Sites Shared Between Individual Cells for Sample G0049
250
200
150
100
50
0
clust_6_CCTATTAGTTCAGCGC-1
clust_6_GCCAAATAGTACGCGA-1
clust_5_CCTAAAGTCCAGATCA-1
clust_5_TGCGTGGCATTAGGCT-1
clust_5_TTTGCGCCACCCATTC-1
clust_5_TAAGTGCTCGAACTGT-1
clust_5_ACAGCTAGTAGGCATG-1
clust_5_GCAAACTAGCTTTGGT-1
clust_2_AGCGGTCAGACTAGAT-1
clust_7_GGCTCGAAGATGTAAC-1
clust_0_CTCTAATTCCTTAATC-1
clust_0_GCGCCAATCTTCGAGA-1
clust_3_GACGTTATCTGTCAAG-1
clust_1_GGGCATCTCTGCGGCA-1
clust_2_CGTAGCGGTCTCCACT-1
clust_7_ACGGAGAAGATCTGCT-1
clust_4_GGGATGATCAACACAC-1
clust_5_GGCCGATTCTTAACCT-1
clust_0_CAACTAGAGCTAAACA-1
clust_0_CATCAAGAGAAACGCC-1
clust_3_CGAGCCACACGAAGCA-1
clust_0_AGATTGCAGATGCGAC-1
clust_0_CCCTCCTAGAATTGTG-1
clust_0_CTAATGGCAATGGAAT-1
clust_2_CCCTCCTGTAAACGCG-1
clust_2_CTTCTCTCAATGTTGC-1
clust_2_TTAACTCCAGCTGTTA-1
clust_1_GATCGCGCATATGAGA-1
clust_4_GTACTCCAGACCGGAT-1
clust_1_AACTTTCGTTTACTCT-1
clust_1_TGAGCATAGTGTACGG-1
clust_4_CTGGTCTTCTGCTGCT-1
clust_4_ACGGGCTCATCTGGTA-1
clust_5_AAGGCAGTCTCTGAGA-1
clust_1_CACATTTTCGCAAGCC-1
clust_1_GCGGGTTCAAGTACCT-1
clust_0_TGCTGCTAGGCATTGG-1
clust_6_TTCGAAGAGGCCATAG-1
clust_2_CGGACACGTAAATACG-1
clust_2_GAATAAGAGCTGAACG-1
clust_3_TGCGGGTAGTGGAGAA-1
clust_7_ATTGGACAGATGTGTA-1
clust_7_CGAGCCATCTCGTTTA-1
clust_1_GGGATGACAAGTAATG-1
clust_5_CCTACACAGTGGTAGC-1
clust_3_ACTATCTGTAGTGAAT-1
clust_6_CTAGCCTTCTGGTGTA-1
clust_6_TAAGCGTTCTAACTGG-1
clust_3_GTAACTGCACCTATCC-1
clust_0_GATGAAAAGTTTCCTT-1
clust_3_GATGAAAGTCTGCGGT-1
clust_4_ACGGAGAAGAGCTTCT-1
clust_1_TAAACCGGTCAAAGAT-1
clust_2_TCTGAGAAGCAGCCTC-1
clust_4_CGTCTACCAATAGCGG-1
clust_1_CGAATGTAGTGGACGT-1
clust_4_CCTCTGACAACGCACC-1
clust_7_CGCTATCTCGTCCAGG-1
clust_6_CAAGATCCACGACGAA-1
clust_6_CTCATTAAGCGTCTAT-1
clust_7_ACGATGTGTTTGACTG-1
clust_7_CTACCCACACATAACC-1
clust_7_CCTTACGAGAACTCGG-1
clust_7_CGTGTAATCTTTAGGG-1
clust_4_TGGCTGGGTAGGCATG-1
clust_7_TGGCGCAGTGCGGTAA-1
clust_2_CCGGTAGTCAAGCCTA-1
clust_2_TTATGCTCAATGAATG-1
clust_3_AAGGAGCGTTTAGGAA-1
clust_3_GCATGATTCGCCGTGA-1
clust_0_GTCCTCAAGATATACG-1
clust_3_CTTTGCGAGTACATGA-1
clust_1_CAGTAACGTGTGAAAT-1
clust_6_CCCAATCGTTCATGGT-1
clust_6_CCTACCATCCACGCAG-1
clust_6_GTGCTTCCACAGGCCT-1
clust_4_TAAACCGCACGTAAGG-1
clust_5_GATCGATTCGTCACGG-1
clust_3_CCTACCAGTGTAAGTA-1
clust_4_GGCAATTTCCAGAAGG-1

**Supplemental Figure 2: Heatmaps of the number of sites where individual cells have shared variant calls**
We calculated the number of sites where each pairwise comparison of individual cells had enough read depth to call variants and either of the cells had a non-reference call. These heatmaps show the number of called variants that would be informative for calculating genetic distance for each sample between pairs of cells. For this calculation we allowed variant calls from sites with as few as two reads remaining after PCR deduplication.

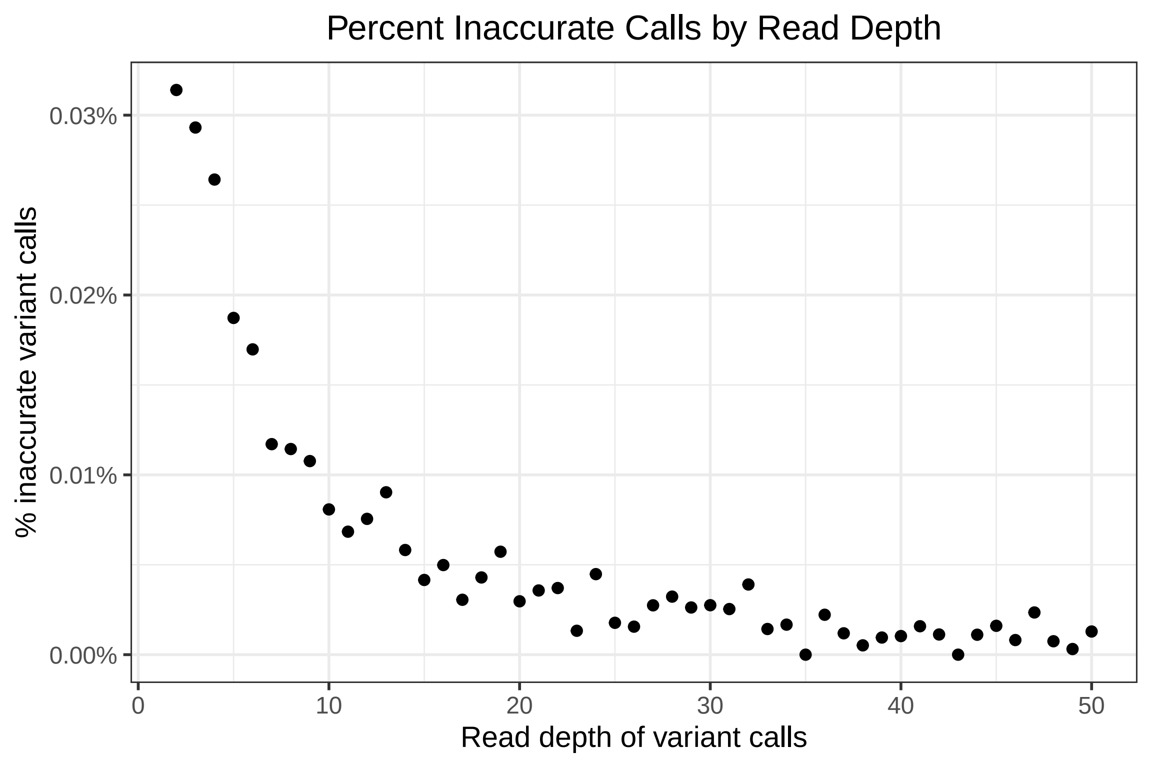

Percent Inaccurate Calls by Read Depth
% inaccurate variant calls
0.03%
0.02%
0.01%
0.00%
0
10
20
30
40
50
Read depth of variant calls

**Supplemental Figure 3: Estimated rates of inaccurately called variants in 10x Flex data**

We called variants on 10x Flex data, where probes matching the reference genome are sequenced instead of the sample RNAs. Any sequence not matching the reference represents an error. We calculated the percent of total covered sites with variant calls which gives an estimate of false positive rates for variant calling.

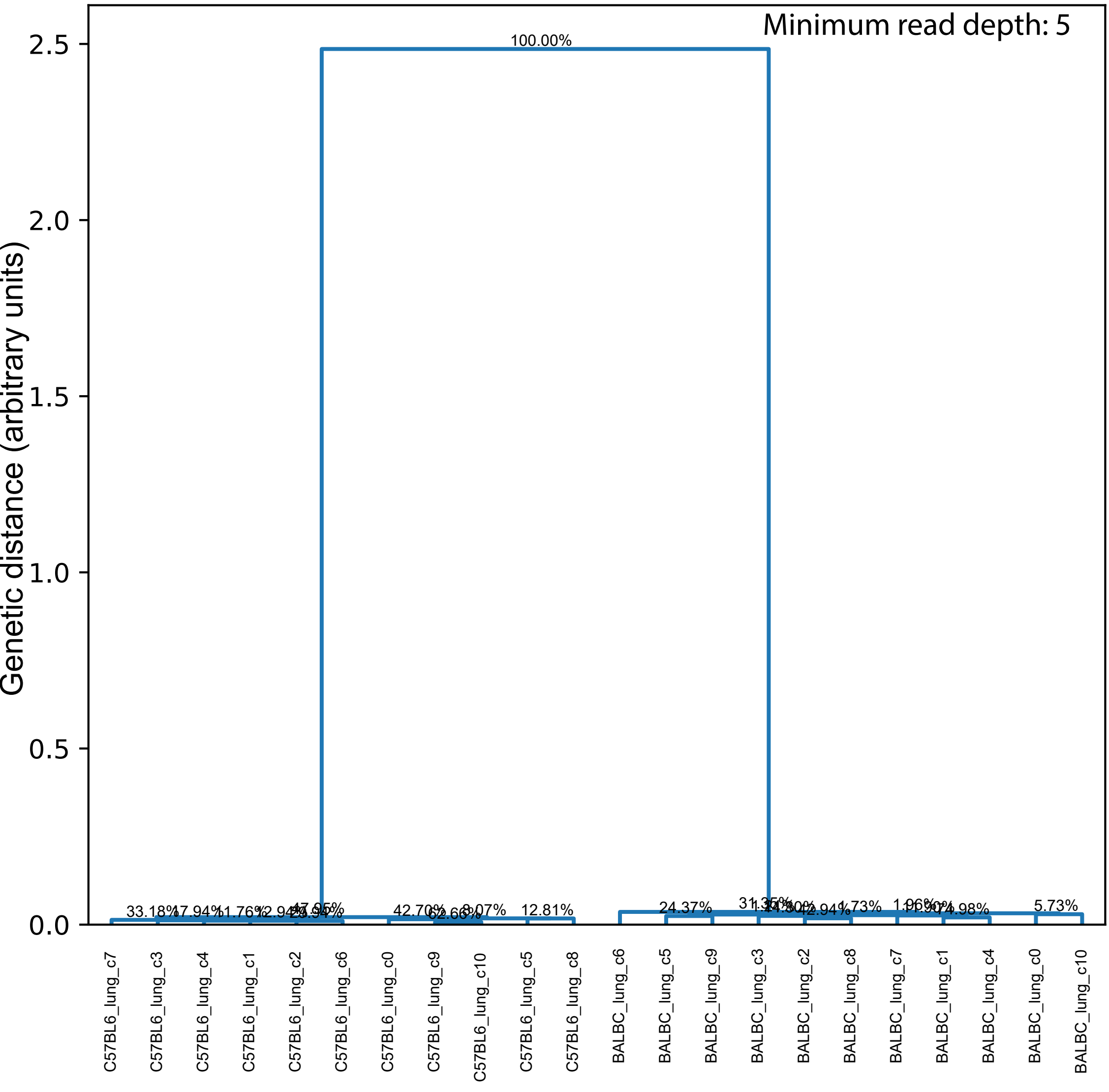

Minimum read depth: 5
100.00%
Genetic distance (arbitrary units)
2.5
2.0
1.5
1.0
0.5
0.0
33.18%
42.70%
62.66%
12.81%
24.37%
5.73%
C57BL6_lung_c7
C57BL6_lung_c3
C57BL6_lung_c4
C57BL6_lung_c1
C57BL6_lung_c2
C57BL6_lung_c6
C57BL6_lung_c0
C57BL6_lung_c9
C57BL6_lung_c10
C57BL6_lung_c5
C57BL6_lung_c8
BALBC_lung_c6
BALBC_lung_c5
BALBC_lung_c9
BALBC_lung_c3
BALBC_lung_c2
BALBC_lung_c8
BALBC_lung_c7
BALBC_lung_c1
BALBC_lung_c4
BALBC_lung_c0
BALBC_lung_c10

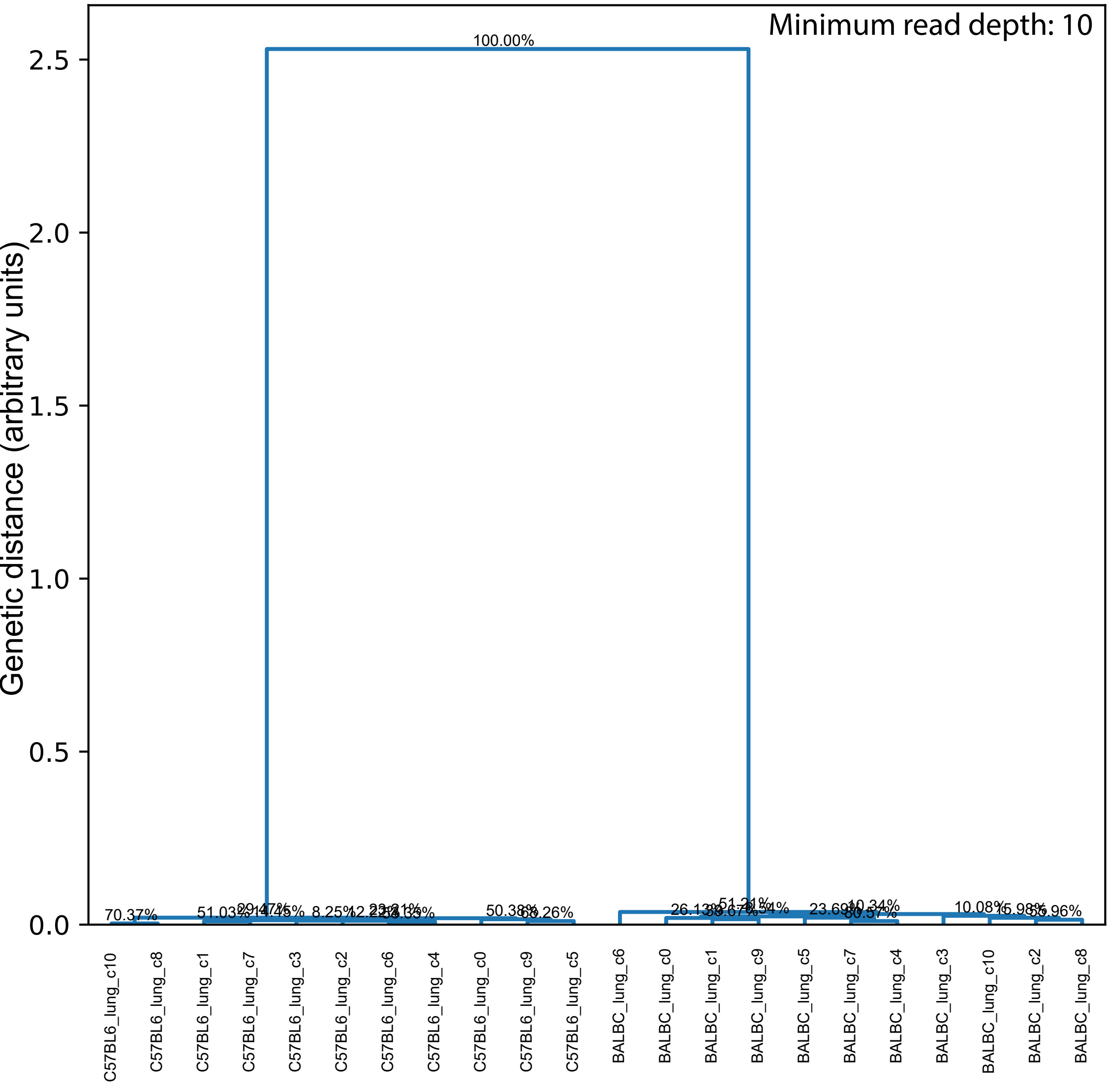
Minimum read depth: 10
100.00%
Genetic distance (arbitrary units)
2.5
2.0
1.5
1.0
0.5
0.0
70.37%
51.03%
8.25%
50.38%
10.34%
10.08%
C57BL6_lung_c10
C57BL6_lung_c8
C57BL6_lung_c1
C57BL6_lung_c7
C57BL6_lung_c3
C57BL6_lung_c2
C57BL6_lung_c6
C57BL6_lung_c4
C57BL6_lung_c0
C57BL6_lung_c9
C57BL6_lung_c5
BALBC_lung_c6
BALBC_lung_c0
BALBC_lung_c1
BALBC_lung_c9
BALBC_lung_c5
BALBC_lung_c7
BALBC_lung_c4
BALBC_lung_c3
BALBC_lung_c10
BALBC_lung_c2
BALBC_lung_c8

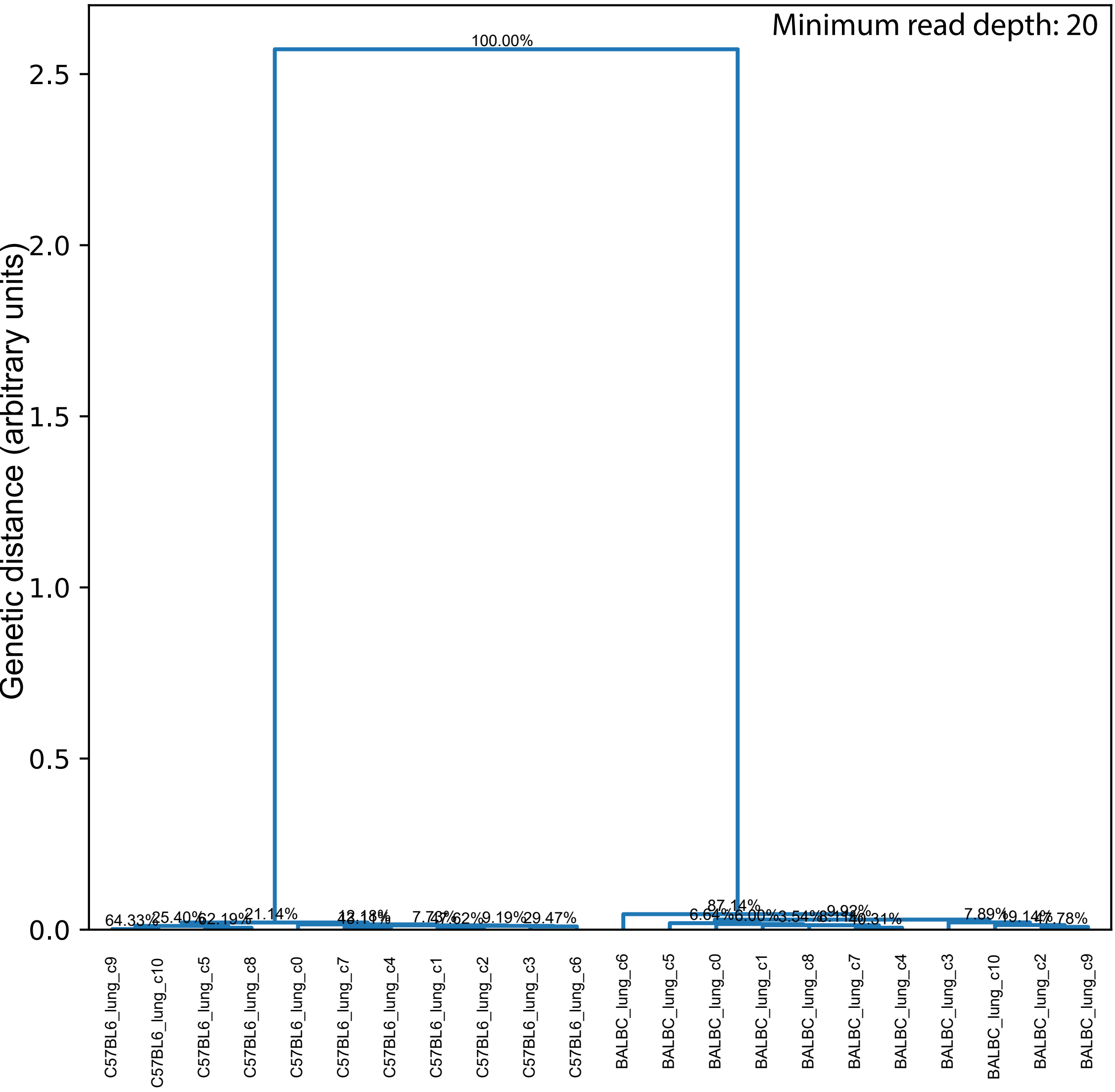

Minimum read depth: 20
100.00%
Genetic distance (arbitrary units)
2.5
2.0
1.5
1.0
0.5
0.0
64.33%
25.40%
62.19%
21.14%
9.19%
29.47%
87.14%
40.31%
7.89%
C57BL6_lung_c9
C57BL6_lung_c10
C57BL6_lung_c5
C57BL6_lung_c8
C57BL6_lung_c0
C57BL6_lung_c7
C57BL6_lung_c4
C57BL6_lung_c1
C57BL6_lung_c2
C57BL6_lung_c3
C57BL6_lung_c6
BALBC_lung_c6
BALBC_lung_c5
BALBC_lung_c0
BALBC_lung_c1
BALBC_lung_c8
BALBC_lung_c7
BALBC_lung_c4
BALBC_lung_c3
BALBC_lung_c10
BALBC_lung_c2
BALBC_lung_c9

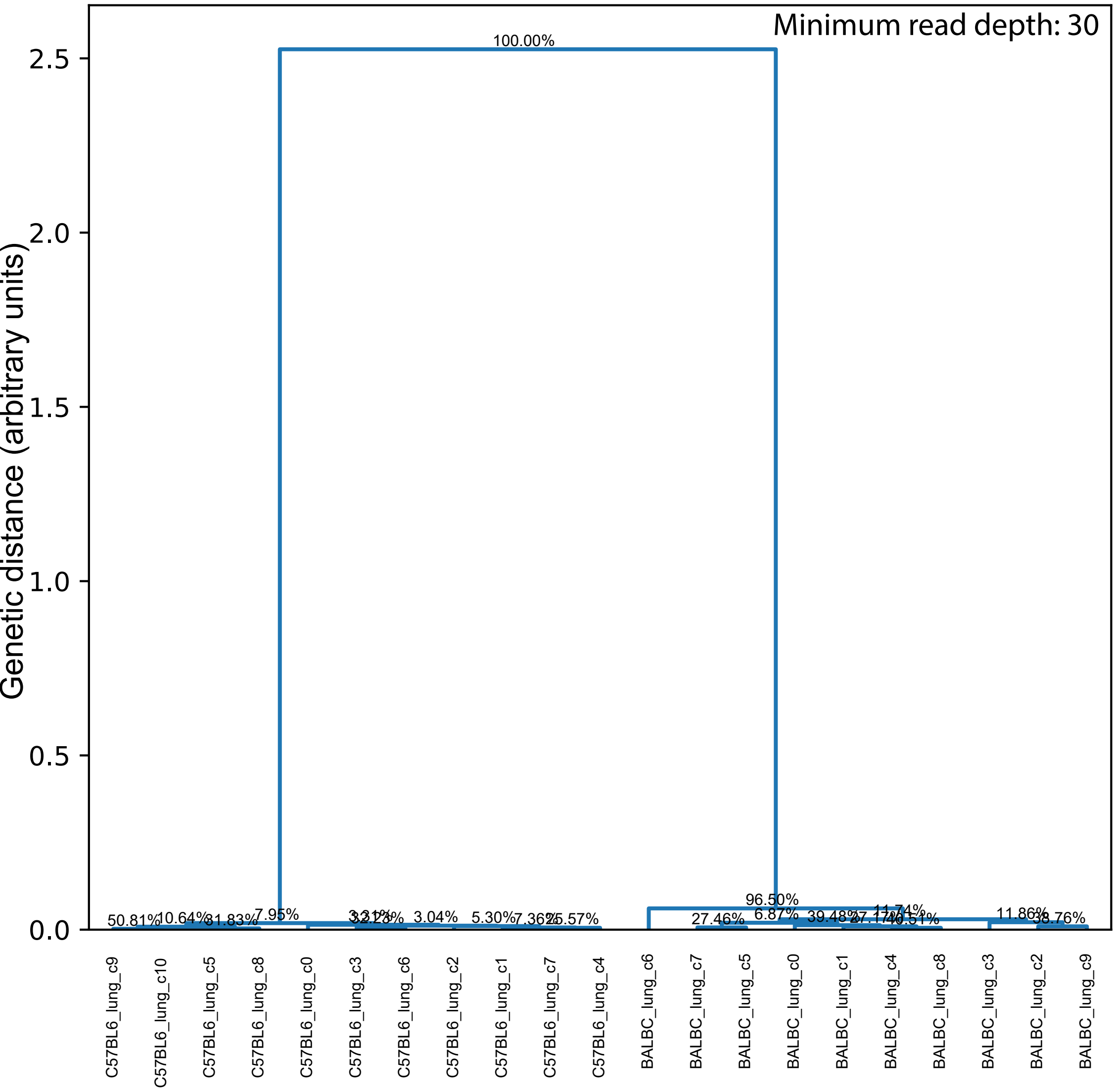

Minimum read depth: 30
Genetic distance (arbitrary units)
0.0
0.5
1.0
1.5
2.0
2.5
100.00%
96.50%
50.81%
10.64%
81.83%
7.95%
3.04%
5.30%
27.46%
6.87%
39.48%
11.74%
40.51%
11.86%
38.76%
C57BL6_lung_c9
C57BL6_lung_c10
C57BL6_lung_c5
C57BL6_lung_c8
C57BL6_lung_c0
C57BL6_lung_c3
C57BL6_lung_c6
C57BL6_lung_c2
C57BL6_lung_c1
C57BL6_lung_c7
C57BL6_lung_c4
BALBC_lung_c6
BALBC_lung_c7
BALBC_lung_c5
BALBC_lung_c0
BALBC_lung_c1
BALBC_lung_c4
BALBC_lung_c8
BALBC_lung_c3
BALBC_lung_c2
BALBC_lung_c9

**Supplemental Figure 4: Hierarchical clustering trees for comparison of C57BL/6 and BALB/c**

We used scanBit to compare C57BL/6 and BALB/c samples to test if variant calls could distinguish the genetically distinct mouse strains. This was tested using minimum read depth thresholds of 5, 10, 20 and 30. Bootstrap values for each node of the tree are provided.

# G0005

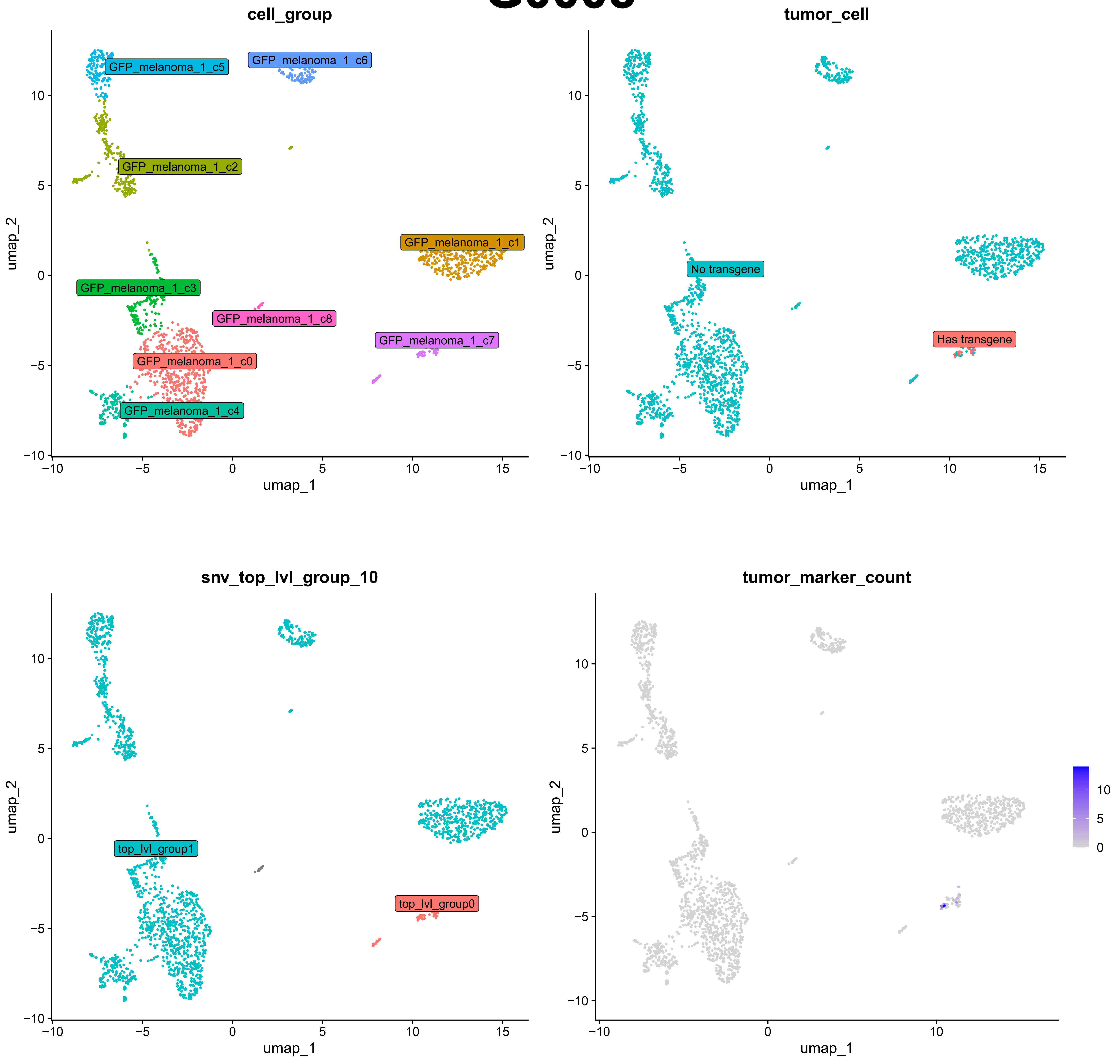

cell_group
GFP_melanoma_1_c5
GFP_melanoma_1_c6
GFP_melanoma_1_c2
GFP_melanoma_1_c1
GFP_melanoma_1_c3
GFP_melanoma_1_c8
GFP_melanoma_1_c7
GFP_melanoma_1_c0
GFP_melanoma_1_c4
tumor_cell
No transgene
Has transgene
snv_top_lvl_group_10
top_lvl_group1
top_lvl_group0
tumor_marker_count
umap_1
umap_2


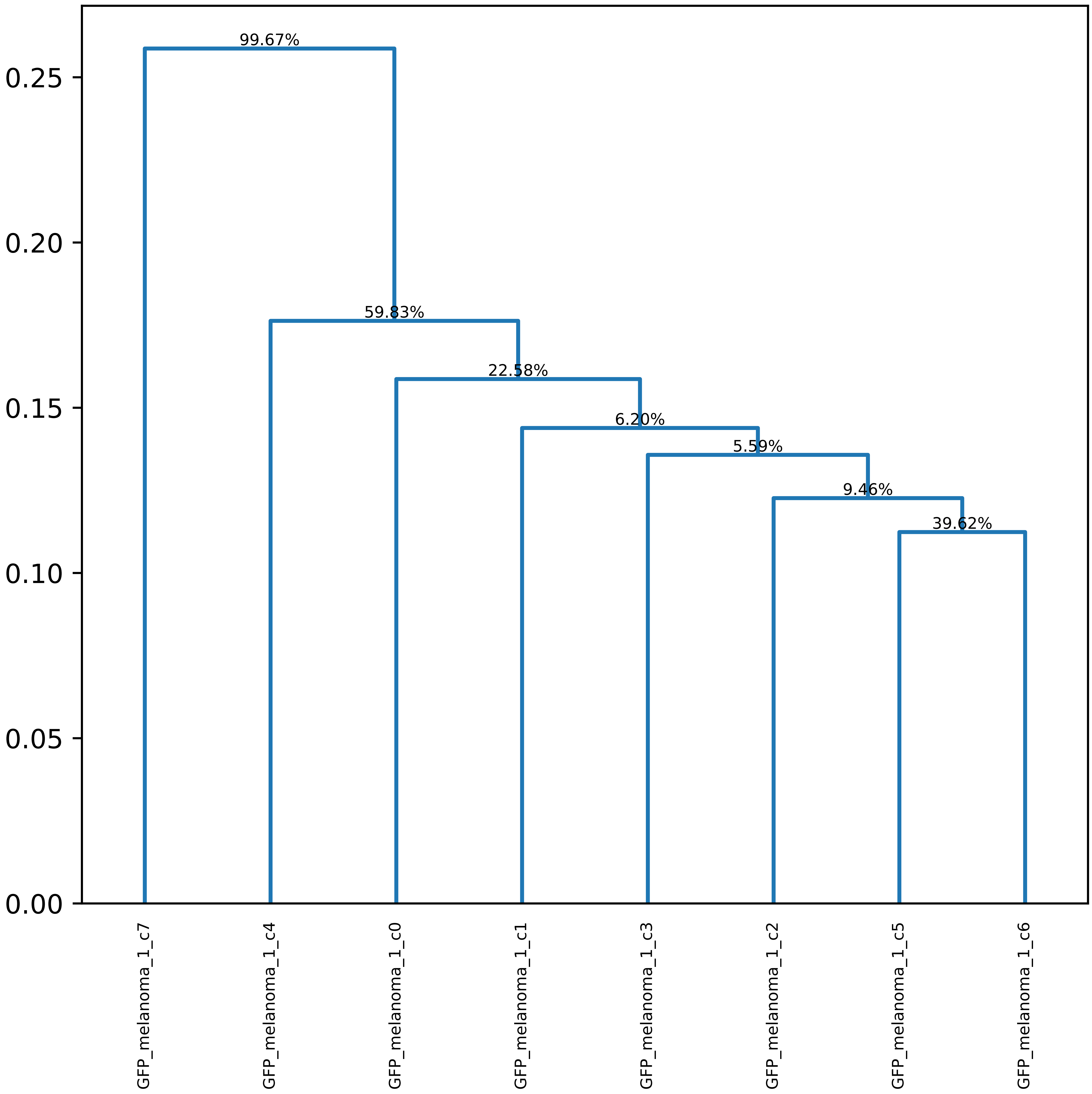

99.67%
59.83%
22.58%
6.20%
5.59%
9.46%
39.62%
0.25
0.20
0.15
0.10
0.05
0.00
GFP_melanoma_1_c7
GFP_melanoma_1_c4
GFP_melanoma_1_c0
GFP_melanoma_1_c1
GFP_melanoma_1_c3
GFP_melanoma_1_c2
GFP_melanoma_1_c5
GFP_melanoma_1_c6

# G0006

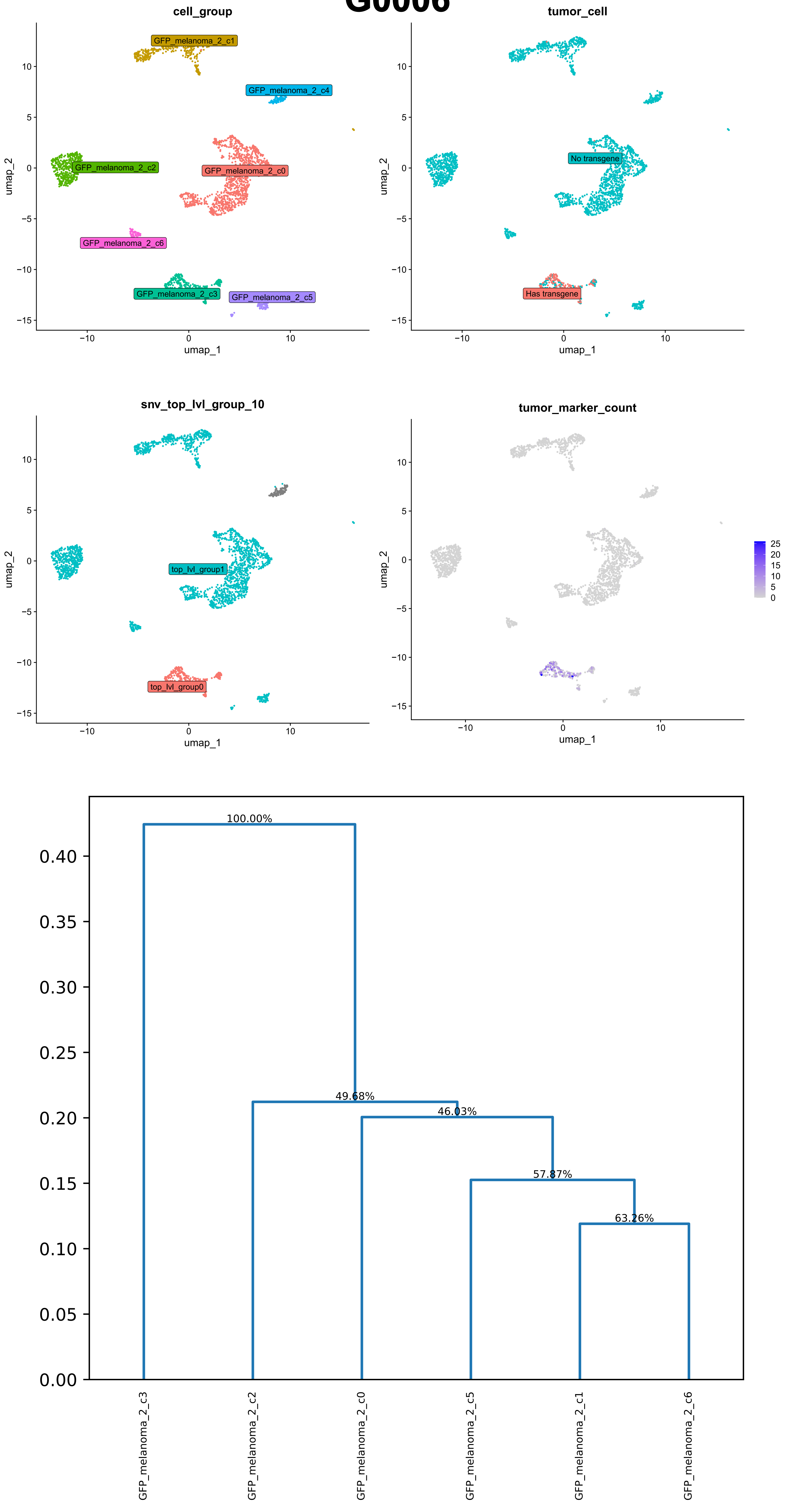
cell_group
GFP_melanoma_2_c1
GFP_melanoma_2_c4
GFP_melanoma_2_c2
GFP_melanoma_2_c0
GFP_melanoma_2_c6
GFP_melanoma_2_c3
GFP_melanoma_2_c5
umap_1
umap_2
tumor_cell
No transgene
Has transgene
snv_top_lvl_group_10
top_lvl_group1
top_lvl_group0
tumor_marker_count
100.00%
49.68%
46.03%
57.87%
63.26%
GFP_melanoma_2_c3
GFP_melanoma_2_c2
GFP_melanoma_2_c0
GFP_melanoma_2_c5
GFP_melanoma_2_c1
GFP_melanoma_2_c6

# G0007

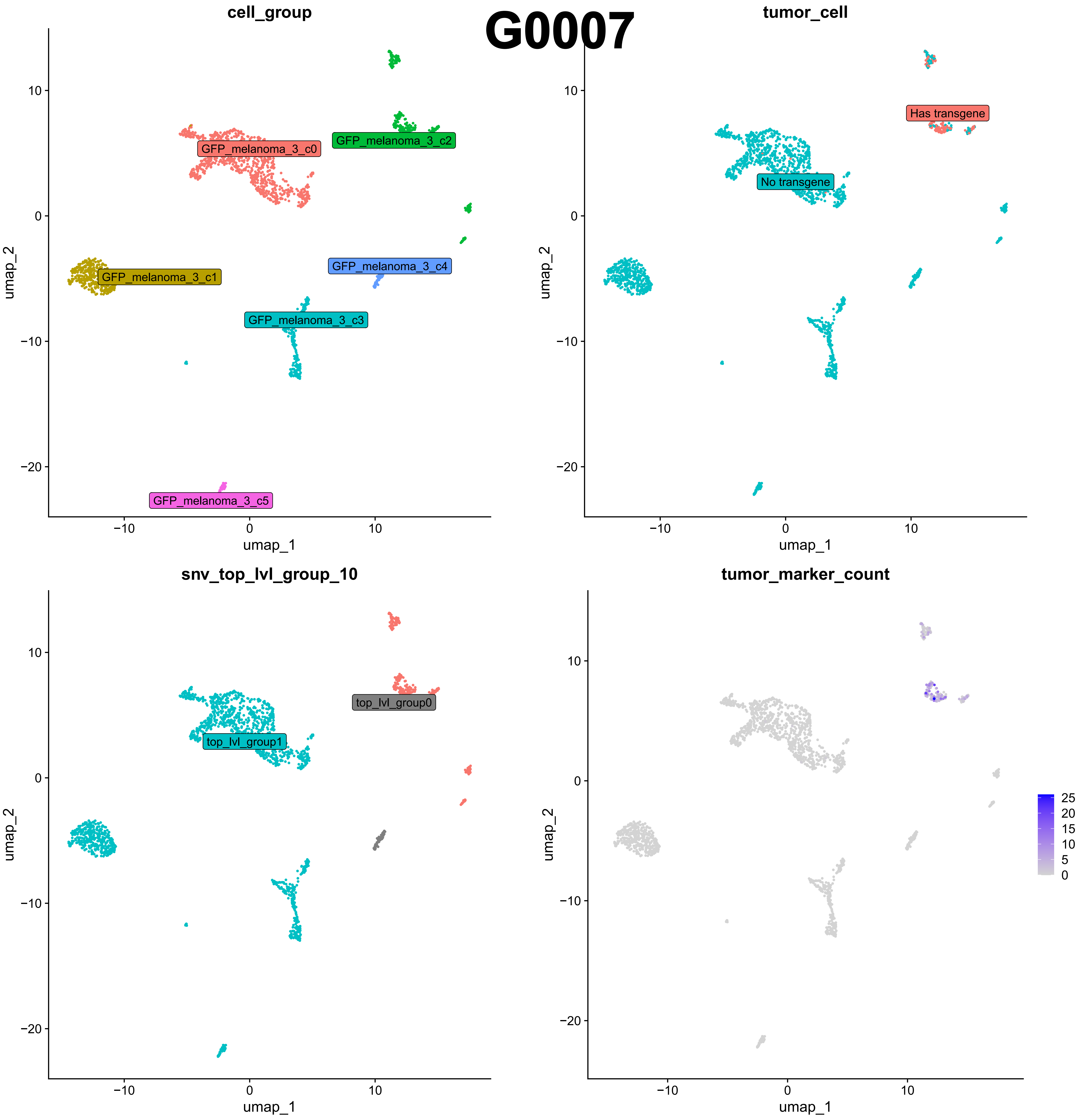

cell_group
GFP_melanoma_3_c0
GFP_melanoma_3_c1
GFP_melanoma_3_c2
GFP_melanoma_3_c3
GFP_melanoma_3_c4
GFP_melanoma_3_c5
tumor_cell
Has transgene
No transgene
snv_top_lvl_group_10
top_lvl_group0
top_lvl_group1
tumor_marker_count
25
20
15
10
5
0
umap_1
umap_2


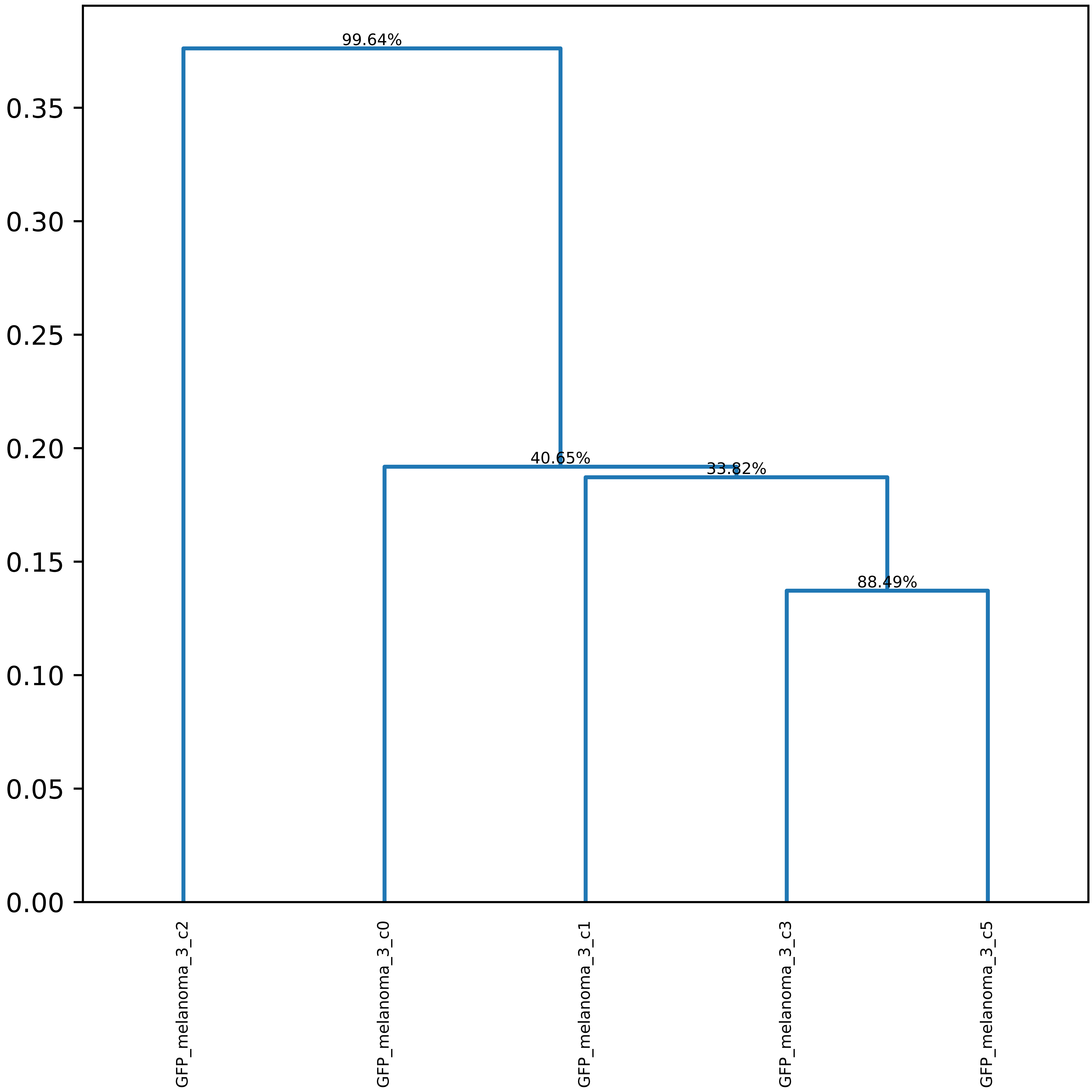

99.64%
40.65%
33.82%
88.49%
0.35
0.30
0.25
0.20
0.15
0.10
0.05
0.00
GFP_melanoma_3_c2
GFP_melanoma_3_c0
GFP_melanoma_3_c1
GFP_melanoma_3_c3
GFP_melanoma_3_c5

# G0008

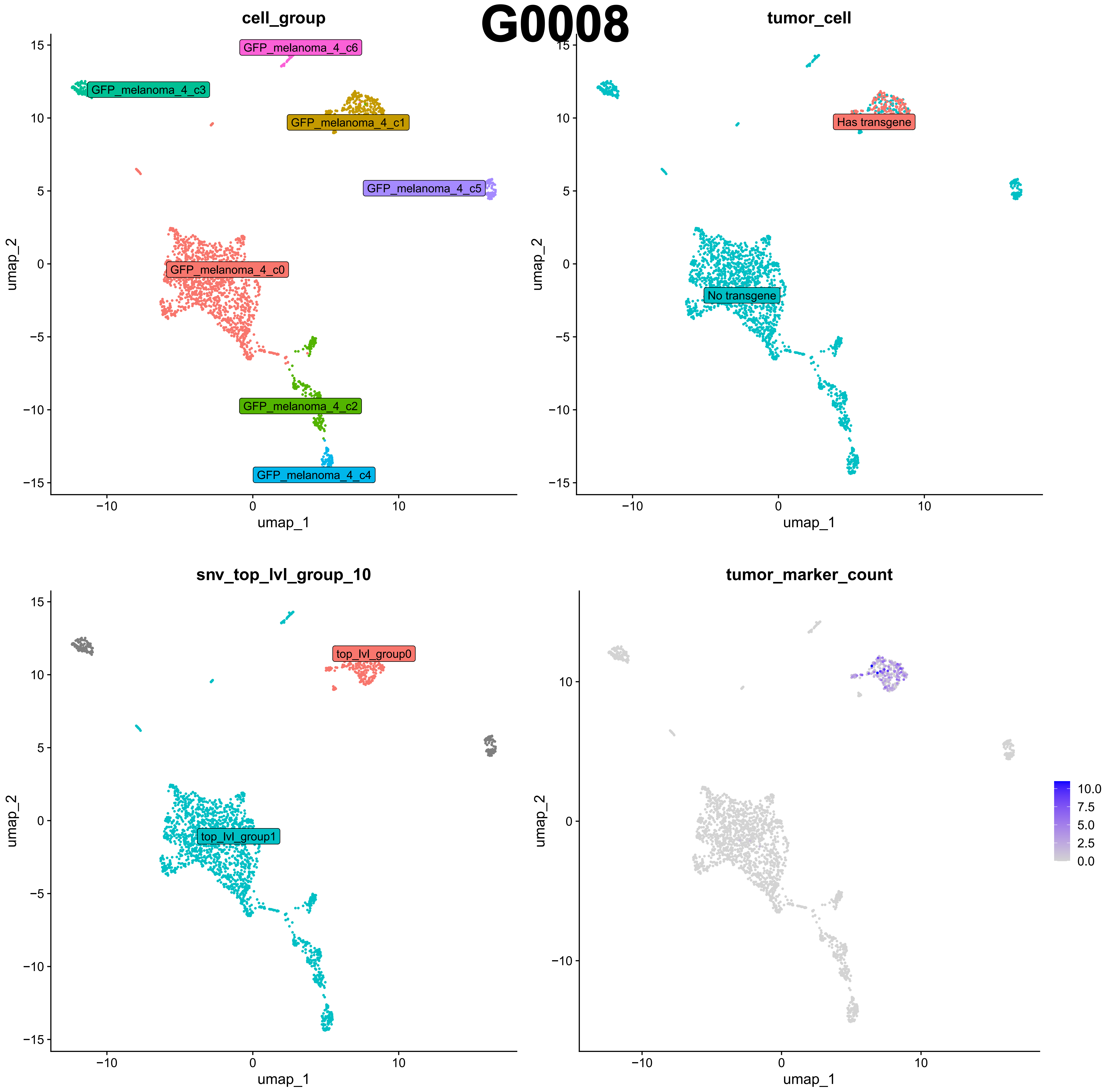
cell_group
GFP_melanoma_4_c6
GFP_melanoma_4_c3
GFP_melanoma_4_c1
GFP_melanoma_4_c5
GFP_melanoma_4_c0
GFP_melanoma_4_c2
GFP_melanoma_4_c4
tumor_cell
Has transgene
No transgene
snv_top_lvl_group_10
top_lvl_group0
top_lvl_group1
tumor_marker_count
10.0
7.5
5.0
2.5
0.0
umap_1
umap_2

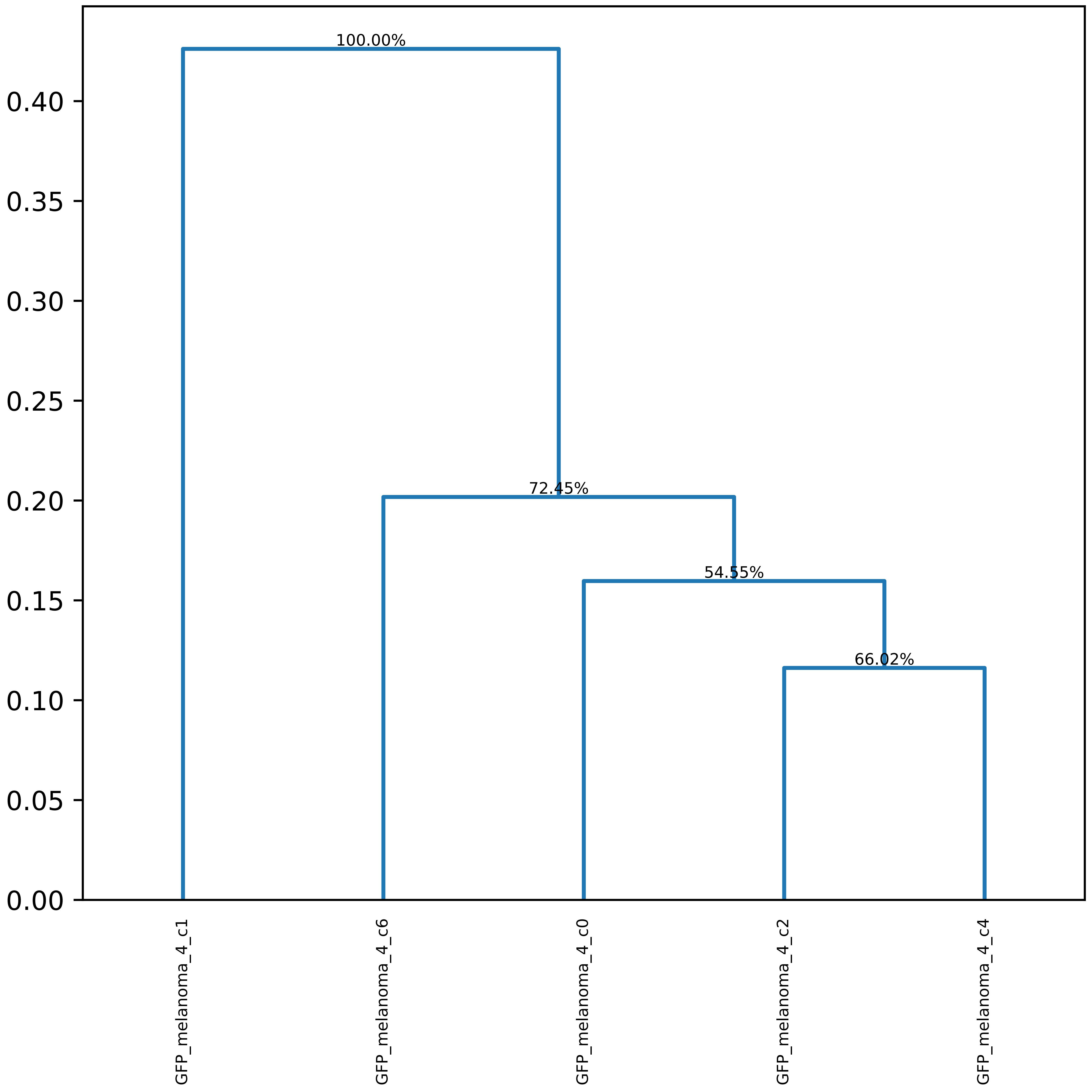
100.00%
72.45%
54.55%
66.02%
0.40
0.35
0.30
0.25
0.20
0.15
0.10
0.05
0.00
GFP_melanoma_4_c1
GFP_melanoma_4_c6
GFP_melanoma_4_c0
GFP_melanoma_4_c2
GFP_melanoma_4_c4

# G0009

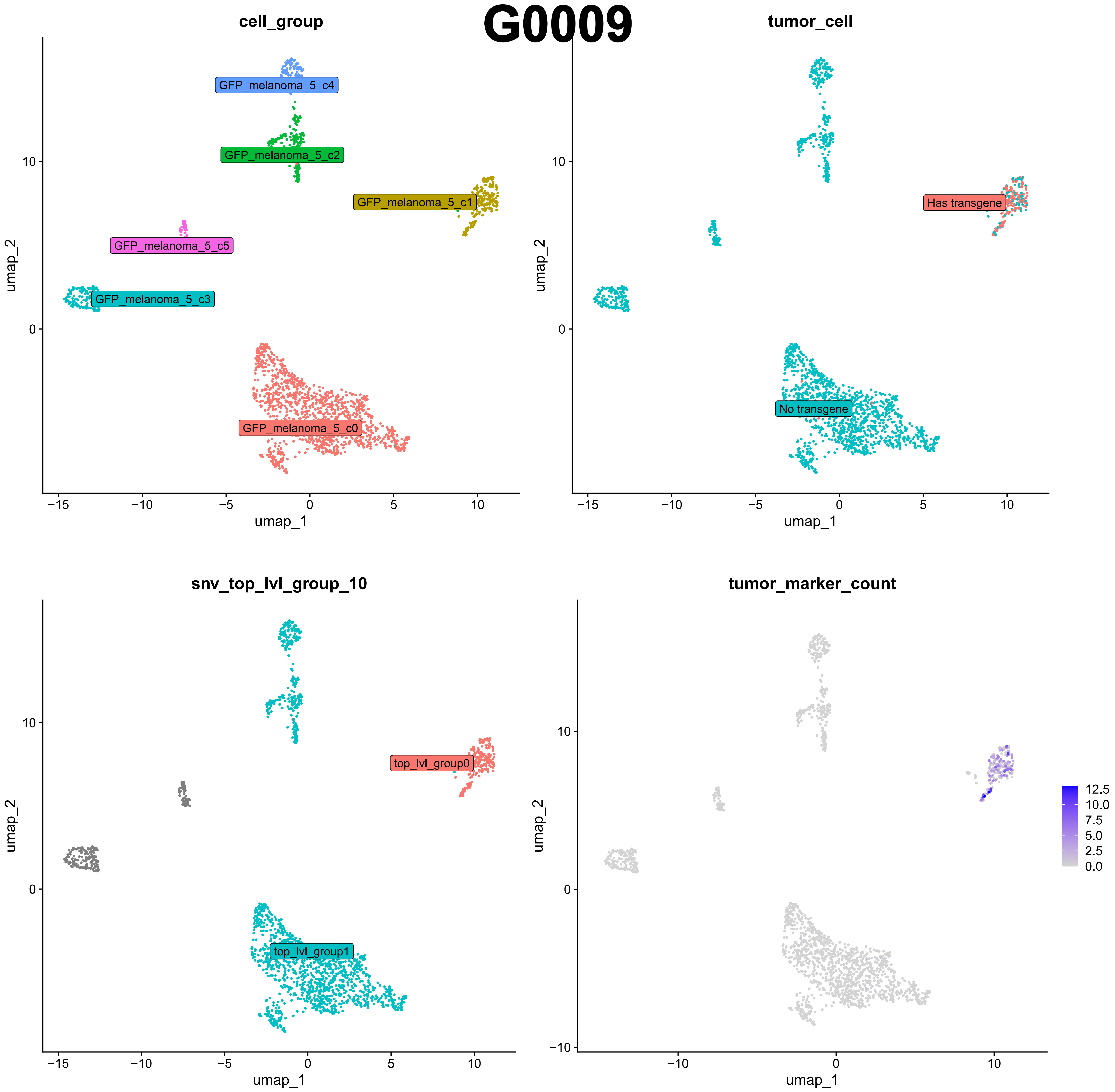
cell_group
GFP_melanoma_5_c4
GFP_melanoma_5_c2
GFP_melanoma_5_c1
GFP_melanoma_5_c5
GFP_melanoma_5_c3
GFP_melanoma_5_c0
umap_1
umap_2
tumor_cell
Has transgene
No transgene
snv_top_lvl_group_10
top_lvl_group0
top_lvl_group1
tumor_marker_count
12.5
10.0
7.5
5.0
2.5
0.0

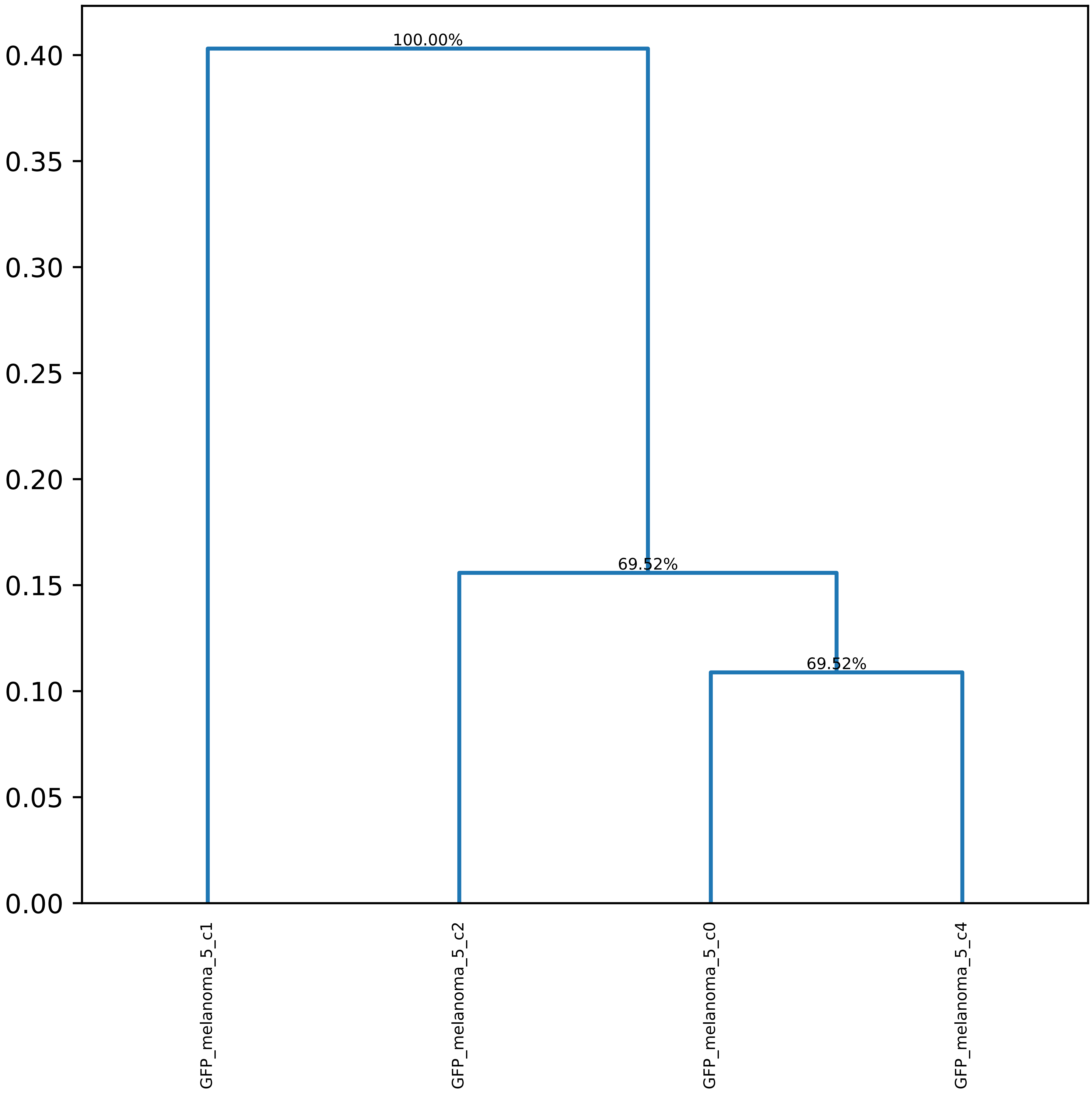
100.00%
69.52%
69.52%
0.40
0.35
0.30
0.25
0.20
0.15
0.10
0.05
0.00
GFP_melanoma_5_c1
GFP_melanoma_5_c2
GFP_melanoma_5_c0
GFP_melanoma_5_c4

# G0010

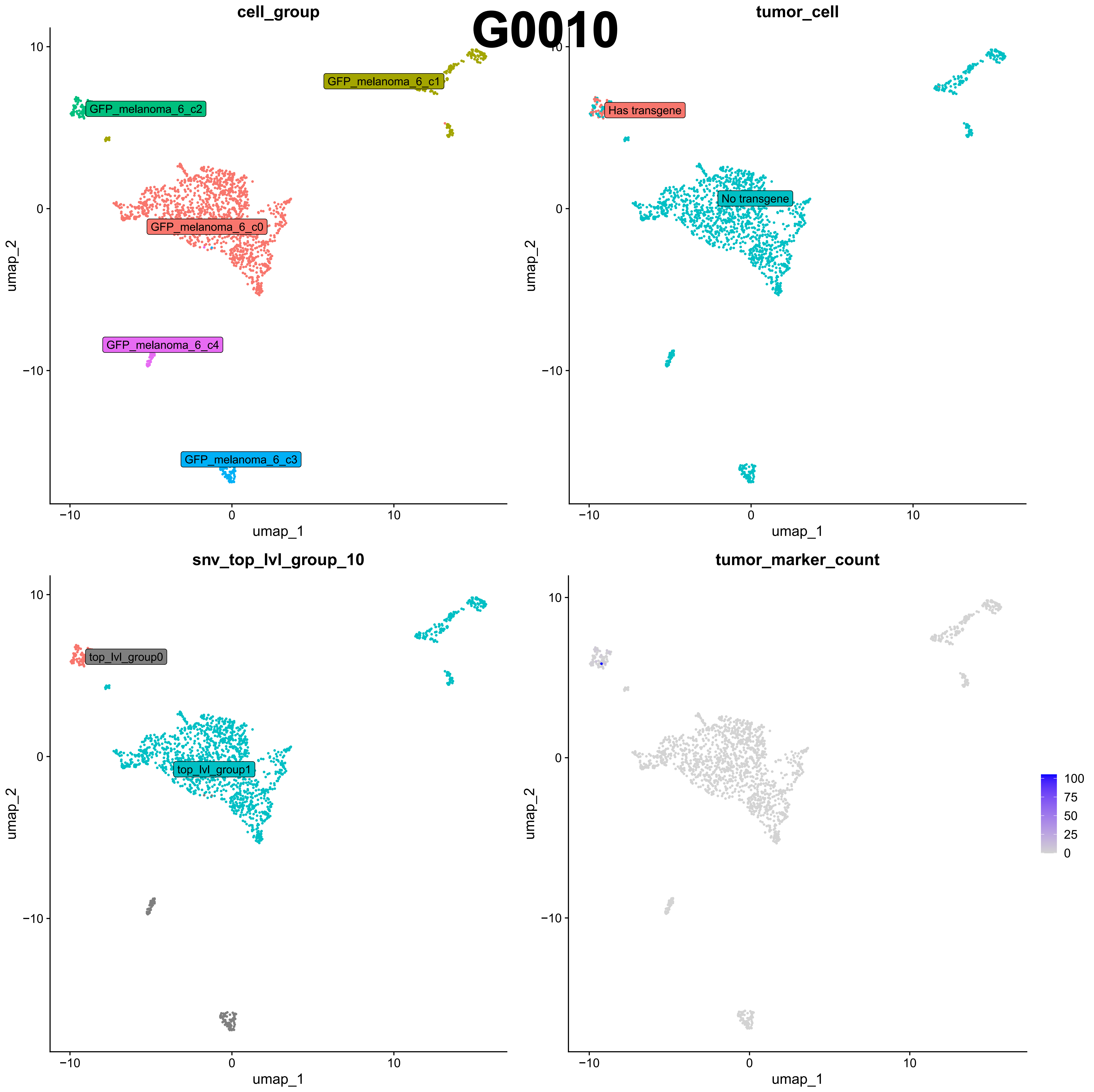

cell_group
GFP_melanoma_6_c2
GFP_melanoma_6_c1
GFP_melanoma_6_c0
GFP_melanoma_6_c4
GFP_melanoma_6_c3
umap_1
umap_2
tumor_cell
Has transgene
No transgene
umap_1
umap_2
snv_top_lvl_group_10
top_lvl_group0
top_lvl_group1
umap_1
umap_2
tumor_marker_count
100
75
50
25
0
umap_1
umap_2


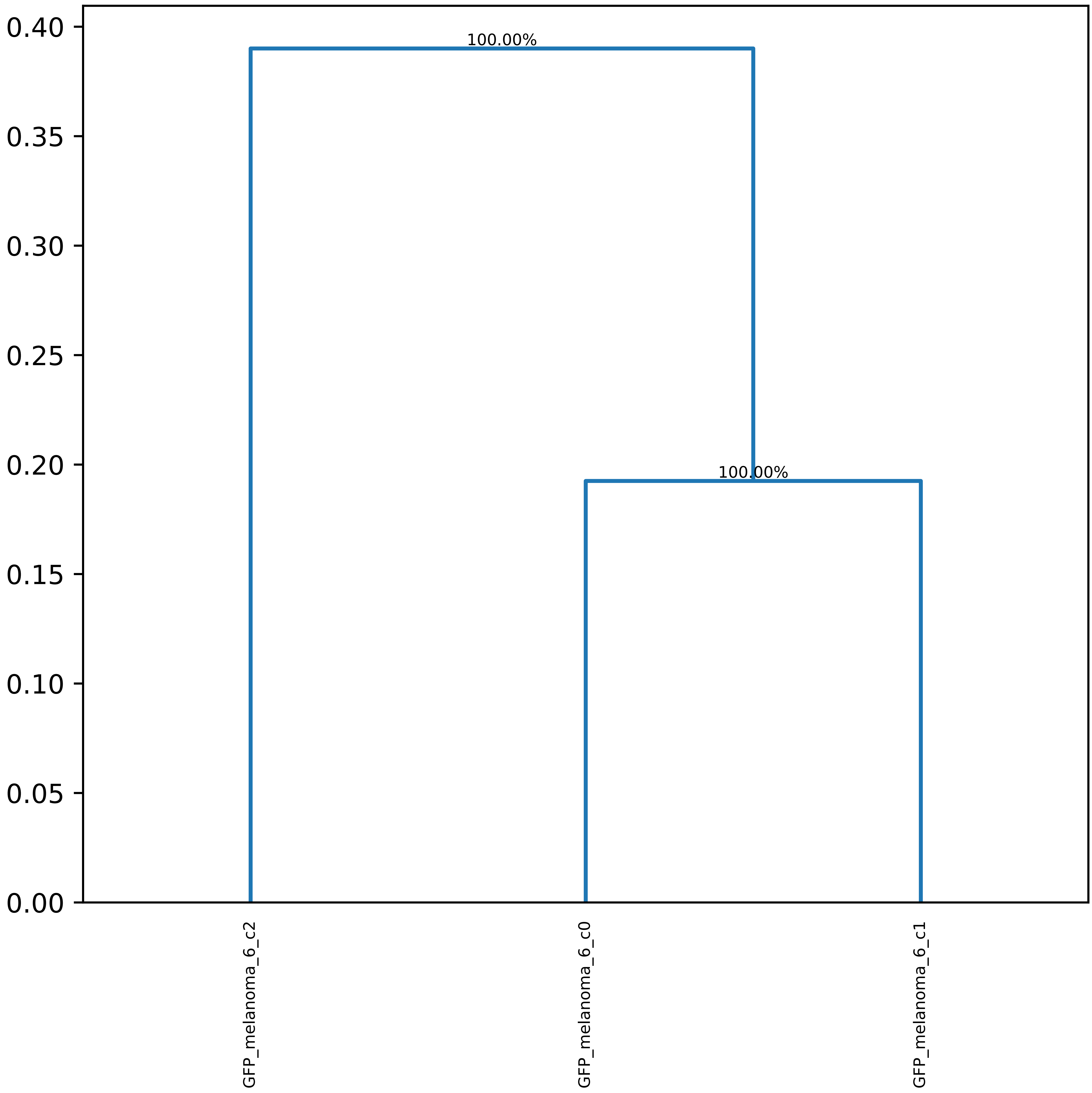

100.00%
100.00%
0.40
0.35
0.30
0.25
0.20
0.15
0.10
0.05
0.00
GFP_melanoma_6_c2
GFP_melanoma_6_c0
GFP_melanoma_6_c1

# G0011

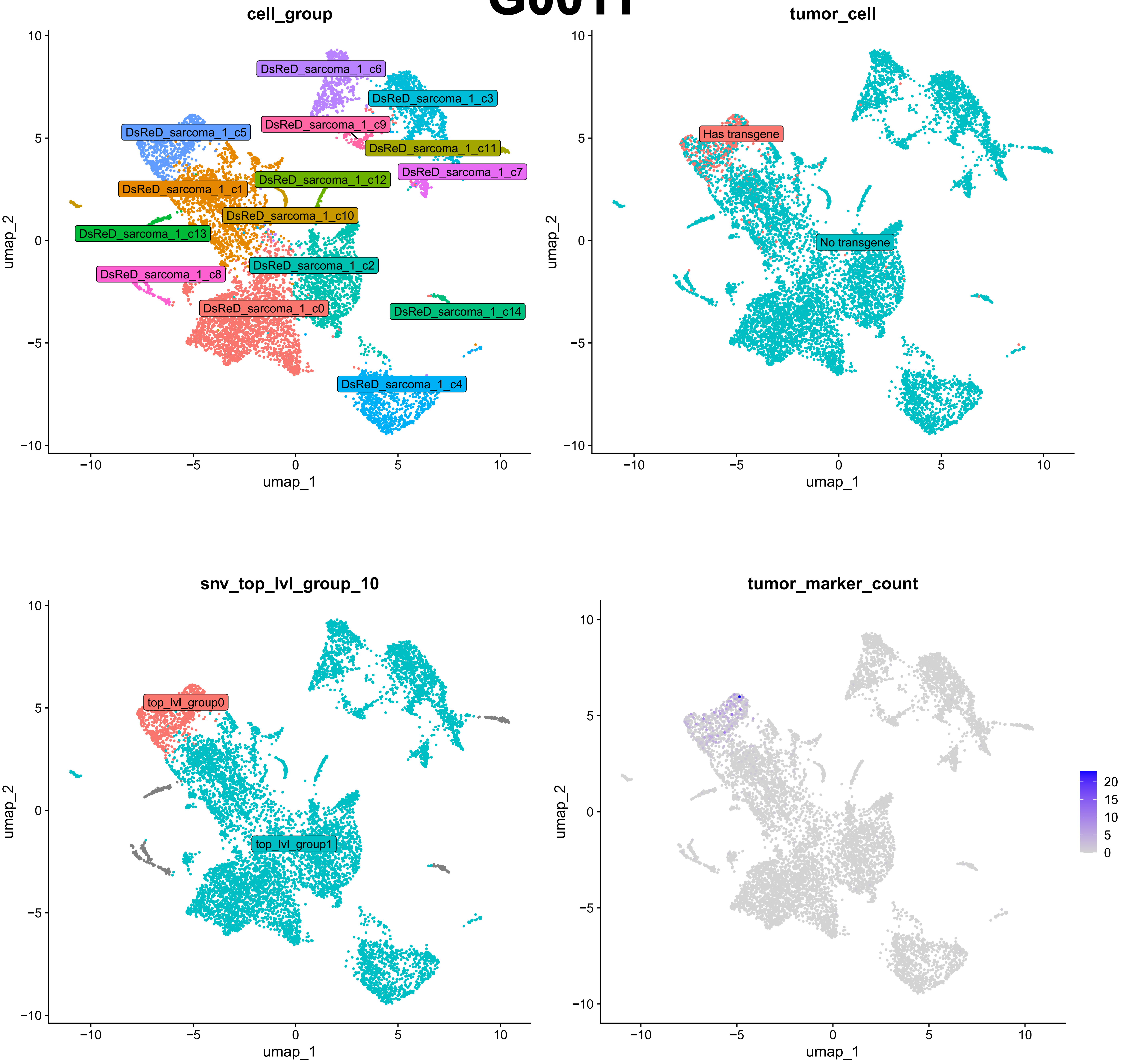
cell_group
DsReD_sarcoma_1_c6
DsReD_sarcoma_1_c3
DsReD_sarcoma_1_c5
DsReD_sarcoma_1_c9
DsReD_sarcoma_1_c11
DsReD_sarcoma_1_c7
DsReD_sarcoma_1_c1
DsReD_sarcoma_1_c12
DsReD_sarcoma_1_c10
DsReD_sarcoma_1_c13
DsReD_sarcoma_1_c2
DsReD_sarcoma_1_c8
DsReD_sarcoma_1_c0
DsReD_sarcoma_1_c14
DsReD_sarcoma_1_c4
tumor_cell
Has transgene
No transgene
snv_top_lvl_group_10
top_lvl_group0
top_lvl_group1
tumor_marker_count
umap_1
umap_2

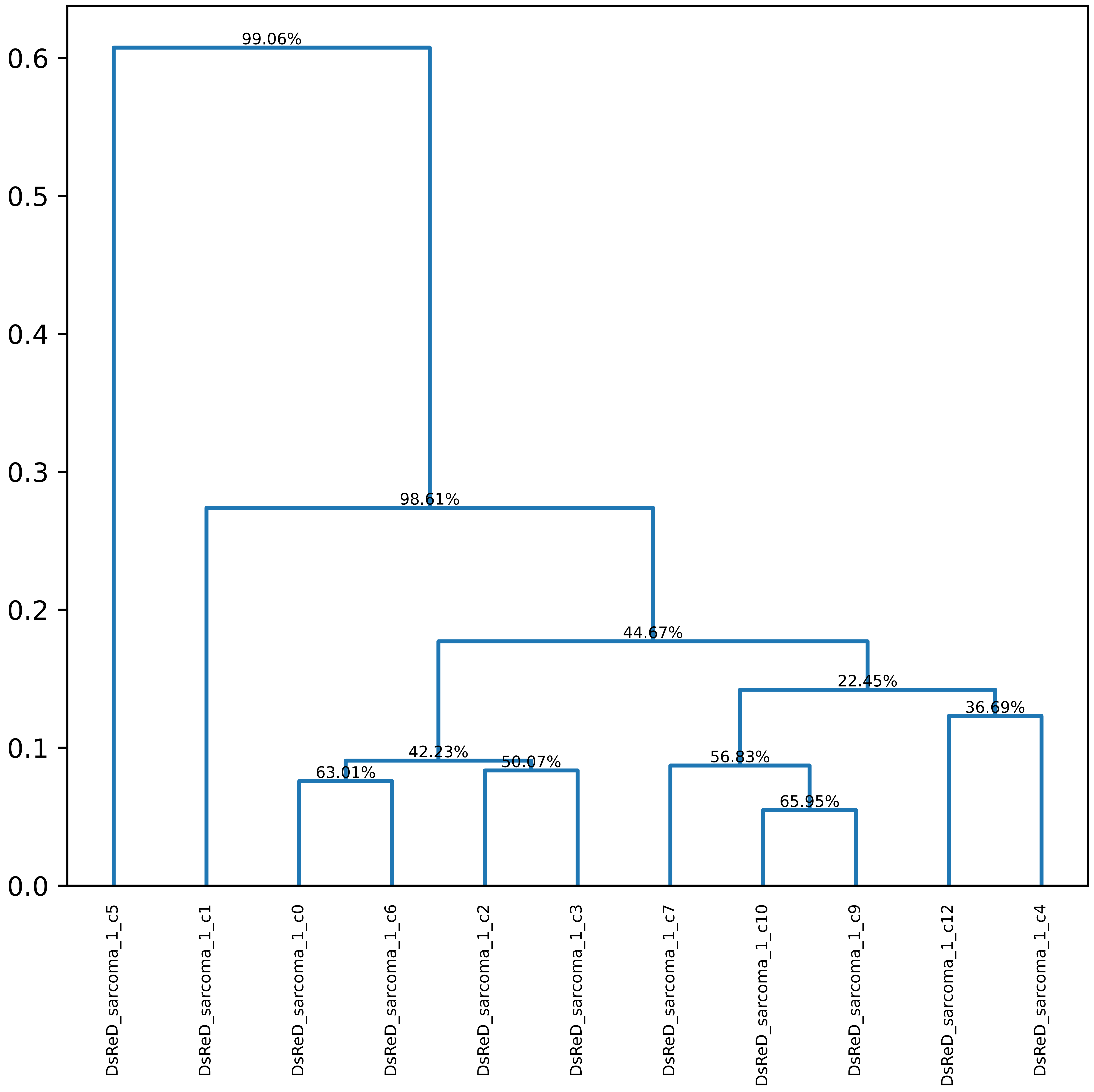
99.06%
98.61%
44.67%
22.45%
36.69%
42.23%
63.01%
50.07%
56.83%
65.95%
DsReD_sarcoma_1_c5
DsReD_sarcoma_1_c1
DsReD_sarcoma_1_c0
DsReD_sarcoma_1_c6
DsReD_sarcoma_1_c2
DsReD_sarcoma_1_c3
DsReD_sarcoma_1_c7
DsReD_sarcoma_1_c10
DsReD_sarcoma_1_c9
DsReD_sarcoma_1_c12
DsReD_sarcoma_1_c4

# G0012

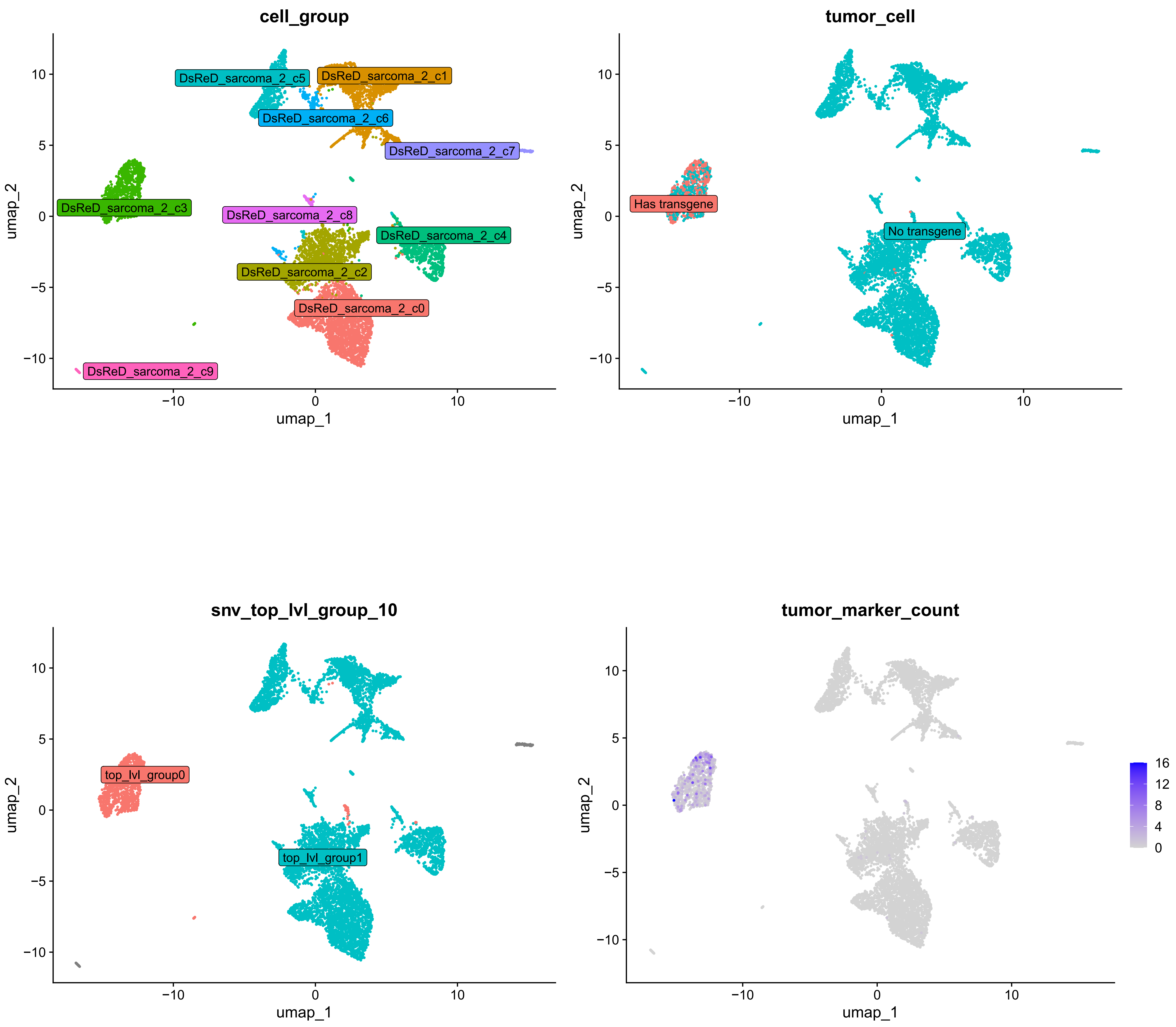

cell_group
DsReD_sarcoma_2_c5
DsReD_sarcoma_2_c1
DsReD_sarcoma_2_c6
DsReD_sarcoma_2_c7
DsReD_sarcoma_2_c3
DsReD_sarcoma_2_c8
DsReD_sarcoma_2_c4
DsReD_sarcoma_2_c2
DsReD_sarcoma_2_c0
DsReD_sarcoma_2_c9
tumor_cell
Has transgene
No transgene
snv_top_lvl_group_10
top_lvl_group0
top_lvl_group1
tumor_marker_count
16
12
8
4
0
umap_1
umap_2
−10
−5
0
5
10


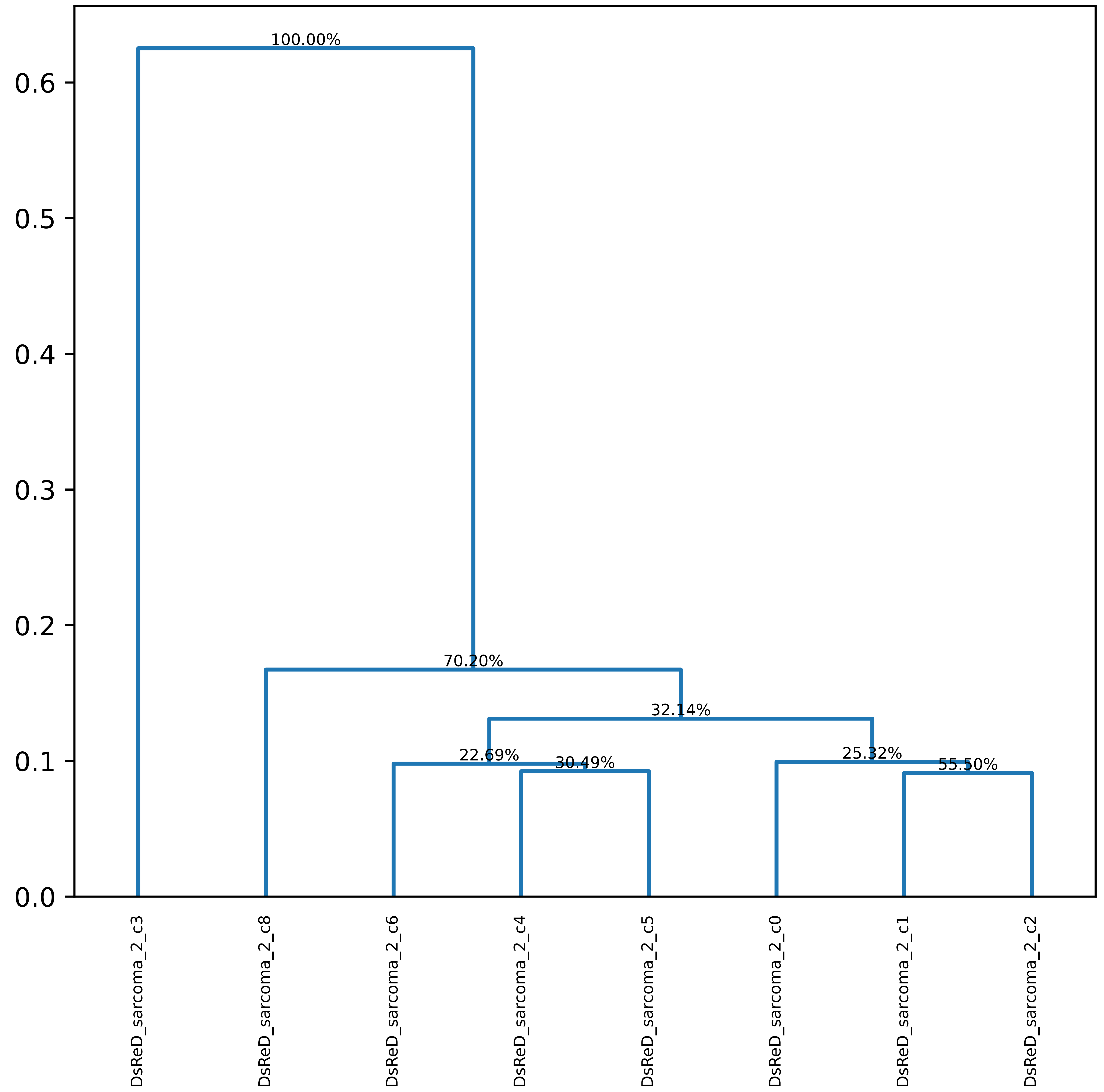

100.00%
70.20%
32.14%
22.69%
30.49%
25.32%
55.50%
0.6
0.5
0.4
0.3
0.2
0.1
0.0
DsReD_sarcoma_2_c3
DsReD_sarcoma_2_c8
DsReD_sarcoma_2_c6
DsReD_sarcoma_2_c4
DsReD_sarcoma_2_c5
DsReD_sarcoma_2_c0
DsReD_sarcoma_2_c1
DsReD_sarcoma_2_c2

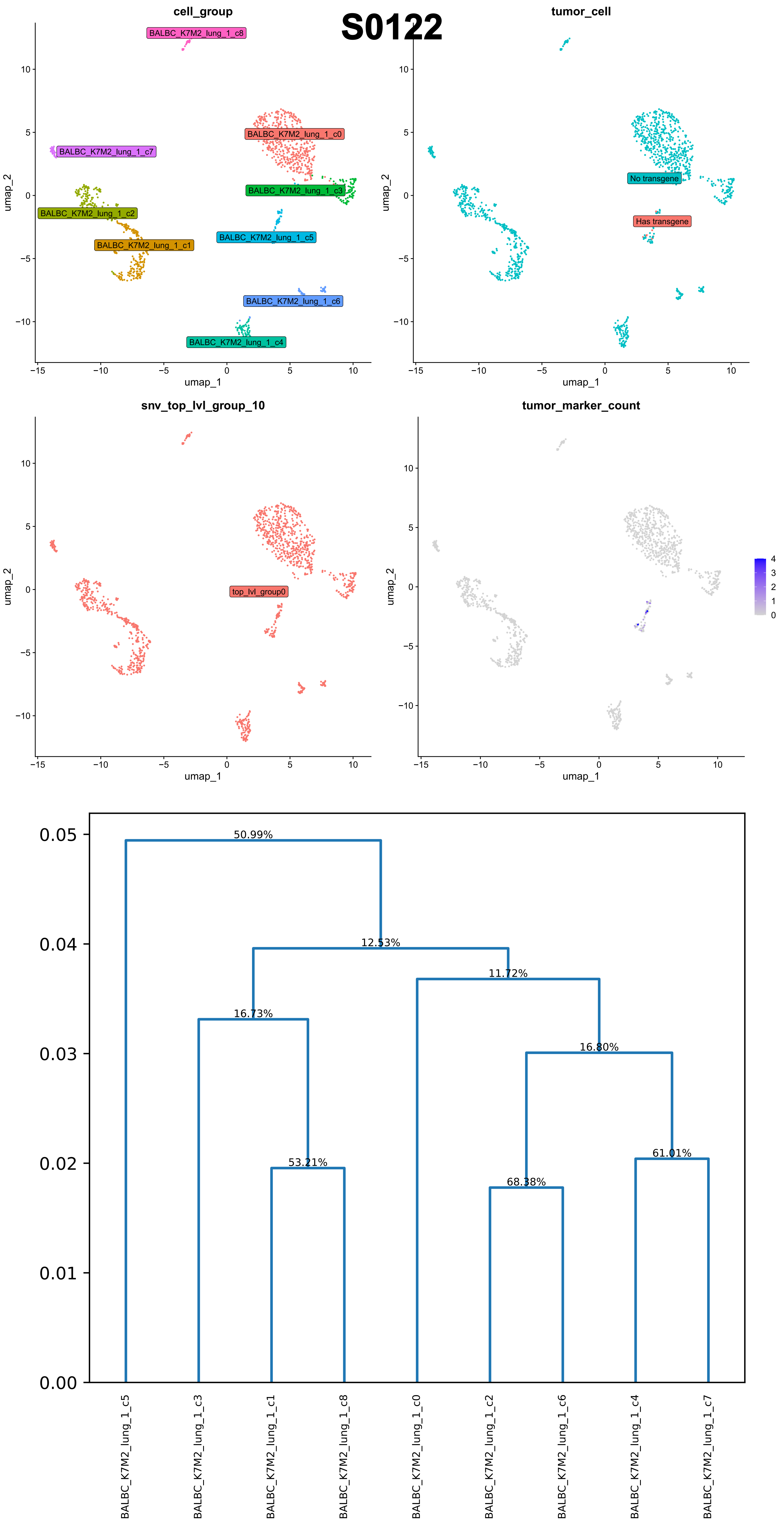
S0122
cell_group
BALBC_K7M2_lung_1_c8
BALBC_K7M2_lung_1_c0
BALBC_K7M2_lung_1_c7
BALBC_K7M2_lung_1_c3
BALBC_K7M2_lung_1_c2
BALBC_K7M2_lung_1_c1
BALBC_K7M2_lung_1_c5
BALBC_K7M2_lung_1_c6
BALBC_K7M2_lung_1_c4
umap_1
umap_2
tumor_cell
No transgene
Has transgene
snv_top_lvl_group_10
top_lvl_group0
tumor_marker_count
50.99%
12.53%
11.72%
16.73%
16.80%
53.21%
68.38%
61.01%
BALBC_K7M2_lung_1_c5
BALBC_K7M2_lung_1_c3
BALBC_K7M2_lung_1_c1
BALBC_K7M2_lung_1_c8
BALBC_K7M2_lung_1_c0
BALBC_K7M2_lung_1_c2
BALBC_K7M2_lung_1_c6
BALBC_K7M2_lung_1_c4
BALBC_K7M2_lung_1_c7

# S0260

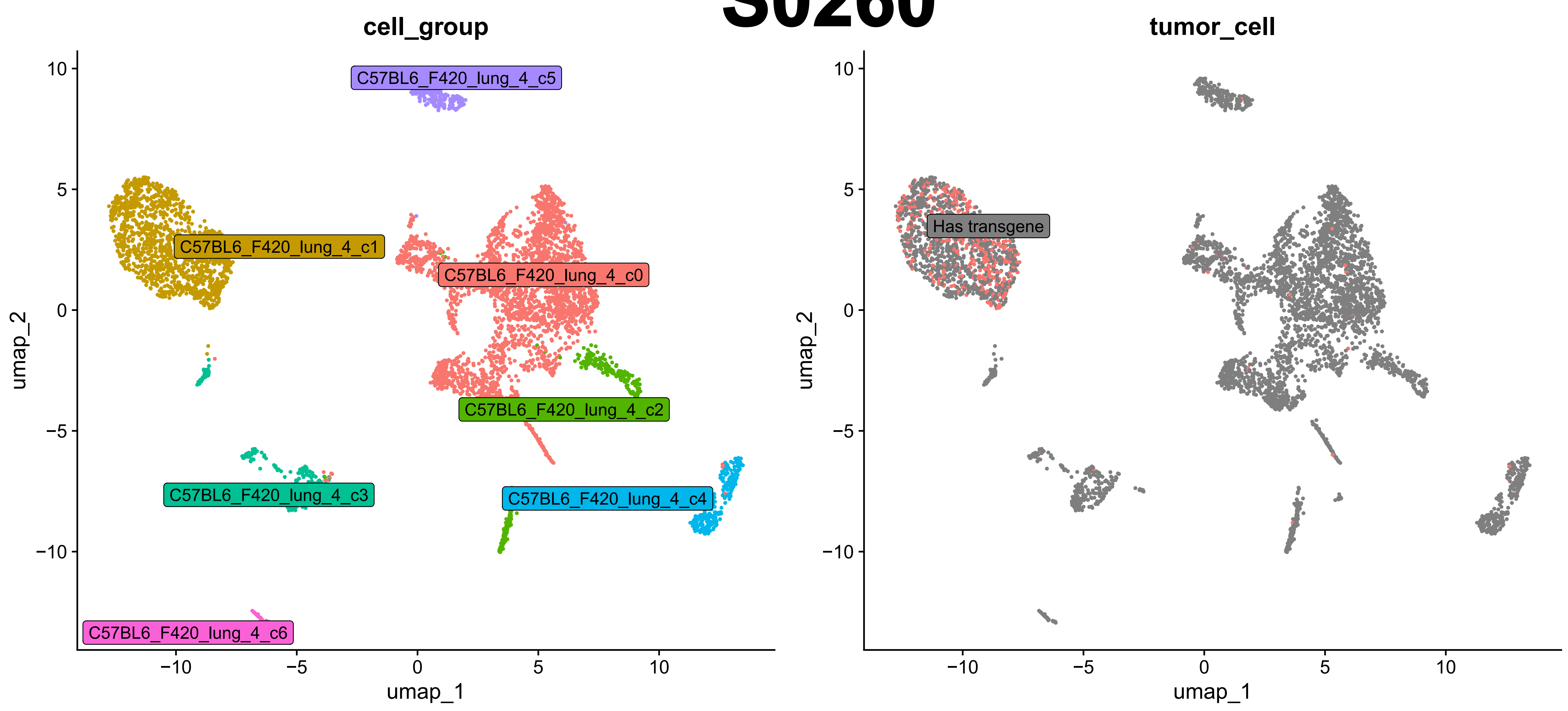

cell_group
C57BL6_F420_lung_4_c5
C57BL6_F420_lung_4_c1
C57BL6_F420_lung_4_c0
C57BL6_F420_lung_4_c2
C57BL6_F420_lung_4_c3
C57BL6_F420_lung_4_c4
C57BL6_F420_lung_4_c6
umap_1
umap_2
tumor_cell
Has transgene
umap_1
umap_2


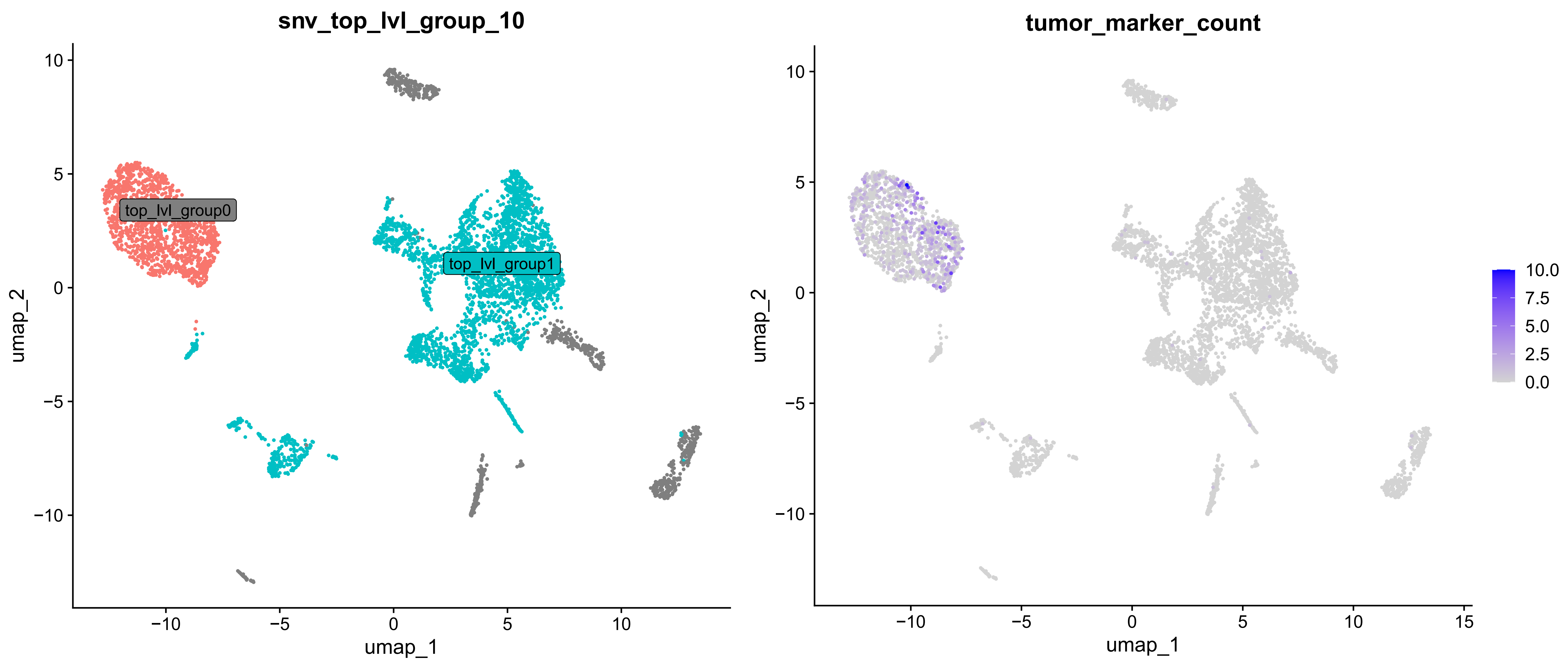

snv_top_lvl_group_10
top_lvl_group0
top_lvl_group1
umap_1
umap_2
tumor_marker_count
10.0
7.5
5.0
2.5
0.0
umap_1
umap_2


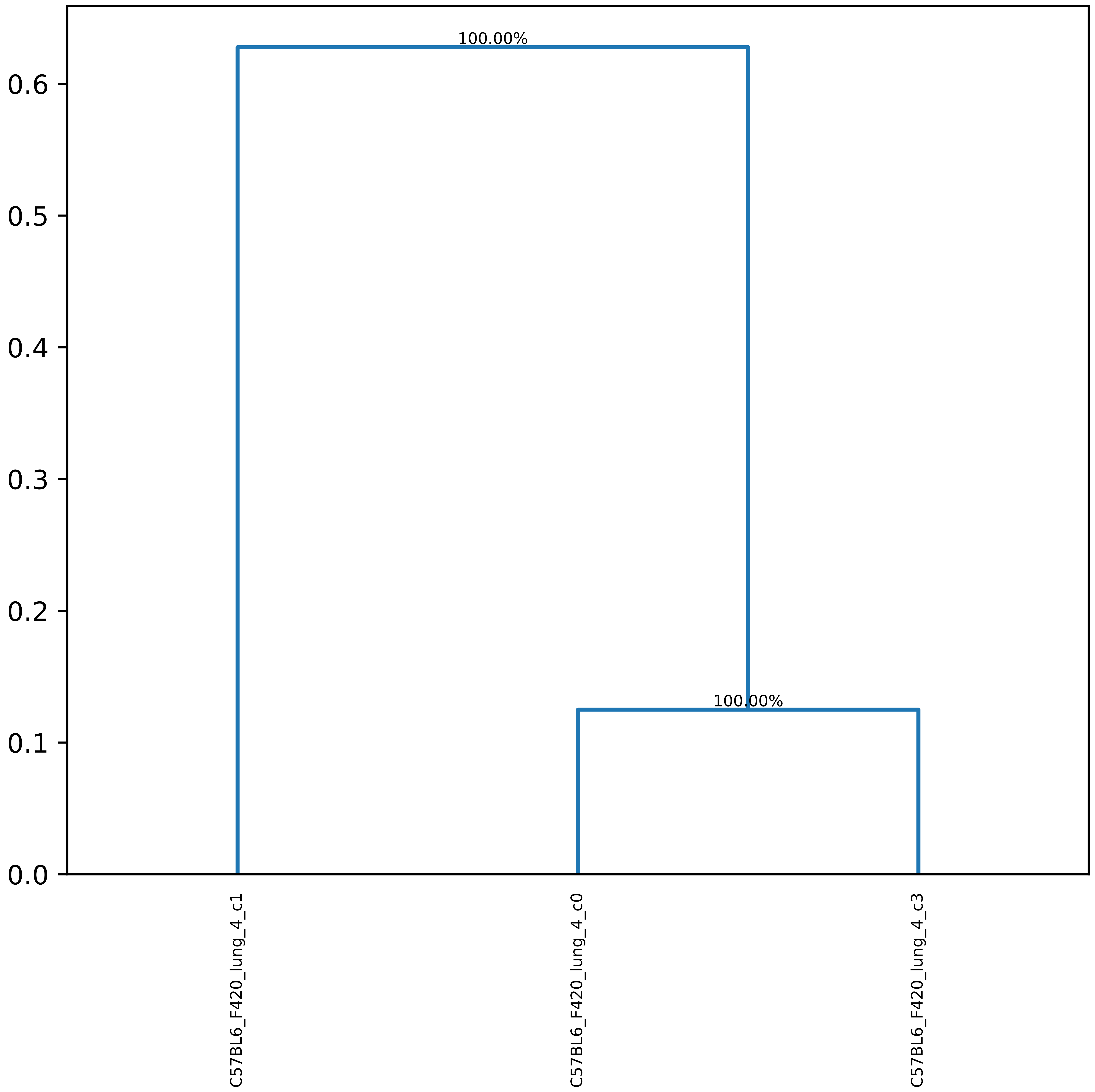

100.00%
100.00%
C57BL6_F420_lung_4_c1
C57BL6_F420_lung_4_c0
C57BL6_F420_lung_4_c3

# S0261

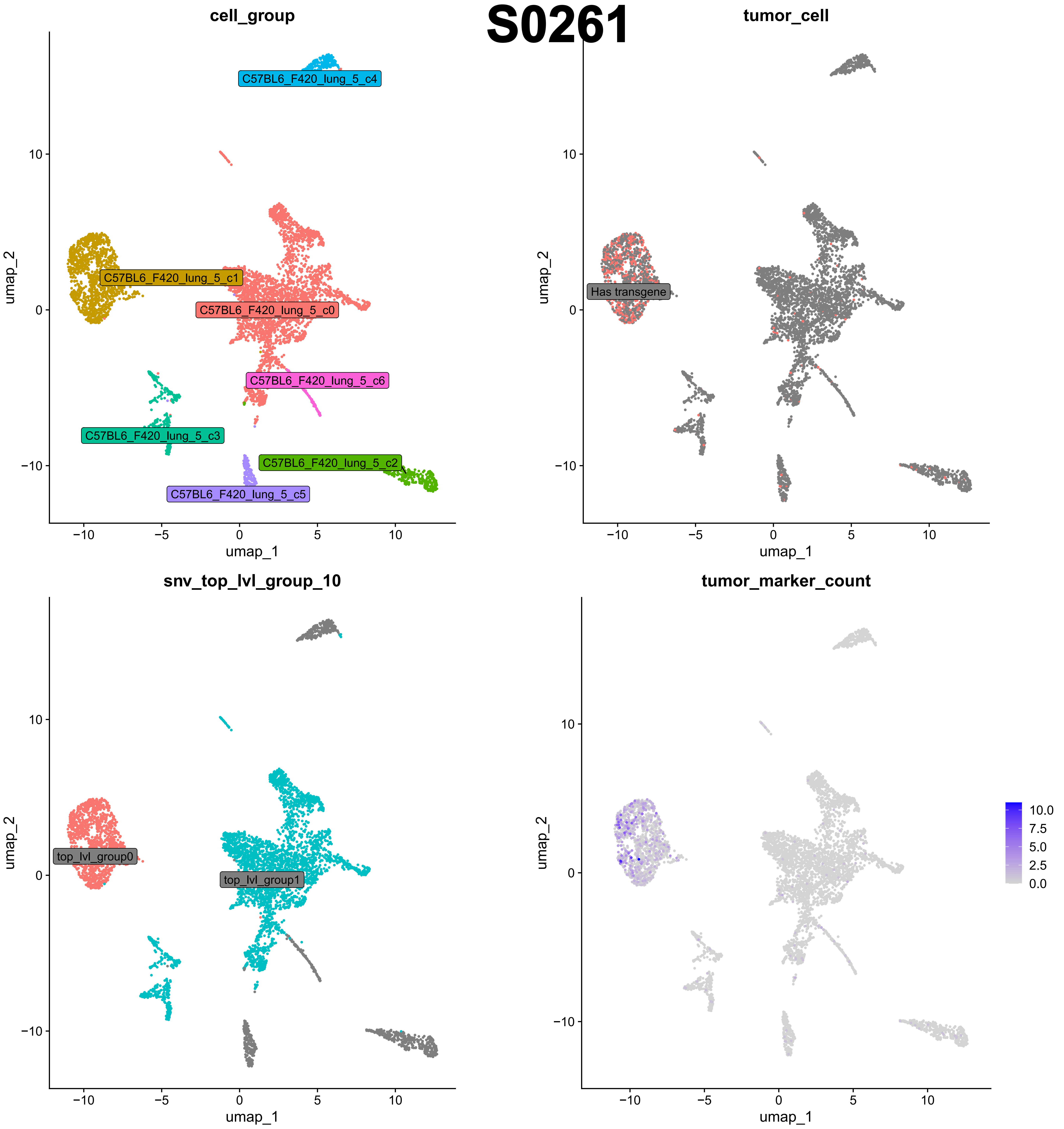

cell_group
C57BL6_F420_lung_5_c4
C57BL6_F420_lung_5_c1
C57BL6_F420_lung_5_c0
C57BL6_F420_lung_5_c6
C57BL6_F420_lung_5_c3
C57BL6_F420_lung_5_c2
C57BL6_F420_lung_5_c5
umap_1
umap_2
tumor_cell
Has transgene
snv_top_lvl_group_10
top_lvl_group0
top_lvl_group1
tumor_marker_count
10.0
7.5
5.0
2.5
0.0


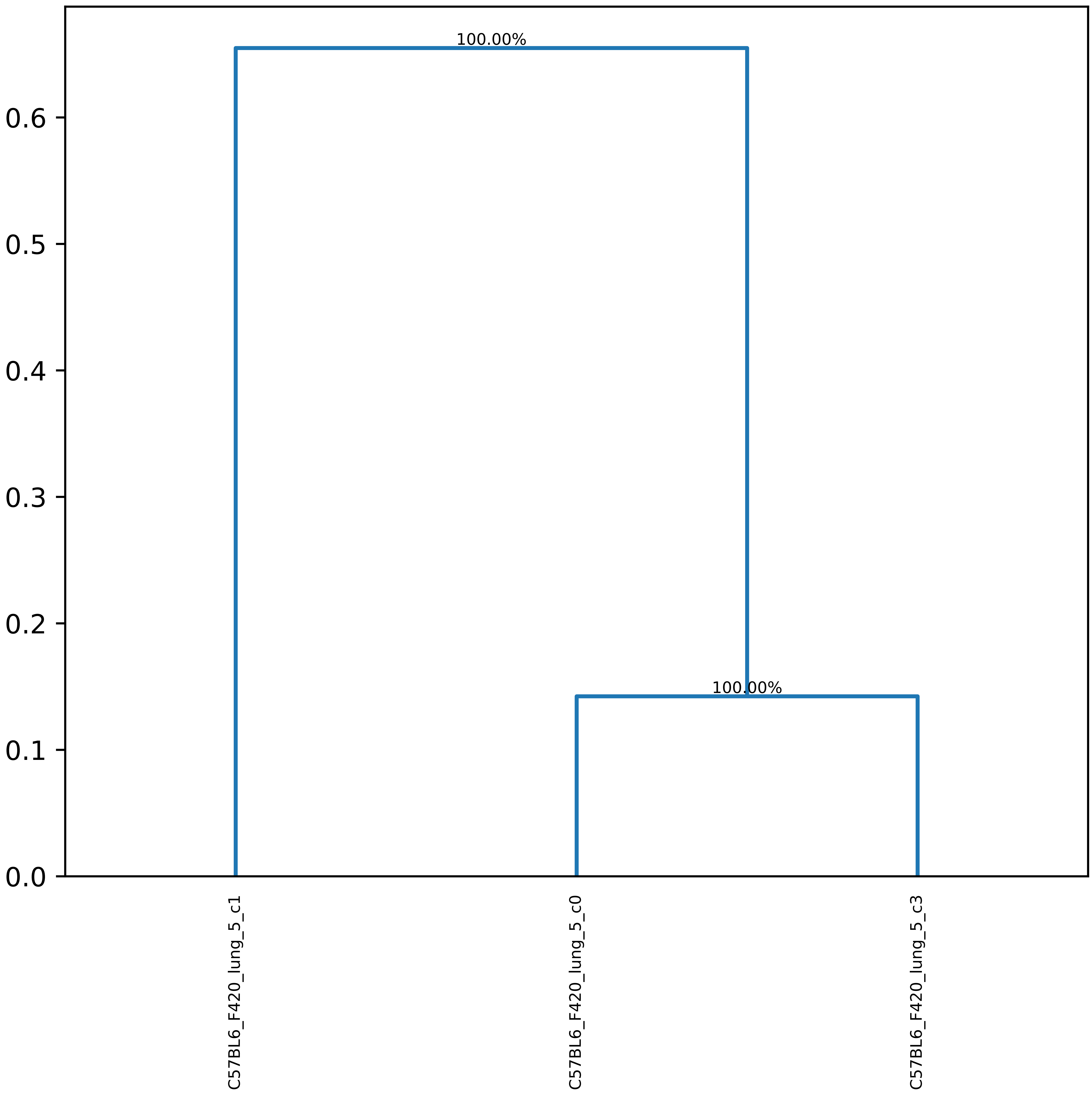

100.00%
100.00%
0.0
0.1
0.2
0.3
0.4
0.5
0.6
C57BL6_F420_lung_5_c1
C57BL6_F420_lung_5_c0
C57BL6_F420_lung_5_c3

**Supplemental Figure 5: Hierarchical clustering trees for comparison of ground truth tumor samples**

We used scanBit on ground truth samples where tumor cells are identifiable by expression of a transgene. We used a minimum read depth threshold of 10 for this analysis. For each of the eleven samples, we plotted the transcriptional clusters (cell_group), cells with detectable transgene reads (tumor_cell), genetically distinct groups as determined by scanBit (snv_top_lvl_group_10), the number of UMIs identified for each transgene (tumor_marker_count) and the resulting hierarchical clustering tree from scanBit with bootstrap values.

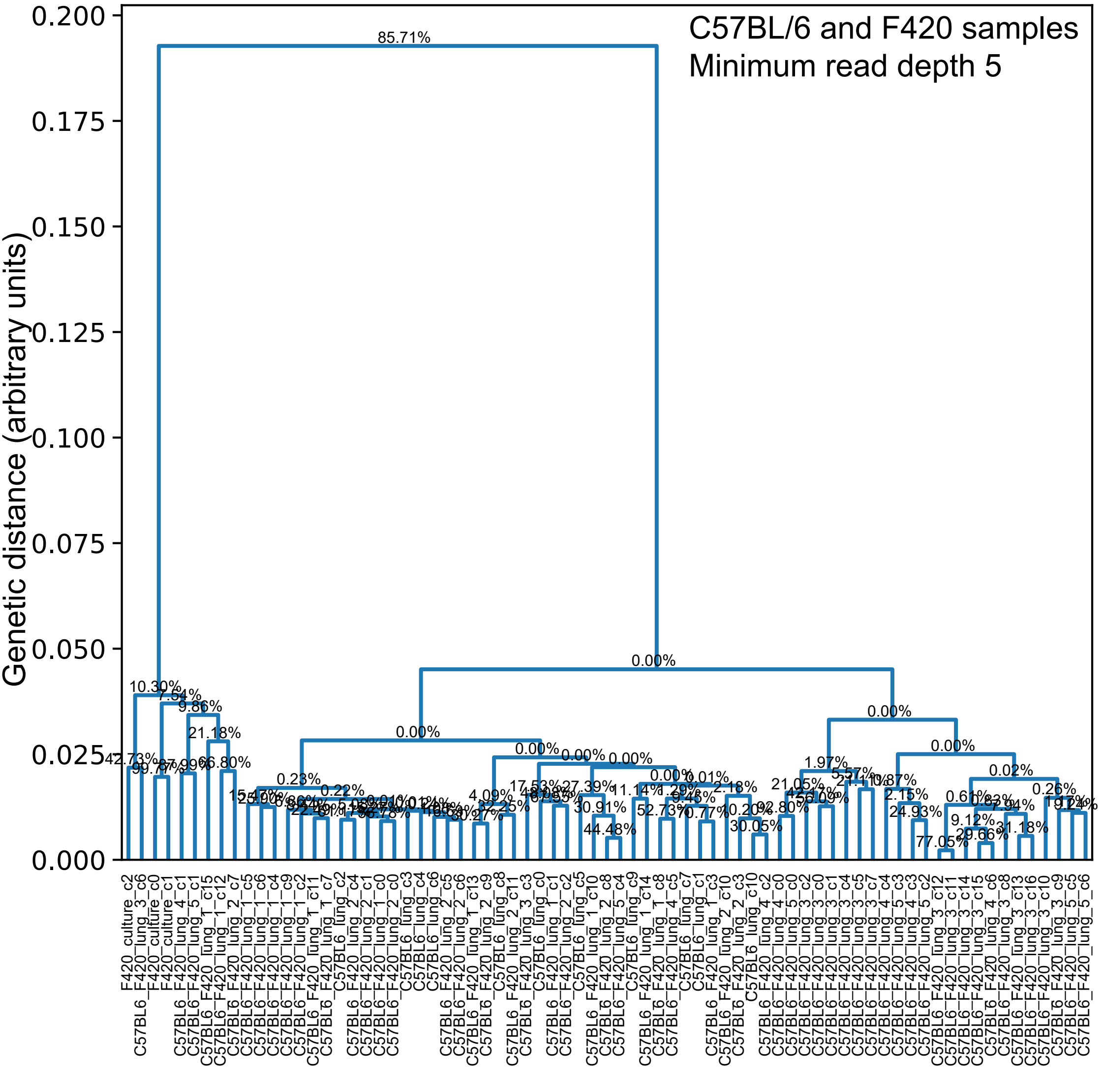
C57BL/6 and F420 samples
Minimum read depth 5
Genetic distance (arbitrary units)
0.200
0.175
0.150
0.125
0.100
0.075
0.050
0.025
0.000
85.71%
0.00%

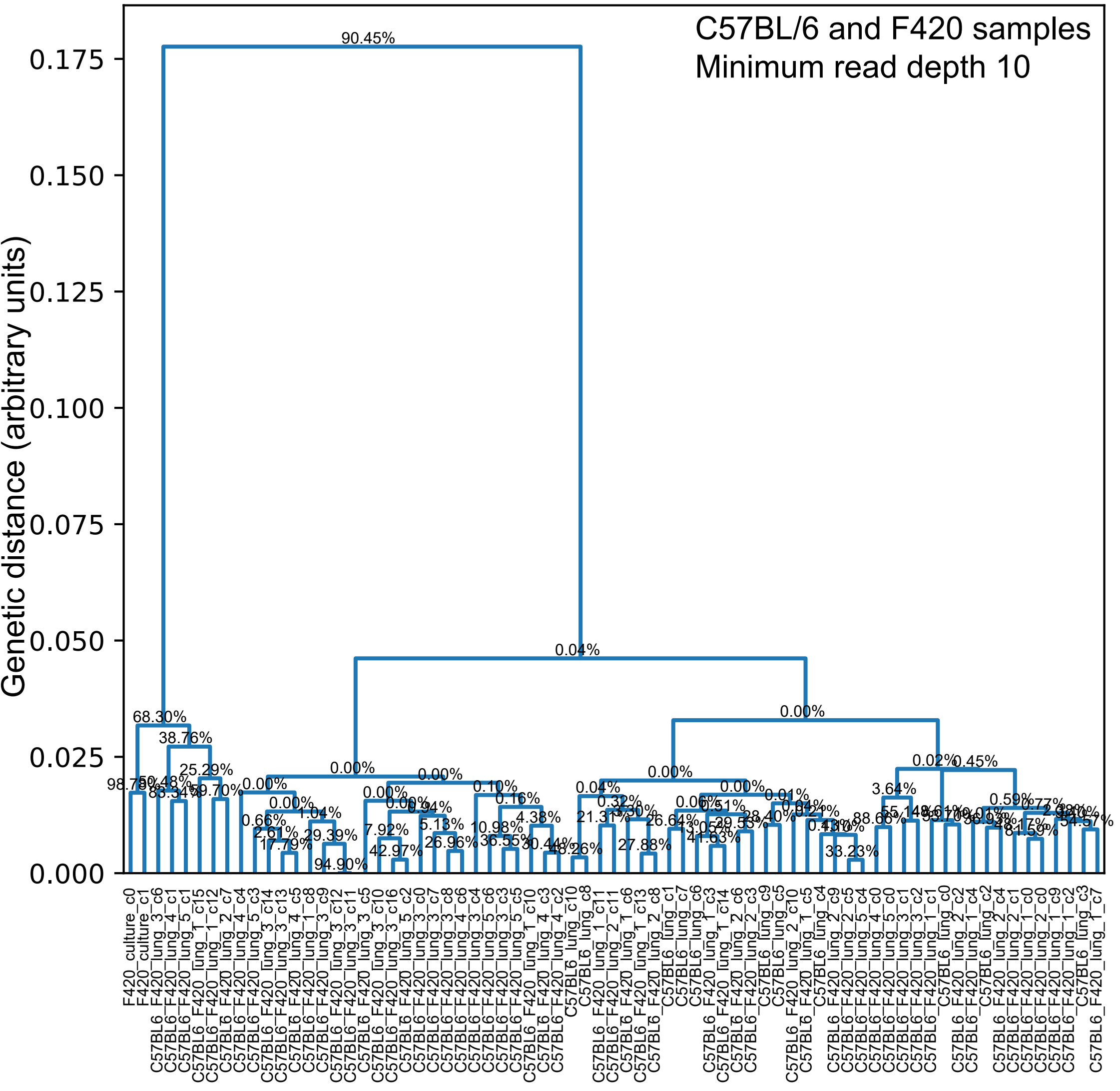
C57BL/6 and F420 samples
Minimum read depth 10
Genetic distance (arbitrary units)
0.175
0.150
0.125
0.100
0.075
0.050
0.025
0.000
90.45%
0.04%
0.00%
68.30%
38.76%
25.29%
0.02%
0.45%
3.64%
F420_culture_c0
F420_culture_c1
C57BL6_F420_lung_3_c6
C57BL6_F420_lung_4_c1
C57BL6_F420_lung_5_c1
C57BL6_F420_lung_T_c15
C57BL6_F420_lung_1_c12
C57BL6_F420_lung_2_c7
C57BL6_F420_lung_4_c4
C57BL6_F420_lung_5_c3
C57BL6_F420_lung_3_c14
C57BL6_F420_lung_3_c13
C57BL6_F420_lung_4_c5
C57BL6_F420_lung_1_c8
C57BL6_F420_lung_3_c9
C57BL6_F420_lung_3_c12
C57BL6_F420_lung_3_c11
C57BL6_F420_lung_3_c5
C57BL6_F420_lung_3_c10
C57BL6_F420_lung_3_c16
C57BL6_F420_lung_5_c2
C57BL6_F420_lung_3_c0
C57BL6_F420_lung_3_c7
C57BL6_F420_lung_3_c8
C57BL6_F420_lung_4_c6
C57BL6_F420_lung_3_c4
C57BL6_F420_lung_5_c6
C57BL6_F420_lung_3_c3
C57BL6_F420_lung_5_c5
C57BL6_F420_lung_T_c10
C57BL6_F420_lung_4_c3
C57BL6_F420_lung_4_c2
C57BL6_lung_c10
C57BL6_lung_c8
C57BL6_F420_lung_1_c11
C57BL6_F420_lung_2_c11
C57BL6_F420_lung_T_c6
C57BL6_F420_lung_T_c13
C57BL6_F420_lung_2_c8
C57BL6_lung_c1
C57BL6_lung_c6
C57BL6_F420_lung_T_c3
C57BL6_F420_lung_T_c14
C57BL6_F420_lung_2_c6
C57BL6_F420_lung_2_c3
C57BL6_lung_c9
C57BL6_lung_c5
C57BL6_F420_lung_2_c10
C57BL6_F420_lung_T_c5
C57BL6_lung_c4
C57BL6_F420_lung_2_c9
C57BL6_F420_lung_2_c5
C57BL6_F420_lung_5_c4
C57BL6_F420_lung_4_c0
C57BL6_F420_lung_5_c0
C57BL6_F420_lung_3_c1
C57BL6_F420_lung_3_c2
C57BL6_F420_lung_1_c1
C57BL6_lung_c0
C57BL6_F420_lung_2_c2
C57BL6_F420_lung_1_c4
C57BL6_lung_c2
C57BL6_F420_lung_2_c4
C57BL6_F420_lung_2_c1
C57BL6_F420_lung_1_c0
C57BL6_F420_lung_2_c0
C57BL6_F420_lung_1_c9
C57BL6_F420_lung_1_c2
C57BL6_lung_c3
C57BL6_F420_lung_1_c7

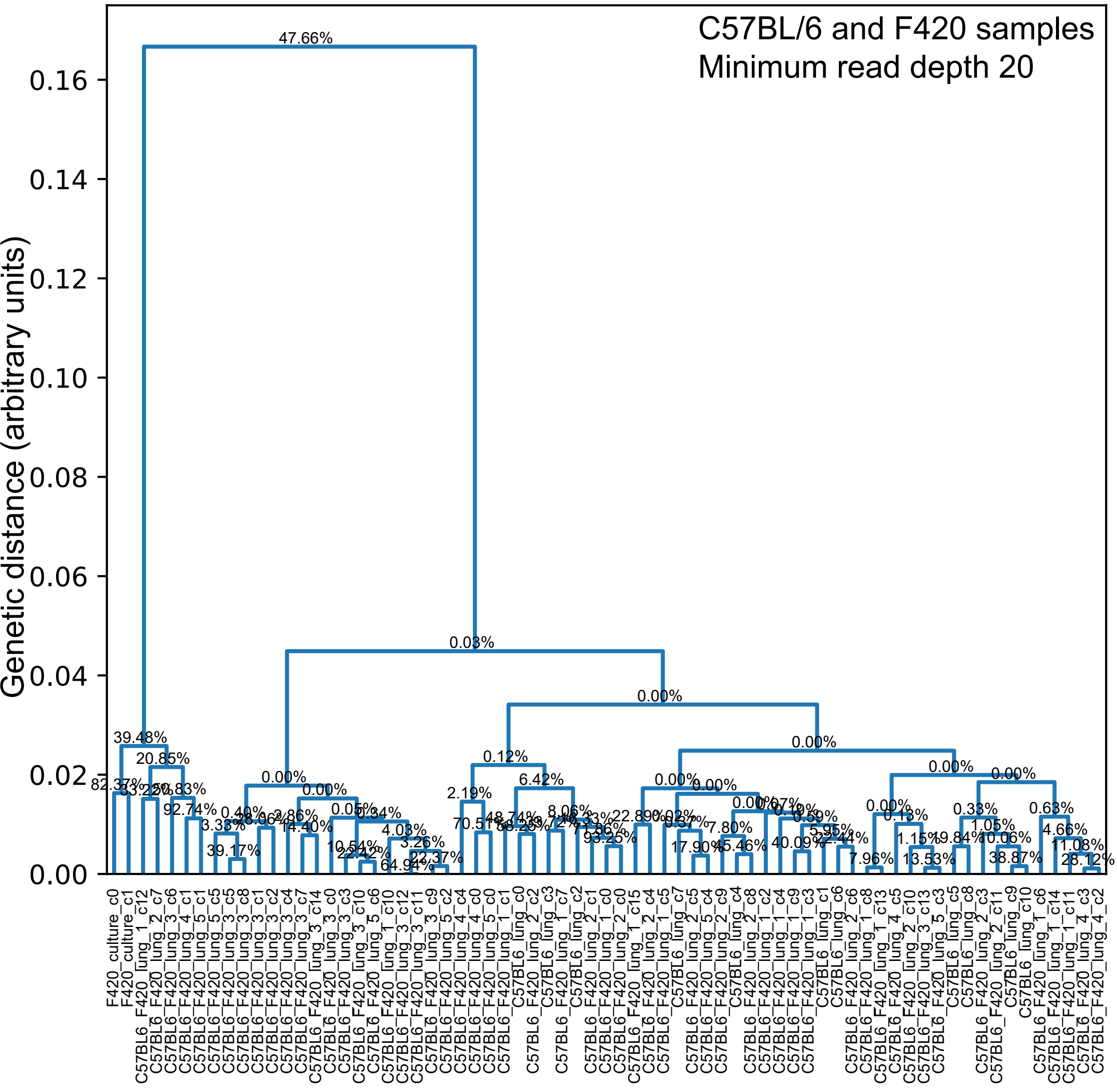
C57BL/6 and F420 samples
Minimum read depth 20
Genetic distance (arbitrary units)
0.00
0.02
0.04
0.06
0.08
0.10
0.12
0.14
0.16
47.66%
0.03%
0.00%
39.48%
20.85%
0.12%
6.42%
2.19%
92.74%
39.17%
14.40%
10.54%
4.03%
3.26%
22.37%
64.94%
70.51%
8.06%
22.89%
93.25%
17.90%
7.80%
45.46%
40.09%
0.59%
5.95%
7.96%
0.13%
1.15%
13.53%
19.84%
0.33%
1.05%
10.06%
38.87%
0.63%
4.66%
11.08%
28.12%
F420_culture_c0
F420_culture_c1
C57BL6_F420_lung_1_c12
C57BL6_F420_lung_2_c7
C57BL6_F420_lung_3_c6
C57BL6_F420_lung_4_c1
C57BL6_F420_lung_5_c1
C57BL6_F420_lung_5_c5
C57BL6_F420_lung_3_c5
C57BL6_F420_lung_3_c8
C57BL6_F420_lung_3_c1
C57BL6_F420_lung_3_c2
C57BL6_F420_lung_3_c4
C57BL6_F420_lung_3_c7
C57BL6_F420_lung_3_c14
C57BL6_F420_lung_3_c0
C57BL6_F420_lung_3_c3
C57BL6_F420_lung_3_c10
C57BL6_F420_lung_5_c6
C57BL6_F420_lung_T_c10
C57BL6_F420_lung_3_c12
C57BL6_F420_lung_3_c11
C57BL6_F420_lung_3_c9
C57BL6_F420_lung_5_c2
C57BL6_F420_lung_4_c4
C57BL6_F420_lung_4_c0
C57BL6_F420_lung_5_c0
C57BL6_F420_lung_1_c1
C57BL6_lung_c0
C57BL6_F420_lung_2_c2
C57BL6_lung_c3
C57BL6_F420_lung_1_c7
C57BL6_lung_c2
C57BL6_F420_lung_2_c1
C57BL6_F420_lung_1_c0
C57BL6_F420_lung_2_c0
C57BL6_F420_lung_T_c15
C57BL6_F420_lung_2_c4
C57BL6_F420_lung_1_c5
C57BL6_lung_c7
C57BL6_F420_lung_2_c5
C57BL6_F420_lung_5_c4
C57BL6_F420_lung_2_c9
C57BL6_lung_c4
C57BL6_F420_lung_2_c8
C57BL6_F420_lung_1_c2
C57BL6_F420_lung_1_c4
C57BL6_F420_lung_1_c9
C57BL6_F420_lung_1_c3
C57BL6_lung_c1
C57BL6_lung_c6
C57BL6_F420_lung_2_c6
C57BL6_F420_lung_1_c8
C57BL6_F420_lung_T_c13
C57BL6_F420_lung_4_c5
C57BL6_F420_lung_2_c10
C57BL6_F420_lung_3_c13
C57BL6_F420_lung_5_c3
C57BL6_lung_c5
C57BL6_lung_c8
C57BL6_F420_lung_2_c3
C57BL6_F420_lung_2_c11
C57BL6_lung_c9
C57BL6_lung_c10
C57BL6_F420_lung_T_c6
C57BL6_F420_lung_T_c14
C57BL6_F420_lung_1_c11
C57BL6_F420_lung_4_c3
C57BL6_F420_lung_4_c2

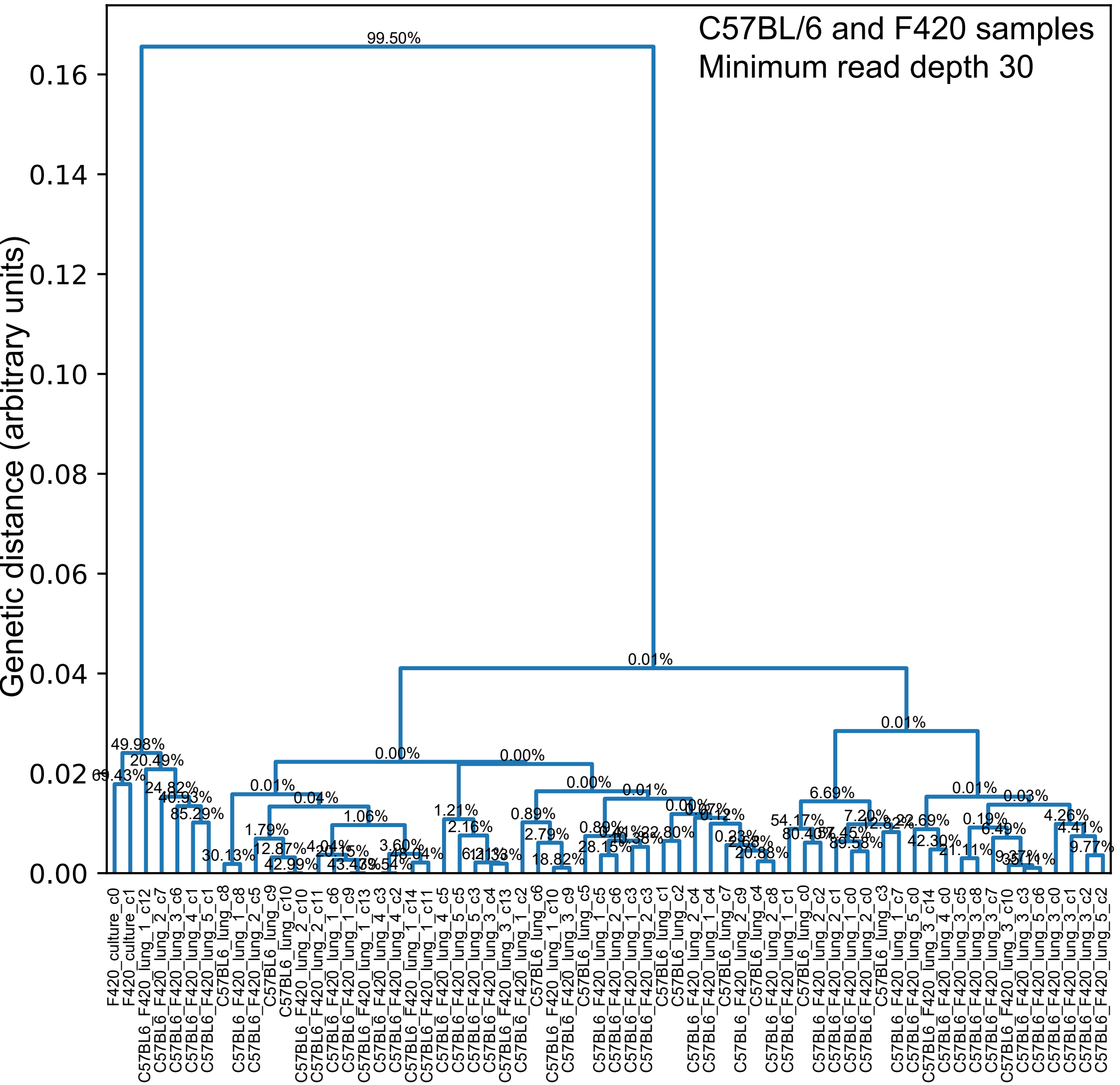
C57BL/6 and F420 samples
Minimum read depth 30
Genetic distance (arbitrary units)
0.00
0.02
0.04
0.06
0.08
0.10
0.12
0.14
0.16
99.50%
0.01%
0.01%
0.00%
0.00%
0.00%
49.98%
20.49%
85.29%
30.13%
1.79%
1.06%
1.21%
2.16%
0.89%
2.79%
18.82%
28.15%
6.69%
54.17%
7.20%
89.58%
42.30%
0.19%
6.49%
0.03%
4.26%
4.41%
9.77%
F420_culture_c0
F420_culture_c1
C57BL6_F420_lung_1_c12
C57BL6_F420_lung_2_c7
C57BL6_F420_lung_3_c6
C57BL6_F420_lung_4_c1
C57BL6_F420_lung_5_c1
C57BL6_lung_c8
C57BL6_F420_lung_1_c8
C57BL6_F420_lung_2_c5
C57BL6_lung_c9
C57BL6_lung_c10
C57BL6_F420_lung_2_c10
C57BL6_F420_lung_2_c11
C57BL6_F420_lung_1_c6
C57BL6_F420_lung_1_c9
C57BL6_F420_lung_1_c13
C57BL6_F420_lung_4_c3
C57BL6_F420_lung_4_c2
C57BL6_F420_lung_1_c14
C57BL6_F420_lung_1_c11
C57BL6_F420_lung_4_c5
C57BL6_F420_lung_5_c5
C57BL6_F420_lung_5_c3
C57BL6_F420_lung_3_c4
C57BL6_F420_lung_3_c13
C57BL6_F420_lung_1_c2
C57BL6_lung_c6
C57BL6_F420_lung_1_c10
C57BL6_F420_lung_3_c9
C57BL6_lung_c5
C57BL6_F420_lung_1_c5
C57BL6_F420_lung_2_c6
C57BL6_F420_lung_1_c3
C57BL6_F420_lung_2_c3
C57BL6_lung_c1
C57BL6_lung_c2
C57BL6_F420_lung_2_c4
C57BL6_F420_lung_1_c4
C57BL6_lung_c7
C57BL6_F420_lung_2_c9
C57BL6_lung_c4
C57BL6_F420_lung_2_c8
C57BL6_F420_lung_1_c1
C57BL6_lung_c0
C57BL6_F420_lung_2_c2
C57BL6_F420_lung_2_c1
C57BL6_F420_lung_1_c0
C57BL6_F420_lung_2_c0
C57BL6_lung_c3
C57BL6_F420_lung_1_c7
C57BL6_F420_lung_5_c0
C57BL6_F420_lung_3_c14
C57BL6_F420_lung_4_c0
C57BL6_F420_lung_3_c5
C57BL6_F420_lung_3_c8
C57BL6_F420_lung_3_c7
C57BL6_F420_lung_3_c10
C57BL6_F420_lung_3_c3
C57BL6_F420_lung_5_c6
C57BL6_F420_lung_3_c0
C57BL6_F420_lung_3_c1
C57BL6_F420_lung_3_c2
C57BL6_F420_lung_5_c2

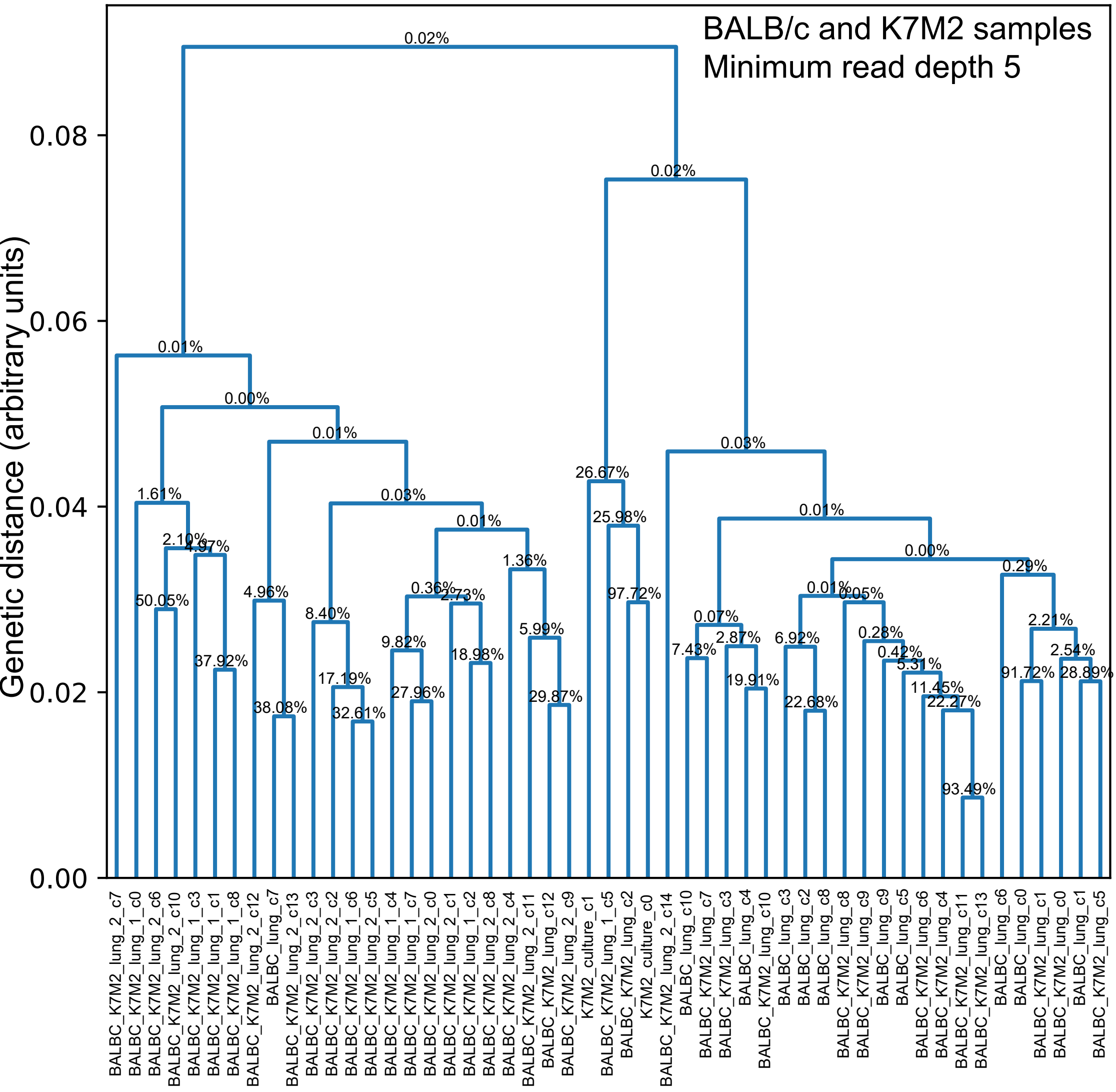
BALB/c and K7M2 samples
Minimum read depth 5
Genetic distance (arbitrary units)
0.00
0.02
0.04
0.06
0.08
0.02%
0.02%
0.01%
0.00%
0.01%
0.03%
0.01%
0.01%
0.00%
0.29%
26.67%
25.98%
97.72%
1.61%
2.10%
4.97%
50.05%
37.92%
4.96%
38.08%
8.40%
17.19%
32.61%
0.36%
2.73%
9.82%
27.96%
18.98%
1.36%
5.99%
29.87%
0.07%
7.43%
2.87%
19.91%
0.01%
0.05%
6.92%
22.68%
0.28%
0.42%
5.31%
11.45%
22.27%
93.49%
2.21%
2.54%
91.72%
28.89%
BALBC_K7M2_lung_2_c7
BALBC_K7M2_lung_1_c0
BALBC_K7M2_lung_2_c6
BALBC_K7M2_lung_2_c10
BALBC_K7M2_lung_1_c3
BALBC_K7M2_lung_1_c1
BALBC_K7M2_lung_1_c8
BALBC_K7M2_lung_2_c12
BALBC_lung_c7
BALBC_K7M2_lung_2_c13
BALBC_K7M2_lung_2_c3
BALBC_K7M2_lung_2_c2
BALBC_K7M2_lung_1_c6
BALBC_K7M2_lung_2_c5
BALBC_K7M2_lung_1_c4
BALBC_K7M2_lung_1_c7
BALBC_K7M2_lung_2_c0
BALBC_K7M2_lung_2_c1
BALBC_K7M2_lung_1_c2
BALBC_K7M2_lung_2_c8
BALBC_K7M2_lung_2_c4
BALBC_K7M2_lung_2_c11
BALBC_K7M2_lung_1_c12
BALBC_K7M2_lung_2_c9
K7M2_culture_c1
BALBC_K7M2_lung_1_c5
BALBC_K7M2_lung_c2
K7M2_culture_c0
BALBC_K7M2_lung_2_c14
BALBC_lung_c10
BALBC_K7M2_lung_c7
BALBC_K7M2_lung_c3
BALBC_lung_c4
BALBC_K7M2_lung_c10
BALBC_lung_c3
BALBC_lung_c2
BALBC_lung_c8
BALBC_K7M2_lung_c8
BALBC_K7M2_lung_c9
BALBC_lung_c9
BALBC_lung_c5
BALBC_K7M2_lung_c6
BALBC_K7M2_lung_c4
BALBC_K7M2_lung_c11
BALBC_K7M2_lung_c13
BALBC_lung_c6
BALBC_lung_c0
BALBC_K7M2_lung_c1
BALBC_K7M2_lung_c0
BALBC_lung_c1
BALBC_K7M2_lung_c5

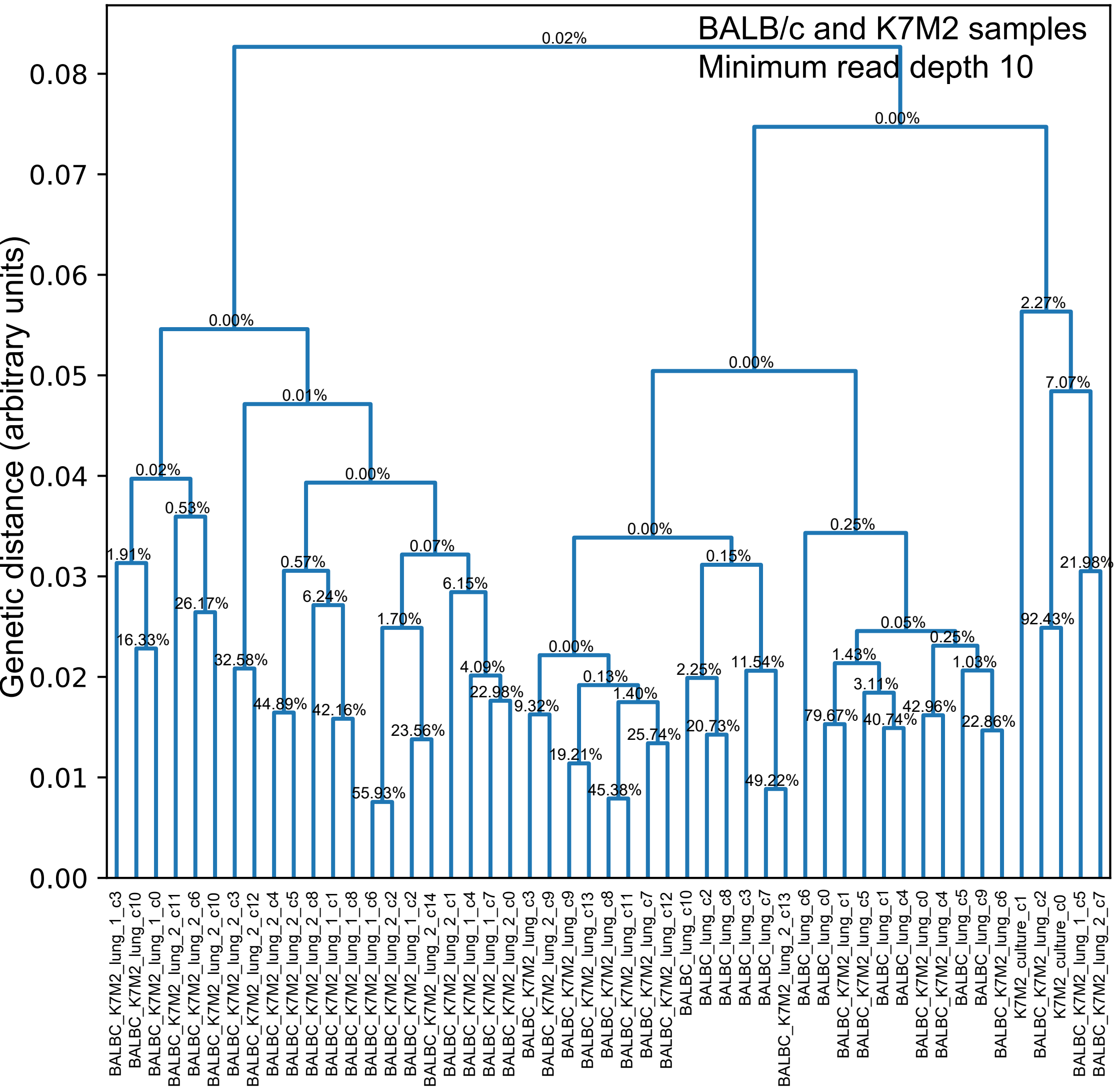

BALB/c and K7M2 samples
Minimum read depth 10
Genetic distance (arbitrary units)
0.00
0.01
0.02
0.03
0.04
0.05
0.06
0.07
0.08
0.02%
0.00%
0.00%
0.01%
0.02%
0.53%
0.00%
0.00%
0.25%
2.27%
7.07%
1.91%
16.33%
26.17%
32.58%
0.57%
6.24%
44.89%
42.16%
0.07%
1.70%
55.93%
23.56%
6.15%
4.09%
22.98%
0.00%
9.32%
0.13%
19.21%
1.40%
45.38%
25.74%
0.15%
2.25%
20.73%
11.54%
49.22%
0.05%
1.43%
79.67%
3.11%
40.74%
0.25%
42.96%
1.03%
22.86%
92.43%
21.98%
BALBC_K7M2_lung_1_c3
BALBC_K7M2_lung_c10
BALBC_K7M2_lung_1_c0
BALBC_K7M2_lung_2_c11
BALBC_K7M2_lung_2_c6
BALBC_K7M2_lung_2_c10
BALBC_K7M2_lung_2_c3
BALBC_K7M2_lung_2_c12
BALBC_K7M2_lung_2_c4
BALBC_K7M2_lung_2_c5
BALBC_K7M2_lung_2_c8
BALBC_K7M2_lung_1_c1
BALBC_K7M2_lung_1_c8
BALBC_K7M2_lung_1_c6
BALBC_K7M2_lung_2_c2
BALBC_K7M2_lung_1_c2
BALBC_K7M2_lung_2_c14
BALBC_K7M2_lung_2_c1
BALBC_K7M2_lung_1_c4
BALBC_K7M2_lung_1_c7
BALBC_K7M2_lung_2_c0
BALBC_K7M2_lung_c3
BALBC_K7M2_lung_2_c9
BALBC_K7M2_lung_c9
BALBC_K7M2_lung_c13
BALBC_K7M2_lung_c8
BALBC_K7M2_lung_c11
BALBC_K7M2_lung_c7
BALBC_K7M2_lung_c12
BALBC_lung_c10
BALBC_lung_c2
BALBC_lung_c8
BALBC_lung_c3
BALBC_lung_c7
BALBC_K7M2_lung_2_c13
BALBC_lung_c6
BALBC_lung_c0
BALBC_K7M2_lung_c1
BALBC_K7M2_lung_c5
BALBC_lung_c1
BALBC_lung_c4
BALBC_K7M2_lung_c0
BALBC_K7M2_lung_c4
BALBC_lung_c5
BALBC_lung_c9
BALBC_K7M2_lung_c6
K7M2_culture_c1
BALBC_K7M2_lung_c2
K7M2_culture_c0
BALBC_K7M2_lung_1_c5
BALBC_K7M2_lung_2_c7

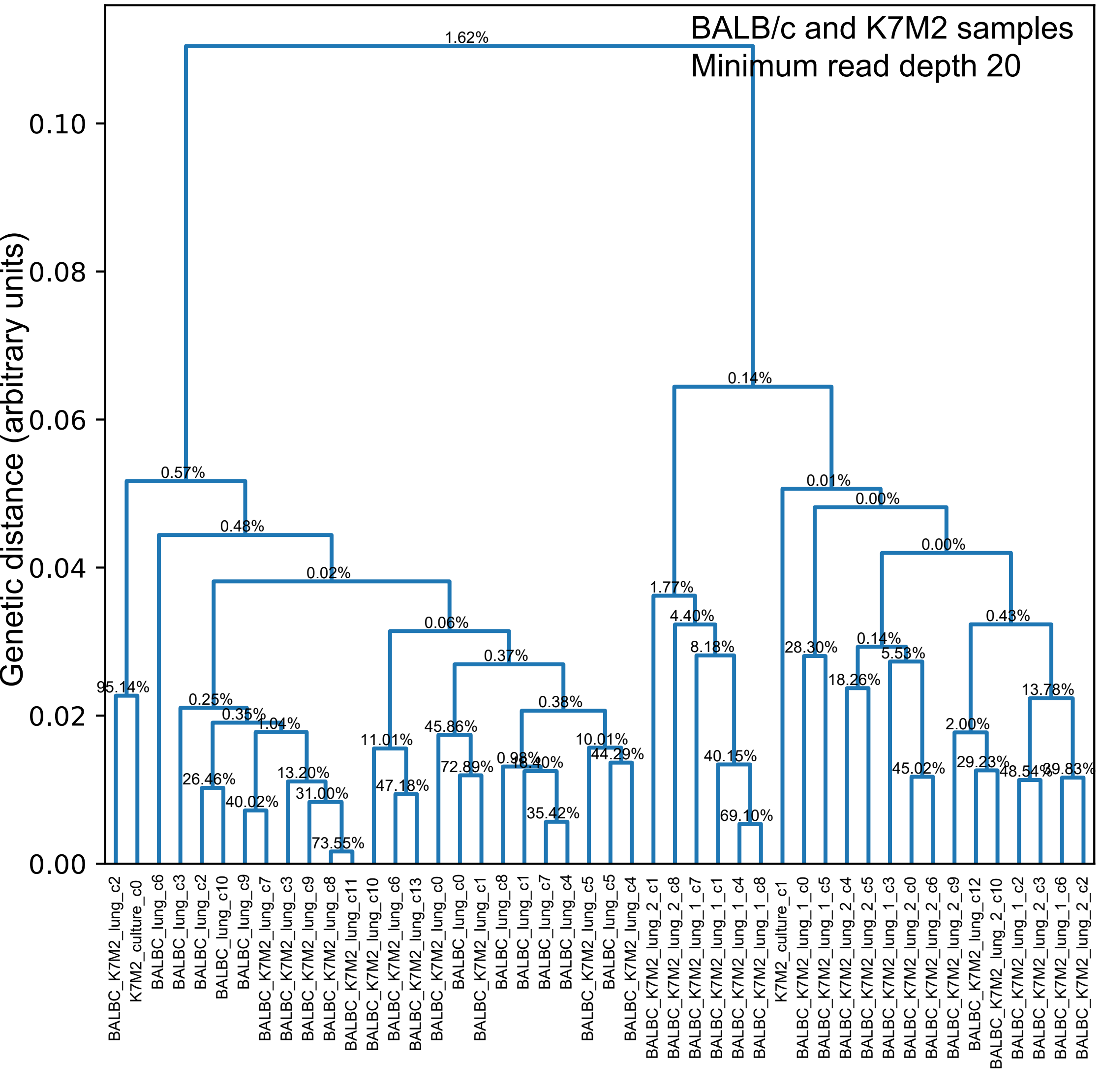
BALB/c and K7M2 samples
Minimum read depth 20
Genetic distance (arbitrary units)
0.00
0.02
0.04
0.06
0.08
0.10
1.62%
0.14%
0.57%
0.48%
0.02%
0.06%
0.37%
0.38%
0.25%
0.35%
1.04%
95.14%
26.46%
40.02%
13.20%
31.00%
73.55%
11.01%
47.18%
45.86%
72.89%
0.98%
18.40%
35.42%
10.01%
44.29%
1.77%
4.40%
8.18%
40.15%
69.10%
0.01%
0.00%
0.00%
28.30%
0.14%
5.53%
18.26%
45.02%
0.43%
2.00%
29.23%
13.78%
48.54%
29.83%
BALBC_K7M2_lung_c2
K7M2_culture_c0
BALBC_lung_c6
BALBC_lung_c3
BALBC_lung_c2
BALBC_lung_c10
BALBC_lung_c9
BALBC_K7M2_lung_c7
BALBC_K7M2_lung_c3
BALBC_K7M2_lung_c9
BALBC_K7M2_lung_c8
BALBC_K7M2_lung_c11
BALBC_K7M2_lung_c10
BALBC_K7M2_lung_c6
BALBC_K7M2_lung_c13
BALBC_K7M2_lung_c0
BALBC_lung_c0
BALBC_K7M2_lung_c1
BALBC_lung_c8
BALBC_lung_c1
BALBC_lung_c7
BALBC_lung_c4
BALBC_K7M2_lung_c5
BALBC_lung_c5
BALBC_K7M2_lung_c4
BALBC_K7M2_lung_2_c1
BALBC_K7M2_lung_2_c8
BALBC_K7M2_lung_1_c7
BALBC_K7M2_lung_1_c1
BALBC_K7M2_lung_1_c4
BALBC_K7M2_lung_1_c8
K7M2_culture_c1
BALBC_K7M2_lung_1_c0
BALBC_K7M2_lung_1_c5
BALBC_K7M2_lung_2_c4
BALBC_K7M2_lung_2_c5
BALBC_K7M2_lung_1_c3
BALBC_K7M2_lung_2_c0
BALBC_K7M2_lung_2_c6
BALBC_K7M2_lung_2_c9
BALBC_K7M2_lung_c12
BALBC_K7M2_lung_2_c10
BALBC_K7M2_lung_1_c2
BALBC_K7M2_lung_2_c3
BALBC_K7M2_lung_1_c6
BALBC_K7M2_lung_2_c2

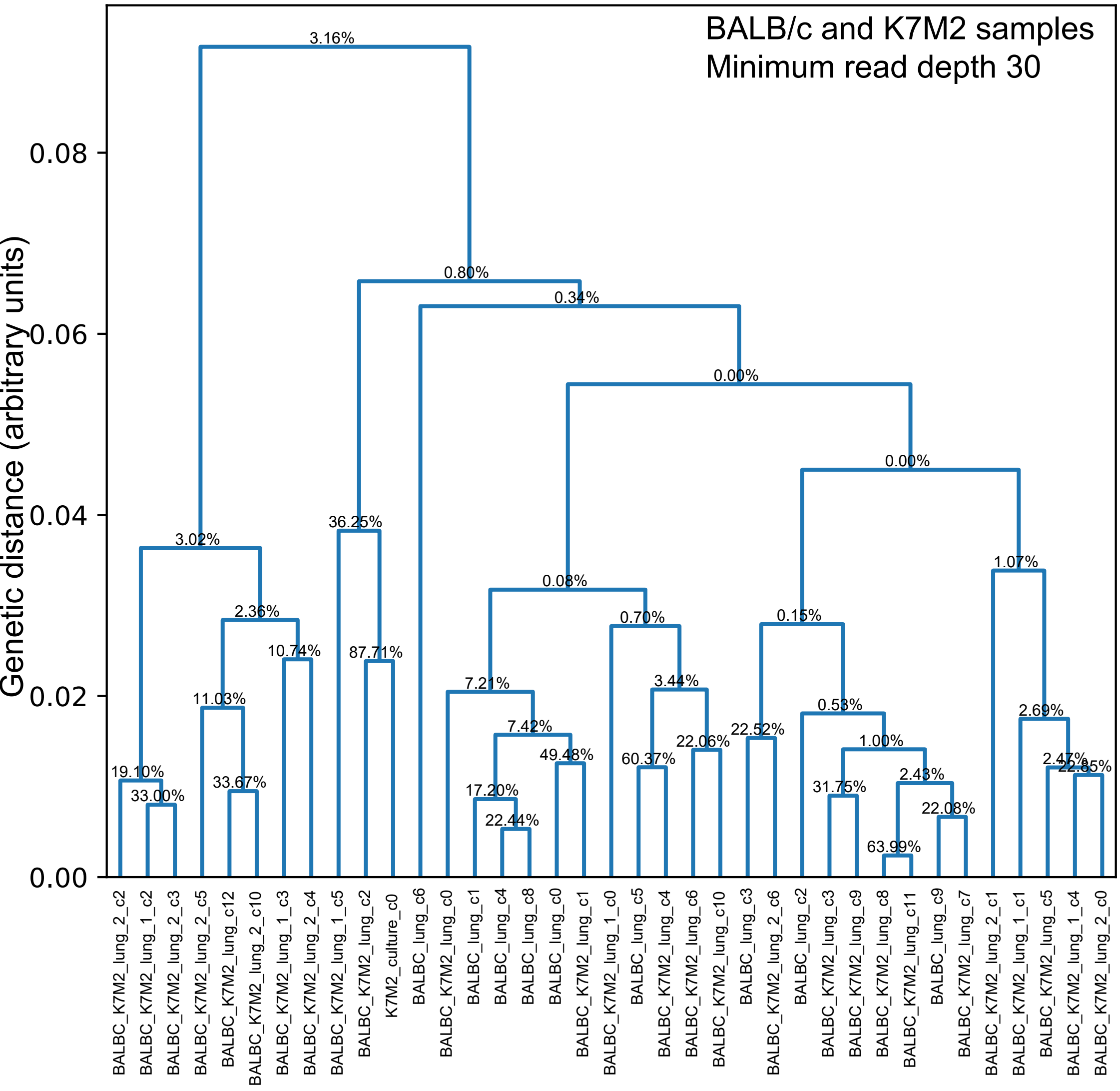

BALB/c and K7M2 samples
Minimum read depth 30
Genetic distance (arbitrary units)
0.00
0.02
0.04
0.06
0.08
3.16%
0.80%
0.34%
0.00%
0.00%
3.02%
2.36%
36.25%
87.71%
10.74%
11.03%
33.67%
19.10%
33.00%
0.08%
7.21%
7.42%
49.48%
17.20%
22.44%
0.70%
3.44%
22.06%
60.37%
0.15%
22.52%
0.53%
1.00%
2.43%
31.75%
22.08%
63.99%
1.07%
2.69%
2.47%
22.85%
BALBC_K7M2_lung_2_c2
BALBC_K7M2_lung_1_c2
BALBC_K7M2_lung_2_c3
BALBC_K7M2_lung_2_c5
BALBC_K7M2_lung_c12
BALBC_K7M2_lung_2_c10
BALBC_K7M2_lung_1_c3
BALBC_K7M2_lung_2_c4
BALBC_K7M2_lung_1_c5
BALBC_K7M2_lung_c2
K7M2_culture_c0
BALBC_lung_c6
BALBC_K7M2_lung_c0
BALBC_lung_c1
BALBC_lung_c4
BALBC_lung_c8
BALBC_lung_c0
BALBC_K7M2_lung_c1
BALBC_K7M2_lung_1_c0
BALBC_lung_c5
BALBC_K7M2_lung_c4
BALBC_K7M2_lung_c6
BALBC_K7M2_lung_c10
BALBC_lung_c3
BALBC_K7M2_lung_2_c6
BALBC_lung_c2
BALBC_K7M2_lung_c3
BALBC_K7M2_lung_c9
BALBC_K7M2_lung_c8
BALBC_K7M2_lung_c11
BALBC_lung_c9
BALBC_K7M2_lung_c7
BALBC_K7M2_lung_2_c1
BALBC_K7M2_lung_1_c1
BALBC_K7M2_lung_c5
BALBC_K7M2_lung_1_c4
BALBC_K7M2_lung_2_c0

**Supplemental Figure 6: Hierarchical clustering trees for comparison of F420 and C57BL/6 as well as K7M2 and BALB/c**

We compiled two datasets consisting of control cell culture tumor cells, control mouse lung and mouse lung metastases from tail vein injected tumor cells. Each dataset had either C57BL/6 and F420 cells or BALB/c and K7M2 cells. To determine if the tumor cells could be accurately distinguished from normal cells, we used scanBit on each dataset to generate hierarchical clustering trees from variant calls. We used minimum read depth thresholds of 5, 10, 20 or 30 for both datasets. Each tree is labeled with the source dataset and minimum read depth threshold applied. Bootstrap values for each node are overlaid in each tree.

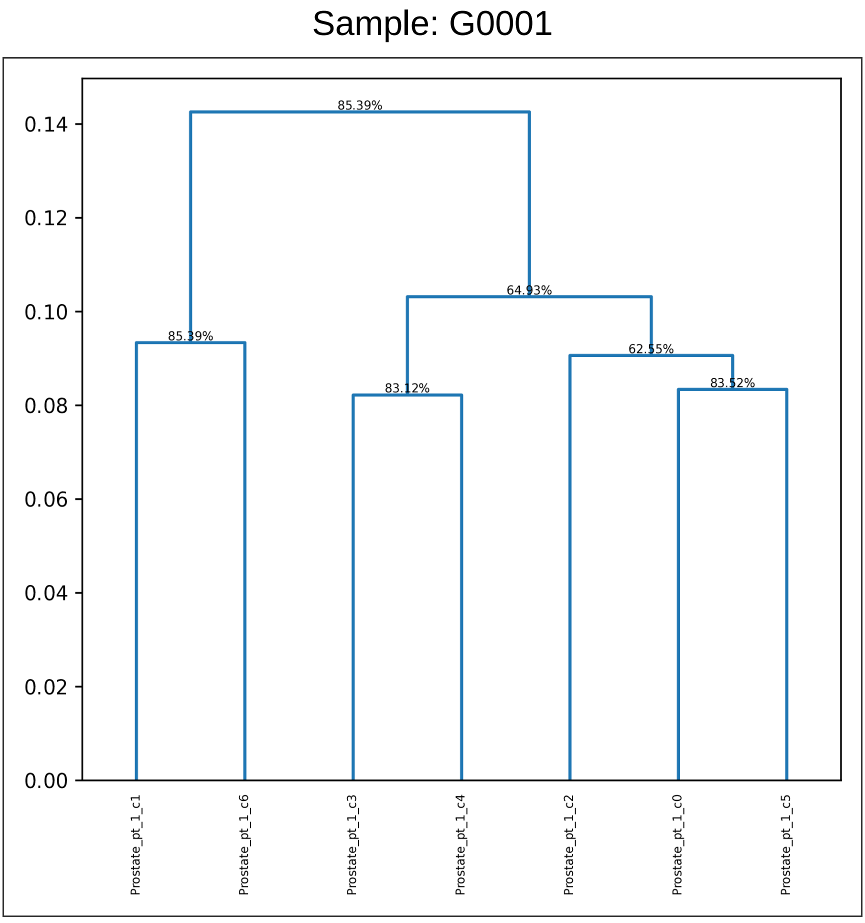
Sample: G0001
85.39%
85.39%
64.93%
83.12%
62.55%
83.52%
0.14
0.12
0.10
0.08
0.06
0.04
0.02
0.00
Prostate_pt_1_c1
Prostate_pt_1_c6
Prostate_pt_1_c3
Prostate_pt_1_c4
Prostate_pt_1_c2
Prostate_pt_1_c0
Prostate_pt_1_c5

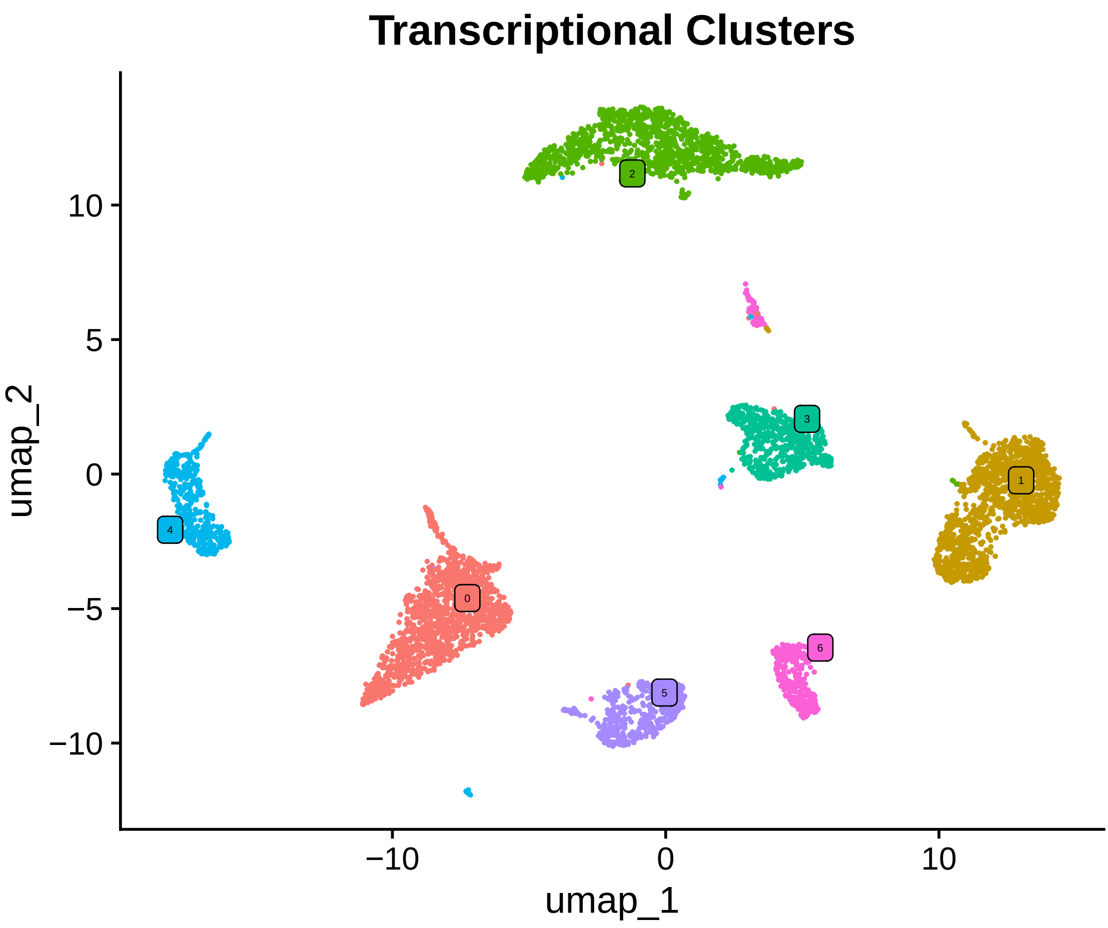
Transcriptional Clusters
umap_2
umap_1
10
5
0
−5
−10
−10
0
10
0
1
2
3
4
5
6

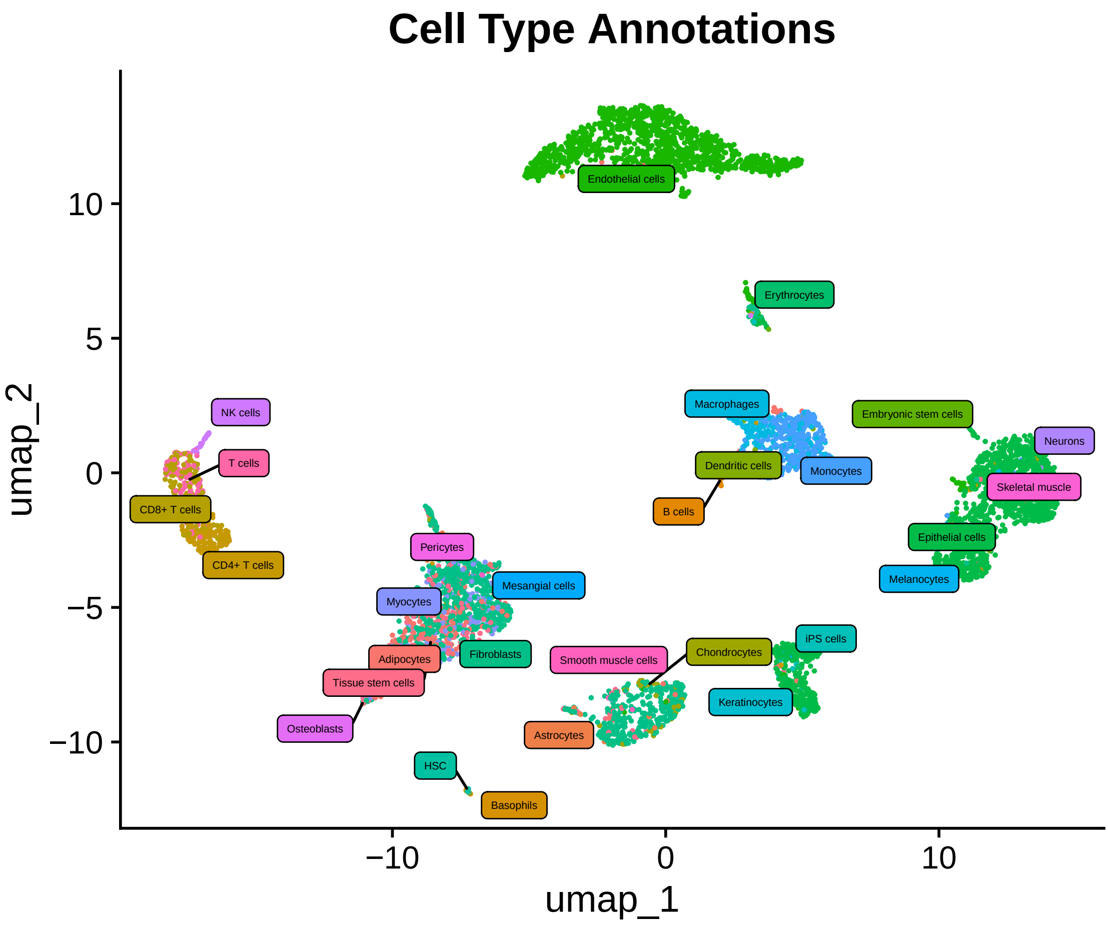
Cell Type Annotations
umap_2
umap_1
10
5
0
−5
−10
−10
0
10
Endothelial cells
Erythrocytes
NK cells
T cells
CD8+ T cells
CD4+ T cells
Macrophages
Dendritic cells
Monocytes
B cells
Embryonic stem cells
Neurons
Skeletal muscle
Epithelial cells
Melanocytes
Pericytes
Myocytes
Mesangial cells
Adipocytes
Fibroblasts
Tissue stem cells
Osteoblasts
Smooth muscle cells
Chondrocytes
iPS cells
Keratinocytes
Astrocytes
HSC
Basophils

Sample: G0002

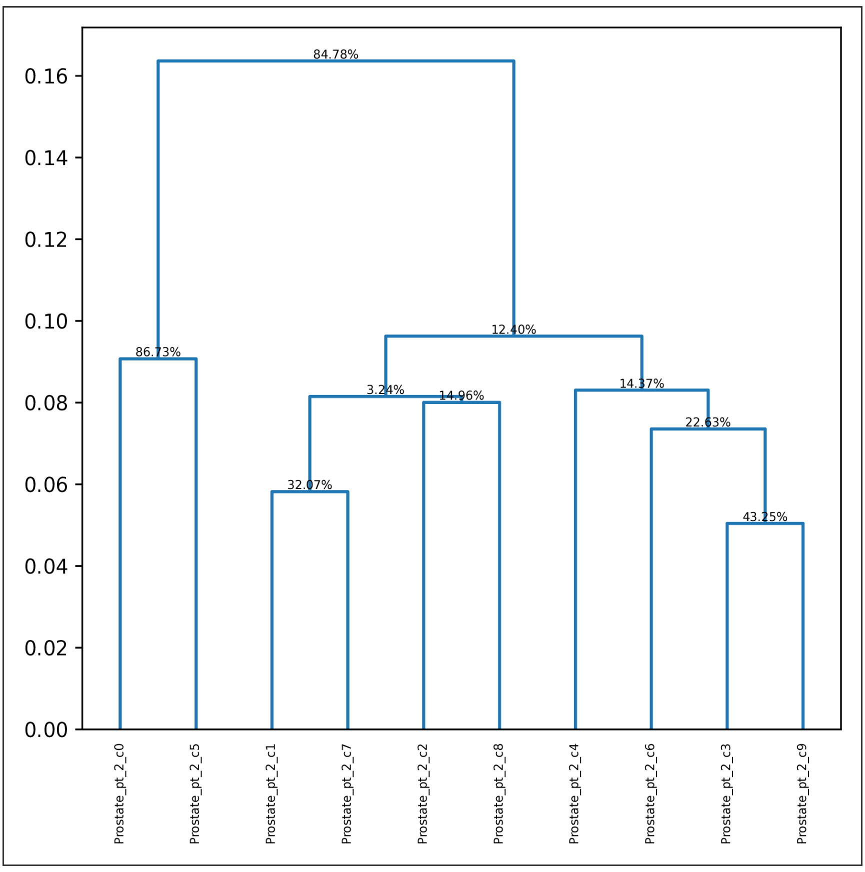


**Transcriptional Clusters**

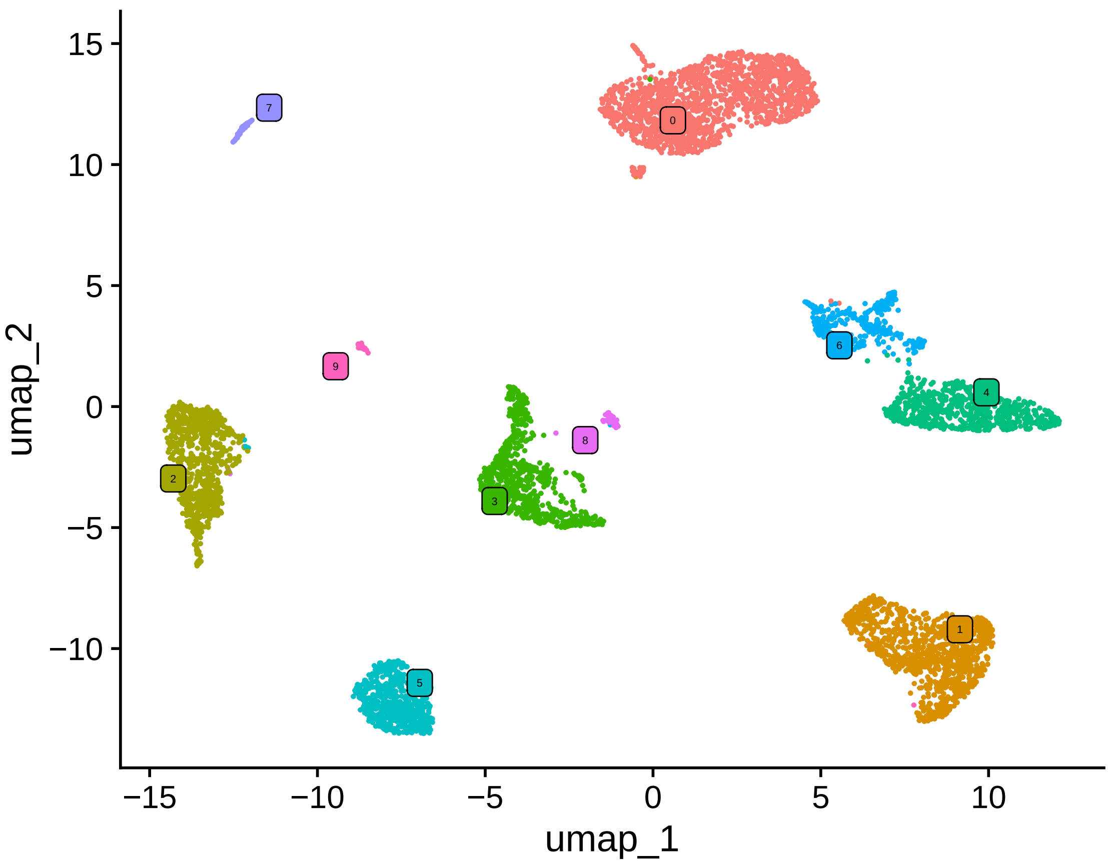


**Cell Type Annotations**

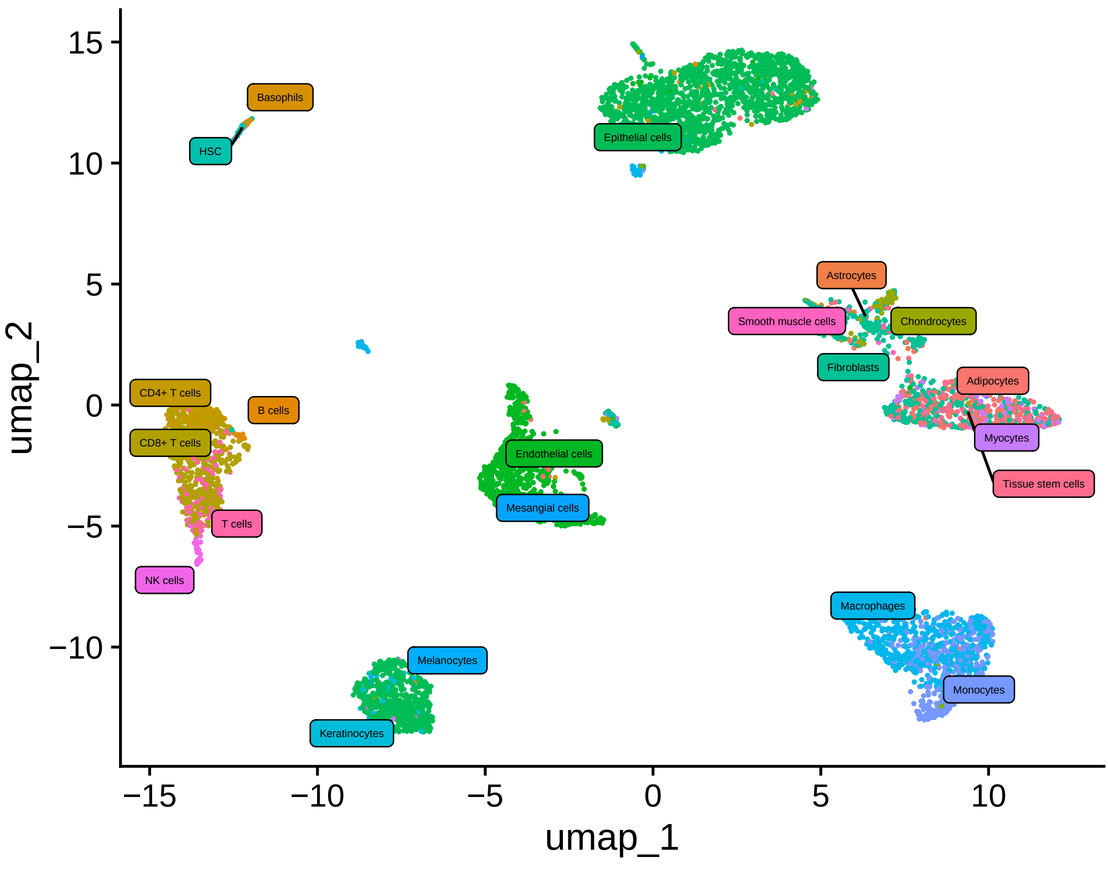

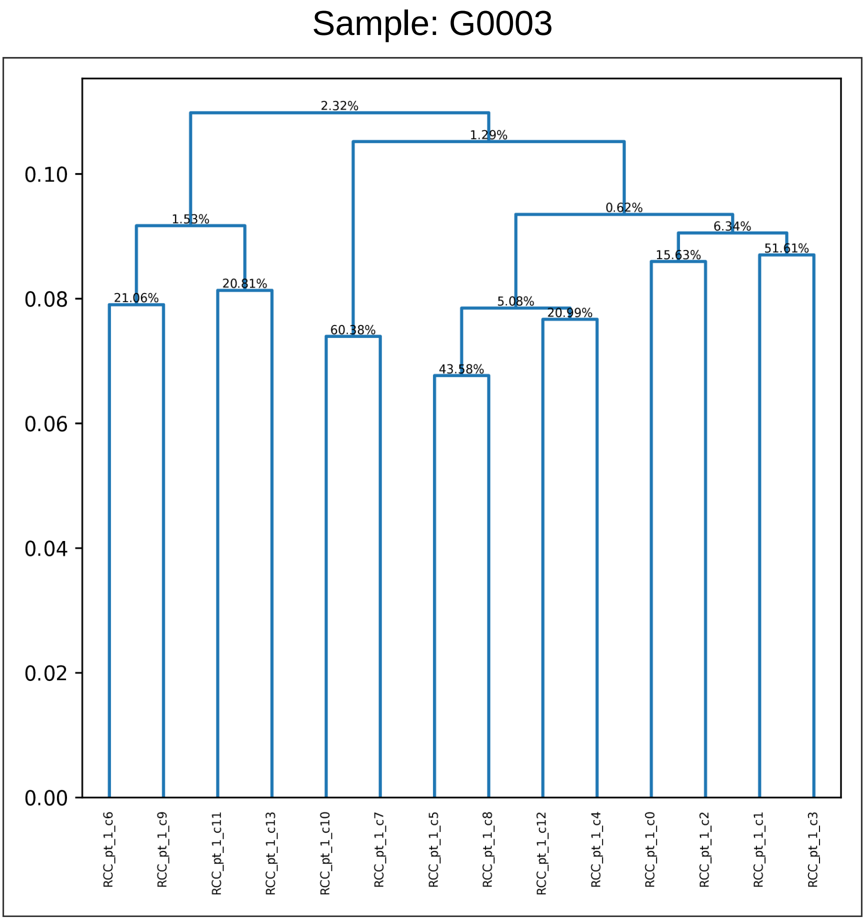
Sample: G0003
0.10
0.08
0.06
0.04
0.02
0.00
2.32%
1.29%
1.53%
0.62%
6.34%
21.06%
20.81%
60.38%
5.08%
20.99%
43.58%
15.63%
51.61%
RCC_pt_1_c6
RCC_pt_1_c9
RCC_pt_1_c11
RCC_pt_1_c13
RCC_pt_1_c10
RCC_pt_1_c7
RCC_pt_1_c5
RCC_pt_1_c8
RCC_pt_1_c12
RCC_pt_1_c4
RCC_pt_1_c0
RCC_pt_1_c2
RCC_pt_1_c1
RCC_pt_1_c3

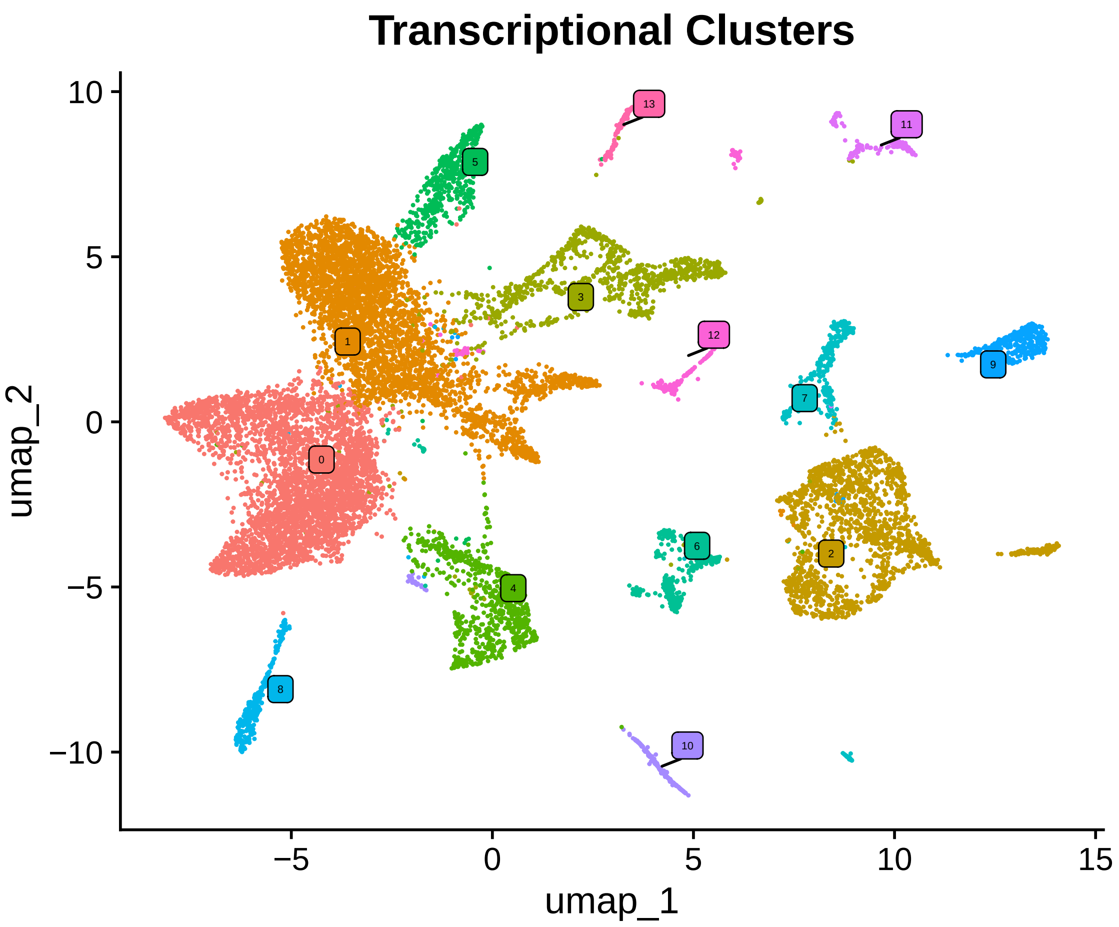
Transcriptional Clusters
umap_2
umap_1
10
5
0
−5
−10
15
0
1
2
3
4
5
6
7
8
9
10
11
12
13

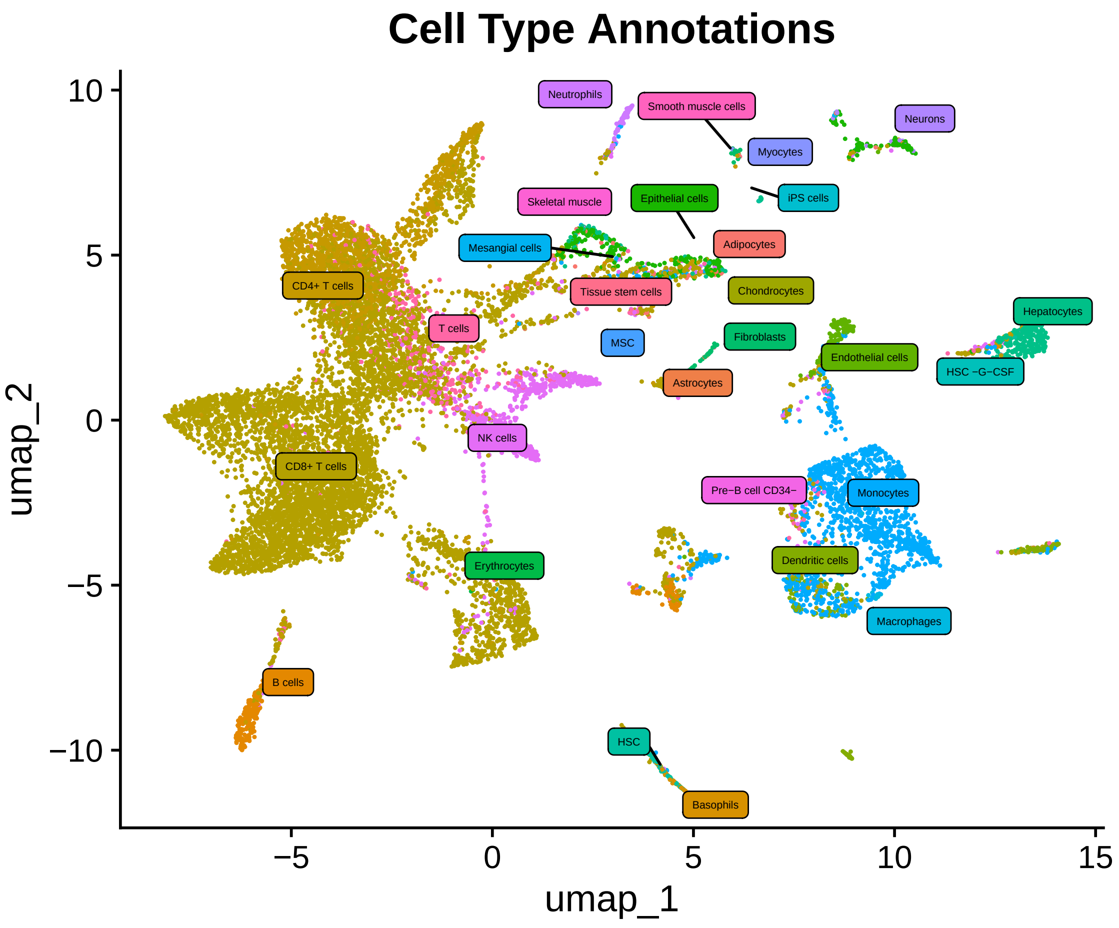
Cell Type Annotations
umap_2
umap_1
10
5
0
−5
−10
15
Neutrophils
Smooth muscle cells
Neurons
Myocytes
Skeletal muscle
Epithelial cells
iPS cells
Mesangial cells
Adipocytes
Tissue stem cells
Chondrocytes
CD4+ T cells
Hepatocytes
T cells
MSC
Fibroblasts
Endothelial cells
HSC -G-CSF
Astrocytes
NK cells
CD8+ T cells
Pre-B cell CD34-
Monocytes
Erythrocytes
Dendritic cells
Macrophages
B cells
HSC
Basophils

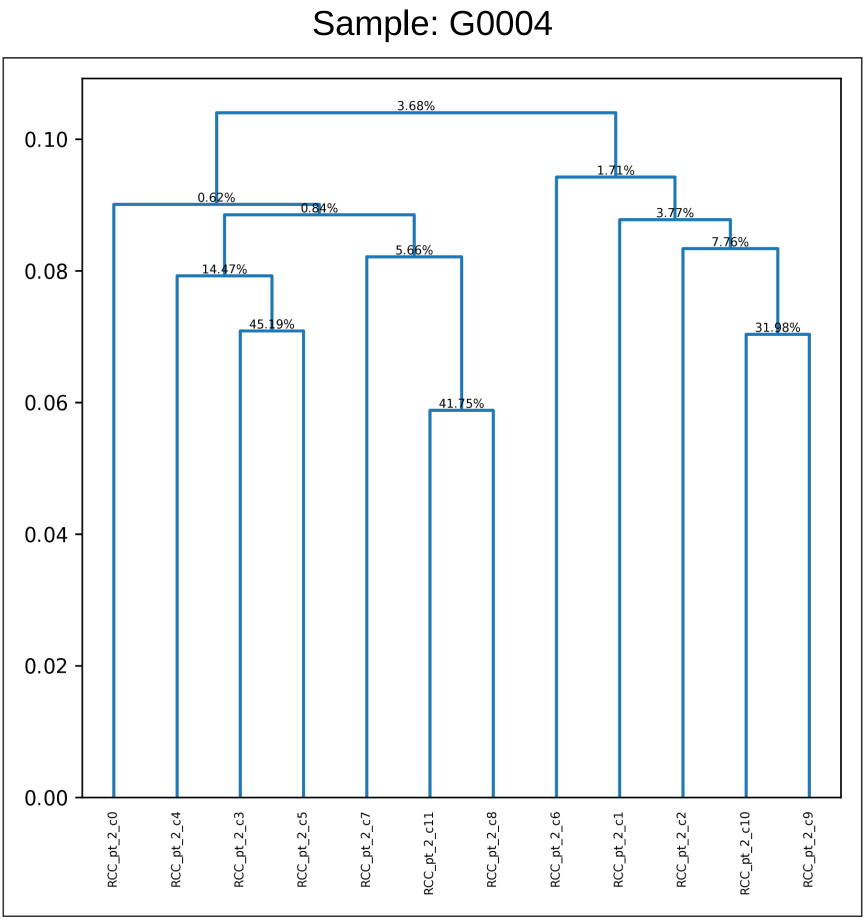
Sample: G0004
3.68%
0.62%
0.84%
1.71%
3.77%
7.76%
14.47%
5.66%
45.19%
31.98%
41.75%
0.10
0.08
0.06
0.04
0.02
0.00
RCC_pt_2_c0
RCC_pt_2_c4
RCC_pt_2_c3
RCC_pt_2_c5
RCC_pt_2_c7
RCC_pt_2_c11
RCC_pt_2_c8
RCC_pt_2_c6
RCC_pt_2_c1
RCC_pt_2_c2
RCC_pt_2_c10
RCC_pt_2_c9

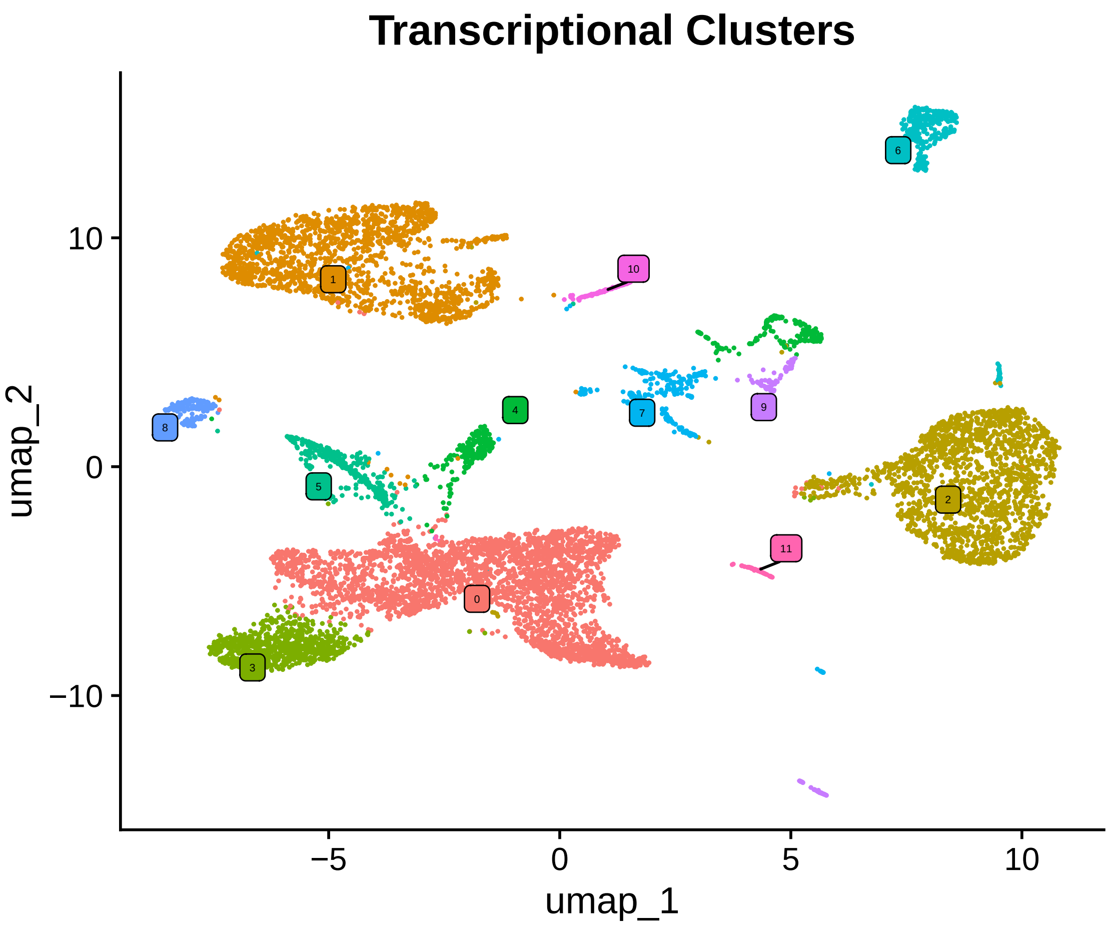
Transcriptional Clusters
umap_2
umap_1
10
0
−10
−5
0
5
10
0
1
2
3
4
5
6
7
8
9
10
11

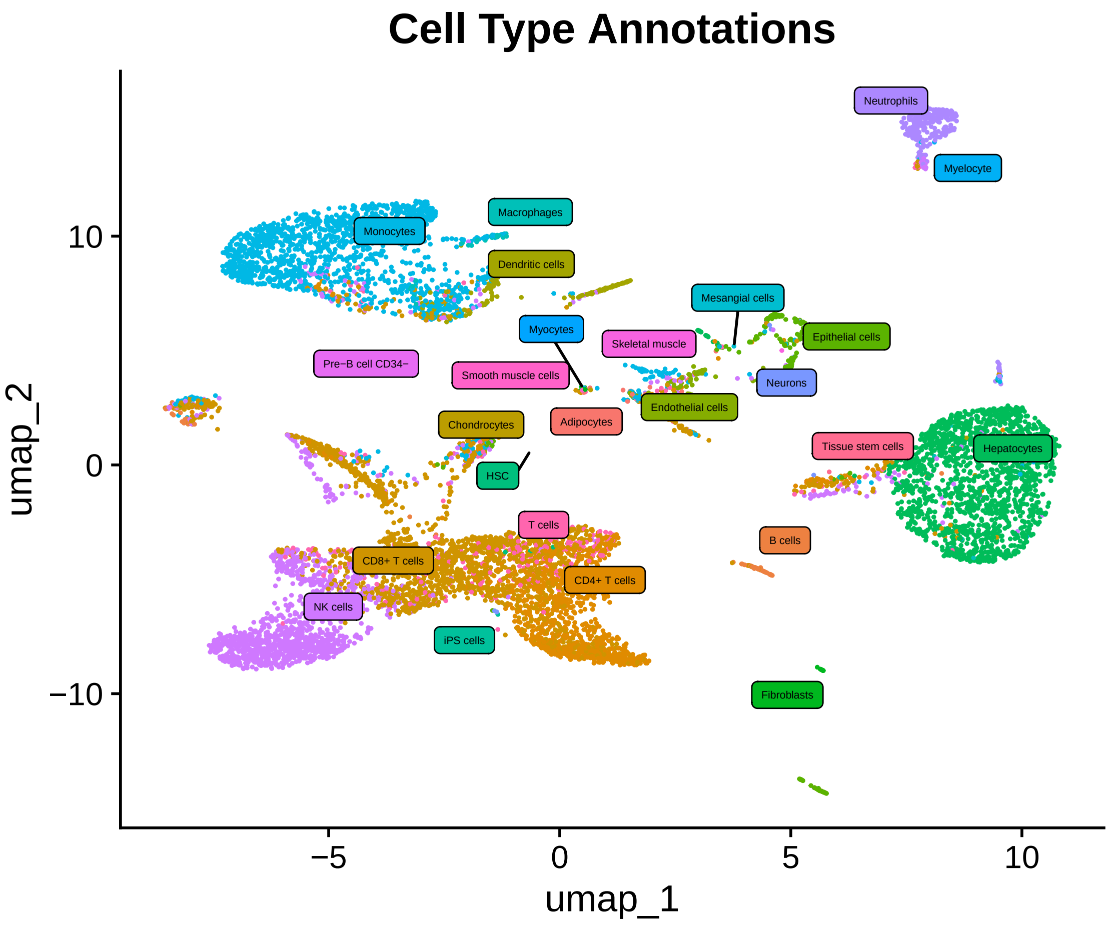
Cell Type Annotations
umap_2
umap_1
10
0
−10
−5
0
5
10
Neutrophils
Myelocyte
Macrophages
Monocytes
Dendritic cells
Mesangial cells
Myocytes
Skeletal muscle
Epithelial cells
Pre−B cell CD34−
Smooth muscle cells
Neurons
Endothelial cells
Chondrocytes
Adipocytes
Tissue stem cells
Hepatocytes
HSC
T cells
B cells
CD8+ T cells
CD4+ T cells
NK cells
iPS cells
Fibroblasts

#### Sample: G0013

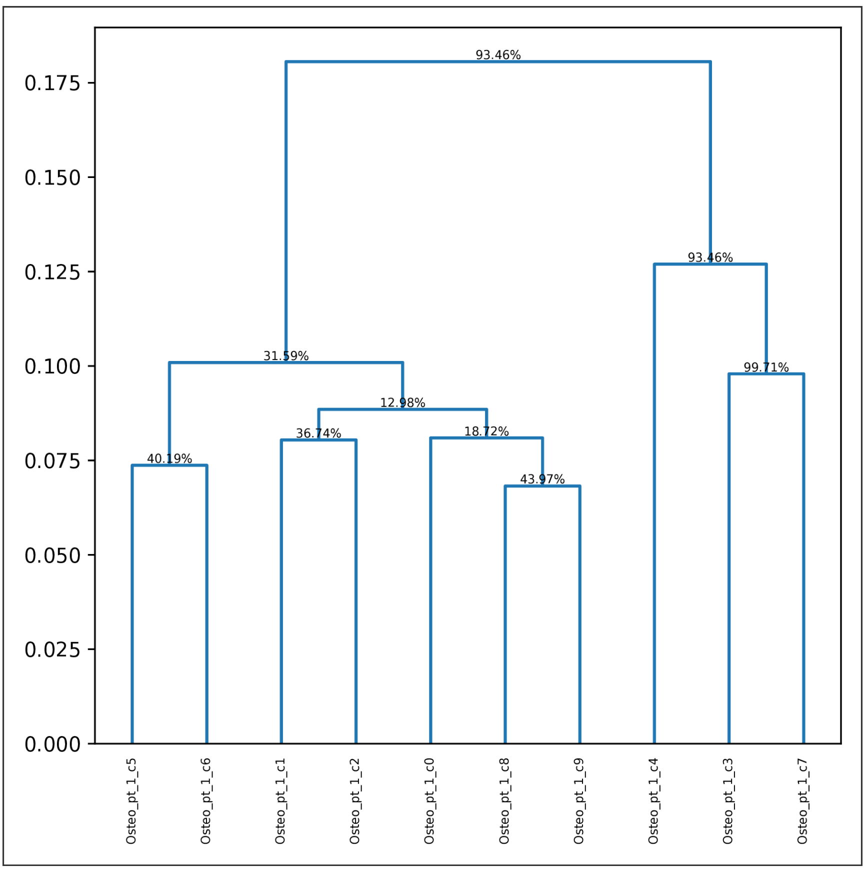


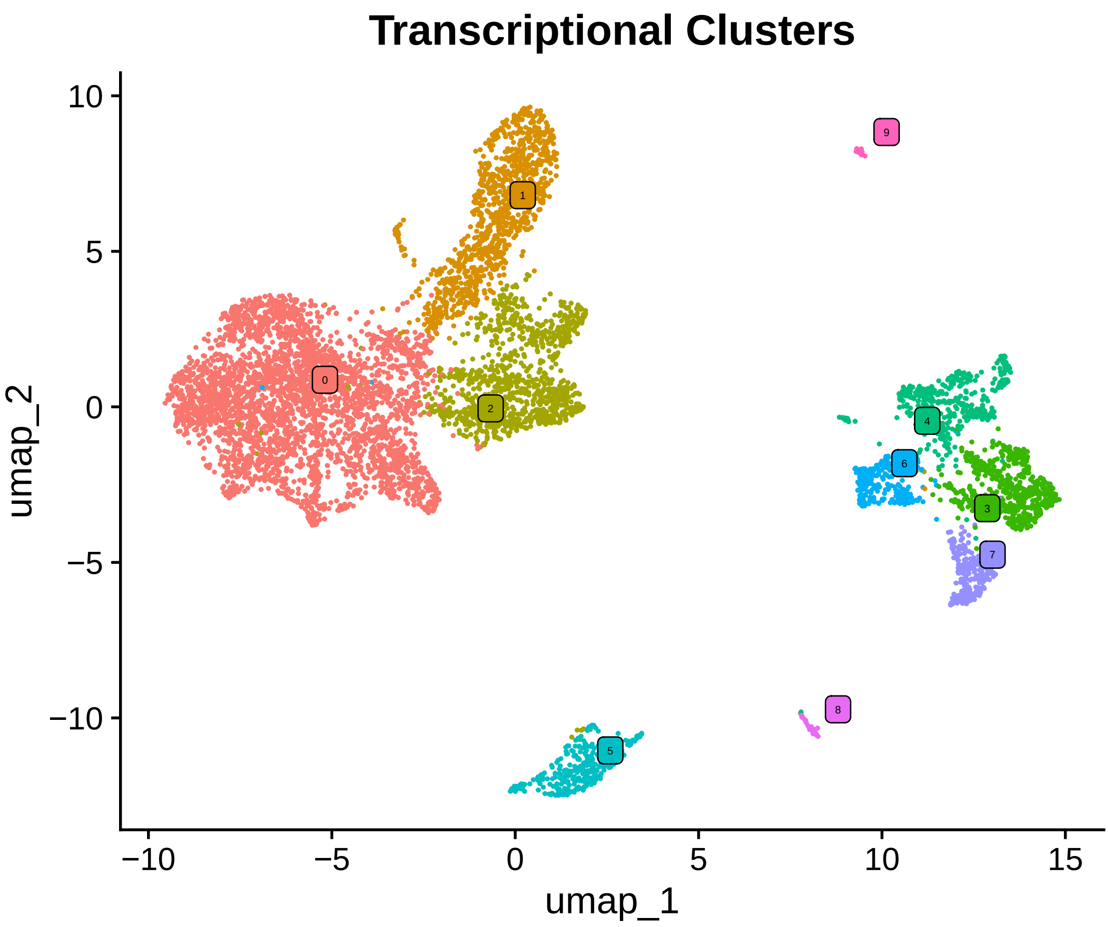


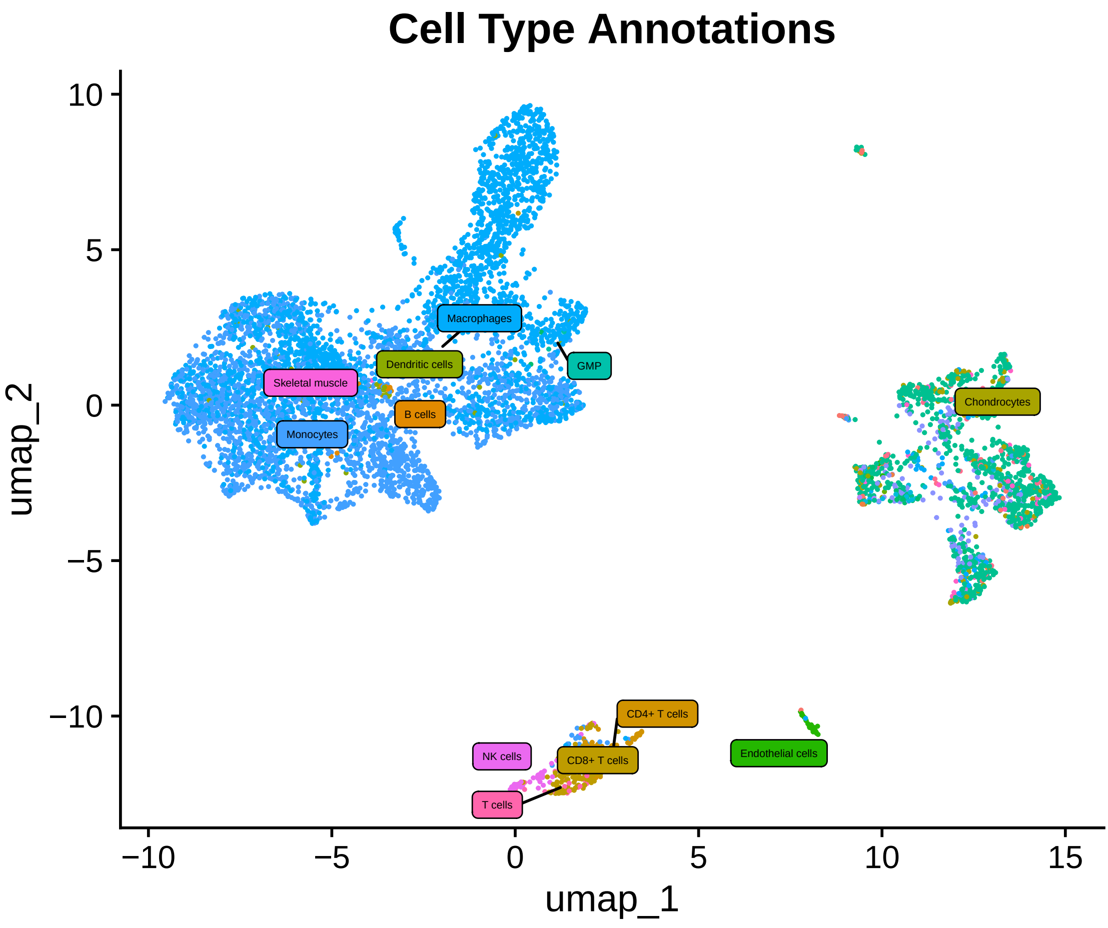

Sample: G0014

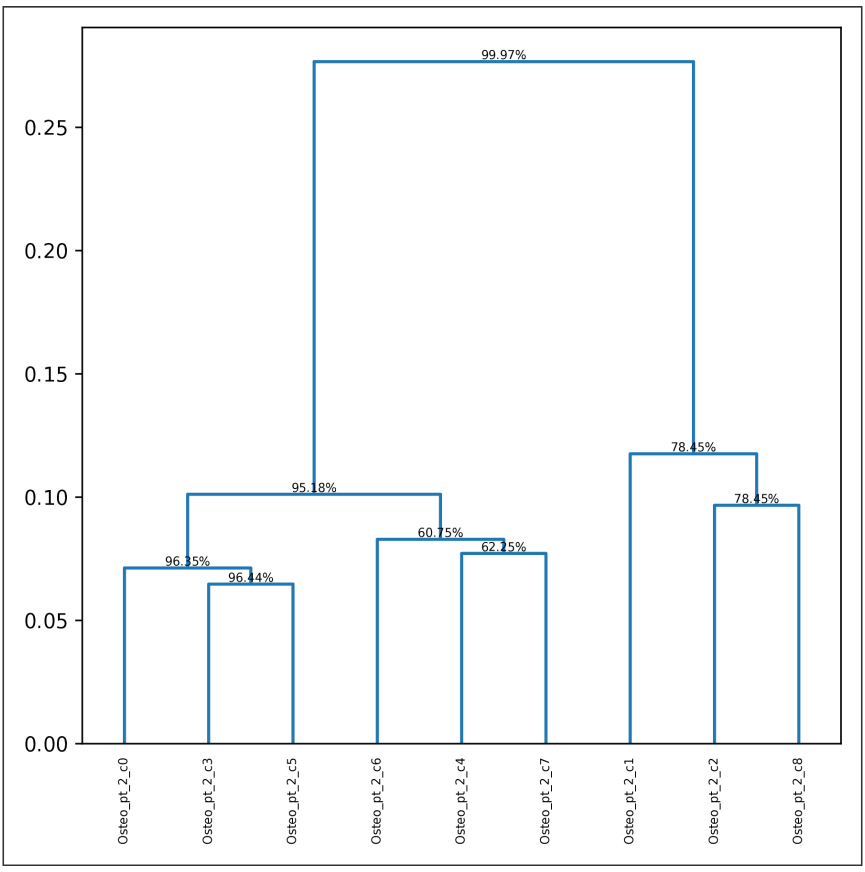


#### Transcriptional Clusters

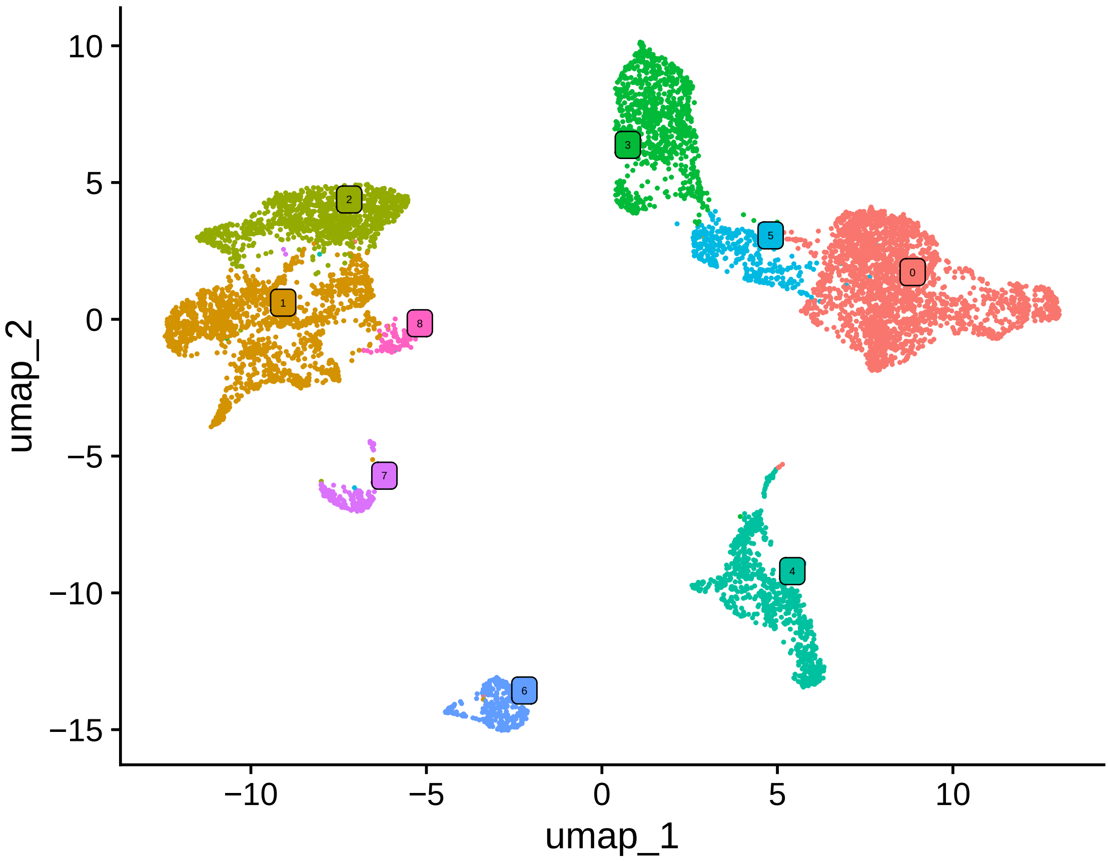


#### Cell Type Annotations

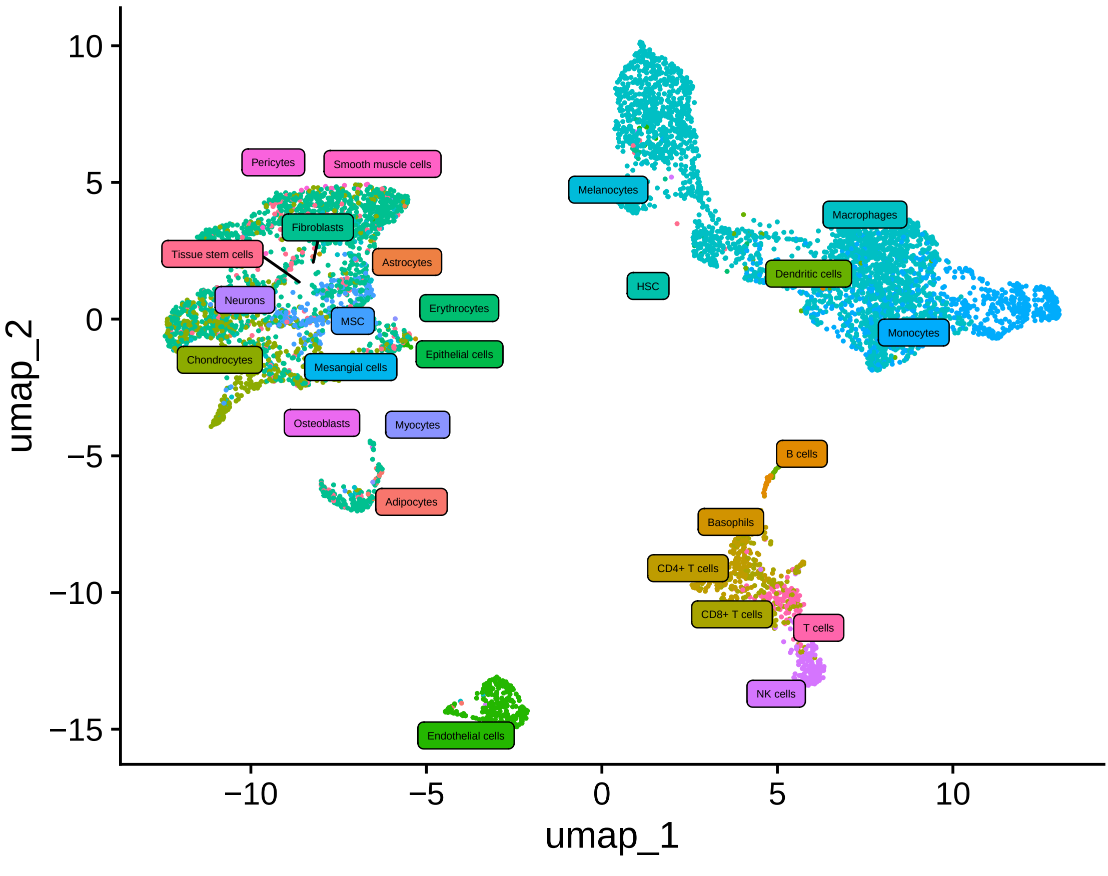

#### Sample: G0015

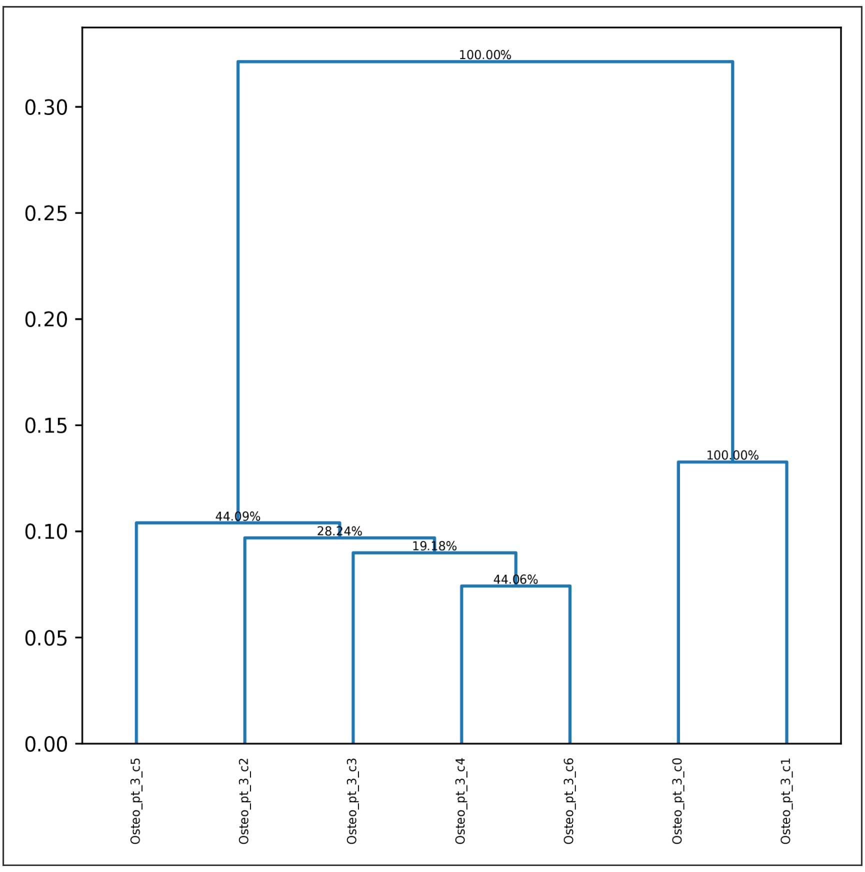


#### Transcriptional Clusters

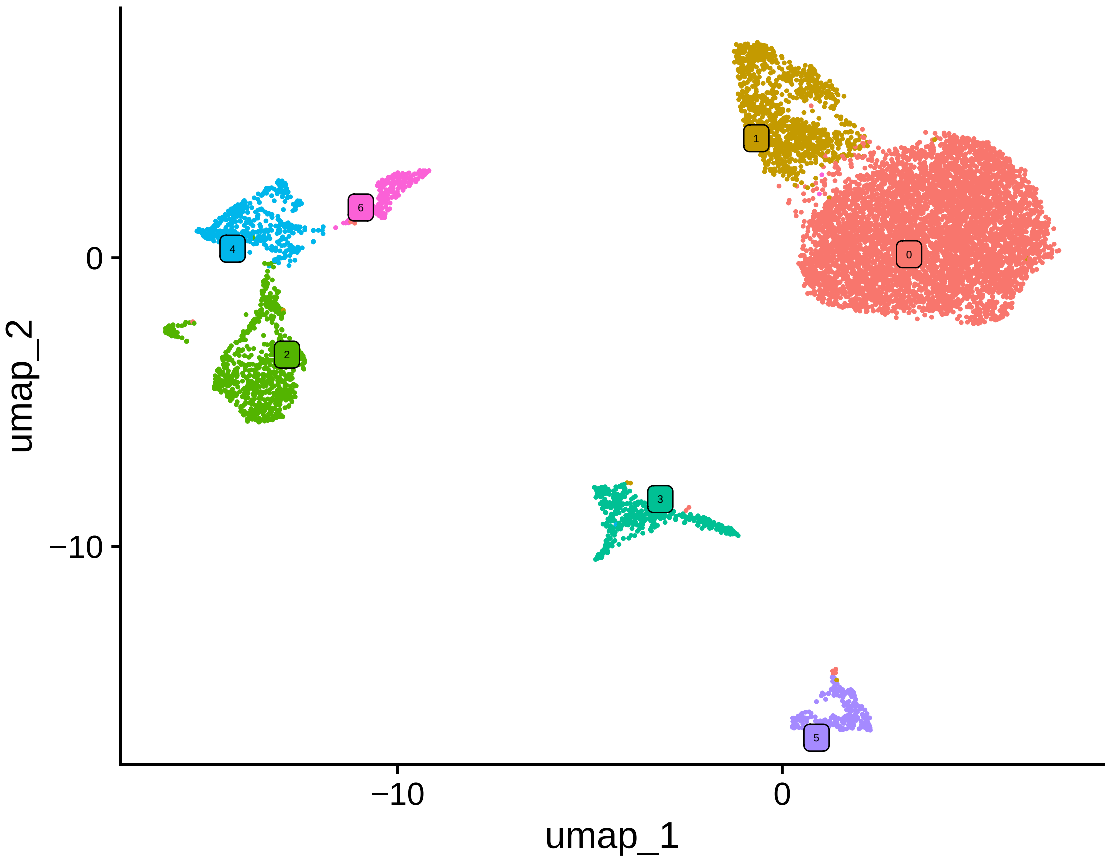


#### Cell Type Annotations

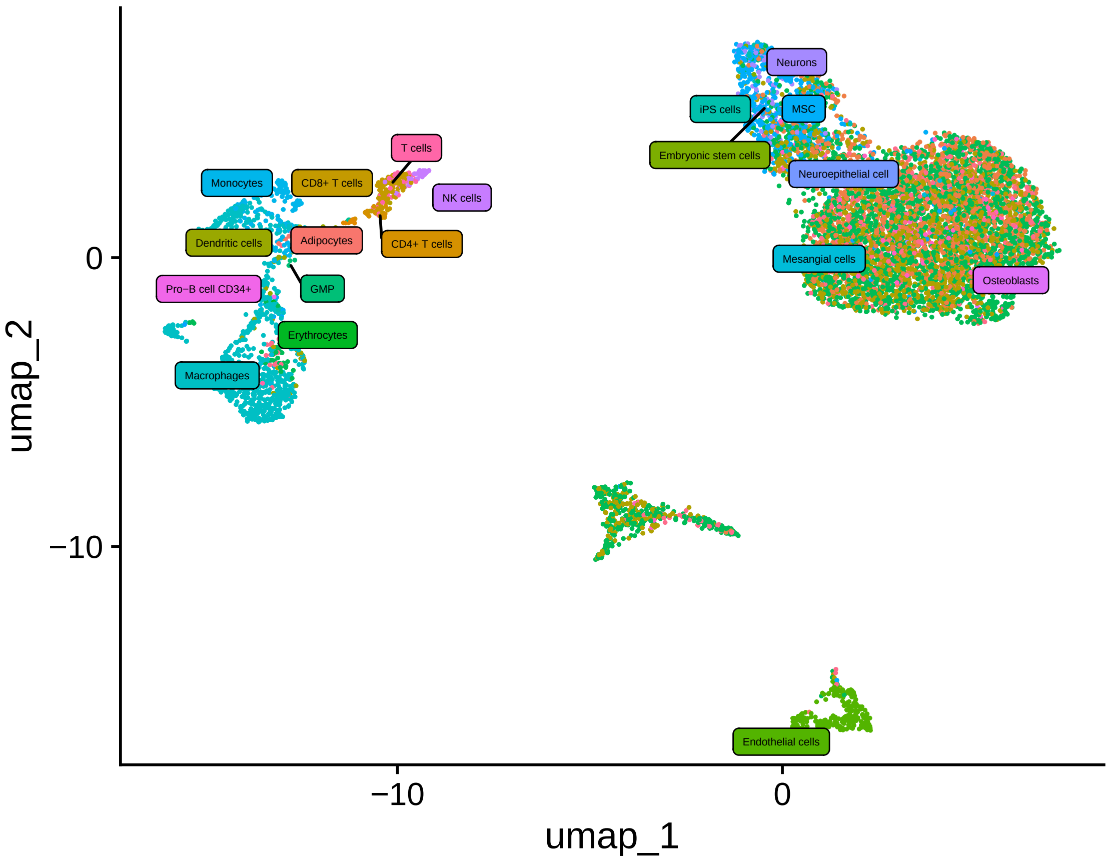

Sample: G0016

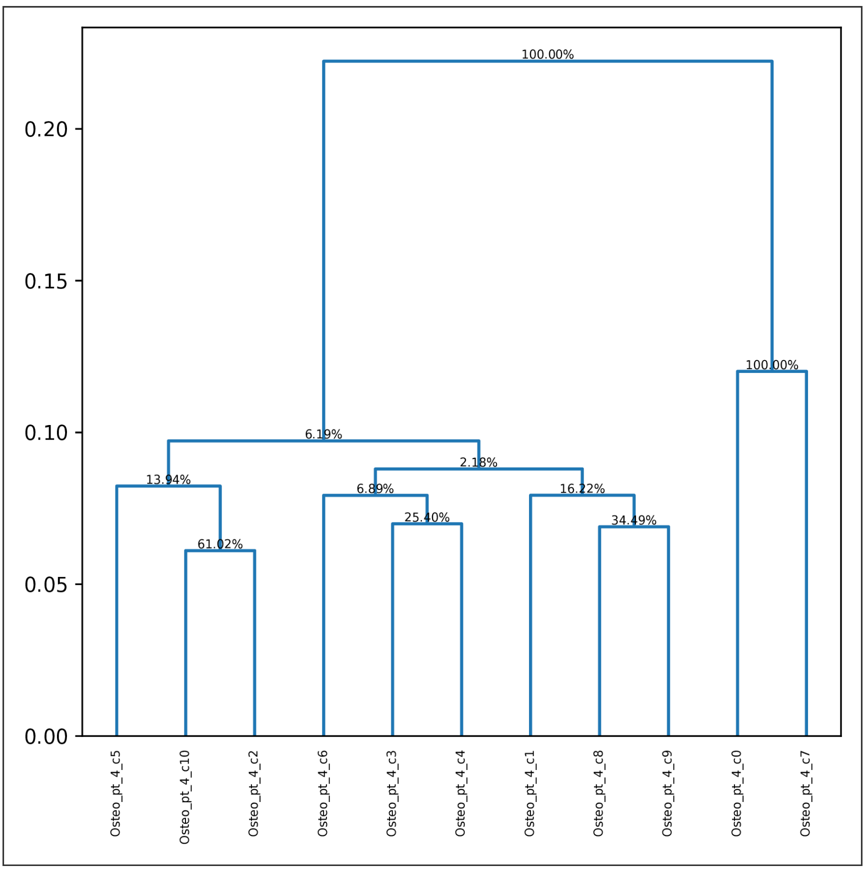


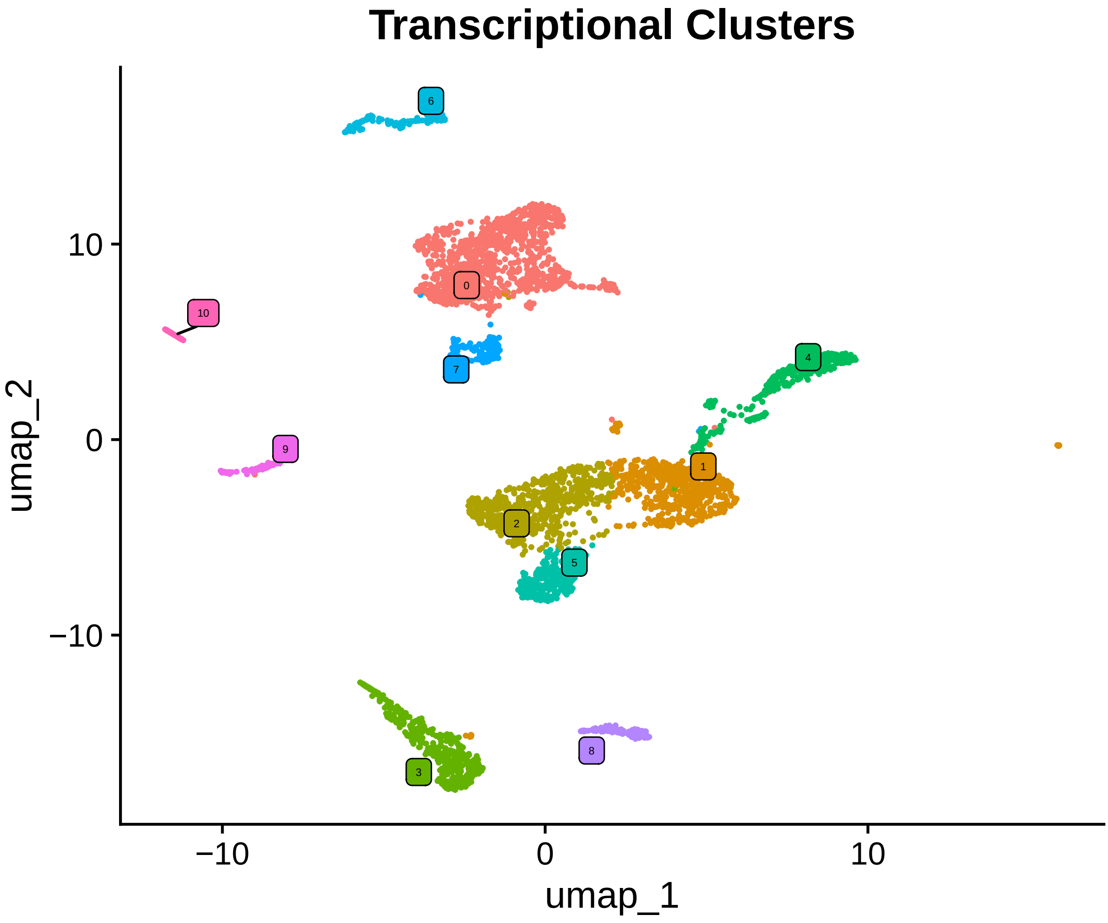


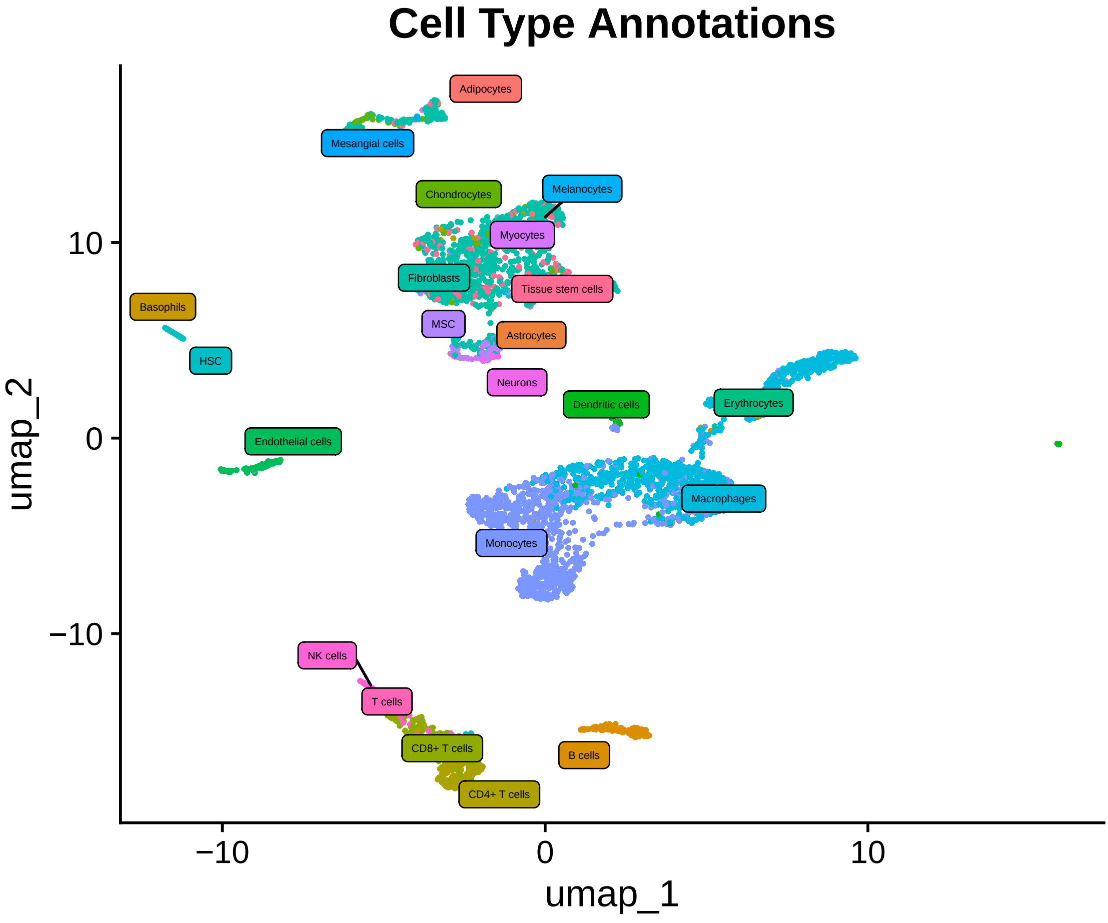

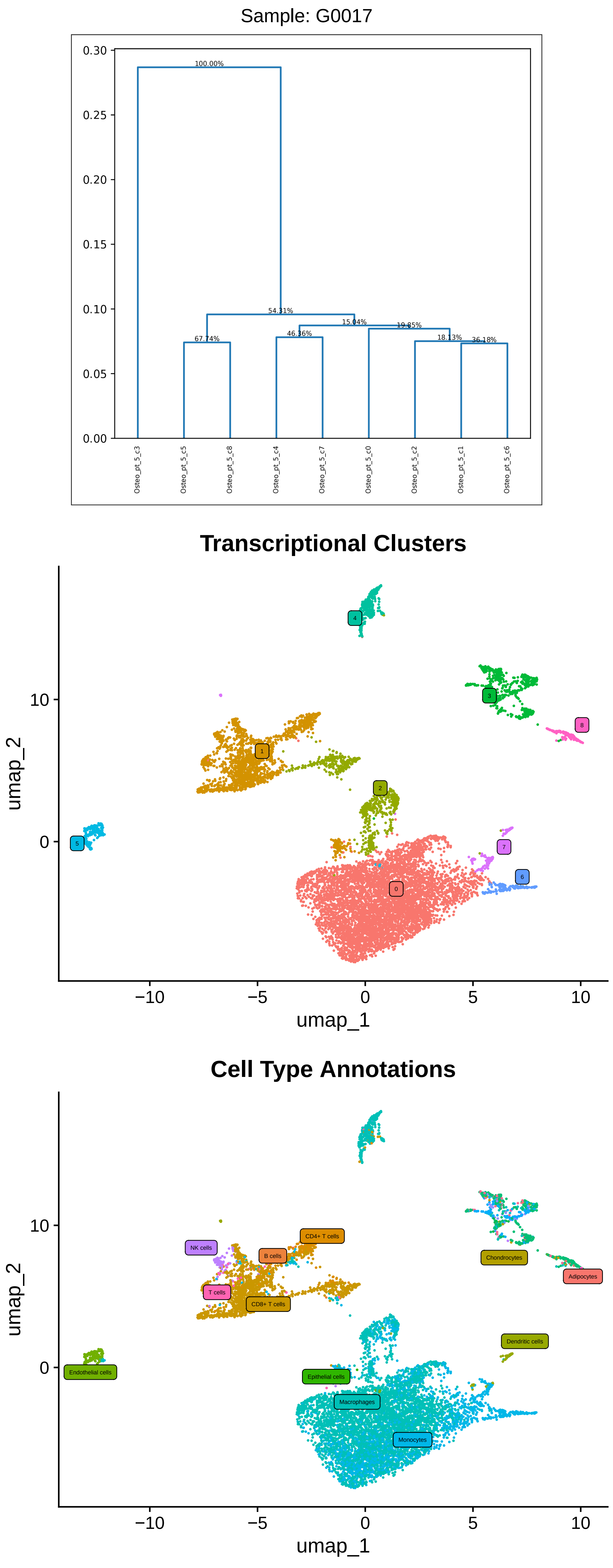
Sample: G0017
0.30
0.25
0.20
0.15
0.10
0.05
0.00
100.00%
54.31%
67.74%
46.36%
15.04%
19.85%
18.13%
36.18%
Osteo_pt_5_c3
Osteo_pt_5_c5
Osteo_pt_5_c8
Osteo_pt_5_c4
Osteo_pt_5_c7
Osteo_pt_5_c0
Osteo_pt_5_c2
Osteo_pt_5_c1
Osteo_pt_5_c6
Transcriptional Clusters
umap_2
umap_1
10
0
−10
−5
0
5
10
0
1
2
3
4
5
6
7
8
Cell Type Annotations
umap_2
umap_1
10
0
−10
−5
0
5
10
NK cells
B cells
CD4+ T cells
T cells
CD8+ T cells
Chondrocytes
Adipocytes
Dendritic cells
Endothelial cells
Epithelial cells
Macrophages
Monocytes

Sample: G0018

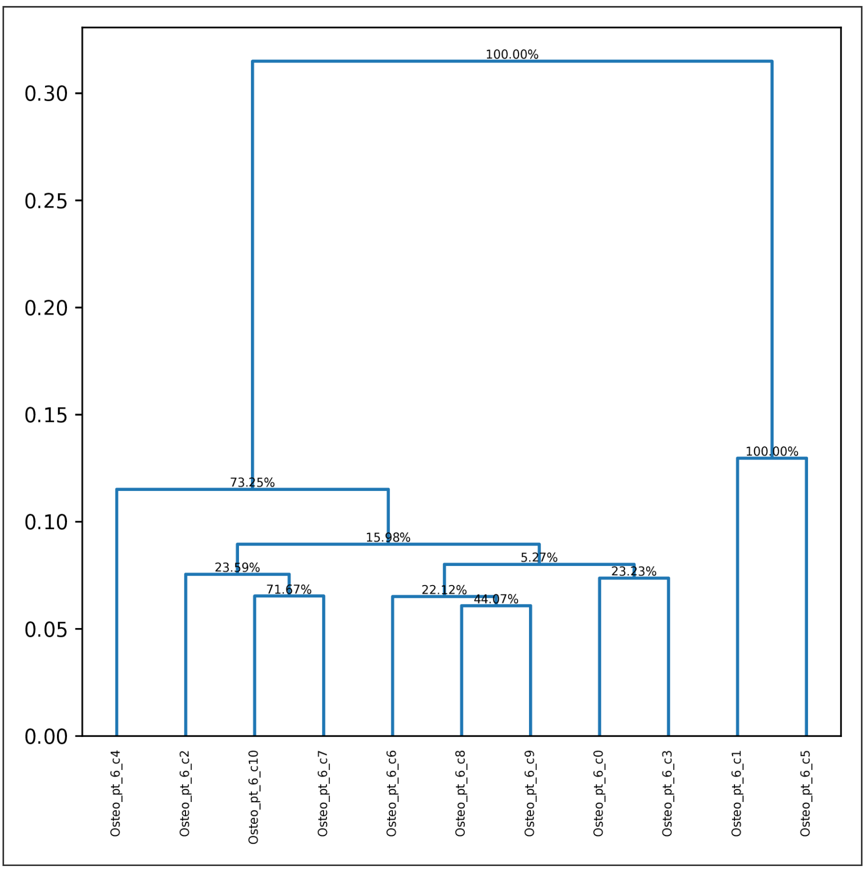


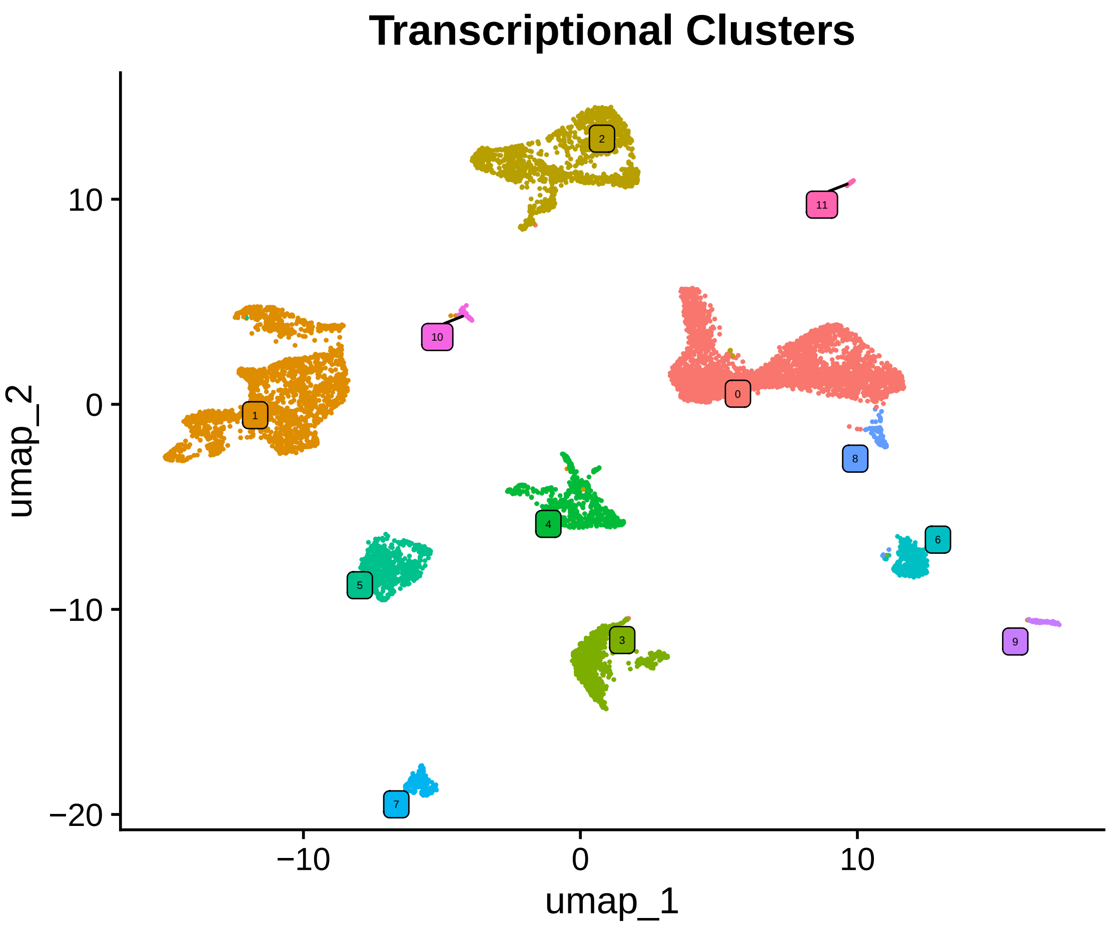


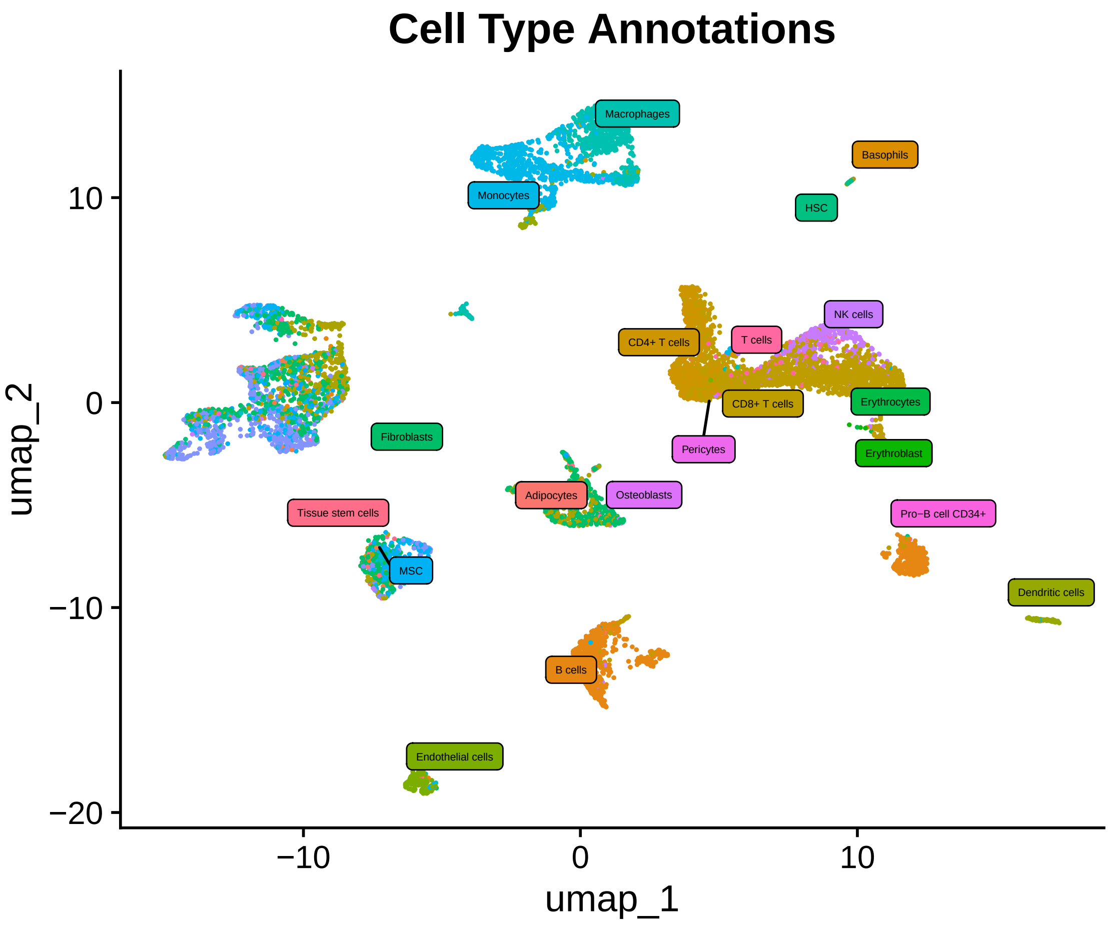

Sample: G0019

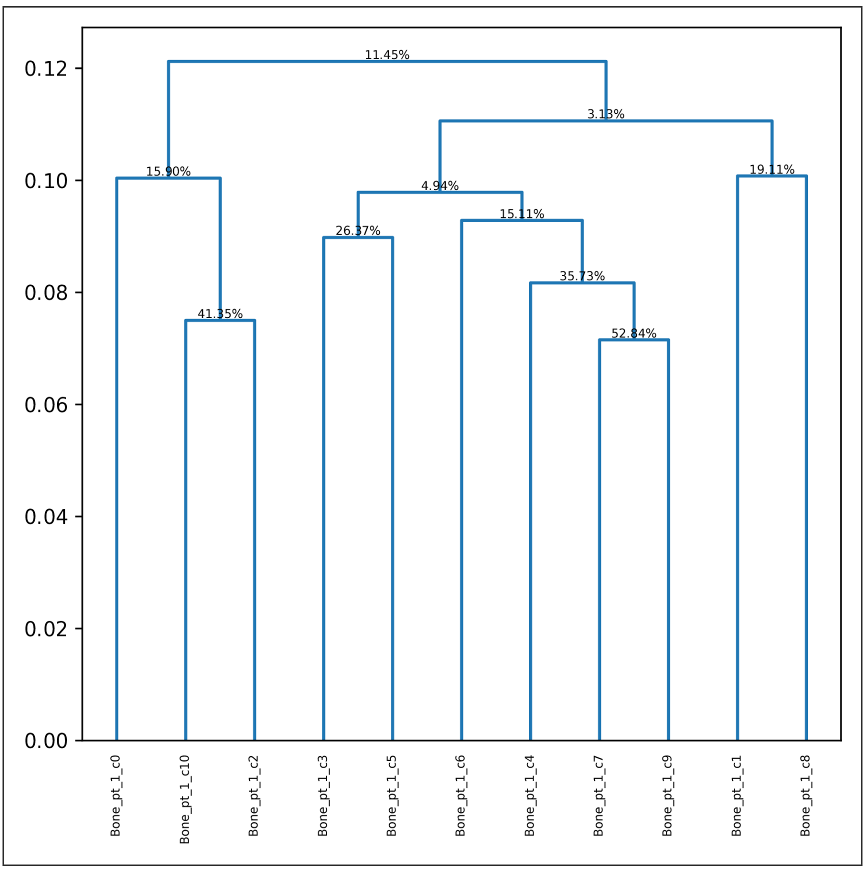


#### Transcriptional Clusters

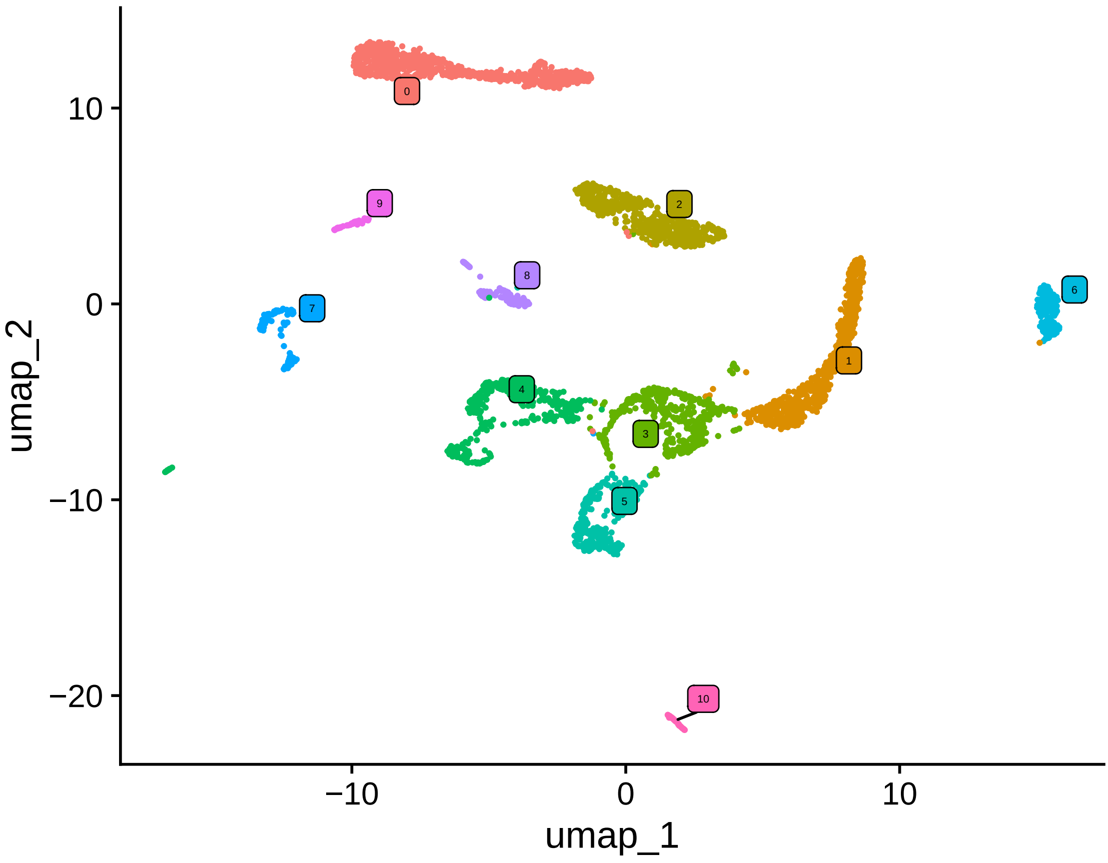


#### Cell Type Annotations

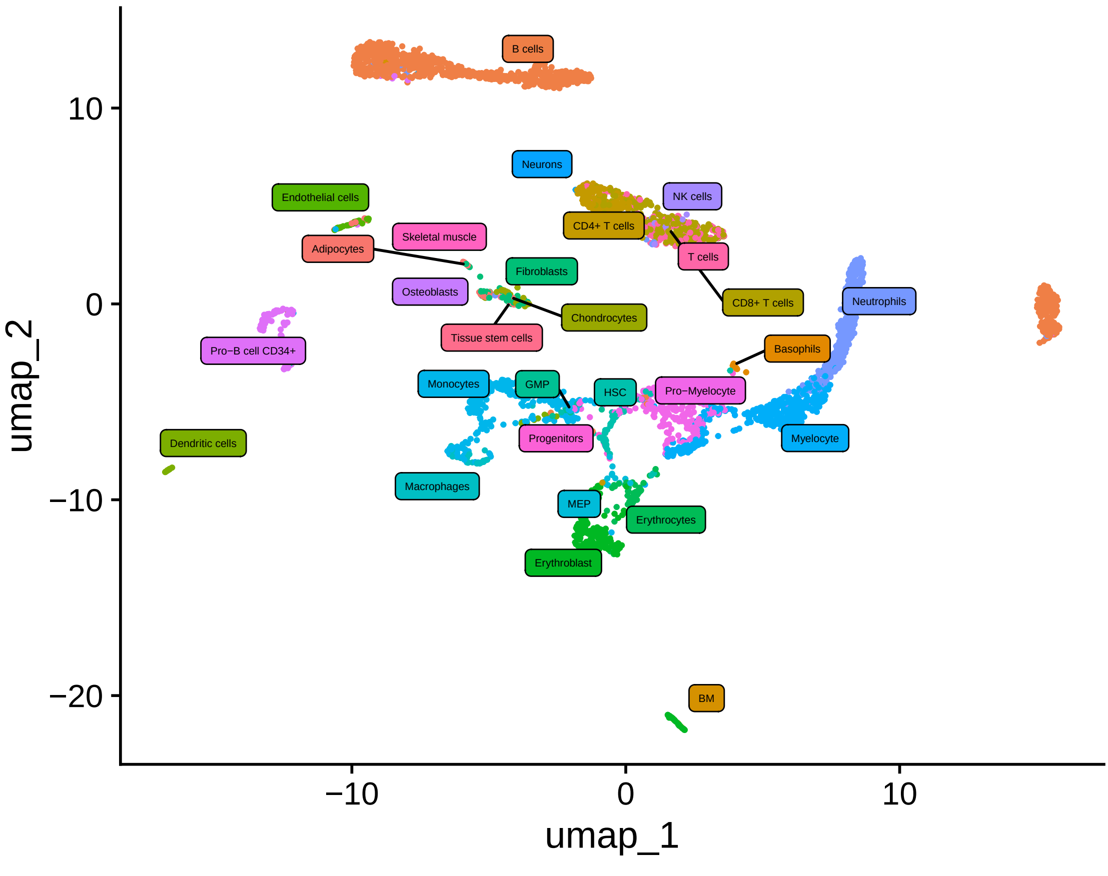

#### Sample: G0020

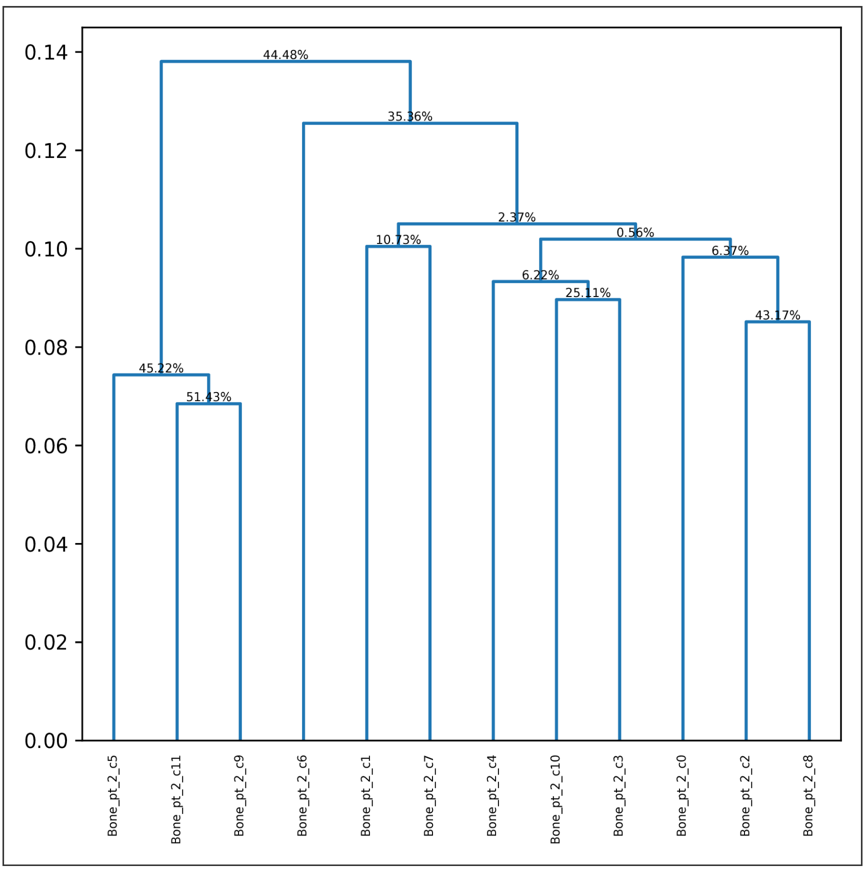


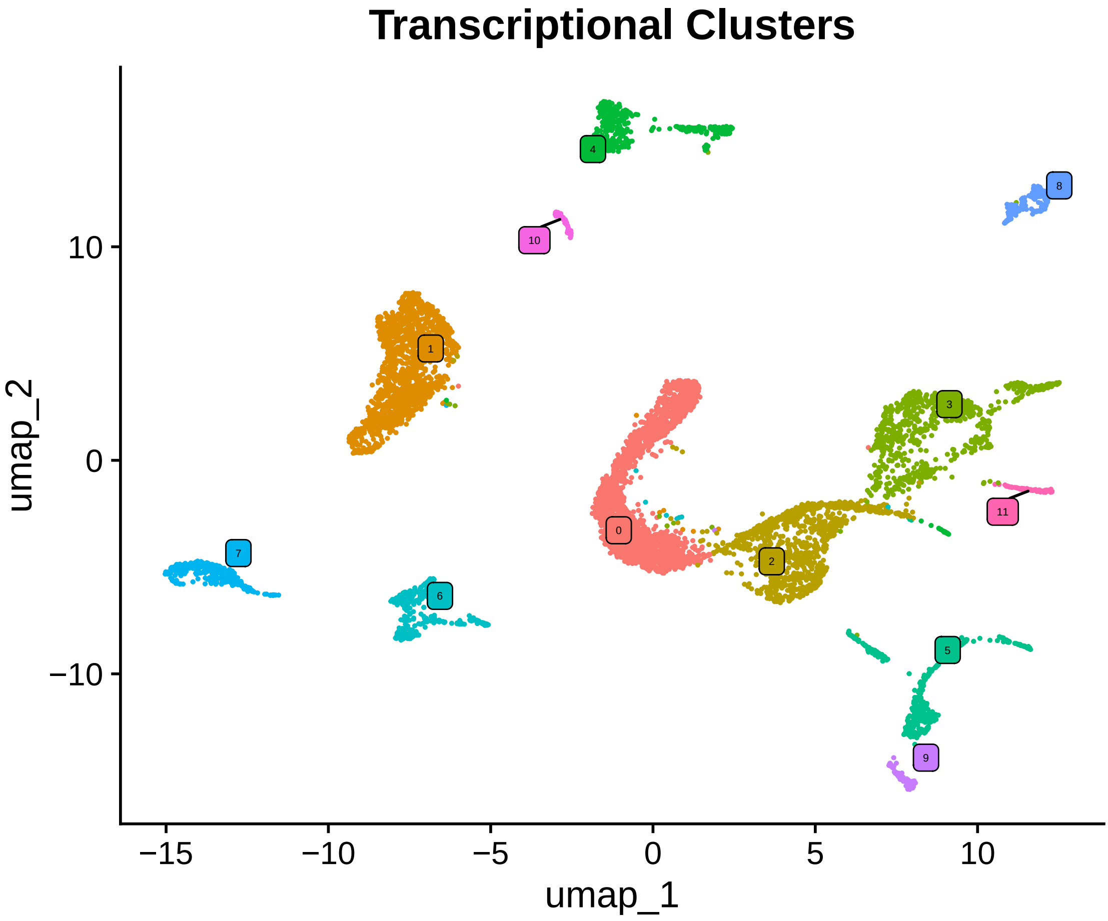


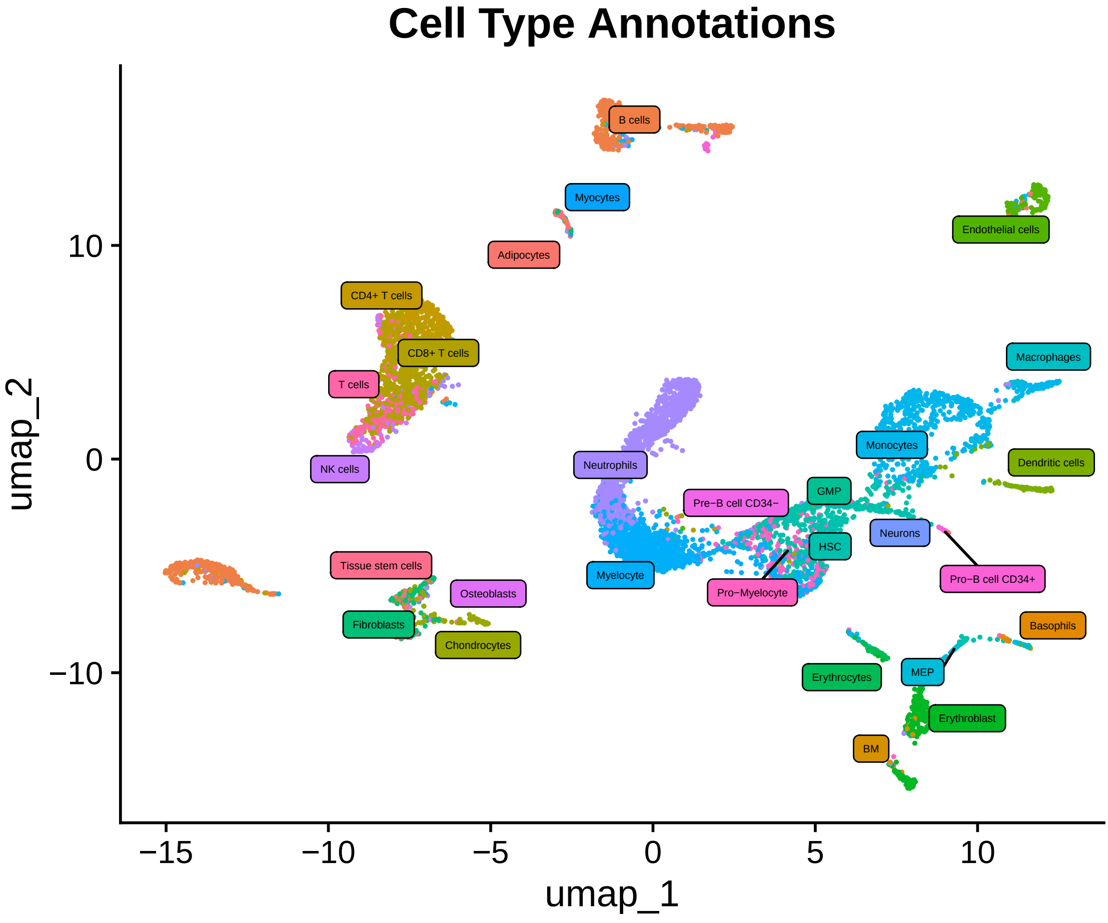

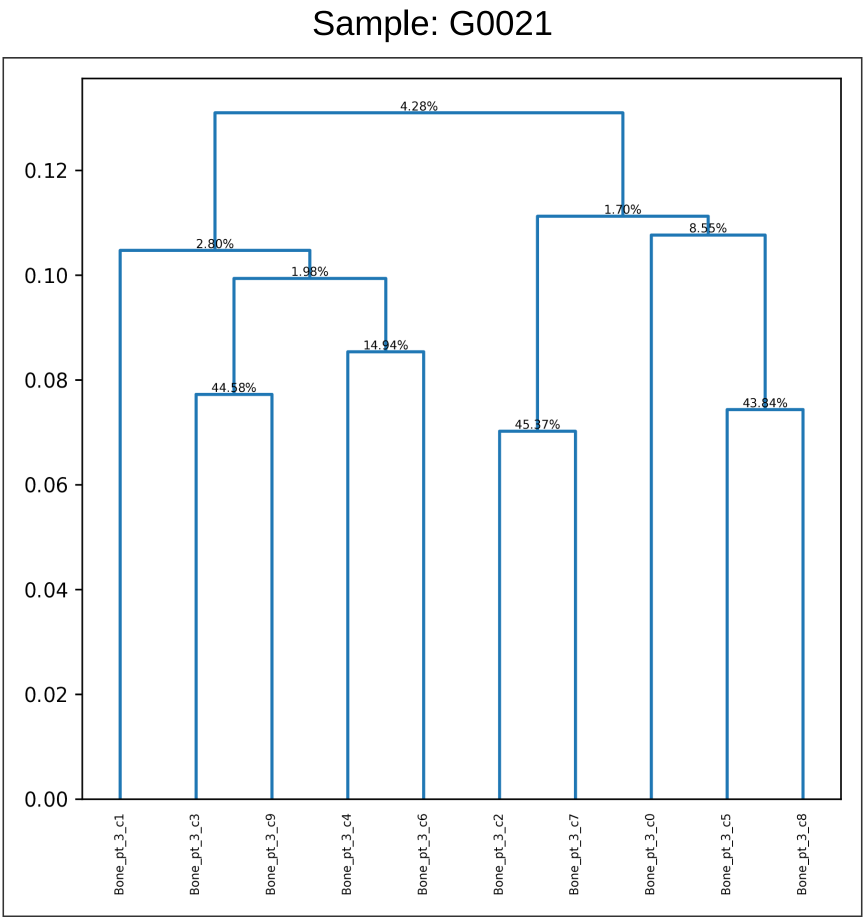
Sample: G0021
4.28%
2.80%
1.98%
14.94%
44.58%
1.70%
8.55%
45.37%
43.84%
0.12
0.10
0.08
0.06
0.04
0.02
0.00
Bone_pt_3_c1
Bone_pt_3_c3
Bone_pt_3_c9
Bone_pt_3_c4
Bone_pt_3_c6
Bone_pt_3_c2
Bone_pt_3_c7
Bone_pt_3_c0
Bone_pt_3_c5
Bone_pt_3_c8

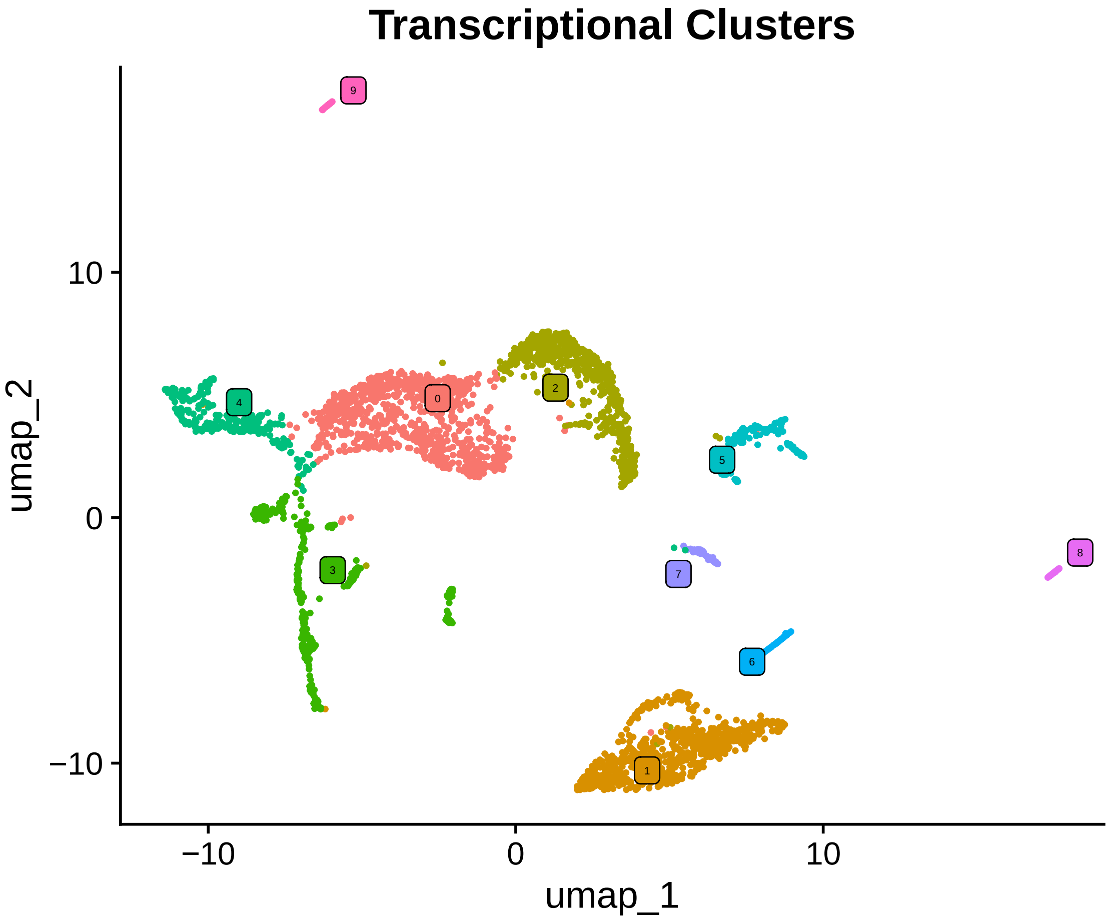
Transcriptional Clusters
9
4
0
2
5
3
7
8
6
1
10
0
−10
umap_2
−10
0
10
umap_1

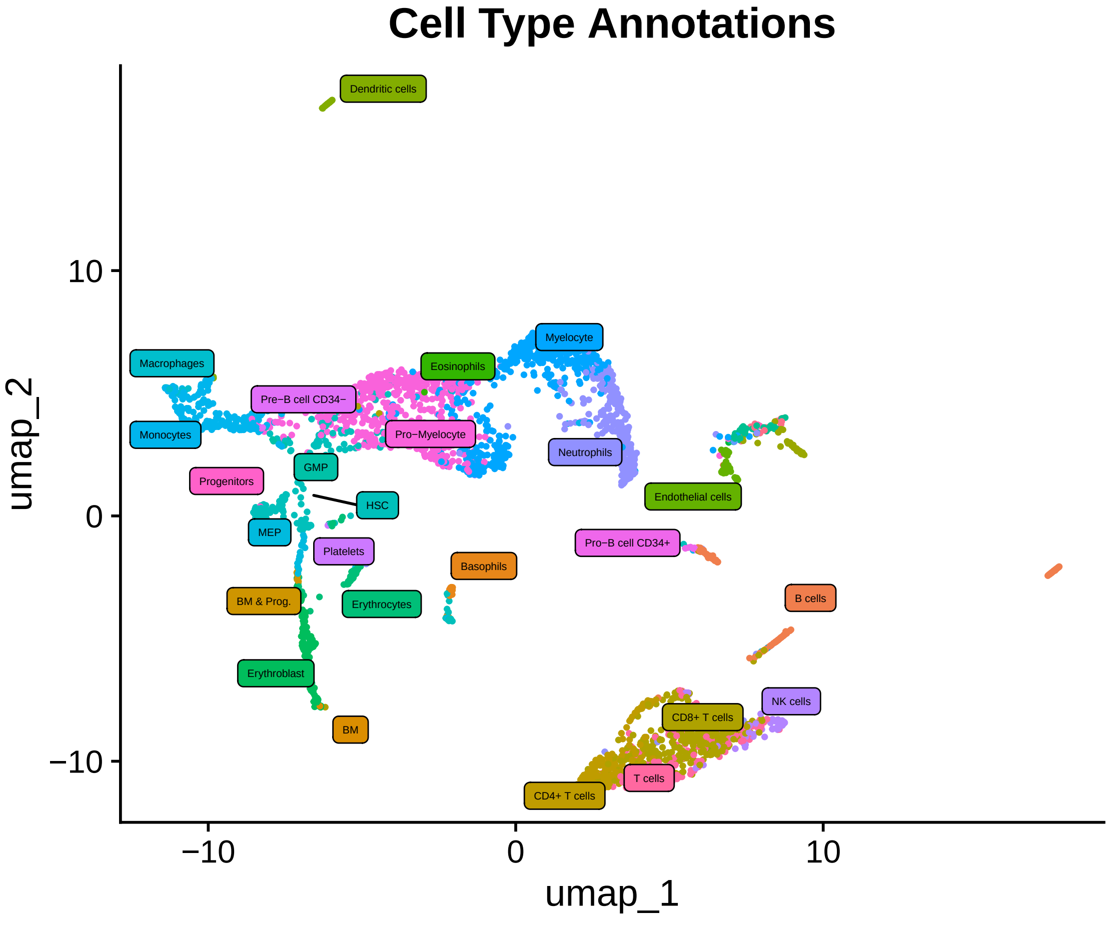
Cell Type Annotations
Dendritic cells
Macrophages
Eosinophils
Myelocyte
Pre−B cell CD34−
Monocytes
Pro−Myelocyte
Neutrophils
GMP
Progenitors
HSC
Endothelial cells
MEP
Platelets
Pro−B cell CD34+
Basophils
BM & Prog.
Erythrocytes
B cells
Erythroblast
NK cells
CD8+ T cells
BM
T cells
CD4+ T cells
10
0
−10
umap_2
−10
0
10
umap_1

Sample: G0022

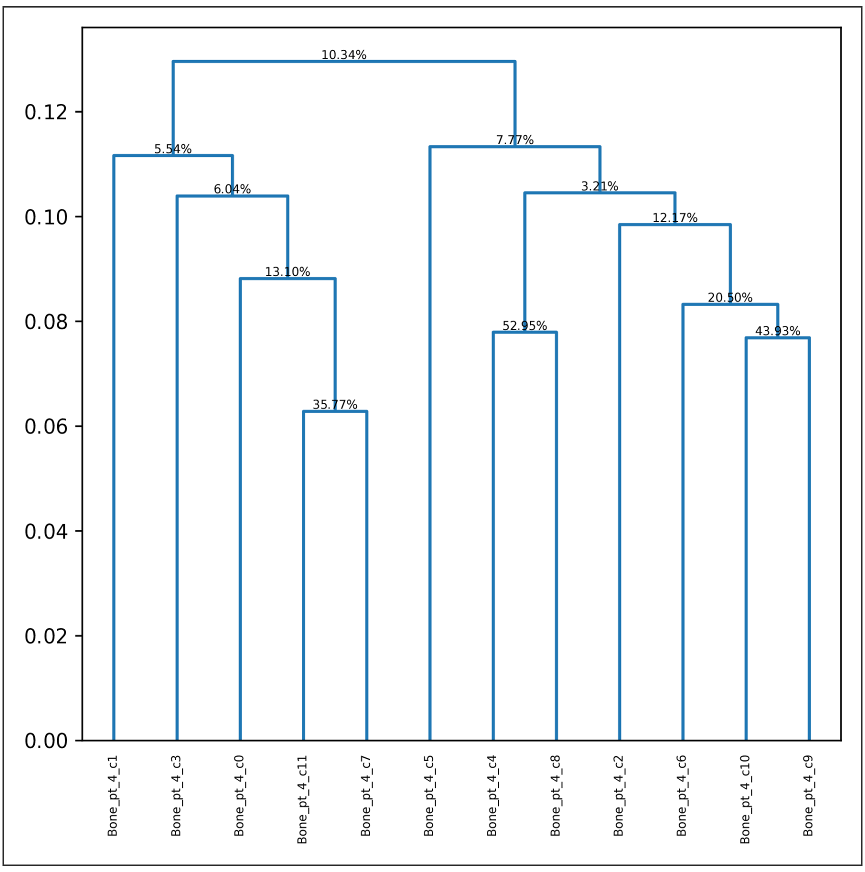


**Transcriptional Clusters**

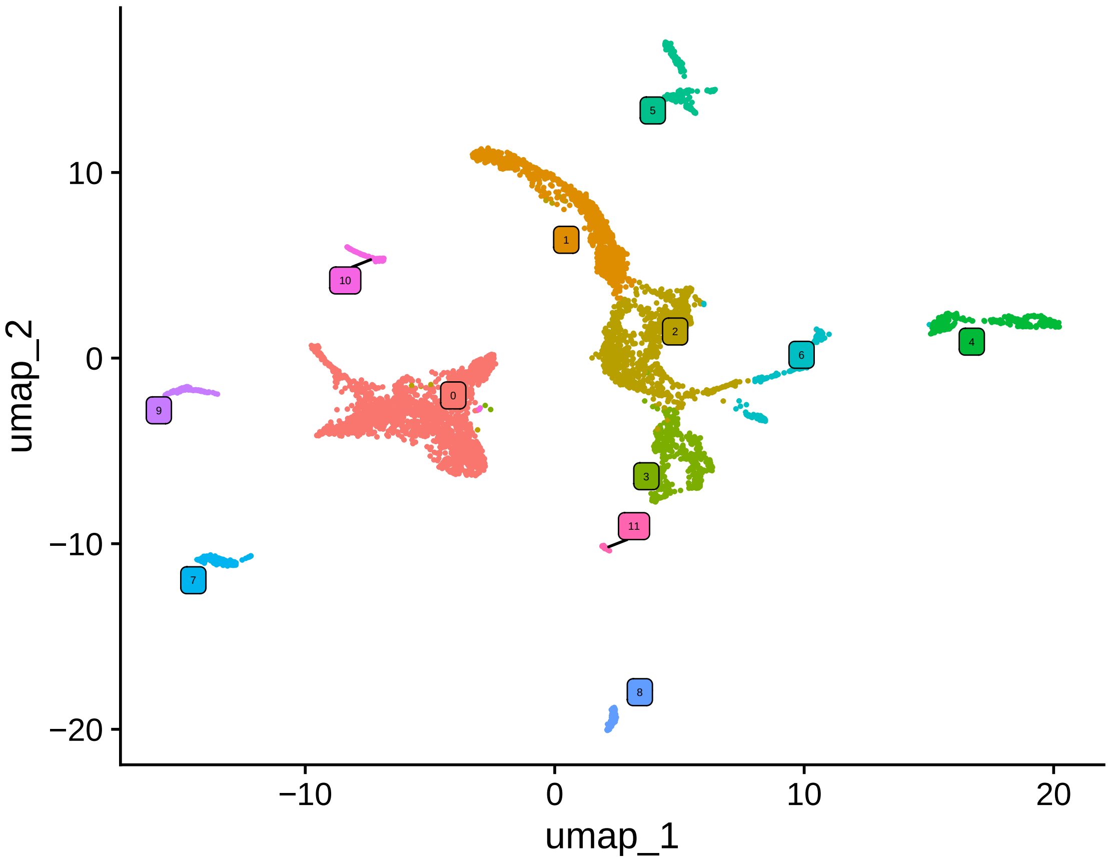


**Cell Type Annotations**

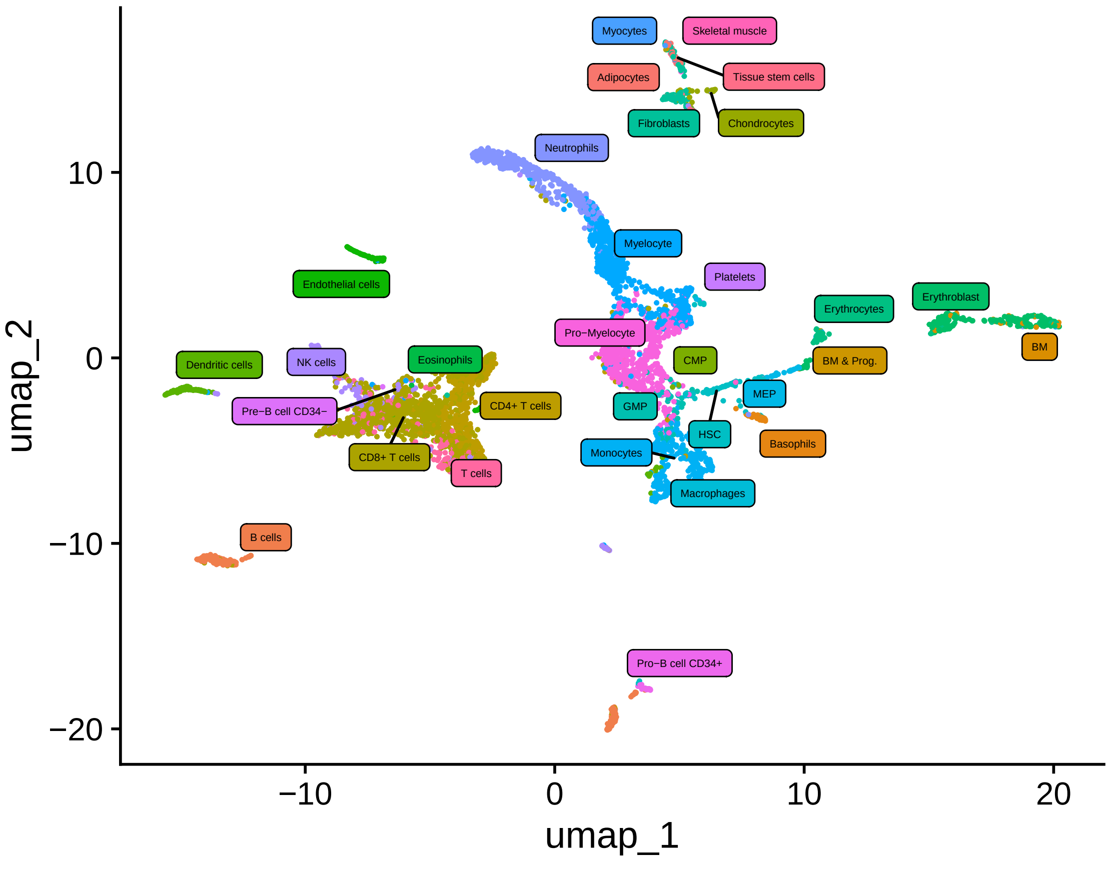

Sample: G0023

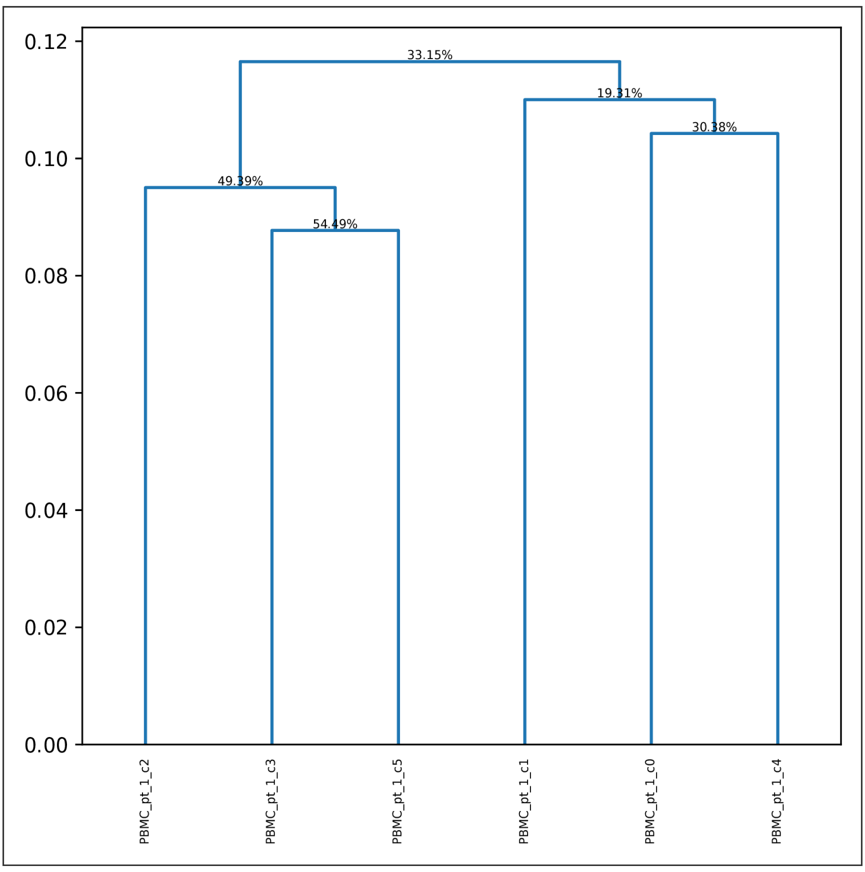


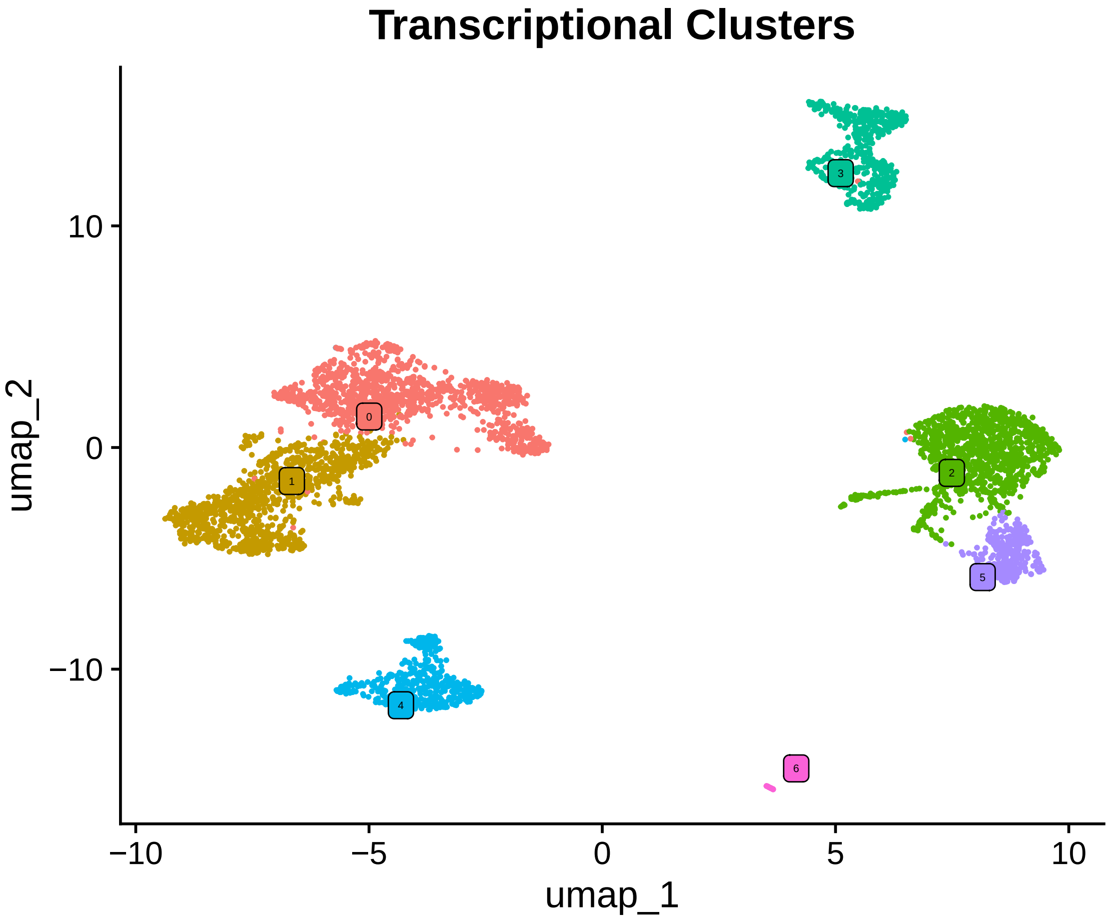


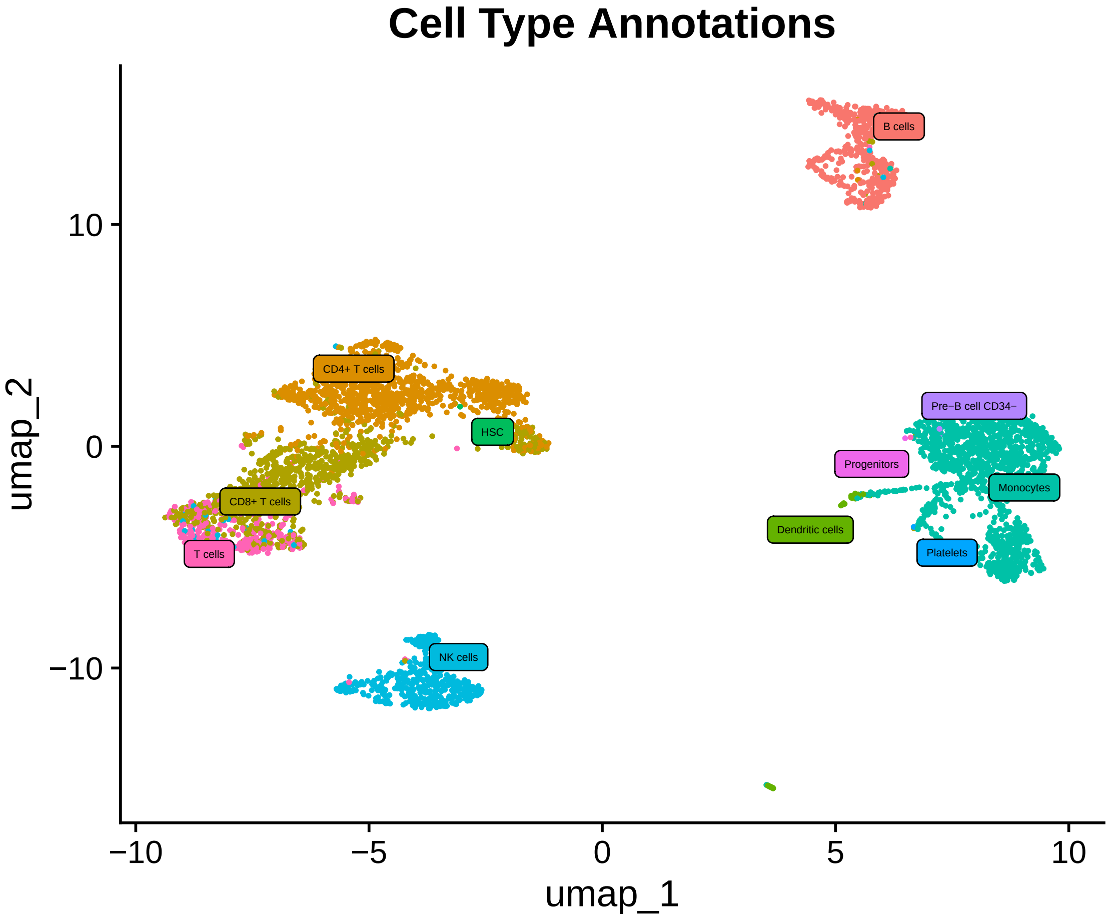

Sample: G0024

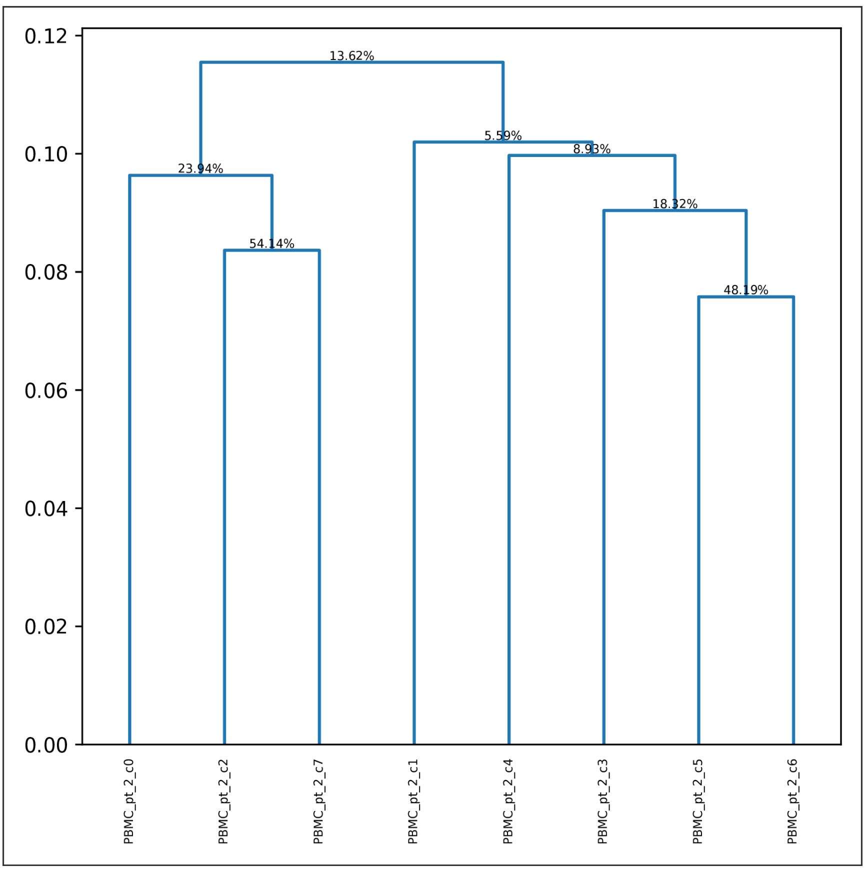


#### Transcriptional Clusters

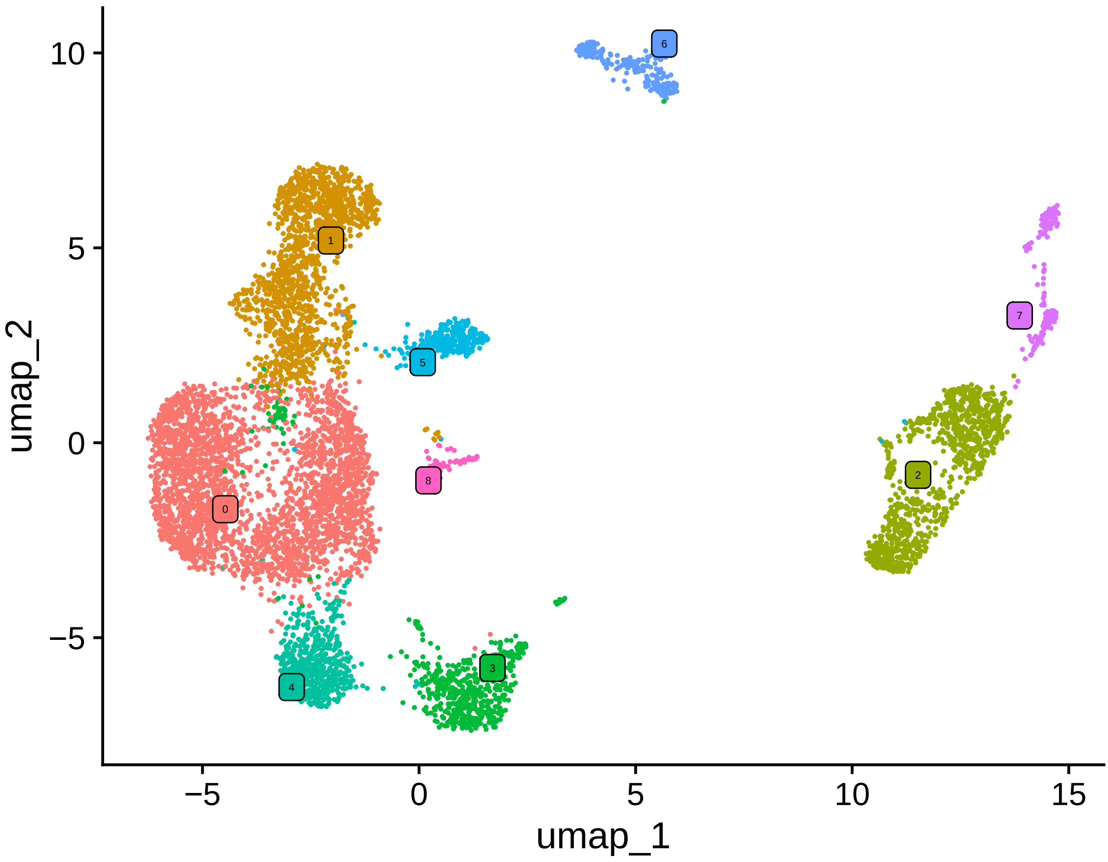


#### Cell Type Annotations

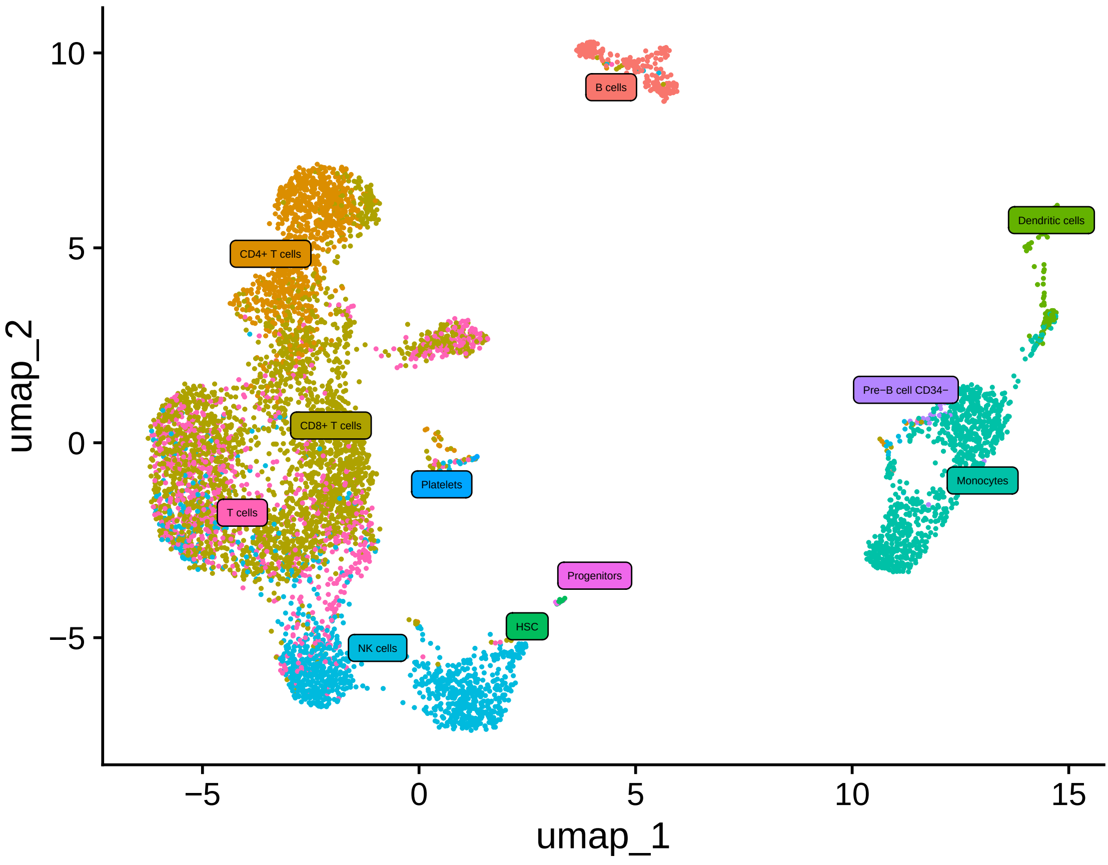

Sample: G0025

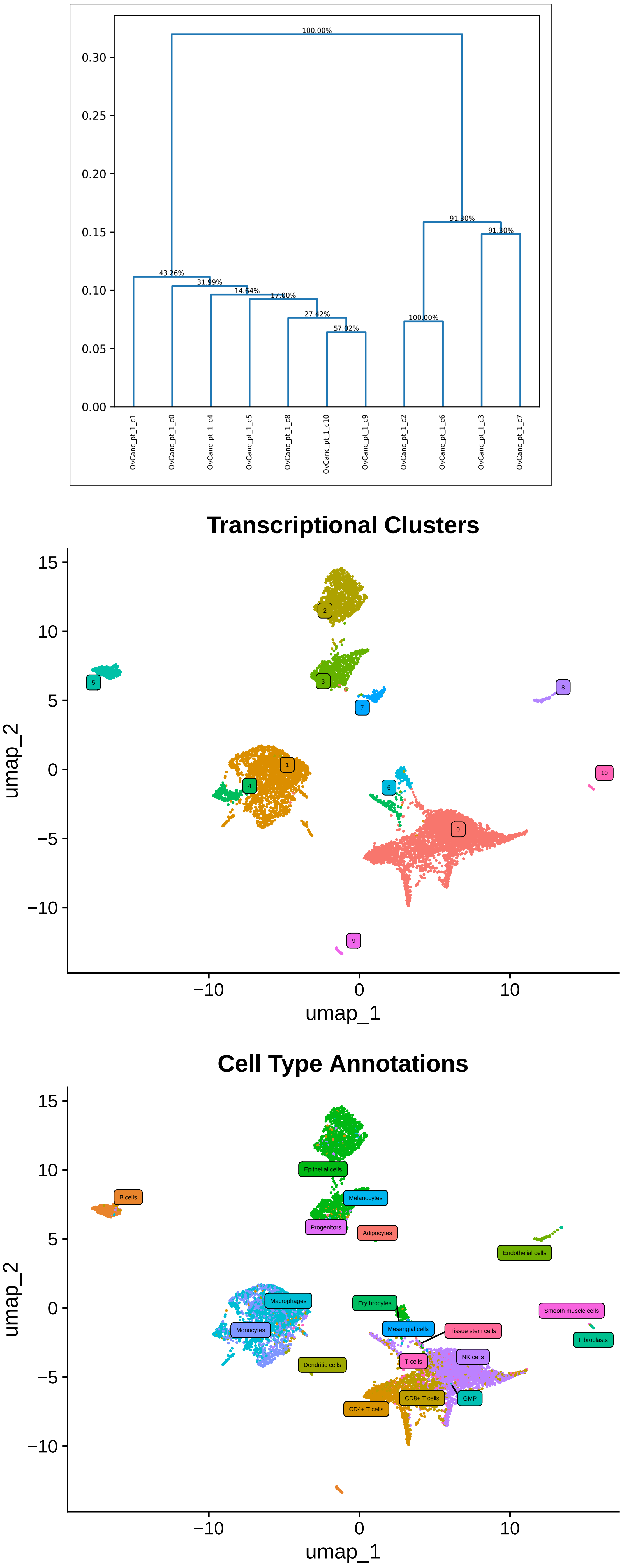

Sample: G0026

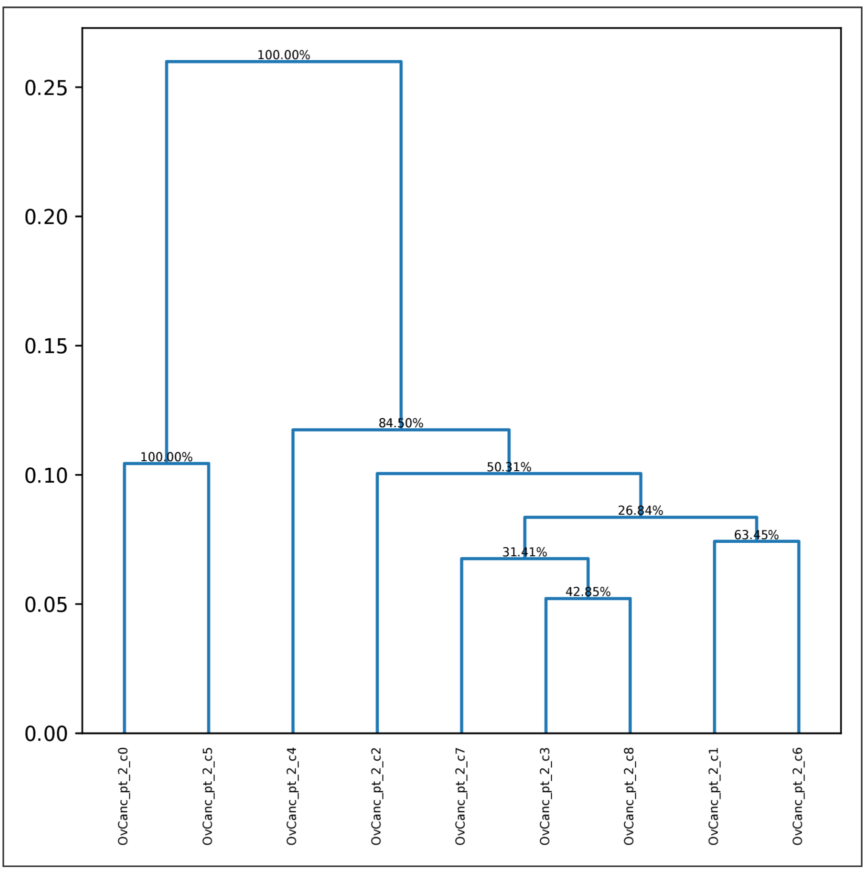


#### Transcriptional Clusters

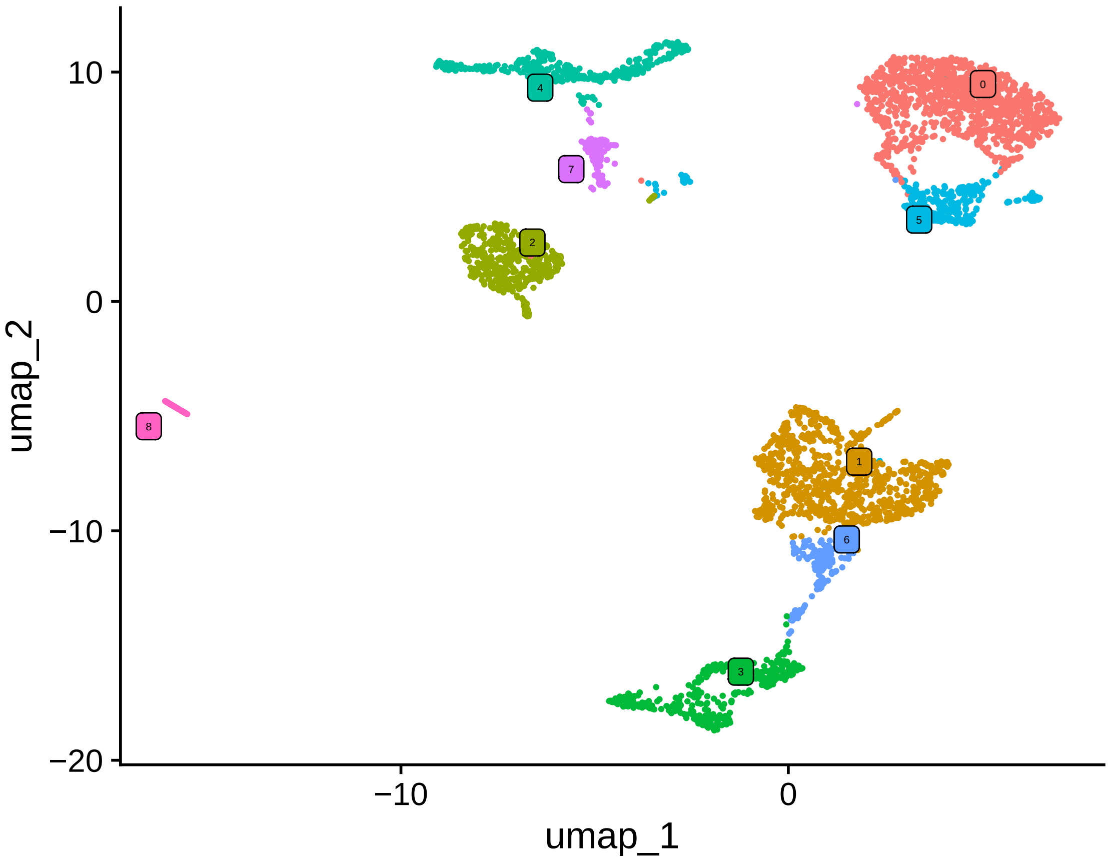


#### Cell Type Annotations

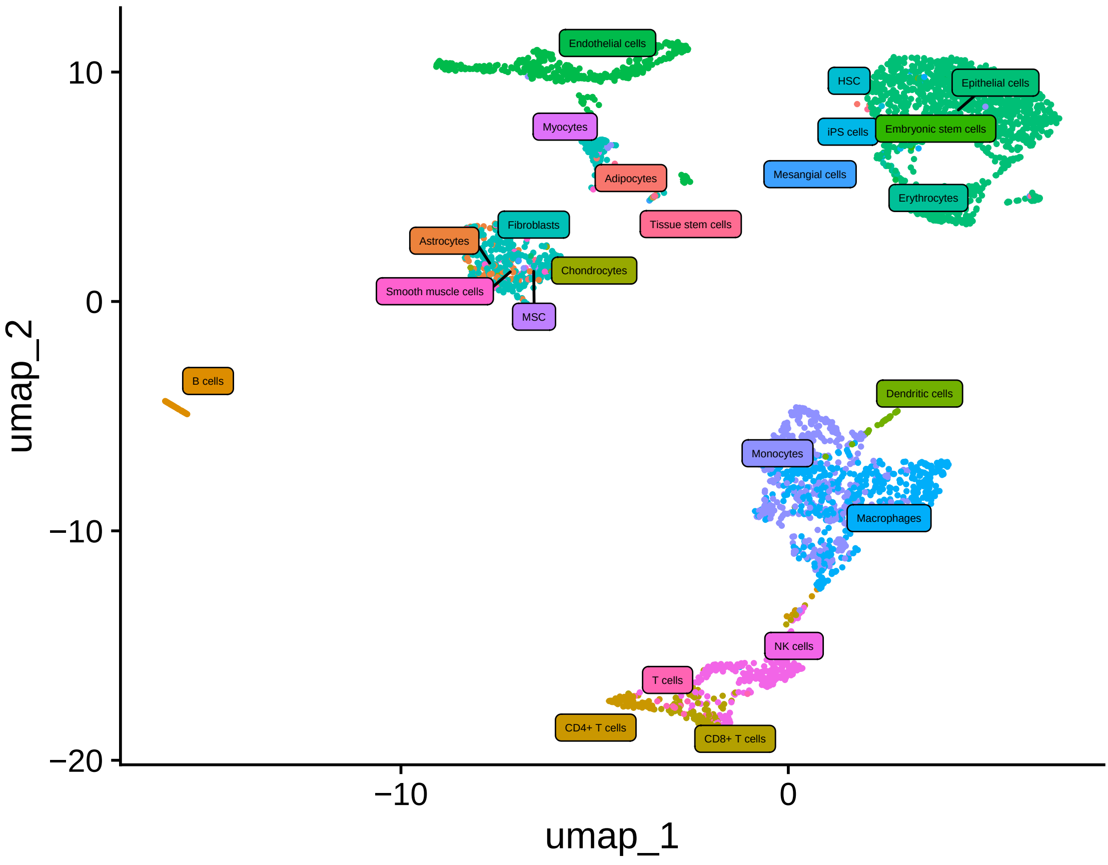

Sample: G0027

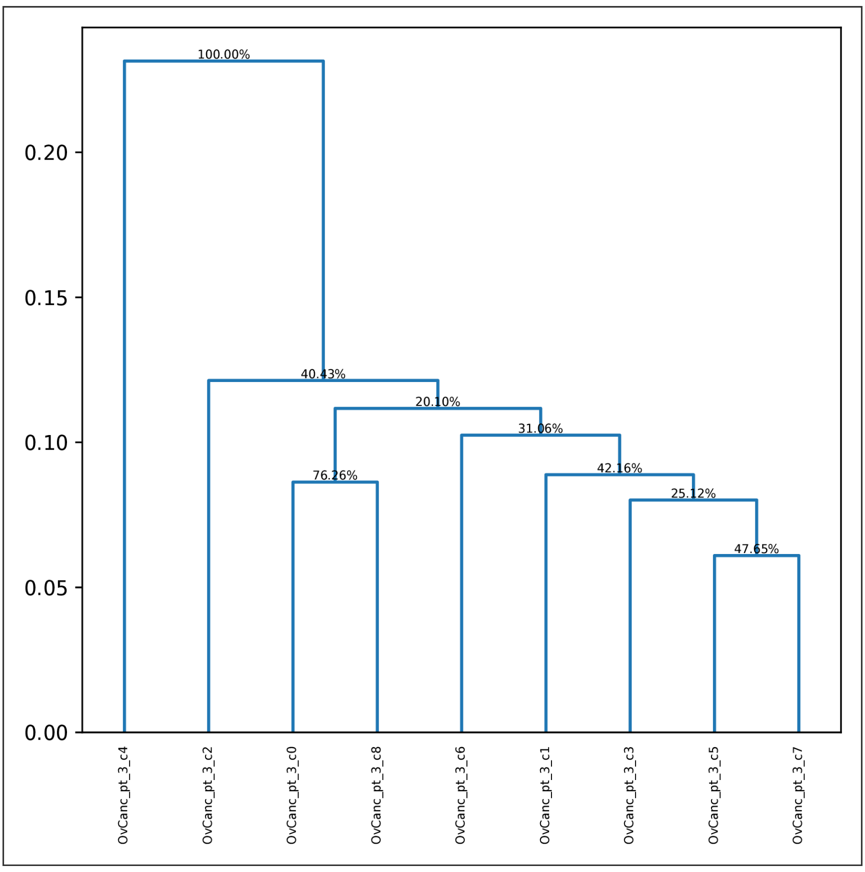


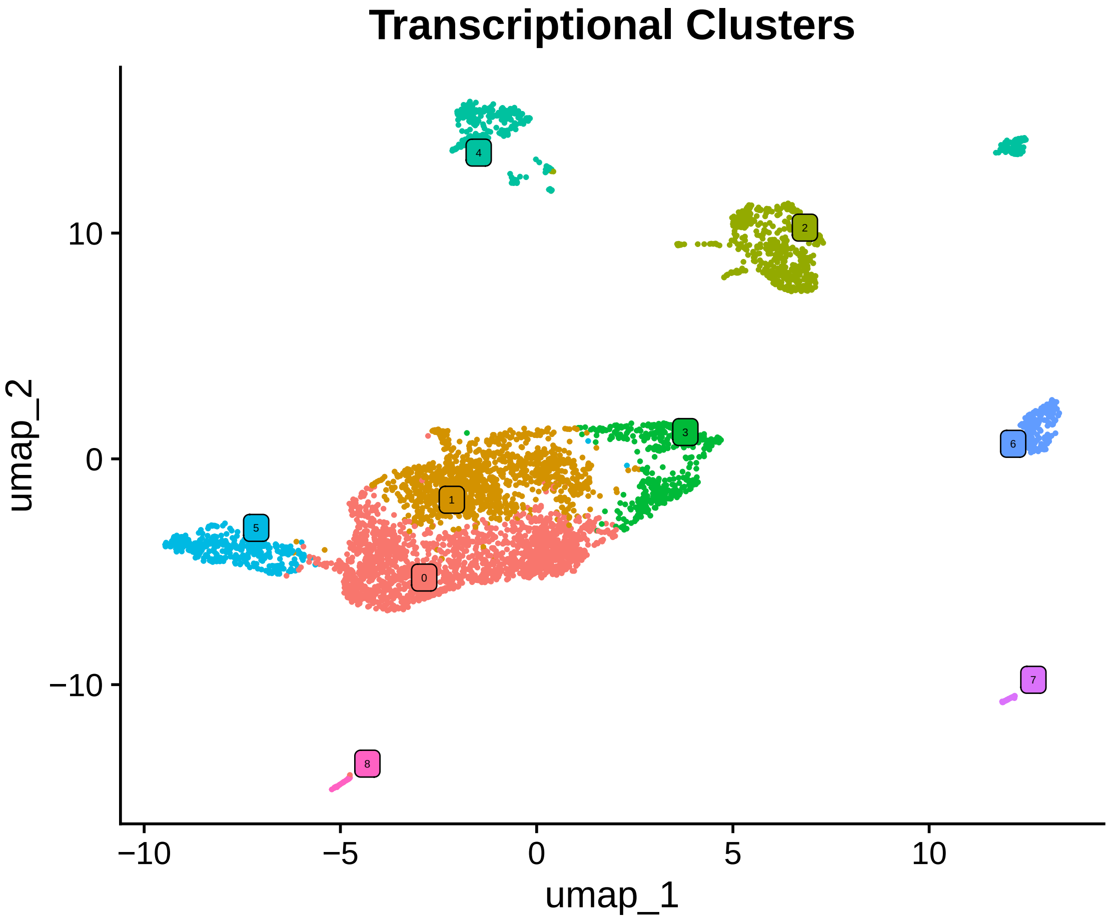


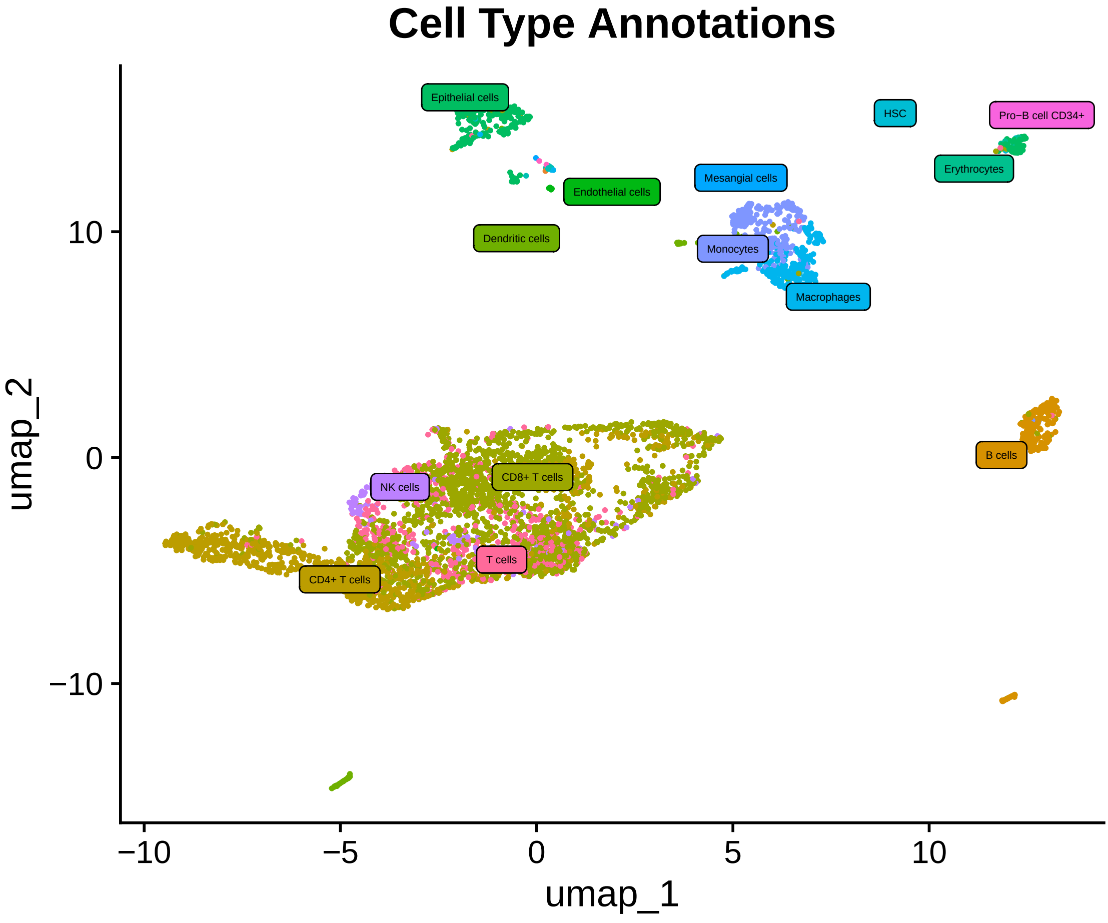

Sample: G0028

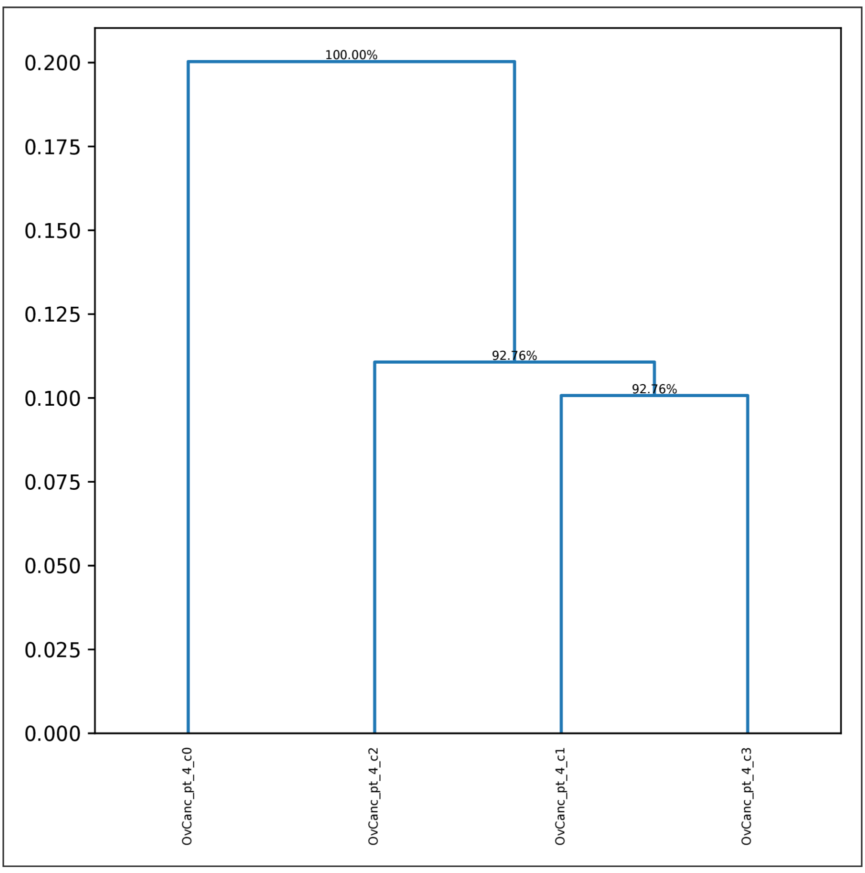


#### Transcriptional Clusters

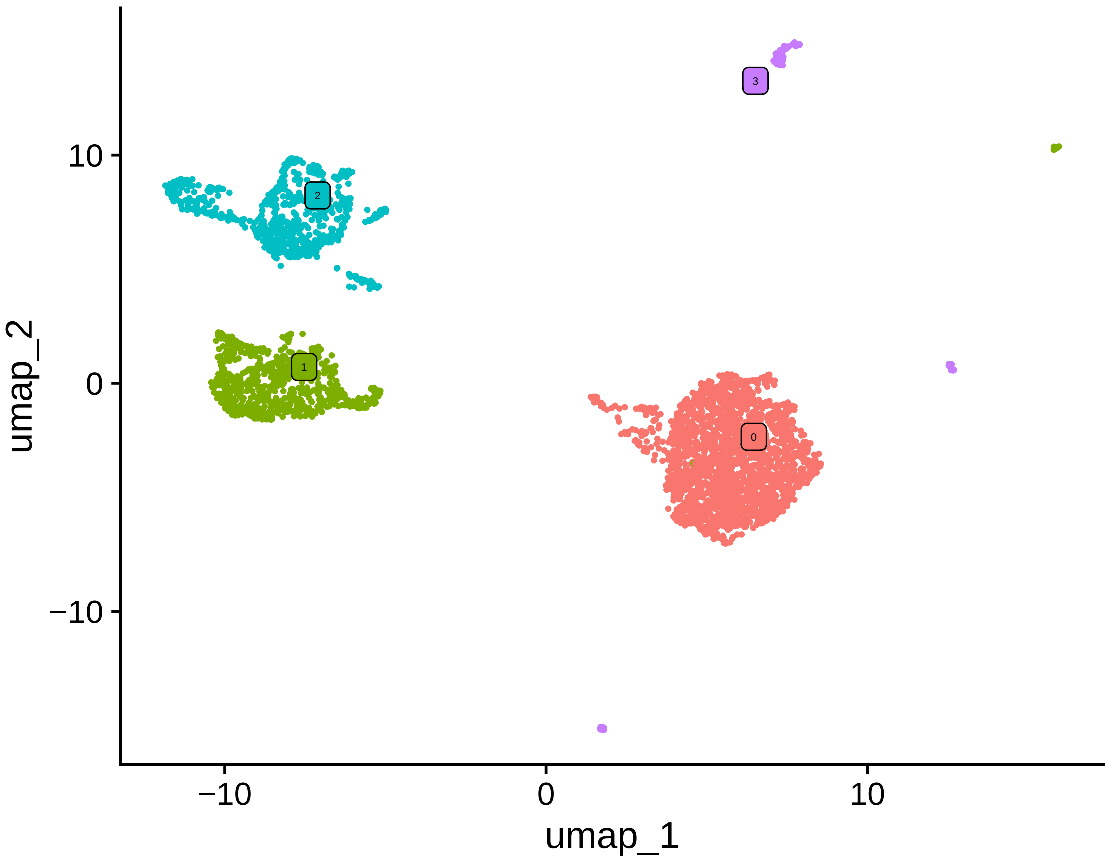


#### Cell Type Annotations

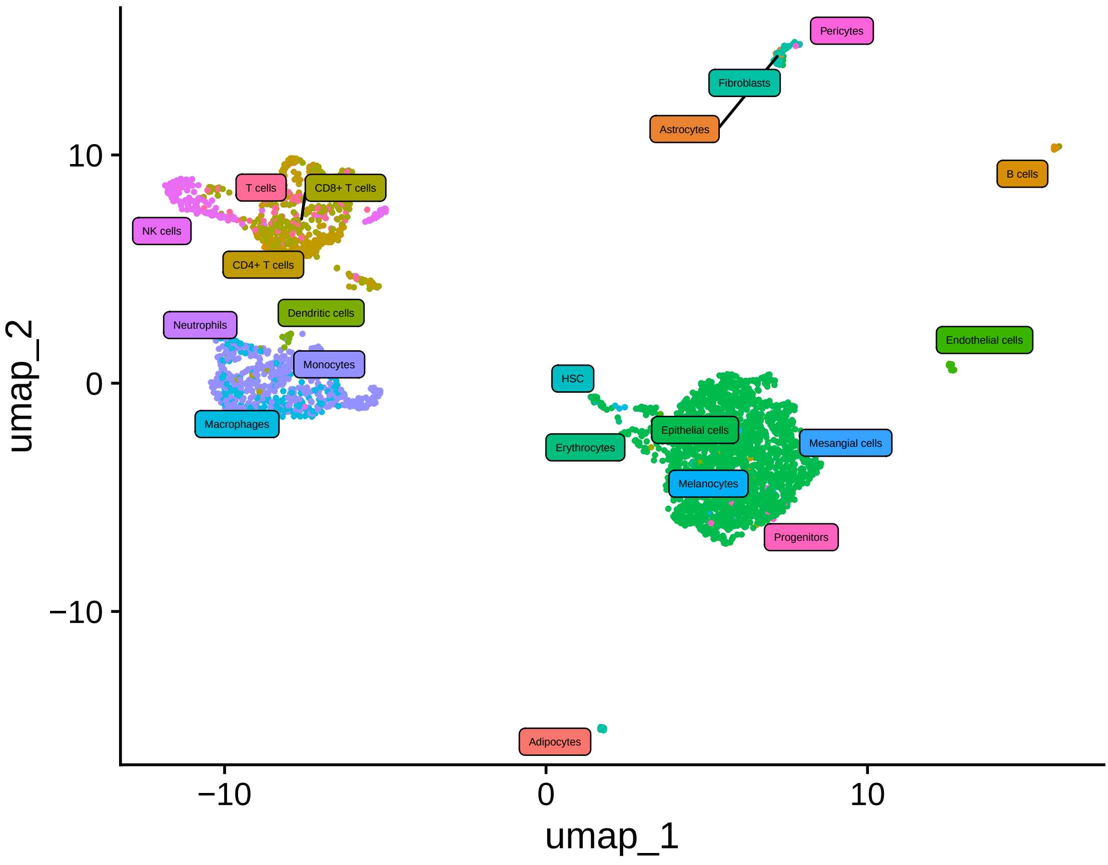

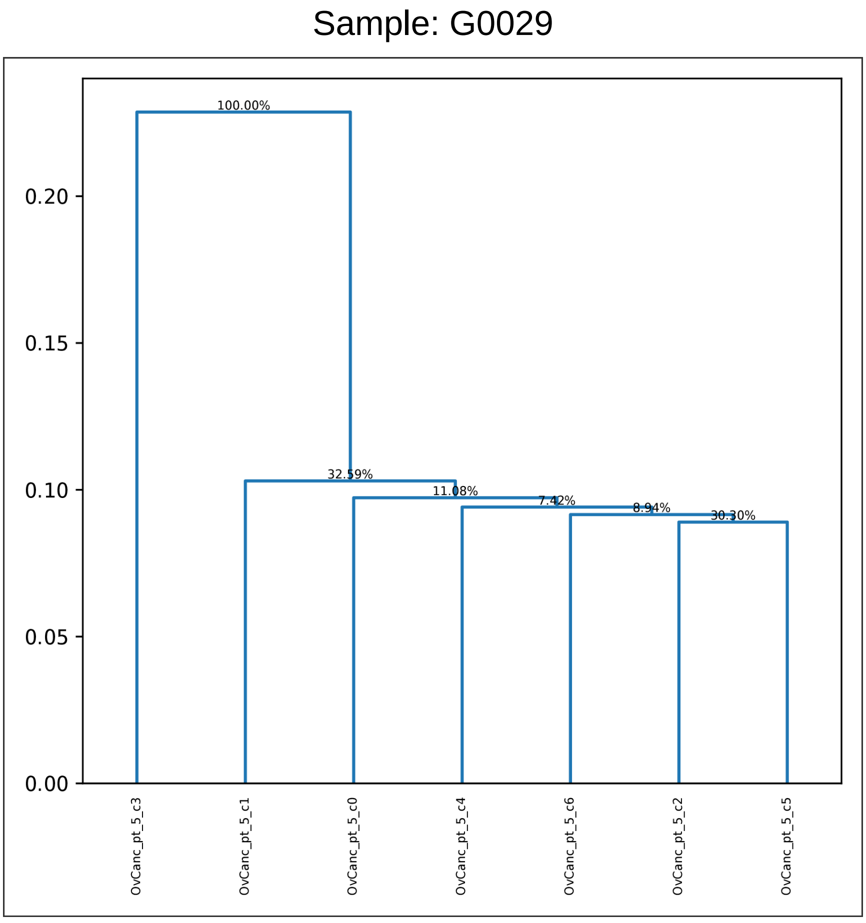

Sample: G0029
100.00%
32.59%
11.08%
7.42%
8.94%
30.30%
0.20
0.15
0.10
0.05
0.00
OvCanc_pt_5_c3
OvCanc_pt_5_c1
OvCanc_pt_5_c0
OvCanc_pt_5_c4
OvCanc_pt_5_c6
OvCanc_pt_5_c2
OvCanc_pt_5_c5


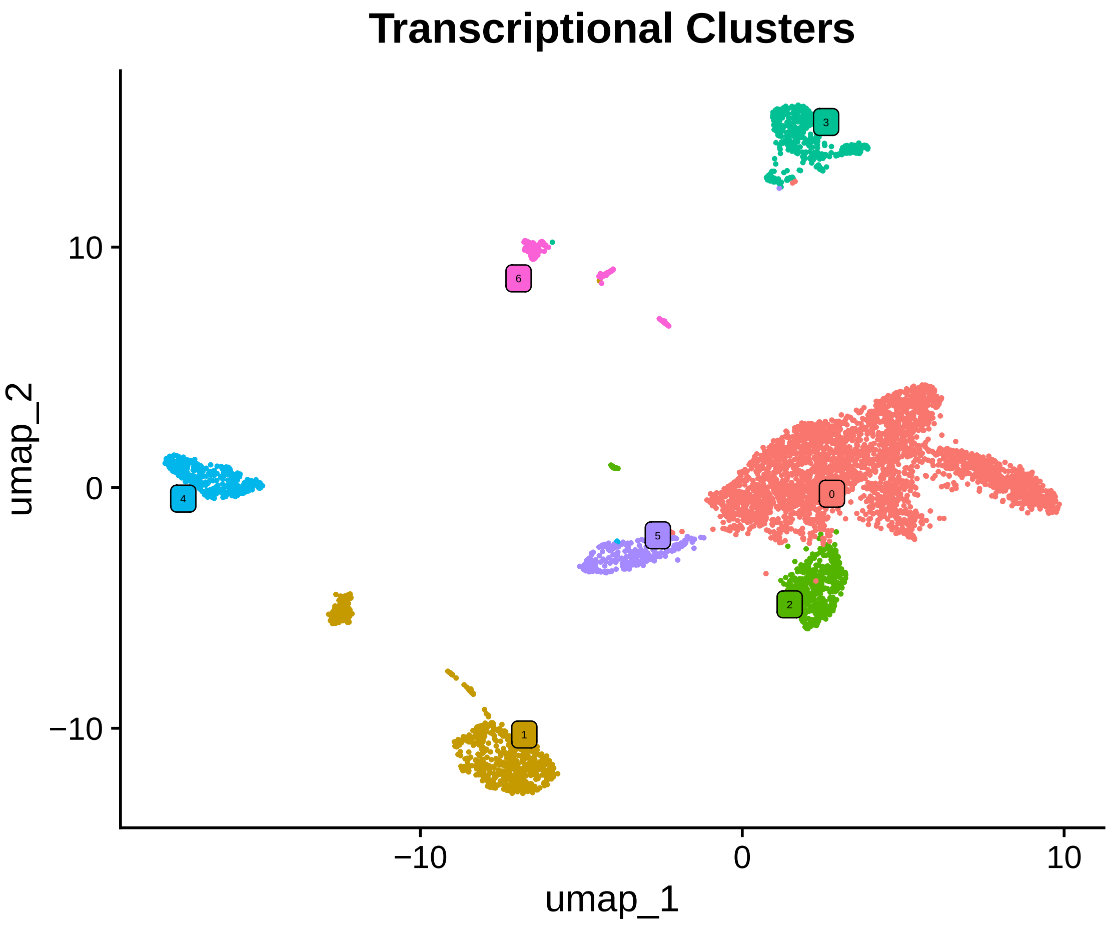

Transcriptional Clusters
umap_2
umap_1
10
0
−10
−10
0
10
3
6
4
5
0
2
1


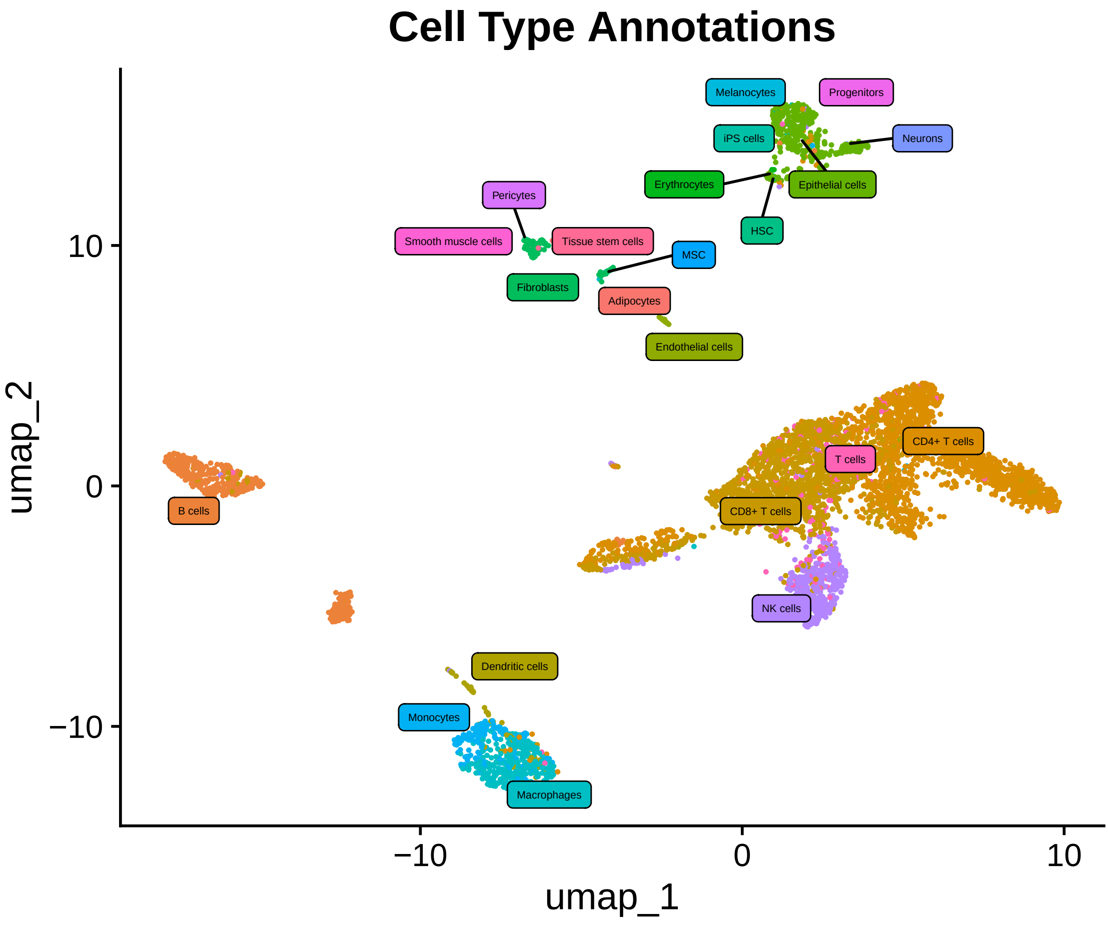

Cell Type Annotations
Melanocytes
Progenitors
iPS cells
Neurons
Erythrocytes
Epithelial cells
HSC
Pericytes
Smooth muscle cells
Tissue stem cells
MSC
Fibroblasts
Adipocytes
Endothelial cells
CD4+ T cells
T cells
B cells
CD8+ T cells
NK cells
Dendritic cells
Monocytes
Macrophages
umap_2
umap_1
10
0
−10
−10
0
10

Sample: G0030

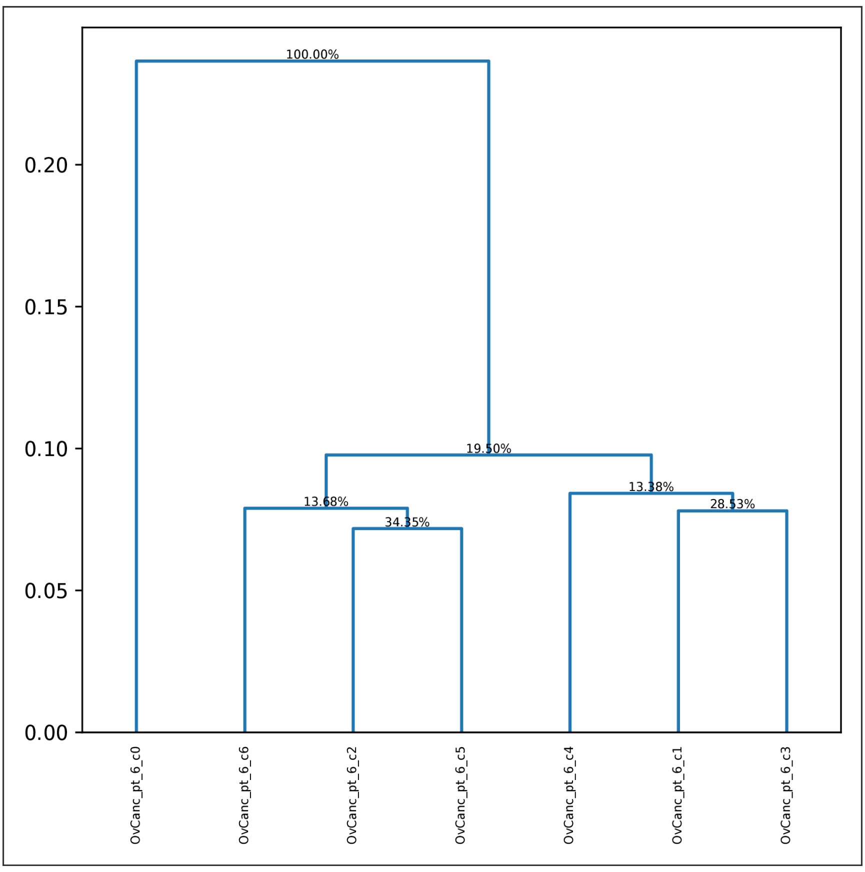


**Transcriptional Clusters**

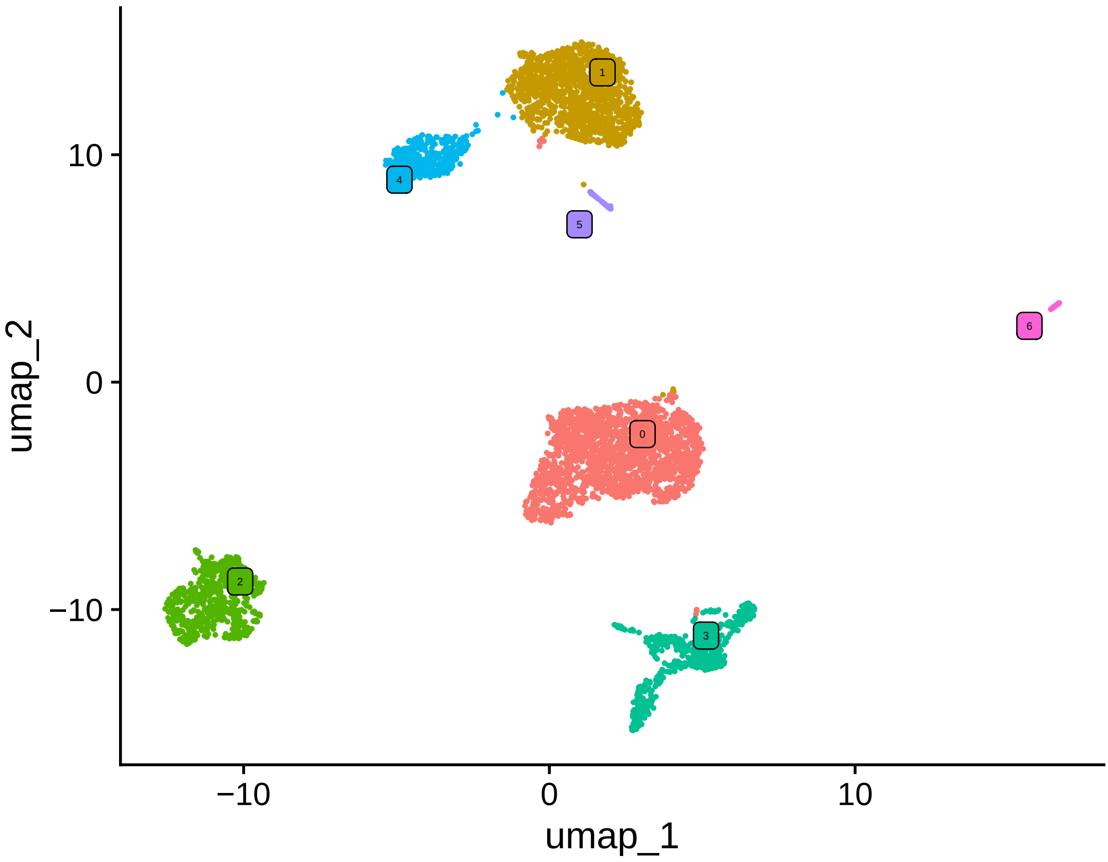


**Cell Type Annotations**

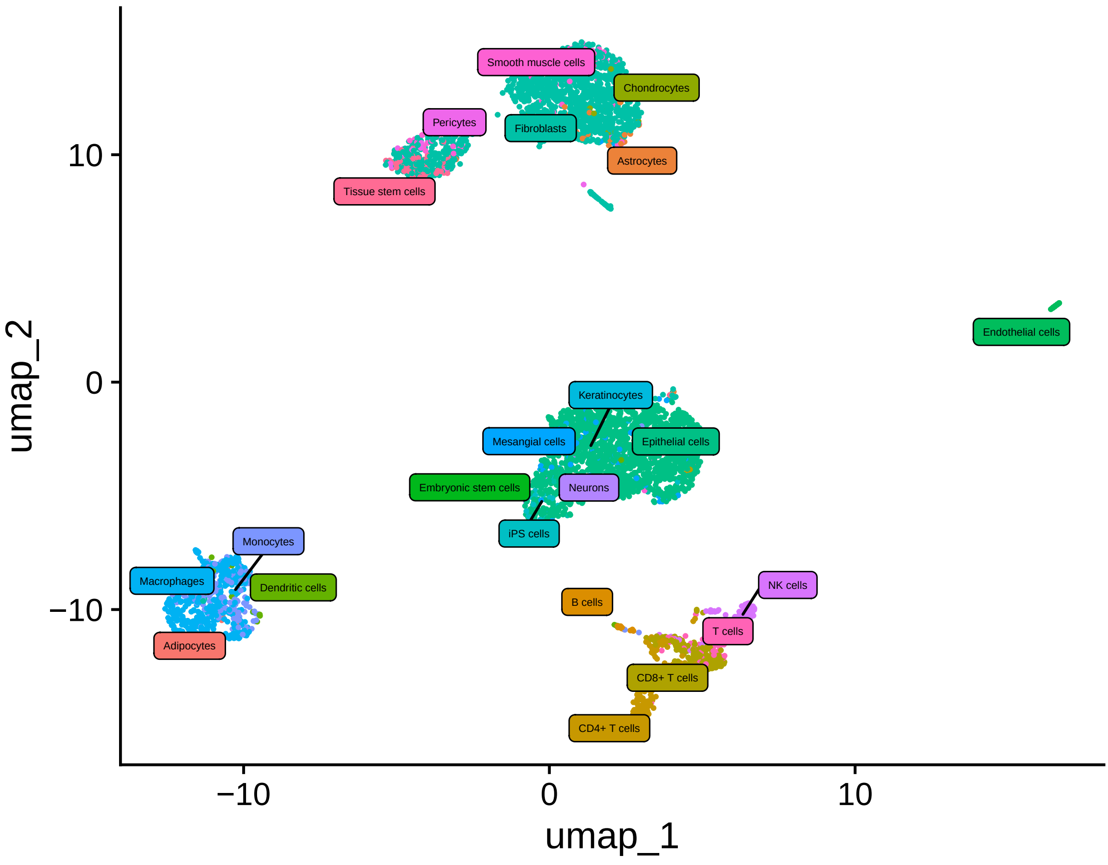

Sample: G0031

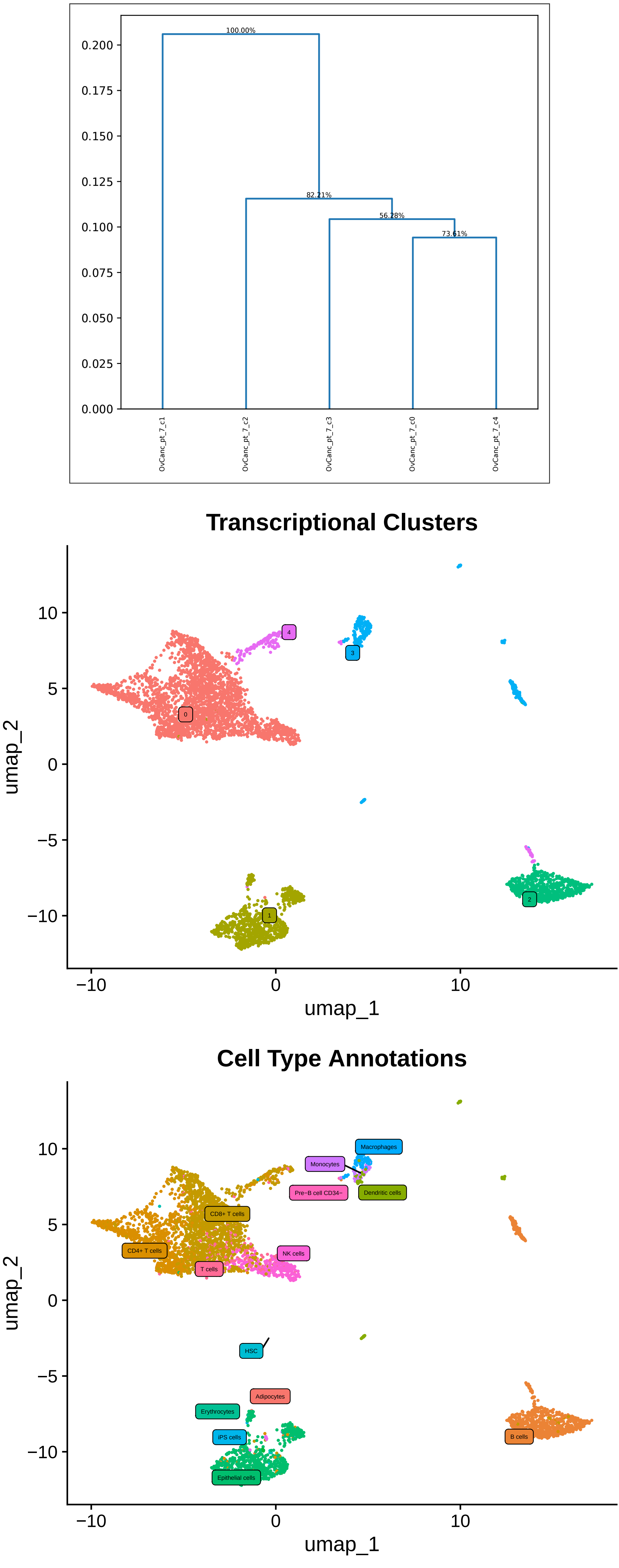

Sample: G0032

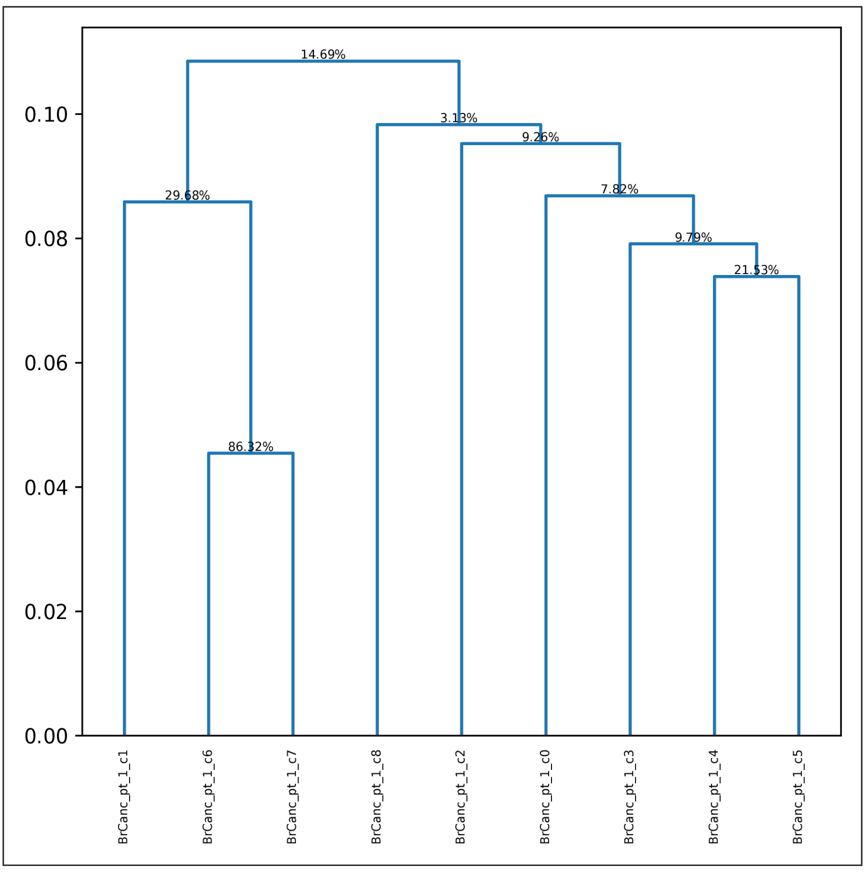


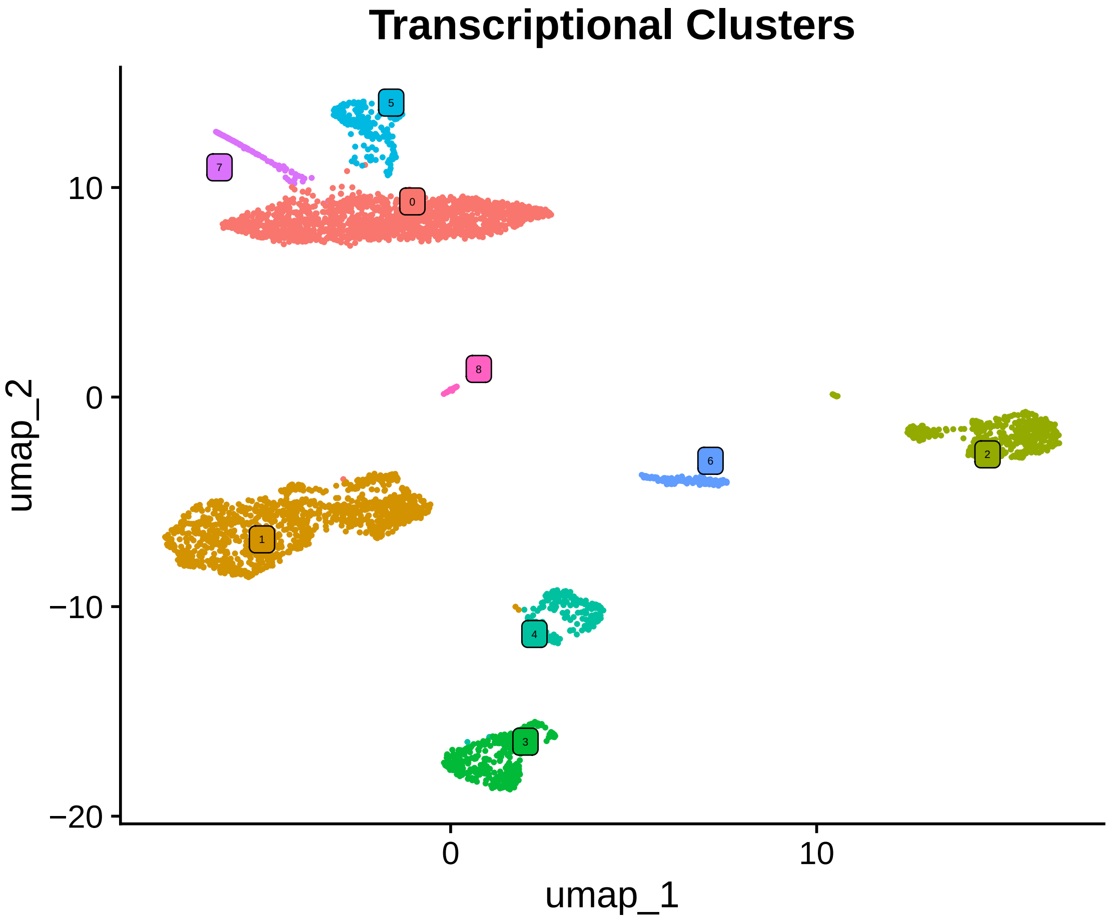


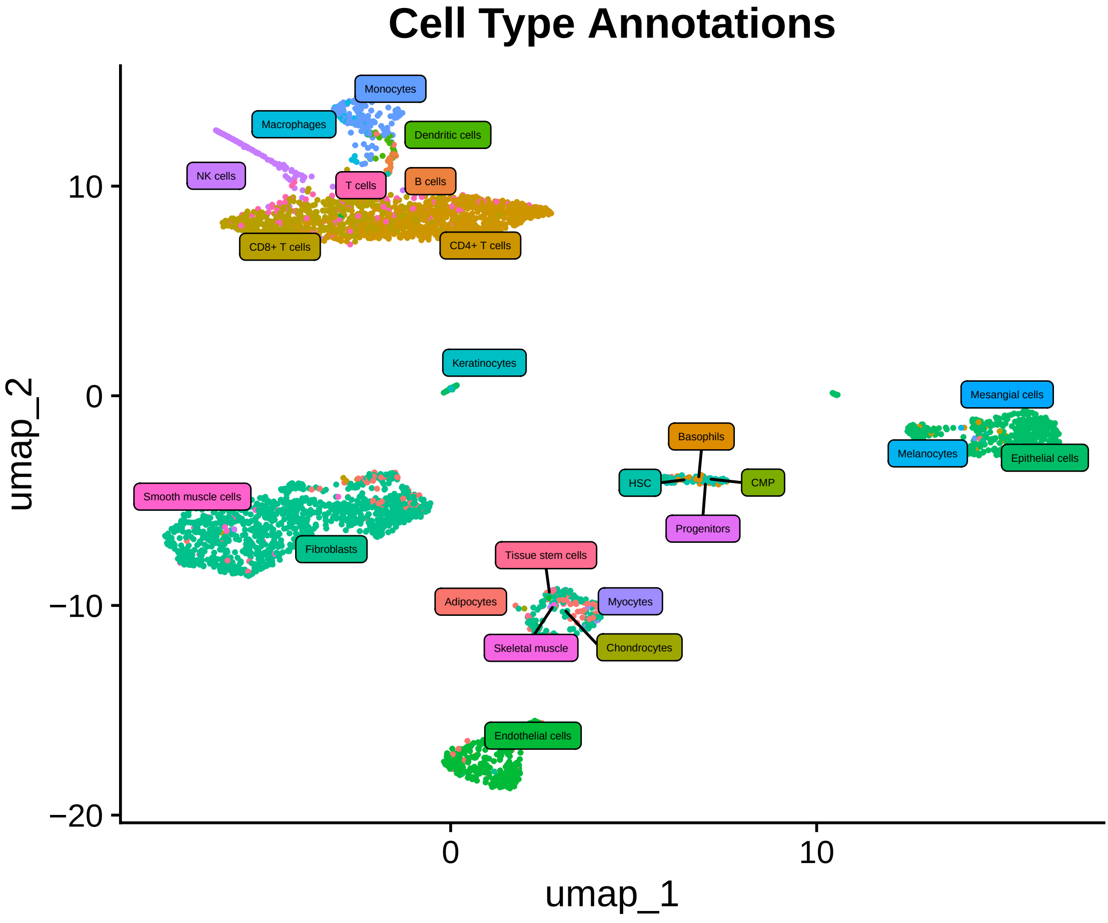

Sample: G0034

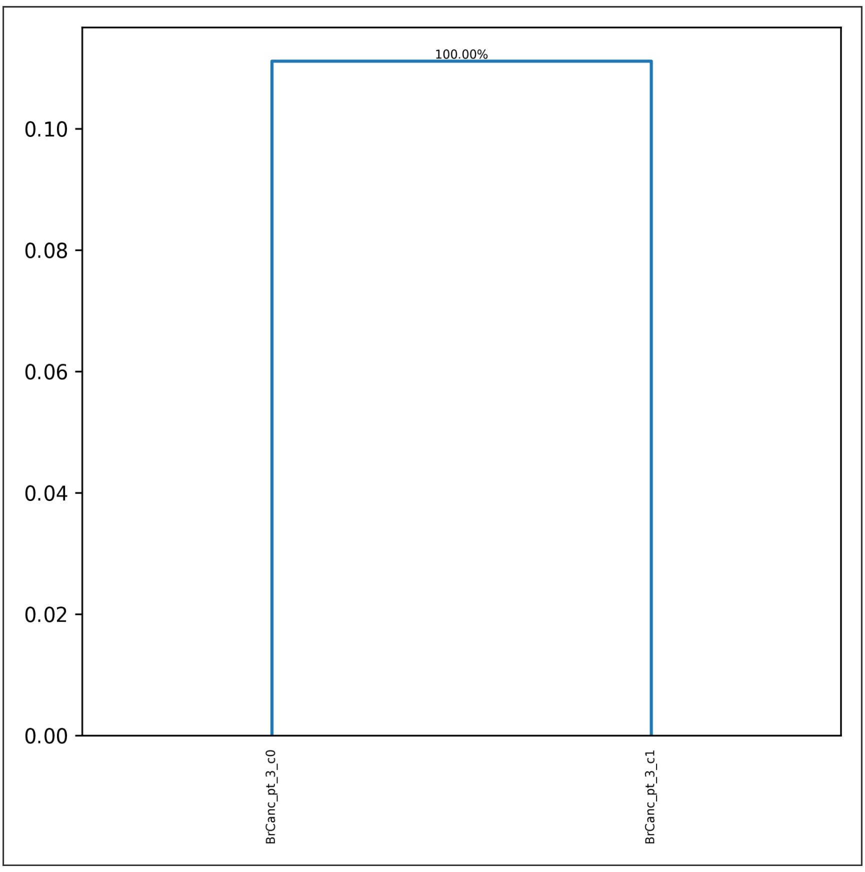


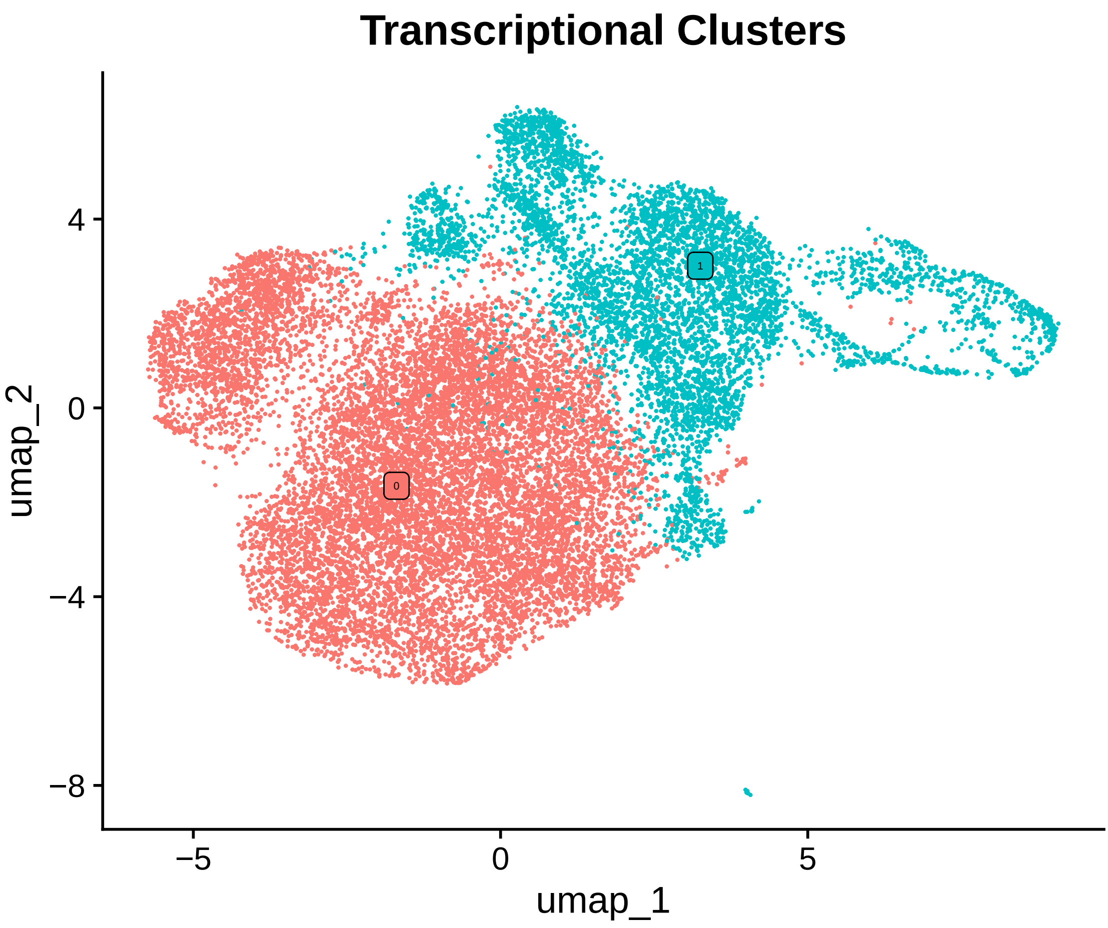


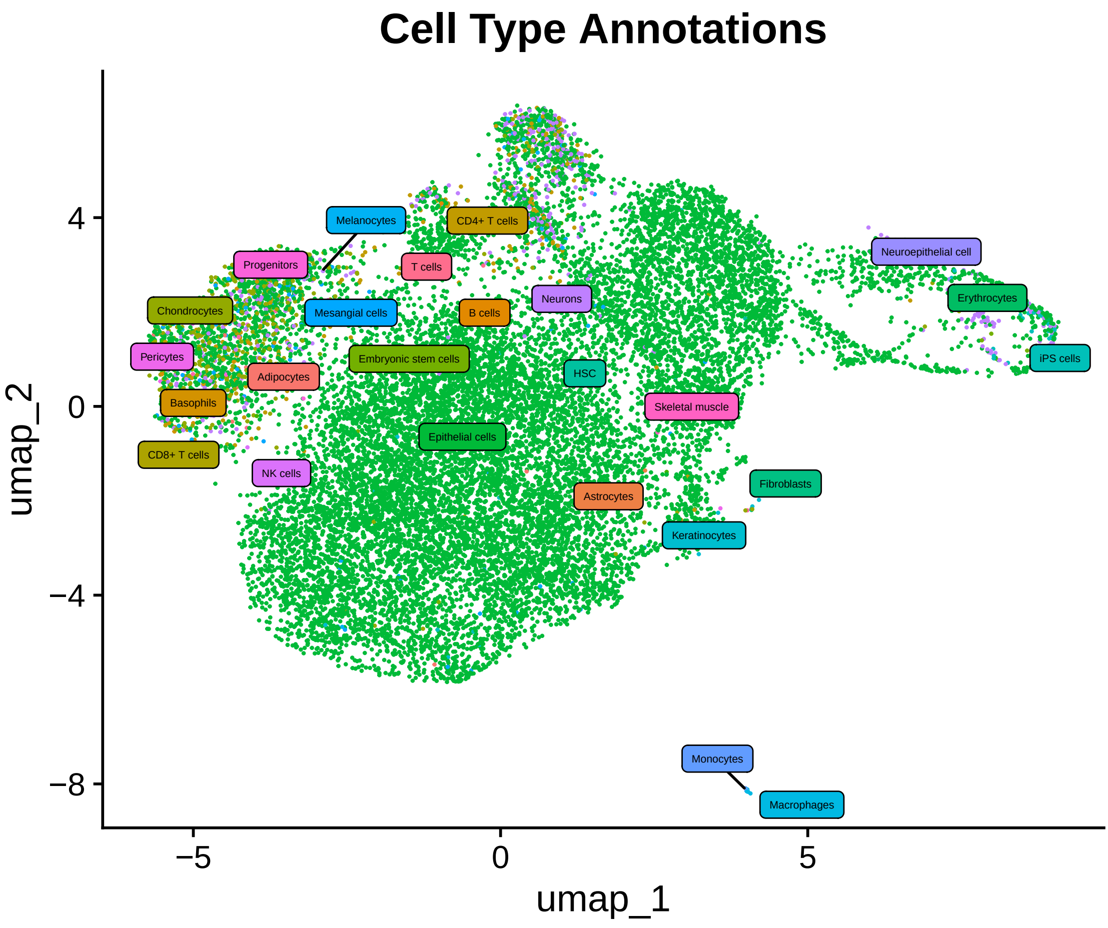

Sample: G0035

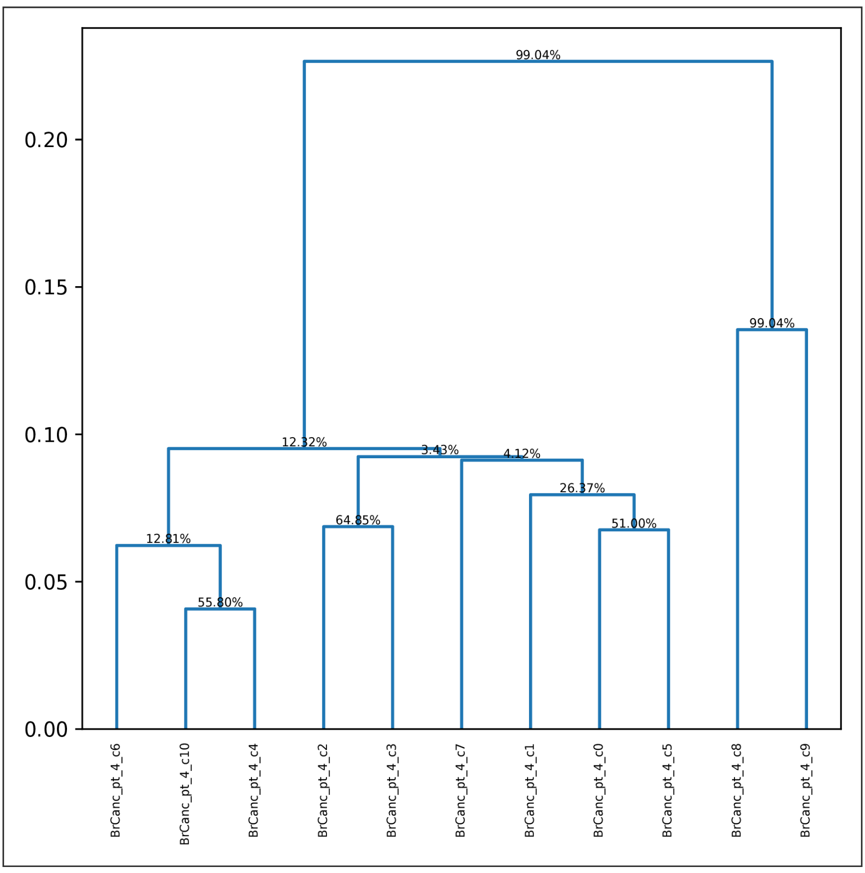


**Transcriptional Clusters**

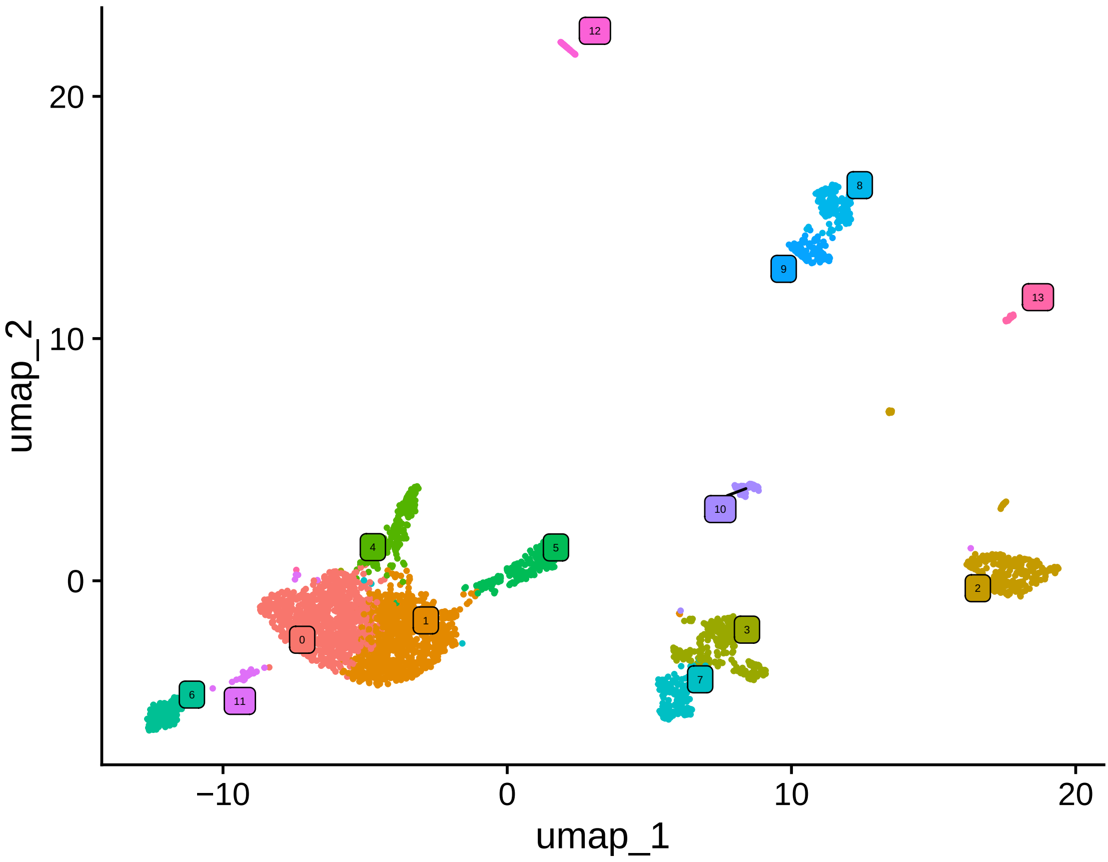


**Cell Type Annotations**

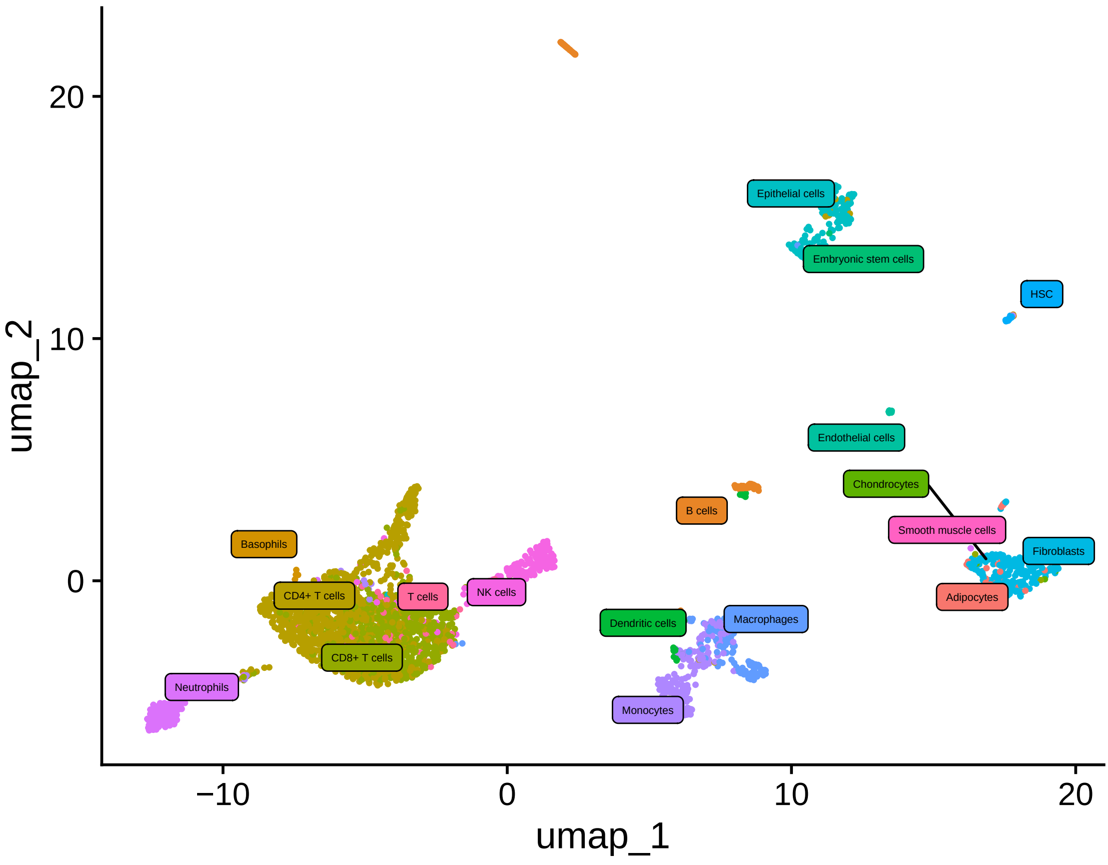

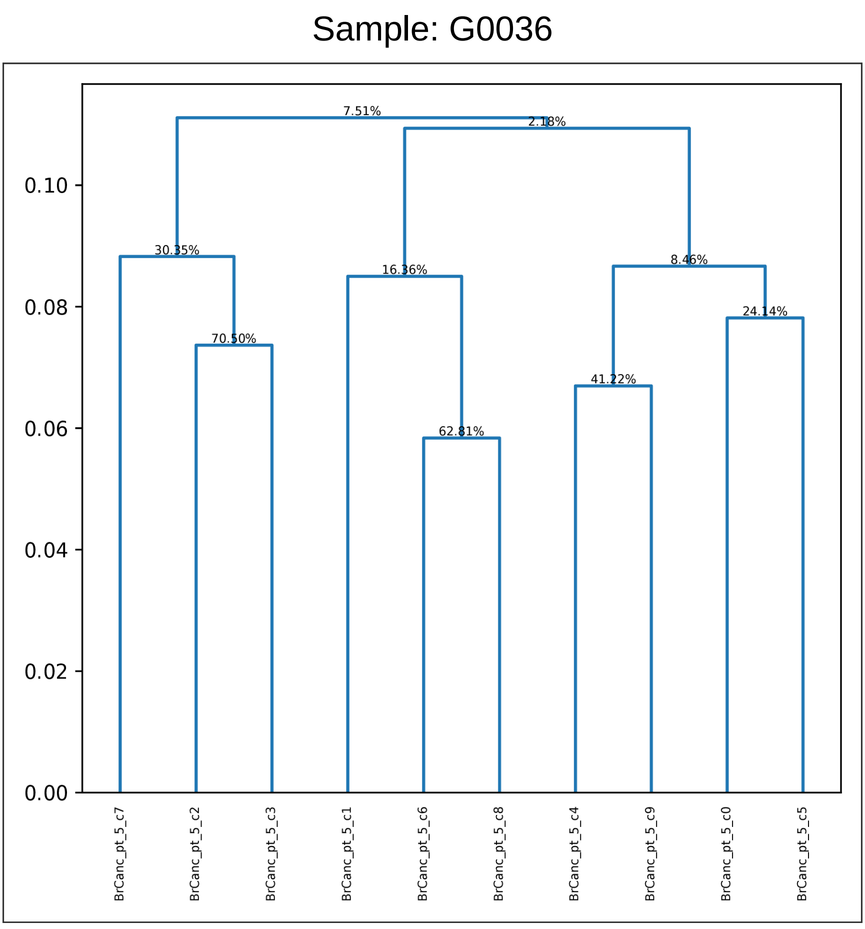
Sample: G0036
7.51%
2.18%
30.35%
16.36%
8.46%
70.50%
24.14%
41.22%
62.81%
0.10
0.08
0.06
0.04
0.02
0.00
BrCanc_pt_5_c7
BrCanc_pt_5_c2
BrCanc_pt_5_c3
BrCanc_pt_5_c1
BrCanc_pt_5_c6
BrCanc_pt_5_c8
BrCanc_pt_5_c4
BrCanc_pt_5_c9
BrCanc_pt_5_c0
BrCanc_pt_5_c5

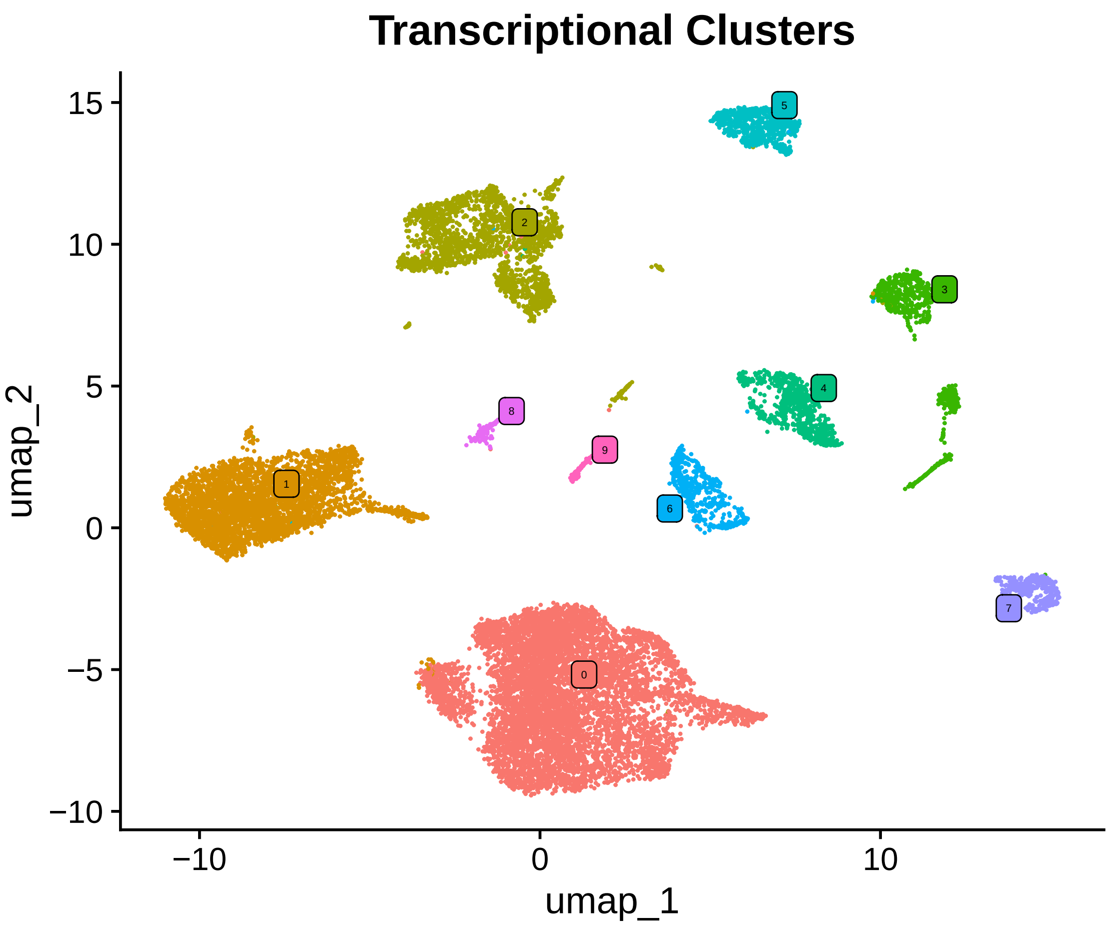
Transcriptional Clusters
umap_2
umap_1
15
10
5
0
−5
−10
−10
0
10
0
1
2
3
4
5
6
7
8
9

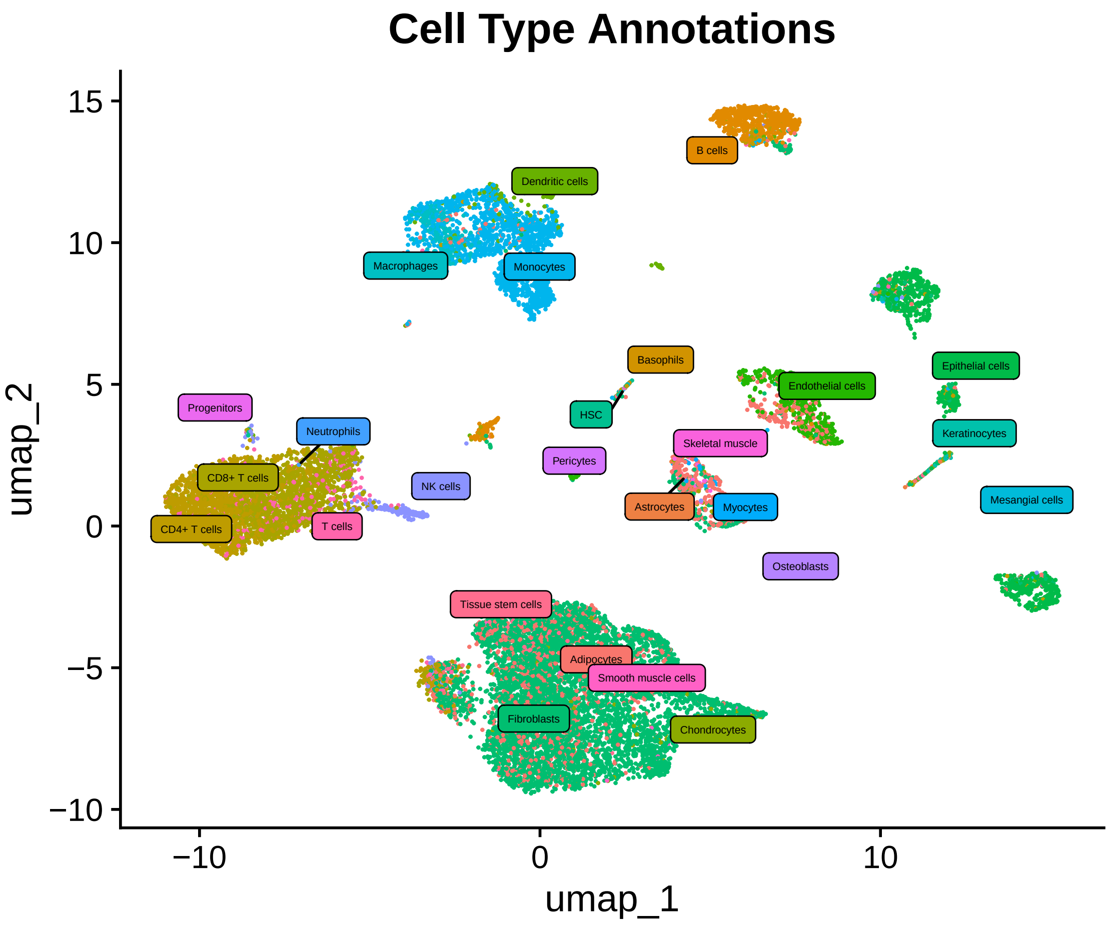
Cell Type Annotations
umap_2
umap_1
15
10
5
0
−5
−10
−10
0
10
B cells
Dendritic cells
Macrophages
Monocytes
Basophils
HSC
Endothelial cells
Epithelial cells
Keratinocytes
Progenitors
Neutrophils
CD8+ T cells
CD4+ T cells
T cells
NK cells
Pericytes
Skeletal muscle
Astrocytes
Myocytes
Mesangial cells
Osteoblasts
Tissue stem cells
Adipocytes
Smooth muscle cells
Fibroblasts
Chondrocytes

Sample: G0038

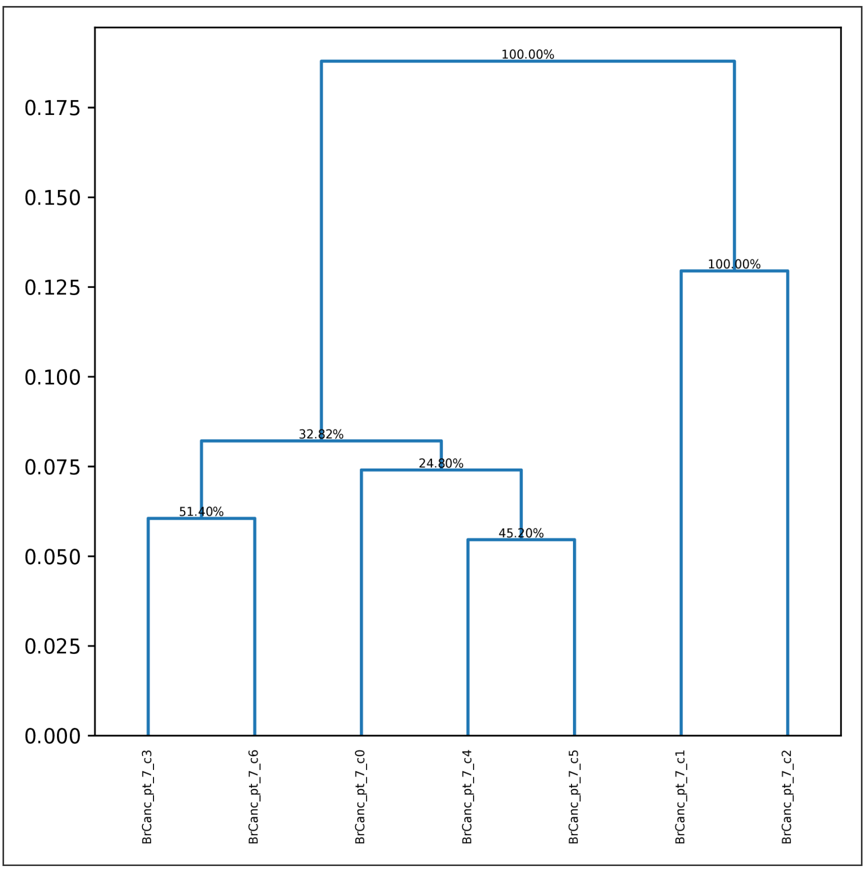


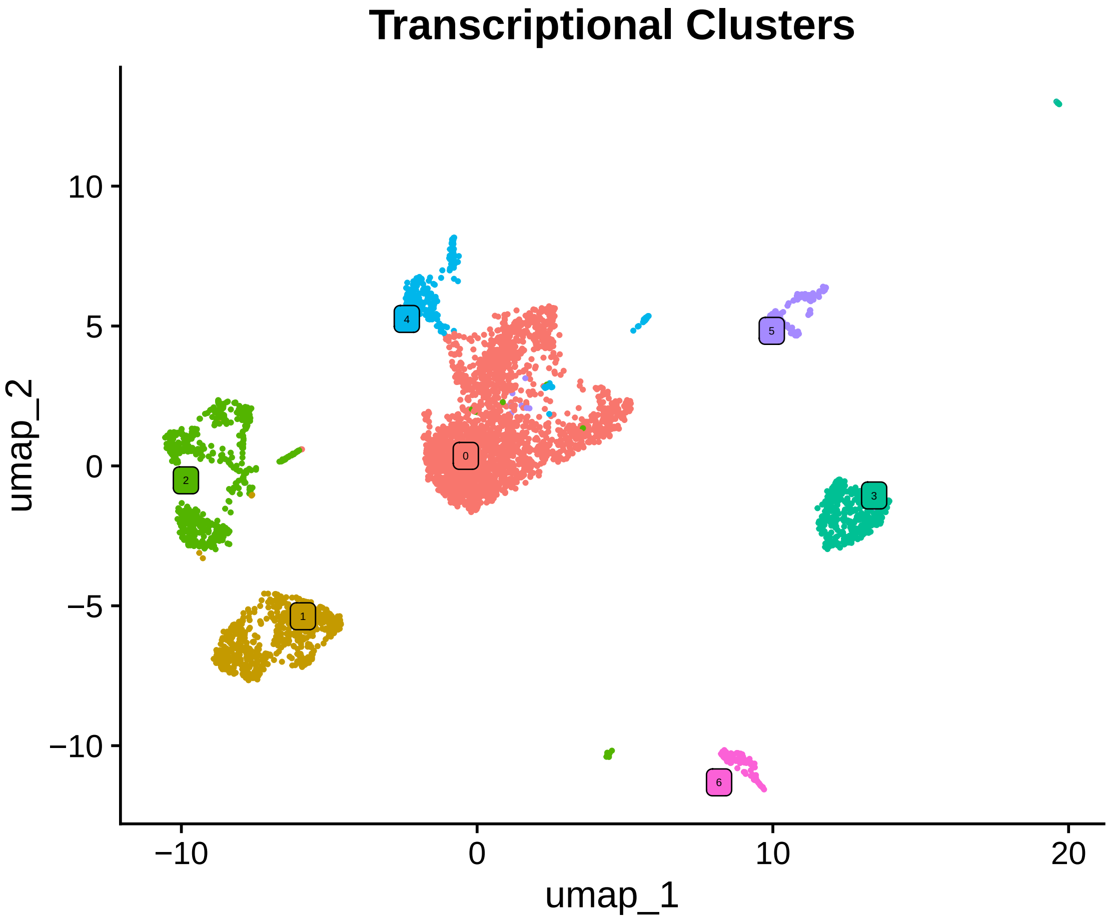


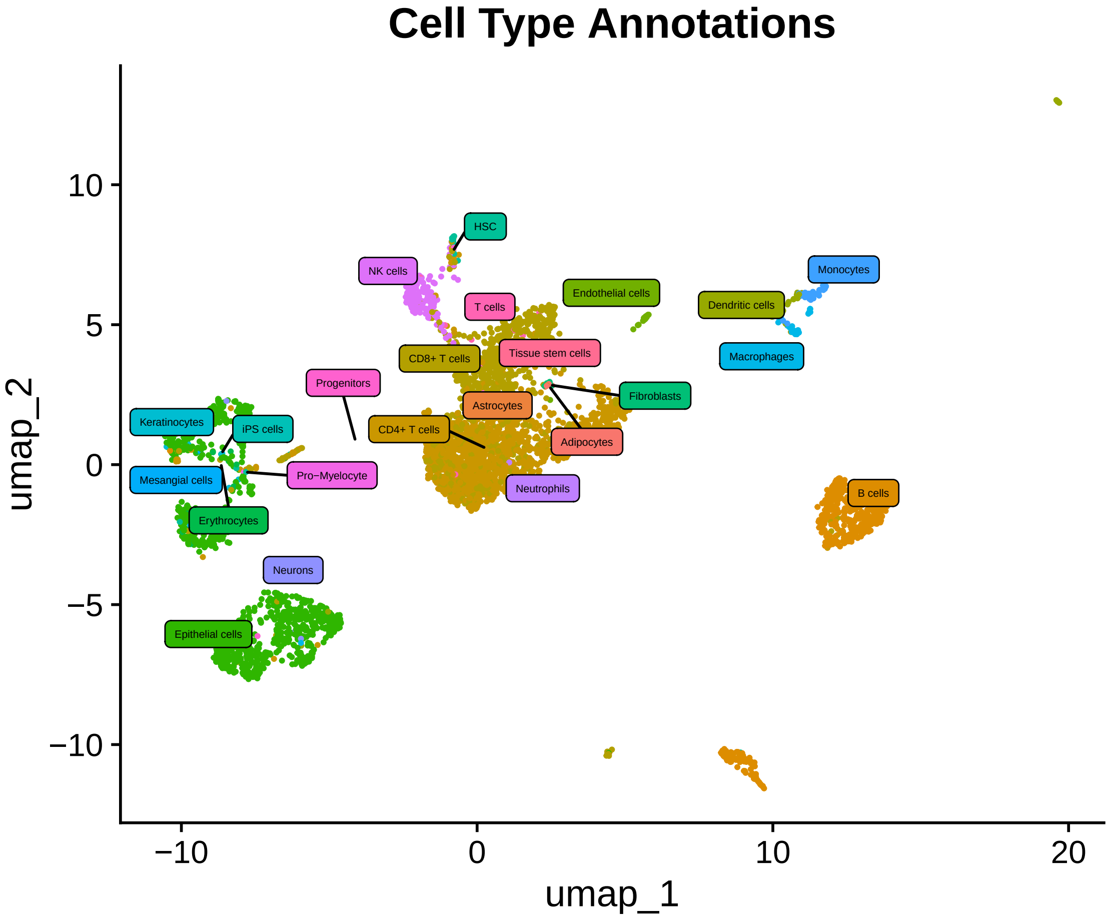

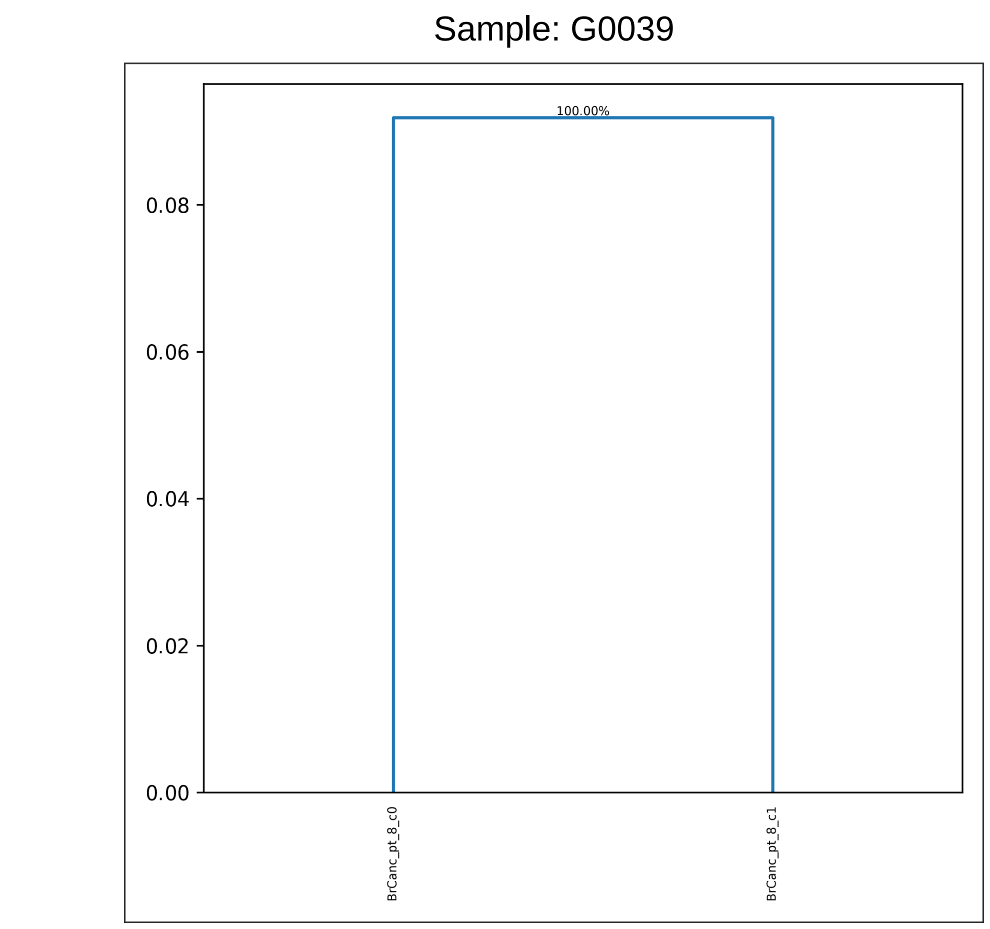

Sample: G0039
100.00%
0.08
0.06
0.04
0.02
0.00
BrCanc_pt_8_c0
BrCanc_pt_8_c1


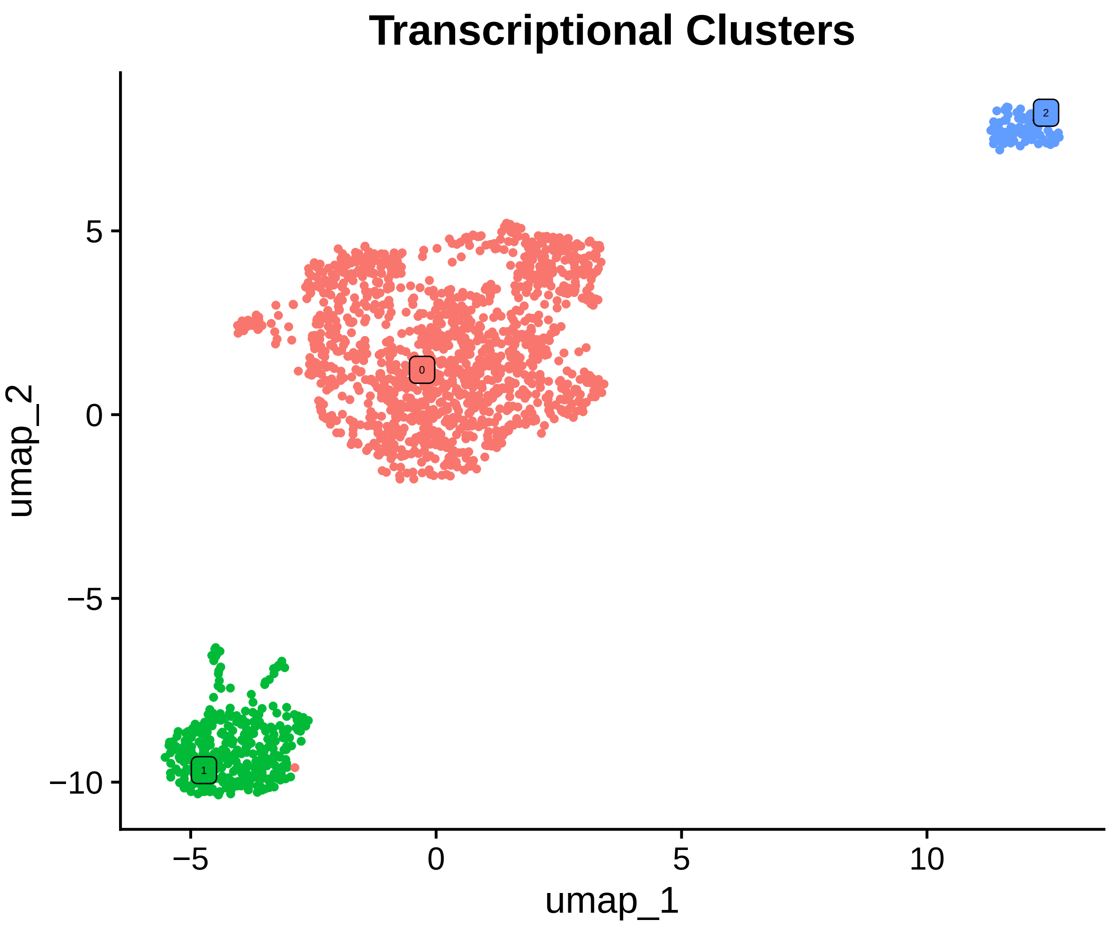

Transcriptional Clusters
umap_2
umap_1
0
1
2
5
0
−5
−10
−5
0
5
10


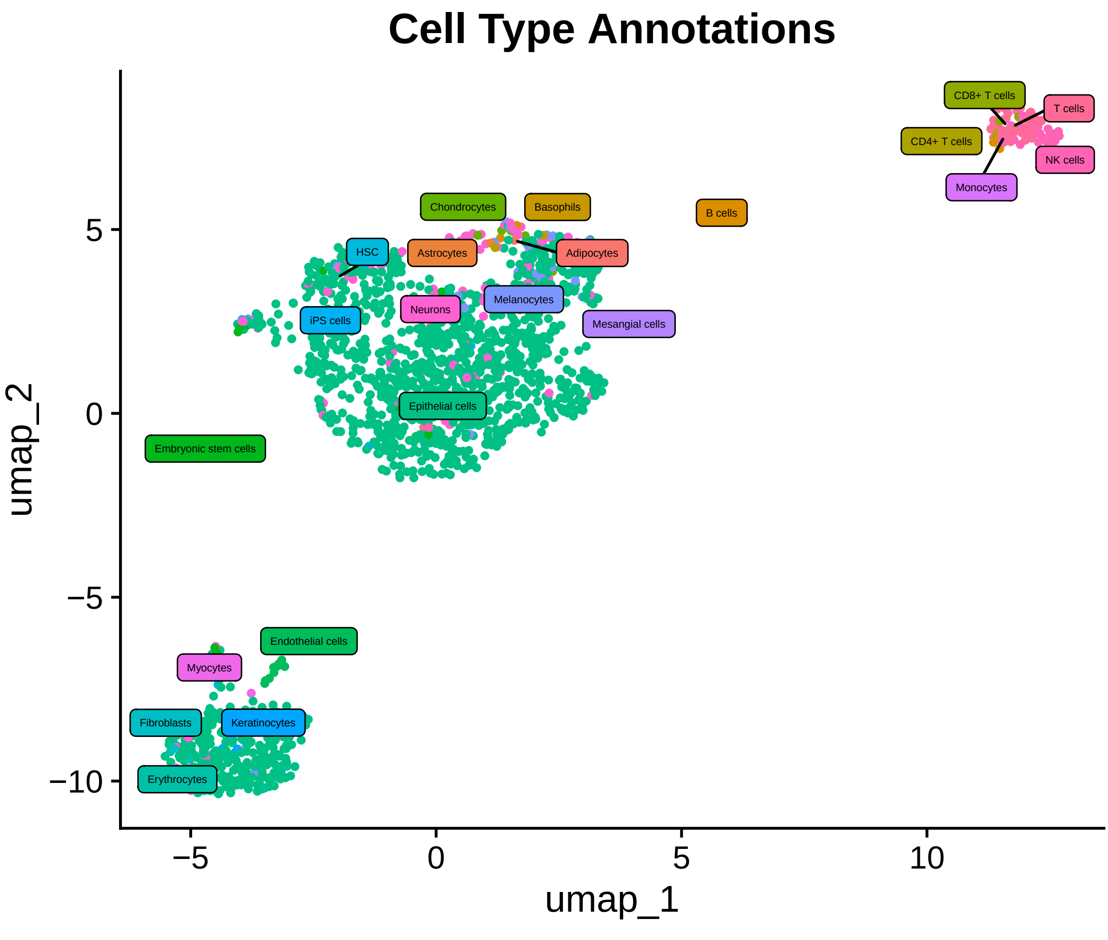

Cell Type Annotations
umap_2
umap_1
CD8+ T cells
T cells
CD4+ T cells
NK cells
Monocytes
Chondrocytes
Basophils
B cells
HSC
Astrocytes
Adipocytes
iPS cells
Neurons
Melanocytes
Mesangial cells
Epithelial cells
Embryonic stem cells
Endothelial cells
Myocytes
Fibroblasts
Keratinocytes
Erythrocytes
5
0
−5
−10
−5
0
5
10

Sample: G0040

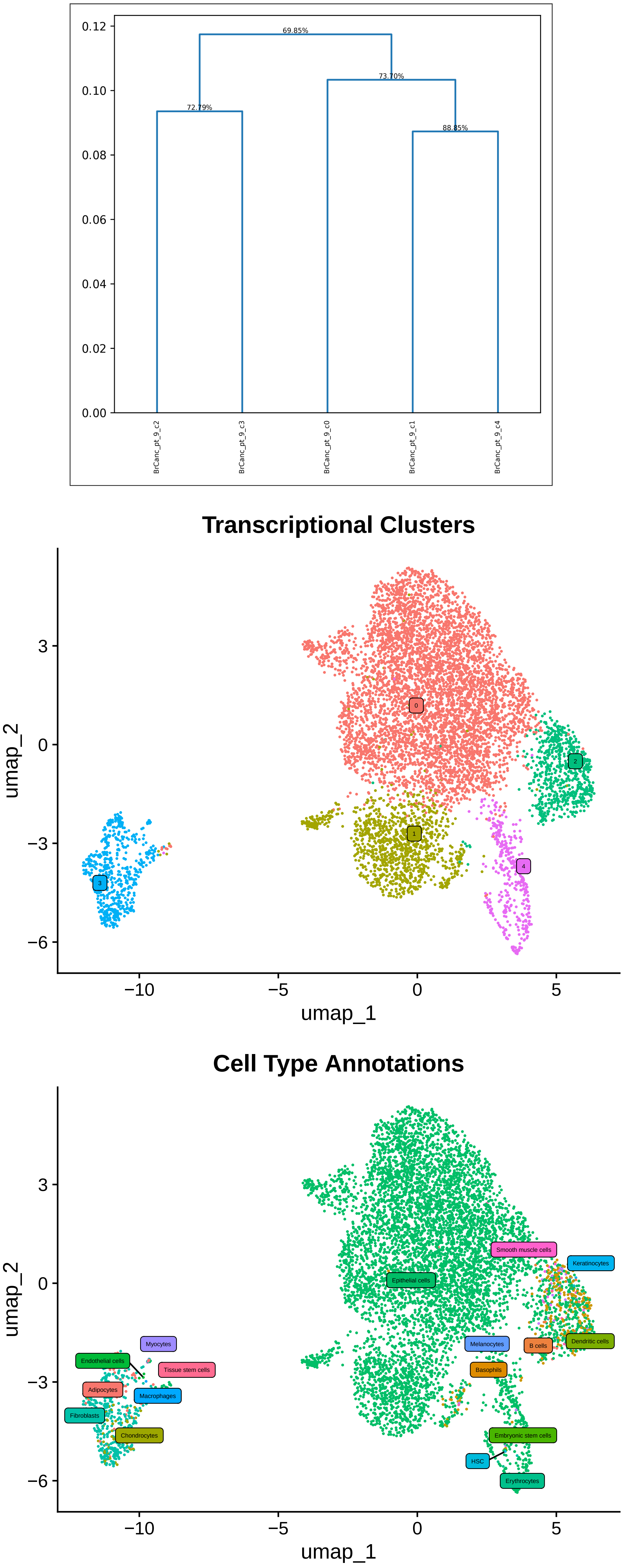

Sample: G0041

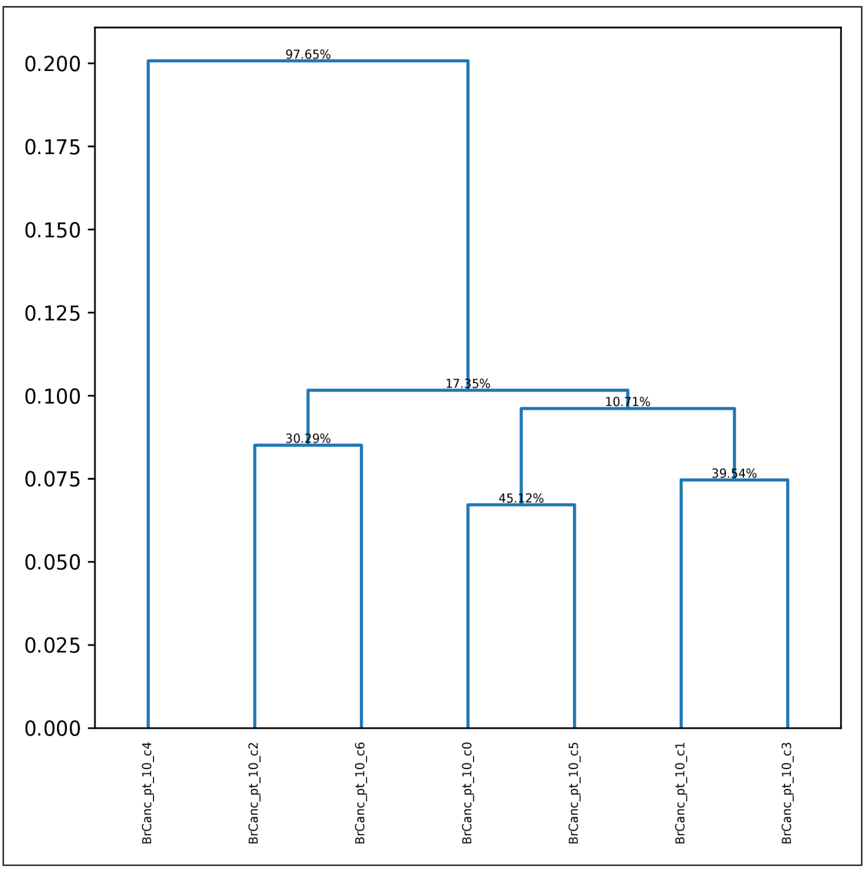


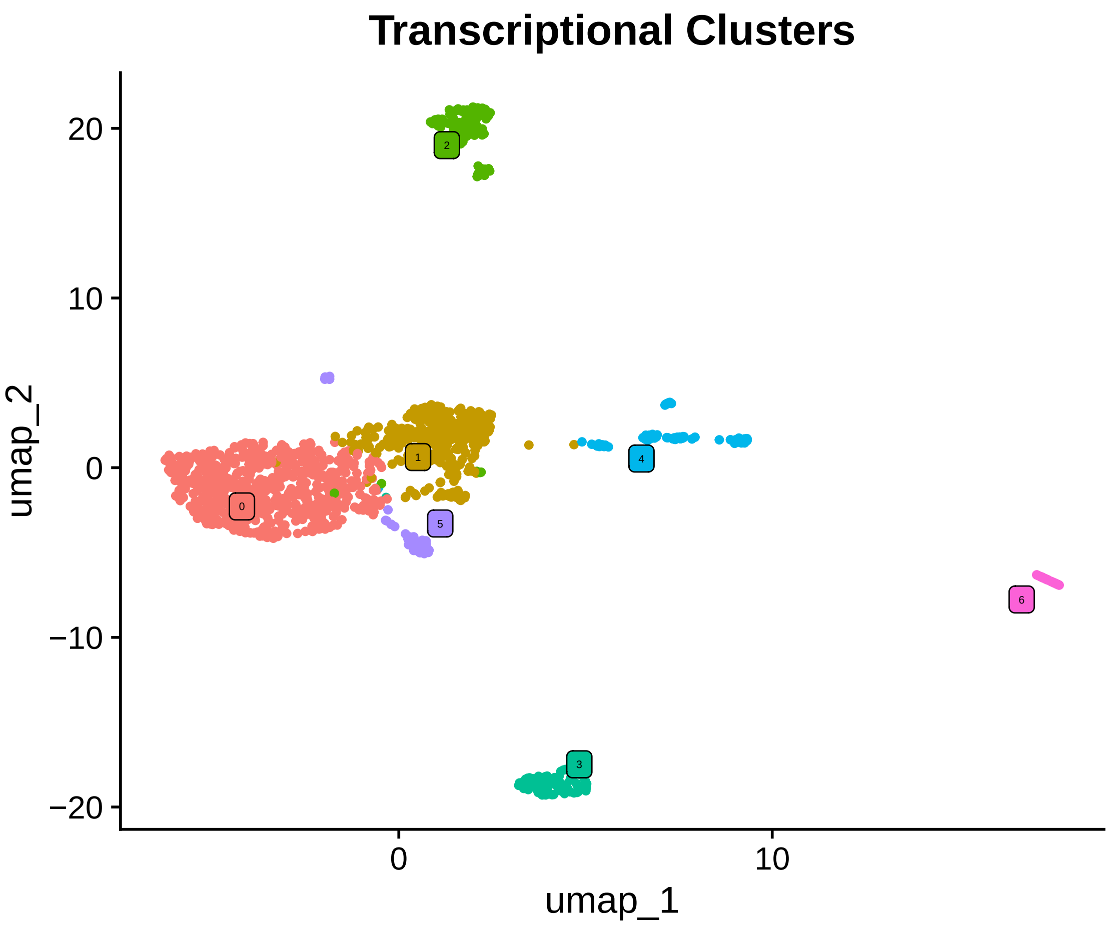


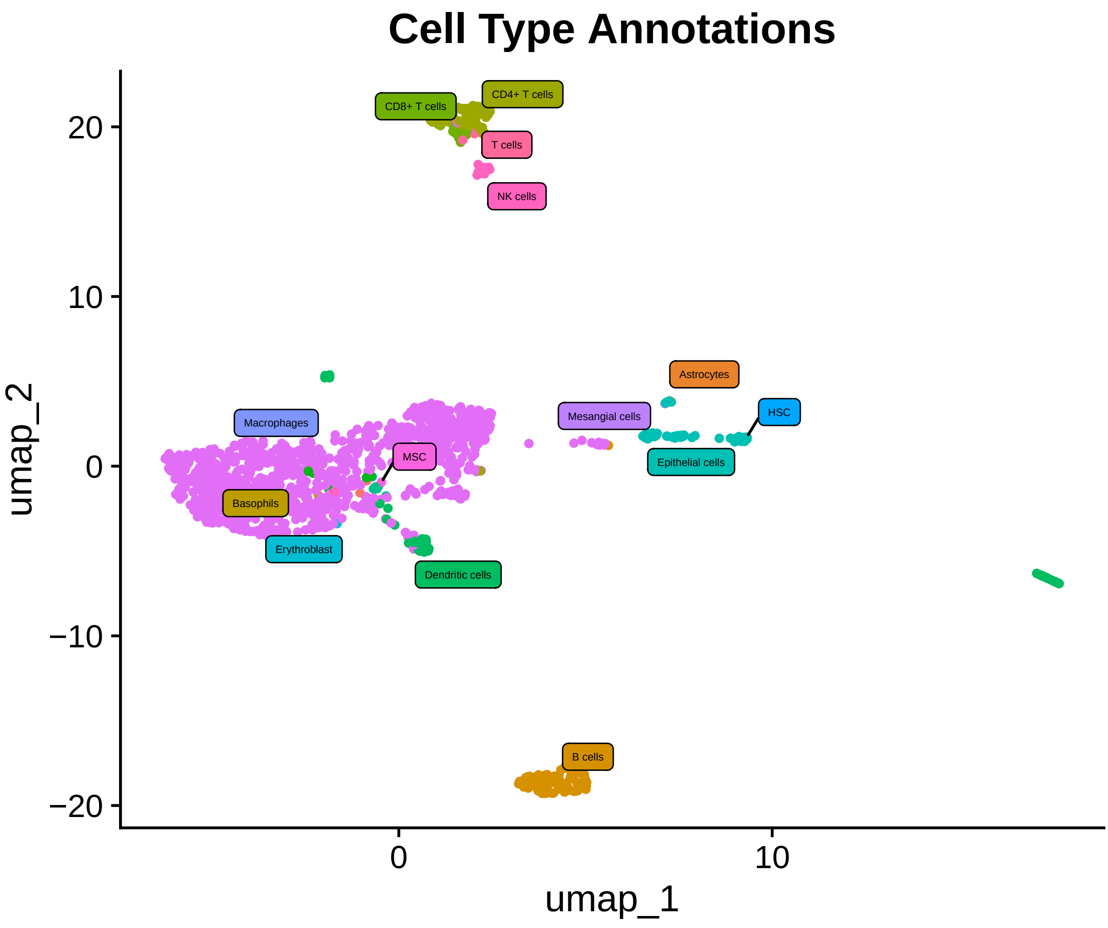

Sample: G0042

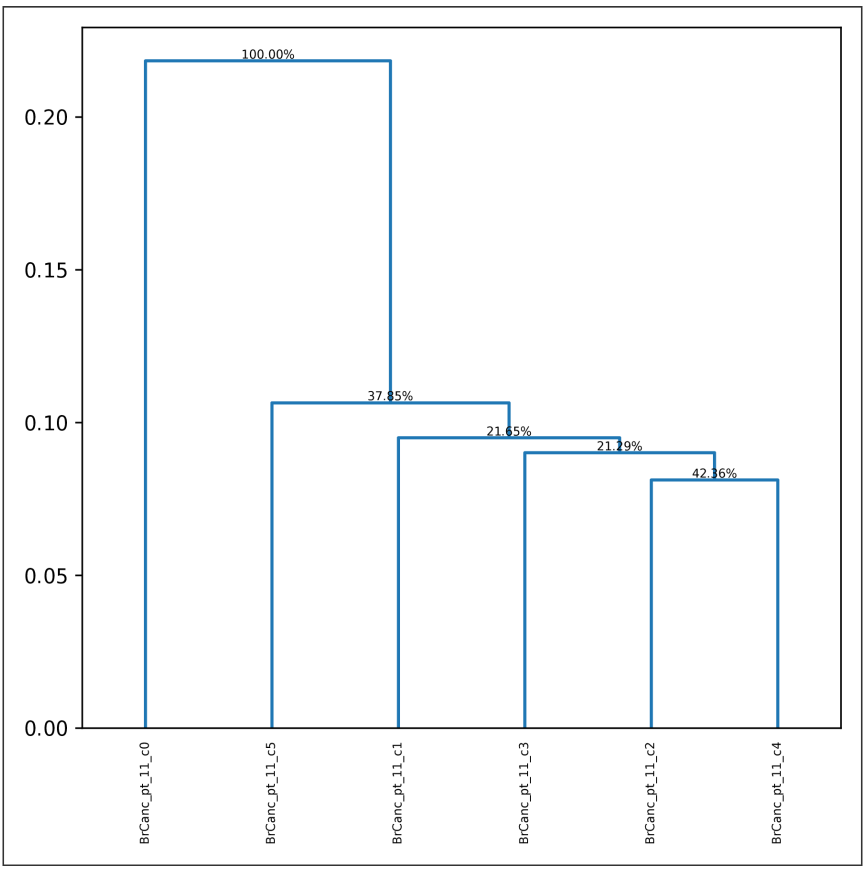


#### Transcriptional Clusters

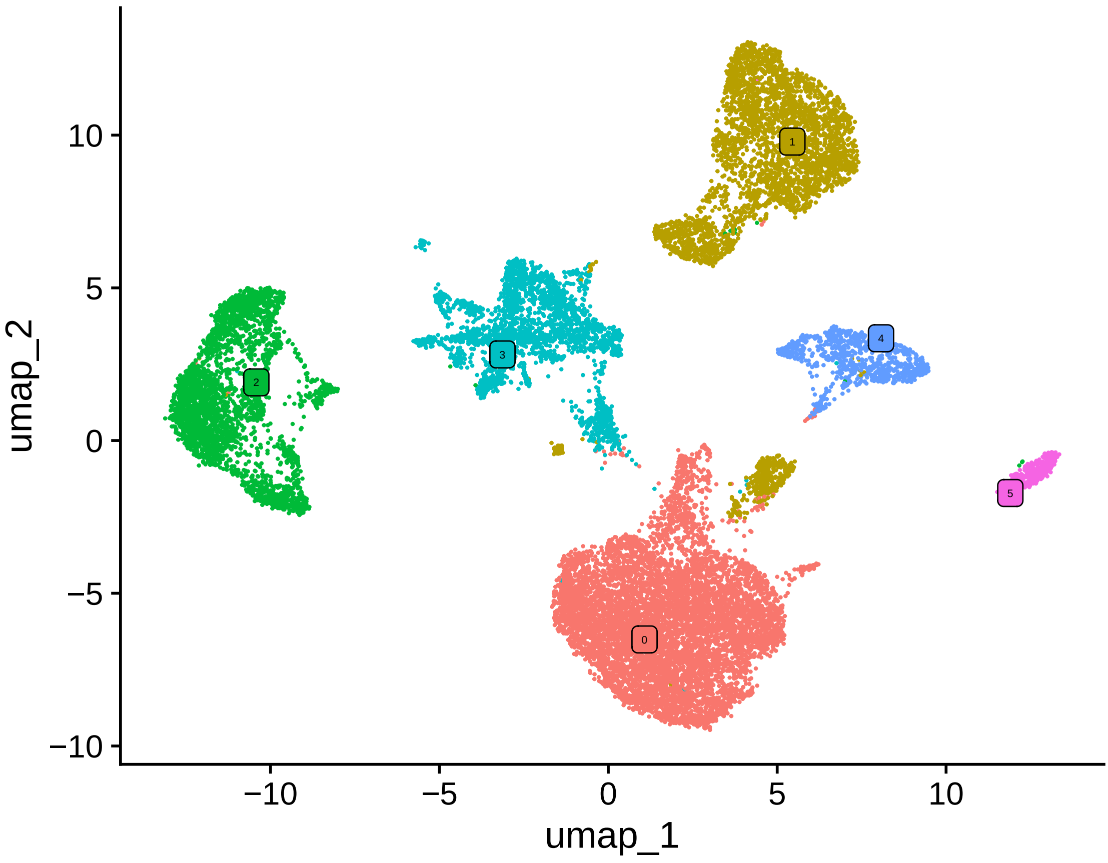


#### Cell Type Annotations

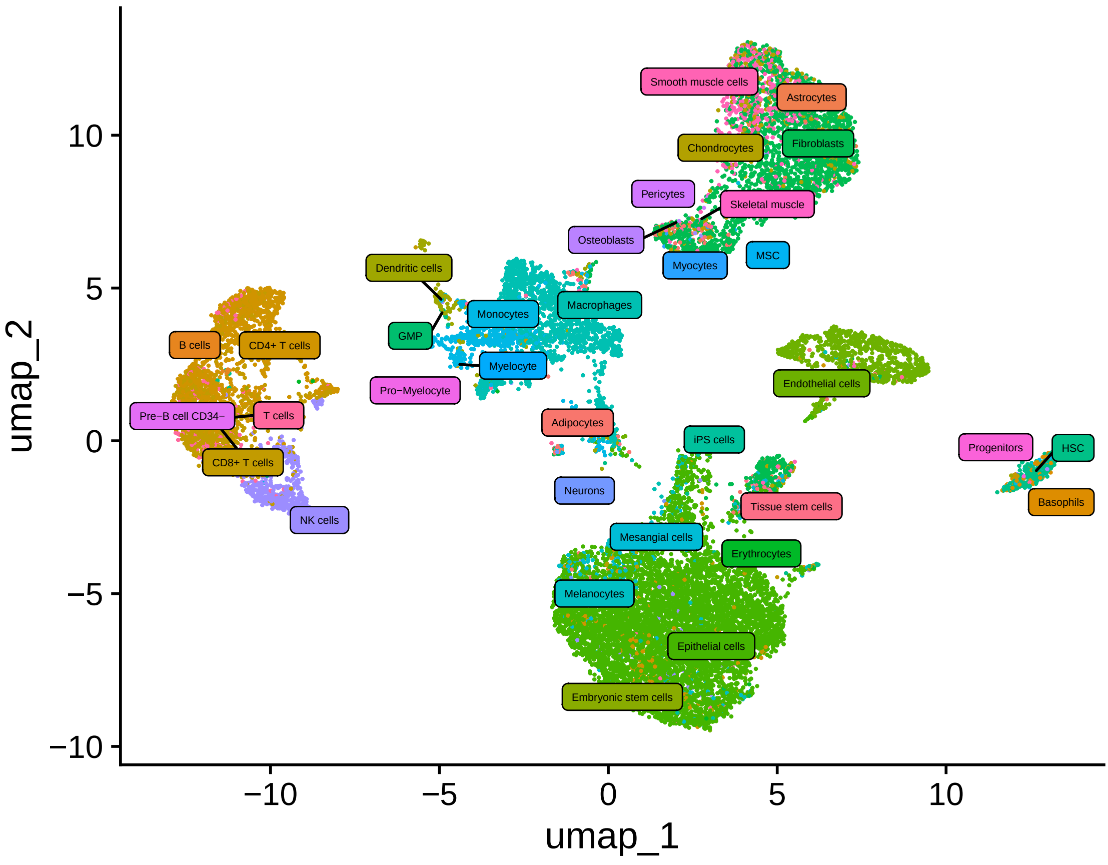

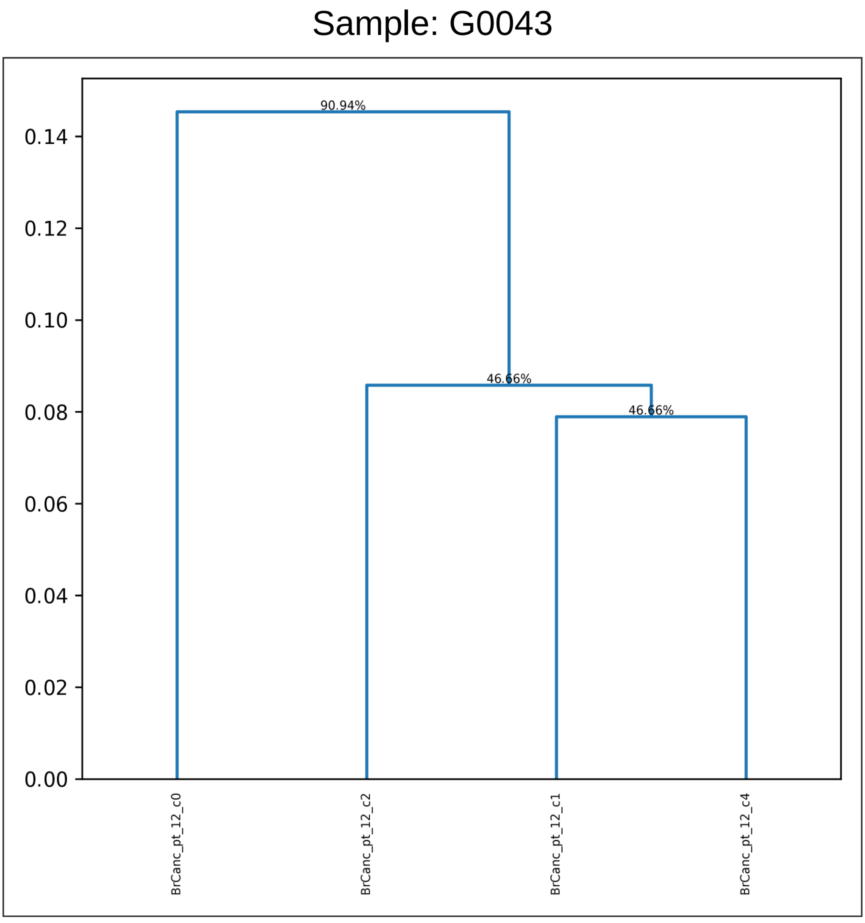
Sample: G0043
90.94%
46.66%
46.66%
0.14
0.12
0.10
0.08
0.06
0.04
0.02
0.00
BrCanc_pt_12_c0
BrCanc_pt_12_c2
BrCanc_pt_12_c1
BrCanc_pt_12_c4

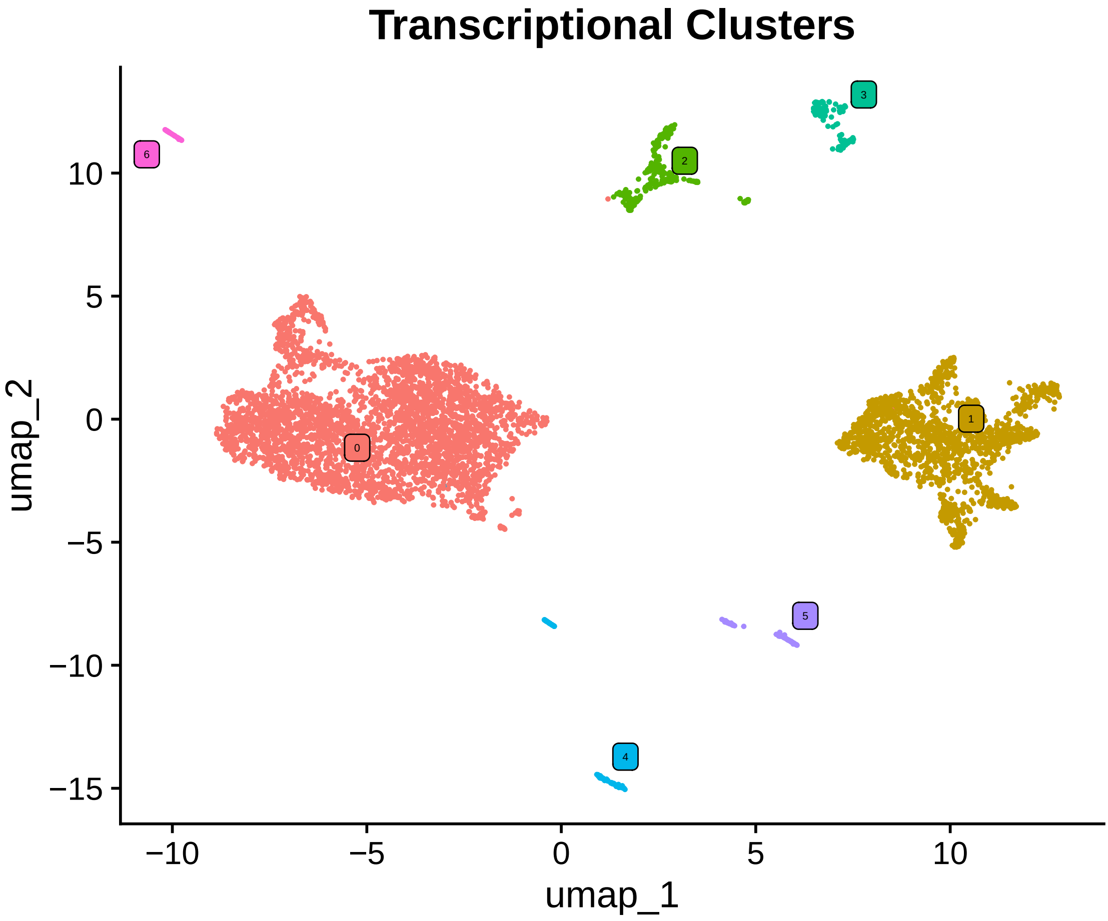
Transcriptional Clusters
umap_2
umap_1
10
5
0
−5
−10
−15
−10
−5
0
5
10
0
1
2
3
4
5
6

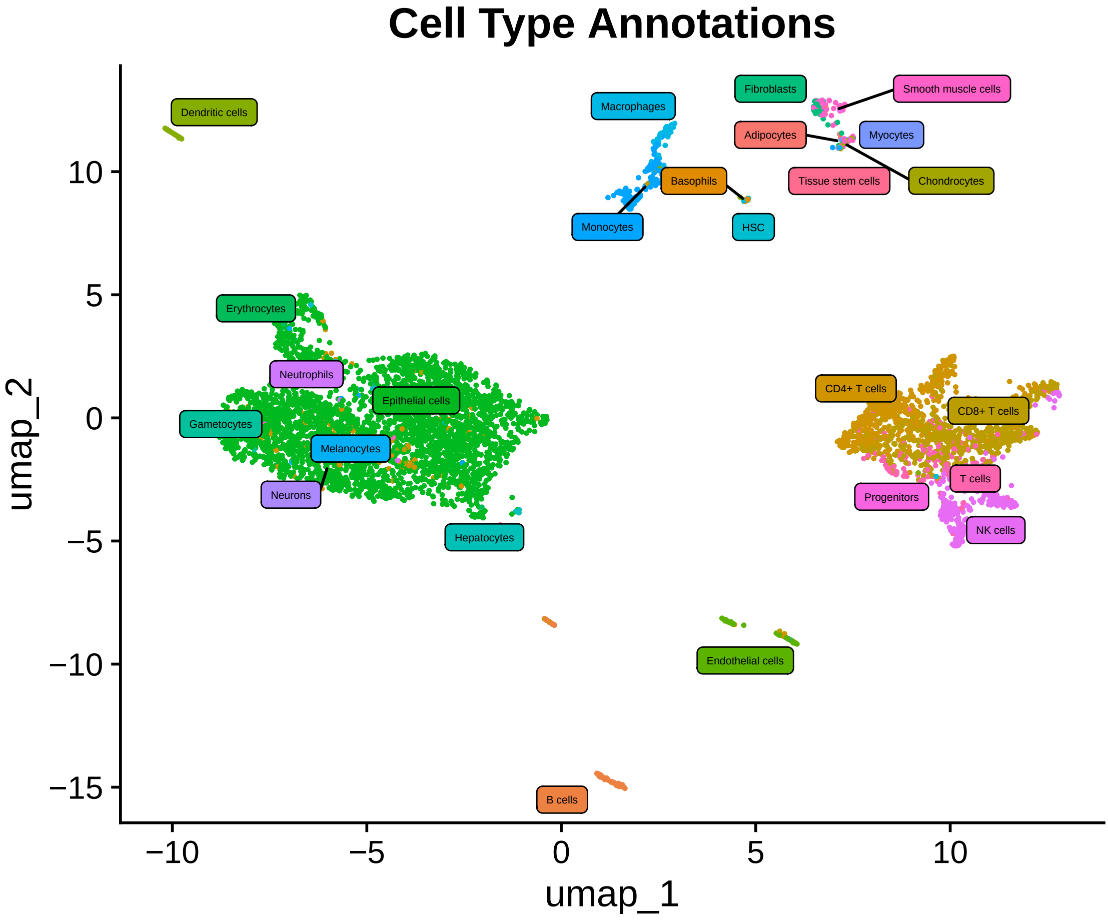
Cell Type Annotations
umap_2
umap_1
Dendritic cells
Macrophages
Fibroblasts
Smooth muscle cells
Adipocytes
Myocytes
Basophils
Tissue stem cells
Chondrocytes
Monocytes
HSC
Erythrocytes
Neutrophils
Epithelial cells
Gametocytes
Melanocytes
Neurons
Hepatocytes
CD4+ T cells
CD8+ T cells
T cells
Progenitors
NK cells
Endothelial cells
B cells
10
5
0
−5
−10
−15
−10
−5
0
5
10

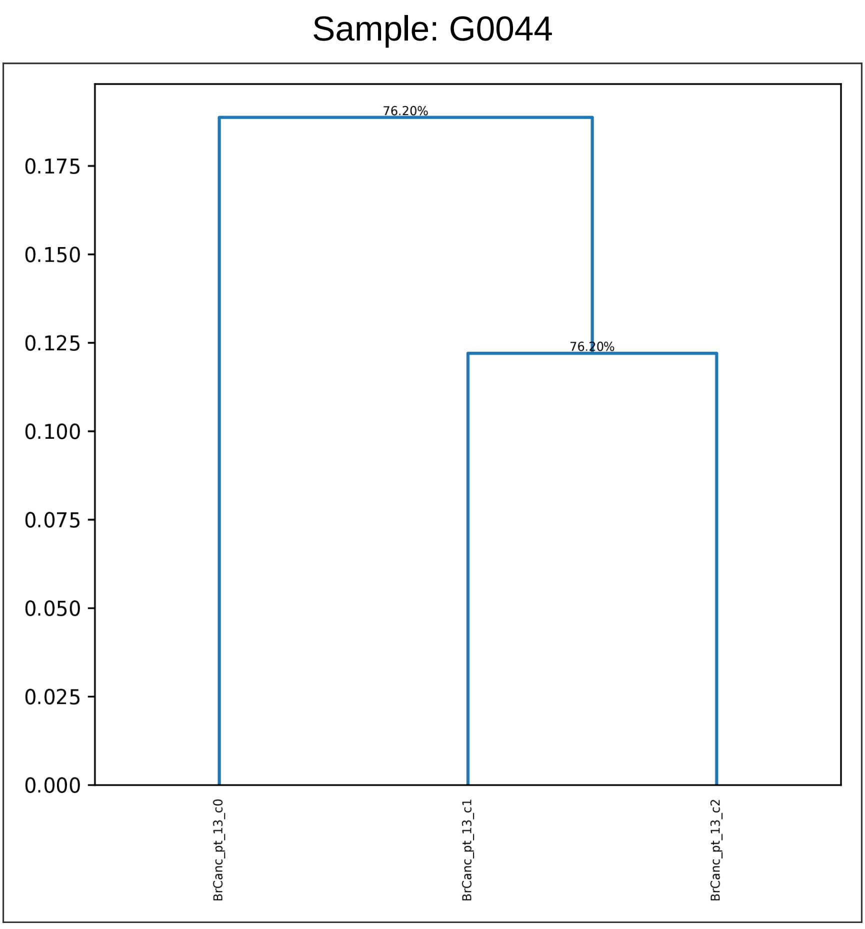
Sample: G0044
76.20%
76.20%
0.175
0.150
0.125
0.100
0.075
0.050
0.025
0.000
BrCanc_pt_13_c0
BrCanc_pt_13_c1
BrCanc_pt_13_c2

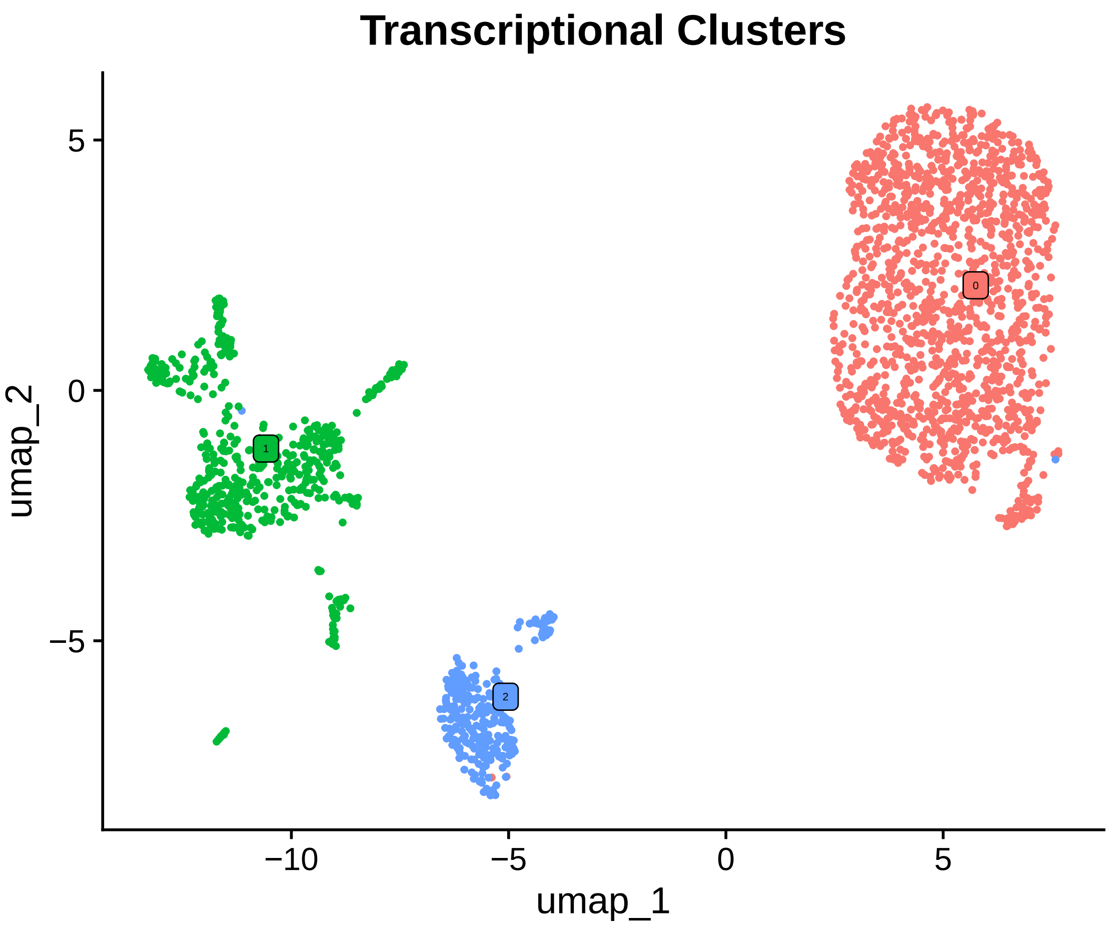
Transcriptional Clusters
umap_2
umap_1
0
1
2

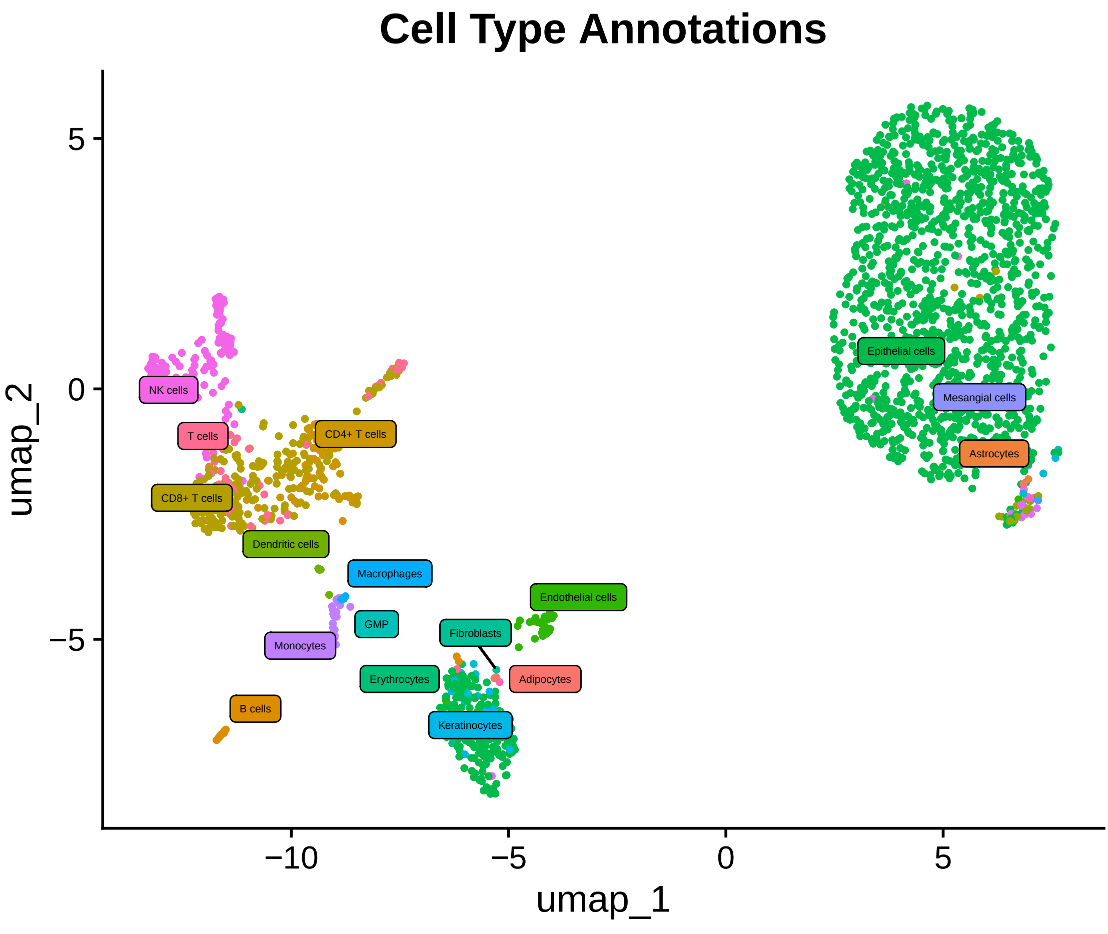
Cell Type Annotations
umap_2
umap_1
NK cells
T cells
CD4+ T cells
CD8+ T cells
Dendritic cells
Macrophages
GMP
Monocytes
B cells
Erythrocytes
Fibroblasts
Adipocytes
Keratinocytes
Endothelial cells
Epithelial cells
Mesangial cells
Astrocytes

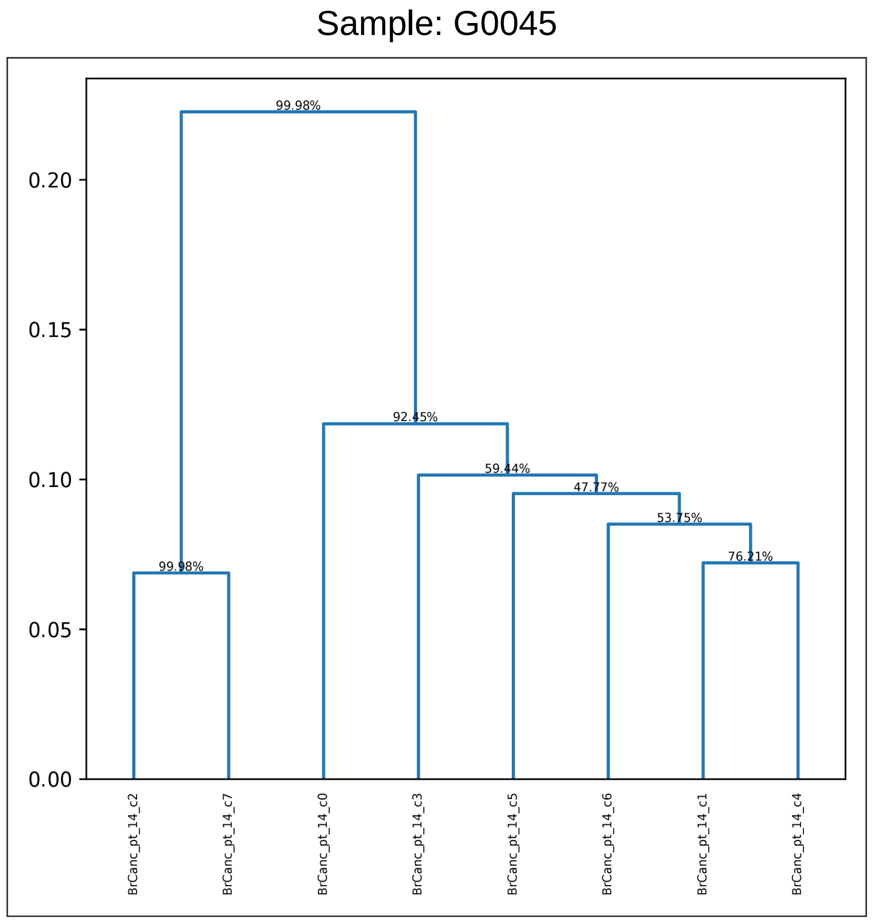
Sample: G0045
0.20
0.15
0.10
0.05
0.00
99.98%
92.45%
59.44%
47.77%
53.75%
76.21%
99.98%
BrCanc_pt_14_c2
BrCanc_pt_14_c7
BrCanc_pt_14_c0
BrCanc_pt_14_c3
BrCanc_pt_14_c5
BrCanc_pt_14_c6
BrCanc_pt_14_c1
BrCanc_pt_14_c4

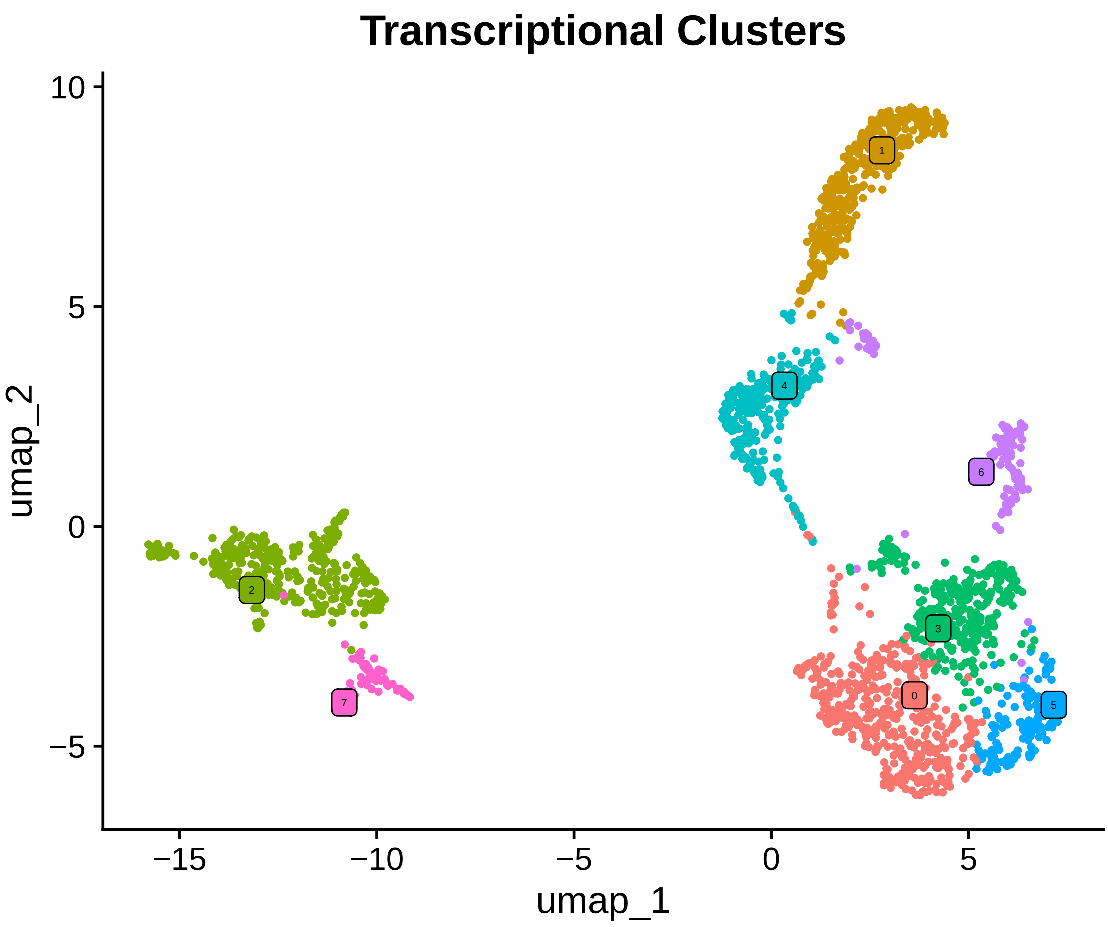
Transcriptional Clusters
10
5
0
−5
umap_2
−15
−10
−5
0
5
umap_1
1
4
6
2
3
7
0
5

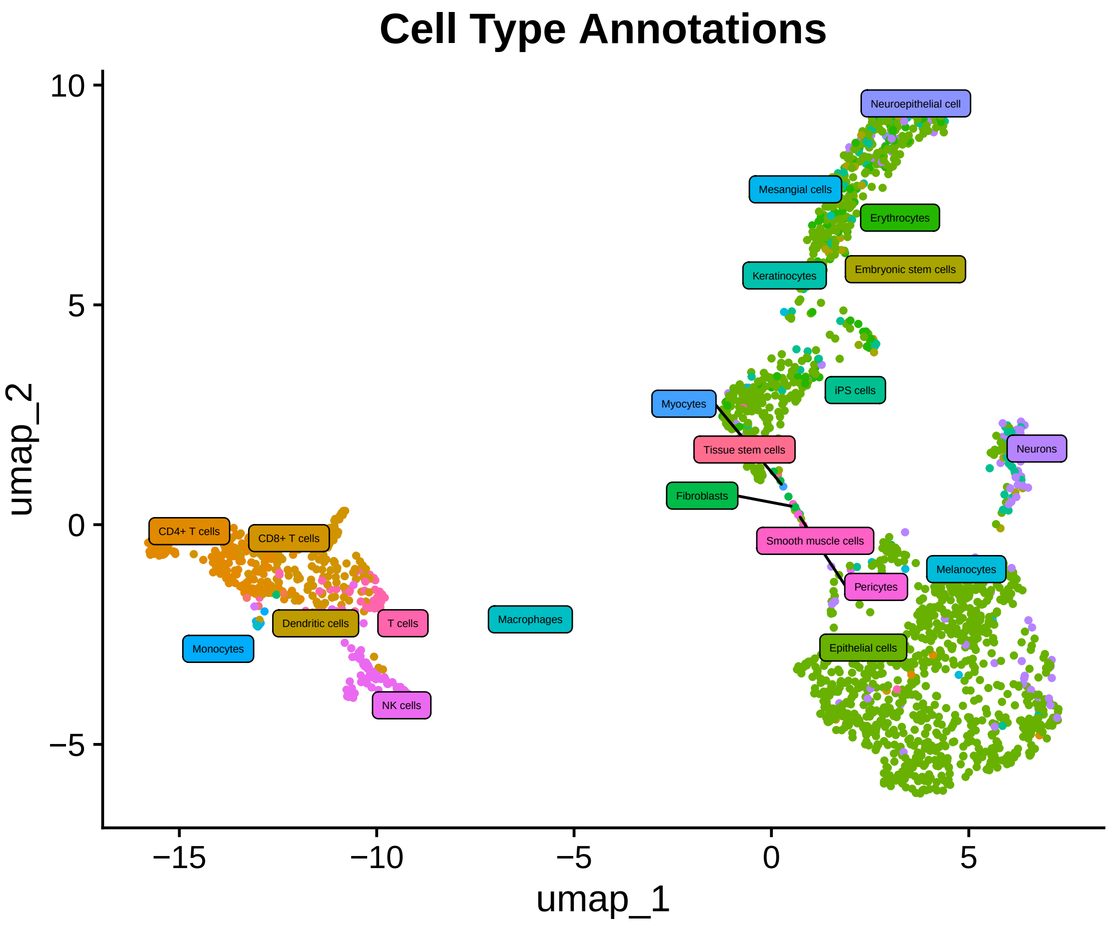
Cell Type Annotations
10
5
0
−5
umap_2
−15
−10
−5
0
5
umap_1
Neuroepithelial cell
Mesangial cells
Erythrocytes
Keratinocytes
Embryonic stem cells
iPS cells
Myocytes
Tissue stem cells
Neurons
Fibroblasts
CD4+ T cells
CD8+ T cells
Smooth muscle cells
Melanocytes
Pericytes
Dendritic cells
T cells
Macrophages
Monocytes
Epithelial cells
NK cells

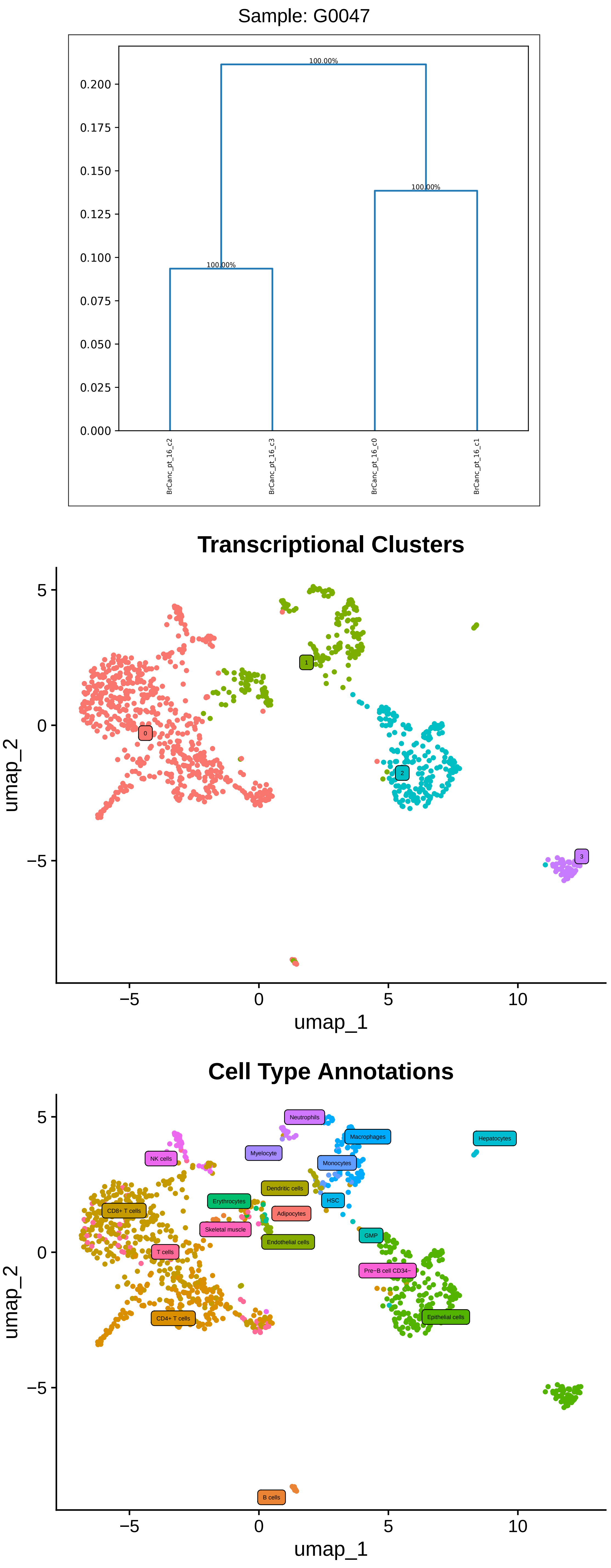

Sample: G0047
100.00%
100.00%
100.00%
BrCanc_pt_16_c2
BrCanc_pt_16_c3
BrCanc_pt_16_c0
BrCanc_pt_16_c1
Transcriptional Clusters
umap_1
umap_2
Cell Type Annotations
Neutrophils
Macrophages
Hepatocytes
NK cells
Myelocyte
Monocytes
Dendritic cells
Erythrocytes
HSC
CD8+ T cells
Adipocytes
Skeletal muscle
Endothelial cells
GMP
T cells
Pre−B cell CD34−
CD4+ T cells
Epithelial cells
B cells

Sample: G0048

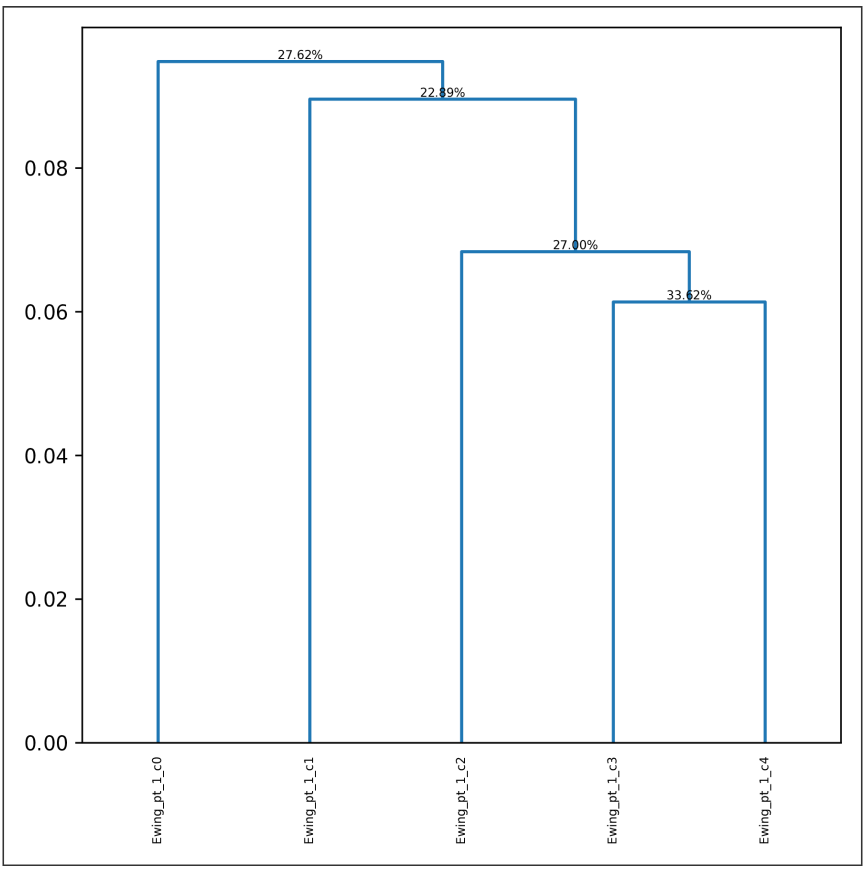


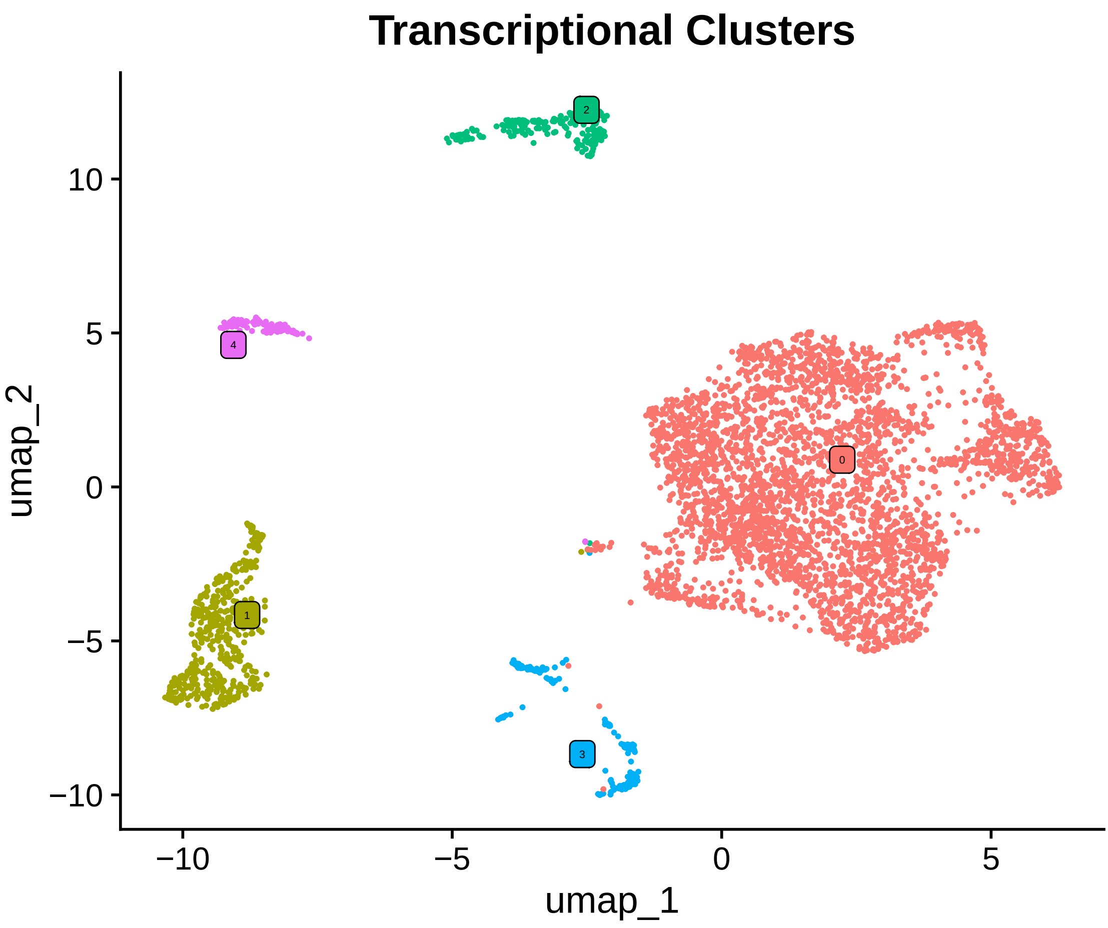


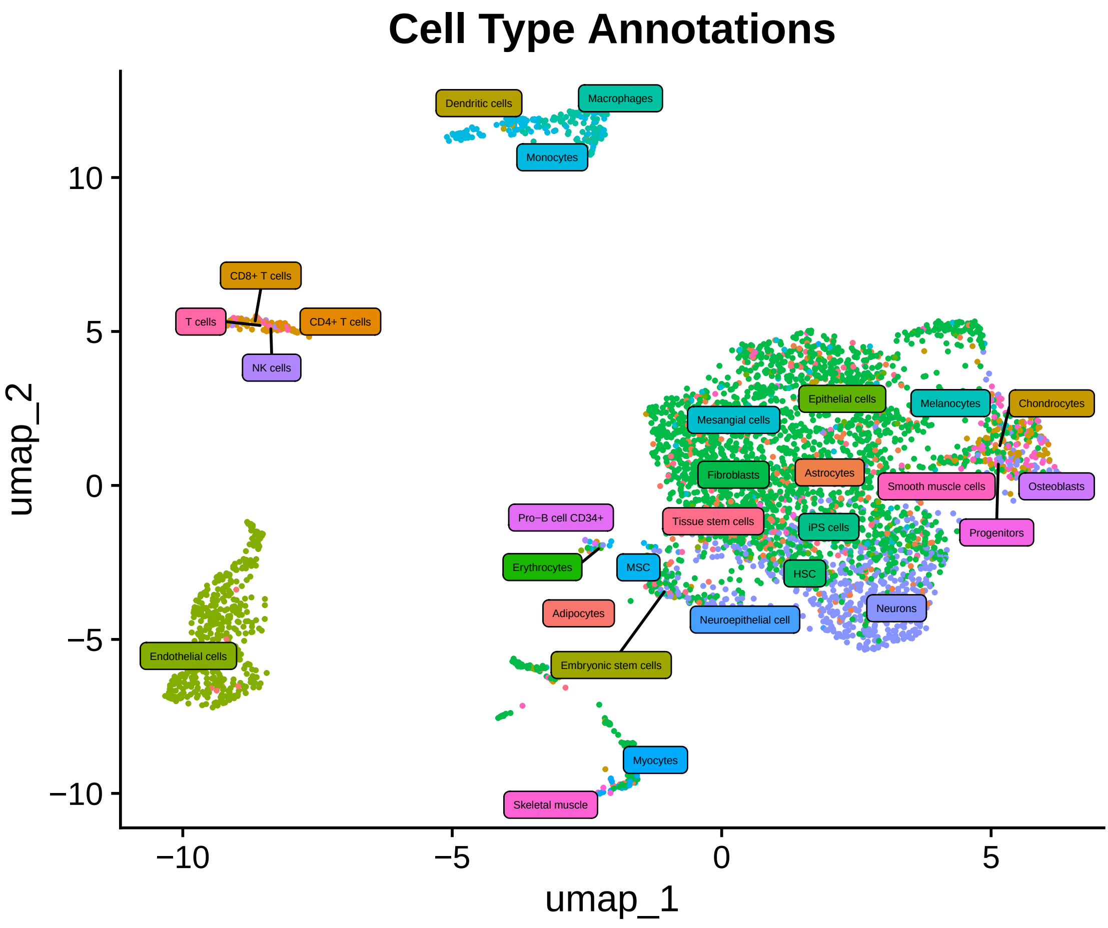

Sample: G0049

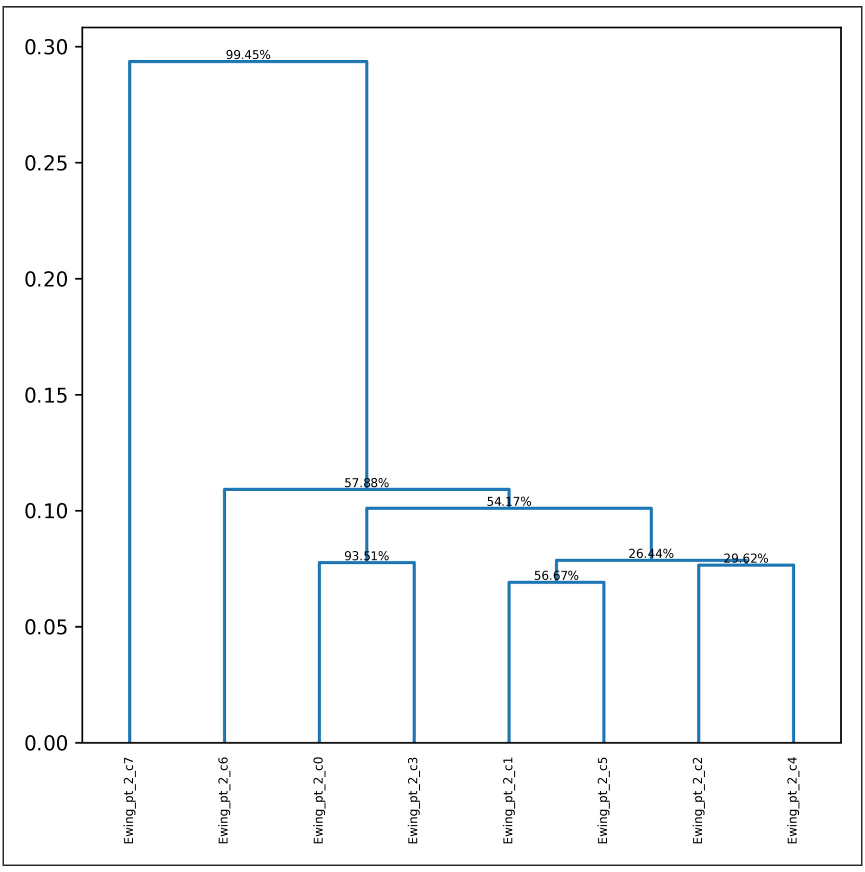


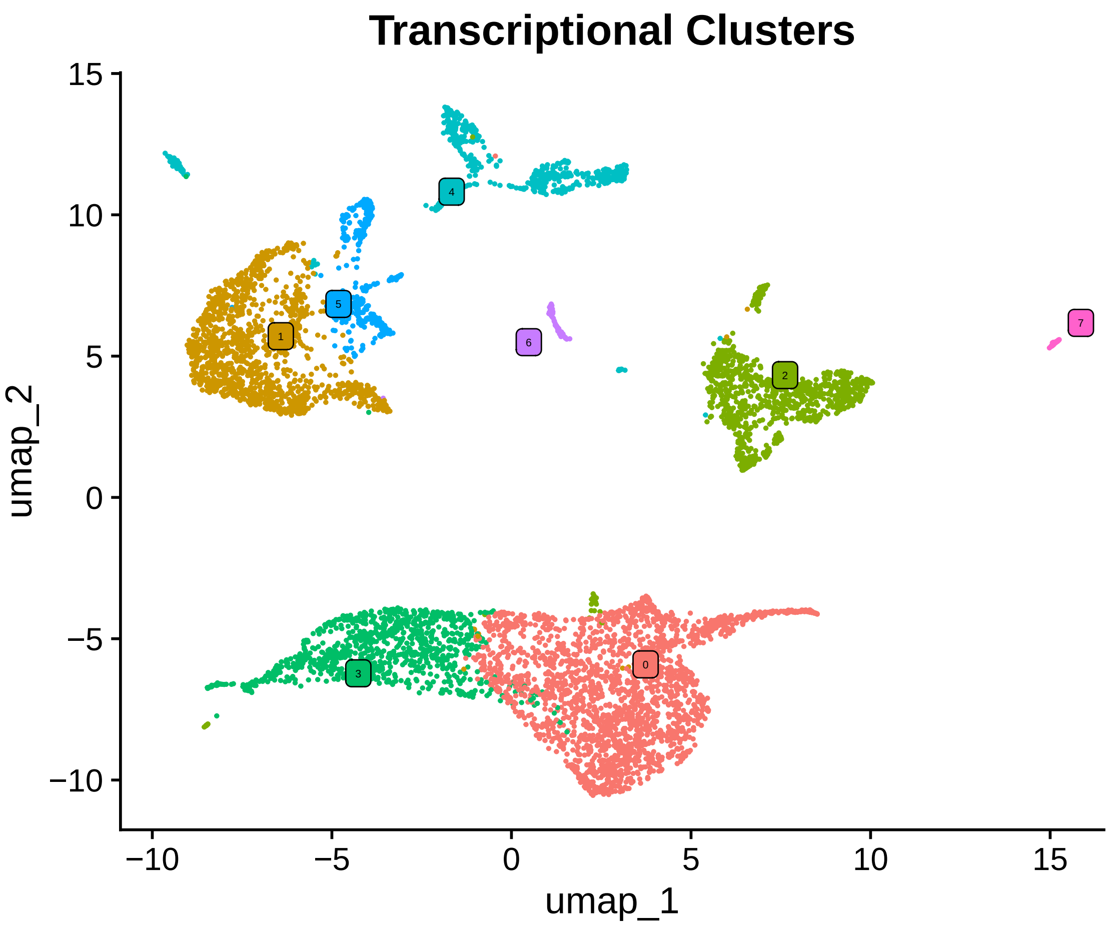


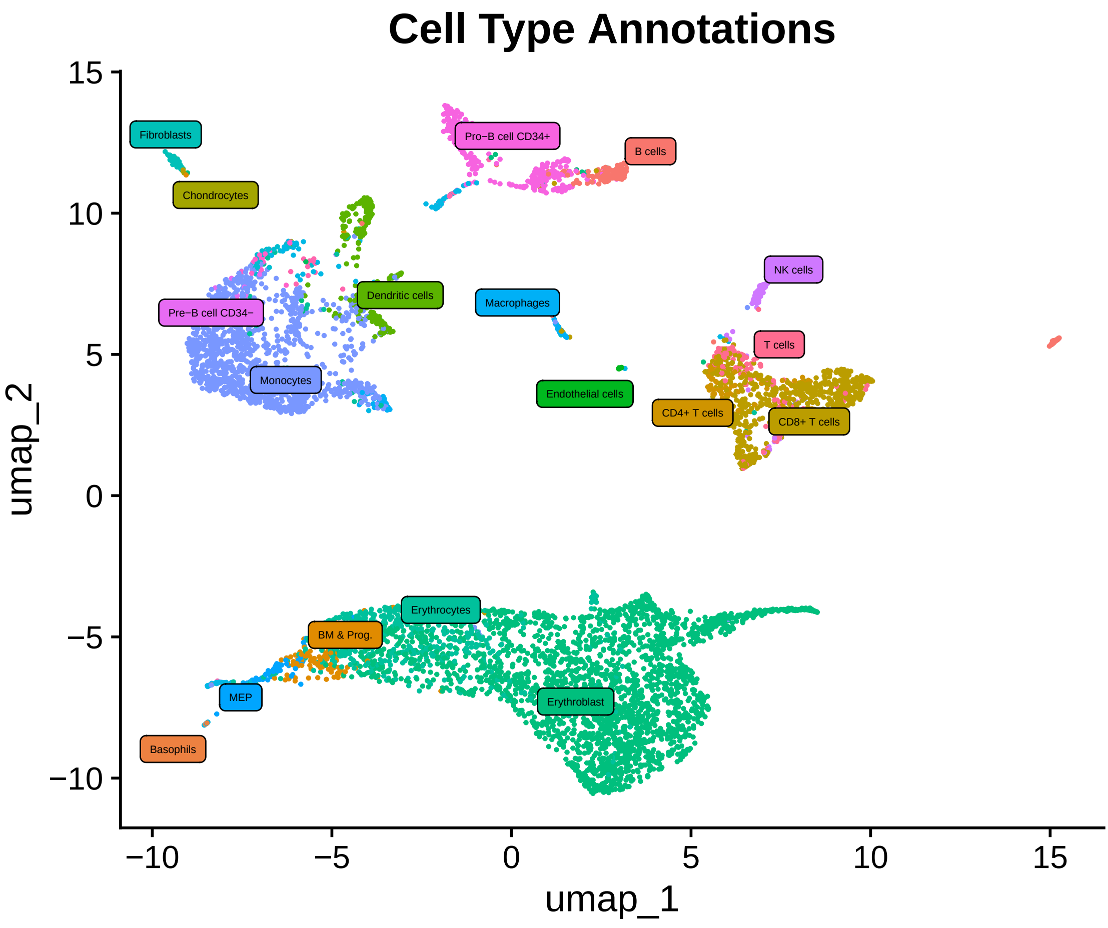

**Supplemental Figure 7: Hierarchical clustering trees on human patient samples**

We tested scanBit on human patient samples including healthy and tumor samples. For each sample we generated a hierarchical clustering tree using a minimum read depth threshold of 10 for variant calling, a UMAP plot showing transcriptional clusters and a UMAP showing reference-based cell type annotation. A table of the metadata for each sample is available in Supplemental Table 1.

Sample: G0001 with all cells per cluster

Min. read depth: 10,  Mean UMIs per cluster:  9,921,855

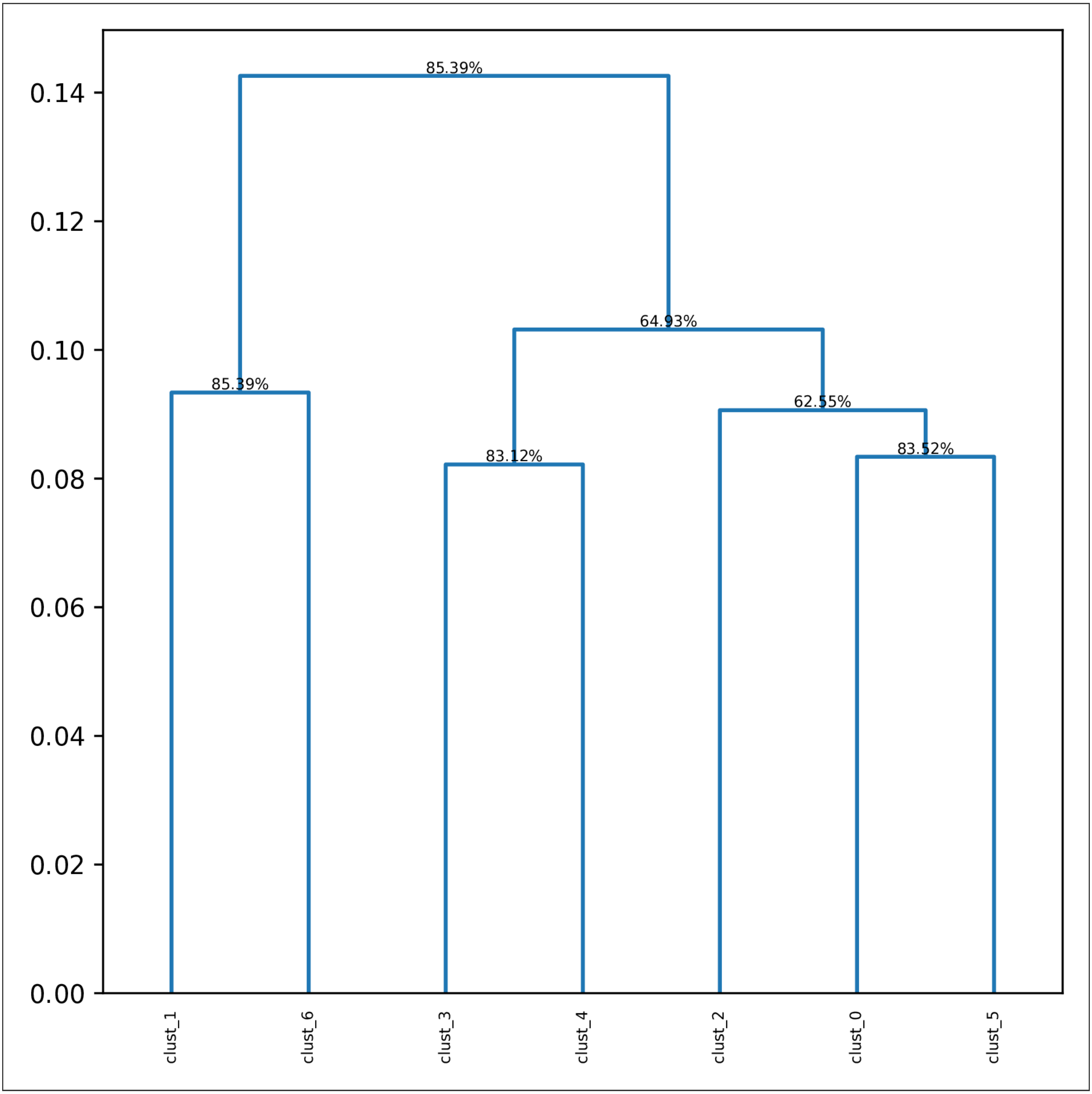

Sample: G0001 with 500 cells per cluster

Min. read depth: 10,  Mean UMIs per cluster:  5,241,268

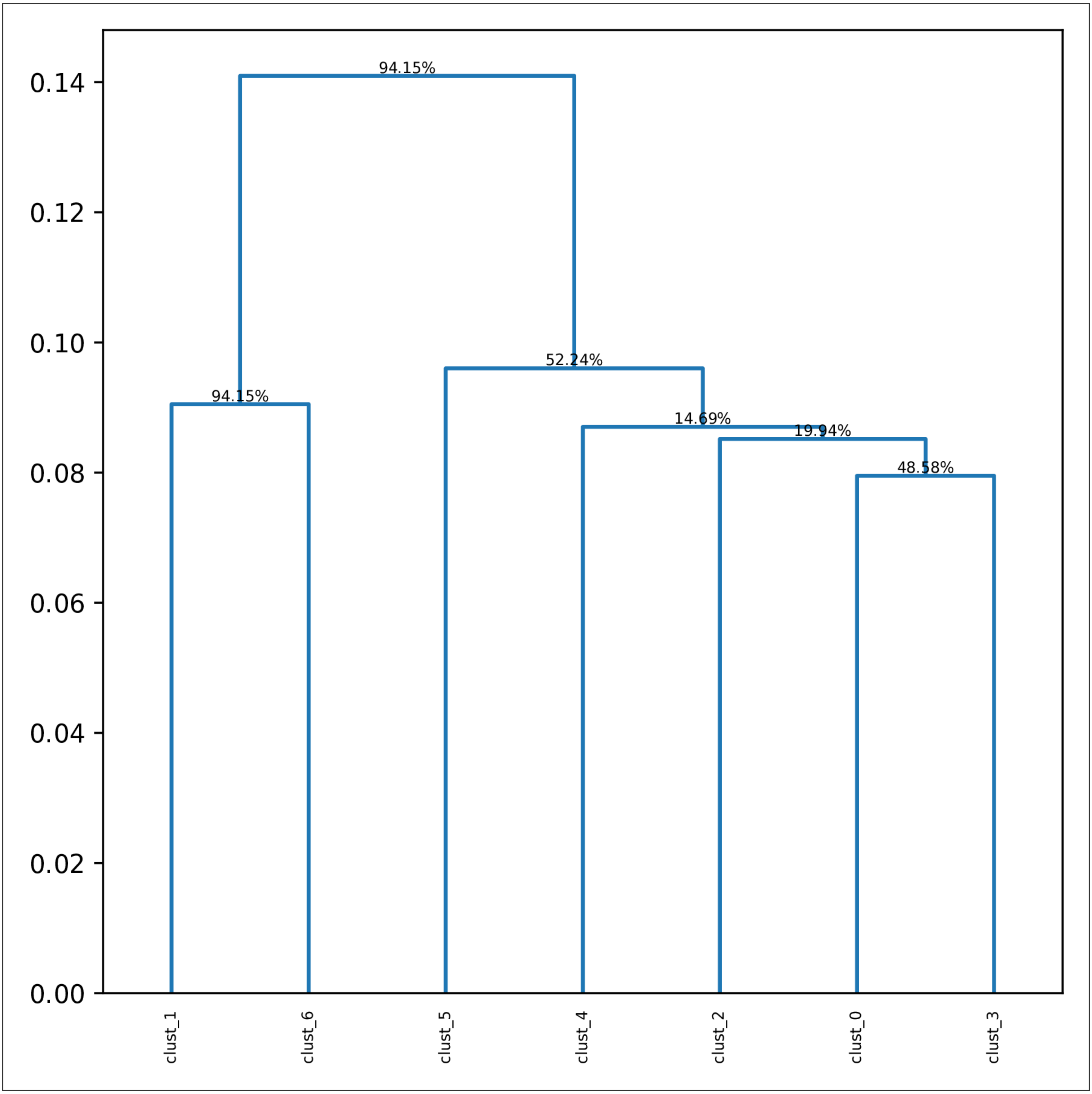

Sample: G0001 with 100 cells per cluster

Min. read depth: 10,  Mean UMIs per cluster:  1,088,514

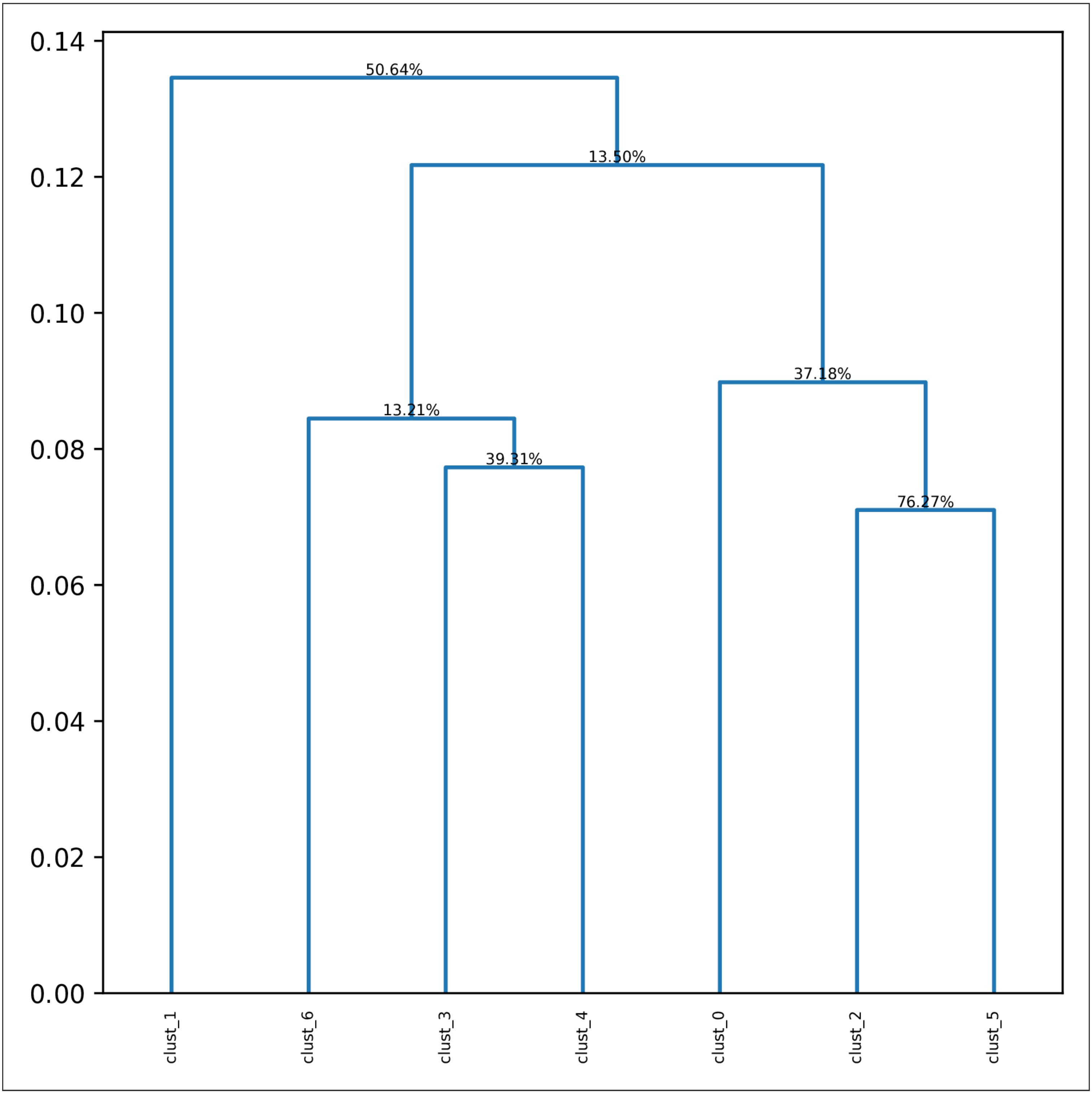

Sample: G0001 with 50 cells per cluster

Min. read depth: 10,  Mean UMIs per cluster:  539,028

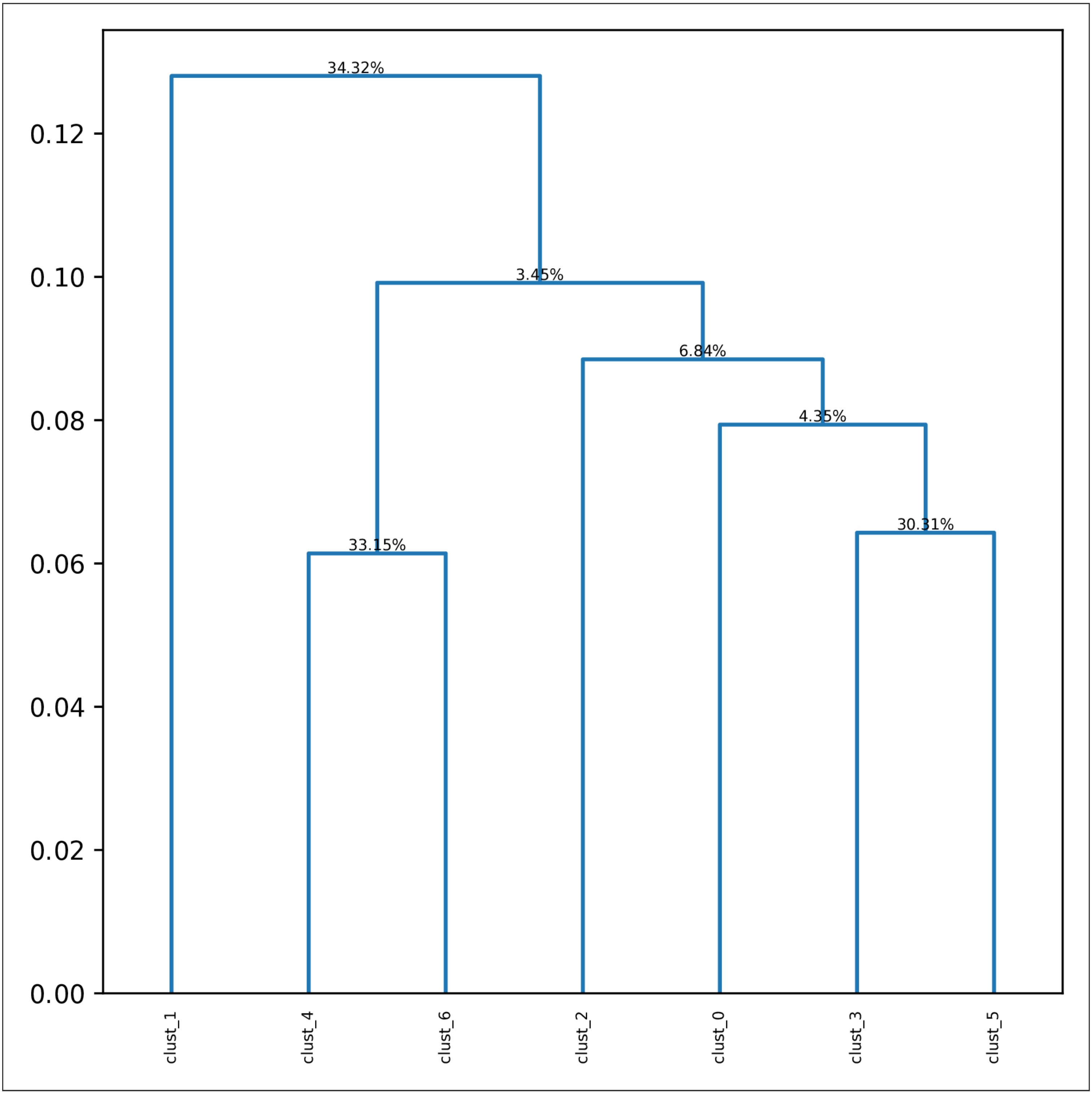

Sample: G0001 with 20 cells per cluster

Min. read depth: 10, Mean UMIs per cluster: 203,656

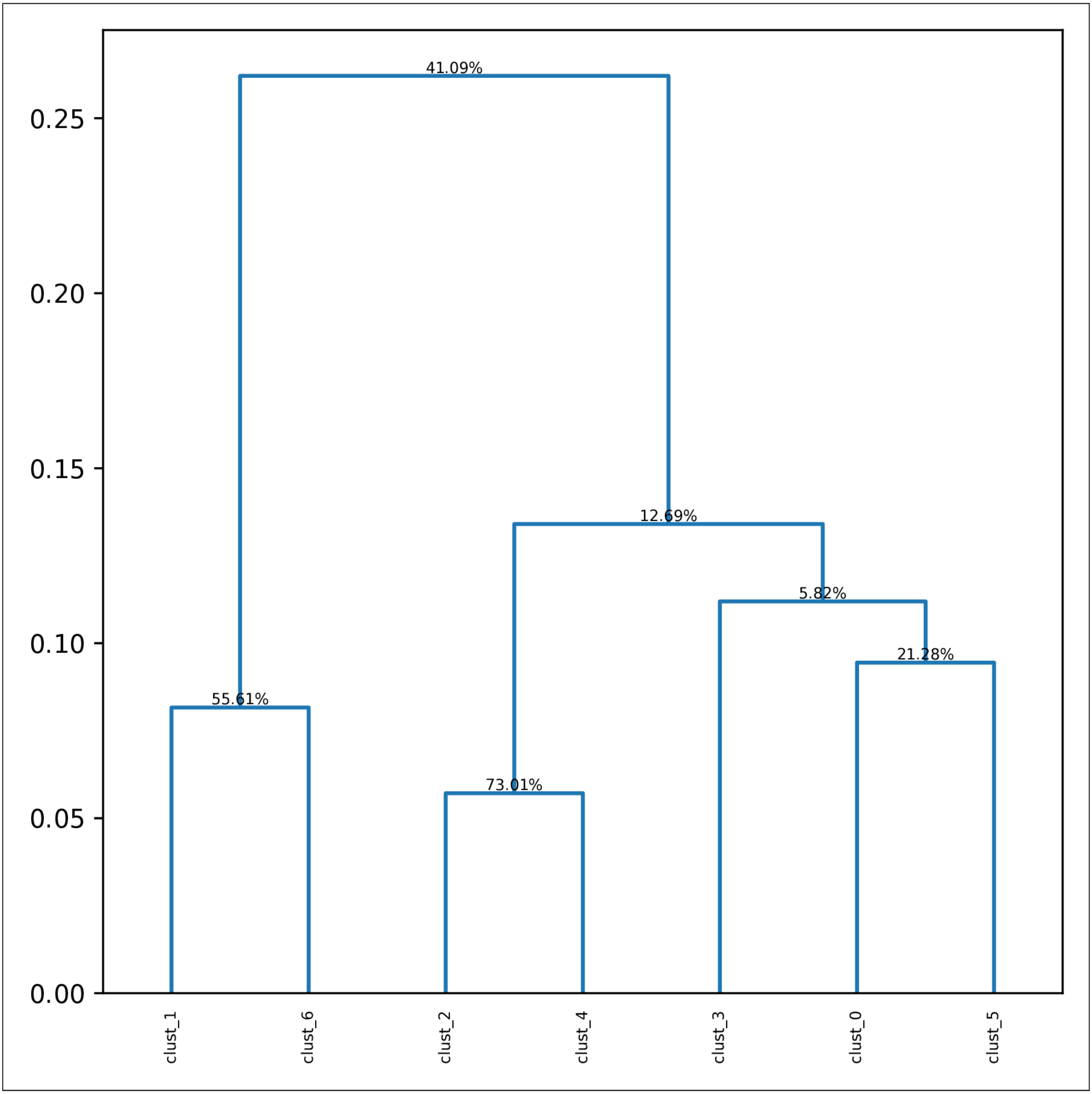

Sample: G0002 with all cells per cluster

Min. read depth: 10,  Mean UMIs per cluster:  9,583,939

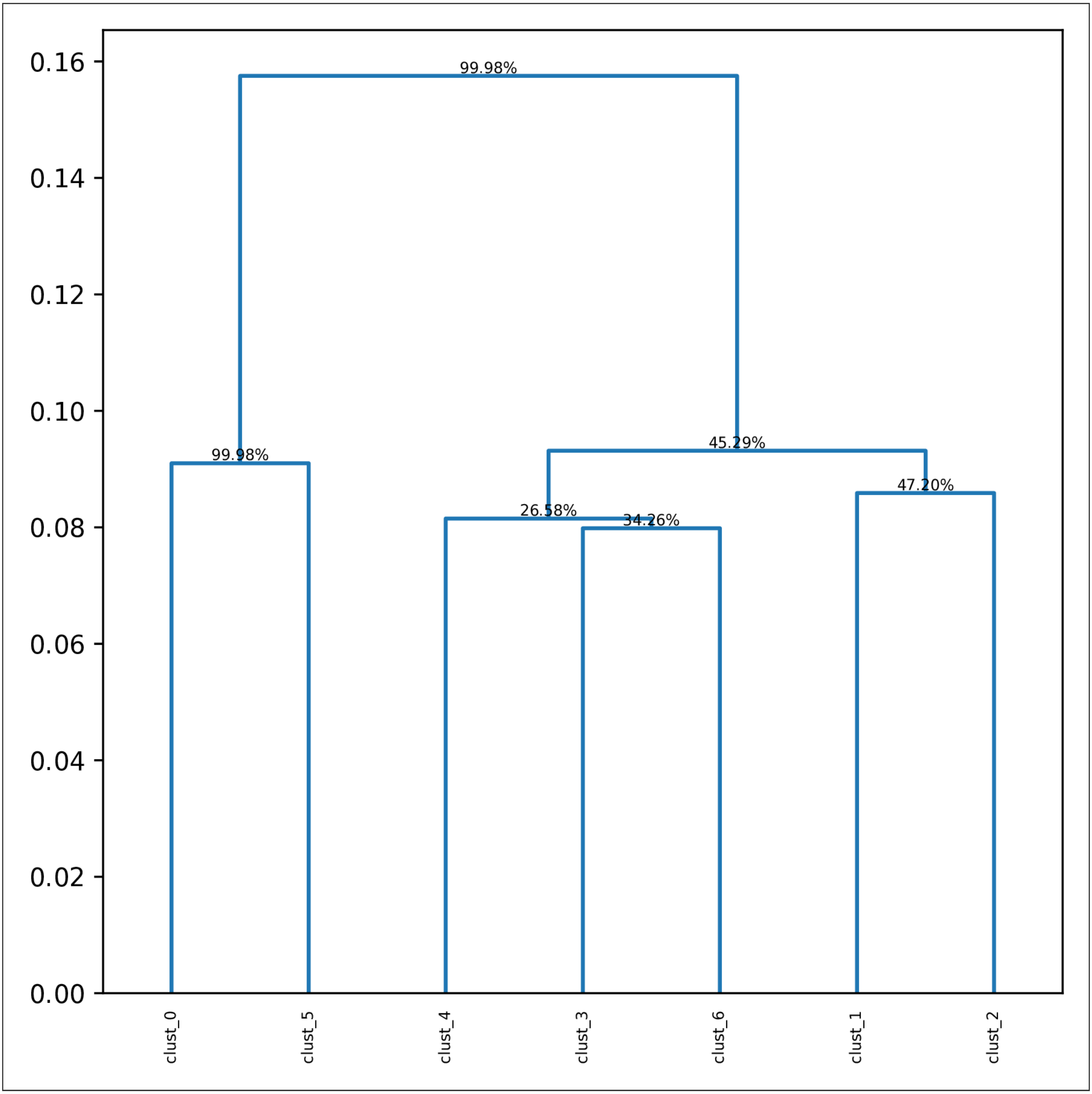

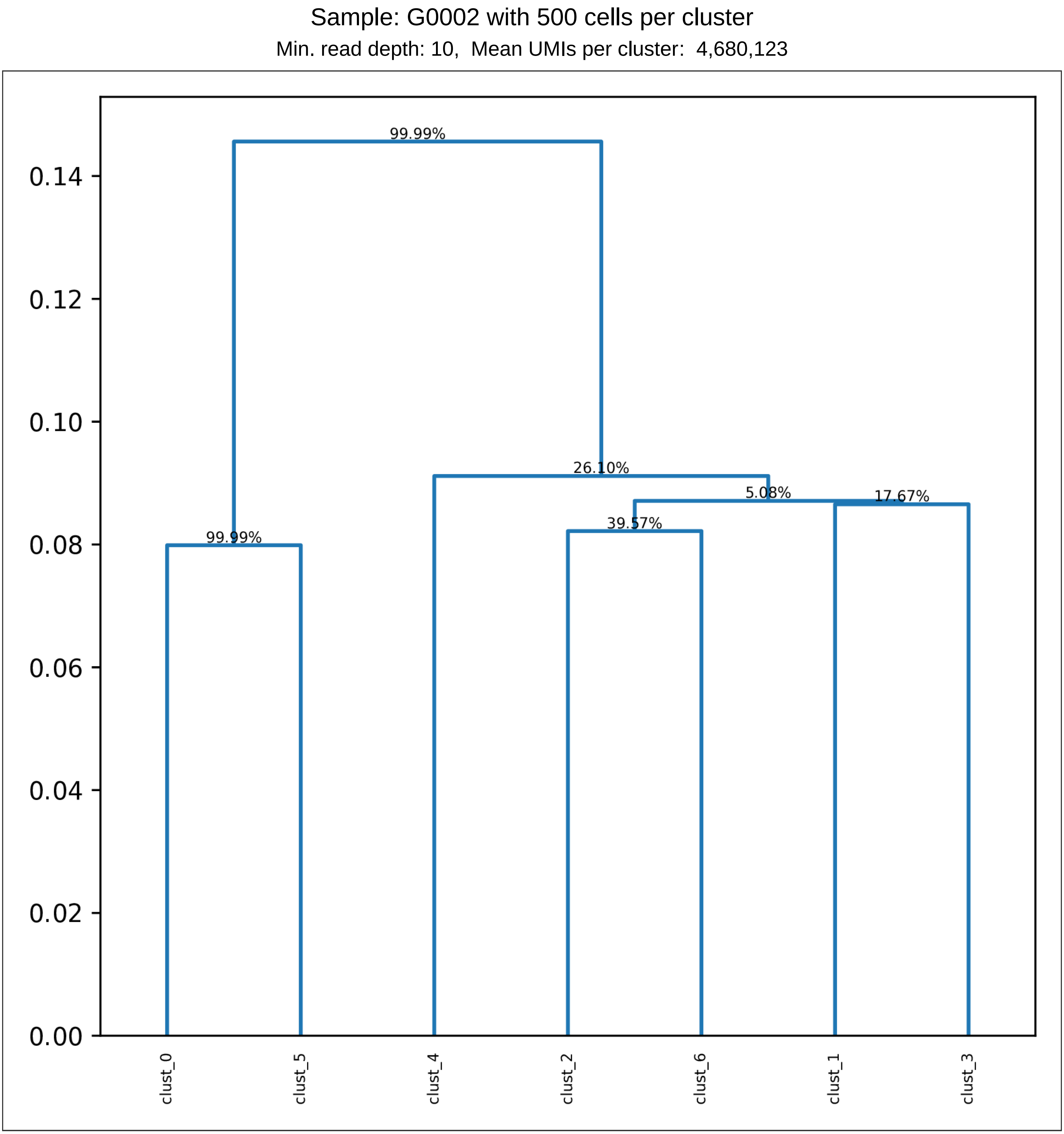
Sample: G0002 with 500 cells per cluster
Min. read depth: 10, Mean UMIs per cluster: 4,680,123
99.99%
26.10%
5.08%
17.67%
39.57%
99.99%
0.14
0.12
0.10
0.08
0.06
0.04
0.02
0.00
clust_0
clust_5
clust_4
clust_2
clust_6
clust_1
clust_3

Sample: G0002 with 100 cells per cluster

Min. read depth: 10,  Mean UMIs per cluster:  969,590

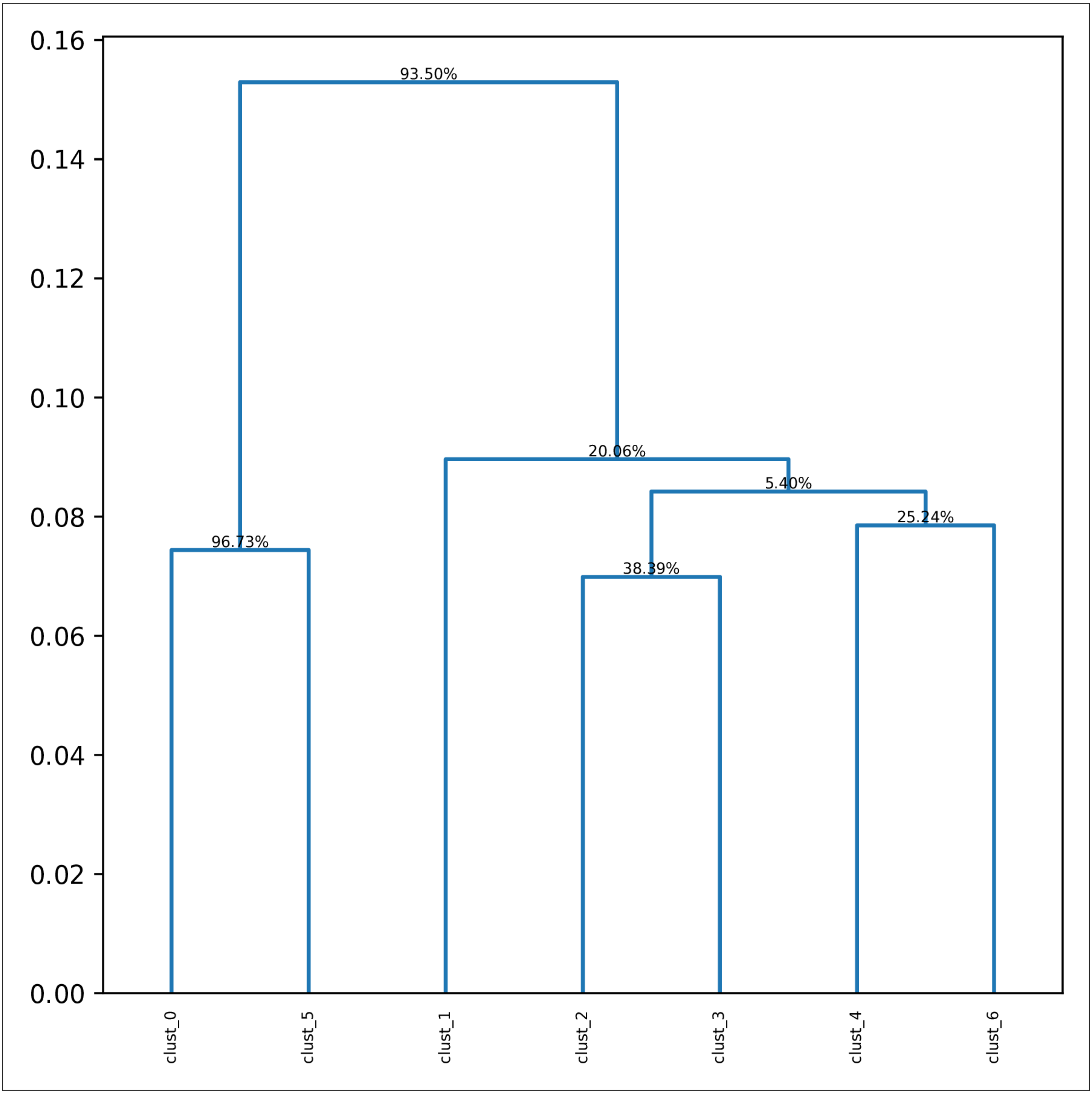

Sample: G0002 with 50 cells per cluster

Min. read depth: 10,  Mean UMIs per cluster:  493,207

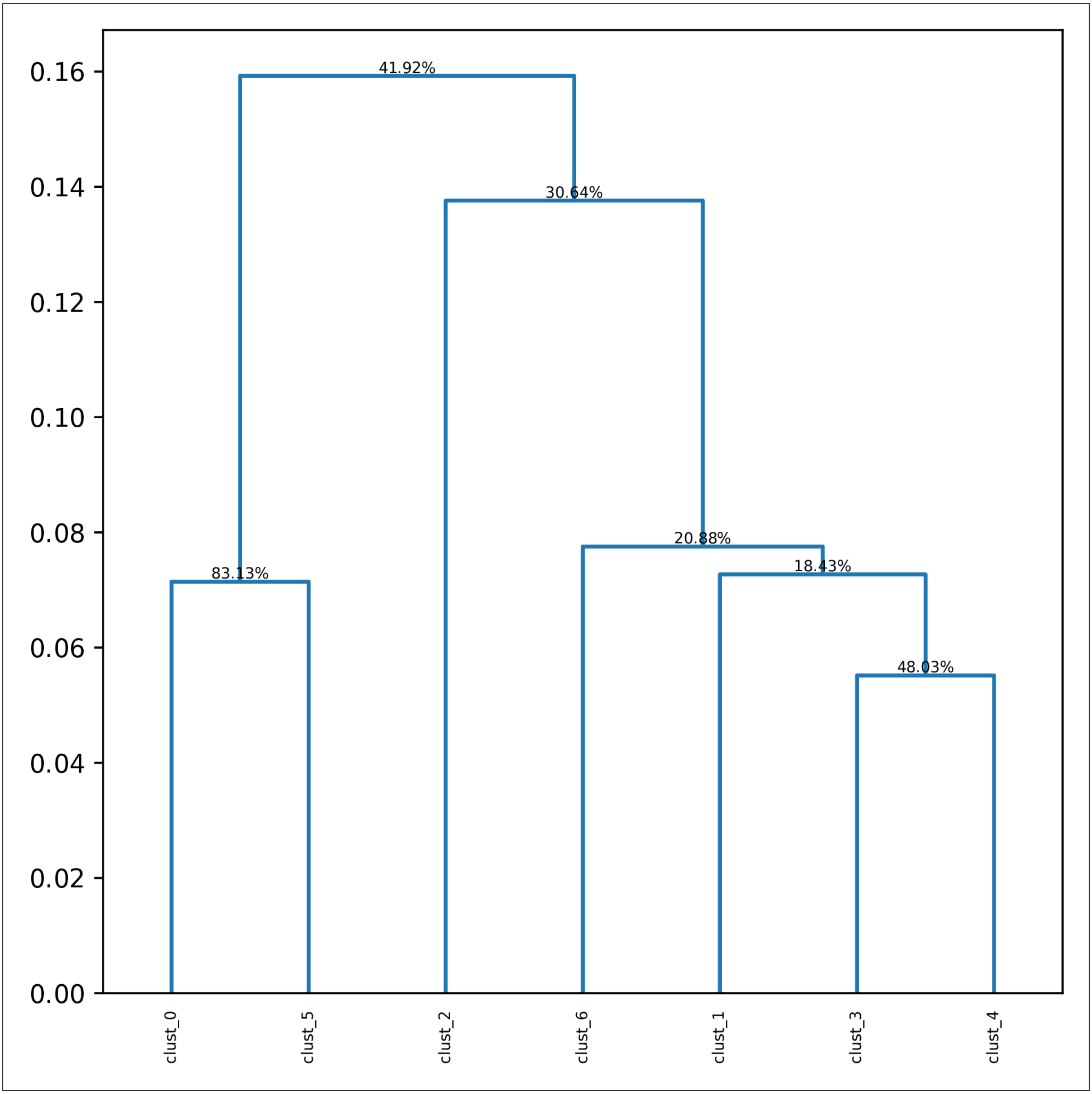

Sample: G0002 with 20 cells per cluster

Min. read depth: 10,  Mean UMIs per cluster:  186,940

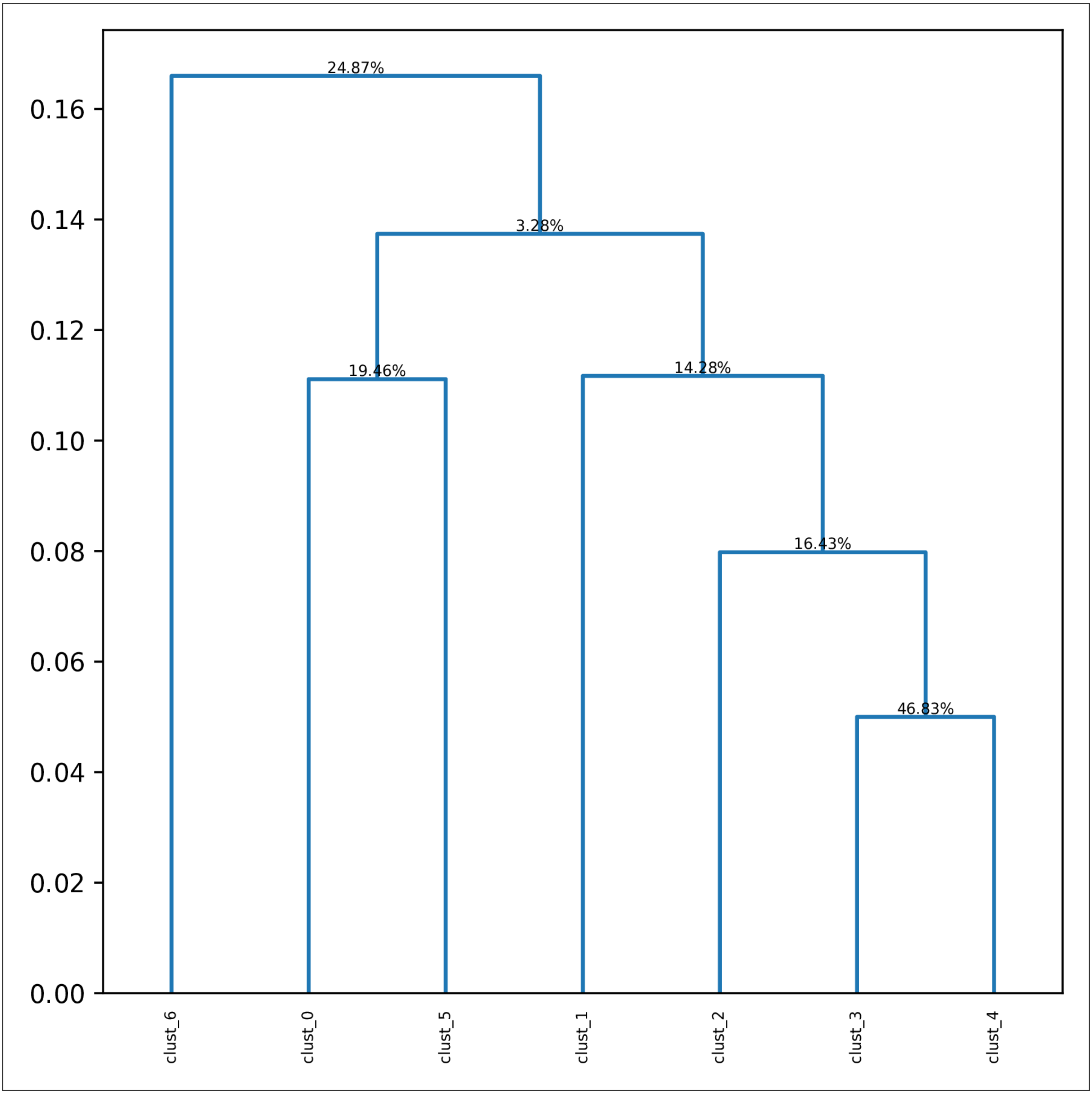

Sample: G0003 with all cells per cluster

Min. read depth: 10,  Mean UMIs per cluster:  9,920,104

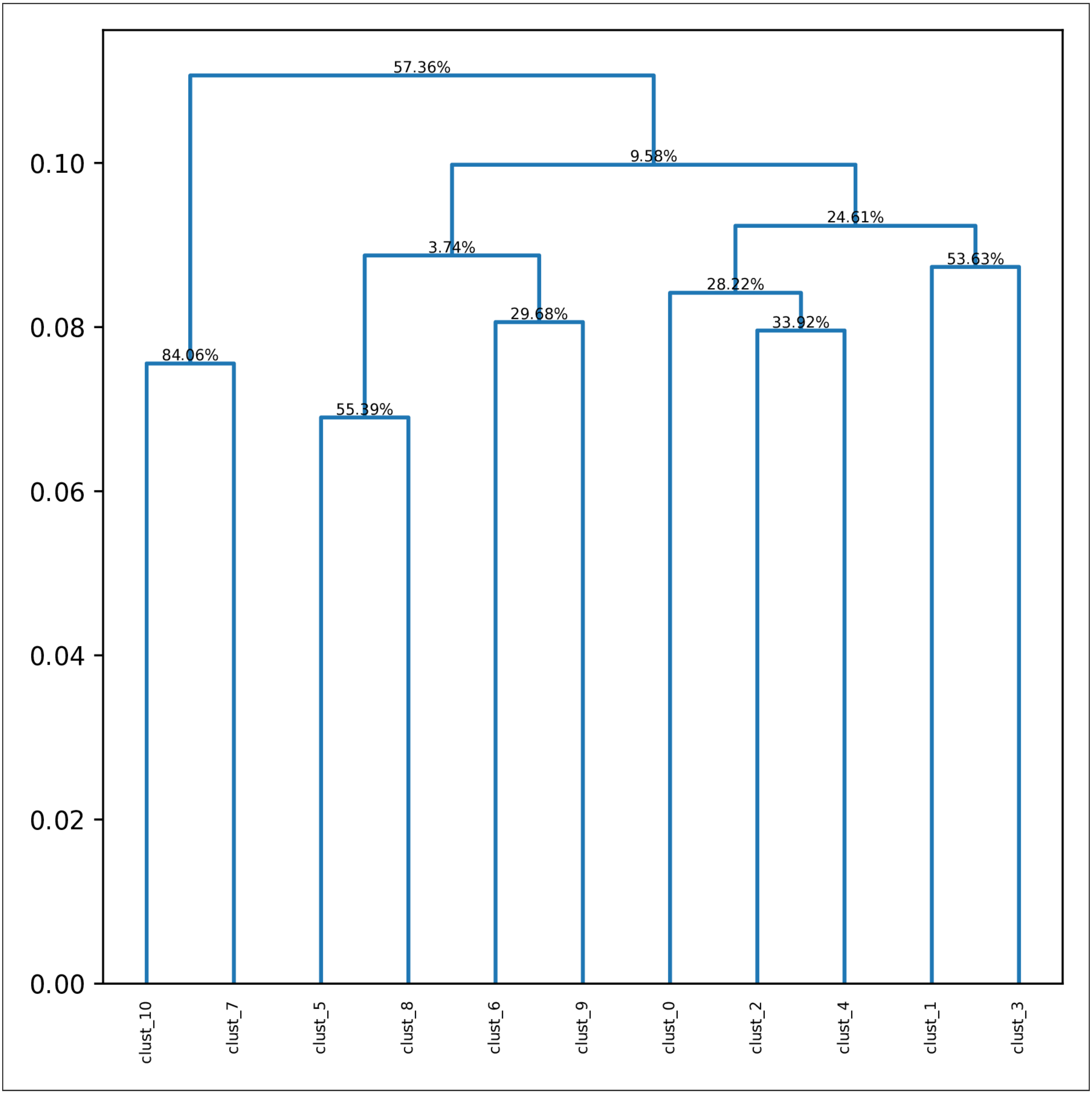

Sample: G0003 with 500 cells per cluster

Min. read depth: 10,  Mean UMIs per cluster:  4,217,144

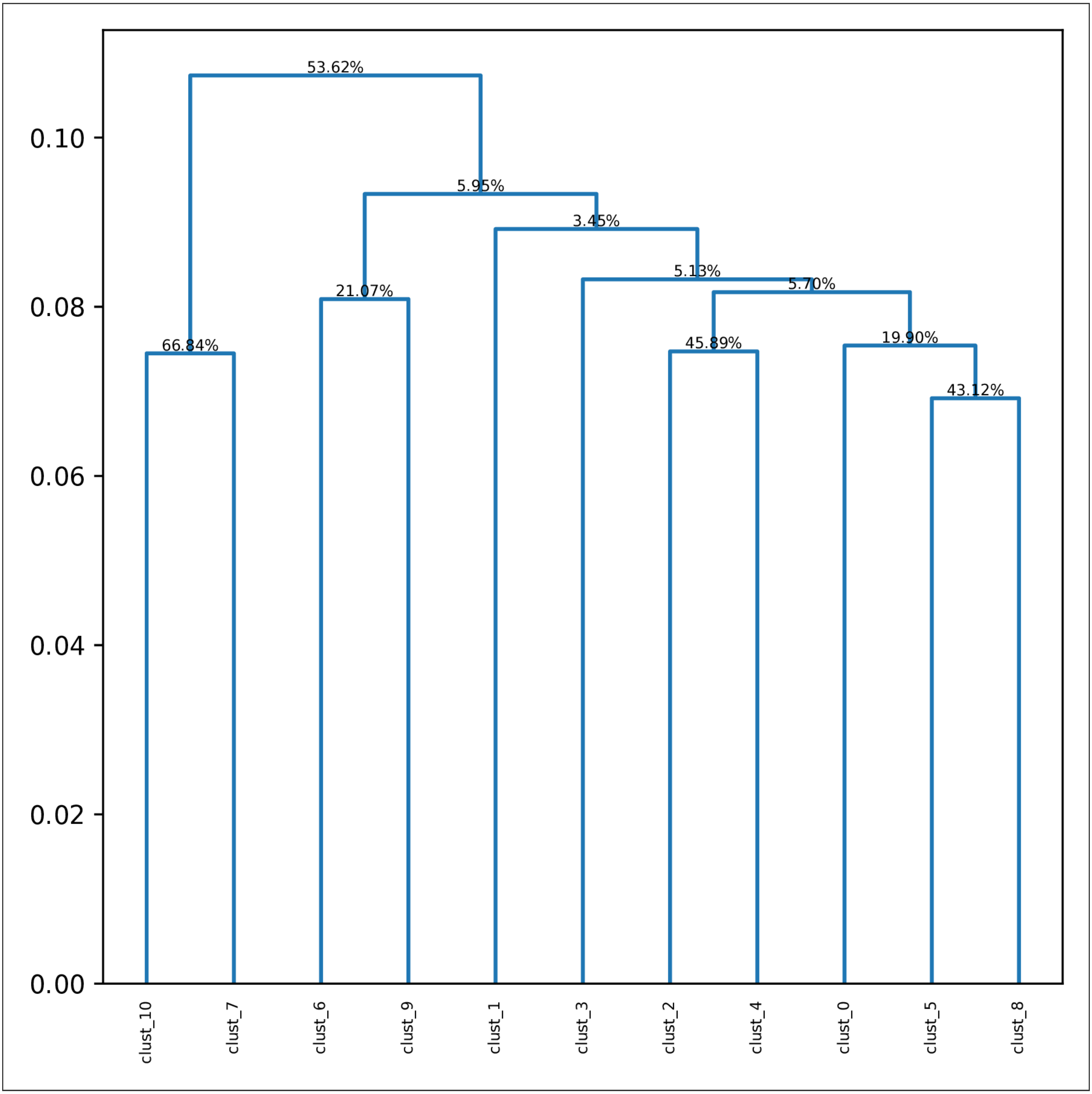

Sample: G0003 with 100 cells per cluster

Min. read depth: 10,  Mean UMIs per cluster:  950,367

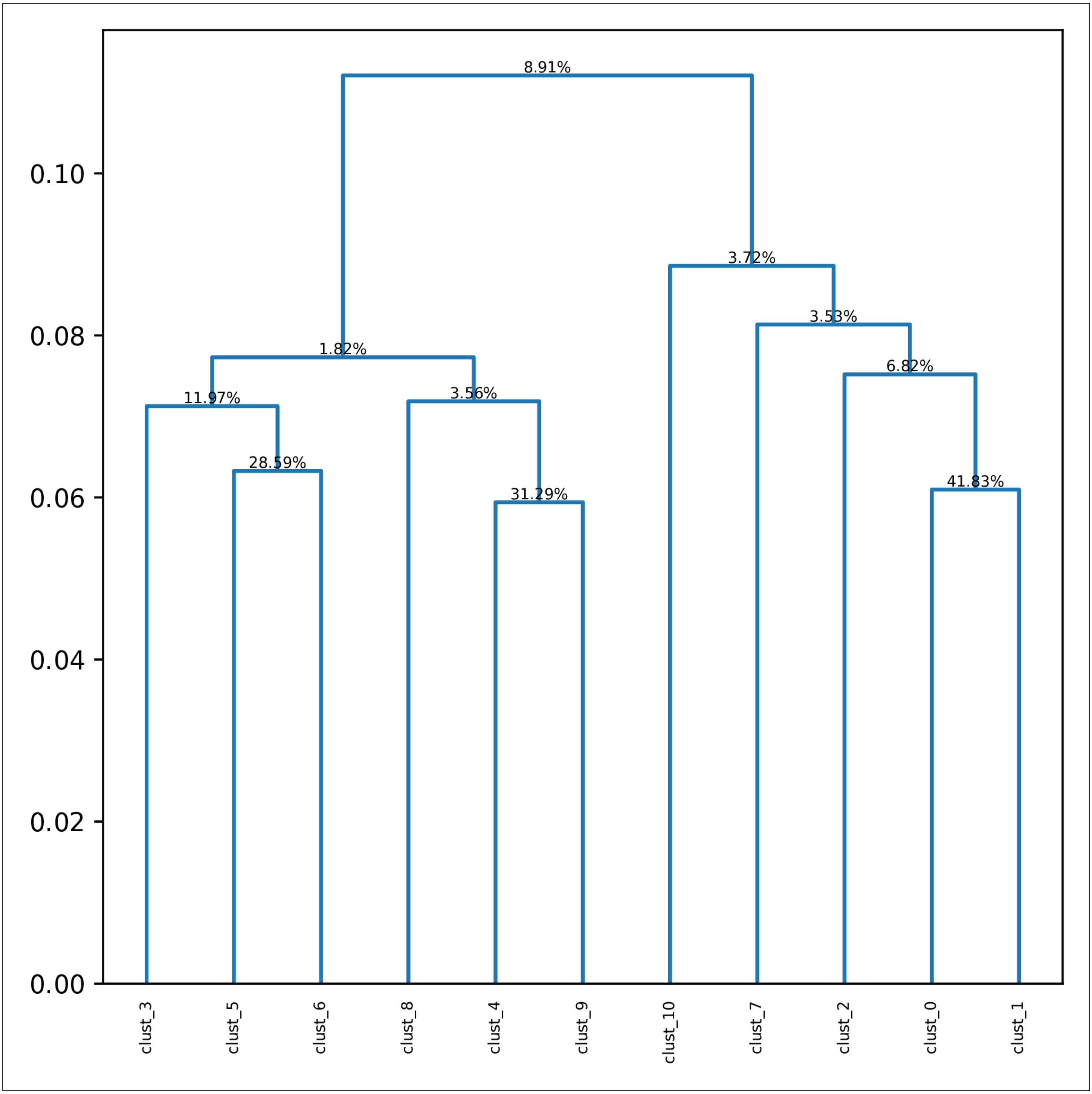

Sample: G0003 with 50 cells per cluster

Min. read depth: 10,  Mean UMIs per cluster:  501,417

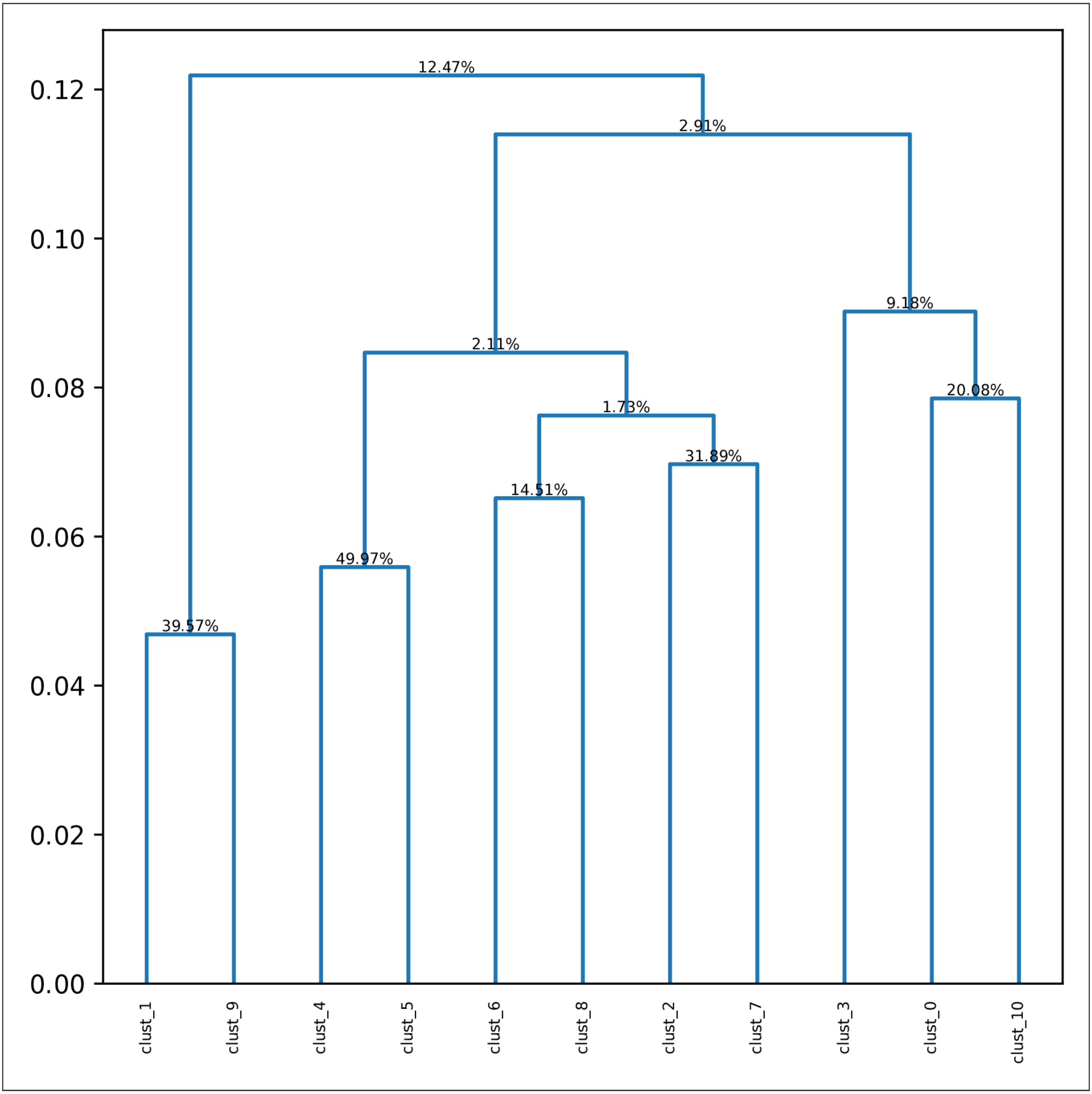

Sample: G0003 with 20 cells per cluster

Min. read depth: 10, Mean UMIs per cluster: 194,344

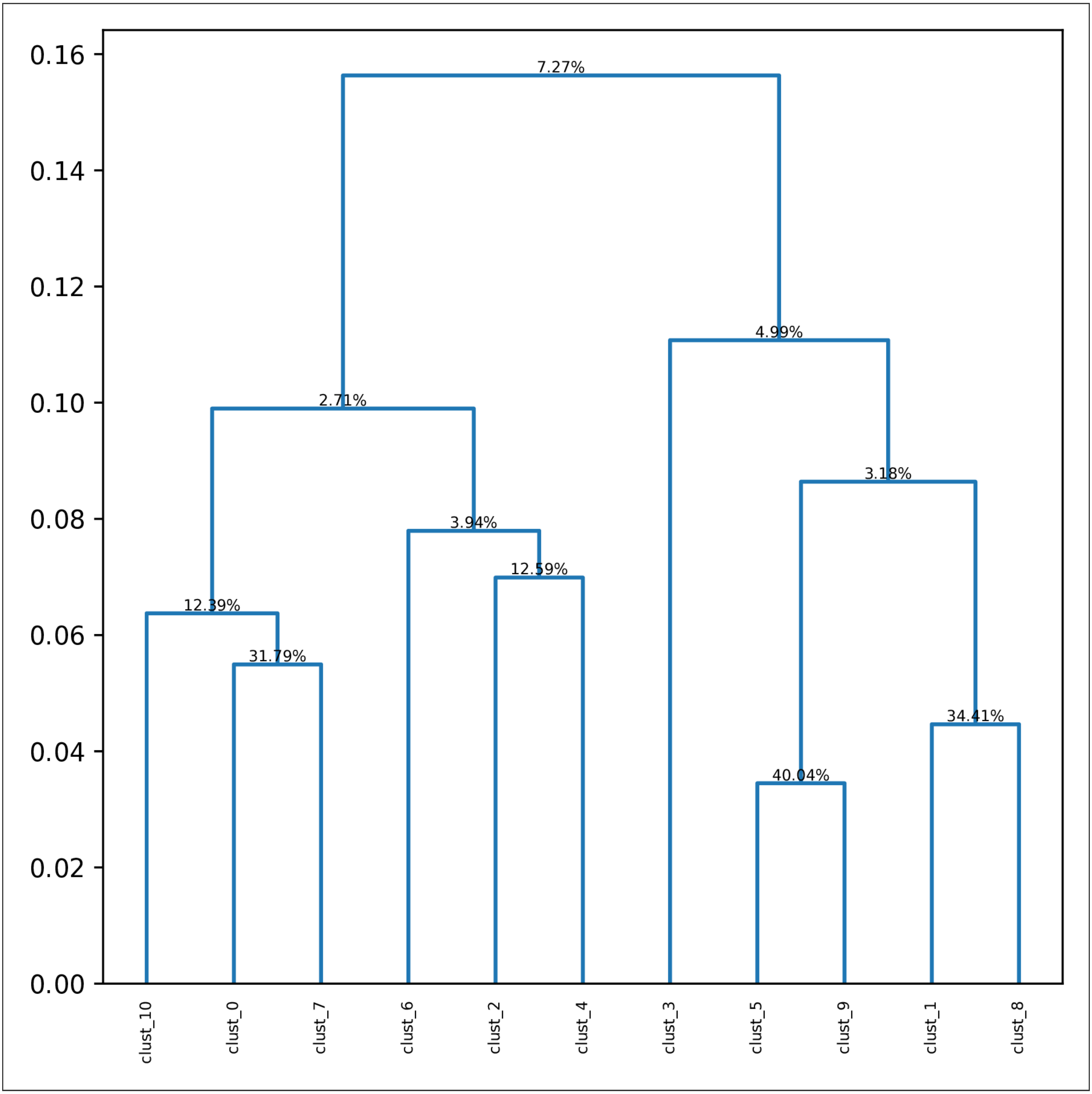

Sample: G0003 with 5 cells per cluster

Min. read depth: 10,  Mean UMIs per cluster:  43,012

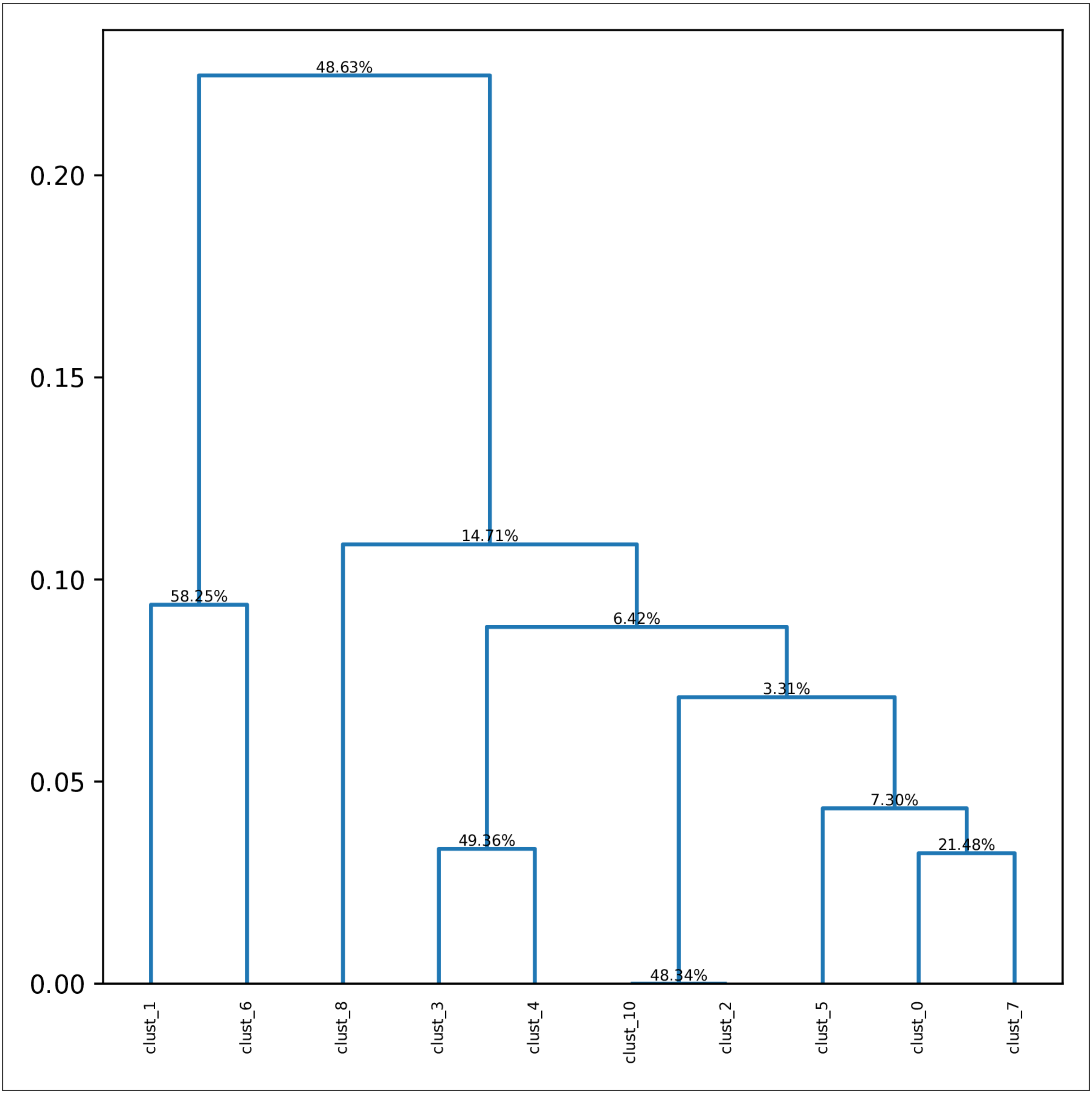

Sample: G0004 with all cells per cluster

Min. read depth: 10,  Mean UMIs per cluster:  7,809,536

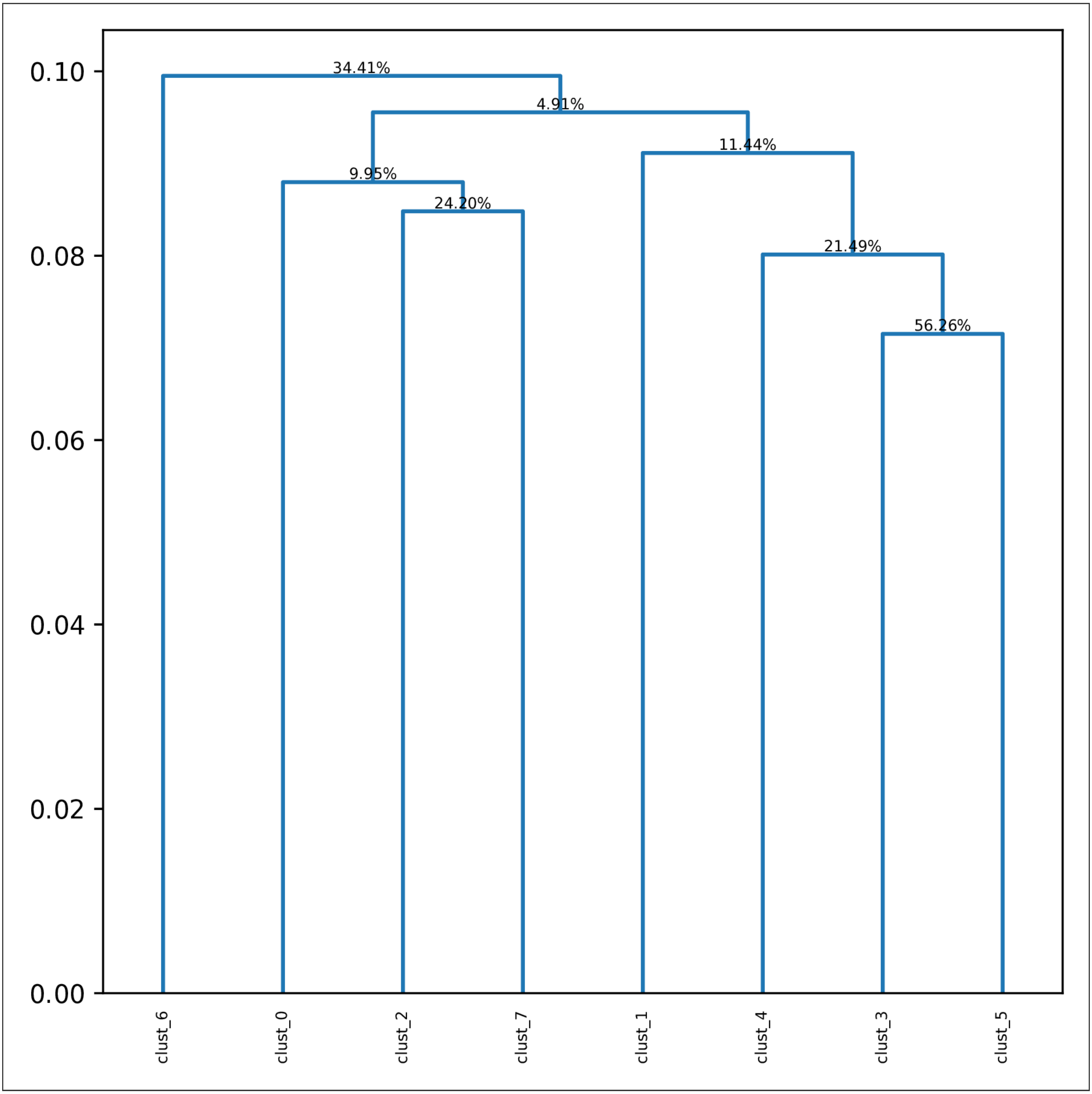

Sample: G0004 with 500 cells per cluster
Min. read depth: 10,  Mean UMIs per cluster:  2,996,617

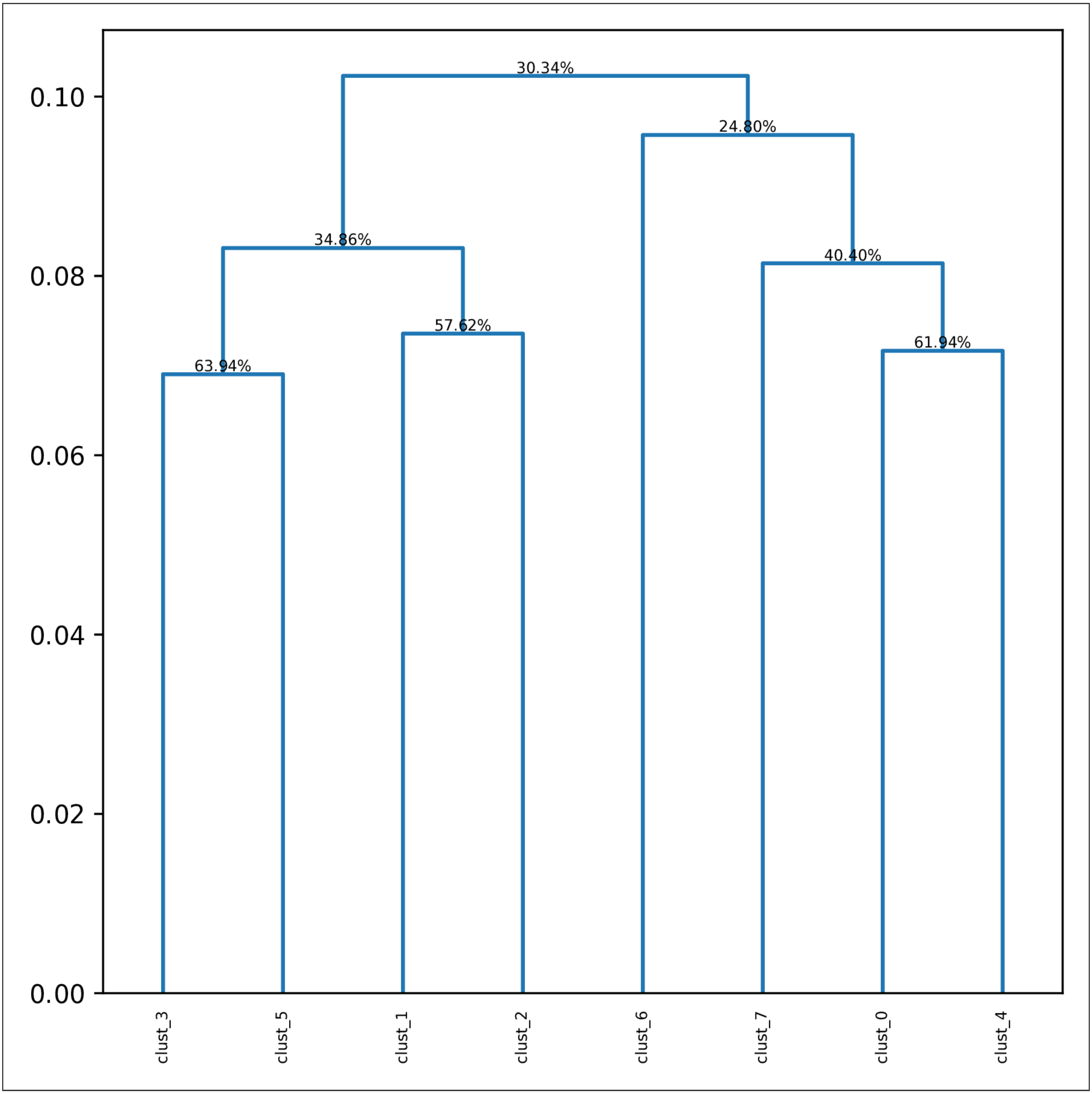

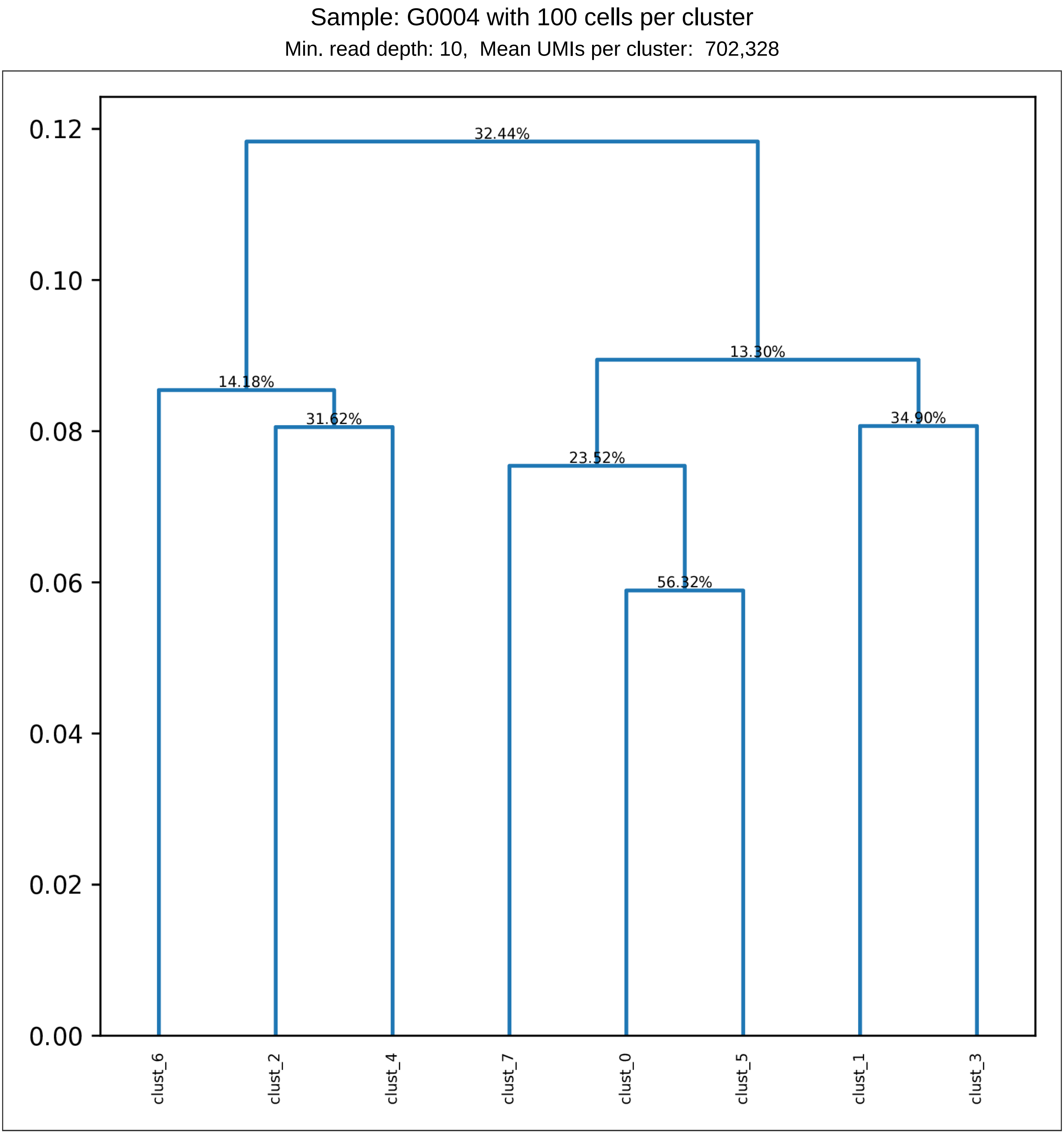

Sample: G0004 with 100 cells per cluster
Min. read depth: 10, Mean UMIs per cluster: 702,328
0.12
0.10
0.08
0.06
0.04
0.02
0.00
32.44%
14.18%
31.62%
13.30%
23.52%
56.32%
34.90%
clust_6
clust_2
clust_4
clust_7
clust_0
clust_5
clust_1
clust_3

Sample: G0004 with 50 cells per cluster

Min. read depth: 10,  Mean UMIs per cluster:  391,221

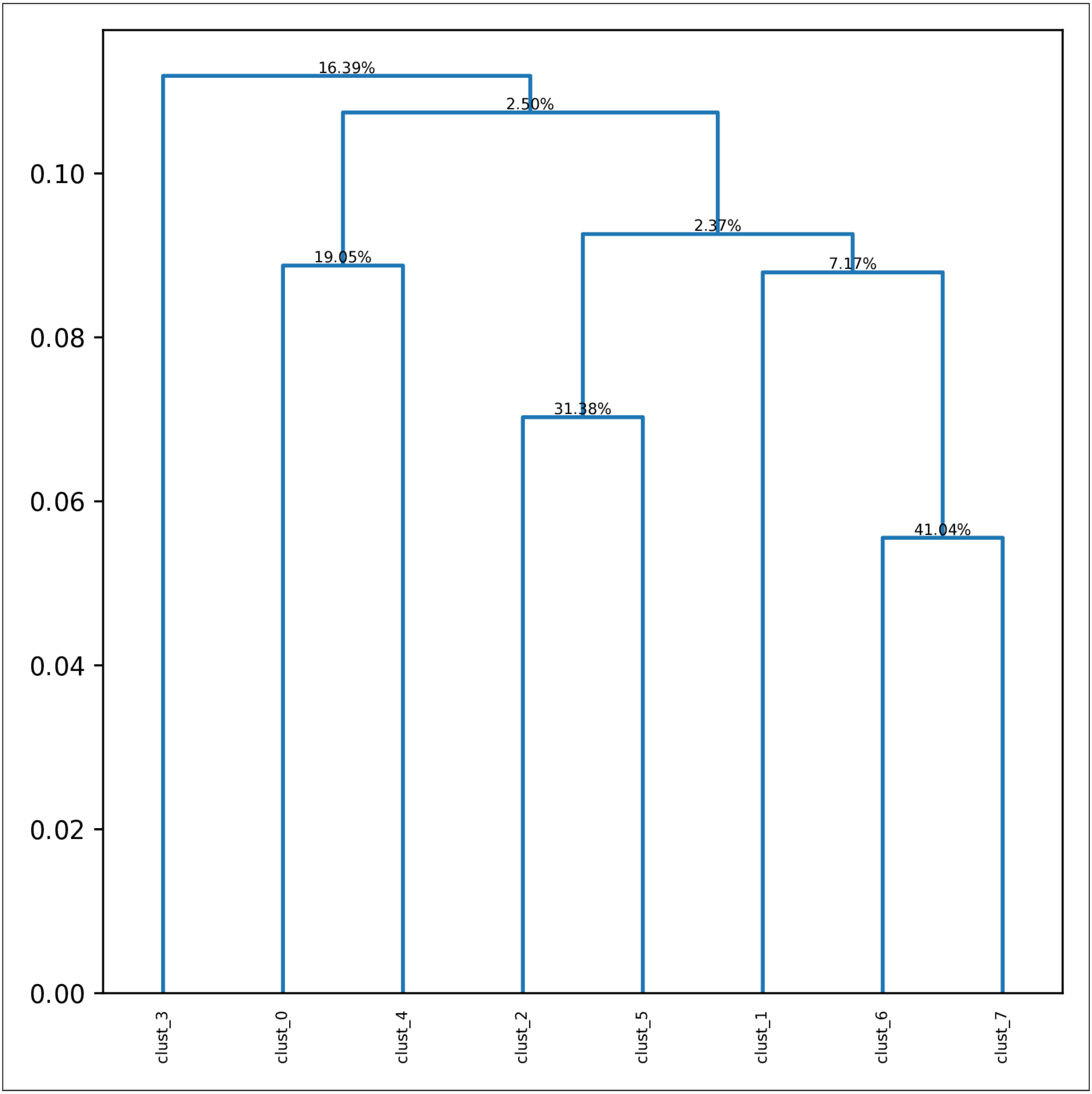

Sample: G0004 with 20 cells per cluster
Min. read depth: 10,  Mean UMIs per cluster:  158,981

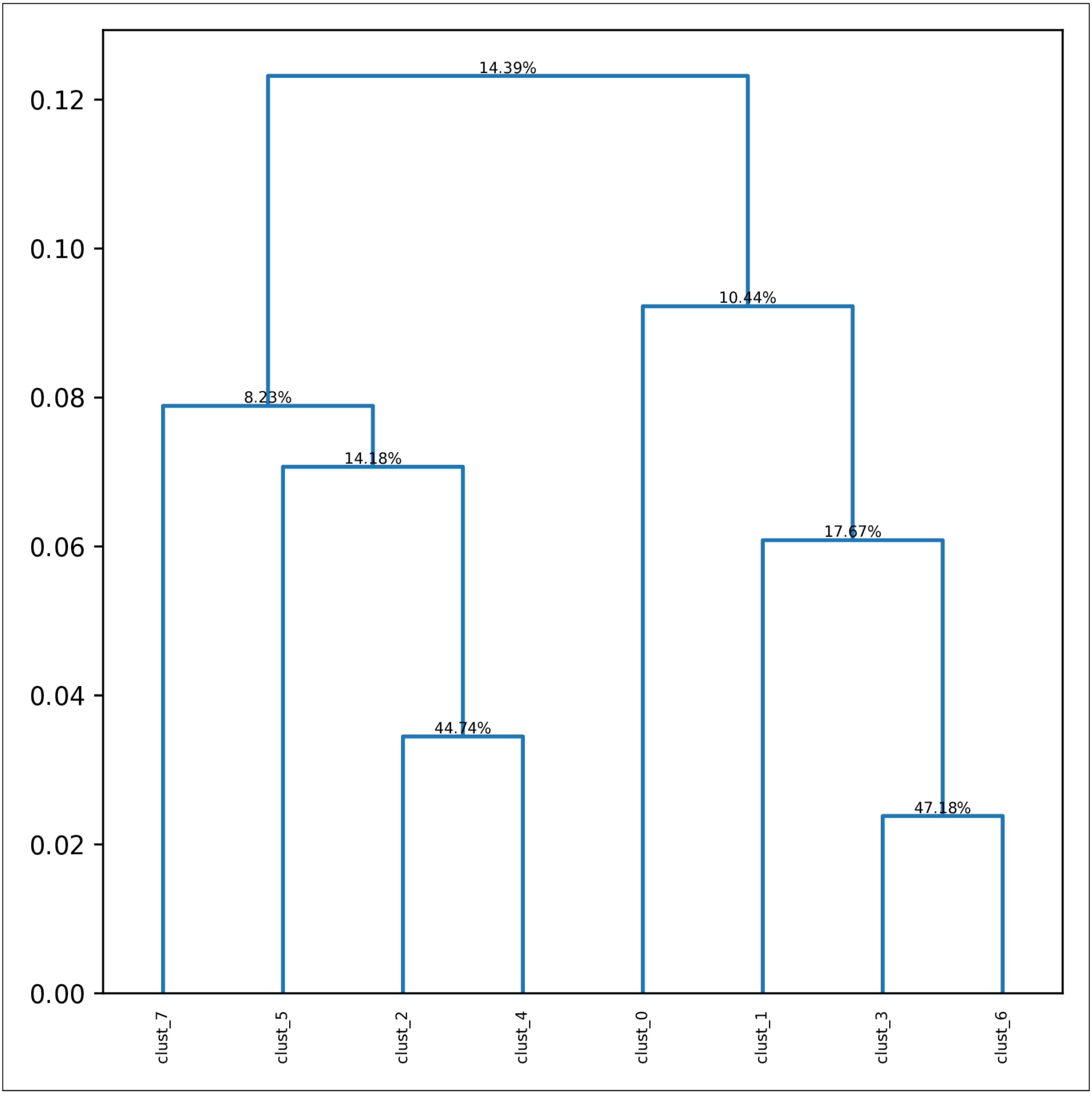

Sample: G0004 with 3 cells per cluster

Min. read depth: 10,  Mean UMIs per cluster:  17,847

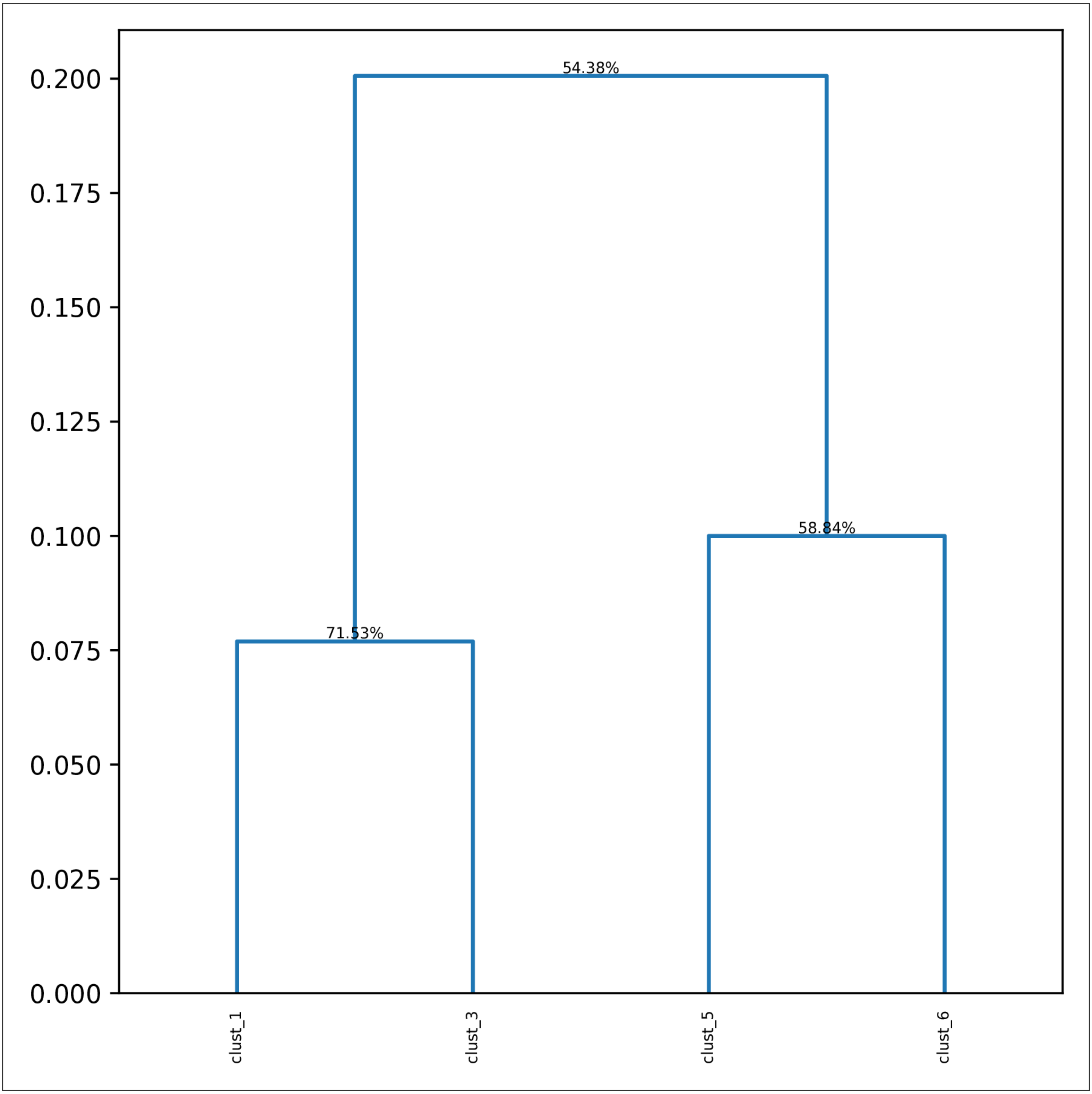

Sample: G0004 with 2 cells per cluster

Min. read depth: 10,  Mean UMIs per cluster:  15,334

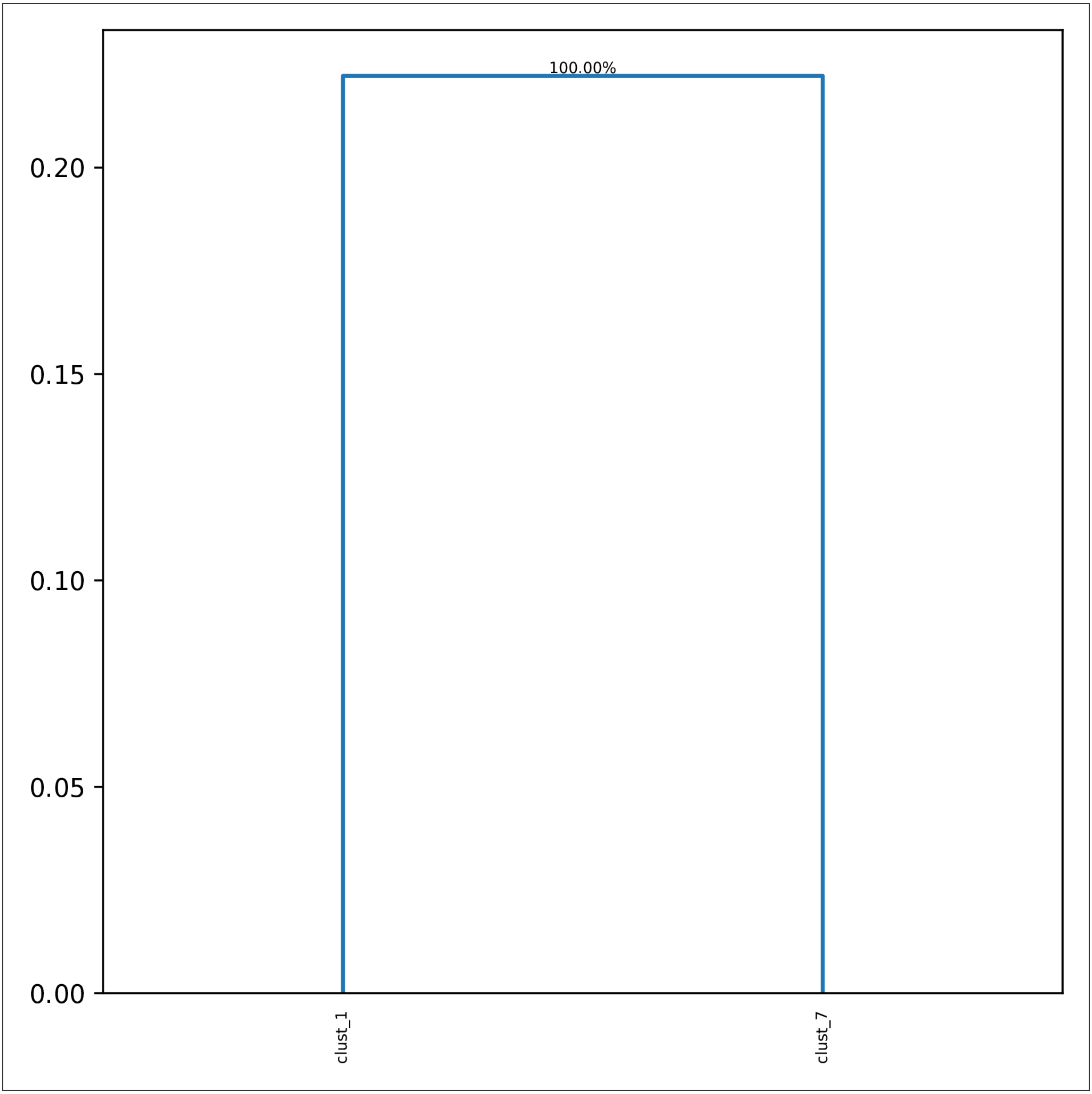

Sample: G0013 with all cells per cluster

Min. read depth: 10,  Mean UMIs per cluster:  21,059,605

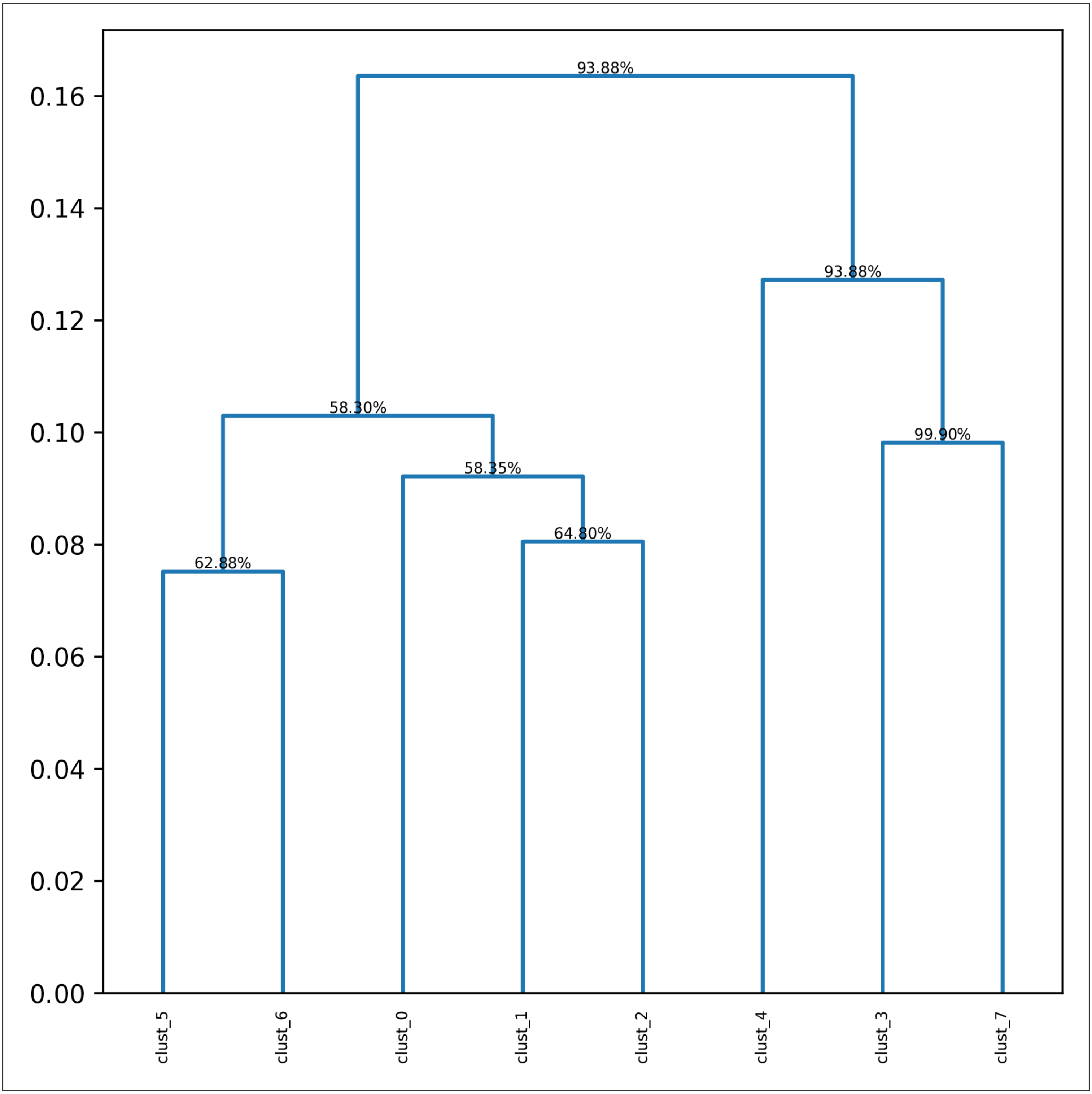

Sample: G0013 with 500 cells per cluster

Min. read depth: 10,  Mean UMIs per cluster:  11,692,211

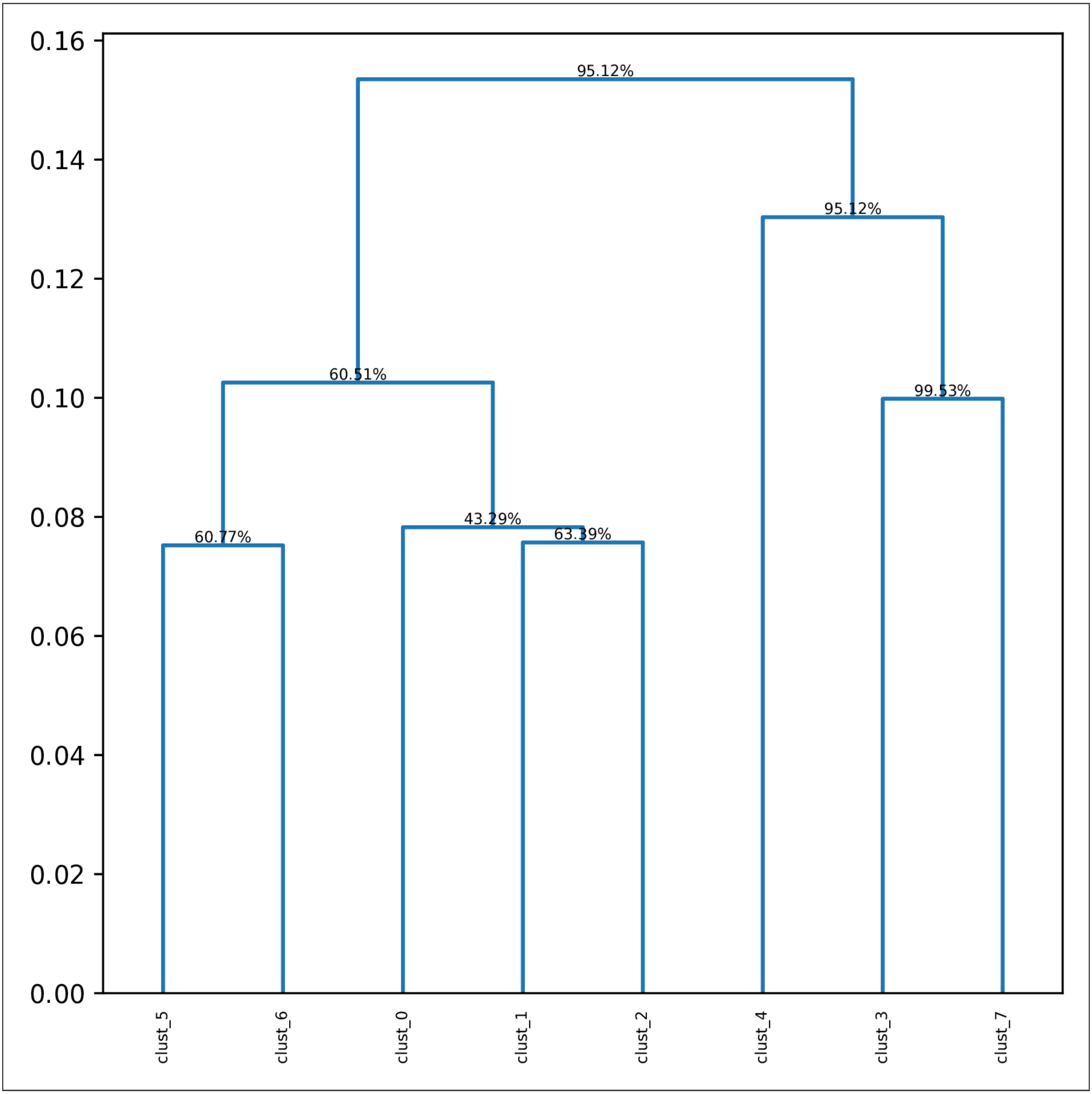

Sample: G0013 with 100 cells per cluster

Min. read depth: 10,  Mean UMIs per cluster:  2,804,627

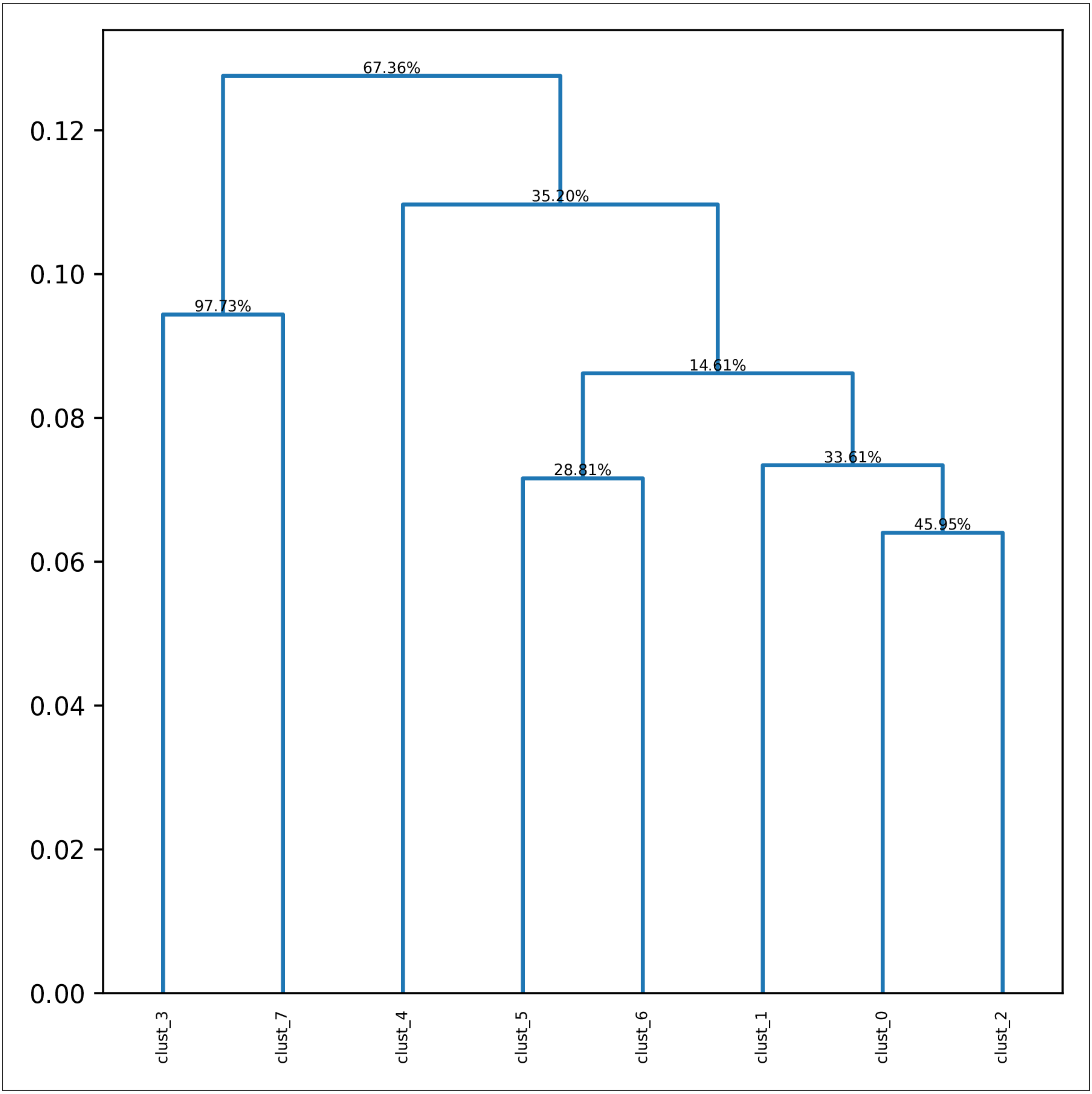

Sample: G0013 with 50 cells per cluster

Min. read depth: 10,  Mean UMIs per cluster:  1,446,237

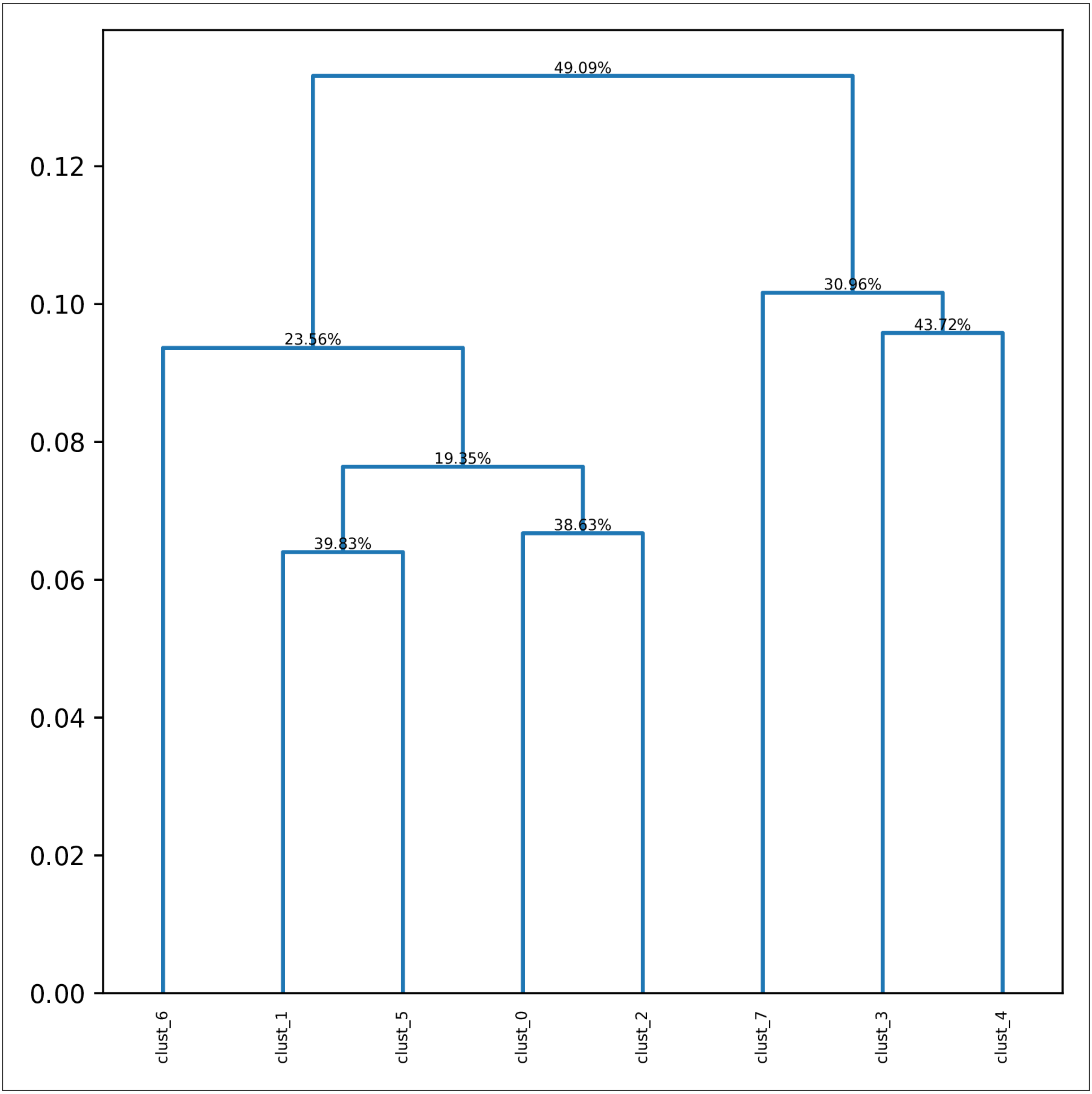

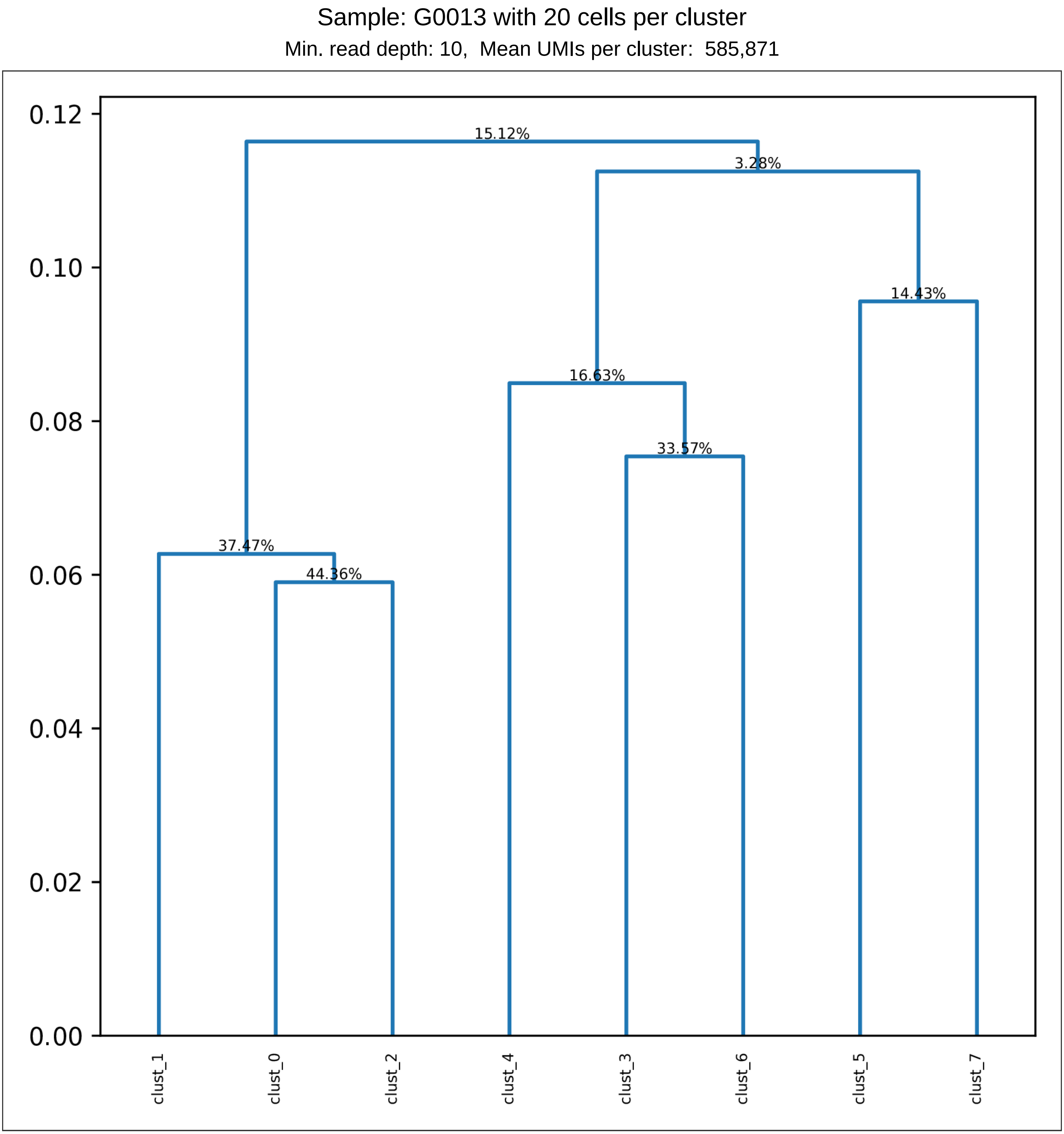
Sample: G0013 with 20 cells per cluster
Min. read depth: 10,  Mean UMIs per cluster:  585,871
0.12
0.10
0.08
0.06
0.04
0.02
0.00
15.12%
3.28%
14.43%
16.63%
33.57%
37.47%
44.36%
clust_1
clust_0
clust_2
clust_4
clust_3
clust_6
clust_5
clust_7

Sample: G0013 with 10 cells per cluster

Min. read depth: 10,  Mean UMIs per cluster:  264,623

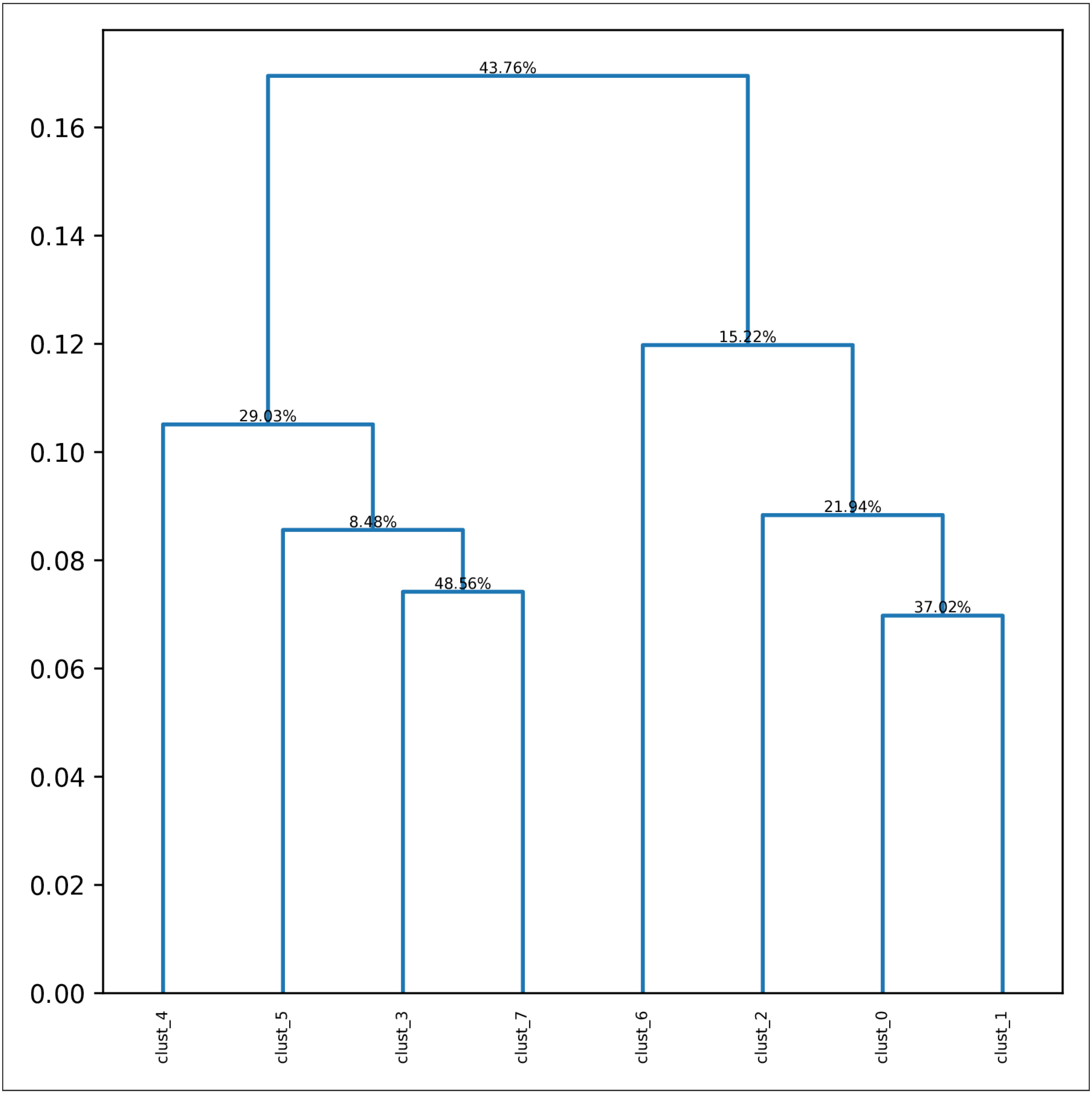

Sample: G0013 with 4 cells per cluster

Min. read depth: 10,  Mean UMIs per cluster:  103,622

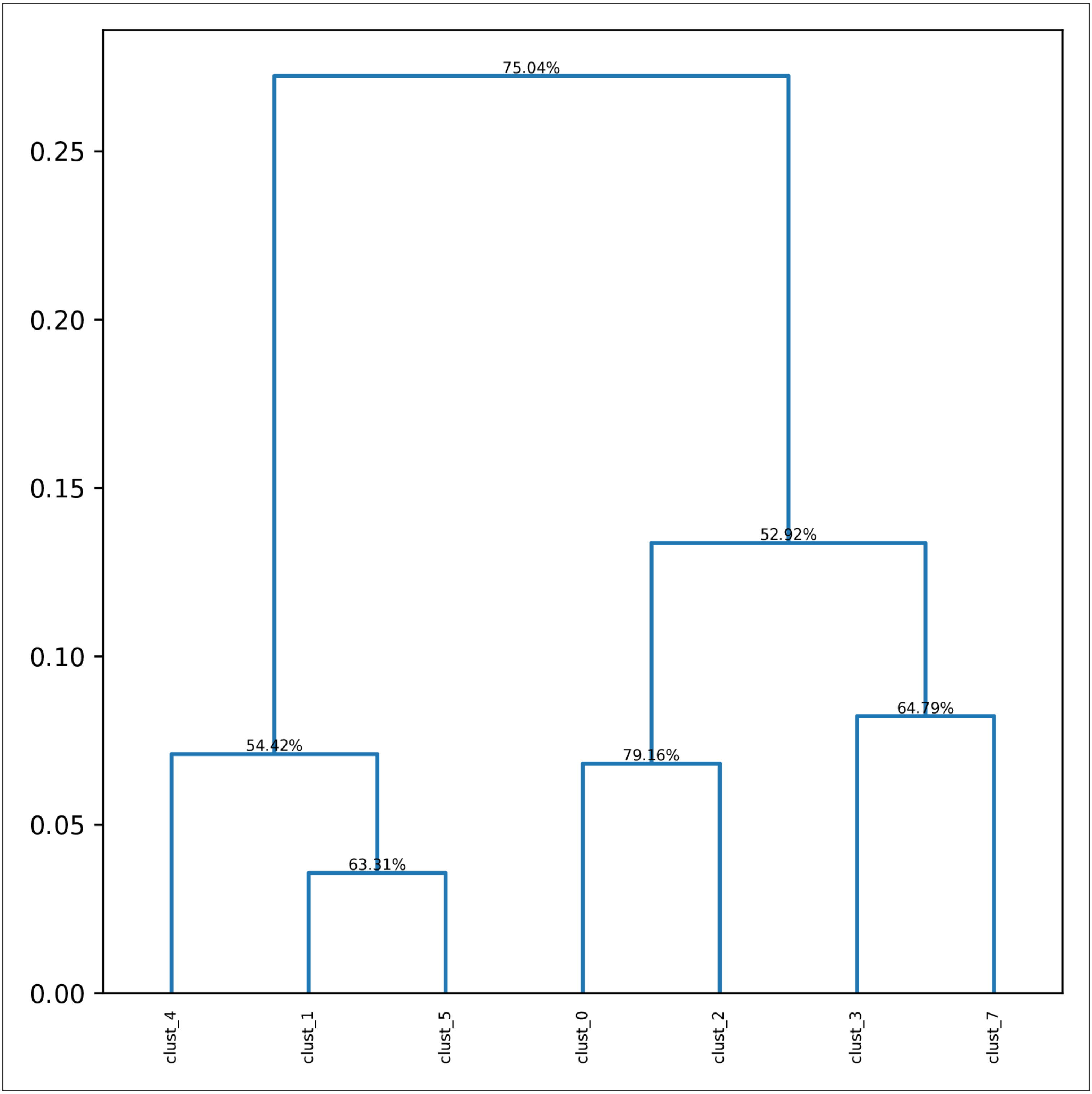

Sample: G0013 with 3 cells per cluster

Min. read depth: 10, Mean UMIs per cluster: 70,706

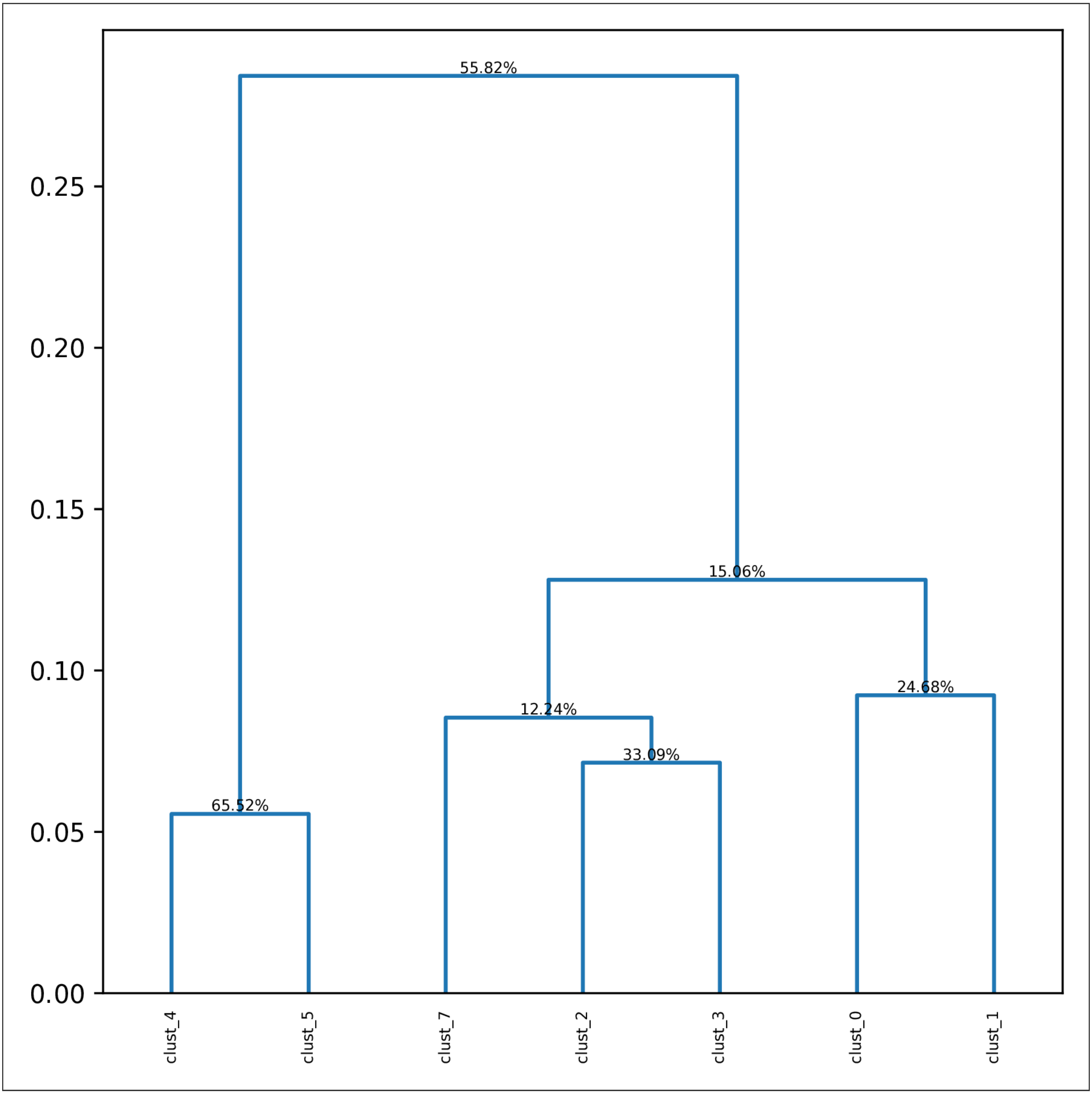

Sample: G0014 with all cells per cluster

Min. read depth: 10,  Mean UMIs per cluster:  18,801,783

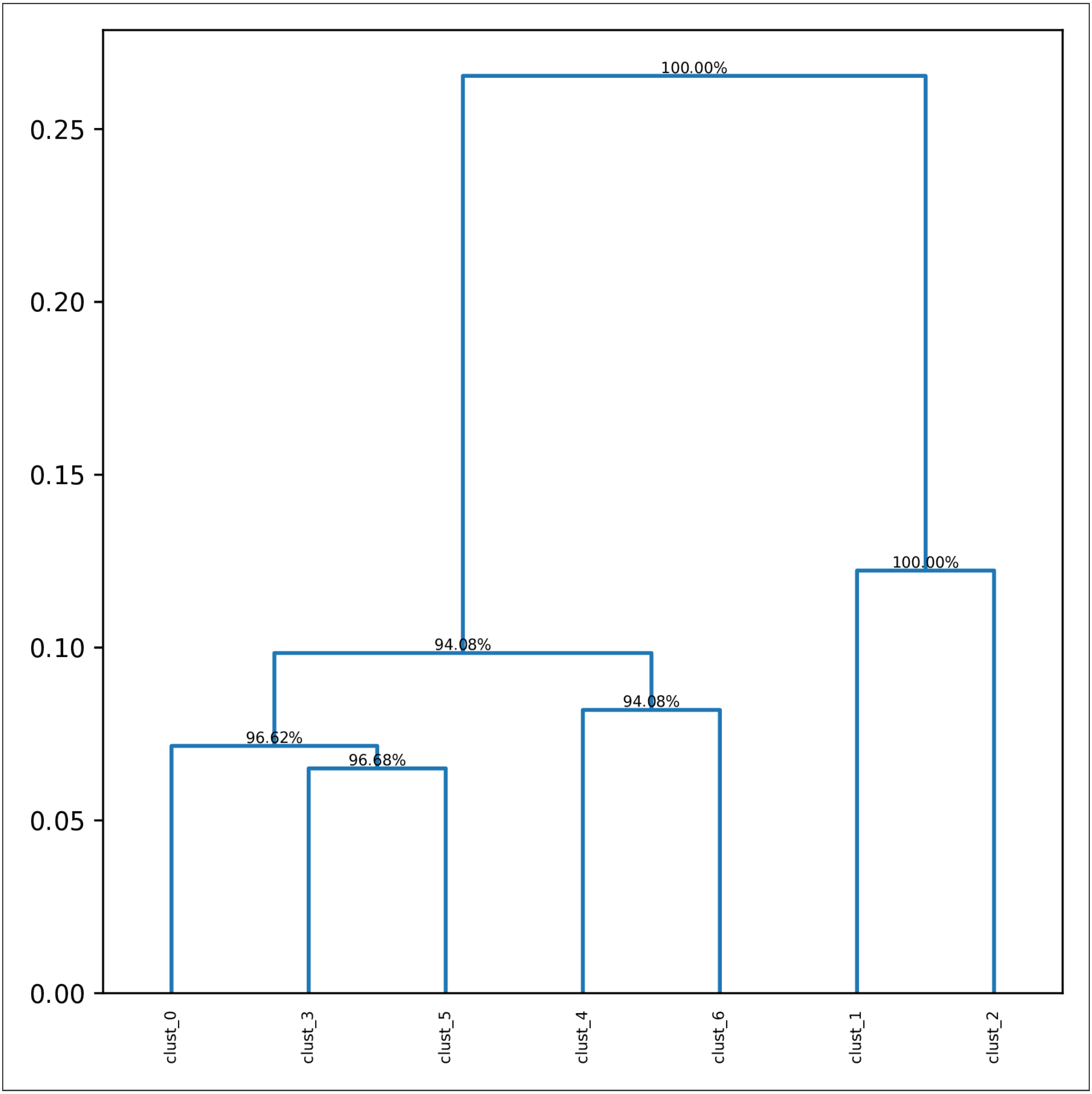

Sample: G0014 with 500 cells per cluster

Min. read depth: 10,  Mean UMIs per cluster:  8,726,497

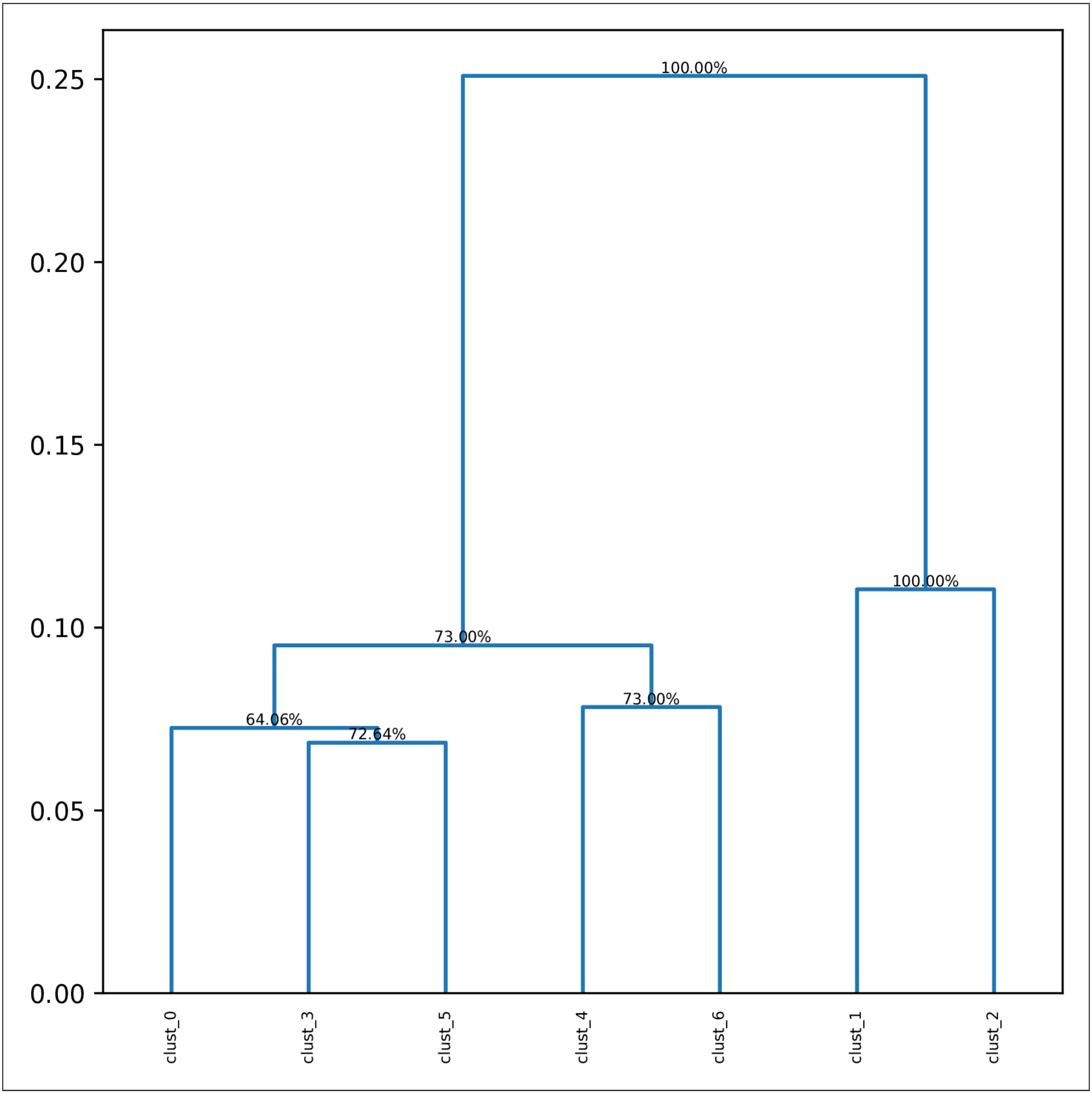

Sample: G0014 with 100 cells per cluster
Min. read depth: 10,  Mean UMIs per cluster:  1,962,408

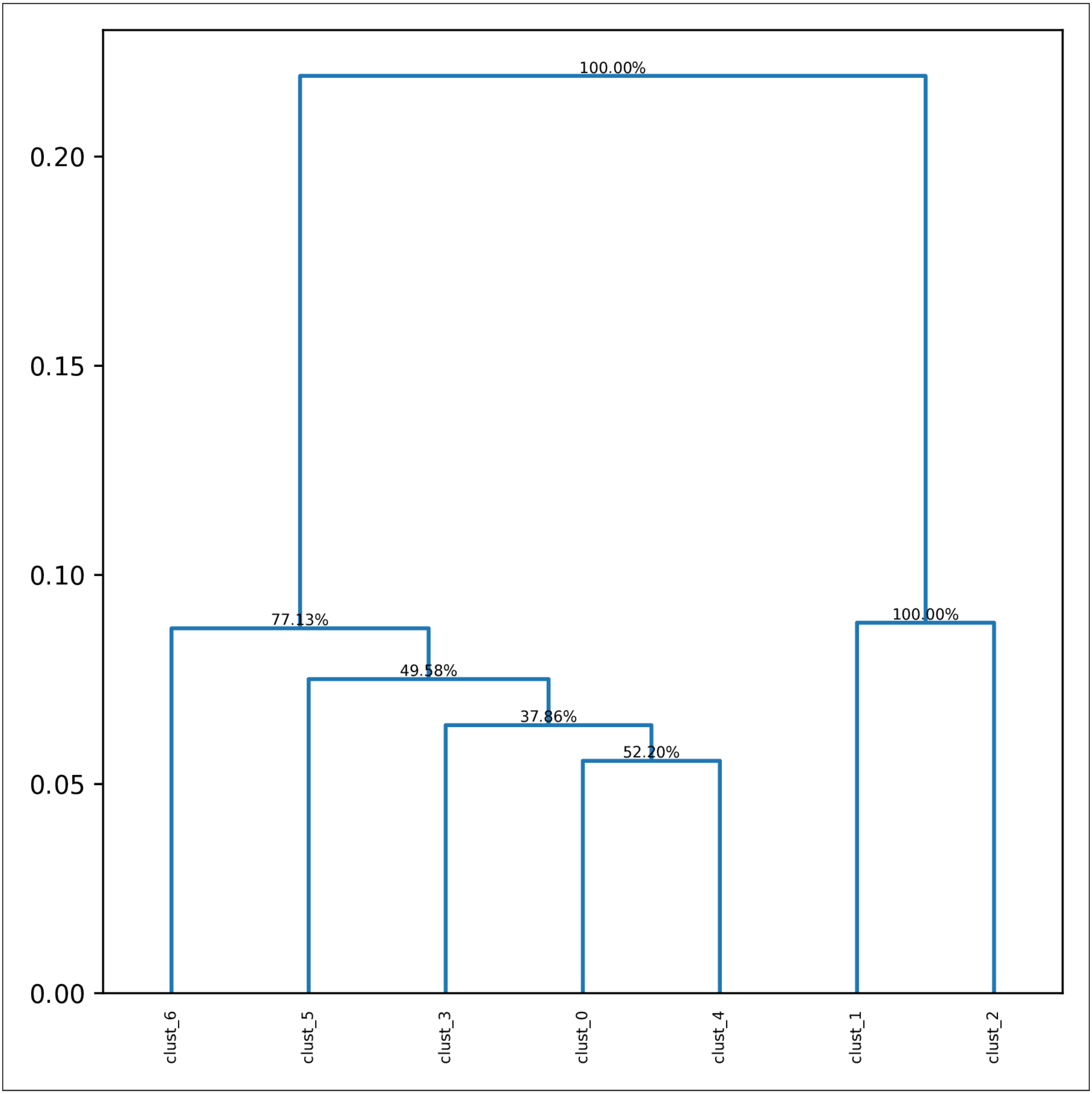

Sample: G0014 with 50 cells per cluster

Min. read depth: 10,  Mean UMIs per cluster:  941,560

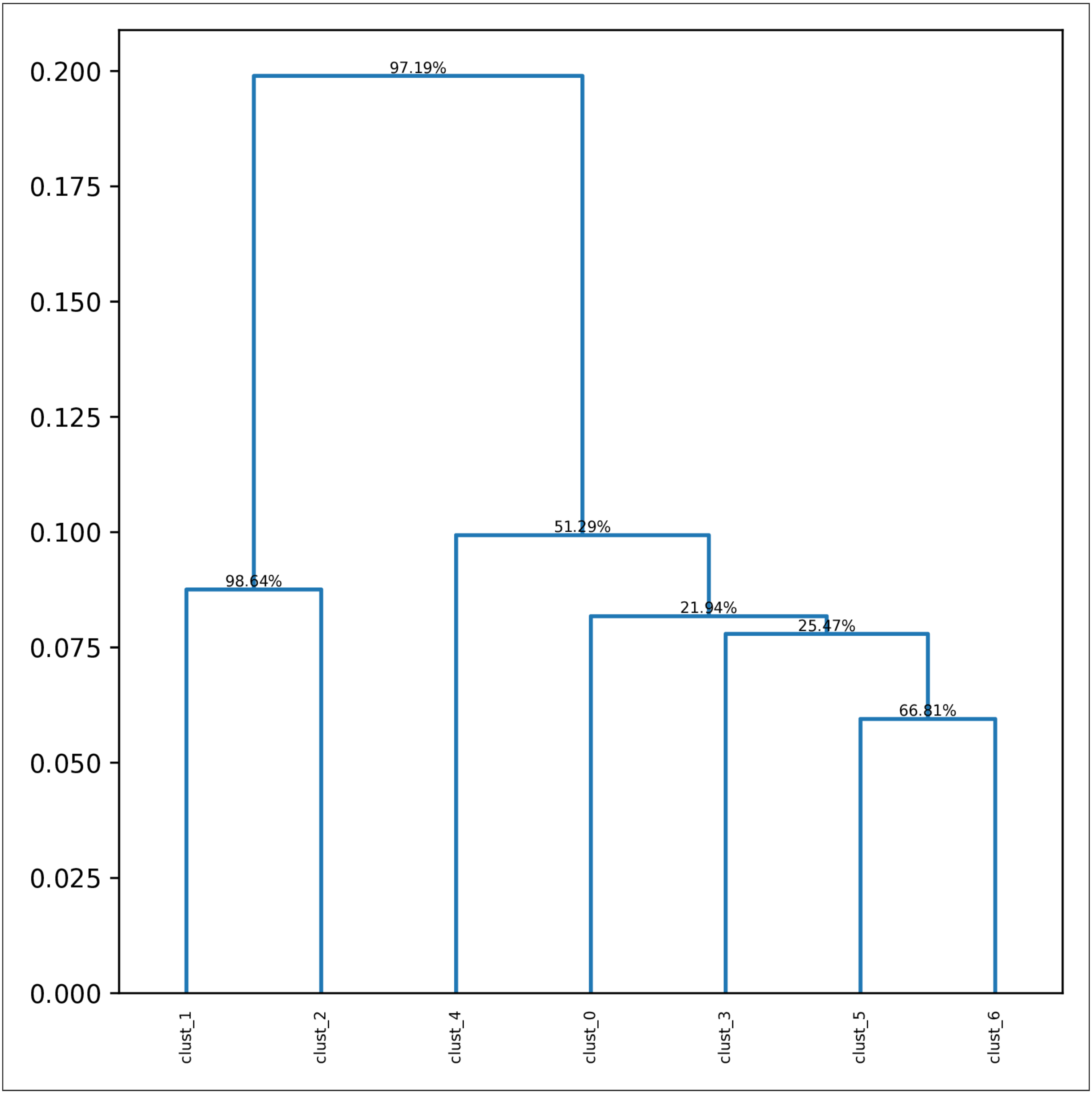

Sample: G0014 with 20 cells per cluster

Min. read depth: 10,  Mean UMIs per cluster:  378,204

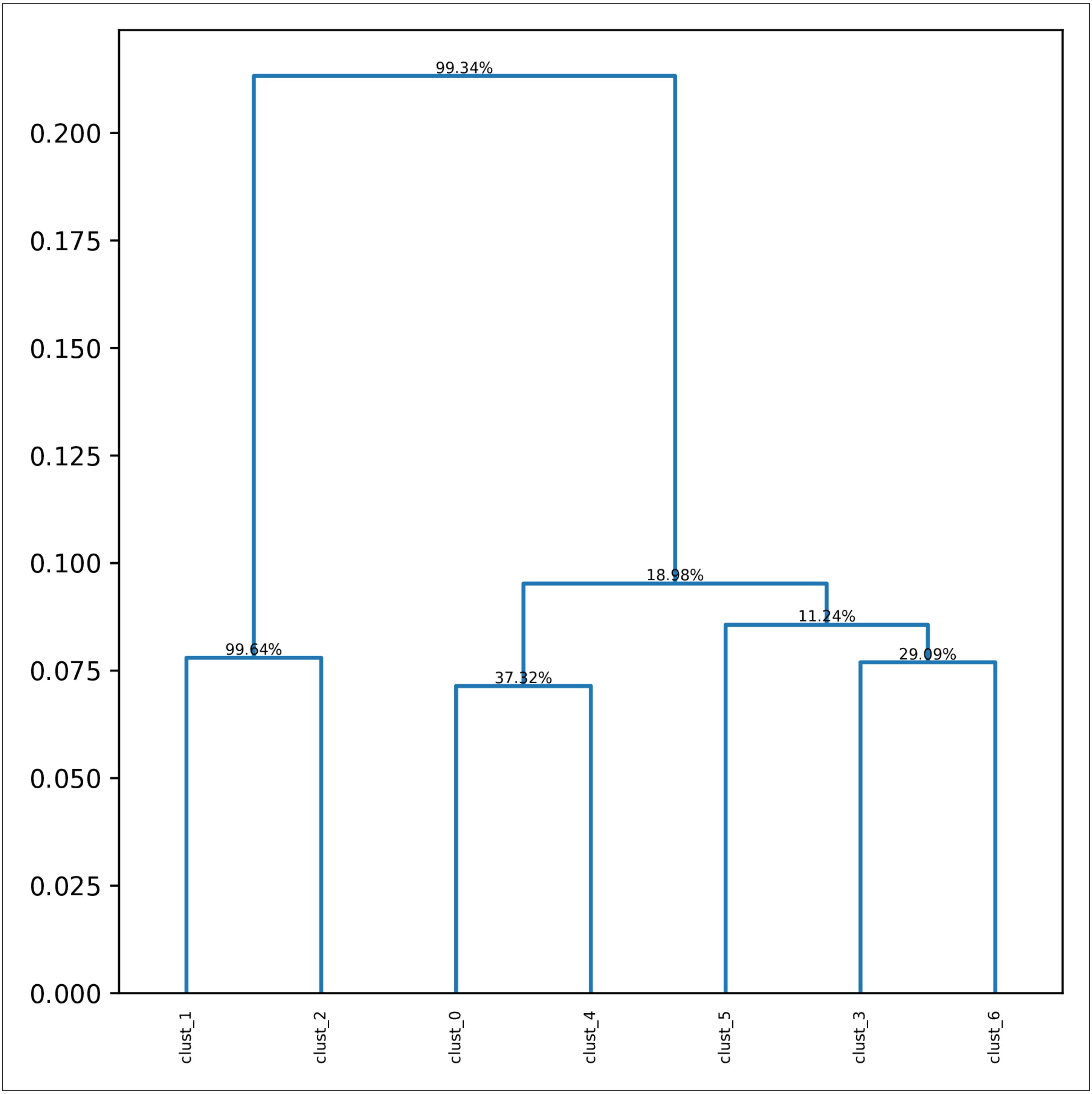

Sample: G0014 with 10 cells per cluster

Min. read depth: 10,  Mean UMIs per cluster:  209,148

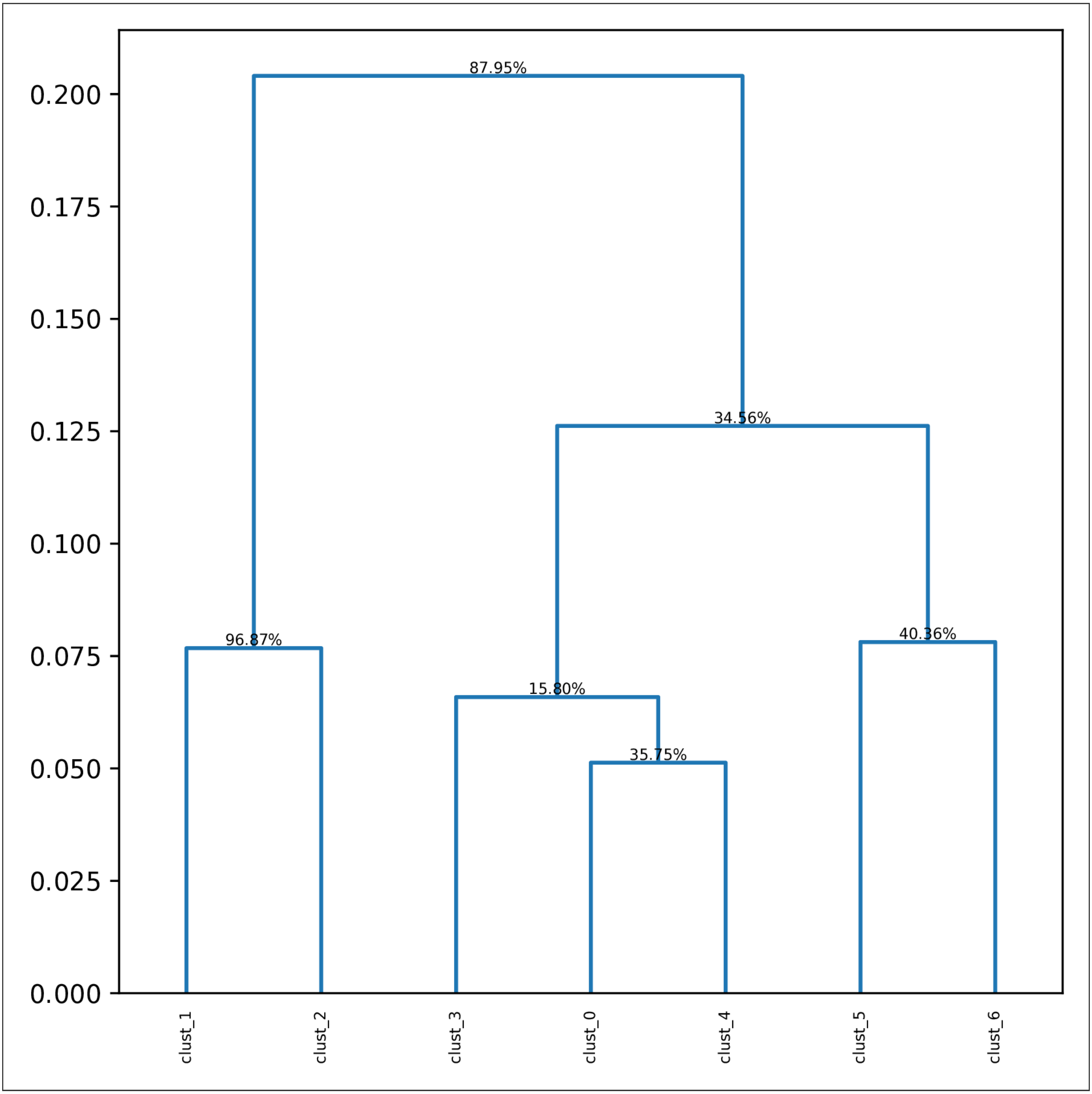

Sample: G0014 with 5 cells per cluster

Min. read depth: 10,  Mean UMIs per cluster:  117,314

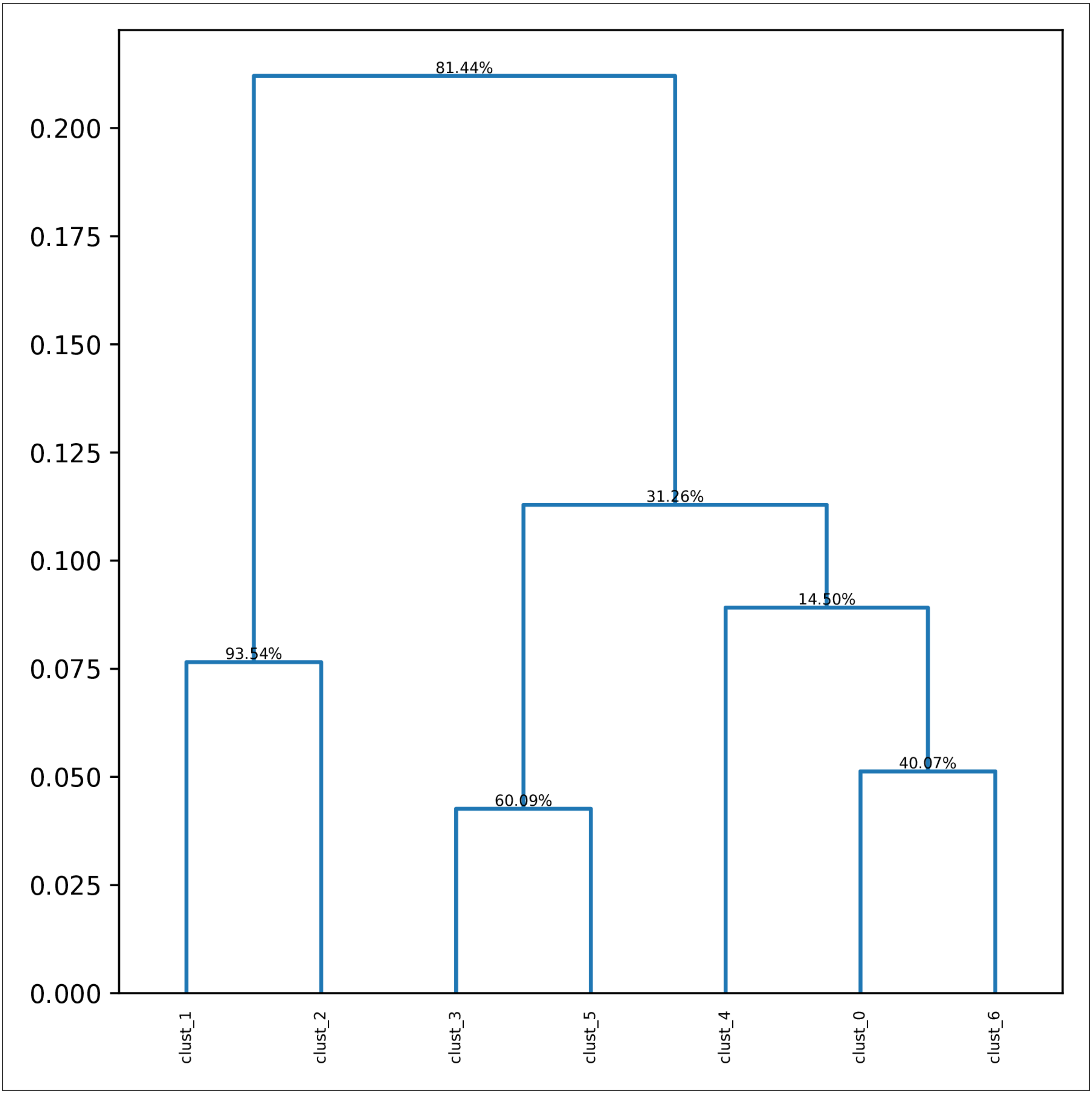

Sample: G0014 with 4 cells per cluster
Min. read depth: 10,  Mean UMIs per cluster:  88,994

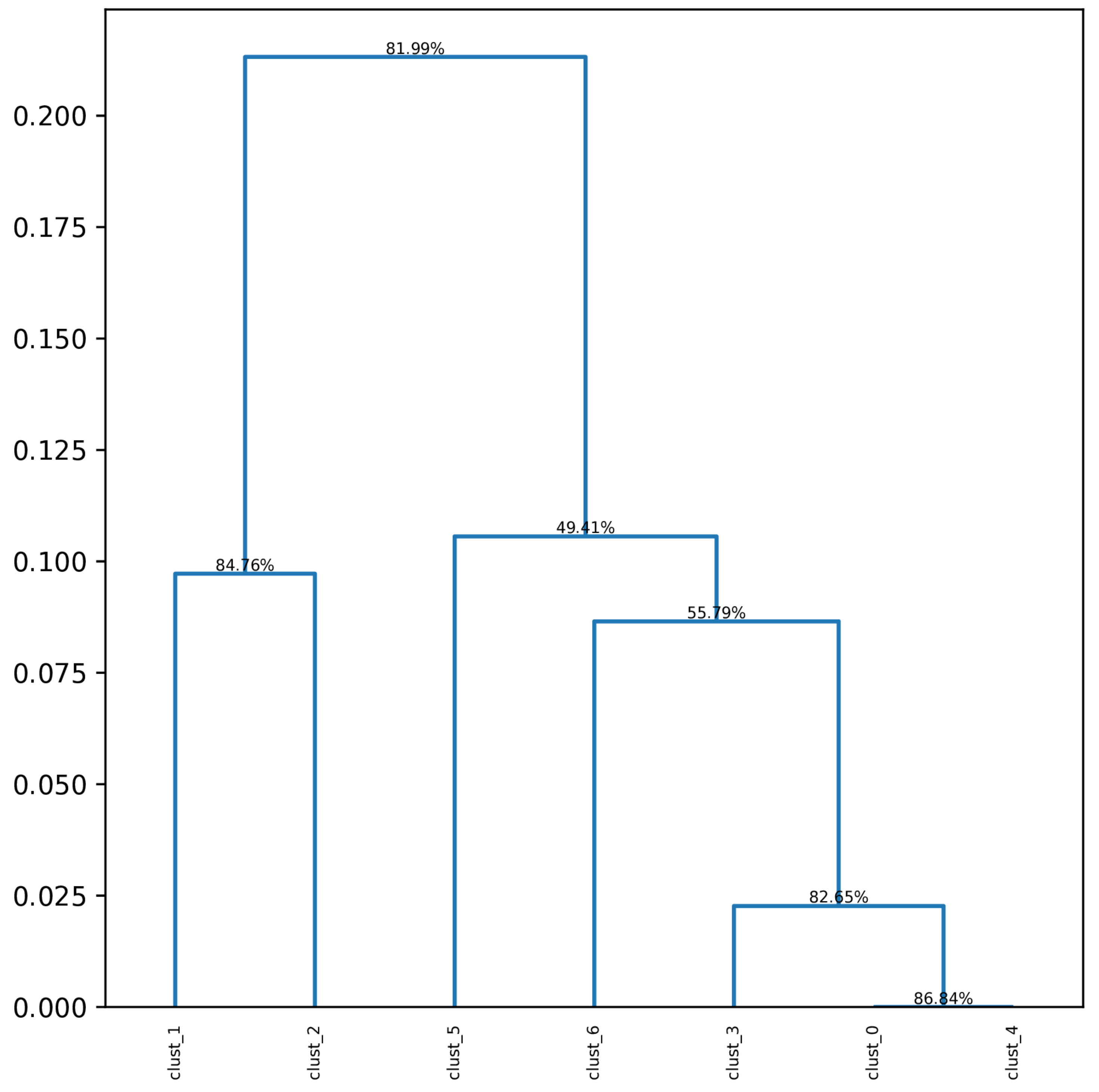

Sample: G0014 with 2 cells per cluster

Min. read depth: 10,  Mean UMIs per cluster:  48,349

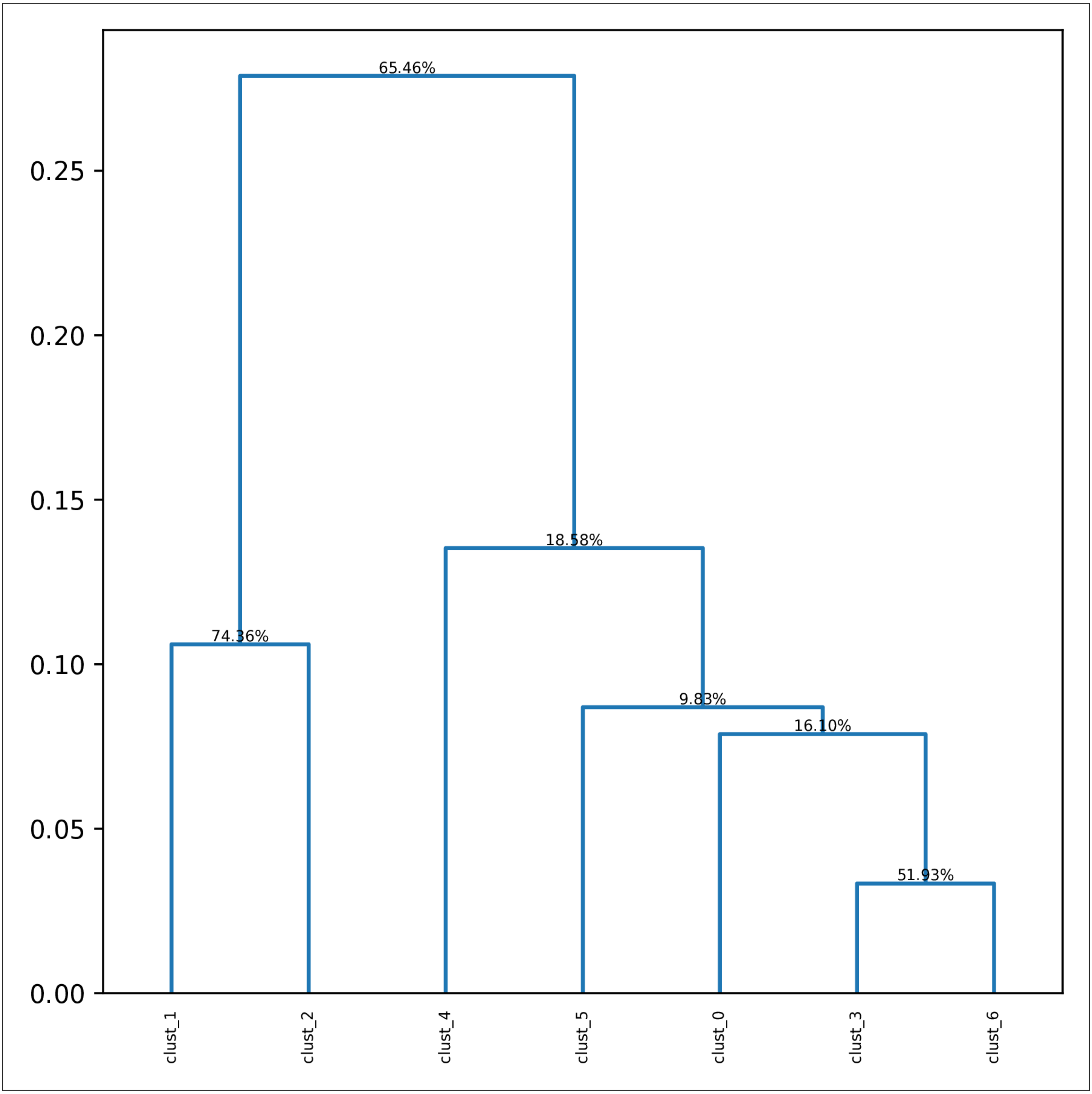

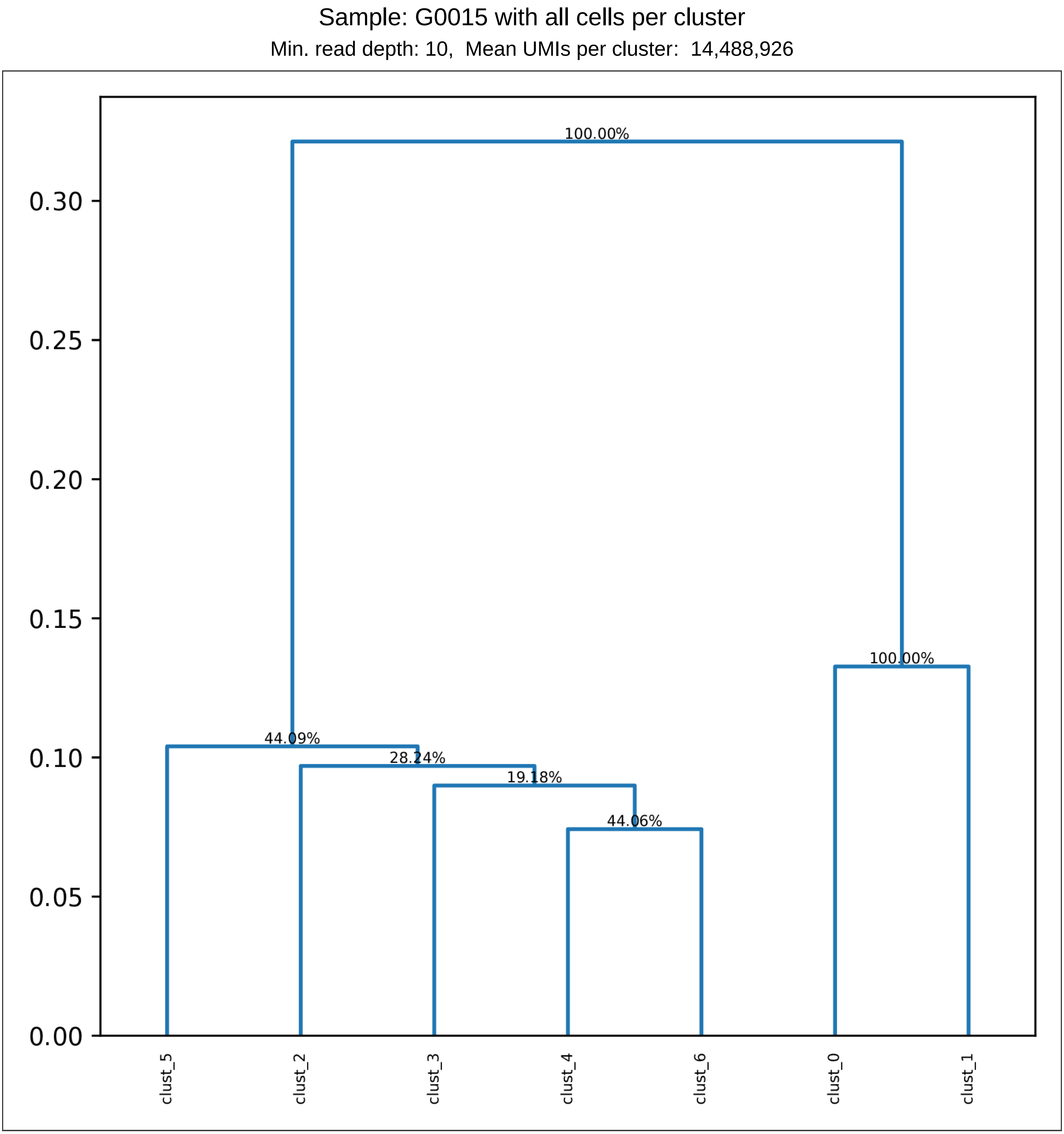

Sample: G0015 with all cells per cluster
Min. read depth: 10, Mean UMIs per cluster: 14,488,926
100.00%
100.00%
44.09%
28.24%
19.18%
44.06%
0.00
0.05
0.10
0.15
0.20
0.25
0.30
clust_5
clust_2
clust_3
clust_4
clust_6
clust_0
clust_1

Sample: G0015 with 500 cells per cluster

Min. read depth: 10,  Mean UMIs per cluster:  4,515,387

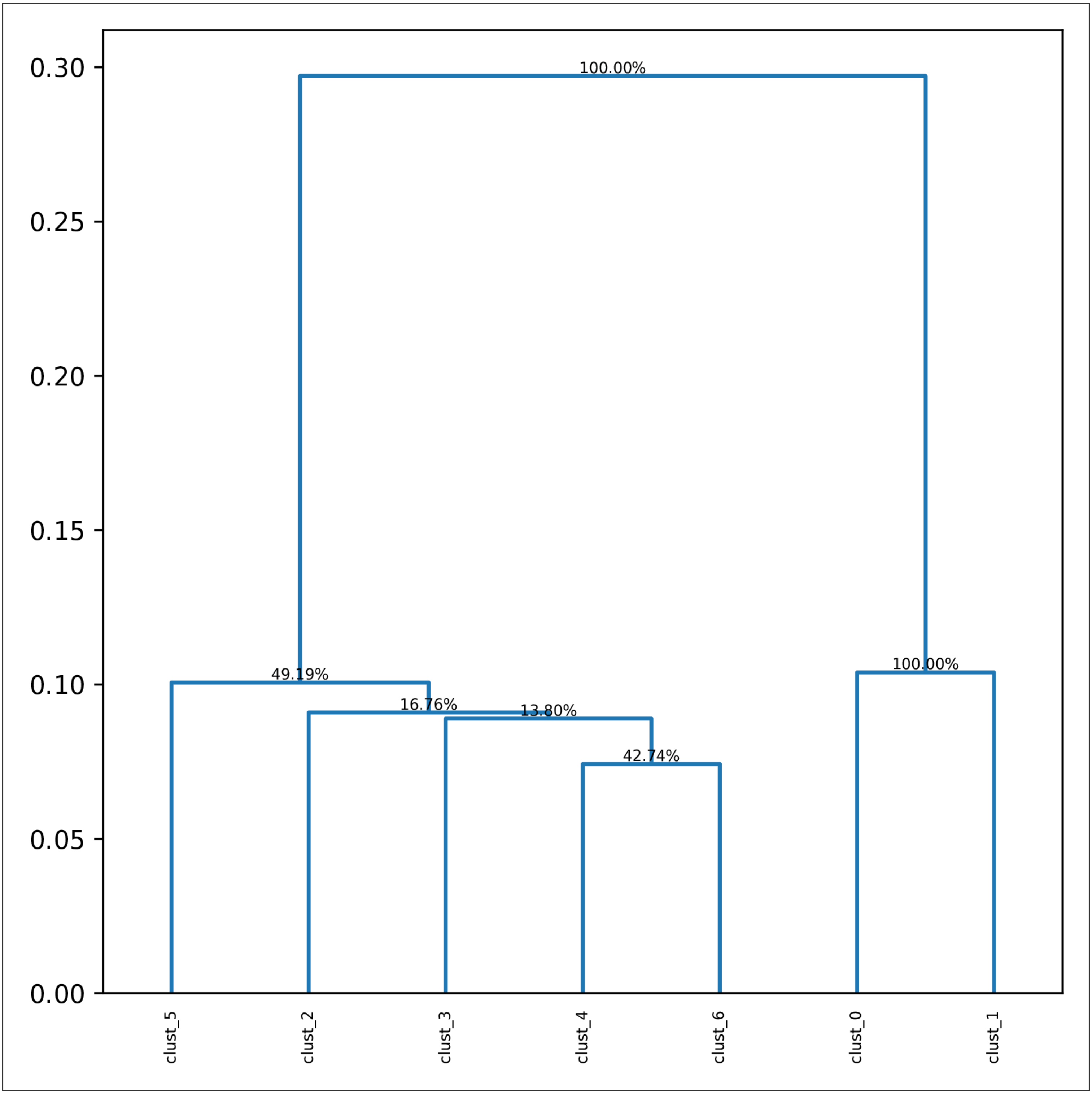

Sample: G0015 with 100 cells per cluster

Min. read depth: 10,  Mean UMIs per cluster:  988,963

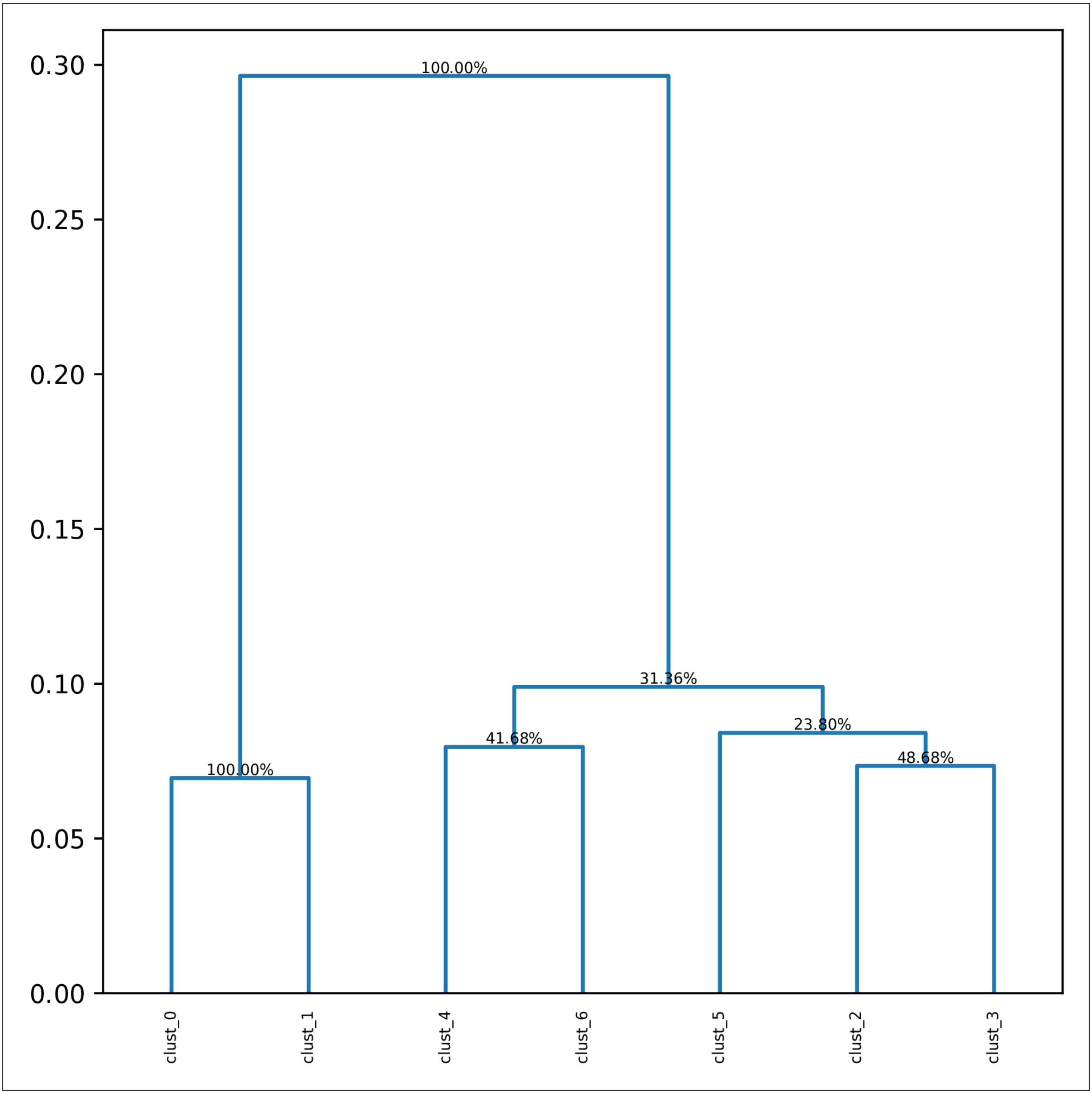

Sample: G0015 with 50 cells per cluster
Min. read depth: 10,  Mean UMIs per cluster:  521,678

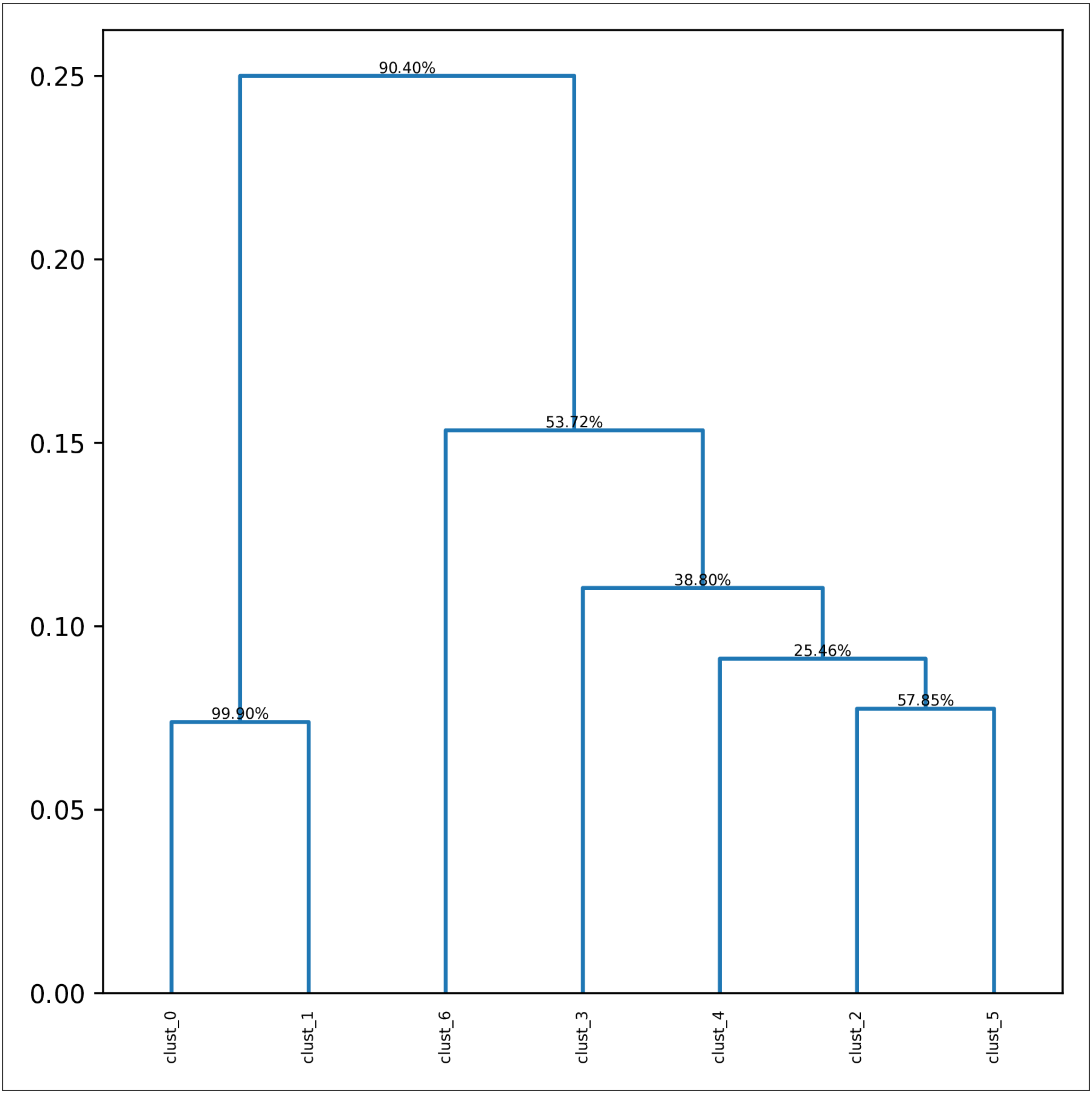

Sample: G0015 with 20 cells per cluster

Min. read depth: 10,  Mean UMIs per cluster:  210,547

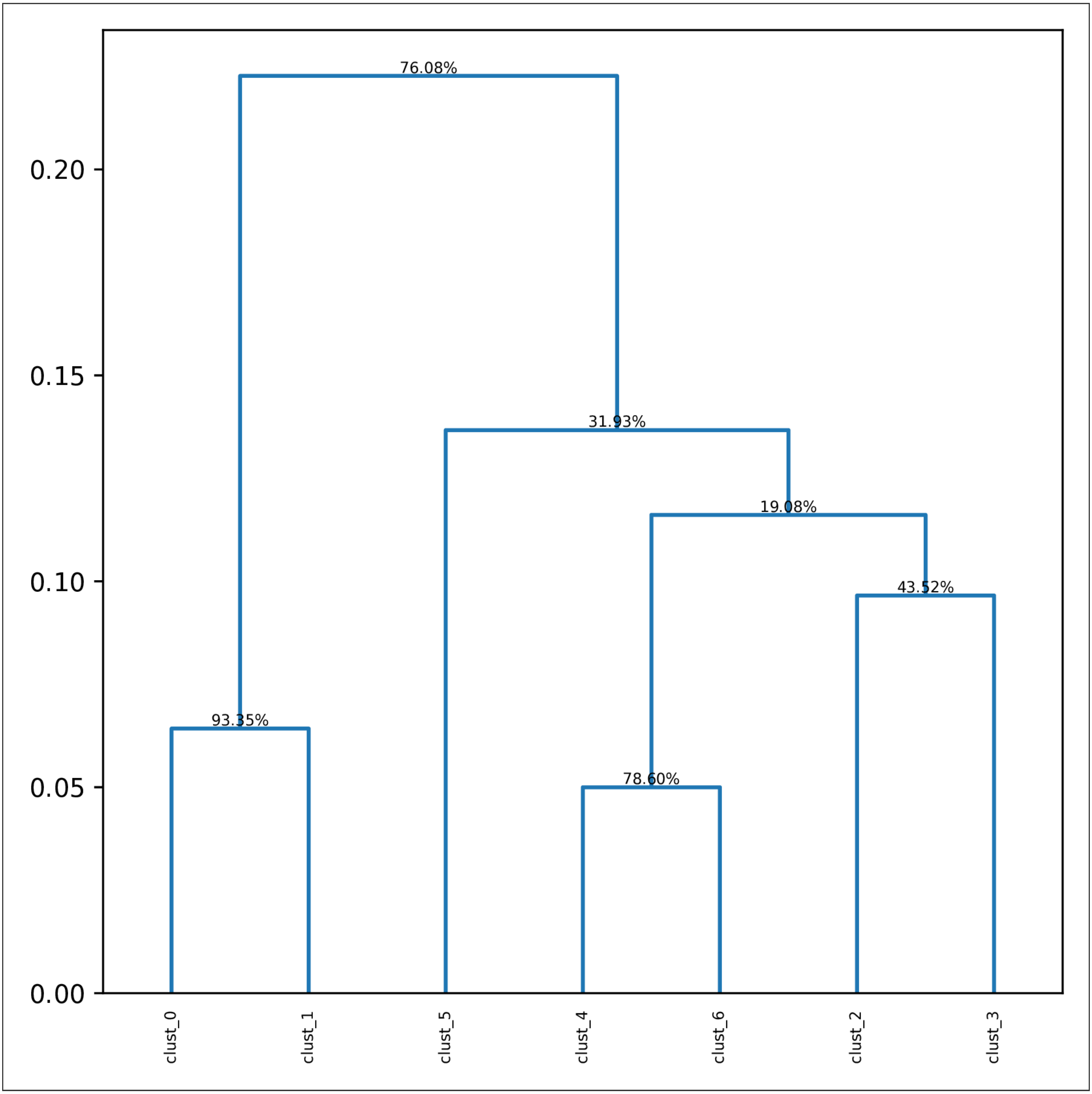

Sample: G0015 with 10 cells per cluster

Min. read depth: 10,  Mean UMIs per cluster:  97,418

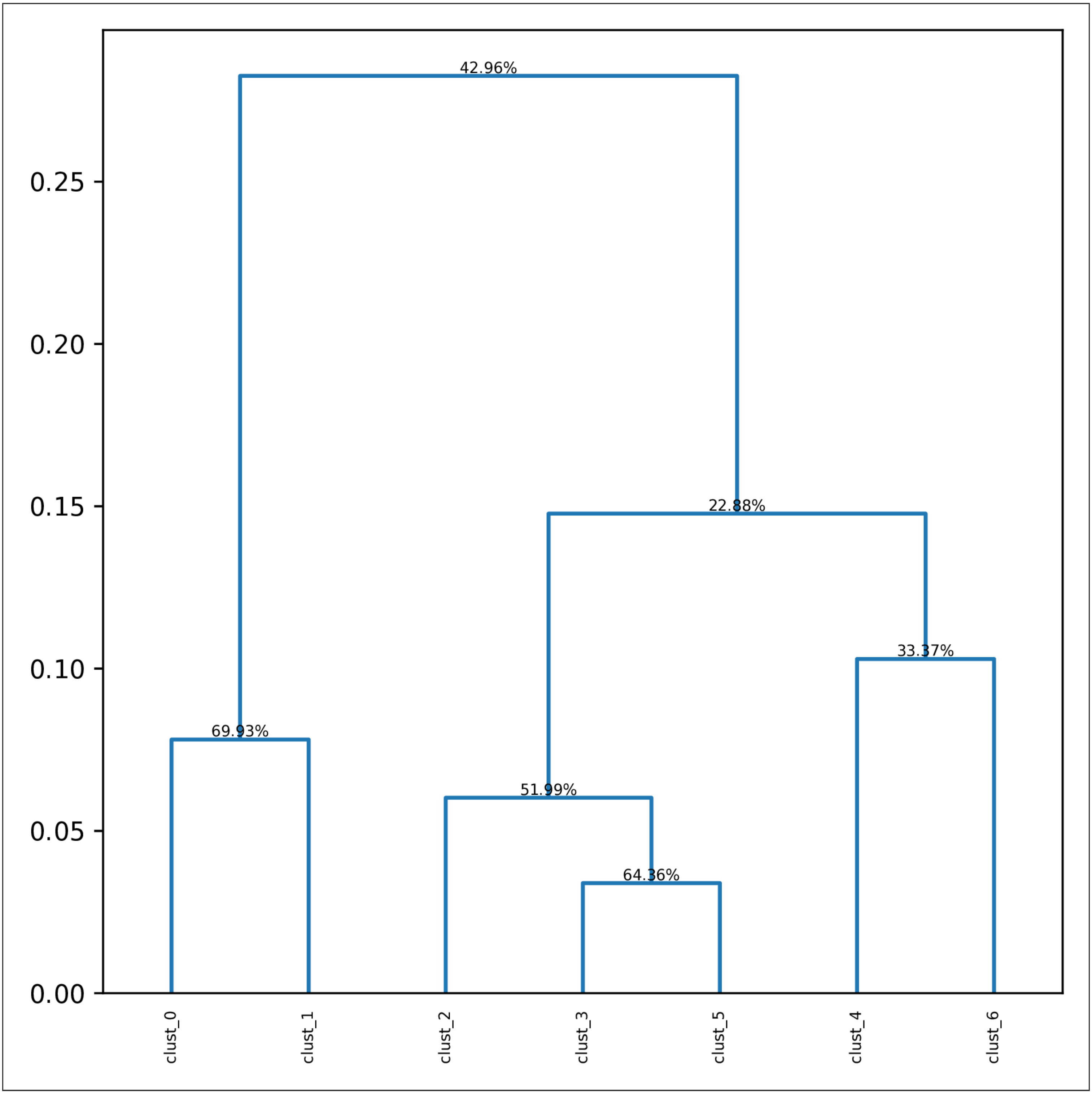

Sample: G0015 with 4 cells per cluster

Min. read depth: 10,  Mean UMIs per cluster:  34,905

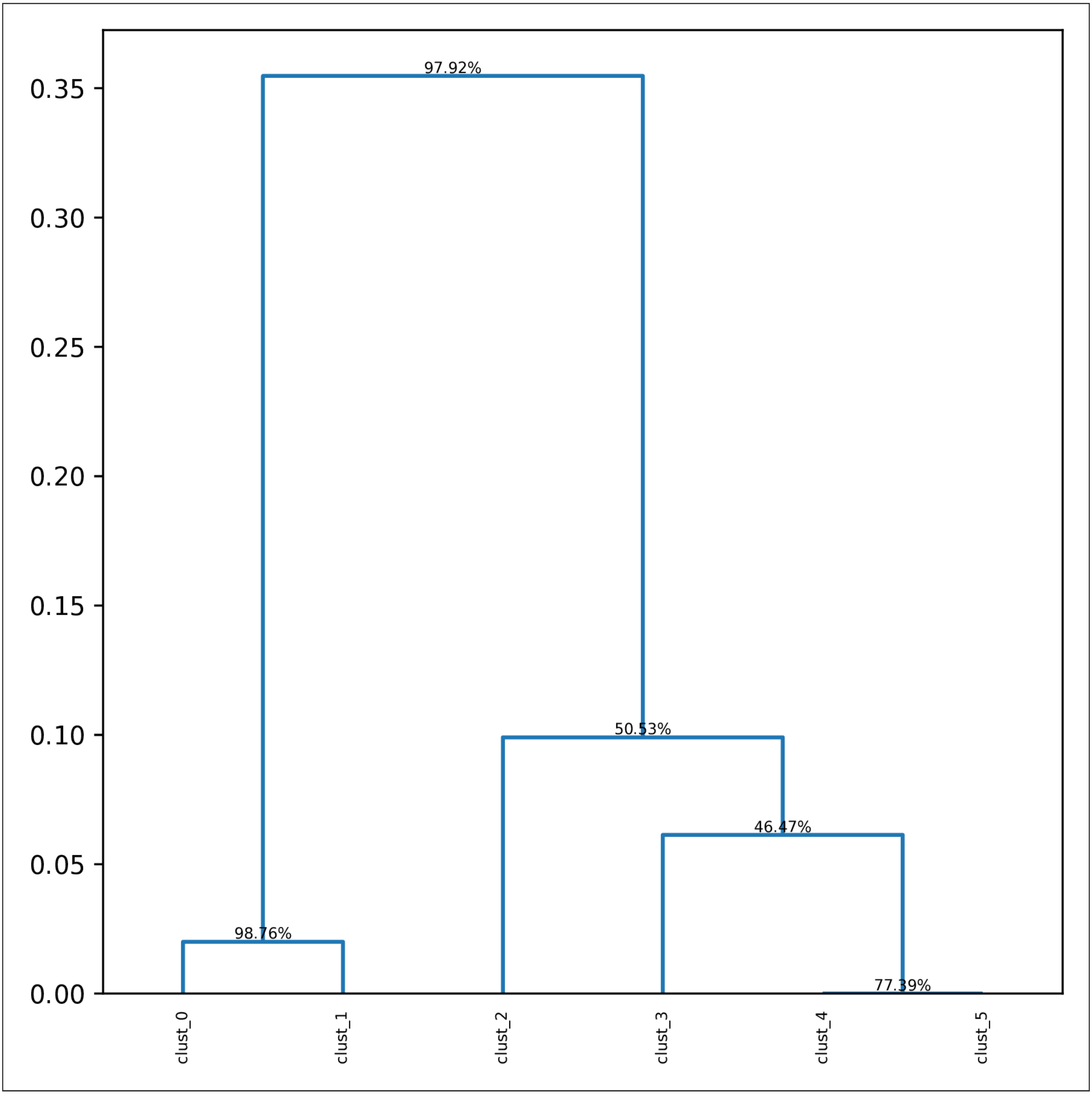

Sample: G0016 with all cells per cluster

Min. read depth: 10,  Mean UMIs per cluster:  12,972,180

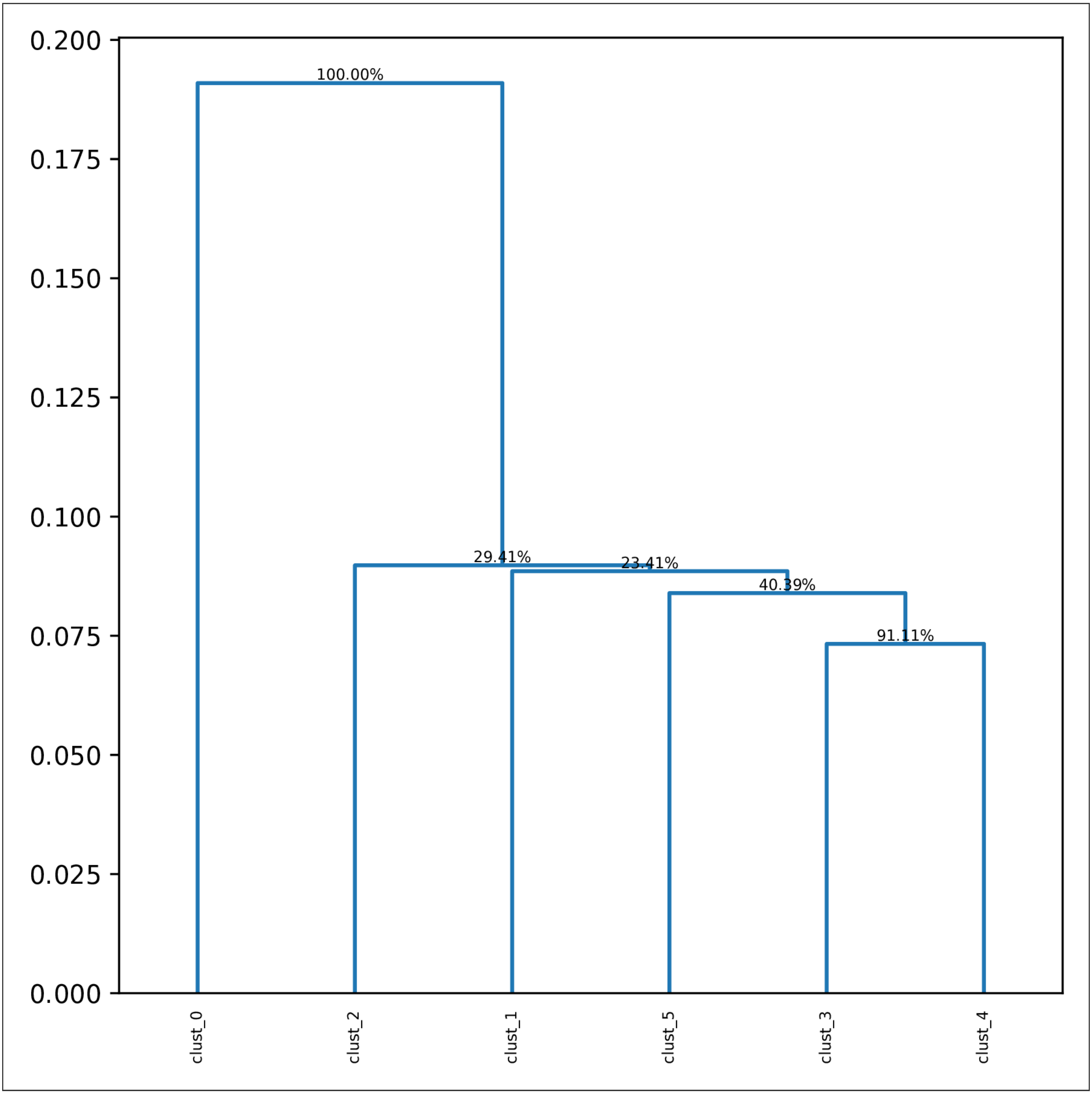

Sample: G0016 with 500 cells per cluster

Min. read depth: 10,  Mean UMIs per cluster:  8,872,918

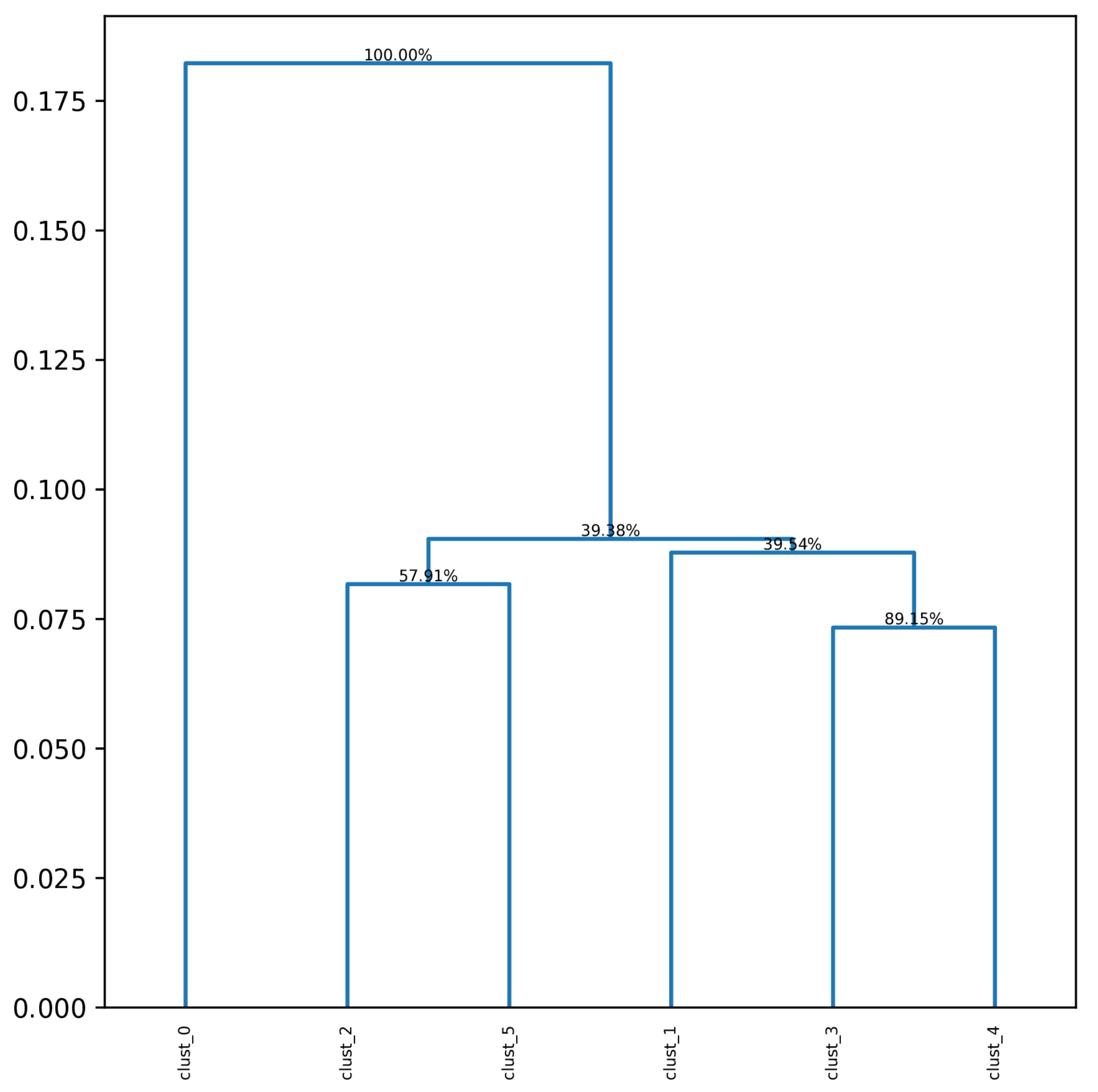

Sample: G0016 with 100 cells per cluster

Min. read depth: 10,  Mean UMIs per cluster:  2,157,994

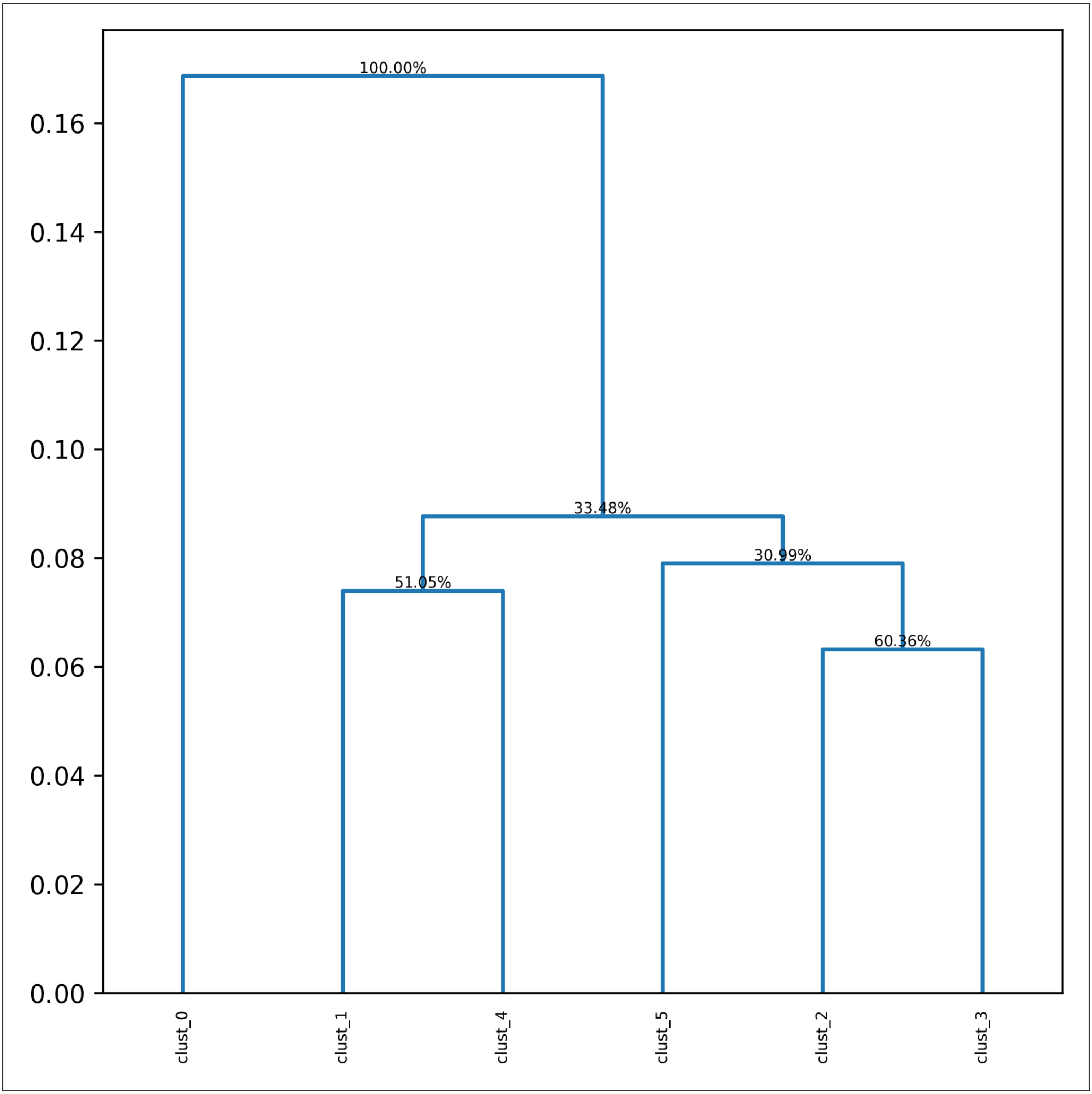

Sample: G0016 with 50 cells per cluster

Min. read depth: 10,  Mean UMIs per cluster:  1,048,666

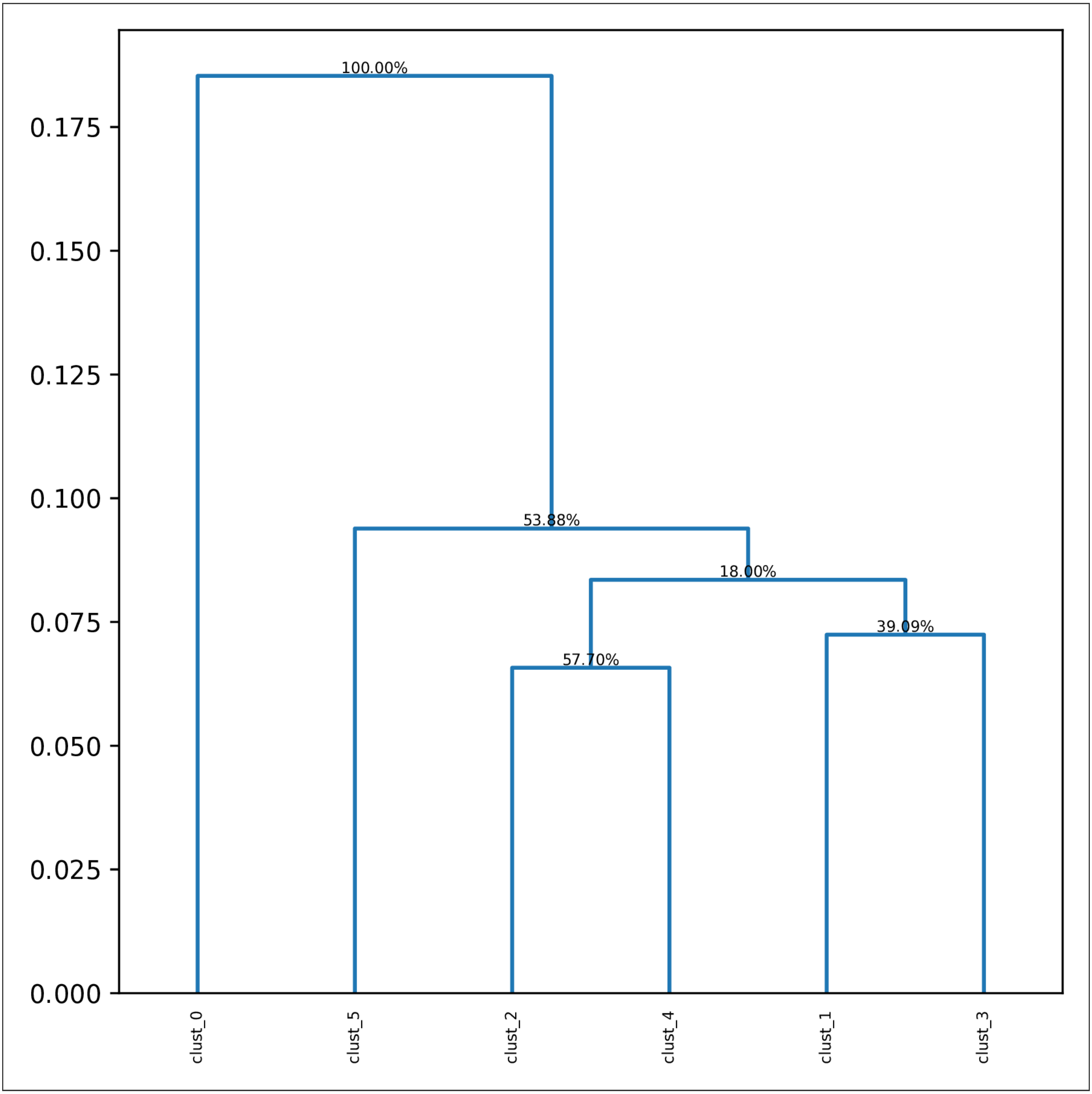

Sample: G0016 with 20 cells per cluster
Min. read depth: 10, Mean UMIs per cluster: 416,649

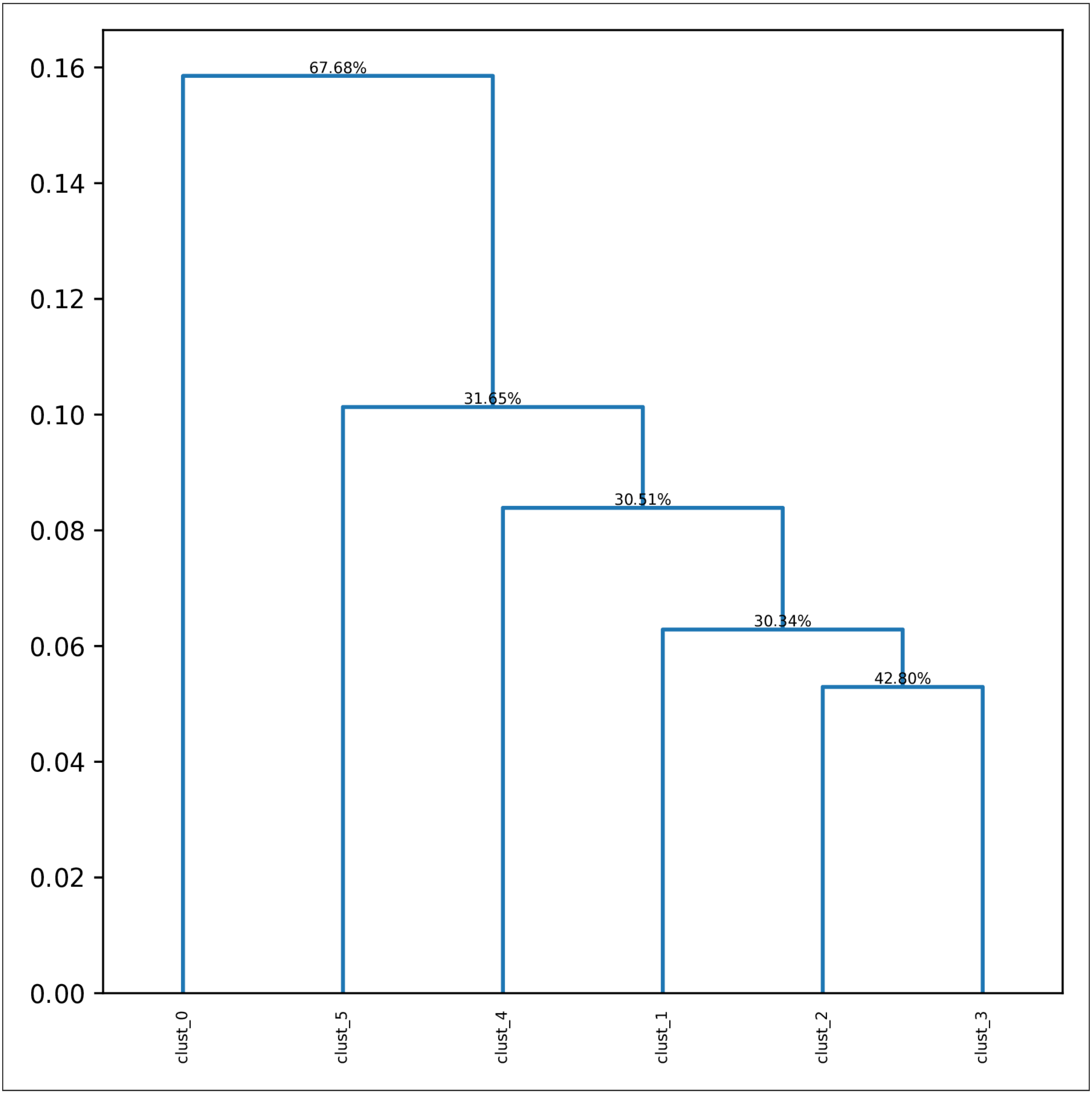

Sample: G0016 with 10 cells per cluster

Min. read depth: 10,  Mean UMIs per cluster:  210,908

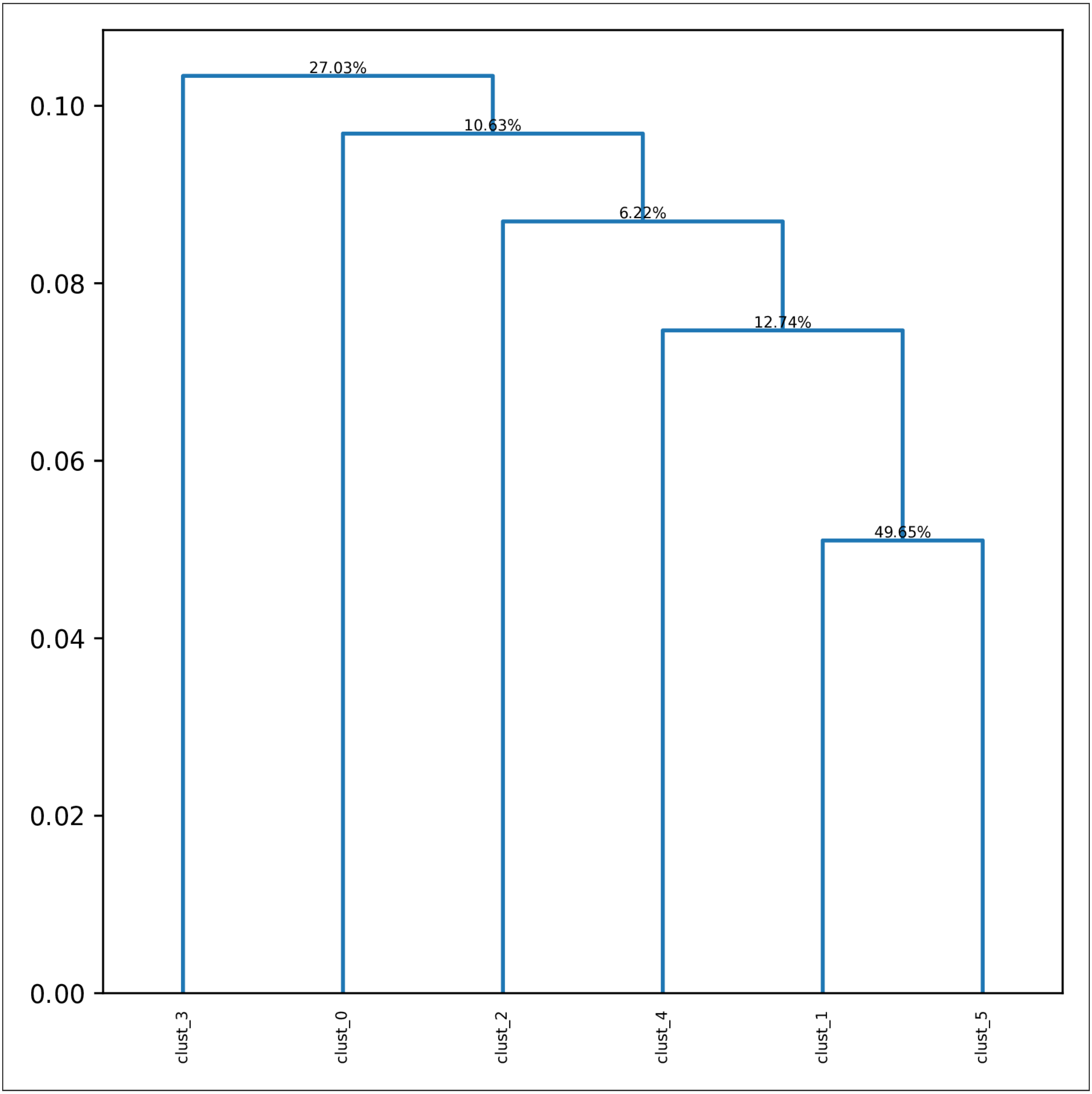

Sample: G0016 with 5 cells per cluster

Min. read depth: 10,  Mean UMIs per cluster:  107,371

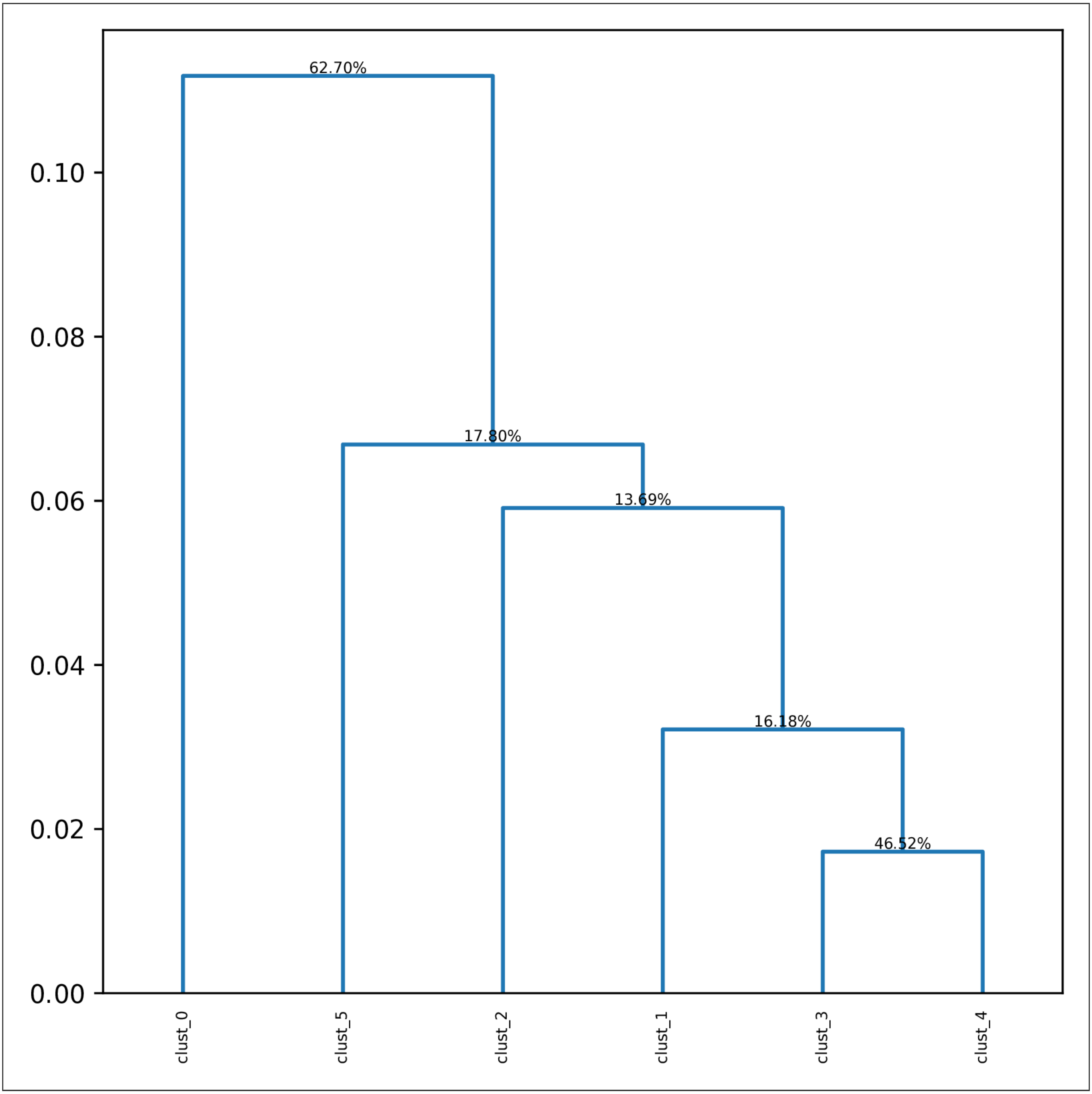

Sample: G0016 with 4 cells per cluster

Min. read depth: 10,  Mean UMIs per cluster:  71,484

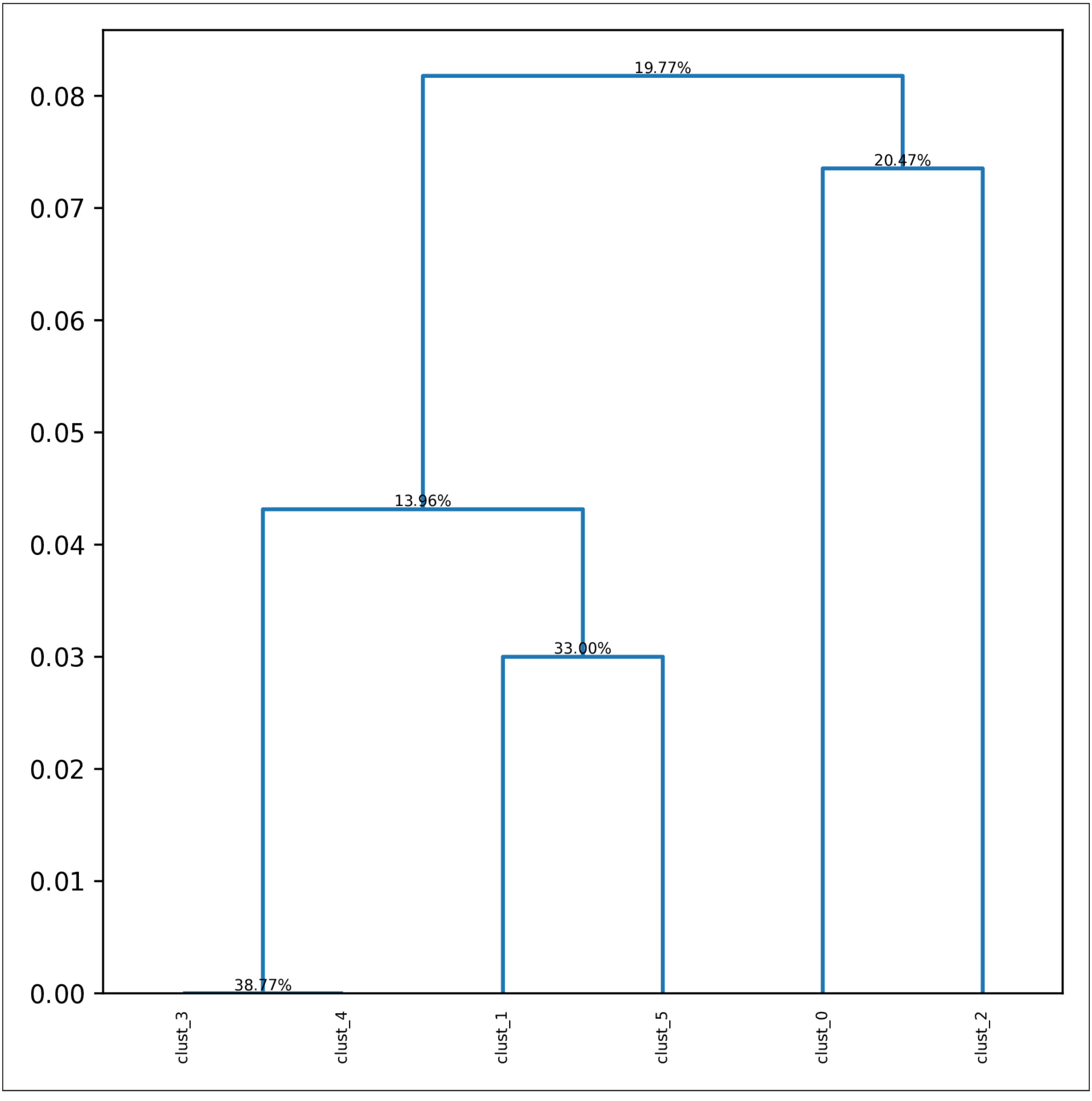

Sample: G0016 with 3 cells per cluster
Min. read depth: 10,  Mean UMIs per cluster:  60,691

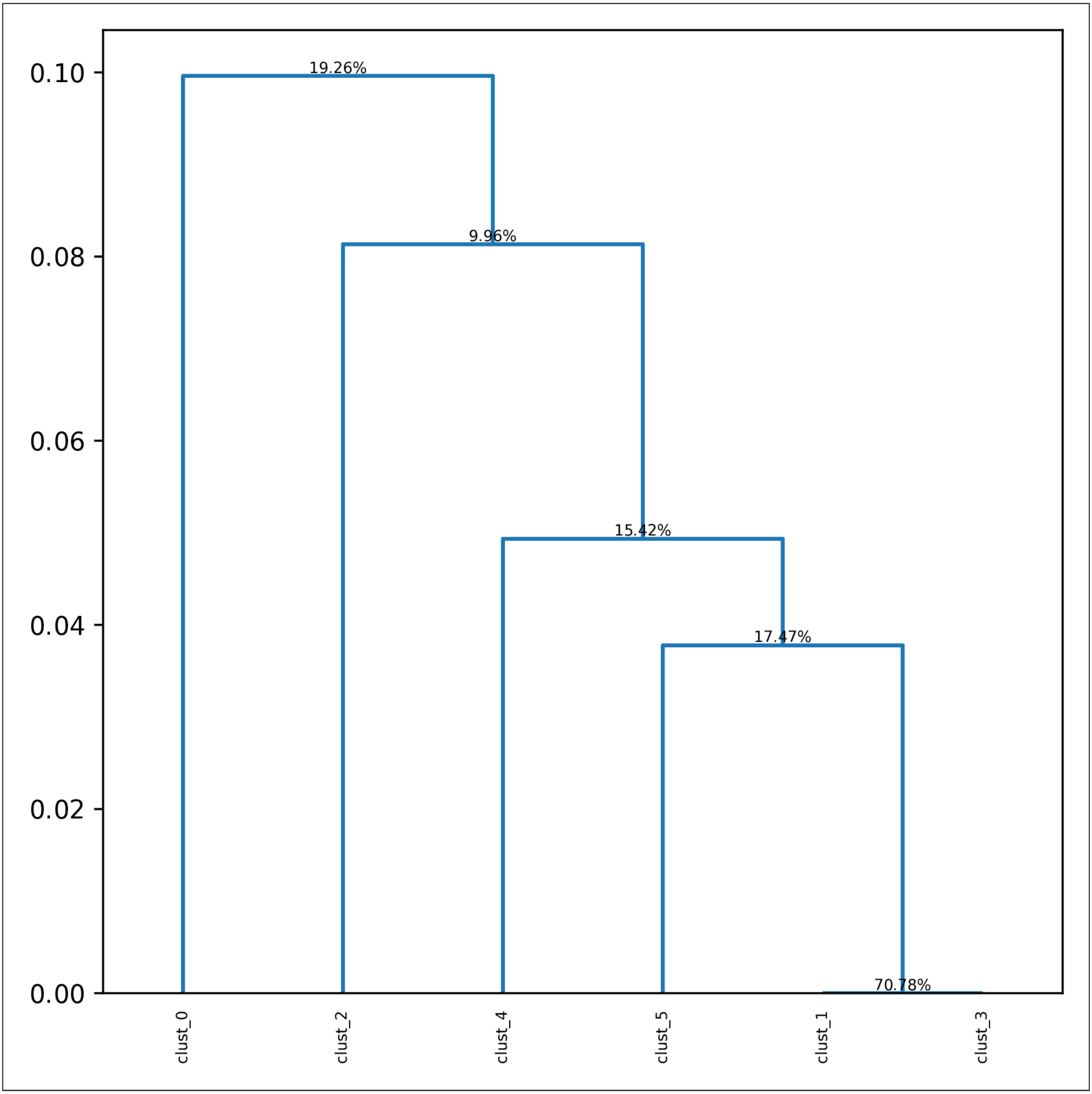

Sample: G0017 with all cells per cluster

Min. read depth: 10,  Mean UMIs per cluster:  25,706,603

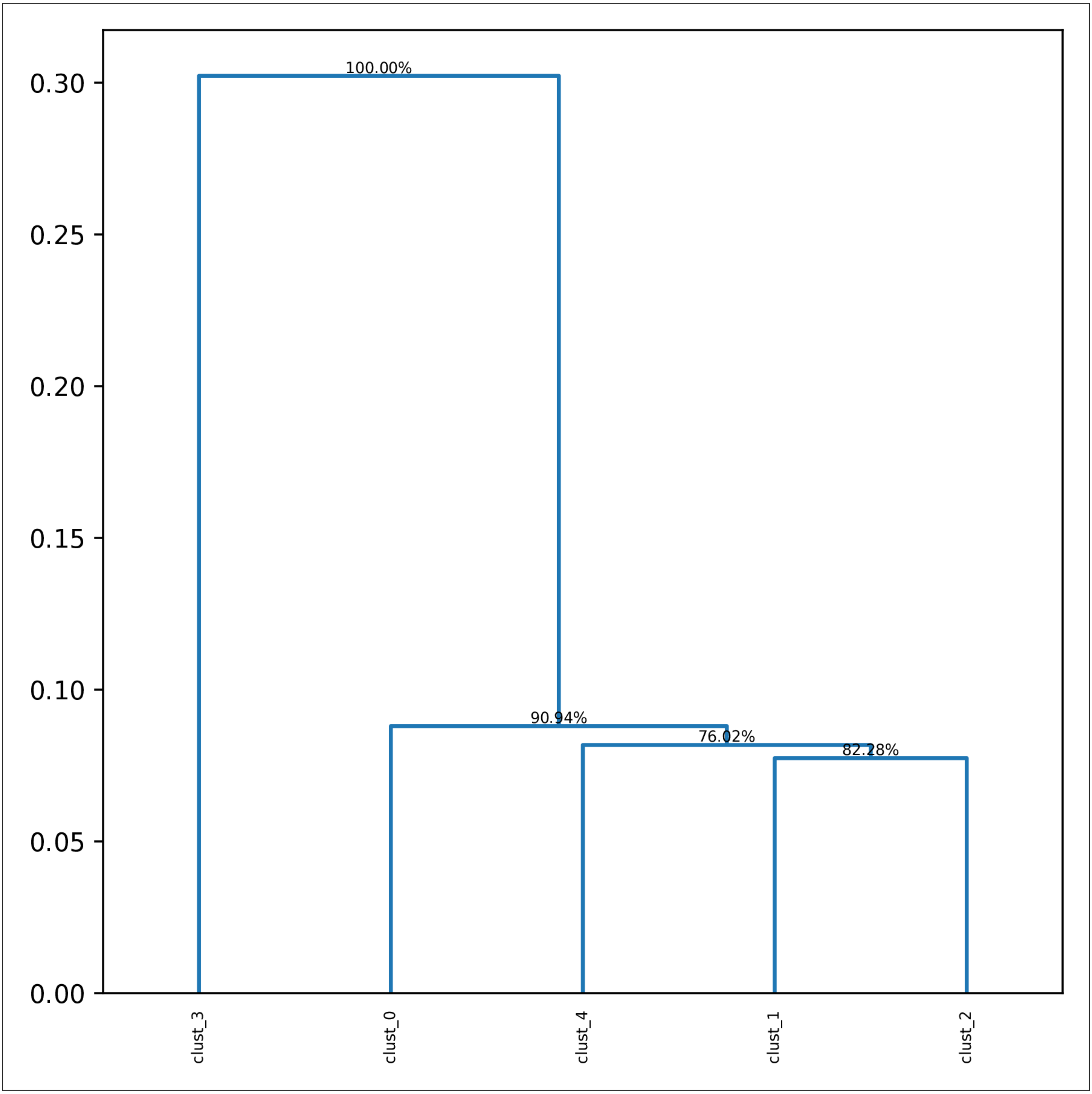

Sample: G0017 with 500 cells per cluster

Min. read depth: 10,  Mean UMIs per cluster:  10,602,599

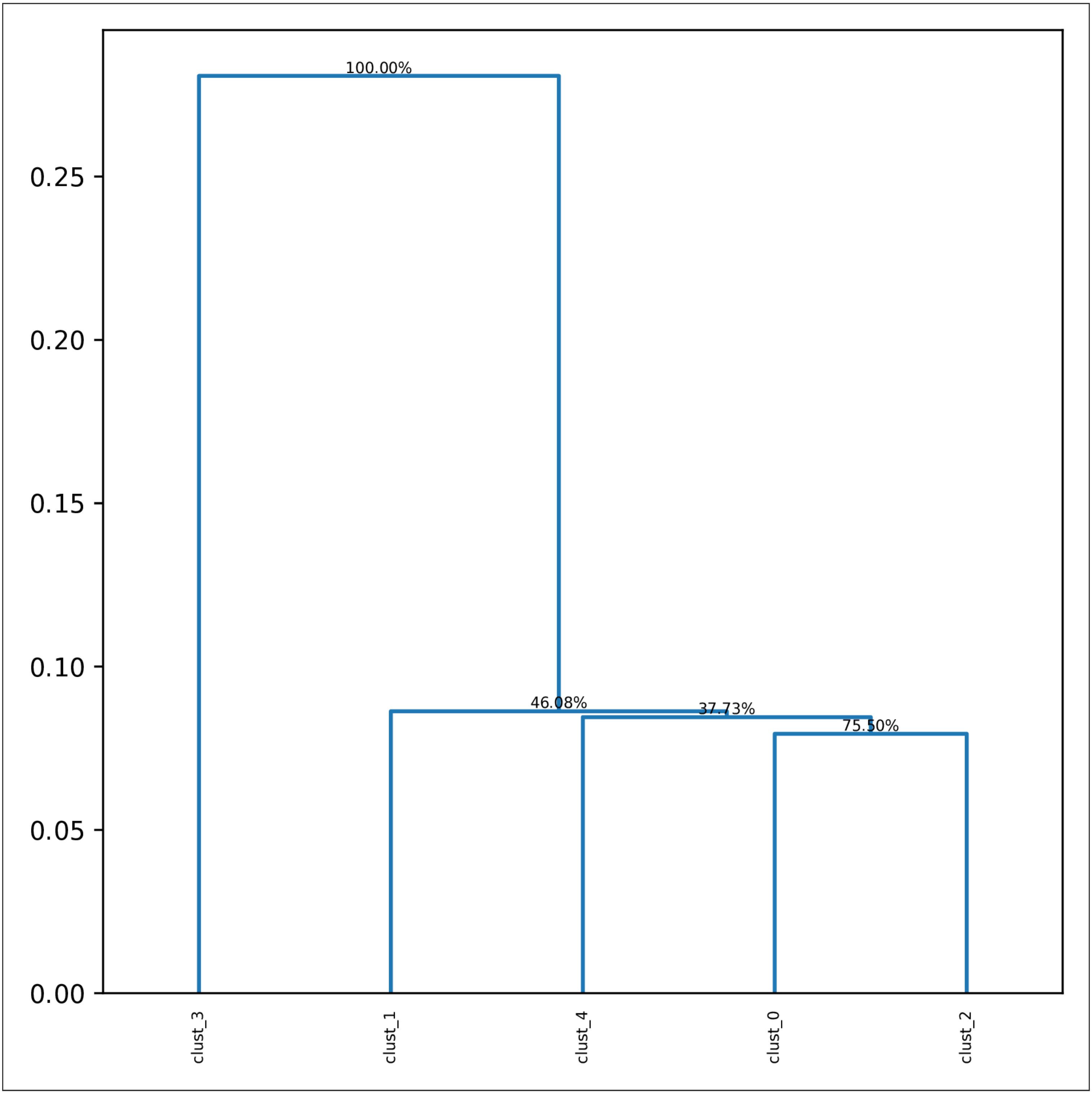

Sample: G0017 with 100 cells per cluster

Min. read depth: 10,  Mean UMIs per cluster:  2,377,534

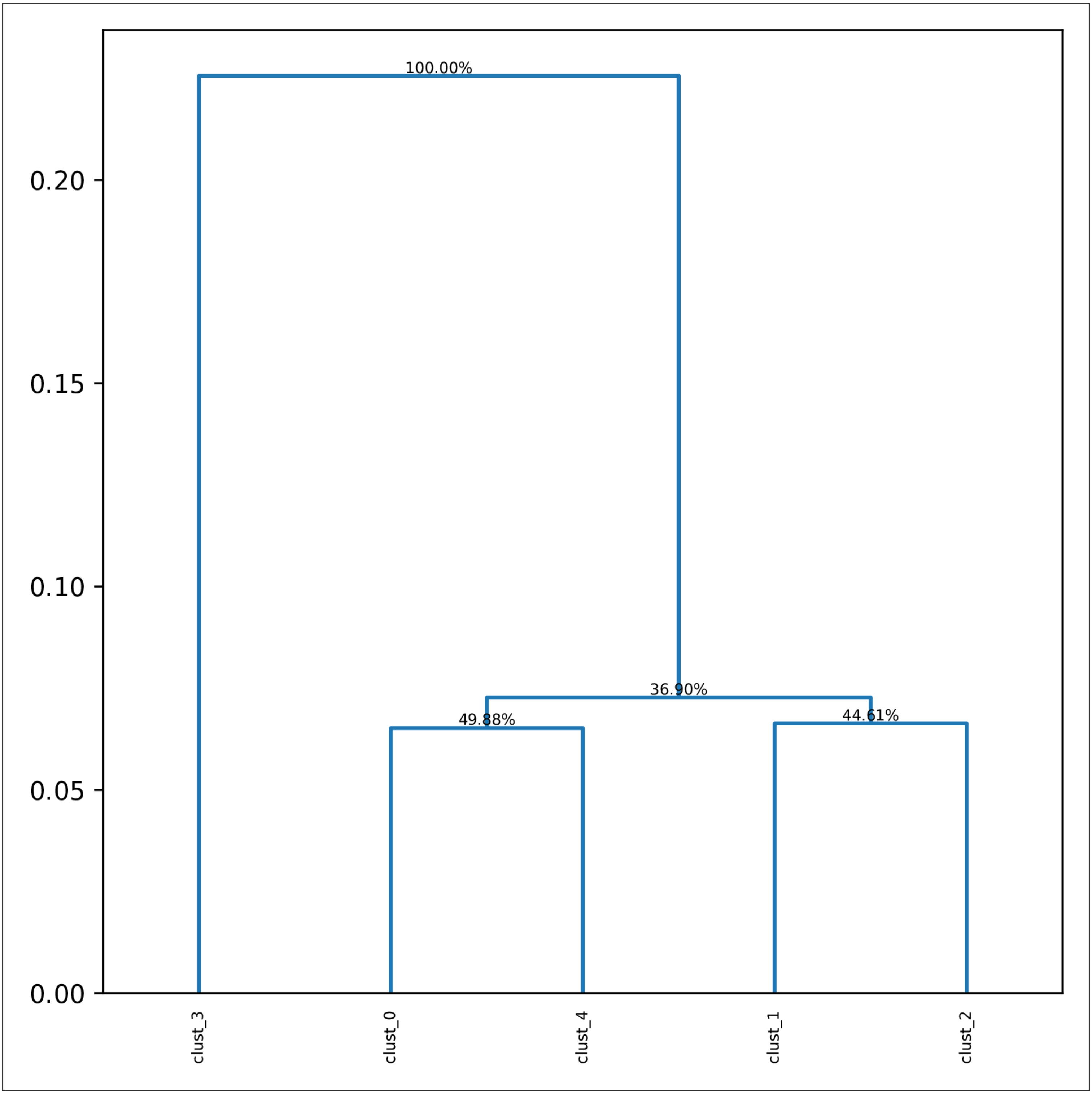

Sample: G0017 with 50 cells per cluster

Min. read depth: 10,  Mean UMIs per cluster:  1,176,397

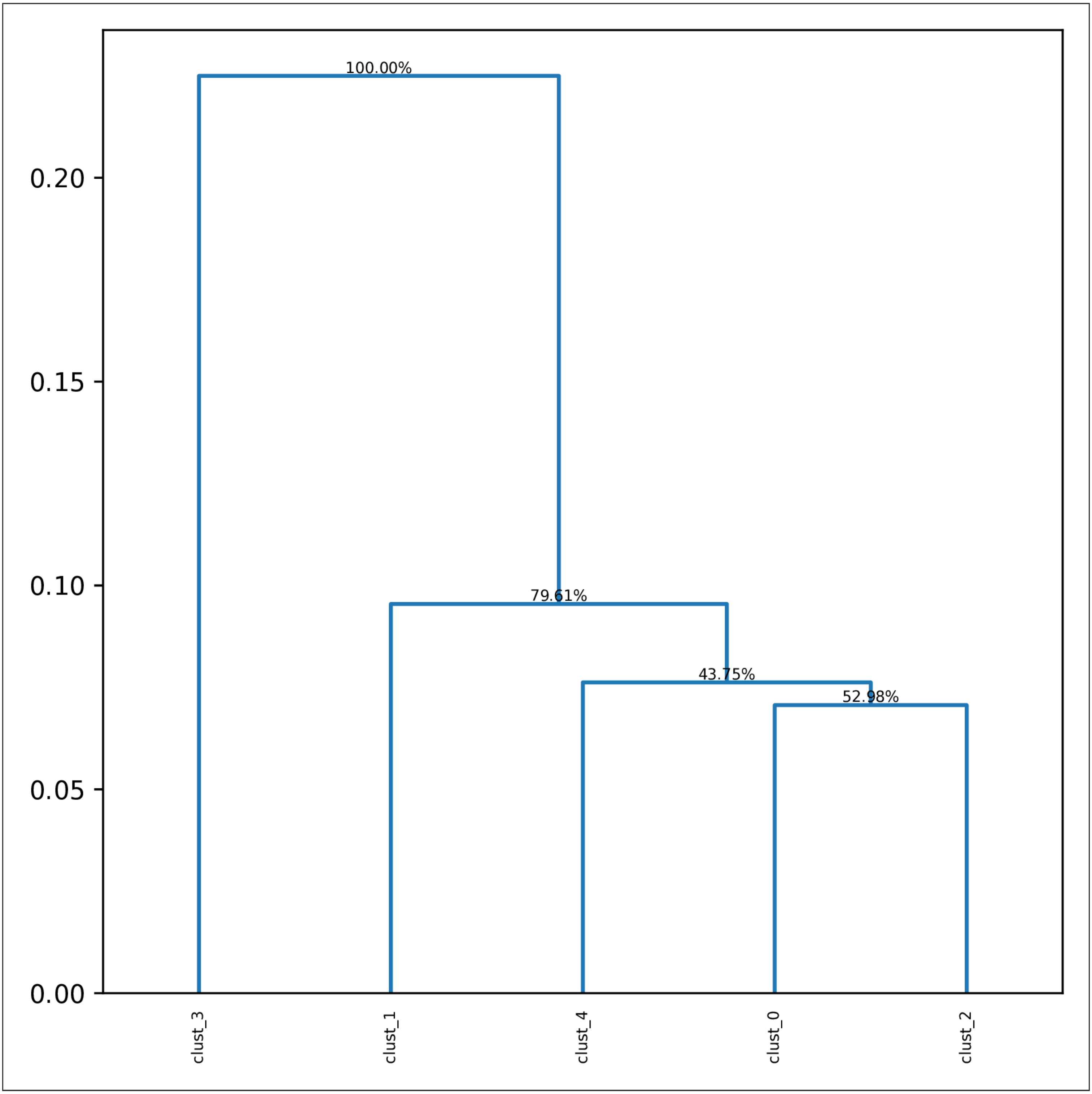

Sample: G0017 with 20 cells per cluster

Min. read depth: 10,  Mean UMIs per cluster:  479,070

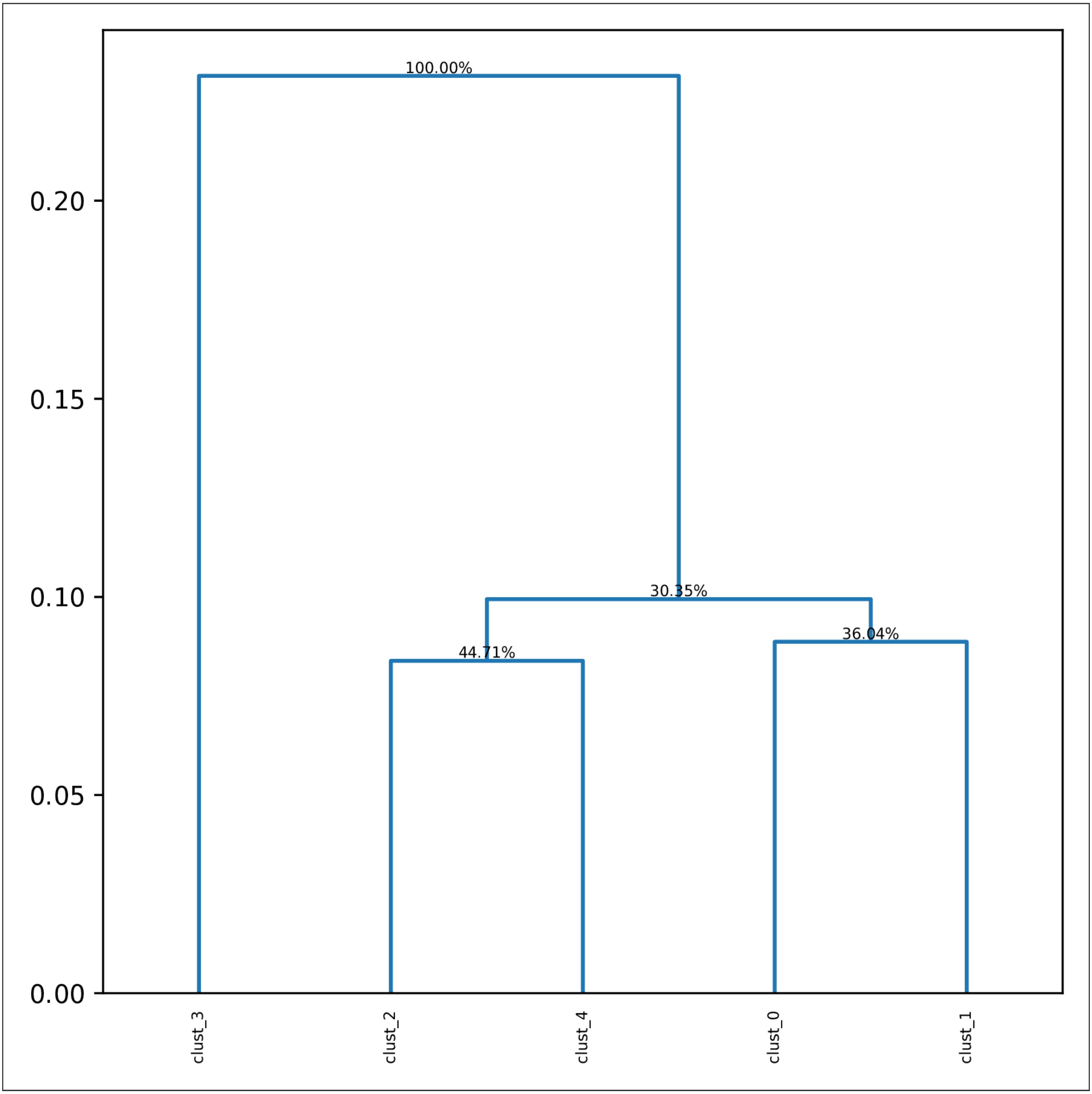

Sample: G0017 with 10 cells per cluster

Min. read depth: 10,  Mean UMIs per cluster:  243,620

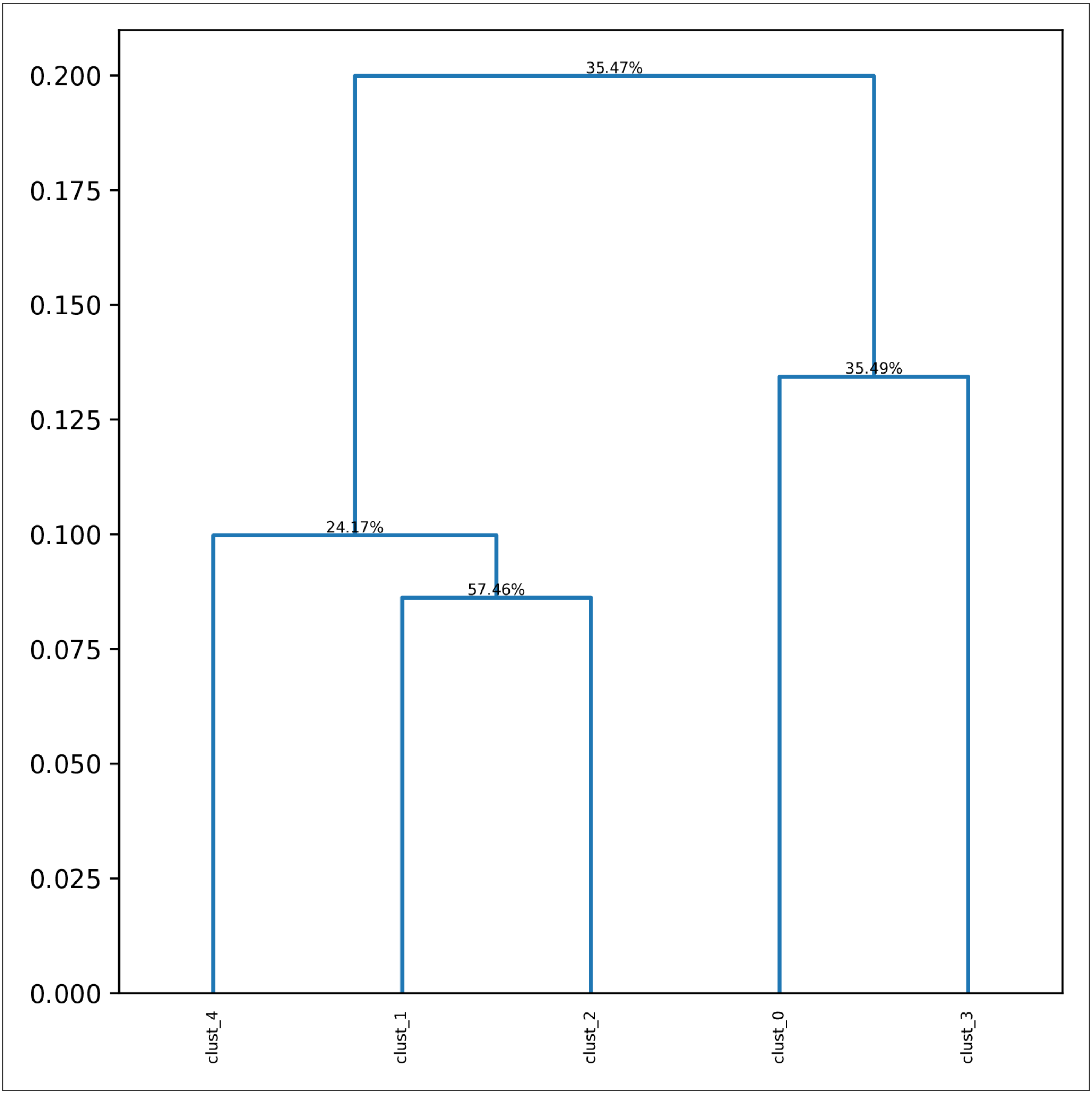

Sample: G0017 with 5 cells per cluster
Min. read depth: 10,  Mean UMIs per cluster:  121,857

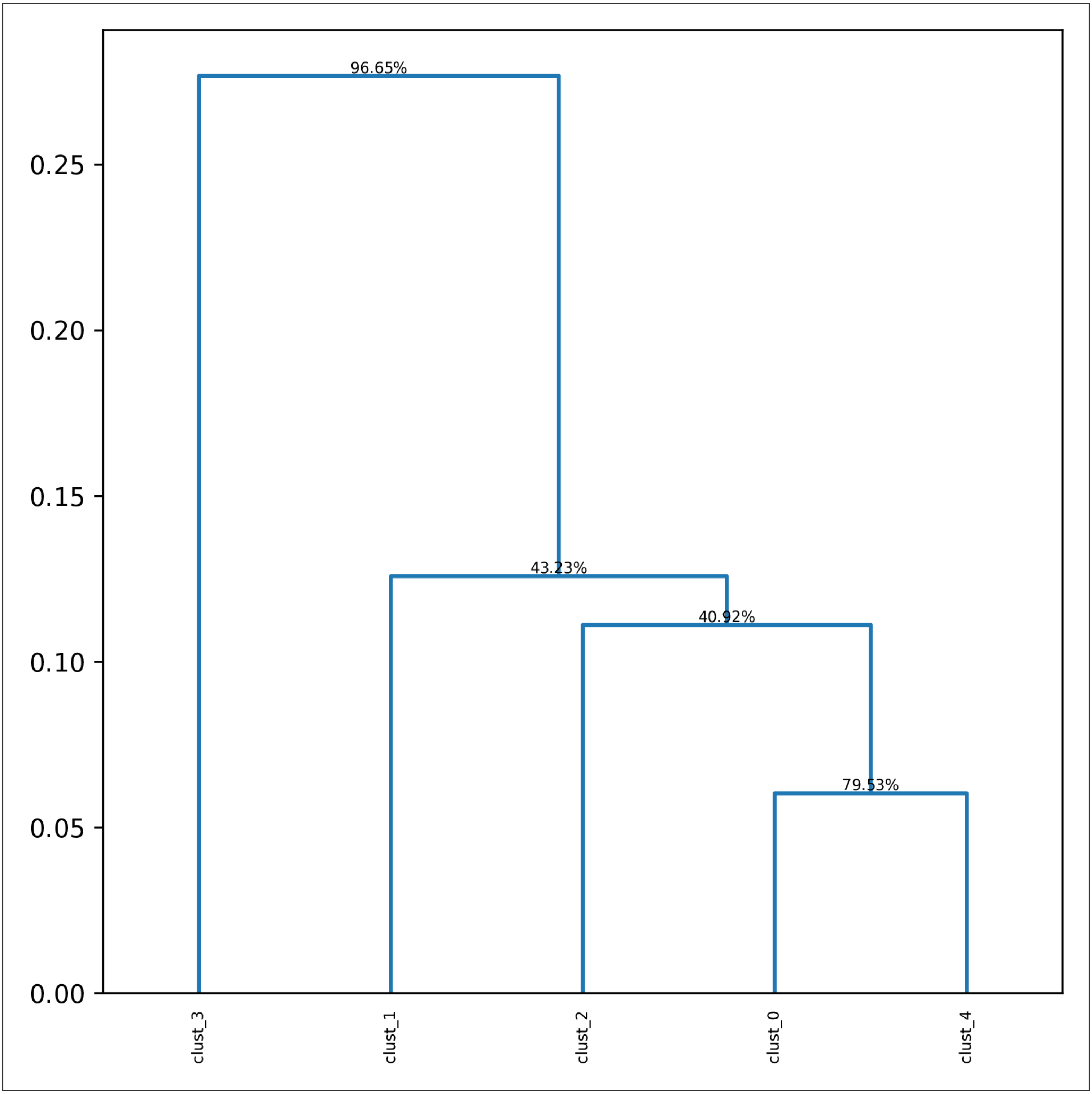

Sample: G0017 with 4 cells per cluster

Min. read depth: 10,  Mean UMIs per cluster:  111,794

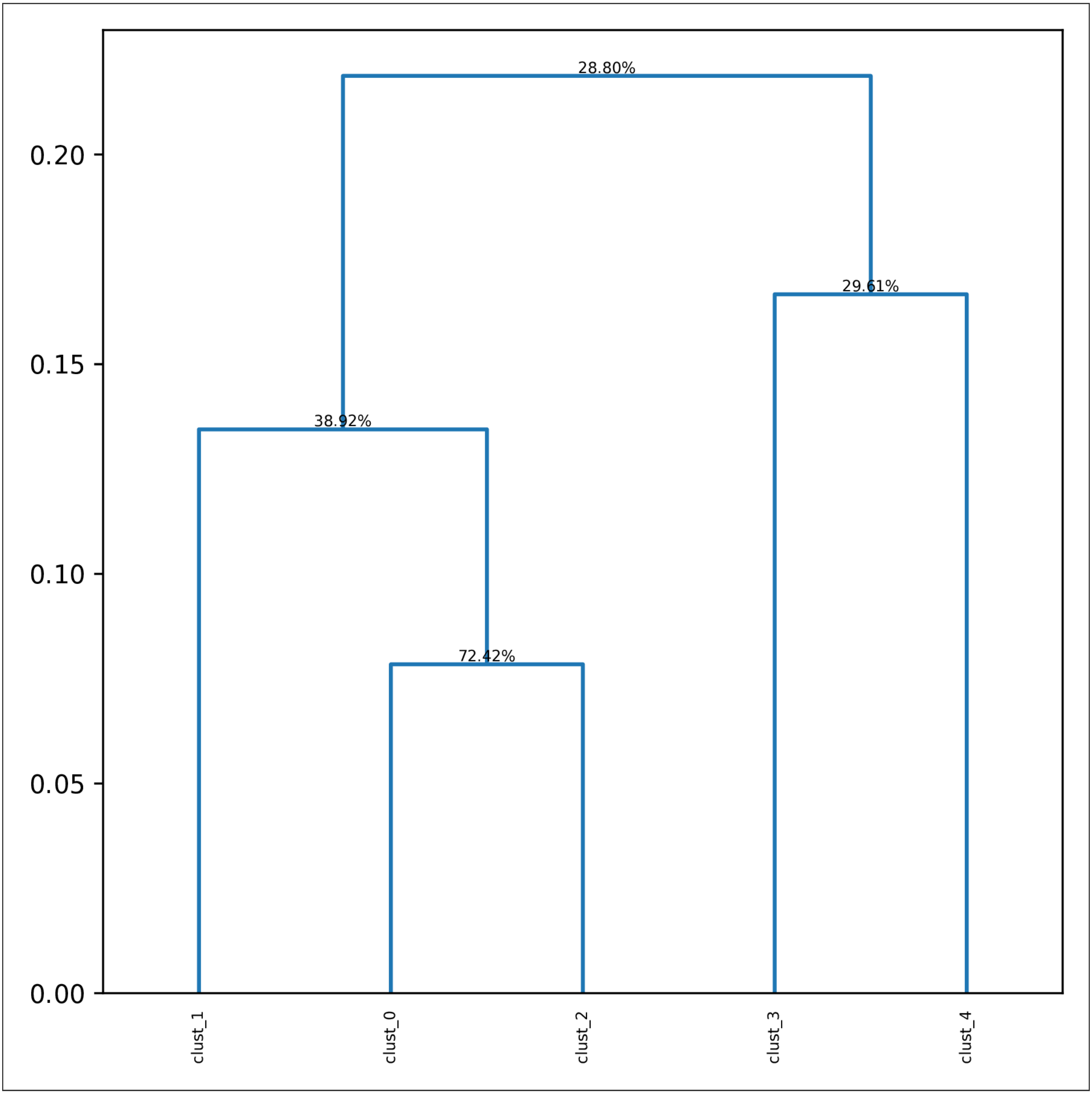

Sample: G0017 with 3 cells per cluster

Min. read depth: 10, Mean UMIs per cluster: 87,989

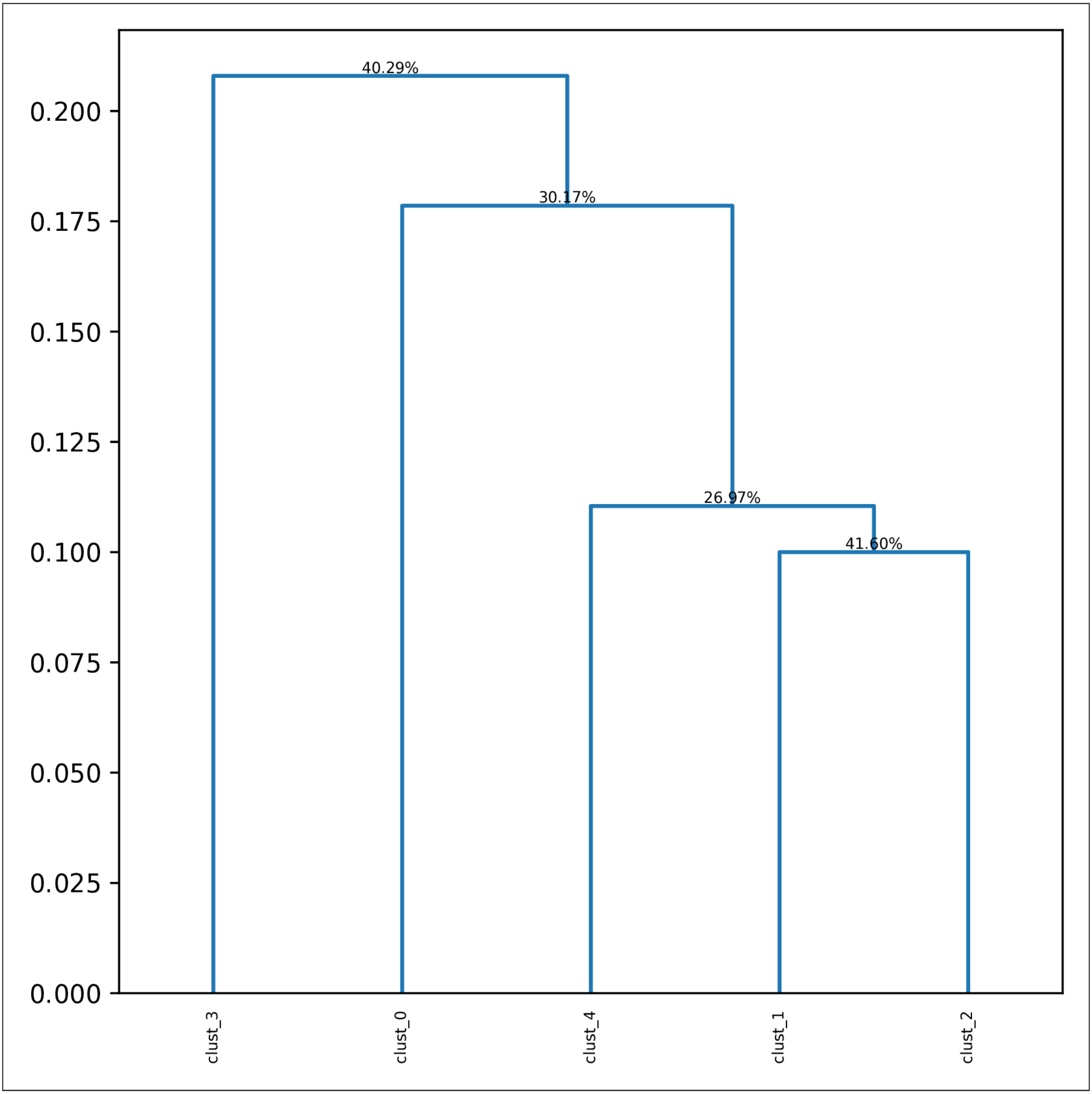

Sample: G0018 with all cells per cluster

Min. read depth: 10,  Mean UMIs per cluster:  17,937,563

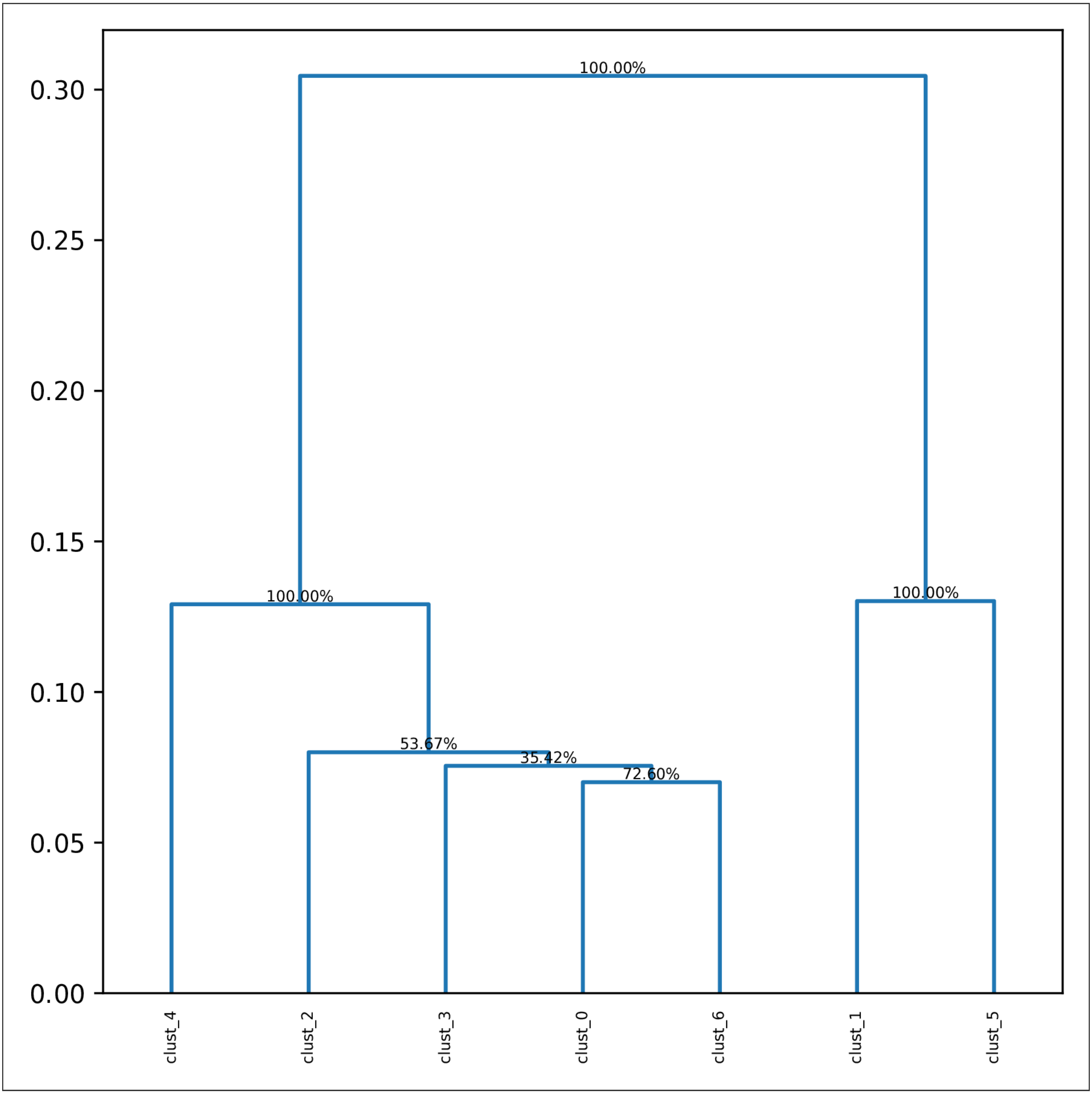

Sample: G0018 with 500 cells per cluster

Min. read depth: 10,  Mean UMIs per cluster:  7,839,699

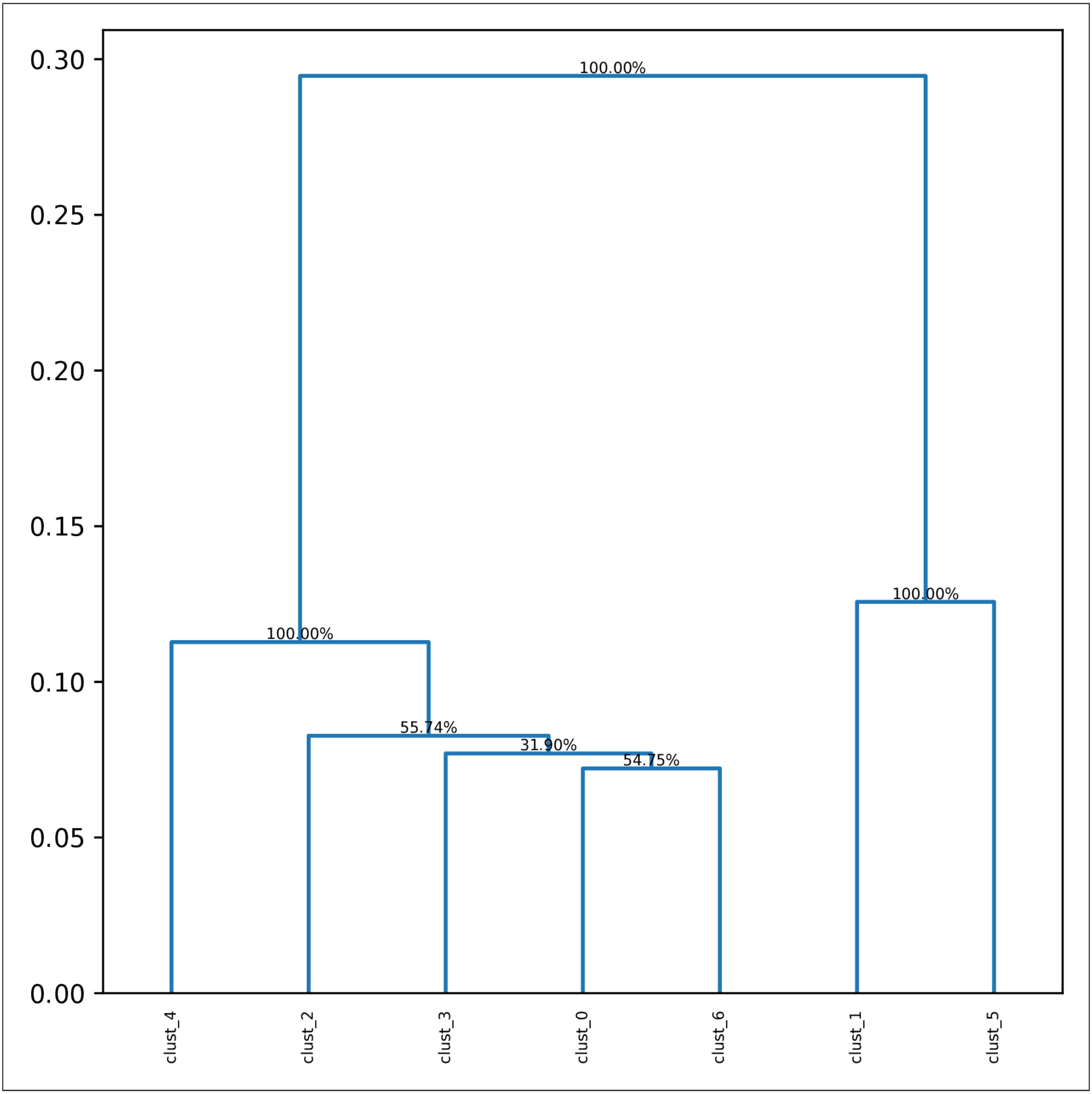

Sample: G0018 with 100 cells per cluster

Min. read depth: 10,  Mean UMIs per cluster:  1,634,743

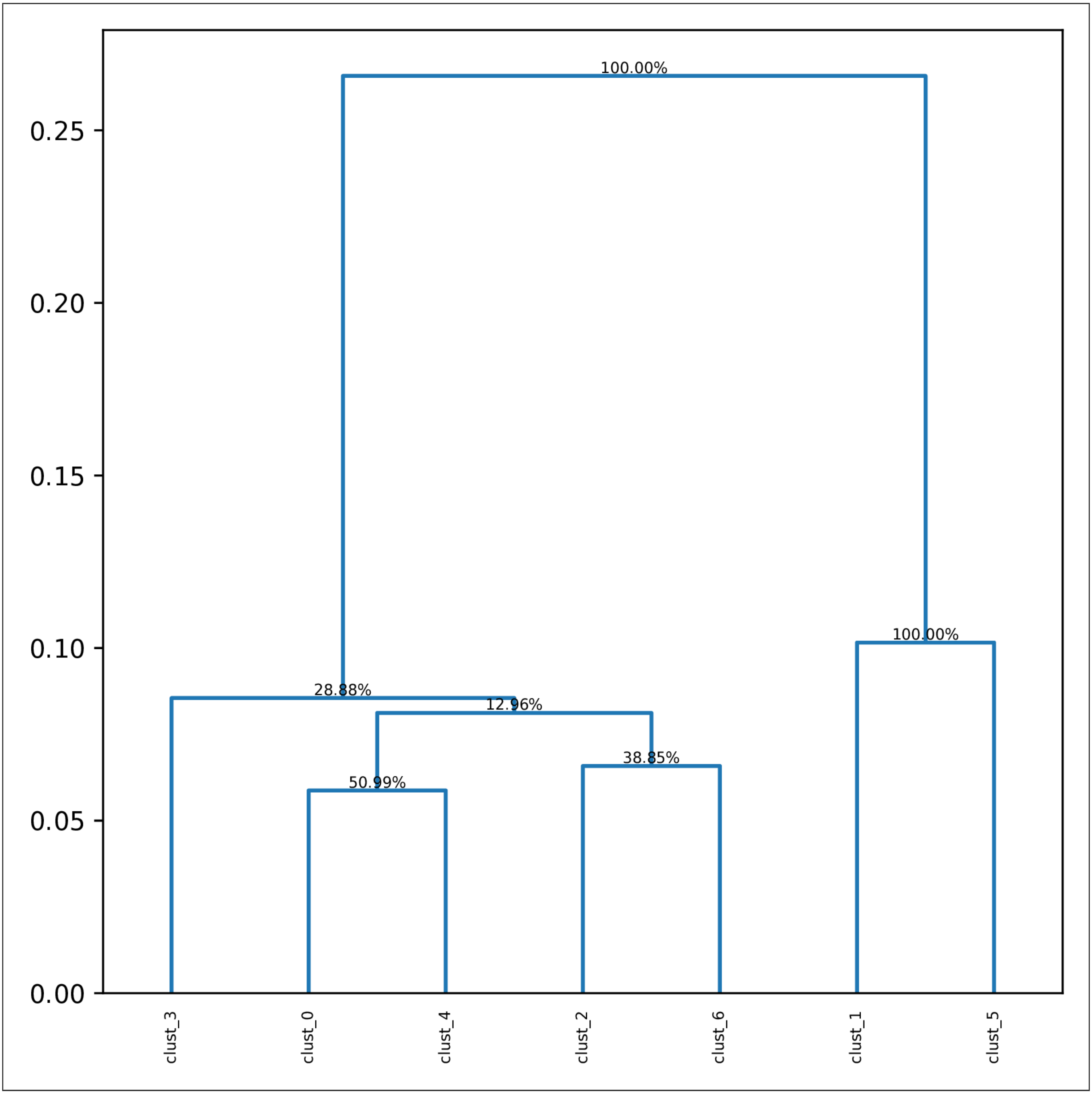

Sample: G0018 with 50 cells per cluster

Min. read depth: 10,  Mean UMIs per cluster:  791,823

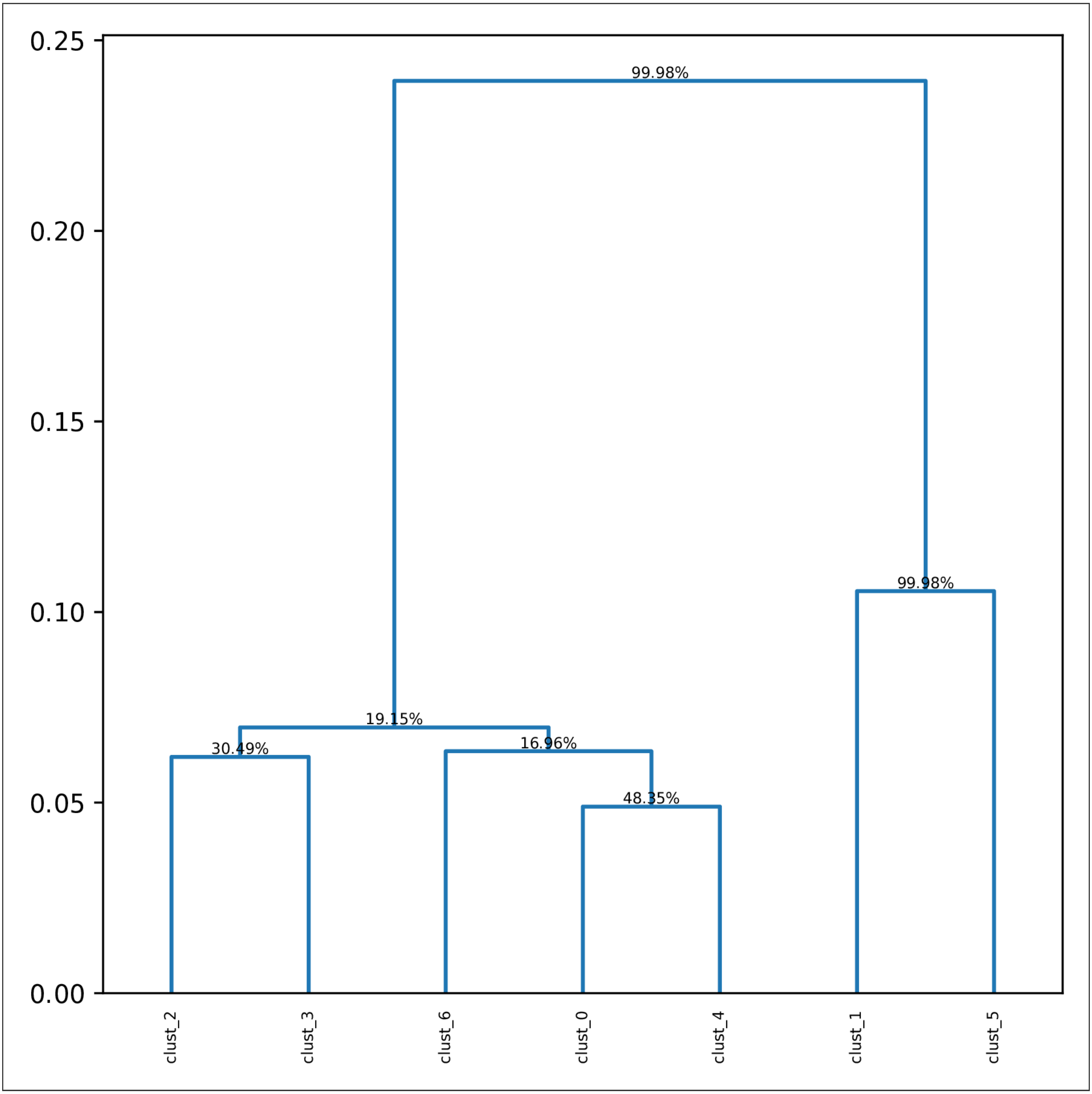

Sample: G0018 with 20 cells per cluster

Min. read depth: 10,  Mean UMIs per cluster:  329,059

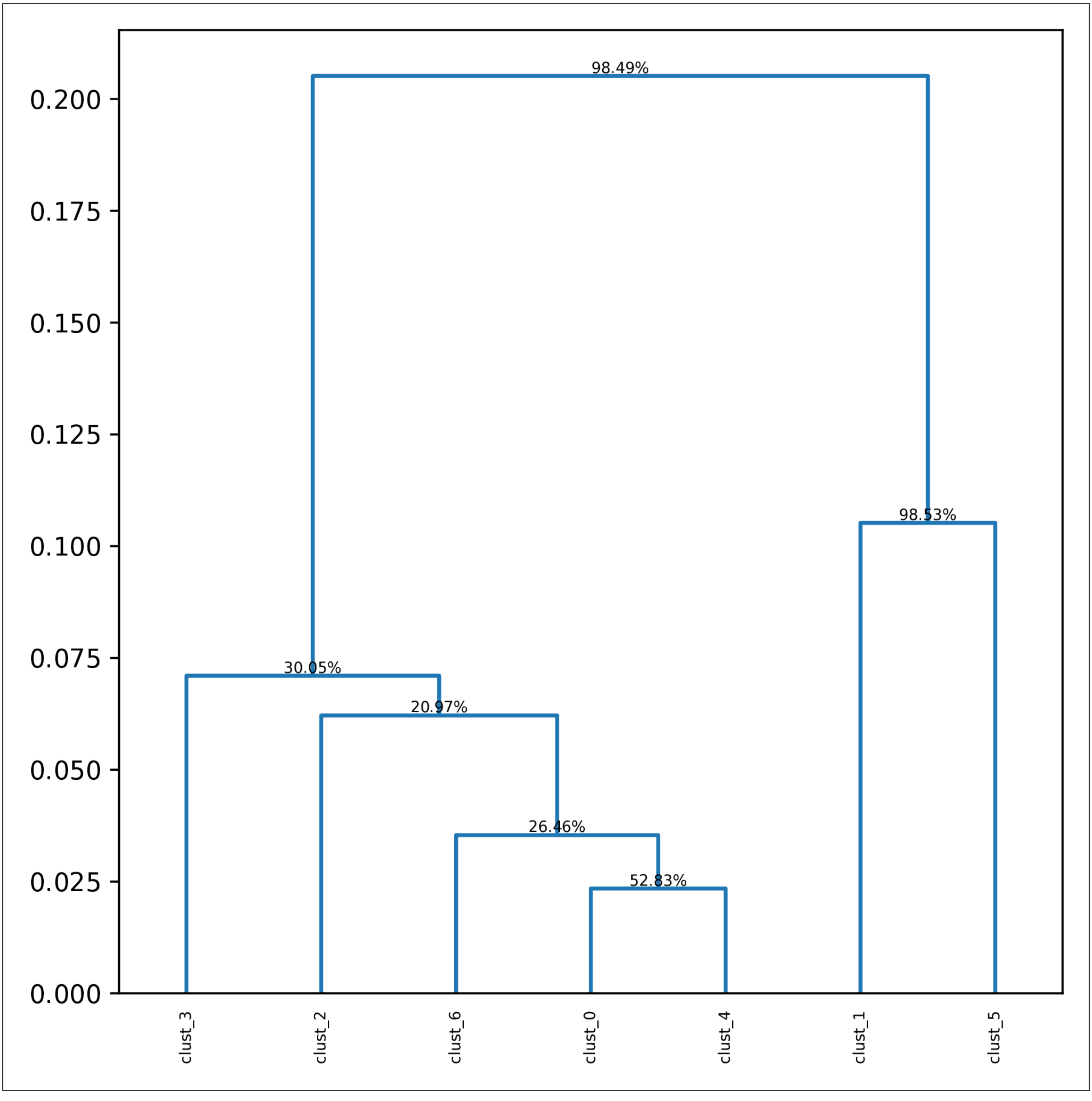

Sample: G0018 with 10 cells per cluster

Min. read depth: 10,  Mean UMIs per cluster:  134,911

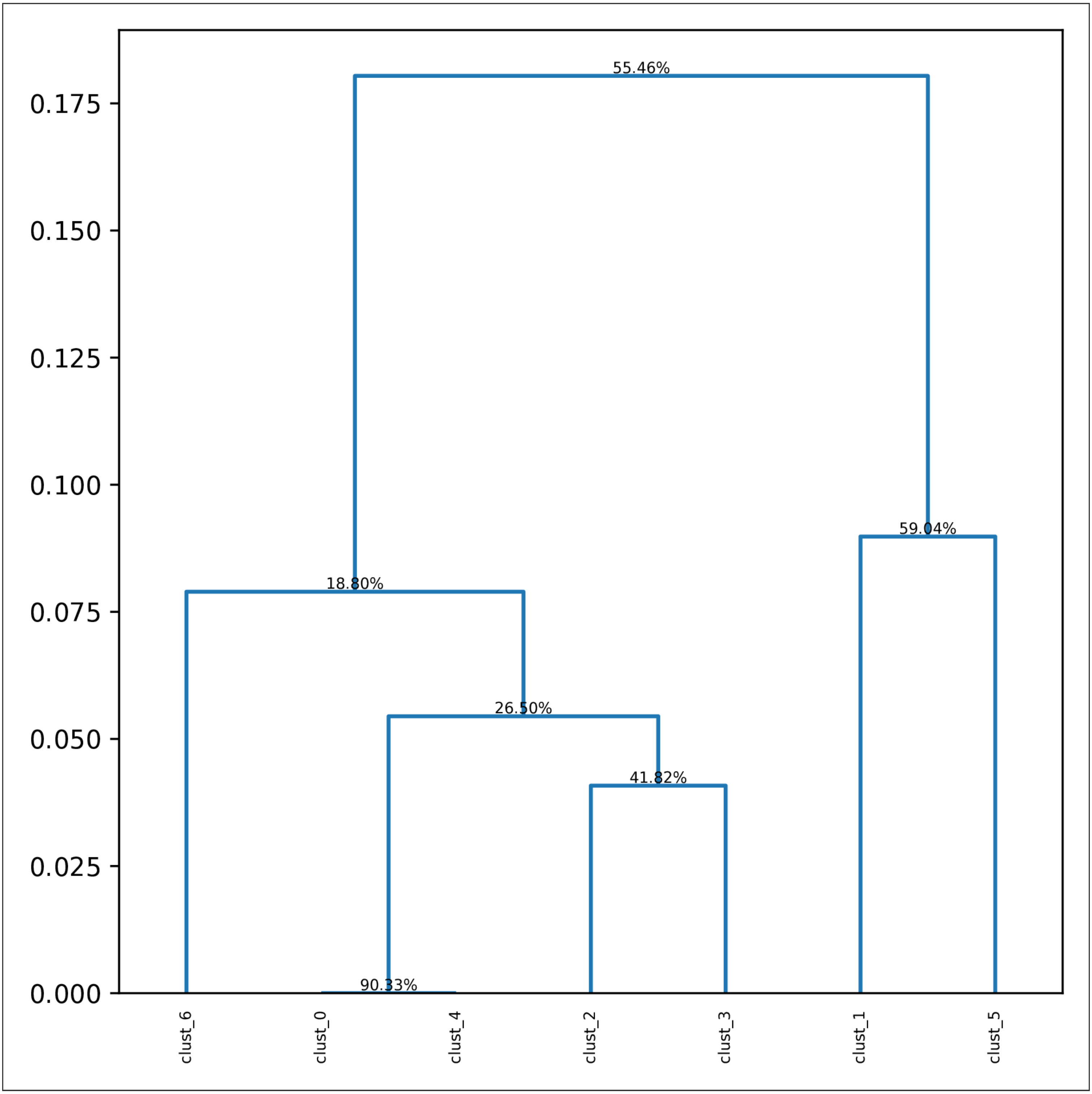

Sample: G0018 with 5 cells per cluster
Min. read depth: 10,  Mean UMIs per cluster:  71,560

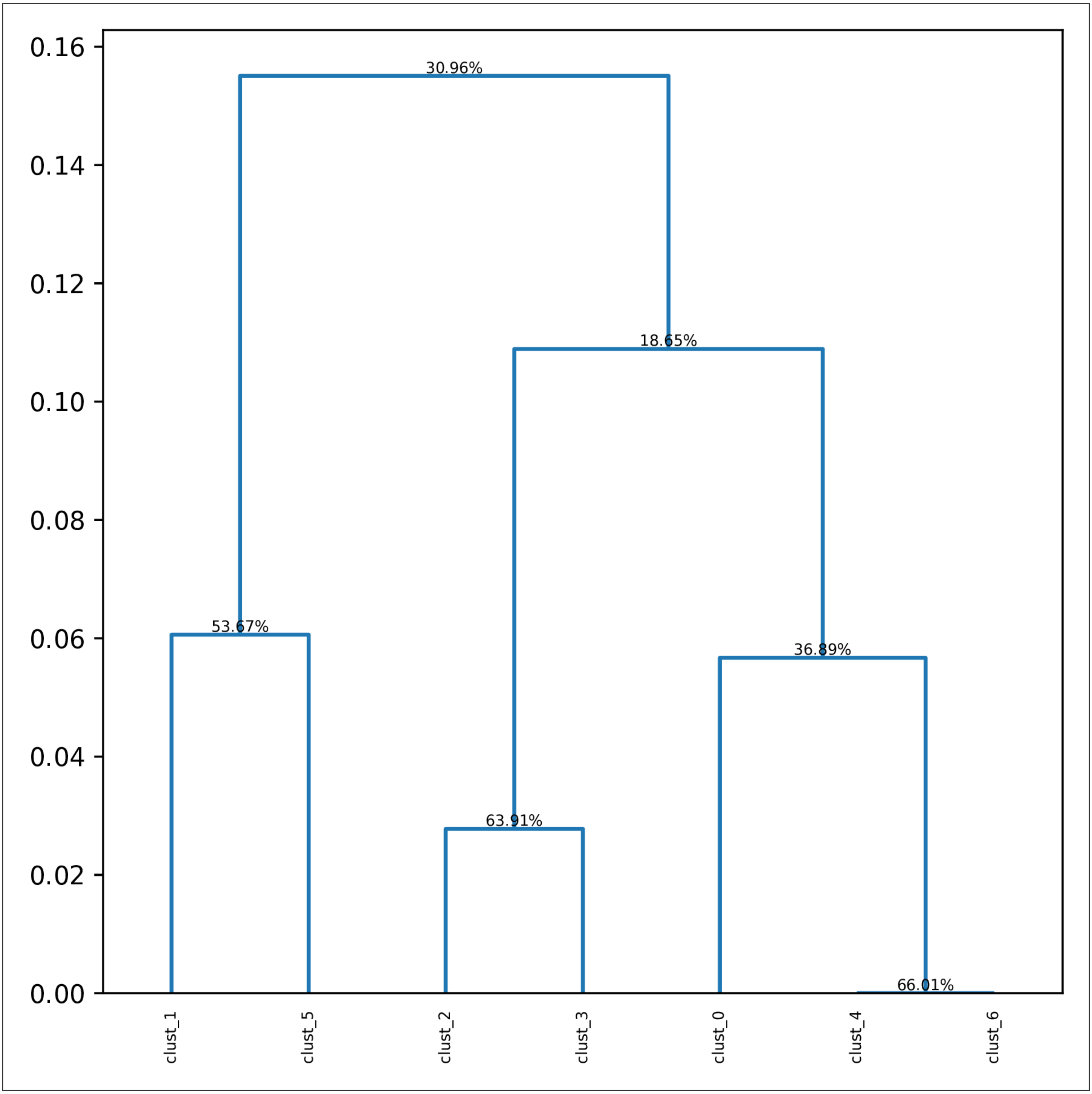

Sample: G0018 with 4 cells per cluster

Min. read depth: 10,  Mean UMIs per cluster:  59,239

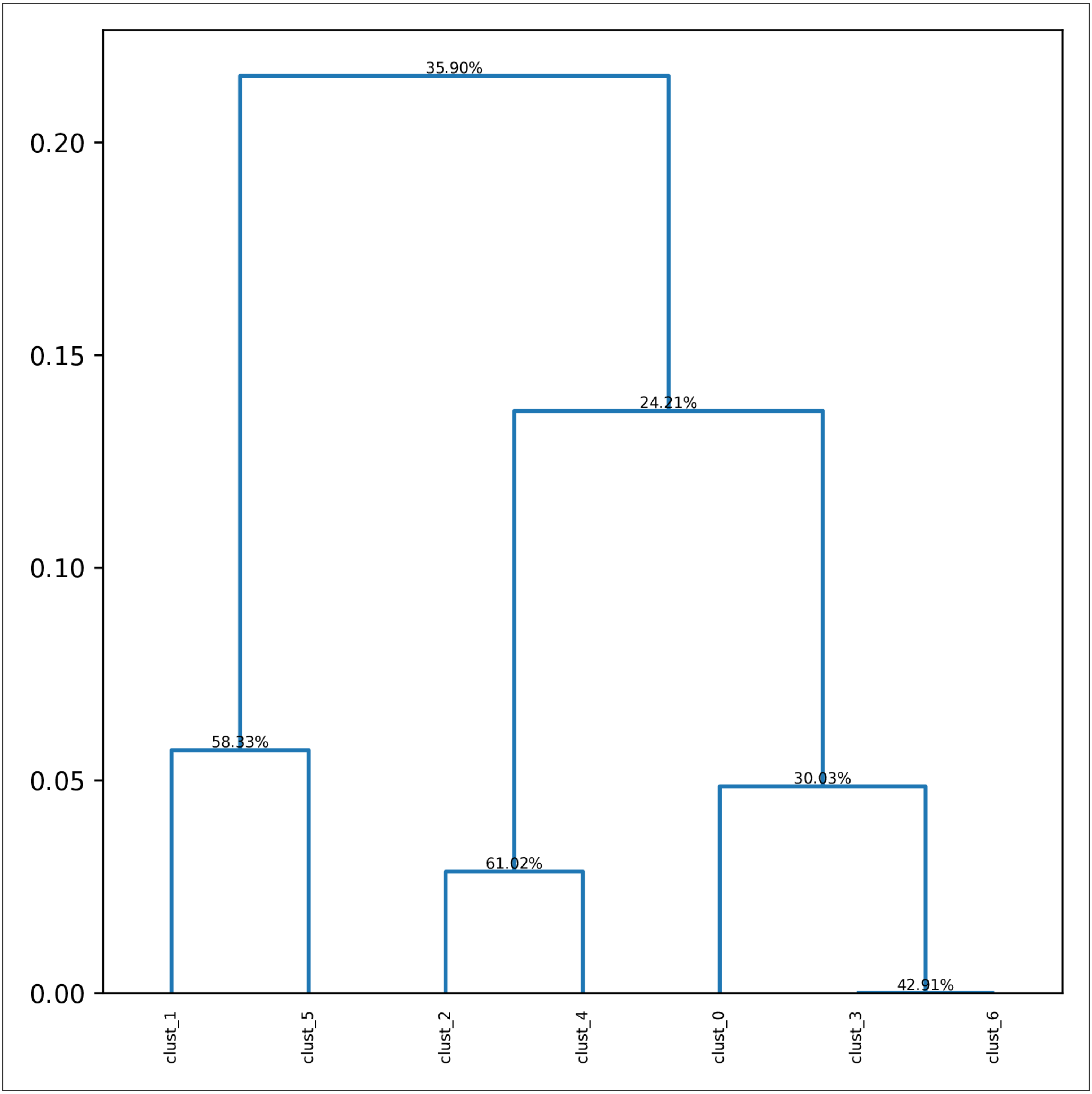

Sample: G0019 with all cells per cluster

Min. read depth: 10,  Mean UMIs per cluster:  5,612,857

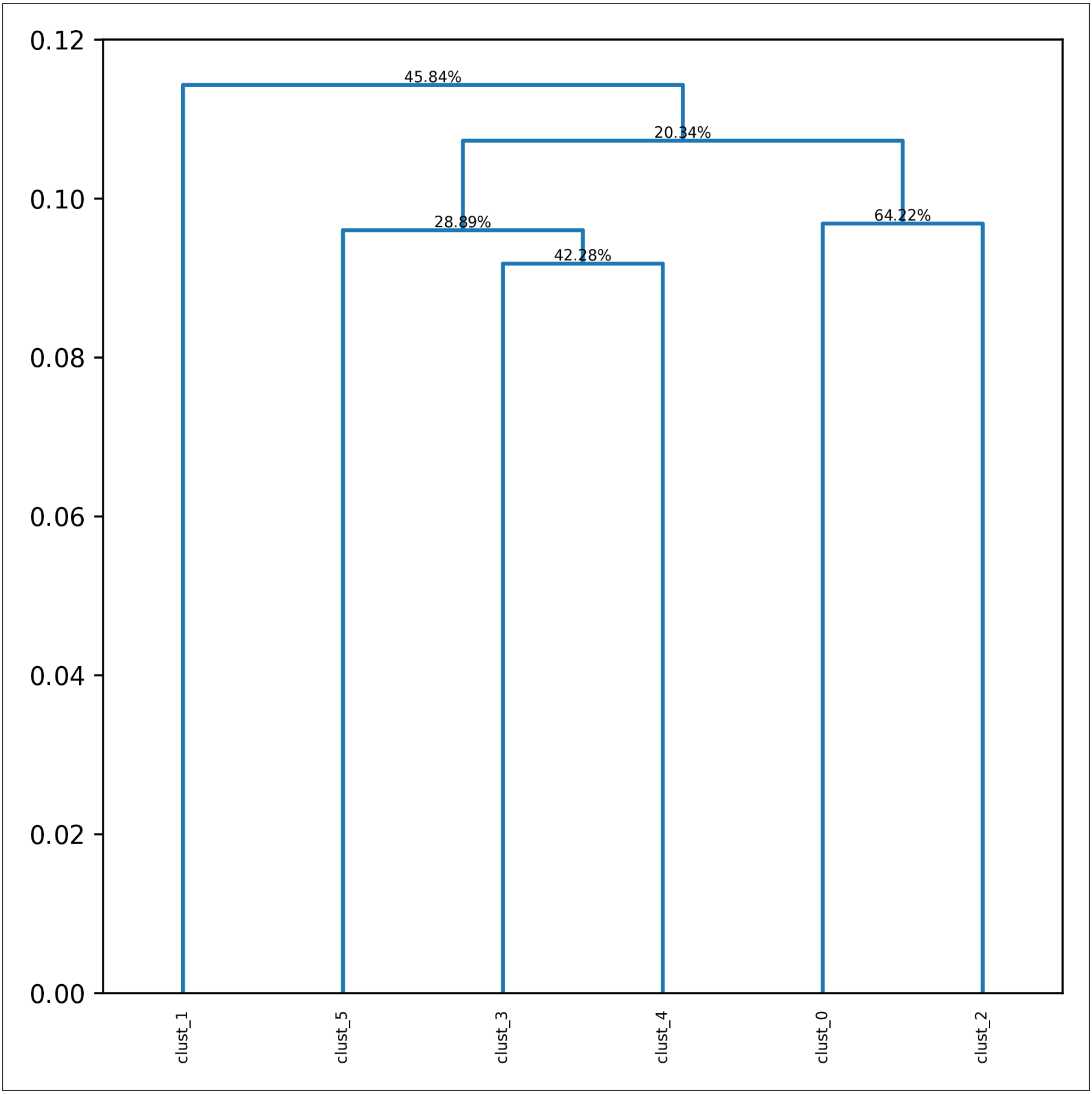

Sample: G0019 with 500 cells per cluster

Min. read depth: 10, Mean UMIs per cluster: 5,049,290

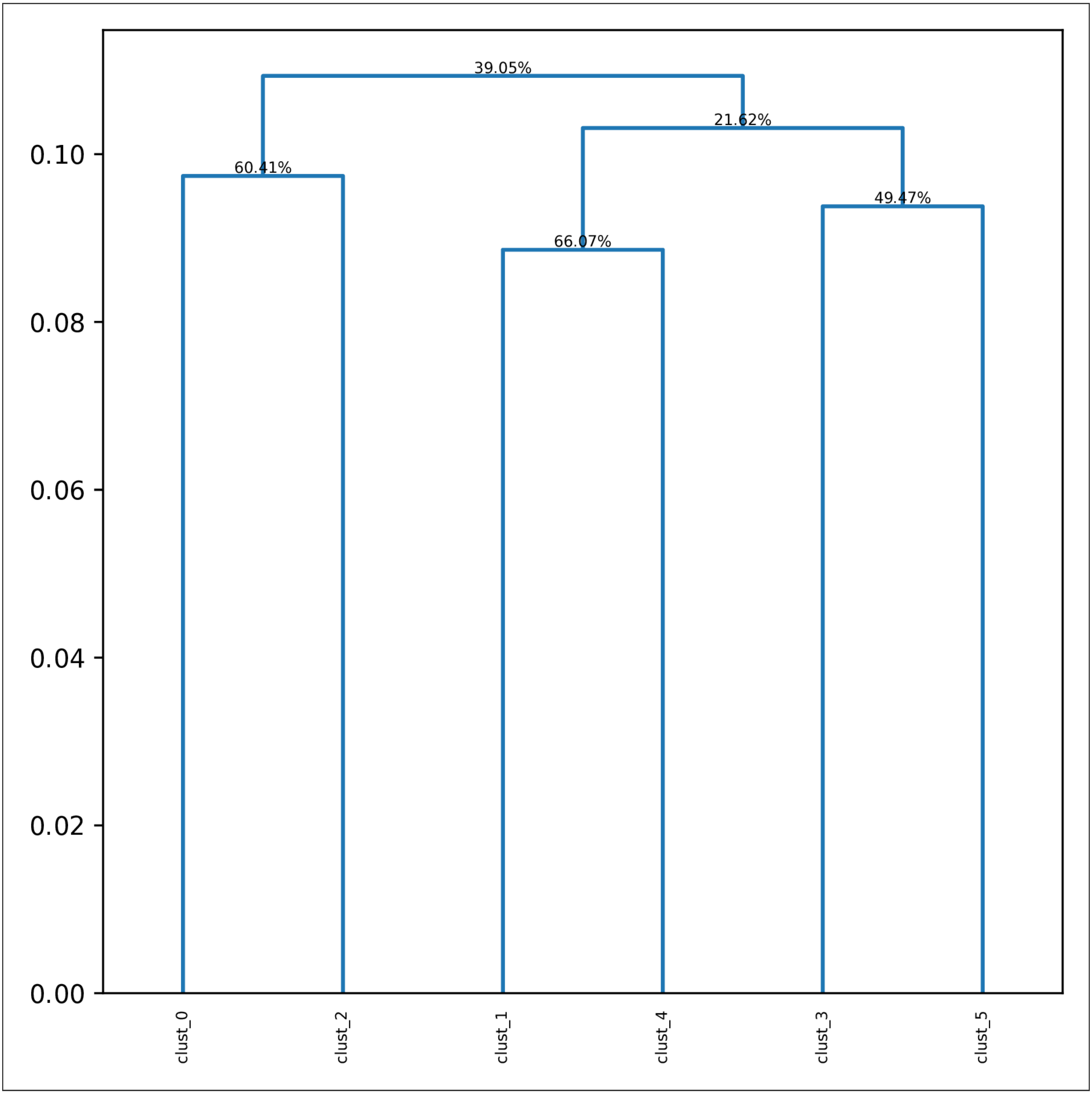

Sample: G0019 with 100 cells per cluster

Min. read depth: 10,  Mean UMIs per cluster:  1,323,846

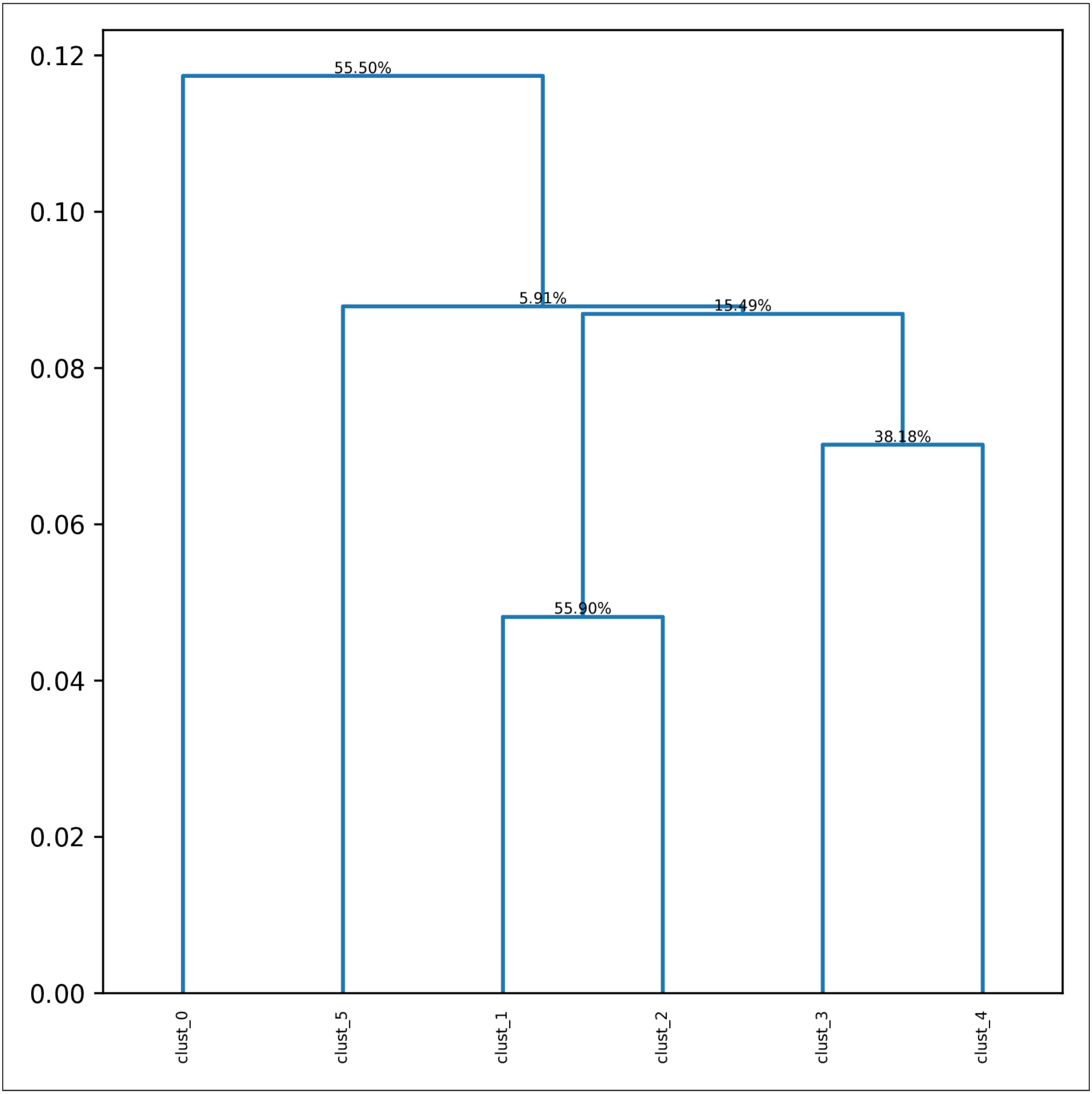

Sample: G0019 with 50 cells per cluster

Min. read depth: 10,  Mean UMIs per cluster:  647,716

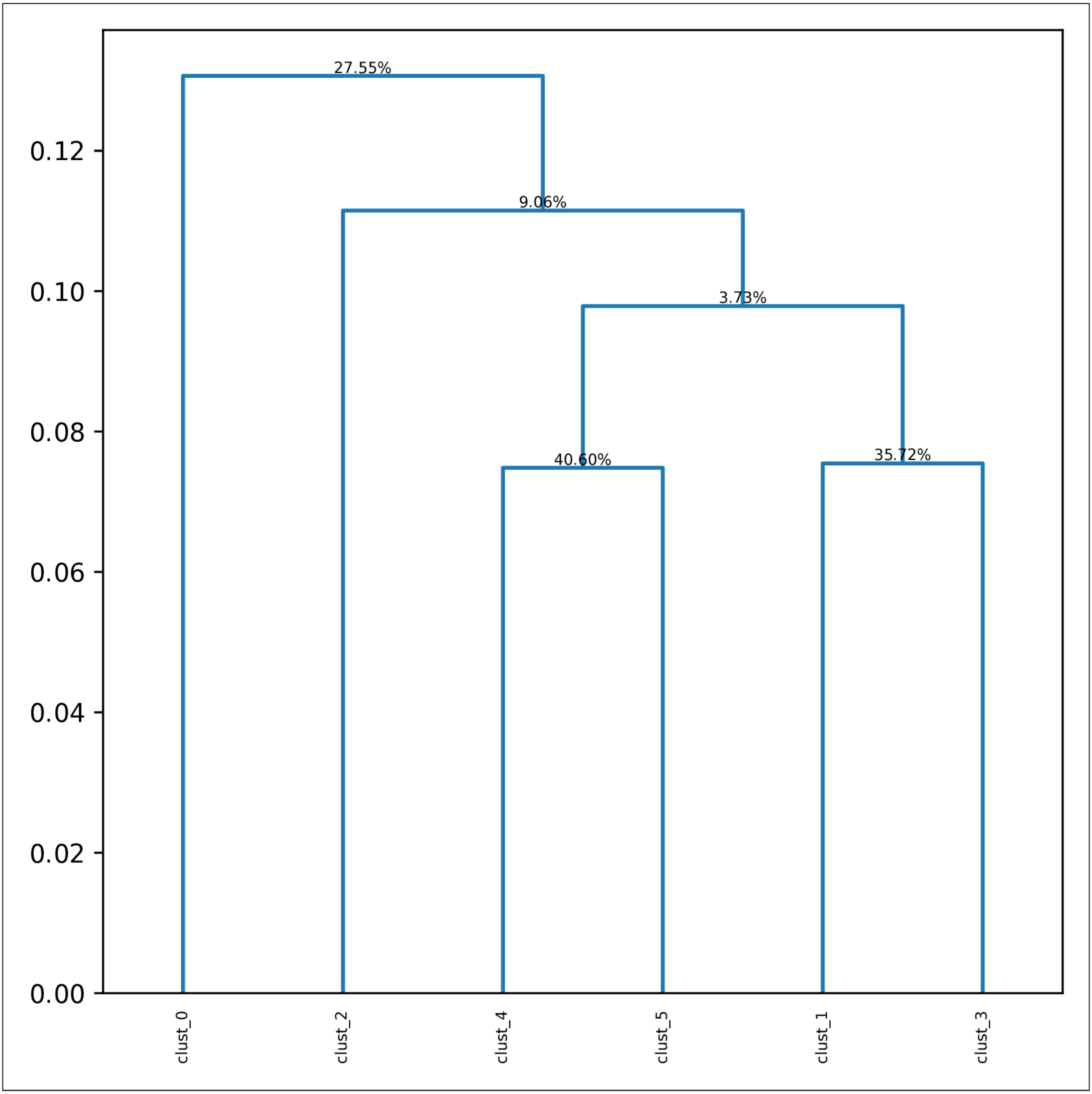

Sample: G0019 with 20 cells per cluster

Min. read depth: 10,  Mean UMIs per cluster:  277,681

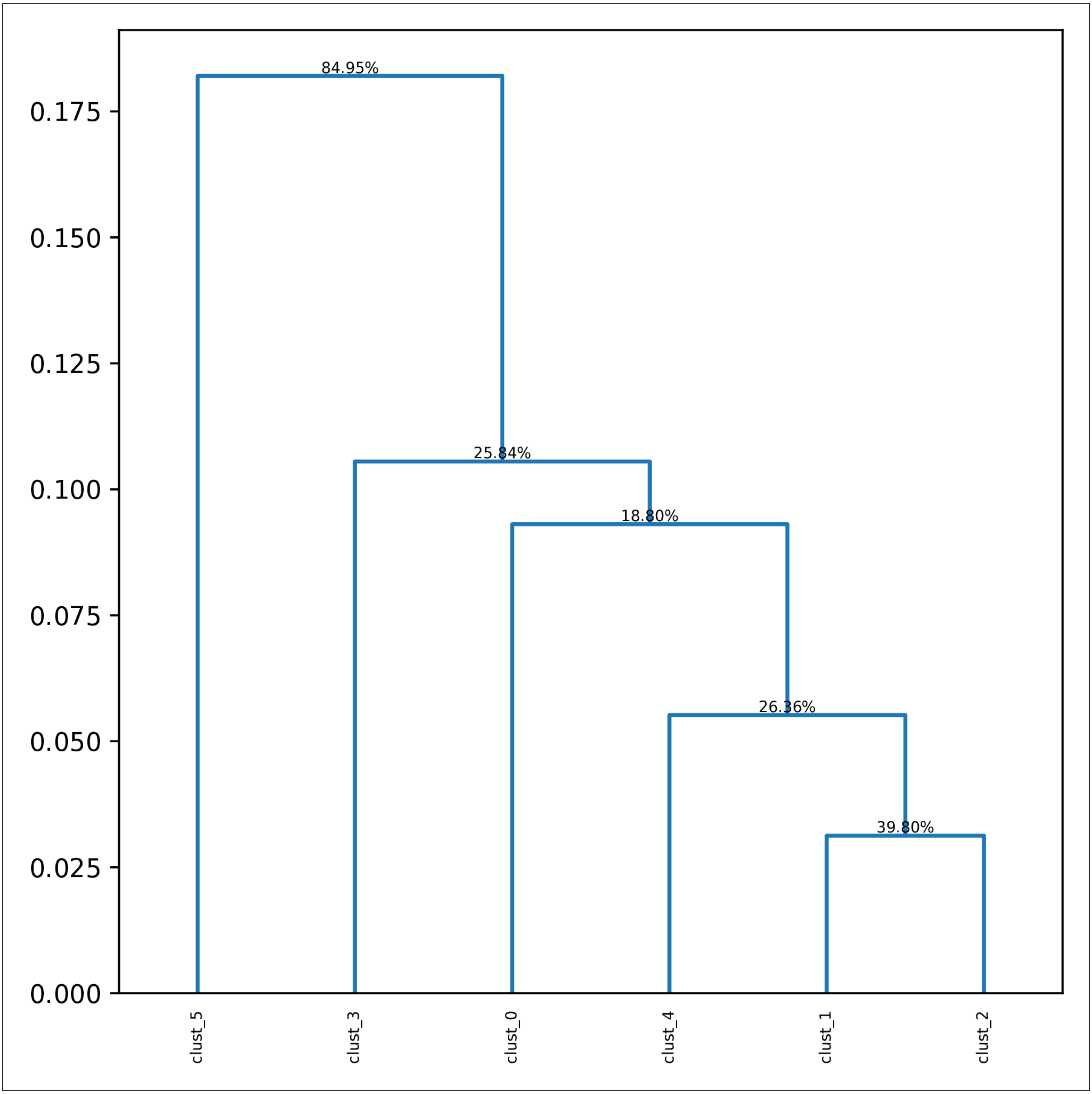

Sample: G0019 with 5 cells per cluster

Min. read depth: 10,  Mean UMIs per cluster:  76,173

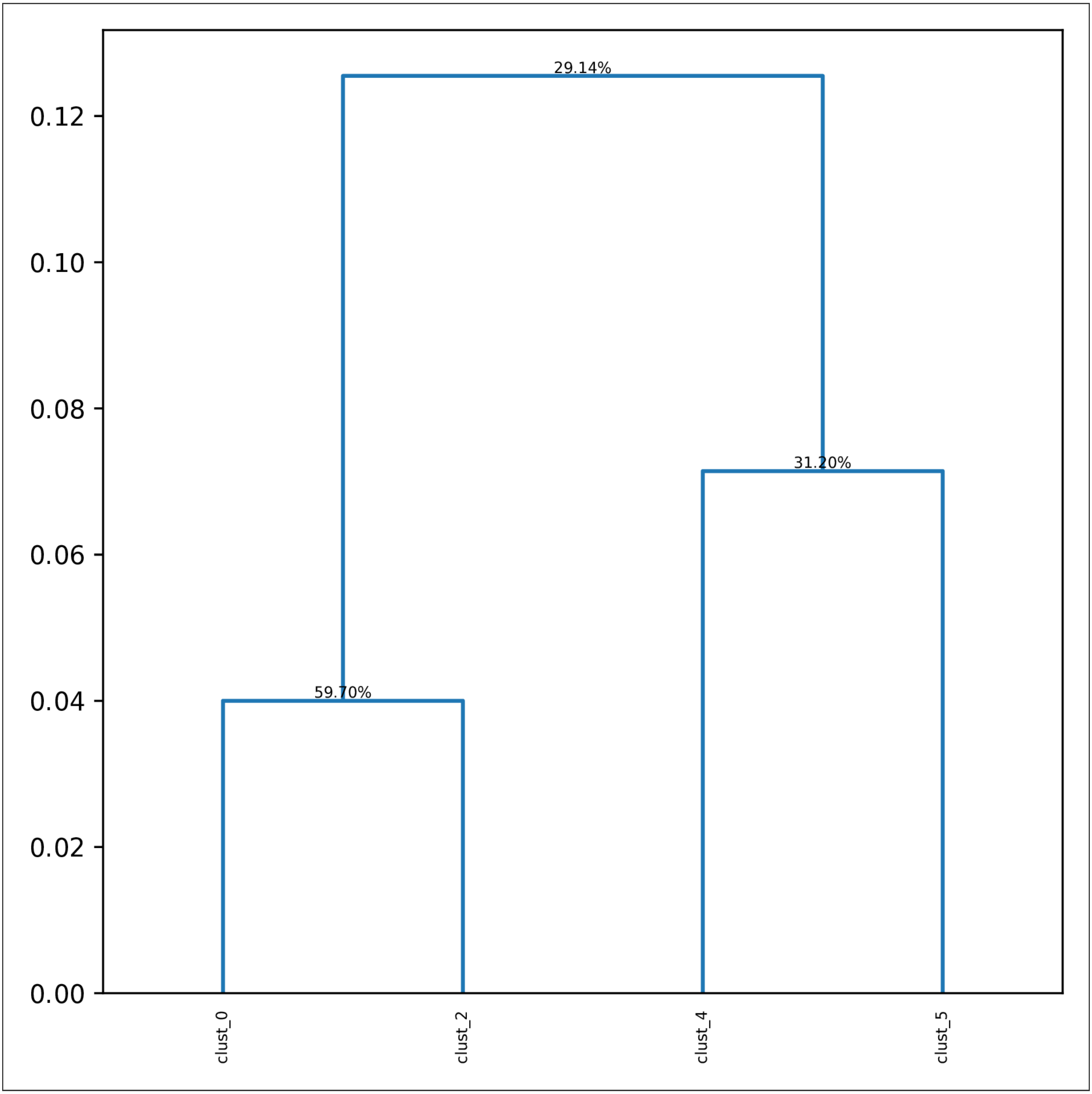

Sample: G0019 with 4 cells per cluster

Min. read depth: 10,  Mean UMIs per cluster:  51,774

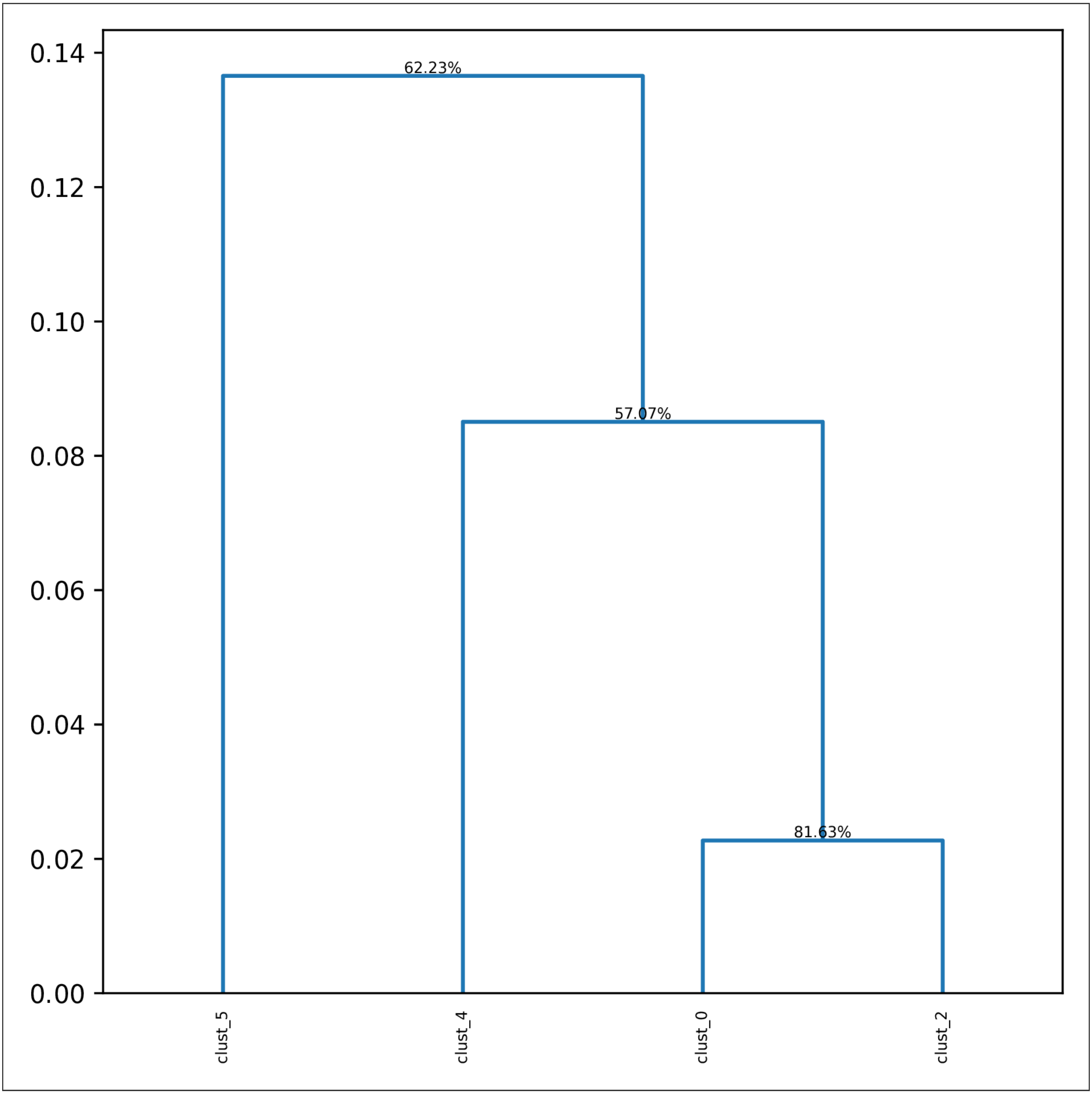

Sample: G0019 with 2 cells per cluster

Min. read depth: 10,  Mean UMIs per cluster:  17,017

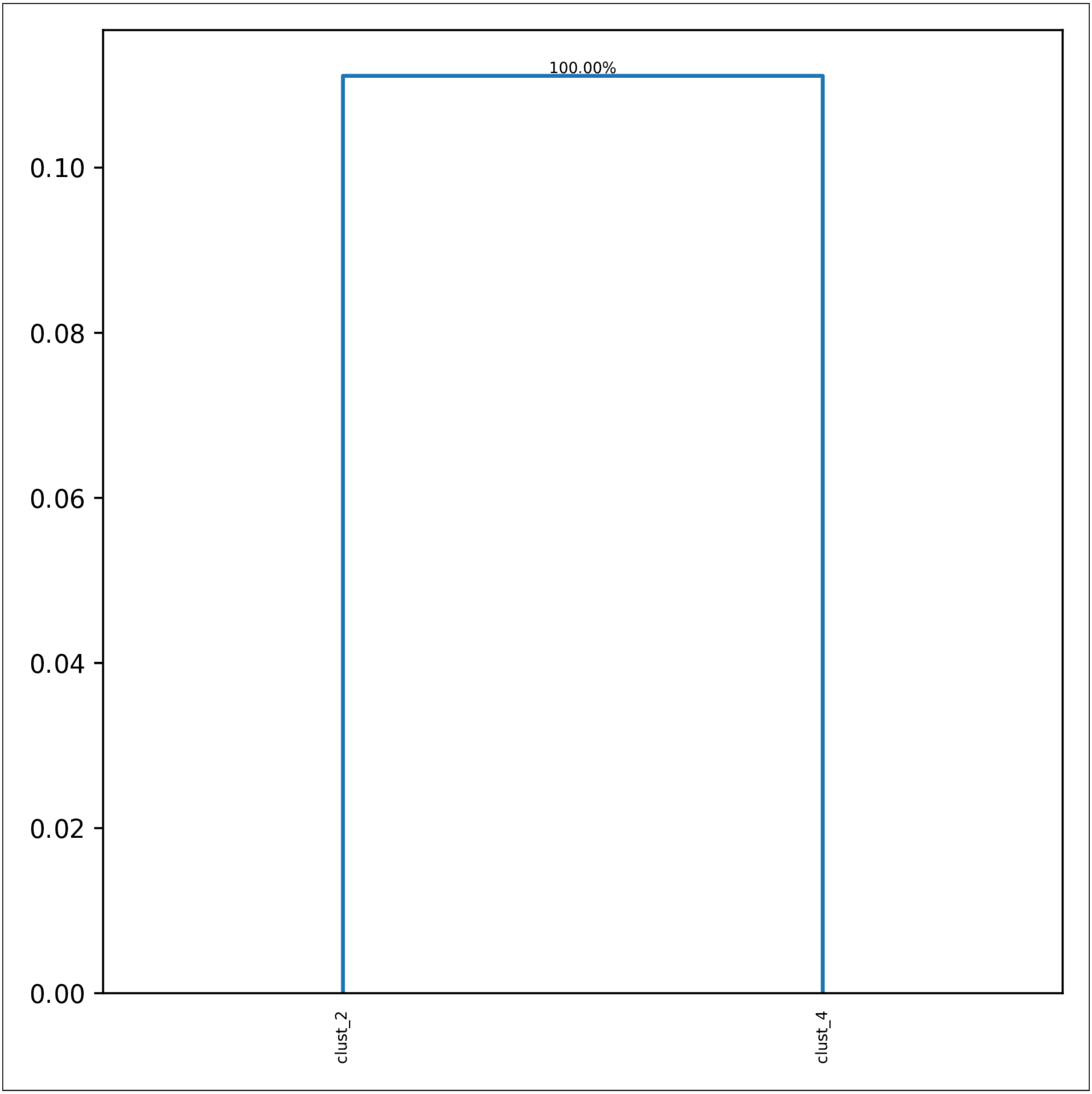

Sample: G0020 with all cells per cluster

Min. read depth: 10,  Mean UMIs per cluster:  7,740,198

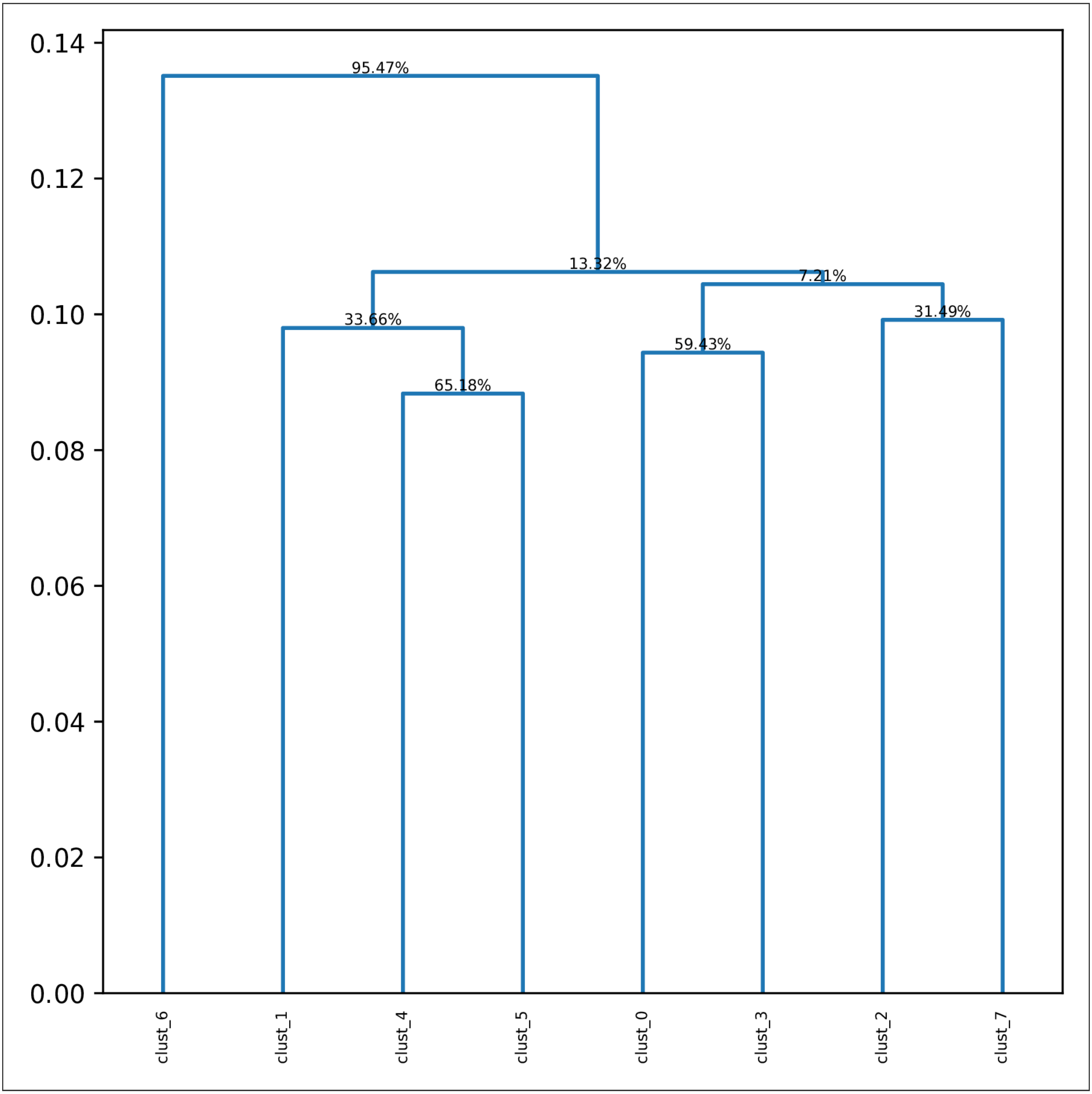

Sample: G0020 with 500 cells per cluster

Min. read depth: 10,  Mean UMIs per cluster:  5,729,651

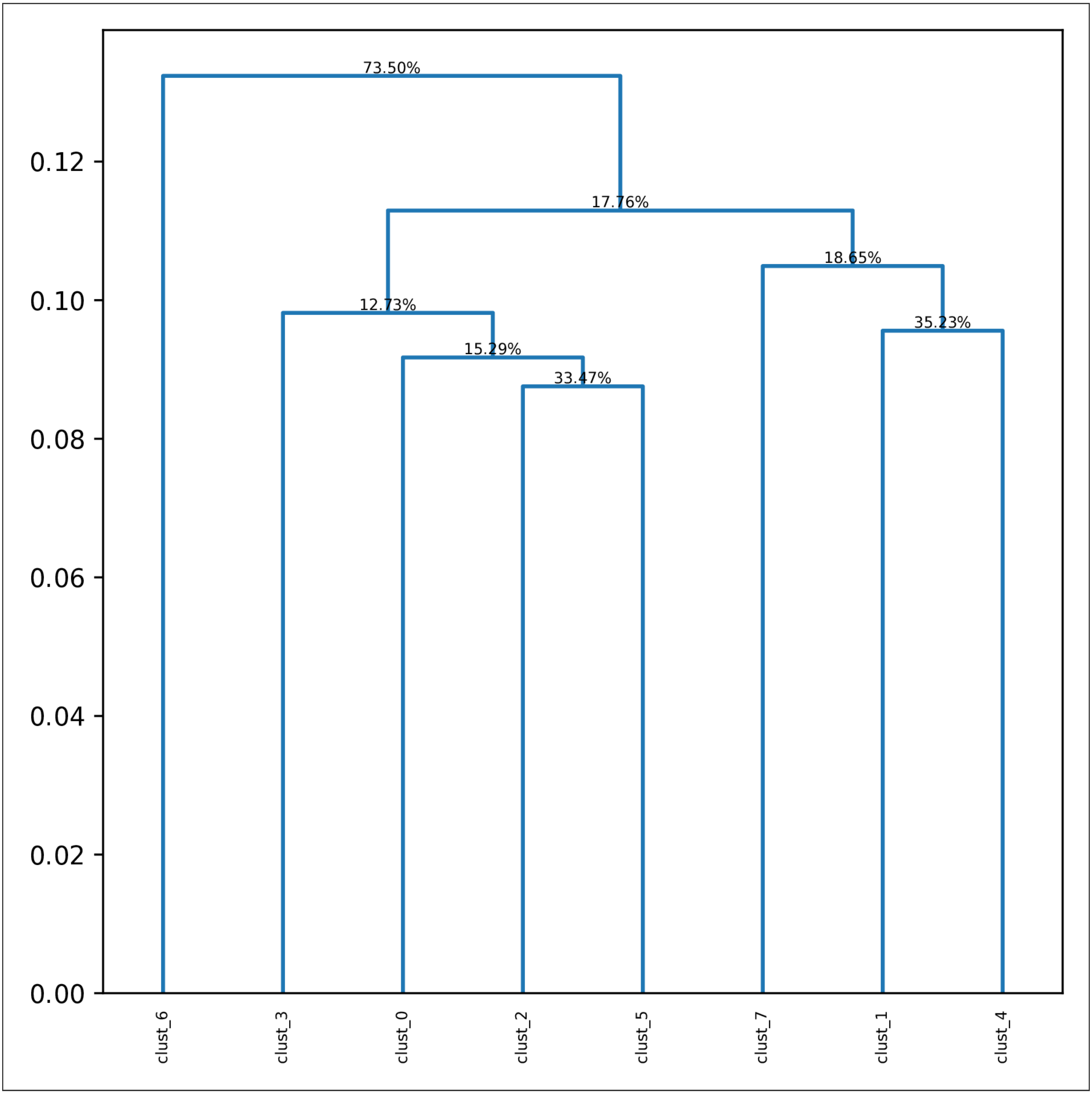

Sample: G0020 with 100 cells per cluster

Min. read depth: 10,  Mean UMIs per cluster:  1,435,312

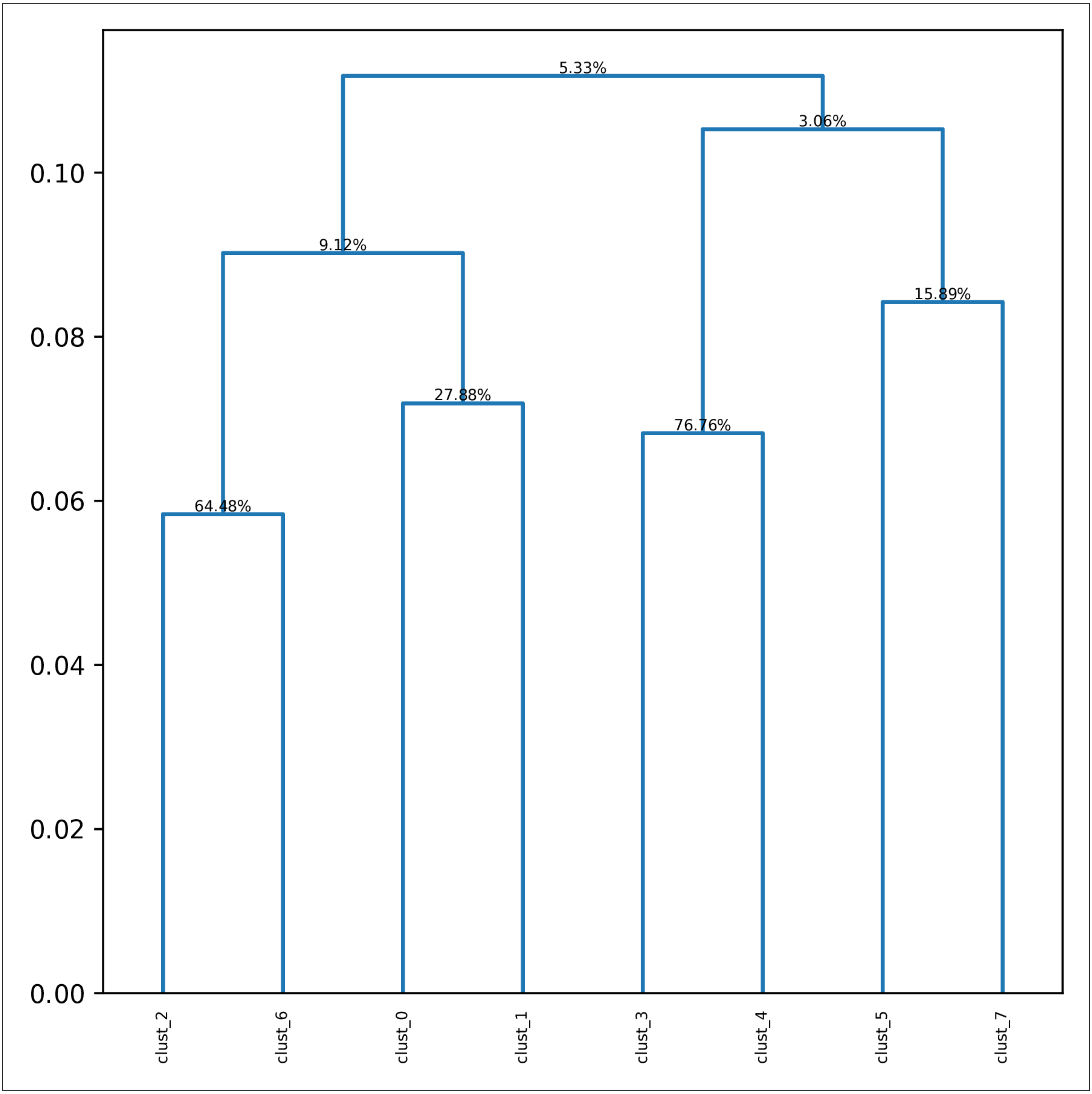

Sample: G0020 with 50 cells per cluster

Min. read depth: 10,  Mean UMIs per cluster:  764,438

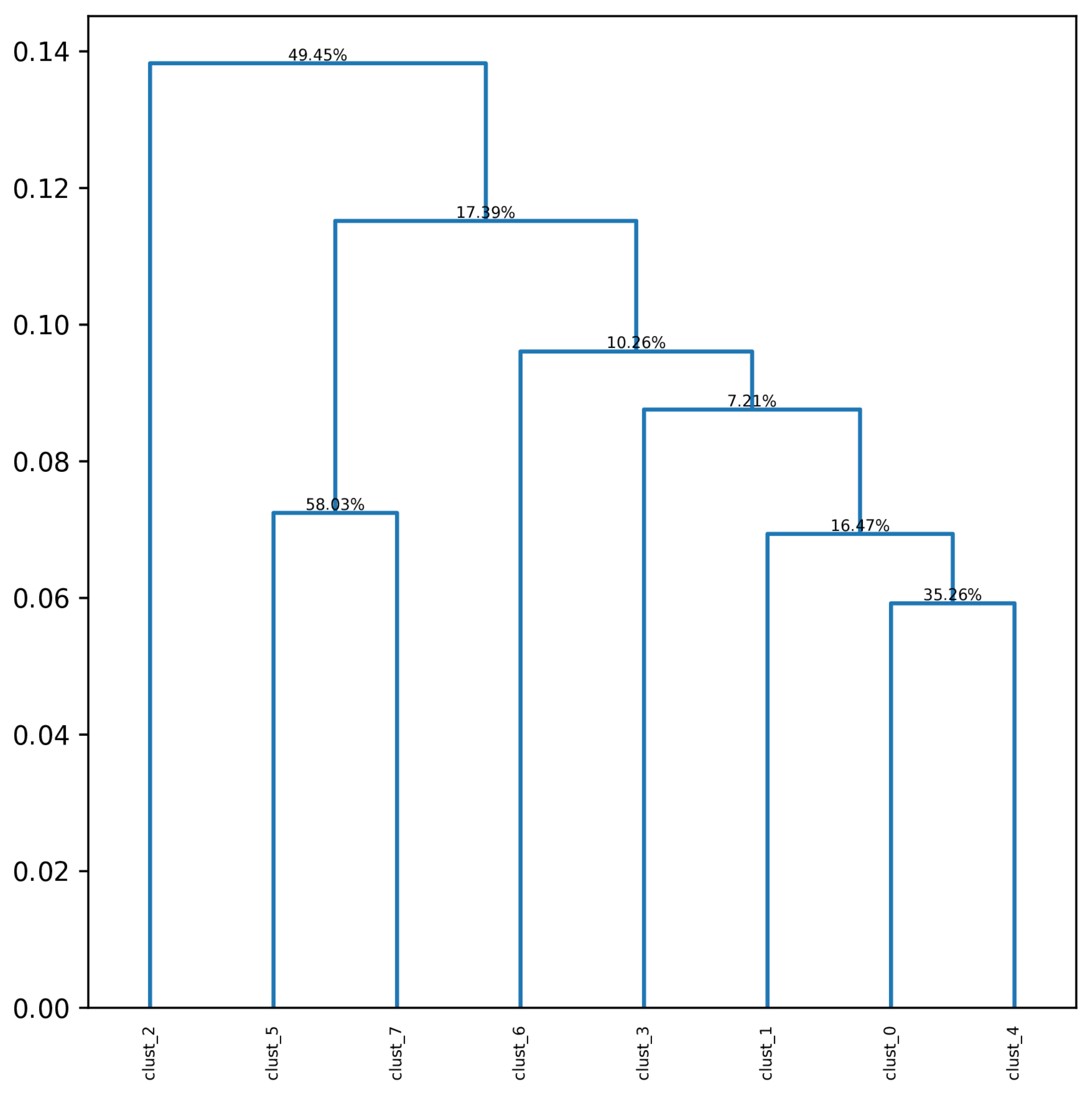

Sample: G0020 with 20 cells per cluster

Min. read depth: 10,  Mean UMIs per cluster:  284,078

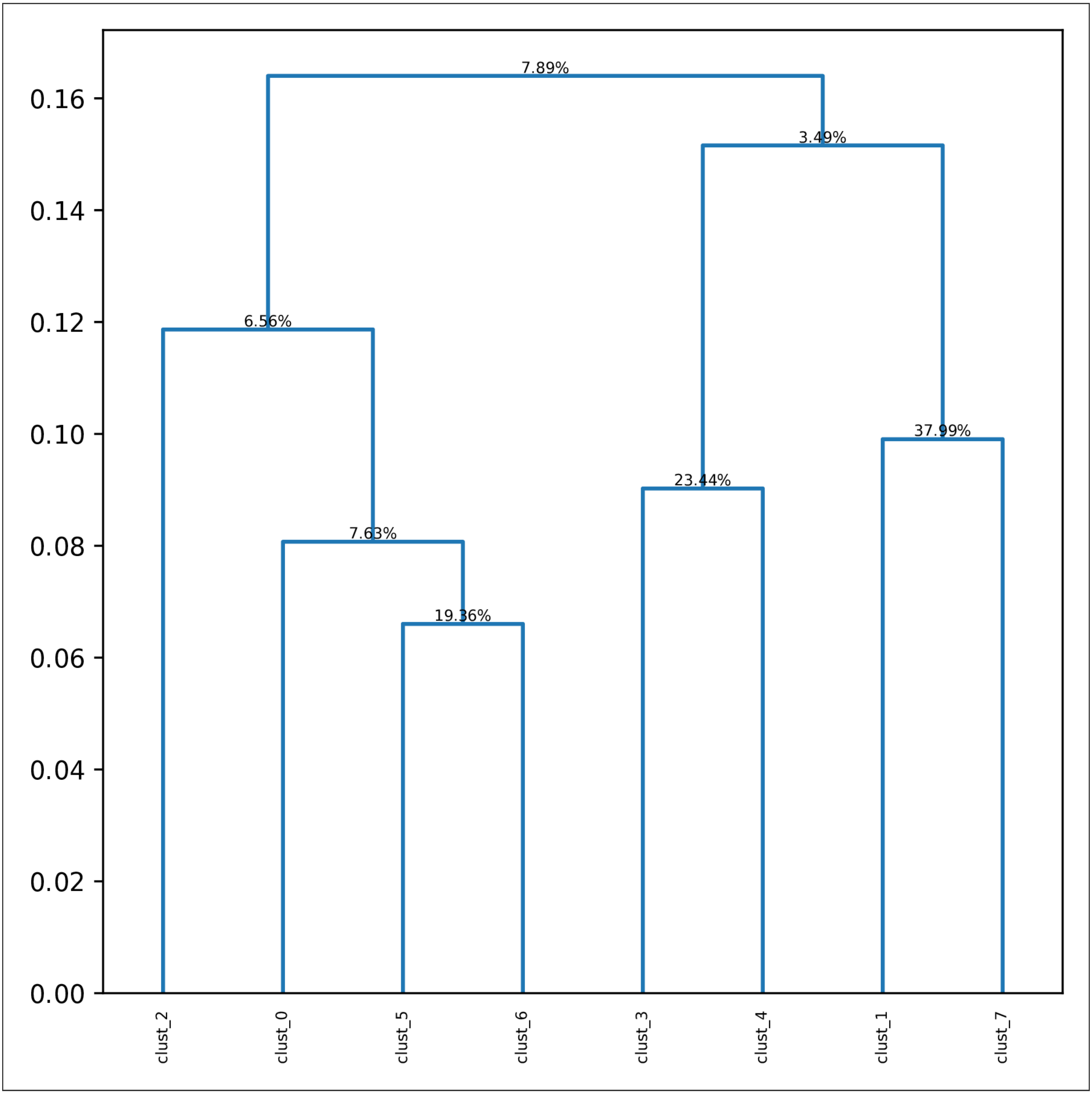

Sample: G0020 with 10 cells per cluster

Min. read depth: 10,  Mean UMIs per cluster:  152,599

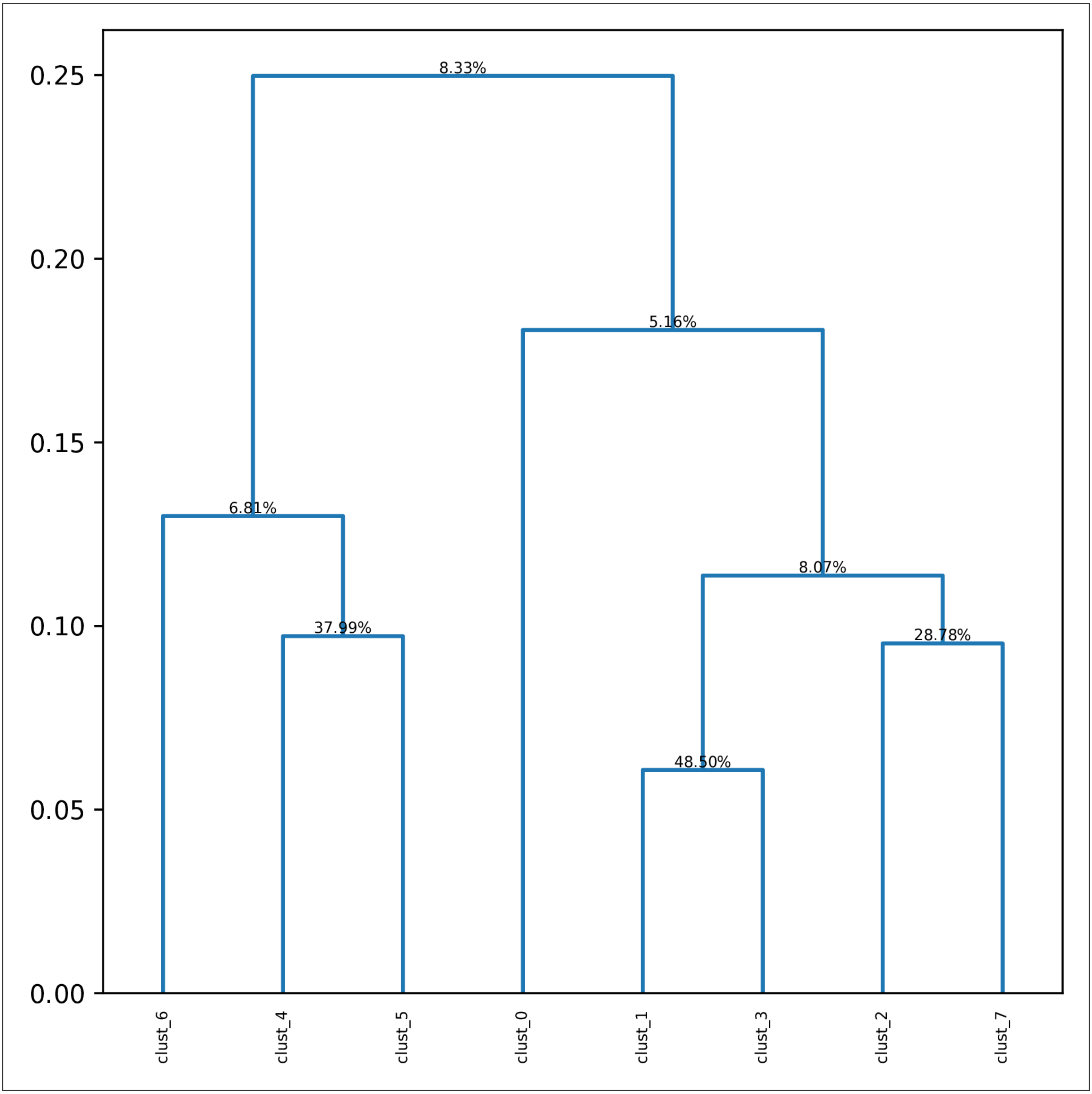

Sample: G0021 with all cells per cluster

Min. read depth: 10,  Mean UMIs per cluster:  4,582,536

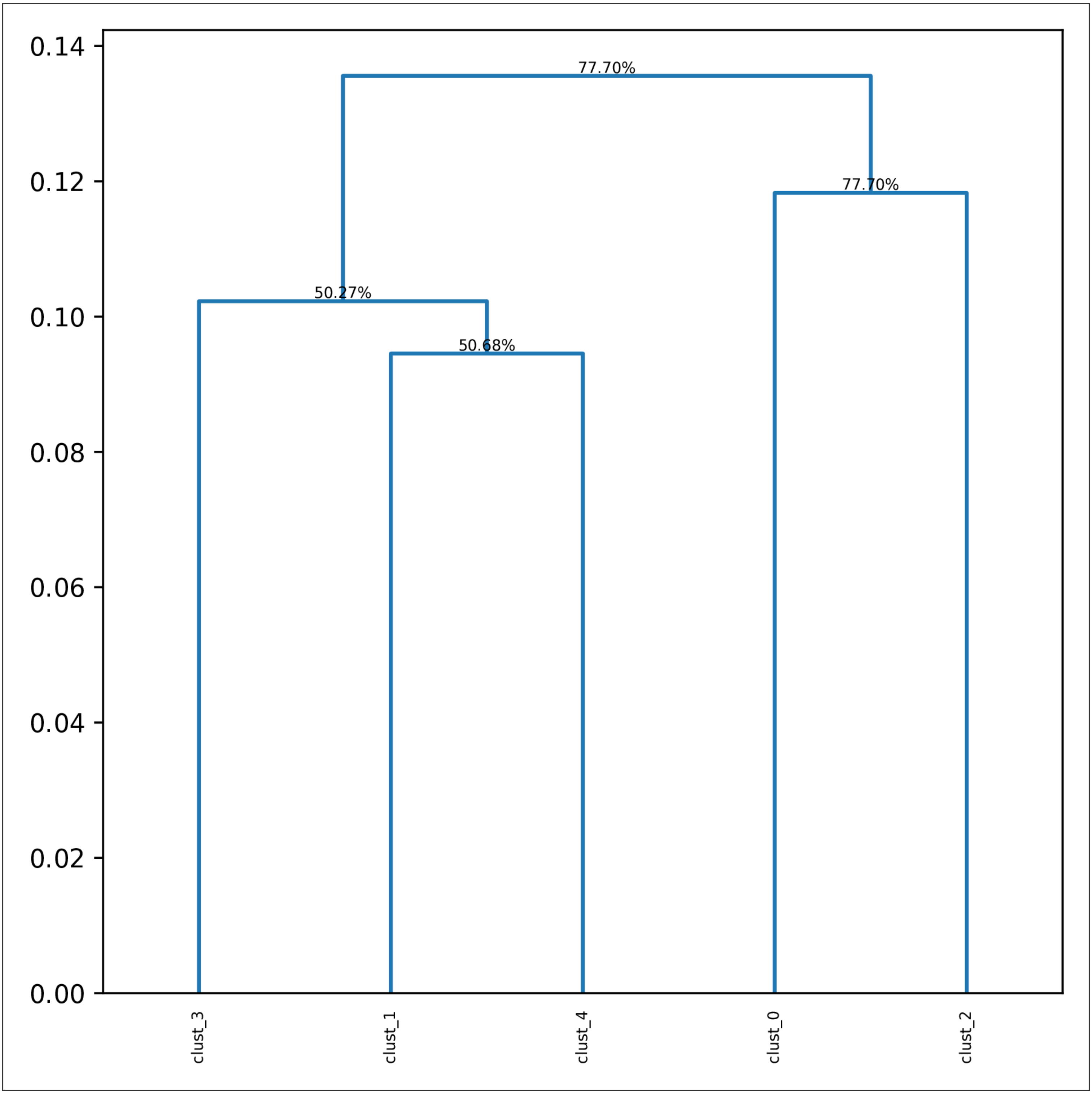

Sample: G0021 with 500 cells per cluster

Min. read depth: 10,  Mean UMIs per cluster:  3,955,092

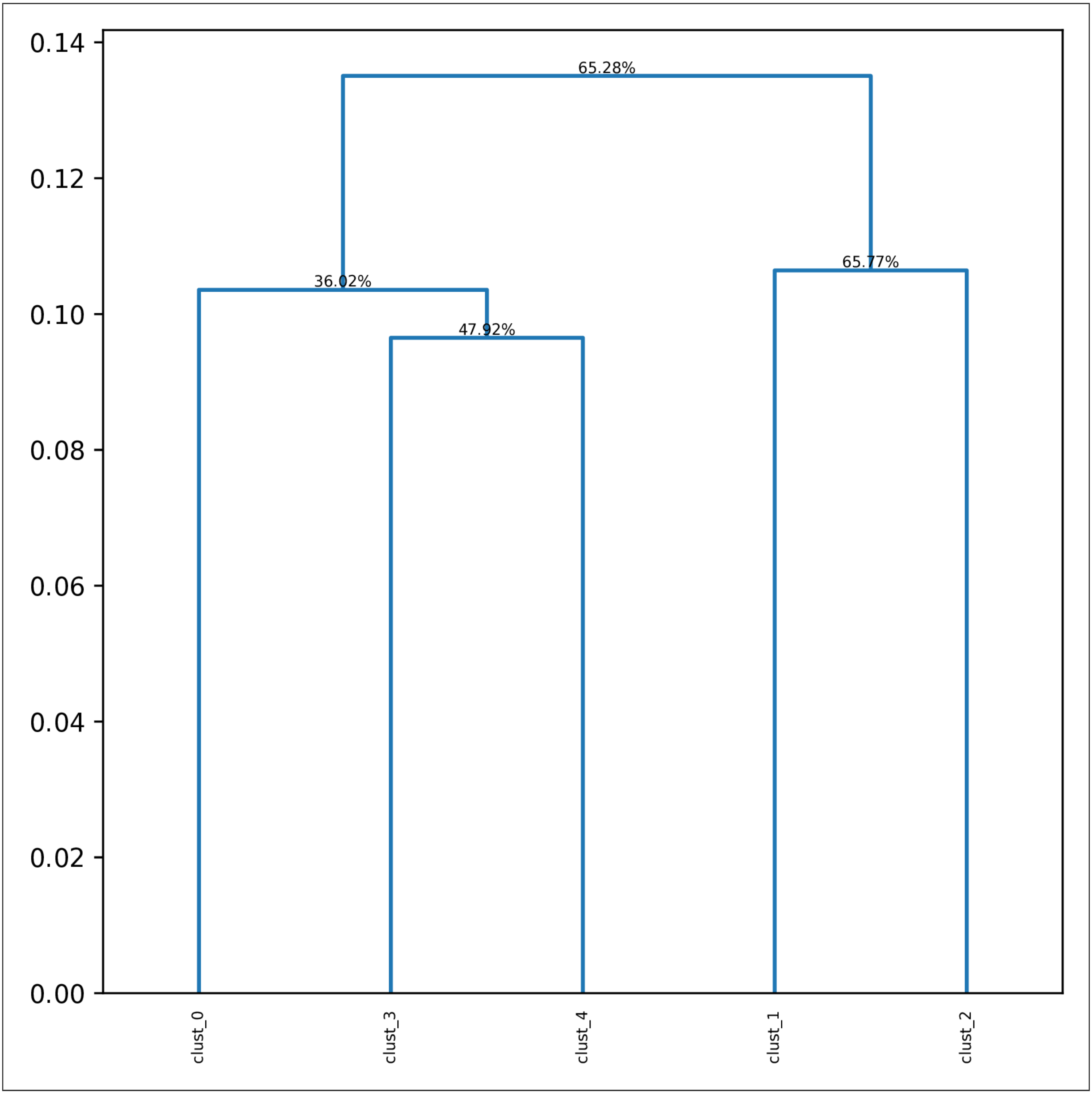

Sample: G0021 with 100 cells per cluster

Min. read depth: 10,  Mean UMIs per cluster:  1,175,811

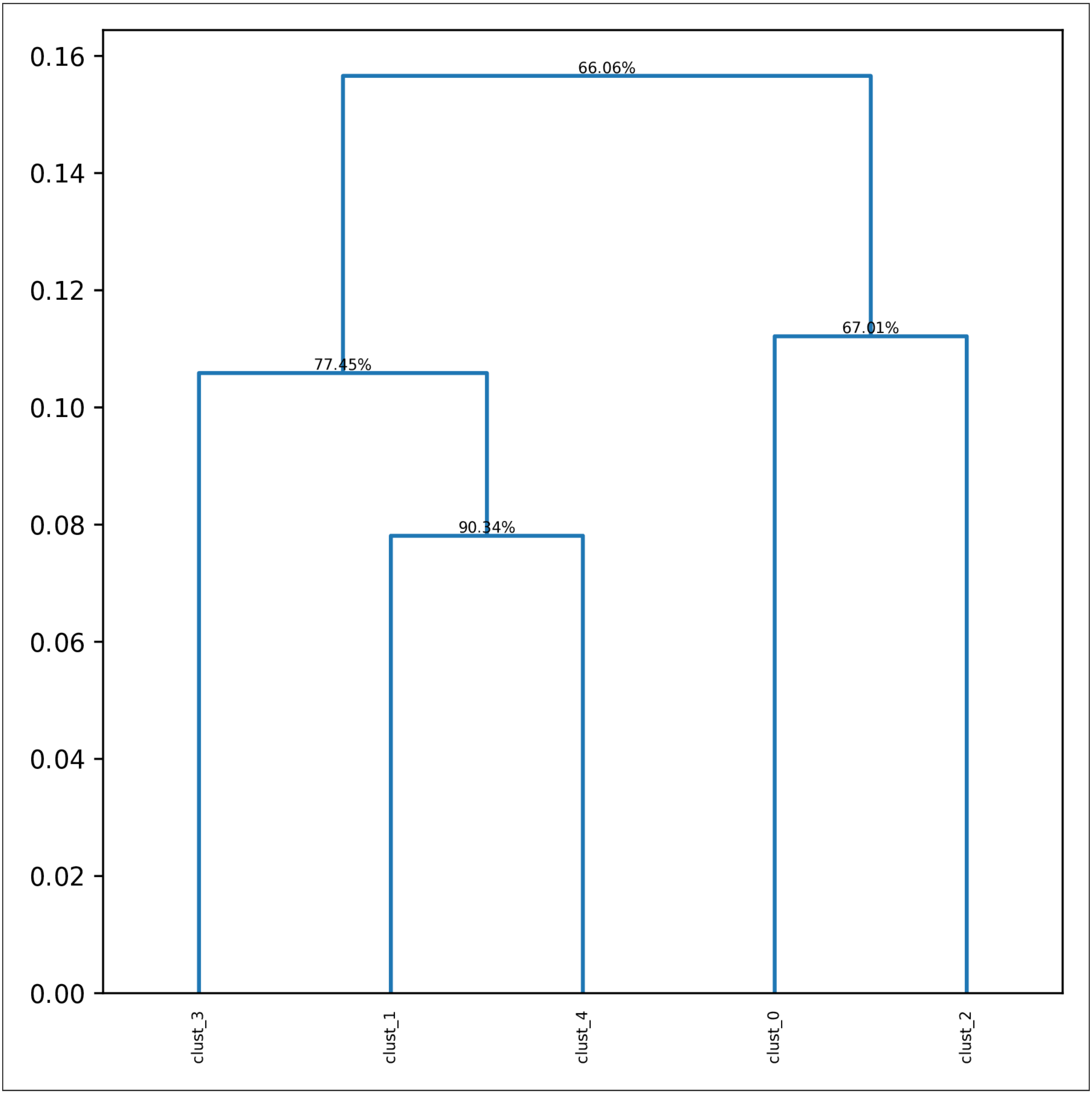

Sample: G0021 with 50 cells per cluster

Min. read depth: 10,  Mean UMIs per cluster:  562,953

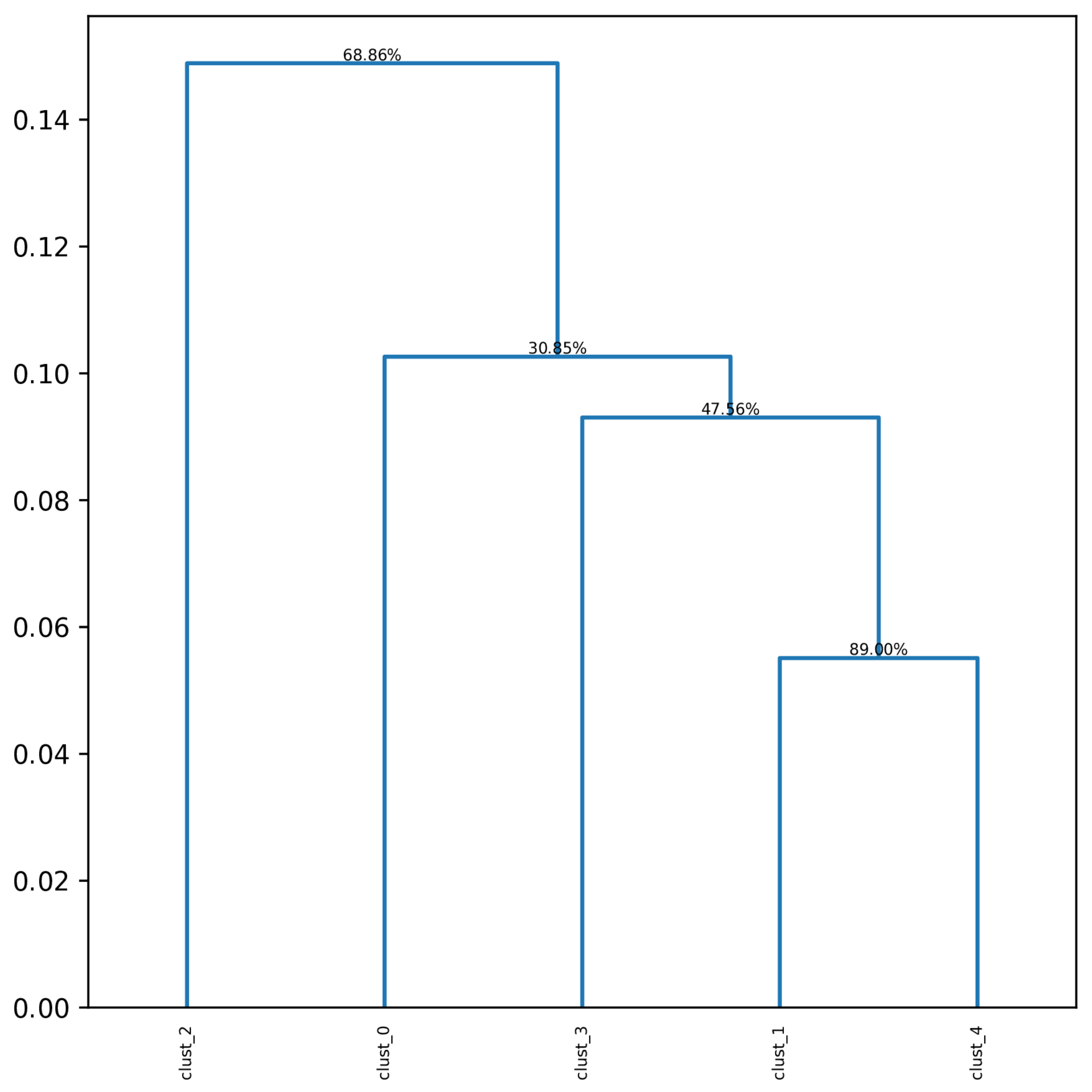

Sample: G0021 with 20 cells per cluster

Min. read depth: 10,  Mean UMIs per cluster:  225,011

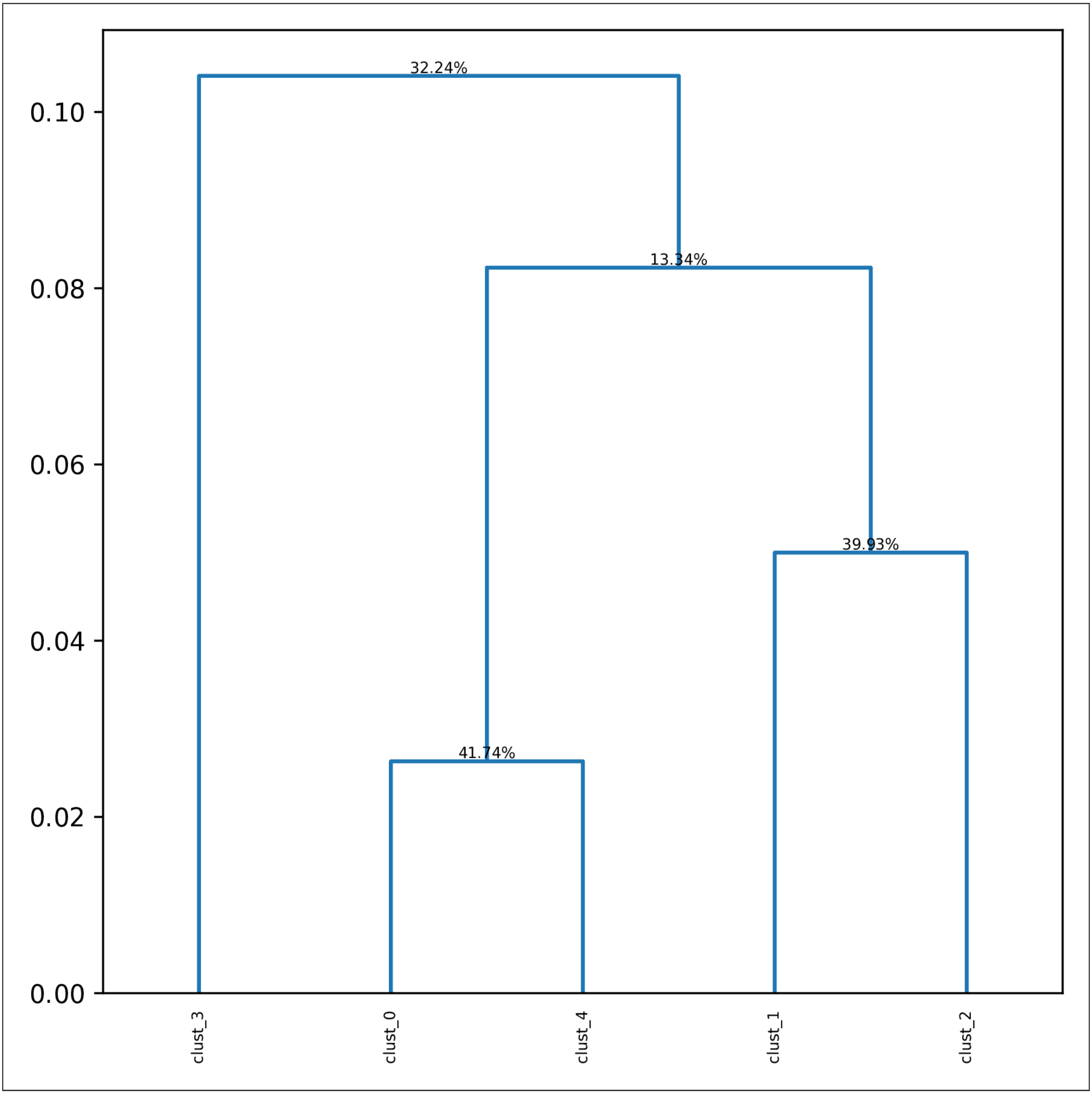

Sample: G0021 with 3 cells per cluster

Min. read depth: 10,  Mean UMIs per cluster:  36,841

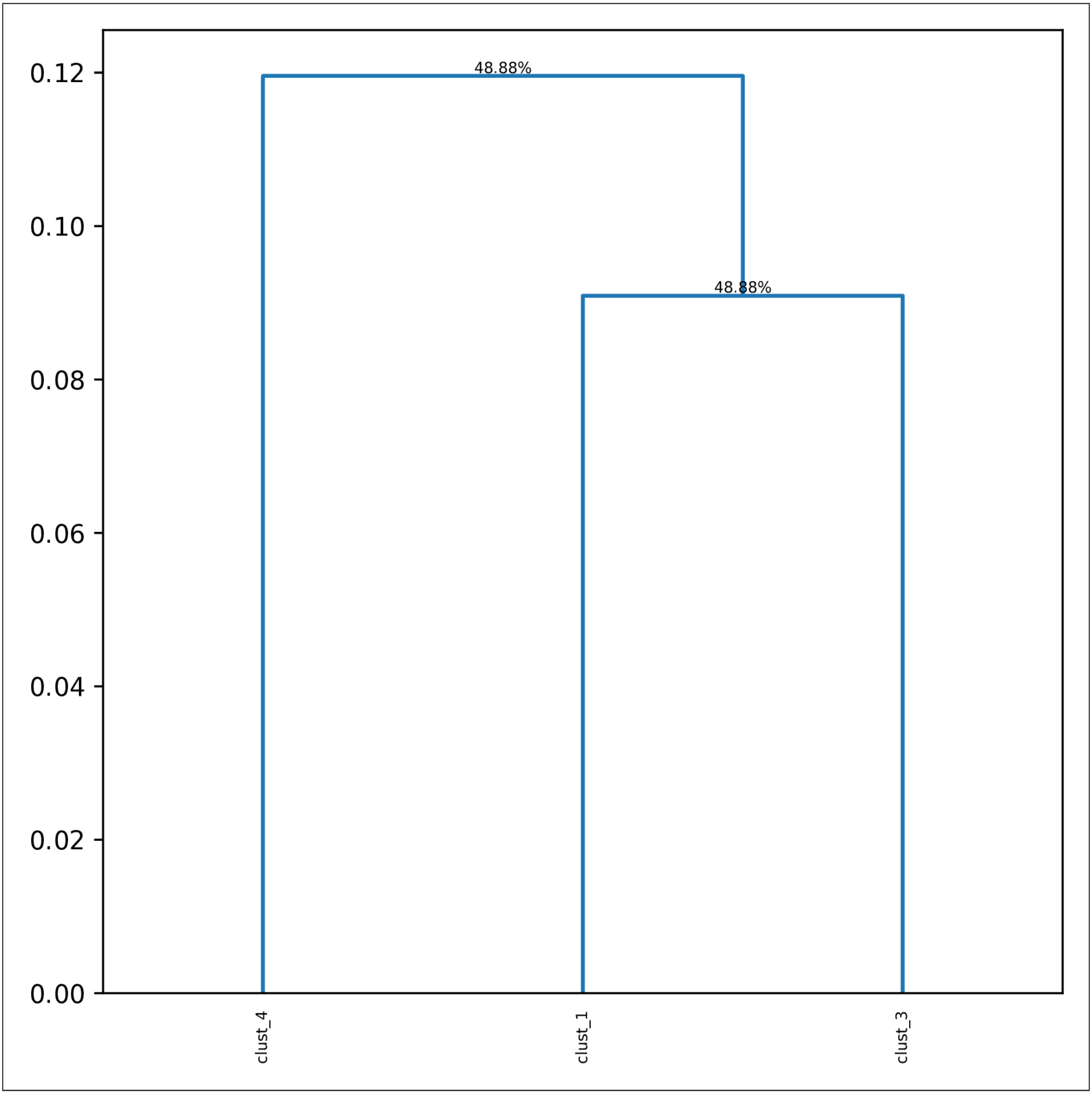

Sample: G0021 with 2 cells per cluster

Min. read depth: 10,  Mean UMIs per cluster:  27,156

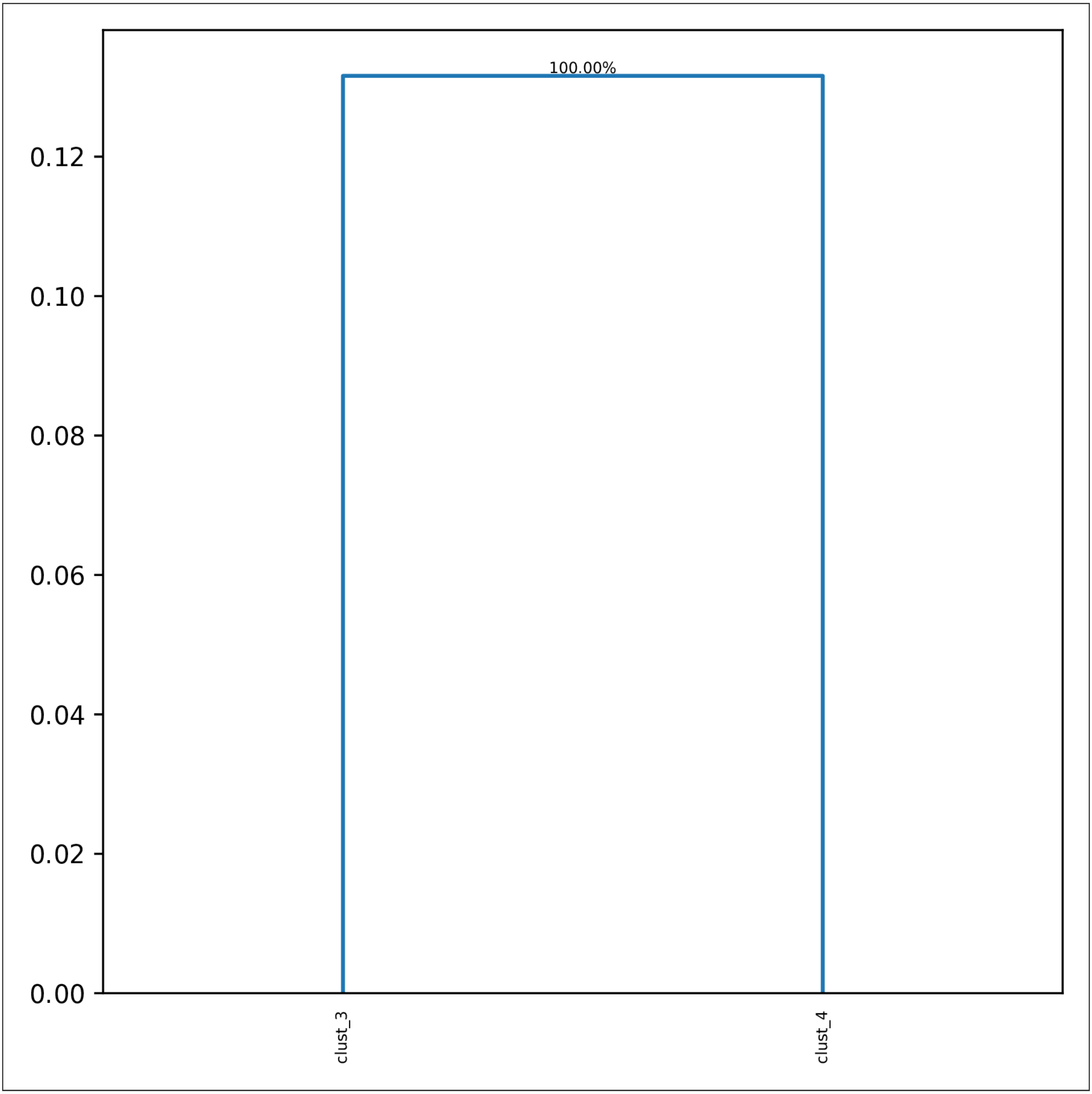

Sample: G0022 with all cells per cluster

Min. read depth: 10,  Mean UMIs per cluster:  6,163,754

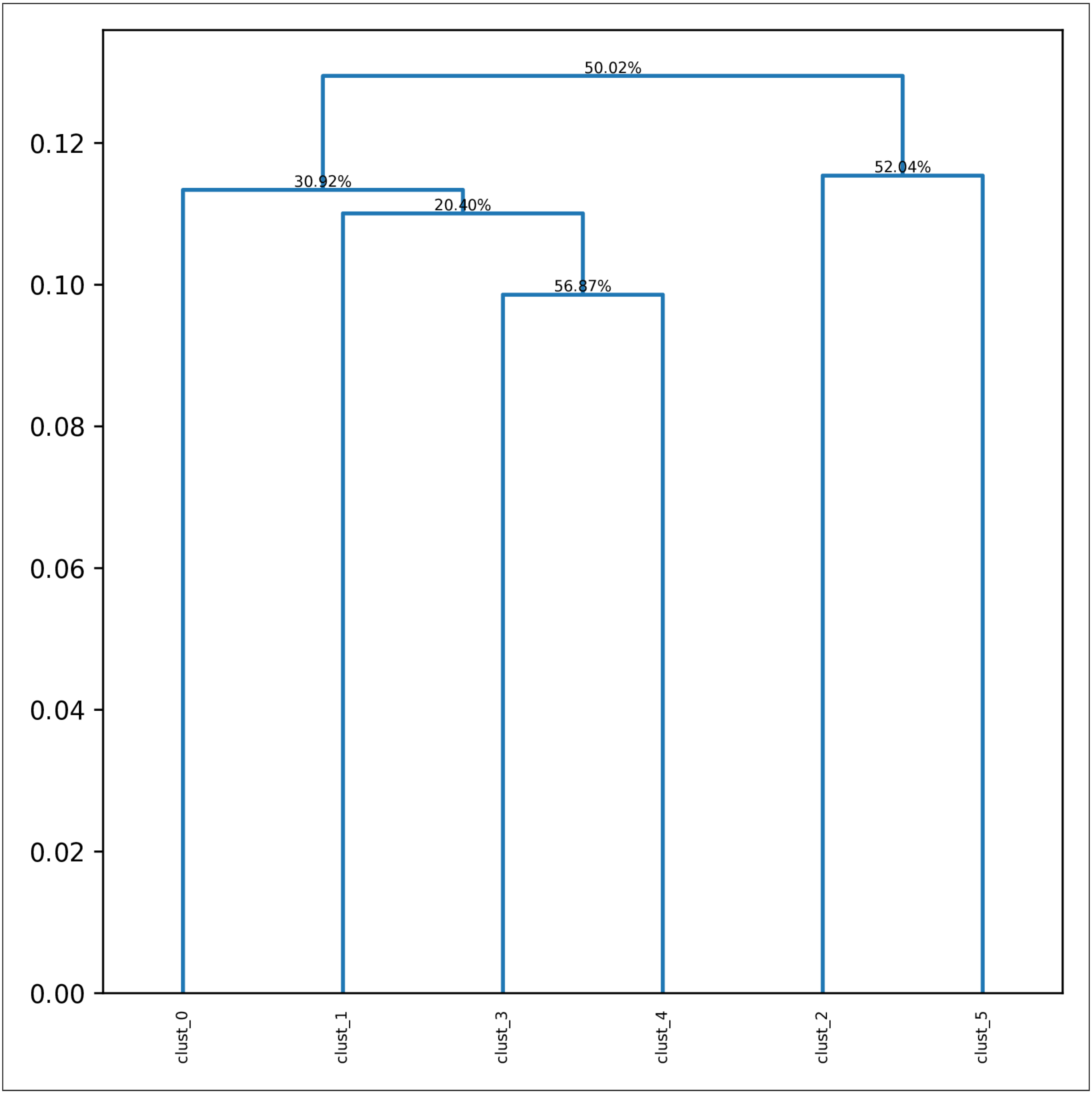

Sample: G0022 with 500 cells per cluster

Min. read depth: 10,  Mean UMIs per cluster:  3,919,936

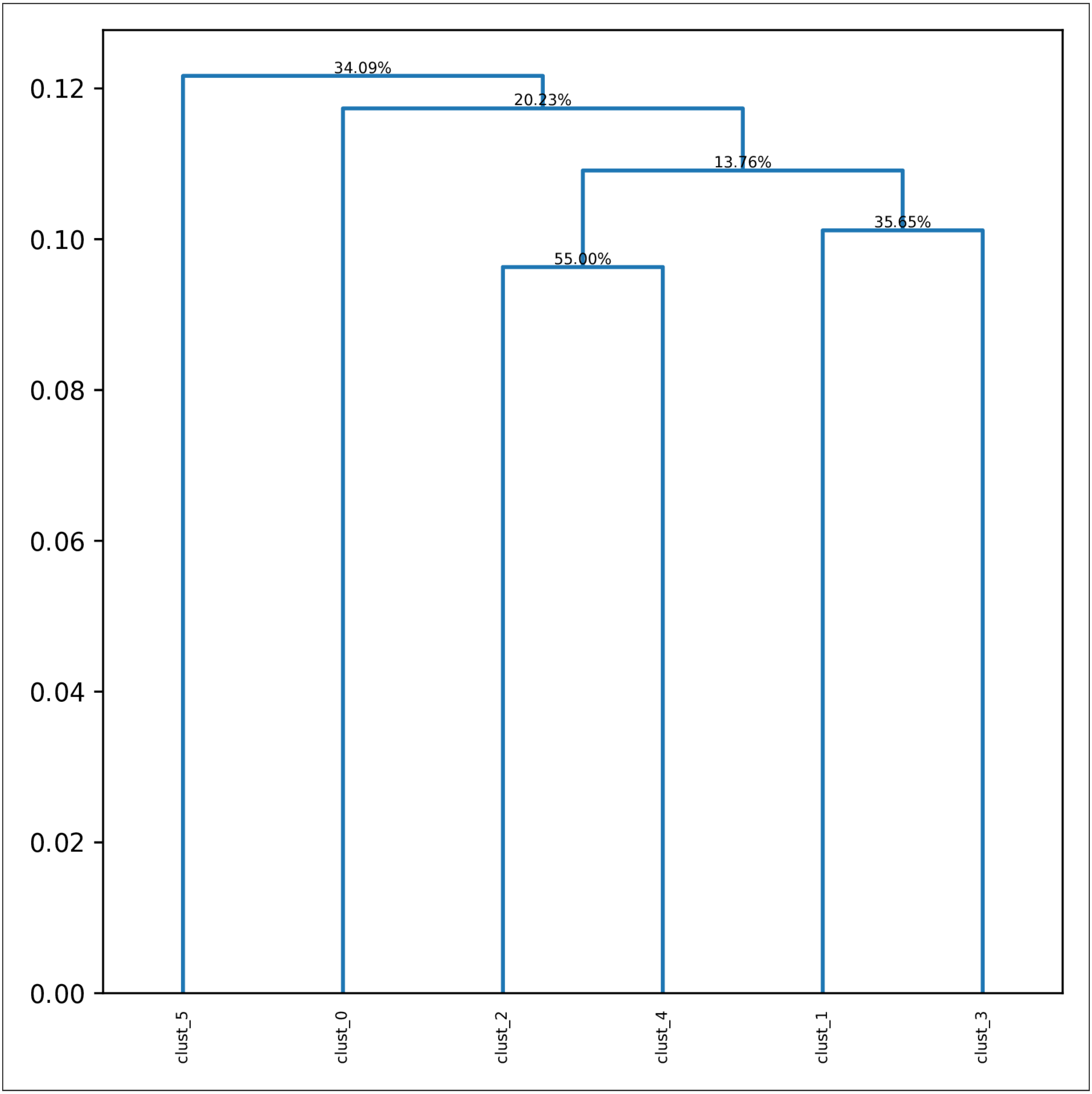

Sample: G0022 with 100 cells per cluster

Min. read depth: 10,  Mean UMIs per cluster:  1,138,864

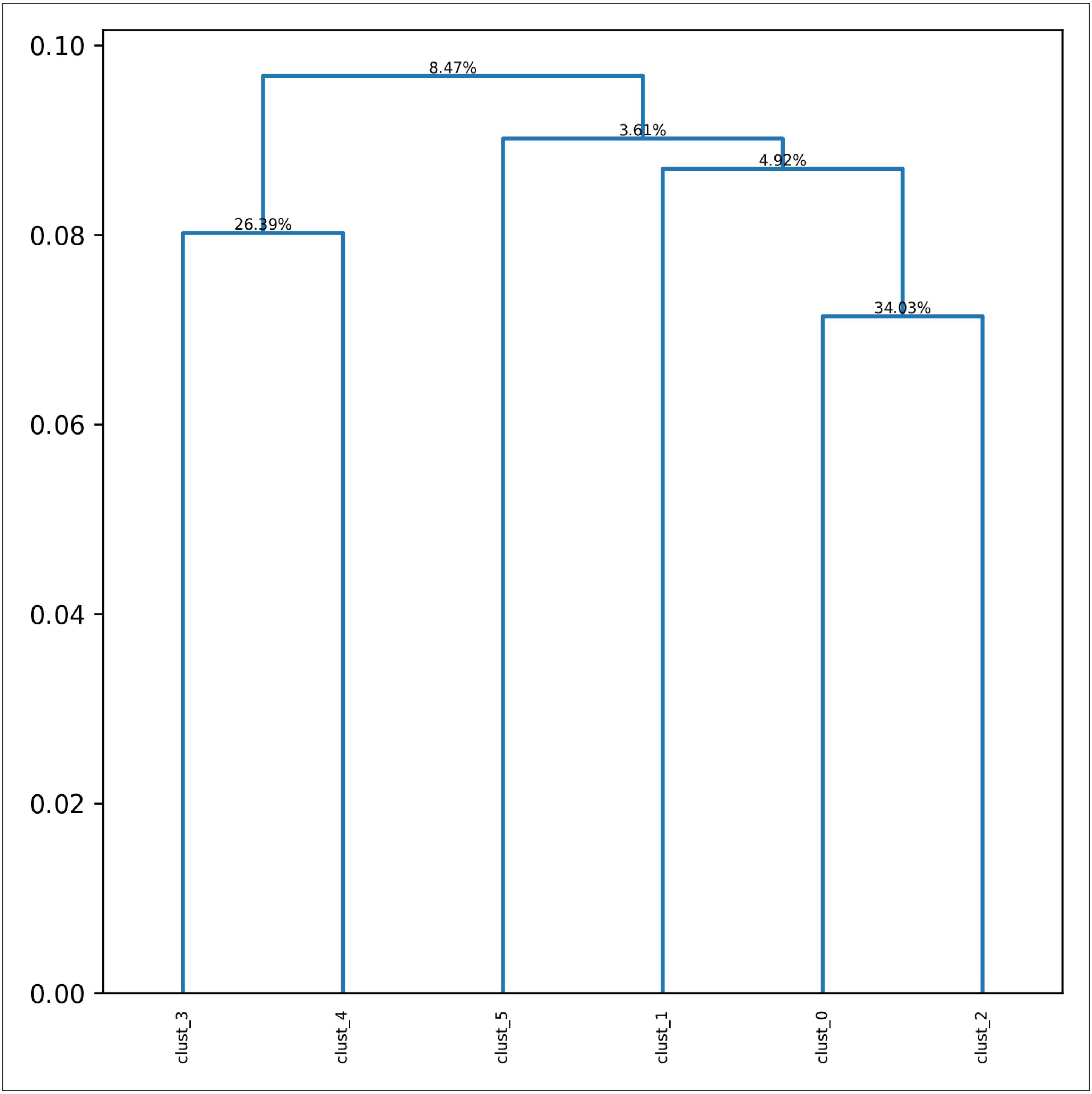

Sample: G0022 with 50 cells per cluster

Min. read depth: 10,  Mean UMIs per cluster:  604,142

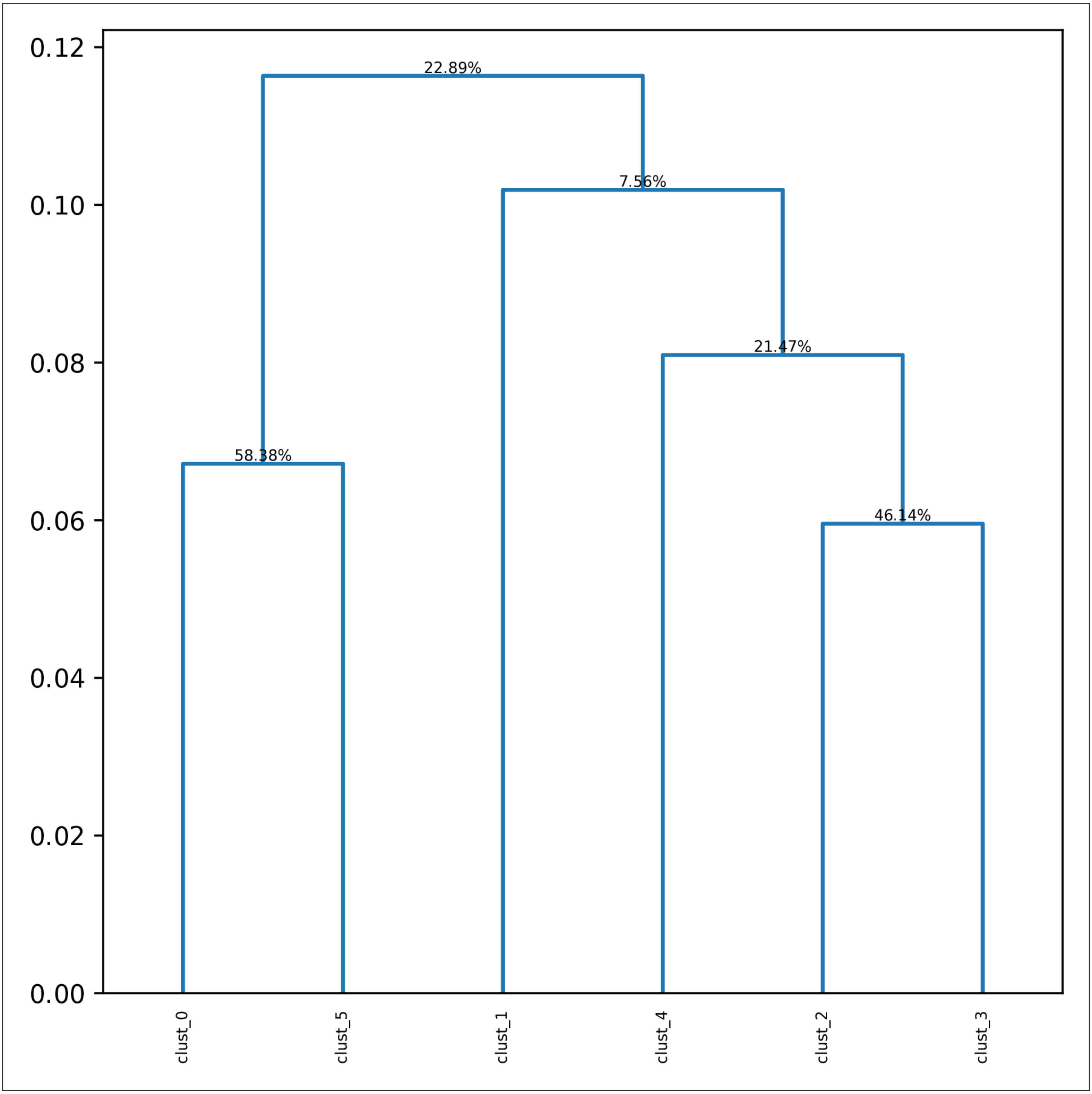

Sample: G0022 with 20 cells per cluster

Min. read depth: 10,  Mean UMIs per cluster:  233,893

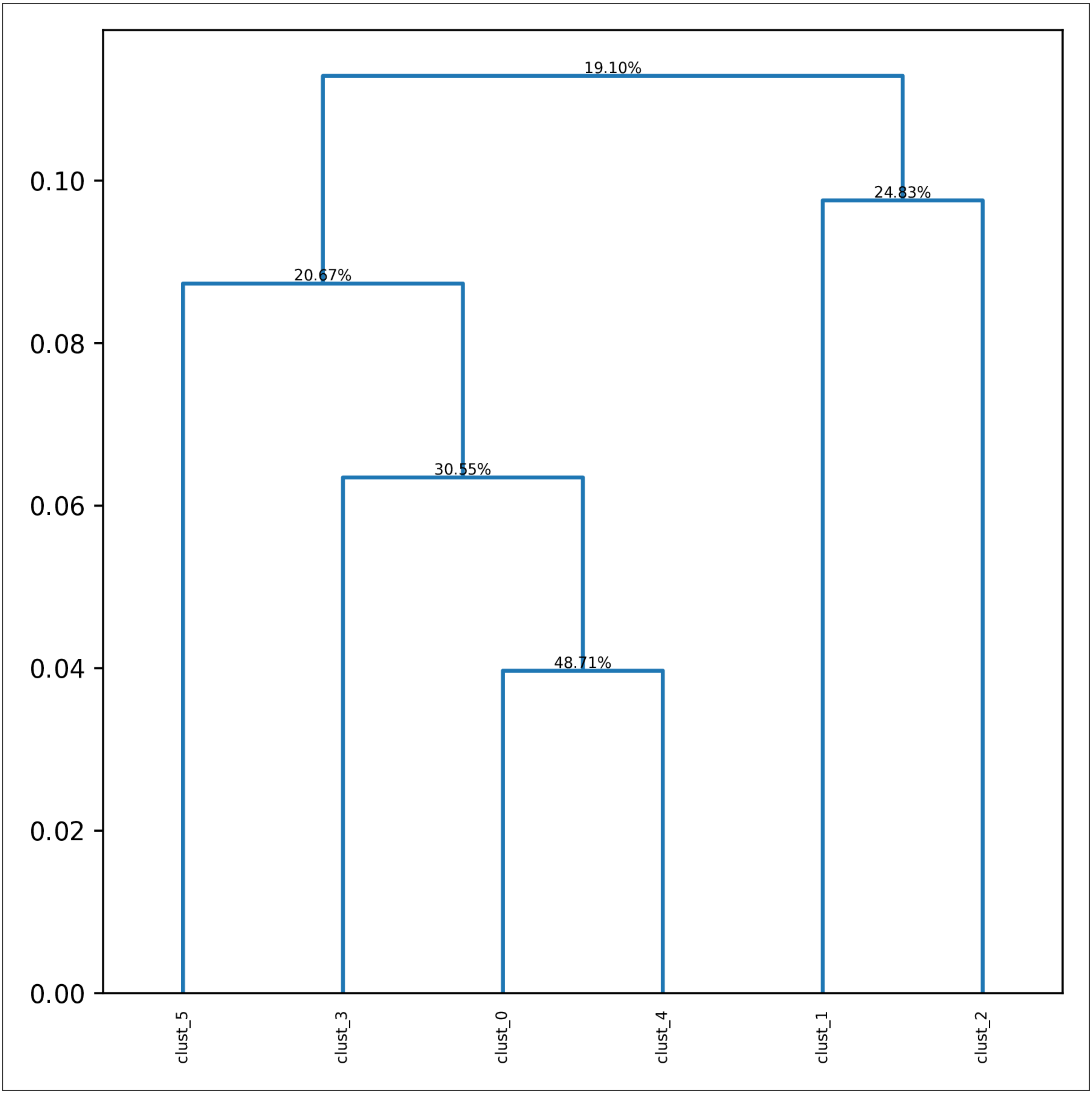

Sample: G0022 with 10 cells per cluster

Min. read depth: 10,  Mean UMIs per cluster:  91,284

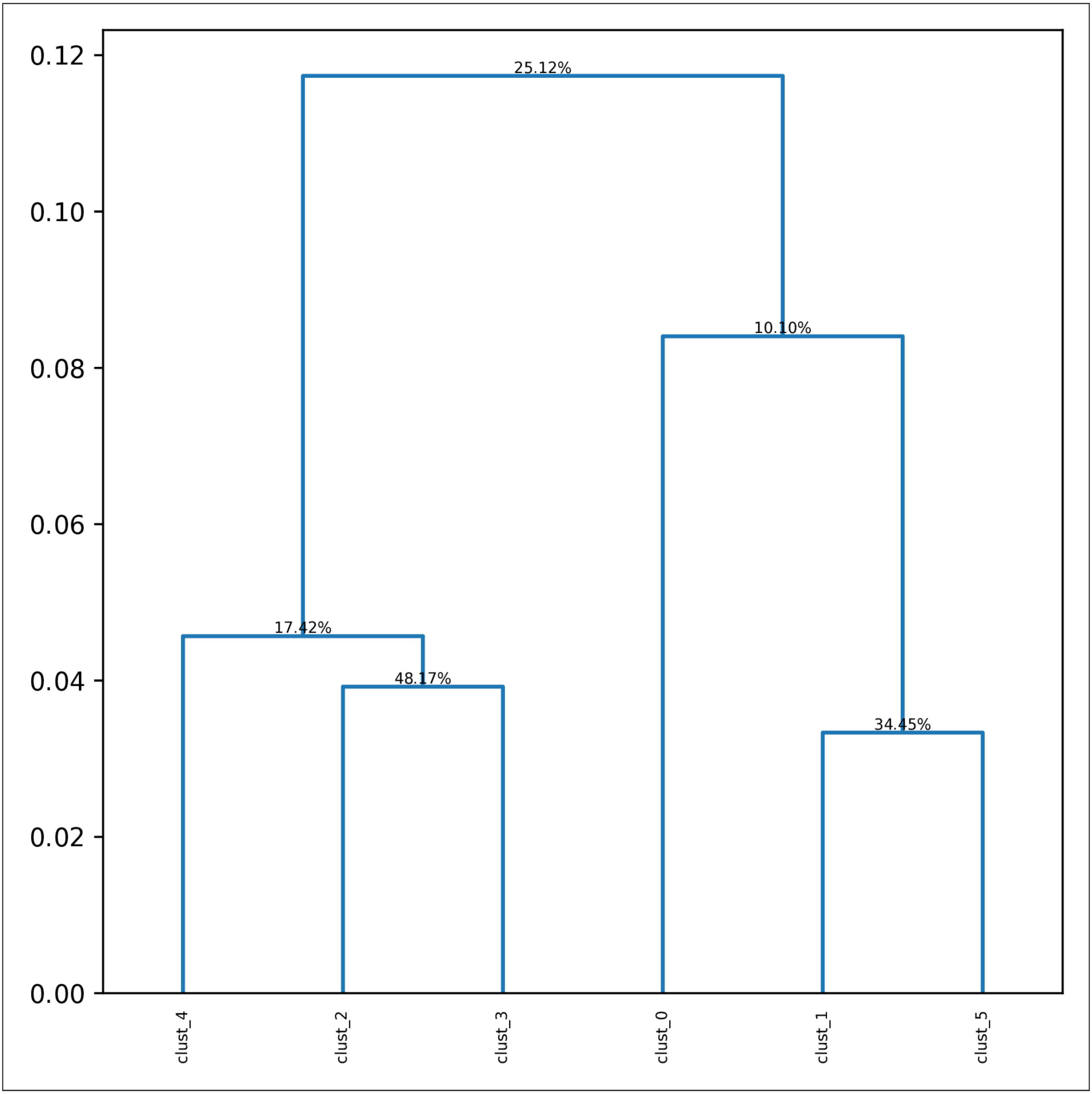

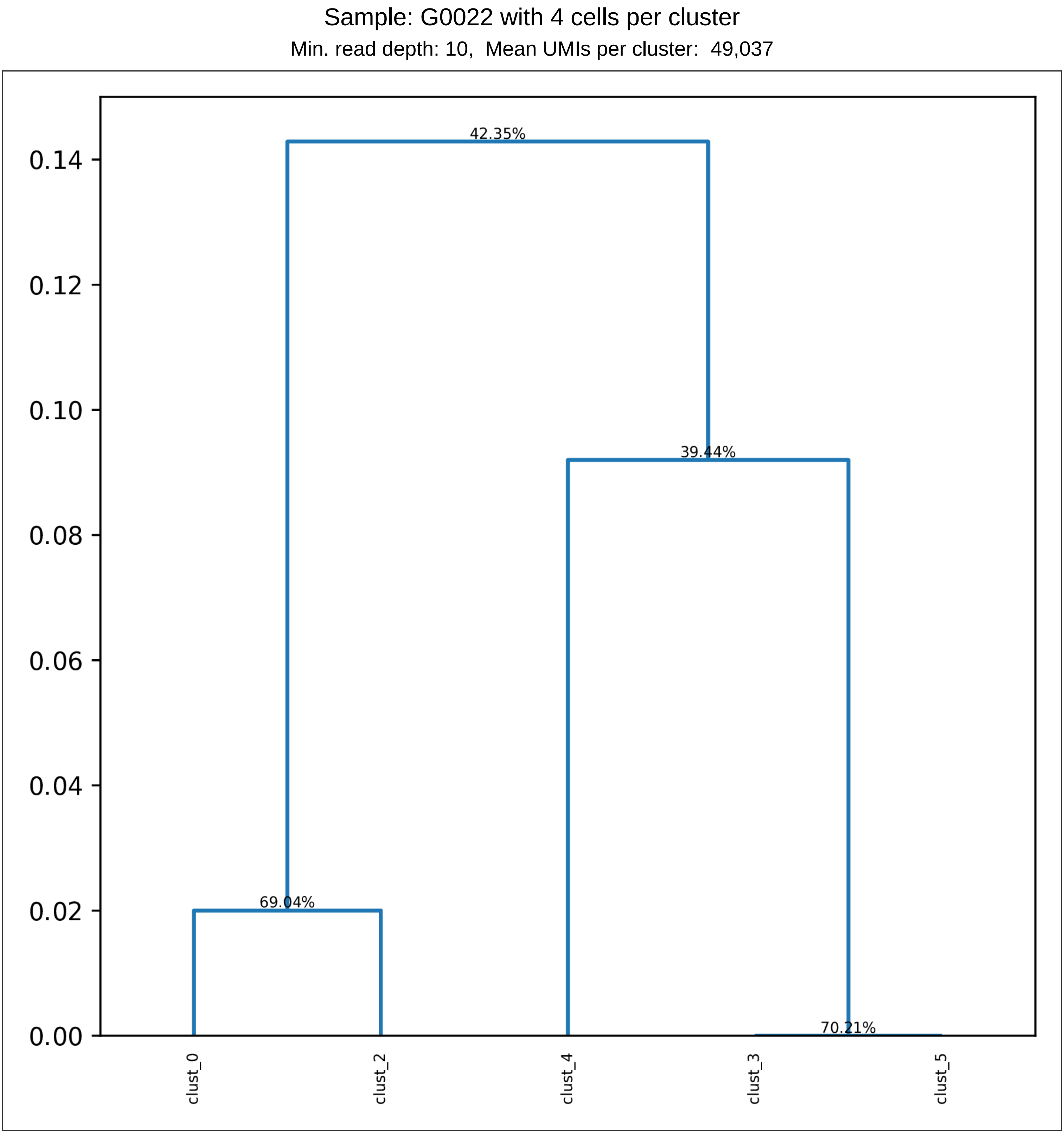
Sample: G0022 with 4 cells per cluster
Min. read depth: 10, Mean UMIs per cluster: 49,037
42.35%
39.44%
69.04%
70.21%
0.14
0.12
0.10
0.08
0.06
0.04
0.02
0.00
clust_0
clust_2
clust_4
clust_3
clust_5

Sample: G0023 with all cells per cluster

Min. read depth: 10,  Mean UMIs per cluster:  3,483,645

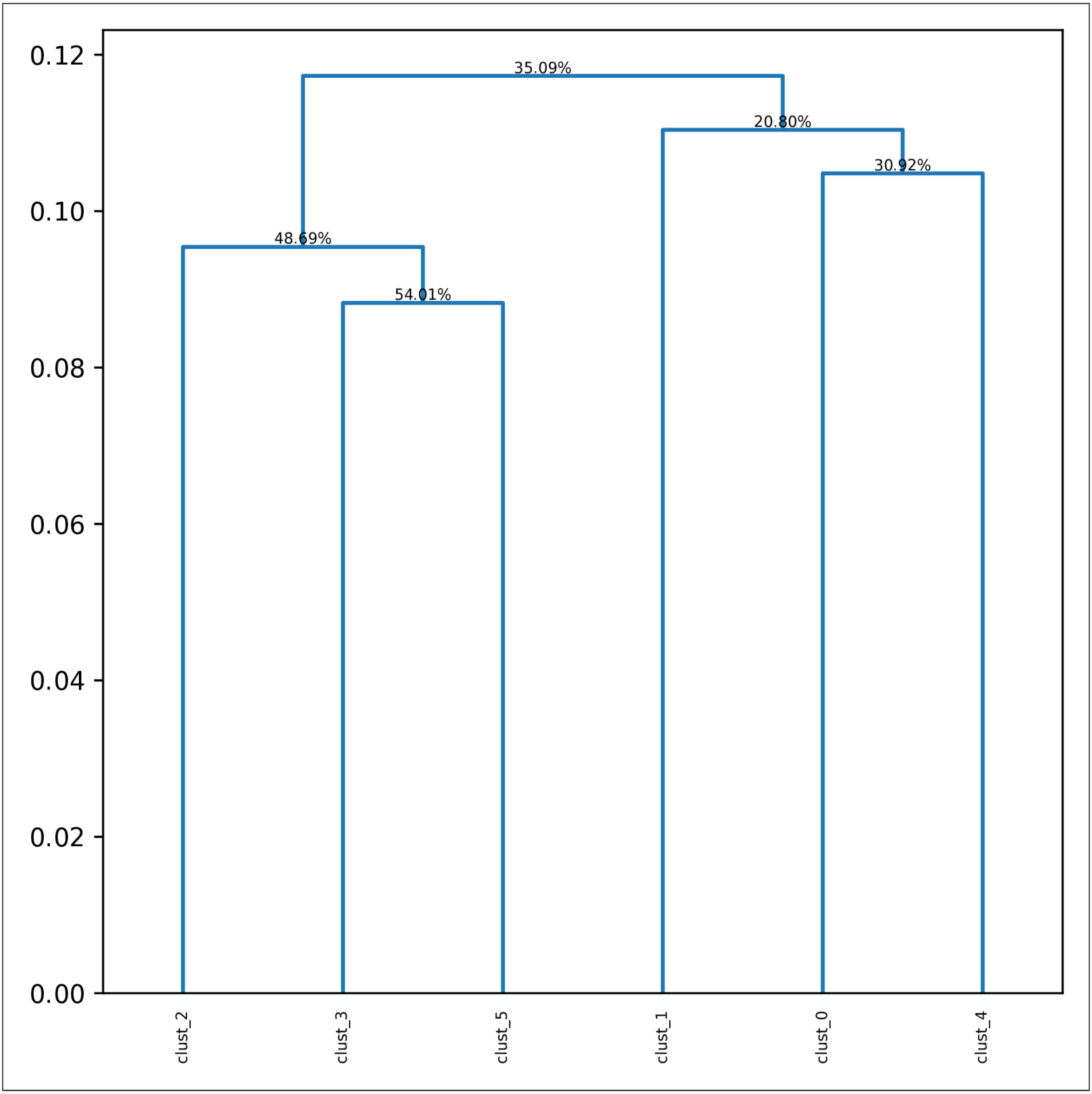

Sample: G0023 with 500 cells per cluster

Min. read depth: 10,  Mean UMIs per cluster:  2,166,358

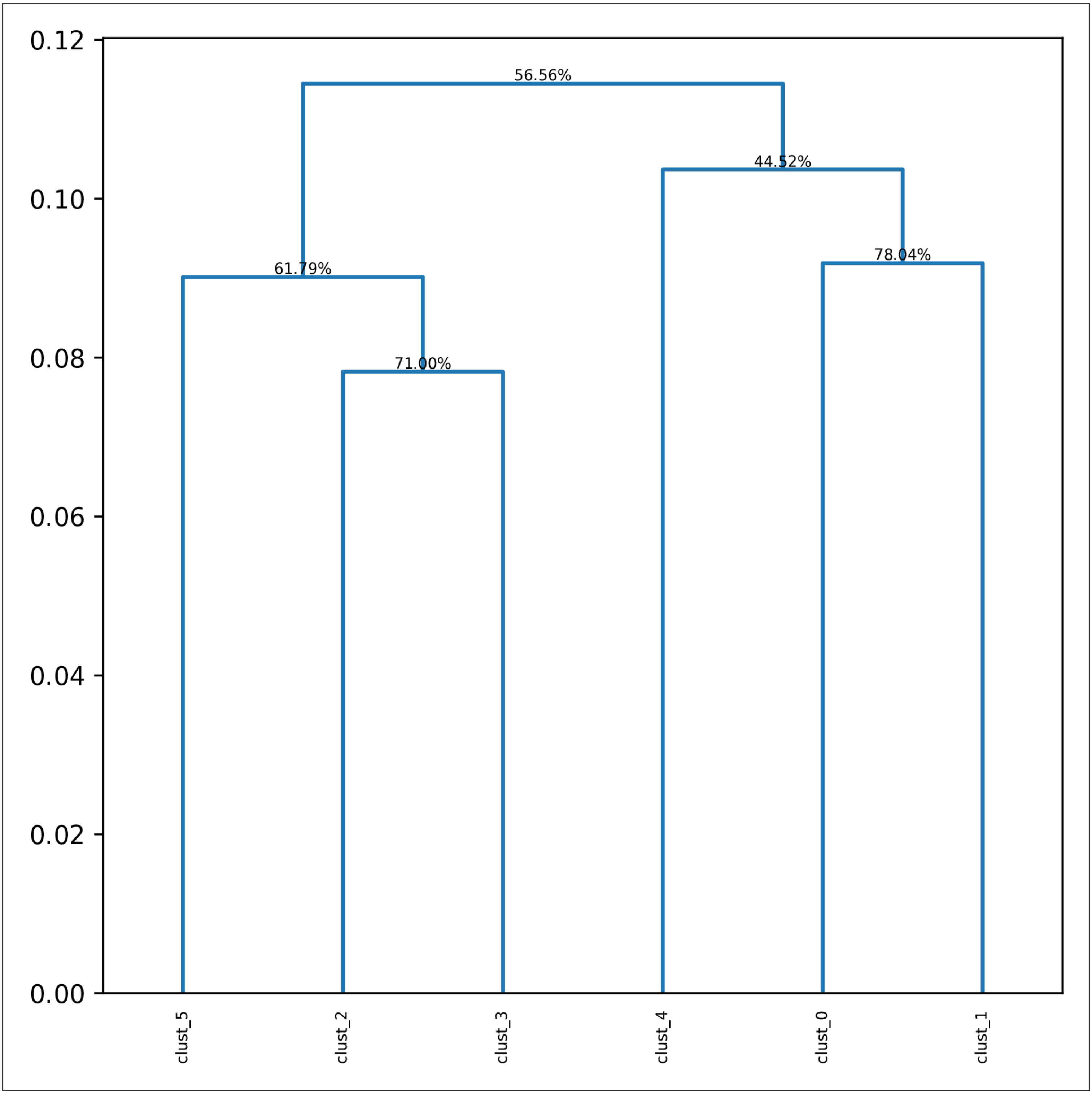

Sample: G0023 with 100 cells per cluster

Min. read depth: 10,  Mean UMIs per cluster:  501,404

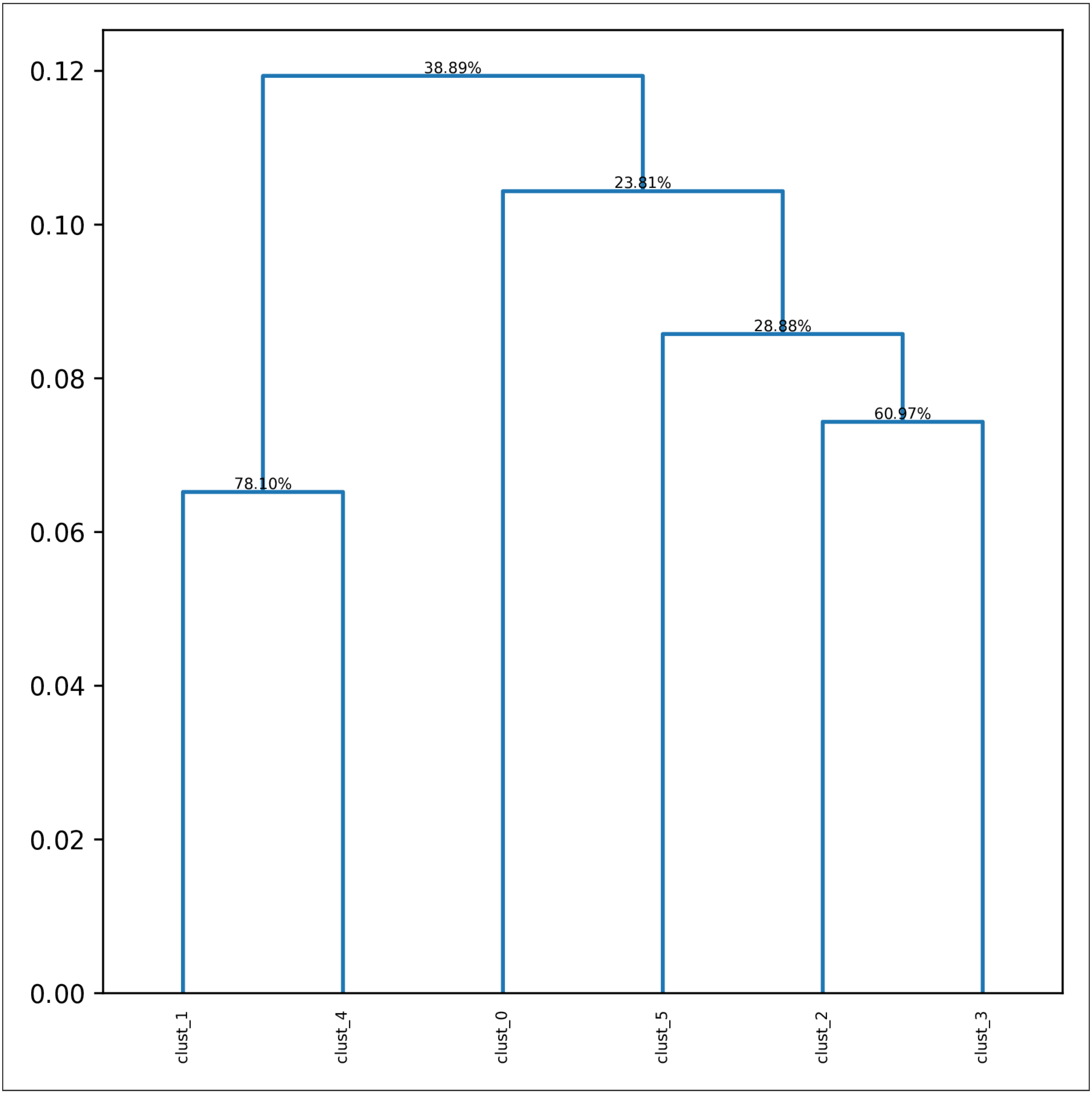

Sample: G0023 with 50 cells per cluster
Min. read depth: 10, Mean UMIs per cluster: 245,781

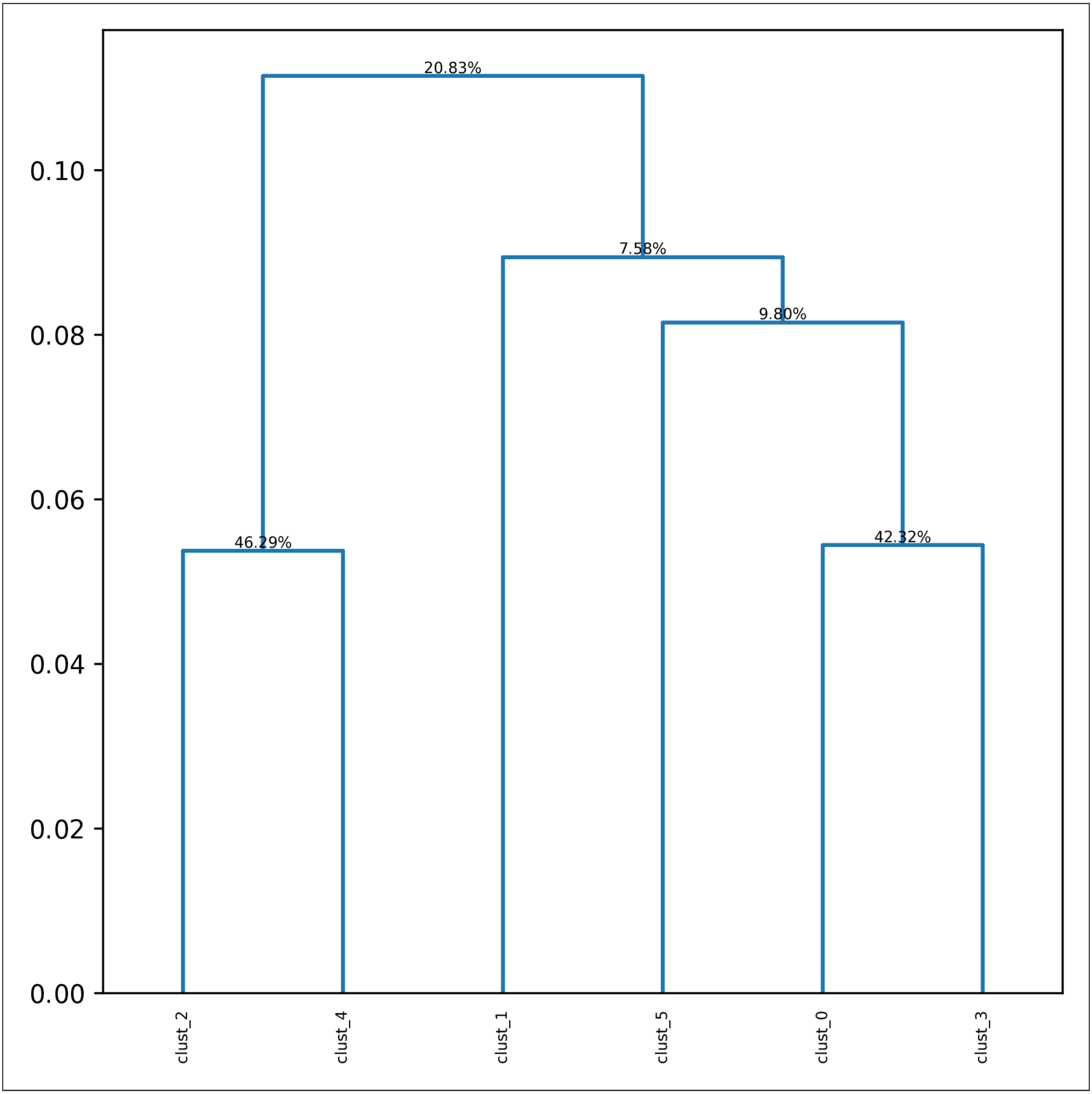

Sample: G0023 with 20 cells per cluster

Min. read depth: 10,  Mean UMIs per cluster:  95,431

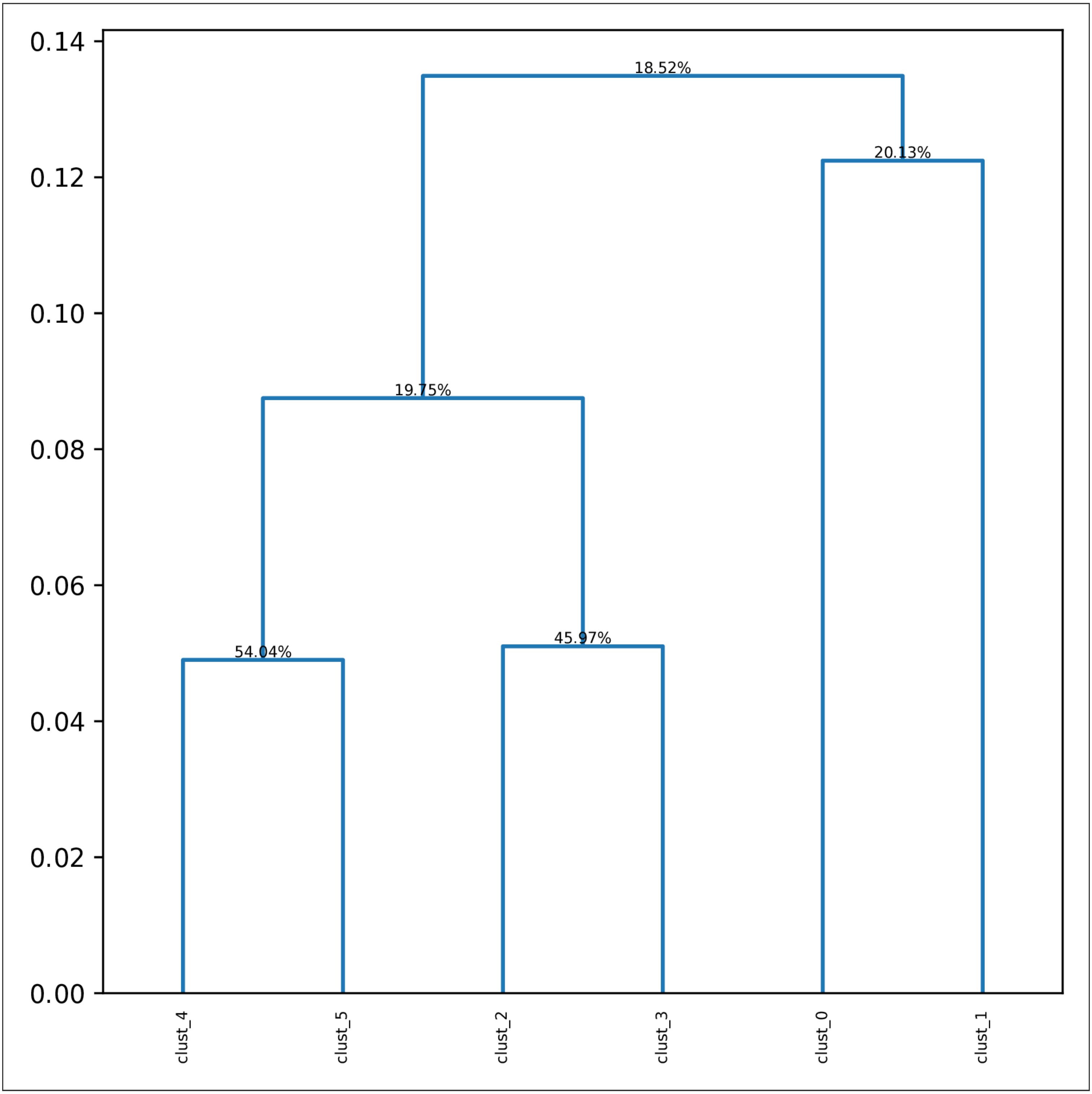

Sample: G0023 with 10 cells per cluster
Min. read depth: 10,  Mean UMIs per cluster:  50,240

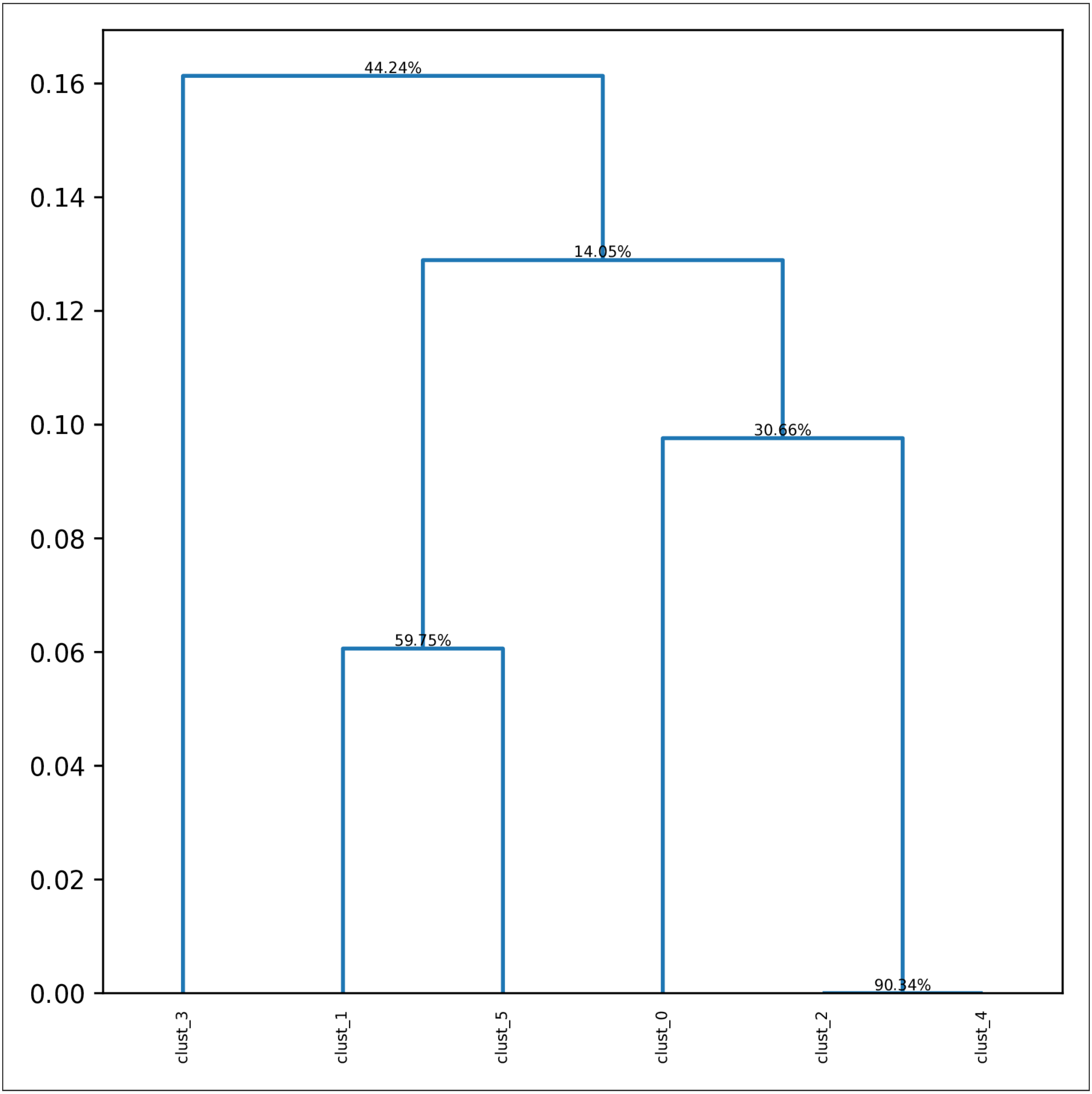

Sample: G0023 with 5 cells per cluster
Min. read depth: 10,  Mean UMIs per cluster:  26,537

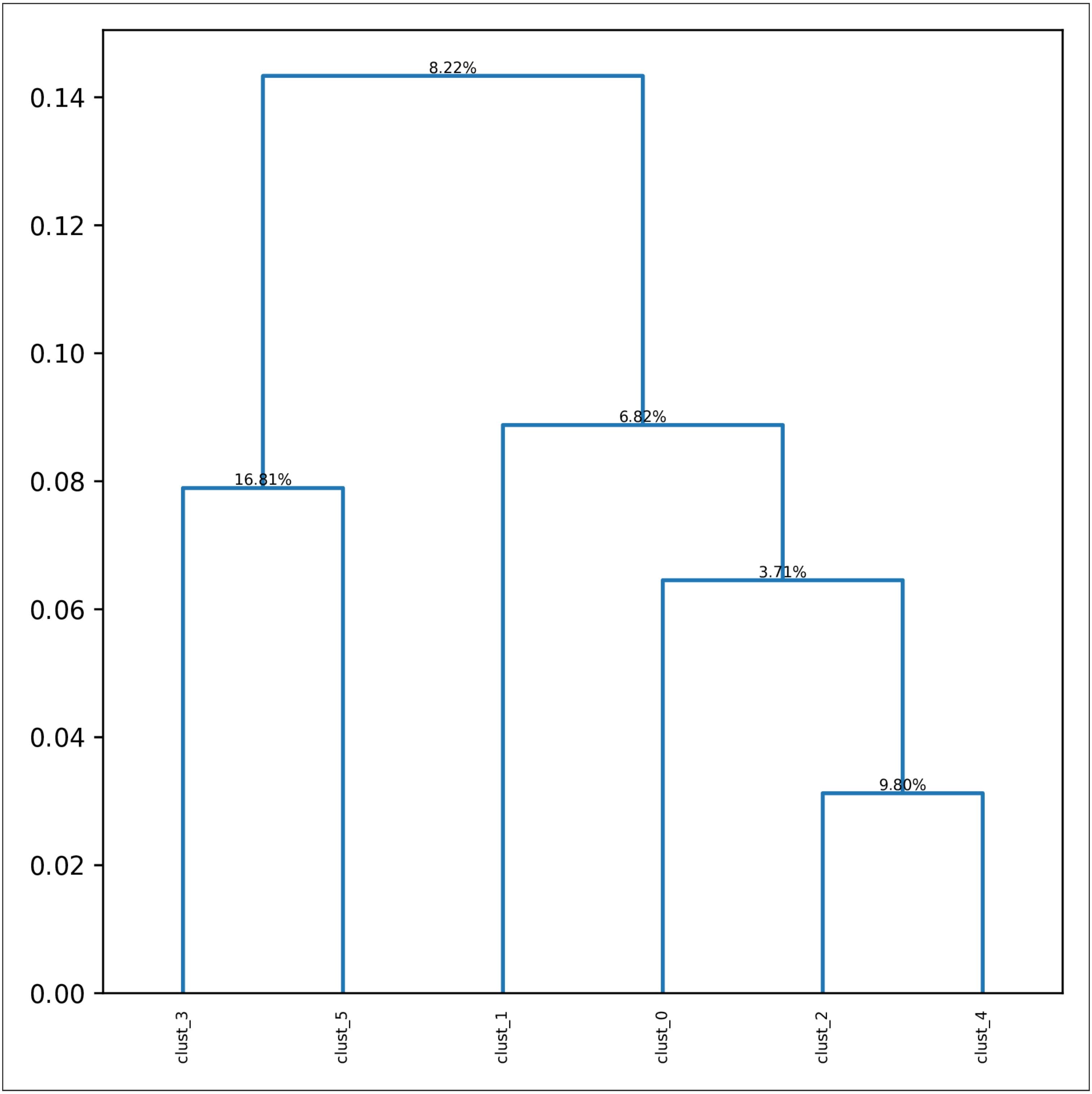

Sample: G0023 with 4 cells per cluster

Min. read depth: 10,  Mean UMIs per cluster:  21,871

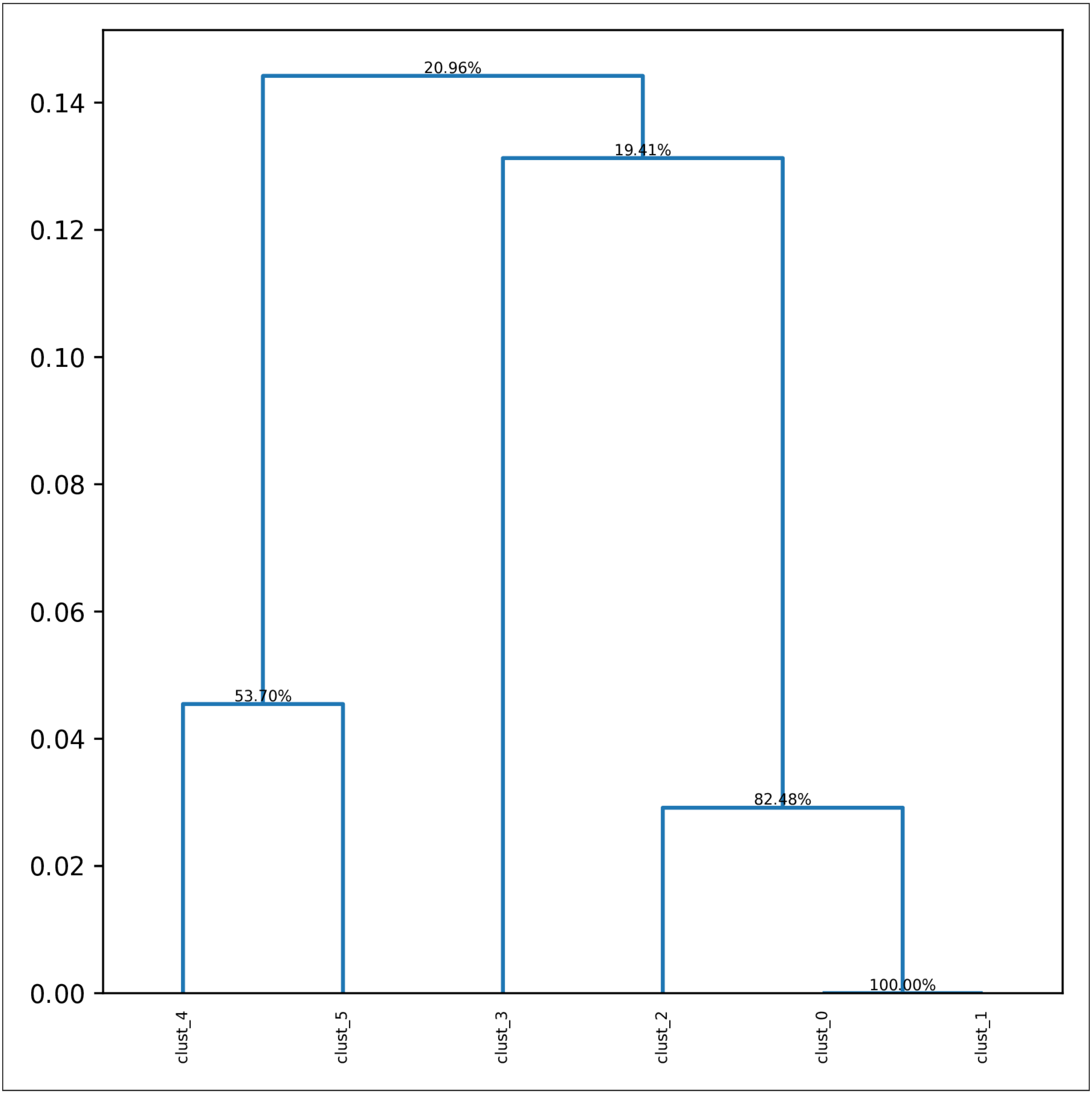

Sample: G0024 with all cells per cluster

Min. read depth: 10,  Mean UMIs per cluster:  3,378,040

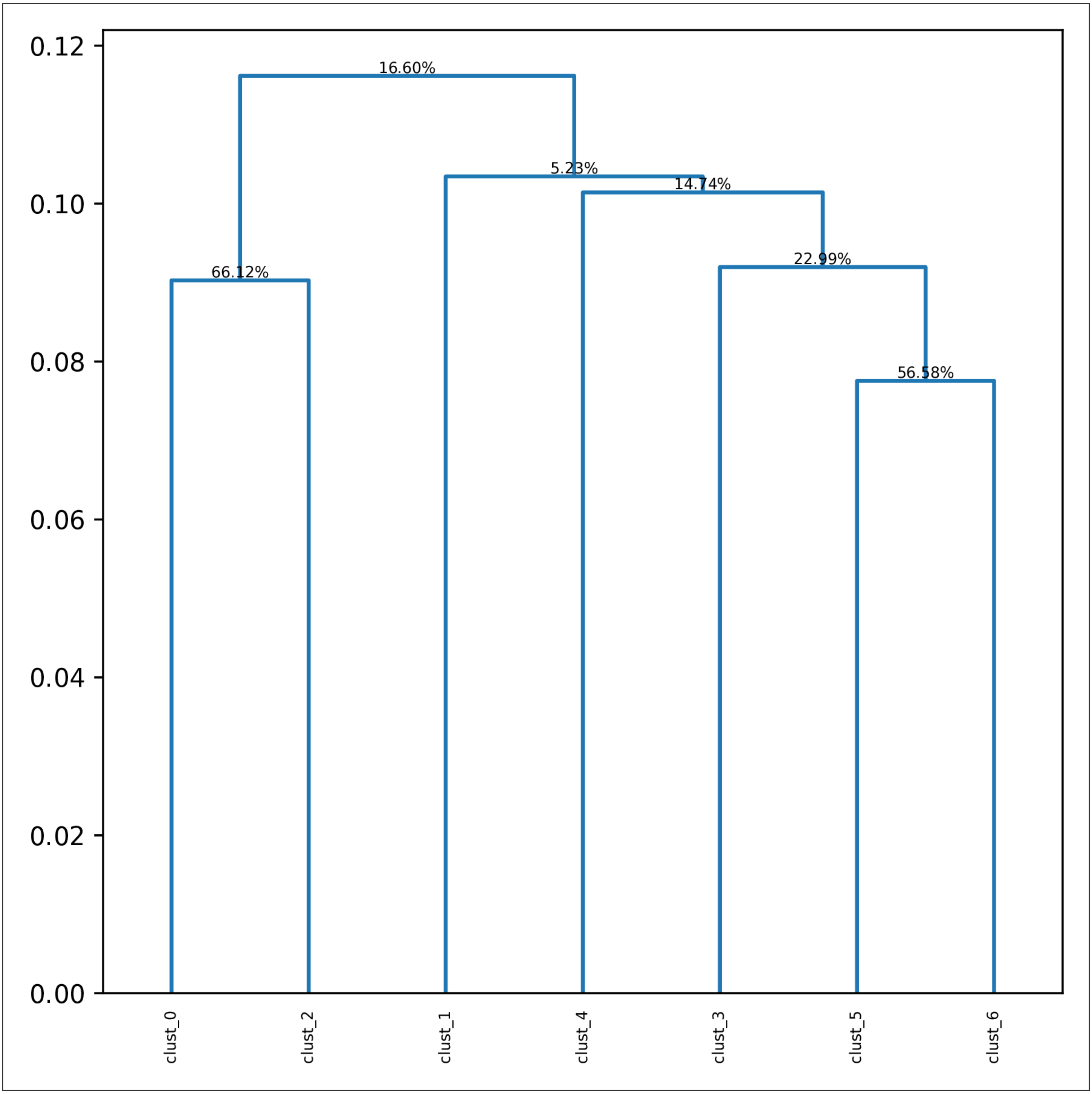

Sample: G0024 with 500 cells per cluster

Min. read depth: 10,  Mean UMIs per cluster:  1,481,688

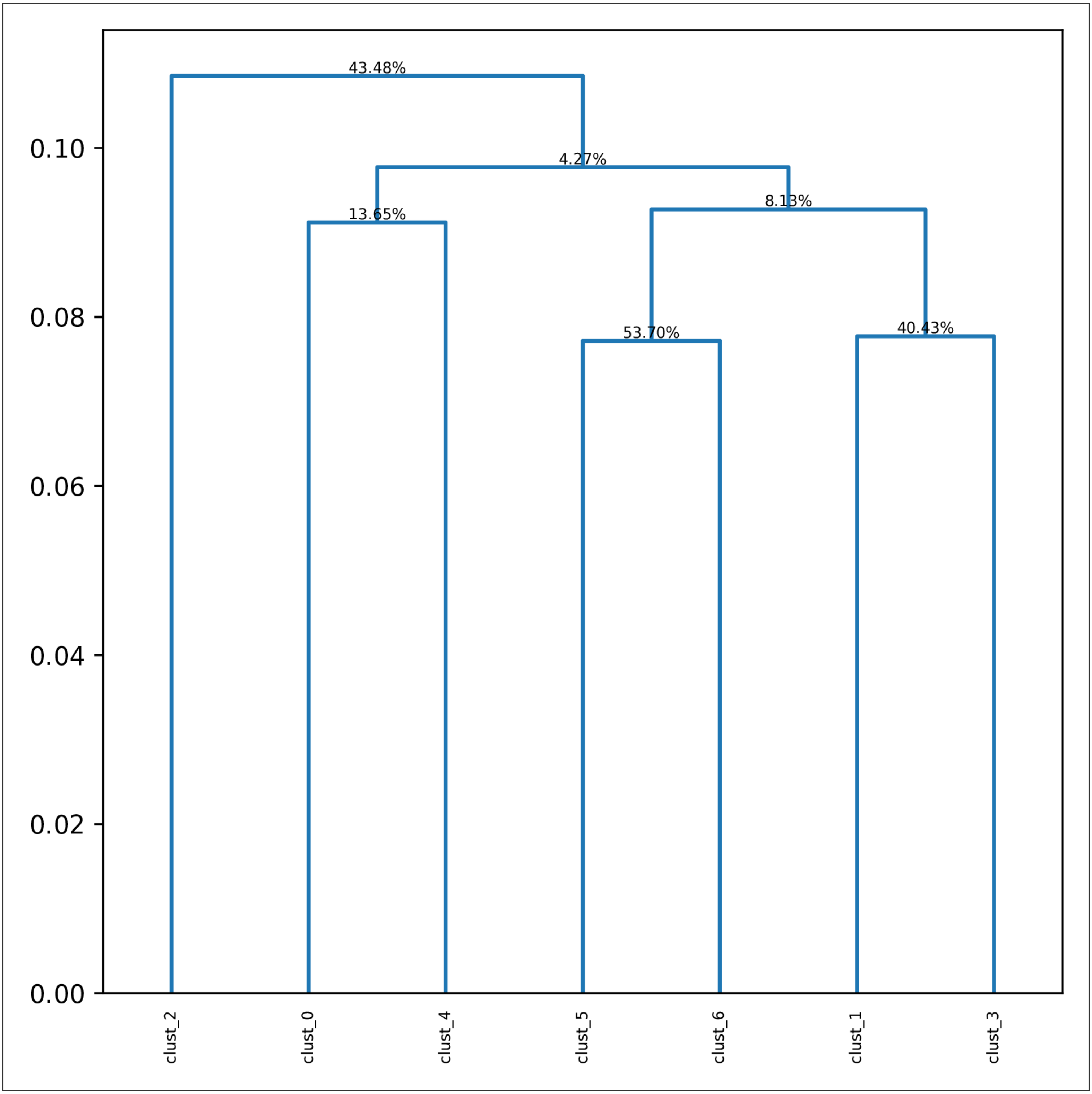

Sample: G0024 with 100 cells per cluster

Min. read depth: 10,  Mean UMIs per cluster:  346,515

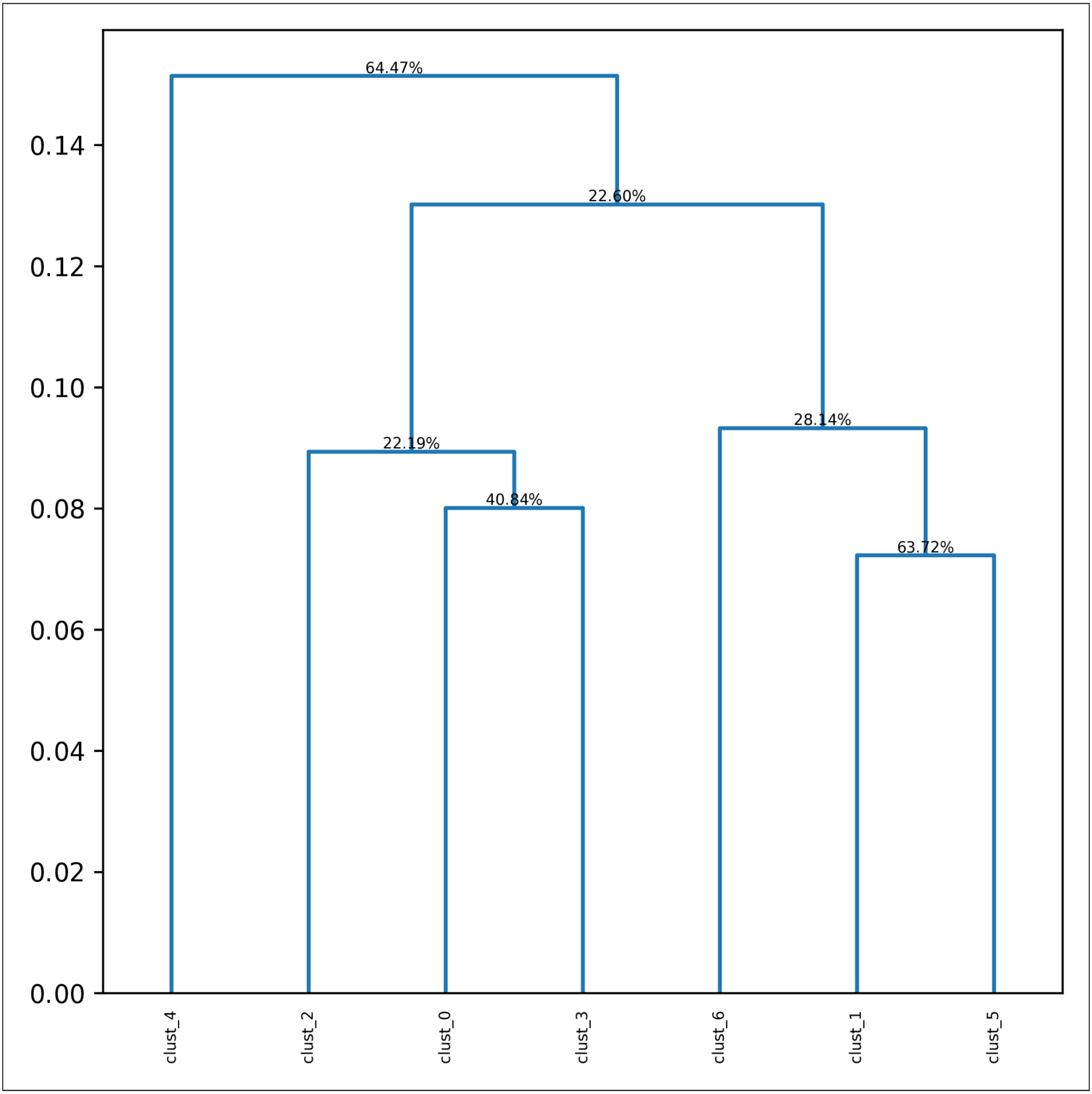

Sample: G0024 with 50 cells per cluster

Min. read depth: 10,  Mean UMIs per cluster:  171,454

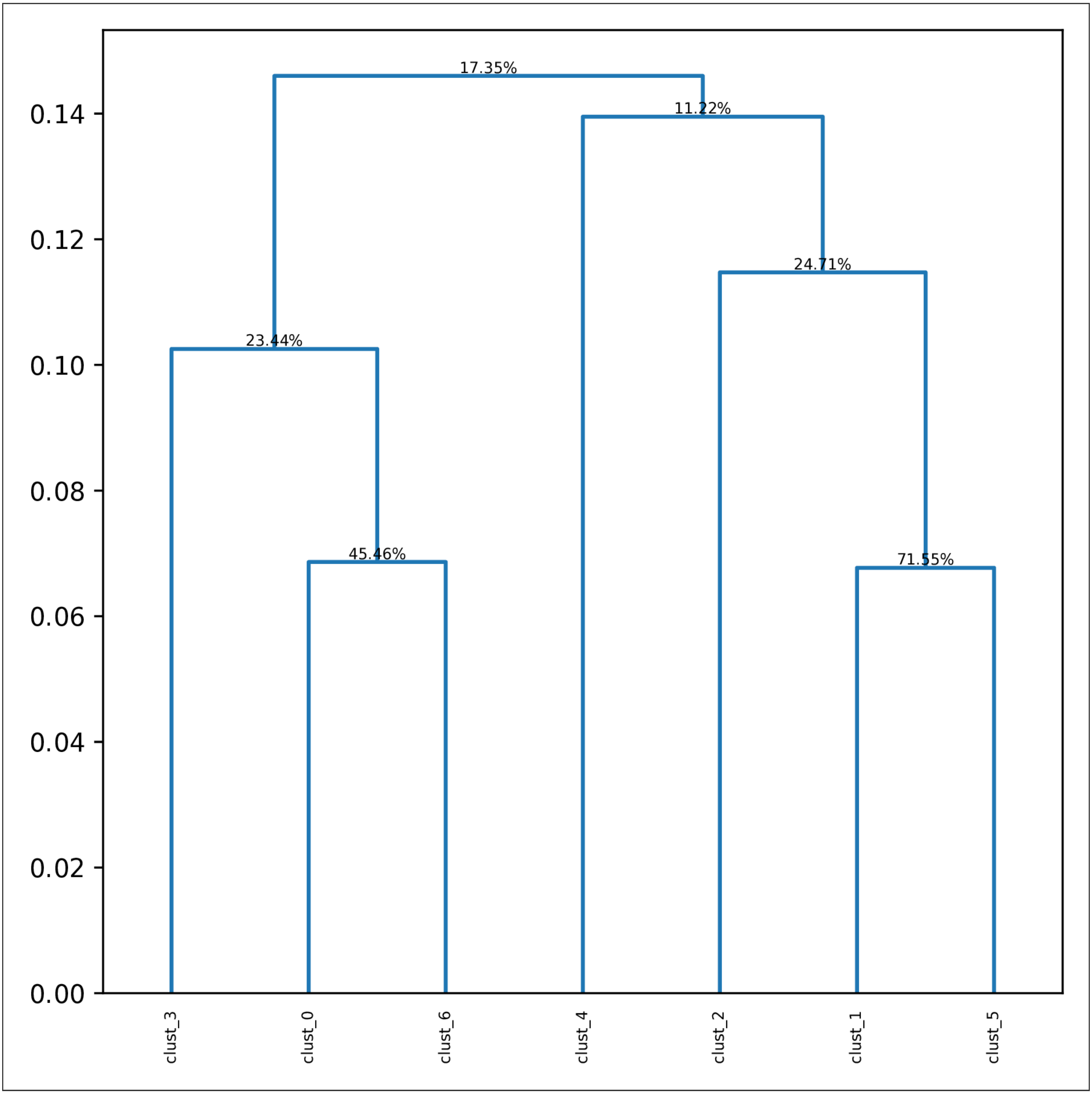

Sample: G0024 with 20 cells per cluster
Min. read depth: 10,  Mean UMIs per cluster:  67,680

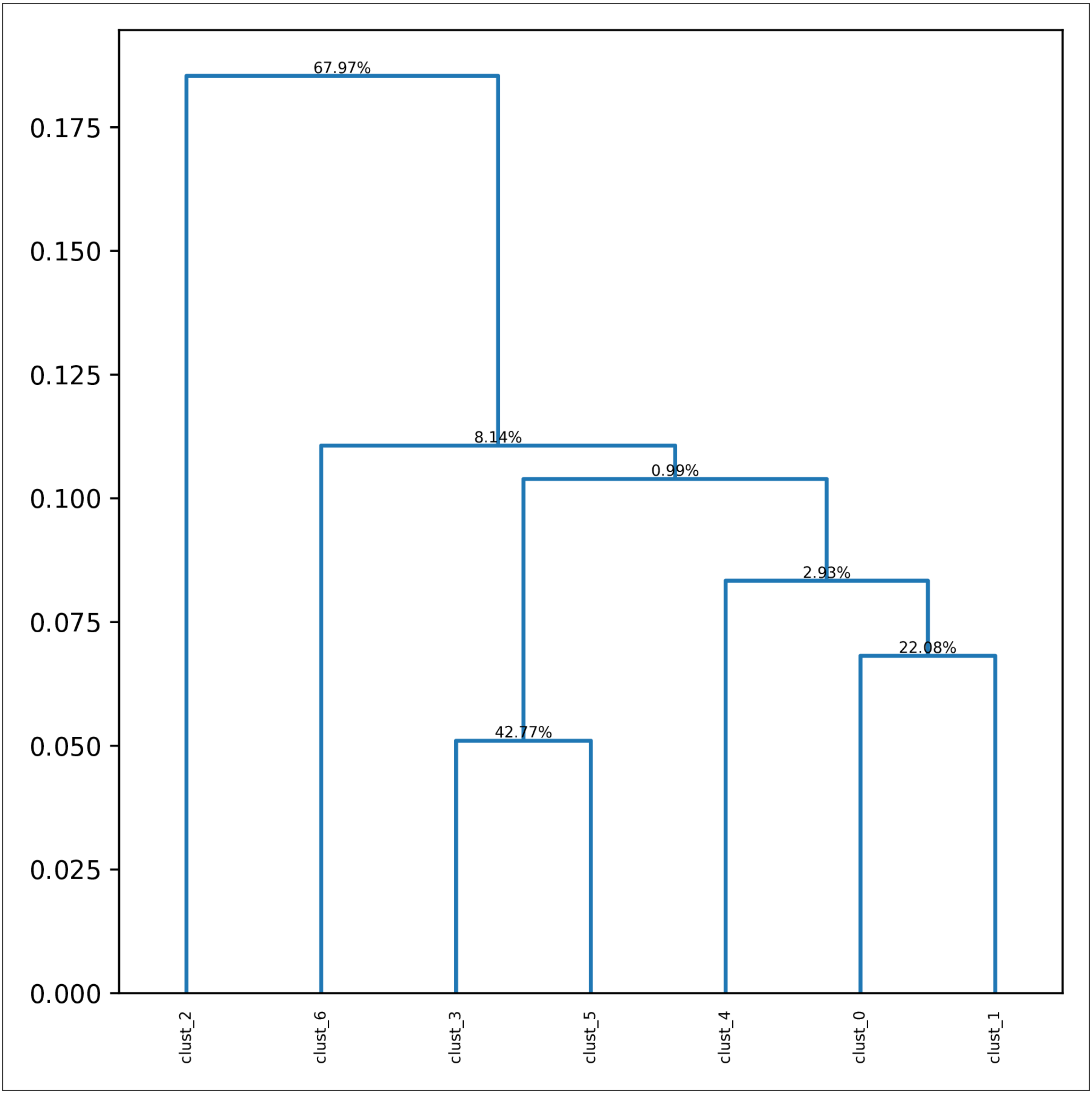

Sample: G0024 with 10 cells per cluster

Min. read depth: 10,  Mean UMIs per cluster:  35,705

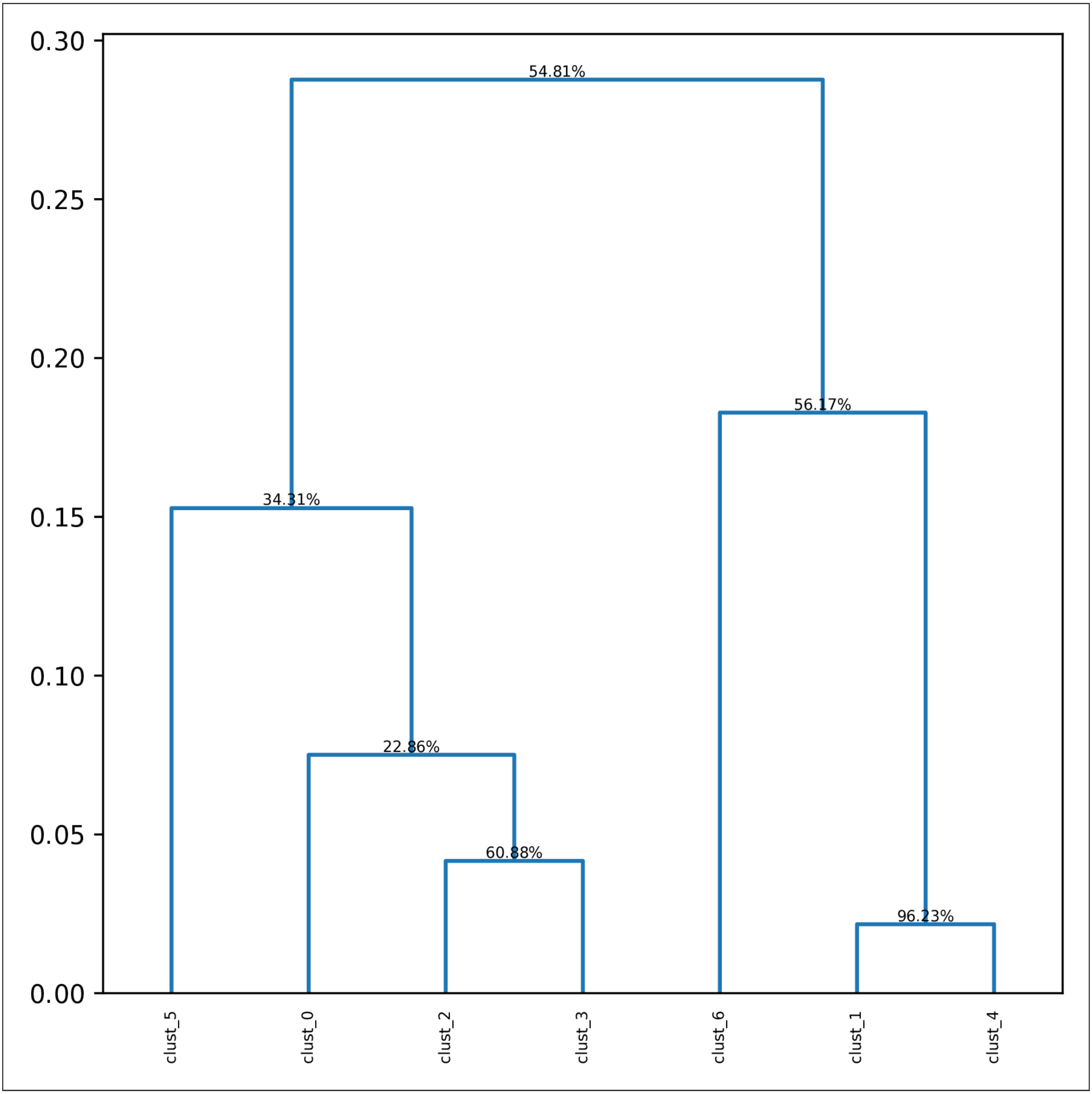

Sample: G0025 with all cells per cluster

Min. read depth: 10,  Mean UMIs per cluster:  28,724,919

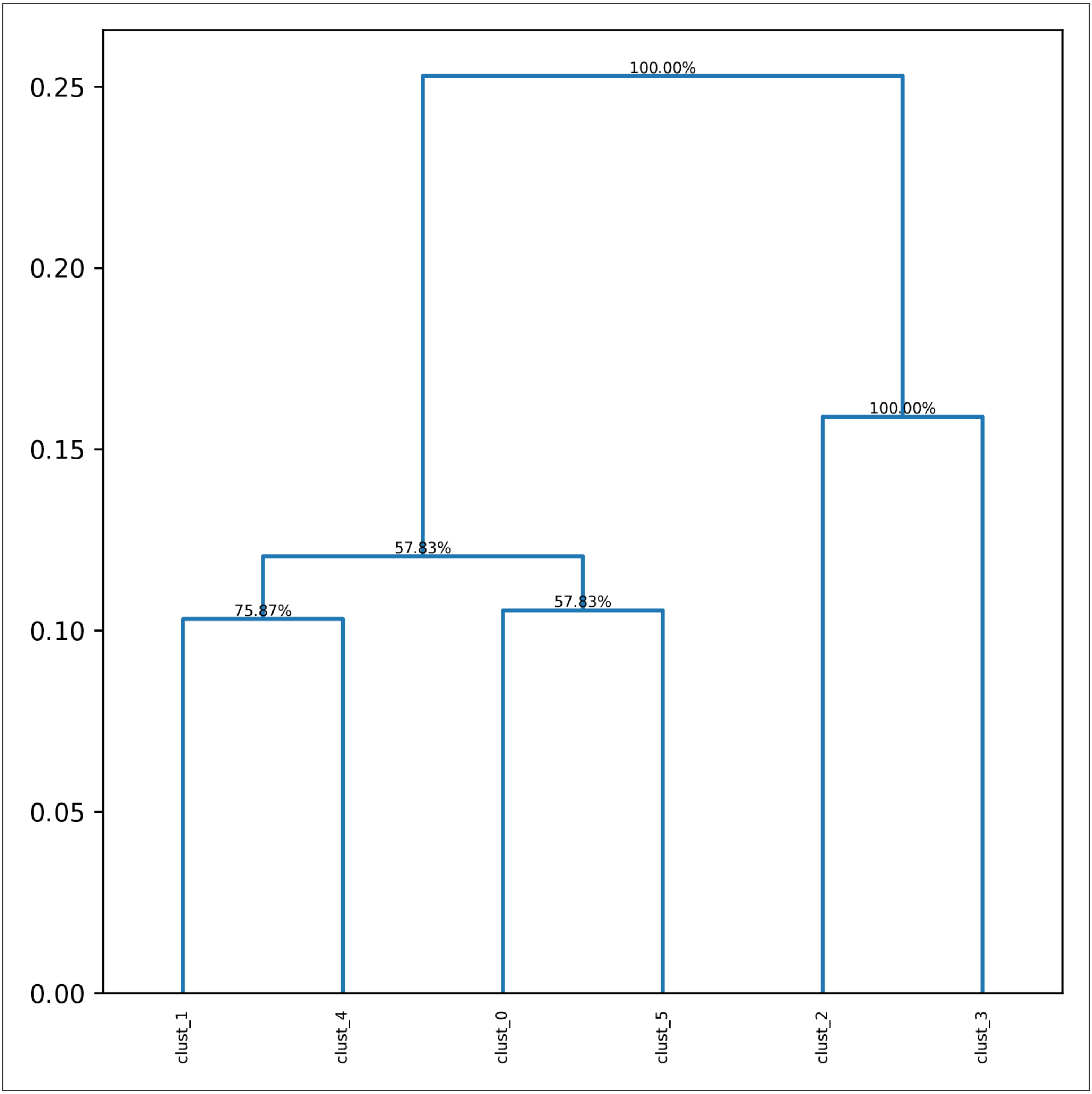

Sample: G0025 with 500 cells per cluster

Min. read depth: 10, Mean UMIs per cluster: 12,672,736

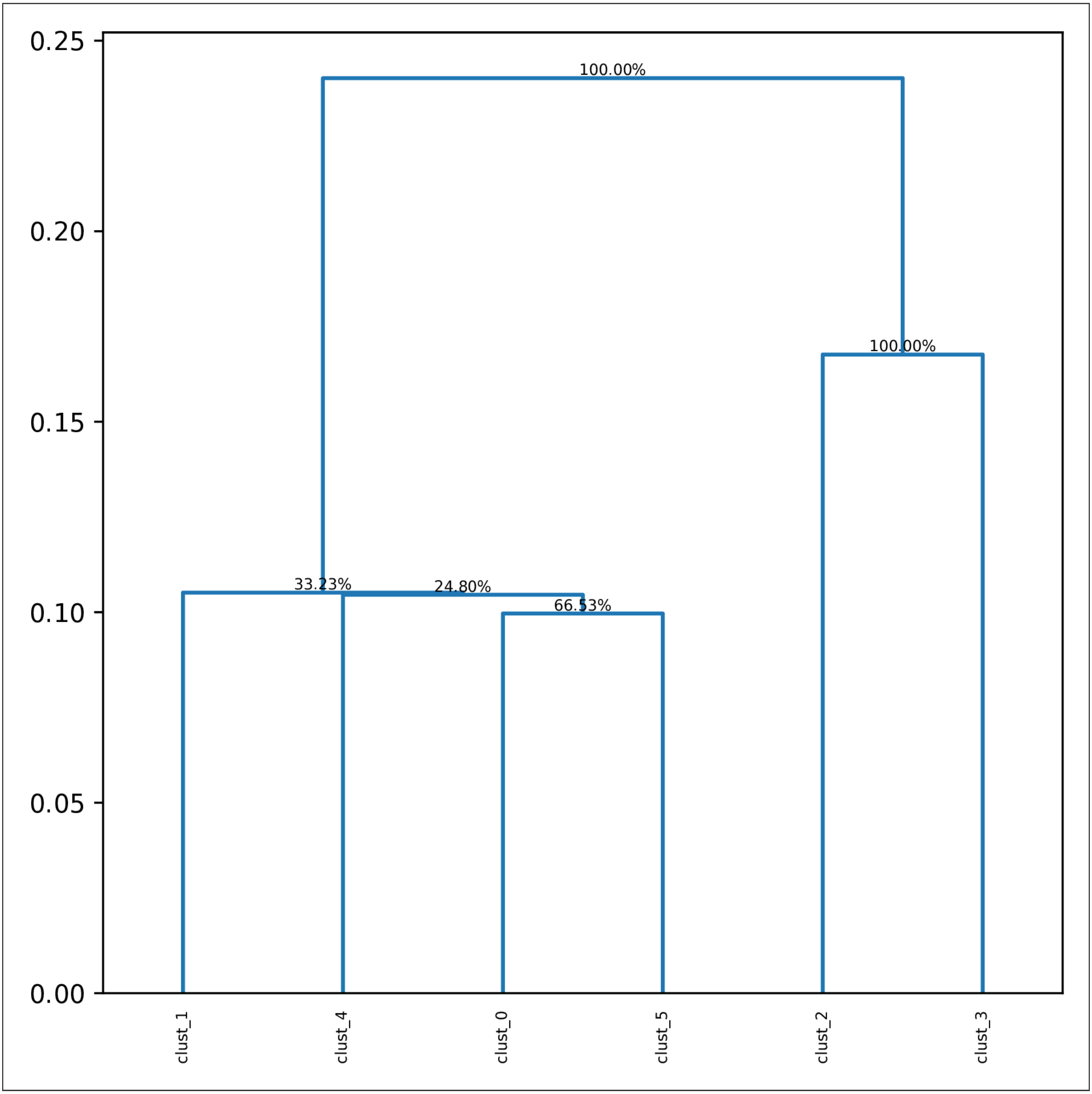

Sample: G0025 with 100 cells per cluster

Min. read depth: 10,  Mean UMIs per cluster:  2,981,537

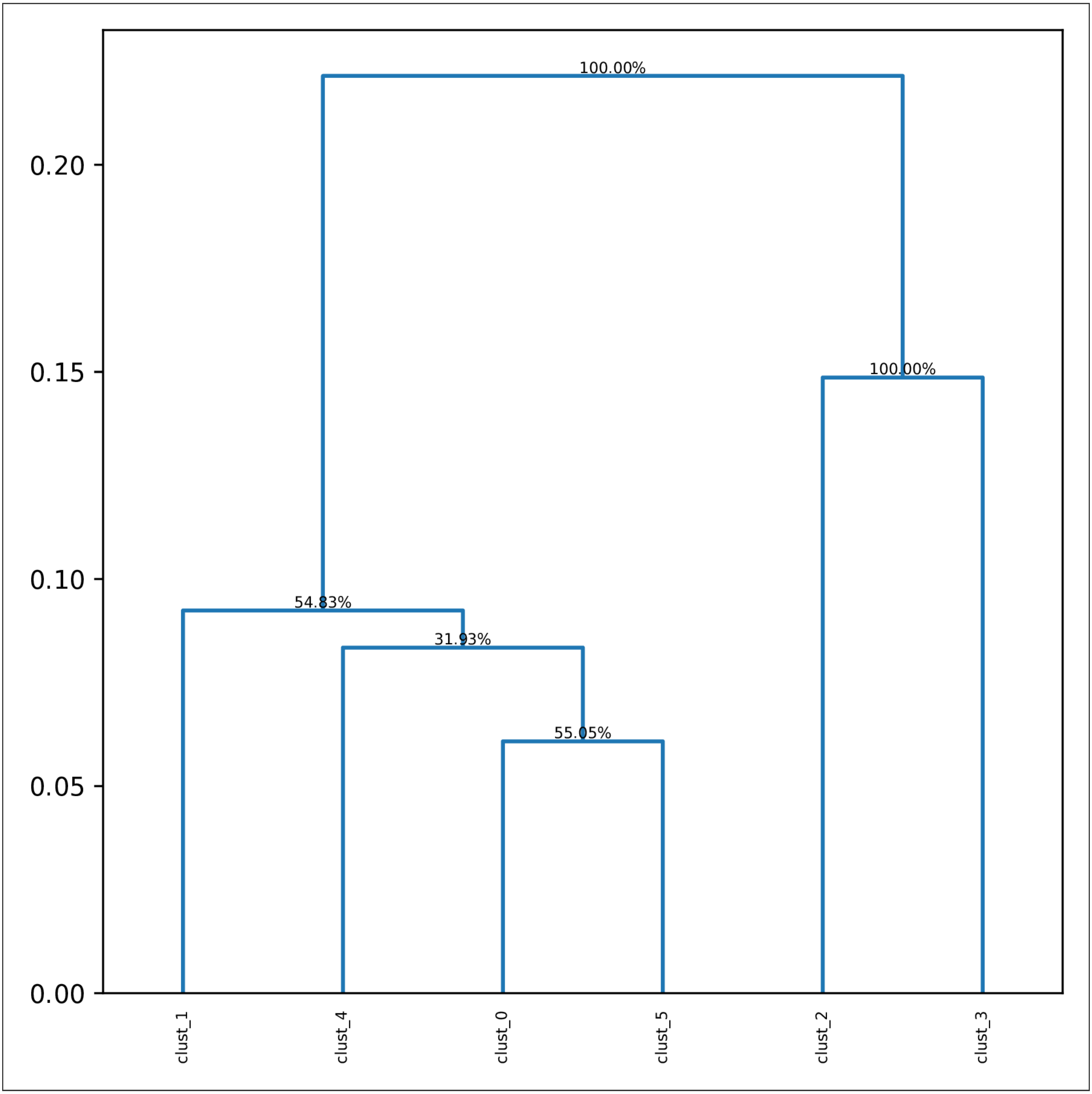

Sample: G0025 with 50 cells per cluster
Min. read depth: 10,  Mean UMIs per cluster:  1,446,721

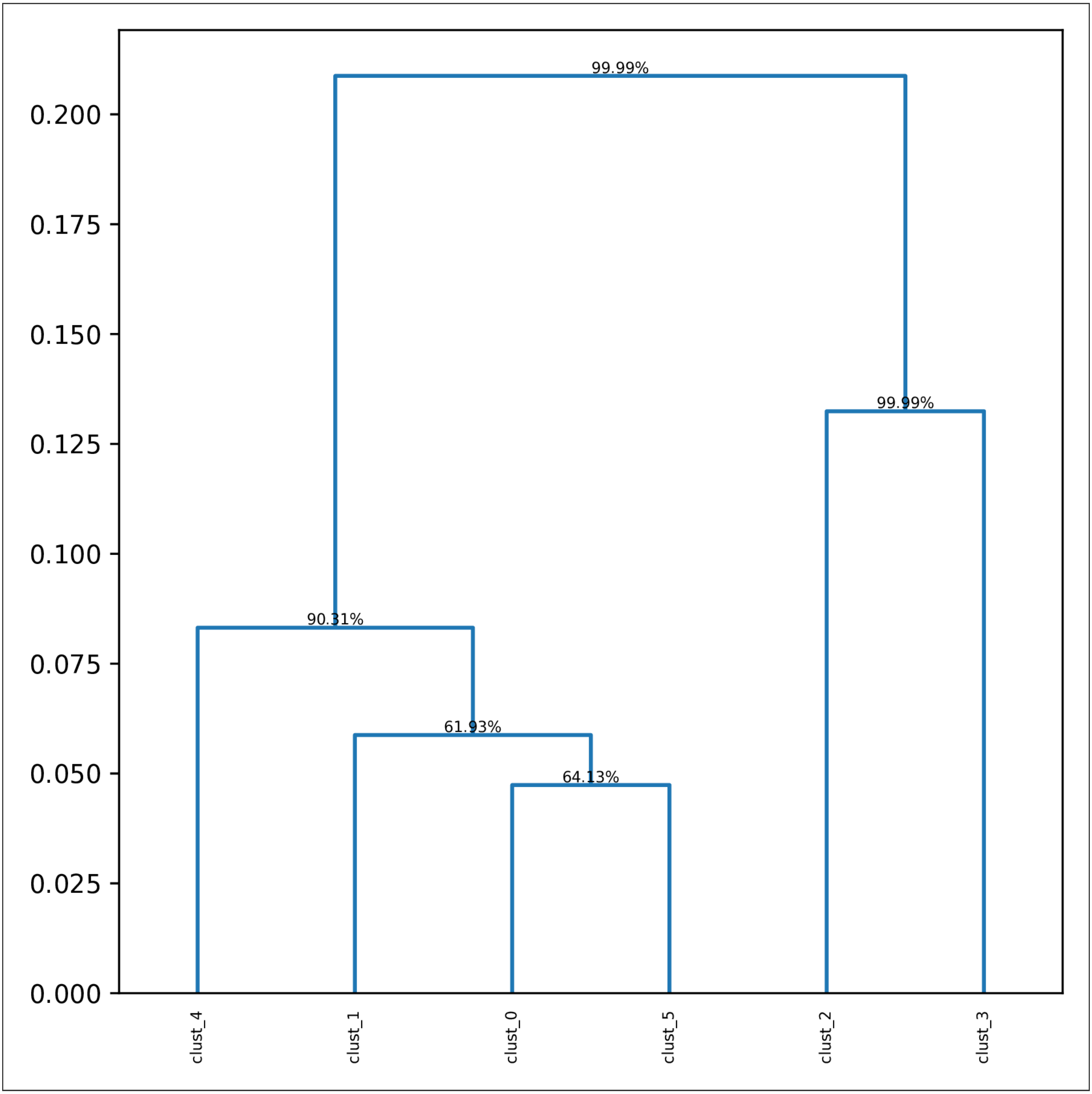

Sample: G0025 with 20 cells per cluster
Min. read depth: 10,  Mean UMIs per cluster:  640,830

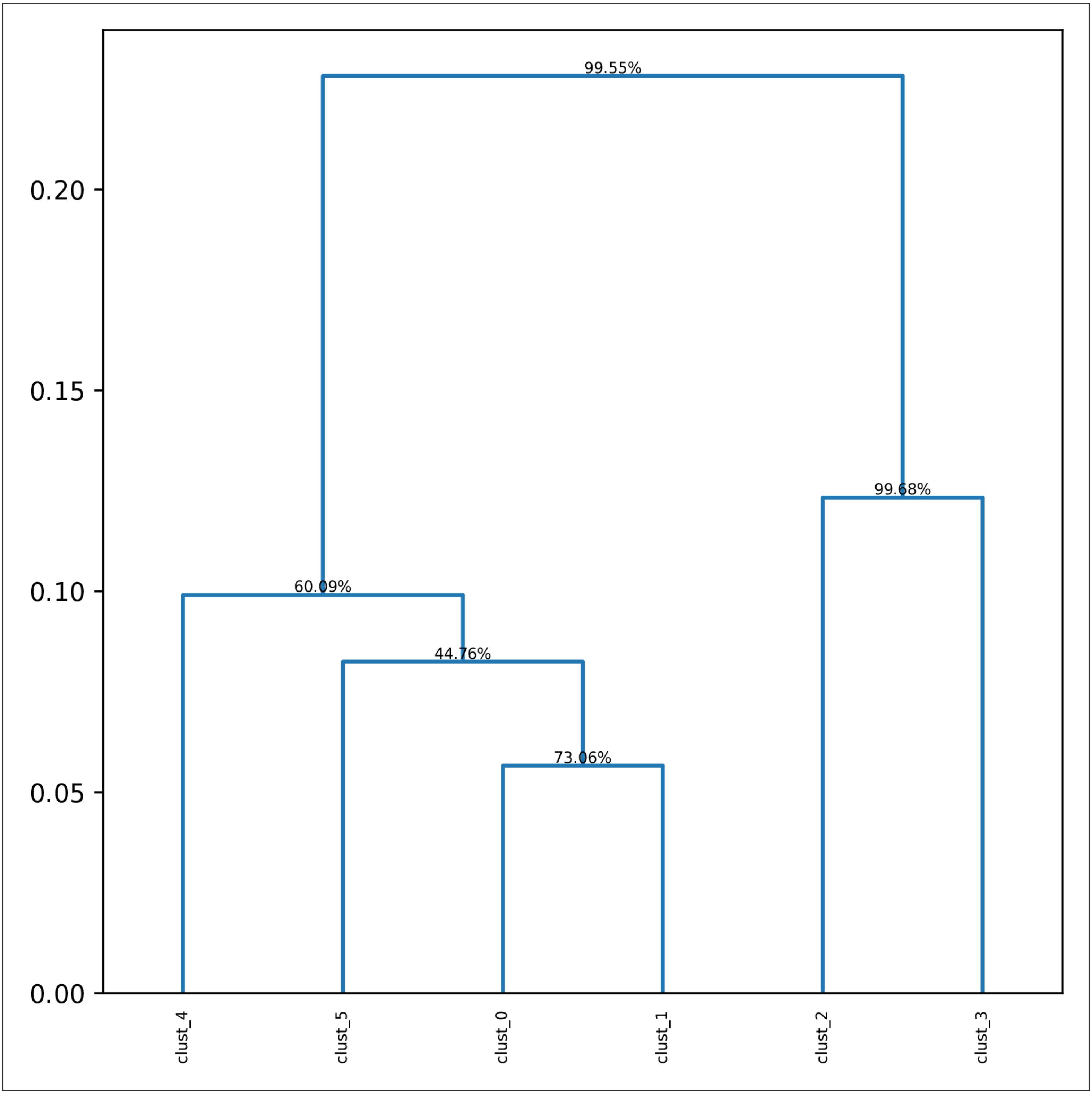

Sample: G0025 with 10 cells per cluster

Min. read depth: 10,  Mean UMIs per cluster:  263,126

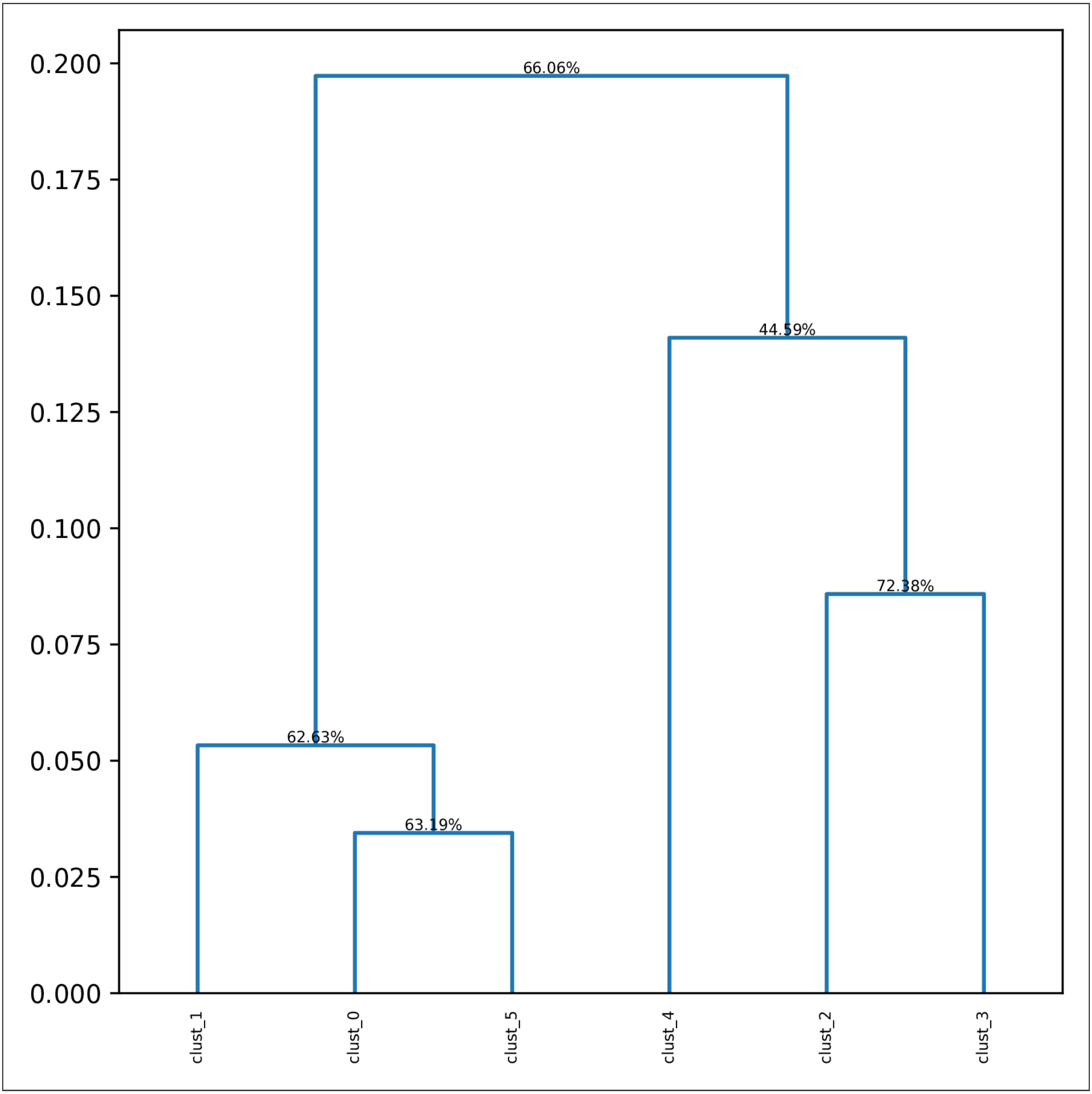

Sample: G0025 with 5 cells per cluster
Min. read depth: 10,  Mean UMIs per cluster:  130,945

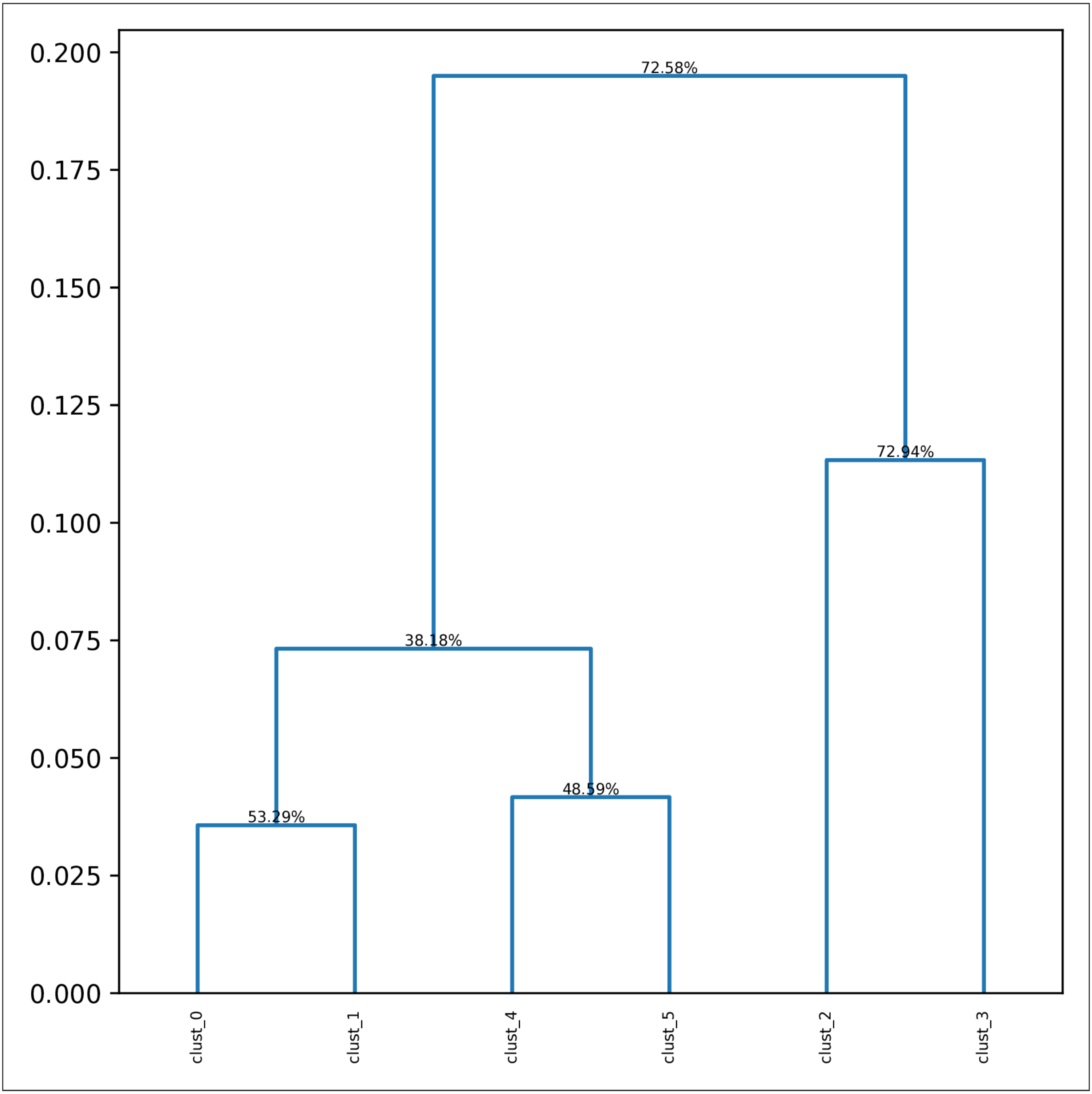

Sample: G0026 with all cells per cluster

Min. read depth: 10,  Mean UMIs per cluster:  16,743,492

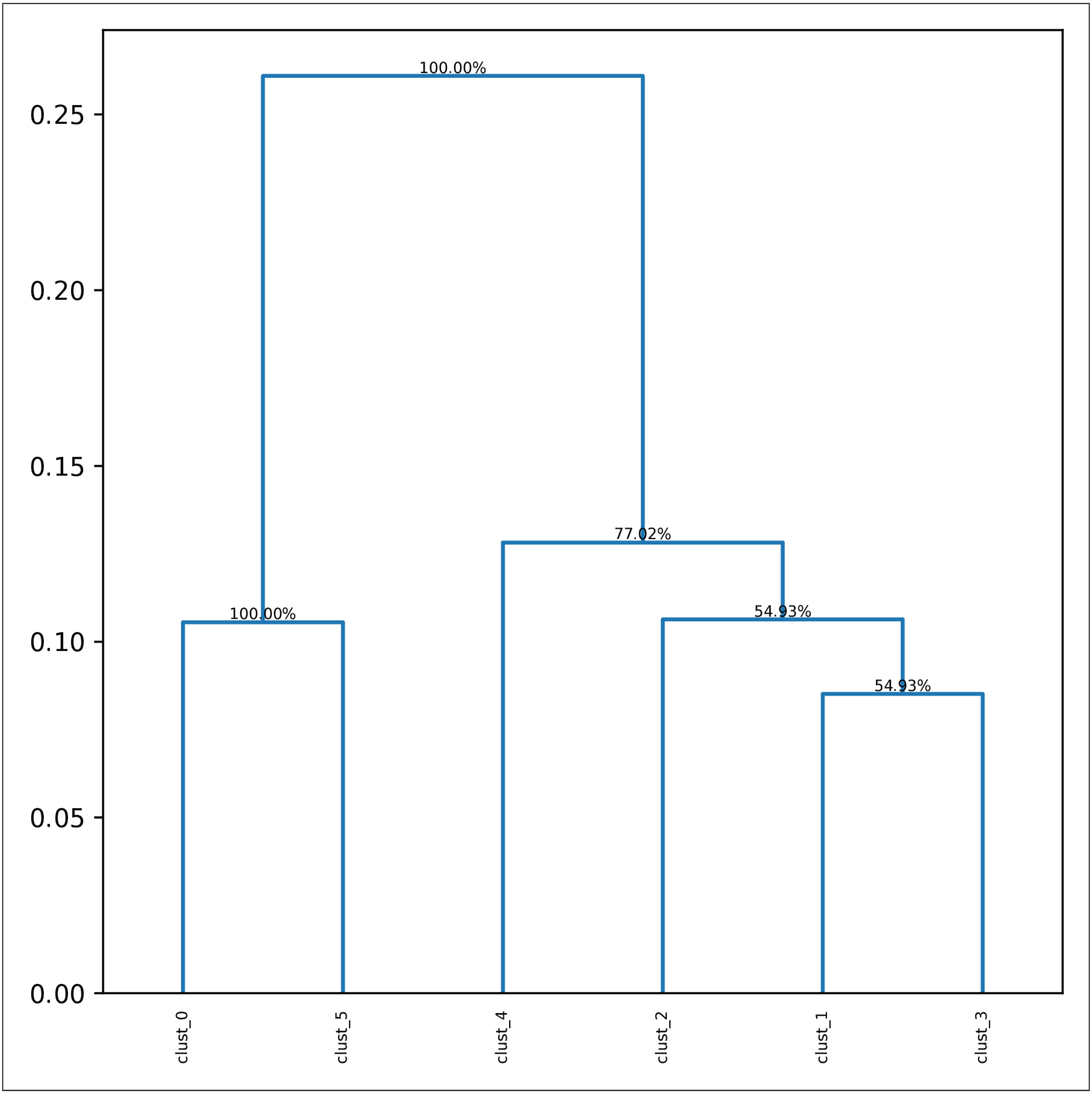

Sample: G0026 with 500 cells per cluster

Min. read depth: 10,  Mean UMIs per cluster:  10,604,950

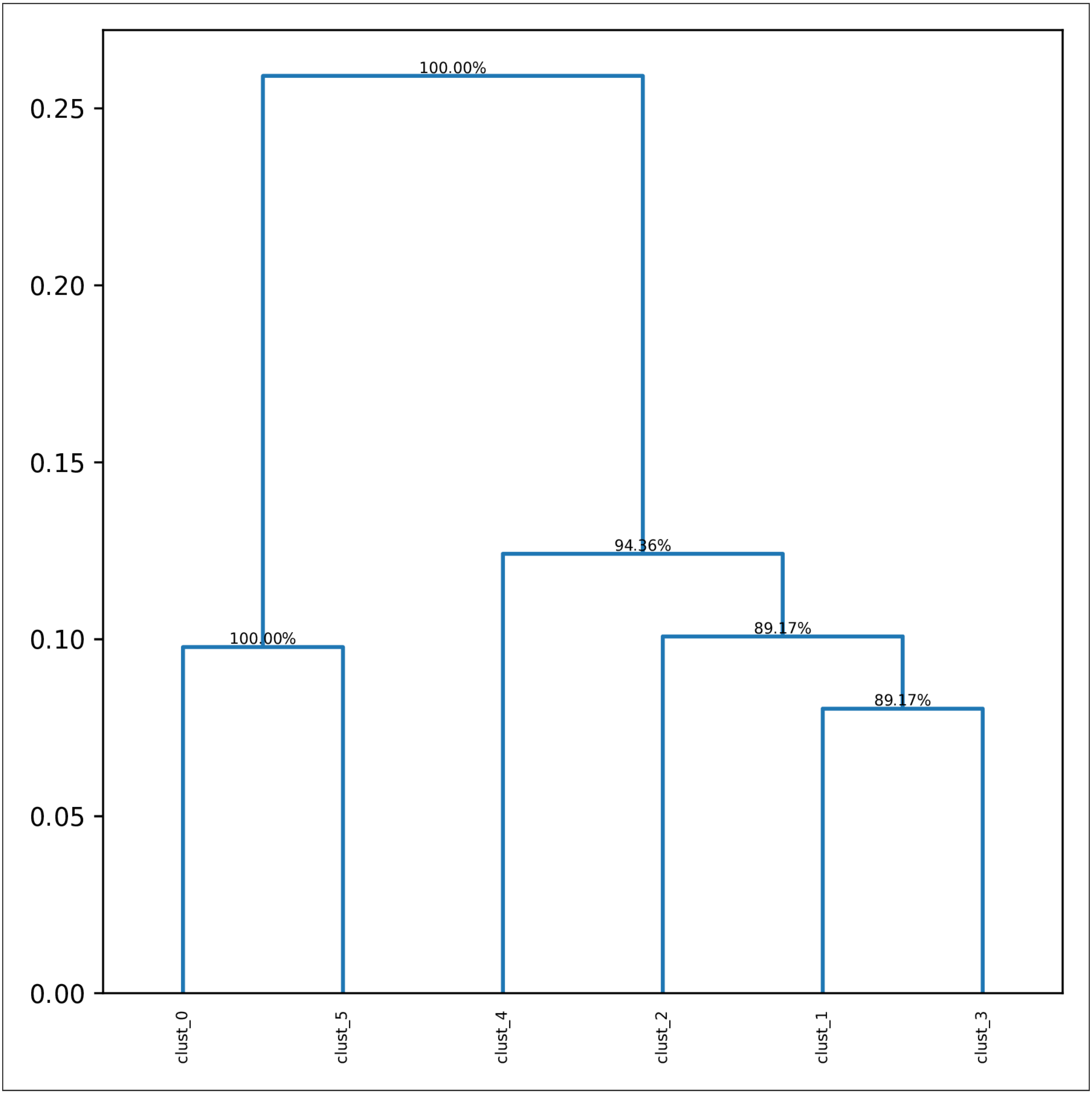

Sample: G0026 with 100 cells per cluster

Min. read depth: 10,  Mean UMIs per cluster:  2,517,773

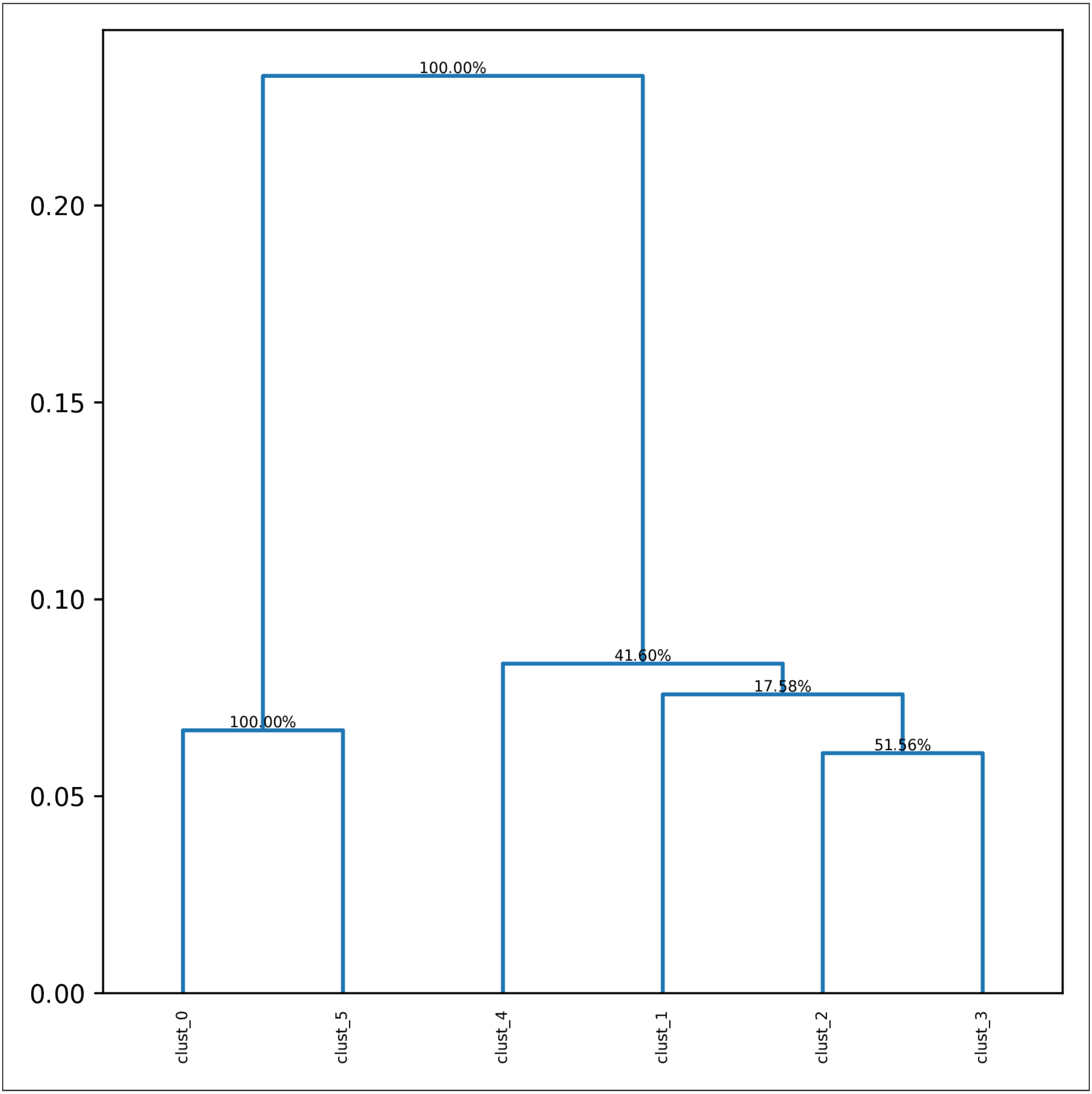

Sample: G0026 with 50 cells per cluster
Min. read depth: 10,  Mean UMIs per cluster:  1,241,452

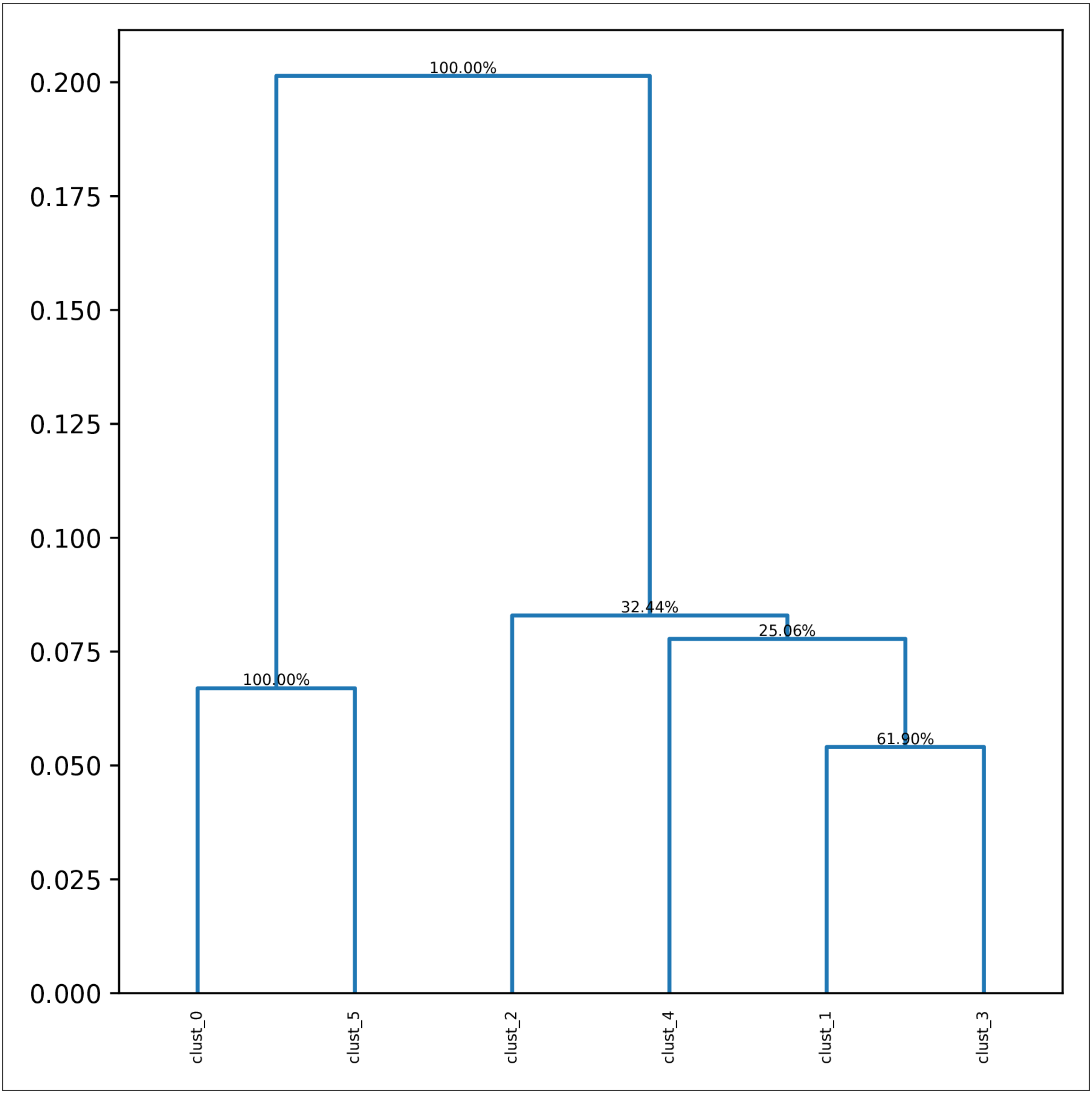

Sample: G0026 with 20 cells per cluster

Min. read depth: 10,  Mean UMIs per cluster:  502,748

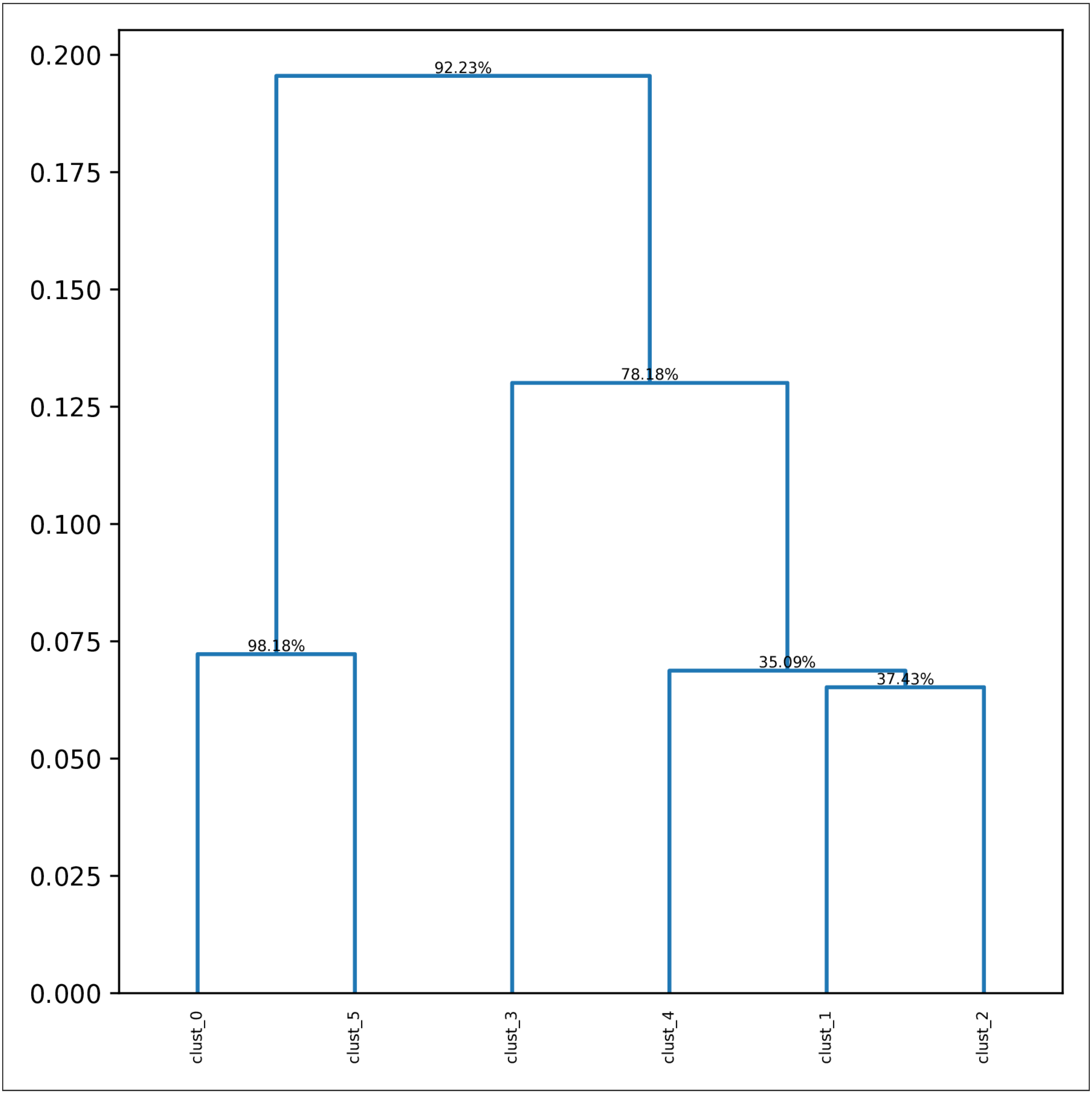

Sample: G0026 with 10 cells per cluster

Min. read depth: 10,  Mean UMIs per cluster:  285,272

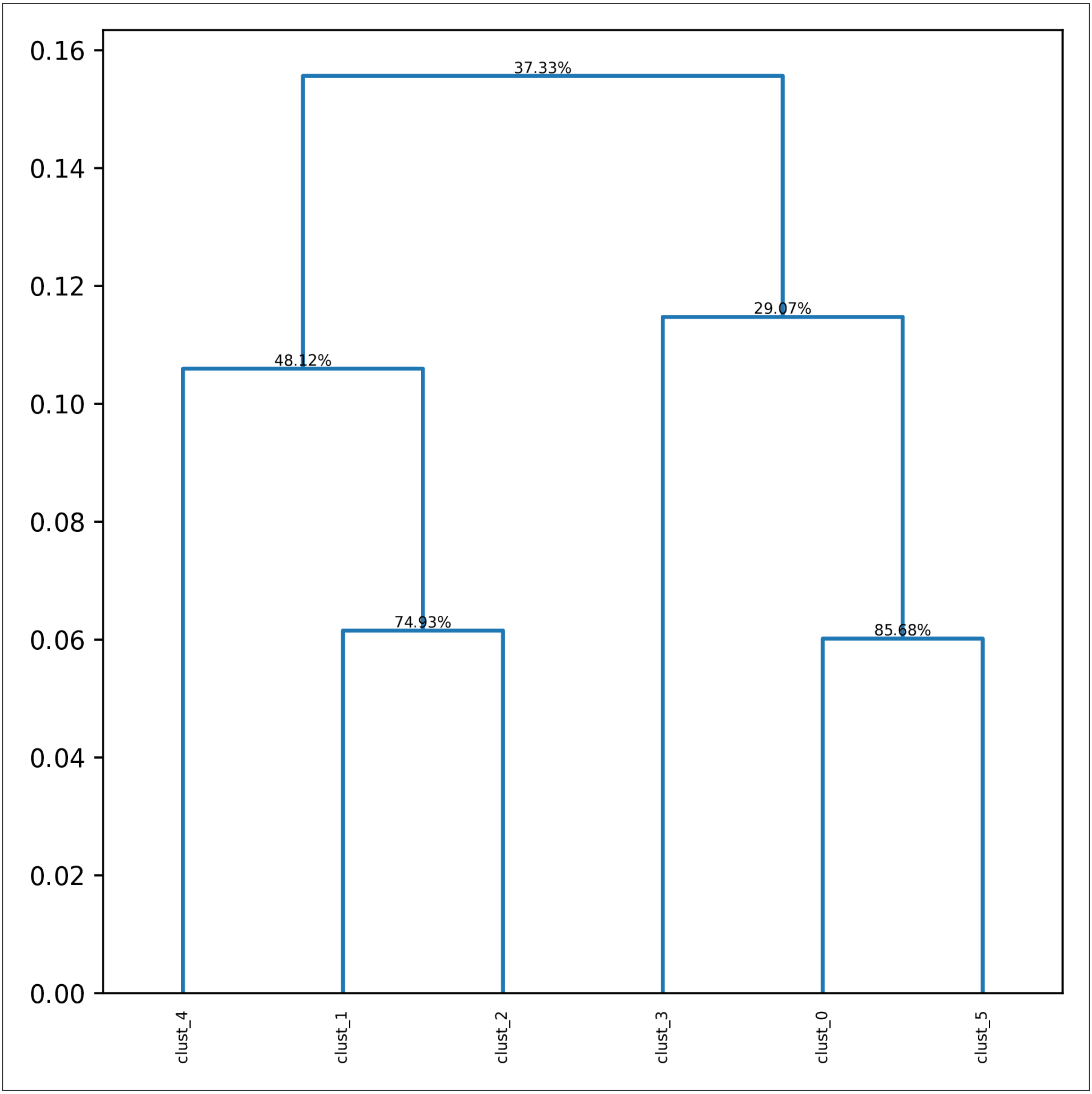

Sample: G0026 with 4 cells per cluster
Min. read depth: 10,  Mean UMIs per cluster:  116,809

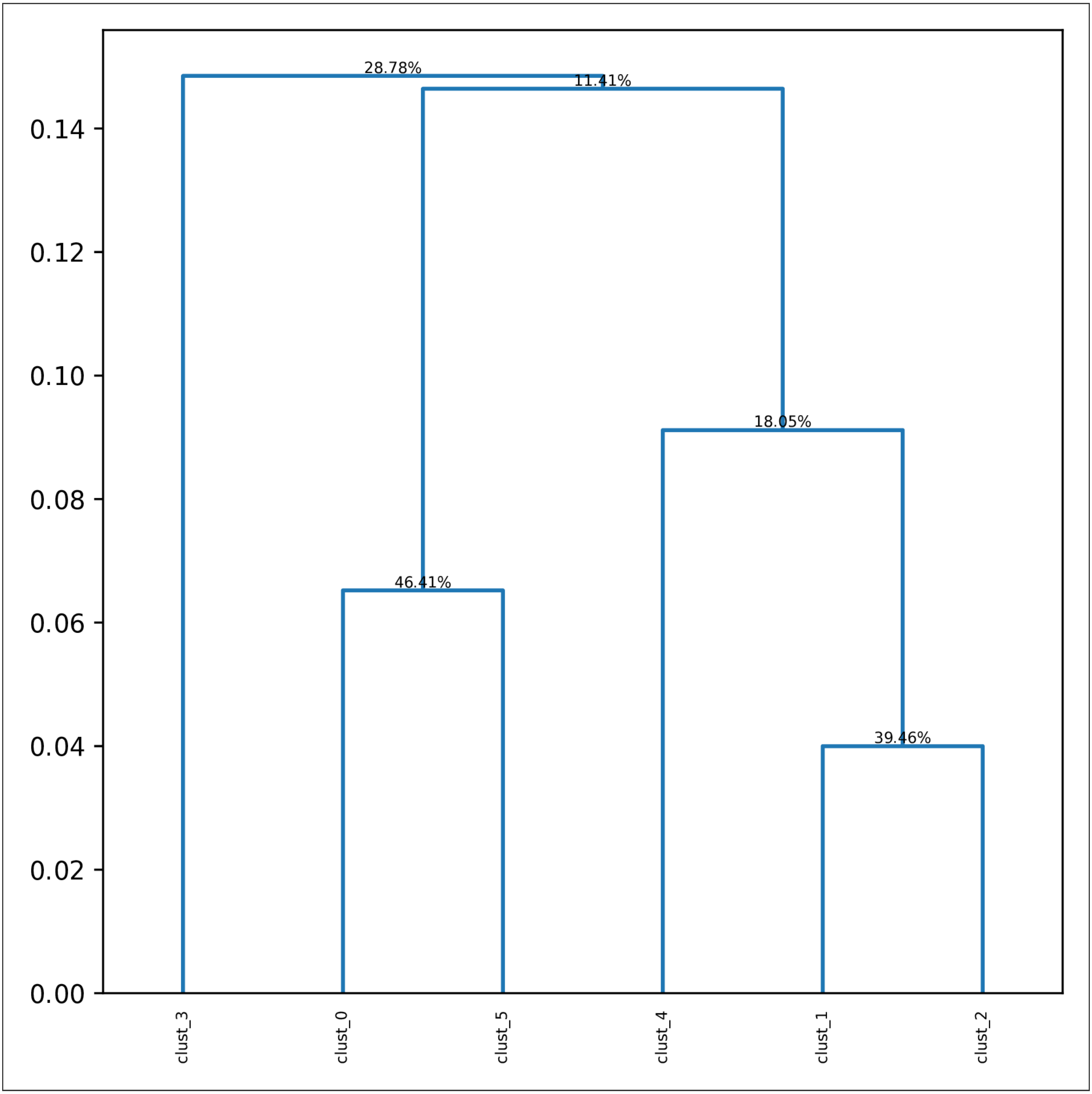

Sample: G0027 with all cells per cluster

Min. read depth: 10,  Mean UMIs per cluster:  9,862,423

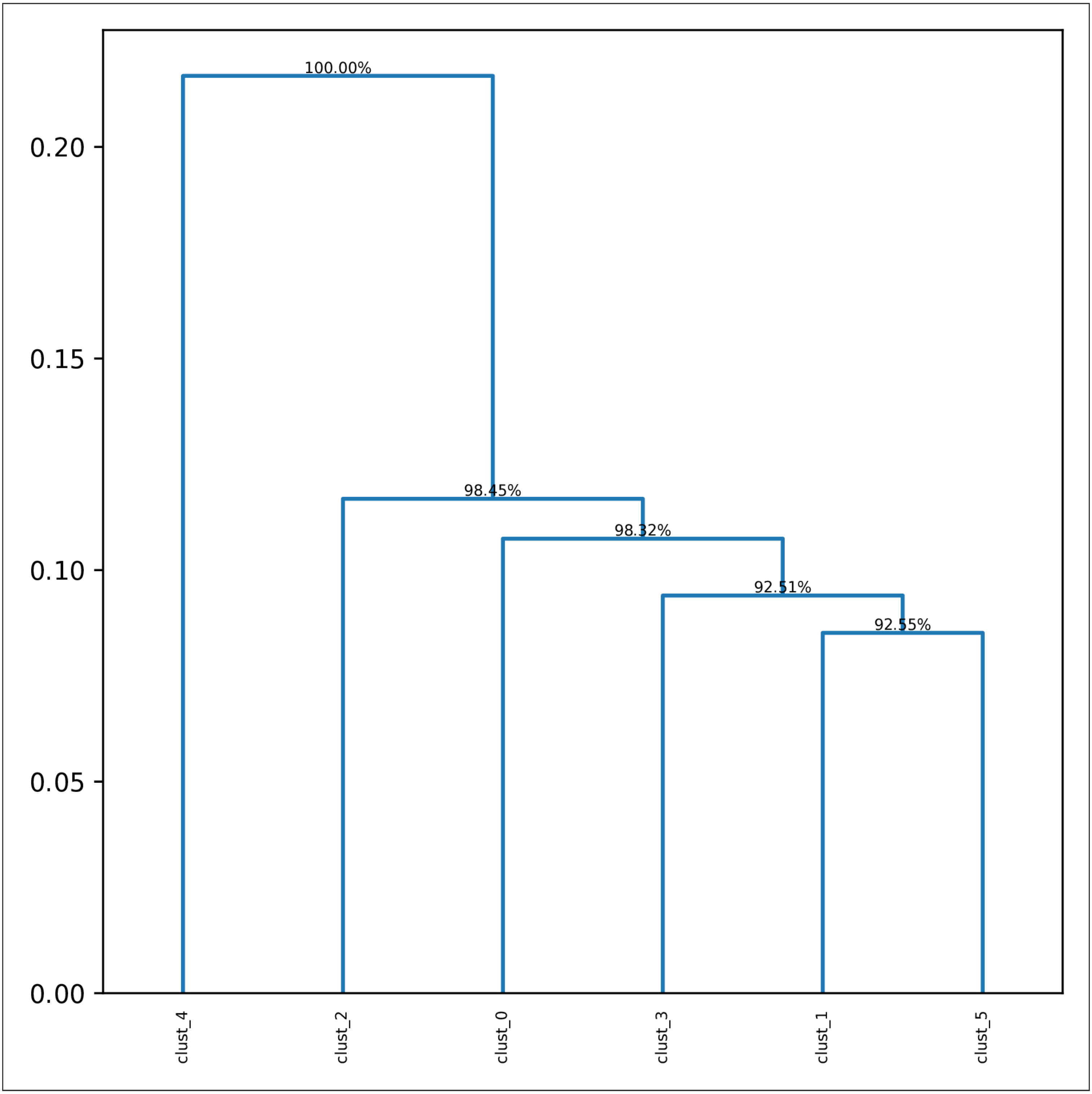

Sample: G0027 with 500 cells per cluster

Min. read depth: 10,  Mean UMIs per cluster:  7,898,730

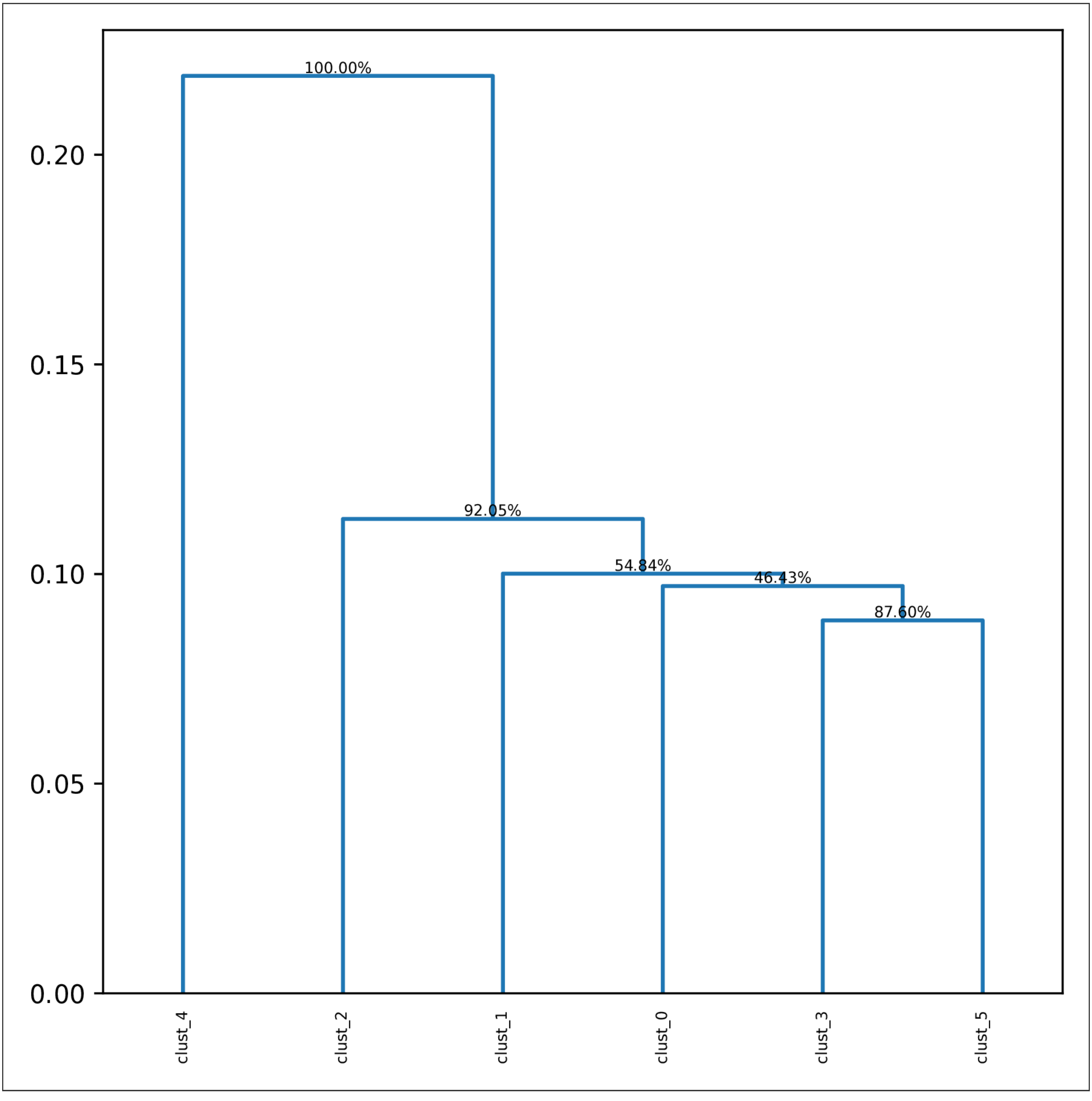

Sample: G0027 with 100 cells per cluster

Min. read depth: 10,  Mean UMIs per cluster:  1,720,702

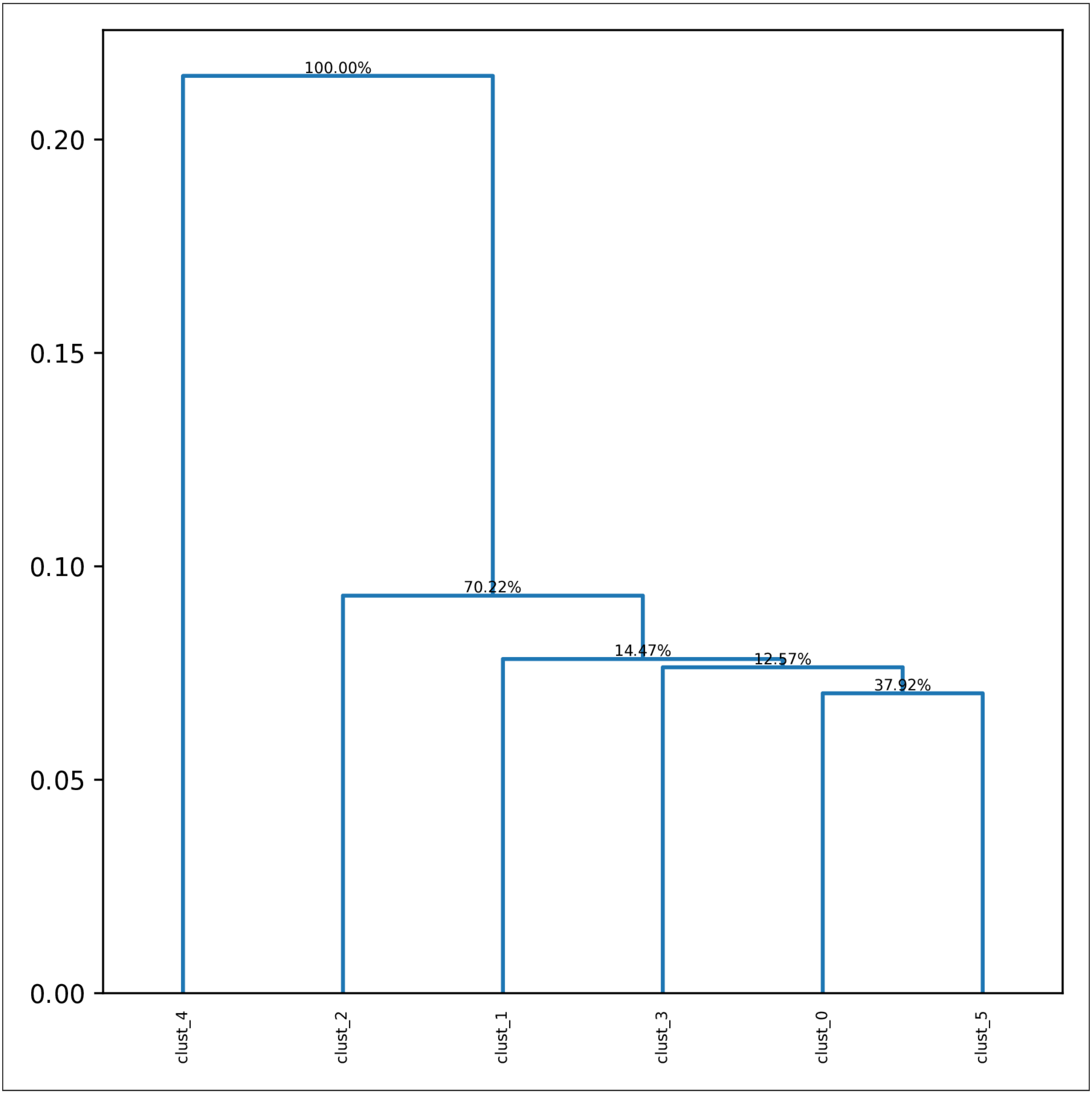

Sample: G0027 with 50 cells per cluster

Min. read depth: 10, Mean UMIs per cluster: 900,081

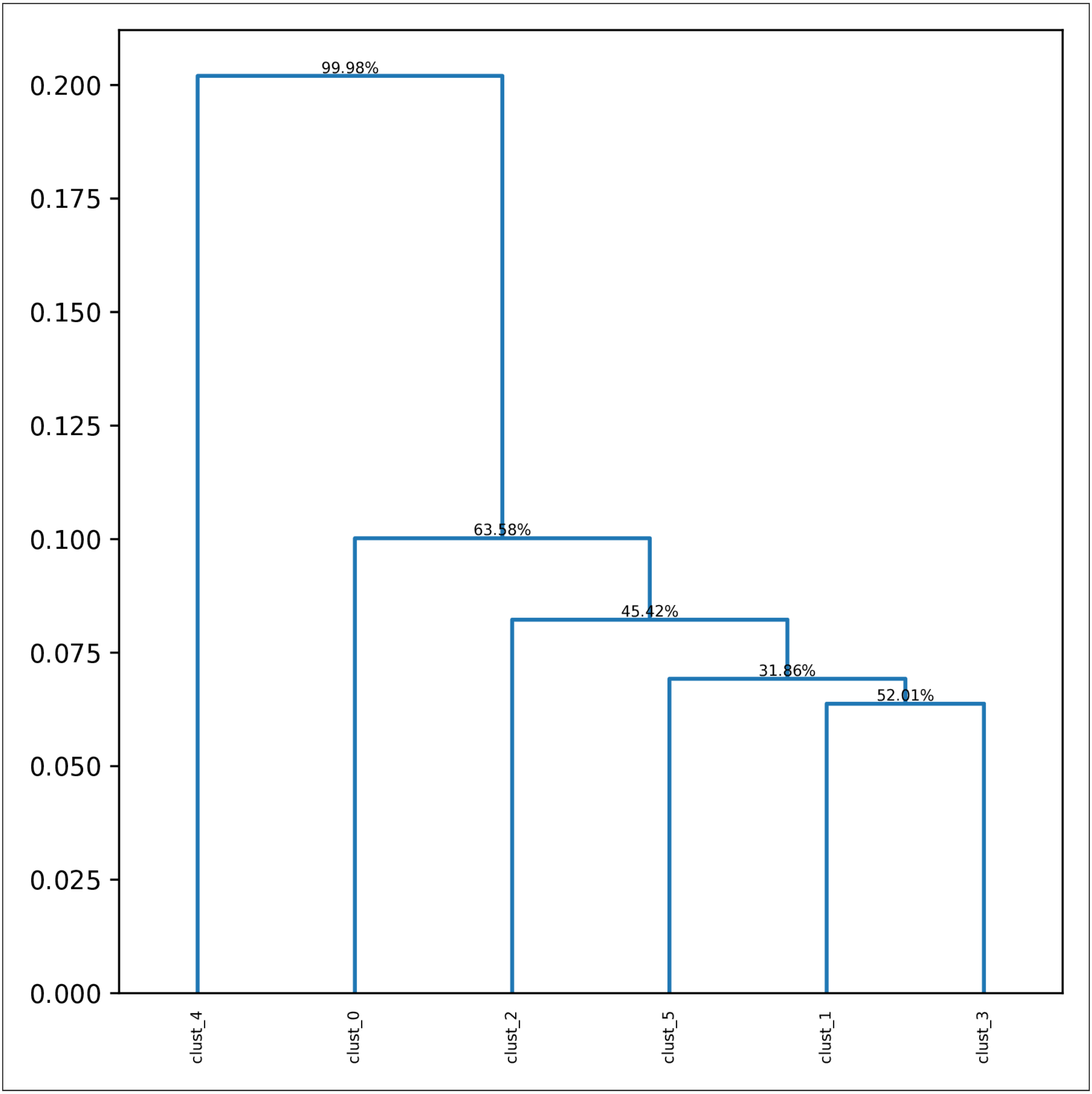

Sample: G0027 with 20 cells per cluster
Min. read depth: 10,  Mean UMIs per cluster:  349,348

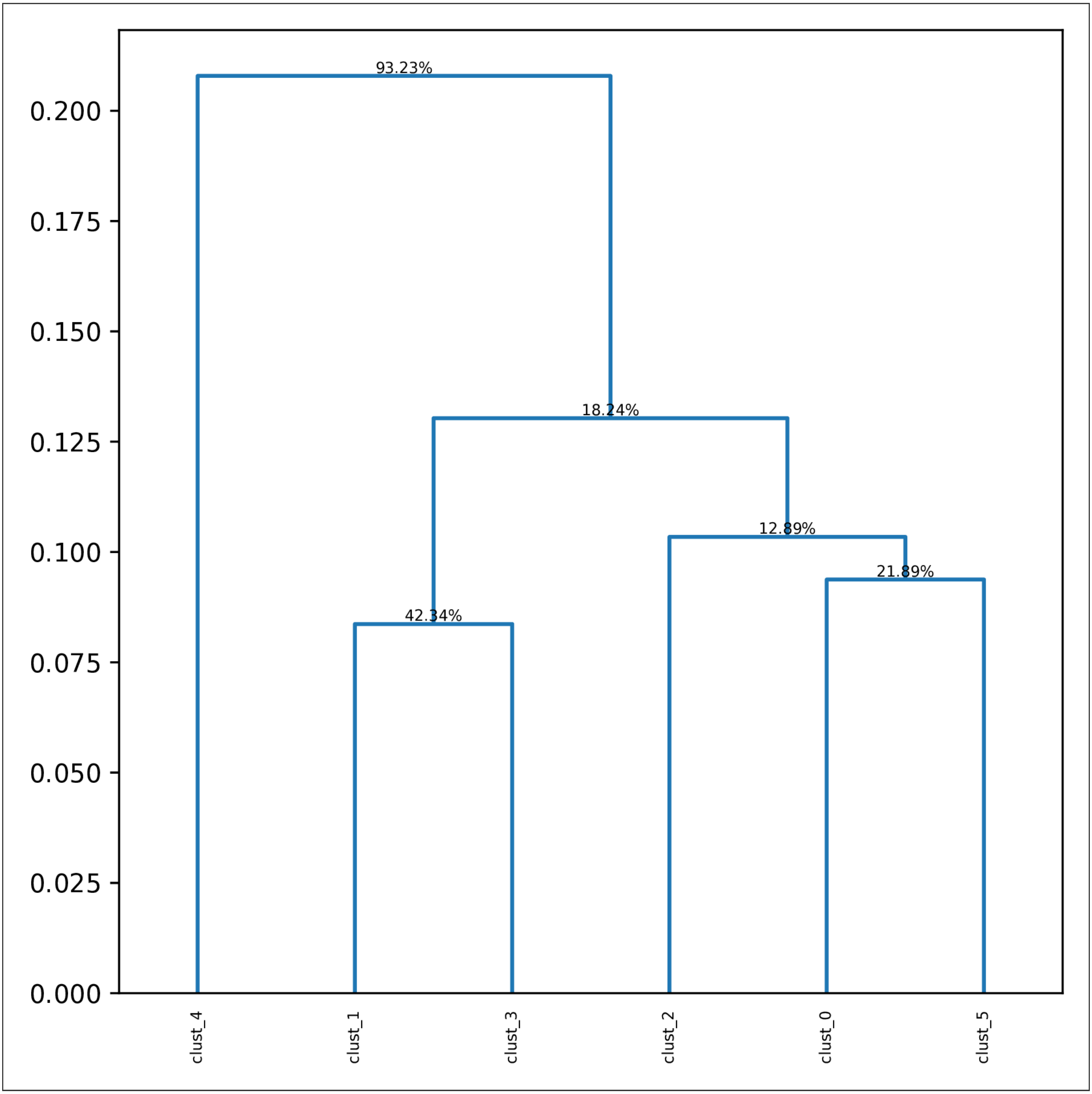

Sample: G0027 with 10 cells per cluster

Min. read depth: 10,  Mean UMIs per cluster:  150,970

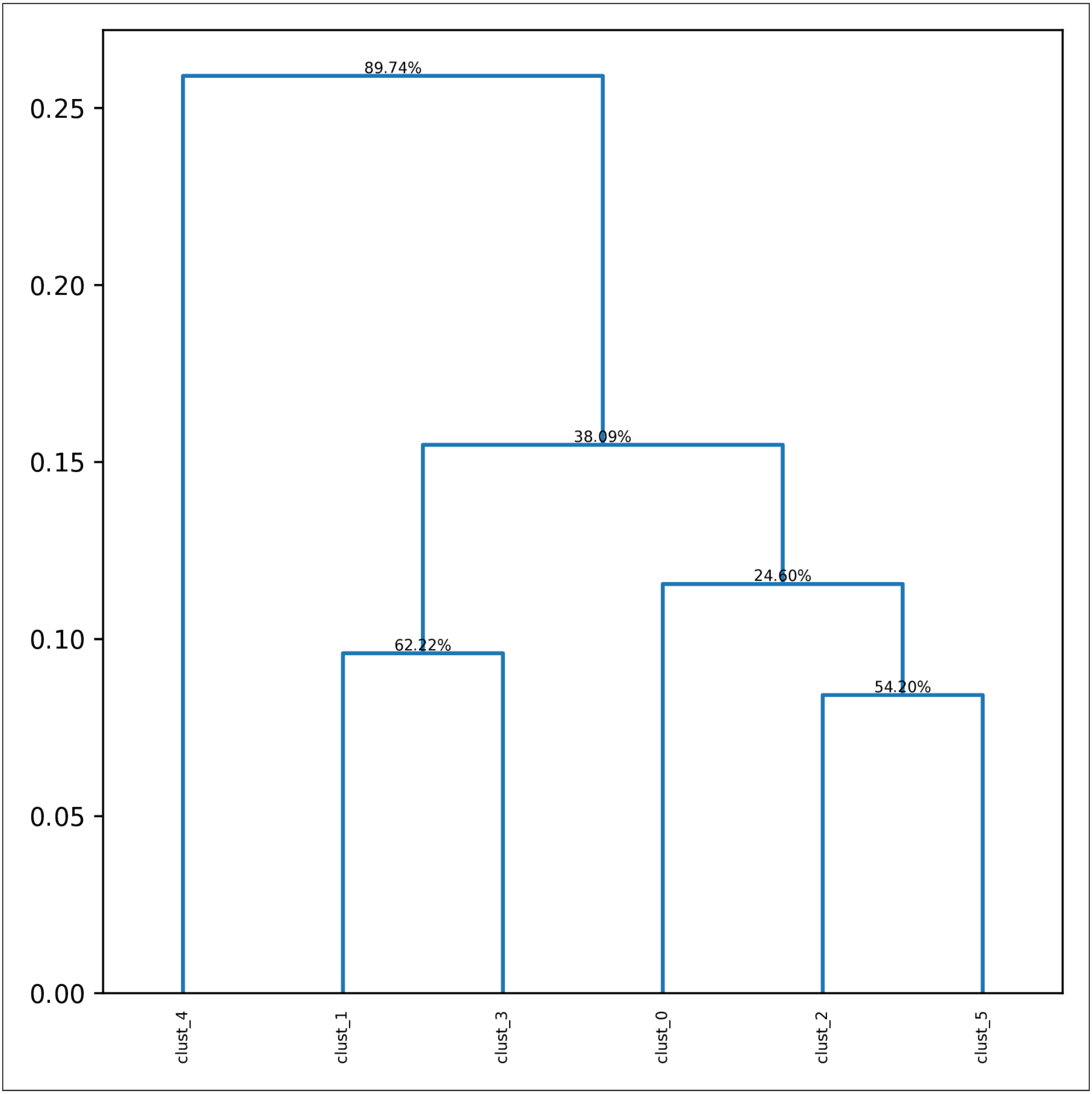

Sample: G0027 with 5 cells per cluster

Min. read depth: 10,  Mean UMIs per cluster:  96,858

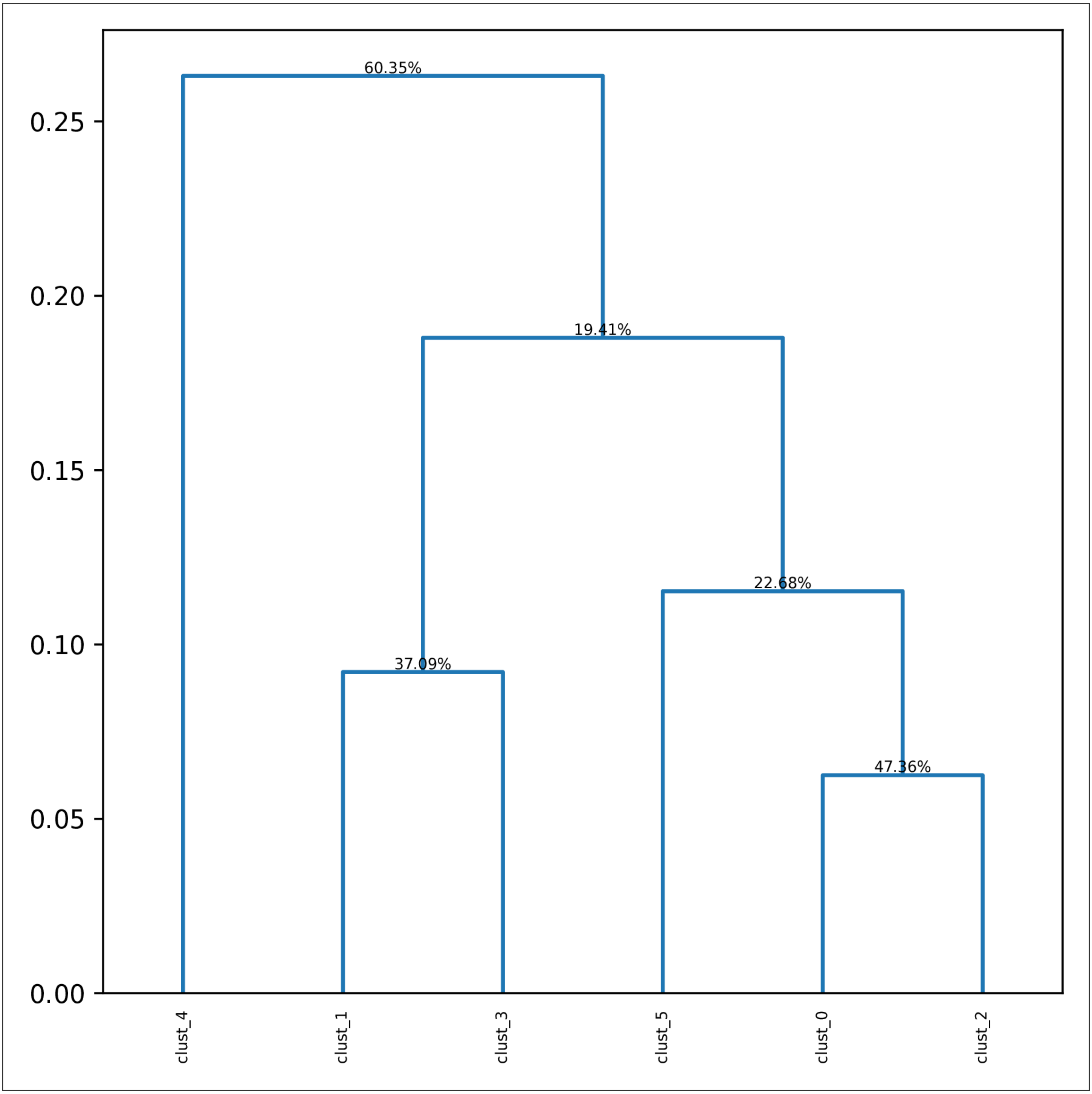

Sample: G0027 with 4 cells per cluster

Min. read depth: 10,  Mean UMIs per cluster:  84,327

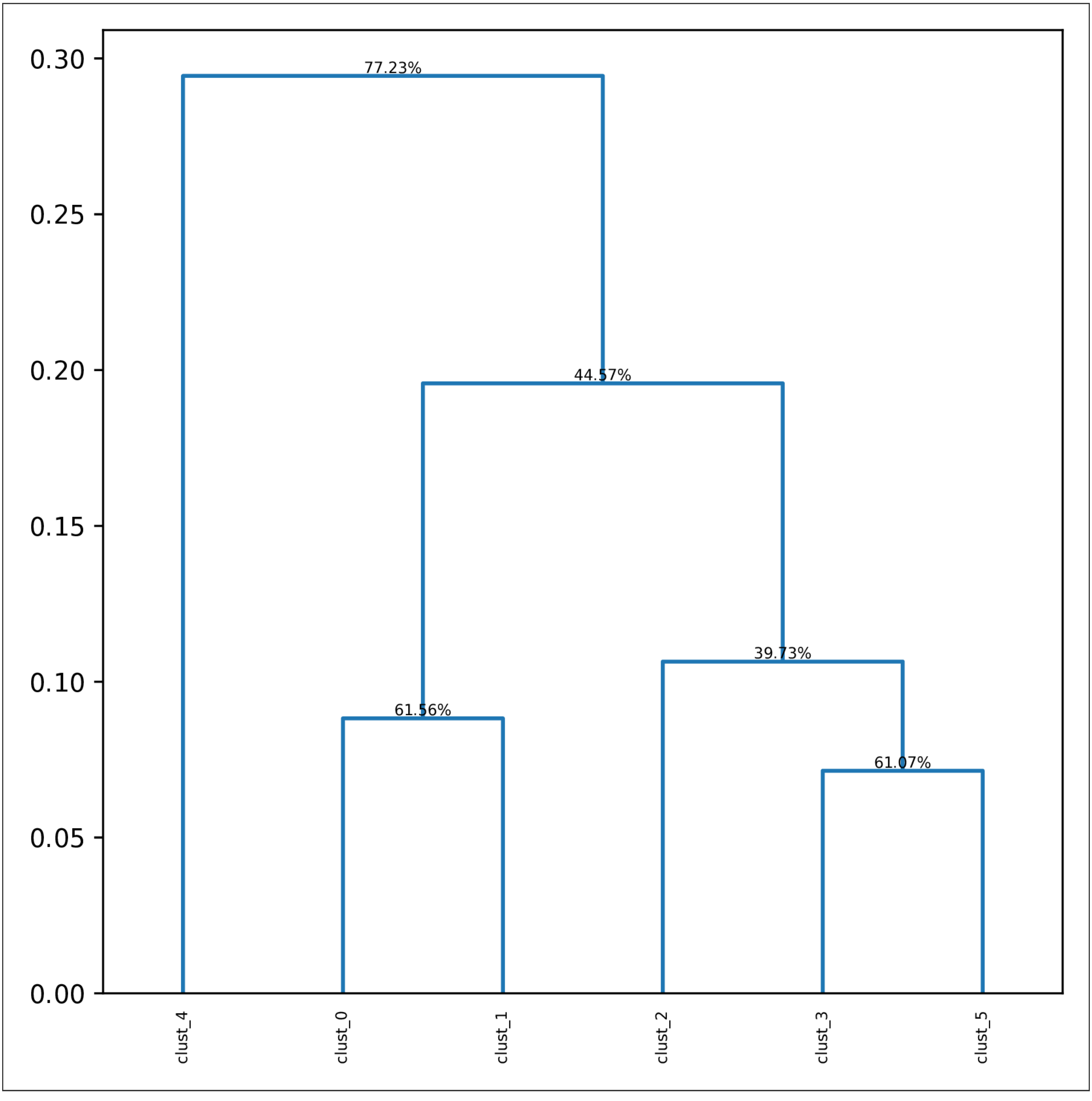

Sample: G0027 with 3 cells per cluster

Min. read depth: 10,  Mean UMIs per cluster:  64,494

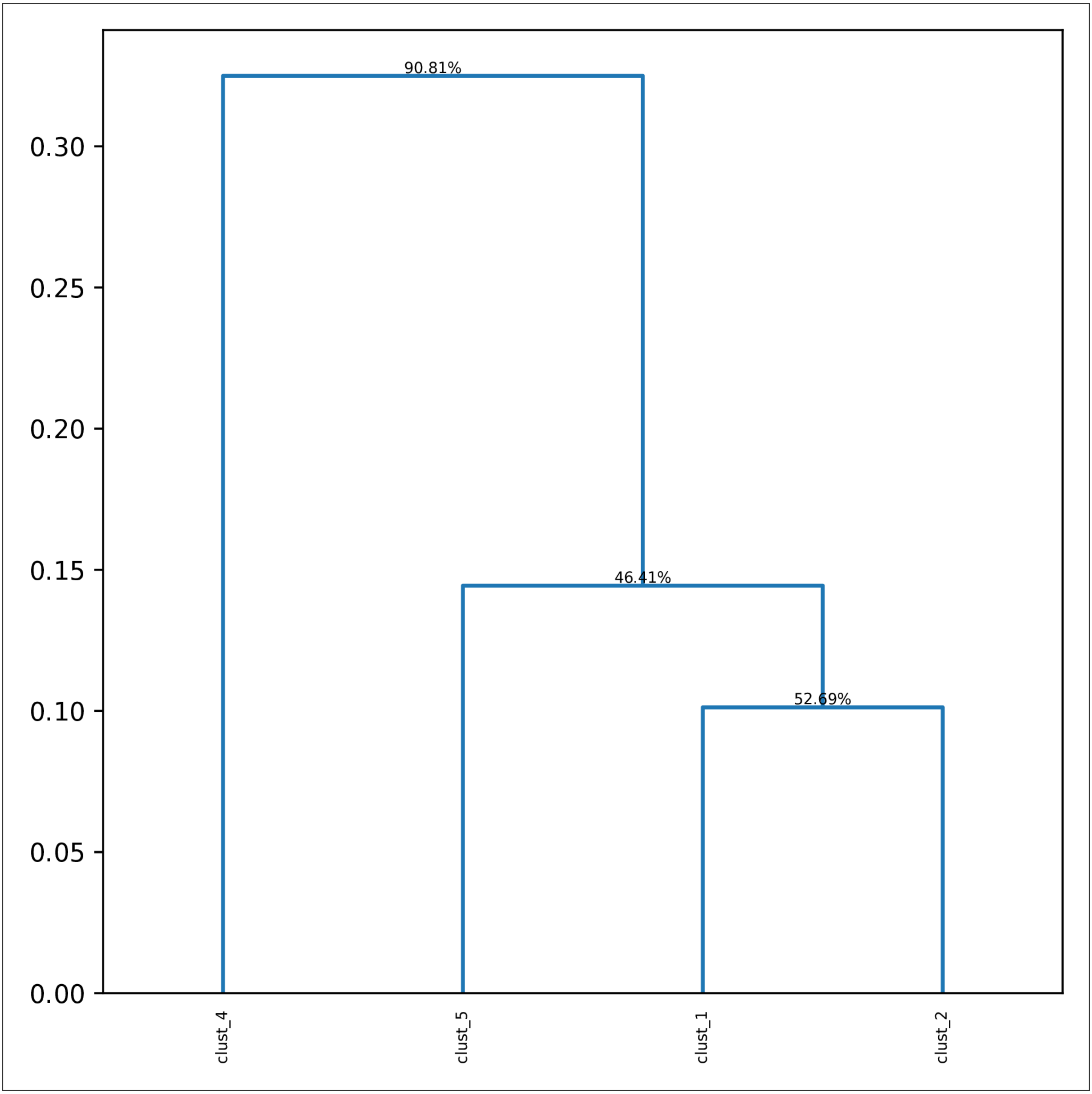

Sample: G0027 with 2 cells per cluster
Min. read depth: 10,  Mean UMIs per cluster:  53,163

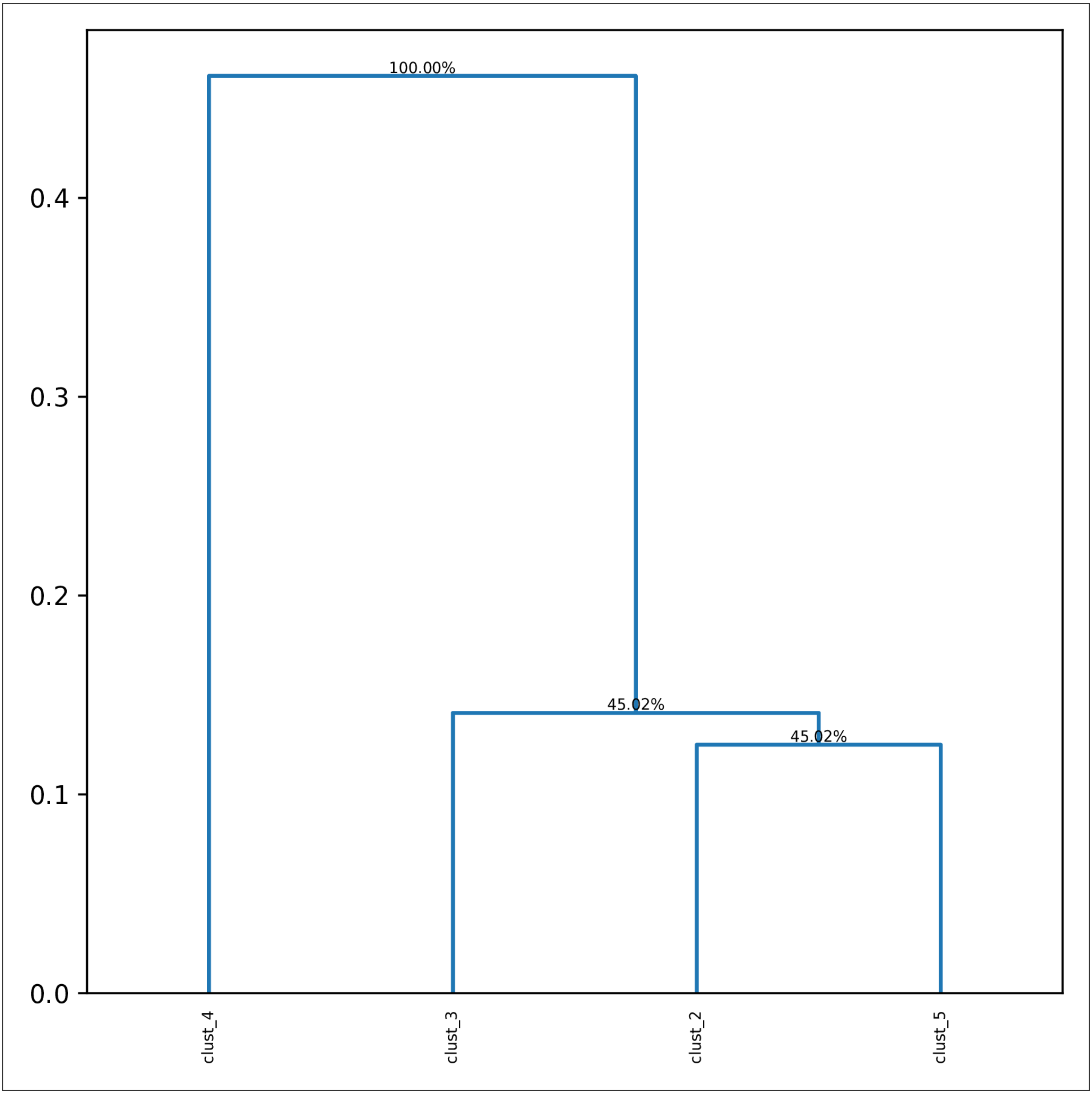

Sample: G0028 with all cells per cluster

Min. read depth: 10,  Mean UMIs per cluster:  25,974,842

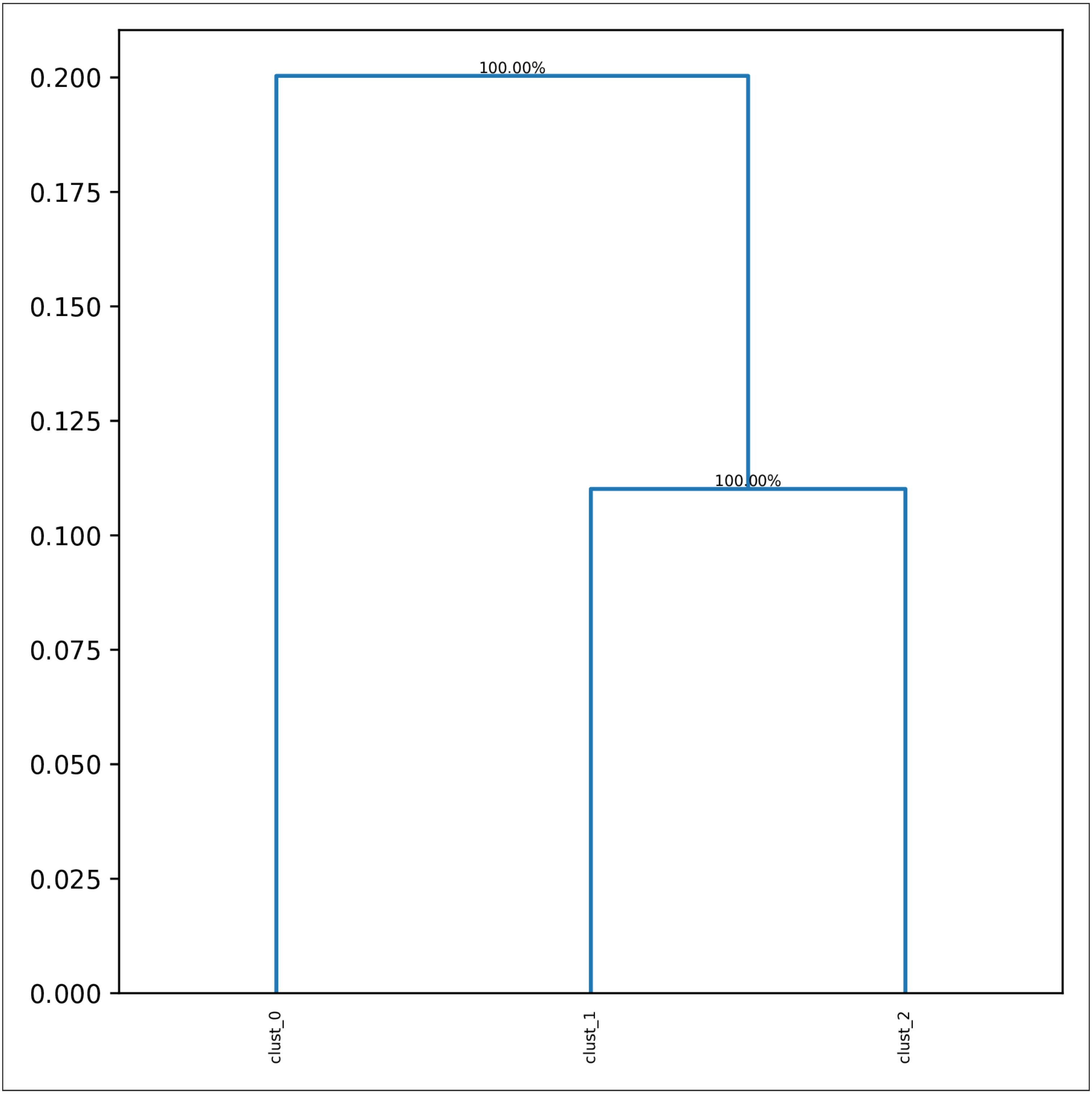

Sample: G0028 with 500 cells per cluster

Min. read depth: 10,  Mean UMIs per cluster:  8,673,360

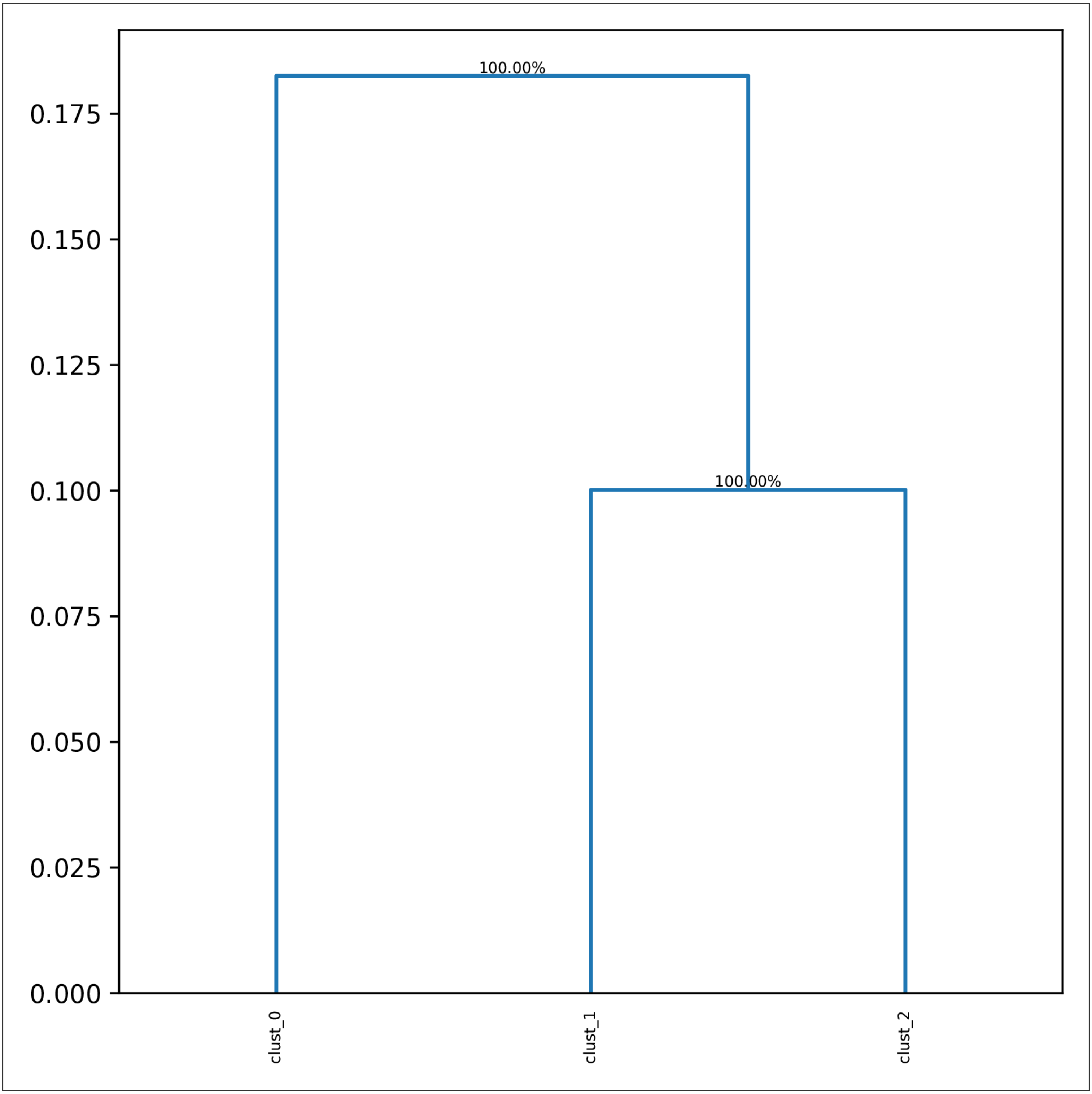

Sample: G0028 with 100 cells per cluster

Min. read depth: 10,  Mean UMIs per cluster:  1,709,076

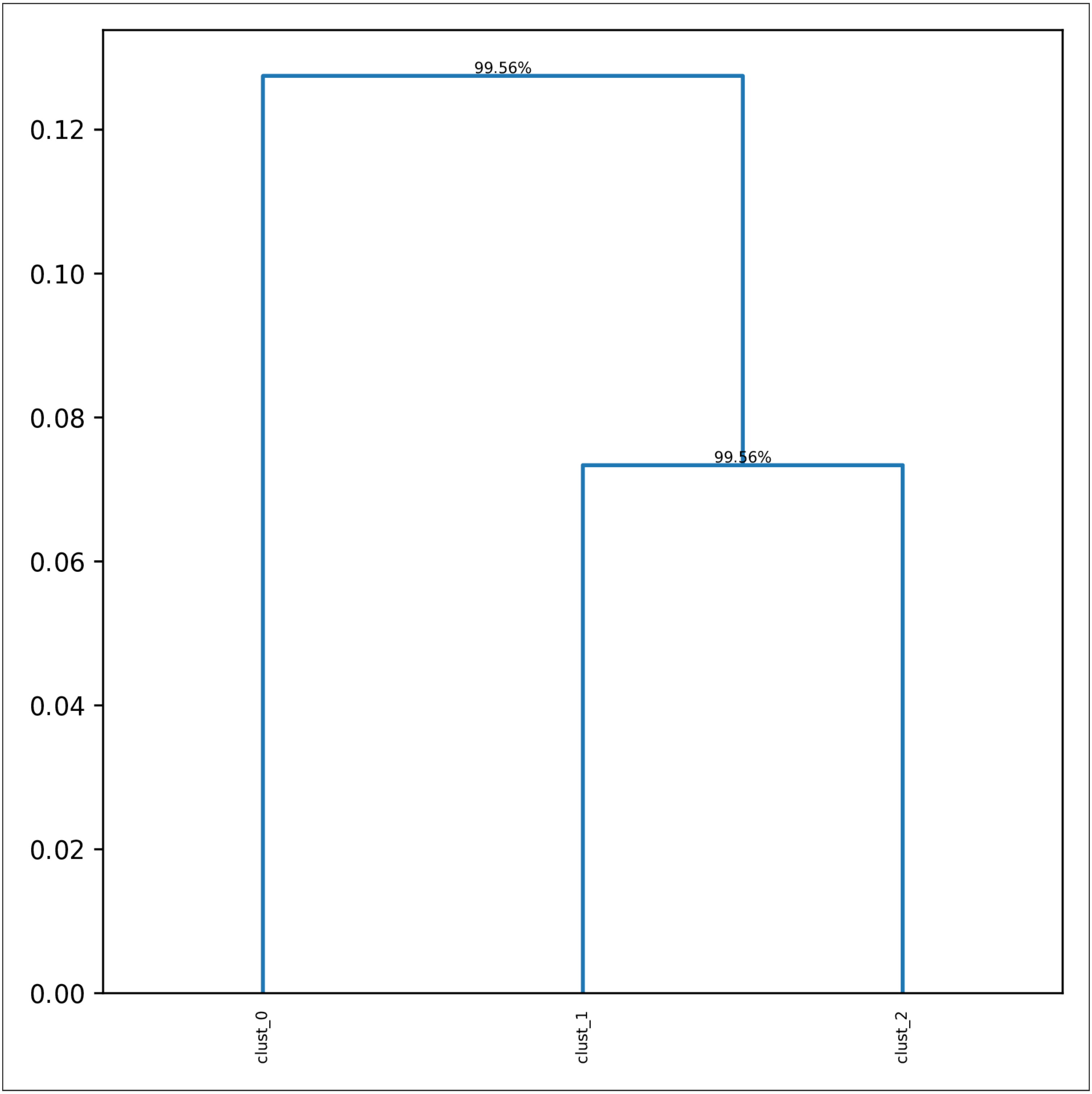

Sample: G0028 with 50 cells per cluster

Min. read depth: 10,  Mean UMIs per cluster:  913,176

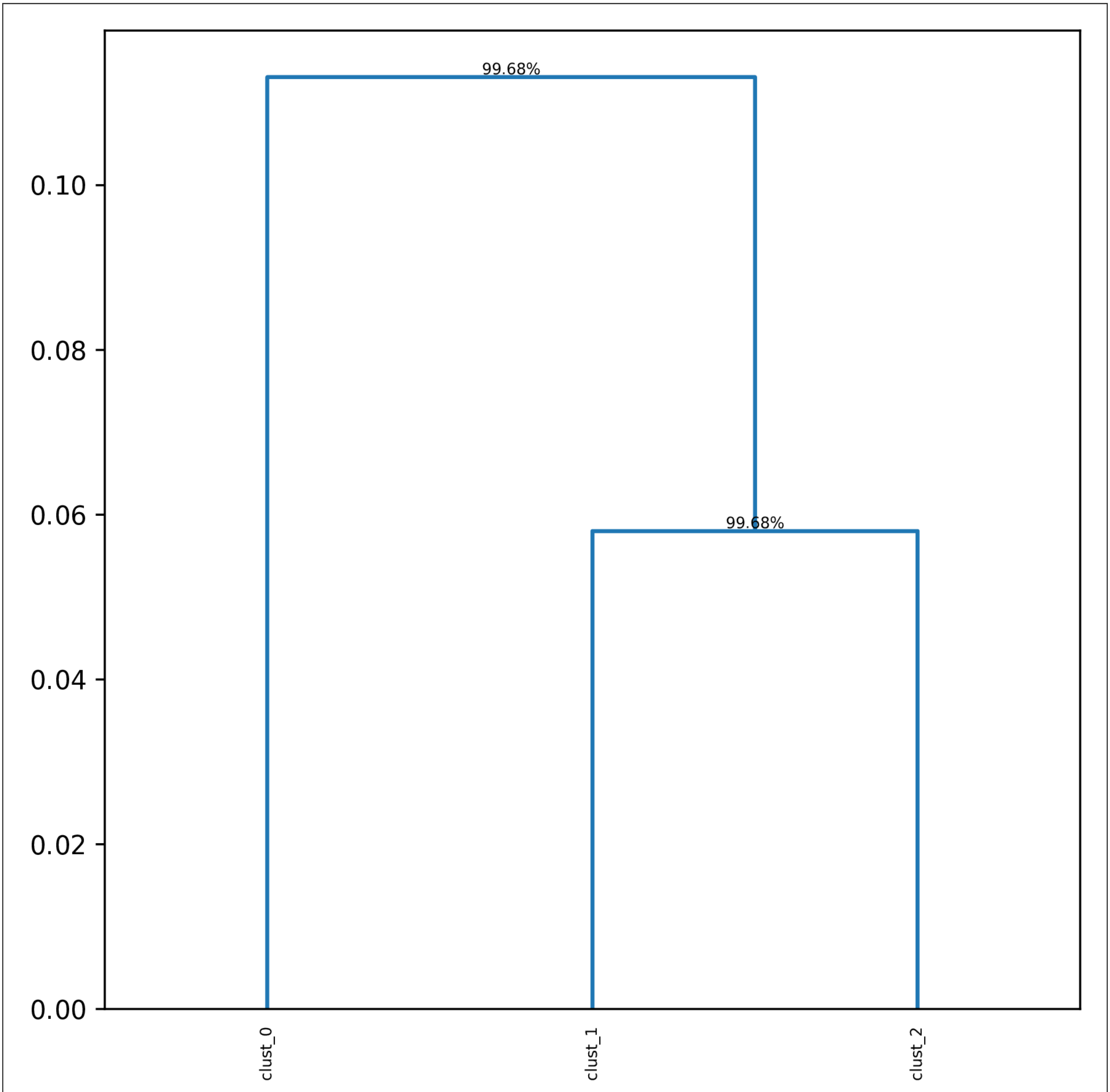

Sample: G0028 with 20 cells per cluster
Min. read depth: 10,  Mean UMIs per cluster:  410,256

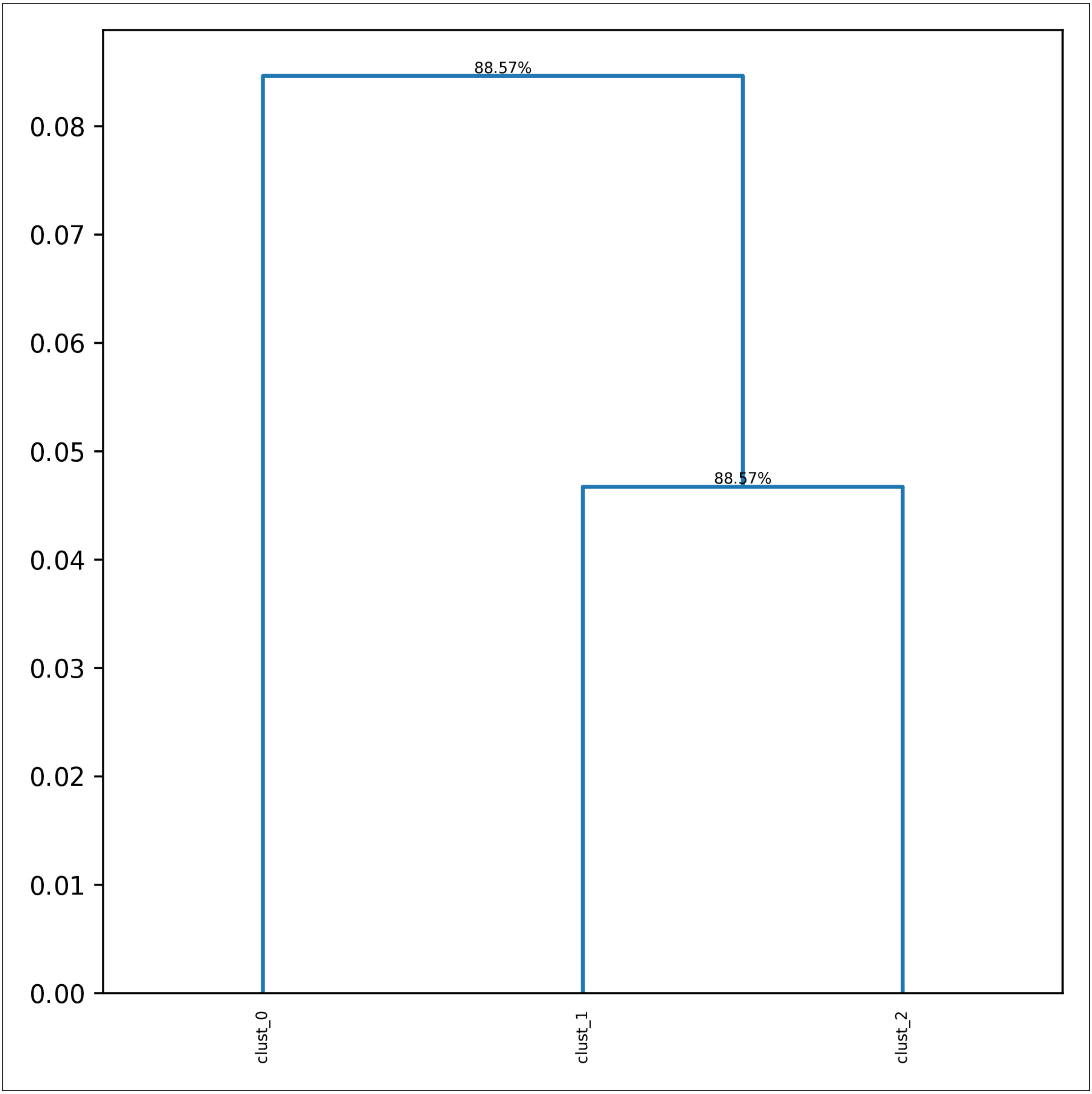

Sample: G0028 with 10 cells per cluster

Min. read depth: 10,  Mean UMIs per cluster:  194,148

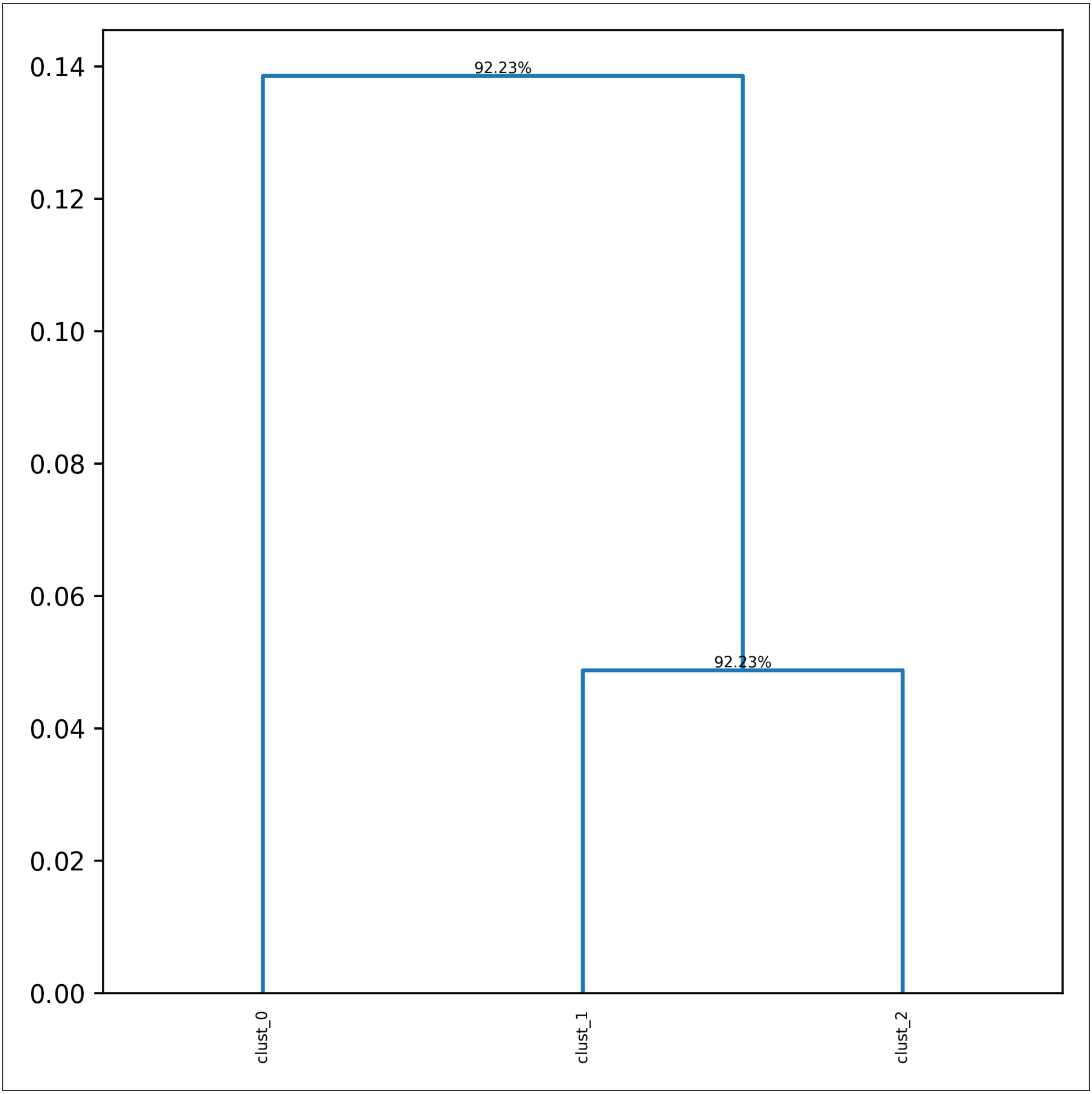

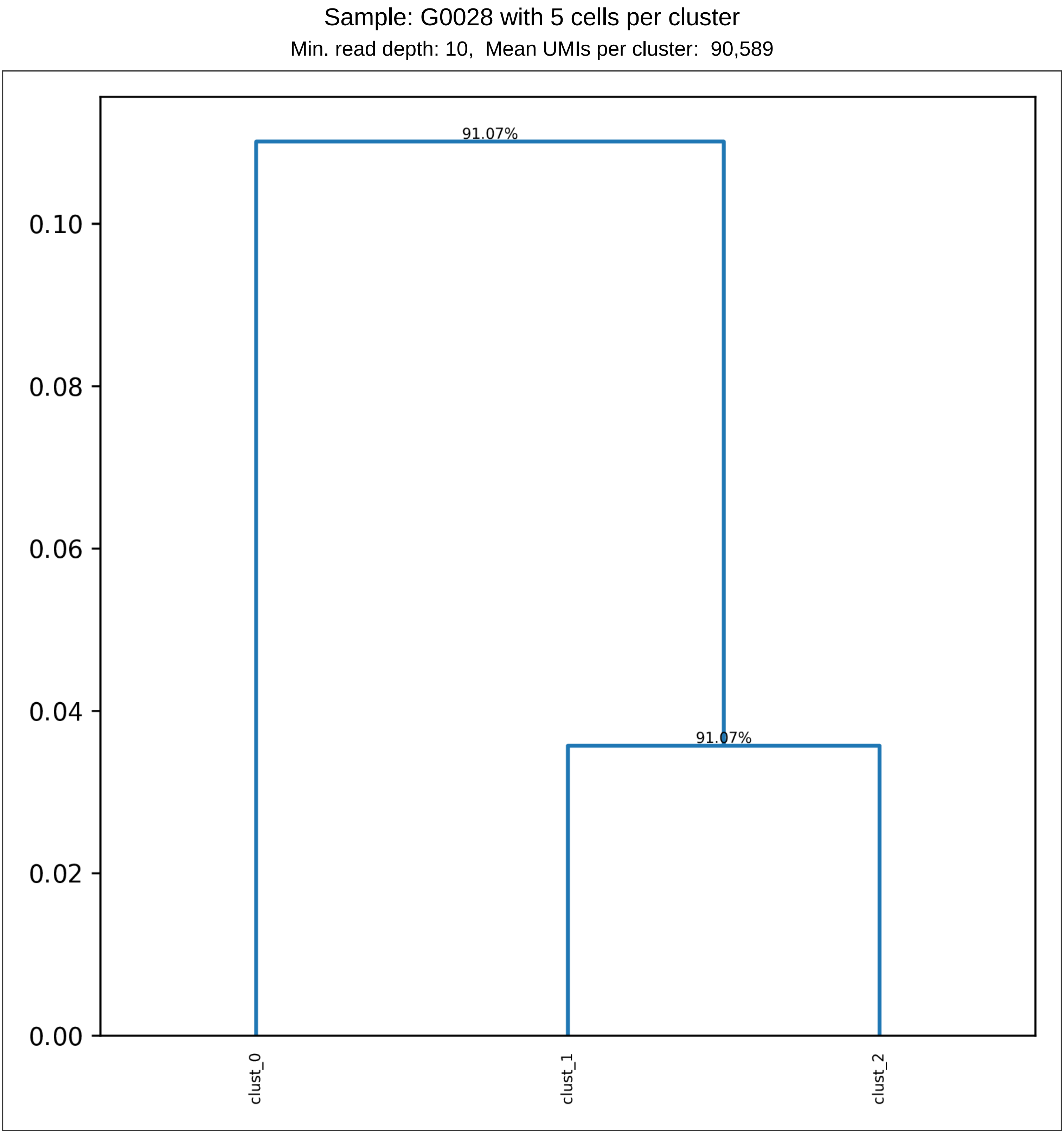

Sample: G0028 with 5 cells per cluster
Min. read depth: 10, Mean UMIs per cluster: 90,589
91.07%
91.07%
0.10
0.08
0.06
0.04
0.02
0.00
clust_0
clust_1
clust_2

Sample: G0028 with 4 cells per cluster

Min. read depth: 10,  Mean UMIs per cluster:  58,267

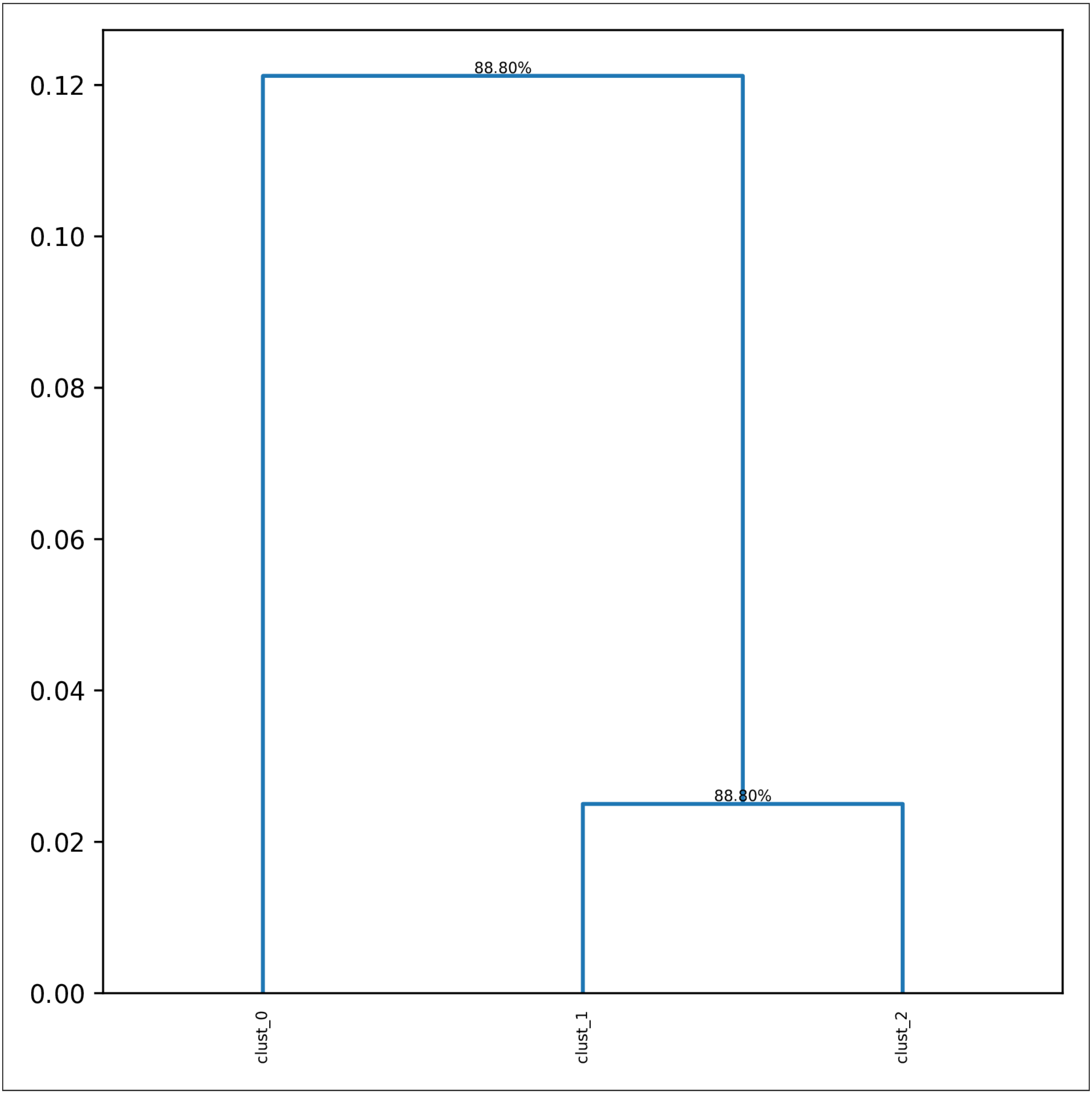

Sample: G0029 with all cells per cluster

Min. read depth: 10,  Mean UMIs per cluster:  11,903,754

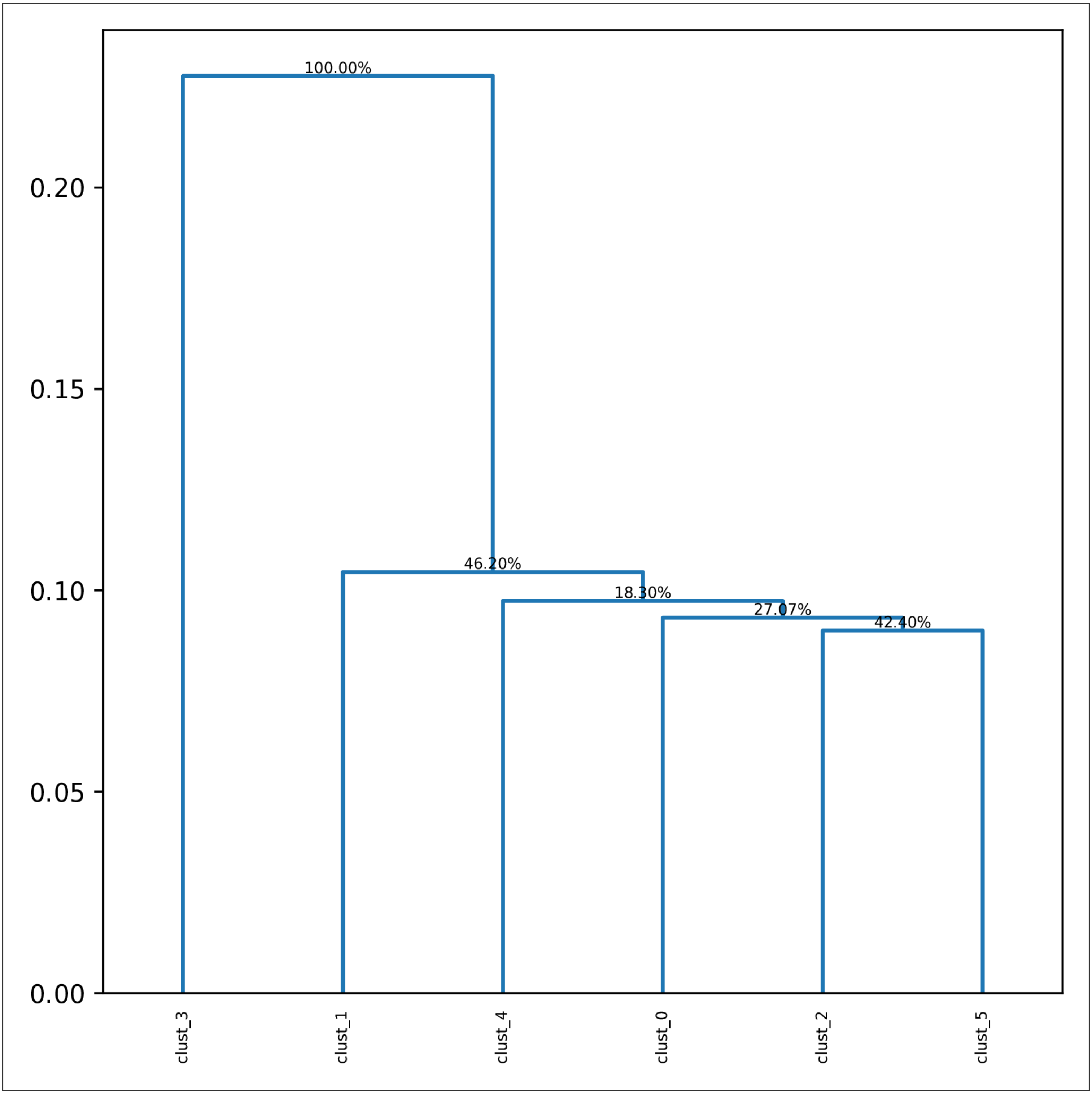

Sample: G0029 with 500 cells per cluster

Min. read depth: 10,  Mean UMIs per cluster:  8,468,528

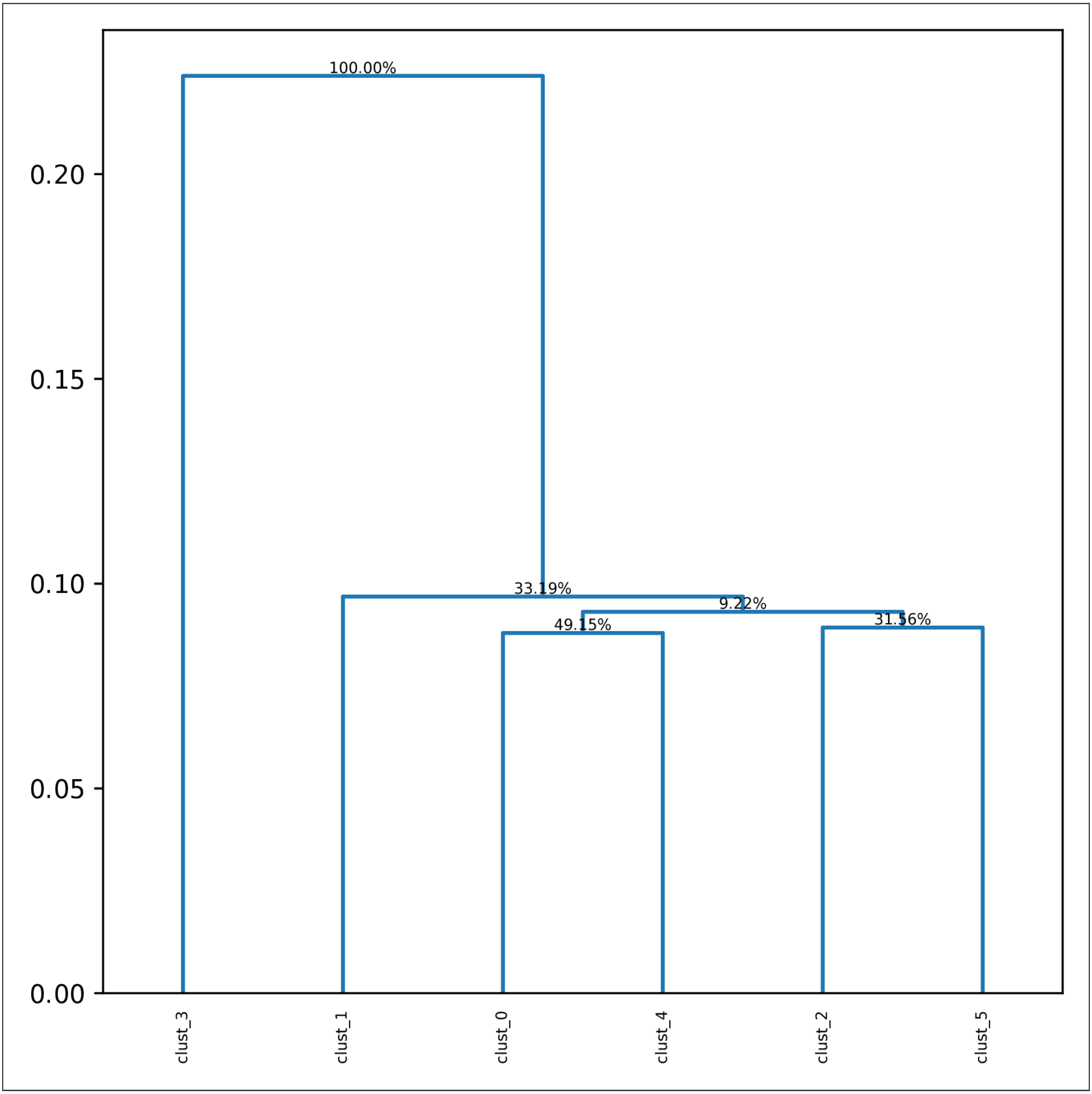

Sample: G0029 with 100 cells per cluster

Min. read depth: 10,  Mean UMIs per cluster:  2,144,514

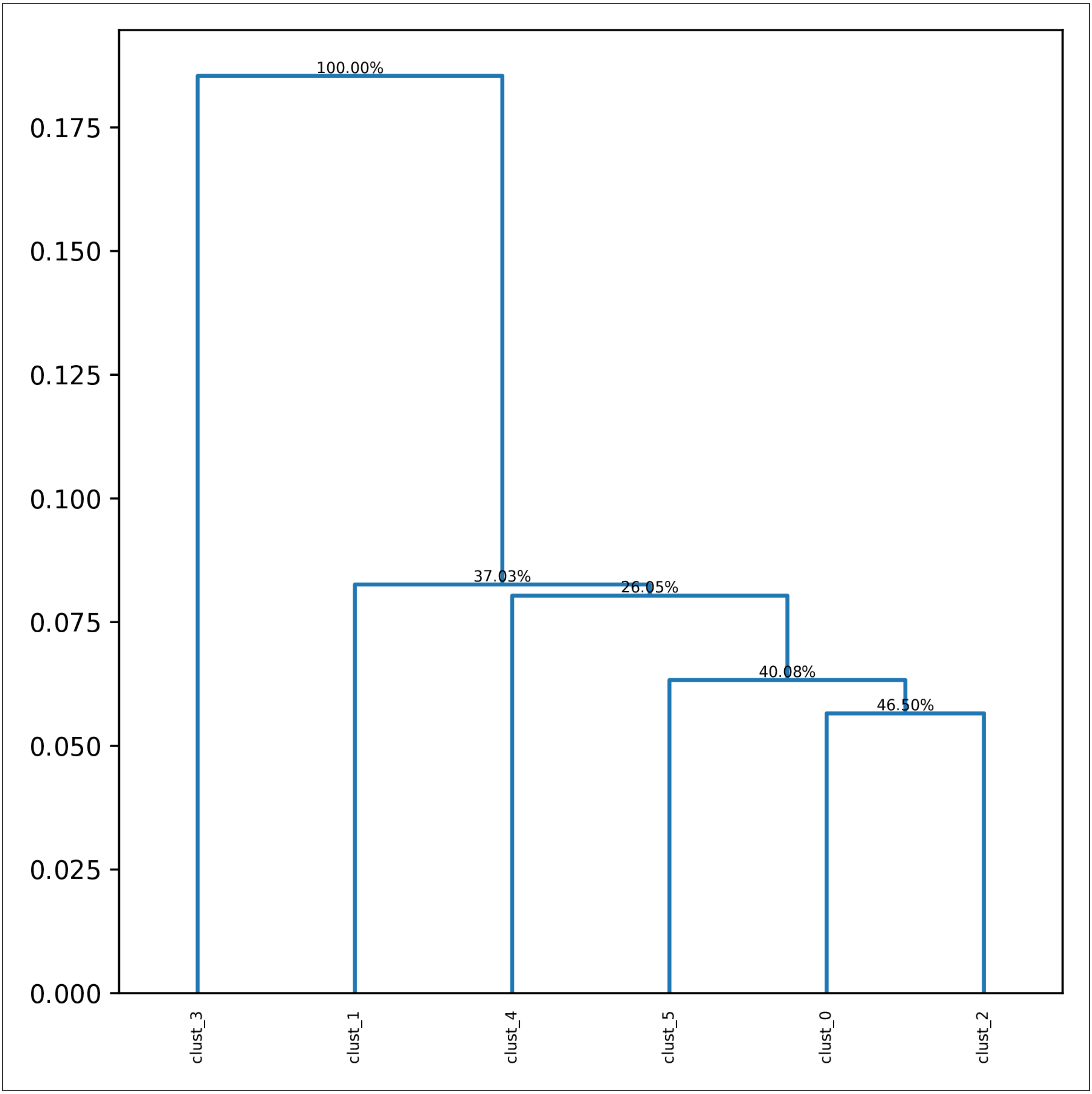

Sample: G0029 with 50 cells per cluster

Min. read depth: 10,  Mean UMIs per cluster:  1,095,023

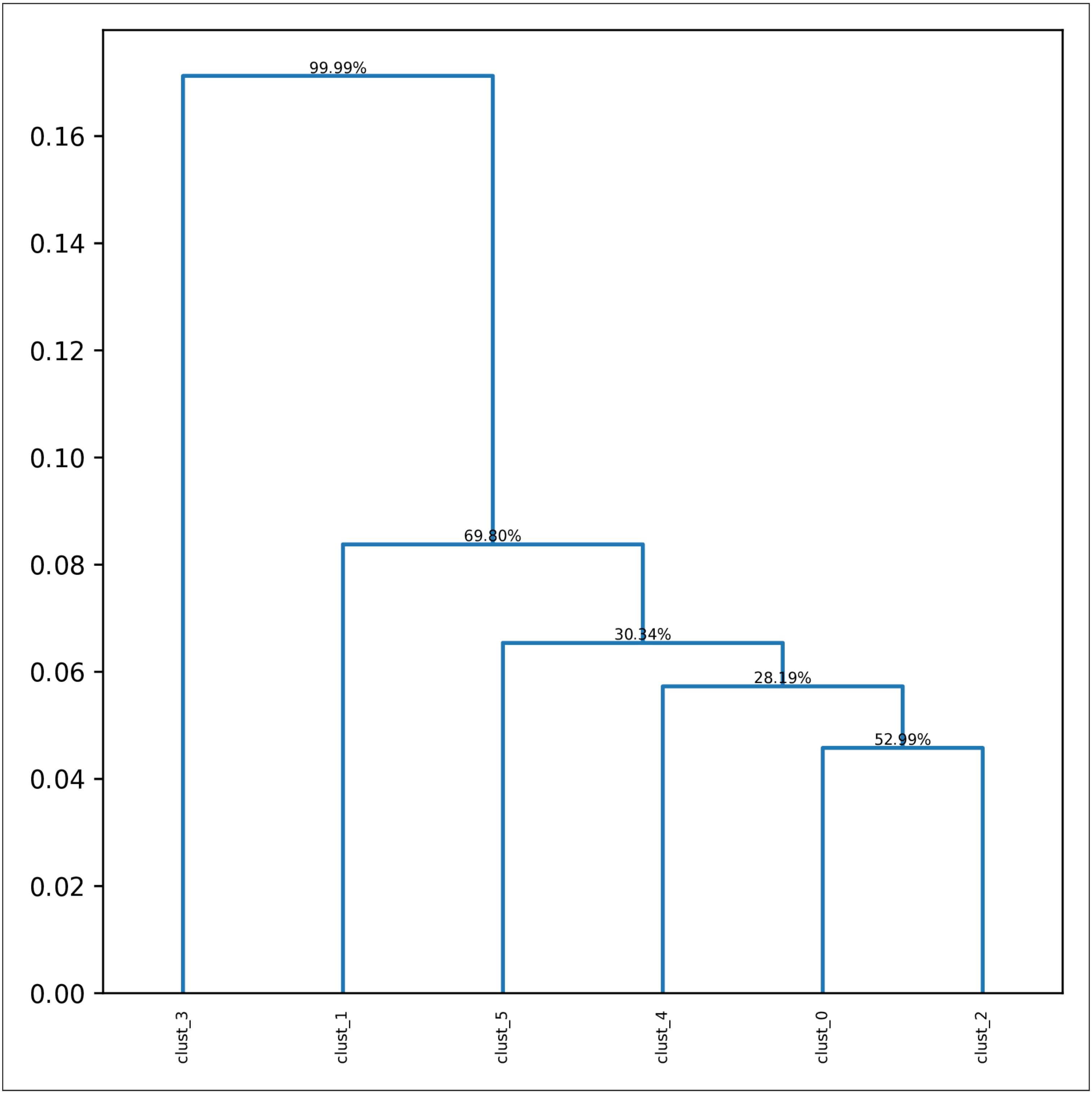

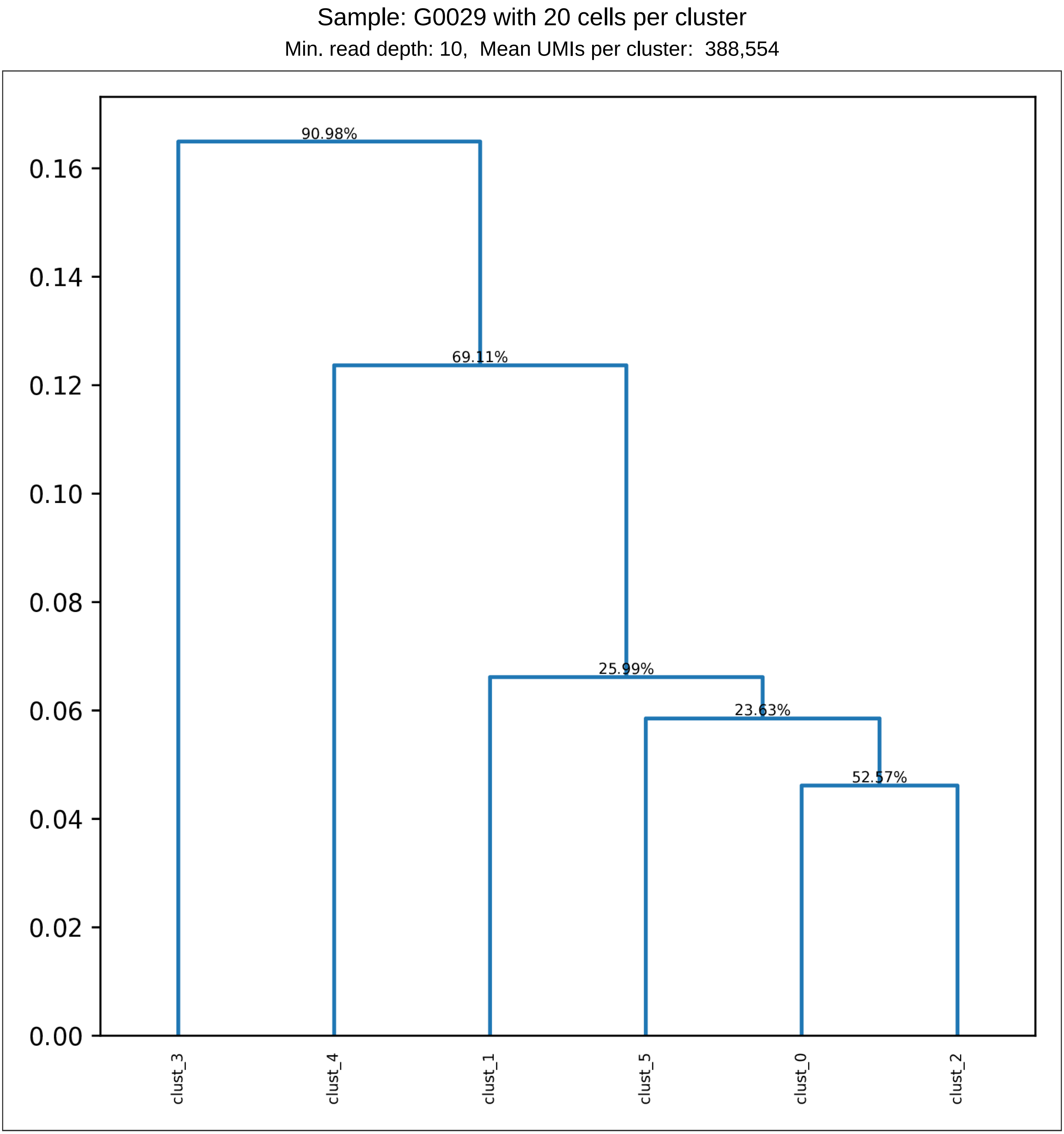
Sample: G0029 with 20 cells per cluster
Min. read depth: 10, Mean UMIs per cluster: 388,554
90.98%
69.11%
25.99%
23.63%
52.57%
0.16
0.14
0.12
0.10
0.08
0.06
0.04
0.02
0.00
clust_3
clust_4
clust_1
clust_5
clust_0
clust_2

Sample: G0029 with 10 cells per cluster

Min. read depth: 10,  Mean UMIs per cluster:  208,033

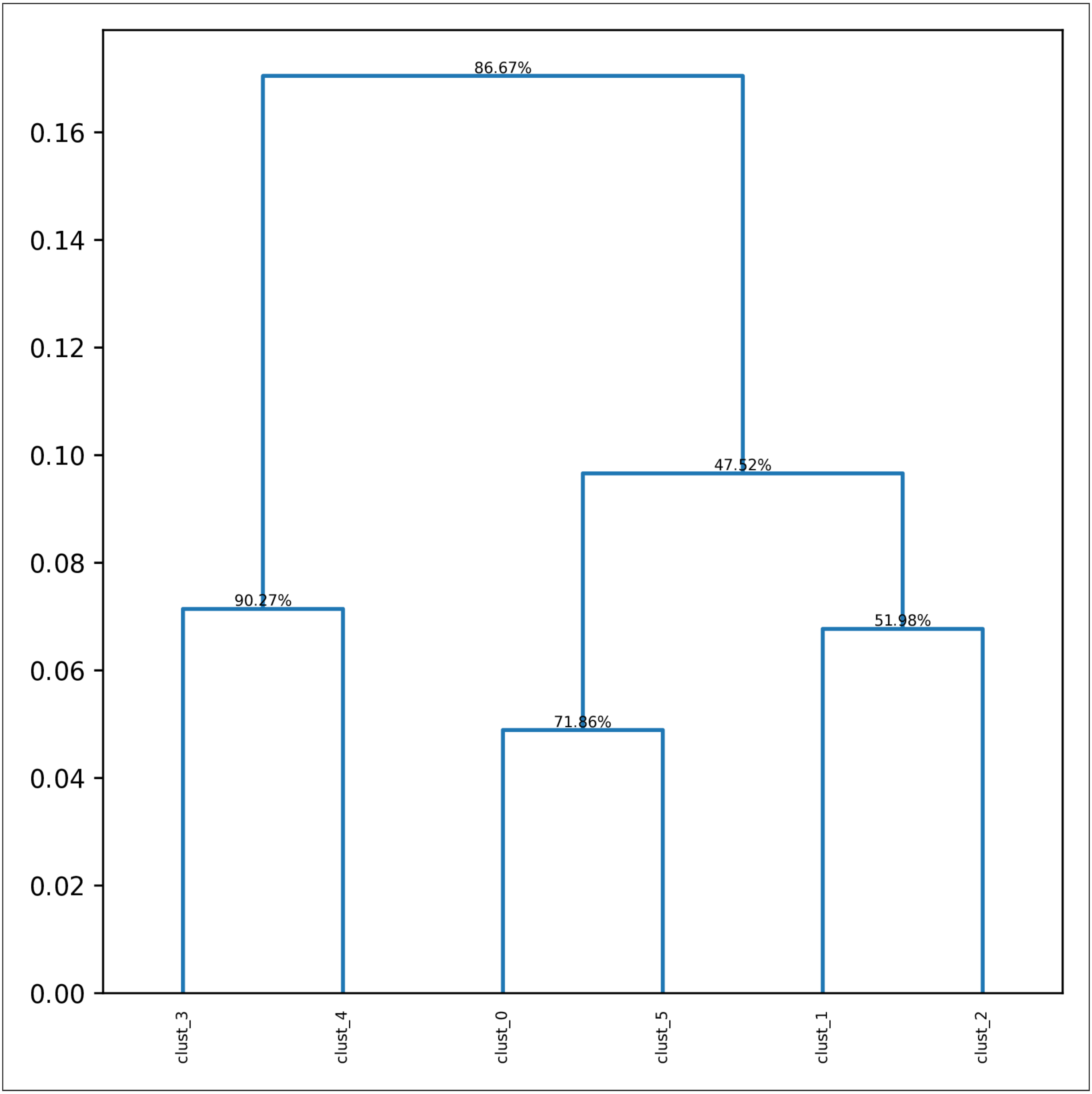

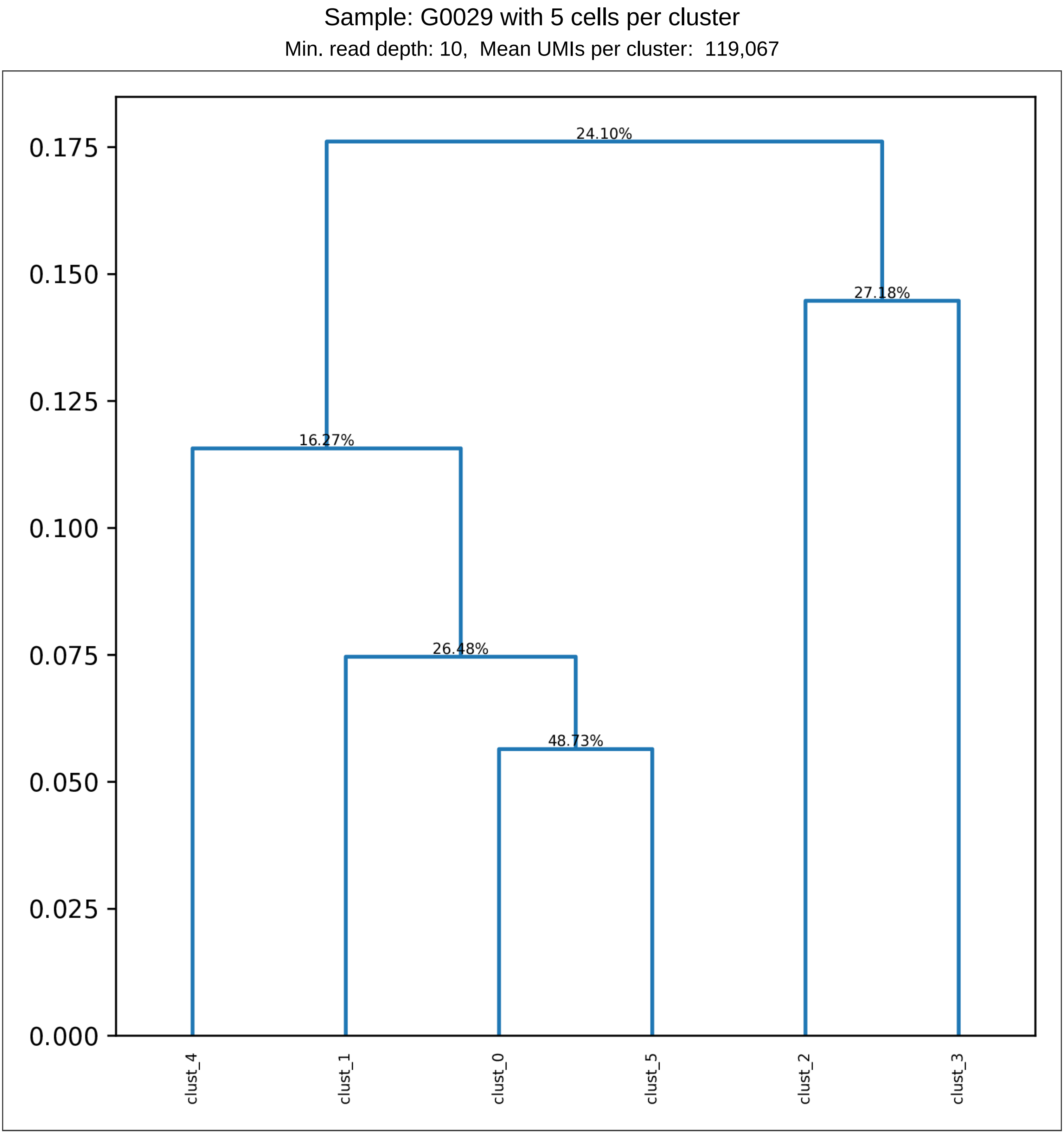
Sample: G0029 with 5 cells per cluster
Min. read depth: 10, Mean UMIs per cluster: 119,067
24.10%
27.18%
16.27%
26.48%
48.73%
0.175
0.150
0.125
0.100
0.075
0.050
0.025
0.000
clust_4
clust_1
clust_0
clust_5
clust_2
clust_3

Sample: G0029 with 4 cells per cluster

Min. read depth: 10,  Mean UMIs per cluster:  73,831

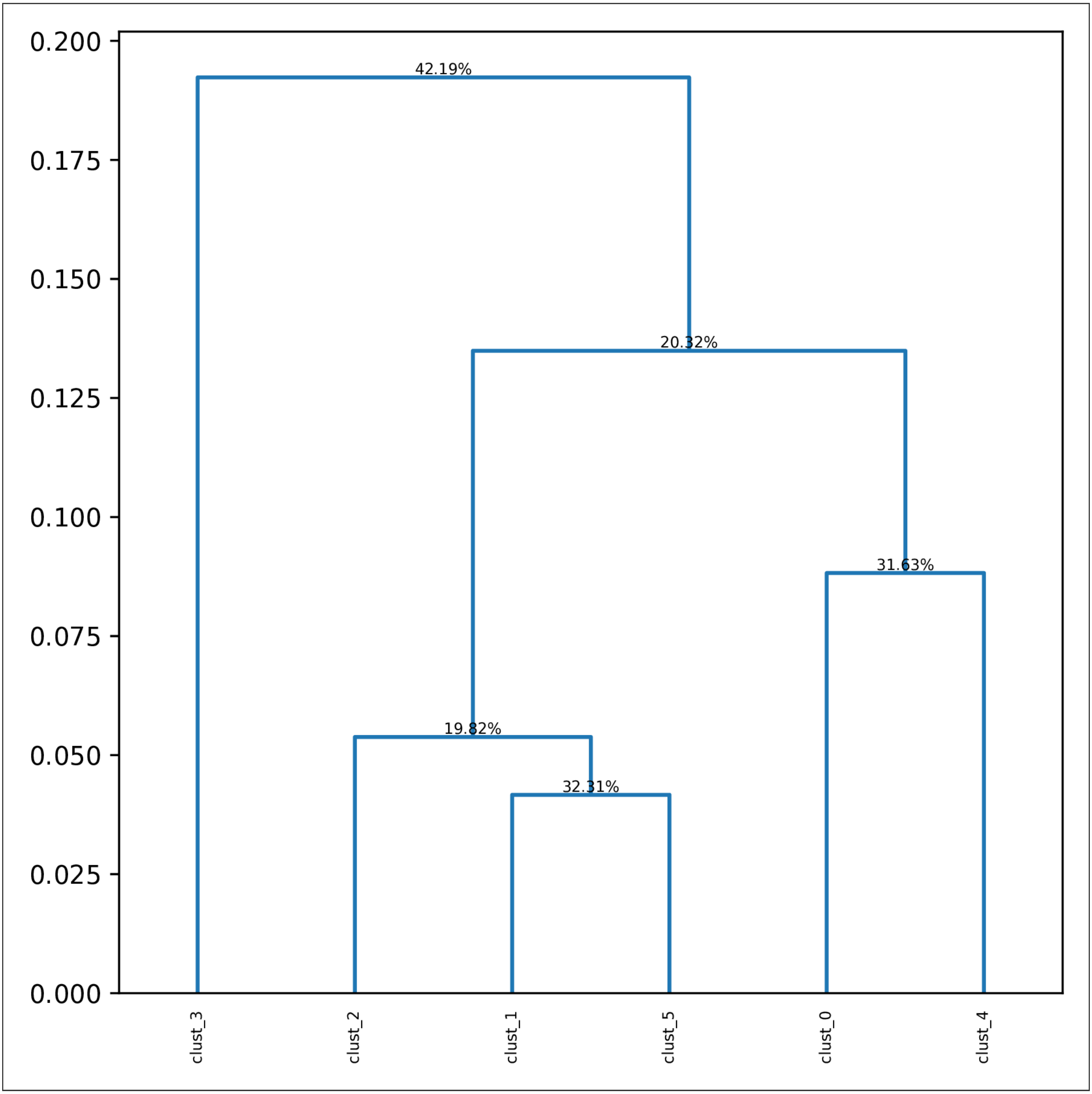

Sample: G0029 with 3 cells per cluster

Min. read depth: 10,  Mean UMIs per cluster:  66,936

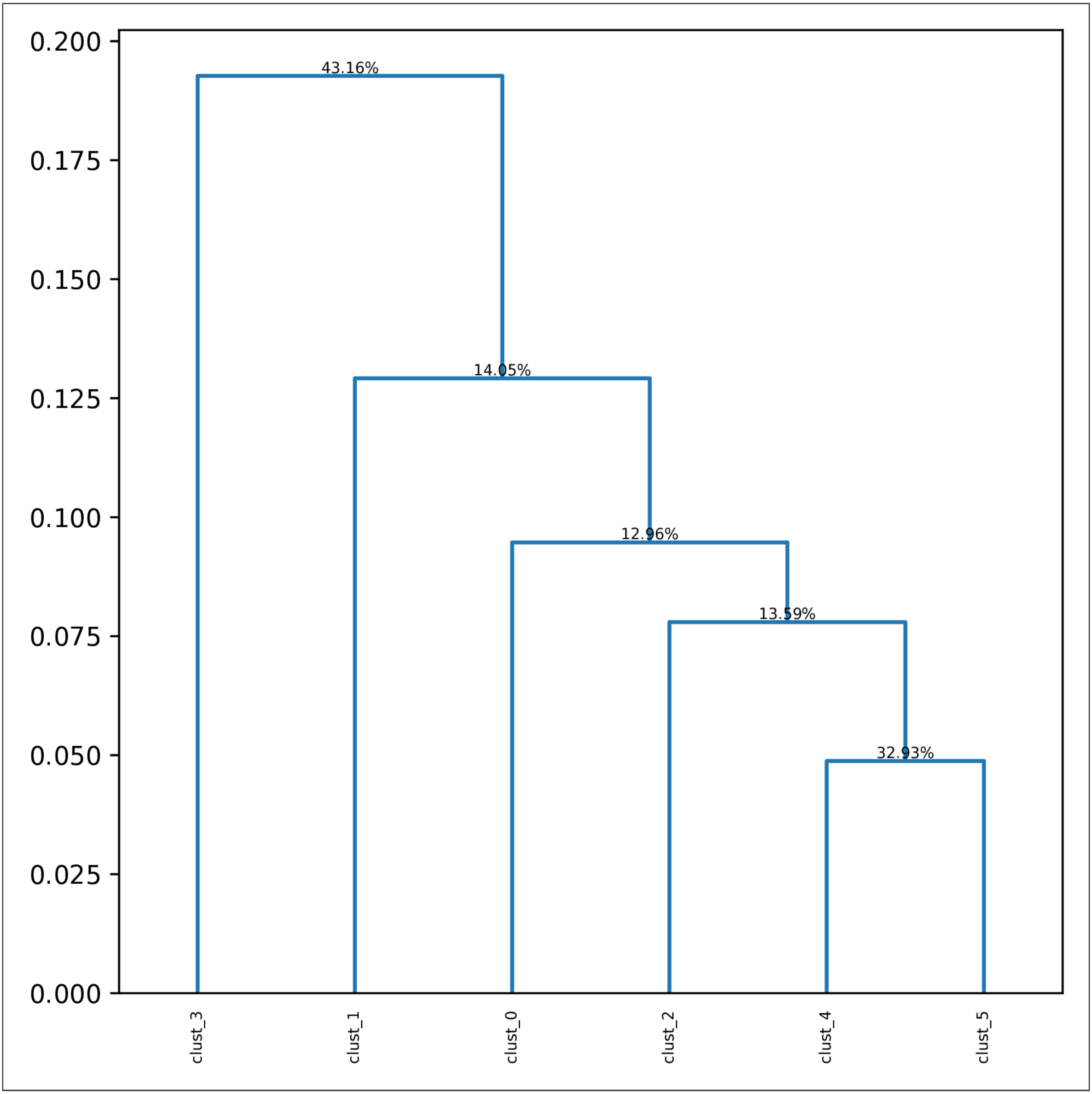

Sample: G0030 with all cells per cluster

Min. read depth: 10, Mean UMIs per cluster: 19,486,335

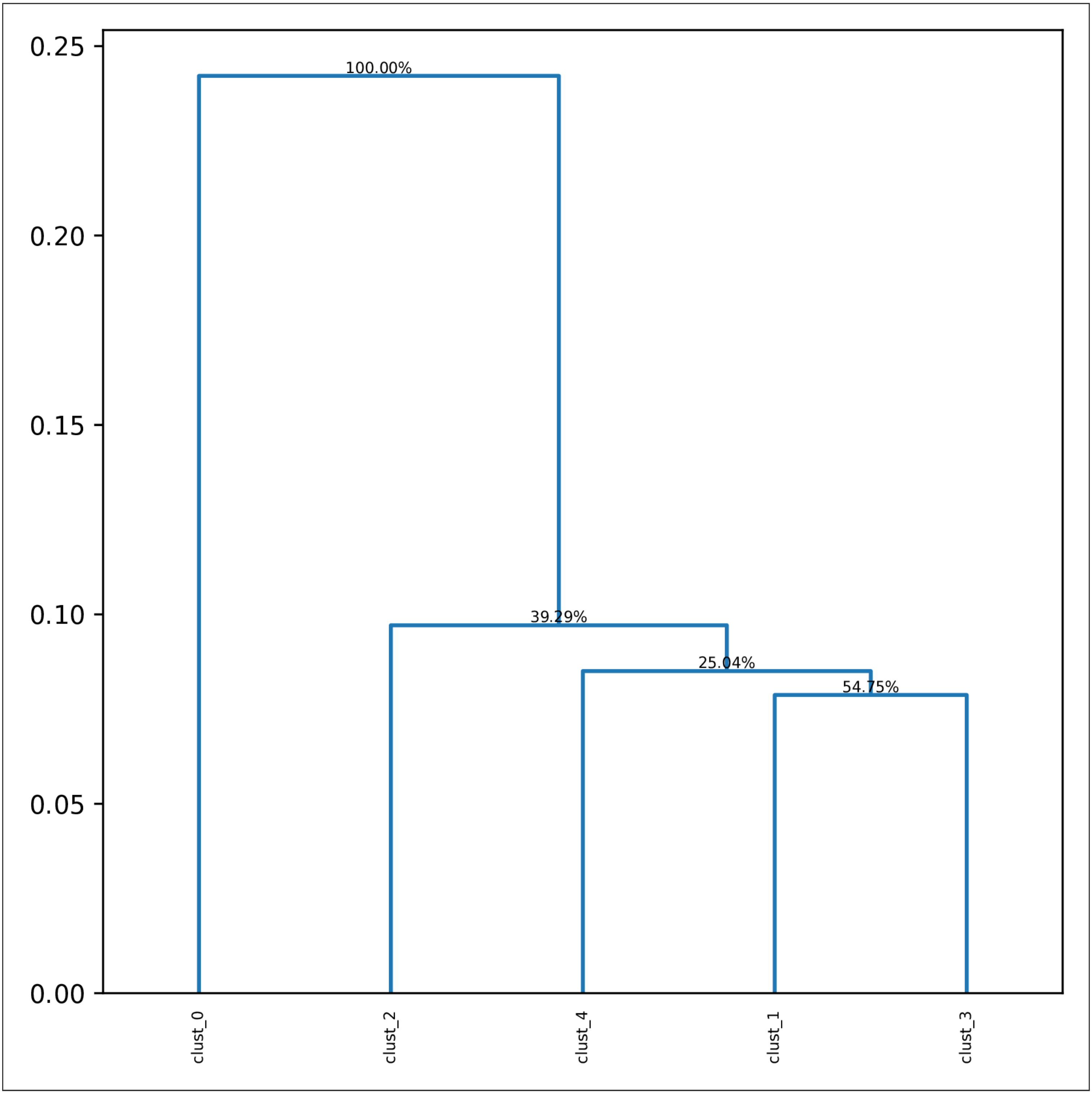

Sample: G0030 with 500 cells per cluster
Min. read depth: 10,  Mean UMIs per cluster:  7,760,997

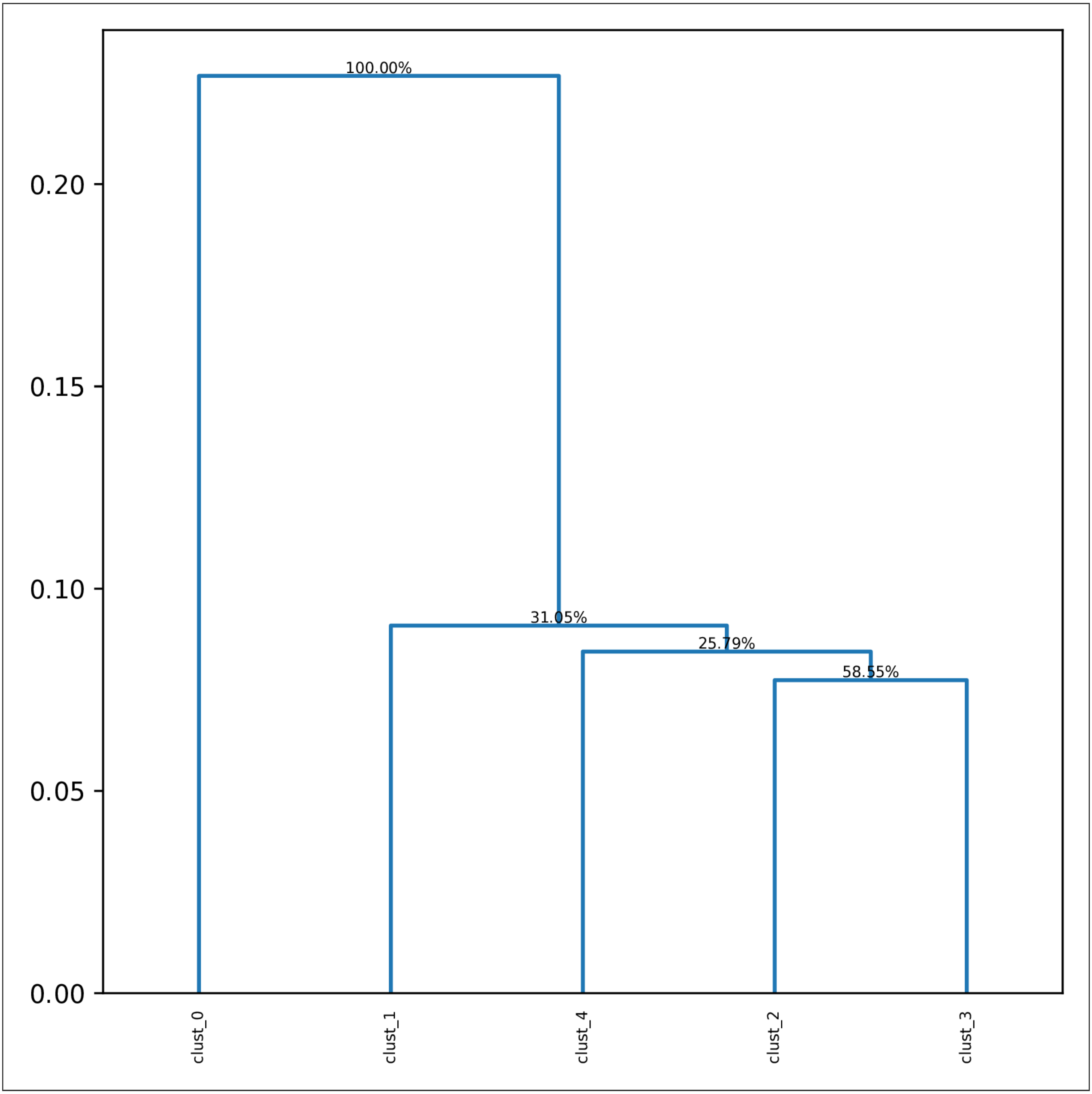

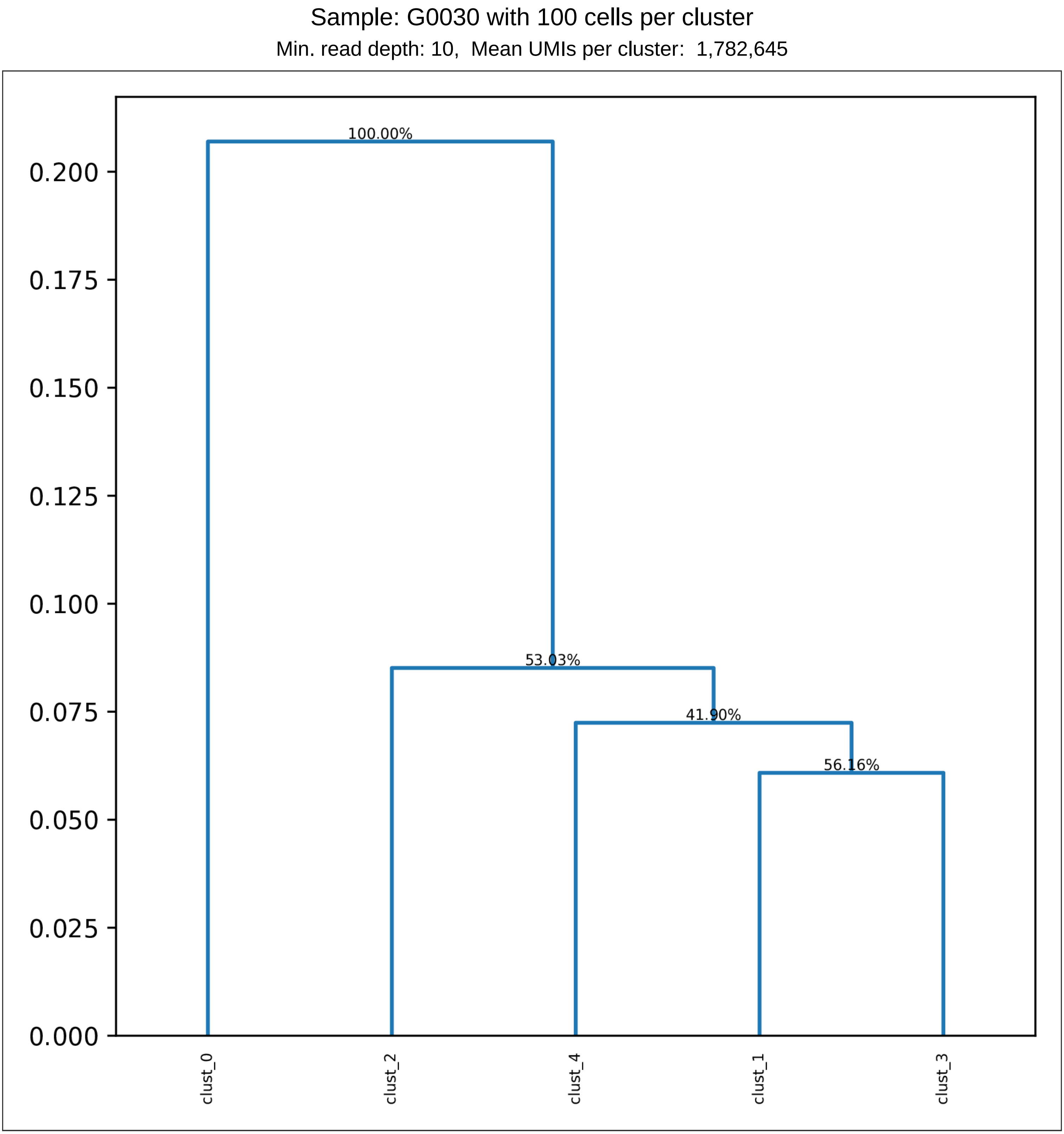

Sample: G0030 with 100 cells per cluster
Min. read depth: 10, Mean UMIs per cluster: 1,782,645
100.00%
53.03%
41.90%
56.16%
0.200
0.175
0.150
0.125
0.100
0.075
0.050
0.025
0.000
clust_0
clust_2
clust_4
clust_1
clust_3

Sample: G0030 with 50 cells per cluster
Min. read depth: 10,  Mean UMIs per cluster:  851,071

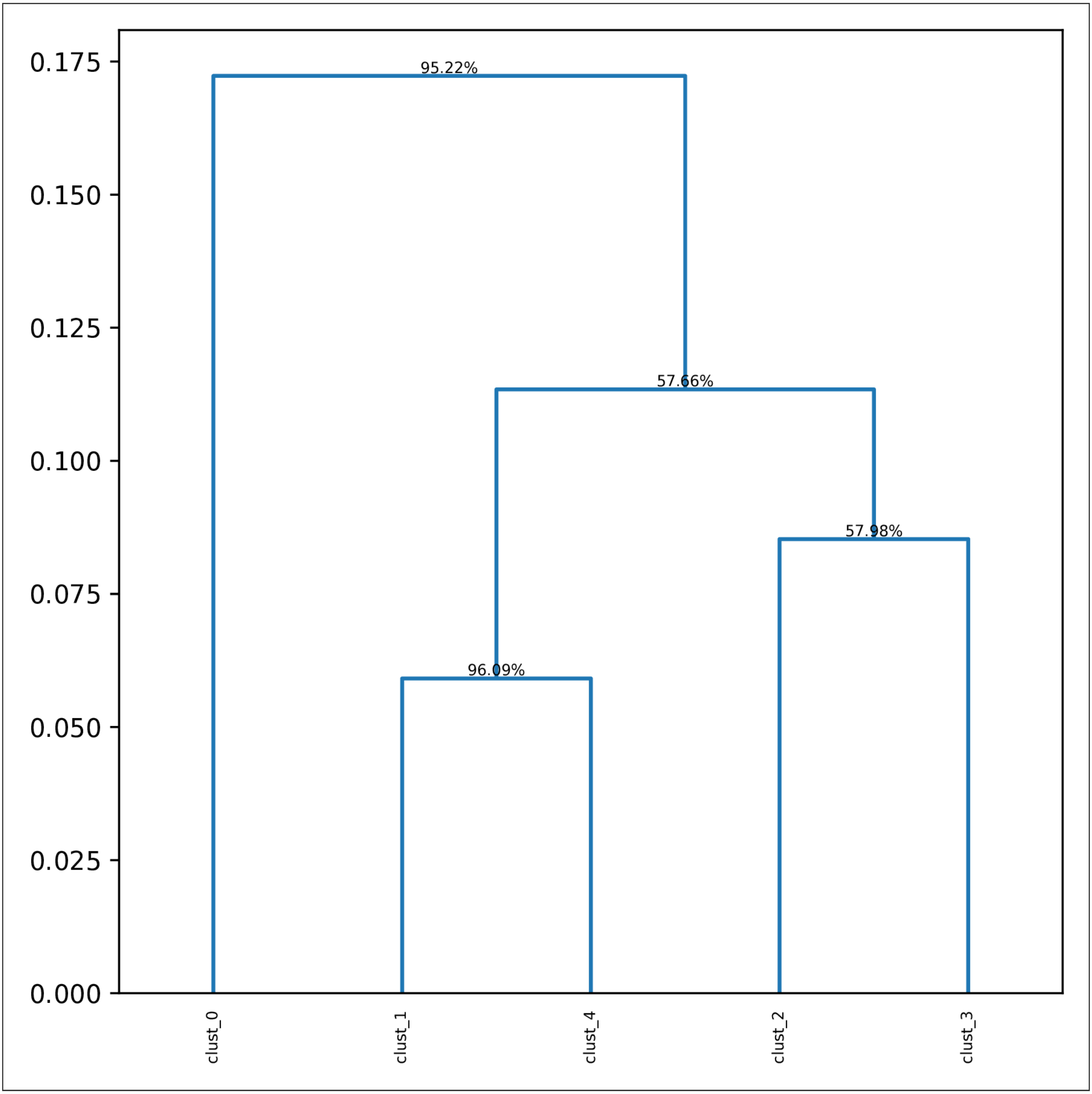

Sample: G0030 with 20 cells per cluster

Min. read depth: 10,  Mean UMIs per cluster:  331,397

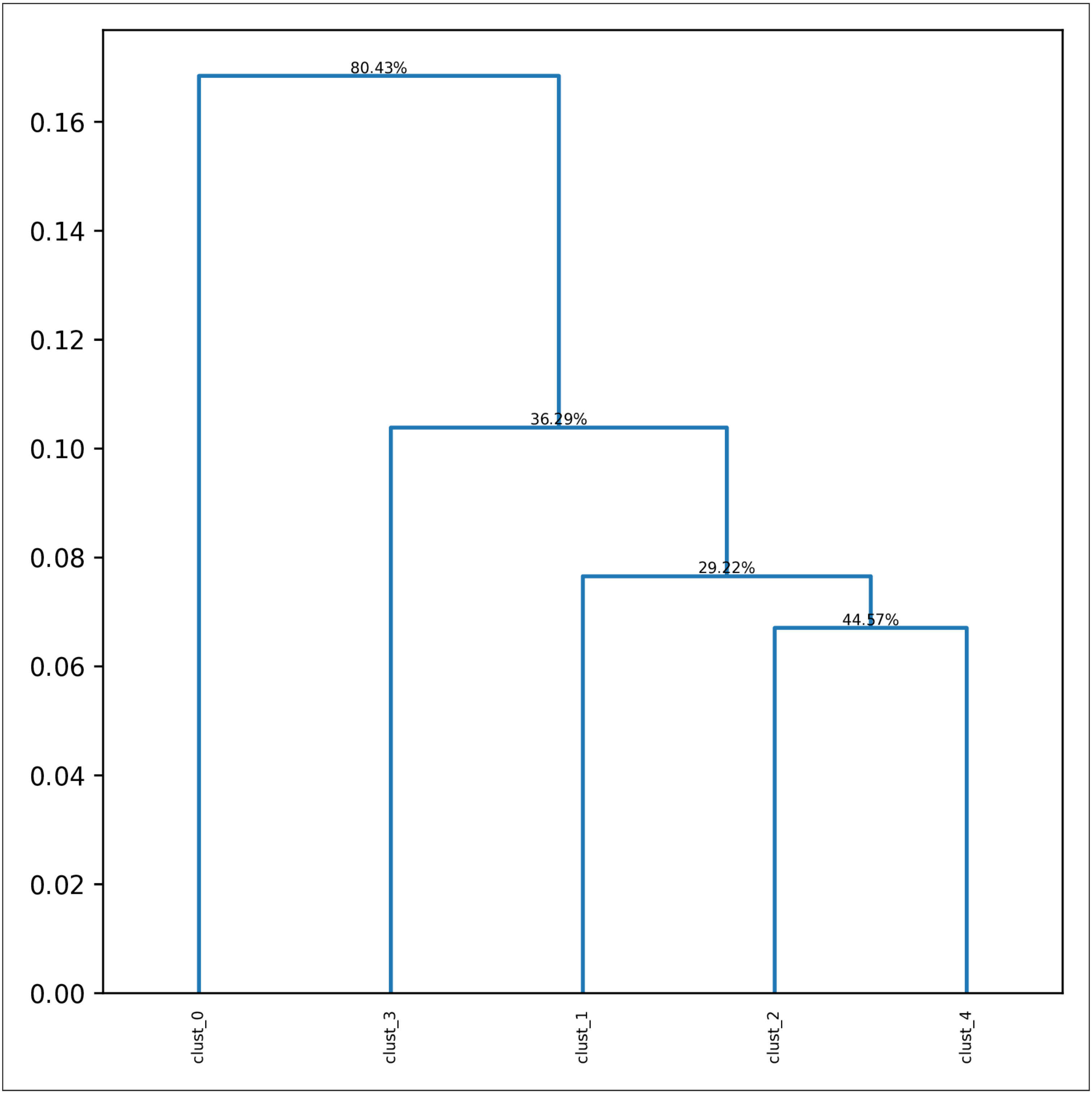

Sample: G0030 with 10 cells per cluster

Min. read depth: 10,  Mean UMIs per cluster:  136,856

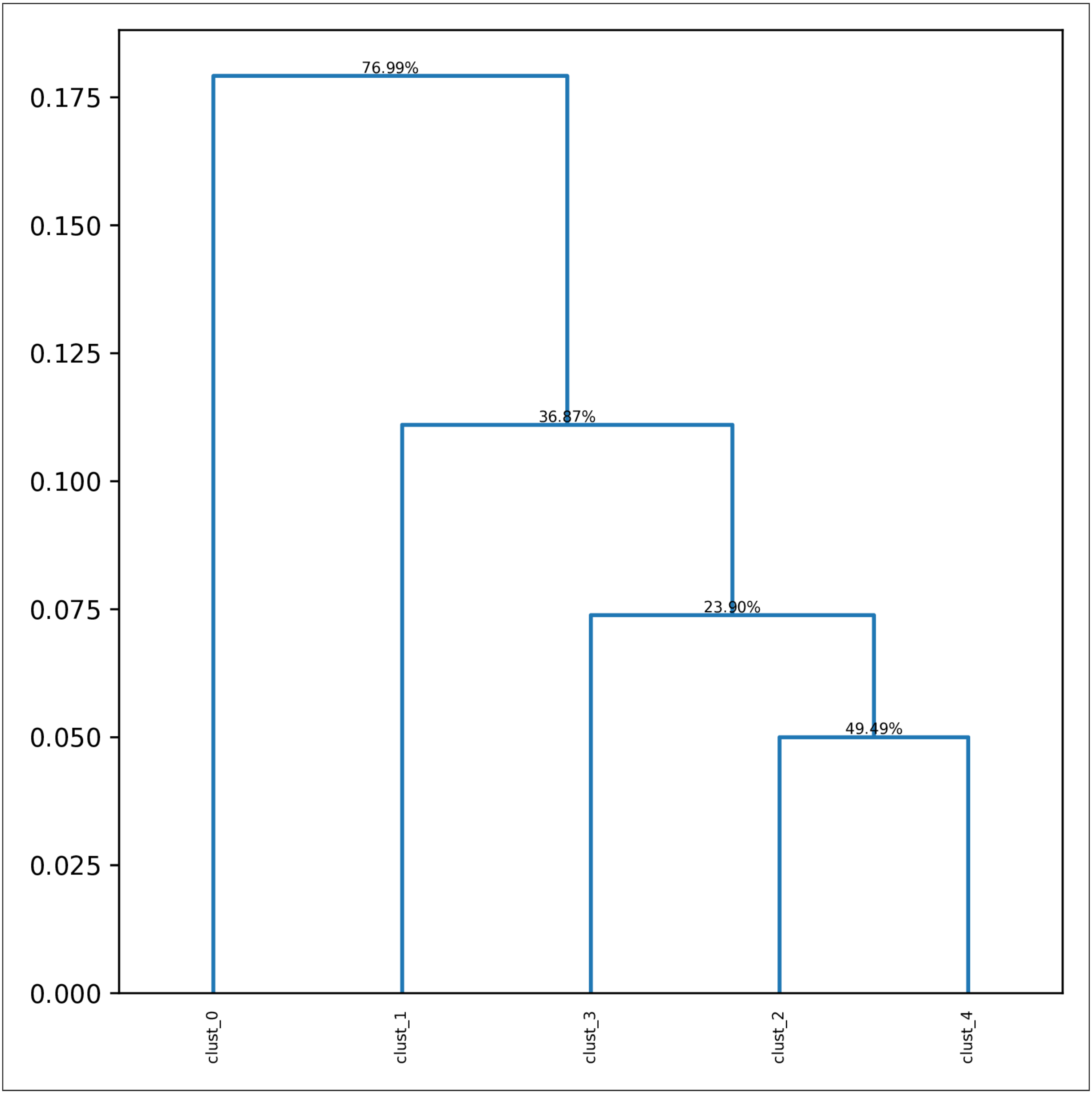

Sample: G0030 with 5 cells per cluster

Min. read depth: 10,  Mean UMIs per cluster:  68,714

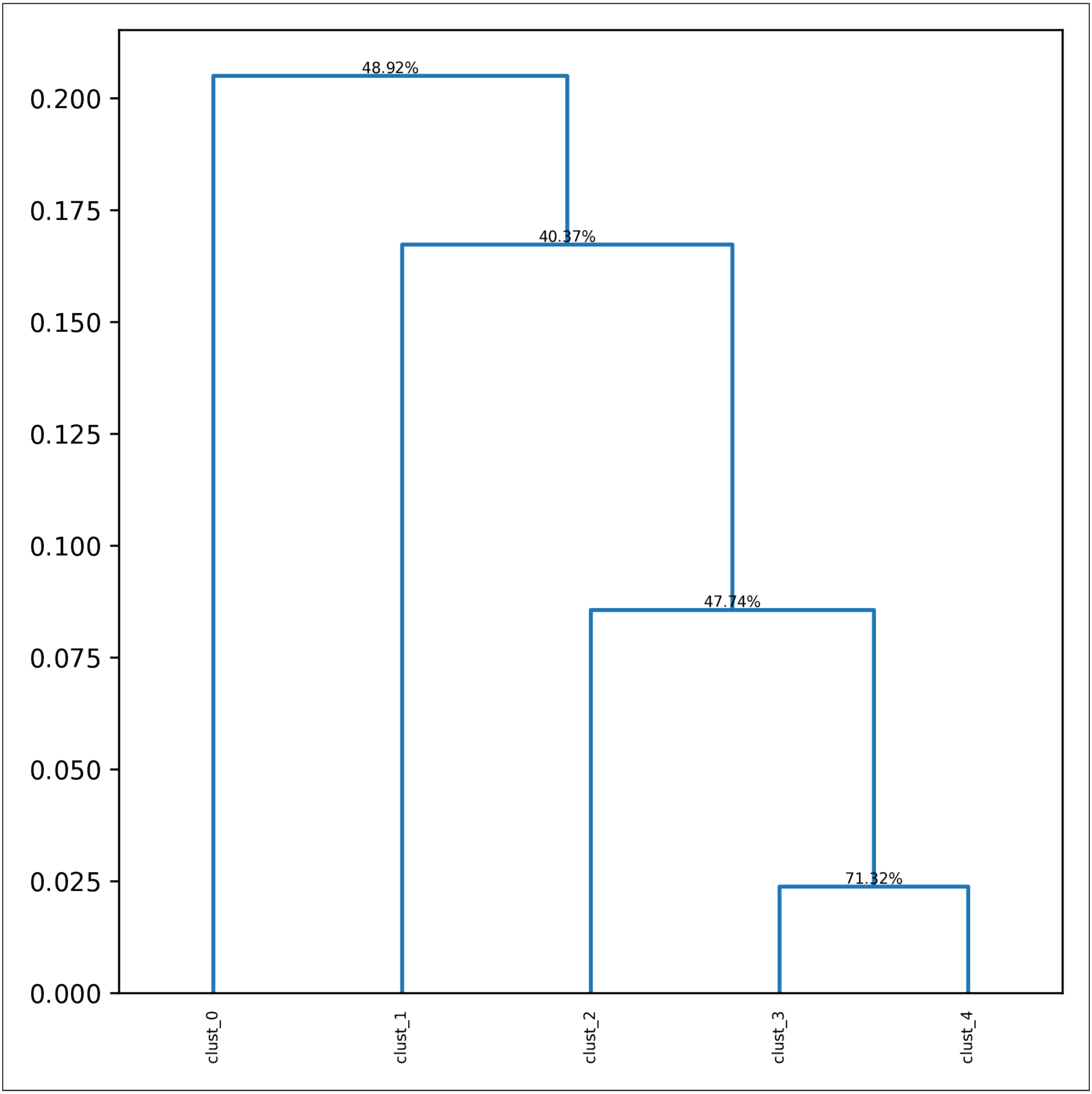

Sample: G0030 with 4 cells per cluster

Min. read depth: 10,  Mean UMIs per cluster:  77,772

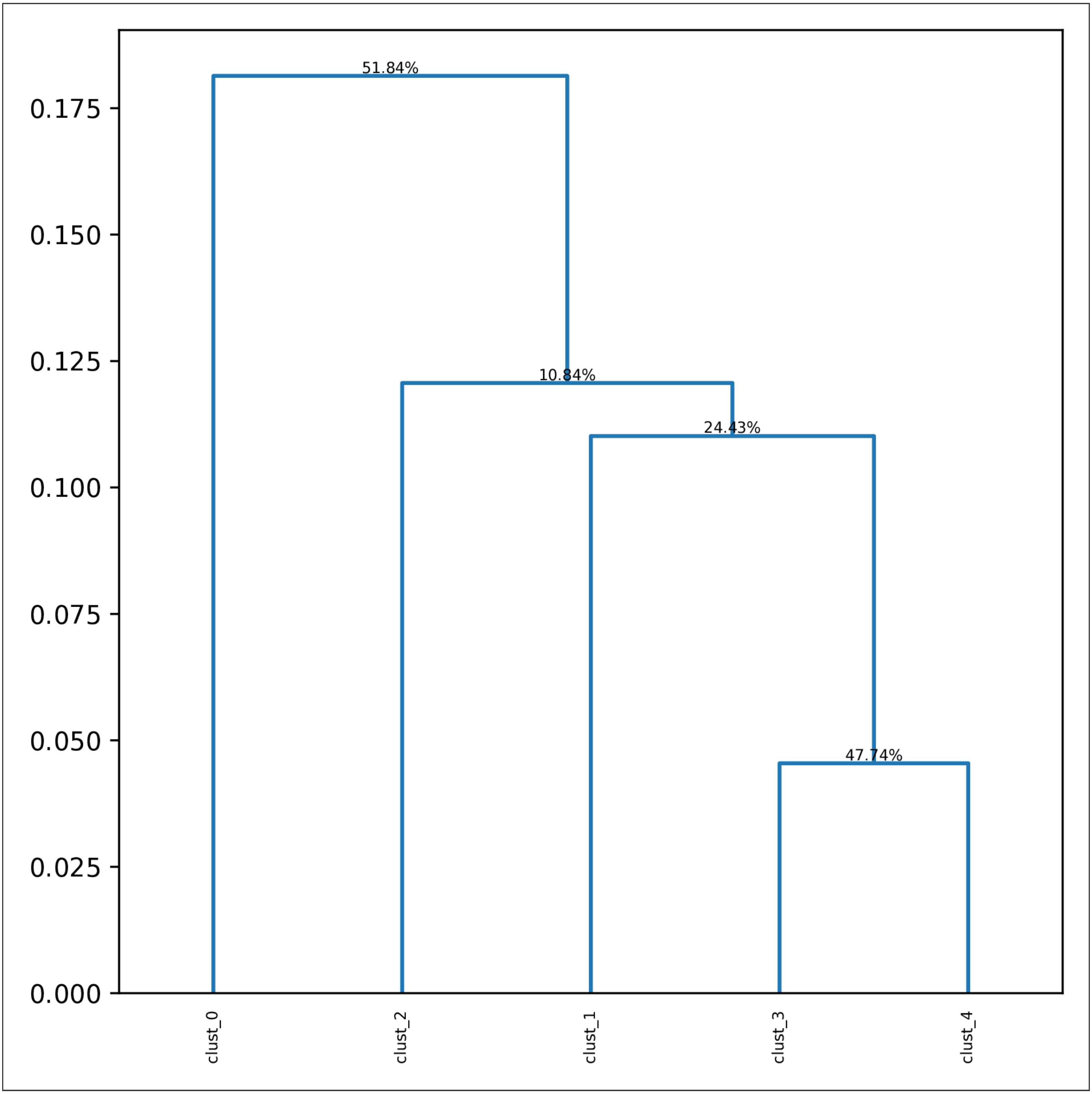

Sample: G0030 with 3 cells per cluster

Min. read depth: 10,  Mean UMIs per cluster:  45,627

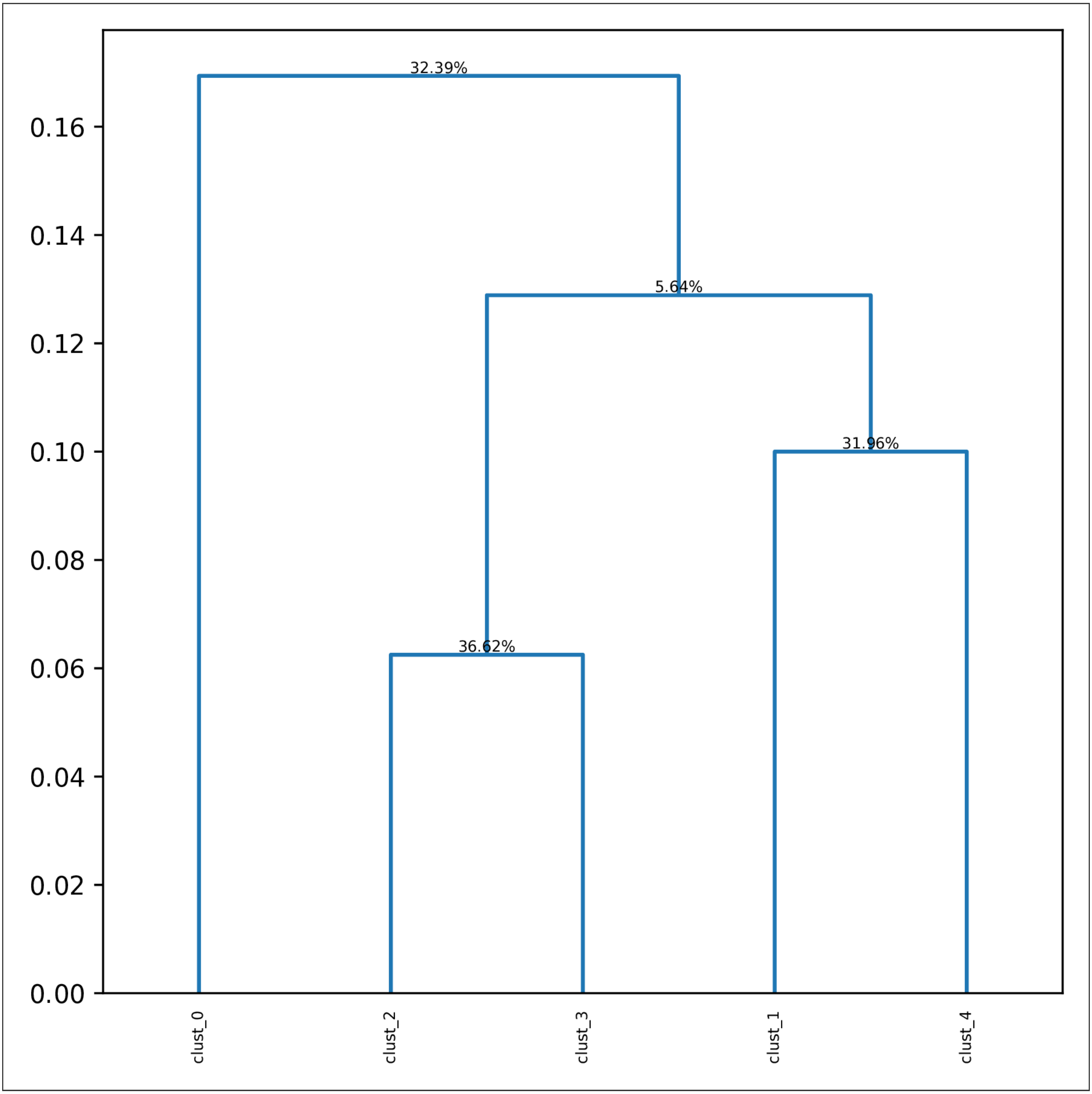

Sample: G0031 with all cells per cluster

Min. read depth: 10,  Mean UMIs per cluster:  15,190,598

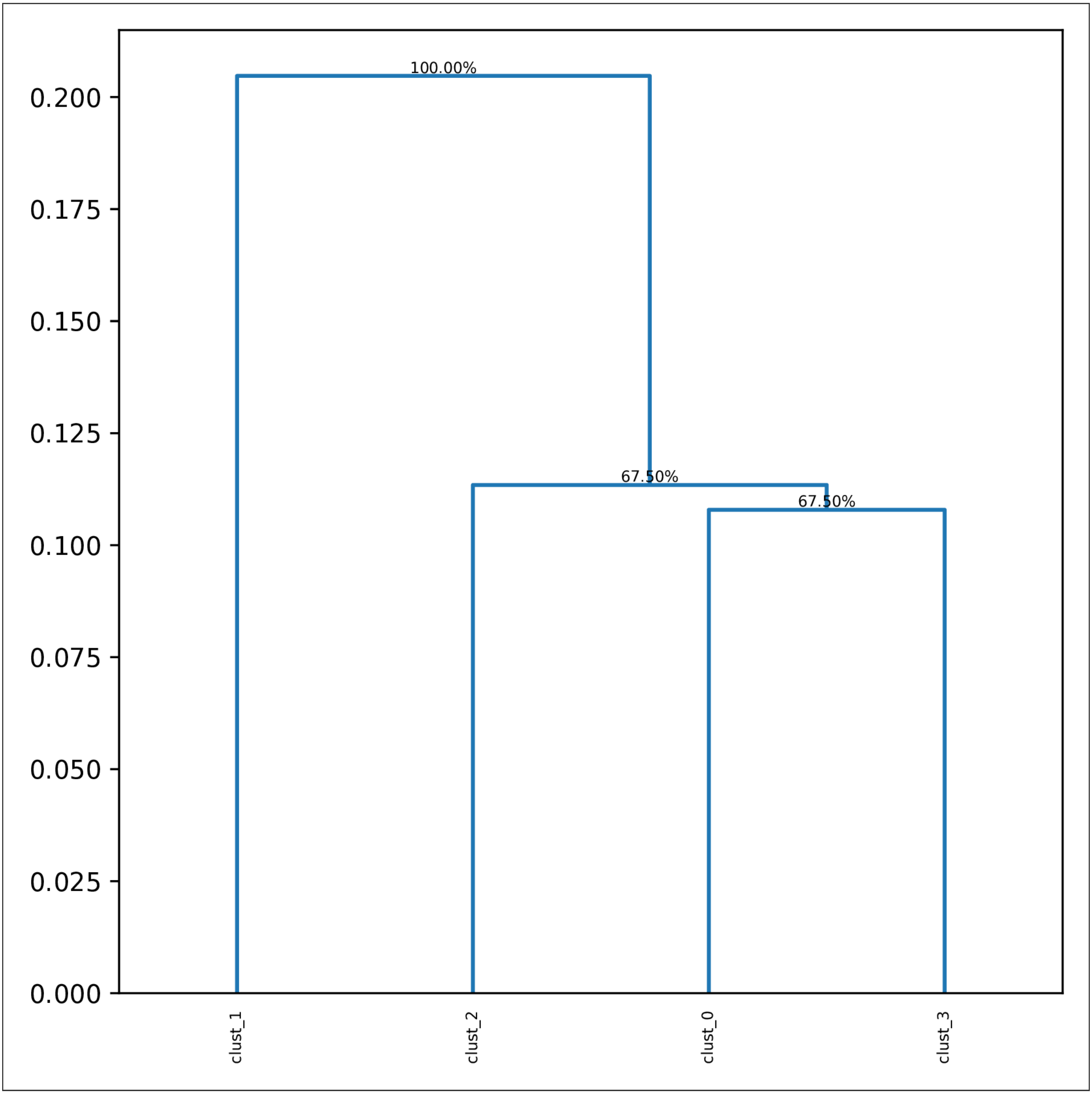

Sample: G0031 with 500 cells per cluster

Min. read depth: 10,  Mean UMIs per cluster:  8,670,030

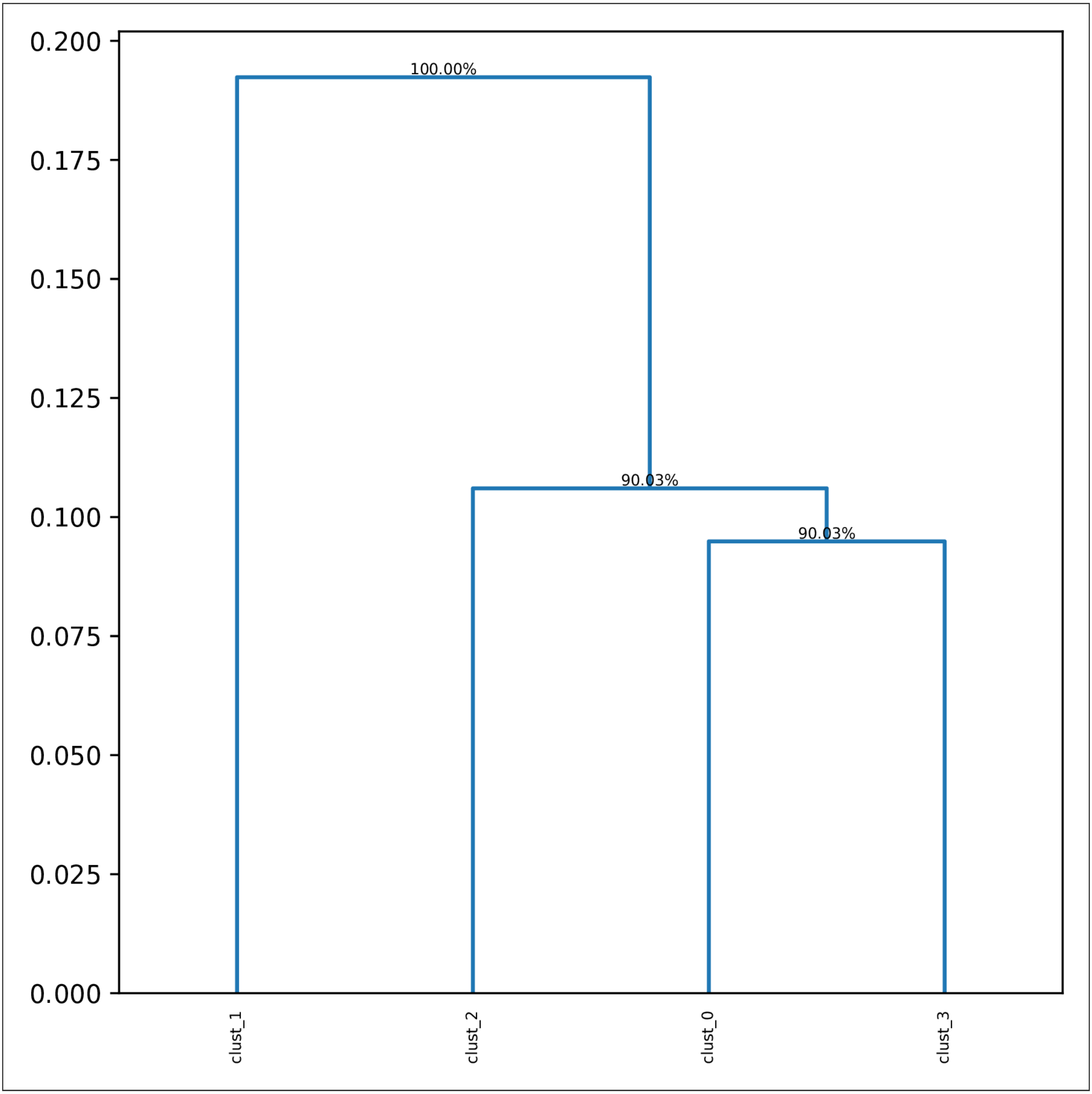

Sample: G0031 with 100 cells per cluster
Min. read depth: 10,  Mean UMIs per cluster:  1,952,992

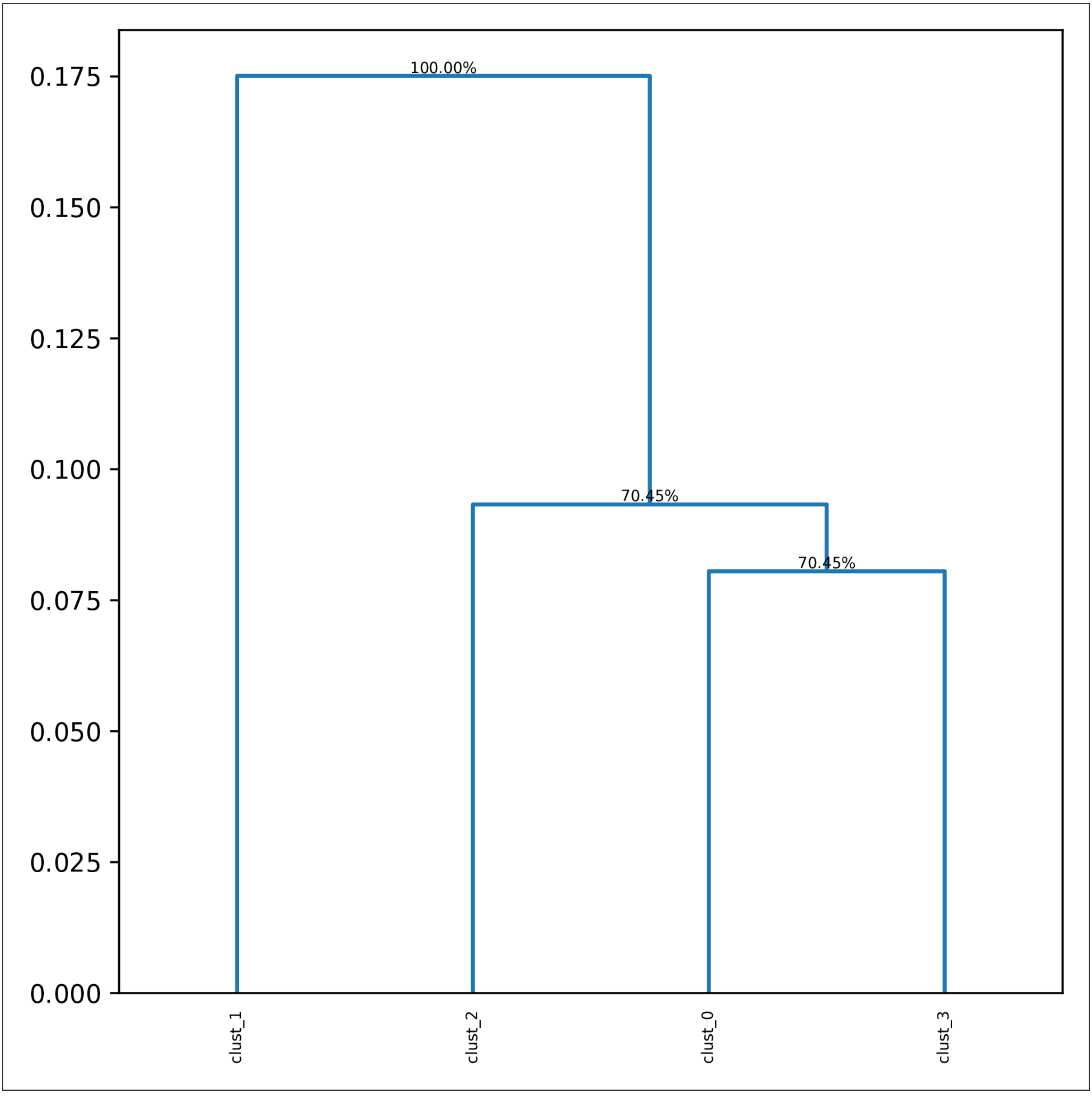

Sample: G0031 with 50 cells per cluster
Min. read depth: 10,  Mean UMIs per cluster:  977,644

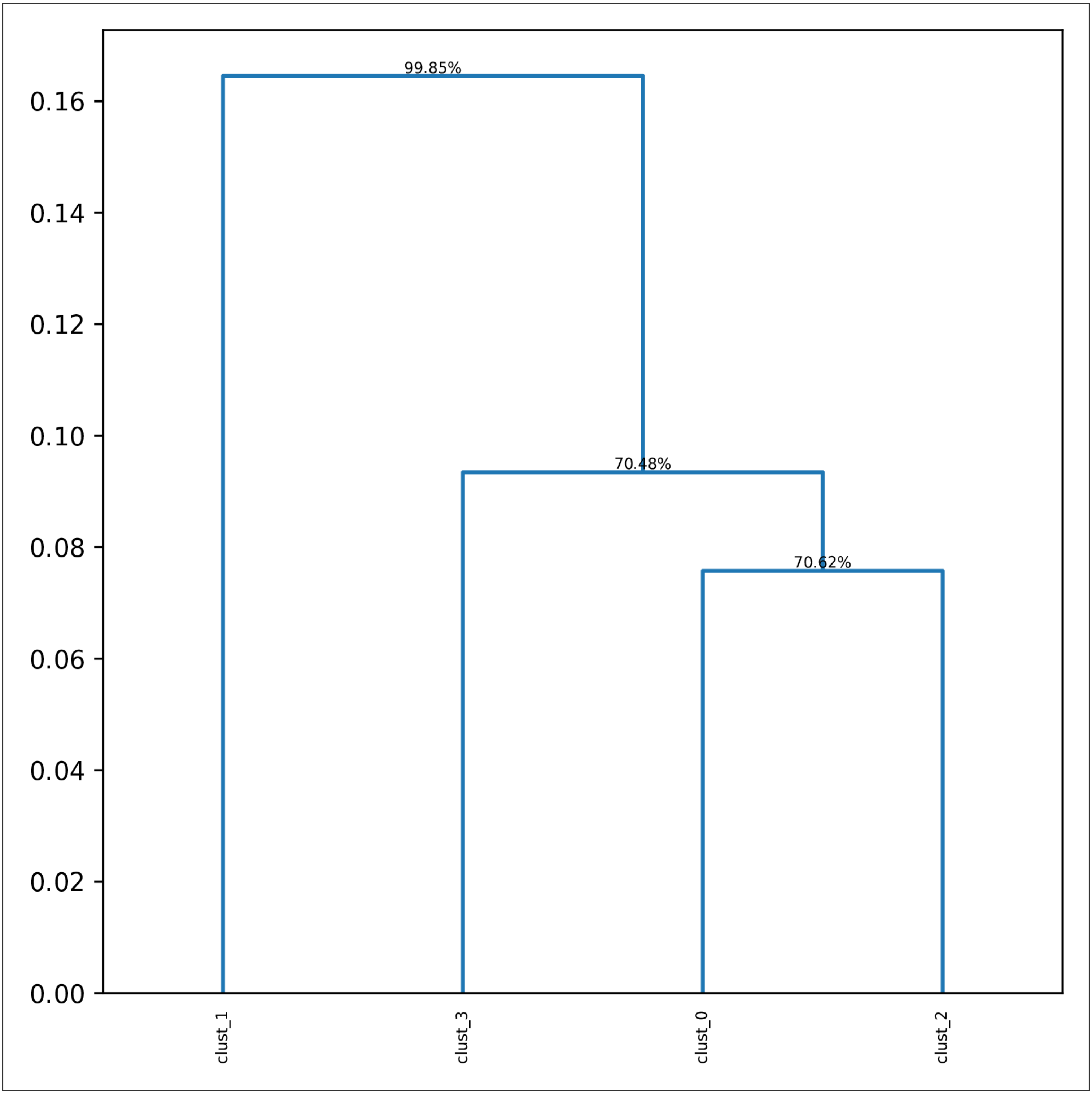

Sample: G0031 with 20 cells per cluster

Min. read depth: 10,  Mean UMIs per cluster:  359,007

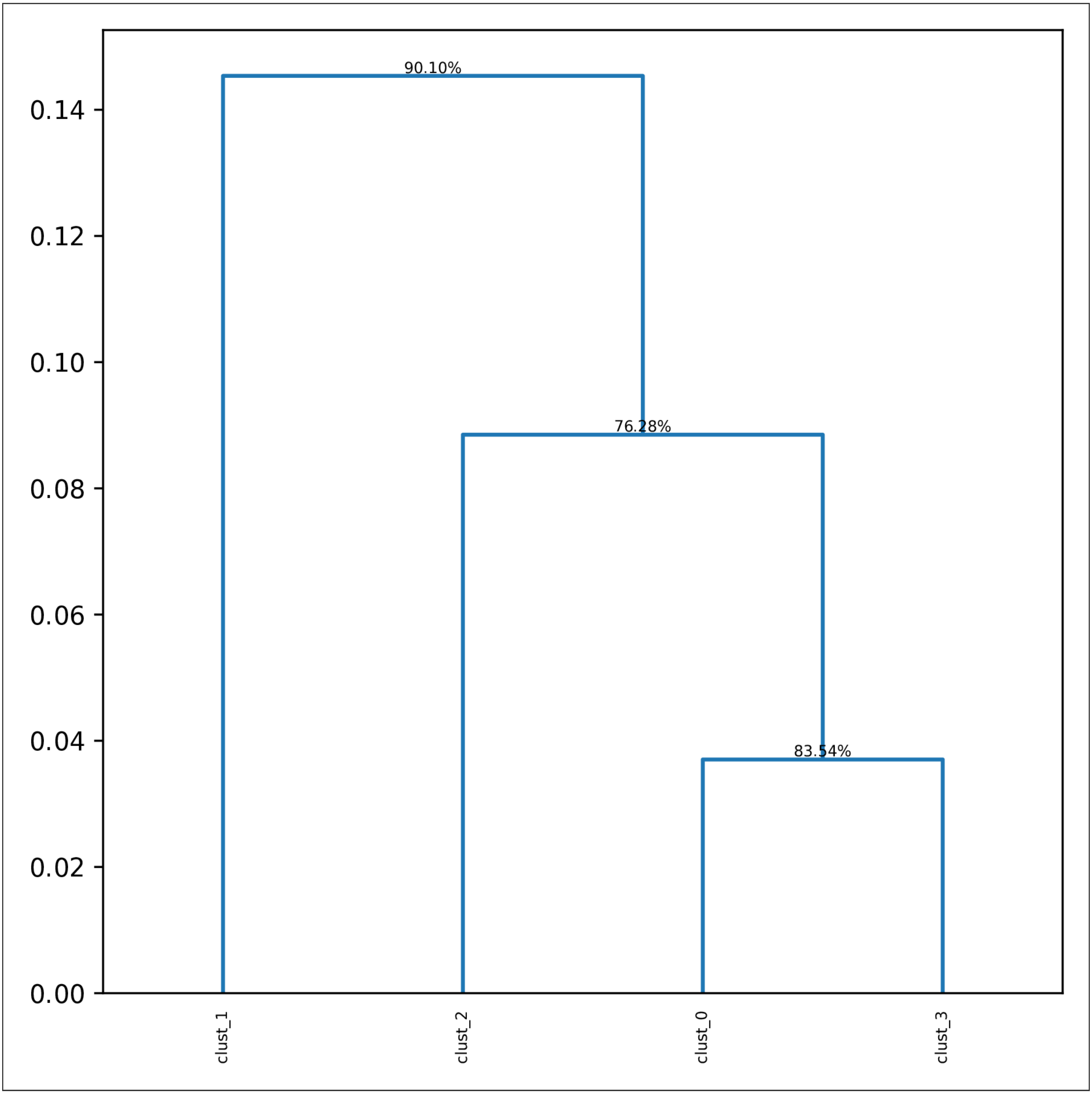

Sample: G0031 with 10 cells per cluster
Min. read depth: 10,  Mean UMIs per cluster:  190,747

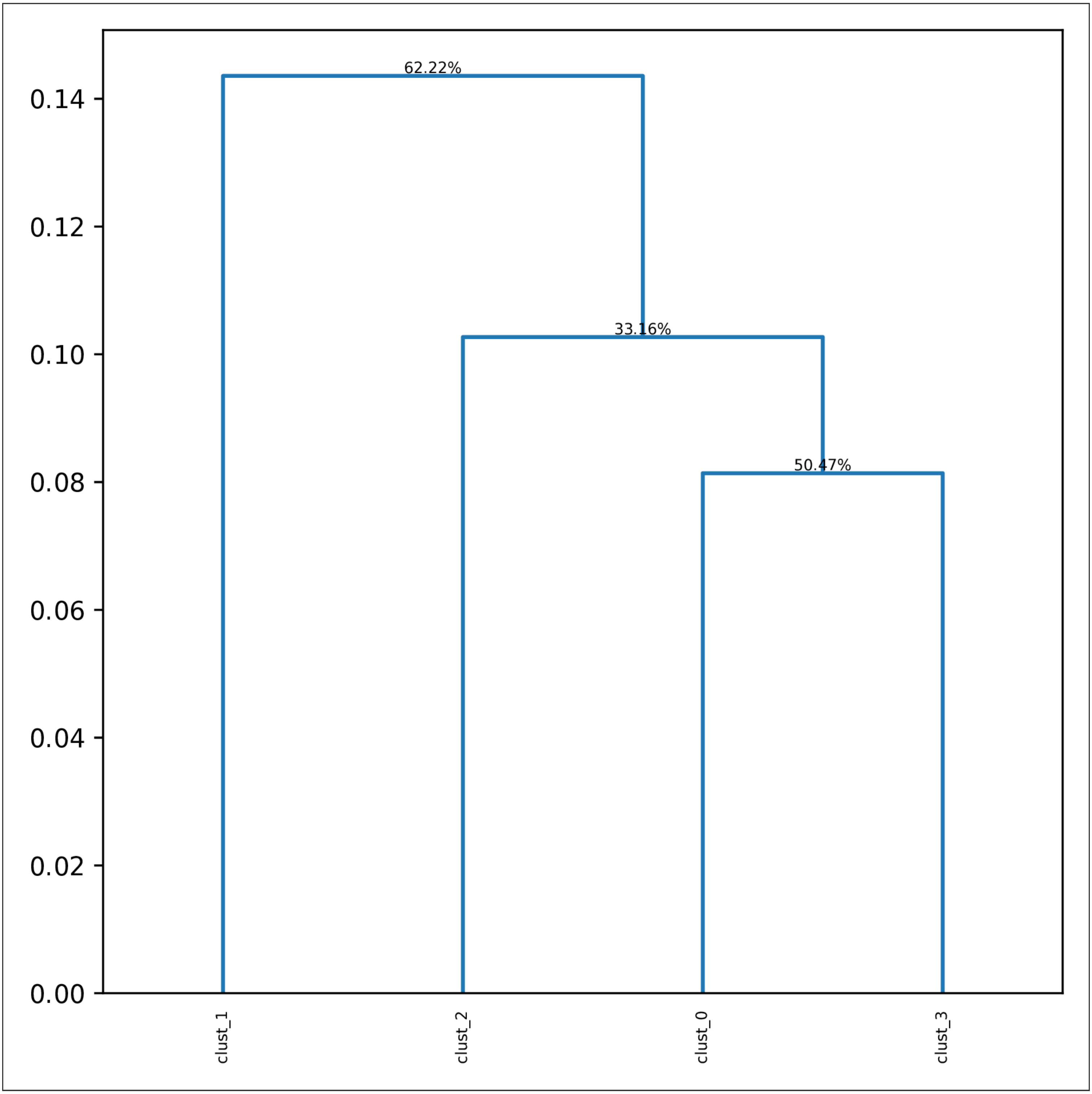

Sample: G0031 with 5 cells per cluster

Min. read depth: 10, Mean UMIs per cluster: 79,498

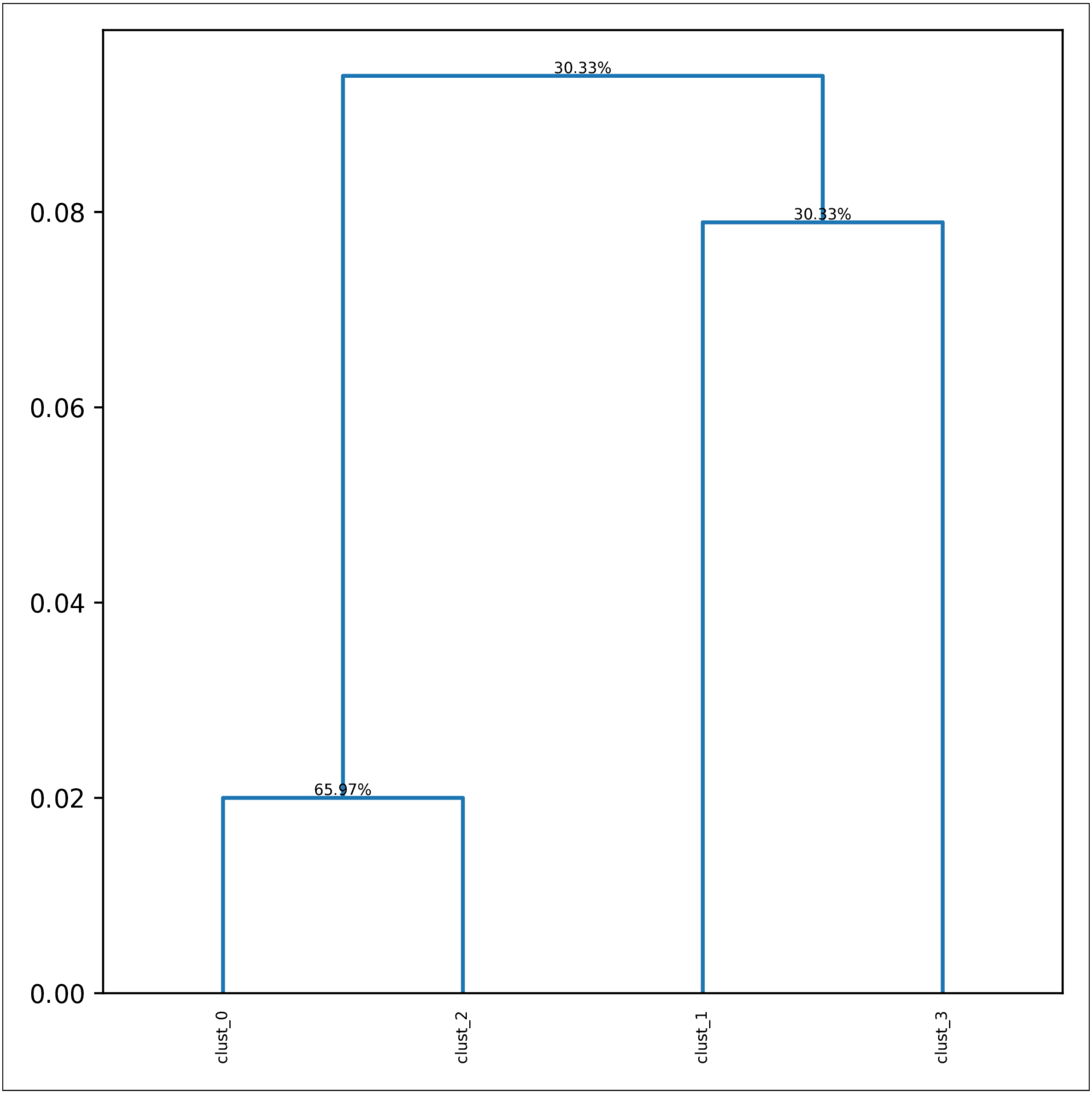

Sample: G0031 with 4 cells per cluster
Min. read depth: 10, Mean UMIs per cluster: 49,218

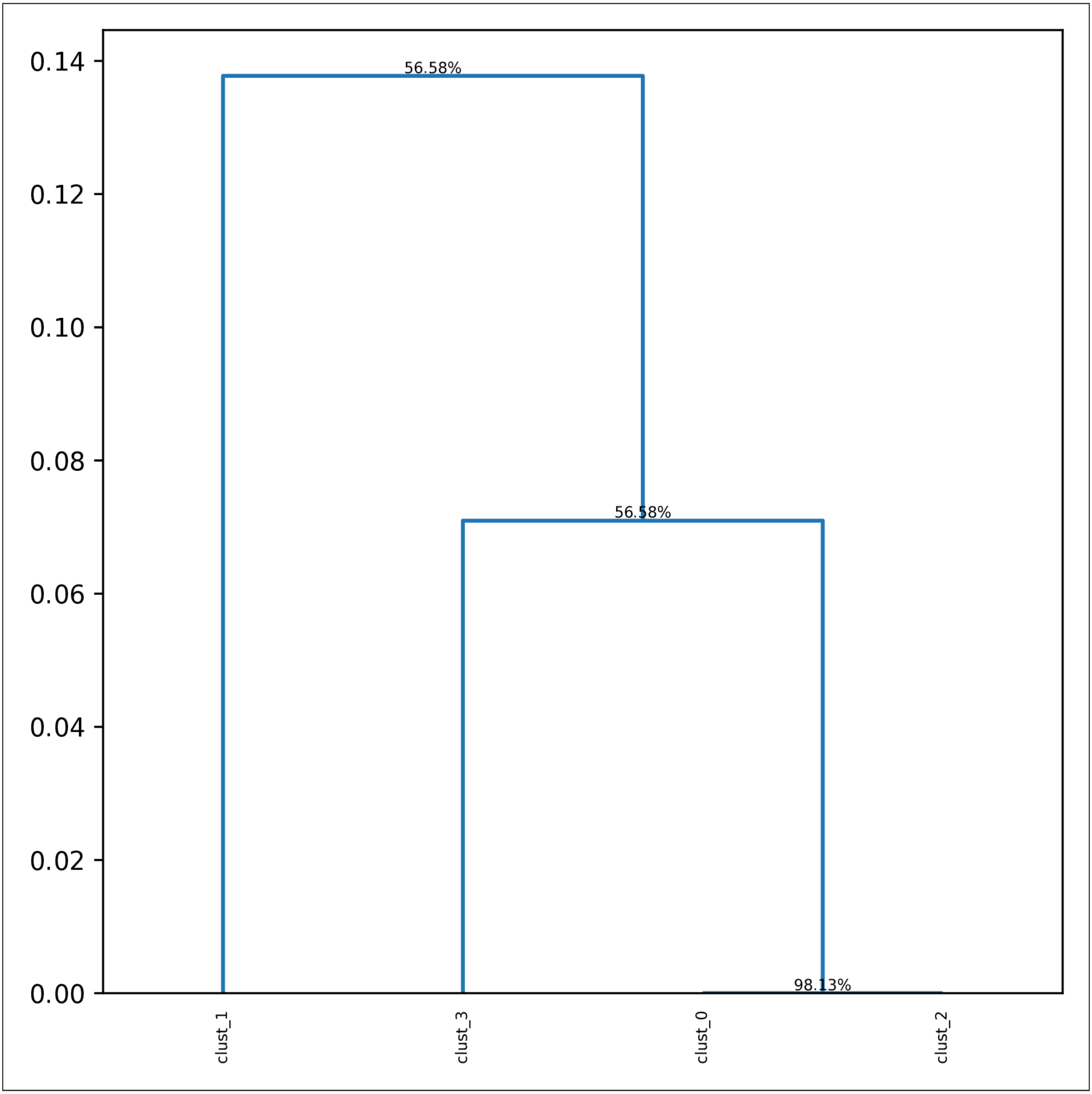

Sample: G0031 with 3 cells per cluster

Min. read depth: 10,  Mean UMIs per cluster:  42,556

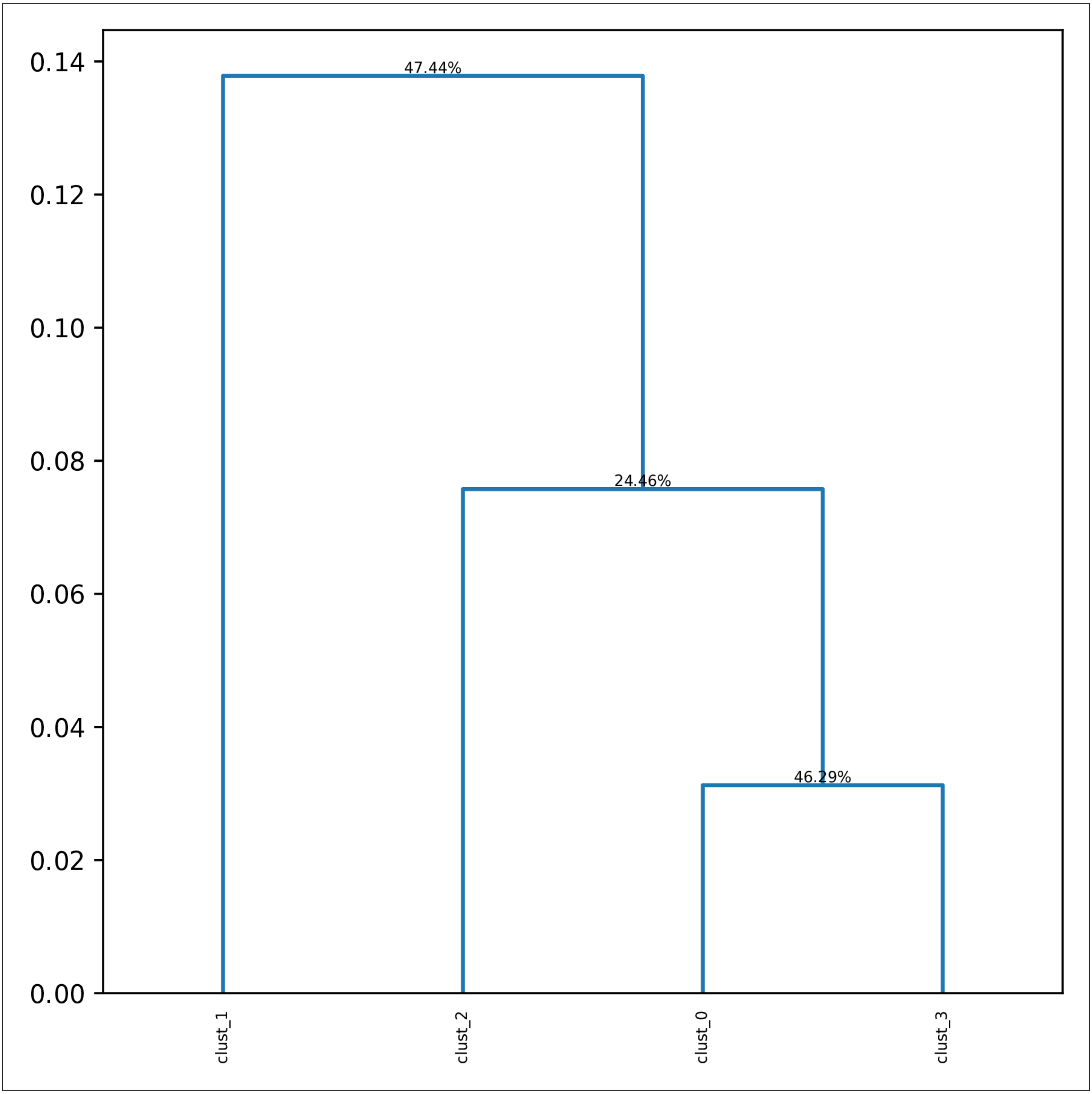

Sample: G0032 with all cells per cluster

Min. read depth: 10,  Mean UMIs per cluster:  5,870,099

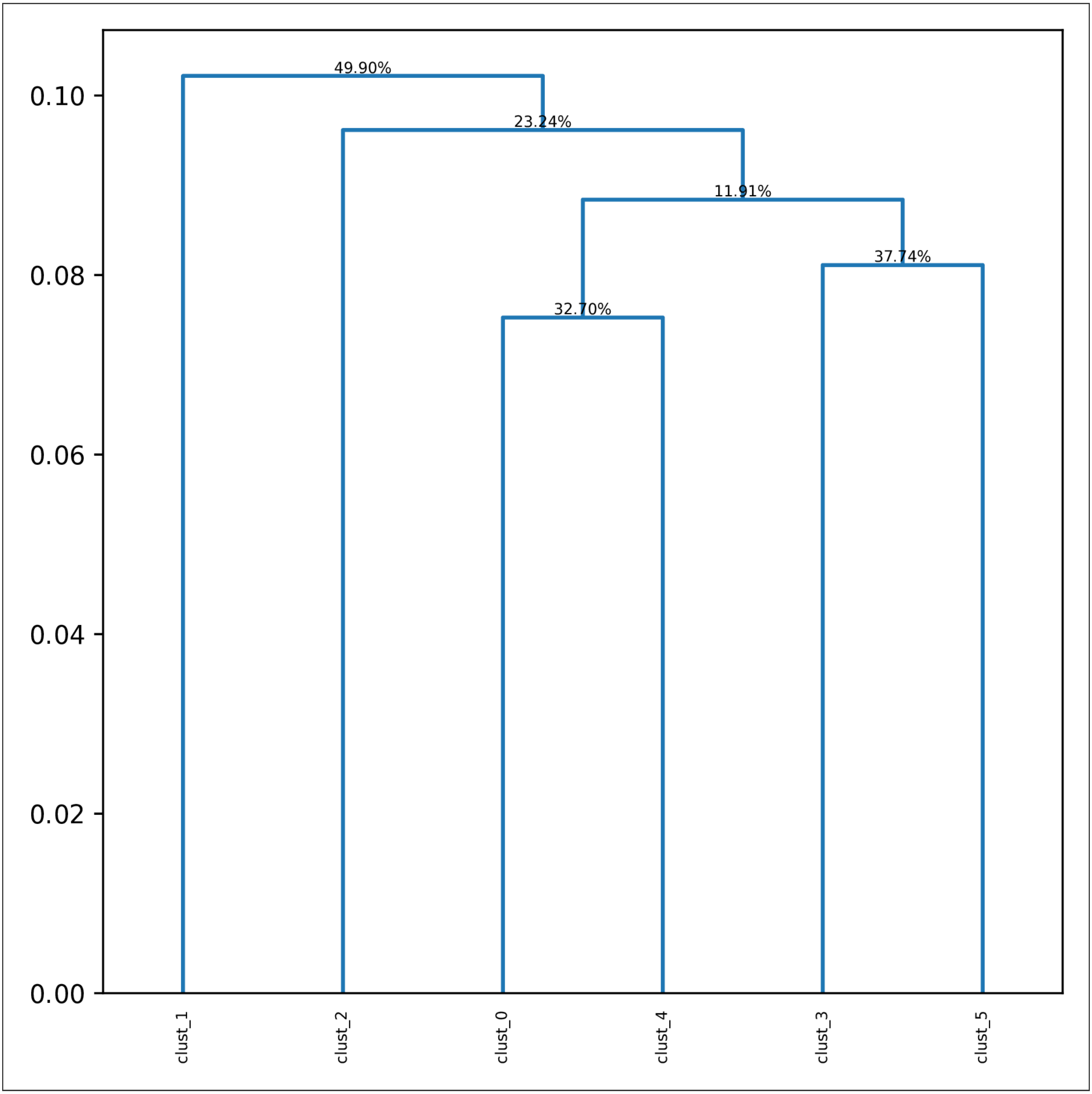

Sample: G0032 with 500 cells per cluster

Min. read depth: 10,  Mean UMIs per cluster:  4,092,682

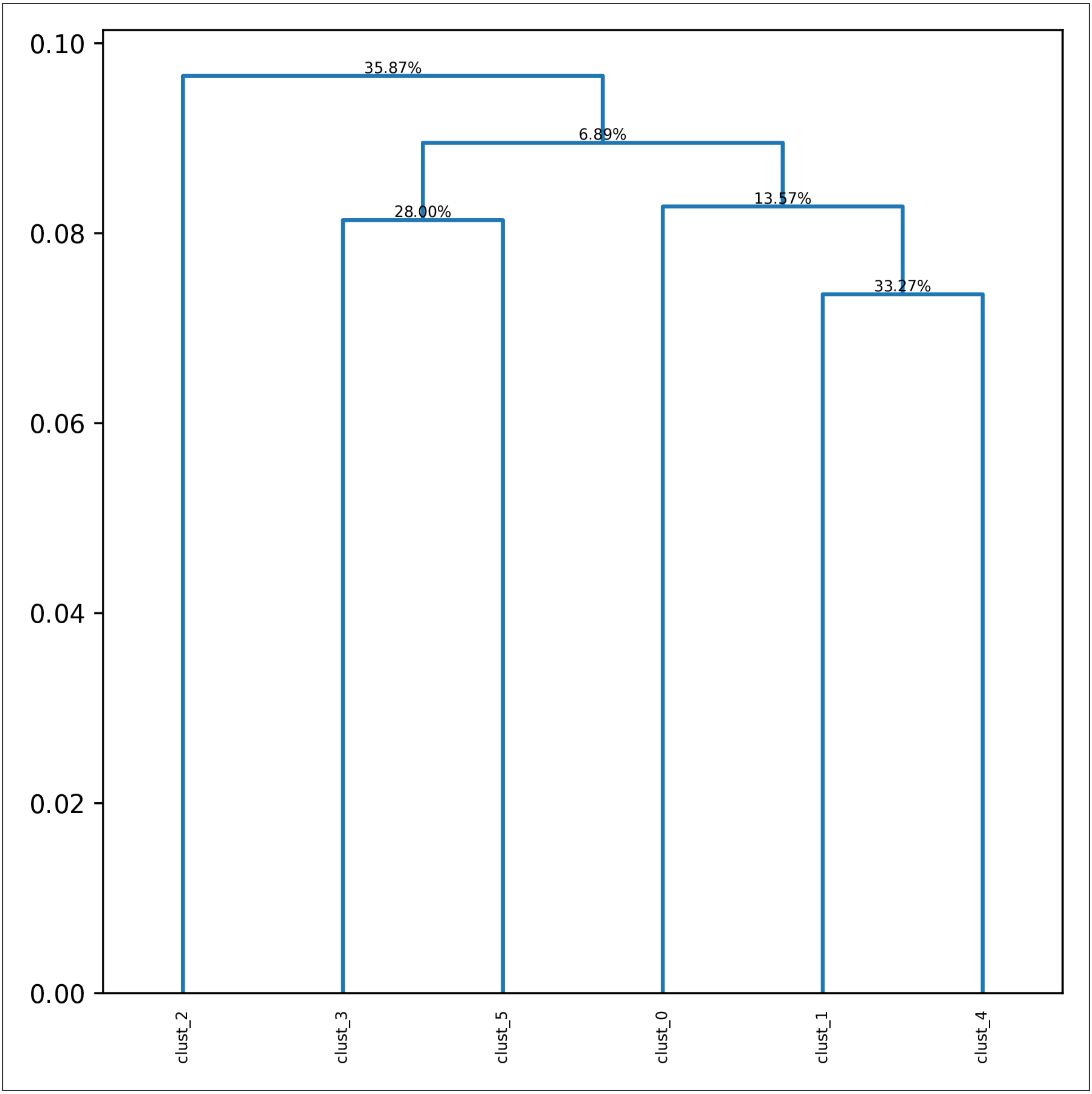

Sample: G0032 with 100 cells per cluster

Min. read depth: 10, Mean UMIs per cluster: 1,138,374

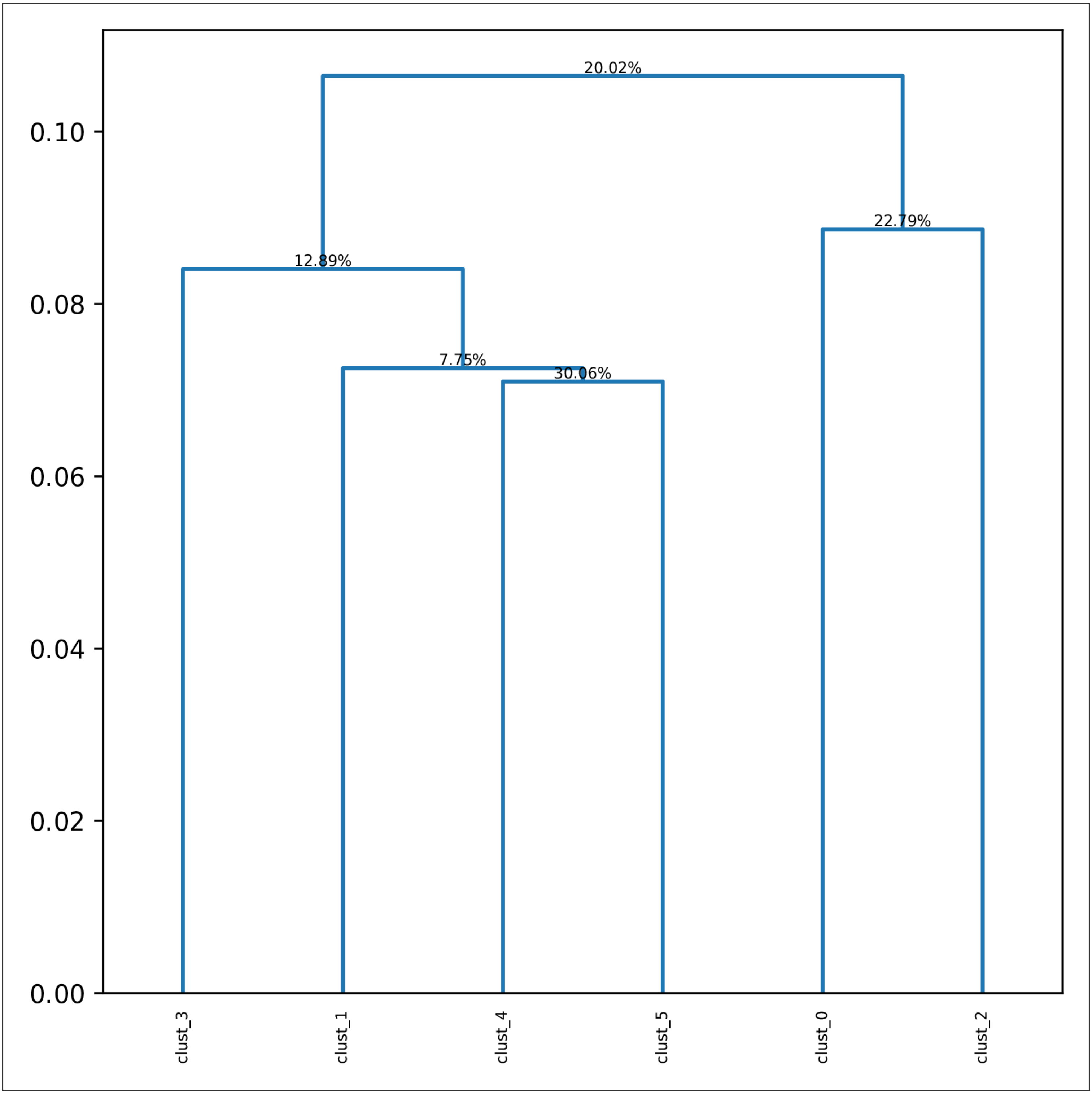

Sample: G0032 with 50 cells per cluster

Min. read depth: 10,  Mean UMIs per cluster:  567,553

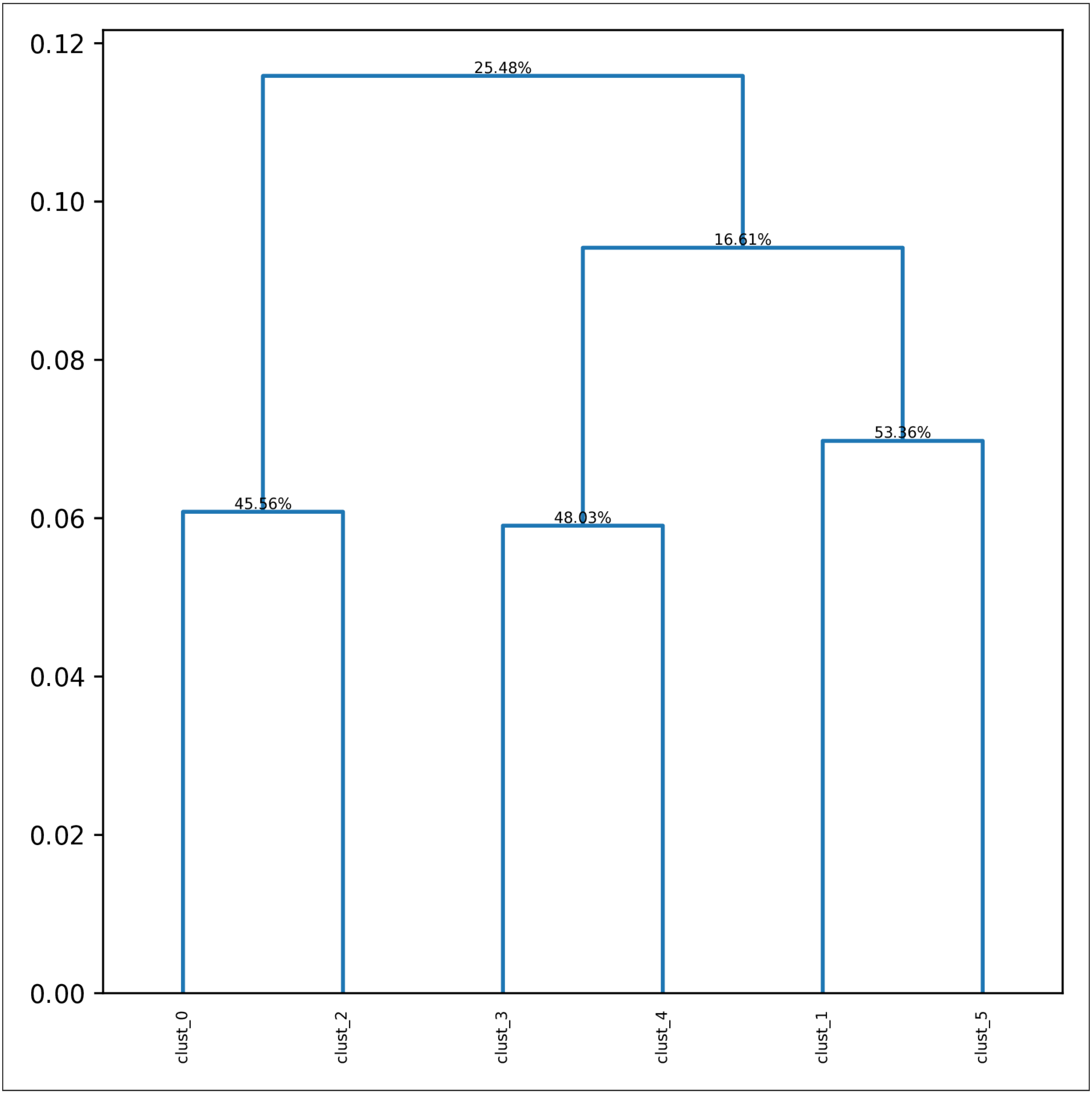

Sample: G0032 with 20 cells per cluster

Min. read depth: 10,  Mean UMIs per cluster:  239,542

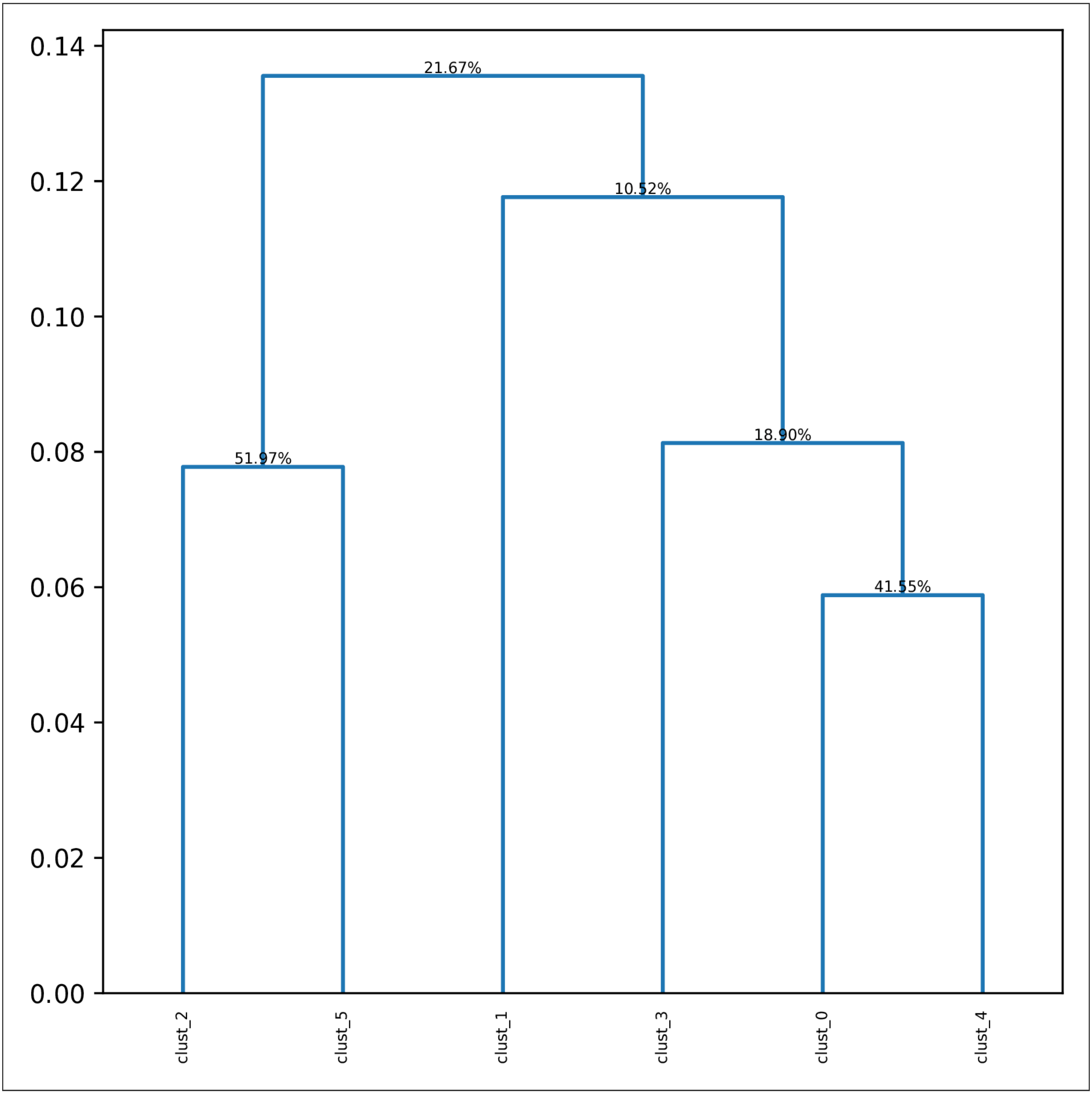

Sample: G0032 with 10 cells per cluster
Min. read depth: 10,  Mean UMIs per cluster:  122,097

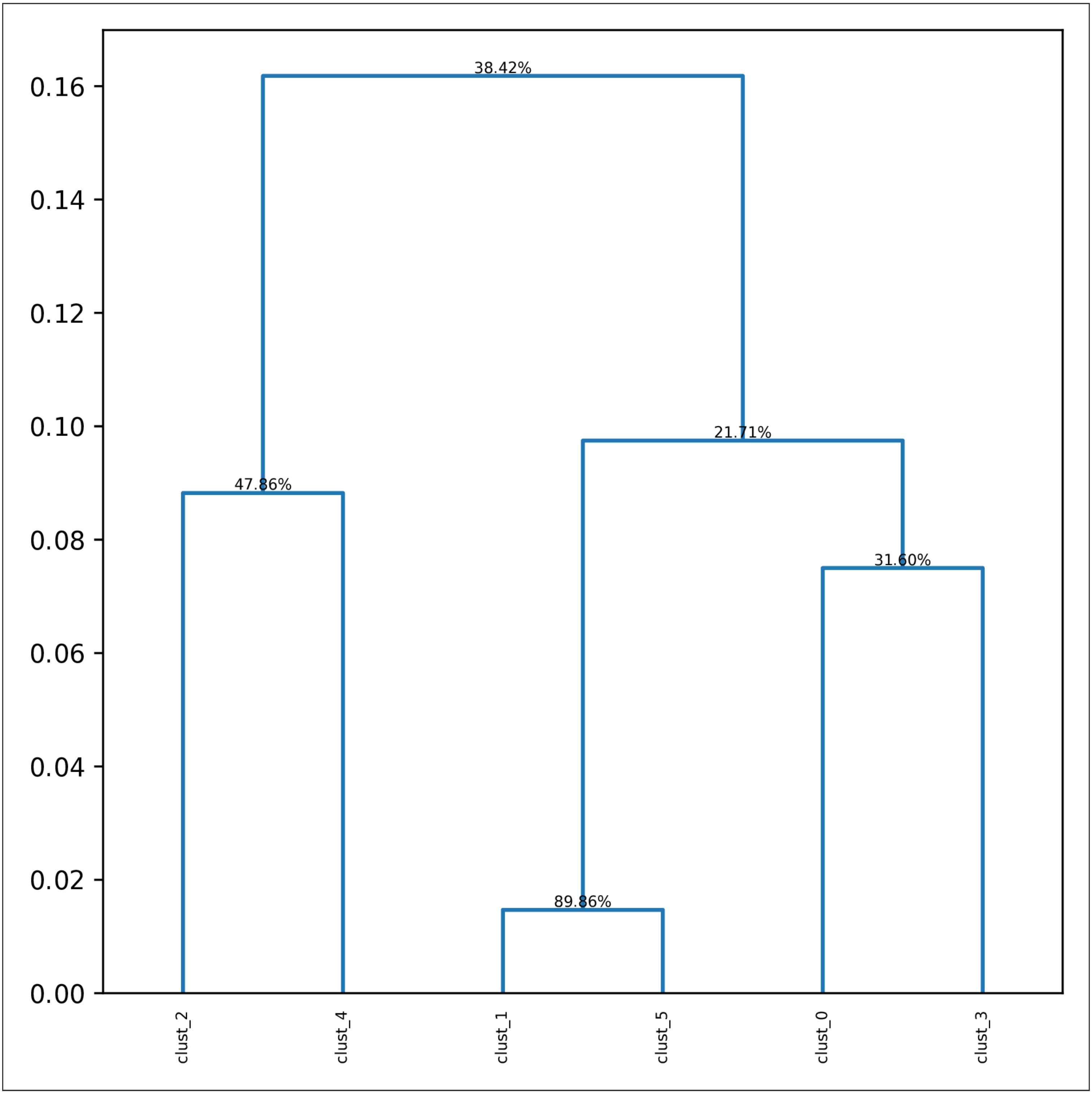

Sample: G0032 with 2 cells per cluster

Min. read depth: 10,  Mean UMIs per cluster:  22,199

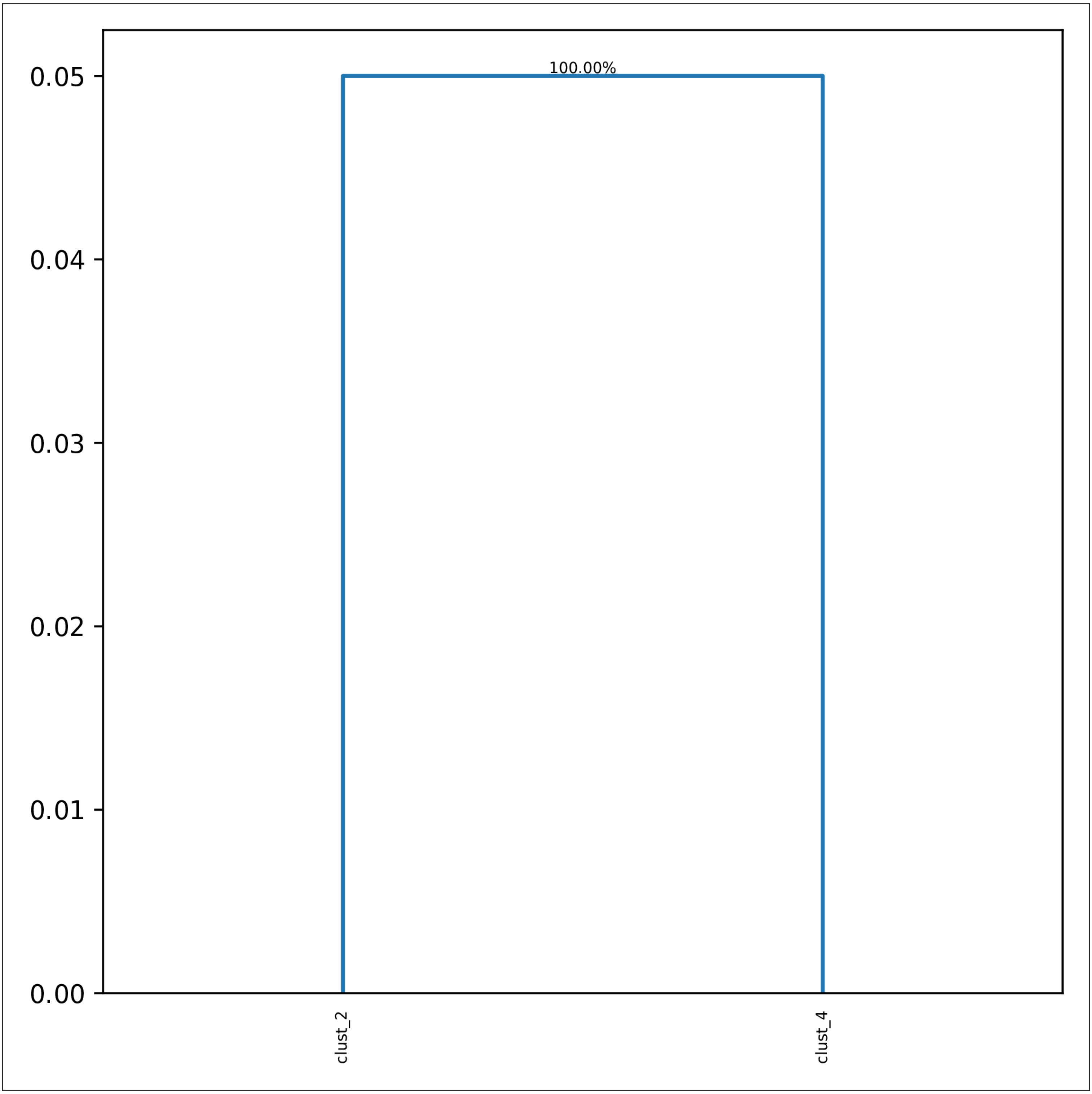

Sample: G0034 with all cells per cluster

Min. read depth: 10,  Mean UMIs per cluster:  72,864,441

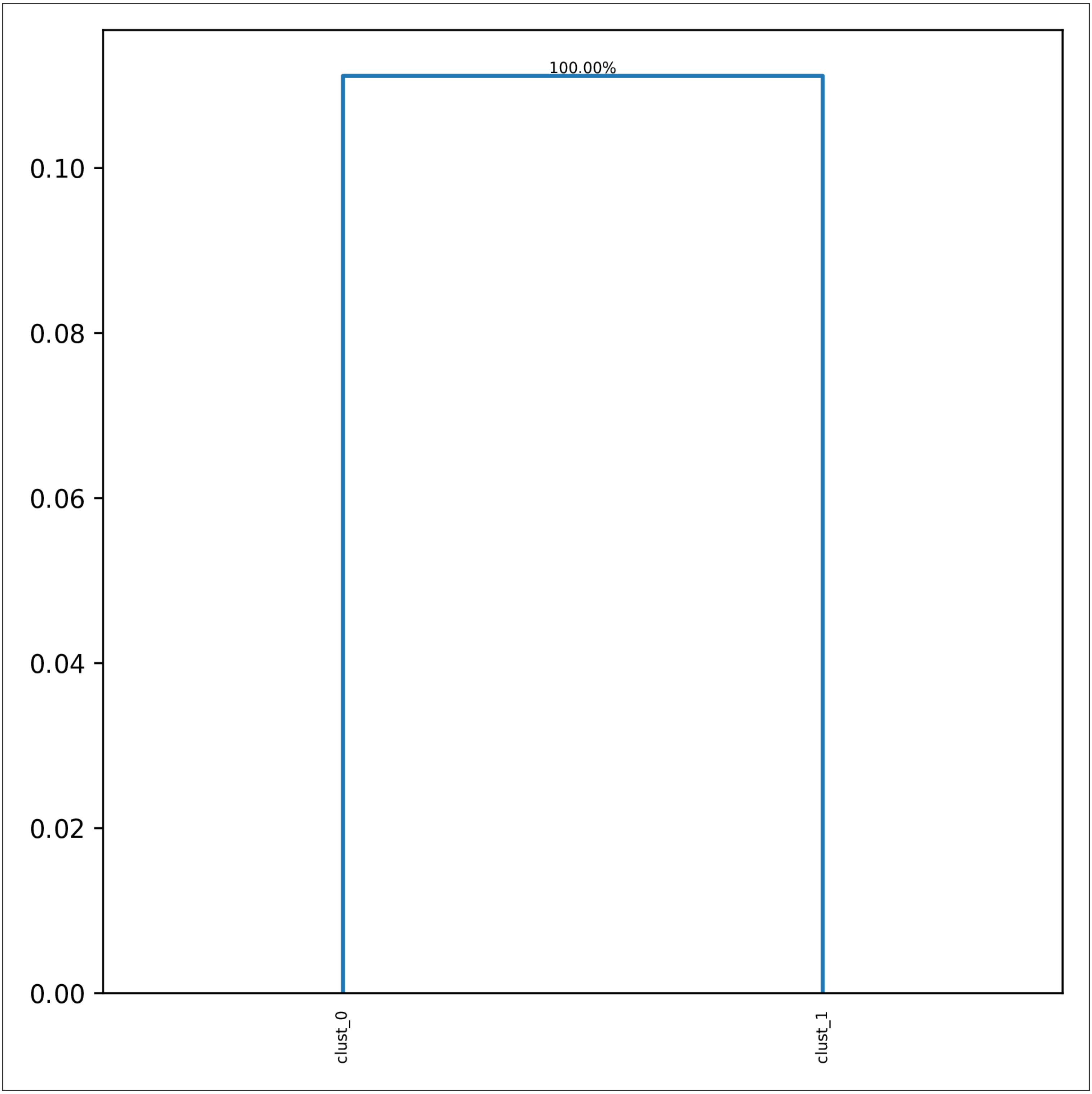

Sample: G0034 with 500 cells per cluster

Min. read depth: 10,  Mean UMIs per cluster:  4,205,904

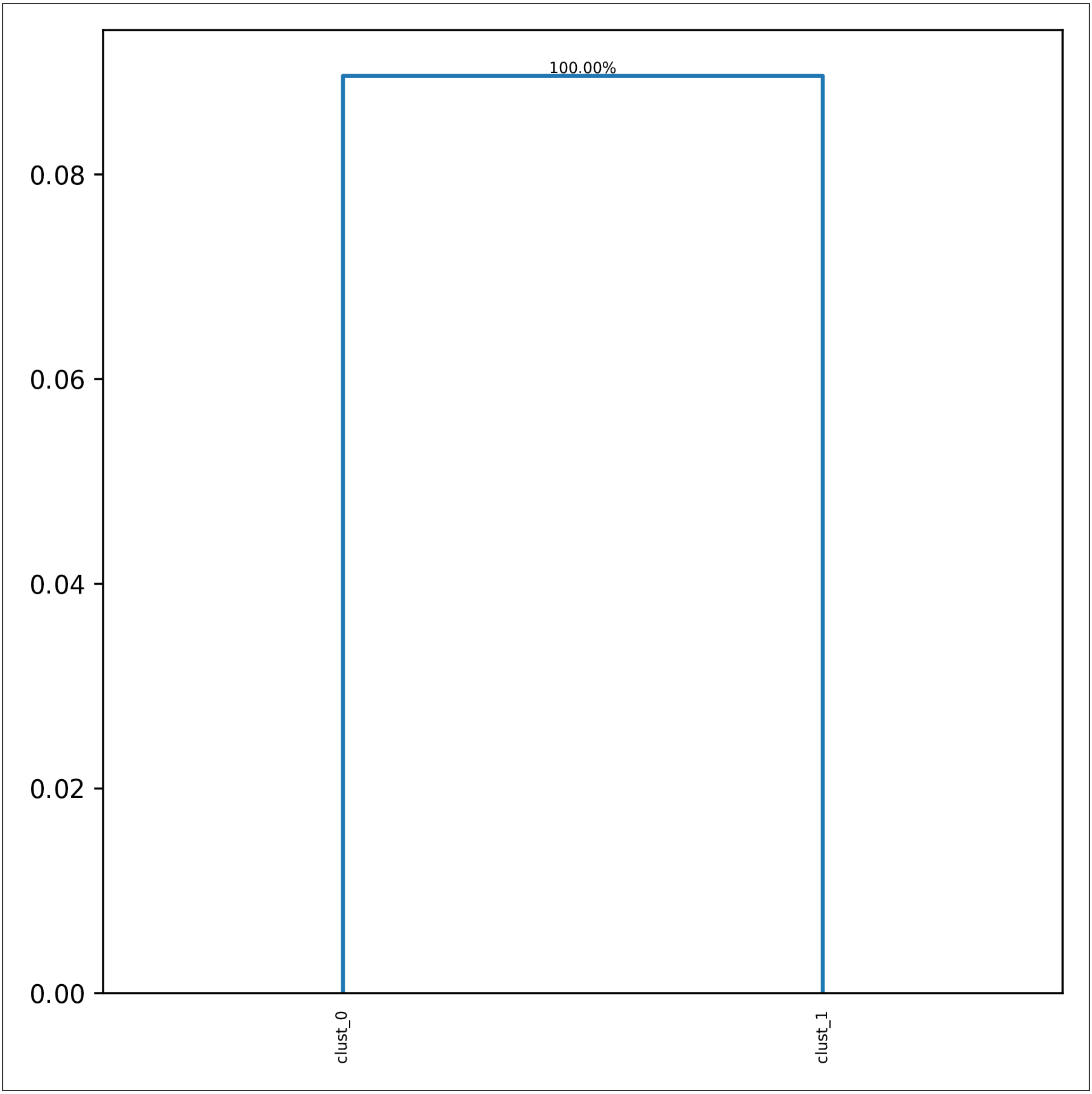

Sample: G0034 with 100 cells per cluster

Min. read depth: 10,  Mean UMIs per cluster:  921,572

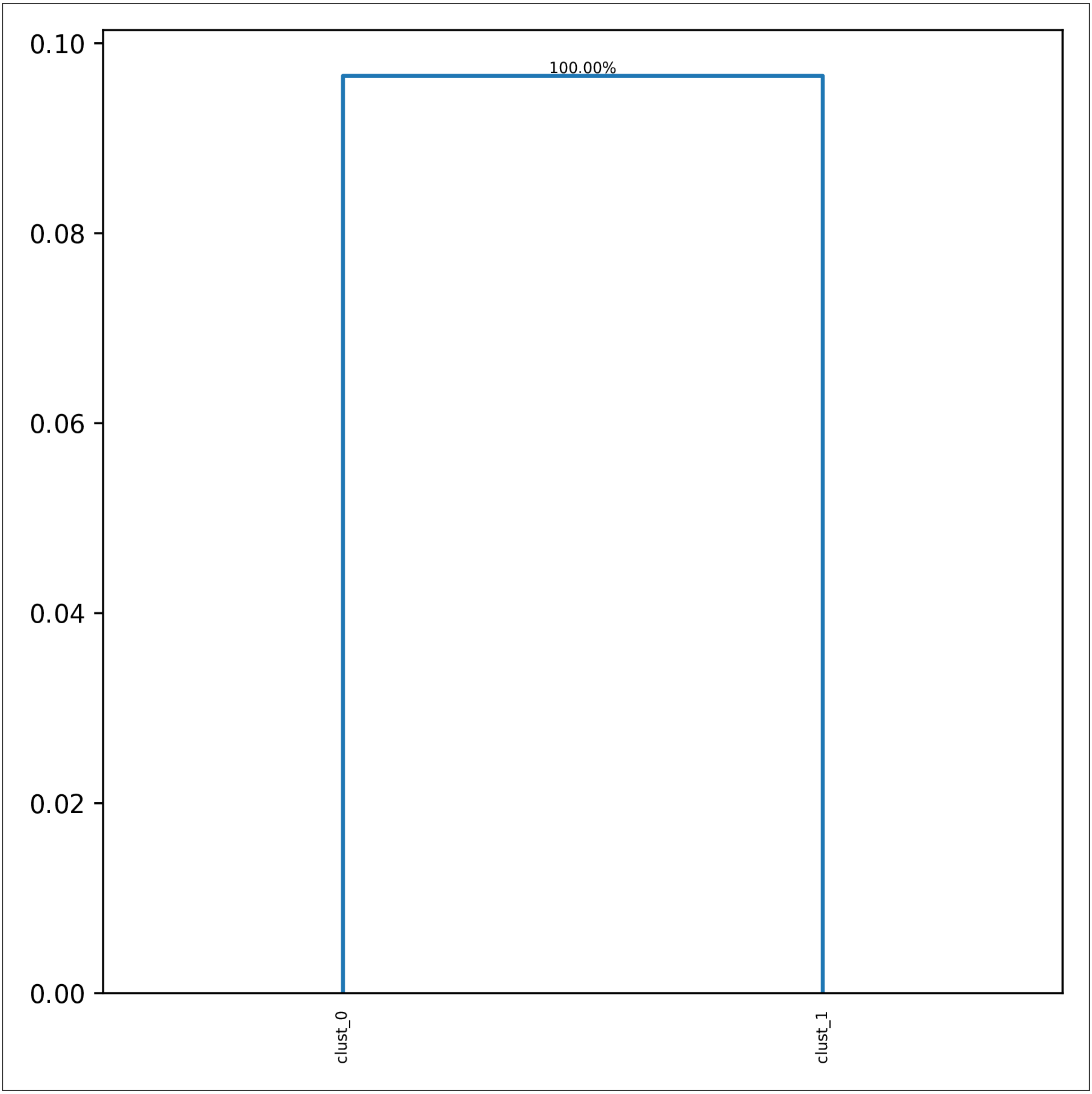

Sample: G0034 with 50 cells per cluster

Min. read depth: 10,  Mean UMIs per cluster:  477,632

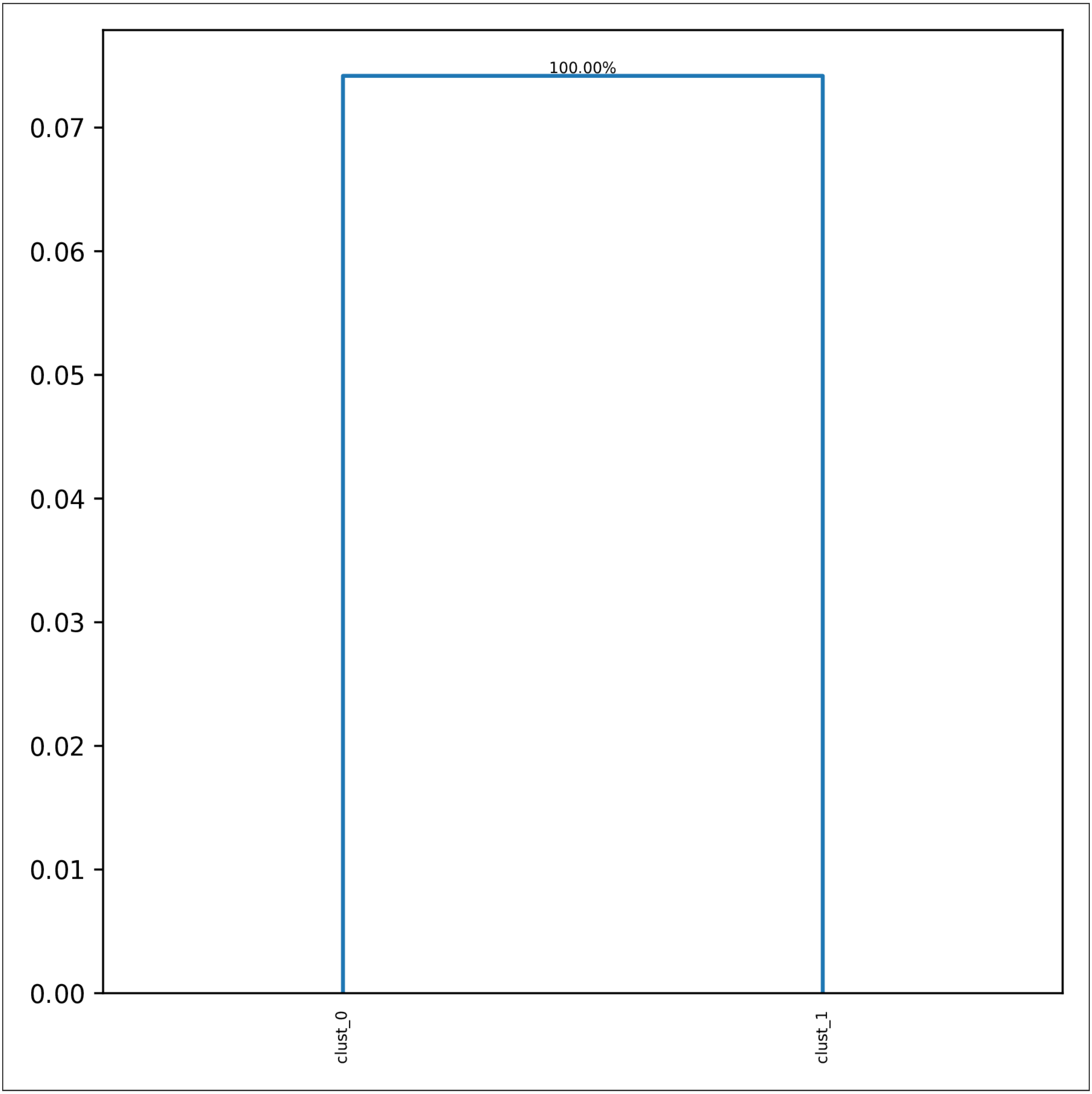

Sample: G0034 with 20 cells per cluster

Min. read depth: 10,  Mean UMIs per cluster:  192,316

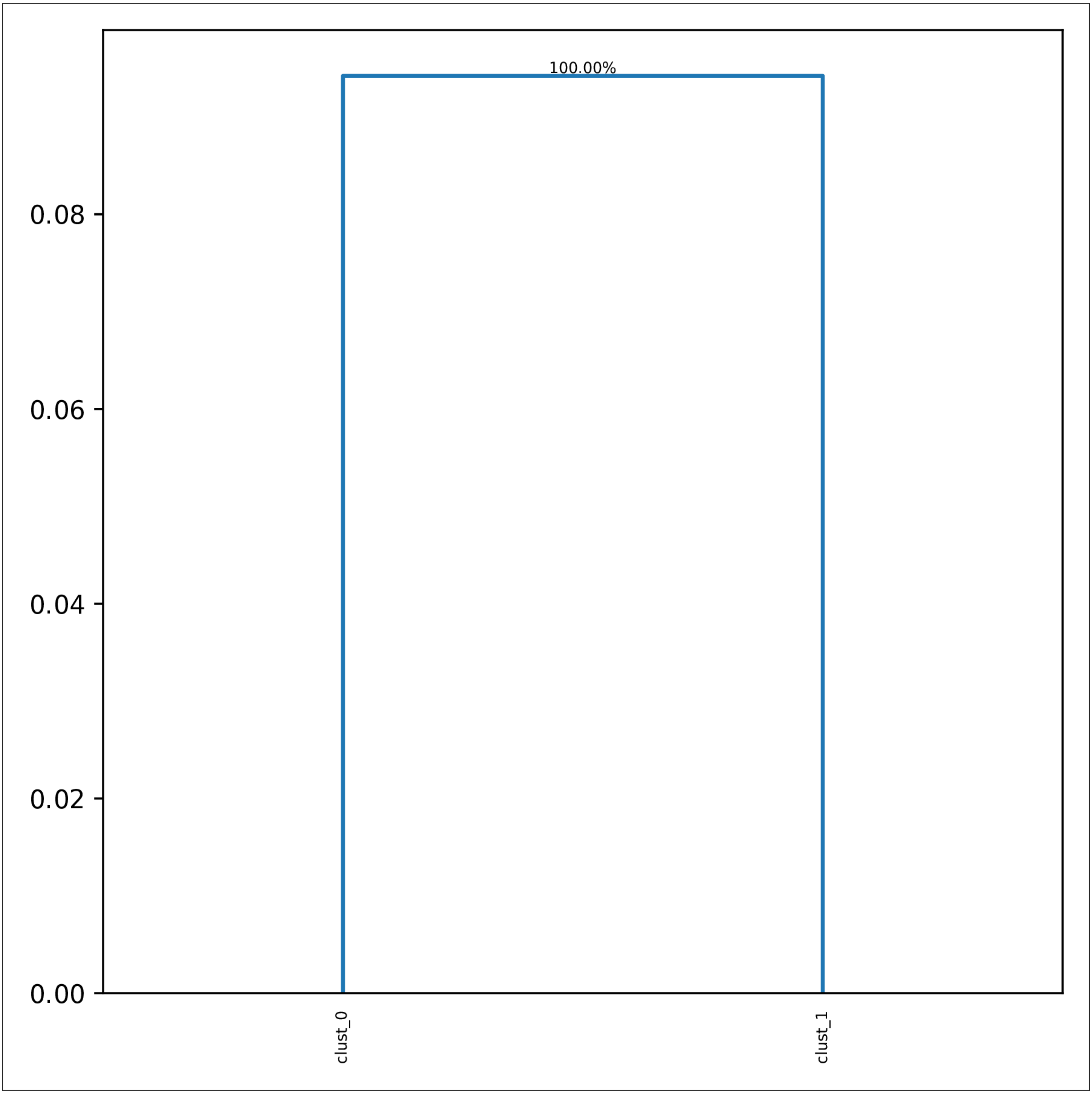

Sample: G0034 with 10 cells per cluster

Min. read depth: 10,  Mean UMIs per cluster:  83,936

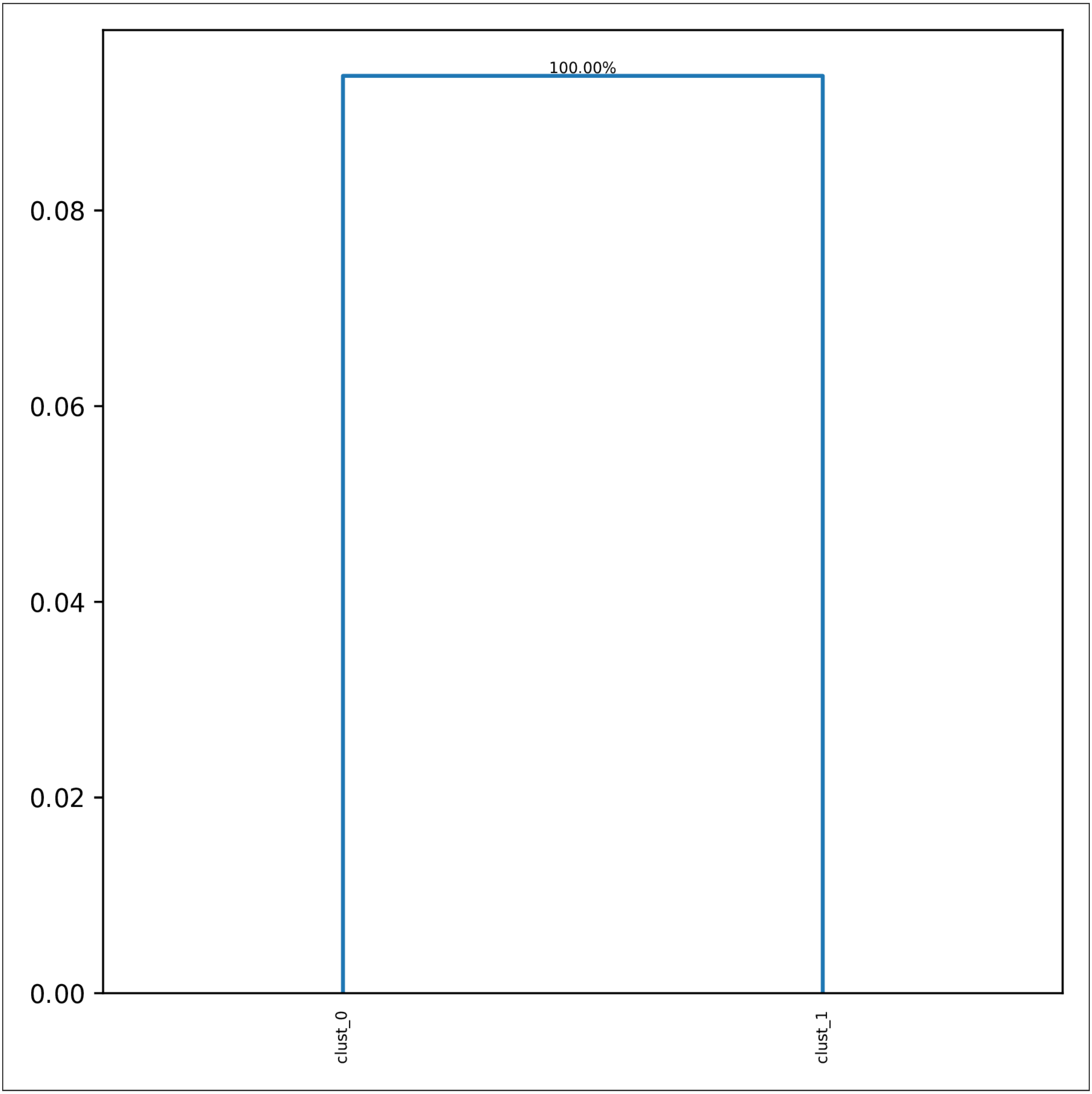

Sample: G0034 with 5 cells per cluster

Min. read depth: 10,  Mean UMIs per cluster:  47,550

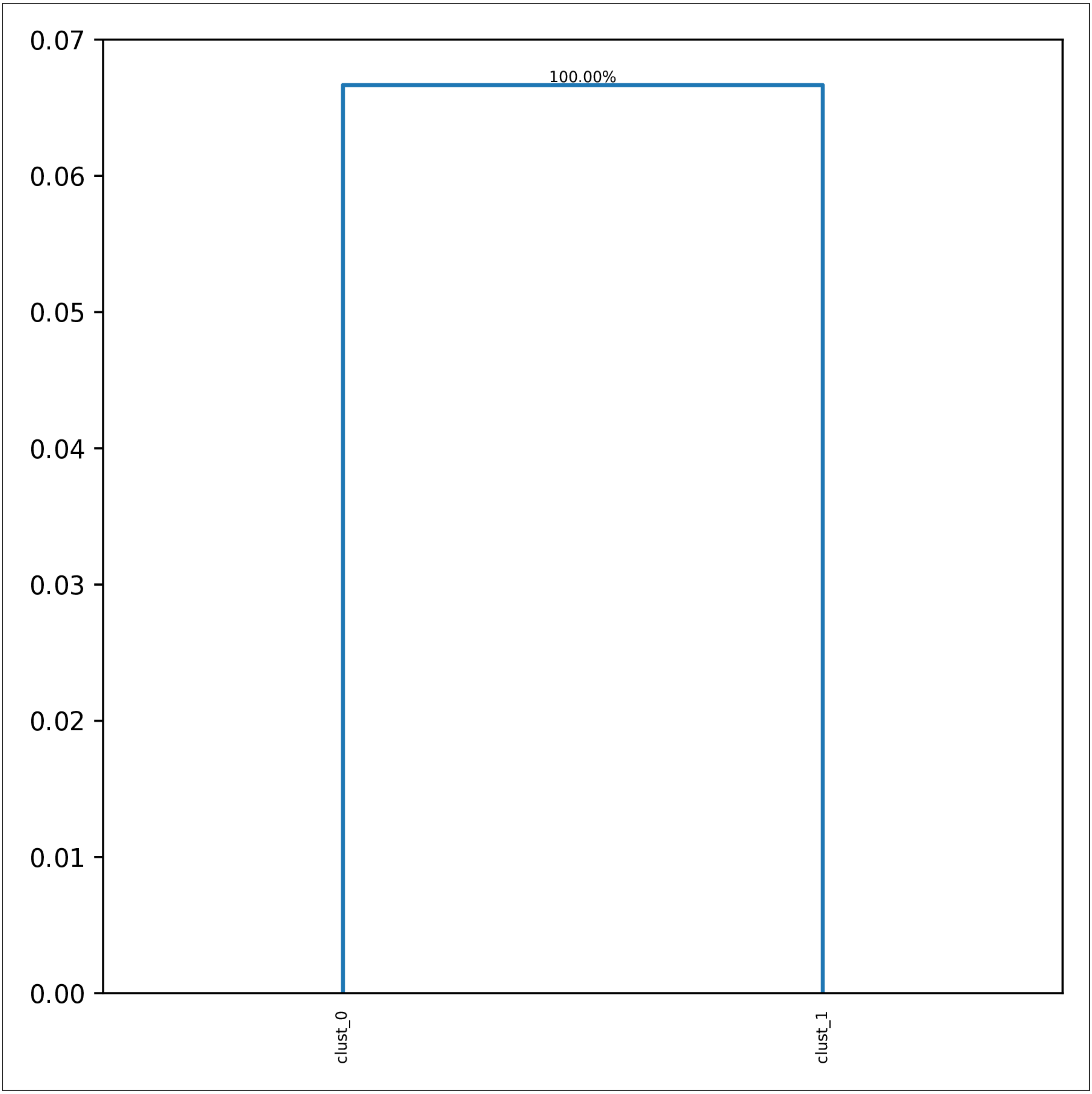

Sample: G0034 with 4 cells per cluster

Min. read depth: 10,  Mean UMIs per cluster:  39,302

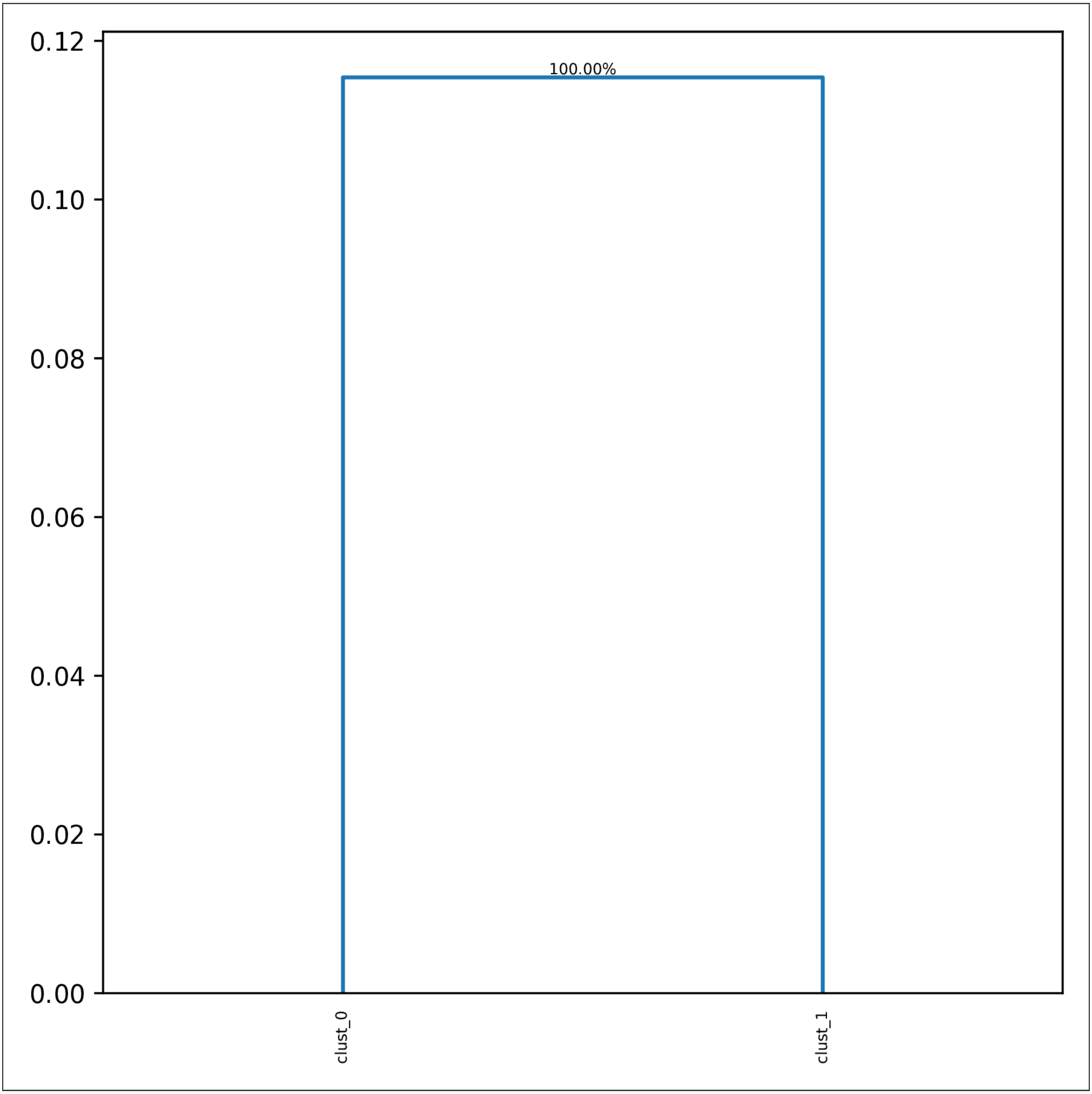

Sample: G0034 with 3 cells per cluster

Min. read depth: 10,  Mean UMIs per cluster:  30,558

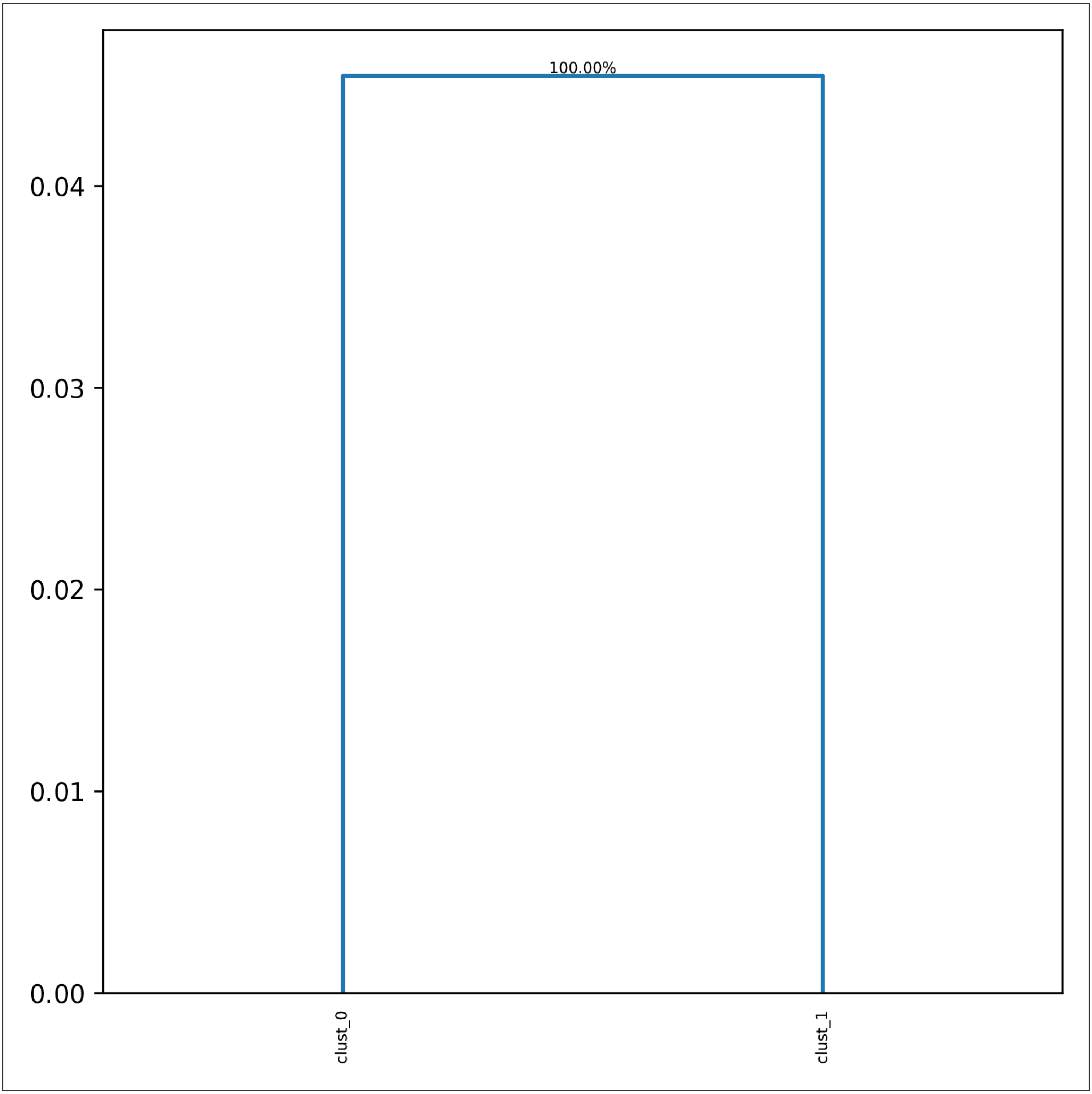

Sample: G0035 with all cells per cluster

Min. read depth: 10,  Mean UMIs per cluster:  4,853,217

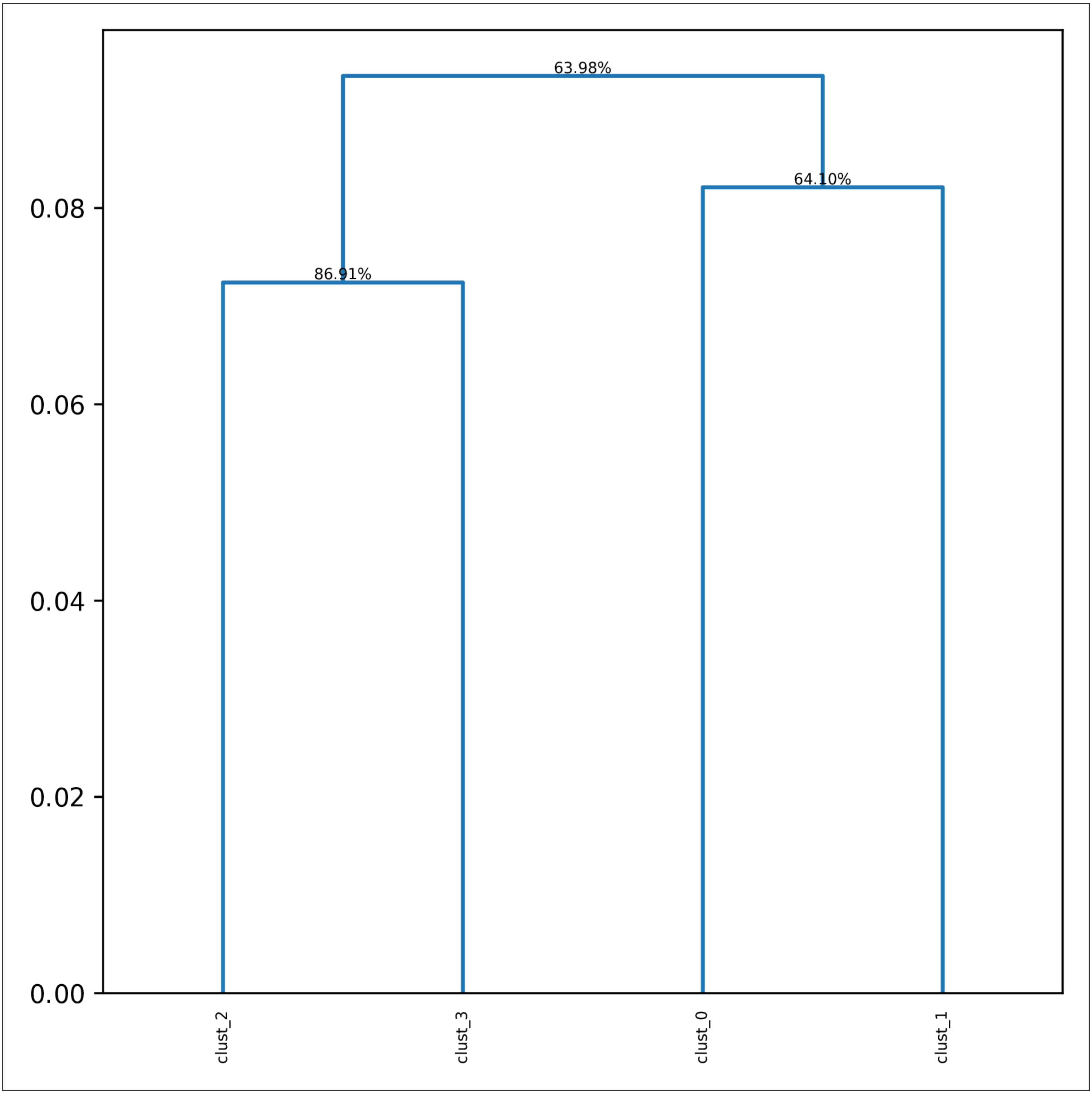

Sample: G0035 with 500 cells per cluster

Min. read depth: 10,  Mean UMIs per cluster:  3,914,214

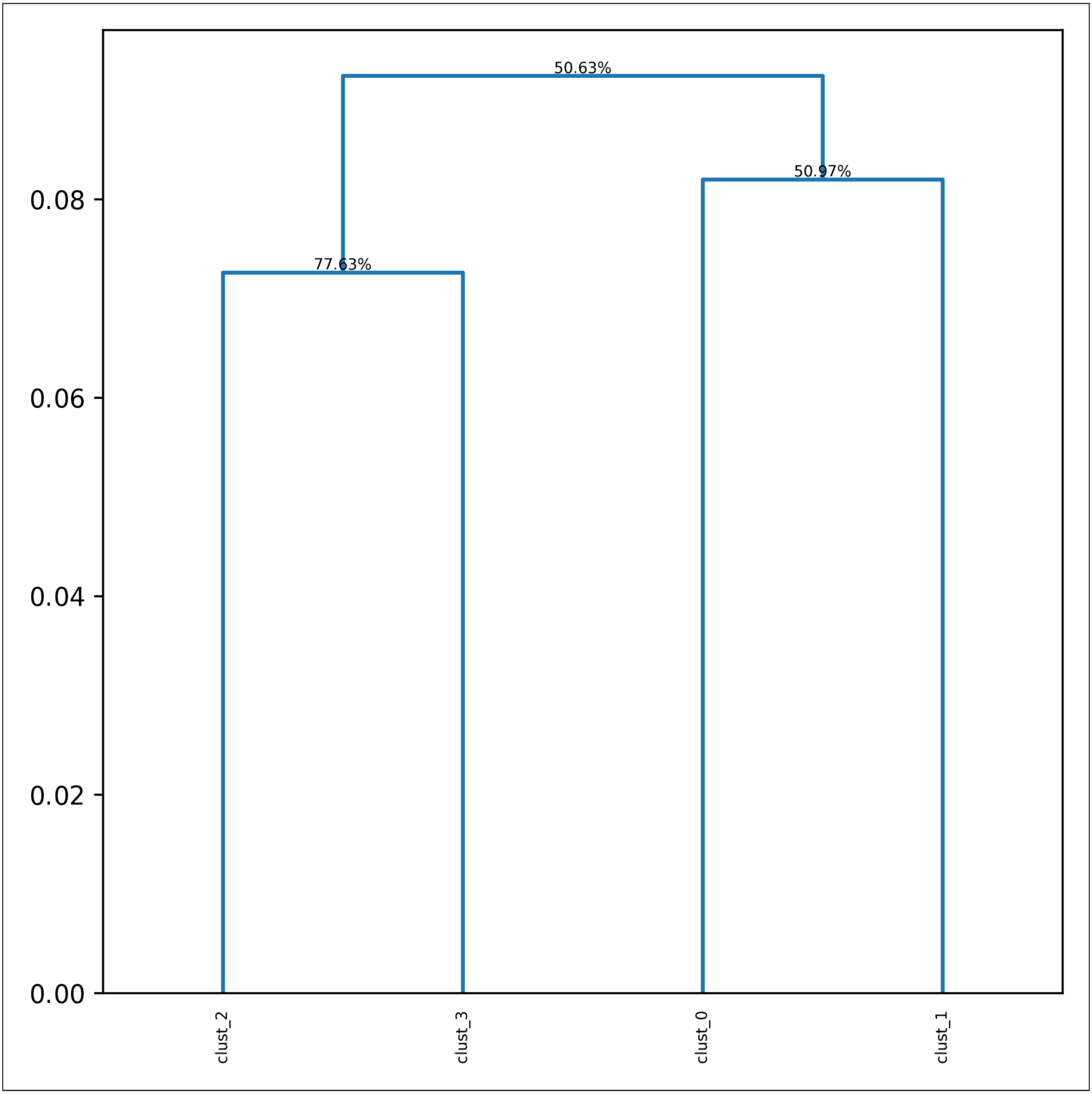

Sample: G0035 with 100 cells per cluster

Min. read depth: 10,  Mean UMIs per cluster:  1,246,536

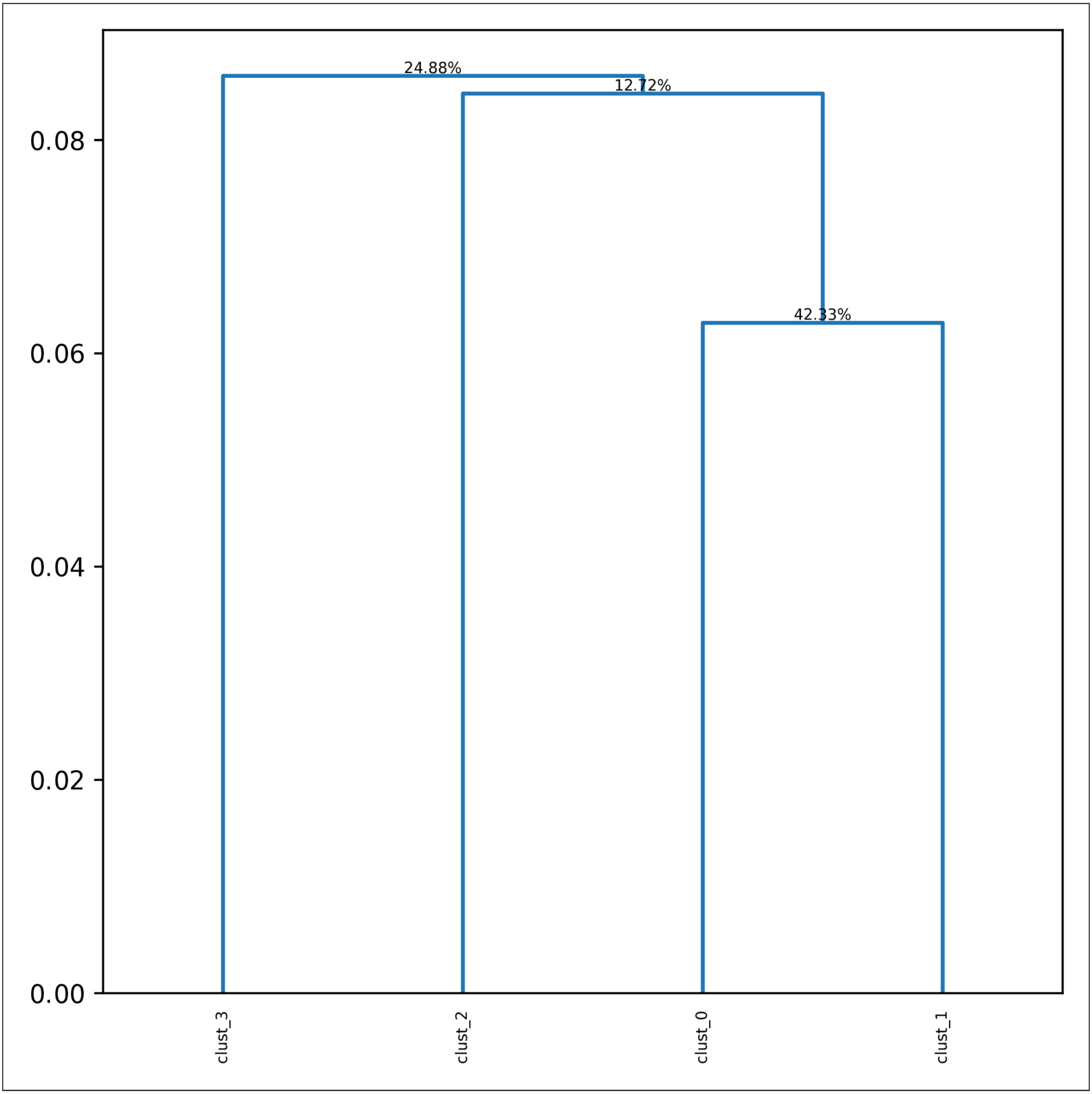

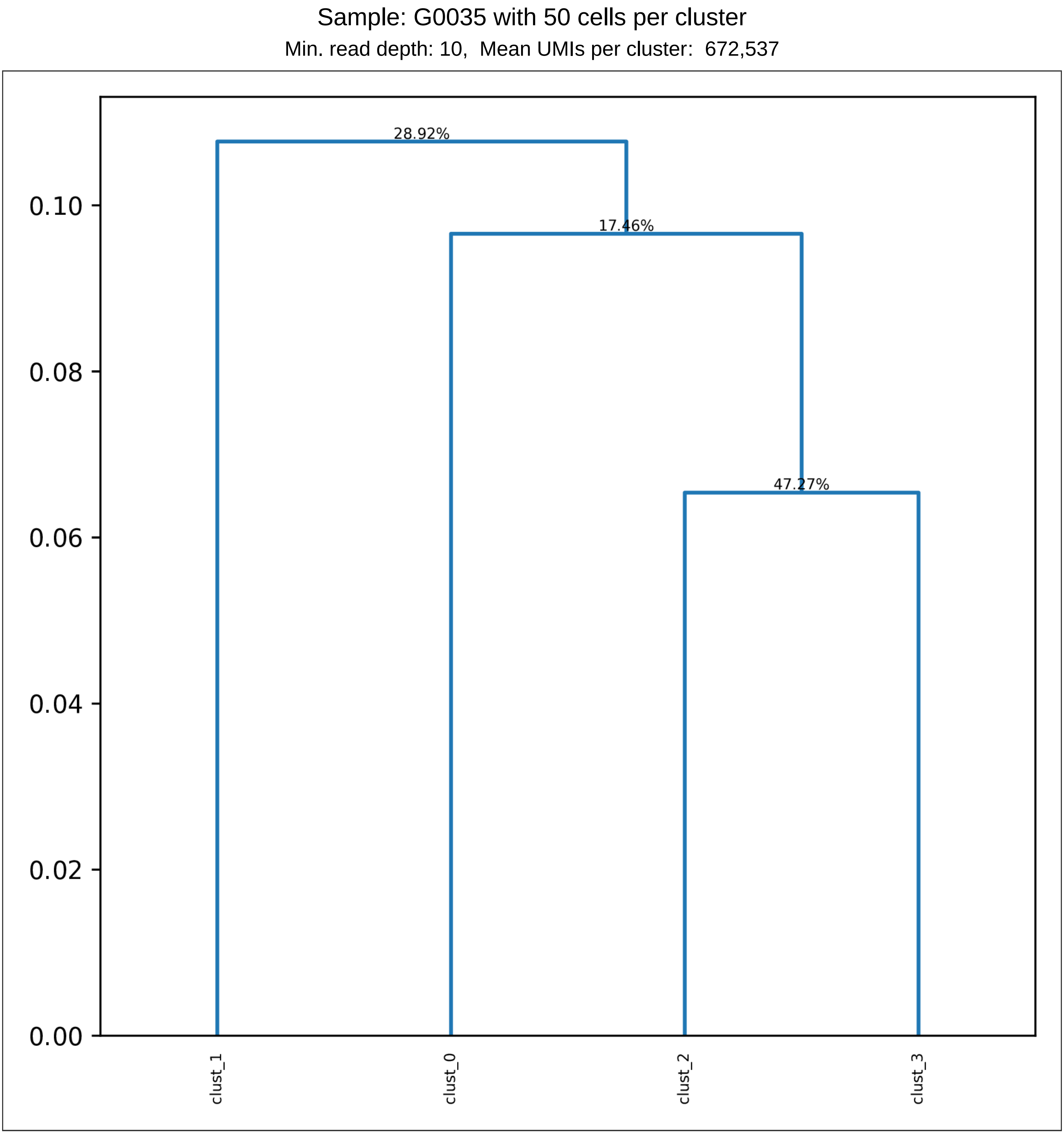
Sample: G0035 with 50 cells per cluster
Min. read depth: 10,  Mean UMIs per cluster:  672,537
28.92%
17.46%
47.27%
0.10
0.08
0.06
0.04
0.02
0.00
clust_1
clust_0
clust_2
clust_3

Sample: G0035 with 20 cells per cluster
Min. read depth: 10,  Mean UMIs per cluster:  249,095

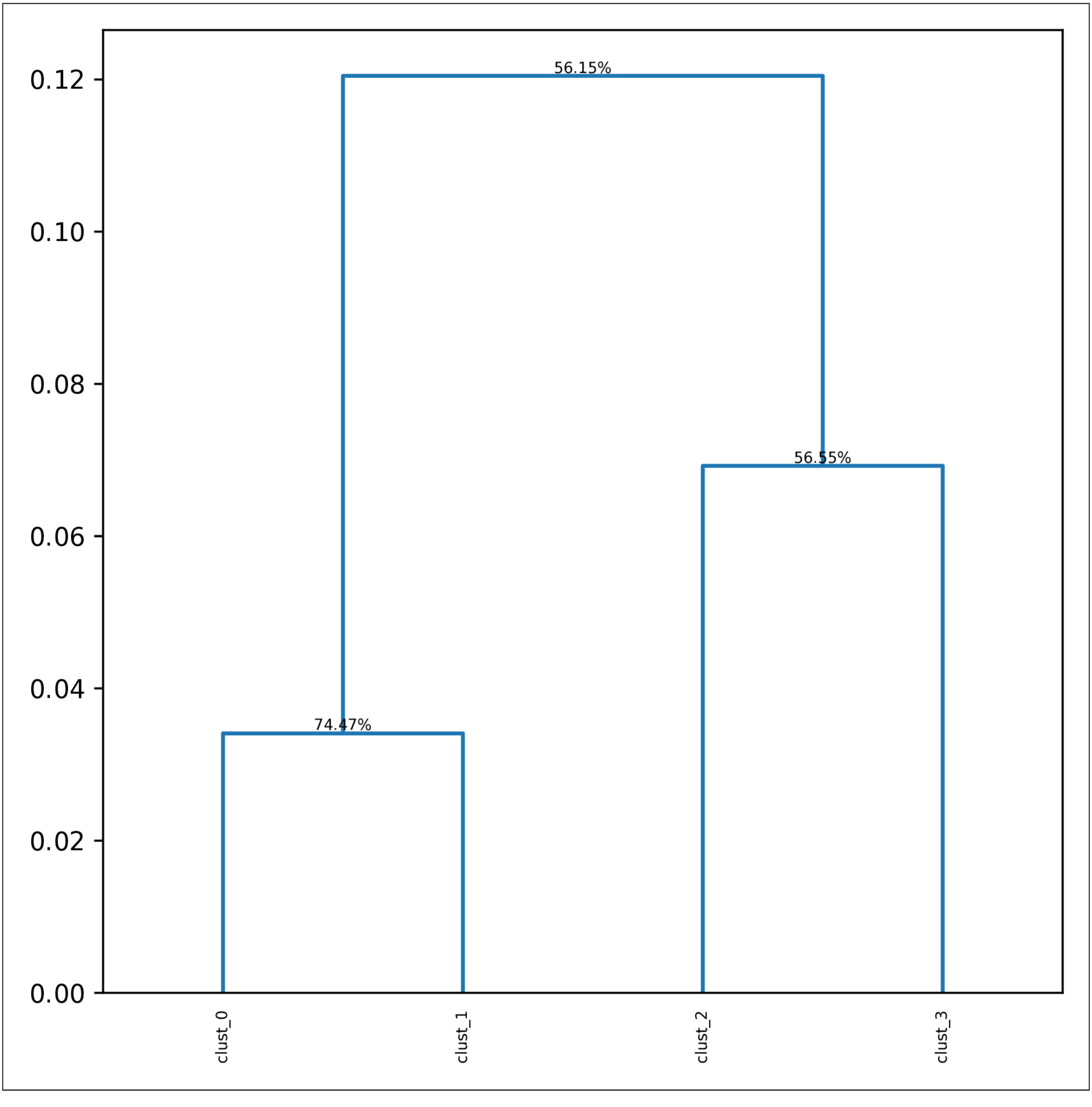

Sample: G0035 with 10 cells per cluster

Min. read depth: 10,  Mean UMIs per cluster:  121,535

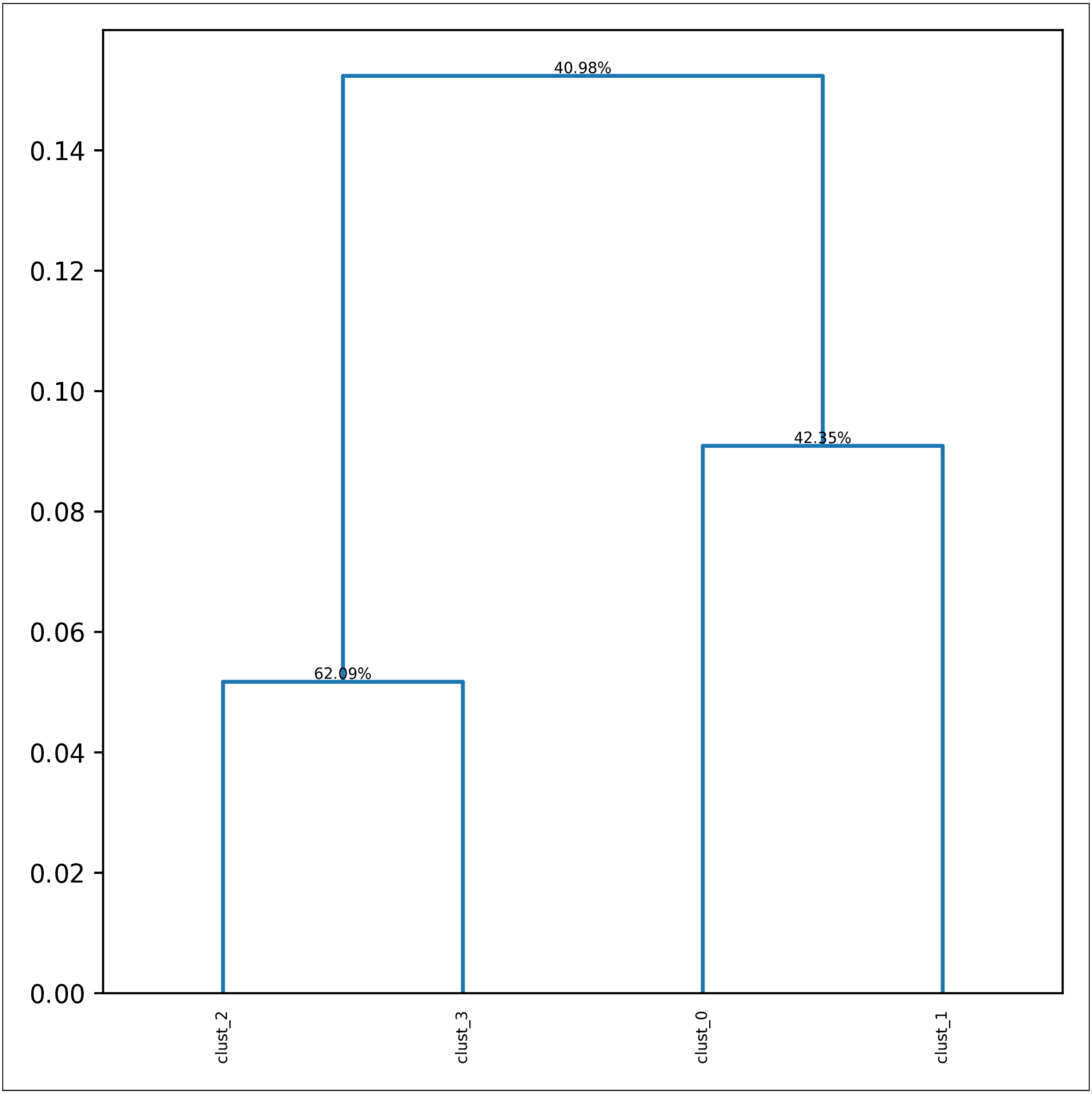

Sample: G0035 with 4 cells per cluster
Min. read depth: 10,  Mean UMIs per cluster:  52,790

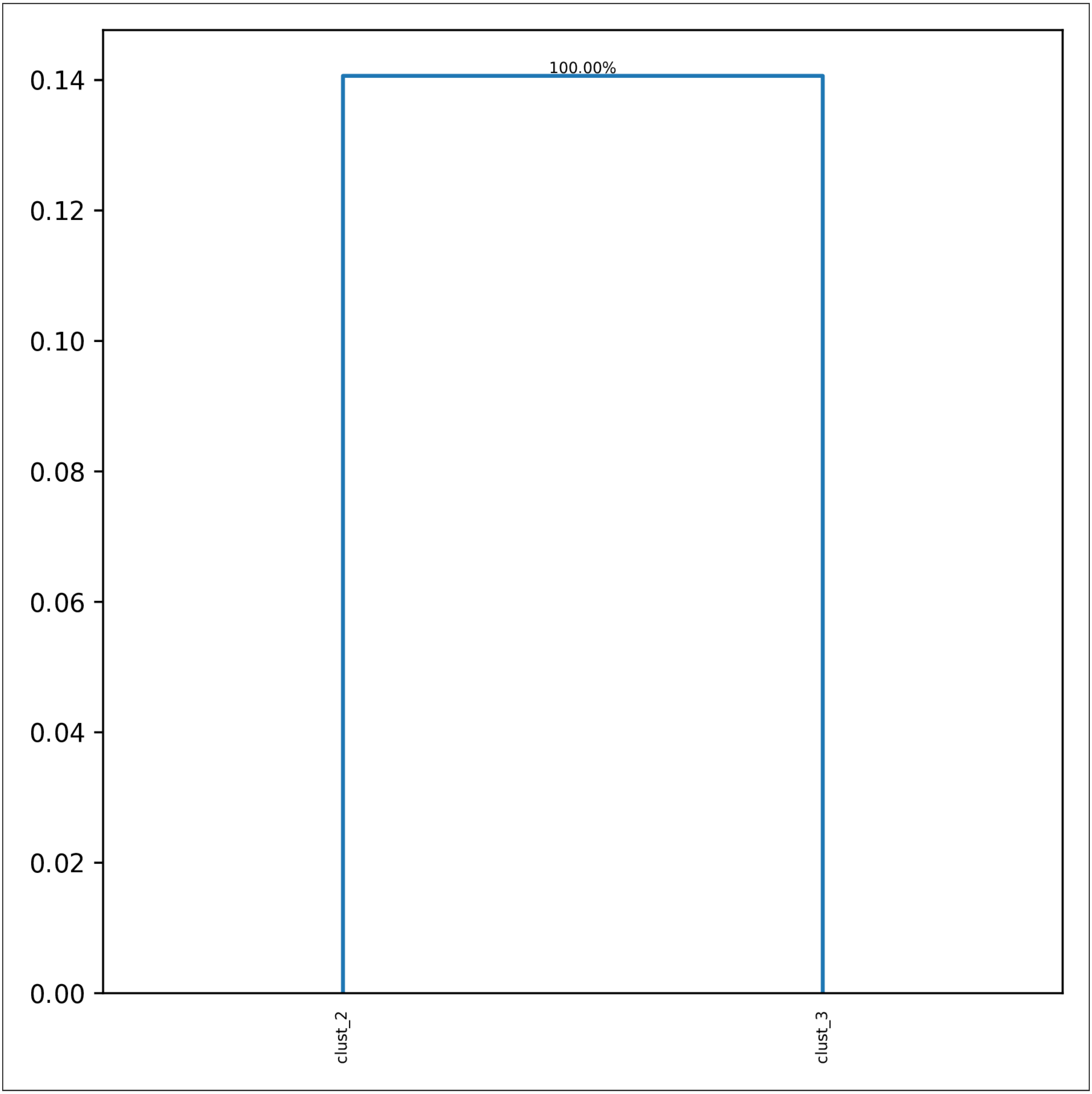

Sample: G0036 with all cells per cluster

Min. read depth: 10,  Mean UMIs per cluster:  7,430,598

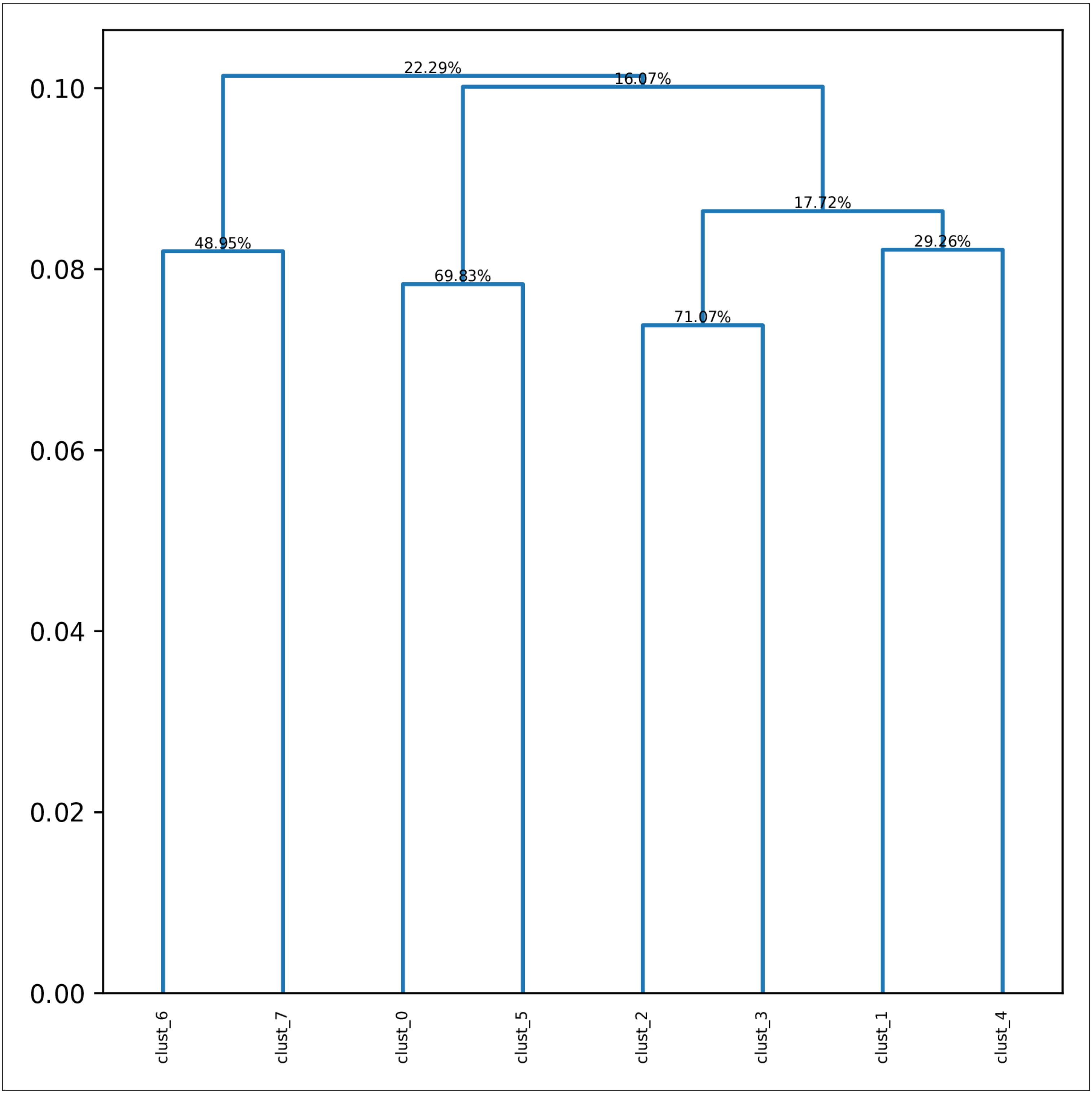

Sample: G0036 with 500 cells per cluster

Min. read depth: 10,  Mean UMIs per cluster:  2,591,780

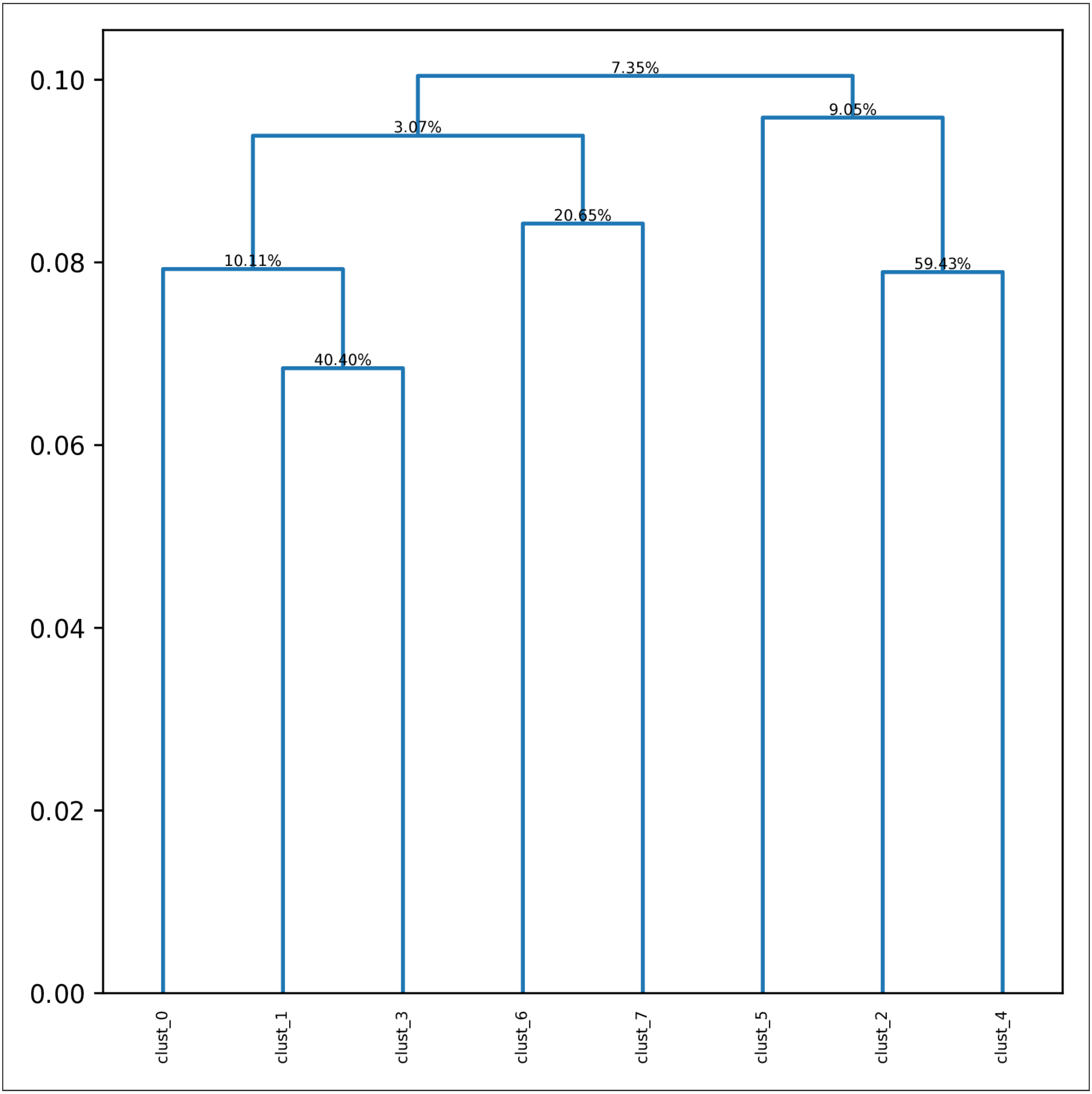

Sample: G0036 with 100 cells per cluster
Min. read depth: 10,  Mean UMIs per cluster:  553,897

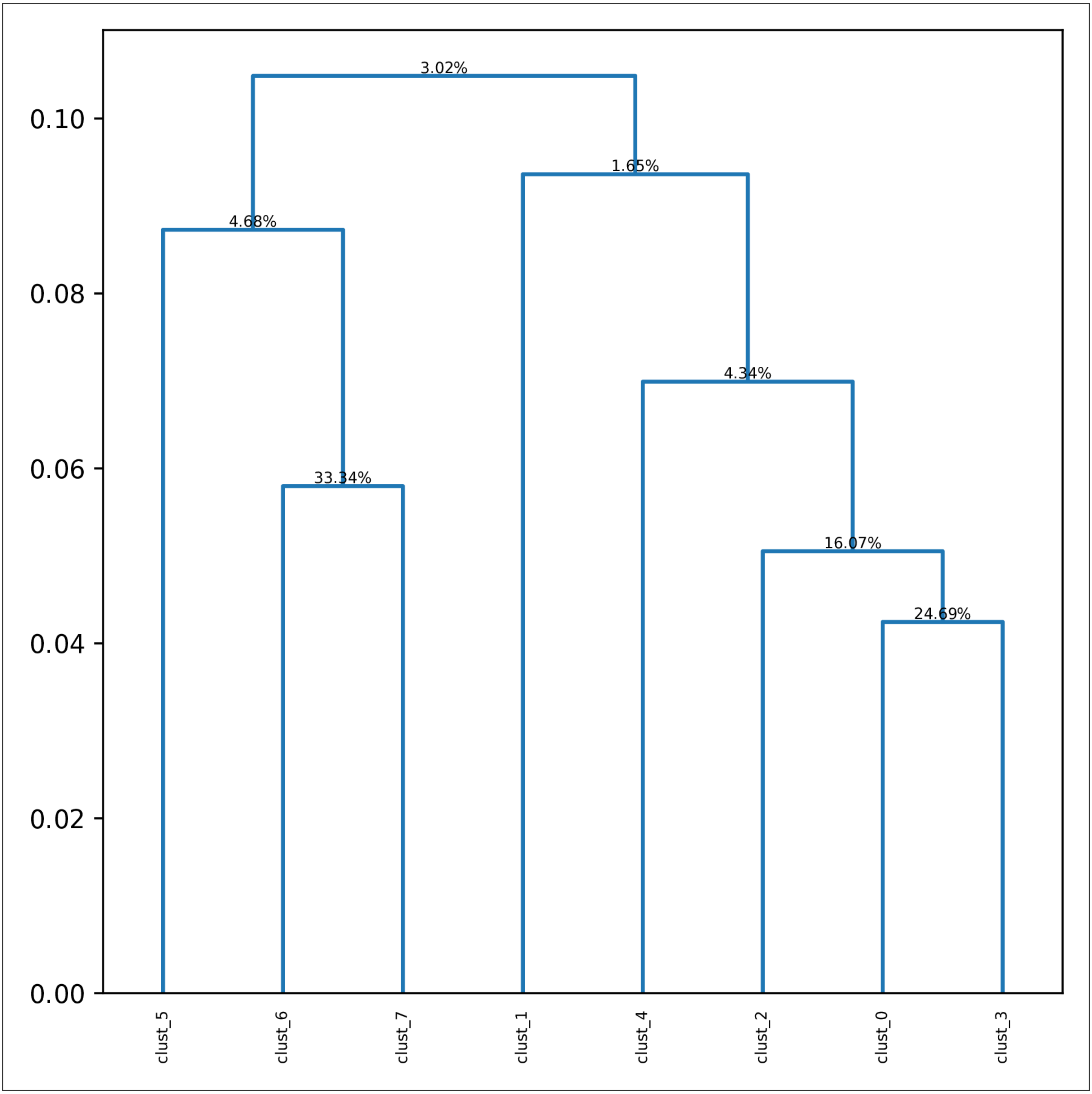

Sample: G0036 with 50 cells per cluster

Min. read depth: 10, Mean UMIs per cluster: 279,442

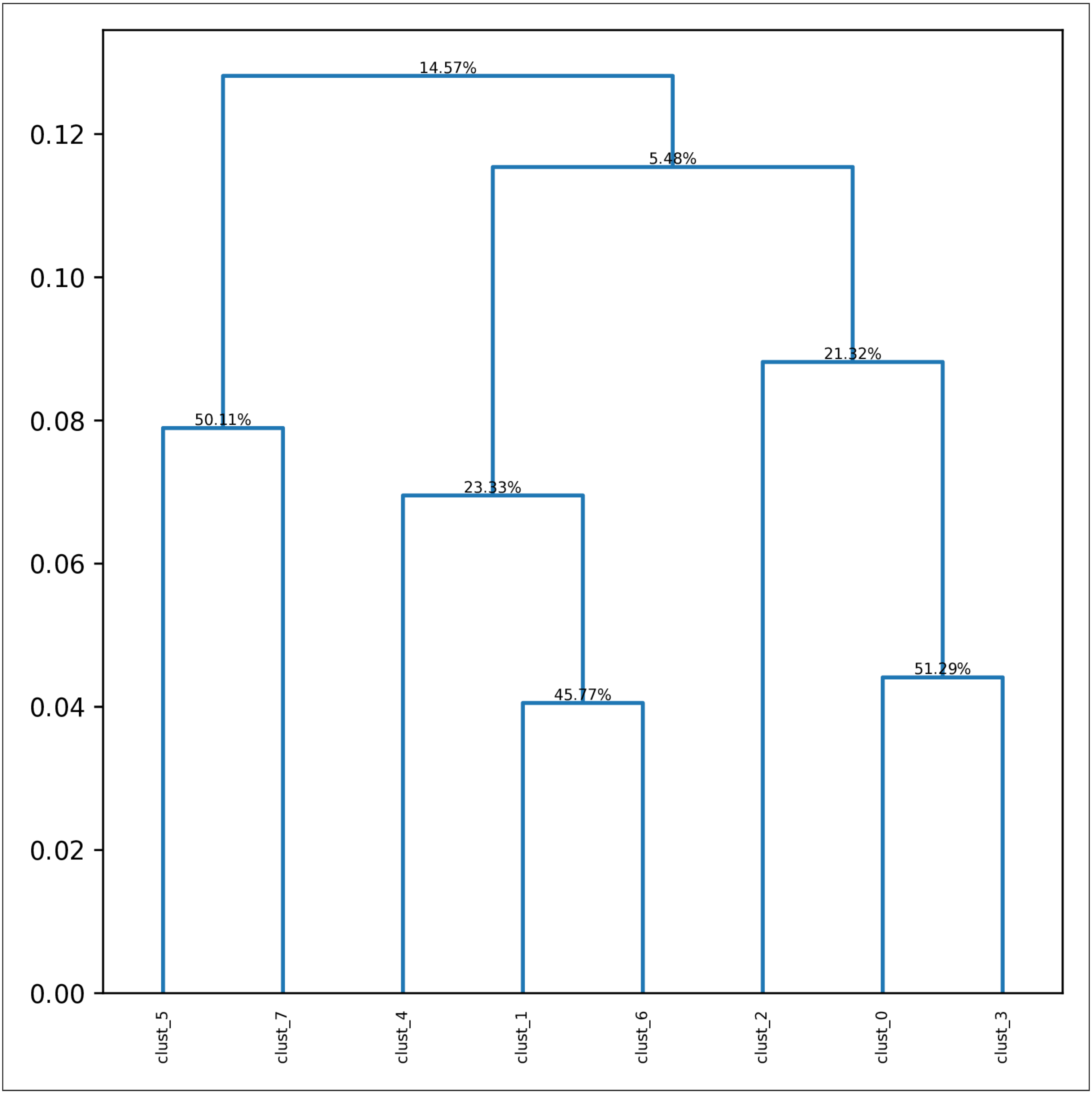

Sample: G0036 with 10 cells per cluster

Min. read depth: 10,  Mean UMIs per cluster:  56,619

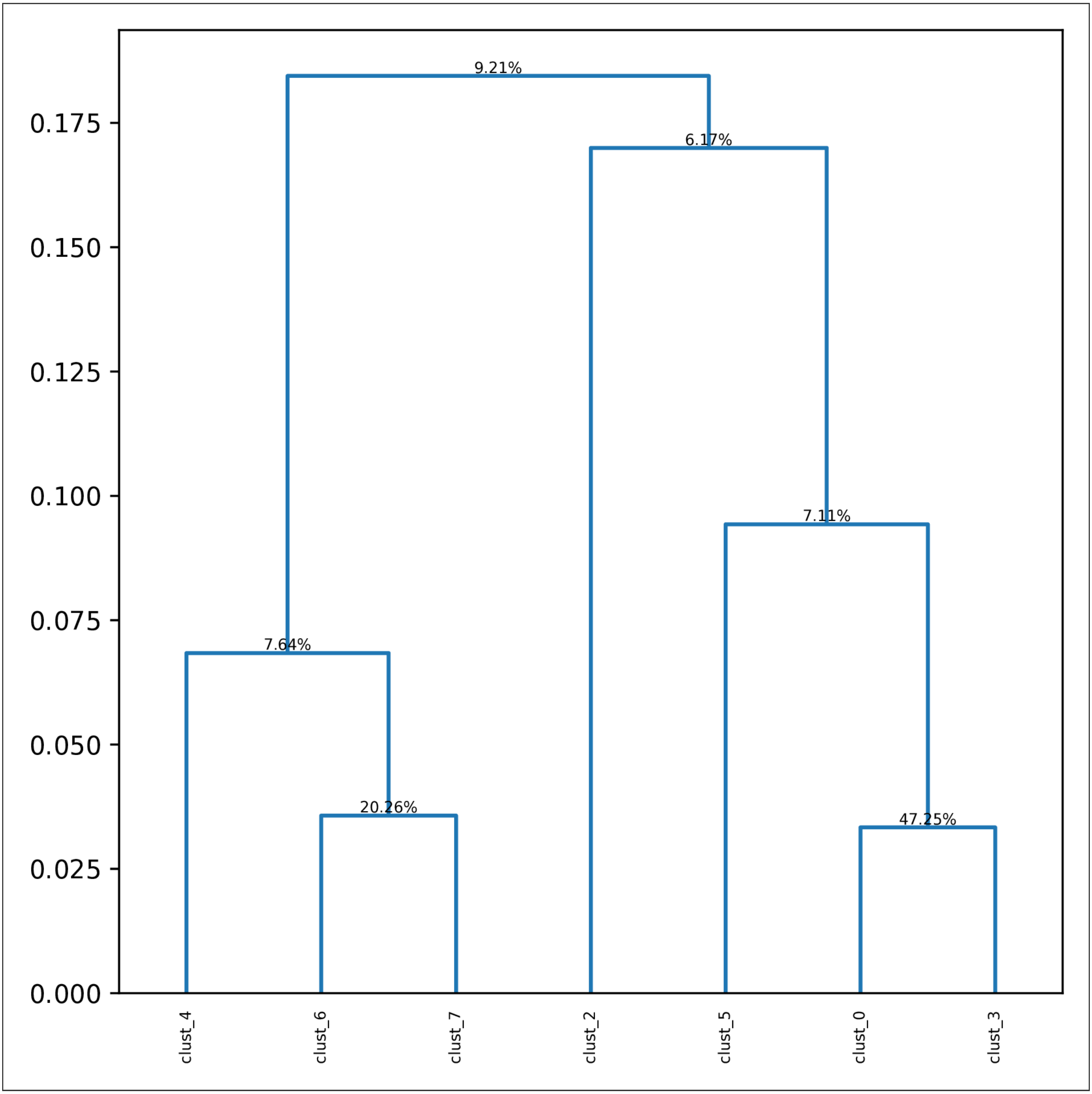

Sample: G0038 with all cells per cluster

Min. read depth: 10,  Mean UMIs per cluster:  7,843,866

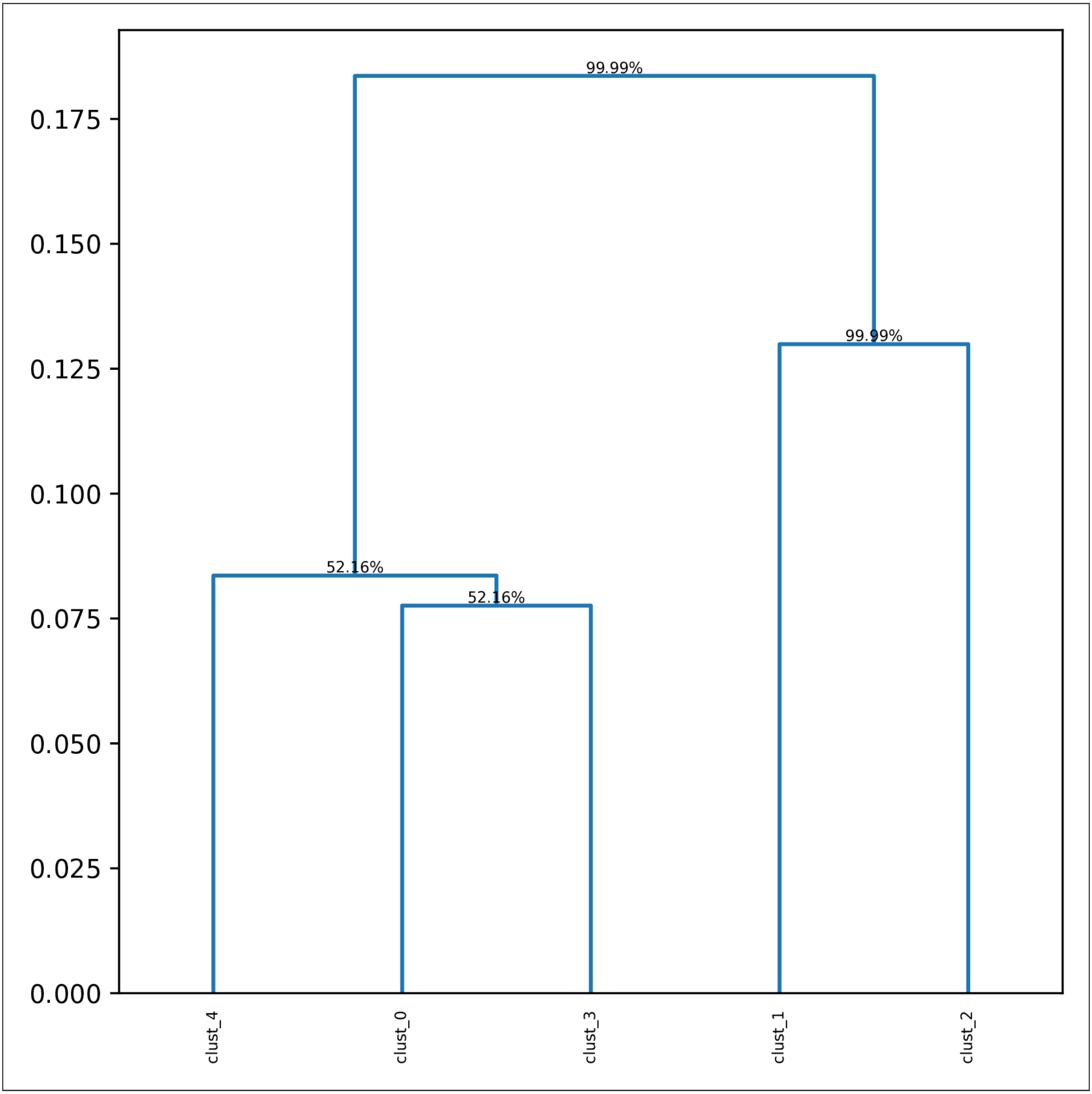

Sample: G0038 with 500 cells per cluster

Min. read depth: 10,  Mean UMIs per cluster:  6,128,216

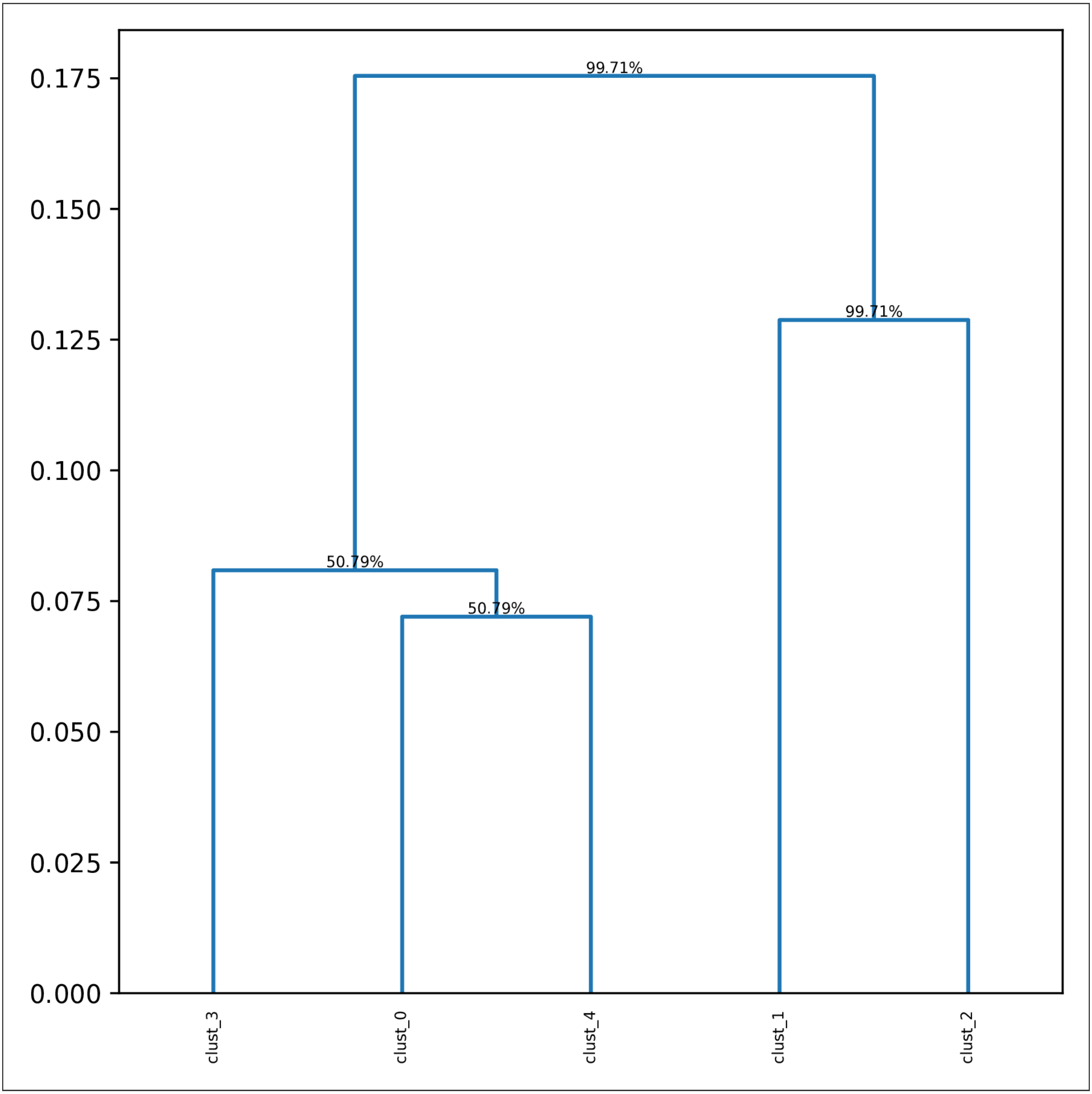

Sample: G0038 with 100 cells per cluster

Min. read depth: 10,  Mean UMIs per cluster:  1,221,695

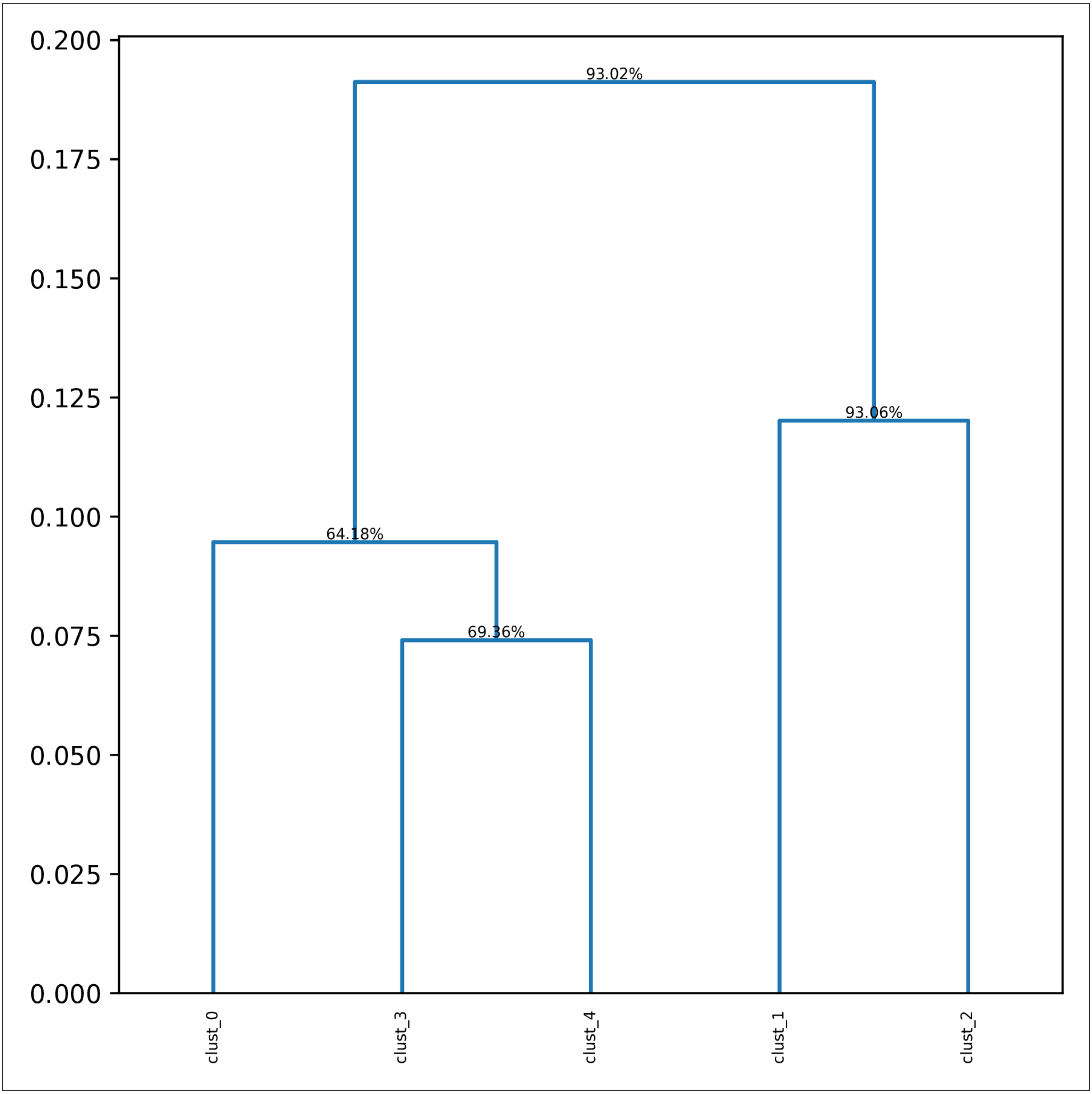

Sample: G0038 with 50 cells per cluster
Min. read depth: 10,  Mean UMIs per cluster:  698,403

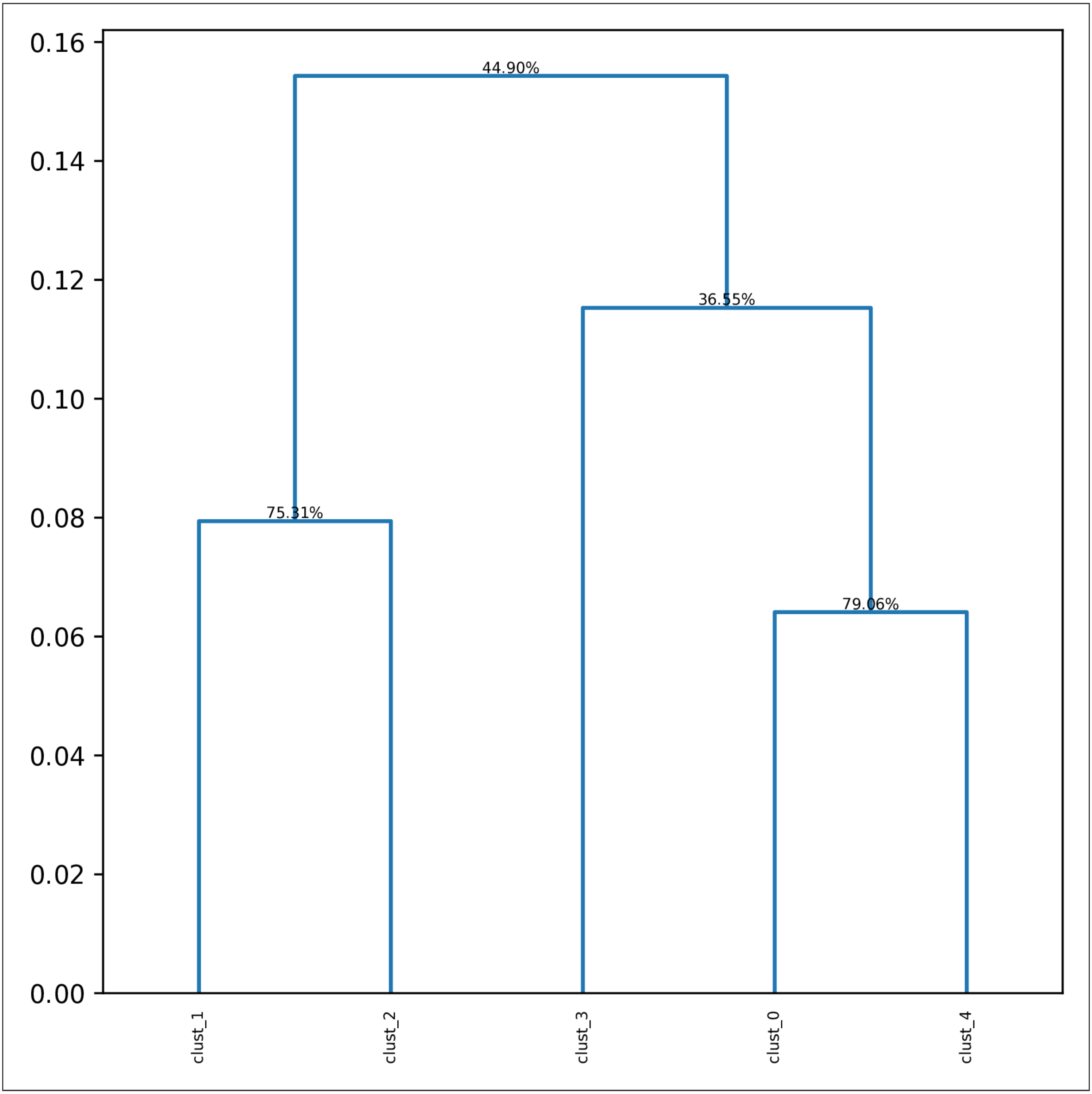

Sample: G0038 with 20 cells per cluster

Min. read depth: 10,  Mean UMIs per cluster:  251,736

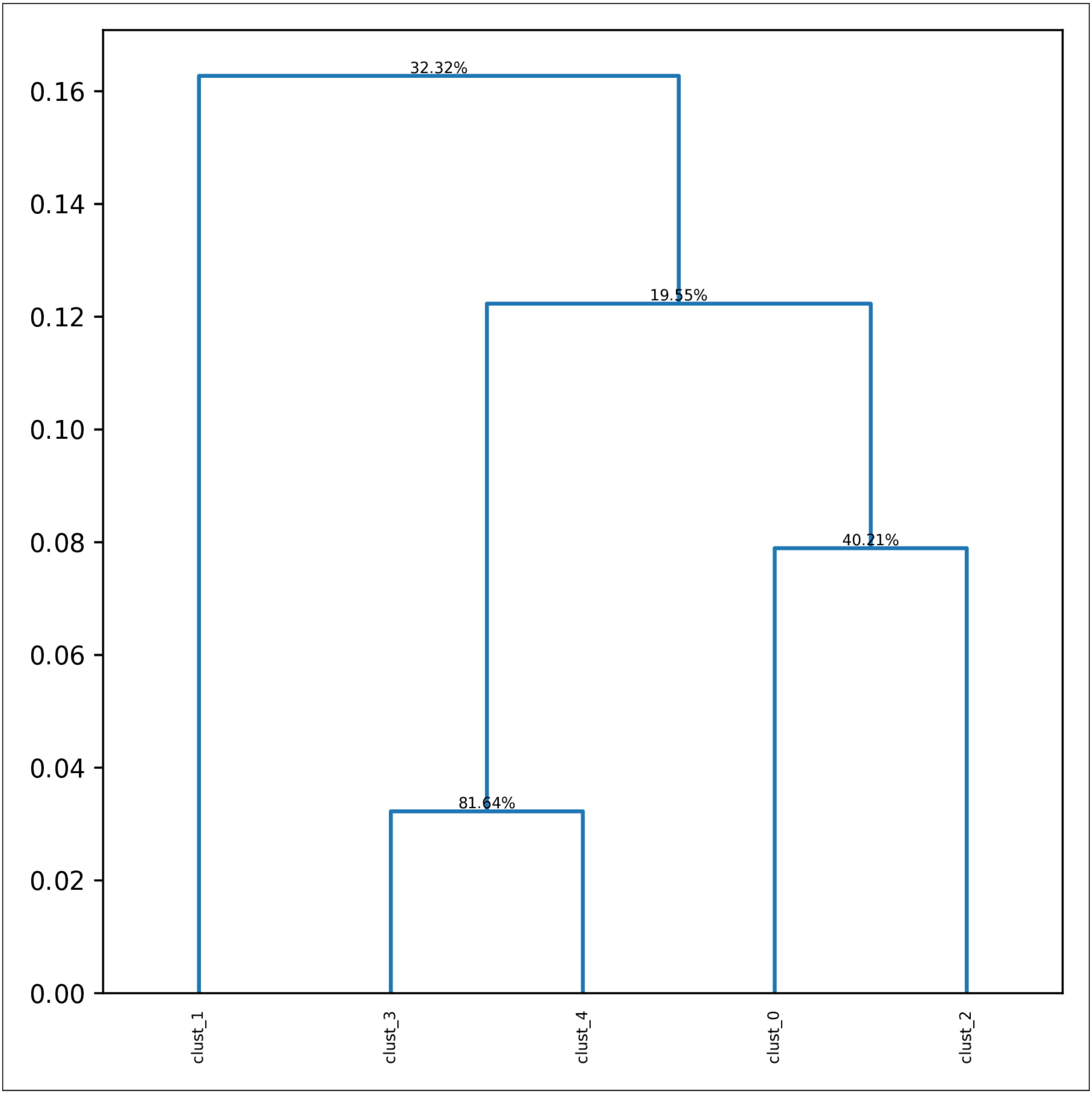

Sample: G0038 with 3 cells per cluster

Min. read depth: 10,  Mean UMIs per cluster:  35,795

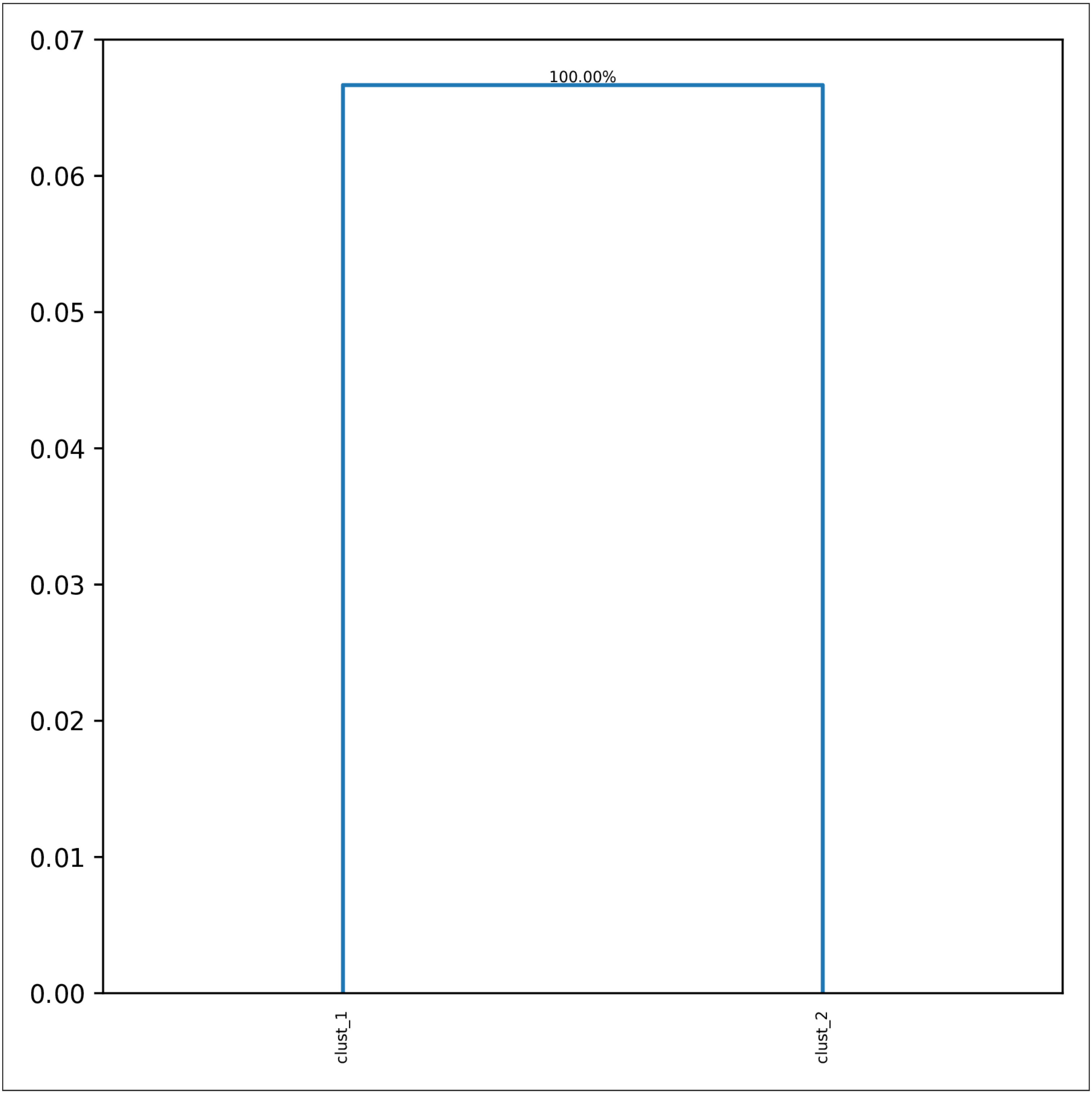

Sample: G0039 with all cells per cluster

Min. read depth: 10,  Mean UMIs per cluster:  6,534,792

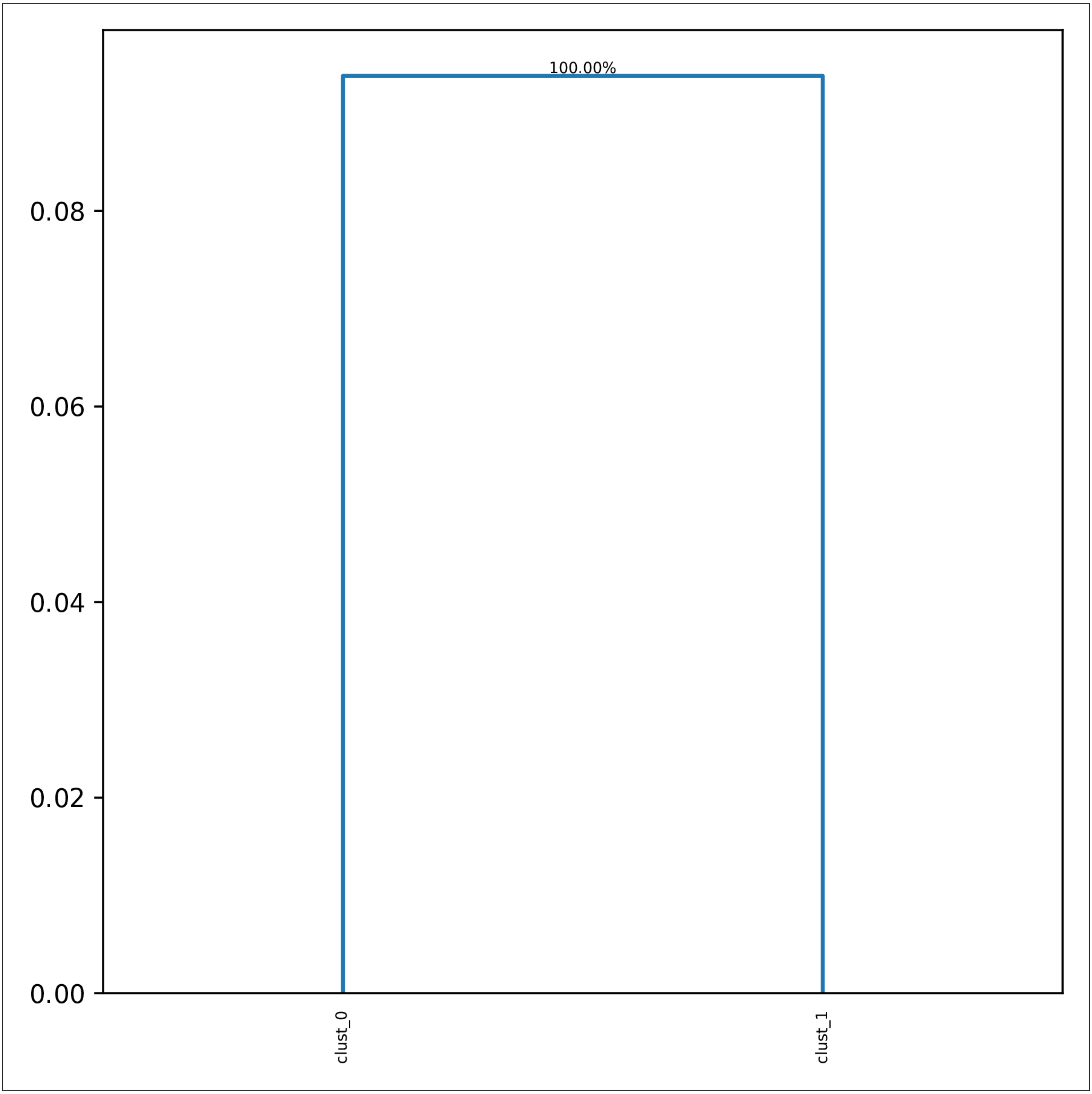

Sample: G0039 with 500 cells per cluster

Min. read depth: 10,  Mean UMIs per cluster:  2,850,004

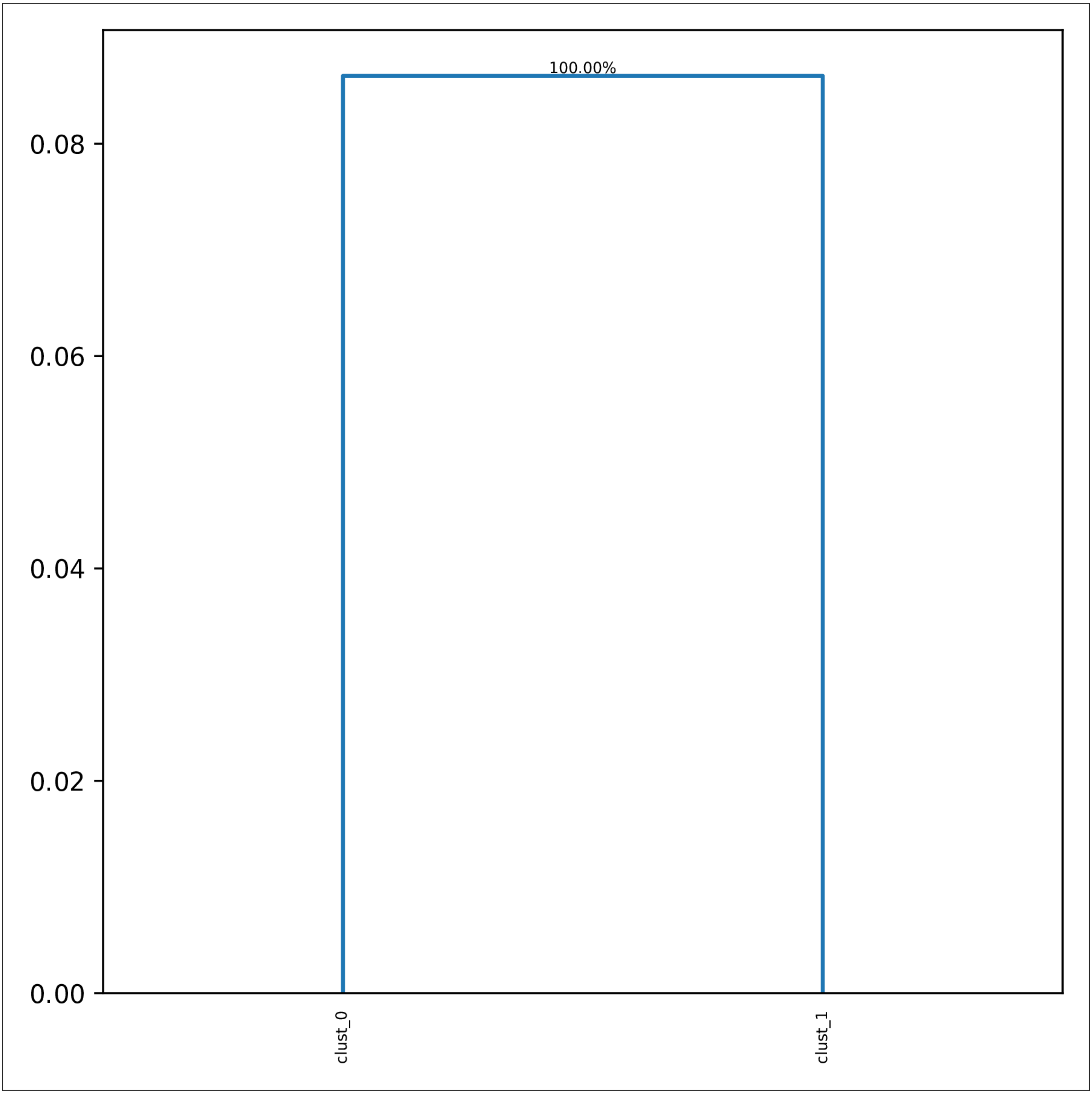

Sample: G0039 with 100 cells per cluster

Min. read depth: 10,  Mean UMIs per cluster:  618,683

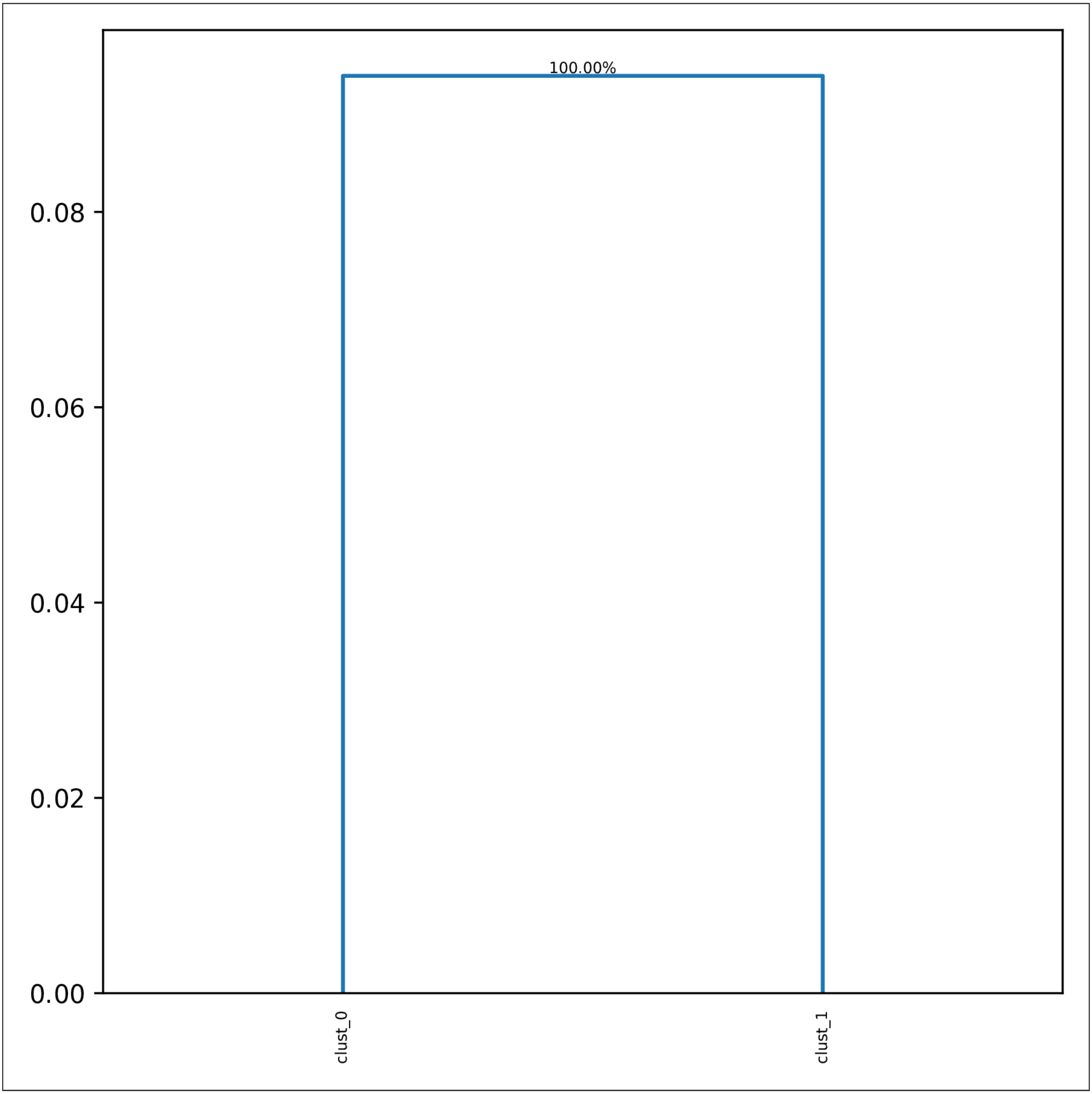

Sample: G0039 with 50 cells per cluster
Min. read depth: 10,  Mean UMIs per cluster:  288,386

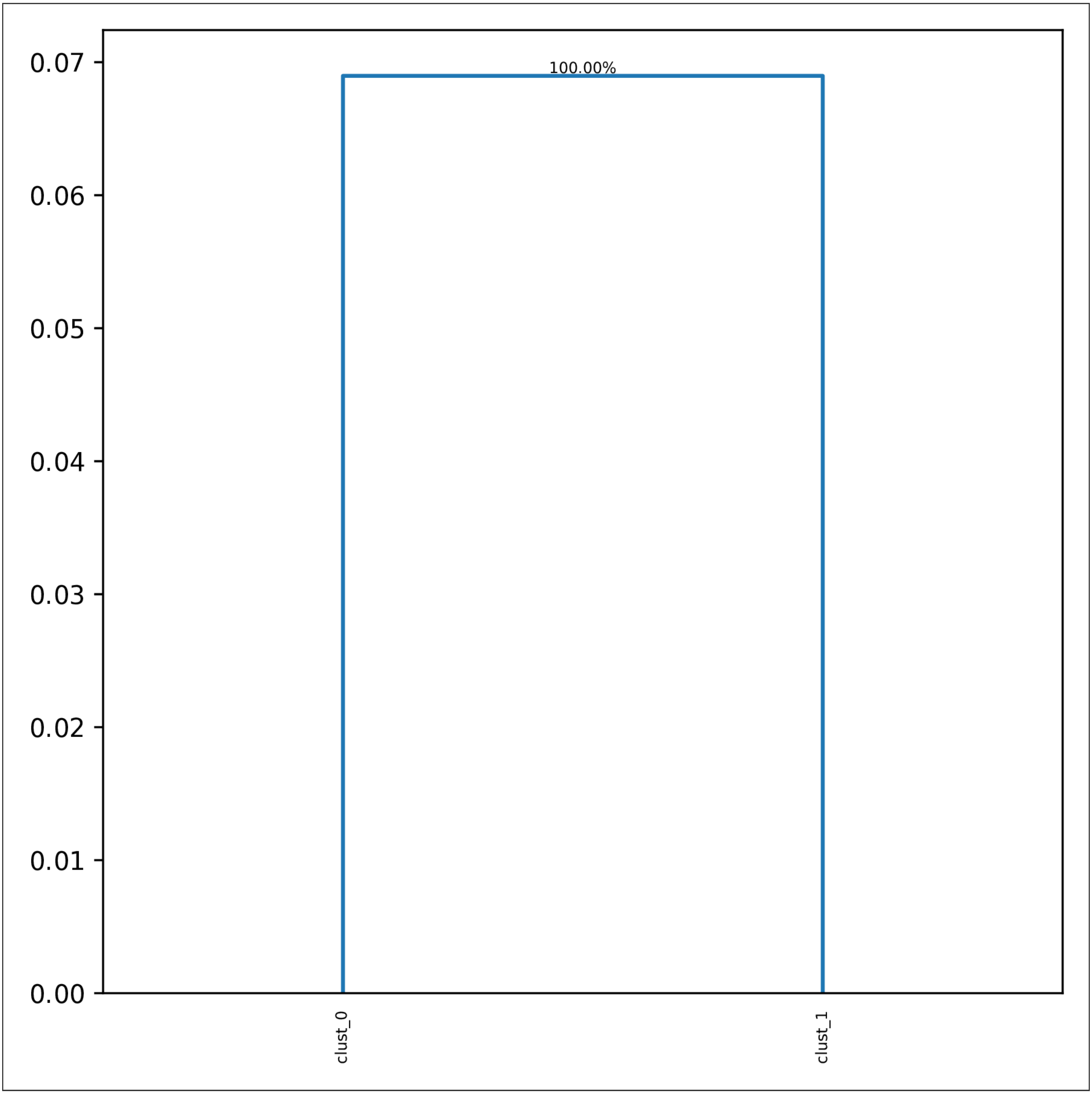

Sample: G0039 with 20 cells per cluster

Min. read depth: 10,  Mean UMIs per cluster:  128,028

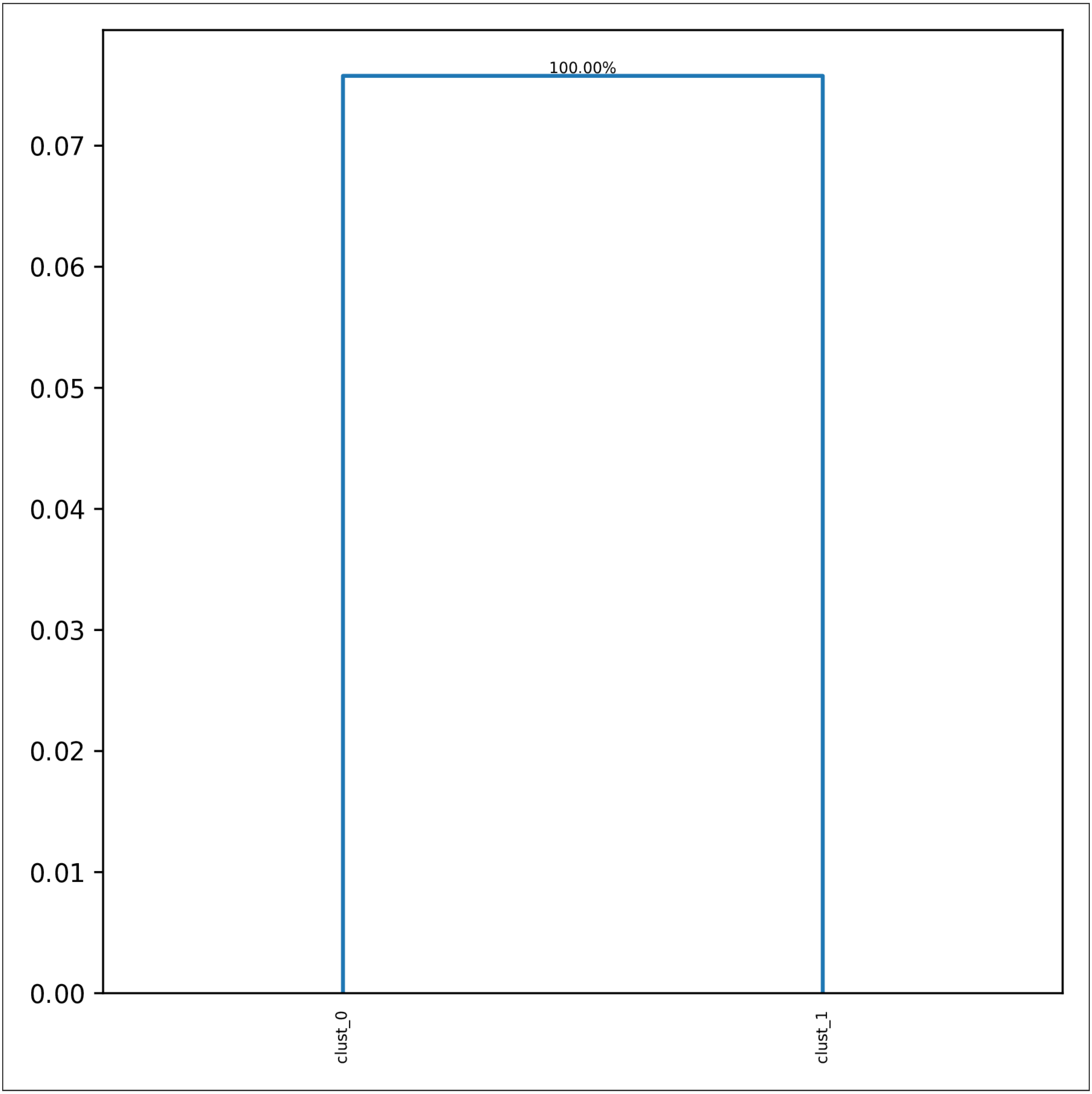

Sample: G0039 with 10 cells per cluster

Min. read depth: 10, Mean UMIs per cluster: 65,074

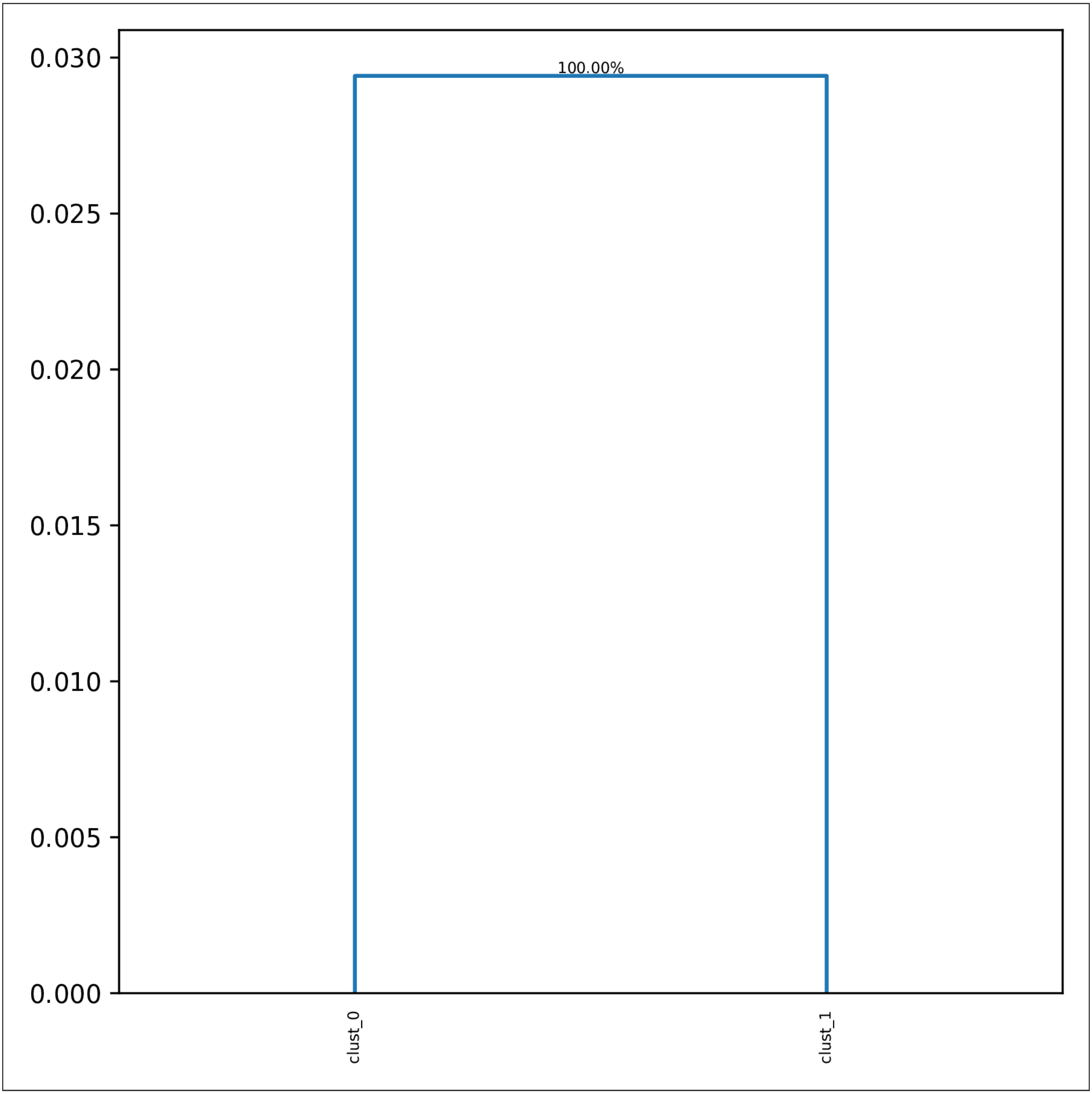

Sample: G0040 with all cells per cluster

Min. read depth: 10,  Mean UMIs per cluster:  15,794,763

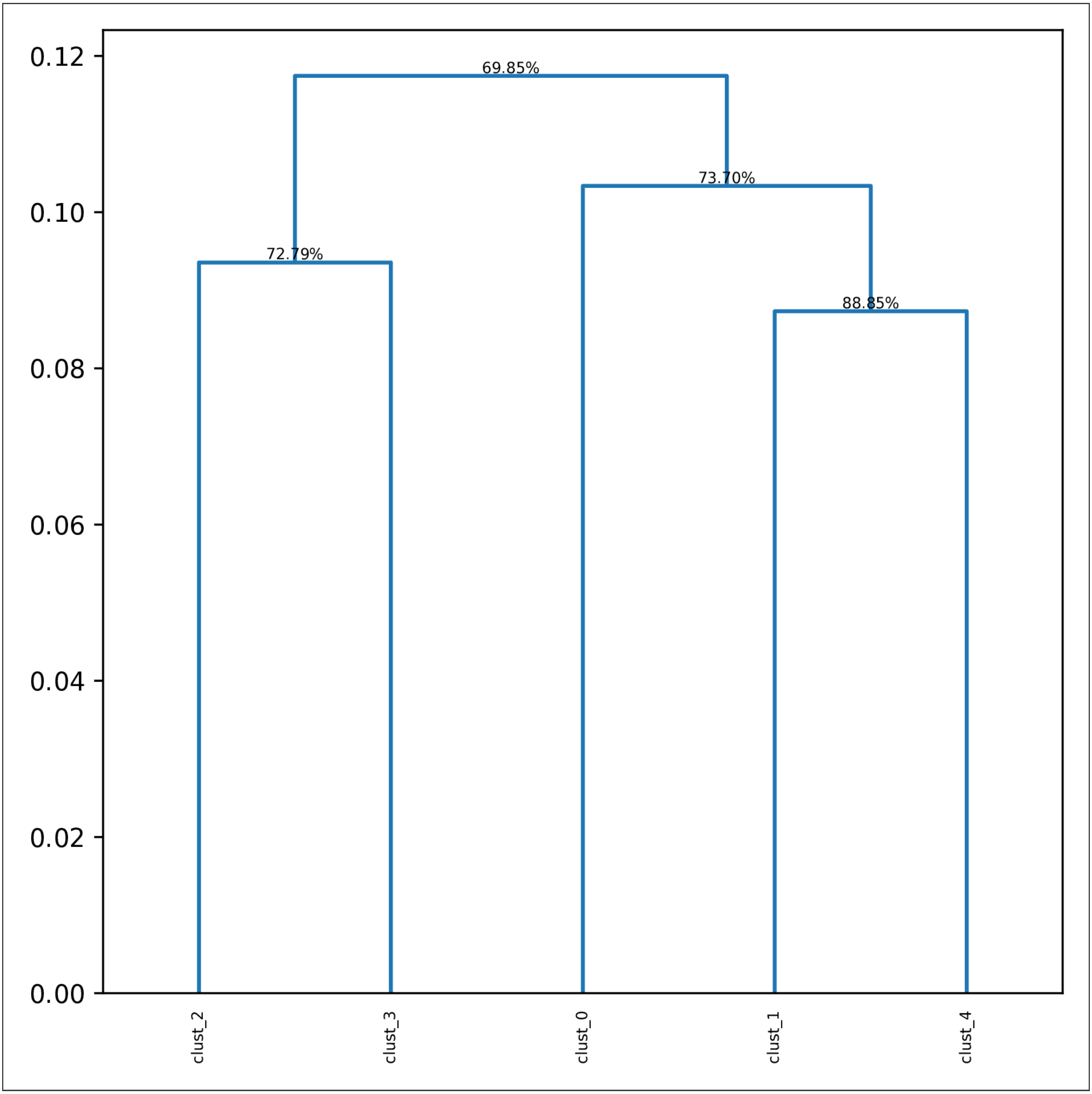

Sample: G0040 with 500 cells per cluster

Min. read depth: 10,  Mean UMIs per cluster:  4,623,819

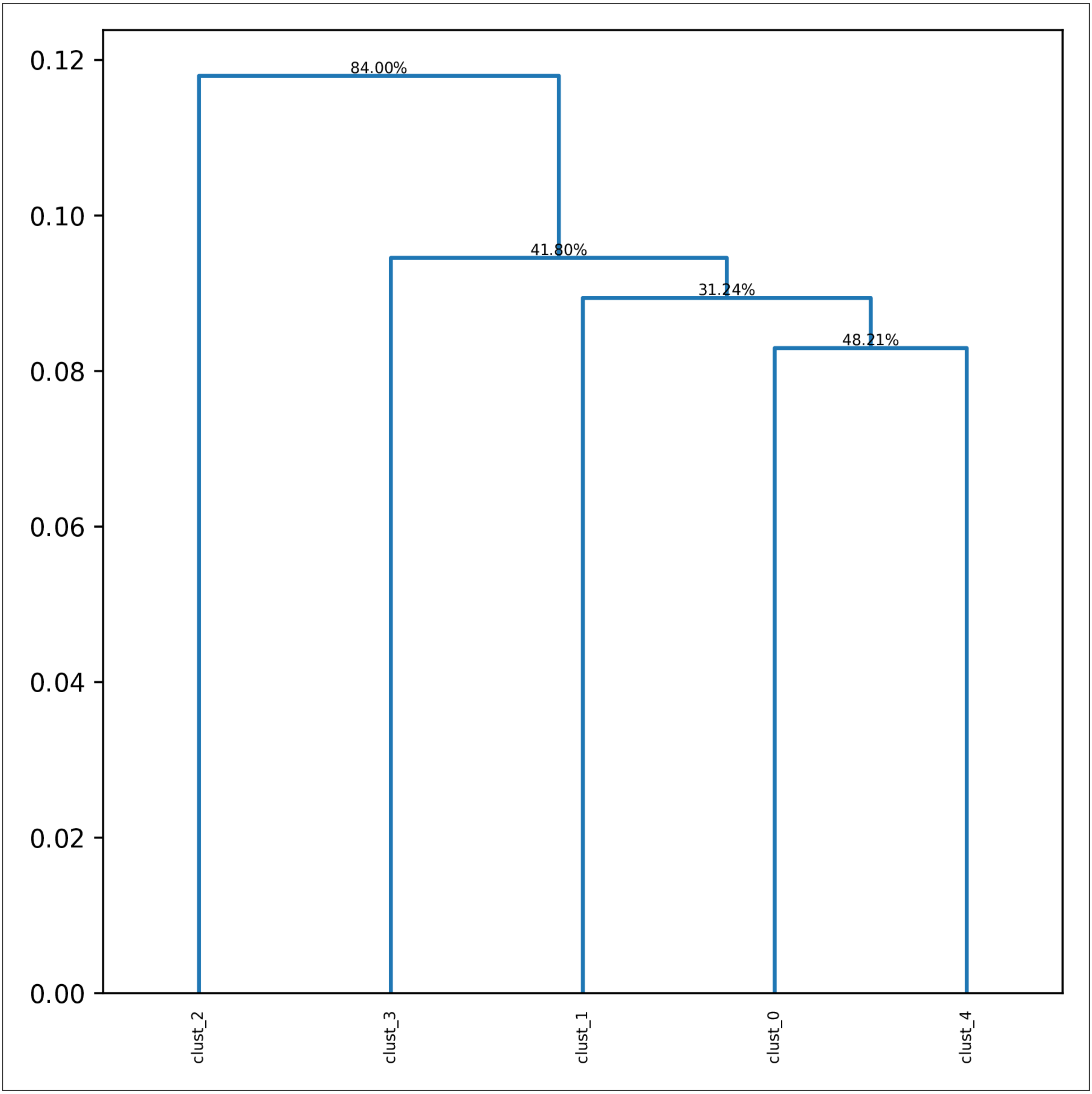

Sample: G0040 with 100 cells per cluster
Min. read depth: 10,  Mean UMIs per cluster:  936,766

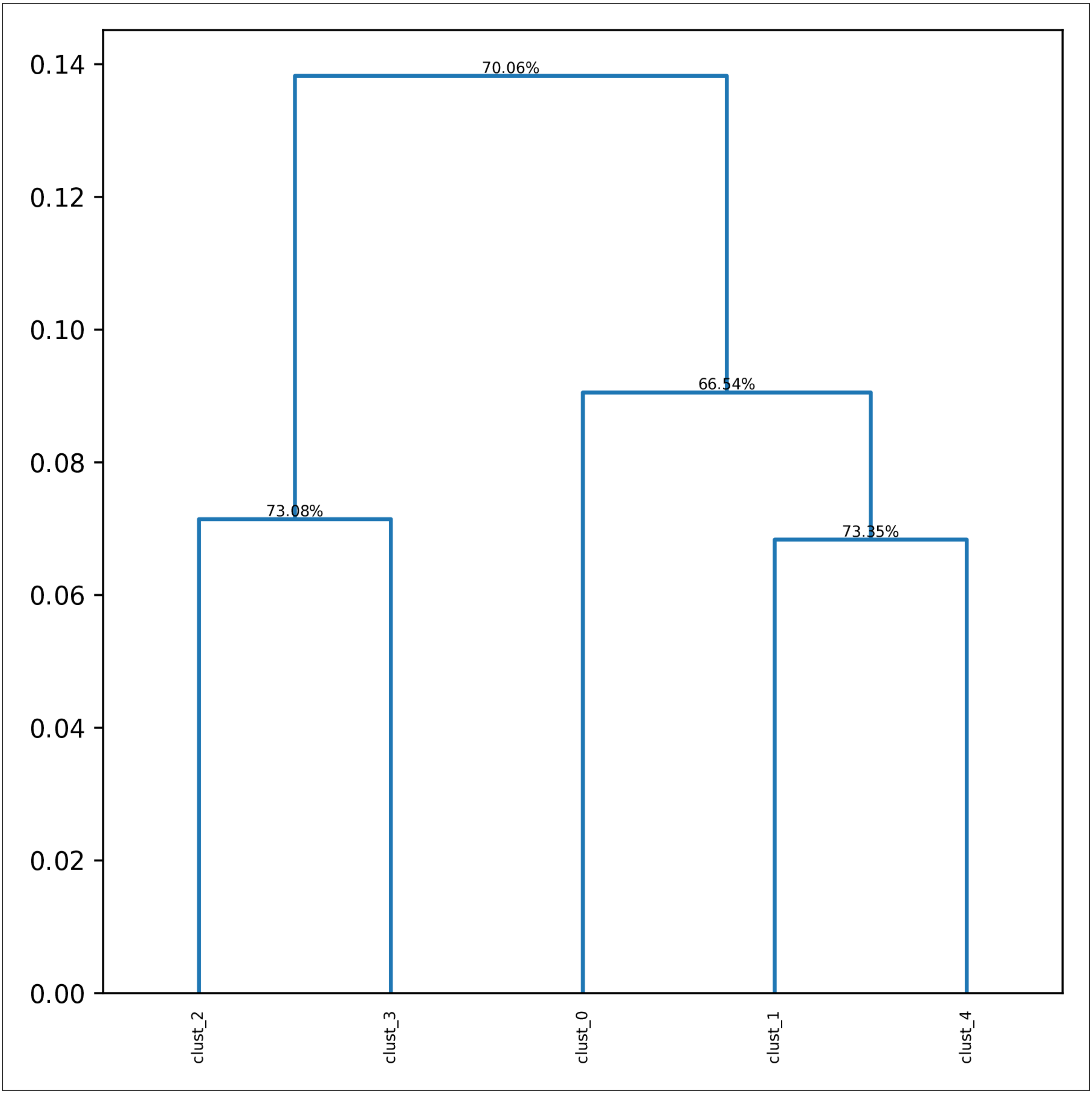

Sample: G0040 with 50 cells per cluster

Min. read depth: 10,  Mean UMIs per cluster:  494,490

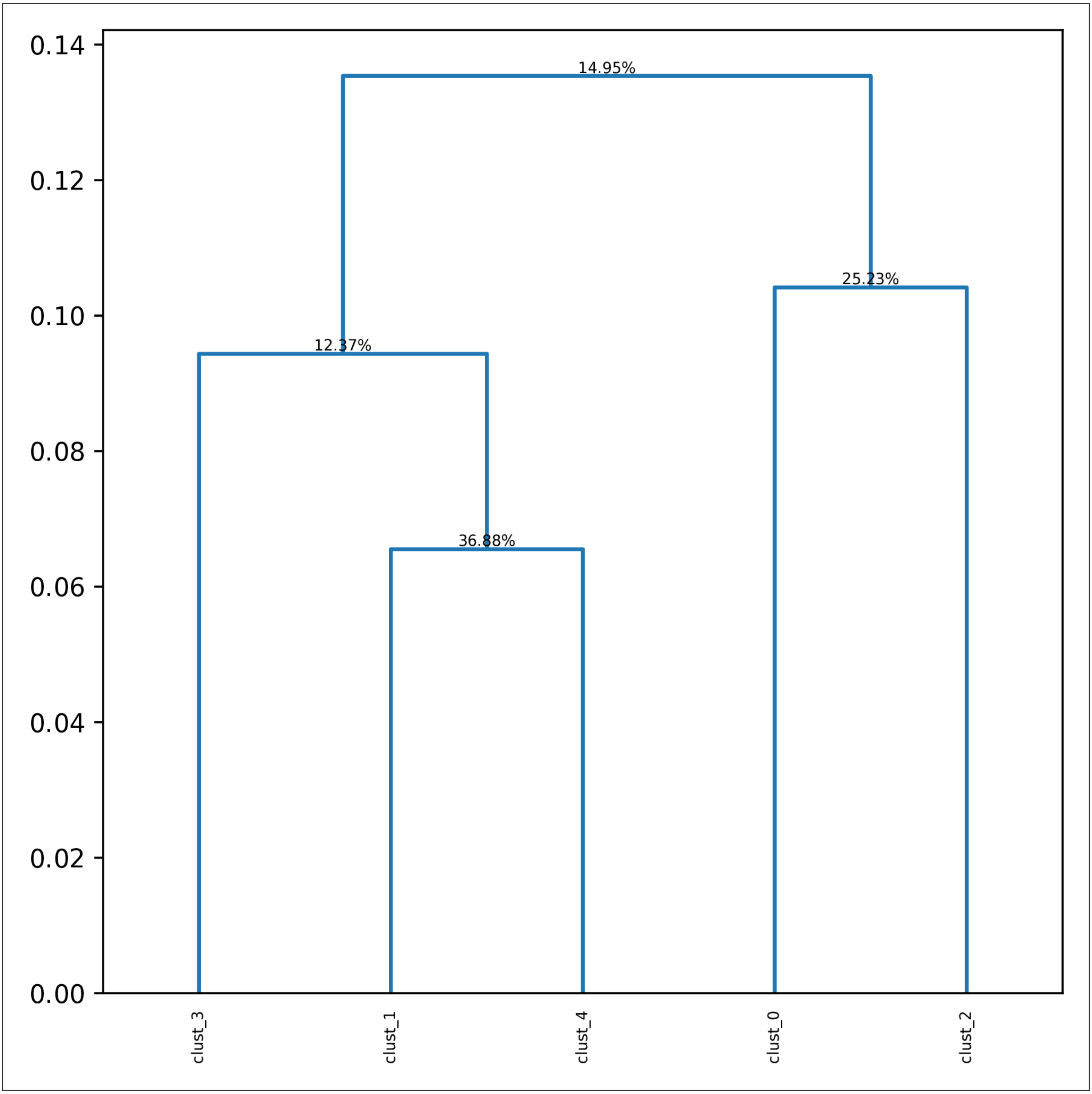

Sample: G0040 with 20 cells per cluster
Min. read depth: 10,  Mean UMIs per cluster:  172,714

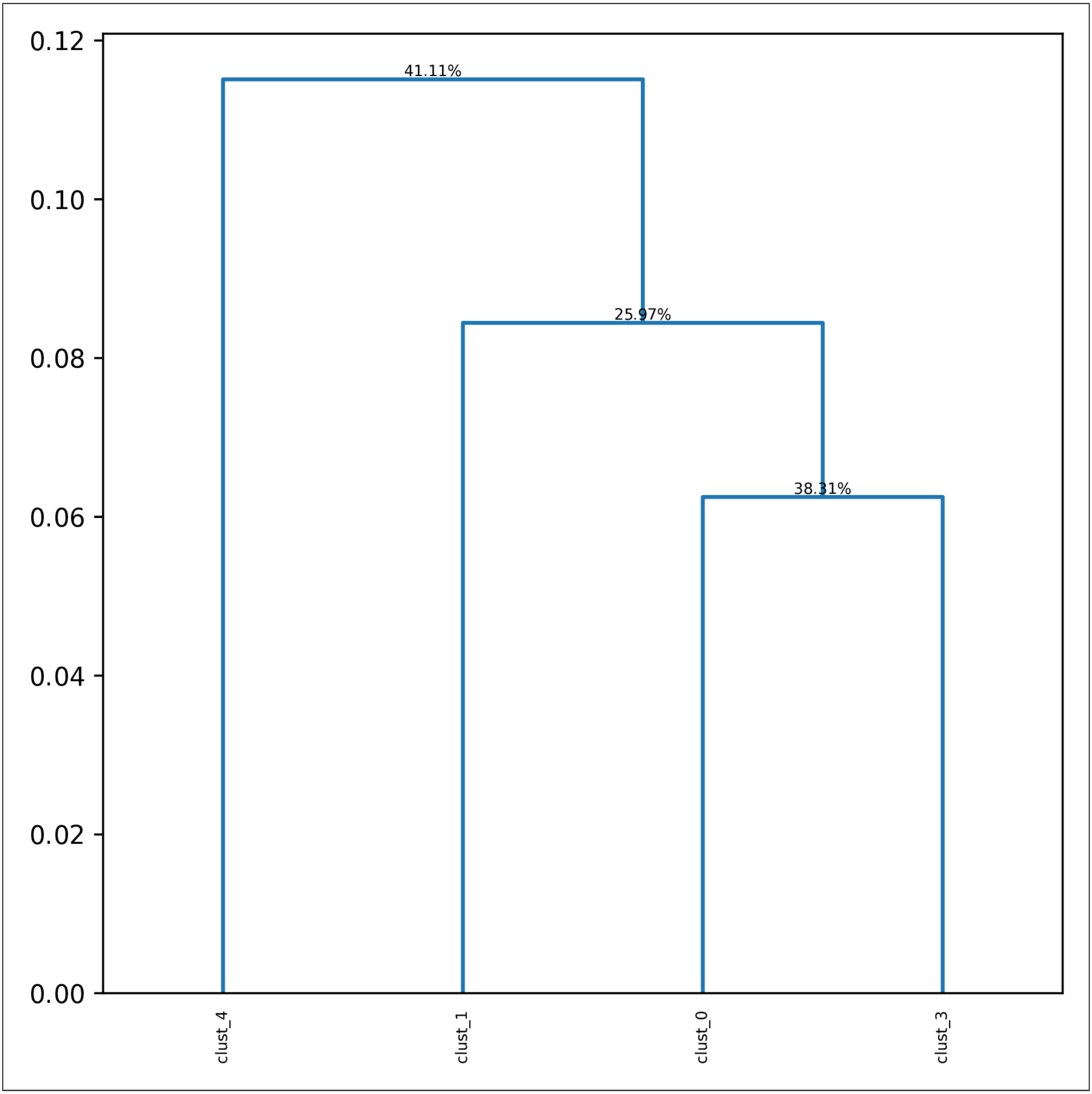

Sample: G0040 with 10 cells per cluster

Min. read depth: 10,  Mean UMIs per cluster:  72,643

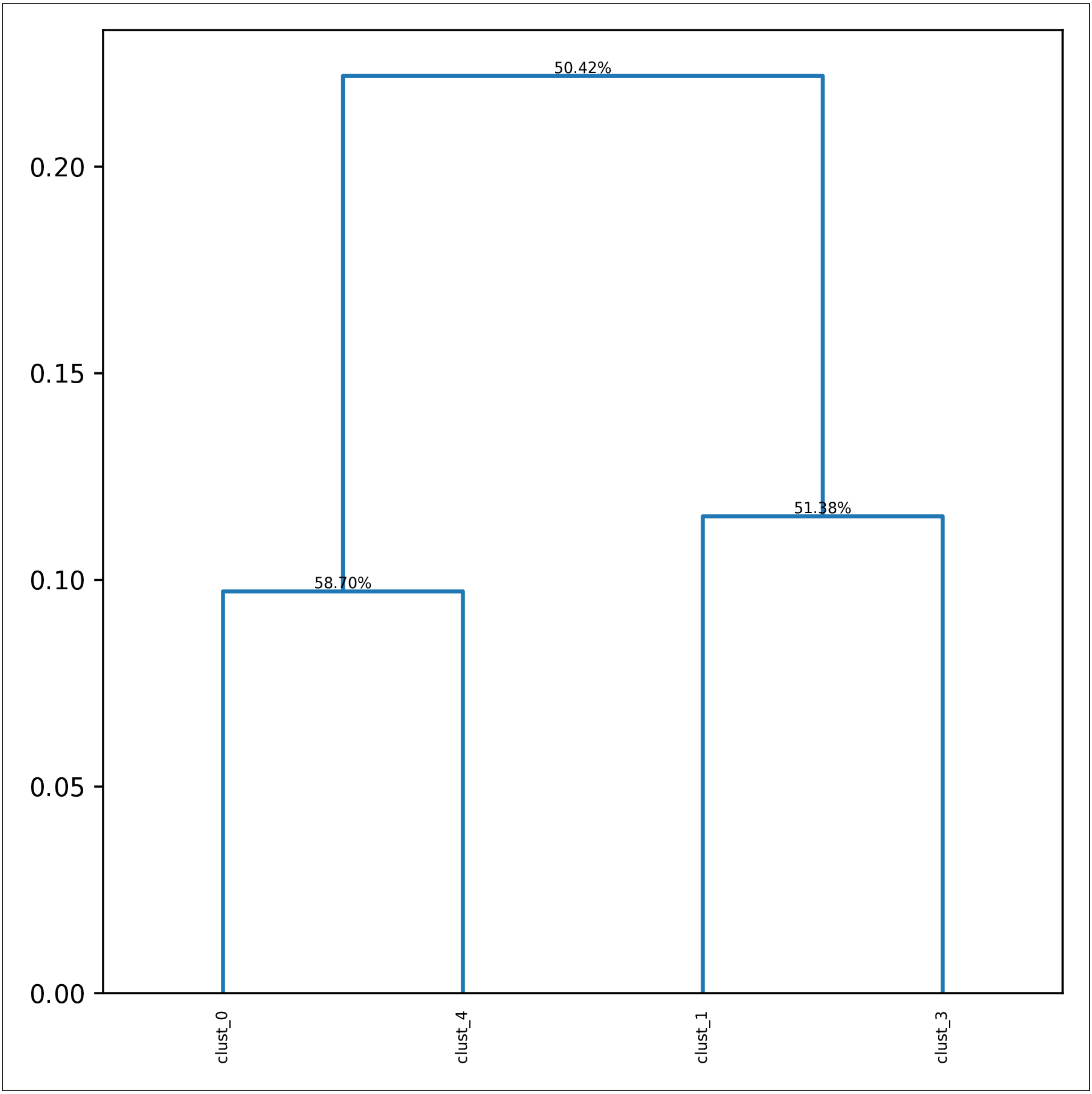

Sample: G0040 with 5 cells per cluster

Min. read depth: 10,  Mean UMIs per cluster:  45,992

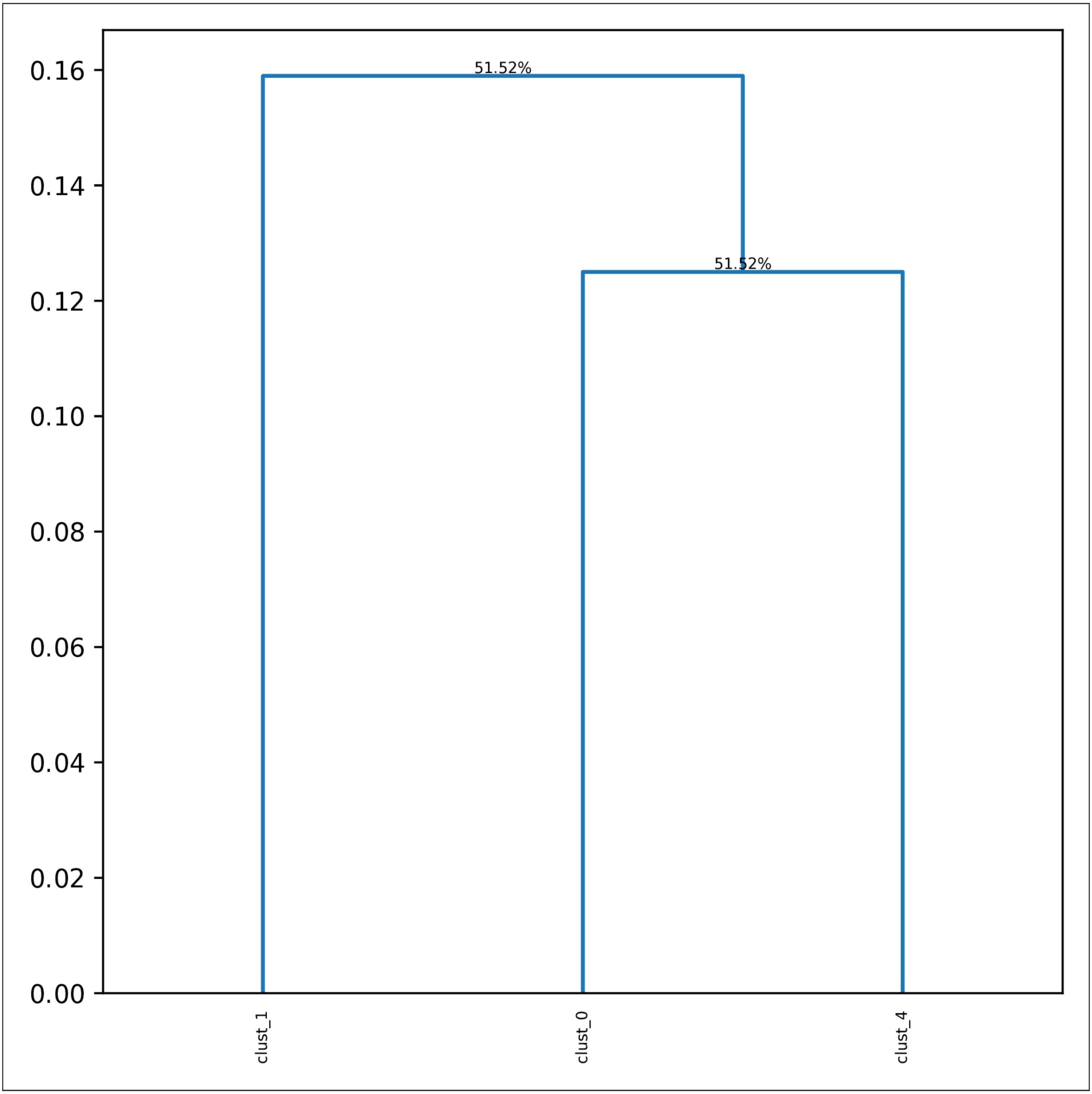

Sample: G0041 with all cells per cluster

Min. read depth: 10,  Mean UMIs per cluster:  4,723,499

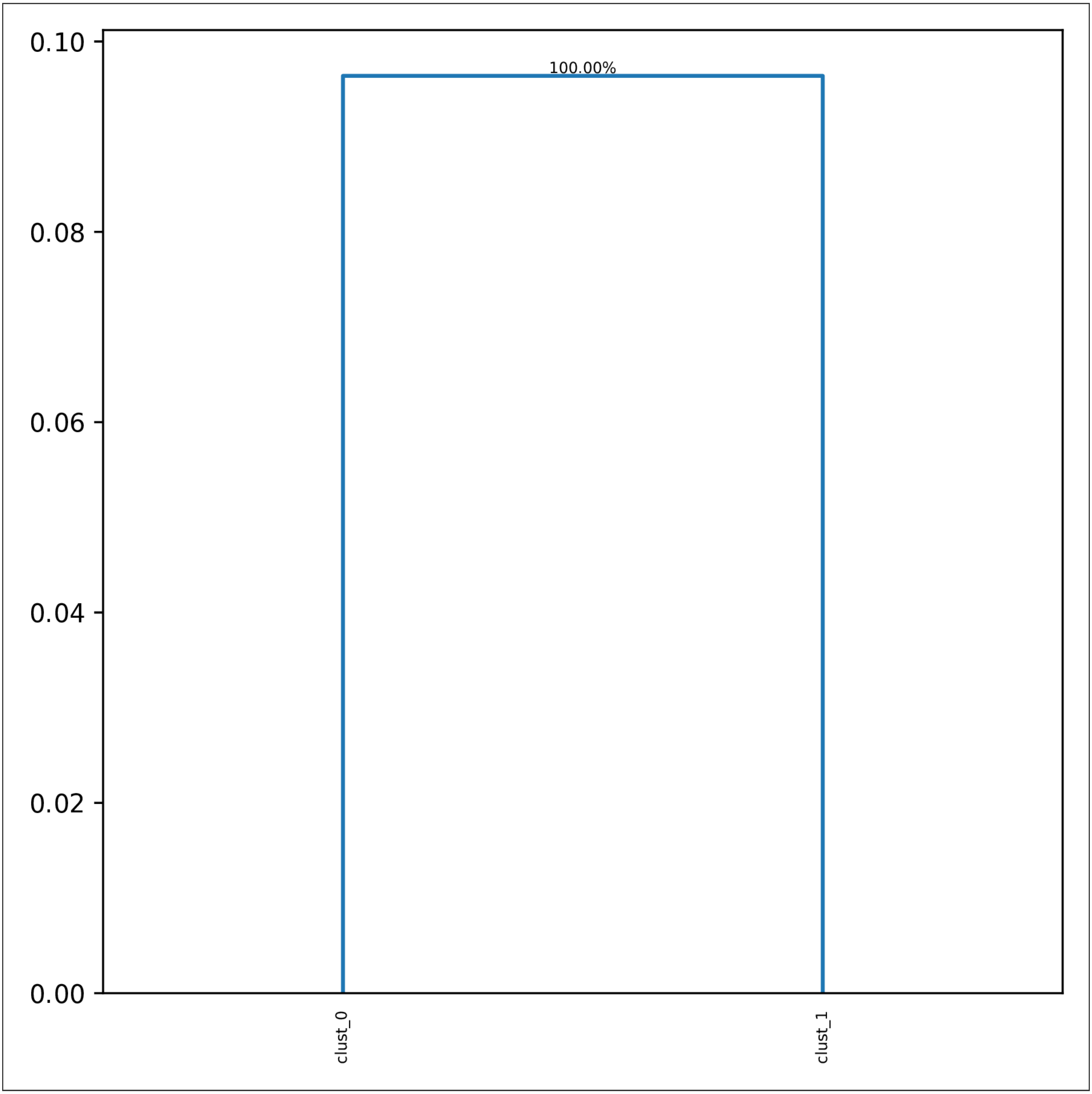

Sample: G0041 with 500 cells per cluster

Min. read depth: 10,  Mean UMIs per cluster:  4,121,158

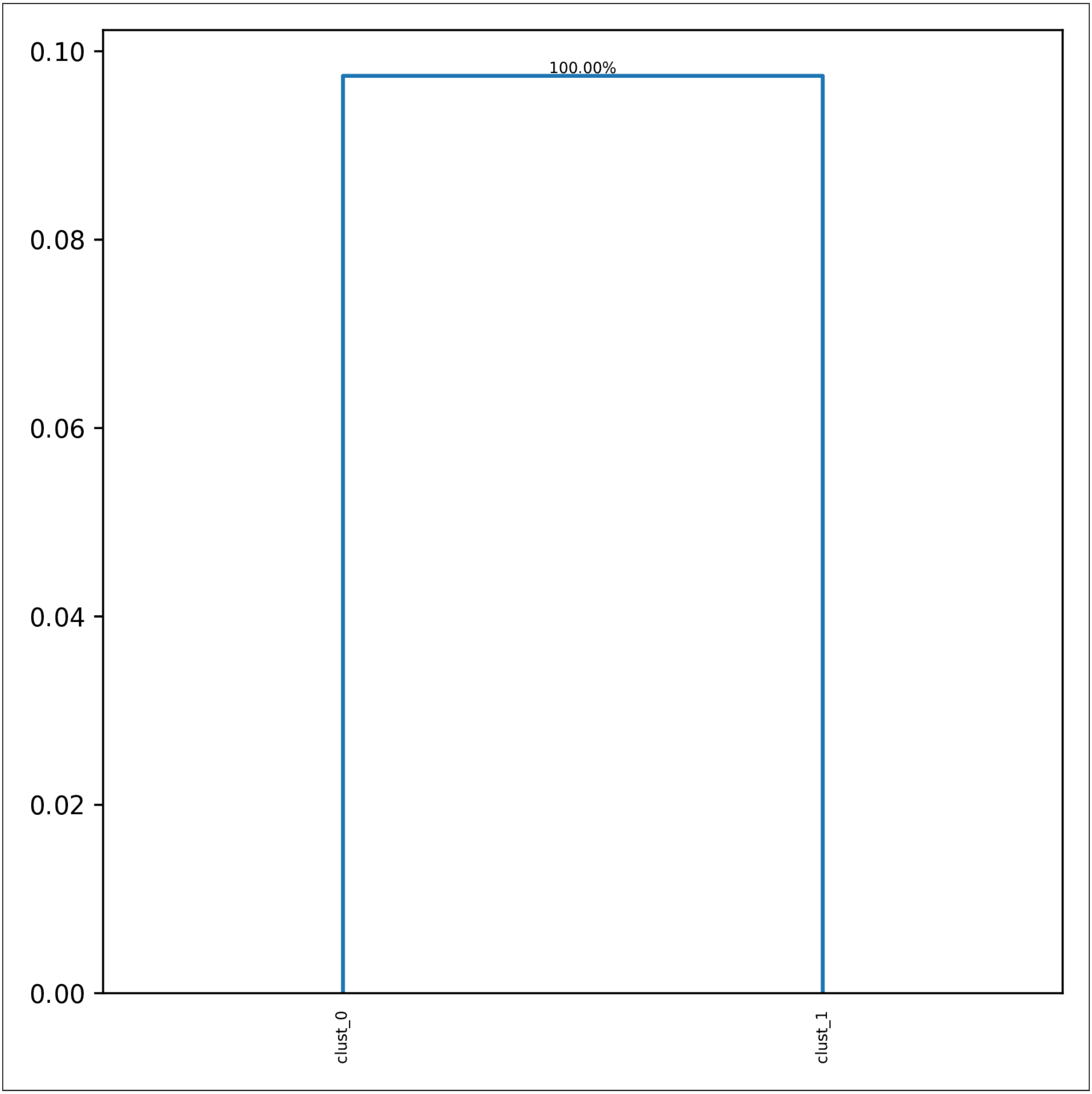

Sample: G0041 with 100 cells per cluster

Min. read depth: 10, Mean UMIs per cluster: 1,042,602

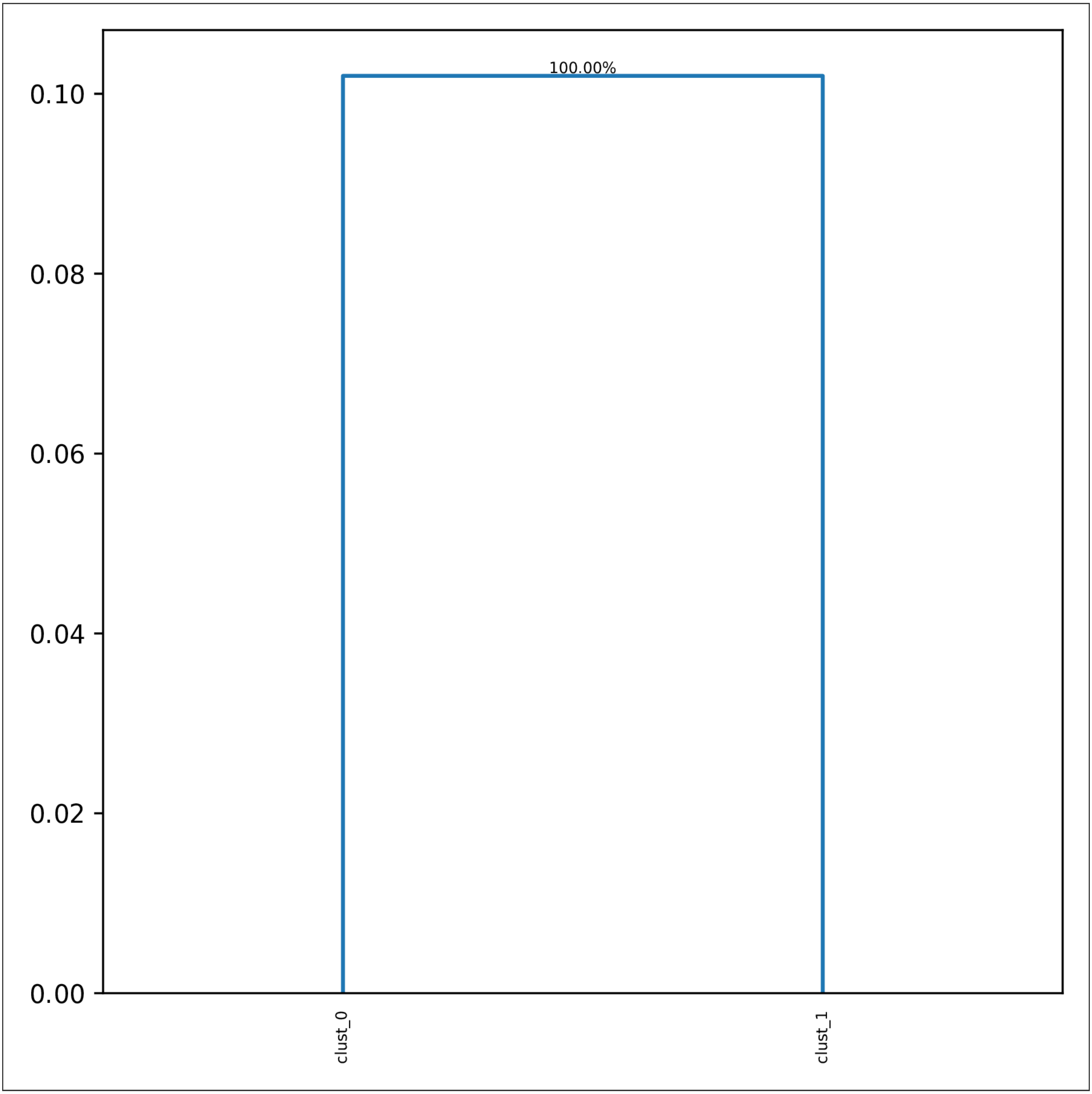

Sample: G0041 with 50 cells per cluster

Min. read depth: 10,  Mean UMIs per cluster:  570,572

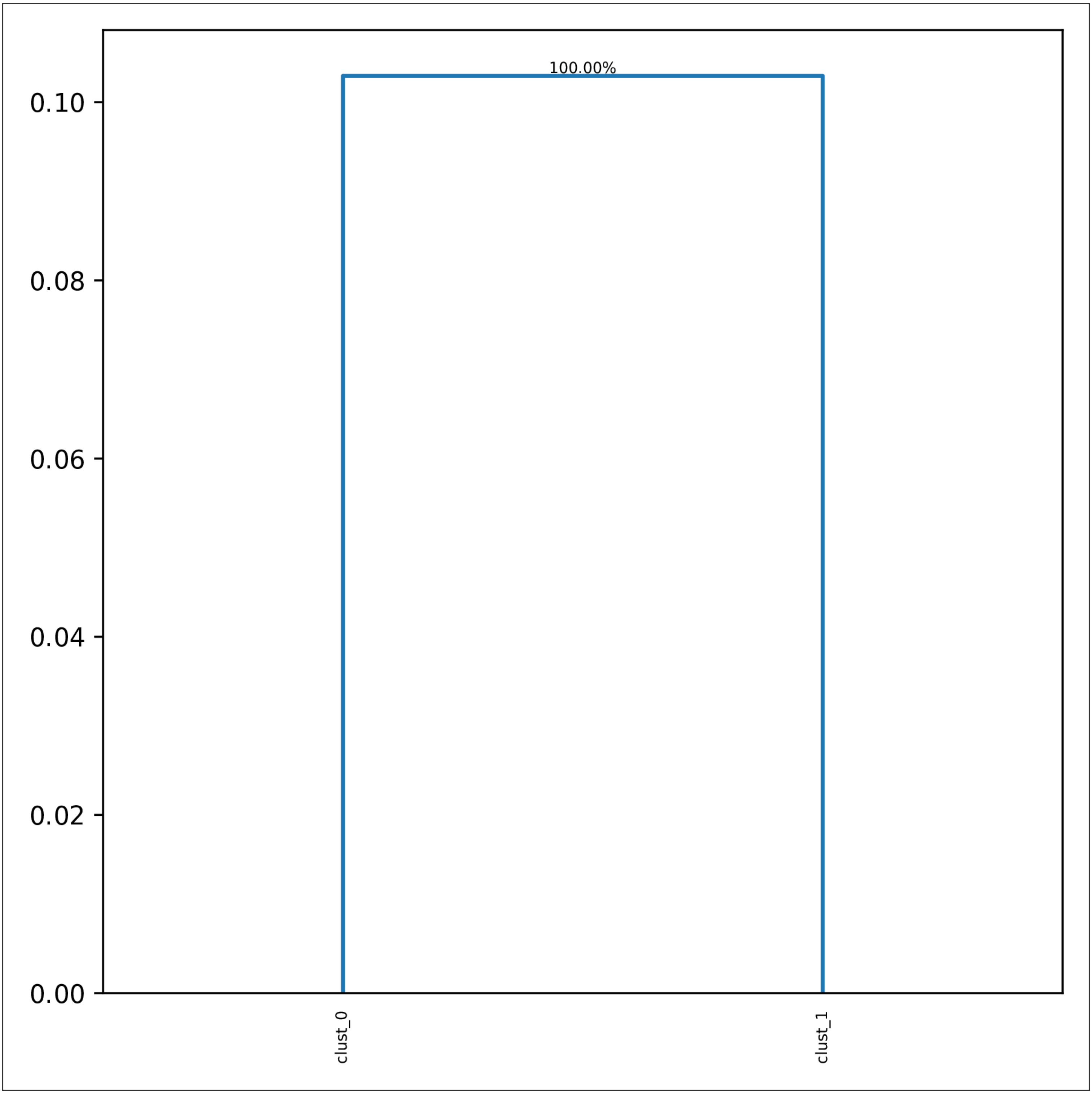

Sample: G0041 with 20 cells per cluster

Min. read depth: 10,  Mean UMIs per cluster:  243,348

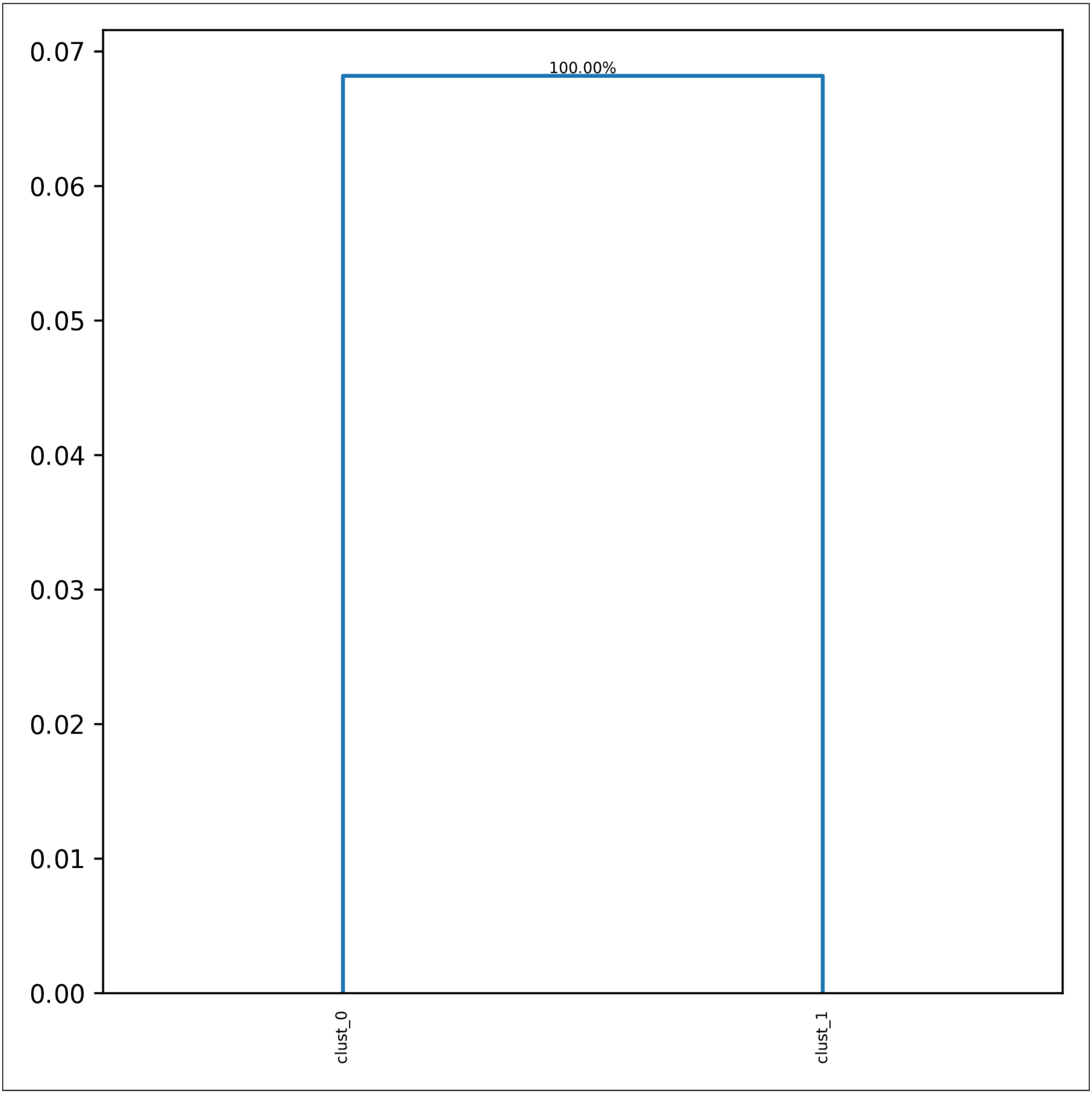

Sample: G0041 with 10 cells per cluster

Min. read depth: 10,  Mean UMIs per cluster:  104,608

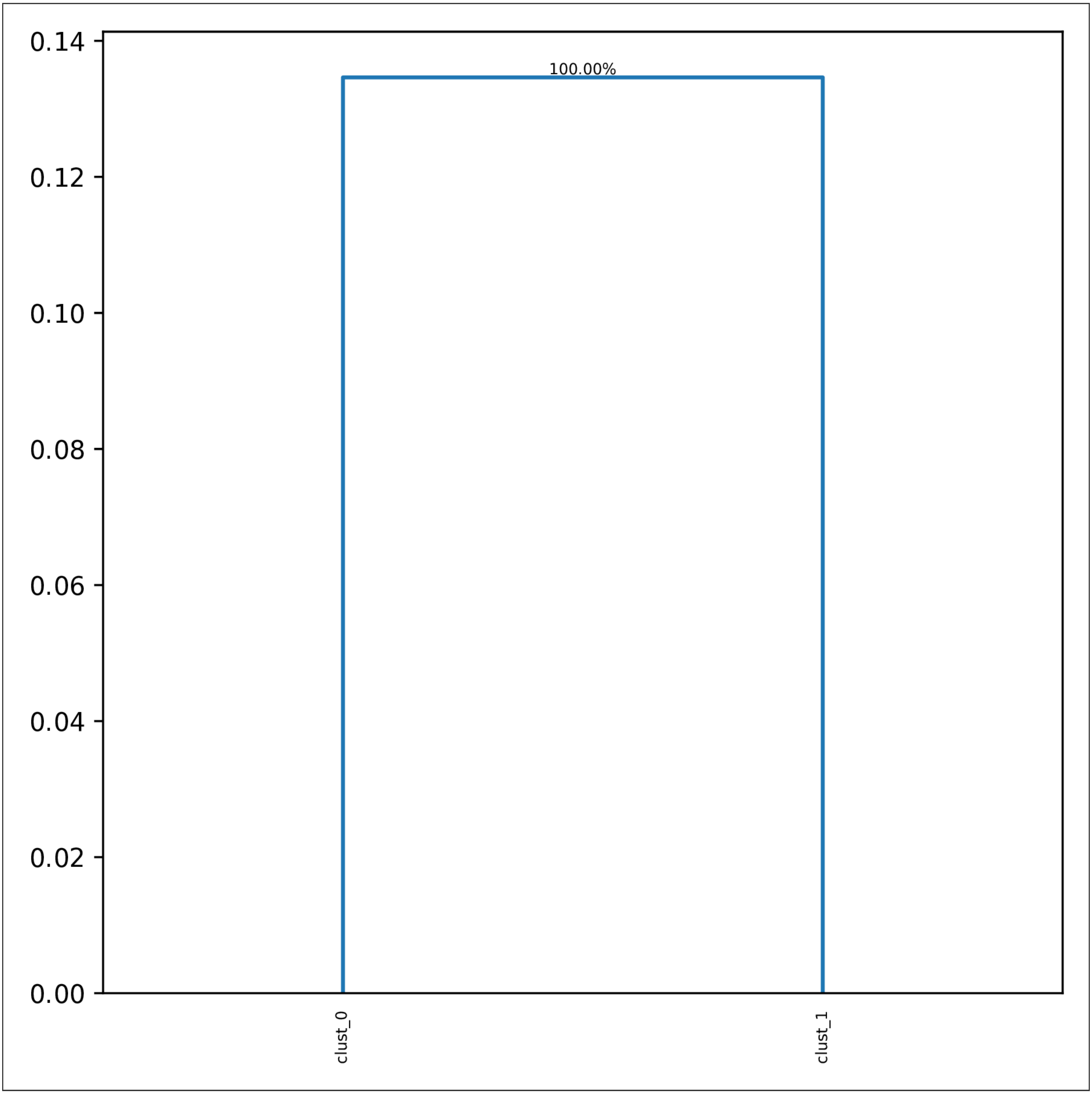

Sample: G0041 with 5 cells per cluster

Min. read depth: 10,  Mean UMIs per cluster:  63,252

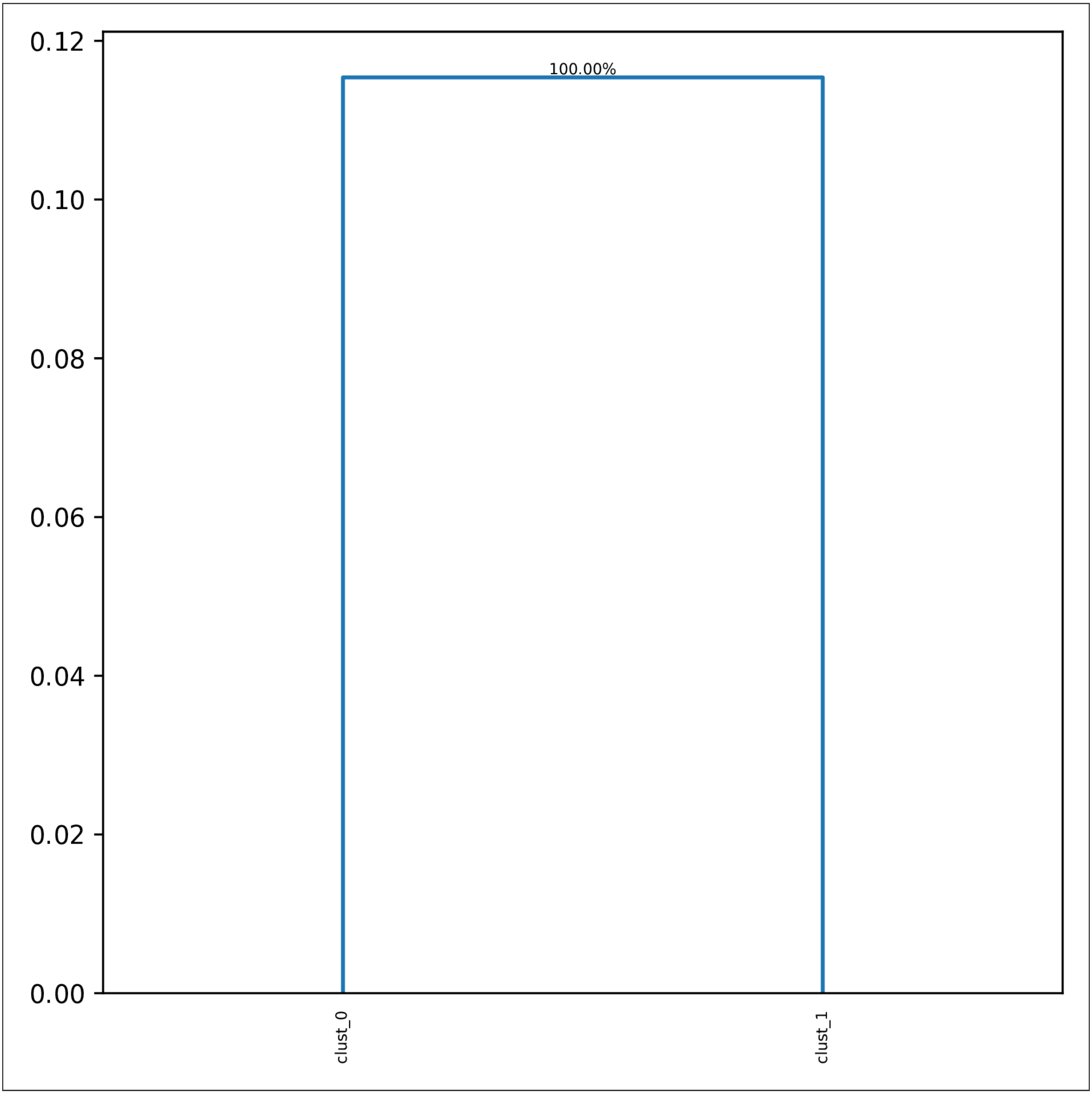

Sample: G0041 with 4 cells per cluster

Min. read depth: 10,  Mean UMIs per cluster:  53,708

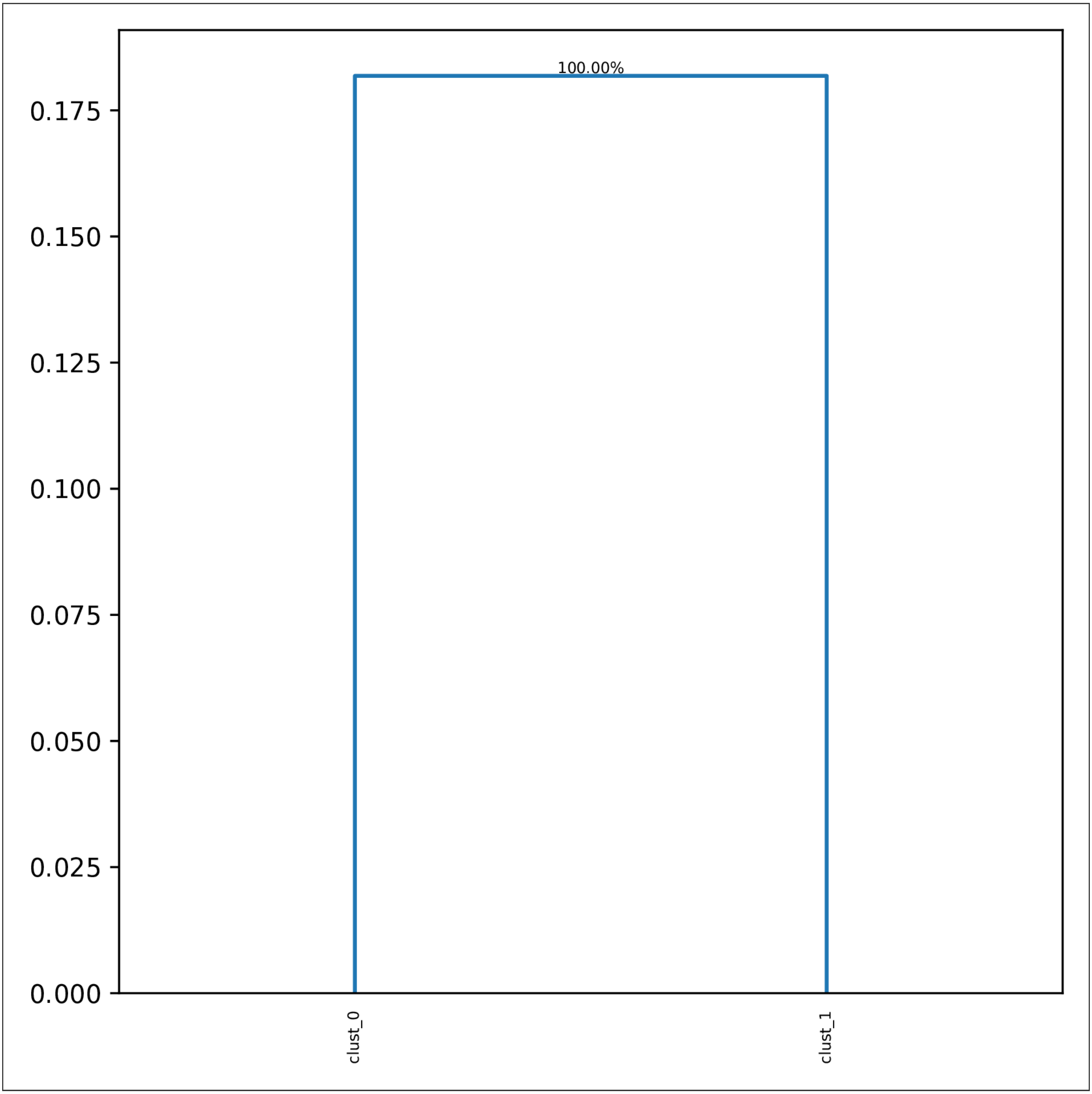

Sample: G0042 with all cells per cluster

Min. read depth: 10, Mean UMIs per cluster: 25,424,346

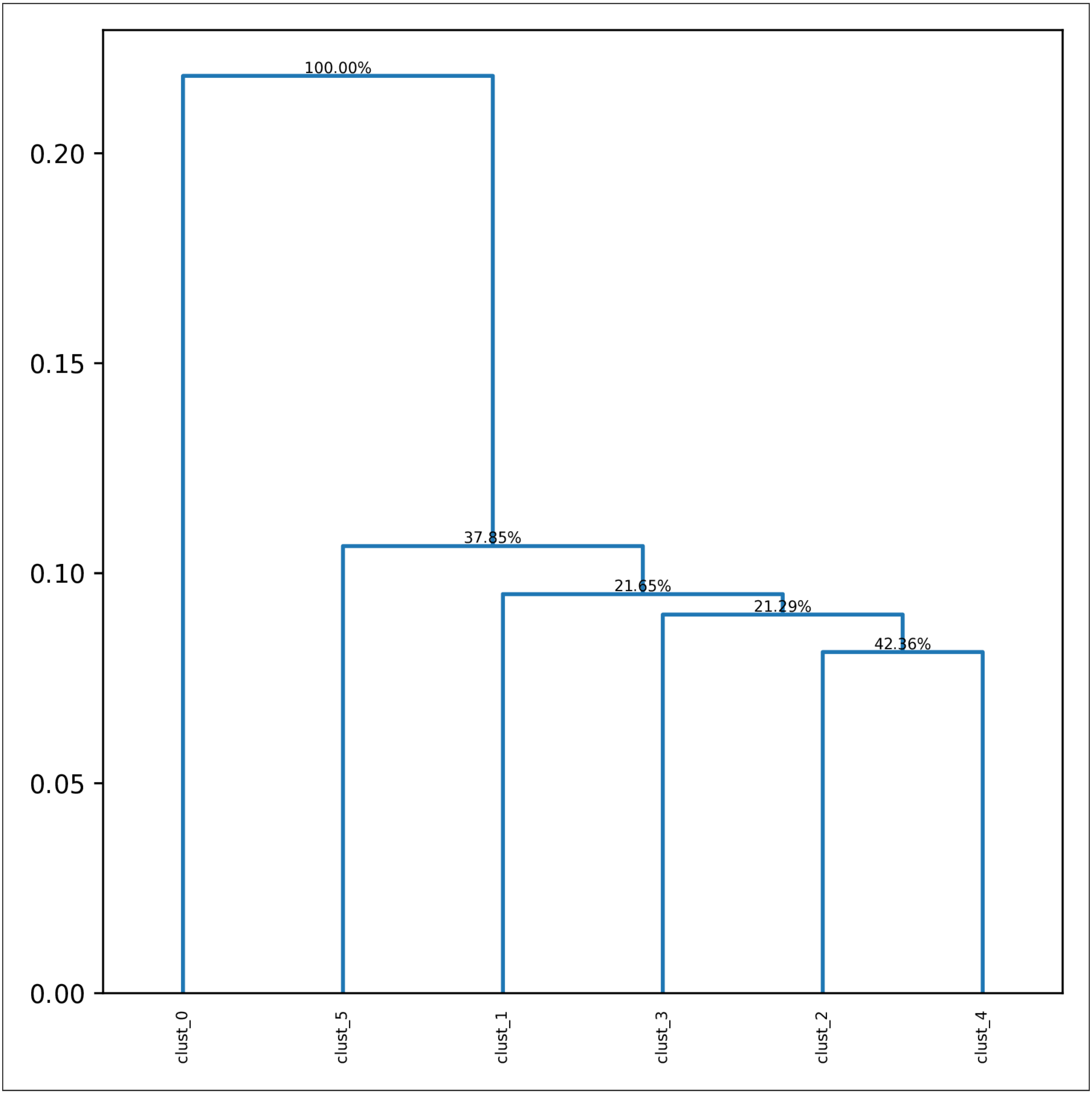

Sample: G0042 with 500 cells per cluster

Min. read depth: 10,  Mean UMIs per cluster:  3,148,425

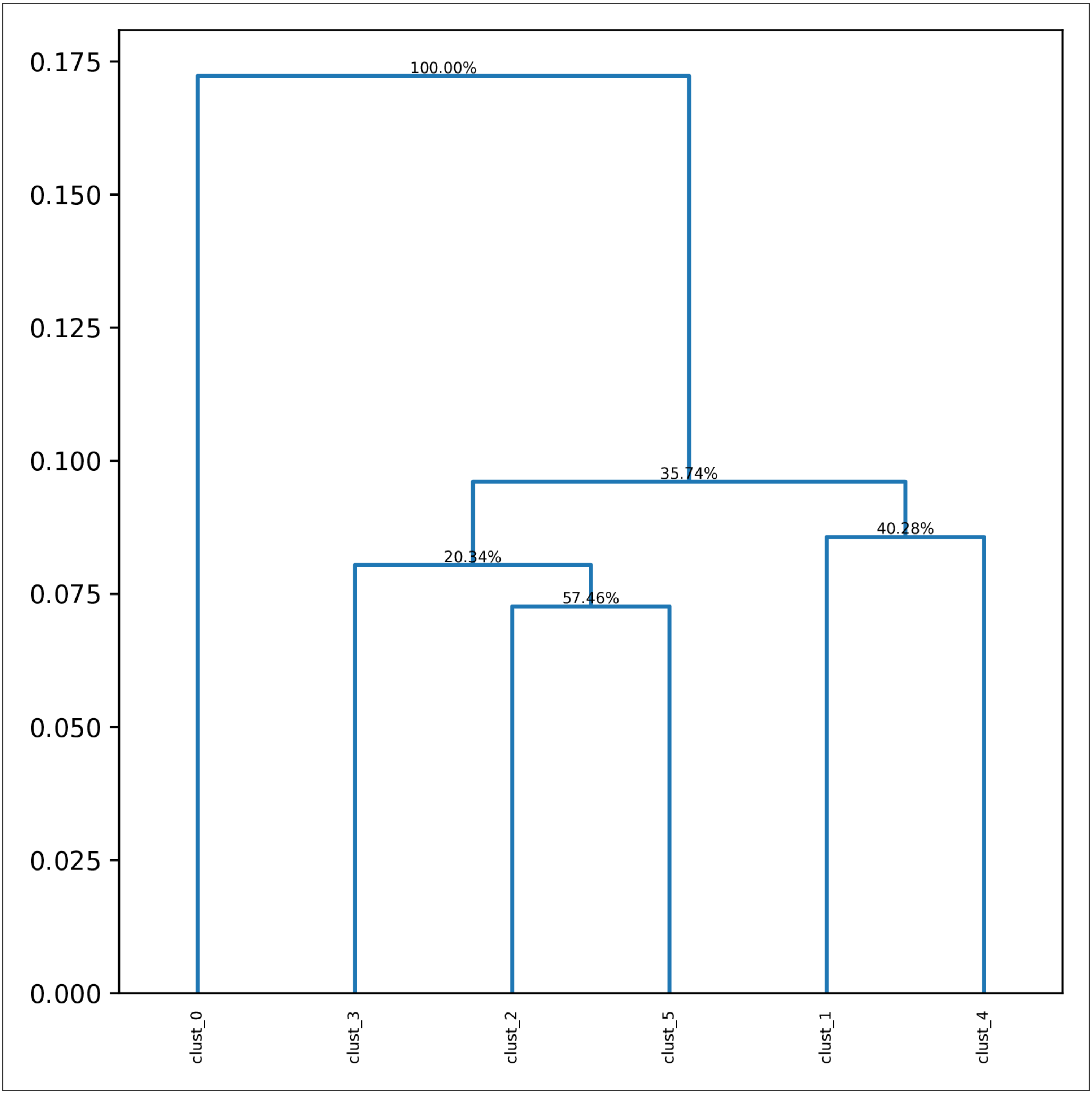

Sample: G0042 with 100 cells per cluster

Min. read depth: 10,  Mean UMIs per cluster:  637,998

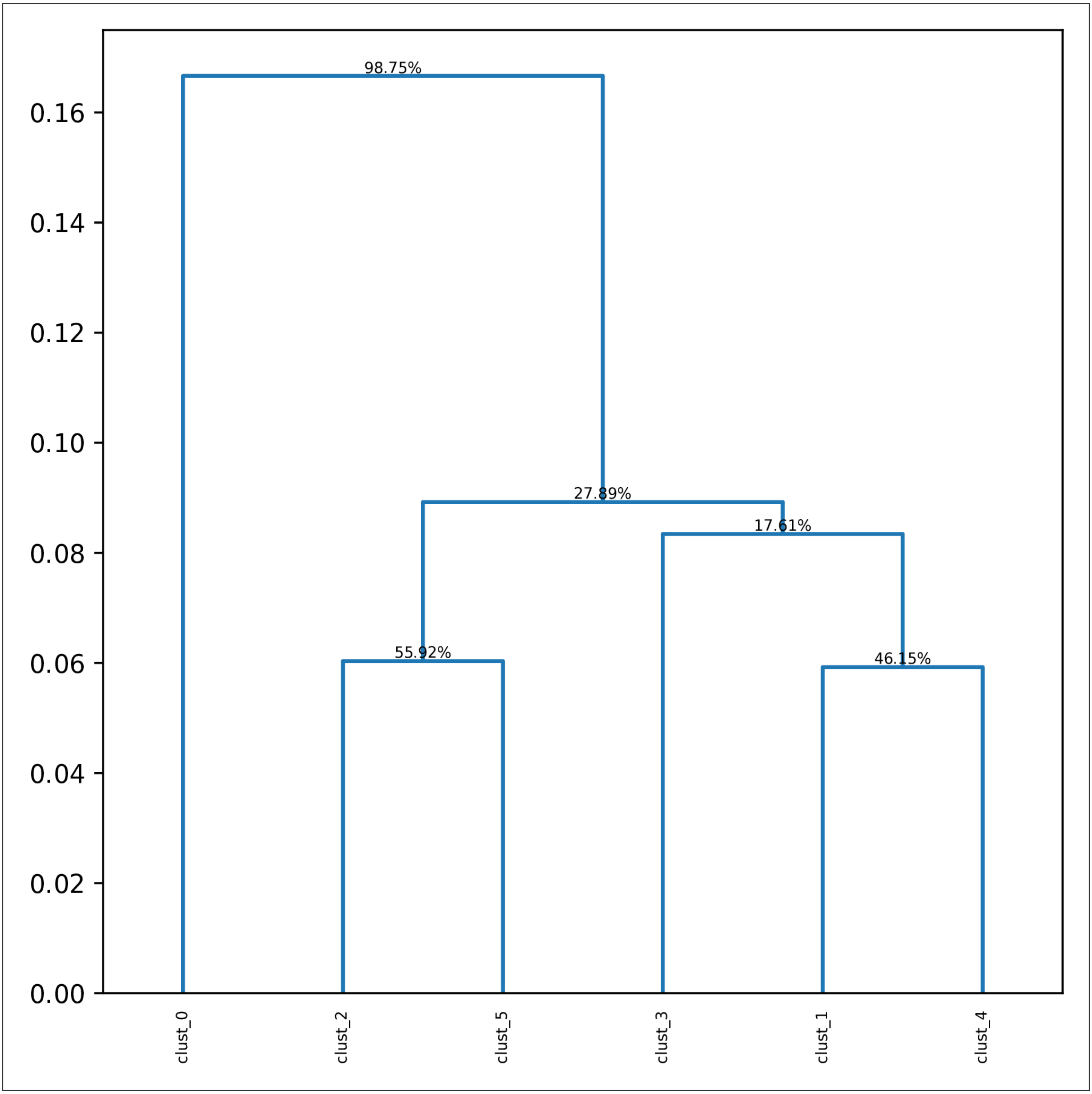

Sample: G0042 with 50 cells per cluster

Min. read depth: 10,  Mean UMIs per cluster:  323,816

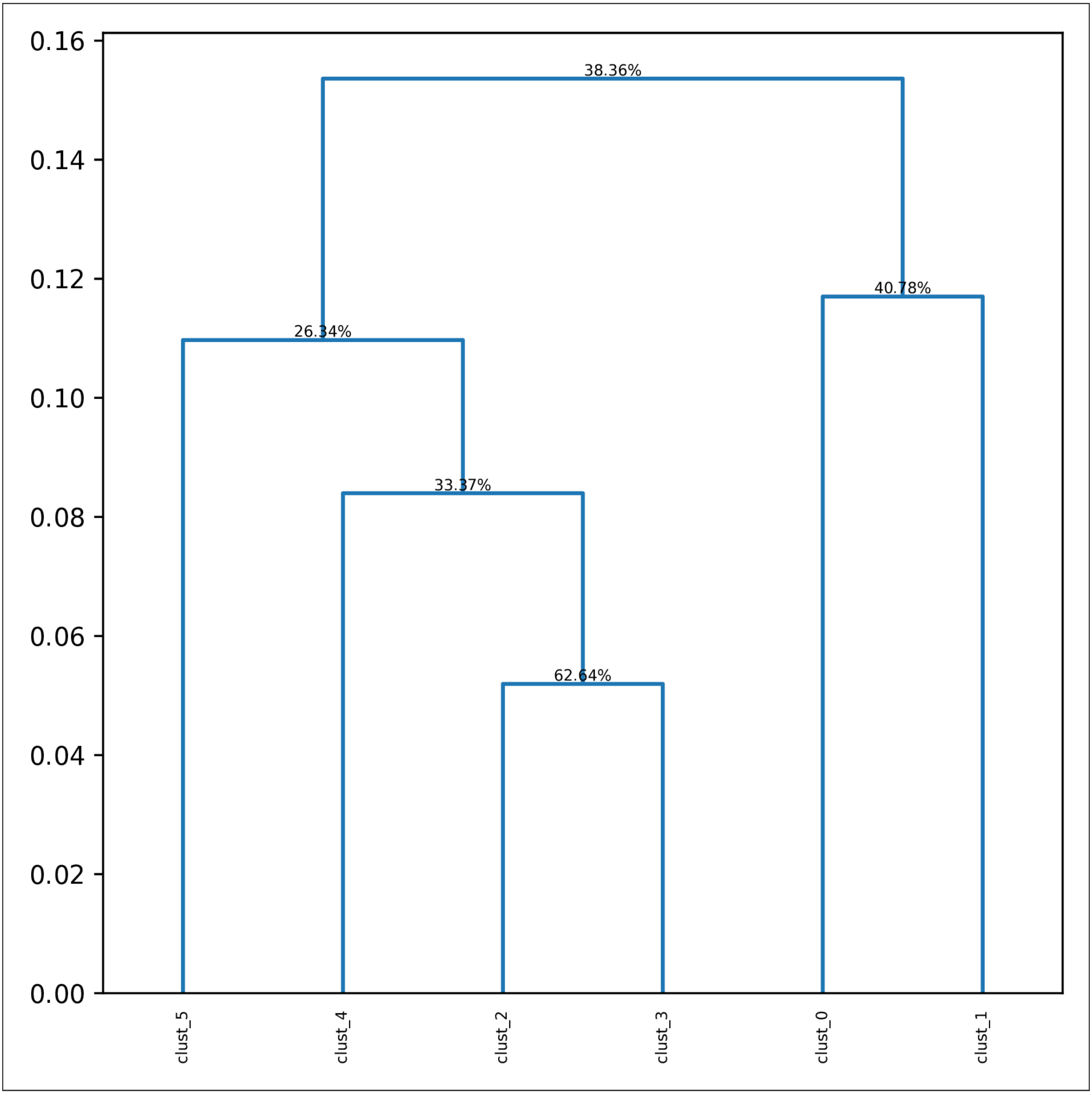

Sample: G0042 with 20 cells per cluster

Min. read depth: 10,  Mean UMIs per cluster:  112,764

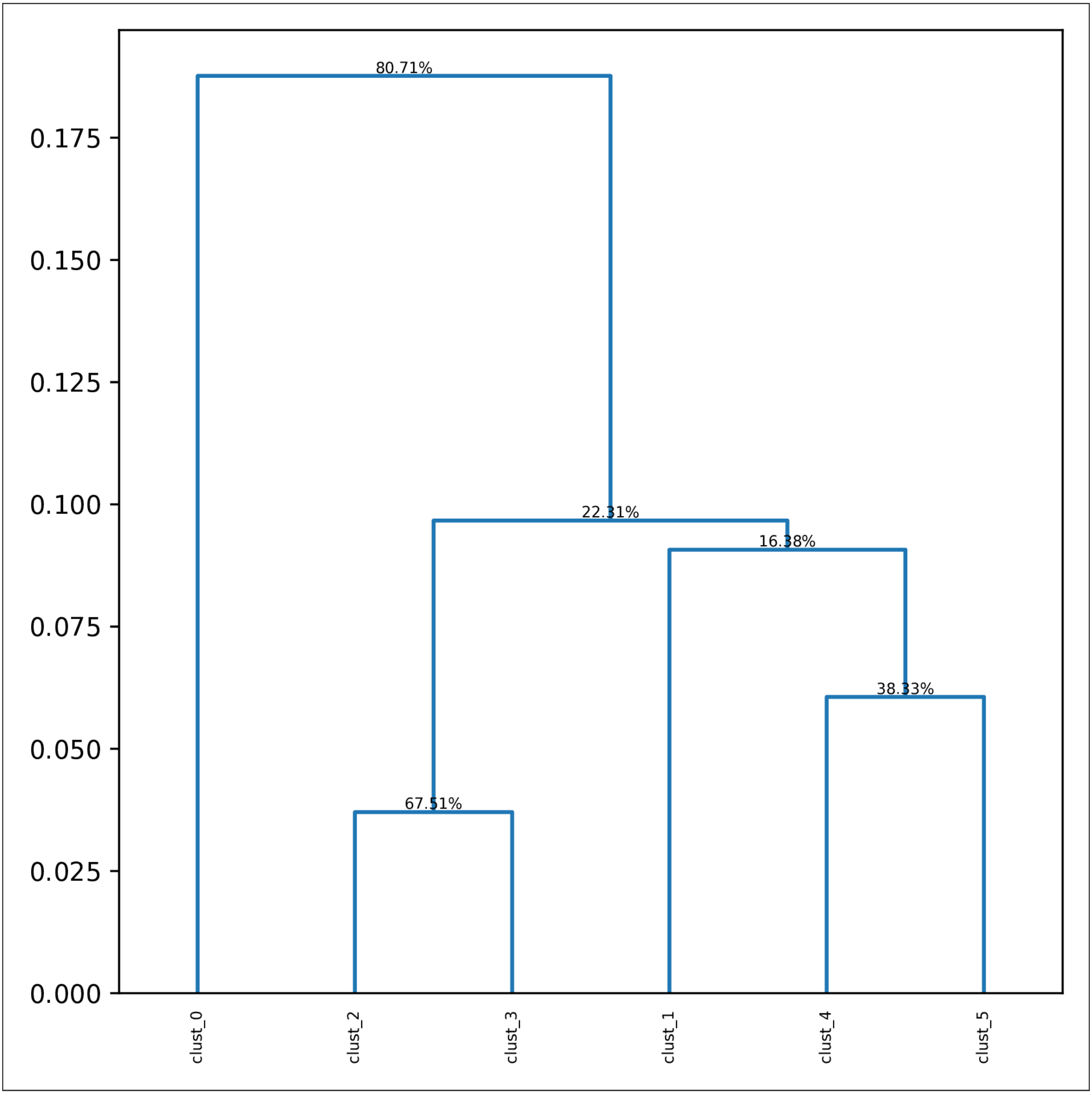

Sample: G0042 with 10 cells per cluster

Min. read depth: 10,  Mean UMIs per cluster:  60,785

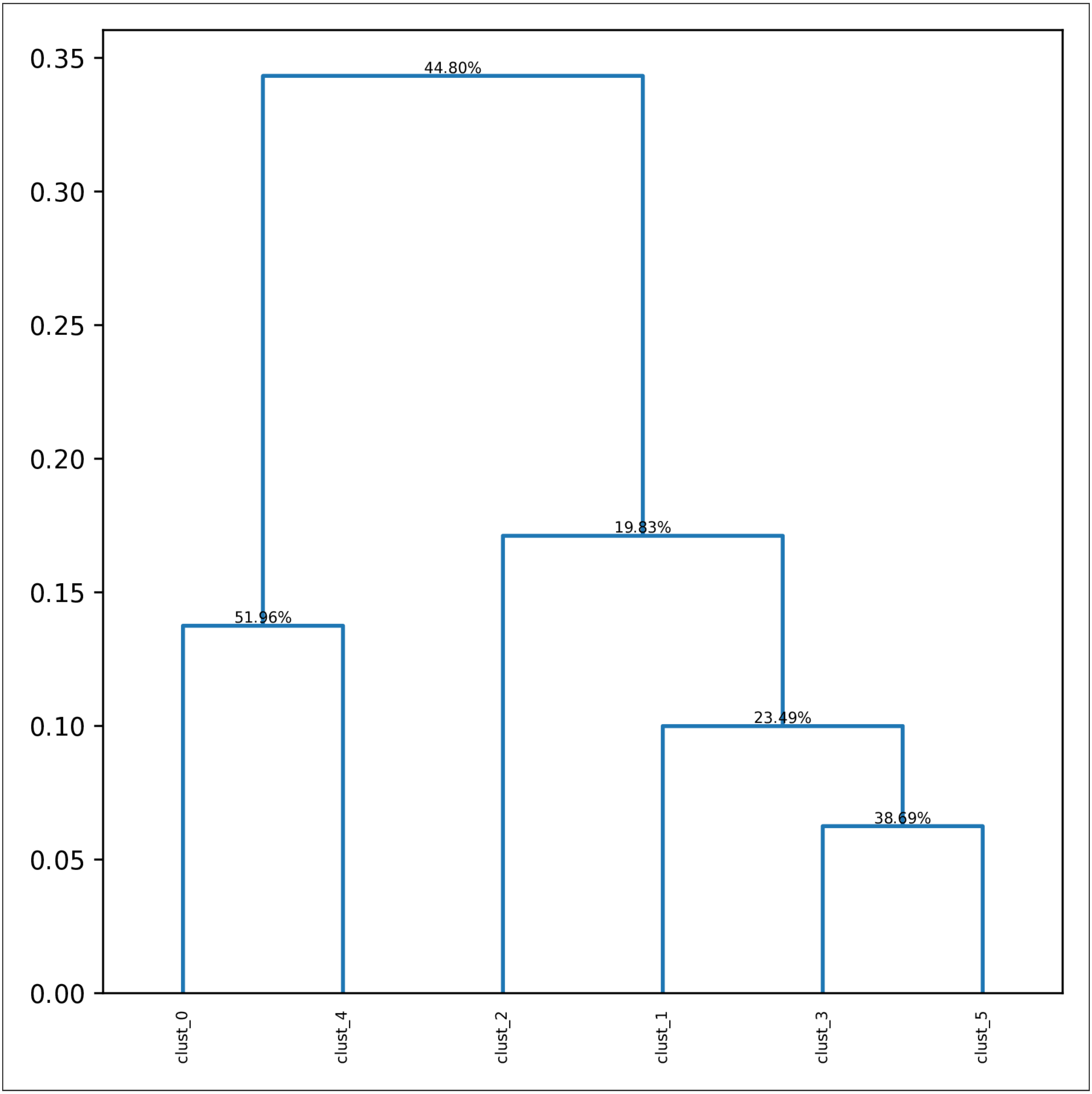

Sample: G0042 with 4 cells per cluster

Min. read depth: 10,  Mean UMIs per cluster:  26,519

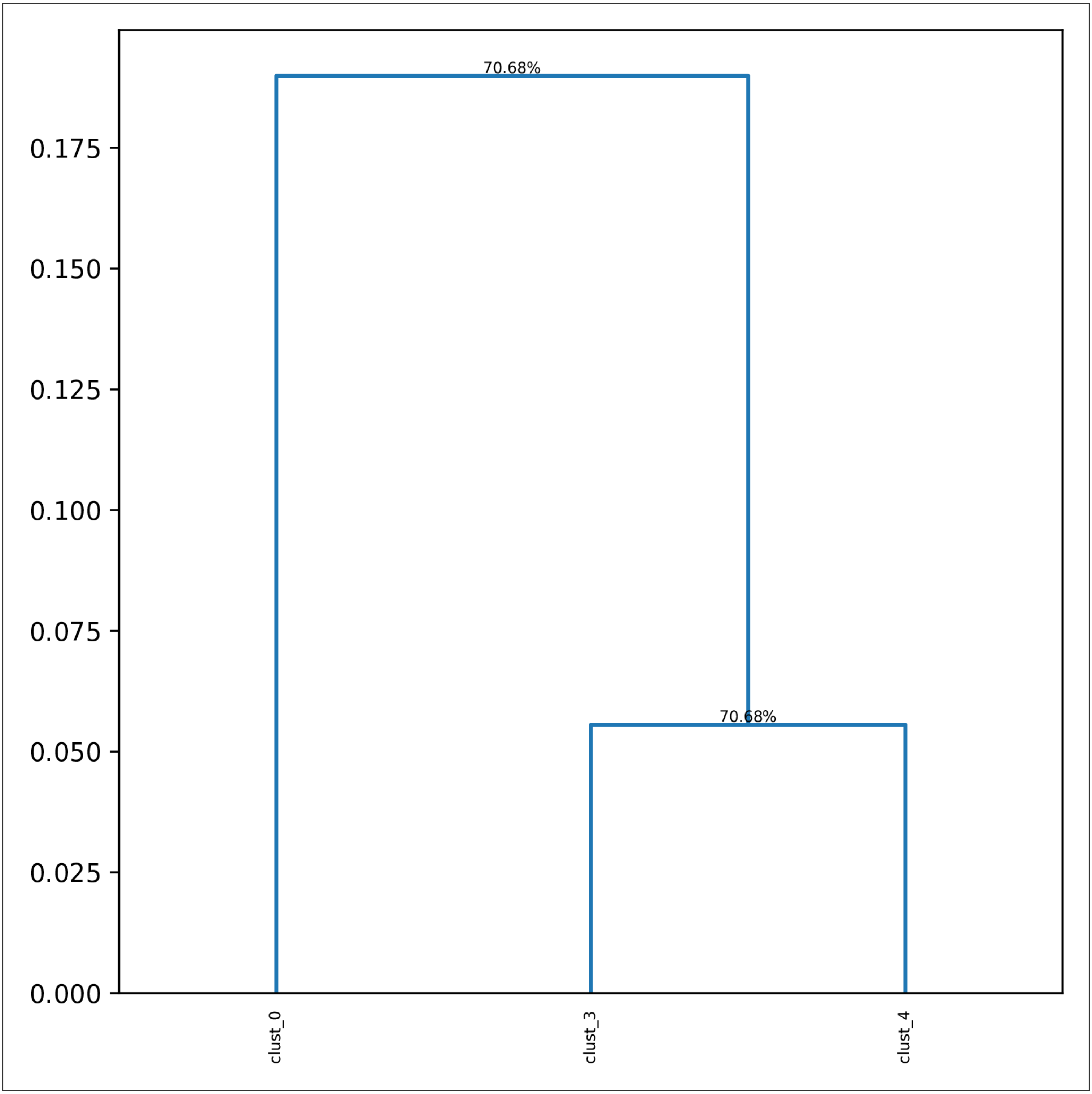

Sample: G0042 with 3 cells per cluster
Min. read depth: 10,  Mean UMIs per cluster:  21,532

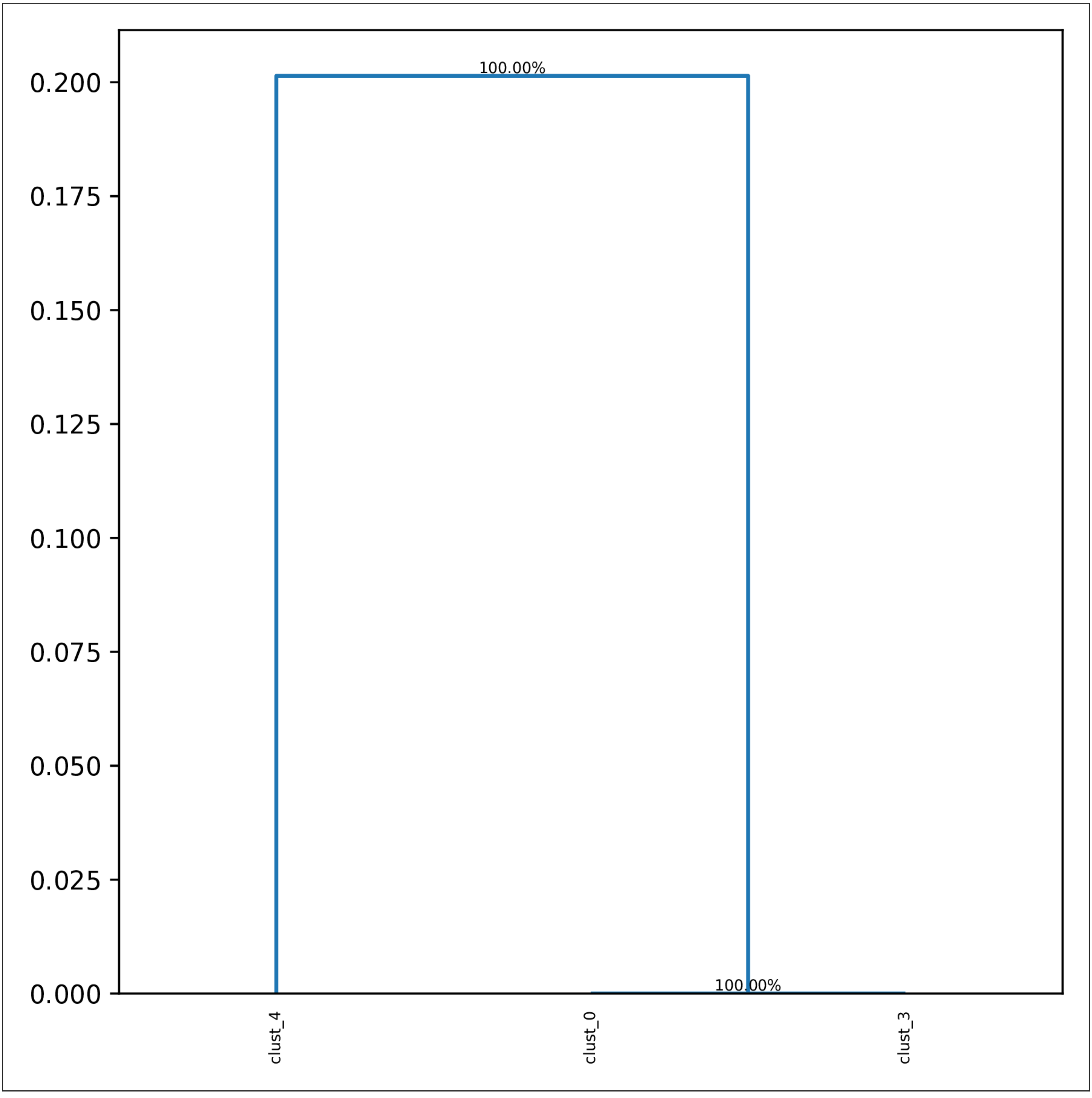

Sample: G0043 with all cells per cluster

Min. read depth: 10,  Mean UMIs per cluster:  9,155,835

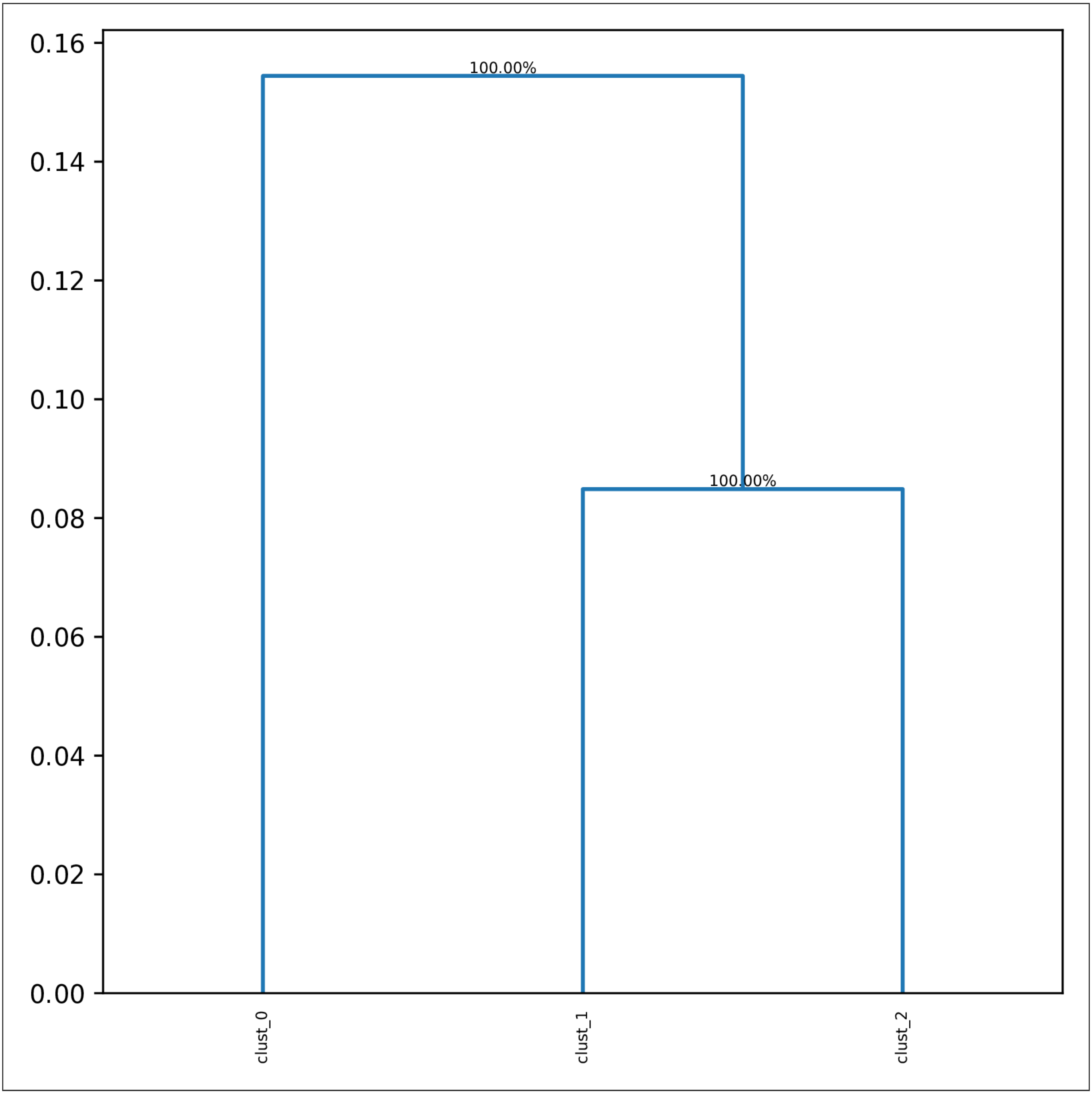

Sample: G0043 with 500 cells per cluster

Min. read depth: 10, Mean UMIs per cluster: 1,778,843

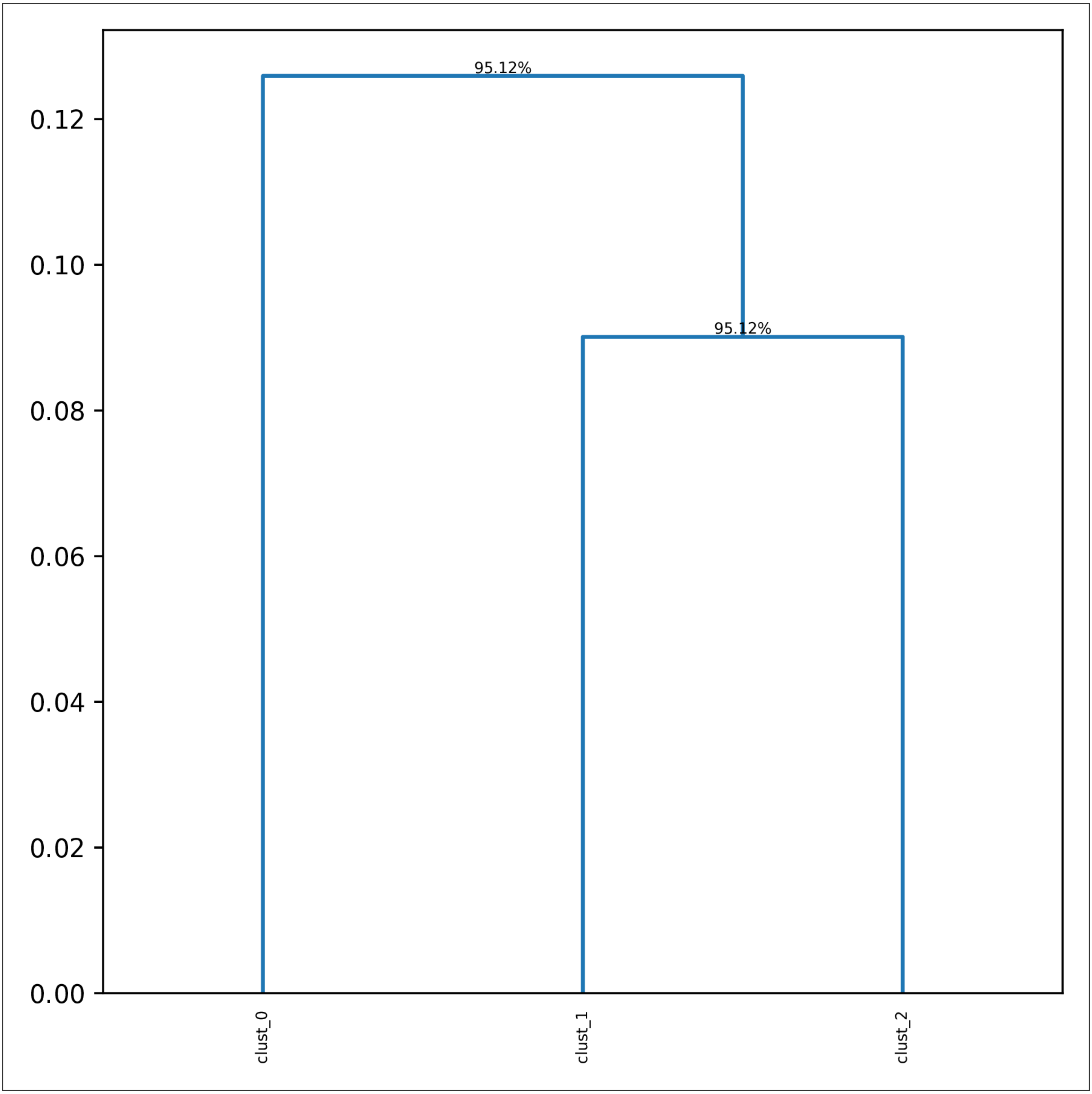

Sample: G0043 with 100 cells per cluster

Min. read depth: 10,  Mean UMIs per cluster:  418,448

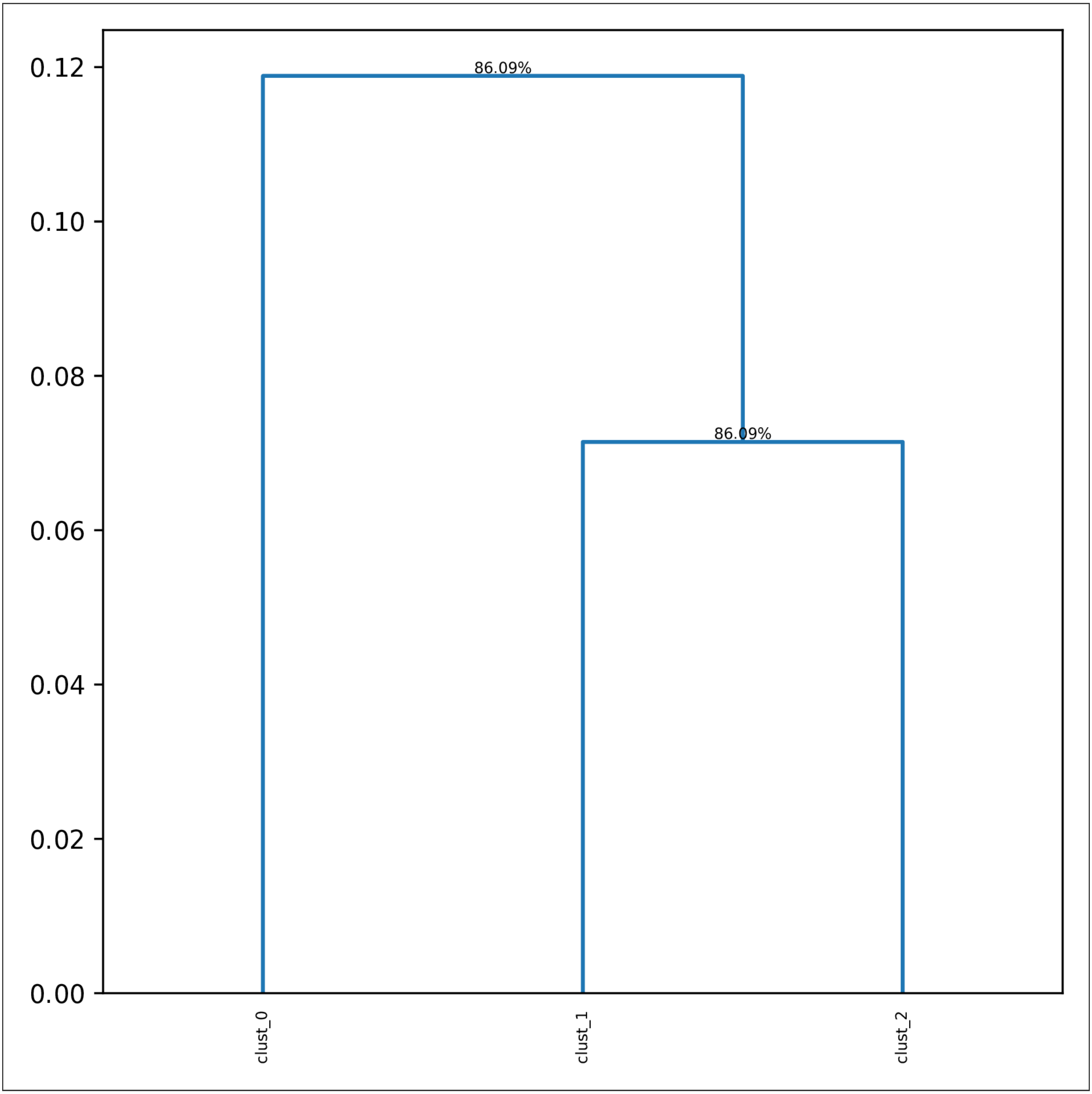

Sample: G0043 with 50 cells per cluster

Min. read depth: 10,  Mean UMIs per cluster:  213,775

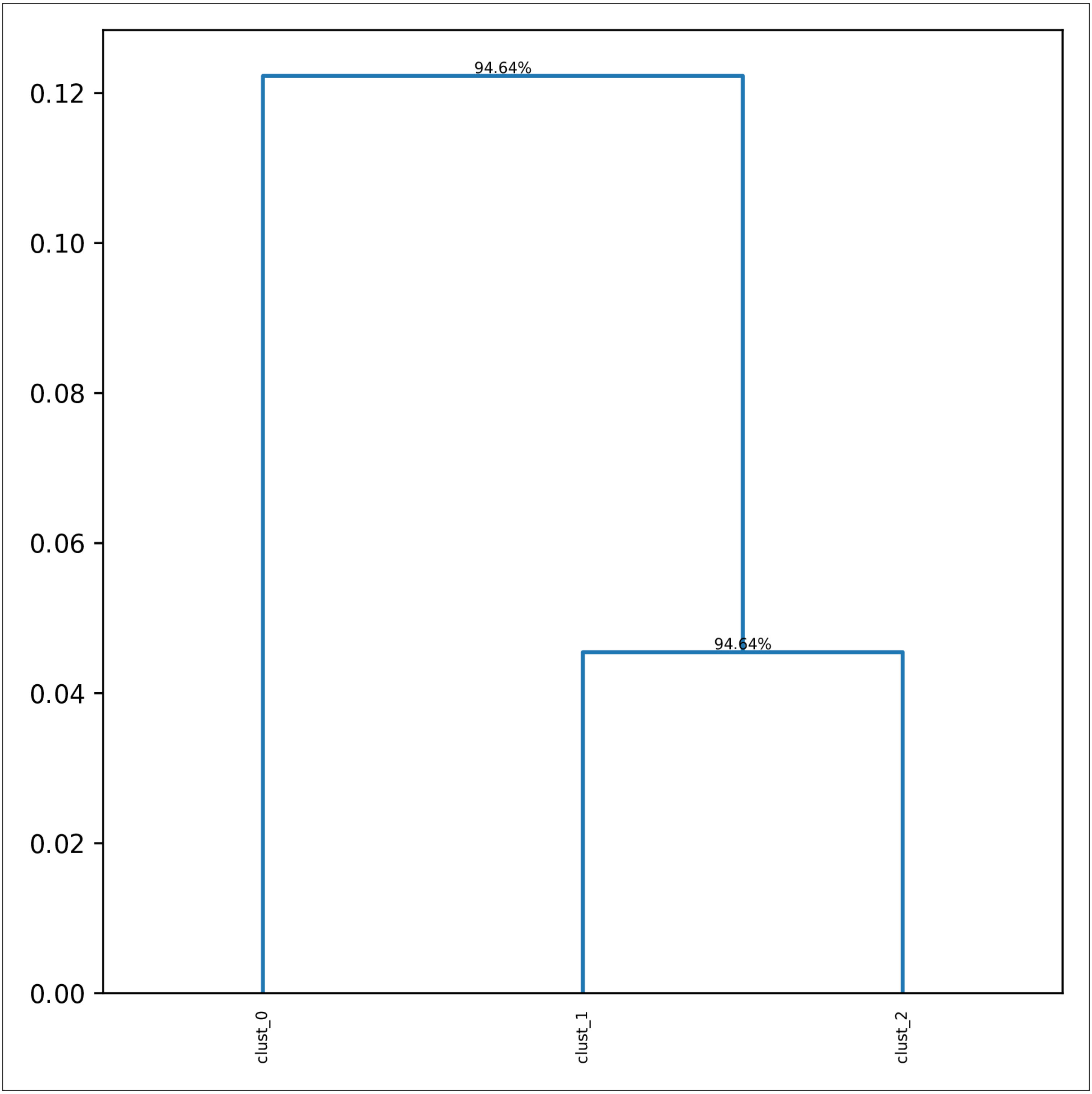

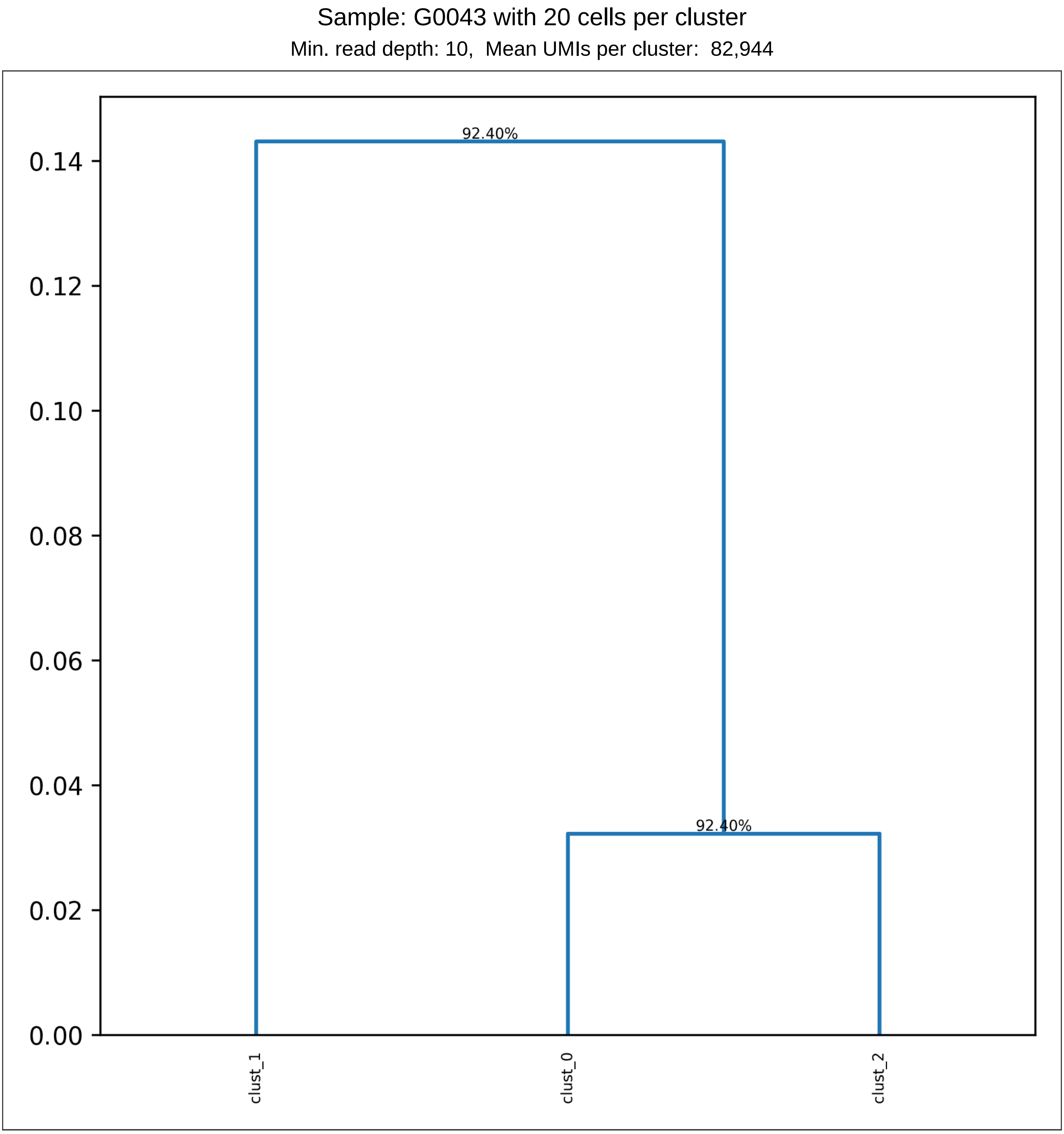

Sample: G0043 with 20 cells per cluster
Min. read depth: 10, Mean UMIs per cluster: 82,944
92.40%
92.40%
0.14
0.12
0.10
0.08
0.06
0.04
0.02
0.00
clust_1
clust_0
clust_2

Sample: G0043 with 10 cells per cluster
Min. read depth: 10,  Mean UMIs per cluster:  48,559

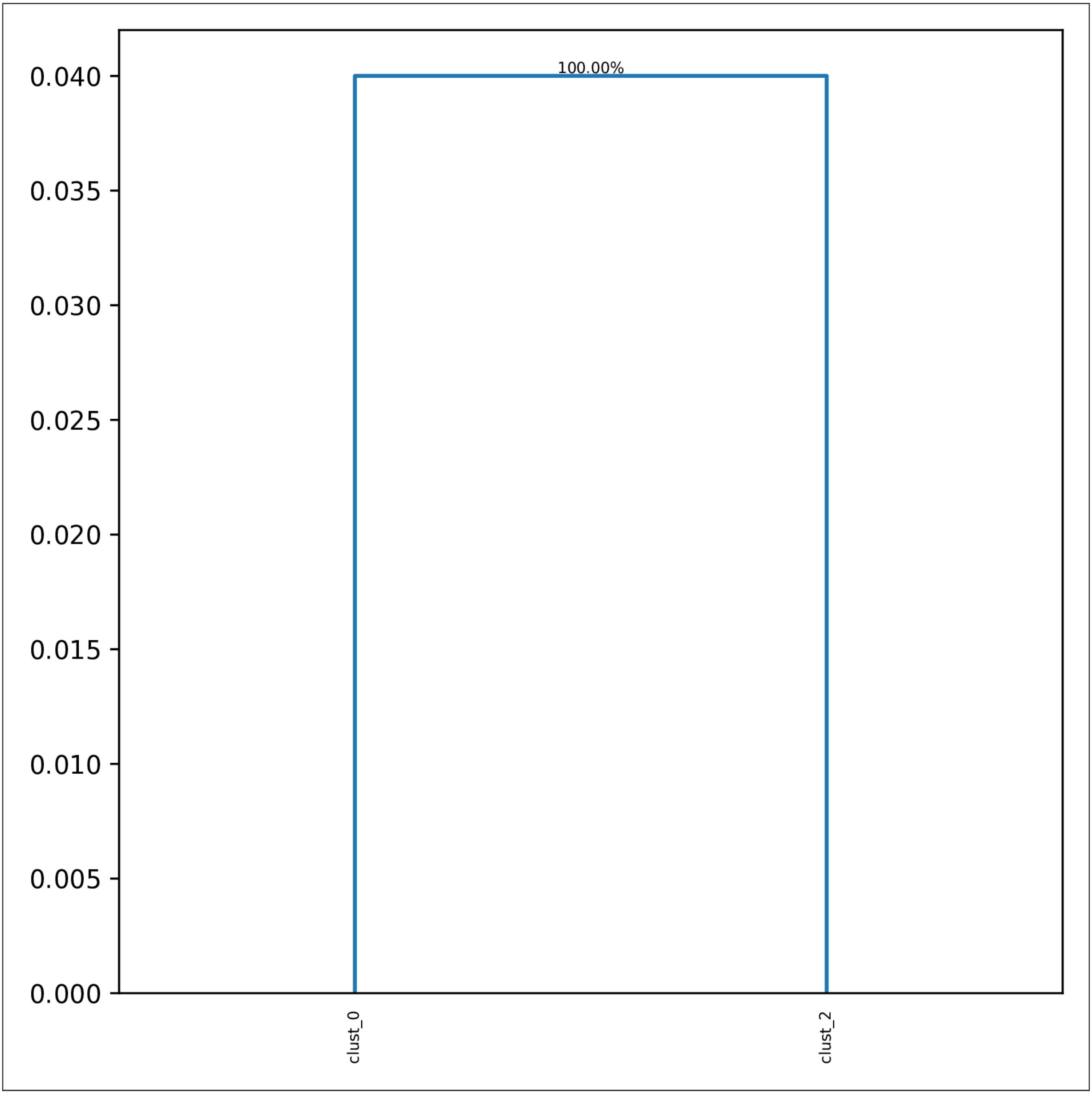

Sample: G0044 with all cells per cluster

Min. read depth: 10,  Mean UMIs per cluster:  11,814,967

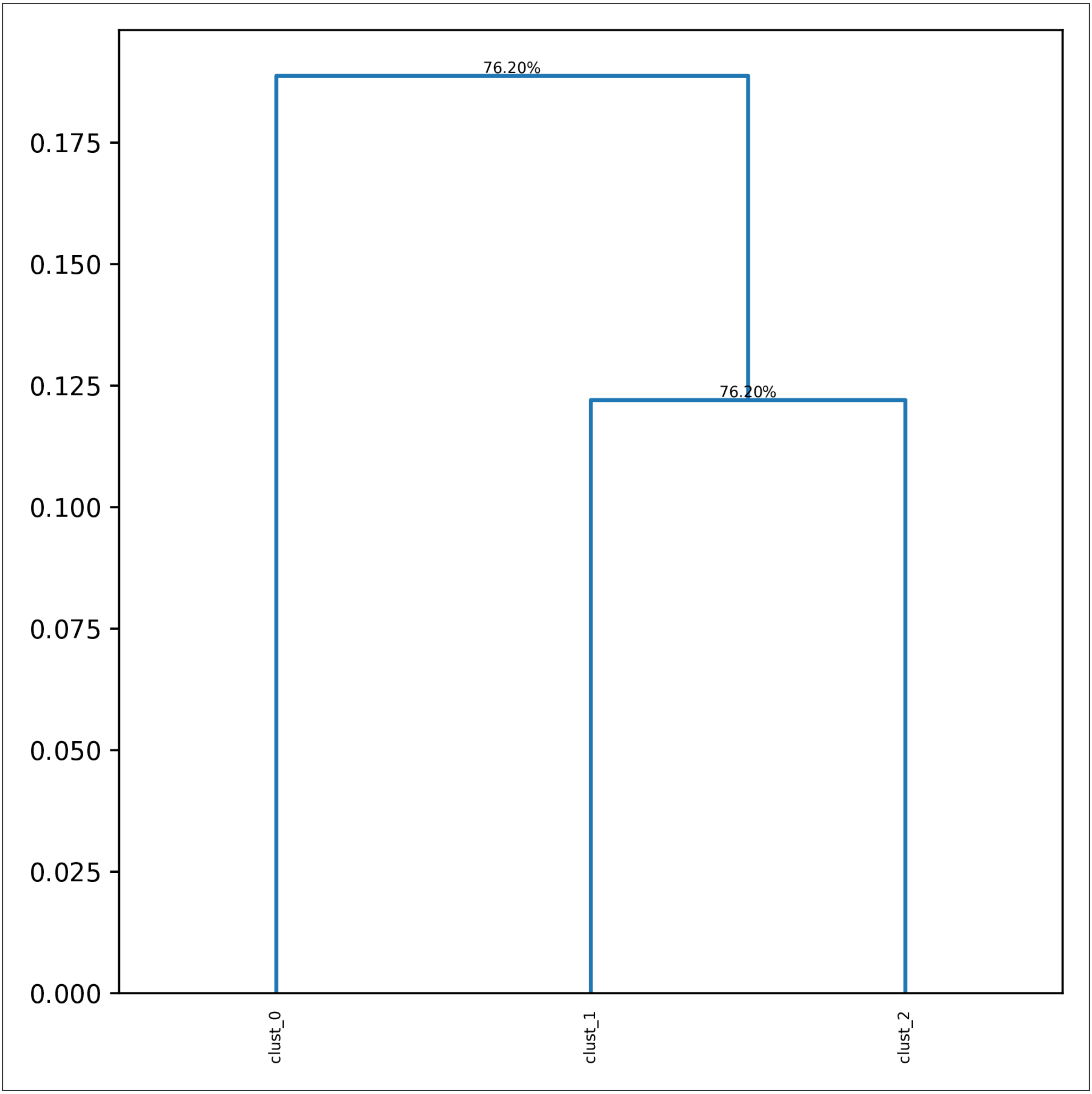

Sample: G0044 with 500 cells per cluster

Min. read depth: 10, Mean UMIs per cluster: 4,877,988

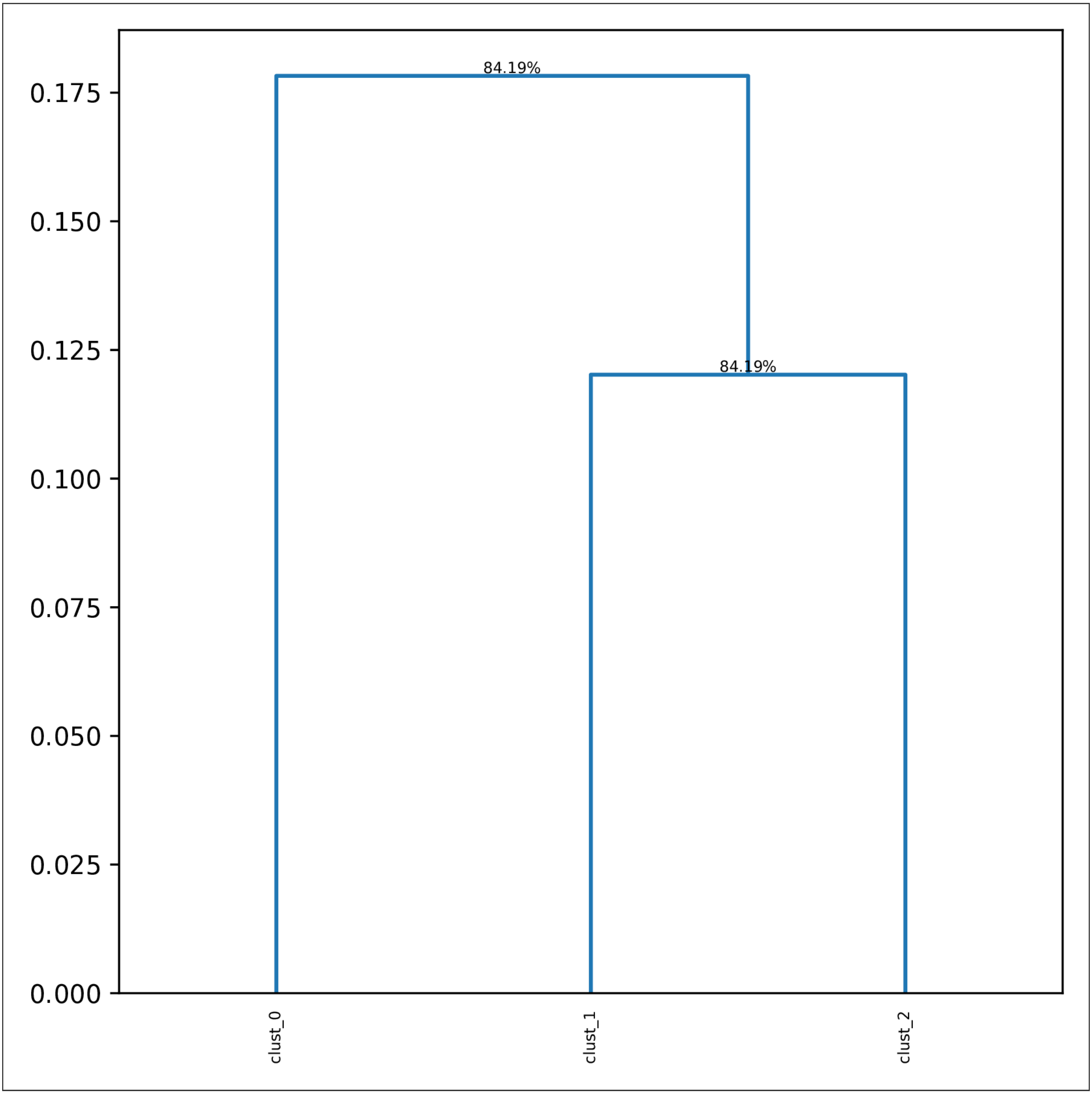

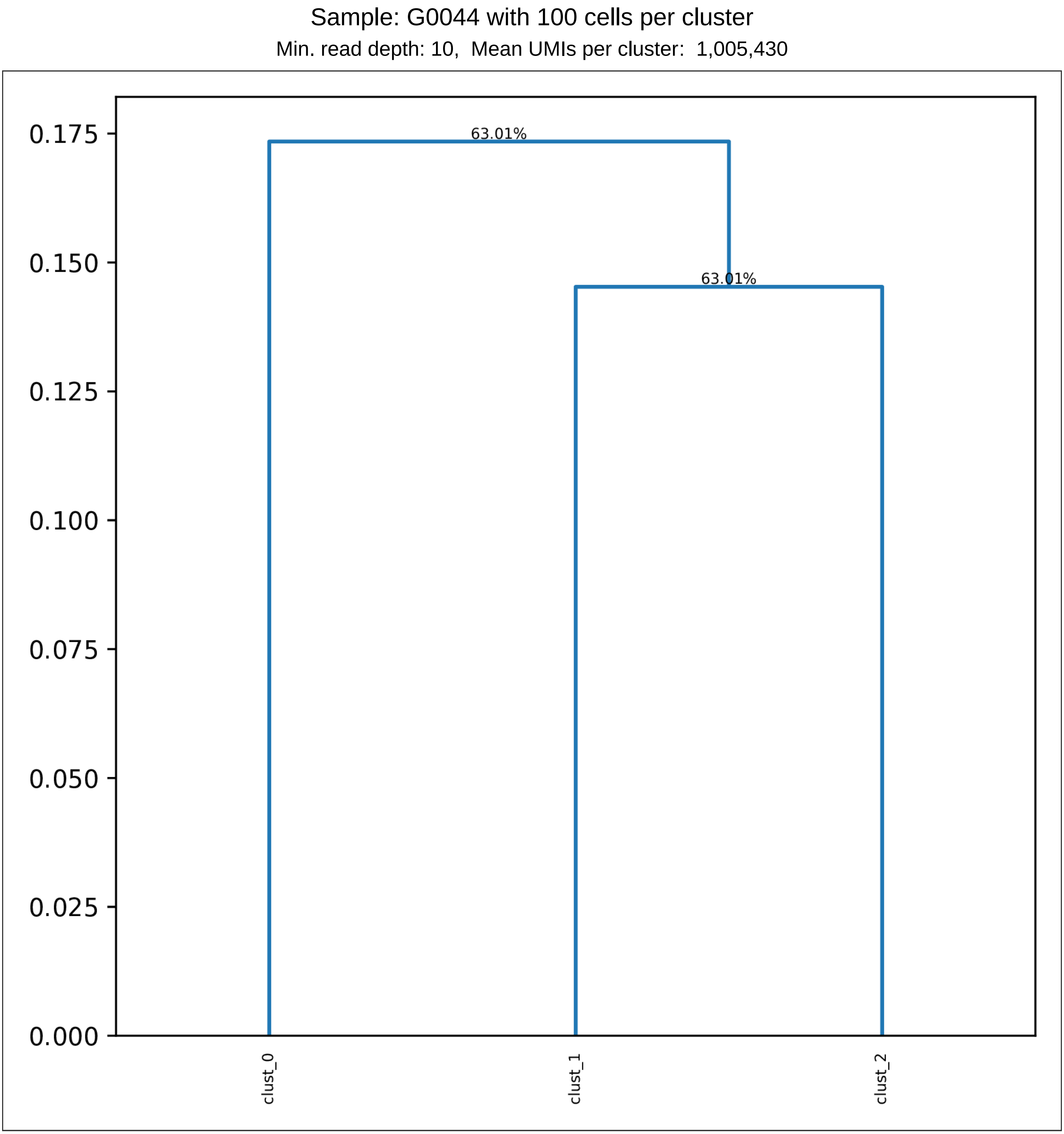
Sample: G0044 with 100 cells per cluster
Min. read depth: 10,  Mean UMIs per cluster:  1,005,430
0.175
0.150
0.125
0.100
0.075
0.050
0.025
0.000
63.01%
63.01%
clust_0
clust_1
clust_2

Sample: G0044 with 50 cells per cluster

Min. read depth: 10,  Mean UMIs per cluster:  514,397

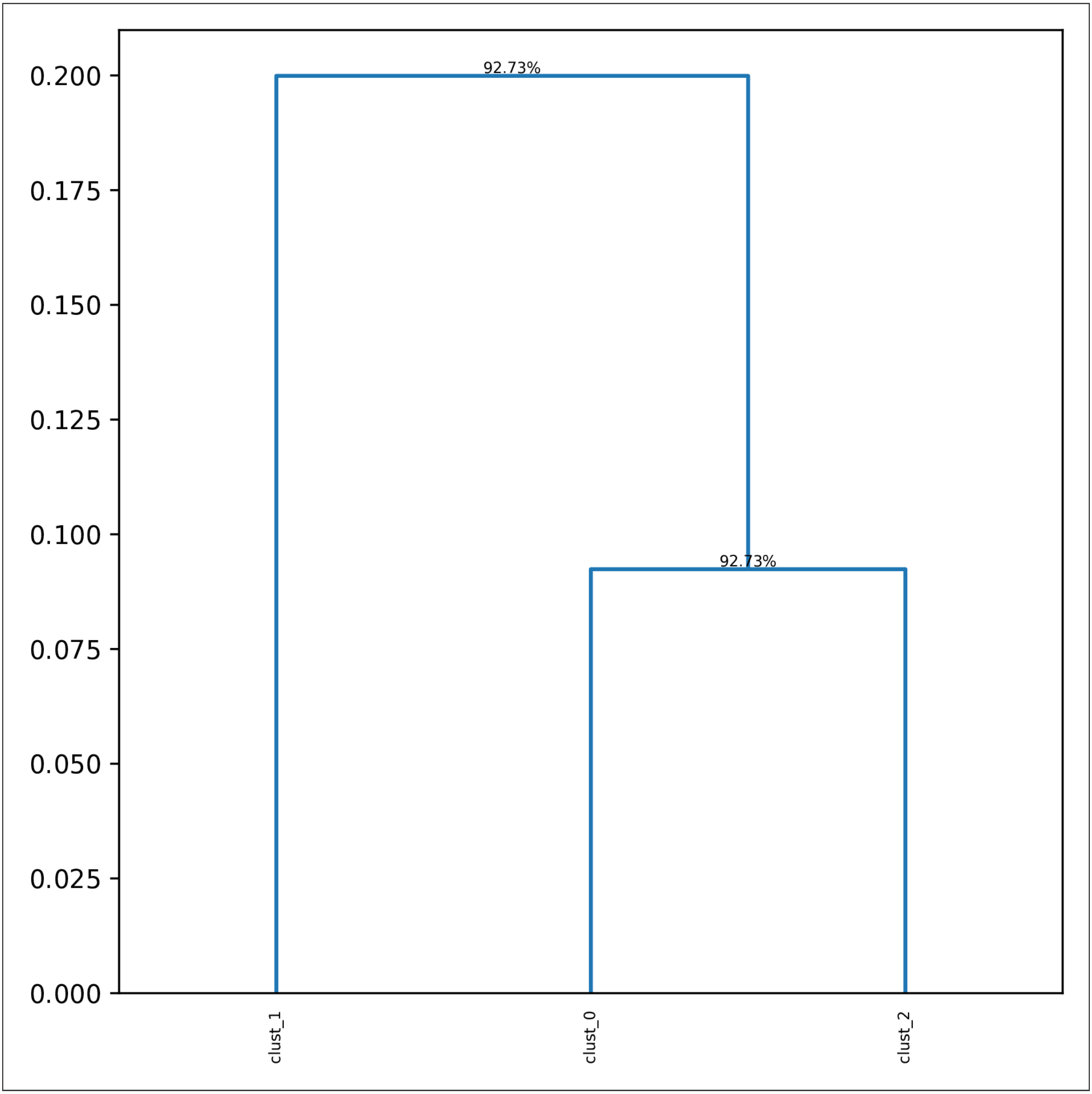

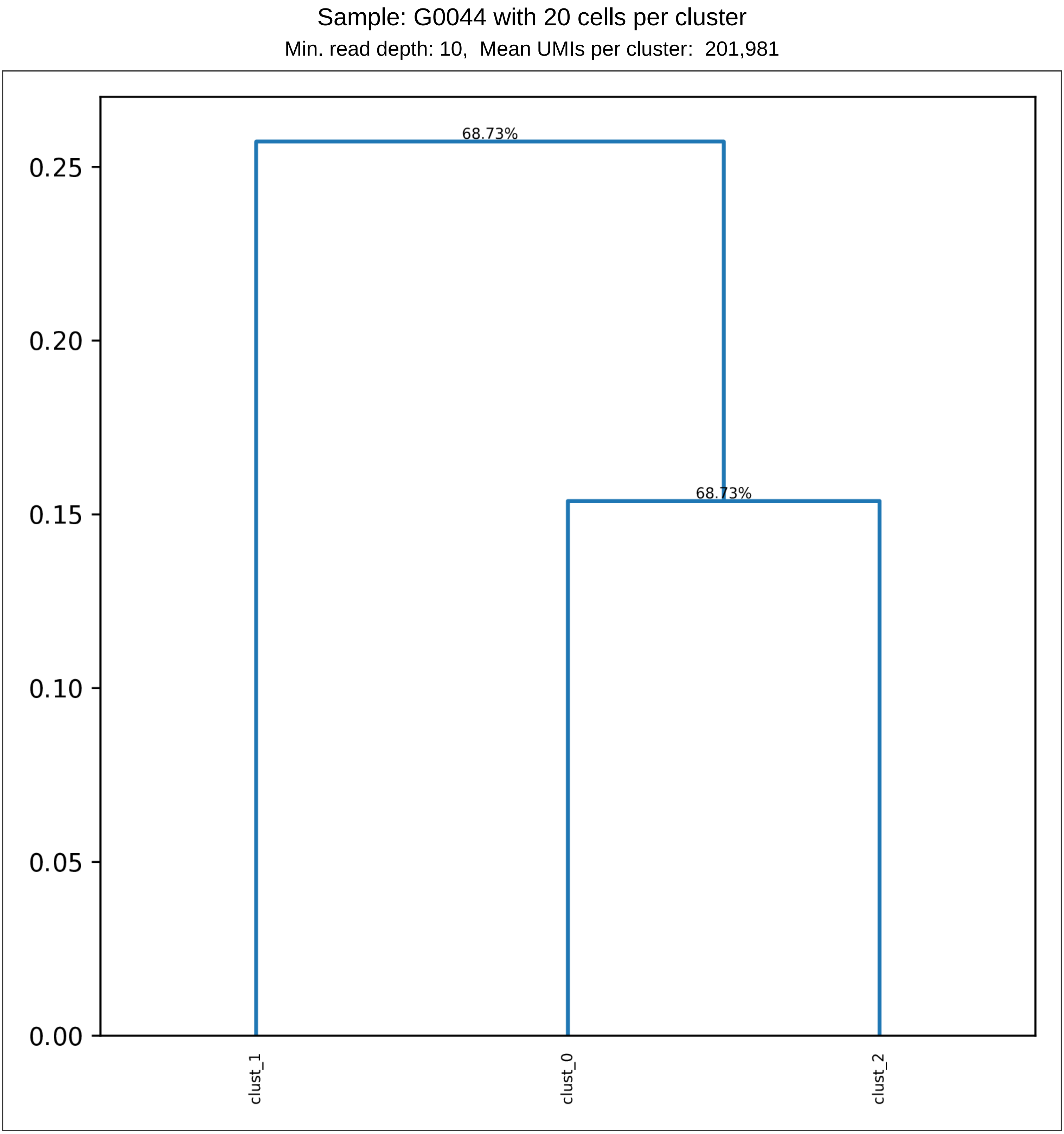
Sample: G0044 with 20 cells per cluster
Min. read depth: 10,  Mean UMIs per cluster:  201,981
68.73%
68.73%
0.25
0.20
0.15
0.10
0.05
0.00
clust_1
clust_0
clust_2

Sample: G0044 with 10 cells per cluster

Min. read depth: 10,  Mean UMIs per cluster:  105,949

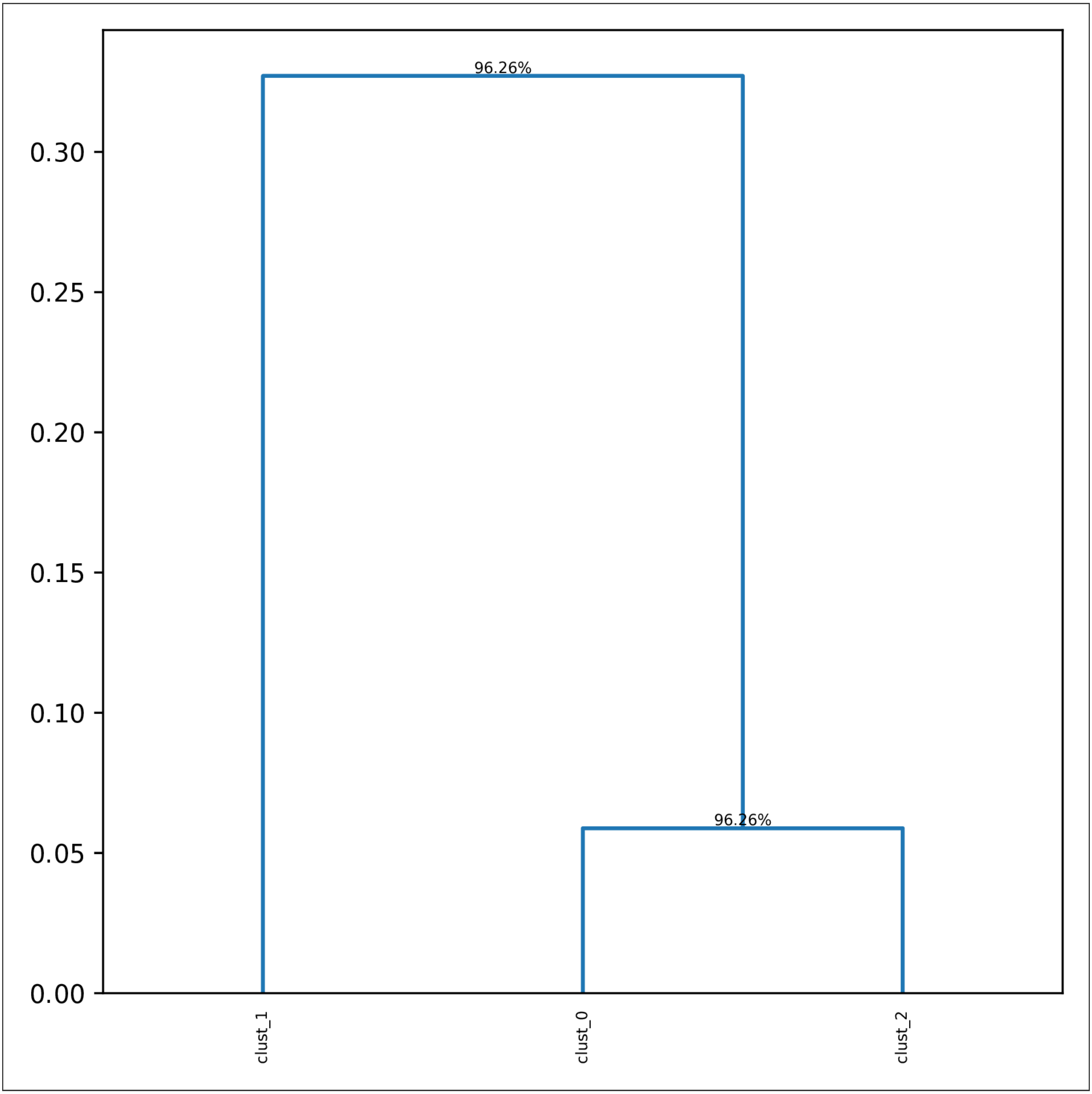

Sample: G0045 with all cells per cluster

Min. read depth: 10,  Mean UMIs per cluster:  6,963,577

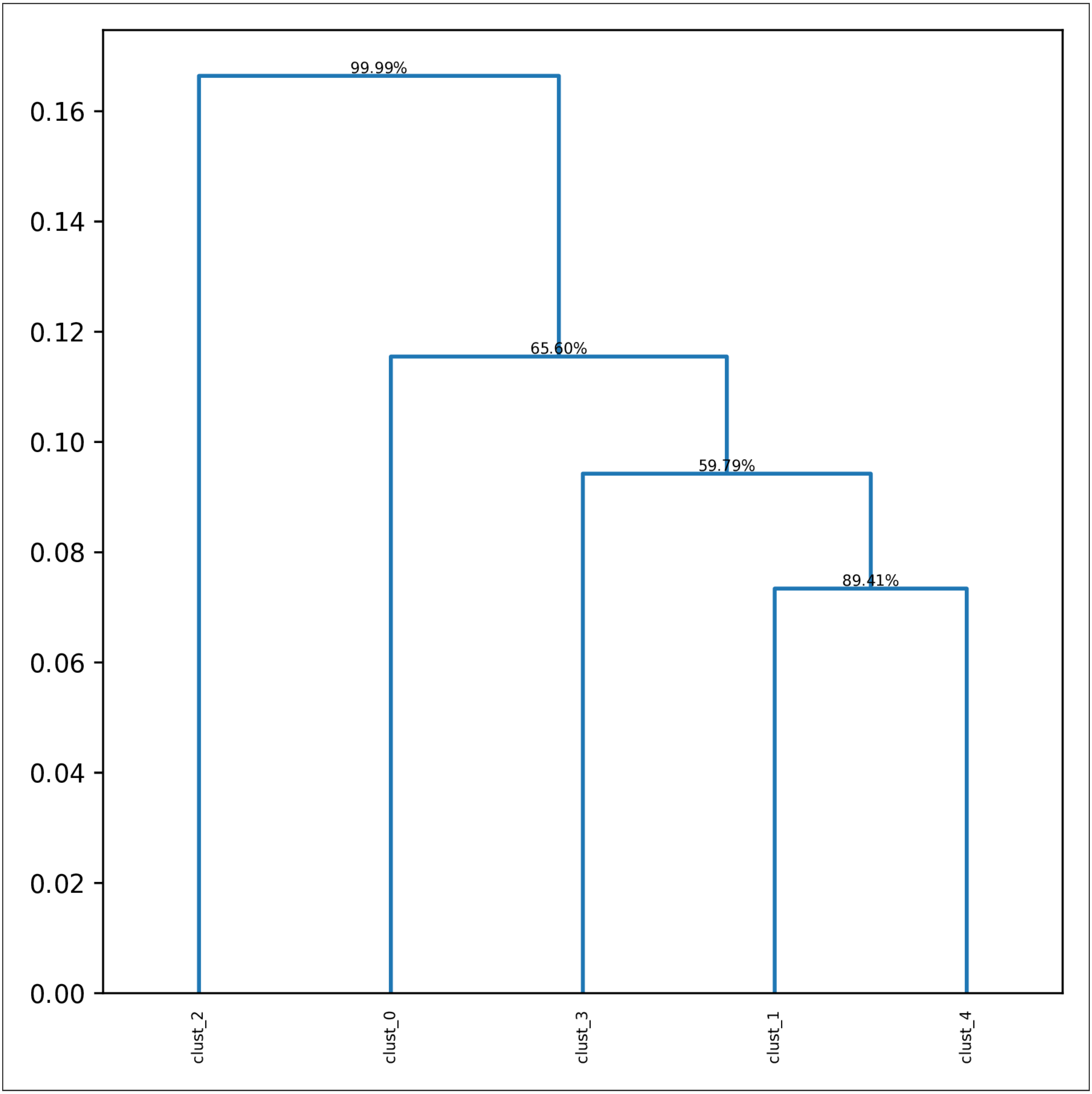

Sample: G0045 with 500 cells per cluster

Min. read depth: 10,  Mean UMIs per cluster:  6,963,577

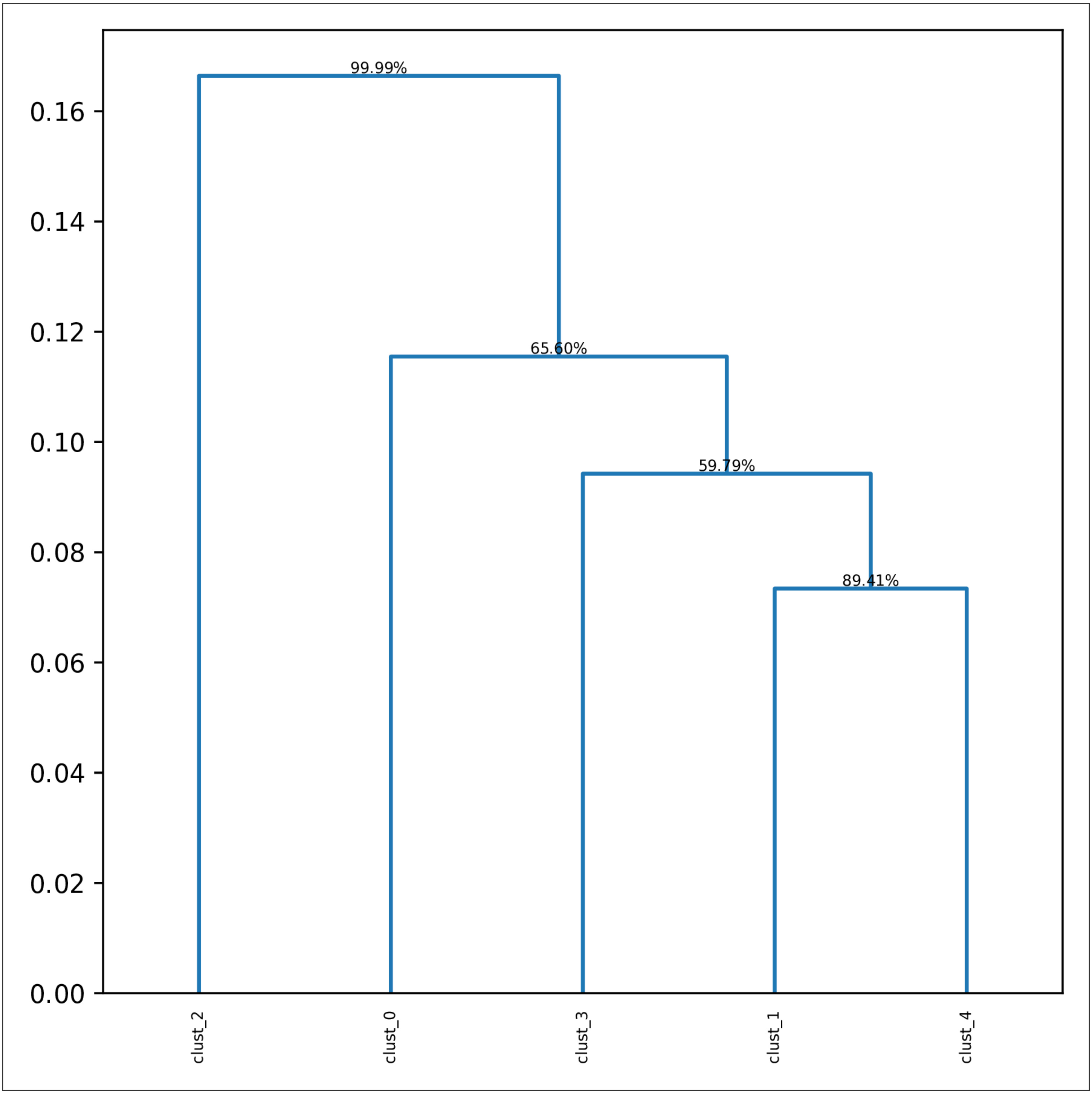

Sample: G0045 with 100 cells per cluster

Min. read depth: 10,  Mean UMIs per cluster:  1,922,758

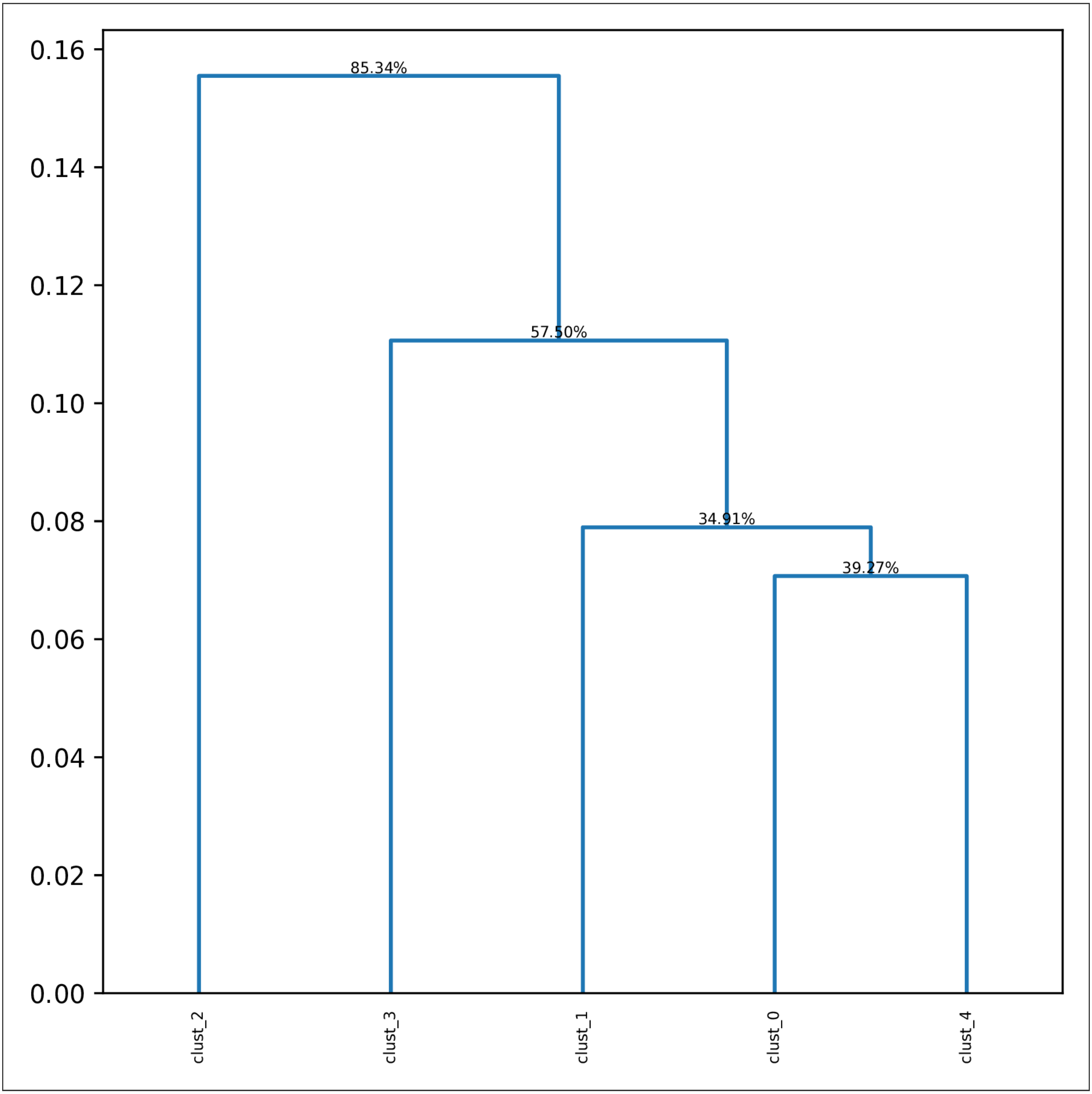

Sample: G0045 with 50 cells per cluster

Min. read depth: 10,  Mean UMIs per cluster:  946,477

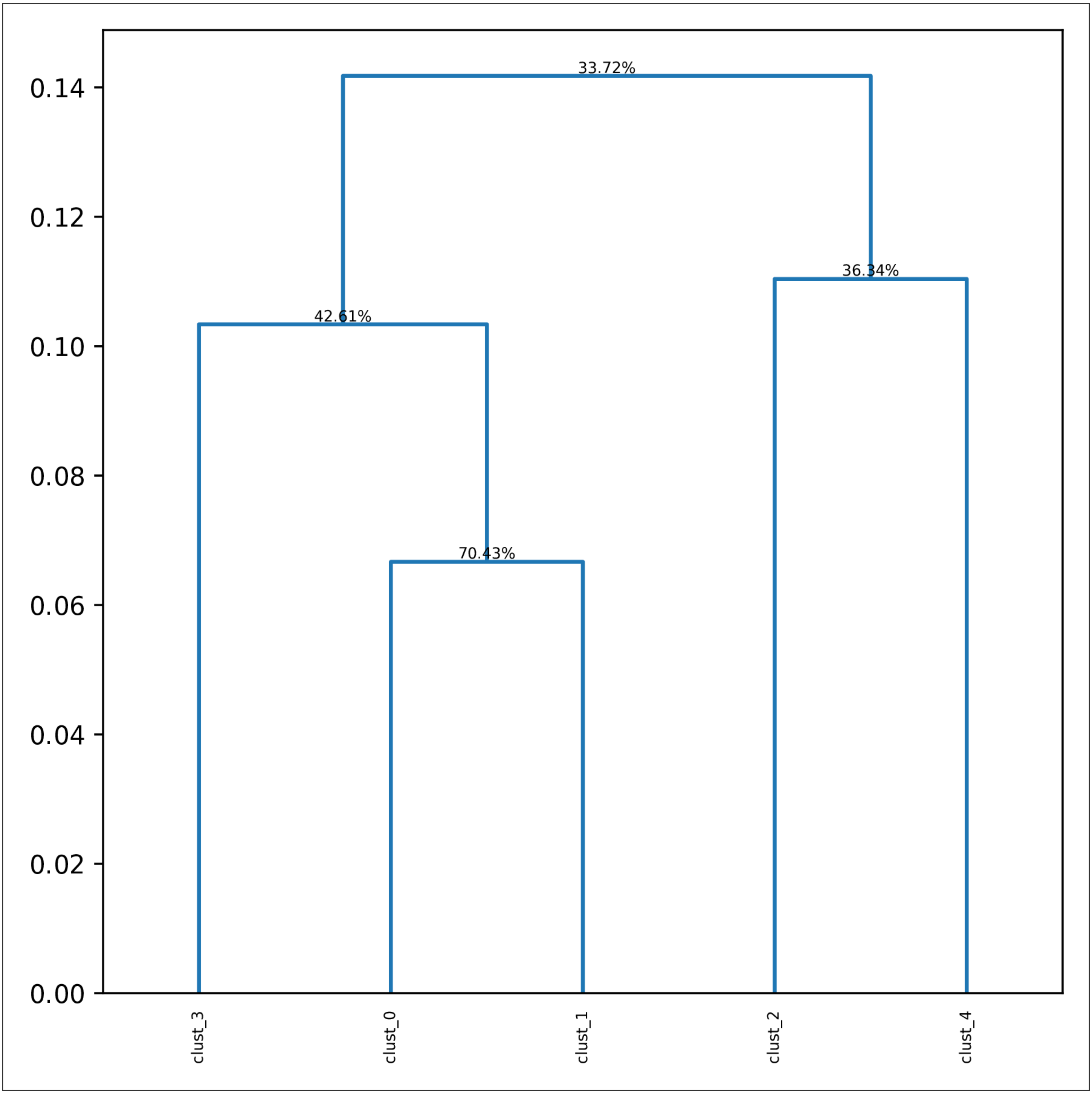

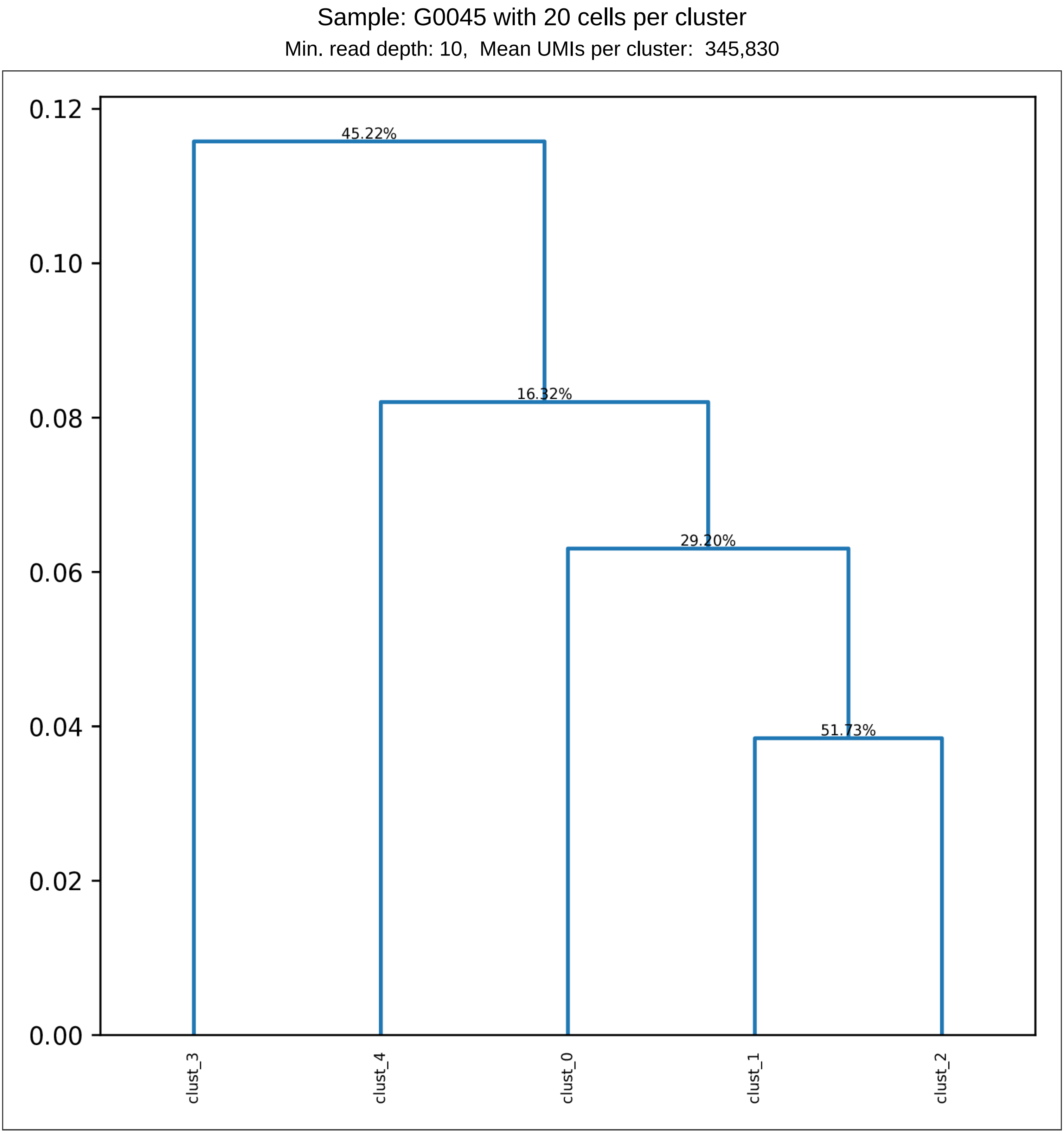
Sample: G0045 with 20 cells per cluster
Min. read depth: 10,  Mean UMIs per cluster:  345,830
0.12
0.10
0.08
0.06
0.04
0.02
0.00
45.22%
16.32%
29.20%
51.73%
clust_3
clust_4
clust_0
clust_1
clust_2

Sample: G0045 with 10 cells per cluster

Min. read depth: 10,  Mean UMIs per cluster:  170,756

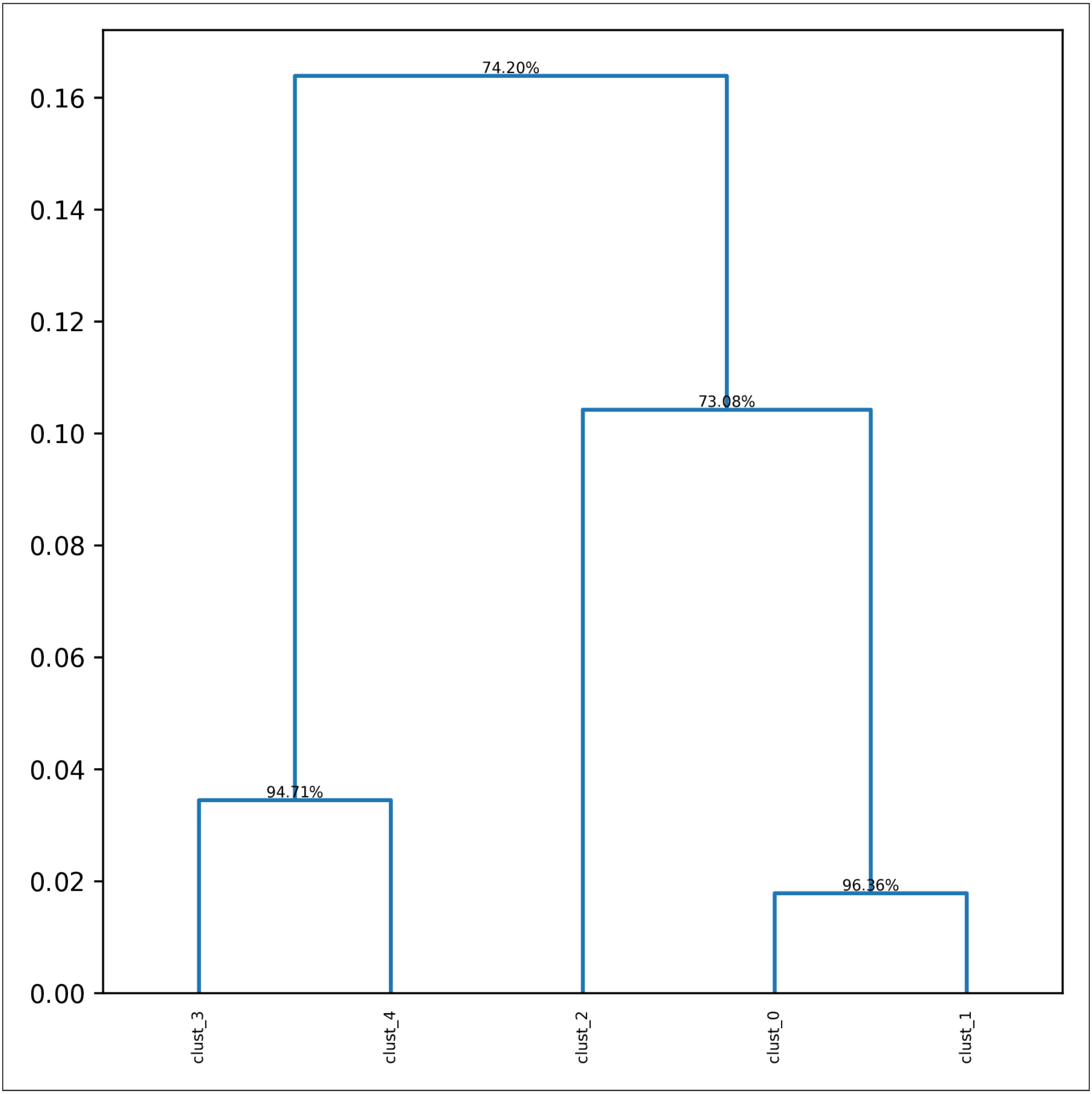

Sample: G0045 with 5 cells per cluster
Min. read depth: 10,  Mean UMIs per cluster:  112,822

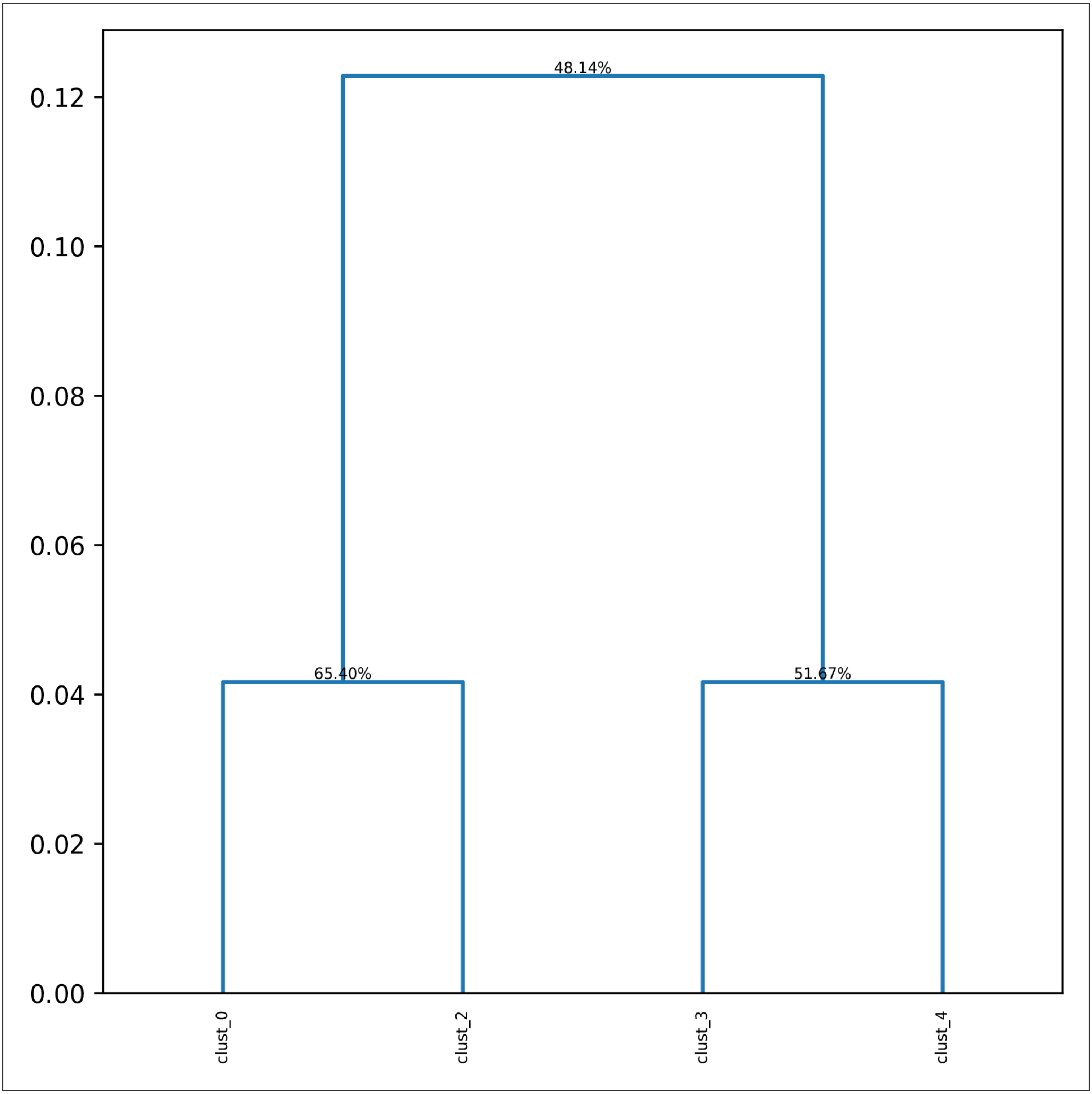

Sample: G0045 with 2 cells per cluster

Min. read depth: 10,  Mean UMIs per cluster:  36,103

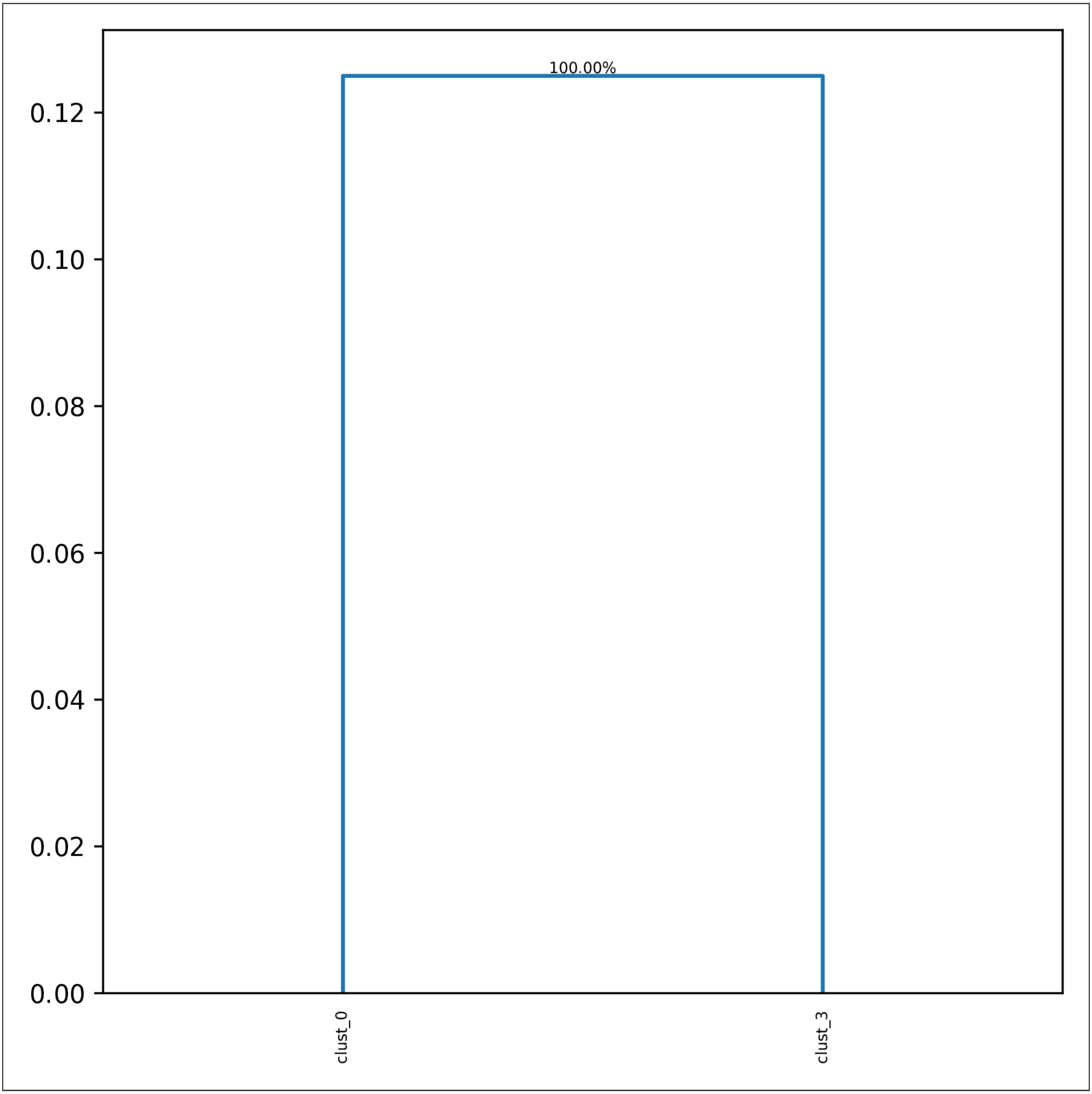

Sample: G0045 with 1 cells per cluster

Min. read depth: 10,  Mean UMIs per cluster:  23,908

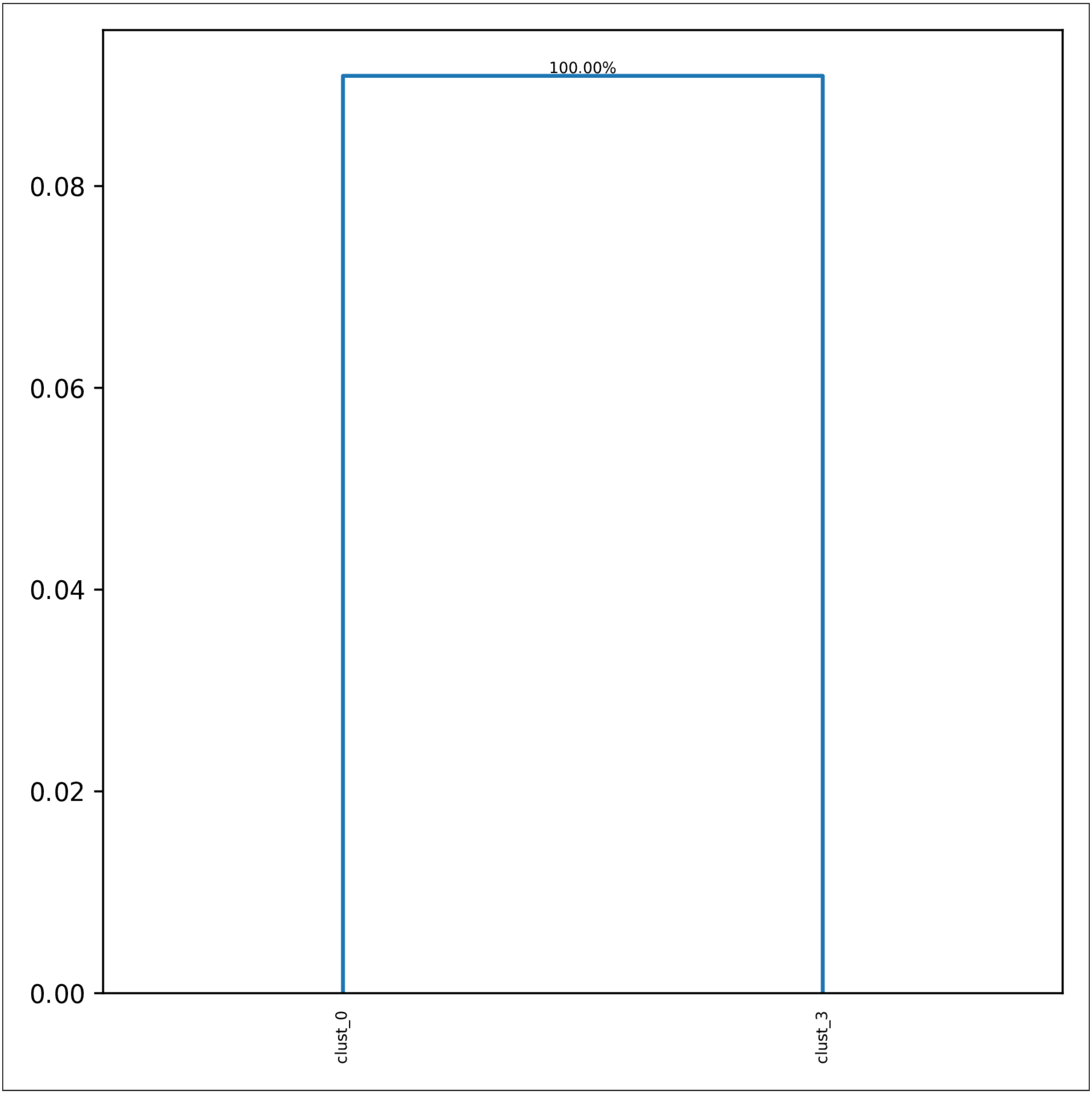

Sample: G0047 with all cells per cluster

Min. read depth: 10, Mean UMIs per cluster: 3,180,556

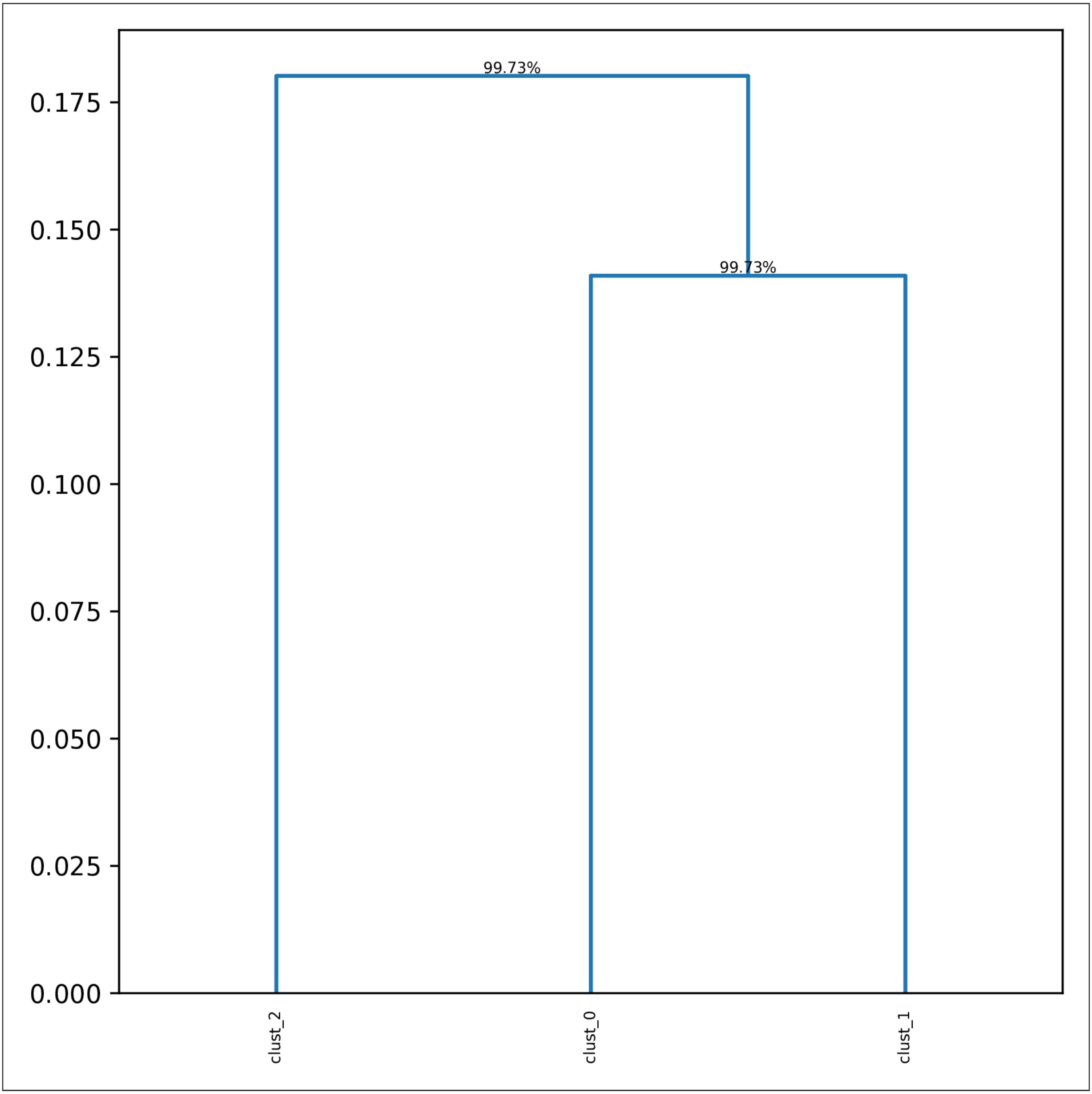

Sample: G0047 with 500 cells per cluster

Min. read depth: 10,  Mean UMIs per cluster:  2,853,721

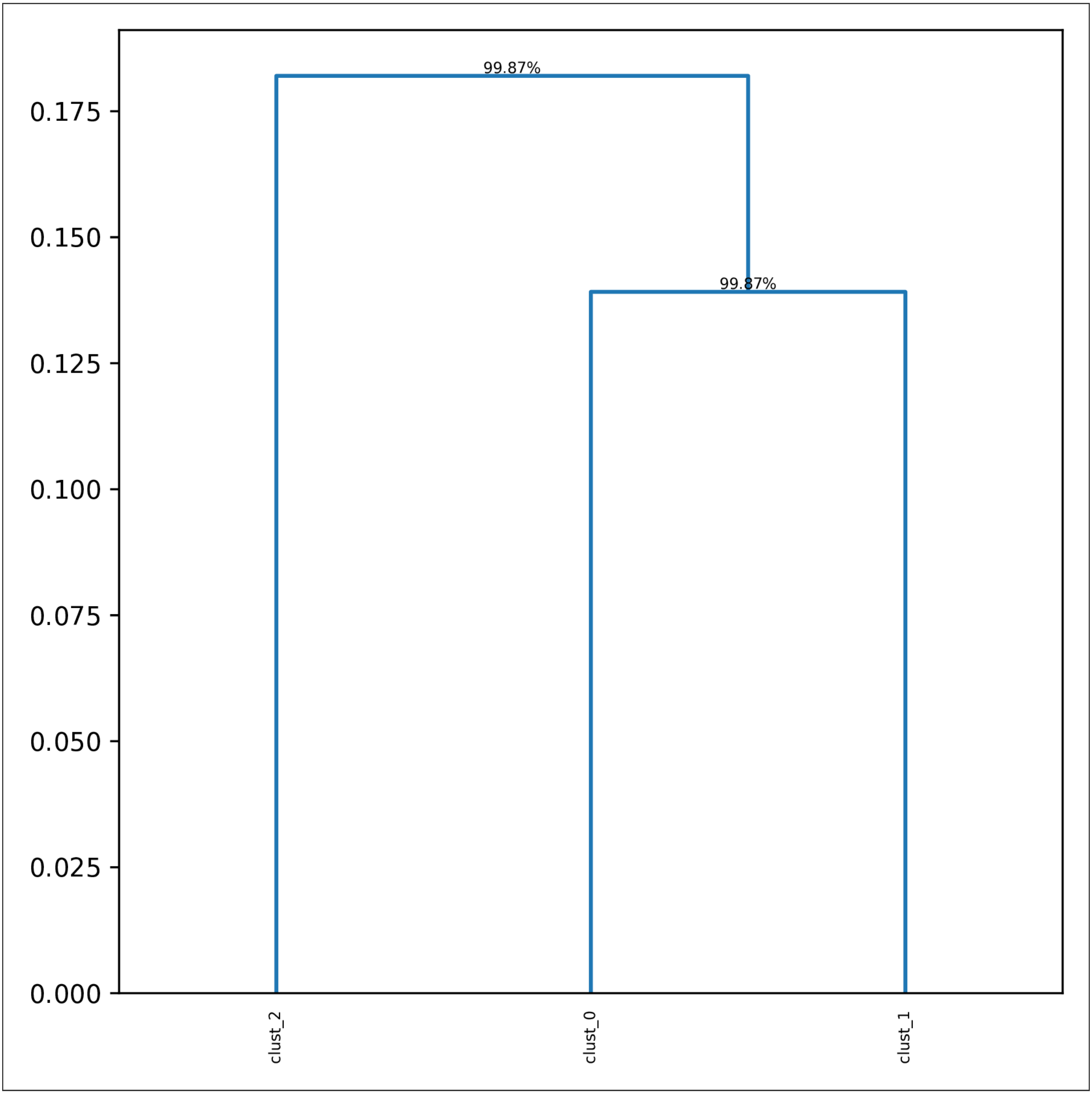

Sample: G0047 with 100 cells per cluster

Min. read depth: 10,  Mean UMIs per cluster:  807,294

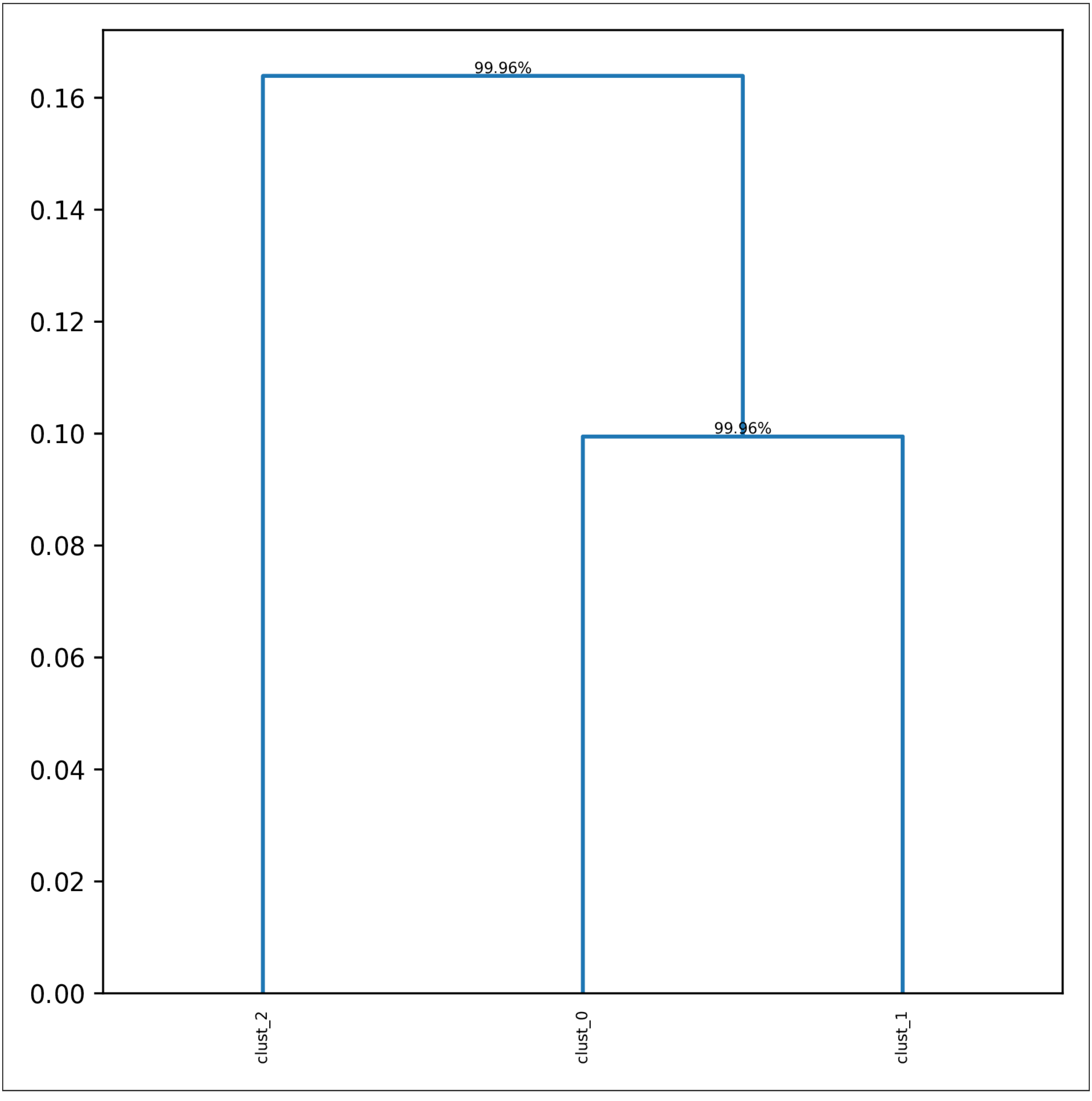

Sample: G0047 with 50 cells per cluster

Min. read depth: 10,  Mean UMIs per cluster:  408,013

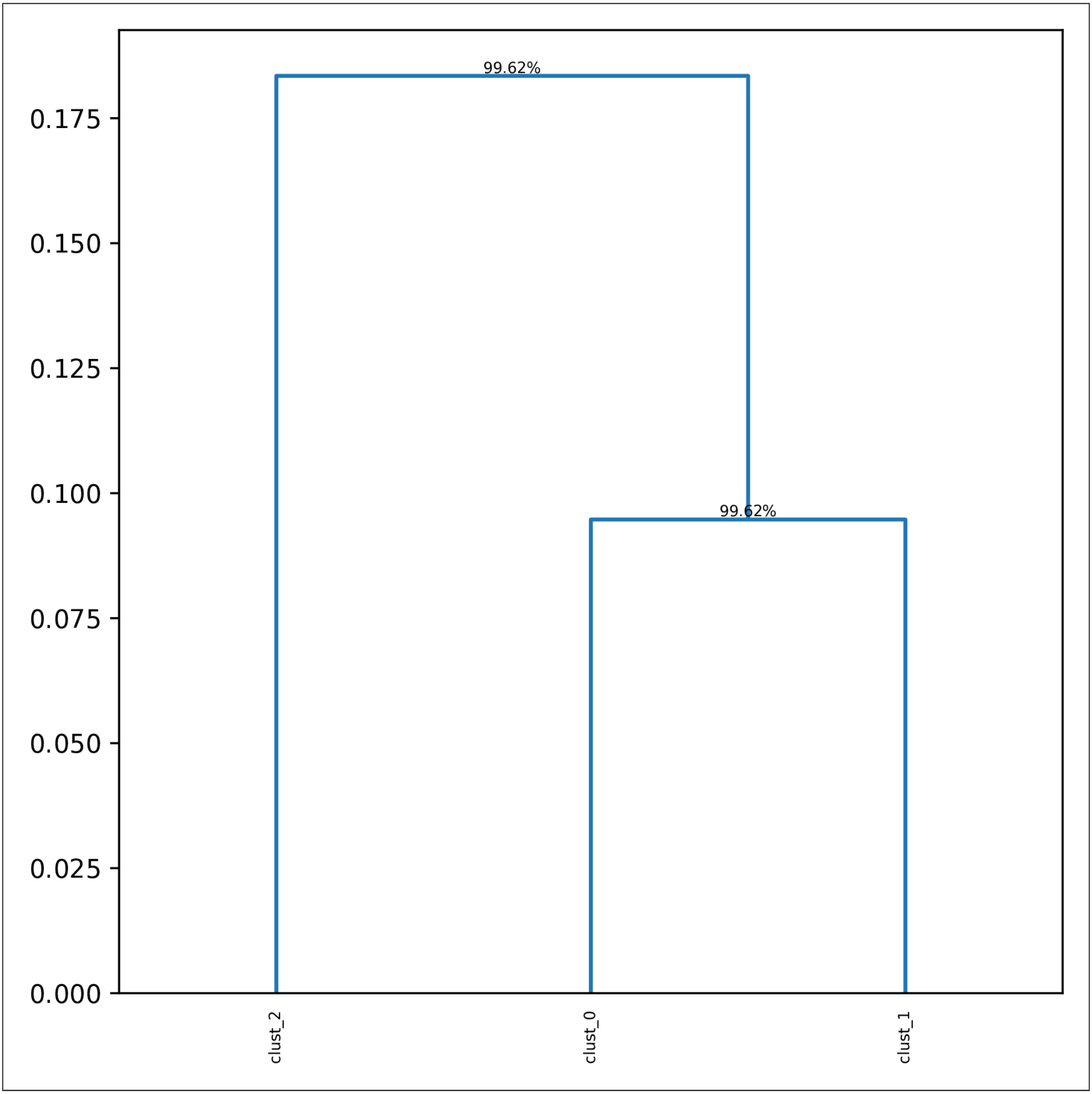

Sample: G0047 with 20 cells per cluster

Min. read depth: 10,  Mean UMIs per cluster:  170,632

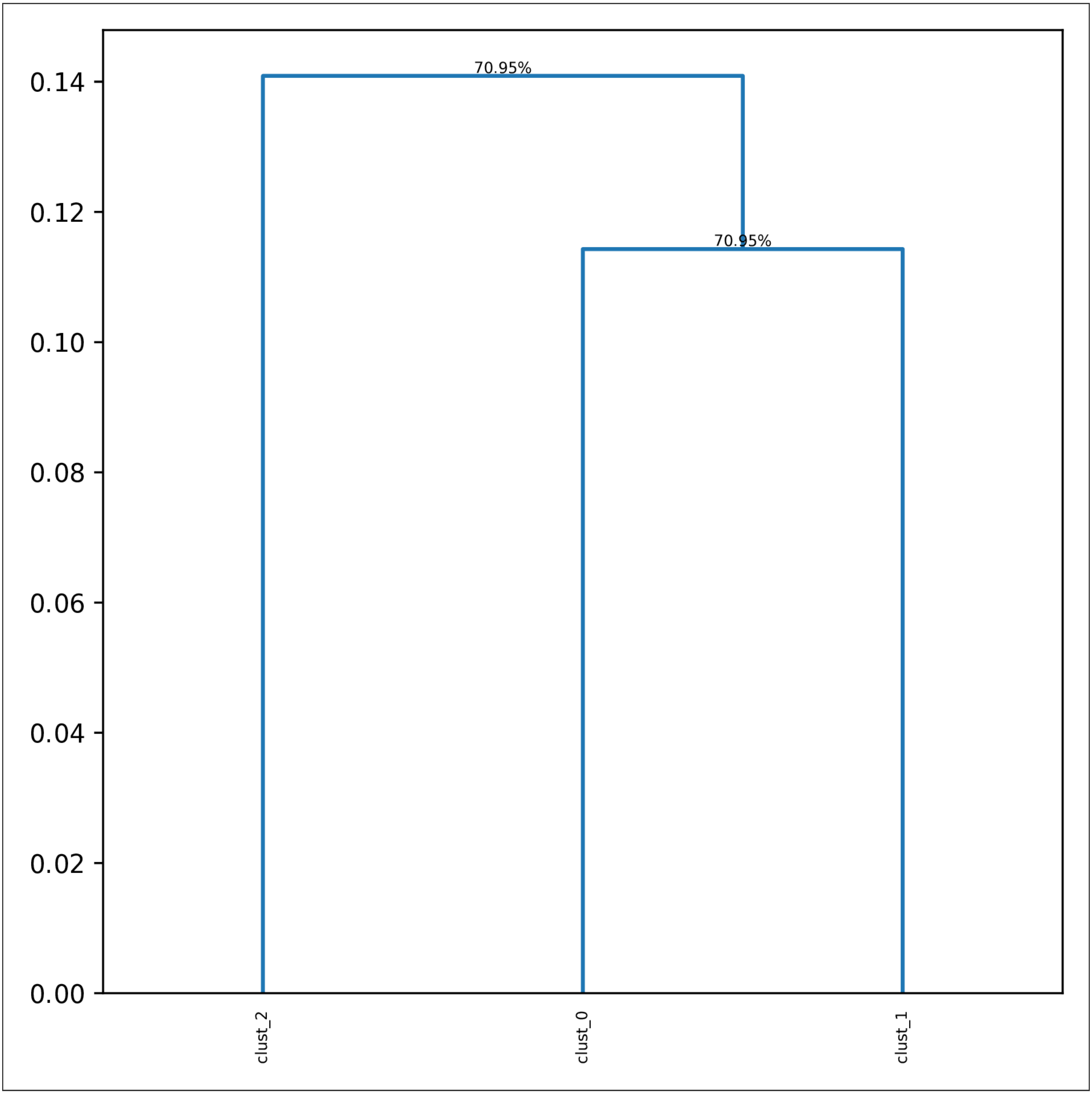

Sample: G0047 with 10 cells per cluster

Min. read depth: 10,  Mean UMIs per cluster:  86,243

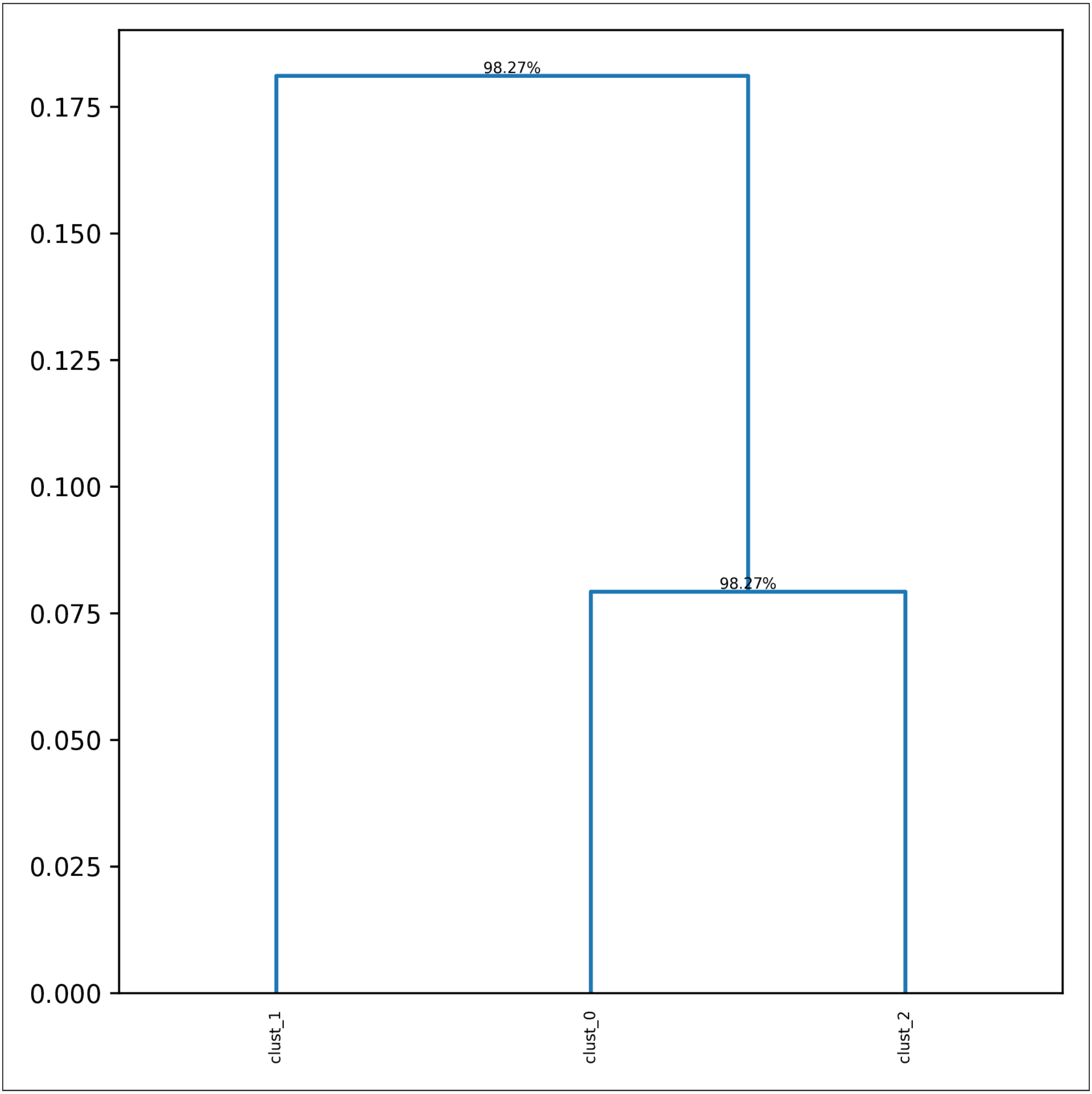

Sample: G0047 with 5 cells per cluster

Min. read depth: 10,  Mean UMIs per cluster:  39,890

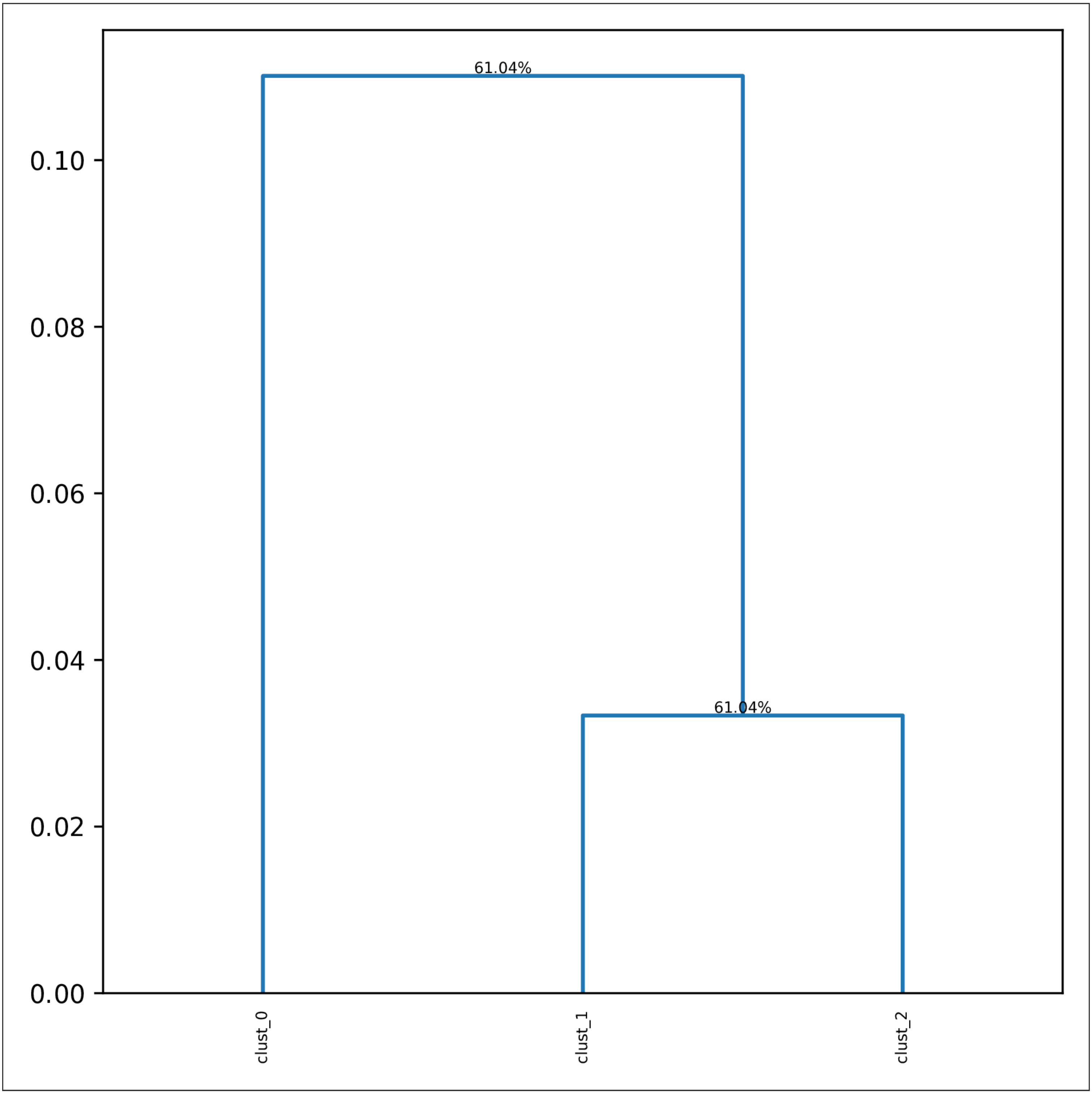

Sample: G0047 with 4 cells per cluster

Min. read depth: 10,  Mean UMIs per cluster:  32,847

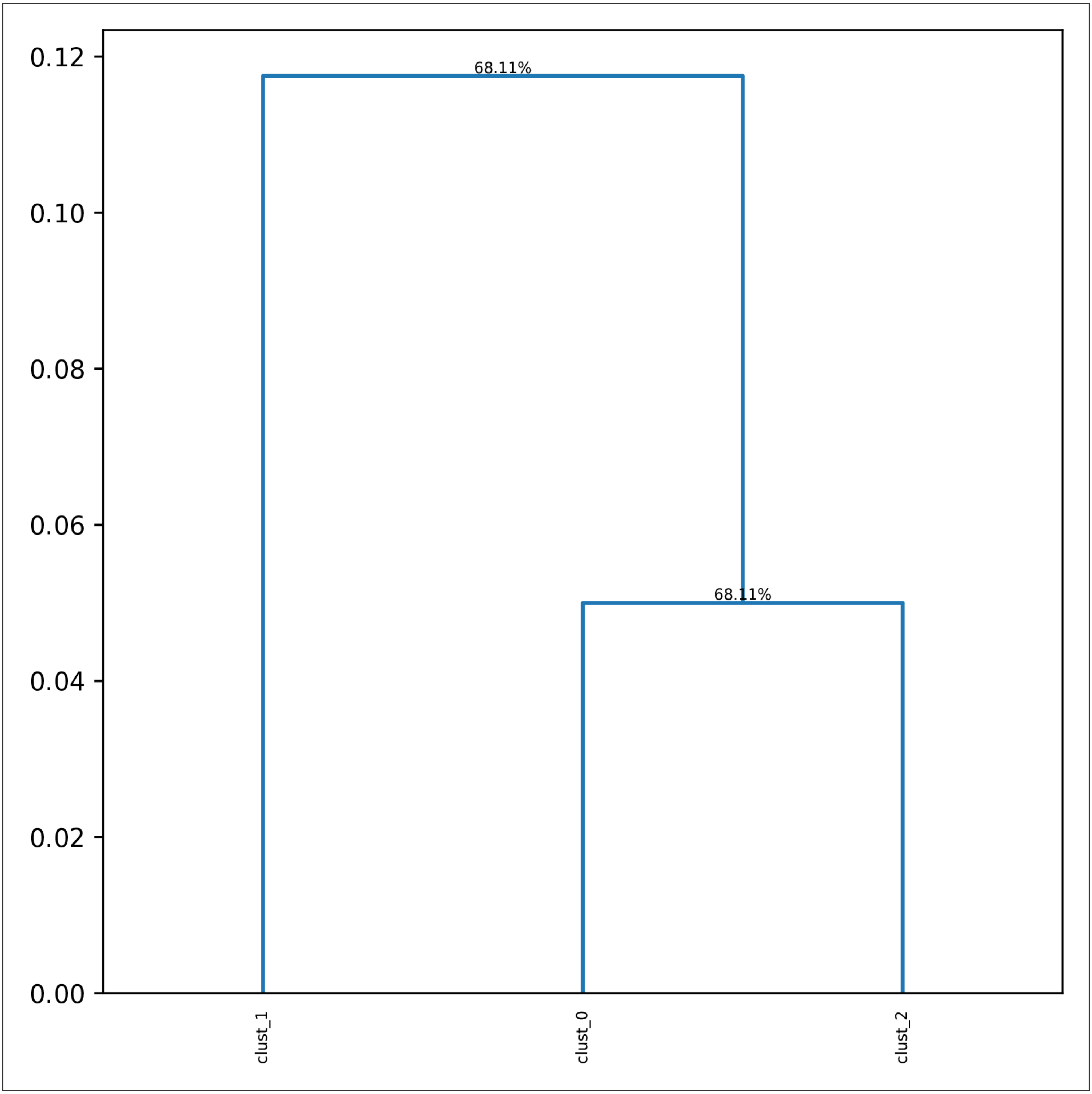

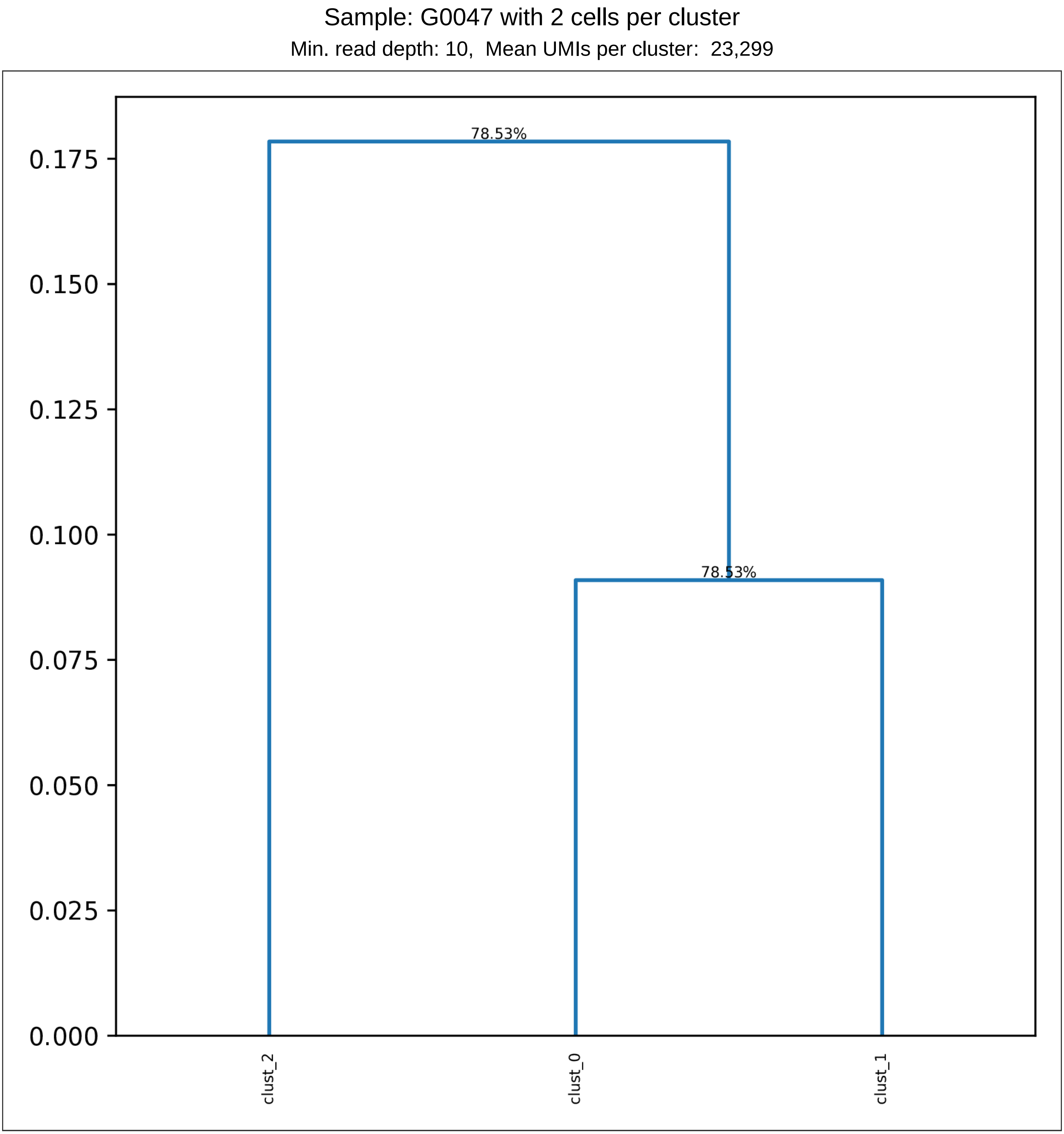

Sample: G0047 with 2 cells per cluster
Min. read depth: 10, Mean UMIs per cluster: 23,299
78.53%
78.53%
0.175
0.150
0.125
0.100
0.075
0.050
0.025
0.000
clust_2
clust_0
clust_1

Sample: G0047 with 1 cells per cluster

Min. read depth: 10,  Mean UMIs per cluster:  17,307

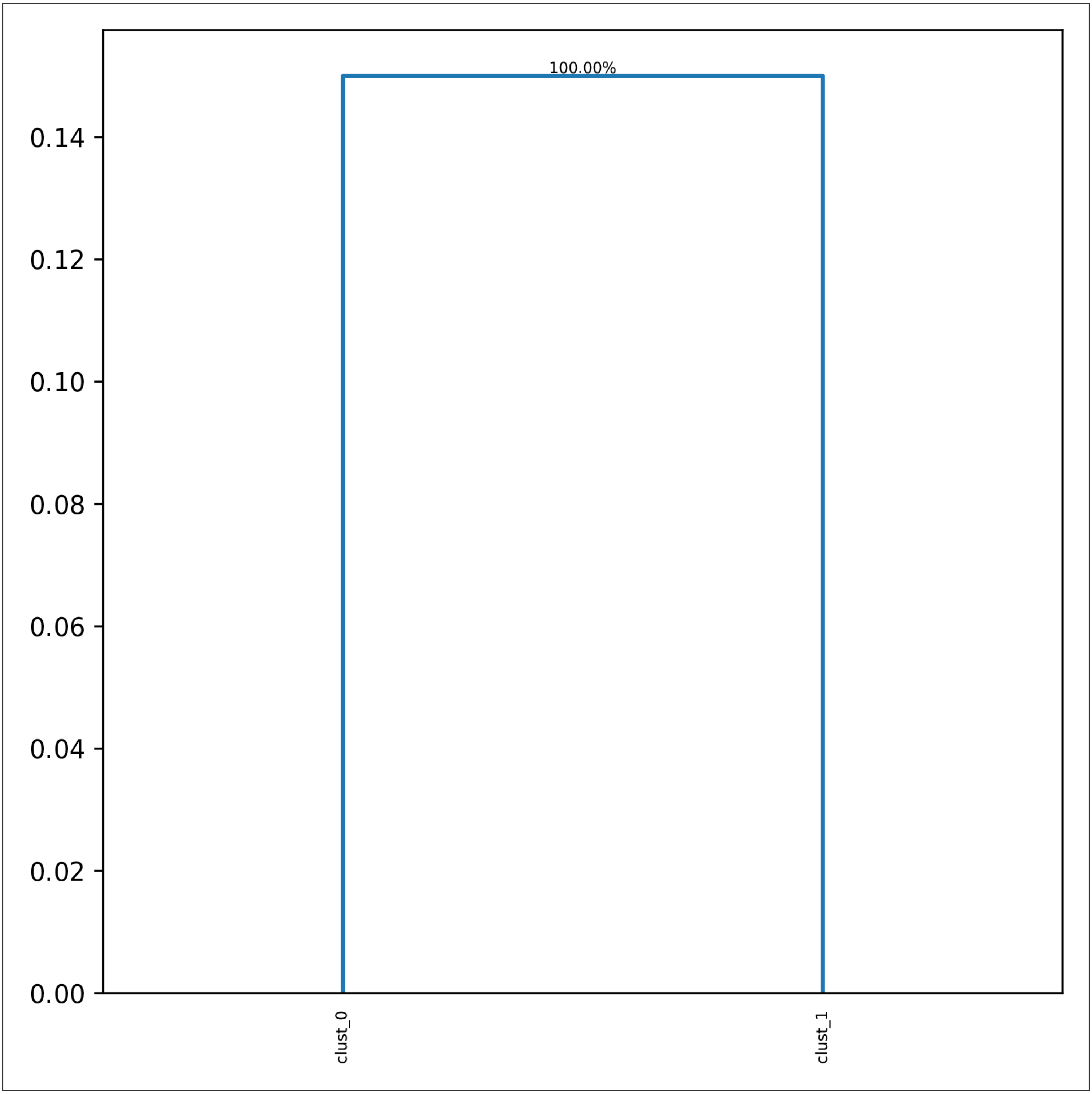

Sample: G0048 with all cells per cluster

Min. read depth: 10,  Mean UMIs per cluster:  12,414,360

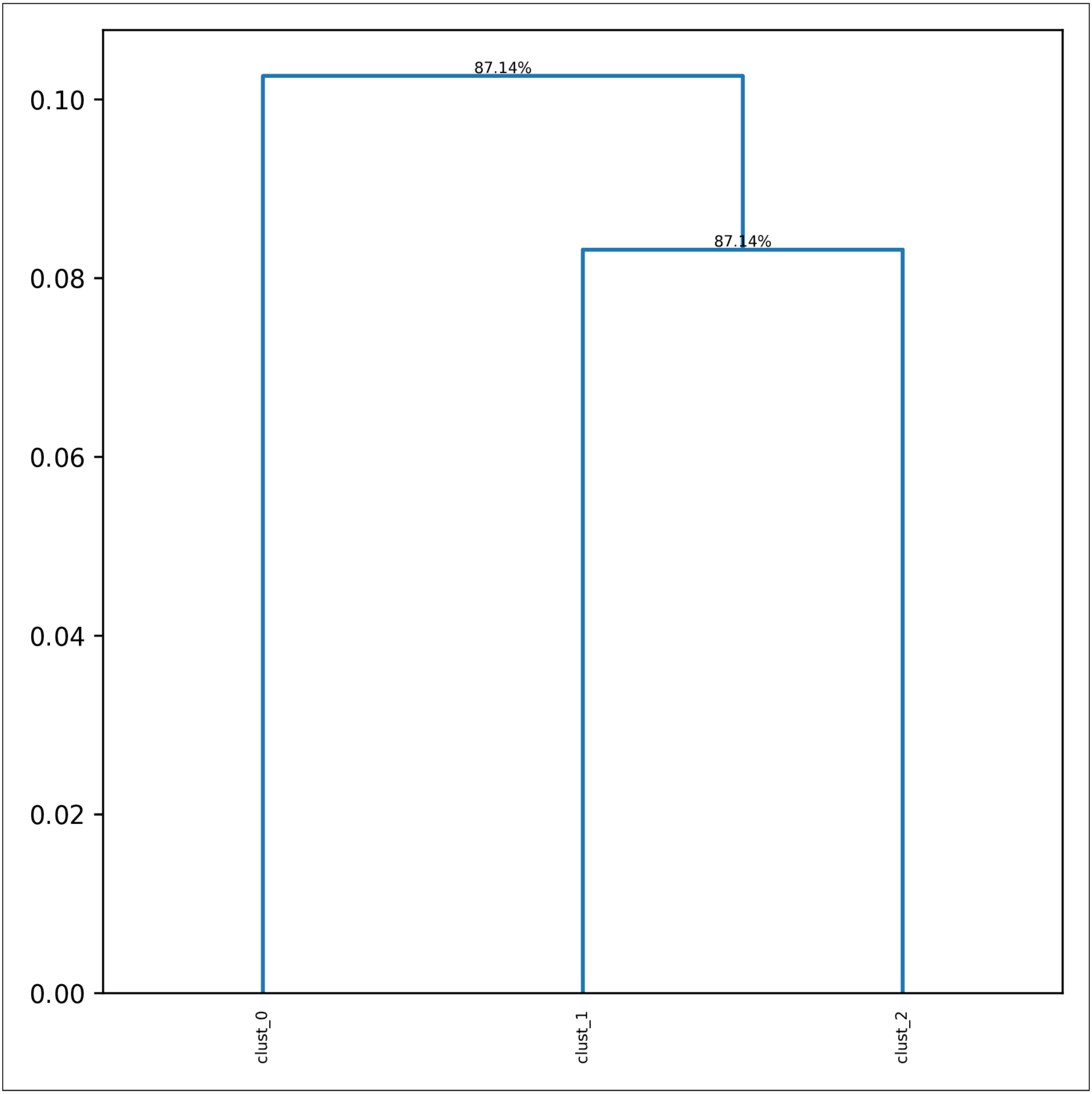

Sample: G0048 with 500 cells per cluster

Min. read depth: 10,  Mean UMIs per cluster:  2,600,307

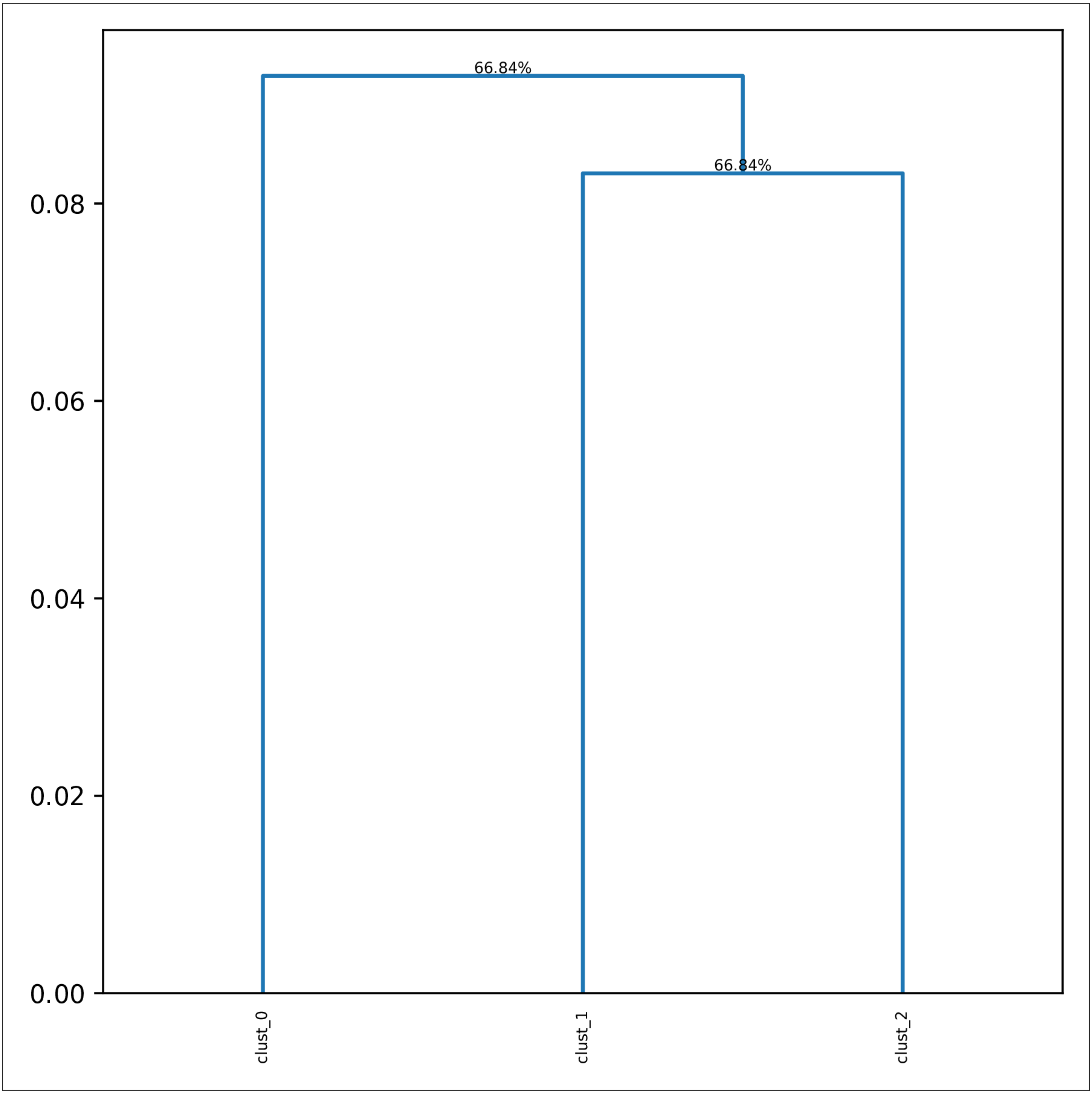

Sample: G0048 with 100 cells per cluster

Min. read depth: 10,  Mean UMIs per cluster:  632,663

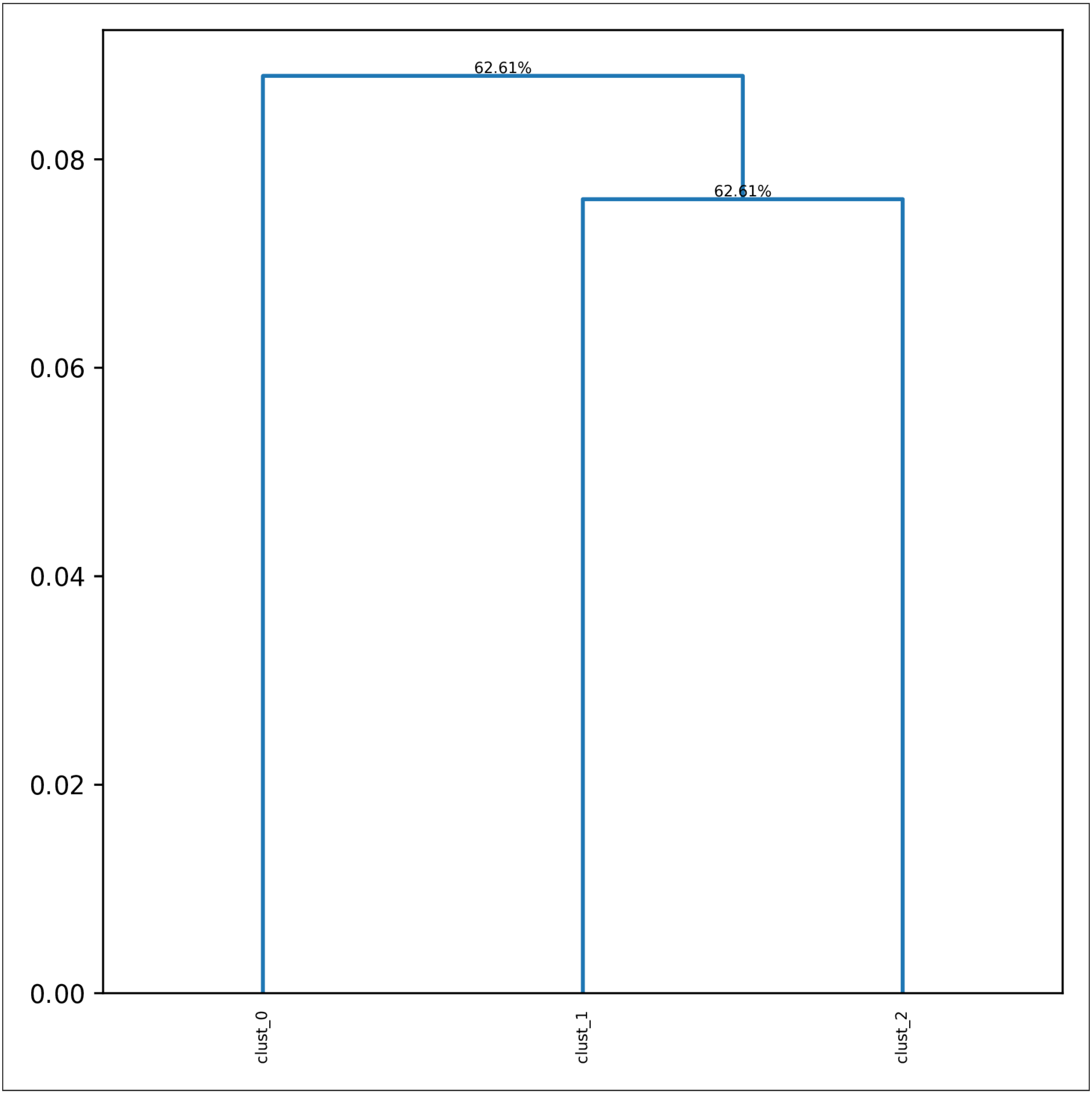

Sample: G0048 with 50 cells per cluster

Min. read depth: 10,  Mean UMIs per cluster:  312,948

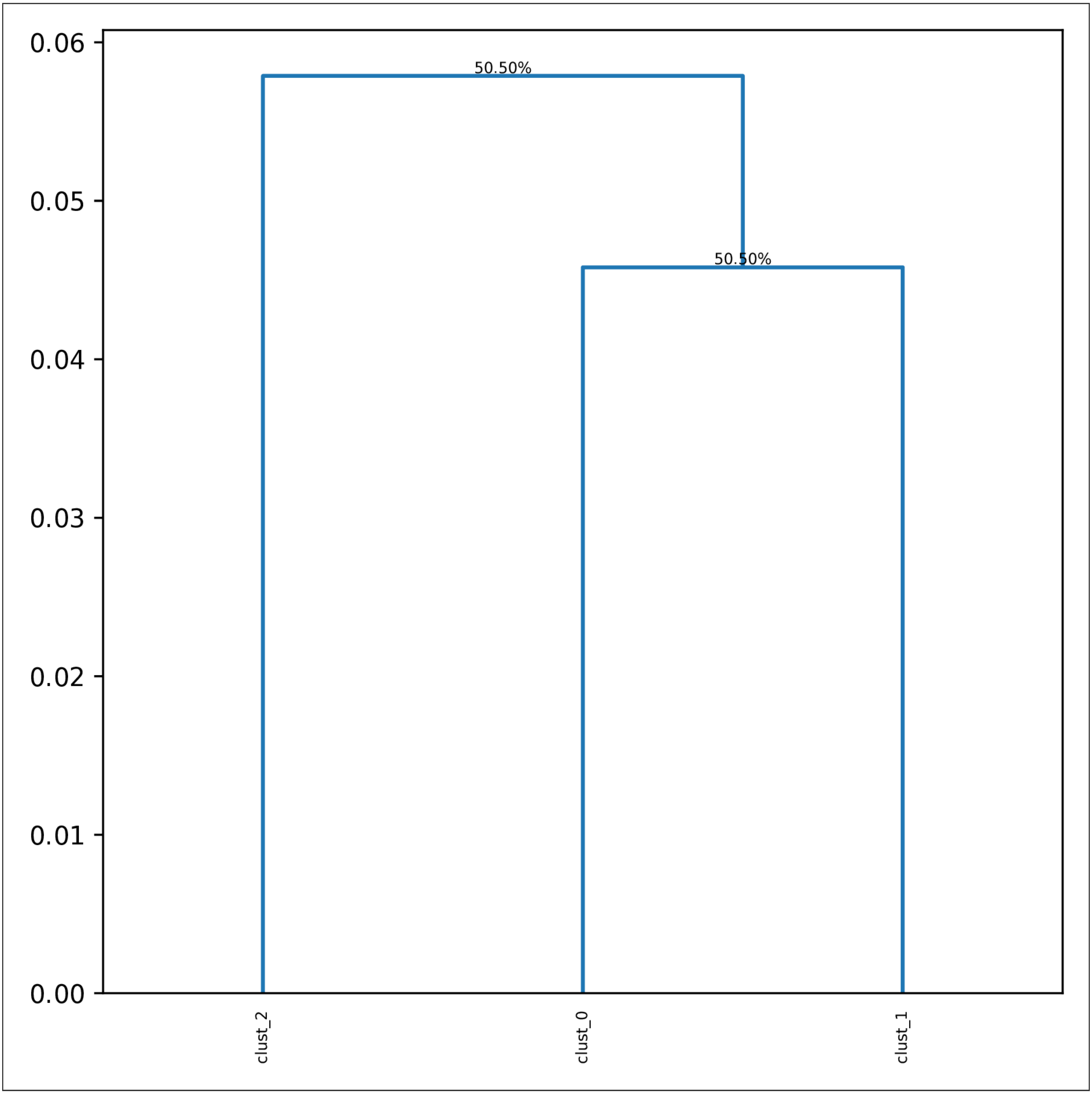

Sample: G0048 with 20 cells per cluster

Min. read depth: 10,  Mean UMIs per cluster:  122,002

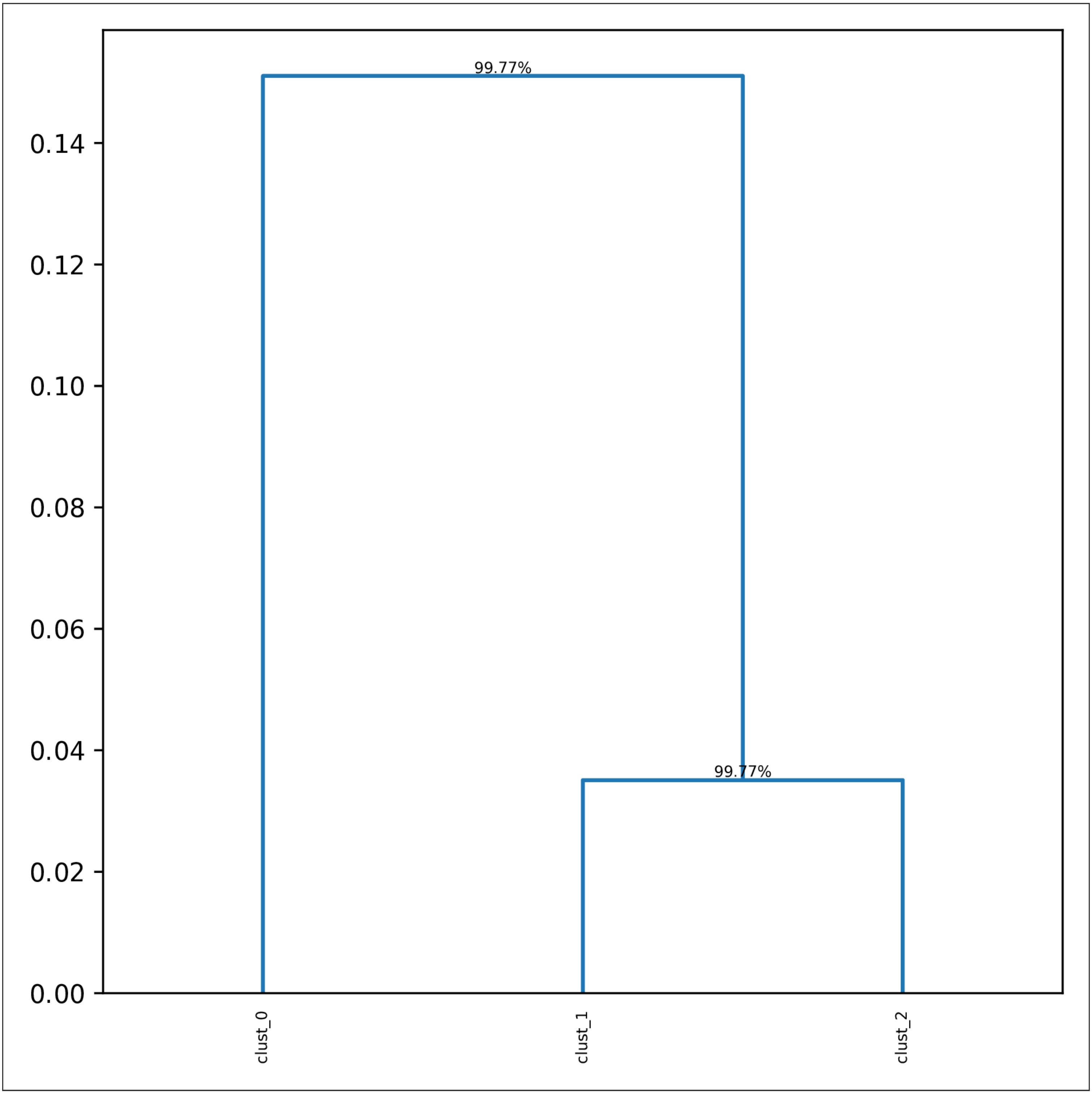

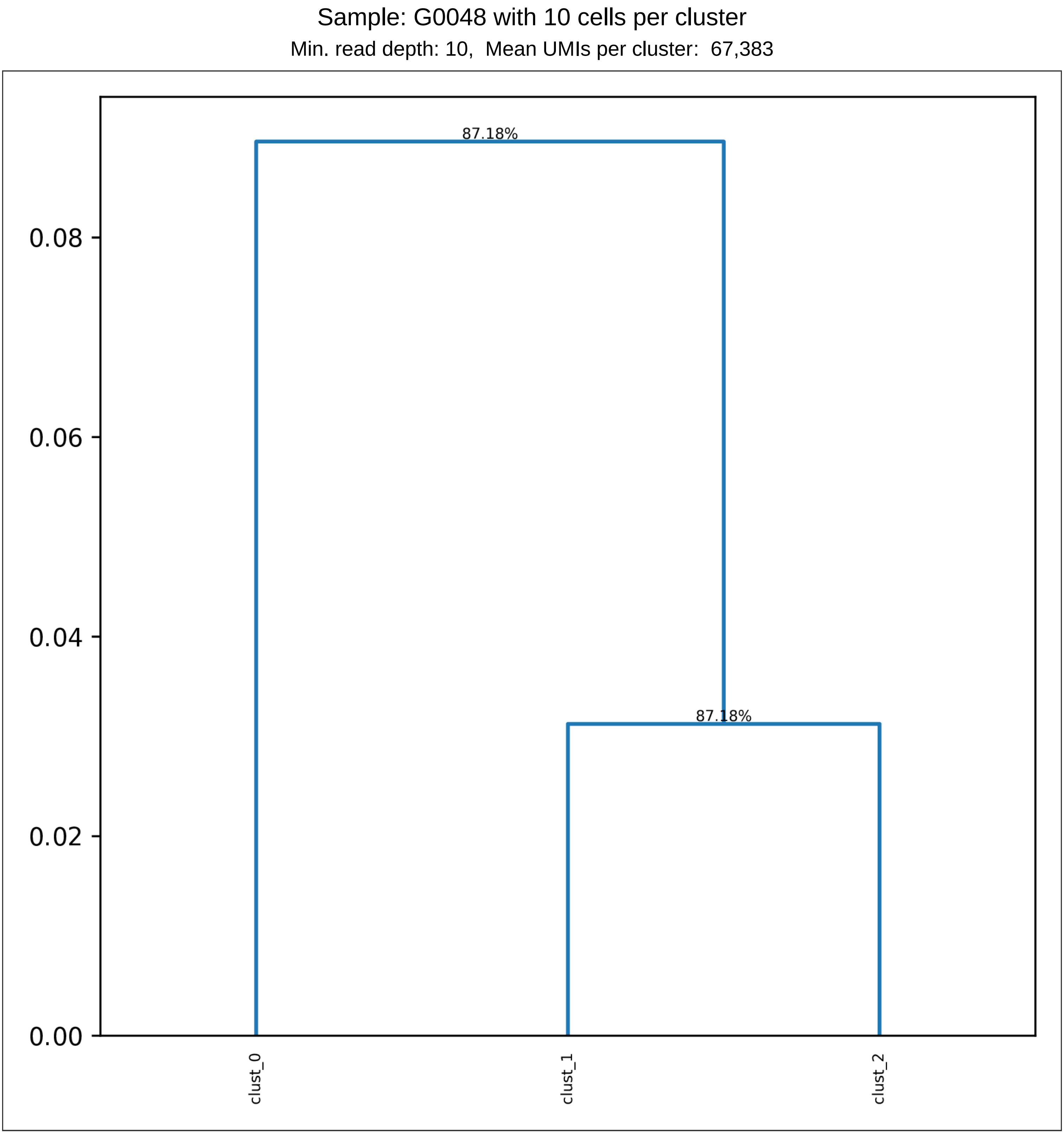

Sample: G0048 with 10 cells per cluster
Min. read depth: 10, Mean UMIs per cluster: 67,383
87.18%
87.18%
0.08
0.06
0.04
0.02
0.00
clust_0
clust_1
clust_2

Sample: G0048 with 5 cells per cluster

Min. read depth: 10,  Mean UMIs per cluster:  29,363

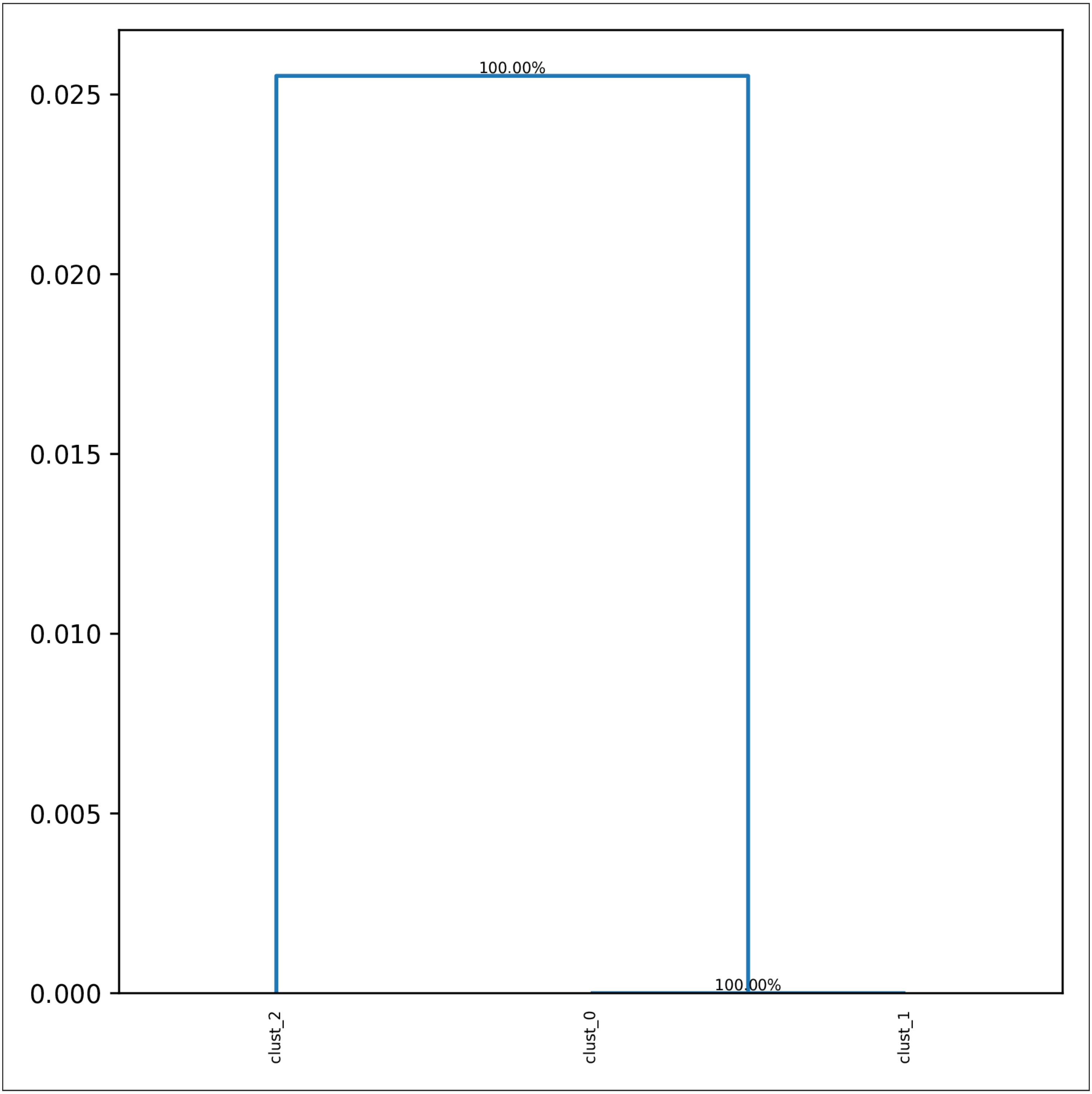

Sample: G0048 with 4 cells per cluster

Min. read depth: 10,  Mean UMIs per cluster:  22,305

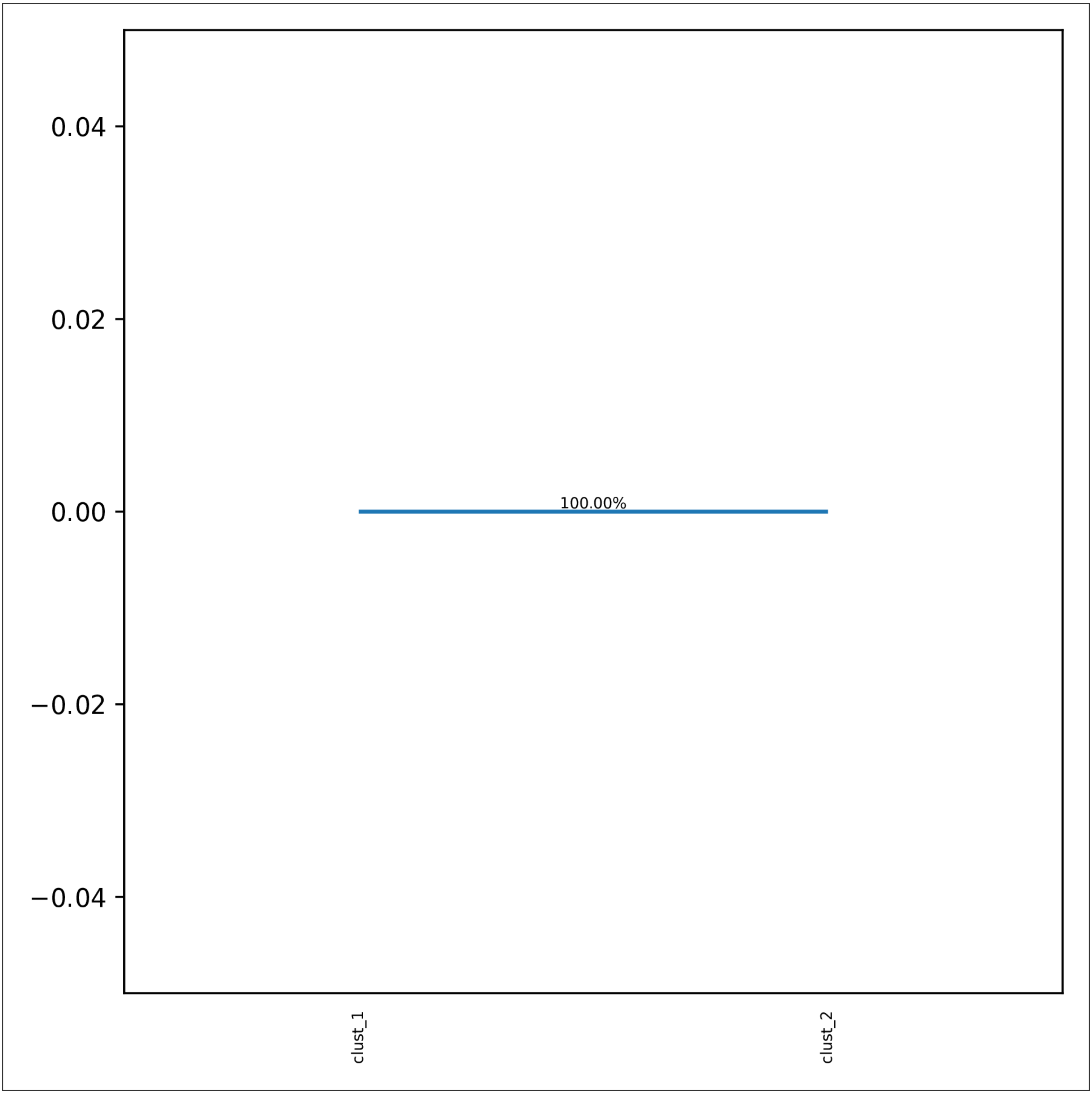

Sample: G0048 with 3 cells per cluster

Min. read depth: 10,  Mean UMIs per cluster:  17,488

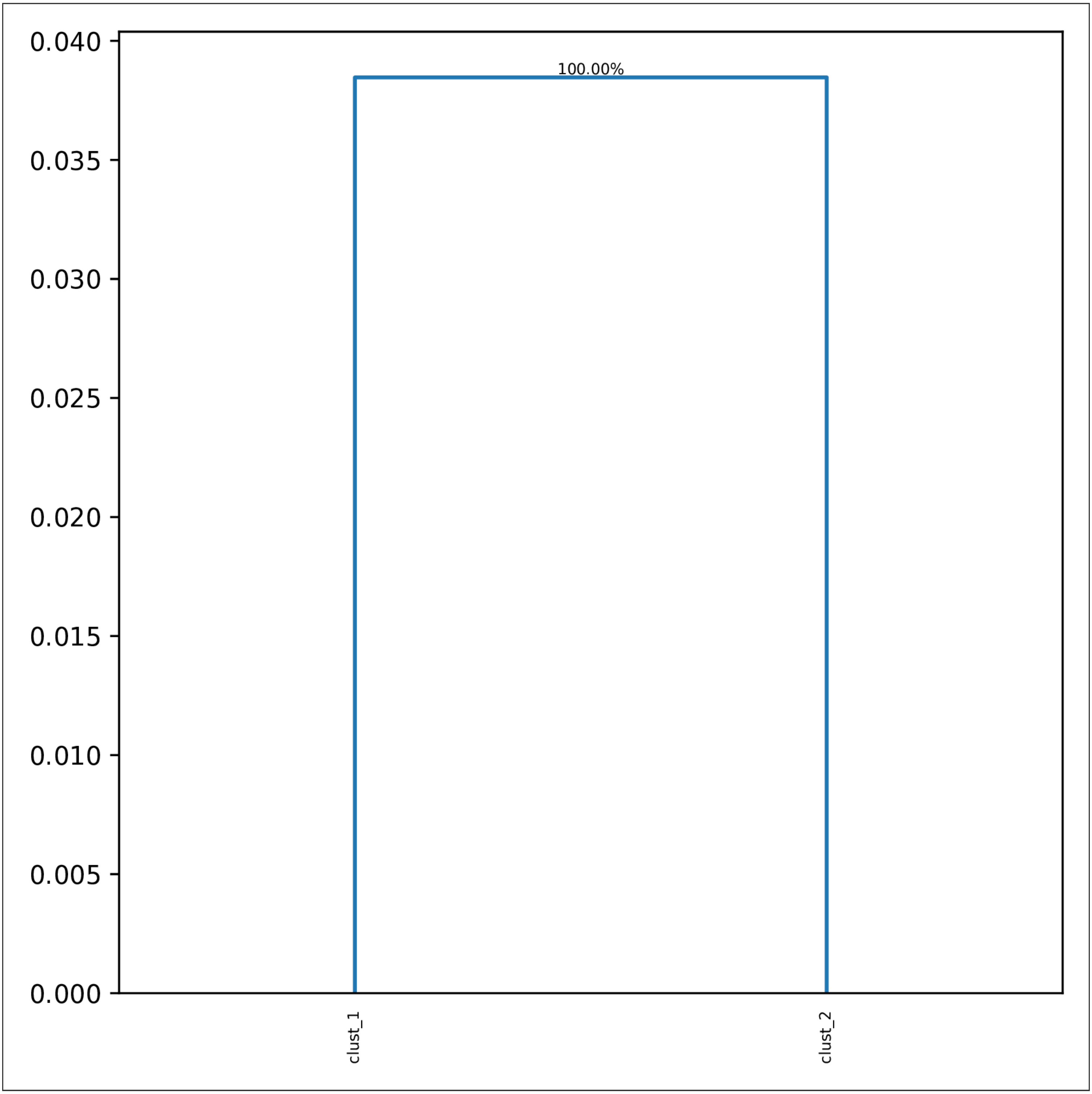

Sample: G0049 with all cells per cluster

Min. read depth: 10,  Mean UMIs per cluster:  11,757,352

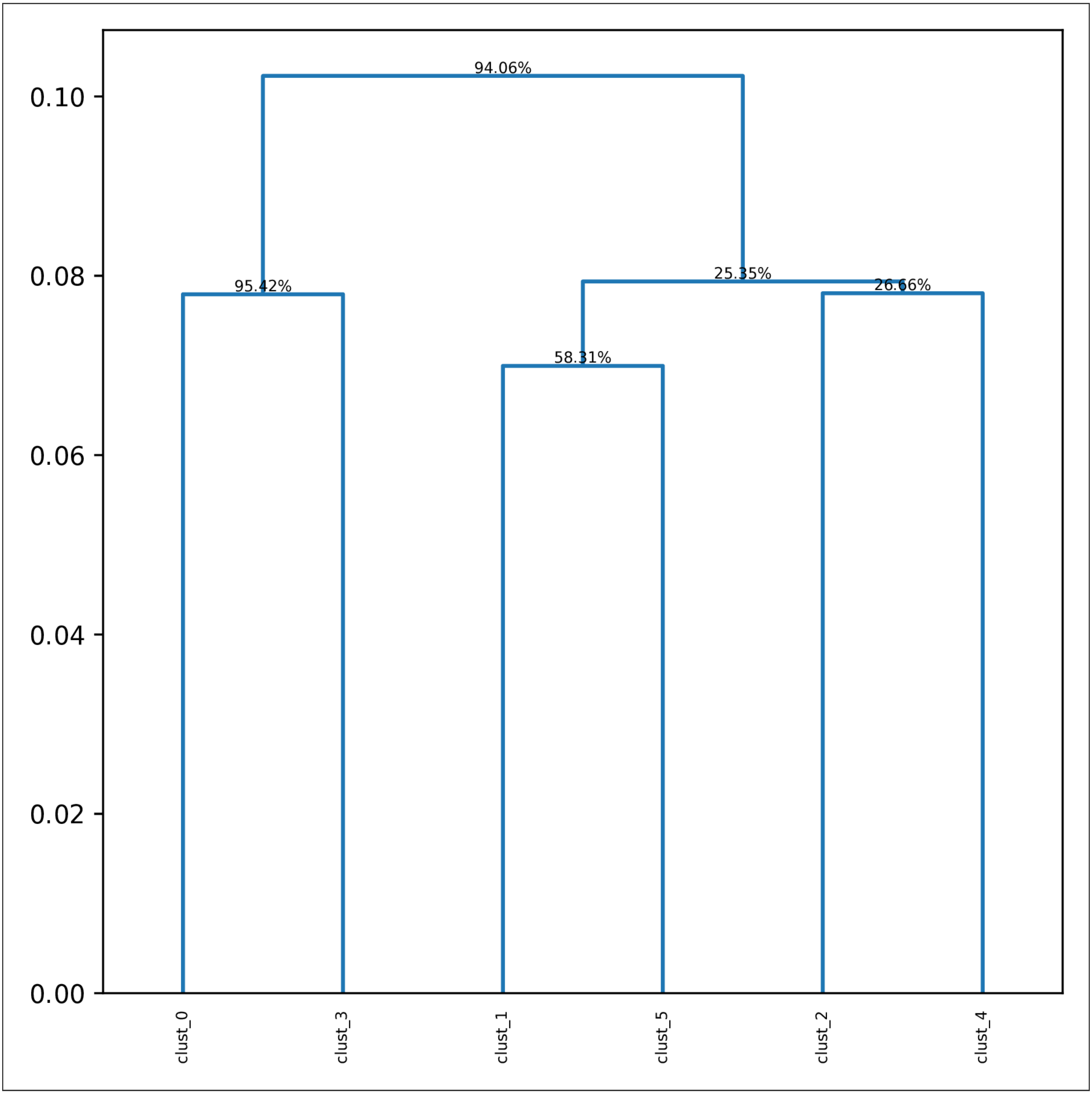

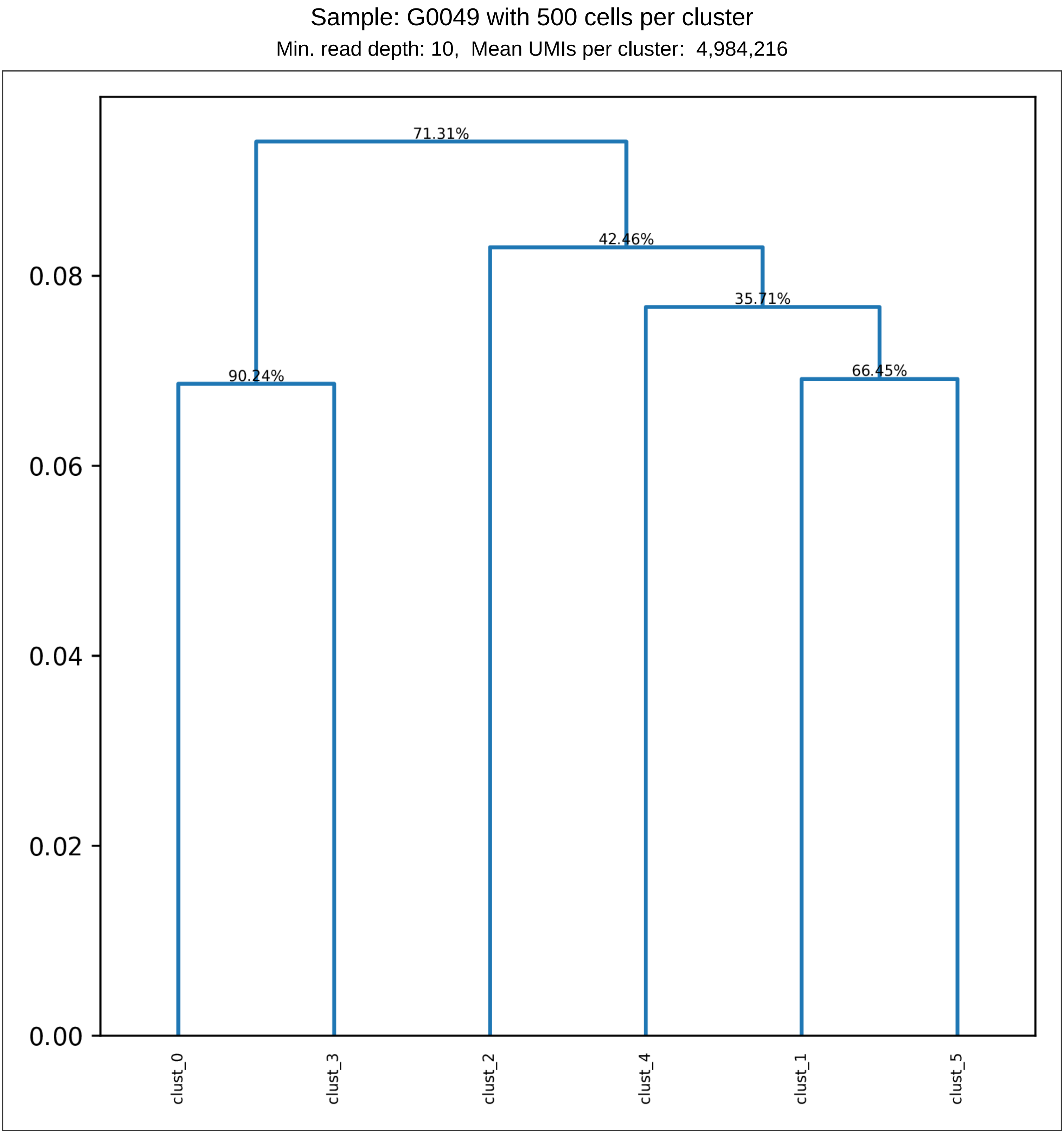
Sample: G0049 with 500 cells per cluster
Min. read depth: 10,  Mean UMIs per cluster:  4,984,216
71.31%
42.46%
35.71%
90.24%
66.45%
0.08
0.06
0.04
0.02
0.00
clust_0
clust_3
clust_2
clust_4
clust_1
clust_5

Sample: G0049 with 100 cells per cluster

Min. read depth: 10,  Mean UMIs per cluster:  1,084,215

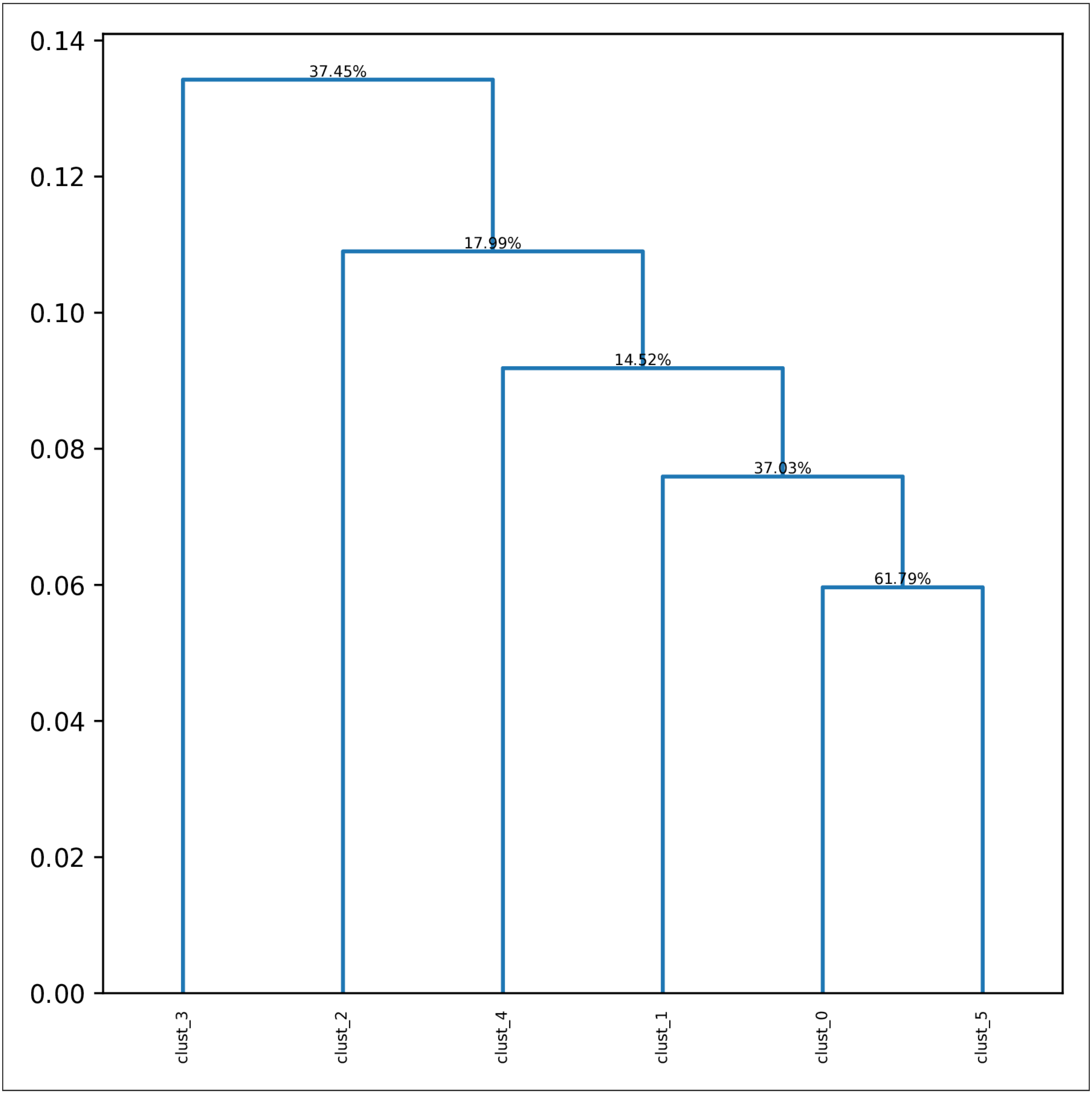

Sample: G0049 with 50 cells per cluster

Min. read depth: 10,  Mean UMIs per cluster:  583,154

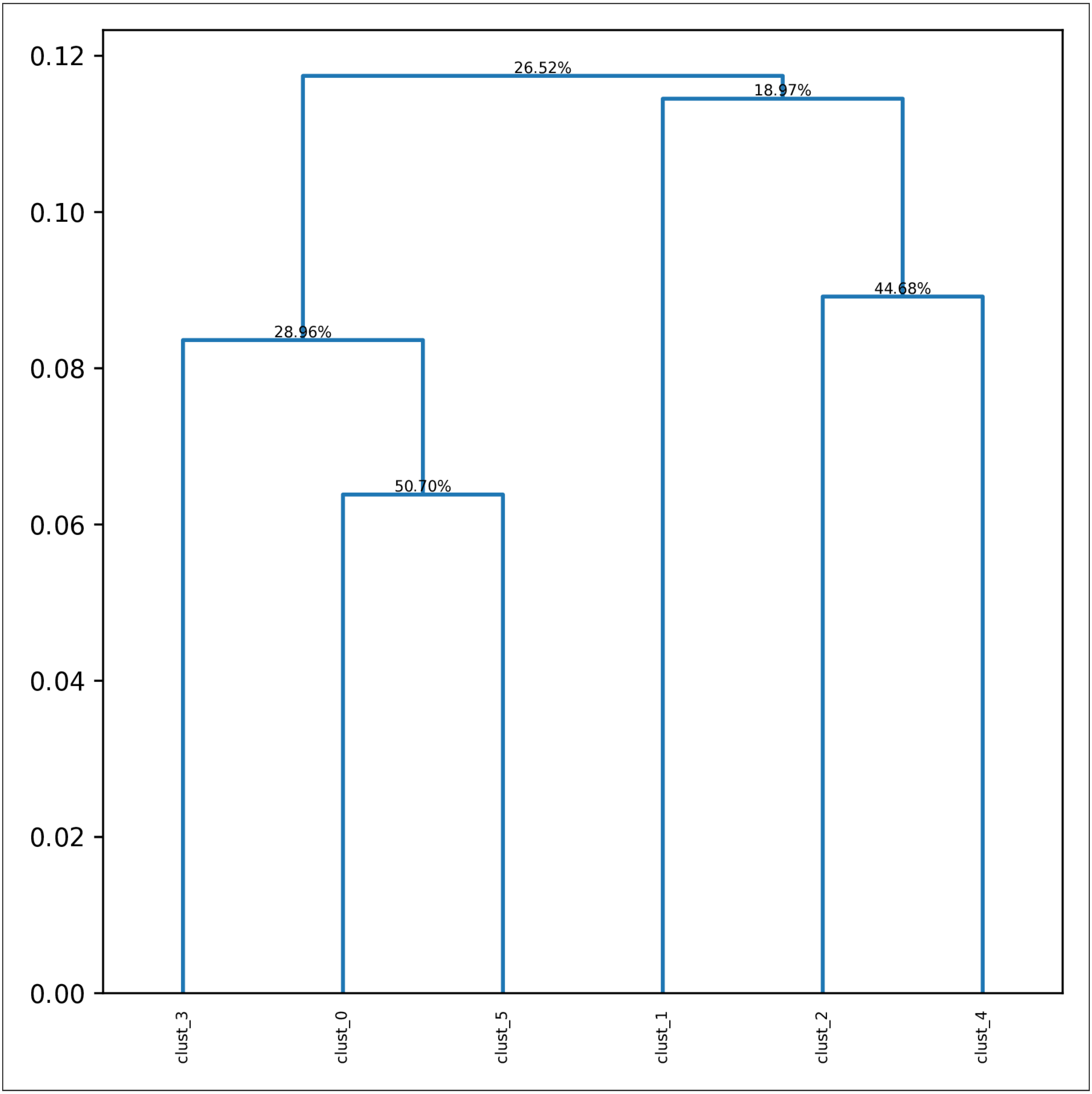

Sample: G0049 with 20 cells per cluster

Min. read depth: 10,  Mean UMIs per cluster:  220,866

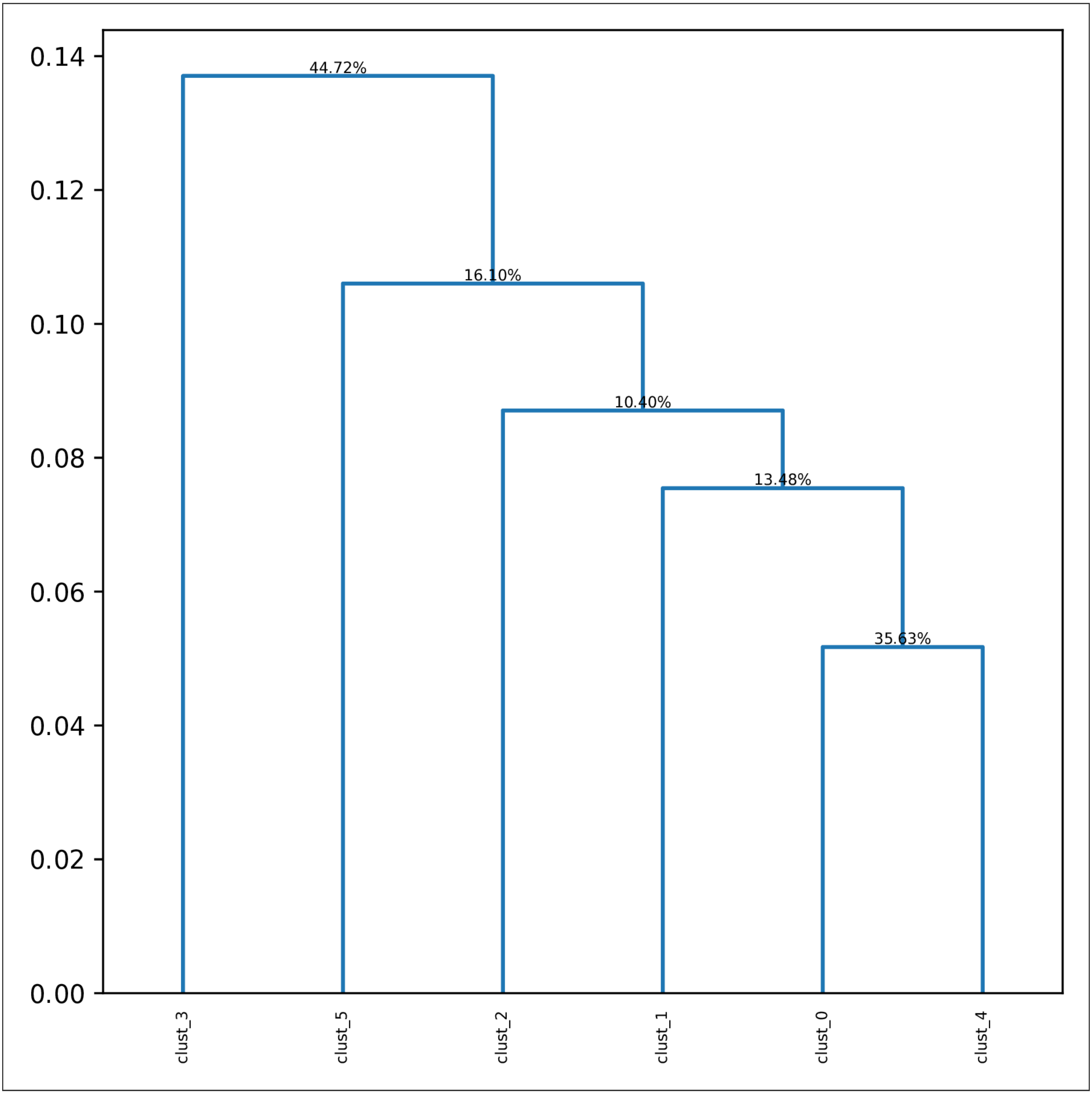

Sample: G0049 with 5 cells per cluster

Min. read depth: 10,  Mean UMIs per cluster:  67,240

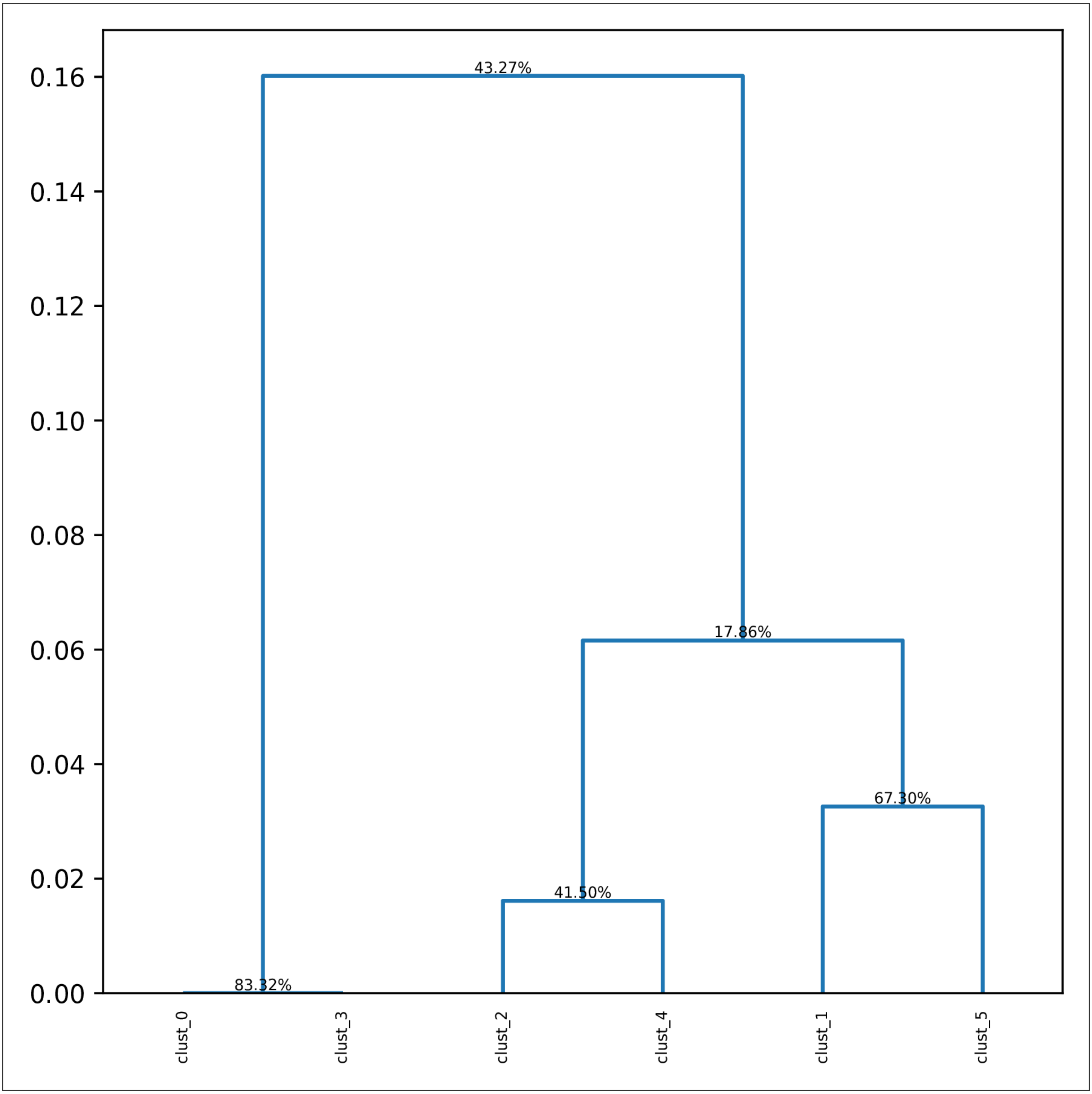

**Supplemental Figure 8: Hierarchical clustering trees from downsampled human patient data**

To determine how many cells or UMIs are needed to accurately distinguish genetically distinct cell populations, we randomly downsampled the human patient data. For each sample, we kept 1, 2, 3, 4, 5, 10, 20, 50, 100, 500, or all cells per transcriptional cluster for all clusters with a minimum of 500 cells in the intact dataset. Each downsampled dataset was analyzed with scanBit to produce a hierarchical clustering tree with bootstrap values. For each tree, we note the number of cells per cluster as well as the average number of UMIs per cluster for that sample.

### Supplemental Table 1: Sample metadata

| geo_id | sra_number | Sample_ID | description | sample_group | tumor_type | align_ref | subset_nCount_RNA_min | subset_nCount_RNA_max | subset_percent.mt_max | cluster_label |
|---|---|---|---|---|---|---|---|---|---|---|
| GSE210358 | SRR20761355 | G0001 | HMP11_A | tumor validation | Prostate cancer | /home/gdrobertslab/lab/GenRef/10x-hg38 | 1000 | 60000 | 25 | Prostate_pt_1 |
| GSE210358 | SRR20761356 | G0001 | HMP11_A | tumor validation | Prostate cancer | /home/gdrobertslab/lab/GenRef/10x-hg38 | 1000 | 60000 | 25 | Prostate_pt_1 |
| GSE210358 | SRR20761353 | G0002 | HMP11_B | tumor validation | Prostate cancer | /home/gdrobertslab/lab/GenRef/10x-hg38 | 1000 | 60000 | 25 | Prostate_pt_2 |
| GSE210358 | SRR20761354 | G0002 | HMP11_B | tumor validation | Prostate cancer | /home/gdrobertslab/lab/GenRef/10x-hg38 | 1000 | 60000 | 25 | Prostate_pt_2 |
| GSE222703 | SRR23063397 | G0003 | ccRCC tumor - patient 1 | tumor validation | Clear cell carcinoma of the kidney | /home/gdrobertslab/lab/GenRef/10x-hg38 | 1000 | 75000 | 30 | RCC_pt_1 |
| GSE222703 | SRR23063398 | G0003 | ccRCC tumor - patient 1 | tumor validation | Clear cell carcinoma of the kidney | /home/gdrobertslab/lab/GenRef/10x-hg38 | 1000 | 75000 | 30 | RCC_pt_1 |
| GSE222703 | SRR23063395 | G0004 | adjacent kidney tissue - patient 1 | healthy samples | healthy | /home/gdrobertslab/lab/GenRef/10x-hg38 | 1000 | 75000 | 30 | RCC_pt_2 |
| GSE222703 | SRR23063396 | G0004 | adjacent kidney tissue - patient 1 | healthy samples | healthy | /home/gdrobertslab/lab/GenRef/10x-hg38 | 1000 | 75000 | 30 | RCC_pt_2 |
| GSE247228 | SRR26705626 | G0005 | YUMM3_3-ActA_GFP_1 | truthset validation | Melanoma | custom_ref/gfp_rfp_mm10 | 1000 | 75000 | 30 | GFP_melanoma_1 |
| GSE247228 | SRR26705627 | G0006 | YUMM3_3-ActA_GFP_2 | truthset validation | Melanoma | custom_ref/gfp_rfp_mm10 | 1000 | 100000 | 30 | GFP_melanoma_2 |
| GSE247228 | SRR26705629 | G0007 | YUMM3_3-ActA_GFP_3 | truthset validation | Melanoma | custom_ref/gfp_rfp_mm10 | 1000 | 100000 | 30 | GFP_melanoma_3 |
| GSE247228 | SRR26705628 | G0008 | YUMM3_3-CTRL_GFP_1 | truthset validation | Melanoma | custom_ref/gfp_rfp_mm10 | 1000 | 80000 | 20 | GFP_melanoma_4 |
| GSE247228 | SRR26705630 | G0009 | YUMM3_3-CTRL_GFP_2 | truthset validation | Melanoma | custom_ref/gfp_rfp_mm10 | 1000 | 75000 | 20 | GFP_melanoma_5 |
| GSE247228 | SRR26705631 | G0010 | YUMM3_3-CTRL_GFP_3 | truthset validation | Melanoma | custom_ref/gfp_rfp_mm10 | 1000 | 60000 | 20 | GFP_melanoma_6 |
| GSE201615 | SRR18932566 | G0011 | Replicate 1, mRNA-derived cDNA | truthset validation | undifferentiated pleiomorphic sarcoma | custom_ref/gfp_rfp_mm10 | 1000 | 60000 | 20 | DsReD_sarcoma_1 |
| GSE201615 | SRR18932567 | G0011 | Replicate 1, mRNA-derived cDNA | truthset validation | undifferentiated pleiomorphic sarcoma | custom_ref/gfp_rfp_mm10 | 1000 | 60000 | 20 | DsReD_sarcoma_1 |
| GSE201615 | SRR18932562 | G0012 | Replicate 2, mRNA-derived cDNA | truthset validation | undifferentiated pleiomorphic sarcoma | custom_ref/gfp_rfp_mm10 | 1000 | 60000 | 20 | DsReD_sarcoma_2 |
| GSE201615 | SRR18932563 | G0012 | Replicate 2, mRNA-derived cDNA | truthset validation | undifferentiated pleiomorphic sarcoma | custom_ref/gfp_rfp_mm10 | 1000 | 60000 | 20 | DsReD_sarcoma_2 |
| GSE162454 | SRR13177101 | G0013 | GSE162454: Osteosarcoma GSM4952363: Patient 1 | tumor validation | Osteosarcoma | /home/gdrobertslab/lab/GenRef/10x-hg38 | 1000 | 140000 | 30 | Osteo_pt_1 |
| GSE162454 | SRR13177102 | G0013 | GSE162454: Osteosarcoma GSM4952363: Patient 1 | tumor validation | Osteosarcoma | /home/gdrobertslab/lab/GenRef/10x-hg38 | 1000 | 140000 | 30 | Osteo_pt_1 |
| GSE162454 | SRR13177103 | G0013 | GSE162454: Osteosarcoma GSM4952363: Patient 1 | tumor validation | Osteosarcoma | /home/gdrobertslab/lab/GenRef/10x-hg38 | 1000 | 140000 | 30 | Osteo_pt_1 |
| GSE162454 | SRR13177104 | G0013 | GSE162454: Osteosarcoma GSM4952363: Patient 1 | tumor validation | Osteosarcoma | /home/gdrobertslab/lab/GenRef/10x-hg38 | 1000 | 140000 | 30 | Osteo_pt_1 |
| GSE162454 | SRR13177105 | G0014 | GSE162454: Osteosarcoma GSM4952364: Patient 2 | tumor validation | Osteosarcoma | /home/gdrobertslab/lab/GenRef/10x-hg38 | 1000 | 100000 | 40 | Osteo_pt_2 |
| GSE162454 | SRR13177106 | G0014 | GSE162454: Osteosarcoma GSM4952364: Patient 2 | tumor validation | Osteosarcoma | /home/gdrobertslab/lab/GenRef/10x-hg38 | 1000 | 100000 | 40 | Osteo_pt_2 |
| GSE162454 | SRR13177107 | G0014 | GSE162454: Osteosarcoma GSM4952364: Patient 2 | tumor validation | Osteosarcoma | /home/gdrobertslab/lab/GenRef/10x-hg38 | 1000 | 100000 | 40 | Osteo_pt_2 |
| GSE162454 | SRR13177108 | G0014 | GSE162454: Osteosarcoma GSM4952364: Patient 2 | tumor validation | Osteosarcoma | /home/gdrobertslab/lab/GenRef/10x-hg38 | 1000 | 100000 | 40 | Osteo_pt_2 |
| GSE162454 | SRR13177109 | G0015 | GSE162454: Osteosarcoma GSM4952365: Patient 3 | tumor validation | Osteosarcoma | /home/gdrobertslab/lab/GenRef/10x-hg38 | 1000 | 50000 | 40 | Osteo_pt_3 |
| GSE162454 | SRR13927112 | G0016 | GSE162454: Osteosarcoma GSM5155198: Patient 4 | tumor validation | Osteosarcoma | /home/gdrobertslab/lab/GenRef/10x-hg38 | 1000 | 120000 | 30 | Osteo_pt_4 |
| GSE162454 | SRR13927113 | G0016 | GSE162454: Osteosarcoma GSM5155198: Patient 4 | tumor validation | Osteosarcoma | /home/gdrobertslab/lab/GenRef/10x-hg38 | 1000 | 120000 | 30 | Osteo_pt_4 |
| GSE162454 | SRR13927114 | G0016 | GSE162454: Osteosarcoma GSM5155198: Patient 4 | tumor validation | Osteosarcoma | /home/gdrobertslab/lab/GenRef/10x-hg38 | 1000 | 120000 | 30 | Osteo_pt_4 |
| GSE162454 | SRR13927115 | G0016 | GSE162454: Osteosarcoma GSM5155198: Patient 4 | tumor validation | Osteosarcoma | /home/gdrobertslab/lab/GenRef/10x-hg38 | 1000 | 120000 | 30 | Osteo_pt_4 |
| GSE162454 | SRR13927116 | G0017 | GSE162454: Osteosarcoma GSM5155199: Patient 5 | tumor validation | Osteosarcoma | /home/gdrobertslab/lab/GenRef/10x-hg38 | 1000 | 120000 | 30 | Osteo_pt_5 |
| GSE162454 | SRR13927117 | G0017 | GSE162454: Osteosarcoma GSM5155199: Patient 5 | tumor validation | Osteosarcoma | /home/gdrobertslab/lab/GenRef/10x-hg38 | 1000 | 120000 | 30 | Osteo_pt_5 |
| GSE162454 | SRR13927118 | G0017 | GSE162454: Osteosarcoma GSM5155199: Patient 5 | tumor validation | Osteosarcoma | /home/gdrobertslab/lab/GenRef/10x-hg38 | 1000 | 120000 | 30 | Osteo_pt_5 |
| GSE162454 | SRR13927119 | G0017 | GSE162454: Osteosarcoma GSM5155199: Patient 5 | tumor validation | Osteosarcoma | /home/gdrobertslab/lab/GenRef/10x-hg38 | 1000 | 120000 | 30 | Osteo_pt_5 |
| GSE162454 | SRR13927120 | G0018 | GSE162454: Osteosarcoma GSM5155200: Patient 6 | tumor validation | Osteosarcoma | /home/gdrobertslab/lab/GenRef/10x-hg38 | 1000 | 100000 | 30 | Osteo_pt_6 |
| GSE162454 | SRR13927121 | G0018 | GSE162454: Osteosarcoma GSM5155200: Patient 6 | tumor validation | Osteosarcoma | /home/gdrobertslab/lab/GenRef/10x-hg38 | 1000 | 100000 | 30 | Osteo_pt_6 |
| GSE162454 | SRR13927122 | G0018 | GSE162454: Osteosarcoma GSM5155200: Patient 6 | tumor validation | Osteosarcoma | /home/gdrobertslab/lab/GenRef/10x-hg38 | 1000 | 100000 | 30 | Osteo_pt_6 |
| GSE162454 | SRR13927123 | G0018 | GSE162454: Osteosarcoma GSM5155200: Patient 6 | tumor validation | Osteosarcoma | /home/gdrobertslab/lab/GenRef/10x-hg38 | 1000 | 100000 | 30 | Osteo_pt_6 |
| GSE169396 | SRR14041330 | G0019 | GSE169396: bone tissue cells-S1 | healthy samples | healthy | /home/gdrobertslab/lab/GenRef/10x-hg38 | 1000 | 100000 | 20 | Bone_pt_1 |
| GSE169396 | SRR14041331 | G0019 | GSE169396: bone tissue cells-S1 | healthy samples | healthy | /home/gdrobertslab/lab/GenRef/10x-hg38 | 1000 | 100000 | 20 | Bone_pt_1 |
| GSE169396 | SRR14041332 | G0019 | GSE169396: bone tissue cells-S1 | healthy samples | healthy | /home/gdrobertslab/lab/GenRef/10x-hg38 | 1000 | 100000 | 20 | Bone_pt_1 |
| GSE169396 | SRR14041333 | G0020 | GSE169396: bone tissue cells-S2 | healthy samples | healthy | /home/gdrobertslab/lab/GenRef/10x-hg38 | 1000 | 80000 | 20 | Bone_pt_2 |
| GSE169396 | SRR14041334 | G0020 | GSE169396: bone tissue cells-S2 | healthy samples | healthy | /home/gdrobertslab/lab/GenRef/10x-hg38 | 1000 | 80000 | 20 | Bone_pt_2 |
| GSE169396 | SRR14041335 | G0020 | GSE169396: bone tissue cells-S2 | healthy samples | healthy | /home/gdrobertslab/lab/GenRef/10x-hg38 | 1000 | 80000 | 20 | Bone_pt_2 |
| GSE169396 | SRR14041336 | G0020 | GSE169396: bone tissue cells-S2 | healthy samples | healthy | /home/gdrobertslab/lab/GenRef/10x-hg38 | 1000 | 80000 | 20 | Bone_pt_2 |
| GSE169396 | SRR14041337 | G0020 | GSE169396: bone tissue cells-S2 | healthy samples | healthy | /home/gdrobertslab/lab/GenRef/10x-hg38 | 1000 | 80000 | 20 | Bone_pt_2 |
| GSE169396 | SRR14041338 | G0020 | GSE169396: bone tissue cells-S2 | healthy samples | healthy | /home/gdrobertslab/lab/GenRef/10x-hg38 | 1000 | 80000 | 20 | Bone_pt_2 |
| GSE169396 | SRR14041339 | G0020 | GSE169396: bone tissue cells-S2 | healthy samples | healthy | /home/gdrobertslab/lab/GenRef/10x-hg38 | 1000 | 80000 | 20 | Bone_pt_2 |
| GSE169396 | SRR14041340 | G0020 | GSE169396: bone tissue cells-S2 | healthy samples | healthy | /home/gdrobertslab/lab/GenRef/10x-hg38 | 1000 | 80000 | 20 | Bone_pt_2 |
| GSE169396 | SRR14041341 | G0020 | GSE169396: bone tissue cells-S2 | healthy samples | healthy | /home/gdrobertslab/lab/GenRef/10x-hg38 | 1000 | 80000 | 20 | Bone_pt_2 |
| GSE169396 | SRR14041342 | G0020 | GSE169396: bone tissue cells-S2 | healthy samples | healthy | /home/gdrobertslab/lab/GenRef/10x-hg38 | 1000 | 80000 | 20 | Bone_pt_2 |
| GSE169396 | SRR14041343 | G0020 | GSE169396: bone tissue cells-S2 | healthy samples | healthy | /home/gdrobertslab/lab/GenRef/10x-hg38 | 1000 | 80000 | 20 | Bone_pt_2 |
| GSE169396 | SRR14041344 | G0021 | GSE169396: bone tissue cells-S3 | healthy samples | healthy | /home/gdrobertslab/lab/GenRef/10x-hg38 | 1000 | 80000 | 20 | Bone_pt_3 |
| GSE169396 | SRR14041345 | G0021 | GSE169396: bone tissue cells-S3 | healthy samples | healthy | /home/gdrobertslab/lab/GenRef/10x-hg38 | 1000 | 80000 | 20 | Bone_pt_3 |
| GSE169396 | SRR14041346 | G0021 | GSE169396: bone tissue cells-S3 | healthy samples | healthy | /home/gdrobertslab/lab/GenRef/10x-hg38 | 1000 | 80000 | 20 | Bone_pt_3 |
| GSE169396 | SRR14041347 | G0021 | GSE169396: bone tissue cells-S3 | healthy samples | healthy | /home/gdrobertslab/lab/GenRef/10x-hg38 | 1000 | 80000 | 20 | Bone_pt_3 |
| GSE169396 | SRR14041348 | G0022 | GSE169396: bone tissue cells-S4 | healthy samples | healthy | /home/gdrobertslab/lab/GenRef/10x-hg38 | 1000 | 80000 | 20 | Bone_pt_4 |
| GSE169396 | SRR14041349 | G0022 | GSE169396: bone tissue cells-S4 | healthy samples | healthy | /home/gdrobertslab/lab/GenRef/10x-hg38 | 1000 | 80000 | 20 | Bone_pt_4 |
| GSE169396 | SRR14041350 | G0022 | GSE169396: bone tissue cells-S4 | healthy samples | healthy | /home/gdrobertslab/lab/GenRef/10x-hg38 | 1000 | 80000 | 20 | Bone_pt_4 |
| GSE169396 | SRR14041351 | G0022 | GSE169396: bone tissue cells-S4 | healthy samples | healthy | /home/gdrobertslab/lab/GenRef/10x-hg38 | 1000 | 80000 | 20 | Bone_pt_4 |
| GSE169396 | SRR14041352 | G0022 | GSE169396: bone tissue cells-S4 | healthy samples | healthy | /home/gdrobertslab/lab/GenRef/10x-hg38 | 1000 | 80000 | 20 | Bone_pt_4 |
| GSE169396 | SRR14041353 | G0022 | GSE169396: bone tissue cells-S4 | healthy samples | healthy | /home/gdrobertslab/lab/GenRef/10x-hg38 | 1000 | 80000 | 20 | Bone_pt_4 |
| GSE169396 | SRR14041354 | G0022 | GSE169396: bone tissue cells-S4 | healthy samples | healthy | /home/gdrobertslab/lab/GenRef/10x-hg38 | 1000 | 80000 | 20 | Bone_pt_4 |
| GSE169396 | SRR14041355 | G0022 | GSE169396: bone tissue cells-S4 | healthy samples | healthy | /home/gdrobertslab/lab/GenRef/10x-hg38 | 1000 | 80000 | 20 | Bone_pt_4 |
| GSE237579 | SRR25308531 | G0023 | scRNA-seq PMBC healthy donor 6 | healthy samples | healthy | /home/gdrobertslab/lab/GenRef/10x-hg38 | 1000 | 20000 | 20 | PBMC_pt_1 |
| GSE237579 | SRR25308533 | G0023 | scRNA-seq PMBC healthy donor 6 | healthy samples | healthy | /home/gdrobertslab/lab/GenRef/10x-hg38 | 1000 | 20000 | 20 | PBMC_pt_1 |
| GSE237579 | SRR25308532 | G0024 | scRNA-seq PMBC healthy donor 7 | healthy samples | healthy | /home/gdrobertslab/lab/GenRef/10x-hg38 | 1000 | 15000 | 20 | PBMC_pt_2 |
| GSE237579 | SRR25308535 | G0024 | scRNA-seq PMBC healthy donor 7 | healthy samples | healthy | /home/gdrobertslab/lab/GenRef/10x-hg38 | 1000 | 15000 | 20 | PBMC_pt_2 |
| GSE184880 | SRR16093349 | G0025 | High-grade serous ovarian cancer tissue | tumor validation | Ovarian cancer | /home/gdrobertslab/lab/GenRef/10x-hg38 | 1000 | 150000 | 40 | OvCanc_pt_1 |
| GSE184880 | SRR16093350 | G0025 | High-grade serous ovarian cancer tissue | tumor validation | Ovarian cancer | /home/gdrobertslab/lab/GenRef/10x-hg38 | 1000 | 150000 | 40 | OvCanc_pt_1 |
| GSE184880 | SRR16093351 | G0025 | High-grade serous ovarian cancer tissue | tumor validation | Ovarian cancer | /home/gdrobertslab/lab/GenRef/10x-hg38 | 1000 | 150000 | 40 | OvCanc_pt_1 |
| GSE184880 | SRR16093352 | G0025 | High-grade serous ovarian cancer tissue | tumor validation | Ovarian cancer | /home/gdrobertslab/lab/GenRef/10x-hg38 | 1000 | 150000 | 40 | OvCanc_pt_1 |
| GSE184880 | SRR16093353 | G0025 | High-grade serous ovarian cancer tissue | tumor validation | Ovarian cancer | /home/gdrobertslab/lab/GenRef/10x-hg38 | 1000 | 150000 | 40 | OvCanc_pt_1 |
| GSE184880 | SRR16093354 | G0025 | High-grade serous ovarian cancer tissue | tumor validation | Ovarian cancer | /home/gdrobertslab/lab/GenRef/10x-hg38 | 1000 | 150000 | 40 | OvCanc_pt_1 |
| GSE184880 | SRR16093355 | G0025 | High-grade serous ovarian cancer tissue | tumor validation | Ovarian cancer | /home/gdrobertslab/lab/GenRef/10x-hg38 | 1000 | 150000 | 40 | OvCanc_pt_1 |
| GSE184880 | SRR16093356 | G0025 | High-grade serous ovarian cancer tissue | tumor validation | Ovarian cancer | /home/gdrobertslab/lab/GenRef/10x-hg38 | 1000 | 150000 | 40 | OvCanc_pt_1 |
| GSE184880 | SRR16093357 | G0026 | High-grade serous ovarian cancer tissue | tumor validation | Ovarian cancer | /home/gdrobertslab/lab/GenRef/10x-hg38 | 1000 | 150000 | 40 | OvCanc_pt_2 |
| GSE184880 | SRR16093358 | G0026 | High-grade serous ovarian cancer tissue | tumor validation | Ovarian cancer | /home/gdrobertslab/lab/GenRef/10x-hg38 | 1000 | 150000 | 40 | OvCanc_pt_2 |
| GSE184880 | SRR16093359 | G0026 | High-grade serous ovarian cancer tissue | tumor validation | Ovarian cancer | /home/gdrobertslab/lab/GenRef/10x-hg38 | 1000 | 150000 | 40 | OvCanc_pt_2 |
| GSE184880 | SRR16093360 | G0026 | High-grade serous ovarian cancer tissue | tumor validation | Ovarian cancer | /home/gdrobertslab/lab/GenRef/10x-hg38 | 1000 | 150000 | 40 | OvCanc_pt_2 |
| GSE184880 | SRR16093361 | G0027 | High-grade serous ovarian cancer tissue | tumor validation | Ovarian cancer | /home/gdrobertslab/lab/GenRef/10x-hg38 | 1000 | 125000 | 40 | OvCanc_pt_3 |
| GSE184880 | SRR16093362 | G0027 | High-grade serous ovarian cancer tissue | tumor validation | Ovarian cancer | /home/gdrobertslab/lab/GenRef/10x-hg38 | 1000 | 125000 | 40 | OvCanc_pt_3 |
| GSE184880 | SRR16093363 | G0027 | High-grade serous ovarian cancer tissue | tumor validation | Ovarian cancer | /home/gdrobertslab/lab/GenRef/10x-hg38 | 1000 | 125000 | 40 | OvCanc_pt_3 |
| GSE184880 | SRR16093364 | G0027 | High-grade serous ovarian cancer tissue | tumor validation | Ovarian cancer | /home/gdrobertslab/lab/GenRef/10x-hg38 | 1000 | 125000 | 40 | OvCanc_pt_3 |
| GSE184880 | SRR16093365 | G0028 | High-grade serous ovarian cancer tissue | tumor validation | Ovarian cancer | /home/gdrobertslab/lab/GenRef/10x-hg38 | 1000 | 150000 | 50 | OvCanc_pt_4 |
| GSE184880 | SRR16093366 | G0028 | High-grade serous ovarian cancer tissue | tumor validation | Ovarian cancer | /home/gdrobertslab/lab/GenRef/10x-hg38 | 1000 | 150000 | 50 | OvCanc_pt_4 |
| GSE184880 | SRR16093367 | G0028 | High-grade serous ovarian cancer tissue | tumor validation | Ovarian cancer | /home/gdrobertslab/lab/GenRef/10x-hg38 | 1000 | 150000 | 50 | OvCanc_pt_4 |
| GSE184880 | SRR16093368 | G0028 | High-grade serous ovarian cancer tissue | tumor validation | Ovarian cancer | /home/gdrobertslab/lab/GenRef/10x-hg38 | 1000 | 150000 | 50 | OvCanc_pt_4 |
| GSE184880 | SRR16093369 | G0029 | High-grade serous ovarian cancer tissue | tumor validation | Ovarian cancer | /home/gdrobertslab/lab/GenRef/10x-hg38 | 1000 | 125000 | 40 | OvCanc_pt_5 |
| GSE184880 | SRR16093370 | G0029 | High-grade serous ovarian cancer tissue | tumor validation | Ovarian cancer | /home/gdrobertslab/lab/GenRef/10x-hg38 | 1000 | 125000 | 40 | OvCanc_pt_5 |
| GSE184880 | SRR16093371 | G0029 | High-grade serous ovarian cancer tissue | tumor validation | Ovarian cancer | /home/gdrobertslab/lab/GenRef/10x-hg38 | 1000 | 125000 | 40 | OvCanc_pt_5 |
| GSE184880 | SRR16093372 | G0029 | High-grade serous ovarian cancer tissue | tumor validation | Ovarian cancer | /home/gdrobertslab/lab/GenRef/10x-hg38 | 1000 | 125000 | 40 | OvCanc_pt_5 |
| GSE184880 | SRR16093373 | G0029 | High-grade serous ovarian cancer tissue | tumor validation | Ovarian cancer | /home/gdrobertslab/lab/GenRef/10x-hg38 | 1000 | 125000 | 40 | OvCanc_pt_5 |
| GSE184880 | SRR16093374 | G0029 | High-grade serous ovarian cancer tissue | tumor validation | Ovarian cancer | /home/gdrobertslab/lab/GenRef/10x-hg38 | 1000 | 125000 | 40 | OvCanc_pt_5 |
| GSE184880 | SRR16093375 | G0029 | High-grade serous ovarian cancer tissue | tumor validation | Ovarian cancer | /home/gdrobertslab/lab/GenRef/10x-hg38 | 1000 | 125000 | 40 | OvCanc_pt_5 |
| GSE184880 | SRR16093376 | G0029 | High-grade serous ovarian cancer tissue | tumor validation | Ovarian cancer | /home/gdrobertslab/lab/GenRef/10x-hg38 | 1000 | 125000 | 40 | OvCanc_pt_5 |
| GSE184880 | SRR16093377 | G0029 | High-grade serous ovarian cancer tissue | tumor validation | Ovarian cancer | /home/gdrobertslab/lab/GenRef/10x-hg38 | 1000 | 125000 | 40 | OvCanc_pt_5 |
| GSE184880 | SRR16093378 | G0029 | High-grade serous ovarian cancer tissue | tumor validation | Ovarian cancer | /home/gdrobertslab/lab/GenRef/10x-hg38 | 1000 | 125000 | 40 | OvCanc_pt_5 |
| GSE184880 | SRR16093379 | G0029 | High-grade serous ovarian cancer tissue | tumor validation | Ovarian cancer | /home/gdrobertslab/lab/GenRef/10x-hg38 | 1000 | 125000 | 40 | OvCanc_pt_5 |
| GSE184880 | SRR16093380 | G0029 | High-grade serous ovarian cancer tissue | tumor validation | Ovarian cancer | /home/gdrobertslab/lab/GenRef/10x-hg38 | 1000 | 125000 | 40 | OvCanc_pt_5 |
| GSE184880 | SRR16093381 | G0030 | High-grade serous ovarian cancer tissue | tumor validation | Ovarian cancer | /home/gdrobertslab/lab/GenRef/10x-hg38 | 1000 | 150000 | 40 | OvCanc_pt_6 |
| GSE184880 | SRR16093382 | G0030 | High-grade serous ovarian cancer tissue | tumor validation | Ovarian cancer | /home/gdrobertslab/lab/GenRef/10x-hg38 | 1000 | 150000 | 40 | OvCanc_pt_6 |
| GSE184880 | SRR16093383 | G0030 | High-grade serous ovarian cancer tissue | tumor validation | Ovarian cancer | /home/gdrobertslab/lab/GenRef/10x-hg38 | 1000 | 150000 | 40 | OvCanc_pt_6 |
| GSE184880 | SRR16093384 | G0030 | High-grade serous ovarian cancer tissue | tumor validation | Ovarian cancer | /home/gdrobertslab/lab/GenRef/10x-hg38 | 1000 | 150000 | 40 | OvCanc_pt_6 |
| GSE184880 | SRR16093385 | G0031 | High-grade serous ovarian cancer tissue | tumor validation | Ovarian cancer | /home/gdrobertslab/lab/GenRef/10x-hg38 | 1000 | 100000 | 40 | OvCanc_pt_7 |
| GSE184880 | SRR16093386 | G0031 | High-grade serous ovarian cancer tissue | tumor validation | Ovarian cancer | /home/gdrobertslab/lab/GenRef/10x-hg38 | 1000 | 100000 | 40 | OvCanc_pt_7 |
| GSE184880 | SRR16093387 | G0031 | High-grade serous ovarian cancer tissue | tumor validation | Ovarian cancer | /home/gdrobertslab/lab/GenRef/10x-hg38 | 1000 | 100000 | 40 | OvCanc_pt_7 |
| GSE184880 | SRR16093388 | G0031 | High-grade serous ovarian cancer tissue | tumor validation | Ovarian cancer | /home/gdrobertslab/lab/GenRef/10x-hg38 | 1000 | 100000 | 40 | OvCanc_pt_7 |
| GSE306201 | SRR35094270 | G0032 | Breast cancer Breast primary | tumor validation | Breast cancer | /home/gdrobertslab/lab/GenRef/10x-hg38 | 1000 | 50000 | 30 | BrCanc_pt_1 |
| GSE306201 | SRR35094266 | G0034 | Breast cancer Breast primary | tumor validation | Breast cancer | /home/gdrobertslab/lab/GenRef/10x-hg38 | 1000 | 30000 | 20 | BrCanc_pt_3 |
| GSE306201 | SRR35094265 | G0035 | Breast cancer Breast primary | tumor validation | Breast cancer | /home/gdrobertslab/lab/GenRef/10x-hg38 | 1000 | 100000 | 20 | BrCanc_pt_4 |
| GSE306201 | SRR35094263 | G0036 | Breast cancer Breast primary | tumor validation | Breast cancer | /home/gdrobertslab/lab/GenRef/10x-hg38 | 1000 | 25000 | 20 | BrCanc_pt_5 |
| GSE306201 | SRR35094264 | G0038 | Breast cancer Lymph Node Metastasis | tumor validation | Breast cancer | /home/gdrobertslab/lab/GenRef/10x-hg38 | 1000 | 90000 | 30 | BrCanc_pt_7 |
| GSE306201 | SRR35094267 | G0039 | Breast cancer Mediastinum Metastasis | tumor validation | Breast cancer | /home/gdrobertslab/lab/GenRef/10x-hg38 | 1000 | 30000 | 25 | BrCanc_pt_8 |
| GSE306201 | SRR35094250 | G0040 | Breast cancer Liver Metastasis | tumor validation | Breast cancer | /home/gdrobertslab/lab/GenRef/10x-hg38 | 1000 | 50000 | 30 | BrCanc_pt_9 |
| GSE306201 | SRR35094252 | G0040 | Breast cancer Liver Metastasis | tumor validation | Breast cancer | /home/gdrobertslab/lab/GenRef/10x-hg38 | 1000 | 50000 | 30 | BrCanc_pt_9 |
| GSE306201 | SRR35094251 | G0041 | Breast cancer Liver Metastasis | tumor validation | Breast cancer | /home/gdrobertslab/lab/GenRef/10x-hg38 | 1000 | 30000 | 20 | BrCanc_pt_10 |
| GSE306201 | SRR35094257 | G0042 | Breast cancer Lymph Node Metastasis | tumor validation | Breast cancer | /home/gdrobertslab/lab/GenRef/10x-hg38 | 1000 | 50000 | 20 | BrCanc_pt_11 |
| GSE306201 | SRR35094255 | G0043 | Breast cancer Liver Metastasis | tumor validation | Breast cancer | /home/gdrobertslab/lab/GenRef/10x-hg38 | 1000 | 25000 | 20 | BrCanc_pt_12 |
| GSE306201 | SRR35094259 | G0043 | Breast cancer Liver Metastasis | tumor validation | Breast cancer | /home/gdrobertslab/lab/GenRef/10x-hg38 | 1000 | 25000 | 20 | BrCanc_pt_12 |
| GSE306201 | SRR35094260 | G0043 | Breast cancer Liver Metastasis | tumor validation | Breast cancer | /home/gdrobertslab/lab/GenRef/10x-hg38 | 1000 | 25000 | 20 | BrCanc_pt_12 |
| GSE306201 | SRR35094261 | G0043 | Breast cancer Liver Metastasis | tumor validation | Breast cancer | /home/gdrobertslab/lab/GenRef/10x-hg38 | 1000 | 25000 | 20 | BrCanc_pt_12 |
| GSE306201 | SRR35094258 | G0044 | Breast cancer Adrenal Gland Metastasis | tumor validation | Breast cancer | /home/gdrobertslab/lab/GenRef/10x-hg38 | 1000 | 60000 | 20 | BrCanc_pt_13 |
| GSE306201 | SRR35094256 | G0045 | Breast cancer Bone Metastasis | tumor validation | Breast cancer | /home/gdrobertslab/lab/GenRef/10x-hg38 | 1000 | 125000 | 40 | BrCanc_pt_14 |
| GSE306201 | SRR35094253 | G0047 | Breast cancer Liver Metastasis | tumor validation | Breast cancer | /home/gdrobertslab/lab/GenRef/10x-hg38 | 1000 | 80000 | 40 | BrCanc_pt_16 |
| GSE261693 | SRR28353274 | G0048 | Ewing2_GEX | tumor validation | Ewing sarcoma | /home/gdrobertslab/lab/GenRef/10x-hg38 | 1000 | 40000 | 35 | Ewing_pt_1 |
| GSE261693 | SRR28353275 | G0049 | Ewing1_GEX | tumor validation | Ewing sarcoma | /home/gdrobertslab/lab/GenRef/10x-hg38 | 1000 | 75000 | 30 | Ewing_pt_2 |

| | | | | | | | | | | |
|---|---|---|---|---|---|---|---|---|---|---|
| GSE316454 | NA | S0122 | sorted Cherry positive fraction 1:1:1 mix of CD45+, Epcam+ and double negative | truthset validation | Osteosarcoma | 10x-mouse | 1000 | 100000 | 30 | BALBC_K7M2_lung_1 |
| GSE316454 | NA | S0123 | sorted Cherry negative fraction 1:1:1 mix of CD45+, Epcam+ and double negative | truthset validation | Osteosarcoma | 10x-mouse | 1000 | 35000 | 30 | BALBC_K7M2_lung_2 |
| GSE252703 | NA | S0066 | B6 ctrl lung Epcam postive:negative 1:1 | mouse validation | healthy | 10x-mouse | 1000 | 90000 | 30 | C57BL6_lung |
| GSE179681 | NA | S0067 | B6, F420 injected, lungs from ROB 273 A, B, D; Epcam | mouse validation | Osteosarcoma | 10x-mouse | 1000 | 100000 | 30 | C57BL6_F420_lung_1 |
| GSE252703 | NA | S0068 | B6, F420 injected, lungs from ROB 273E and ROB 274A; Epcam postive:negative 1:1 | mouse validation | Osteosarcoma | 10x-mouse | 1000 | 120000 | 30 | C57BL6_F420_lung_2 |
| GSE252703 | NA | S0074 | Balb/c ctrl lung Epcam postive:negative 1:1 | mouse validation | healthy | 10x-mouse | 1000 | 75000 | 20 | BALBC_lung |
| GSE252703 | NA | S0075 | Balb/c, K7M2 injected, lungs; Epcam postive:negative 1:1 | mouse validation | Osteosarcoma | 10x-mouse | 1000 | 120000 | 30 | BALBC_K7M2_lung |
| GSE179681 | NA | S0200 | F420 cellculture p17 | mouse validation | Osteosarcoma | 10x-mouse | 1000 | 75000 | 20 | F420_culture |
| GSE179681 | NA | S0201 | K7M2 cellculture p17 | mouse validation | Osteosarcoma | 10x-mouse | 1000 | 80000 | 20 | K7M2_culture |
| GSE316454 | NA | S0246 | F420 lung mets and niche cells ctrl | mouse validation | Osteosarcoma | 10x-mouse | 1000 | 50000 | 20 | C57BL6_F420_lung_3 |
| GSE316454 | NA | S0260 | F420, barcoded, 46 days post TV injection | mouse validation | Osteosarcoma | 10x-mouse | 1000 | 80000 | 20 | C57BL6_F420_lung_4 |
| GSE316454 | NA | S0261 | F420, barcoded, 46 days post TV injection | mouse validation | Osteosarcoma | 10x-mouse | 1000 | 90000 | 20 | C57BL6_F420_lung_5 |
| NA | NA | flex_eye | Eye mouse flex data | flex validation | healthy | mm10 | 0 | 75000 | 20 | Flex_eye |
| NA | NA | flex_intestine | Intestine mouse flex data | flex validation | healthy | mm11 | 0 | 100000 | 20 | Flex_intestine |
| NA | NA | flex_ovary | Ovary mouse flex data | flex validation | healthy | mm12 | 0 | 150000 | 20 | Flex_ovary |
| NA | NA | flex_stomach | Stomach mouse flex data | flex validation | healthy | mm13 | 0 | 150000 | 20 | Flex_stomach |

#### Supplemental Table 2: Genomic coverage for individual cells

| depth | mean_count | median_count_var | se_count | cum_count_var |
|---|---|---|---|---|
| 2 | 69,025.9295 | 32,827.00 | 1,774.65 | 137,352.66 |
| 3 | 22,432.2593 | 10,605.50 | 584.08 | 68,326.73 |
| 4 | 11,258.4776 | 5,657.50 | 289.19 | 45,894.47 |
| 5 | 6,829.5942 | 3,718.00 | 172.47 | 34,636.00 |
| 6 | 4,616.8391 | 2,631.00 | 115.04 | 27,806.40 |
| 7 | 3,337.8276 | 1,980.50 | 82.24 | 23,189.56 |
| 8 | 2,533.5673 | 1,515.00 | 62.11 | 19,851.73 |
| 9 | 1,988.3391 | 1,173.50 | 48.56 | 17,318.17 |
| 10 | 1,609.5016 | 931.00 | 39.01 | 15,329.83 |
| 11 | 1,328.2413 | 730.50 | 32.12 | 13,720.33 |
| 12 | 1,106.5228 | 581.50 | 27.00 | 12,392.09 |
| 13 | 940.5929 | 466.50 | 22.98 | 11,285.56 |
| 14 | 812.6856 | 387.00 | 19.92 | 10,344.97 |
| 15 | 708.3708 | 329.50 | 17.54 | 9,532.28 |
| 16 | 620.7276 | 268.00 | 15.39 | 8,823.91 |
| 17 | 547.9804 | 238.50 | 13.73 | 8,203.19 |
| 18 | 488.0971 | 202.50 | 12.35 | 7,655.21 |
| 19 | 436.1090 | 176.00 | 11.13 | 7,167.11 |
| 20 | 395.0356 | 152.00 | 10.15 | 6,731.00 |
| 21 | 358.3045 | 137.00 | 9.28 | 6,335.96 |
| 22 | 324.0462 | 115.00 | 8.53 | 5,977.66 |
| 23 | 292.8811 | 105.00 | 7.79 | 5,653.61 |
| 24 | 268.8449 | 98.00 | 7.19 | 5,360.73 |
| 25 | 245.3423 | 84.00 | 6.63 | 5,091.89 |
| 26 | 227.6888 | 80.00 | 6.19 | 4,846.54 |
| 27 | 210.9865 | 71.00 | 5.79 | 4,618.86 |
| 28 | 195.7250 | 67.00 | 5.44 | 4,407.87 |
| 29 | 182.0439 | 61.00 | 5.12 | 4,212.14 |
| 30 | 169.1747 | 55.00 | 4.82 | 4,030.10 |
| 31 | 157.7622 | 52.50 | 4.50 | 3,860.93 |
| 32 | 146.8215 | 47.00 | 4.23 | 3,703.16 |
| 33 | 138.3856 | 46.00 | 3.98 | 3,556.34 |
| 34 | 129.2093 | 41.00 | 3.80 | 3,417.96 |
| 35 | 121.9599 | 39.00 | 3.59 | 3,288.75 |
| 36 | 115.1587 | 36.00 | 3.41 | 3,166.79 |
| 37 | 108.9635 | 33.00 | 3.27 | 3,051.63 |
| 38 | 102.8135 | 31.00 | 3.11 | 2,942.67 |
| 39 | 97.0212 | 31.00 | 2.94 | 2,839.85 |
| 40 | 91.4833 | 28.00 | 2.80 | 2,742.83 |
| 41 | 86.6888 | 25.00 | 2.68 | 2,651.35 |
| 42 | 82.2785 | 24.00 | 2.58 | 2,564.66 |
| 43 | 77.8179 | 22.00 | 2.48 | 2,482.38 |
| 44 | 74.2333 | 21.00 | 2.37 | 2,404.56 |

| | | | | |
|---|---|---|---|---|
| 45 | 69.8183 | 20.00 | 2.23 | 2,330.33 |
| 46 | 67.3506 | 18.00 | 2.20 | 2,260.51 |
| 47 | 64.0878 | 18.00 | 2.08 | 2,193.16 |
| 48 | 61.0413 | 17.00 | 2.00 | 2,129.07 |
| 49 | 58.2596 | 15.00 | 1.95 | 2,068.03 |
| 50 | 55.3571 | 15.00 | 1.84 | 2,009.77 |
| 51 | 53.4112 | 14.00 | 1.79 | 1,954.41 |
| 52 | 50.7740 | 13.00 | 1.72 | 1,901.00 |
| 53 | 48.6923 | 13.00 | 1.67 | 1,850.23 |
| 54 | 46.7064 | 11.00 | 1.59 | 1,801.54 |
| 55 | 45.0026 | 11.00 | 1.55 | 1,754.83 |
| 56 | 43.2974 | 11.00 | 1.49 | 1,709.83 |
| 57 | 41.2404 | 10.00 | 1.43 | 1,666.53 |
| 58 | 40.0516 | 9.00 | 1.41 | 1,625.29 |
| 59 | 37.9913 | 8.00 | 1.34 | 1,585.24 |
| 60 | 36.6968 | 9.00 | 1.31 | 1,547.25 |
| 61 | 35.6167 | 8.00 | 1.27 | 1,510.55 |
| 62 | 34.2397 | 7.00 | 1.24 | 1,474.93 |
| 63 | 32.9221 | 8.00 | 1.19 | 1,440.69 |
| 64 | 31.9394 | 7.00 | 1.16 | 1,407.77 |
| 65 | 30.2747 | 6.00 | 1.12 | 1,375.83 |
| 66 | 29.2176 | 6.00 | 1.08 | 1,345.56 |
| 67 | 28.4337 | 6.00 | 1.06 | 1,316.34 |
| 68 | 27.7513 | 5.00 | 1.04 | 1,287.91 |
| 69 | 26.6160 | 5.00 | 1.00 | 1,260.15 |
| 70 | 25.6859 | 5.00 | 0.96 | 1,233.54 |
| 71 | 24.8907 | 5.00 | 0.95 | 1,207.85 |
| 72 | 24.0769 | 4.50 | 0.91 | 1,182.96 |
| 73 | 23.3962 | 4.00 | 0.90 | 1,158.88 |
| 74 | 22.6843 | 4.00 | 0.87 | 1,135.49 |
| 75 | 21.8026 | 4.00 | 0.86 | 1,112.80 |
| 76 | 21.0006 | 4.00 | 0.82 | 1,091.00 |
| 77 | 20.4228 | 3.00 | 0.80 | 1,070.00 |
| 78 | 19.8795 | 3.00 | 0.79 | 1,049.58 |
| 79 | 19.2237 | 3.00 | 0.77 | 1,029.70 |
| 80 | 18.9125 | 3.00 | 0.75 | 1,010.48 |
| 81 | 18.2731 | 2.00 | 0.73 | 991.56 |
| 82 | 17.7026 | 2.00 | 0.71 | 973.29 |
| 83 | 17.2218 | 2.00 | 0.70 | 955.59 |
| 84 | 16.8535 | 2.00 | 0.69 | 938.37 |
| 85 | 16.3929 | 2.00 | 0.67 | 921.51 |
| 86 | 15.9535 | 2.00 | 0.65 | 905.12 |
| 87 | 15.4769 | 2.00 | 0.63 | 889.17 |
| 88 | 14.9721 | 1.00 | 0.63 | 873.69 |

| | | | | |
|---|---|---|---|---|
| 89 | 14.6103 | 1.00 | 0.60 | 858.72 |
| 90 | 14.3141 | 1.00 | 0.59 | 844.11 |
| 91 | 13.7535 | 1.00 | 0.56 | 829.79 |
| 92 | 13.4481 | 1.00 | 0.56 | 816.04 |
| 93 | 13.1112 | 1.00 | 0.55 | 802.59 |
| 94 | 12.7981 | 1.00 | 0.54 | 789.48 |
| 95 | 12.4468 | 1.00 | 0.53 | 776.68 |
| 96 | 12.2327 | 1.00 | 0.52 | 764.23 |
| 97 | 11.8237 | 1.00 | 0.50 | 752.00 |
| 98 | 11.5093 | 1.00 | 0.49 | 740.18 |
| 99 | 11.0958 | 0.00 | 0.48 | 728.67 |
| 100 | 10.7715 | 0.00 | 0.47 | 717.57 |
| 101 | 10.3603 | 0.00 | 0.45 | 706.80 |
| 102 | 10.1513 | 0.00 | 0.44 | 696.44 |
| 103 | 9.9074 | 0.00 | 0.43 | 686.29 |
| 104 | 9.7026 | 0.00 | 0.42 | 676.38 |
| 105 | 9.5125 | 0.00 | 0.41 | 666.68 |
| 106 | 9.2933 | 0.00 | 0.40 | 657.17 |
| 107 | 9.0865 | 0.00 | 0.39 | 647.87 |
| 108 | 8.7946 | 0.00 | 0.39 | 638.79 |
| 109 | 8.7122 | 0.00 | 0.39 | 629.99 |
| 110 | 8.5561 | 0.00 | 0.38 | 621.28 |
| 111 | 8.5609 | 0.00 | 0.38 | 612.72 |
| 112 | 8.3231 | 0.00 | 0.37 | 604.16 |
| 113 | 7.8984 | 0.00 | 0.35 | 595.84 |
| 114 | 7.8058 | 0.00 | 0.35 | 587.94 |
| 115 | 7.6542 | 0.00 | 0.34 | 580.14 |
| 116 | 7.4061 | 0.00 | 0.33 | 572.48 |
| 117 | 7.2901 | 0.00 | 0.33 | 565.08 |
| 118 | 7.2266 | 0.00 | 0.32 | 557.79 |
| 119 | 7.0141 | 0.00 | 0.31 | 550.56 |
| 120 | 6.8994 | 0.00 | 0.31 | 543.55 |
| 121 | 6.7397 | 0.00 | 0.31 | 536.65 |
| 122 | 6.6625 | 0.00 | 0.31 | 529.91 |
| 123 | 6.4862 | 0.00 | 0.30 | 523.24 |
| 124 | 6.4933 | 0.00 | 0.30 | 516.76 |
| 125 | 6.3240 | 0.00 | 0.29 | 510.26 |
| 126 | 6.1904 | 0.00 | 0.29 | 503.94 |
| 127 | 5.9433 | 0.00 | 0.27 | 497.75 |
| 128 | 5.8449 | 0.00 | 0.27 | 491.81 |
| 129 | 5.8192 | 0.00 | 0.27 | 485.96 |
| 130 | 5.7647 | 0.00 | 0.26 | 480.14 |
| 131 | 5.5003 | 0.00 | 0.25 | 474.38 |
| 132 | 5.4032 | 0.00 | 0.25 | 468.88 |

| | | | | |
|---|---|---|---|---|
| 133 | 5.2635 | 0.00 | 0.24 | 463.47 |
| 134 | 5.1458 | 0.00 | 0.24 | 458.21 |
| 135 | 4.9942 | 0.00 | 0.23 | 453.07 |
| 136 | 4.9660 | 0.00 | 0.23 | 448.07 |
| 137 | 4.9298 | 0.00 | 0.23 | 443.10 |
| 138 | 4.7728 | 0.00 | 0.22 | 438.18 |
| 139 | 4.7452 | 0.00 | 0.22 | 433.40 |
| 140 | 4.6689 | 0.00 | 0.22 | 428.66 |
| 141 | 4.4907 | 0.00 | 0.21 | 423.99 |
| 142 | 4.4442 | 0.00 | 0.21 | 419.50 |
| 143 | 4.4276 | 0.00 | 0.21 | 415.05 |
| 144 | 4.2776 | 0.00 | 0.20 | 410.63 |
| 145 | 4.2718 | 0.00 | 0.20 | 406.35 |
| 146 | 4.2186 | 0.00 | 0.20 | 402.08 |
| 147 | 4.2071 | 0.00 | 0.20 | 397.86 |
| 148 | 3.9968 | 0.00 | 0.19 | 393.65 |
| 149 | 3.8929 | 0.00 | 0.18 | 389.65 |
| 150 | 3.8872 | 0.00 | 0.19 | 385.76 |
| 151 | 3.7676 | 0.00 | 0.18 | 381.87 |
| 152 | 3.7096 | 0.00 | 0.17 | 378.11 |
| 153 | 3.6112 | 0.00 | 0.17 | 374.40 |
| 154 | 3.6022 | 0.00 | 0.17 | 370.79 |
| 155 | 3.5522 | 0.00 | 0.17 | 367.18 |
| 156 | 3.4356 | 0.00 | 0.16 | 363.63 |
| 157 | 3.4183 | 0.00 | 0.16 | 360.20 |
| 158 | 3.4000 | 0.00 | 0.17 | 356.78 |
| 159 | 3.3256 | 0.00 | 0.16 | 353.38 |
| 160 | 3.3228 | 0.00 | 0.16 | 350.05 |
| 161 | 3.2878 | 0.00 | 0.16 | 346.73 |
| 162 | 3.1045 | 0.00 | 0.15 | 343.44 |
| 163 | 3.1904 | 0.00 | 0.16 | 340.34 |
| 164 | 3.1202 | 0.00 | 0.15 | 337.15 |
| 165 | 3.1471 | 0.00 | 0.15 | 334.03 |
| 166 | 2.9542 | 0.00 | 0.15 | 330.88 |
| 167 | 2.9897 | 0.00 | 0.14 | 327.92 |
| 168 | 2.9413 | 0.00 | 0.15 | 324.93 |
| 169 | 2.9115 | 0.00 | 0.14 | 321.99 |
| 170 | 2.8567 | 0.00 | 0.14 | 319.08 |
| 171 | 2.7413 | 0.00 | 0.14 | 316.23 |
| 172 | 2.6875 | 0.00 | 0.13 | 313.48 |
| 173 | 2.7439 | 0.00 | 0.13 | 310.80 |
| 174 | 2.7022 | 0.00 | 0.13 | 308.05 |
| 175 | 2.6340 | 0.00 | 0.13 | 305.35 |
| 176 | 2.5516 | 0.00 | 0.13 | 302.72 |

| | | | | |
|---|---|---|---|---|
| 177 | 2.5247 | 0.00 | 0.13 | 300.16 |
| 178 | 2.4788 | 0.00 | 0.13 | 297.64 |
| 179 | 2.4071 | 0.00 | 0.12 | 295.16 |
| 180 | 2.3817 | 0.00 | 0.12 | 292.75 |
| 181 | 2.3965 | 0.00 | 0.12 | 290.37 |
| 182 | 2.2734 | 0.00 | 0.12 | 287.98 |
| 183 | 2.2686 | 0.00 | 0.12 | 285.70 |
| 184 | 2.2712 | 0.00 | 0.12 | 283.43 |
| 185 | 2.2087 | 0.00 | 0.12 | 281.16 |
| 186 | 2.1750 | 0.00 | 0.11 | 278.95 |
| 187 | 2.0641 | 0.00 | 0.11 | 276.78 |
| 188 | 2.0609 | 0.00 | 0.11 | 274.71 |
| 189 | 2.0276 | 0.00 | 0.11 | 272.65 |
| 190 | 2.0744 | 0.00 | 0.11 | 270.63 |
| 191 | 2.0881 | 0.00 | 0.11 | 268.55 |
| 192 | 2.0510 | 0.00 | 0.10 | 266.46 |
| 193 | 2.0295 | 0.00 | 0.10 | 264.41 |
| 194 | 1.9715 | 0.00 | 0.10 | 262.38 |
| 195 | 2.0346 | 0.00 | 0.11 | 260.41 |
| 196 | 1.9795 | 0.00 | 0.10 | 258.38 |
| 197 | 1.9429 | 0.00 | 0.10 | 256.40 |
| 198 | 1.9067 | 0.00 | 0.10 | 254.45 |
| 199 | 1.9692 | 0.00 | 0.10 | 252.55 |
| 200 | 1.8657 | 0.00 | 0.10 | 250.58 |
| 201 | 1.9000 | 0.00 | 0.10 | 248.71 |
| 202 | 1.8224 | 0.00 | 0.10 | 246.81 |
| 203 | 1.8029 | 0.00 | 0.10 | 244.99 |
| 204 | 1.7234 | 0.00 | 0.09 | 243.19 |
| 205 | 1.7375 | 0.00 | 0.09 | 241.46 |
| 206 | 1.7567 | 0.00 | 0.09 | 239.73 |
| 207 | 1.7128 | 0.00 | 0.09 | 237.97 |
| 208 | 1.6696 | 0.00 | 0.08 | 236.26 |
| 209 | 1.6667 | 0.00 | 0.09 | 234.59 |
| 210 | 1.6891 | 0.00 | 0.09 | 232.92 |
| 211 | 1.6224 | 0.00 | 0.08 | 231.23 |
| 212 | 1.6228 | 0.00 | 0.08 | 229.61 |
| 213 | 1.5917 | 0.00 | 0.08 | 227.99 |
| 214 | 1.5673 | 0.00 | 0.08 | 226.40 |
| 215 | 1.5484 | 0.00 | 0.08 | 224.83 |
| 216 | 1.6035 | 0.00 | 0.09 | 223.28 |
| 217 | 1.5635 | 0.00 | 0.08 | 221.68 |
| 218 | 1.4984 | 0.00 | 0.08 | 220.11 |
| 219 | 1.5256 | 0.00 | 0.08 | 218.61 |
| 220 | 1.4667 | 0.00 | 0.08 | 217.09 |

| | | | | |
|---|---|---|---|---|
| 221 | 1.4417 | 0.00 | 0.08 | 215.62 |
| 222 | 1.4308 | 0.00 | 0.07 | 214.18 |
| 223 | 1.4151 | 0.00 | 0.08 | 212.75 |
| 224 | 1.3913 | 0.00 | 0.07 | 211.33 |
| 225 | 1.3115 | 0.00 | 0.07 | 209.94 |
| 226 | 1.3314 | 0.00 | 0.07 | 208.63 |
| 227 | 1.3532 | 0.00 | 0.07 | 207.30 |
| 228 | 1.3593 | 0.00 | 0.07 | 205.95 |
| 229 | 1.3269 | 0.00 | 0.08 | 204.59 |
| 230 | 1.3147 | 0.00 | 0.07 | 203.26 |
| 231 | 1.2628 | 0.00 | 0.07 | 201.95 |
| 232 | 1.3000 | 0.00 | 0.07 | 200.68 |
| 233 | 1.2641 | 0.00 | 0.07 | 199.38 |
| 234 | 1.2282 | 0.00 | 0.07 | 198.12 |
| 235 | 1.2228 | 0.00 | 0.06 | 196.89 |
| 236 | 1.2417 | 0.00 | 0.07 | 195.67 |
| 237 | 1.2574 | 0.00 | 0.07 | 194.43 |
| 238 | 1.1635 | 0.00 | 0.06 | 193.17 |
| 239 | 1.1712 | 0.00 | 0.06 | 192.01 |
| 240 | 1.1692 | 0.00 | 0.07 | 190.83 |
| 241 | 1.1830 | 0.00 | 0.07 | 189.67 |
| 242 | 1.1657 | 0.00 | 0.07 | 188.48 |
| 243 | 1.1247 | 0.00 | 0.06 | 187.32 |
| 244 | 1.0942 | 0.00 | 0.06 | 186.19 |
| 245 | 1.0917 | 0.00 | 0.06 | 185.10 |
| 246 | 1.1369 | 0.00 | 0.07 | 184.01 |
| 247 | 1.0635 | 0.00 | 0.06 | 182.87 |
| 248 | 1.0933 | 0.00 | 0.06 | 181.81 |
| 249 | 1.1048 | 0.00 | 0.06 | 180.71 |
| 250 | 1.0785 | 0.00 | 0.06 | 179.61 |
| 251 | 1.0651 | 0.00 | 0.06 | 178.53 |
| 252 | 1.0247 | 0.00 | 0.06 | 177.46 |
| 253 | 1.0026 | 0.00 | 0.05 | 176.44 |
| 254 | 0.9891 | 0.00 | 0.06 | 175.44 |
| 255 | 0.9817 | 0.00 | 0.05 | 174.45 |
| 256 | 1.0077 | 0.00 | 0.06 | 173.47 |
| 257 | 0.9907 | 0.00 | 0.06 | 172.46 |
| 258 | 0.9545 | 0.00 | 0.05 | 171.47 |
| 259 | 0.9170 | 0.00 | 0.05 | 170.51 |
| 260 | 0.9385 | 0.00 | 0.05 | 169.60 |
| 261 | 0.9330 | 0.00 | 0.05 | 168.66 |
| 262 | 0.9250 | 0.00 | 0.05 | 167.72 |
| 263 | 0.8875 | 0.00 | 0.05 | 166.80 |
| 264 | 0.9147 | 0.00 | 0.05 | 165.91 |

| | | | | |
|---|---|---|---|---|
| 265 | 0.8696 | 0.00 | 0.05 | 165.00 |
| 266 | 0.9202 | 0.00 | 0.05 | 164.13 |
| 267 | 0.8965 | 0.00 | 0.05 | 163.21 |
| 268 | 0.8375 | 0.00 | 0.05 | 162.31 |
| 269 | 0.8276 | 0.00 | 0.05 | 161.47 |
| 270 | 0.8465 | 0.00 | 0.05 | 160.65 |
| 271 | 0.8609 | 0.00 | 0.05 | 159.80 |
| 272 | 0.8446 | 0.00 | 0.05 | 158.94 |
| 273 | 0.8574 | 0.00 | 0.05 | 158.09 |
| 274 | 0.8144 | 0.00 | 0.05 | 157.24 |
| 275 | 0.7689 | 0.00 | 0.04 | 156.42 |
| 276 | 0.7728 | 0.00 | 0.05 | 155.65 |
| 277 | 0.8167 | 0.00 | 0.05 | 154.88 |
| 278 | 0.7827 | 0.00 | 0.05 | 154.06 |
| 279 | 0.8080 | 0.00 | 0.05 | 153.28 |
| 280 | 0.8228 | 0.00 | 0.05 | 152.47 |
| 281 | 0.7776 | 0.00 | 0.05 | 151.65 |
| 282 | 0.7744 | 0.00 | 0.04 | 150.87 |
| 283 | 0.7183 | 0.00 | 0.04 | 150.10 |
| 284 | 0.7692 | 0.00 | 0.05 | 149.38 |
| 285 | 0.7615 | 0.00 | 0.04 | 148.61 |
| 286 | 0.7567 | 0.00 | 0.04 | 147.85 |
| 287 | 0.6942 | 0.00 | 0.04 | 147.09 |
| 288 | 0.6804 | 0.00 | 0.04 | 146.40 |
| 289 | 0.7061 | 0.00 | 0.04 | 145.72 |
| 290 | 0.6397 | 0.00 | 0.04 | 145.01 |
| 291 | 0.6577 | 0.00 | 0.04 | 144.37 |
| 292 | 0.6535 | 0.00 | 0.04 | 143.71 |
| 293 | 0.7253 | 0.00 | 0.04 | 143.06 |
| 294 | 0.6779 | 0.00 | 0.04 | 142.34 |
| 295 | 0.6699 | 0.00 | 0.04 | 141.66 |
| 296 | 0.6548 | 0.00 | 0.04 | 140.99 |
| 297 | 0.6853 | 0.00 | 0.05 | 140.33 |
| 298 | 0.6417 | 0.00 | 0.04 | 139.65 |
| 299 | 0.6631 | 0.00 | 0.04 | 139.01 |
| 300 | 0.6263 | 0.00 | 0.04 | 138.34 |
| 301 | 0.6119 | 0.00 | 0.04 | 137.72 |
| 302 | 0.6016 | 0.00 | 0.04 | 137.10 |
| 303 | 0.6000 | 0.00 | 0.04 | 136.50 |
| 304 | 0.5744 | 0.00 | 0.04 | 135.90 |
| 305 | 0.6240 | 0.00 | 0.04 | 135.33 |
| 306 | 0.5952 | 0.00 | 0.04 | 134.70 |
| 307 | 0.5824 | 0.00 | 0.04 | 134.11 |
| 308 | 0.5913 | 0.00 | 0.04 | 133.53 |

| | | | | |
|---|---|---|---|---|
| 309 | 0.5888 | 0.00 | 0.04 | 132.94 |
| 310 | 0.5657 | 0.00 | 0.04 | 132.35 |
| 311 | 0.5923 | 0.00 | 0.04 | 131.78 |
| 312 | 0.5862 | 0.00 | 0.04 | 131.19 |
| 313 | 0.5603 | 0.00 | 0.04 | 130.60 |
| 314 | 0.5452 | 0.00 | 0.04 | 130.04 |
| 315 | 0.5859 | 0.00 | 0.04 | 129.50 |
| 316 | 0.5660 | 0.00 | 0.04 | 128.91 |
| 317 | 0.5593 | 0.00 | 0.04 | 128.35 |
| 318 | 0.5894 | 0.00 | 0.04 | 127.79 |
| 319 | 0.5481 | 0.00 | 0.04 | 127.20 |
| 320 | 0.5433 | 0.00 | 0.03 | 126.65 |
| 321 | 0.5554 | 0.00 | 0.04 | 126.11 |
| 322 | 0.5446 | 0.00 | 0.03 | 125.55 |
| 323 | 0.5327 | 0.00 | 0.03 | 125.01 |
| 324 | 0.5119 | 0.00 | 0.03 | 124.47 |
| 325 | 0.5006 | 0.00 | 0.03 | 123.96 |
| 326 | 0.5215 | 0.00 | 0.03 | 123.46 |
| 327 | 0.4978 | 0.00 | 0.03 | 122.94 |
| 328 | 0.5276 | 0.00 | 0.03 | 122.44 |
| 329 | 0.5311 | 0.00 | 0.03 | 121.91 |
| 330 | 0.5103 | 0.00 | 0.03 | 121.38 |
| 331 | 0.5006 | 0.00 | 0.03 | 120.87 |
| 332 | 0.5016 | 0.00 | 0.03 | 120.37 |
| 333 | 0.4750 | 0.00 | 0.03 | 119.87 |
| 334 | 0.4897 | 0.00 | 0.03 | 119.39 |
| 335 | 0.4660 | 0.00 | 0.03 | 118.91 |
| 336 | 0.4740 | 0.00 | 0.03 | 118.44 |
| 337 | 0.4968 | 0.00 | 0.03 | 117.97 |
| 338 | 0.5170 | 0.00 | 0.04 | 117.47 |
| 339 | 0.5234 | 0.00 | 0.04 | 116.95 |
| 340 | 0.4939 | 0.00 | 0.03 | 116.43 |
| 341 | 0.4497 | 0.00 | 0.03 | 115.93 |
| 342 | 0.4474 | 0.00 | 0.03 | 115.48 |
| 343 | 0.4724 | 0.00 | 0.03 | 115.04 |
| 344 | 0.4817 | 0.00 | 0.04 | 114.56 |
| 345 | 0.4468 | 0.00 | 0.03 | 114.08 |
| 346 | 0.4670 | 0.00 | 0.03 | 113.64 |
| 347 | 0.4500 | 0.00 | 0.03 | 113.17 |
| 348 | 0.4510 | 0.00 | 0.03 | 112.72 |
| 349 | 0.4397 | 0.00 | 0.03 | 112.27 |
| 350 | 0.4324 | 0.00 | 0.03 | 111.83 |
| 351 | 0.4212 | 0.00 | 0.03 | 111.40 |
| 352 | 0.4401 | 0.00 | 0.03 | 110.97 |

| | | | | |
|---|---|---|---|---|
| 353 | 0.4163 | 0.00 | 0.03 | 110.53 |
| 354 | 0.4179 | 0.00 | 0.03 | 110.12 |
| 355 | 0.4250 | 0.00 | 0.03 | 109.70 |
| 356 | 0.3974 | 0.00 | 0.03 | 109.28 |
| 357 | 0.4173 | 0.00 | 0.03 | 108.88 |
| 358 | 0.4096 | 0.00 | 0.03 | 108.46 |
| 359 | 0.4029 | 0.00 | 0.03 | 108.05 |
| 360 | 0.4061 | 0.00 | 0.03 | 107.65 |
| 361 | 0.4163 | 0.00 | 0.03 | 107.24 |
| 362 | 0.3994 | 0.00 | 0.03 | 106.83 |
| 363 | 0.3901 | 0.00 | 0.03 | 106.43 |
| 364 | 0.4022 | 0.00 | 0.03 | 106.04 |
| 365 | 0.3686 | 0.00 | 0.03 | 105.63 |
| 366 | 0.3631 | 0.00 | 0.03 | 105.27 |
| 367 | 0.4138 | 0.00 | 0.03 | 104.90 |
| 368 | 0.3728 | 0.00 | 0.03 | 104.49 |
| 369 | 0.3872 | 0.00 | 0.03 | 104.12 |
| 370 | 0.3663 | 0.00 | 0.02 | 103.73 |
| 371 | 0.3667 | 0.00 | 0.03 | 103.36 |
| 372 | 0.3660 | 0.00 | 0.02 | 103.00 |
| 373 | 0.3644 | 0.00 | 0.03 | 102.63 |
| 374 | 0.3548 | 0.00 | 0.02 | 102.27 |
| 375 | 0.3567 | 0.00 | 0.02 | 101.91 |
| 376 | 0.3612 | 0.00 | 0.02 | 101.55 |
| 377 | 0.3651 | 0.00 | 0.03 | 101.19 |
| 378 | 0.3471 | 0.00 | 0.02 | 100.83 |
| 379 | 0.3462 | 0.00 | 0.02 | 100.48 |
| 380 | 0.3481 | 0.00 | 0.03 | 100.13 |
| 381 | 0.3606 | 0.00 | 0.02 | 99.79 |
| 382 | 0.3423 | 0.00 | 0.02 | 99.43 |
| 383 | 0.3465 | 0.00 | 0.02 | 99.08 |
| 384 | 0.3577 | 0.00 | 0.03 | 98.74 |
| 385 | 0.3314 | 0.00 | 0.02 | 98.38 |
| 386 | 0.3301 | 0.00 | 0.02 | 98.05 |
| 387 | 0.3314 | 0.00 | 0.02 | 97.72 |
| 388 | 0.3160 | 0.00 | 0.02 | 97.39 |
| 389 | 0.3272 | 0.00 | 0.02 | 97.07 |
| 390 | 0.3385 | 0.00 | 0.02 | 96.74 |
| 391 | 0.3337 | 0.00 | 0.02 | 96.40 |
| 392 | 0.3433 | 0.00 | 0.02 | 96.07 |
| 393 | 0.3314 | 0.00 | 0.02 | 95.73 |
| 394 | 0.3529 | 0.00 | 0.02 | 95.40 |
| 395 | 0.3401 | 0.00 | 0.02 | 95.04 |
| 396 | 0.3263 | 0.00 | 0.02 | 94.70 |

| | | | | |
|---|---|---|---|---|
| 397 | 0.3224 | 0.00 | 0.02 | 94.38 |
| 398 | 0.3087 | 0.00 | 0.02 | 94.05 |
| 399 | 0.3276 | 0.00 | 0.02 | 93.75 |
| 400 | 0.3359 | 0.00 | 0.02 | 93.42 |
| 401 | 0.3250 | 0.00 | 0.02 | 93.08 |
| 402 | 0.3205 | 0.00 | 0.02 | 92.76 |
| 403 | 0.3215 | 0.00 | 0.02 | 92.44 |
| 404 | 0.3228 | 0.00 | 0.02 | 92.12 |
| 405 | 0.2859 | 0.00 | 0.02 | 91.79 |
| 406 | 0.2885 | 0.00 | 0.02 | 91.51 |
| 407 | 0.2875 | 0.00 | 0.02 | 91.22 |
| 408 | 0.3042 | 0.00 | 0.02 | 90.93 |
| 409 | 0.2939 | 0.00 | 0.02 | 90.63 |
| 410 | 0.2837 | 0.00 | 0.02 | 90.33 |
| 411 | 0.2872 | 0.00 | 0.02 | 90.05 |
| 412 | 0.2702 | 0.00 | 0.02 | 89.76 |
| 413 | 0.2881 | 0.00 | 0.02 | 89.49 |
| 414 | 0.2885 | 0.00 | 0.02 | 89.20 |
| 415 | 0.2894 | 0.00 | 0.02 | 88.91 |
| 416 | 0.2788 | 0.00 | 0.02 | 88.63 |
| 417 | 0.2913 | 0.00 | 0.02 | 88.35 |
| 418 | 0.2724 | 0.00 | 0.02 | 88.06 |
| 419 | 0.2548 | 0.00 | 0.02 | 87.78 |
| 420 | 0.2628 | 0.00 | 0.02 | 87.53 |
| 421 | 0.2580 | 0.00 | 0.02 | 87.27 |
| 422 | 0.2593 | 0.00 | 0.02 | 87.01 |
| 423 | 0.2577 | 0.00 | 0.02 | 86.75 |
| 424 | 0.2631 | 0.00 | 0.02 | 86.49 |
| 425 | 0.2631 | 0.00 | 0.02 | 86.23 |
| 426 | 0.2580 | 0.00 | 0.02 | 85.96 |
| 427 | 0.2529 | 0.00 | 0.02 | 85.71 |
| 428 | 0.2487 | 0.00 | 0.02 | 85.45 |
| 429 | 0.2641 | 0.00 | 0.02 | 85.20 |
| 430 | 0.2596 | 0.00 | 0.02 | 84.94 |
| 431 | 0.2683 | 0.00 | 0.02 | 84.68 |
| 432 | 0.2647 | 0.00 | 0.02 | 84.41 |
| 433 | 0.2244 | 0.00 | 0.02 | 84.15 |
| 434 | 0.2324 | 0.00 | 0.02 | 83.92 |
| 435 | 0.2522 | 0.00 | 0.02 | 83.69 |
| 436 | 0.2372 | 0.00 | 0.02 | 83.44 |
| 437 | 0.2542 | 0.00 | 0.02 | 83.20 |
| 438 | 0.2369 | 0.00 | 0.02 | 82.95 |
| 439 | 0.2548 | 0.00 | 0.02 | 82.71 |
| 440 | 0.2532 | 0.00 | 0.02 | 82.46 |

| | | | | |
|---|---|---|---|---|
| 441 | 0.2462 | 0.00 | 0.02 | 82.20 |
| 442 | 0.2522 | 0.00 | 0.02 | 81.96 |
| 443 | 0.2365 | 0.00 | 0.02 | 81.70 |
| 444 | 0.2269 | 0.00 | 0.02 | 81.47 |
| 445 | 0.2266 | 0.00 | 0.02 | 81.24 |
| 446 | 0.2285 | 0.00 | 0.02 | 81.01 |
| 447 | 0.2436 | 0.00 | 0.02 | 80.79 |
| 448 | 0.2359 | 0.00 | 0.02 | 80.54 |
| 449 | 0.2266 | 0.00 | 0.02 | 80.31 |
| 450 | 0.2346 | 0.00 | 0.02 | 80.08 |
| 451 | 0.2266 | 0.00 | 0.02 | 79.84 |
| 452 | 0.2516 | 0.00 | 0.02 | 79.62 |
| 453 | 0.2212 | 0.00 | 0.02 | 79.37 |
| 454 | 0.2381 | 0.00 | 0.02 | 79.15 |
| 455 | 0.2349 | 0.00 | 0.02 | 78.91 |
| 456 | 0.2298 | 0.00 | 0.02 | 78.67 |
| 457 | 0.2231 | 0.00 | 0.02 | 78.44 |
| 458 | 0.2029 | 0.00 | 0.02 | 78.22 |
| 459 | 0.2189 | 0.00 | 0.02 | 78.02 |
| 460 | 0.2311 | 0.00 | 0.02 | 77.80 |
| 461 | 0.2189 | 0.00 | 0.02 | 77.57 |
| 462 | 0.2096 | 0.00 | 0.02 | 77.35 |
| 463 | 0.2054 | 0.00 | 0.02 | 77.14 |
| 464 | 0.2221 | 0.00 | 0.02 | 76.93 |
| 465 | 0.2170 | 0.00 | 0.02 | 76.71 |
| 466 | 0.2051 | 0.00 | 0.02 | 76.49 |
| 467 | 0.2083 | 0.00 | 0.02 | 76.29 |
| 468 | 0.2051 | 0.00 | 0.02 | 76.08 |
| 469 | 0.2051 | 0.00 | 0.02 | 75.87 |
| 470 | 0.2006 | 0.00 | 0.02 | 75.67 |
| 471 | 0.2010 | 0.00 | 0.02 | 75.47 |
| 472 | 0.1974 | 0.00 | 0.01 | 75.27 |
| 473 | 0.1917 | 0.00 | 0.01 | 75.07 |
| 474 | 0.2042 | 0.00 | 0.02 | 74.88 |
| 475 | 0.1881 | 0.00 | 0.02 | 74.67 |
| 476 | 0.2032 | 0.00 | 0.02 | 74.49 |
| 477 | 0.1987 | 0.00 | 0.02 | 74.28 |
| 478 | 0.1974 | 0.00 | 0.02 | 74.08 |
| 479 | 0.2071 | 0.00 | 0.02 | 73.89 |
| 480 | 0.2029 | 0.00 | 0.02 | 73.68 |
| 481 | 0.2022 | 0.00 | 0.02 | 73.48 |
| 482 | 0.1904 | 0.00 | 0.02 | 73.28 |
| 483 | 0.1885 | 0.00 | 0.02 | 73.08 |
| 484 | 0.2026 | 0.00 | 0.02 | 72.90 |

| | | | | |
|---|---|---|---|---|
| 485 | 0.1904 | 0.00 | 0.02 | 72.69 |
| 486 | 0.1946 | 0.00 | 0.02 | 72.50 |
| 487 | 0.1923 | 0.00 | 0.02 | 72.31 |
| 488 | 0.1994 | 0.00 | 0.02 | 72.12 |
| 489 | 0.1785 | 0.00 | 0.02 | 71.92 |
| 490 | 0.1885 | 0.00 | 0.02 | 71.74 |
| 491 | 0.2019 | 0.00 | 0.02 | 71.55 |
| 492 | 0.1788 | 0.00 | 0.02 | 71.35 |
| 493 | 0.1699 | 0.00 | 0.01 | 71.17 |
| 494 | 0.1891 | 0.00 | 0.02 | 71.00 |
| 495 | 0.1939 | 0.00 | 0.02 | 70.81 |
| 496 | 0.1679 | 0.00 | 0.01 | 70.62 |
| 497 | 0.1853 | 0.00 | 0.02 | 70.45 |
| 498 | 0.1683 | 0.00 | 0.01 | 70.26 |
| 499 | 0.1769 | 0.00 | 0.02 | 70.09 |
| 500 | 0.1667 | 0.00 | 0.01 | 69.92 |
| 501 | 0.1683 | 0.00 | 0.01 | 69.75 |
| 502 | 0.1731 | 0.00 | 0.01 | 69.58 |
| 503 | 0.1772 | 0.00 | 0.02 | 69.41 |
| 504 | 0.1833 | 0.00 | 0.01 | 69.23 |
| 505 | 0.1753 | 0.00 | 0.01 | 69.05 |
| 506 | 0.1712 | 0.00 | 0.01 | 68.87 |
| 507 | 0.1542 | 0.00 | 0.01 | 68.70 |
| 508 | 0.1840 | 0.00 | 0.02 | 68.55 |
| 509 | 0.1663 | 0.00 | 0.01 | 68.36 |
| 510 | 0.1590 | 0.00 | 0.01 | 68.20 |
| 511 | 0.1494 | 0.00 | 0.01 | 68.04 |
| 512 | 0.1692 | 0.00 | 0.02 | 67.89 |
| 513 | 0.1663 | 0.00 | 0.01 | 67.72 |
| 514 | 0.1712 | 0.00 | 0.01 | 67.55 |
| 515 | 0.1529 | 0.00 | 0.01 | 67.38 |
| 516 | 0.1545 | 0.00 | 0.01 | 67.23 |
| 517 | 0.1689 | 0.00 | 0.01 | 67.08 |
| 518 | 0.1494 | 0.00 | 0.01 | 66.91 |
| 519 | 0.1452 | 0.00 | 0.01 | 66.76 |
| 520 | 0.1631 | 0.00 | 0.01 | 66.61 |
| 521 | 0.1660 | 0.00 | 0.01 | 66.45 |
| 522 | 0.1574 | 0.00 | 0.01 | 66.28 |
| 523 | 0.1538 | 0.00 | 0.01 | 66.13 |
| 524 | 0.1538 | 0.00 | 0.01 | 65.97 |
| 525 | 0.1651 | 0.00 | 0.01 | 65.82 |
| 526 | 0.1673 | 0.00 | 0.01 | 65.65 |
| 527 | 0.1564 | 0.00 | 0.01 | 65.49 |
| 528 | 0.1580 | 0.00 | 0.01 | 65.33 |

| | | | | |
|---|---|---|---|---|
| 529 | 0.1442 | 0.00 | 0.01 | 65.17 |
| 530 | 0.1696 | 0.00 | 0.01 | 65.03 |
| 531 | 0.1571 | 0.00 | 0.01 | 64.86 |
| 532 | 0.1686 | 0.00 | 0.01 | 64.70 |
| 533 | 0.1478 | 0.00 | 0.01 | 64.53 |
| 534 | 0.1679 | 0.00 | 0.01 | 64.38 |
| 535 | 0.1583 | 0.00 | 0.01 | 64.22 |
| 536 | 0.1712 | 0.00 | 0.01 | 64.06 |
| 537 | 0.1596 | 0.00 | 0.02 | 63.89 |
| 538 | 0.1580 | 0.00 | 0.01 | 63.73 |
| 539 | 0.1548 | 0.00 | 0.01 | 63.57 |
| 540 | 0.1426 | 0.00 | 0.01 | 63.41 |
| 541 | 0.1715 | 0.00 | 0.02 | 63.27 |
| 542 | 0.1622 | 0.00 | 0.02 | 63.10 |
| 543 | 0.1587 | 0.00 | 0.01 | 62.94 |
| 544 | 0.1545 | 0.00 | 0.01 | 62.78 |
| 545 | 0.1494 | 0.00 | 0.01 | 62.63 |
| 546 | 0.1526 | 0.00 | 0.01 | 62.48 |
| 547 | 0.1619 | 0.00 | 0.02 | 62.32 |
| 548 | 0.1631 | 0.00 | 0.01 | 62.16 |
| 549 | 0.1628 | 0.00 | 0.02 | 62.00 |
| 550 | 0.1571 | 0.00 | 0.02 | 61.84 |
| 551 | 0.1446 | 0.00 | 0.01 | 61.68 |
| 552 | 0.1212 | 0.00 | 0.01 | 61.53 |
| 553 | 0.1503 | 0.00 | 0.01 | 61.41 |
| 554 | 0.1503 | 0.00 | 0.01 | 61.26 |
| 555 | 0.1455 | 0.00 | 0.02 | 61.11 |
| 556 | 0.1417 | 0.00 | 0.01 | 60.97 |
| 557 | 0.1462 | 0.00 | 0.01 | 60.82 |
| 558 | 0.1516 | 0.00 | 0.01 | 60.68 |
| 559 | 0.1385 | 0.00 | 0.01 | 60.53 |
| 560 | 0.1423 | 0.00 | 0.01 | 60.39 |
| 561 | 0.1330 | 0.00 | 0.01 | 60.25 |
| 562 | 0.1468 | 0.00 | 0.01 | 60.11 |
| 563 | 0.1282 | 0.00 | 0.01 | 59.97 |
| 564 | 0.1375 | 0.00 | 0.01 | 59.84 |
| 565 | 0.1372 | 0.00 | 0.01 | 59.70 |
| 566 | 0.1356 | 0.00 | 0.01 | 59.56 |
| 567 | 0.1224 | 0.00 | 0.01 | 59.43 |
| 568 | 0.1330 | 0.00 | 0.01 | 59.31 |
| 569 | 0.1359 | 0.00 | 0.01 | 59.17 |
| 570 | 0.1298 | 0.00 | 0.01 | 59.04 |
| 571 | 0.1503 | 0.00 | 0.01 | 58.91 |
| 572 | 0.1308 | 0.00 | 0.01 | 58.76 |

| | | | | |
|---|---|---|---|---|
| 573 | 0.1340 | 0.00 | 0.01 | 58.63 |
| 574 | 0.1256 | 0.00 | 0.01 | 58.49 |
| 575 | 0.1446 | 0.00 | 0.02 | 58.37 |
| 576 | 0.1311 | 0.00 | 0.01 | 58.22 |
| 577 | 0.1199 | 0.00 | 0.01 | 58.09 |
| 578 | 0.1170 | 0.00 | 0.01 | 57.97 |
| 579 | 0.1256 | 0.00 | 0.01 | 57.85 |
| 580 | 0.1163 | 0.00 | 0.01 | 57.73 |
| 581 | 0.1231 | 0.00 | 0.01 | 57.61 |
| 582 | 0.1256 | 0.00 | 0.01 | 57.49 |
| 583 | 0.1231 | 0.00 | 0.01 | 57.36 |
| 584 | 0.1138 | 0.00 | 0.01 | 57.24 |
| 585 | 0.1135 | 0.00 | 0.01 | 57.13 |
| 586 | 0.1215 | 0.00 | 0.01 | 57.01 |
| 587 | 0.1119 | 0.00 | 0.01 | 56.89 |
| 588 | 0.1176 | 0.00 | 0.01 | 56.78 |
| 589 | 0.1119 | 0.00 | 0.01 | 56.66 |
| 590 | 0.1173 | 0.00 | 0.01 | 56.55 |
| 591 | 0.1119 | 0.00 | 0.01 | 56.43 |
| 592 | 0.1192 | 0.00 | 0.01 | 56.32 |
| 593 | 0.1071 | 0.00 | 0.01 | 56.20 |
| 594 | 0.0946 | 0.00 | 0.01 | 56.09 |
| 595 | 0.1163 | 0.00 | 0.01 | 56.00 |
| 596 | 0.1192 | 0.00 | 0.01 | 55.88 |
| 597 | 0.1125 | 0.00 | 0.01 | 55.76 |
| 598 | 0.1138 | 0.00 | 0.01 | 55.65 |
| 599 | 0.1099 | 0.00 | 0.01 | 55.54 |
| 600 | 0.1167 | 0.00 | 0.01 | 55.43 |
| 601 | 0.1324 | 0.00 | 0.02 | 55.31 |
| 602 | 0.1029 | 0.00 | 0.01 | 55.18 |
| 603 | 0.1013 | 0.00 | 0.01 | 55.08 |
| 604 | 0.1115 | 0.00 | 0.01 | 54.97 |
| 605 | 0.0971 | 0.00 | 0.01 | 54.86 |
| 606 | 0.1288 | 0.00 | 0.02 | 54.77 |
| 607 | 0.0968 | 0.00 | 0.01 | 54.64 |
| 608 | 0.1163 | 0.00 | 0.01 | 54.54 |
| 609 | 0.1058 | 0.00 | 0.01 | 54.42 |
| 610 | 0.1096 | 0.00 | 0.01 | 54.32 |
| 611 | 0.1067 | 0.00 | 0.01 | 54.21 |
| 612 | 0.1067 | 0.00 | 0.01 | 54.10 |
| 613 | 0.0939 | 0.00 | 0.01 | 54.00 |
| 614 | 0.0907 | 0.00 | 0.01 | 53.90 |
| 615 | 0.1141 | 0.00 | 0.01 | 53.81 |
| 616 | 0.1112 | 0.00 | 0.01 | 53.70 |

| | | | | |
|---|---|---|---|---|
| 617 | 0.1173 | 0.00 | 0.01 | 53.59 |
| 618 | 0.1035 | 0.00 | 0.01 | 53.47 |
| 619 | 0.0984 | 0.00 | 0.01 | 53.36 |
| 620 | 0.1163 | 0.00 | 0.01 | 53.27 |
| 621 | 0.1093 | 0.00 | 0.01 | 53.15 |
| 622 | 0.1029 | 0.00 | 0.01 | 53.04 |
| 623 | 0.1016 | 0.00 | 0.01 | 52.94 |
| 624 | 0.1051 | 0.00 | 0.01 | 52.84 |
| 625 | 0.0968 | 0.00 | 0.01 | 52.73 |
| 626 | 0.0933 | 0.00 | 0.01 | 52.63 |
| 627 | 0.0936 | 0.00 | 0.01 | 52.54 |
| 628 | 0.1000 | 0.00 | 0.01 | 52.45 |
| 629 | 0.0981 | 0.00 | 0.01 | 52.35 |
| 630 | 0.0987 | 0.00 | 0.01 | 52.25 |
| 631 | 0.1125 | 0.00 | 0.01 | 52.15 |
| 632 | 0.1038 | 0.00 | 0.01 | 52.04 |
| 633 | 0.0952 | 0.00 | 0.01 | 51.93 |
| 634 | 0.0990 | 0.00 | 0.01 | 51.84 |
| 635 | 0.1013 | 0.00 | 0.01 | 51.74 |
| 636 | 0.1061 | 0.00 | 0.01 | 51.64 |
| 637 | 0.1006 | 0.00 | 0.01 | 51.53 |
| 638 | 0.0926 | 0.00 | 0.01 | 51.43 |
| 639 | 0.1038 | 0.00 | 0.01 | 51.34 |
| 640 | 0.0984 | 0.00 | 0.01 | 51.24 |
| 641 | 0.0872 | 0.00 | 0.01 | 51.14 |
| 642 | 0.1016 | 0.00 | 0.01 | 51.05 |
| 643 | 0.0987 | 0.00 | 0.01 | 50.95 |
| 644 | 0.0808 | 0.00 | 0.01 | 50.85 |
| 645 | 0.0875 | 0.00 | 0.01 | 50.77 |
| 646 | 0.0856 | 0.00 | 0.01 | 50.68 |
| 647 | 0.0888 | 0.00 | 0.01 | 50.60 |
| 648 | 0.0939 | 0.00 | 0.01 | 50.51 |
| 649 | 0.1054 | 0.00 | 0.02 | 50.41 |
| 650 | 0.0881 | 0.00 | 0.01 | 50.31 |
| 651 | 0.0917 | 0.00 | 0.01 | 50.22 |
| 652 | 0.0785 | 0.00 | 0.01 | 50.13 |
| 653 | 0.0830 | 0.00 | 0.01 | 50.05 |
| 654 | 0.0894 | 0.00 | 0.01 | 49.97 |
| 655 | 0.0843 | 0.00 | 0.01 | 49.88 |
| 656 | 0.0862 | 0.00 | 0.01 | 49.79 |
| 657 | 0.0792 | 0.00 | 0.01 | 49.71 |
| 658 | 0.0833 | 0.00 | 0.01 | 49.63 |
| 659 | 0.0840 | 0.00 | 0.01 | 49.54 |
| 660 | 0.0753 | 0.00 | 0.01 | 49.46 |

| | | | | |
|---|---|---|---|---|
| 661 | 0.0715 | 0.00 | 0.01 | 49.38 |
| 662 | 0.0776 | 0.00 | 0.01 | 49.31 |
| 663 | 0.0795 | 0.00 | 0.01 | 49.24 |
| 664 | 0.0804 | 0.00 | 0.01 | 49.16 |
| 665 | 0.0821 | 0.00 | 0.01 | 49.08 |
| 666 | 0.0942 | 0.00 | 0.01 | 48.99 |
| 667 | 0.0923 | 0.00 | 0.02 | 48.90 |
| 668 | 0.0862 | 0.00 | 0.01 | 48.81 |
| 669 | 0.0772 | 0.00 | 0.01 | 48.72 |
| 670 | 0.0667 | 0.00 | 0.01 | 48.64 |
| 671 | 0.0801 | 0.00 | 0.01 | 48.58 |
| 672 | 0.0808 | 0.00 | 0.01 | 48.50 |
| 673 | 0.0737 | 0.00 | 0.01 | 48.42 |
| 674 | 0.0766 | 0.00 | 0.01 | 48.34 |
| 675 | 0.0840 | 0.00 | 0.01 | 48.27 |
| 676 | 0.0702 | 0.00 | 0.01 | 48.18 |
| 677 | 0.0811 | 0.00 | 0.01 | 48.11 |
| 678 | 0.0705 | 0.00 | 0.01 | 48.03 |
| 679 | 0.0753 | 0.00 | 0.01 | 47.96 |
| 680 | 0.0798 | 0.00 | 0.01 | 47.88 |
| 681 | 0.0833 | 0.00 | 0.01 | 47.80 |
| 682 | 0.0734 | 0.00 | 0.01 | 47.72 |
| 683 | 0.0926 | 0.00 | 0.02 | 47.65 |
| 684 | 0.0663 | 0.00 | 0.01 | 47.56 |
| 685 | 0.0676 | 0.00 | 0.01 | 47.49 |
| 686 | 0.0654 | 0.00 | 0.01 | 47.42 |
| 687 | 0.0663 | 0.00 | 0.01 | 47.36 |
| 688 | 0.0756 | 0.00 | 0.01 | 47.29 |
| 689 | 0.0779 | 0.00 | 0.01 | 47.21 |
| 690 | 0.0897 | 0.00 | 0.02 | 47.14 |
| 691 | 0.0708 | 0.00 | 0.01 | 47.05 |
| 692 | 0.0750 | 0.00 | 0.01 | 46.98 |
| 693 | 0.0718 | 0.00 | 0.01 | 46.90 |
| 694 | 0.0766 | 0.00 | 0.01 | 46.83 |
| 695 | 0.0724 | 0.00 | 0.01 | 46.75 |
| 696 | 0.0718 | 0.00 | 0.01 | 46.68 |
| 697 | 0.0679 | 0.00 | 0.01 | 46.61 |
| 698 | 0.0641 | 0.00 | 0.01 | 46.54 |
| 699 | 0.0827 | 0.00 | 0.01 | 46.48 |
| 700 | 0.0769 | 0.00 | 0.01 | 46.39 |
| 701 | 0.0647 | 0.00 | 0.01 | 46.32 |
| 702 | 0.0654 | 0.00 | 0.01 | 46.25 |
| 703 | 0.0667 | 0.00 | 0.01 | 46.19 |
| 704 | 0.0663 | 0.00 | 0.01 | 46.12 |

| | | | | |
|---|---|---|---|---|
| 705 | 0.0692 | 0.00 | 0.01 | 46.05 |
| 706 | 0.0756 | 0.00 | 0.01 | 45.98 |
| 707 | 0.0660 | 0.00 | 0.01 | 45.91 |
| 708 | 0.0641 | 0.00 | 0.01 | 45.84 |
| 709 | 0.0673 | 0.00 | 0.01 | 45.78 |
| 710 | 0.0577 | 0.00 | 0.01 | 45.71 |
| 711 | 0.0638 | 0.00 | 0.01 | 45.65 |
| 712 | 0.0615 | 0.00 | 0.01 | 45.59 |
| 713 | 0.0644 | 0.00 | 0.01 | 45.53 |
| 714 | 0.0651 | 0.00 | 0.01 | 45.46 |
| 715 | 0.0561 | 0.00 | 0.01 | 45.40 |
| 716 | 0.0644 | 0.00 | 0.01 | 45.34 |
| 717 | 0.0679 | 0.00 | 0.01 | 45.28 |
| 718 | 0.0647 | 0.00 | 0.01 | 45.21 |
| 719 | 0.0712 | 0.00 | 0.01 | 45.14 |
| 720 | 0.0718 | 0.00 | 0.01 | 45.07 |
| 721 | 0.0788 | 0.00 | 0.01 | 45.00 |
| 722 | 0.0843 | 0.00 | 0.01 | 44.92 |
| 723 | 0.0702 | 0.00 | 0.01 | 44.84 |
| 724 | 0.0638 | 0.00 | 0.01 | 44.77 |
| 725 | 0.0625 | 0.00 | 0.01 | 44.70 |
| 726 | 0.0689 | 0.00 | 0.01 | 44.64 |
| 727 | 0.0670 | 0.00 | 0.01 | 44.57 |
| 728 | 0.0622 | 0.00 | 0.01 | 44.51 |
| 729 | 0.0679 | 0.00 | 0.01 | 44.44 |
| 730 | 0.0657 | 0.00 | 0.01 | 44.38 |
| 731 | 0.0875 | 0.00 | 0.03 | 44.31 |
| 732 | 0.0712 | 0.00 | 0.01 | 44.22 |
| 733 | 0.0679 | 0.00 | 0.01 | 44.15 |
| 734 | 0.0692 | 0.00 | 0.01 | 44.08 |
| 735 | 0.0724 | 0.00 | 0.01 | 44.01 |
| 736 | 0.0686 | 0.00 | 0.01 | 43.94 |
| 737 | 0.0744 | 0.00 | 0.01 | 43.87 |
| 738 | 0.0574 | 0.00 | 0.01 | 43.80 |
| 739 | 0.0683 | 0.00 | 0.01 | 43.74 |
| 740 | 0.0564 | 0.00 | 0.01 | 43.67 |
| 741 | 0.0651 | 0.00 | 0.01 | 43.62 |
| 742 | 0.0792 | 0.00 | 0.02 | 43.55 |
| 743 | 0.0712 | 0.00 | 0.01 | 43.47 |
| 744 | 0.0766 | 0.00 | 0.01 | 43.40 |
| 745 | 0.0696 | 0.00 | 0.01 | 43.33 |
| 746 | 0.0638 | 0.00 | 0.01 | 43.26 |
| 747 | 0.0538 | 0.00 | 0.01 | 43.19 |
| 748 | 0.0574 | 0.00 | 0.01 | 43.14 |

| | | | | |
|---|---|---|---|---|
| 749 | 0.0814 | 0.00 | 0.01 | 43.08 |
| 750 | 0.0724 | 0.00 | 0.01 | 43.00 |
| 751 | 0.0615 | 0.00 | 0.01 | 42.93 |
| 752 | 0.0756 | 0.00 | 0.01 | 42.87 |
| 753 | 0.0606 | 0.00 | 0.01 | 42.79 |
| 754 | 0.0619 | 0.00 | 0.01 | 42.73 |
| 755 | 0.0625 | 0.00 | 0.01 | 42.67 |
| 756 | 0.0593 | 0.00 | 0.01 | 42.60 |
| 757 | 0.0718 | 0.00 | 0.01 | 42.55 |
| 758 | 0.0622 | 0.00 | 0.01 | 42.47 |
| 759 | 0.0587 | 0.00 | 0.01 | 42.41 |
| 760 | 0.0811 | 0.00 | 0.01 | 42.35 |
| 761 | 0.0654 | 0.00 | 0.01 | 42.27 |
| 762 | 0.0641 | 0.00 | 0.01 | 42.21 |
| 763 | 0.0615 | 0.00 | 0.01 | 42.14 |
| 764 | 0.0628 | 0.00 | 0.01 | 42.08 |
| 765 | 0.0574 | 0.00 | 0.01 | 42.02 |
| 766 | 0.0609 | 0.00 | 0.01 | 41.96 |
| 767 | 0.0628 | 0.00 | 0.01 | 41.90 |
| 768 | 0.0612 | 0.00 | 0.01 | 41.84 |
| 769 | 0.0628 | 0.00 | 0.01 | 41.78 |
| 770 | 0.0580 | 0.00 | 0.01 | 41.71 |
| 771 | 0.0558 | 0.00 | 0.01 | 41.65 |
| 772 | 0.0596 | 0.00 | 0.01 | 41.60 |
| 773 | 0.0535 | 0.00 | 0.01 | 41.54 |
| 774 | 0.0590 | 0.00 | 0.01 | 41.49 |
| 775 | 0.0689 | 0.00 | 0.01 | 41.43 |
| 776 | 0.0574 | 0.00 | 0.01 | 41.36 |
| 777 | 0.0542 | 0.00 | 0.01 | 41.30 |
| 778 | 0.0548 | 0.00 | 0.01 | 41.25 |
| 779 | 0.0615 | 0.00 | 0.01 | 41.19 |
| 780 | 0.0667 | 0.00 | 0.01 | 41.13 |
| 781 | 0.0548 | 0.00 | 0.01 | 41.06 |
| 782 | 0.0635 | 0.00 | 0.01 | 41.01 |
| 783 | 0.0522 | 0.00 | 0.01 | 40.95 |
| 784 | 0.0506 | 0.00 | 0.01 | 40.89 |
| 785 | 0.0603 | 0.00 | 0.01 | 40.84 |
| 786 | 0.0606 | 0.00 | 0.01 | 40.78 |
| 787 | 0.0487 | 0.00 | 0.01 | 40.72 |
| 788 | 0.0577 | 0.00 | 0.01 | 40.67 |
| 789 | 0.0538 | 0.00 | 0.01 | 40.62 |
| 790 | 0.0538 | 0.00 | 0.01 | 40.56 |
| 791 | 0.0503 | 0.00 | 0.01 | 40.51 |
| 792 | 0.0551 | 0.00 | 0.01 | 40.46 |

| | | | | |
|---|---|---|---|---|
| 793 | 0.0510 | 0.00 | 0.01 | 40.40 |
| 794 | 0.0484 | 0.00 | 0.01 | 40.35 |
| 795 | 0.0651 | 0.00 | 0.01 | 40.30 |
| 796 | 0.0558 | 0.00 | 0.01 | 40.24 |
| 797 | 0.0526 | 0.00 | 0.01 | 40.18 |
| 798 | 0.0516 | 0.00 | 0.01 | 40.13 |
| 799 | 0.0535 | 0.00 | 0.01 | 40.08 |
| 800 | 0.0561 | 0.00 | 0.01 | 40.02 |
| 801 | 0.0580 | 0.00 | 0.01 | 39.97 |
| 802 | 0.0506 | 0.00 | 0.01 | 39.91 |
| 803 | 0.0494 | 0.00 | 0.01 | 39.86 |
| 804 | 0.0535 | 0.00 | 0.01 | 39.81 |
| 805 | 0.0554 | 0.00 | 0.01 | 39.76 |
| 806 | 0.0606 | 0.00 | 0.01 | 39.70 |
| 807 | 0.0538 | 0.00 | 0.01 | 39.64 |
| 808 | 0.0554 | 0.00 | 0.01 | 39.59 |
| 809 | 0.0577 | 0.00 | 0.01 | 39.53 |
| 810 | 0.0558 | 0.00 | 0.01 | 39.47 |
| 811 | 0.0603 | 0.00 | 0.01 | 39.42 |
| 812 | 0.0462 | 0.00 | 0.01 | 39.36 |
| 813 | 0.0500 | 0.00 | 0.01 | 39.31 |
| 814 | 0.0478 | 0.00 | 0.01 | 39.26 |
| 815 | 0.0529 | 0.00 | 0.01 | 39.21 |
| 816 | 0.0484 | 0.00 | 0.01 | 39.16 |
| 817 | 0.0497 | 0.00 | 0.01 | 39.11 |
| 818 | 0.0471 | 0.00 | 0.01 | 39.06 |
| 819 | 0.0484 | 0.00 | 0.01 | 39.02 |
| 820 | 0.0529 | 0.00 | 0.01 | 38.97 |
| 821 | 0.0452 | 0.00 | 0.01 | 38.91 |
| 822 | 0.0372 | 0.00 | 0.00 | 38.87 |
| 823 | 0.0449 | 0.00 | 0.01 | 38.83 |
| 824 | 0.0474 | 0.00 | 0.01 | 38.79 |
| 825 | 0.0490 | 0.00 | 0.01 | 38.74 |
| 826 | 0.0426 | 0.00 | 0.01 | 38.69 |
| 827 | 0.0446 | 0.00 | 0.01 | 38.65 |
| 828 | 0.0503 | 0.00 | 0.01 | 38.60 |
| 829 | 0.0404 | 0.00 | 0.00 | 38.55 |
| 830 | 0.0471 | 0.00 | 0.01 | 38.51 |
| 831 | 0.0458 | 0.00 | 0.01 | 38.47 |
| 832 | 0.0433 | 0.00 | 0.01 | 38.42 |
| 833 | 0.0478 | 0.00 | 0.01 | 38.38 |
| 834 | 0.0471 | 0.00 | 0.01 | 38.33 |
| 835 | 0.0503 | 0.00 | 0.01 | 38.28 |
| 836 | 0.0478 | 0.00 | 0.01 | 38.23 |

| | | | | |
|---|---|---|---|---|
| 837 | 0.0526 | 0.00 | 0.01 | 38.18 |
| 838 | 0.0487 | 0.00 | 0.01 | 38.13 |
| 839 | 0.0548 | 0.00 | 0.01 | 38.08 |
| 840 | 0.0474 | 0.00 | 0.01 | 38.03 |
| 841 | 0.0446 | 0.00 | 0.01 | 37.98 |
| 842 | 0.0558 | 0.00 | 0.01 | 37.94 |
| 843 | 0.0487 | 0.00 | 0.01 | 37.88 |
| 844 | 0.0436 | 0.00 | 0.01 | 37.83 |
| 845 | 0.0385 | 0.00 | 0.00 | 37.79 |
| 846 | 0.0567 | 0.00 | 0.01 | 37.75 |
| 847 | 0.0635 | 0.00 | 0.02 | 37.69 |
| 848 | 0.0446 | 0.00 | 0.01 | 37.63 |
| 849 | 0.0442 | 0.00 | 0.01 | 37.58 |
| 850 | 0.0532 | 0.00 | 0.01 | 37.54 |
| 851 | 0.0481 | 0.00 | 0.01 | 37.49 |
| 852 | 0.0522 | 0.00 | 0.01 | 37.44 |
| 853 | 0.0500 | 0.00 | 0.01 | 37.39 |
| 854 | 0.0468 | 0.00 | 0.01 | 37.34 |
| 855 | 0.0481 | 0.00 | 0.01 | 37.29 |
| 856 | 0.0426 | 0.00 | 0.01 | 37.24 |
| 857 | 0.0449 | 0.00 | 0.01 | 37.20 |
| 858 | 0.0522 | 0.00 | 0.01 | 37.15 |
| 859 | 0.0449 | 0.00 | 0.01 | 37.10 |
| 860 | 0.0481 | 0.00 | 0.01 | 37.06 |
| 861 | 0.0500 | 0.00 | 0.01 | 37.01 |
| 862 | 0.0478 | 0.00 | 0.01 | 36.96 |
| 863 | 0.0401 | 0.00 | 0.01 | 36.91 |
| 864 | 0.0429 | 0.00 | 0.01 | 36.87 |
| 865 | 0.0429 | 0.00 | 0.01 | 36.83 |
| 866 | 0.0439 | 0.00 | 0.01 | 36.78 |
| 867 | 0.0494 | 0.00 | 0.01 | 36.74 |
| 868 | 0.0462 | 0.00 | 0.01 | 36.69 |
| 869 | 0.0423 | 0.00 | 0.01 | 36.65 |
| 870 | 0.0349 | 0.00 | 0.00 | 36.60 |
| 871 | 0.0471 | 0.00 | 0.01 | 36.57 |
| 872 | 0.0413 | 0.00 | 0.01 | 36.52 |
| 873 | 0.0436 | 0.00 | 0.01 | 36.48 |
| 874 | 0.0375 | 0.00 | 0.00 | 36.44 |
| 875 | 0.0356 | 0.00 | 0.00 | 36.40 |
| 876 | 0.0381 | 0.00 | 0.01 | 36.36 |
| 877 | 0.0410 | 0.00 | 0.01 | 36.33 |
| 878 | 0.0410 | 0.00 | 0.01 | 36.28 |
| 879 | 0.0372 | 0.00 | 0.01 | 36.24 |
| 880 | 0.0369 | 0.00 | 0.00 | 36.21 |

| | | | | |
|---|---|---|---|---|
| 881 | 0.0394 | 0.00 | 0.00 | 36.17 |
| 882 | 0.0436 | 0.00 | 0.01 | 36.13 |
| 883 | 0.0429 | 0.00 | 0.01 | 36.09 |
| 884 | 0.0481 | 0.00 | 0.01 | 36.04 |
| 885 | 0.0423 | 0.00 | 0.01 | 35.99 |
| 886 | 0.0494 | 0.00 | 0.01 | 35.95 |
| 887 | 0.0426 | 0.00 | 0.01 | 35.90 |
| 888 | 0.0381 | 0.00 | 0.00 | 35.86 |
| 889 | 0.0394 | 0.00 | 0.01 | 35.82 |
| 890 | 0.0433 | 0.00 | 0.01 | 35.78 |
| 891 | 0.0391 | 0.00 | 0.01 | 35.74 |
| 892 | 0.0417 | 0.00 | 0.00 | 35.70 |
| 893 | 0.0337 | 0.00 | 0.00 | 35.66 |
| 894 | 0.0391 | 0.00 | 0.00 | 35.63 |
| 895 | 0.0340 | 0.00 | 0.00 | 35.59 |
| 896 | 0.0349 | 0.00 | 0.00 | 35.55 |
| 897 | 0.0401 | 0.00 | 0.01 | 35.52 |
| 898 | 0.0407 | 0.00 | 0.00 | 35.48 |
| 899 | 0.0420 | 0.00 | 0.01 | 35.44 |
| 900 | 0.0436 | 0.00 | 0.01 | 35.39 |
| 901 | 0.0417 | 0.00 | 0.01 | 35.35 |
| 902 | 0.0378 | 0.00 | 0.00 | 35.31 |
| 903 | 0.0375 | 0.00 | 0.00 | 35.27 |
| 904 | 0.0452 | 0.00 | 0.01 | 35.23 |
| 905 | 0.0391 | 0.00 | 0.00 | 35.19 |
| 906 | 0.0436 | 0.00 | 0.01 | 35.15 |
| 907 | 0.0356 | 0.00 | 0.00 | 35.11 |
| 908 | 0.0330 | 0.00 | 0.00 | 35.07 |
| 909 | 0.0404 | 0.00 | 0.01 | 35.04 |
| 910 | 0.0333 | 0.00 | 0.00 | 35.00 |
| 911 | 0.0295 | 0.00 | 0.00 | 34.96 |
| 912 | 0.0356 | 0.00 | 0.00 | 34.93 |
| 913 | 0.0381 | 0.00 | 0.00 | 34.90 |
| 914 | 0.0298 | 0.00 | 0.00 | 34.86 |
| 915 | 0.0365 | 0.00 | 0.00 | 34.83 |
| 916 | 0.0308 | 0.00 | 0.00 | 34.79 |
| 917 | 0.0359 | 0.00 | 0.00 | 34.76 |
| 918 | 0.0365 | 0.00 | 0.00 | 34.73 |
| 919 | 0.0324 | 0.00 | 0.00 | 34.69 |
| 920 | 0.0365 | 0.00 | 0.01 | 34.66 |
| 921 | 0.0343 | 0.00 | 0.00 | 34.62 |
| 922 | 0.0372 | 0.00 | 0.00 | 34.59 |
| 923 | 0.0394 | 0.00 | 0.01 | 34.55 |
| 924 | 0.0401 | 0.00 | 0.01 | 34.51 |

| | | | | |
|---|---|---|---|---|
| 925 | 0.0417 | 0.00 | 0.01 | 34.47 |
| 926 | 0.0429 | 0.00 | 0.01 | 34.43 |
| 927 | 0.0301 | 0.00 | 0.00 | 34.39 |
| 928 | 0.0321 | 0.00 | 0.00 | 34.36 |
| 929 | 0.0337 | 0.00 | 0.00 | 34.32 |
| 930 | 0.0375 | 0.00 | 0.00 | 34.29 |
| 931 | 0.0337 | 0.00 | 0.00 | 34.25 |
| 932 | 0.0404 | 0.00 | 0.01 | 34.22 |
| 933 | 0.0394 | 0.00 | 0.01 | 34.18 |
| 934 | 0.0394 | 0.00 | 0.01 | 34.14 |
| 935 | 0.0327 | 0.00 | 0.00 | 34.10 |
| 936 | 0.0298 | 0.00 | 0.00 | 34.07 |
| 937 | 0.0327 | 0.00 | 0.00 | 34.04 |
| 938 | 0.0349 | 0.00 | 0.00 | 34.01 |
| 939 | 0.0324 | 0.00 | 0.00 | 33.97 |
| 940 | 0.0362 | 0.00 | 0.01 | 33.94 |
| 941 | 0.0304 | 0.00 | 0.00 | 33.90 |
| 942 | 0.0285 | 0.00 | 0.00 | 33.87 |
| 943 | 0.0298 | 0.00 | 0.00 | 33.84 |
| 944 | 0.0250 | 0.00 | 0.00 | 33.81 |
| 945 | 0.0385 | 0.00 | 0.00 | 33.79 |
| 946 | 0.0324 | 0.00 | 0.00 | 33.75 |
| 947 | 0.0337 | 0.00 | 0.00 | 33.72 |
| 948 | 0.0346 | 0.00 | 0.00 | 33.68 |
| 949 | 0.0301 | 0.00 | 0.00 | 33.65 |
| 950 | 0.0413 | 0.00 | 0.01 | 33.62 |
| 951 | 0.0327 | 0.00 | 0.00 | 33.58 |
| 952 | 0.0349 | 0.00 | 0.00 | 33.54 |
| 953 | 0.0304 | 0.00 | 0.00 | 33.51 |
| 954 | 0.0381 | 0.00 | 0.01 | 33.48 |
| 955 | 0.0311 | 0.00 | 0.00 | 33.44 |
| 956 | 0.0288 | 0.00 | 0.00 | 33.41 |
| 957 | 0.0346 | 0.00 | 0.00 | 33.38 |
| 958 | 0.0378 | 0.00 | 0.00 | 33.35 |
| 959 | 0.0308 | 0.00 | 0.00 | 33.31 |
| 960 | 0.0362 | 0.00 | 0.00 | 33.28 |
| 961 | 0.0359 | 0.00 | 0.00 | 33.24 |
| 962 | 0.0349 | 0.00 | 0.00 | 33.21 |
| 963 | 0.0324 | 0.00 | 0.00 | 33.17 |
| 964 | 0.0337 | 0.00 | 0.00 | 33.14 |
| 965 | 0.0359 | 0.00 | 0.00 | 33.10 |
| 966 | 0.0343 | 0.00 | 0.00 | 33.07 |
| 967 | 0.0301 | 0.00 | 0.00 | 33.03 |
| 968 | 0.0269 | 0.00 | 0.00 | 33.00 |

| | | | | |
|---|---|---|---|---|
| 969 | 0.0353 | 0.00 | 0.00 | 32.98 |
| 970 | 0.0327 | 0.00 | 0.00 | 32.94 |
| 971 | 0.0346 | 0.00 | 0.00 | 32.91 |
| 972 | 0.0356 | 0.00 | 0.00 | 32.88 |
| 973 | 0.0442 | 0.00 | 0.01 | 32.84 |
| 974 | 0.0365 | 0.00 | 0.00 | 32.80 |
| 975 | 0.0385 | 0.00 | 0.01 | 32.76 |
| 976 | 0.0298 | 0.00 | 0.00 | 32.72 |
| 977 | 0.0317 | 0.00 | 0.00 | 32.69 |
| 978 | 0.0256 | 0.00 | 0.00 | 32.66 |
| 979 | 0.0301 | 0.00 | 0.00 | 32.63 |
| 980 | 0.0298 | 0.00 | 0.00 | 32.60 |
| 981 | 0.0346 | 0.00 | 0.00 | 32.57 |
| 982 | 0.0279 | 0.00 | 0.00 | 32.54 |
| 983 | 0.0340 | 0.00 | 0.00 | 32.51 |
| 984 | 0.0321 | 0.00 | 0.00 | 32.48 |
| 985 | 0.0327 | 0.00 | 0.00 | 32.44 |
| 986 | 0.0272 | 0.00 | 0.00 | 32.41 |
| 987 | 0.0353 | 0.00 | 0.00 | 32.38 |
| 988 | 0.0324 | 0.00 | 0.00 | 32.35 |
| 989 | 0.0321 | 0.00 | 0.00 | 32.32 |
| 990 | 0.0324 | 0.00 | 0.00 | 32.28 |
| 991 | 0.0333 | 0.00 | 0.00 | 32.25 |
| 992 | 0.0330 | 0.00 | 0.01 | 32.22 |
| 993 | 0.0244 | 0.00 | 0.00 | 32.19 |
| 994 | 0.0314 | 0.00 | 0.00 | 32.16 |
| 995 | 0.0292 | 0.00 | 0.00 | 32.13 |
| 996 | 0.0401 | 0.00 | 0.01 | 32.10 |
| 997 | 0.0333 | 0.00 | 0.01 | 32.06 |
| 998 | 0.0378 | 0.00 | 0.00 | 32.03 |
| 999 | 0.0327 | 0.00 | 0.00 | 31.99 |
| 1000 | 0.0292 | 0.00 | 0.00 | 31.96 |
| 1001 | 0.0279 | 0.00 | 0.00 | 31.93 |
| 1002 | 0.0346 | 0.00 | 0.00 | 31.90 |
| 1003 | 0.0285 | 0.00 | 0.00 | 31.87 |
| 1004 | 0.0356 | 0.00 | 0.00 | 31.84 |
| 1005 | 0.0362 | 0.00 | 0.01 | 31.80 |
| 1006 | 0.0330 | 0.00 | 0.01 | 31.77 |
| 1007 | 0.0301 | 0.00 | 0.00 | 31.73 |
| 1008 | 0.0276 | 0.00 | 0.00 | 31.70 |
| 1009 | 0.0321 | 0.00 | 0.00 | 31.67 |
| 1010 | 0.0282 | 0.00 | 0.00 | 31.64 |
| 1011 | 0.0349 | 0.00 | 0.00 | 31.61 |
| 1012 | 0.0269 | 0.00 | 0.00 | 31.58 |

| | | | | |
|---|---|---|---|---|
| 1013 | 0.0333 | 0.00 | 0.00 | 31.55 |
| 1014 | 0.0272 | 0.00 | 0.00 | 31.52 |
| 1015 | 0.0346 | 0.00 | 0.00 | 31.49 |
| 1016 | 0.0285 | 0.00 | 0.00 | 31.46 |
| 1017 | 0.0308 | 0.00 | 0.00 | 31.43 |
| 1018 | 0.0337 | 0.00 | 0.00 | 31.40 |
| 1019 | 0.0234 | 0.00 | 0.00 | 31.36 |
| 1020 | 0.0301 | 0.00 | 0.00 | 31.34 |
| 1021 | 0.0285 | 0.00 | 0.00 | 31.31 |
| 1022 | 0.0288 | 0.00 | 0.00 | 31.28 |
| 1023 | 0.0317 | 0.00 | 0.00 | 31.25 |
| 1024 | 0.0308 | 0.00 | 0.00 | 31.22 |
| 1025 | 0.0301 | 0.00 | 0.00 | 31.19 |
| 1026 | 0.0256 | 0.00 | 0.00 | 31.16 |
| 1027 | 0.0276 | 0.00 | 0.00 | 31.14 |
| 1028 | 0.0202 | 0.00 | 0.00 | 31.11 |
| 1029 | 0.0285 | 0.00 | 0.00 | 31.09 |
| 1030 | 0.0292 | 0.00 | 0.00 | 31.06 |
| 1031 | 0.0269 | 0.00 | 0.00 | 31.03 |
| 1032 | 0.0292 | 0.00 | 0.00 | 31.00 |
| 1033 | 0.0317 | 0.00 | 0.00 | 30.97 |
| 1034 | 0.0340 | 0.00 | 0.00 | 30.94 |
| 1035 | 0.0276 | 0.00 | 0.00 | 30.91 |
| 1036 | 0.0324 | 0.00 | 0.00 | 30.88 |
| 1037 | 0.0301 | 0.00 | 0.00 | 30.85 |
| 1038 | 0.0362 | 0.00 | 0.01 | 30.82 |
| 1039 | 0.0343 | 0.00 | 0.00 | 30.78 |
| 1040 | 0.0343 | 0.00 | 0.00 | 30.75 |
| 1041 | 0.0420 | 0.00 | 0.01 | 30.71 |
| 1042 | 0.0349 | 0.00 | 0.01 | 30.67 |
| 1043 | 0.0279 | 0.00 | 0.00 | 30.64 |
| 1044 | 0.0388 | 0.00 | 0.01 | 30.61 |
| 1045 | 0.0298 | 0.00 | 0.00 | 30.57 |
| 1046 | 0.0327 | 0.00 | 0.00 | 30.54 |
| 1047 | 0.0327 | 0.00 | 0.00 | 30.51 |
| 1048 | 0.0266 | 0.00 | 0.00 | 30.47 |
| 1049 | 0.0301 | 0.00 | 0.00 | 30.45 |
| 1050 | 0.0250 | 0.00 | 0.00 | 30.42 |
| 1051 | 0.0330 | 0.00 | 0.00 | 30.39 |
| 1052 | 0.0304 | 0.00 | 0.01 | 30.36 |
| 1053 | 0.0218 | 0.00 | 0.00 | 30.33 |
| 1054 | 0.0327 | 0.00 | 0.00 | 30.31 |
| 1055 | 0.0292 | 0.00 | 0.00 | 30.27 |
| 1056 | 0.0272 | 0.00 | 0.00 | 30.25 |

| | | | | |
|---|---|---|---|---|
| 1057 | 0.0269 | 0.00 | 0.00 | 30.22 |
| 1058 | 0.0298 | 0.00 | 0.00 | 30.19 |
| 1059 | 0.0298 | 0.00 | 0.00 | 30.16 |
| 1060 | 0.0301 | 0.00 | 0.00 | 30.13 |
| 1061 | 0.0247 | 0.00 | 0.00 | 30.10 |
| 1062 | 0.0301 | 0.00 | 0.01 | 30.08 |
| 1063 | 0.0353 | 0.00 | 0.01 | 30.05 |
| 1064 | 0.0298 | 0.00 | 0.00 | 30.01 |
| 1065 | 0.0321 | 0.00 | 0.00 | 29.98 |
| 1066 | 0.0282 | 0.00 | 0.00 | 29.95 |
| 1067 | 0.0224 | 0.00 | 0.00 | 29.92 |
| 1068 | 0.0263 | 0.00 | 0.00 | 29.90 |
| 1069 | 0.0288 | 0.00 | 0.00 | 29.87 |
| 1070 | 0.0301 | 0.00 | 0.00 | 29.84 |
| 1071 | 0.0288 | 0.00 | 0.00 | 29.81 |
| 1072 | 0.0301 | 0.00 | 0.01 | 29.78 |
| 1073 | 0.0343 | 0.00 | 0.01 | 29.75 |
| 1074 | 0.0231 | 0.00 | 0.00 | 29.72 |
| 1075 | 0.0240 | 0.00 | 0.00 | 29.70 |
| 1076 | 0.0208 | 0.00 | 0.00 | 29.67 |
| 1077 | 0.0183 | 0.00 | 0.00 | 29.65 |
| 1078 | 0.0247 | 0.00 | 0.00 | 29.63 |
| 1079 | 0.0269 | 0.00 | 0.00 | 29.61 |
| 1080 | 0.0314 | 0.00 | 0.00 | 29.58 |
| 1081 | 0.0266 | 0.00 | 0.00 | 29.55 |
| 1082 | 0.0266 | 0.00 | 0.00 | 29.52 |
| 1083 | 0.0282 | 0.00 | 0.00 | 29.50 |
| 1084 | 0.0231 | 0.00 | 0.00 | 29.47 |
| 1085 | 0.0301 | 0.00 | 0.00 | 29.45 |
| 1086 | 0.0266 | 0.00 | 0.00 | 29.42 |
| 1087 | 0.0228 | 0.00 | 0.00 | 29.39 |
| 1088 | 0.0244 | 0.00 | 0.00 | 29.37 |
| 1089 | 0.0292 | 0.00 | 0.00 | 29.34 |
| 1090 | 0.0288 | 0.00 | 0.00 | 29.31 |
| 1091 | 0.0212 | 0.00 | 0.00 | 29.28 |
| 1092 | 0.0272 | 0.00 | 0.00 | 29.26 |
| 1093 | 0.0234 | 0.00 | 0.00 | 29.24 |
| 1094 | 0.0234 | 0.00 | 0.00 | 29.21 |
| 1095 | 0.0202 | 0.00 | 0.00 | 29.19 |
| 1096 | 0.0308 | 0.00 | 0.00 | 29.17 |
| 1097 | 0.0269 | 0.00 | 0.00 | 29.14 |
| 1098 | 0.0266 | 0.00 | 0.00 | 29.11 |
| 1099 | 0.0282 | 0.00 | 0.00 | 29.09 |
| 1100 | 0.0311 | 0.00 | 0.00 | 29.06 |

| | | | | |
|---|---|---|---|---|
| 1101 | 0.0282 | 0.00 | 0.00 | 29.03 |
| 1102 | 0.0272 | 0.00 | 0.00 | 29.00 |
| 1103 | 0.0279 | 0.00 | 0.00 | 28.97 |
| 1104 | 0.0304 | 0.00 | 0.01 | 28.94 |
| 1105 | 0.0288 | 0.00 | 0.00 | 28.91 |
| 1106 | 0.0228 | 0.00 | 0.00 | 28.88 |
| 1107 | 0.0240 | 0.00 | 0.00 | 28.86 |
| 1108 | 0.0253 | 0.00 | 0.00 | 28.84 |
| 1109 | 0.0250 | 0.00 | 0.00 | 28.81 |
| 1110 | 0.0272 | 0.00 | 0.00 | 28.79 |
| 1111 | 0.0228 | 0.00 | 0.00 | 28.76 |
| 1112 | 0.0282 | 0.00 | 0.00 | 28.74 |
| 1113 | 0.0199 | 0.00 | 0.00 | 28.71 |
| 1114 | 0.0231 | 0.00 | 0.00 | 28.69 |
| 1115 | 0.0279 | 0.00 | 0.00 | 28.67 |
| 1116 | 0.0250 | 0.00 | 0.00 | 28.64 |
| 1117 | 0.0237 | 0.00 | 0.00 | 28.61 |
| 1118 | 0.0272 | 0.00 | 0.00 | 28.59 |
| 1119 | 0.0260 | 0.00 | 0.00 | 28.56 |
| 1120 | 0.0295 | 0.00 | 0.00 | 28.54 |
| 1121 | 0.0218 | 0.00 | 0.00 | 28.51 |
| 1122 | 0.0218 | 0.00 | 0.00 | 28.48 |
| 1123 | 0.0240 | 0.00 | 0.00 | 28.46 |
| 1124 | 0.0224 | 0.00 | 0.00 | 28.44 |
| 1125 | 0.0221 | 0.00 | 0.00 | 28.42 |
| 1126 | 0.0237 | 0.00 | 0.00 | 28.39 |
| 1127 | 0.0186 | 0.00 | 0.00 | 28.37 |
| 1128 | 0.0199 | 0.00 | 0.00 | 28.35 |
| 1129 | 0.0292 | 0.00 | 0.00 | 28.33 |
| 1130 | 0.0272 | 0.00 | 0.00 | 28.30 |
| 1131 | 0.0218 | 0.00 | 0.00 | 28.28 |
| 1132 | 0.0224 | 0.00 | 0.00 | 28.25 |
| 1133 | 0.0250 | 0.00 | 0.00 | 28.23 |
| 1134 | 0.0189 | 0.00 | 0.00 | 28.21 |
| 1135 | 0.0231 | 0.00 | 0.00 | 28.19 |
| 1136 | 0.0263 | 0.00 | 0.00 | 28.16 |
| 1137 | 0.0228 | 0.00 | 0.00 | 28.14 |
| 1138 | 0.0260 | 0.00 | 0.00 | 28.11 |
| 1139 | 0.0260 | 0.00 | 0.00 | 28.09 |
| 1140 | 0.0215 | 0.00 | 0.00 | 28.06 |
| 1141 | 0.0221 | 0.00 | 0.00 | 28.04 |
| 1142 | 0.0218 | 0.00 | 0.00 | 28.02 |
| 1143 | 0.0199 | 0.00 | 0.00 | 28.00 |
| 1144 | 0.0260 | 0.00 | 0.00 | 27.98 |

| | | | | |
|---|---|---|---|---|
| 1145 | 0.0212 | 0.00 | 0.00 | 27.95 |
| 1146 | 0.0212 | 0.00 | 0.00 | 27.93 |
| 1147 | 0.0215 | 0.00 | 0.00 | 27.91 |
| 1148 | 0.0212 | 0.00 | 0.00 | 27.89 |
| 1149 | 0.0183 | 0.00 | 0.00 | 27.87 |
| 1150 | 0.0253 | 0.00 | 0.00 | 27.85 |
| 1151 | 0.0256 | 0.00 | 0.00 | 27.82 |
| 1152 | 0.0260 | 0.00 | 0.00 | 27.80 |
| 1153 | 0.0189 | 0.00 | 0.00 | 27.77 |
| 1154 | 0.0224 | 0.00 | 0.00 | 27.75 |
| 1155 | 0.0192 | 0.00 | 0.00 | 27.73 |
| 1156 | 0.0279 | 0.00 | 0.00 | 27.71 |
| 1157 | 0.0228 | 0.00 | 0.00 | 27.68 |
| 1158 | 0.0276 | 0.00 | 0.00 | 27.66 |
| 1159 | 0.0212 | 0.00 | 0.00 | 27.63 |
| 1160 | 0.0192 | 0.00 | 0.00 | 27.61 |
| 1161 | 0.0167 | 0.00 | 0.00 | 27.59 |
| 1162 | 0.0215 | 0.00 | 0.00 | 27.58 |
| 1163 | 0.0215 | 0.00 | 0.00 | 27.55 |
| 1164 | 0.0224 | 0.00 | 0.00 | 27.53 |
| 1165 | 0.0186 | 0.00 | 0.00 | 27.51 |
| 1166 | 0.0266 | 0.00 | 0.00 | 27.49 |
| 1167 | 0.0253 | 0.00 | 0.00 | 27.47 |
| 1168 | 0.0212 | 0.00 | 0.00 | 27.44 |
| 1169 | 0.0224 | 0.00 | 0.00 | 27.42 |
| 1170 | 0.0253 | 0.00 | 0.00 | 27.40 |
| 1171 | 0.0224 | 0.00 | 0.00 | 27.37 |
| 1172 | 0.0212 | 0.00 | 0.00 | 27.35 |
| 1173 | 0.0212 | 0.00 | 0.00 | 27.33 |
| 1174 | 0.0266 | 0.00 | 0.00 | 27.31 |
| 1175 | 0.0234 | 0.00 | 0.00 | 27.28 |
| 1176 | 0.0295 | 0.00 | 0.01 | 27.26 |
| 1177 | 0.0224 | 0.00 | 0.00 | 27.23 |
| 1178 | 0.0263 | 0.00 | 0.00 | 27.20 |
| 1179 | 0.0186 | 0.00 | 0.00 | 27.18 |
| 1180 | 0.0208 | 0.00 | 0.00 | 27.16 |
| 1181 | 0.0199 | 0.00 | 0.00 | 27.14 |
| 1182 | 0.0244 | 0.00 | 0.00 | 27.12 |
| 1183 | 0.0183 | 0.00 | 0.00 | 27.09 |
| 1184 | 0.0202 | 0.00 | 0.00 | 27.08 |
| 1185 | 0.0244 | 0.00 | 0.00 | 27.06 |
| 1186 | 0.0237 | 0.00 | 0.00 | 27.03 |
| 1187 | 0.0224 | 0.00 | 0.00 | 27.01 |
| 1188 | 0.0215 | 0.00 | 0.00 | 26.99 |

| | | | | |
|---|---|---|---|---|
| 1189 | 0.0247 | 0.00 | 0.00 | 26.96 |
| 1190 | 0.0253 | 0.00 | 0.00 | 26.94 |
| 1191 | 0.0202 | 0.00 | 0.00 | 26.91 |
| 1192 | 0.0208 | 0.00 | 0.00 | 26.89 |
| 1193 | 0.0234 | 0.00 | 0.00 | 26.87 |
| 1194 | 0.0208 | 0.00 | 0.00 | 26.85 |
| 1195 | 0.0183 | 0.00 | 0.00 | 26.83 |
| 1196 | 0.0170 | 0.00 | 0.00 | 26.81 |
| 1197 | 0.0212 | 0.00 | 0.00 | 26.79 |
| 1198 | 0.0196 | 0.00 | 0.00 | 26.77 |
| 1199 | 0.0279 | 0.00 | 0.00 | 26.75 |
| 1200 | 0.0138 | 0.00 | 0.00 | 26.72 |
| 1201 | 0.0179 | 0.00 | 0.00 | 26.71 |
| 1202 | 0.0228 | 0.00 | 0.00 | 26.69 |
| 1203 | 0.0218 | 0.00 | 0.00 | 26.67 |
| 1204 | 0.0186 | 0.00 | 0.00 | 26.65 |
| 1205 | 0.0183 | 0.00 | 0.00 | 26.63 |
| 1206 | 0.0250 | 0.00 | 0.00 | 26.61 |
| 1207 | 0.0192 | 0.00 | 0.00 | 26.59 |
| 1208 | 0.0234 | 0.00 | 0.00 | 26.57 |
| 1209 | 0.0272 | 0.00 | 0.00 | 26.54 |
| 1210 | 0.0196 | 0.00 | 0.00 | 26.52 |
| 1211 | 0.0221 | 0.00 | 0.00 | 26.50 |
| 1212 | 0.0208 | 0.00 | 0.00 | 26.48 |
| 1213 | 0.0218 | 0.00 | 0.00 | 26.45 |
| 1214 | 0.0199 | 0.00 | 0.00 | 26.43 |
| 1215 | 0.0173 | 0.00 | 0.00 | 26.41 |
| 1216 | 0.0179 | 0.00 | 0.00 | 26.40 |
| 1217 | 0.0208 | 0.00 | 0.00 | 26.38 |
| 1218 | 0.0192 | 0.00 | 0.00 | 26.36 |
| 1219 | 0.0218 | 0.00 | 0.00 | 26.34 |
| 1220 | 0.0199 | 0.00 | 0.00 | 26.32 |
| 1221 | 0.0208 | 0.00 | 0.00 | 26.30 |
| 1222 | 0.0141 | 0.00 | 0.00 | 26.27 |
| 1223 | 0.0163 | 0.00 | 0.00 | 26.26 |
| 1224 | 0.0151 | 0.00 | 0.00 | 26.24 |
| 1225 | 0.0160 | 0.00 | 0.00 | 26.23 |
| 1226 | 0.0205 | 0.00 | 0.00 | 26.21 |
| 1227 | 0.0208 | 0.00 | 0.00 | 26.19 |
| 1228 | 0.0208 | 0.00 | 0.00 | 26.17 |
| 1229 | 0.0167 | 0.00 | 0.00 | 26.15 |
| 1230 | 0.0199 | 0.00 | 0.00 | 26.13 |
| 1231 | 0.0215 | 0.00 | 0.00 | 26.11 |
| 1232 | 0.0186 | 0.00 | 0.00 | 26.09 |

| | | | | |
|---|---|---|---|---|
| 1233 | 0.0208 | 0.00 | 0.00 | 26.07 |
| 1234 | 0.0208 | 0.00 | 0.00 | 26.05 |
| 1235 | 0.0221 | 0.00 | 0.00 | 26.03 |
| 1236 | 0.0234 | 0.00 | 0.00 | 26.01 |
| 1237 | 0.0388 | 0.00 | 0.02 | 25.99 |
| 1238 | 0.0247 | 0.00 | 0.00 | 25.95 |
| 1239 | 0.0208 | 0.00 | 0.00 | 25.92 |
| 1240 | 0.0263 | 0.00 | 0.01 | 25.90 |
| 1241 | 0.0202 | 0.00 | 0.00 | 25.88 |
| 1242 | 0.0196 | 0.00 | 0.00 | 25.86 |
| 1243 | 0.0186 | 0.00 | 0.00 | 25.84 |
| 1244 | 0.0192 | 0.00 | 0.00 | 25.82 |
| 1245 | 0.0228 | 0.00 | 0.00 | 25.80 |
| 1246 | 0.0170 | 0.00 | 0.00 | 25.78 |
| 1247 | 0.0215 | 0.00 | 0.00 | 25.76 |
| 1248 | 0.0186 | 0.00 | 0.00 | 25.74 |
| 1249 | 0.0202 | 0.00 | 0.00 | 25.72 |
| 1250 | 0.0205 | 0.00 | 0.00 | 25.70 |
| 1251 | 0.0163 | 0.00 | 0.00 | 25.68 |
| 1252 | 0.0183 | 0.00 | 0.00 | 25.66 |
| 1253 | 0.0170 | 0.00 | 0.00 | 25.64 |
| 1254 | 0.0192 | 0.00 | 0.00 | 25.63 |
| 1255 | 0.0199 | 0.00 | 0.00 | 25.61 |
| 1256 | 0.0199 | 0.00 | 0.00 | 25.59 |
| 1257 | 0.0224 | 0.00 | 0.00 | 25.57 |
| 1258 | 0.0173 | 0.00 | 0.00 | 25.55 |
| 1259 | 0.0183 | 0.00 | 0.00 | 25.53 |
| 1260 | 0.0173 | 0.00 | 0.00 | 25.51 |
| 1261 | 0.0186 | 0.00 | 0.00 | 25.49 |
| 1262 | 0.0205 | 0.00 | 0.00 | 25.47 |
| 1263 | 0.0176 | 0.00 | 0.00 | 25.45 |
| 1264 | 0.0202 | 0.00 | 0.00 | 25.44 |
| 1265 | 0.0215 | 0.00 | 0.00 | 25.42 |
| 1266 | 0.0192 | 0.00 | 0.00 | 25.39 |
| 1267 | 0.0192 | 0.00 | 0.00 | 25.38 |
| 1268 | 0.0189 | 0.00 | 0.00 | 25.36 |
| 1269 | 0.0202 | 0.00 | 0.00 | 25.34 |
| 1270 | 0.0167 | 0.00 | 0.00 | 25.32 |
| 1271 | 0.0199 | 0.00 | 0.00 | 25.30 |
| 1272 | 0.0202 | 0.00 | 0.00 | 25.28 |
| 1273 | 0.0240 | 0.00 | 0.00 | 25.26 |
| 1274 | 0.0167 | 0.00 | 0.00 | 25.24 |
| 1275 | 0.0170 | 0.00 | 0.00 | 25.22 |
| 1276 | 0.0192 | 0.00 | 0.00 | 25.20 |

| | | | | |
|---|---|---|---|---|
| 1277 | 0.0173 | 0.00 | 0.00 | 25.18 |
| 1278 | 0.0144 | 0.00 | 0.00 | 25.17 |
| 1279 | 0.0176 | 0.00 | 0.00 | 25.15 |
| 1280 | 0.0192 | 0.00 | 0.00 | 25.13 |
| 1281 | 0.0179 | 0.00 | 0.00 | 25.11 |
| 1282 | 0.0202 | 0.00 | 0.00 | 25.10 |
| 1283 | 0.0138 | 0.00 | 0.00 | 25.08 |
| 1284 | 0.0224 | 0.00 | 0.00 | 25.06 |
| 1285 | 0.0167 | 0.00 | 0.00 | 25.04 |
| 1286 | 0.0228 | 0.00 | 0.00 | 25.02 |
| 1287 | 0.0151 | 0.00 | 0.00 | 25.00 |
| 1288 | 0.0154 | 0.00 | 0.00 | 24.99 |
| 1289 | 0.0199 | 0.00 | 0.00 | 24.97 |
| 1290 | 0.0160 | 0.00 | 0.00 | 24.95 |
| 1291 | 0.0173 | 0.00 | 0.00 | 24.93 |
| 1292 | 0.0163 | 0.00 | 0.00 | 24.92 |
| 1293 | 0.0192 | 0.00 | 0.00 | 24.90 |
| 1294 | 0.0186 | 0.00 | 0.00 | 24.88 |
| 1295 | 0.0163 | 0.00 | 0.00 | 24.86 |
| 1296 | 0.0163 | 0.00 | 0.00 | 24.85 |
| 1297 | 0.0186 | 0.00 | 0.00 | 24.83 |
| 1298 | 0.0151 | 0.00 | 0.00 | 24.81 |
| 1299 | 0.0170 | 0.00 | 0.00 | 24.80 |
| 1300 | 0.0170 | 0.00 | 0.00 | 24.78 |
| 1301 | 0.0173 | 0.00 | 0.00 | 24.76 |
| 1302 | 0.0160 | 0.00 | 0.00 | 24.75 |
| 1303 | 0.0192 | 0.00 | 0.00 | 24.73 |
| 1304 | 0.0189 | 0.00 | 0.00 | 24.71 |
| 1305 | 0.0170 | 0.00 | 0.00 | 24.69 |
| 1306 | 0.0154 | 0.00 | 0.00 | 24.67 |
| 1307 | 0.0151 | 0.00 | 0.00 | 24.66 |
| 1308 | 0.0176 | 0.00 | 0.00 | 24.64 |
| 1309 | 0.0186 | 0.00 | 0.00 | 24.63 |
| 1310 | 0.0151 | 0.00 | 0.00 | 24.61 |
| 1311 | 0.0179 | 0.00 | 0.00 | 24.59 |
| 1312 | 0.0163 | 0.00 | 0.00 | 24.57 |
| 1313 | 0.0183 | 0.00 | 0.00 | 24.56 |
| 1314 | 0.0157 | 0.00 | 0.00 | 24.54 |
| 1315 | 0.0186 | 0.00 | 0.00 | 24.52 |
| 1316 | 0.0135 | 0.00 | 0.00 | 24.51 |
| 1317 | 0.0141 | 0.00 | 0.00 | 24.49 |
| 1318 | 0.0154 | 0.00 | 0.00 | 24.48 |
| 1319 | 0.0125 | 0.00 | 0.00 | 24.46 |
| 1320 | 0.0186 | 0.00 | 0.00 | 24.45 |

| | | | | |
|---|---|---|---|---|
| 1321 | 0.0157 | 0.00 | 0.00 | 24.43 |
| 1322 | 0.0221 | 0.00 | 0.00 | 24.42 |
| 1323 | 0.0173 | 0.00 | 0.00 | 24.39 |
| 1324 | 0.0170 | 0.00 | 0.00 | 24.38 |
| 1325 | 0.0160 | 0.00 | 0.00 | 24.36 |
| 1326 | 0.0199 | 0.00 | 0.00 | 24.34 |
| 1327 | 0.0141 | 0.00 | 0.00 | 24.32 |
| 1328 | 0.0131 | 0.00 | 0.00 | 24.31 |
| 1329 | 0.0157 | 0.00 | 0.00 | 24.30 |
| 1330 | 0.0157 | 0.00 | 0.00 | 24.28 |
| 1331 | 0.0141 | 0.00 | 0.00 | 24.26 |
| 1332 | 0.0167 | 0.00 | 0.00 | 24.25 |
| 1333 | 0.0135 | 0.00 | 0.00 | 24.23 |
| 1334 | 0.0189 | 0.00 | 0.00 | 24.22 |
| 1335 | 0.0186 | 0.00 | 0.00 | 24.20 |
| 1336 | 0.0167 | 0.00 | 0.00 | 24.18 |
| 1337 | 0.0183 | 0.00 | 0.00 | 24.17 |
| 1338 | 0.0183 | 0.00 | 0.00 | 24.15 |
| 1339 | 0.0205 | 0.00 | 0.00 | 24.13 |
| 1340 | 0.0176 | 0.00 | 0.00 | 24.11 |
| 1341 | 0.0135 | 0.00 | 0.00 | 24.09 |
| 1342 | 0.0135 | 0.00 | 0.00 | 24.08 |
| 1343 | 0.0160 | 0.00 | 0.00 | 24.06 |
| 1344 | 0.0154 | 0.00 | 0.00 | 24.05 |
| 1345 | 0.0125 | 0.00 | 0.00 | 24.03 |
| 1346 | 0.0160 | 0.00 | 0.00 | 24.02 |
| 1347 | 0.0176 | 0.00 | 0.00 | 24.00 |
| 1348 | 0.0119 | 0.00 | 0.00 | 23.99 |
| 1349 | 0.0196 | 0.00 | 0.01 | 23.98 |
| 1350 | 0.0196 | 0.00 | 0.00 | 23.96 |
| 1351 | 0.0154 | 0.00 | 0.00 | 23.94 |
| 1352 | 0.0205 | 0.00 | 0.00 | 23.92 |
| 1353 | 0.0138 | 0.00 | 0.00 | 23.90 |
| 1354 | 0.0125 | 0.00 | 0.00 | 23.89 |
| 1355 | 0.0128 | 0.00 | 0.00 | 23.87 |
| 1356 | 0.0138 | 0.00 | 0.00 | 23.86 |
| 1357 | 0.0151 | 0.00 | 0.00 | 23.85 |
| 1358 | 0.0144 | 0.00 | 0.00 | 23.83 |
| 1359 | 0.0122 | 0.00 | 0.00 | 23.82 |
| 1360 | 0.0160 | 0.00 | 0.00 | 23.81 |
| 1361 | 0.0170 | 0.00 | 0.00 | 23.79 |
| 1362 | 0.0141 | 0.00 | 0.00 | 23.77 |
| 1363 | 0.0167 | 0.00 | 0.00 | 23.76 |
| 1364 | 0.0179 | 0.00 | 0.00 | 23.74 |

| | | | | |
|---|---|---|---|---|
| 1365 | 0.0208 | 0.00 | 0.00 | 23.72 |
| 1366 | 0.0141 | 0.00 | 0.00 | 23.70 |
| 1367 | 0.0167 | 0.00 | 0.00 | 23.69 |
| 1368 | 0.0141 | 0.00 | 0.00 | 23.67 |
| 1369 | 0.0135 | 0.00 | 0.00 | 23.66 |
| 1370 | 0.0157 | 0.00 | 0.00 | 23.64 |
| 1371 | 0.0147 | 0.00 | 0.00 | 23.63 |
| 1372 | 0.0125 | 0.00 | 0.00 | 23.61 |
| 1373 | 0.0109 | 0.00 | 0.00 | 23.60 |
| 1374 | 0.0096 | 0.00 | 0.00 | 23.59 |
| 1375 | 0.0167 | 0.00 | 0.00 | 23.58 |
| 1376 | 0.0109 | 0.00 | 0.00 | 23.56 |
| 1377 | 0.0125 | 0.00 | 0.00 | 23.55 |
| 1378 | 0.0106 | 0.00 | 0.00 | 23.54 |
| 1379 | 0.0154 | 0.00 | 0.00 | 23.53 |
| 1380 | 0.0138 | 0.00 | 0.00 | 23.52 |
| 1381 | 0.0160 | 0.00 | 0.00 | 23.50 |
| 1382 | 0.0115 | 0.00 | 0.00 | 23.49 |
| 1383 | 0.0212 | 0.00 | 0.00 | 23.47 |
| 1384 | 0.0125 | 0.00 | 0.00 | 23.45 |
| 1385 | 0.0141 | 0.00 | 0.00 | 23.44 |
| 1386 | 0.0170 | 0.00 | 0.00 | 23.43 |
| 1387 | 0.0128 | 0.00 | 0.00 | 23.41 |
| 1388 | 0.0154 | 0.00 | 0.00 | 23.40 |
| 1389 | 0.0131 | 0.00 | 0.00 | 23.38 |
| 1390 | 0.0183 | 0.00 | 0.00 | 23.37 |
| 1391 | 0.0167 | 0.00 | 0.00 | 23.35 |
| 1392 | 0.0173 | 0.00 | 0.00 | 23.33 |
| 1393 | 0.0176 | 0.00 | 0.00 | 23.32 |
| 1394 | 0.0144 | 0.00 | 0.00 | 23.30 |
| 1395 | 0.0112 | 0.00 | 0.00 | 23.28 |
| 1396 | 0.0128 | 0.00 | 0.00 | 23.27 |
| 1397 | 0.0176 | 0.00 | 0.00 | 23.26 |
| 1398 | 0.0128 | 0.00 | 0.00 | 23.24 |
| 1399 | 0.0205 | 0.00 | 0.00 | 23.23 |
| 1400 | 0.0138 | 0.00 | 0.00 | 23.21 |
| 1401 | 0.0183 | 0.00 | 0.00 | 23.19 |
| 1402 | 0.0157 | 0.00 | 0.00 | 23.18 |
| 1403 | 0.0170 | 0.00 | 0.00 | 23.16 |
| 1404 | 0.0170 | 0.00 | 0.00 | 23.14 |
| 1405 | 0.0135 | 0.00 | 0.00 | 23.13 |
| 1406 | 0.0176 | 0.00 | 0.00 | 23.11 |
| 1407 | 0.0096 | 0.00 | 0.00 | 23.10 |
| 1408 | 0.0147 | 0.00 | 0.00 | 23.09 |

| | | | | |
|---|---|---|---|---|
| 1409 | 0.0163 | 0.00 | 0.00 | 23.07 |
| 1410 | 0.0151 | 0.00 | 0.00 | 23.06 |
| 1411 | 0.0141 | 0.00 | 0.00 | 23.04 |
| 1412 | 0.0138 | 0.00 | 0.00 | 23.03 |
| 1413 | 0.0170 | 0.00 | 0.00 | 23.01 |
| 1414 | 0.0154 | 0.00 | 0.00 | 23.00 |
| 1415 | 0.0173 | 0.00 | 0.00 | 22.98 |
| 1416 | 0.0176 | 0.00 | 0.00 | 22.96 |
| 1417 | 0.0141 | 0.00 | 0.00 | 22.94 |
| 1418 | 0.0125 | 0.00 | 0.00 | 22.93 |
| 1419 | 0.0138 | 0.00 | 0.00 | 22.92 |
| 1420 | 0.0157 | 0.00 | 0.00 | 22.90 |
| 1421 | 0.0154 | 0.00 | 0.00 | 22.89 |
| 1422 | 0.0157 | 0.00 | 0.00 | 22.87 |
| 1423 | 0.0141 | 0.00 | 0.00 | 22.86 |
| 1424 | 0.0160 | 0.00 | 0.00 | 22.84 |
| 1425 | 0.0138 | 0.00 | 0.00 | 22.83 |
| 1426 | 0.0154 | 0.00 | 0.00 | 22.81 |
| 1427 | 0.0154 | 0.00 | 0.00 | 22.80 |
| 1428 | 0.0119 | 0.00 | 0.00 | 22.78 |
| 1429 | 0.0115 | 0.00 | 0.00 | 22.77 |
| 1430 | 0.0151 | 0.00 | 0.00 | 22.76 |
| 1431 | 0.0151 | 0.00 | 0.00 | 22.74 |
| 1432 | 0.0122 | 0.00 | 0.00 | 22.73 |
| 1433 | 0.0144 | 0.00 | 0.00 | 22.72 |
| 1434 | 0.0186 | 0.00 | 0.00 | 22.70 |
| 1435 | 0.0157 | 0.00 | 0.00 | 22.68 |
| 1436 | 0.0183 | 0.00 | 0.00 | 22.67 |
| 1437 | 0.0167 | 0.00 | 0.00 | 22.65 |
| 1438 | 0.0128 | 0.00 | 0.00 | 22.63 |
| 1439 | 0.0176 | 0.00 | 0.00 | 22.62 |
| 1440 | 0.0144 | 0.00 | 0.00 | 22.60 |
| 1441 | 0.0160 | 0.00 | 0.00 | 22.59 |
| 1442 | 0.0154 | 0.00 | 0.00 | 22.57 |
| 1443 | 0.0141 | 0.00 | 0.00 | 22.56 |
| 1444 | 0.0179 | 0.00 | 0.00 | 22.54 |
| 1445 | 0.0147 | 0.00 | 0.00 | 22.53 |
| 1446 | 0.0115 | 0.00 | 0.00 | 22.51 |
| 1447 | 0.0103 | 0.00 | 0.00 | 22.50 |
| 1448 | 0.0163 | 0.00 | 0.00 | 22.49 |
| 1449 | 0.0112 | 0.00 | 0.00 | 22.47 |
| 1450 | 0.0144 | 0.00 | 0.00 | 22.46 |
| 1451 | 0.0135 | 0.00 | 0.00 | 22.45 |
| 1452 | 0.0103 | 0.00 | 0.00 | 22.43 |

| | | | | |
|---|---|---|---|---|
| 1453 | 0.0176 | 0.00 | 0.00 | 22.42 |
| 1454 | 0.0128 | 0.00 | 0.00 | 22.41 |
| 1455 | 0.0144 | 0.00 | 0.00 | 22.39 |
| 1456 | 0.0125 | 0.00 | 0.00 | 22.38 |
| 1457 | 0.0109 | 0.00 | 0.00 | 22.37 |
| 1458 | 0.0154 | 0.00 | 0.00 | 22.35 |
| 1459 | 0.0115 | 0.00 | 0.00 | 22.34 |
| 1460 | 0.0160 | 0.00 | 0.00 | 22.33 |
| 1461 | 0.0109 | 0.00 | 0.00 | 22.31 |
| 1462 | 0.0103 | 0.00 | 0.00 | 22.30 |
| 1463 | 0.0122 | 0.00 | 0.00 | 22.29 |
| 1464 | 0.0109 | 0.00 | 0.00 | 22.28 |
| 1465 | 0.0138 | 0.00 | 0.00 | 22.27 |
| 1466 | 0.0144 | 0.00 | 0.00 | 22.25 |
| 1467 | 0.0128 | 0.00 | 0.00 | 22.24 |
| 1468 | 0.0109 | 0.00 | 0.00 | 22.23 |
| 1469 | 0.0151 | 0.00 | 0.00 | 22.22 |
| 1470 | 0.0131 | 0.00 | 0.00 | 22.20 |
| 1471 | 0.0131 | 0.00 | 0.00 | 22.19 |
| 1472 | 0.0115 | 0.00 | 0.00 | 22.17 |
| 1473 | 0.0125 | 0.00 | 0.00 | 22.16 |
| 1474 | 0.0112 | 0.00 | 0.00 | 22.15 |
| 1475 | 0.0119 | 0.00 | 0.00 | 22.14 |
| 1476 | 0.0125 | 0.00 | 0.00 | 22.13 |
| 1477 | 0.0128 | 0.00 | 0.00 | 22.11 |
| 1478 | 0.0125 | 0.00 | 0.00 | 22.10 |
| 1479 | 0.0115 | 0.00 | 0.00 | 22.09 |
| 1480 | 0.0122 | 0.00 | 0.00 | 22.08 |
| 1481 | 0.0112 | 0.00 | 0.00 | 22.07 |
| 1482 | 0.0173 | 0.00 | 0.00 | 22.05 |
| 1483 | 0.0144 | 0.00 | 0.00 | 22.04 |
| 1484 | 0.0112 | 0.00 | 0.00 | 22.02 |
| 1485 | 0.0109 | 0.00 | 0.00 | 22.01 |
| 1486 | 0.0160 | 0.00 | 0.00 | 22.00 |
| 1487 | 0.0154 | 0.00 | 0.00 | 21.98 |
| 1488 | 0.0154 | 0.00 | 0.00 | 21.97 |
| 1489 | 0.0167 | 0.00 | 0.00 | 21.95 |
| 1490 | 0.0144 | 0.00 | 0.00 | 21.94 |
| 1491 | 0.0131 | 0.00 | 0.00 | 21.92 |
| 1492 | 0.0115 | 0.00 | 0.00 | 21.91 |
| 1493 | 0.0103 | 0.00 | 0.00 | 21.90 |
| 1494 | 0.0135 | 0.00 | 0.00 | 21.89 |
| 1495 | 0.0128 | 0.00 | 0.00 | 21.87 |
| 1496 | 0.0115 | 0.00 | 0.00 | 21.86 |

| | | | | |
|---|---|---|---|---|
| 1497 | 0.0109 | 0.00 | 0.00 | 21.85 |
| 1498 | 0.0131 | 0.00 | 0.00 | 21.84 |
| 1499 | 0.0138 | 0.00 | 0.00 | 21.83 |
| 1500 | 0.0135 | 0.00 | 0.00 | 21.81 |
| 1501 | 0.0138 | 0.00 | 0.00 | 21.80 |
| 1502 | 0.0151 | 0.00 | 0.00 | 21.78 |
| 1503 | 0.0186 | 0.00 | 0.00 | 21.77 |
| 1504 | 0.0128 | 0.00 | 0.00 | 21.75 |
| 1505 | 0.0096 | 0.00 | 0.00 | 21.74 |
| 1506 | 0.0125 | 0.00 | 0.00 | 21.73 |
| 1507 | 0.0131 | 0.00 | 0.00 | 21.72 |
| 1508 | 0.0144 | 0.00 | 0.00 | 21.70 |
| 1509 | 0.0144 | 0.00 | 0.00 | 21.69 |
| 1510 | 0.0135 | 0.00 | 0.00 | 21.67 |
| 1511 | 0.0125 | 0.00 | 0.00 | 21.66 |
| 1512 | 0.0112 | 0.00 | 0.00 | 21.65 |
| 1513 | 0.0144 | 0.00 | 0.00 | 21.64 |
| 1514 | 0.0170 | 0.00 | 0.00 | 21.62 |
| 1515 | 0.0109 | 0.00 | 0.00 | 21.61 |
| 1516 | 0.0170 | 0.00 | 0.00 | 21.59 |
| 1517 | 0.0147 | 0.00 | 0.00 | 21.58 |
| 1518 | 0.0125 | 0.00 | 0.00 | 21.56 |
| 1519 | 0.0183 | 0.00 | 0.00 | 21.55 |
| 1520 | 0.0135 | 0.00 | 0.00 | 21.53 |
| 1521 | 0.0141 | 0.00 | 0.00 | 21.52 |
| 1522 | 0.0167 | 0.00 | 0.00 | 21.50 |
| 1523 | 0.0170 | 0.00 | 0.00 | 21.49 |
| 1524 | 0.0173 | 0.00 | 0.00 | 21.47 |
| 1525 | 0.0131 | 0.00 | 0.00 | 21.45 |
| 1526 | 0.0125 | 0.00 | 0.00 | 21.44 |
| 1527 | 0.0122 | 0.00 | 0.00 | 21.43 |
| 1528 | 0.0144 | 0.00 | 0.00 | 21.42 |
| 1529 | 0.0151 | 0.00 | 0.00 | 21.40 |
| 1530 | 0.0119 | 0.00 | 0.00 | 21.39 |
| 1531 | 0.0151 | 0.00 | 0.00 | 21.37 |
| 1532 | 0.0186 | 0.00 | 0.00 | 21.36 |
| 1533 | 0.0119 | 0.00 | 0.00 | 21.34 |
| 1534 | 0.0196 | 0.00 | 0.00 | 21.33 |
| 1535 | 0.0163 | 0.00 | 0.00 | 21.31 |
| 1536 | 0.0160 | 0.00 | 0.00 | 21.29 |
| 1537 | 0.0135 | 0.00 | 0.00 | 21.28 |
| 1538 | 0.0147 | 0.00 | 0.00 | 21.26 |
| 1539 | 0.0163 | 0.00 | 0.00 | 21.25 |
| 1540 | 0.0128 | 0.00 | 0.00 | 21.23 |

| | | | | |
|---|---|---|---|---|
| 1541 | 0.0160 | 0.00 | 0.00 | 21.22 |
| 1542 | 0.0154 | 0.00 | 0.00 | 21.20 |
| 1543 | 0.0144 | 0.00 | 0.00 | 21.19 |
| 1544 | 0.0147 | 0.00 | 0.00 | 21.17 |
| 1545 | 0.0176 | 0.00 | 0.00 | 21.16 |
| 1546 | 0.0144 | 0.00 | 0.00 | 21.14 |
| 1547 | 0.0115 | 0.00 | 0.00 | 21.13 |
| 1548 | 0.0122 | 0.00 | 0.00 | 21.12 |
| 1549 | 0.0135 | 0.00 | 0.00 | 21.10 |
| 1550 | 0.0135 | 0.00 | 0.00 | 21.09 |
| 1551 | 0.0131 | 0.00 | 0.00 | 21.08 |
| 1552 | 0.0144 | 0.00 | 0.00 | 21.06 |
| 1553 | 0.0157 | 0.00 | 0.00 | 21.05 |
| 1554 | 0.0125 | 0.00 | 0.00 | 21.03 |
| 1555 | 0.0106 | 0.00 | 0.00 | 21.02 |
| 1556 | 0.0141 | 0.00 | 0.00 | 21.01 |
| 1557 | 0.0099 | 0.00 | 0.00 | 21.00 |
| 1558 | 0.0125 | 0.00 | 0.00 | 20.99 |
| 1559 | 0.0157 | 0.00 | 0.00 | 20.97 |
| 1560 | 0.0141 | 0.00 | 0.00 | 20.96 |
| 1561 | 0.0109 | 0.00 | 0.00 | 20.94 |
| 1562 | 0.0157 | 0.00 | 0.00 | 20.93 |
| 1563 | 0.0141 | 0.00 | 0.00 | 20.92 |
| 1564 | 0.0131 | 0.00 | 0.00 | 20.90 |
| 1565 | 0.0135 | 0.00 | 0.00 | 20.89 |
| 1566 | 0.0157 | 0.00 | 0.00 | 20.88 |
| 1567 | 0.0135 | 0.00 | 0.00 | 20.86 |
| 1568 | 0.0157 | 0.00 | 0.00 | 20.85 |
| 1569 | 0.0144 | 0.00 | 0.00 | 20.83 |
| 1570 | 0.0163 | 0.00 | 0.00 | 20.82 |
| 1571 | 0.0083 | 0.00 | 0.00 | 20.80 |
| 1572 | 0.0106 | 0.00 | 0.00 | 20.79 |
| 1573 | 0.0141 | 0.00 | 0.00 | 20.78 |
| 1574 | 0.0112 | 0.00 | 0.00 | 20.77 |
| 1575 | 0.0128 | 0.00 | 0.00 | 20.76 |
| 1576 | 0.0163 | 0.00 | 0.00 | 20.74 |
| 1577 | 0.0144 | 0.00 | 0.00 | 20.73 |
| 1578 | 0.0119 | 0.00 | 0.00 | 20.71 |
| 1579 | 0.0087 | 0.00 | 0.00 | 20.70 |
| 1580 | 0.0122 | 0.00 | 0.00 | 20.69 |
| 1581 | 0.0138 | 0.00 | 0.00 | 20.68 |
| 1582 | 0.0080 | 0.00 | 0.00 | 20.67 |
| 1583 | 0.0112 | 0.00 | 0.00 | 20.66 |
| 1584 | 0.0087 | 0.00 | 0.00 | 20.65 |

| | | | | |
|---|---|---|---|---|
| 1585 | 0.0154 | 0.00 | 0.00 | 20.64 |
| 1586 | 0.0131 | 0.00 | 0.00 | 20.62 |
| 1587 | 0.0115 | 0.00 | 0.00 | 20.61 |
| 1588 | 0.0157 | 0.00 | 0.00 | 20.60 |
| 1589 | 0.0115 | 0.00 | 0.00 | 20.58 |
| 1590 | 0.0135 | 0.00 | 0.00 | 20.57 |
| 1591 | 0.0115 | 0.00 | 0.00 | 20.56 |
| 1592 | 0.0122 | 0.00 | 0.00 | 20.55 |
| 1593 | 0.0112 | 0.00 | 0.00 | 20.53 |
| 1594 | 0.0099 | 0.00 | 0.00 | 20.52 |
| 1595 | 0.0115 | 0.00 | 0.00 | 20.51 |
| 1596 | 0.0109 | 0.00 | 0.00 | 20.50 |
| 1597 | 0.0147 | 0.00 | 0.00 | 20.49 |
| 1598 | 0.0112 | 0.00 | 0.00 | 20.48 |
| 1599 | 0.0122 | 0.00 | 0.00 | 20.46 |
| 1600 | 0.0103 | 0.00 | 0.00 | 20.45 |
| 1601 | 0.0093 | 0.00 | 0.00 | 20.44 |
| 1602 | 0.0119 | 0.00 | 0.00 | 20.43 |
| 1603 | 0.0122 | 0.00 | 0.00 | 20.42 |
| 1604 | 0.0125 | 0.00 | 0.00 | 20.41 |
| 1605 | 0.0112 | 0.00 | 0.00 | 20.40 |
| 1606 | 0.0087 | 0.00 | 0.00 | 20.39 |
| 1607 | 0.0135 | 0.00 | 0.00 | 20.38 |
| 1608 | 0.0128 | 0.00 | 0.00 | 20.36 |
| 1609 | 0.0138 | 0.00 | 0.00 | 20.35 |
| 1610 | 0.0115 | 0.00 | 0.00 | 20.34 |
| 1611 | 0.0115 | 0.00 | 0.00 | 20.33 |
| 1612 | 0.0147 | 0.00 | 0.00 | 20.31 |
| 1613 | 0.0067 | 0.00 | 0.00 | 20.30 |
| 1614 | 0.0099 | 0.00 | 0.00 | 20.29 |
| 1615 | 0.0119 | 0.00 | 0.00 | 20.28 |
| 1616 | 0.0080 | 0.00 | 0.00 | 20.27 |
| 1617 | 0.0144 | 0.00 | 0.00 | 20.26 |
| 1618 | 0.0093 | 0.00 | 0.00 | 20.25 |
| 1619 | 0.0119 | 0.00 | 0.00 | 20.24 |
| 1620 | 0.0141 | 0.00 | 0.00 | 20.23 |
| 1621 | 0.0093 | 0.00 | 0.00 | 20.21 |
| 1622 | 0.0122 | 0.00 | 0.00 | 20.20 |
| 1623 | 0.0119 | 0.00 | 0.00 | 20.19 |
| 1624 | 0.0144 | 0.00 | 0.00 | 20.18 |
| 1625 | 0.0099 | 0.00 | 0.00 | 20.16 |
| 1626 | 0.0099 | 0.00 | 0.00 | 20.15 |
| 1627 | 0.0141 | 0.00 | 0.00 | 20.14 |
| 1628 | 0.0135 | 0.00 | 0.00 | 20.13 |

| | | | | |
|---|---|---|---|---|
| 1629 | 0.0106 | 0.00 | 0.00 | 20.12 |
| 1630 | 0.0096 | 0.00 | 0.00 | 20.11 |
| 1631 | 0.0131 | 0.00 | 0.00 | 20.10 |
| 1632 | 0.0109 | 0.00 | 0.00 | 20.08 |
| 1633 | 0.0141 | 0.00 | 0.00 | 20.07 |
| 1634 | 0.0109 | 0.00 | 0.00 | 20.06 |
| 1635 | 0.0122 | 0.00 | 0.00 | 20.05 |
| 1636 | 0.0109 | 0.00 | 0.00 | 20.04 |
| 1637 | 0.0122 | 0.00 | 0.00 | 20.03 |
| 1638 | 0.0138 | 0.00 | 0.00 | 20.01 |
| 1639 | 0.0099 | 0.00 | 0.00 | 20.00 |
| 1640 | 0.0119 | 0.00 | 0.00 | 19.99 |
| 1641 | 0.0141 | 0.00 | 0.00 | 19.98 |
| 1642 | 0.0163 | 0.00 | 0.00 | 19.96 |
| 1643 | 0.0096 | 0.00 | 0.00 | 19.95 |
| 1644 | 0.0096 | 0.00 | 0.00 | 19.94 |
| 1645 | 0.0163 | 0.00 | 0.00 | 19.93 |
| 1646 | 0.0099 | 0.00 | 0.00 | 19.91 |
| 1647 | 0.0090 | 0.00 | 0.00 | 19.90 |
| 1648 | 0.0147 | 0.00 | 0.00 | 19.89 |
| 1649 | 0.0103 | 0.00 | 0.00 | 19.88 |
| 1650 | 0.0125 | 0.00 | 0.00 | 19.87 |
| 1651 | 0.0119 | 0.00 | 0.00 | 19.85 |
| 1652 | 0.0109 | 0.00 | 0.00 | 19.84 |
| 1653 | 0.0151 | 0.00 | 0.00 | 19.83 |
| 1654 | 0.0135 | 0.00 | 0.00 | 19.82 |
| 1655 | 0.0128 | 0.00 | 0.00 | 19.80 |
| 1656 | 0.0103 | 0.00 | 0.00 | 19.79 |
| 1657 | 0.0115 | 0.00 | 0.00 | 19.78 |
| 1658 | 0.0112 | 0.00 | 0.00 | 19.77 |
| 1659 | 0.0128 | 0.00 | 0.00 | 19.76 |
| 1660 | 0.0096 | 0.00 | 0.00 | 19.74 |
| 1661 | 0.0128 | 0.00 | 0.00 | 19.74 |
| 1662 | 0.0112 | 0.00 | 0.00 | 19.72 |
| 1663 | 0.0103 | 0.00 | 0.00 | 19.71 |
| 1664 | 0.0138 | 0.00 | 0.00 | 19.70 |
| 1665 | 0.0112 | 0.00 | 0.00 | 19.69 |
| 1666 | 0.0087 | 0.00 | 0.00 | 19.68 |
| 1667 | 0.0151 | 0.00 | 0.00 | 19.67 |
| 1668 | 0.0099 | 0.00 | 0.00 | 19.65 |
| 1669 | 0.0144 | 0.00 | 0.00 | 19.64 |
| 1670 | 0.0106 | 0.00 | 0.00 | 19.63 |
| 1671 | 0.0119 | 0.00 | 0.00 | 19.62 |
| 1672 | 0.0112 | 0.00 | 0.00 | 19.61 |

| | | | | |
|---|---|---|---|---|
| 1673 | 0.0093 | 0.00 | 0.00 | 19.59 |
| 1674 | 0.0112 | 0.00 | 0.00 | 19.58 |
| 1675 | 0.0122 | 0.00 | 0.00 | 19.57 |
| 1676 | 0.0128 | 0.00 | 0.00 | 19.56 |
| 1677 | 0.0128 | 0.00 | 0.00 | 19.55 |
| 1678 | 0.0112 | 0.00 | 0.00 | 19.54 |
| 1679 | 0.0167 | 0.00 | 0.00 | 19.52 |
| 1680 | 0.0096 | 0.00 | 0.00 | 19.51 |
| 1681 | 0.0103 | 0.00 | 0.00 | 19.50 |
| 1682 | 0.0125 | 0.00 | 0.00 | 19.49 |
| 1683 | 0.0167 | 0.00 | 0.00 | 19.48 |
| 1684 | 0.0144 | 0.00 | 0.00 | 19.46 |
| 1685 | 0.0240 | 0.00 | 0.01 | 19.44 |
| 1686 | 0.0160 | 0.00 | 0.00 | 19.42 |
| 1687 | 0.0115 | 0.00 | 0.00 | 19.40 |
| 1688 | 0.0093 | 0.00 | 0.00 | 19.39 |
| 1689 | 0.0093 | 0.00 | 0.00 | 19.38 |
| 1690 | 0.0099 | 0.00 | 0.00 | 19.37 |
| 1691 | 0.0096 | 0.00 | 0.00 | 19.36 |
| 1692 | 0.0115 | 0.00 | 0.00 | 19.35 |
| 1693 | 0.0109 | 0.00 | 0.00 | 19.34 |
| 1694 | 0.0138 | 0.00 | 0.00 | 19.33 |
| 1695 | 0.0160 | 0.00 | 0.01 | 19.32 |
| 1696 | 0.0109 | 0.00 | 0.00 | 19.30 |
| 1697 | 0.0119 | 0.00 | 0.00 | 19.29 |
| 1698 | 0.0115 | 0.00 | 0.00 | 19.28 |
| 1699 | 0.0096 | 0.00 | 0.00 | 19.27 |
| 1700 | 0.0103 | 0.00 | 0.00 | 19.26 |
| 1701 | 0.0083 | 0.00 | 0.00 | 19.25 |
| 1702 | 0.0099 | 0.00 | 0.00 | 19.24 |
| 1703 | 0.0135 | 0.00 | 0.00 | 19.23 |
| 1704 | 0.0071 | 0.00 | 0.00 | 19.22 |
| 1705 | 0.0087 | 0.00 | 0.00 | 19.21 |
| 1706 | 0.0064 | 0.00 | 0.00 | 19.20 |
| 1707 | 0.0093 | 0.00 | 0.00 | 19.19 |
| 1708 | 0.0122 | 0.00 | 0.00 | 19.19 |
| 1709 | 0.0112 | 0.00 | 0.00 | 19.17 |
| 1710 | 0.0112 | 0.00 | 0.00 | 19.16 |
| 1711 | 0.0122 | 0.00 | 0.00 | 19.15 |
| 1712 | 0.0115 | 0.00 | 0.00 | 19.14 |
| 1713 | 0.0096 | 0.00 | 0.00 | 19.13 |
| 1714 | 0.0087 | 0.00 | 0.00 | 19.12 |
| 1715 | 0.0109 | 0.00 | 0.00 | 19.11 |
| 1716 | 0.0135 | 0.00 | 0.00 | 19.10 |

| | | | | |
|---|---|---|---|---|
| 1717 | 0.0080 | 0.00 | 0.00 | 19.08 |
| 1718 | 0.0103 | 0.00 | 0.00 | 19.08 |
| 1719 | 0.0103 | 0.00 | 0.00 | 19.07 |
| 1720 | 0.0093 | 0.00 | 0.00 | 19.06 |
| 1721 | 0.0106 | 0.00 | 0.00 | 19.05 |
| 1722 | 0.0074 | 0.00 | 0.00 | 19.04 |
| 1723 | 0.0112 | 0.00 | 0.00 | 19.03 |
| 1724 | 0.0106 | 0.00 | 0.00 | 19.02 |
| 1725 | 0.0106 | 0.00 | 0.00 | 19.01 |
| 1726 | 0.0115 | 0.00 | 0.00 | 19.00 |
| 1727 | 0.0119 | 0.00 | 0.00 | 18.98 |
| 1728 | 0.0099 | 0.00 | 0.00 | 18.97 |
| 1729 | 0.0103 | 0.00 | 0.00 | 18.96 |
| 1730 | 0.0090 | 0.00 | 0.00 | 18.95 |
| 1731 | 0.0080 | 0.00 | 0.00 | 18.94 |
| 1732 | 0.0109 | 0.00 | 0.00 | 18.94 |
| 1733 | 0.0090 | 0.00 | 0.00 | 18.92 |
| 1734 | 0.0122 | 0.00 | 0.00 | 18.92 |
| 1735 | 0.0099 | 0.00 | 0.00 | 18.90 |
| 1736 | 0.0103 | 0.00 | 0.00 | 18.89 |
| 1737 | 0.0093 | 0.00 | 0.00 | 18.88 |
| 1738 | 0.0087 | 0.00 | 0.00 | 18.87 |
| 1739 | 0.0115 | 0.00 | 0.00 | 18.87 |
| 1740 | 0.0103 | 0.00 | 0.00 | 18.85 |
| 1741 | 0.0077 | 0.00 | 0.00 | 18.84 |
| 1742 | 0.0080 | 0.00 | 0.00 | 18.84 |
| 1743 | 0.0128 | 0.00 | 0.00 | 18.83 |
| 1744 | 0.0087 | 0.00 | 0.00 | 18.82 |
| 1745 | 0.0131 | 0.00 | 0.00 | 18.81 |
| 1746 | 0.0115 | 0.00 | 0.00 | 18.79 |
| 1747 | 0.0125 | 0.00 | 0.00 | 18.78 |
| 1748 | 0.0125 | 0.00 | 0.00 | 18.77 |
| 1749 | 0.0128 | 0.00 | 0.00 | 18.76 |
| 1750 | 0.0109 | 0.00 | 0.00 | 18.74 |
| 1751 | 0.0087 | 0.00 | 0.00 | 18.73 |
| 1752 | 0.0071 | 0.00 | 0.00 | 18.72 |
| 1753 | 0.0099 | 0.00 | 0.00 | 18.72 |
| 1754 | 0.0144 | 0.00 | 0.00 | 18.71 |
| 1755 | 0.0096 | 0.00 | 0.00 | 18.69 |
| 1756 | 0.0106 | 0.00 | 0.00 | 18.68 |
| 1757 | 0.0087 | 0.00 | 0.00 | 18.67 |
| 1758 | 0.0103 | 0.00 | 0.00 | 18.66 |
| 1759 | 0.0109 | 0.00 | 0.00 | 18.65 |
| 1760 | 0.0119 | 0.00 | 0.00 | 18.64 |

| | | | | |
|---|---|---|---|---|
| 1761 | 0.0115 | 0.00 | 0.00 | 18.63 |
| 1762 | 0.0122 | 0.00 | 0.00 | 18.62 |
| 1763 | 0.0122 | 0.00 | 0.00 | 18.61 |
| 1764 | 0.0109 | 0.00 | 0.00 | 18.60 |
| 1765 | 0.0090 | 0.00 | 0.00 | 18.58 |
| 1766 | 0.0106 | 0.00 | 0.00 | 18.58 |
| 1767 | 0.0087 | 0.00 | 0.00 | 18.56 |
| 1768 | 0.0103 | 0.00 | 0.00 | 18.56 |
| 1769 | 0.0122 | 0.00 | 0.00 | 18.55 |
| 1770 | 0.0141 | 0.00 | 0.00 | 18.53 |
| 1771 | 0.0112 | 0.00 | 0.00 | 18.52 |
| 1772 | 0.0071 | 0.00 | 0.00 | 18.51 |
| 1773 | 0.0128 | 0.00 | 0.00 | 18.50 |
| 1774 | 0.0122 | 0.00 | 0.00 | 18.49 |
| 1775 | 0.0138 | 0.00 | 0.00 | 18.48 |
| 1776 | 0.0122 | 0.00 | 0.00 | 18.46 |
| 1777 | 0.0122 | 0.00 | 0.00 | 18.45 |
| 1778 | 0.0119 | 0.00 | 0.00 | 18.44 |
| 1779 | 0.0115 | 0.00 | 0.00 | 18.43 |
| 1780 | 0.0109 | 0.00 | 0.00 | 18.41 |
| 1781 | 0.0106 | 0.00 | 0.00 | 18.40 |
| 1782 | 0.0096 | 0.00 | 0.00 | 18.39 |
| 1783 | 0.0106 | 0.00 | 0.00 | 18.38 |
| 1784 | 0.0071 | 0.00 | 0.00 | 18.37 |
| 1785 | 0.0096 | 0.00 | 0.00 | 18.37 |
| 1786 | 0.0103 | 0.00 | 0.00 | 18.36 |
| 1787 | 0.0115 | 0.00 | 0.00 | 18.35 |
| 1788 | 0.0083 | 0.00 | 0.00 | 18.33 |
| 1789 | 0.0138 | 0.00 | 0.00 | 18.33 |
| 1790 | 0.0099 | 0.00 | 0.00 | 18.31 |
| 1791 | 0.0131 | 0.00 | 0.00 | 18.30 |
| 1792 | 0.0093 | 0.00 | 0.00 | 18.29 |
| 1793 | 0.0090 | 0.00 | 0.00 | 18.28 |
| 1794 | 0.0067 | 0.00 | 0.00 | 18.27 |
| 1795 | 0.0099 | 0.00 | 0.00 | 18.26 |
| 1796 | 0.0093 | 0.00 | 0.00 | 18.25 |
| 1797 | 0.0141 | 0.00 | 0.00 | 18.25 |
| 1798 | 0.0112 | 0.00 | 0.00 | 18.23 |
| 1799 | 0.0106 | 0.00 | 0.00 | 18.22 |
| 1800 | 0.0067 | 0.00 | 0.00 | 18.21 |
| 1801 | 0.0122 | 0.00 | 0.00 | 18.20 |
| 1802 | 0.0131 | 0.00 | 0.00 | 18.19 |
| 1803 | 0.0131 | 0.00 | 0.00 | 18.18 |
| 1804 | 0.0115 | 0.00 | 0.00 | 18.16 |

| | | | | |
|---|---|---|---|---|
| 1805 | 0.0099 | 0.00 | 0.00 | 18.15 |
| 1806 | 0.0093 | 0.00 | 0.00 | 18.14 |
| 1807 | 0.0125 | 0.00 | 0.00 | 18.13 |
| 1808 | 0.0112 | 0.00 | 0.00 | 18.12 |
| 1809 | 0.0087 | 0.00 | 0.00 | 18.11 |
| 1810 | 0.0109 | 0.00 | 0.00 | 18.10 |
| 1811 | 0.0099 | 0.00 | 0.00 | 18.09 |
| 1812 | 0.0090 | 0.00 | 0.00 | 18.08 |
| 1813 | 0.0080 | 0.00 | 0.00 | 18.07 |
| 1814 | 0.0109 | 0.00 | 0.00 | 18.06 |
| 1815 | 0.0103 | 0.00 | 0.00 | 18.05 |
| 1816 | 0.0231 | 0.00 | 0.01 | 18.04 |
| 1817 | 0.0122 | 0.00 | 0.00 | 18.02 |
| 1818 | 0.0083 | 0.00 | 0.00 | 18.01 |
| 1819 | 0.0106 | 0.00 | 0.00 | 18.00 |
| 1820 | 0.0096 | 0.00 | 0.00 | 17.99 |
| 1821 | 0.0064 | 0.00 | 0.00 | 17.98 |
| 1822 | 0.0106 | 0.00 | 0.00 | 17.97 |
| 1823 | 0.0096 | 0.00 | 0.00 | 17.96 |
| 1824 | 0.0093 | 0.00 | 0.00 | 17.95 |
| 1825 | 0.0090 | 0.00 | 0.00 | 17.94 |
| 1826 | 0.0096 | 0.00 | 0.00 | 17.93 |
| 1827 | 0.0103 | 0.00 | 0.00 | 17.92 |
| 1828 | 0.0077 | 0.00 | 0.00 | 17.91 |
| 1829 | 0.0099 | 0.00 | 0.00 | 17.91 |
| 1830 | 0.0074 | 0.00 | 0.00 | 17.90 |
| 1831 | 0.0109 | 0.00 | 0.00 | 17.89 |
| 1832 | 0.0077 | 0.00 | 0.00 | 17.88 |
| 1833 | 0.0093 | 0.00 | 0.00 | 17.87 |
| 1834 | 0.0087 | 0.00 | 0.00 | 17.86 |
| 1835 | 0.0115 | 0.00 | 0.00 | 17.85 |
| 1836 | 0.0080 | 0.00 | 0.00 | 17.84 |
| 1837 | 0.0131 | 0.00 | 0.00 | 17.83 |
| 1838 | 0.0087 | 0.00 | 0.00 | 17.82 |
| 1839 | 0.0096 | 0.00 | 0.00 | 17.81 |
| 1840 | 0.0093 | 0.00 | 0.00 | 17.80 |
| 1841 | 0.0080 | 0.00 | 0.00 | 17.79 |
| 1842 | 0.0106 | 0.00 | 0.00 | 17.78 |
| 1843 | 0.0083 | 0.00 | 0.00 | 17.77 |
| 1844 | 0.0103 | 0.00 | 0.00 | 17.76 |
| 1845 | 0.0119 | 0.00 | 0.00 | 17.75 |
| 1846 | 0.0099 | 0.00 | 0.00 | 17.74 |
| 1847 | 0.0087 | 0.00 | 0.00 | 17.73 |
| 1848 | 0.0112 | 0.00 | 0.00 | 17.72 |

| | | | | |
|---|---|---|---|---|
| 1849 | 0.0112 | 0.00 | 0.00 | 17.71 |
| 1850 | 0.0087 | 0.00 | 0.00 | 17.70 |
| 1851 | 0.0090 | 0.00 | 0.00 | 17.69 |
| 1852 | 0.0112 | 0.00 | 0.00 | 17.68 |
| 1853 | 0.0099 | 0.00 | 0.00 | 17.67 |
| 1854 | 0.0080 | 0.00 | 0.00 | 17.66 |
| 1855 | 0.0115 | 0.00 | 0.00 | 17.65 |
| 1856 | 0.0103 | 0.00 | 0.00 | 17.64 |
| 1857 | 0.0080 | 0.00 | 0.00 | 17.63 |
| 1858 | 0.0131 | 0.00 | 0.00 | 17.63 |
| 1859 | 0.0058 | 0.00 | 0.00 | 17.61 |
| 1860 | 0.0080 | 0.00 | 0.00 | 17.61 |
| 1861 | 0.0080 | 0.00 | 0.00 | 17.60 |
| 1862 | 0.0099 | 0.00 | 0.00 | 17.59 |
| 1863 | 0.0083 | 0.00 | 0.00 | 17.58 |
| 1864 | 0.0083 | 0.00 | 0.00 | 17.57 |
| 1865 | 0.0090 | 0.00 | 0.00 | 17.56 |
| 1866 | 0.0115 | 0.00 | 0.00 | 17.55 |
| 1867 | 0.0106 | 0.00 | 0.00 | 17.54 |
| 1868 | 0.0071 | 0.00 | 0.00 | 17.53 |
| 1869 | 0.0131 | 0.00 | 0.00 | 17.53 |
| 1870 | 0.0083 | 0.00 | 0.00 | 17.51 |
| 1871 | 0.0112 | 0.00 | 0.00 | 17.50 |
| 1872 | 0.0083 | 0.00 | 0.00 | 17.49 |
| 1873 | 0.0064 | 0.00 | 0.00 | 17.48 |
| 1874 | 0.0074 | 0.00 | 0.00 | 17.48 |
| 1875 | 0.0061 | 0.00 | 0.00 | 17.47 |
| 1876 | 0.0106 | 0.00 | 0.00 | 17.46 |
| 1877 | 0.0083 | 0.00 | 0.00 | 17.45 |
| 1878 | 0.0083 | 0.00 | 0.00 | 17.45 |
| 1879 | 0.0109 | 0.00 | 0.00 | 17.44 |
| 1880 | 0.0099 | 0.00 | 0.00 | 17.43 |
| 1881 | 0.0090 | 0.00 | 0.00 | 17.42 |
| 1882 | 0.0119 | 0.00 | 0.00 | 17.41 |
| 1883 | 0.0074 | 0.00 | 0.00 | 17.40 |
| 1884 | 0.0093 | 0.00 | 0.00 | 17.39 |
| 1885 | 0.0093 | 0.00 | 0.00 | 17.38 |
| 1886 | 0.0083 | 0.00 | 0.00 | 17.37 |
| 1887 | 0.0067 | 0.00 | 0.00 | 17.36 |
| 1888 | 0.0096 | 0.00 | 0.00 | 17.35 |
| 1889 | 0.0087 | 0.00 | 0.00 | 17.34 |
| 1890 | 0.0112 | 0.00 | 0.00 | 17.34 |
| 1891 | 0.0112 | 0.00 | 0.00 | 17.33 |
| 1892 | 0.0083 | 0.00 | 0.00 | 17.31 |

| | | | | |
|---|---|---|---|---|
| 1893 | 0.0093 | 0.00 | 0.00 | 17.31 |
| 1894 | 0.0064 | 0.00 | 0.00 | 17.30 |
| 1895 | 0.0090 | 0.00 | 0.00 | 17.29 |
| 1896 | 0.0106 | 0.00 | 0.00 | 17.28 |
| 1897 | 0.0099 | 0.00 | 0.00 | 17.27 |
| 1898 | 0.0087 | 0.00 | 0.00 | 17.26 |
| 1899 | 0.0096 | 0.00 | 0.00 | 17.25 |
| 1900 | 0.0087 | 0.00 | 0.00 | 17.24 |
| 1901 | 0.0099 | 0.00 | 0.00 | 17.23 |
| 1902 | 0.0090 | 0.00 | 0.00 | 17.22 |
| 1903 | 0.0106 | 0.00 | 0.00 | 17.21 |
| 1904 | 0.0083 | 0.00 | 0.00 | 17.20 |
| 1905 | 0.0071 | 0.00 | 0.00 | 17.20 |
| 1906 | 0.0064 | 0.00 | 0.00 | 17.19 |
| 1907 | 0.0093 | 0.00 | 0.00 | 17.18 |
| 1908 | 0.0093 | 0.00 | 0.00 | 17.17 |
| 1909 | 0.0074 | 0.00 | 0.00 | 17.16 |
| 1910 | 0.0109 | 0.00 | 0.00 | 17.16 |
| 1911 | 0.0090 | 0.00 | 0.00 | 17.15 |
| 1912 | 0.0077 | 0.00 | 0.00 | 17.14 |
| 1913 | 0.0087 | 0.00 | 0.00 | 17.13 |
| 1914 | 0.0135 | 0.00 | 0.00 | 17.12 |
| 1915 | 0.0106 | 0.00 | 0.00 | 17.11 |
| 1916 | 0.0115 | 0.00 | 0.00 | 17.10 |
| 1917 | 0.0080 | 0.00 | 0.00 | 17.08 |
| 1918 | 0.0090 | 0.00 | 0.00 | 17.08 |
| 1919 | 0.0058 | 0.00 | 0.00 | 17.07 |
| 1920 | 0.0058 | 0.00 | 0.00 | 17.06 |
| 1921 | 0.0099 | 0.00 | 0.00 | 17.06 |
| 1922 | 0.0080 | 0.00 | 0.00 | 17.05 |
| 1923 | 0.0112 | 0.00 | 0.00 | 17.04 |
| 1924 | 0.0074 | 0.00 | 0.00 | 17.03 |
| 1925 | 0.0125 | 0.00 | 0.01 | 17.02 |
| 1926 | 0.0090 | 0.00 | 0.00 | 17.01 |
| 1927 | 0.0058 | 0.00 | 0.00 | 17.00 |
| 1928 | 0.0090 | 0.00 | 0.00 | 16.99 |
| 1929 | 0.0087 | 0.00 | 0.00 | 16.98 |
| 1930 | 0.0074 | 0.00 | 0.00 | 16.97 |
| 1931 | 0.0064 | 0.00 | 0.00 | 16.97 |
| 1932 | 0.0103 | 0.00 | 0.00 | 16.96 |
| 1933 | 0.0074 | 0.00 | 0.00 | 16.95 |
| 1934 | 0.0106 | 0.00 | 0.00 | 16.94 |
| 1935 | 0.0058 | 0.00 | 0.00 | 16.93 |
| 1936 | 0.0074 | 0.00 | 0.00 | 16.93 |

| | | | | |
|---|---|---|---|---|
| 1937 | 0.0067 | 0.00 | 0.00 | 16.92 |
| 1938 | 0.0115 | 0.00 | 0.00 | 16.91 |
| 1939 | 0.0071 | 0.00 | 0.00 | 16.90 |
| 1940 | 0.0080 | 0.00 | 0.00 | 16.89 |
| 1941 | 0.0071 | 0.00 | 0.00 | 16.89 |
| 1942 | 0.0074 | 0.00 | 0.00 | 16.88 |
| 1943 | 0.0074 | 0.00 | 0.00 | 16.87 |
| 1944 | 0.0067 | 0.00 | 0.00 | 16.86 |
| 1945 | 0.0093 | 0.00 | 0.00 | 16.86 |
| 1946 | 0.0106 | 0.00 | 0.00 | 16.85 |
| 1947 | 0.0048 | 0.00 | 0.00 | 16.84 |
| 1948 | 0.0083 | 0.00 | 0.00 | 16.83 |
| 1949 | 0.0103 | 0.00 | 0.00 | 16.82 |
| 1950 | 0.0080 | 0.00 | 0.00 | 16.81 |
| 1951 | 0.0138 | 0.00 | 0.00 | 16.81 |
| 1952 | 0.0080 | 0.00 | 0.00 | 16.79 |
| 1953 | 0.0067 | 0.00 | 0.00 | 16.78 |
| 1954 | 0.0083 | 0.00 | 0.00 | 16.78 |
| 1955 | 0.0080 | 0.00 | 0.00 | 16.77 |
| 1956 | 0.0067 | 0.00 | 0.00 | 16.76 |
| 1957 | 0.0077 | 0.00 | 0.00 | 16.75 |
| 1958 | 0.0103 | 0.00 | 0.00 | 16.75 |
| 1959 | 0.0067 | 0.00 | 0.00 | 16.74 |
| 1960 | 0.0090 | 0.00 | 0.00 | 16.73 |
| 1961 | 0.0067 | 0.00 | 0.00 | 16.72 |
| 1962 | 0.0074 | 0.00 | 0.00 | 16.71 |
| 1963 | 0.0048 | 0.00 | 0.00 | 16.71 |
| 1964 | 0.0058 | 0.00 | 0.00 | 16.70 |
| 1965 | 0.0087 | 0.00 | 0.00 | 16.70 |
| 1966 | 0.0106 | 0.00 | 0.00 | 16.69 |
| 1967 | 0.0061 | 0.00 | 0.00 | 16.68 |
| 1968 | 0.0067 | 0.00 | 0.00 | 16.67 |
| 1969 | 0.0071 | 0.00 | 0.00 | 16.66 |
| 1970 | 0.0071 | 0.00 | 0.00 | 16.66 |
| 1971 | 0.0087 | 0.00 | 0.00 | 16.65 |
| 1972 | 0.0051 | 0.00 | 0.00 | 16.64 |
| 1973 | 0.0074 | 0.00 | 0.00 | 16.64 |
| 1974 | 0.0054 | 0.00 | 0.00 | 16.63 |
| 1975 | 0.0048 | 0.00 | 0.00 | 16.62 |
| 1976 | 0.0080 | 0.00 | 0.00 | 16.62 |
| 1977 | 0.0083 | 0.00 | 0.00 | 16.61 |
| 1978 | 0.0071 | 0.00 | 0.00 | 16.60 |
| 1979 | 0.0083 | 0.00 | 0.00 | 16.60 |
| 1980 | 0.0064 | 0.00 | 0.00 | 16.59 |

| | | | | |
|---|---|---|---|---|
| 1981 | 0.0071 | 0.00 | 0.00 | 16.58 |
| 1982 | 0.0080 | 0.00 | 0.00 | 16.57 |
| 1983 | 0.0074 | 0.00 | 0.00 | 16.57 |
| 1984 | 0.0074 | 0.00 | 0.00 | 16.56 |
| 1985 | 0.0074 | 0.00 | 0.00 | 16.55 |
| 1986 | 0.0093 | 0.00 | 0.00 | 16.54 |
| 1987 | 0.0064 | 0.00 | 0.00 | 16.53 |
| 1988 | 0.0074 | 0.00 | 0.00 | 16.53 |
| 1989 | 0.0093 | 0.00 | 0.00 | 16.52 |
| 1990 | 0.0064 | 0.00 | 0.00 | 16.51 |
| 1991 | 0.0083 | 0.00 | 0.00 | 16.50 |
| 1992 | 0.0087 | 0.00 | 0.00 | 16.50 |
| 1993 | 0.0083 | 0.00 | 0.00 | 16.49 |
| 1994 | 0.0064 | 0.00 | 0.00 | 16.48 |
| 1995 | 0.0061 | 0.00 | 0.00 | 16.47 |
| 1996 | 0.0099 | 0.00 | 0.00 | 16.47 |
| 1997 | 0.0106 | 0.00 | 0.00 | 16.46 |
| 1998 | 0.0064 | 0.00 | 0.00 | 16.45 |
| 1999 | 0.0080 | 0.00 | 0.00 | 16.44 |
| 2000 | 0.0090 | 0.00 | 0.00 | 16.43 |
| 2001 | 0.0077 | 0.00 | 0.00 | 16.42 |
| 2002 | 0.0074 | 0.00 | 0.00 | 16.42 |
| 2003 | 0.0071 | 0.00 | 0.00 | 16.41 |
| 2004 | 0.0067 | 0.00 | 0.00 | 16.40 |
| 2005 | 0.0051 | 0.00 | 0.00 | 16.39 |
| 2006 | 0.0071 | 0.00 | 0.00 | 16.39 |
| 2007 | 0.0083 | 0.00 | 0.00 | 16.38 |
| 2008 | 0.0058 | 0.00 | 0.00 | 16.37 |
| 2009 | 0.0083 | 0.00 | 0.00 | 16.37 |
| 2010 | 0.0067 | 0.00 | 0.00 | 16.36 |
| 2011 | 0.0061 | 0.00 | 0.00 | 16.35 |
| 2012 | 0.0096 | 0.00 | 0.00 | 16.35 |
| 2013 | 0.0083 | 0.00 | 0.00 | 16.34 |
| 2014 | 0.0077 | 0.00 | 0.00 | 16.33 |
| 2015 | 0.0064 | 0.00 | 0.00 | 16.32 |
| 2016 | 0.0077 | 0.00 | 0.00 | 16.31 |
| 2017 | 0.0061 | 0.00 | 0.00 | 16.31 |
| 2018 | 0.0093 | 0.00 | 0.00 | 16.30 |
| 2019 | 0.0071 | 0.00 | 0.00 | 16.29 |
| 2020 | 0.0074 | 0.00 | 0.00 | 16.28 |
| 2021 | 0.0087 | 0.00 | 0.00 | 16.28 |
| 2022 | 0.0087 | 0.00 | 0.00 | 16.27 |
| 2023 | 0.0122 | 0.00 | 0.00 | 16.26 |
| 2024 | 0.0099 | 0.00 | 0.00 | 16.25 |

| | | | | |
|---|---|---|---|---|
| 2025 | 0.0093 | 0.00 | 0.00 | 16.24 |
| 2026 | 0.0096 | 0.00 | 0.00 | 16.23 |
| 2027 | 0.0077 | 0.00 | 0.00 | 16.22 |
| 2028 | 0.0087 | 0.00 | 0.00 | 16.21 |
| 2029 | 0.0083 | 0.00 | 0.00 | 16.20 |
| 2030 | 0.0054 | 0.00 | 0.00 | 16.19 |
| 2031 | 0.0103 | 0.00 | 0.00 | 16.19 |
| 2032 | 0.0074 | 0.00 | 0.00 | 16.18 |
| 2033 | 0.0077 | 0.00 | 0.00 | 16.17 |
| 2034 | 0.0067 | 0.00 | 0.00 | 16.16 |
| 2035 | 0.0087 | 0.00 | 0.00 | 16.16 |
| 2036 | 0.0106 | 0.00 | 0.00 | 16.15 |
| 2037 | 0.0099 | 0.00 | 0.00 | 16.14 |
| 2038 | 0.0080 | 0.00 | 0.00 | 16.13 |
| 2039 | 0.0096 | 0.00 | 0.00 | 16.12 |
| 2040 | 0.0074 | 0.00 | 0.00 | 16.11 |
| 2041 | 0.0042 | 0.00 | 0.00 | 16.10 |
| 2042 | 0.0106 | 0.00 | 0.00 | 16.10 |
| 2043 | 0.0083 | 0.00 | 0.00 | 16.09 |
| 2044 | 0.0090 | 0.00 | 0.00 | 16.08 |
| 2045 | 0.0071 | 0.00 | 0.00 | 16.07 |
| 2046 | 0.0096 | 0.00 | 0.00 | 16.06 |
| 2047 | 0.0054 | 0.00 | 0.00 | 16.05 |
| 2048 | 0.0054 | 0.00 | 0.00 | 16.05 |
| 2049 | 0.0067 | 0.00 | 0.00 | 16.04 |
| 2050 | 0.0083 | 0.00 | 0.00 | 16.04 |
| 2051 | 0.0096 | 0.00 | 0.00 | 16.03 |
| 2052 | 0.0077 | 0.00 | 0.00 | 16.02 |
| 2053 | 0.0071 | 0.00 | 0.00 | 16.01 |
| 2054 | 0.0074 | 0.00 | 0.00 | 16.00 |
| 2055 | 0.0074 | 0.00 | 0.00 | 16.00 |
| 2056 | 0.0074 | 0.00 | 0.00 | 15.99 |
| 2057 | 0.0071 | 0.00 | 0.00 | 15.98 |
| 2058 | 0.0077 | 0.00 | 0.00 | 15.97 |
| 2059 | 0.0058 | 0.00 | 0.00 | 15.97 |
| 2060 | 0.0077 | 0.00 | 0.00 | 15.96 |
| 2061 | 0.0090 | 0.00 | 0.00 | 15.95 |
| 2062 | 0.0058 | 0.00 | 0.00 | 15.94 |
| 2063 | 0.0054 | 0.00 | 0.00 | 15.94 |
| 2064 | 0.0096 | 0.00 | 0.00 | 15.93 |
| 2065 | 0.0064 | 0.00 | 0.00 | 15.92 |
| 2066 | 0.0087 | 0.00 | 0.00 | 15.92 |
| 2067 | 0.0074 | 0.00 | 0.00 | 15.91 |
| 2068 | 0.0090 | 0.00 | 0.00 | 15.90 |

| | | | | |
|---|---|---|---|---|
| 2069 | 0.0074 | 0.00 | 0.00 | 15.89 |
| 2070 | 0.0090 | 0.00 | 0.00 | 15.88 |
| 2071 | 0.0077 | 0.00 | 0.00 | 15.88 |
| 2072 | 0.0093 | 0.00 | 0.00 | 15.87 |
| 2073 | 0.0083 | 0.00 | 0.00 | 15.86 |
| 2074 | 0.0119 | 0.00 | 0.00 | 15.85 |
| 2075 | 0.0054 | 0.00 | 0.00 | 15.84 |
| 2076 | 0.0077 | 0.00 | 0.00 | 15.83 |
| 2077 | 0.0083 | 0.00 | 0.00 | 15.83 |
| 2078 | 0.0074 | 0.00 | 0.00 | 15.82 |
| 2079 | 0.0054 | 0.00 | 0.00 | 15.81 |
| 2080 | 0.0061 | 0.00 | 0.00 | 15.80 |
| 2081 | 0.0083 | 0.00 | 0.00 | 15.80 |
| 2082 | 0.0058 | 0.00 | 0.00 | 15.79 |
| 2083 | 0.0067 | 0.00 | 0.00 | 15.78 |
| 2084 | 0.0054 | 0.00 | 0.00 | 15.78 |
| 2085 | 0.0071 | 0.00 | 0.00 | 15.77 |
| 2086 | 0.0054 | 0.00 | 0.00 | 15.76 |
| 2087 | 0.0048 | 0.00 | 0.00 | 15.76 |
| 2088 | 0.0061 | 0.00 | 0.00 | 15.75 |
| 2089 | 0.0061 | 0.00 | 0.00 | 15.75 |
| 2090 | 0.0071 | 0.00 | 0.00 | 15.74 |
| 2091 | 0.0071 | 0.00 | 0.00 | 15.73 |
| 2092 | 0.0067 | 0.00 | 0.00 | 15.73 |
| 2093 | 0.0051 | 0.00 | 0.00 | 15.72 |
| 2094 | 0.0077 | 0.00 | 0.00 | 15.72 |
| 2095 | 0.0087 | 0.00 | 0.00 | 15.71 |
| 2096 | 0.0074 | 0.00 | 0.00 | 15.70 |
| 2097 | 0.0048 | 0.00 | 0.00 | 15.69 |
| 2098 | 0.0071 | 0.00 | 0.00 | 15.69 |
| 2099 | 0.0067 | 0.00 | 0.00 | 15.68 |
| 2100 | 0.0071 | 0.00 | 0.00 | 15.67 |
| 2101 | 0.0087 | 0.00 | 0.00 | 15.67 |
| 2102 | 0.0071 | 0.00 | 0.00 | 15.66 |
| 2103 | 0.0080 | 0.00 | 0.00 | 15.65 |
| 2104 | 0.0054 | 0.00 | 0.00 | 15.64 |
| 2105 | 0.0038 | 0.00 | 0.00 | 15.64 |
| 2106 | 0.0067 | 0.00 | 0.00 | 15.63 |
| 2107 | 0.0077 | 0.00 | 0.00 | 15.63 |
| 2108 | 0.0051 | 0.00 | 0.00 | 15.62 |
| 2109 | 0.0096 | 0.00 | 0.00 | 15.61 |
| 2110 | 0.0067 | 0.00 | 0.00 | 15.60 |
| 2111 | 0.0064 | 0.00 | 0.00 | 15.60 |
| 2112 | 0.0093 | 0.00 | 0.00 | 15.59 |

| | | | | |
|---|---|---|---|---|
| 2113 | 0.0093 | 0.00 | 0.00 | 15.58 |
| 2114 | 0.0083 | 0.00 | 0.00 | 15.57 |
| 2115 | 0.0042 | 0.00 | 0.00 | 15.56 |
| 2116 | 0.0061 | 0.00 | 0.00 | 15.56 |
| 2117 | 0.0067 | 0.00 | 0.00 | 15.55 |
| 2118 | 0.0058 | 0.00 | 0.00 | 15.55 |
| 2119 | 0.0064 | 0.00 | 0.00 | 15.54 |
| 2120 | 0.0064 | 0.00 | 0.00 | 15.54 |
| 2121 | 0.0077 | 0.00 | 0.00 | 15.53 |
| 2122 | 0.0051 | 0.00 | 0.00 | 15.52 |
| 2123 | 0.0054 | 0.00 | 0.00 | 15.52 |
| 2124 | 0.0048 | 0.00 | 0.00 | 15.51 |
| 2125 | 0.0083 | 0.00 | 0.00 | 15.51 |
| 2126 | 0.0042 | 0.00 | 0.00 | 15.50 |
| 2127 | 0.0061 | 0.00 | 0.00 | 15.49 |
| 2128 | 0.0042 | 0.00 | 0.00 | 15.49 |
| 2129 | 0.0083 | 0.00 | 0.00 | 15.48 |
| 2130 | 0.0074 | 0.00 | 0.00 | 15.47 |
| 2131 | 0.0054 | 0.00 | 0.00 | 15.47 |
| 2132 | 0.0071 | 0.00 | 0.00 | 15.46 |
| 2133 | 0.0058 | 0.00 | 0.00 | 15.45 |
| 2134 | 0.0048 | 0.00 | 0.00 | 15.45 |
| 2135 | 0.0071 | 0.00 | 0.00 | 15.44 |
| 2136 | 0.0077 | 0.00 | 0.00 | 15.44 |
| 2137 | 0.0067 | 0.00 | 0.00 | 15.43 |
| 2138 | 0.0071 | 0.00 | 0.00 | 15.42 |
| 2139 | 0.0087 | 0.00 | 0.00 | 15.42 |
| 2140 | 0.0067 | 0.00 | 0.00 | 15.41 |
| 2141 | 0.0042 | 0.00 | 0.00 | 15.40 |
| 2142 | 0.0048 | 0.00 | 0.00 | 15.40 |
| 2143 | 0.0048 | 0.00 | 0.00 | 15.39 |
| 2144 | 0.0067 | 0.00 | 0.00 | 15.39 |
| 2145 | 0.0064 | 0.00 | 0.00 | 15.38 |
| 2146 | 0.0064 | 0.00 | 0.00 | 15.37 |
| 2147 | 0.0051 | 0.00 | 0.00 | 15.37 |
| 2148 | 0.0093 | 0.00 | 0.00 | 15.36 |
| 2149 | 0.0061 | 0.00 | 0.00 | 15.35 |
| 2150 | 0.0064 | 0.00 | 0.00 | 15.35 |
| 2151 | 0.0074 | 0.00 | 0.00 | 15.34 |
| 2152 | 0.0054 | 0.00 | 0.00 | 15.33 |
| 2153 | 0.0058 | 0.00 | 0.00 | 15.33 |
| 2154 | 0.0071 | 0.00 | 0.00 | 15.32 |
| 2155 | 0.0064 | 0.00 | 0.00 | 15.31 |
| 2156 | 0.0067 | 0.00 | 0.00 | 15.31 |

| | | | | |
|---|---|---|---|---|
| 2157 | 0.0054 | 0.00 | 0.00 | 15.30 |
| 2158 | 0.0071 | 0.00 | 0.00 | 15.30 |
| 2159 | 0.0045 | 0.00 | 0.00 | 15.29 |
| 2160 | 0.0058 | 0.00 | 0.00 | 15.28 |
| 2161 | 0.0058 | 0.00 | 0.00 | 15.28 |
| 2162 | 0.0071 | 0.00 | 0.00 | 15.27 |
| 2163 | 0.0054 | 0.00 | 0.00 | 15.27 |
| 2164 | 0.0077 | 0.00 | 0.00 | 15.26 |
| 2165 | 0.0077 | 0.00 | 0.00 | 15.25 |
| 2166 | 0.0067 | 0.00 | 0.00 | 15.24 |
| 2167 | 0.0083 | 0.00 | 0.00 | 15.24 |
| 2168 | 0.0071 | 0.00 | 0.00 | 15.23 |
| 2169 | 0.0067 | 0.00 | 0.00 | 15.22 |
| 2170 | 0.0061 | 0.00 | 0.00 | 15.22 |
| 2171 | 0.0038 | 0.00 | 0.00 | 15.21 |
| 2172 | 0.0074 | 0.00 | 0.00 | 15.21 |
| 2173 | 0.0071 | 0.00 | 0.00 | 15.20 |
| 2174 | 0.0087 | 0.00 | 0.00 | 15.19 |
| 2175 | 0.0048 | 0.00 | 0.00 | 15.18 |
| 2176 | 0.0061 | 0.00 | 0.00 | 15.18 |
| 2177 | 0.0064 | 0.00 | 0.00 | 15.17 |
| 2178 | 0.0064 | 0.00 | 0.00 | 15.17 |
| 2179 | 0.0064 | 0.00 | 0.00 | 15.16 |
| 2180 | 0.0087 | 0.00 | 0.00 | 15.15 |
| 2181 | 0.0067 | 0.00 | 0.00 | 15.14 |
| 2182 | 0.0051 | 0.00 | 0.00 | 15.14 |
| 2183 | 0.0074 | 0.00 | 0.00 | 15.13 |
| 2184 | 0.0077 | 0.00 | 0.00 | 15.13 |
| 2185 | 0.0051 | 0.00 | 0.00 | 15.12 |
| 2186 | 0.0064 | 0.00 | 0.00 | 15.11 |
| 2187 | 0.0074 | 0.00 | 0.00 | 15.11 |
| 2188 | 0.0038 | 0.00 | 0.00 | 15.10 |
| 2189 | 0.0051 | 0.00 | 0.00 | 15.09 |
| 2190 | 0.0103 | 0.00 | 0.00 | 15.09 |
| 2191 | 0.0054 | 0.00 | 0.00 | 15.08 |
| 2192 | 0.0080 | 0.00 | 0.00 | 15.07 |
| 2193 | 0.0058 | 0.00 | 0.00 | 15.07 |
| 2194 | 0.0054 | 0.00 | 0.00 | 15.06 |
| 2195 | 0.0077 | 0.00 | 0.00 | 15.05 |
| 2196 | 0.0125 | 0.00 | 0.01 | 15.05 |
| 2197 | 0.0074 | 0.00 | 0.00 | 15.03 |
| 2198 | 0.0071 | 0.00 | 0.00 | 15.03 |
| 2199 | 0.0067 | 0.00 | 0.00 | 15.02 |
| 2200 | 0.0071 | 0.00 | 0.00 | 15.01 |

| | | | | |
|---|---|---|---|---|
| 2201 | 0.0090 | 0.00 | 0.00 | 15.01 |
| 2202 | 0.0074 | 0.00 | 0.00 | 15.00 |
| 2203 | 0.0042 | 0.00 | 0.00 | 14.99 |
| 2204 | 0.0080 | 0.00 | 0.00 | 14.99 |
| 2205 | 0.0077 | 0.00 | 0.00 | 14.98 |
| 2206 | 0.0083 | 0.00 | 0.00 | 14.97 |
| 2207 | 0.0087 | 0.00 | 0.00 | 14.96 |
| 2208 | 0.0054 | 0.00 | 0.00 | 14.95 |
| 2209 | 0.0054 | 0.00 | 0.00 | 14.95 |
| 2210 | 0.0054 | 0.00 | 0.00 | 14.94 |
| 2211 | 0.0035 | 0.00 | 0.00 | 14.94 |
| 2212 | 0.0077 | 0.00 | 0.00 | 14.93 |
| 2213 | 0.0099 | 0.00 | 0.00 | 14.93 |
| 2214 | 0.0045 | 0.00 | 0.00 | 14.92 |
| 2215 | 0.0103 | 0.00 | 0.00 | 14.91 |
| 2216 | 0.0048 | 0.00 | 0.00 | 14.90 |
| 2217 | 0.0071 | 0.00 | 0.00 | 14.90 |
| 2218 | 0.0077 | 0.00 | 0.00 | 14.89 |
| 2219 | 0.0067 | 0.00 | 0.00 | 14.88 |
| 2220 | 0.0096 | 0.00 | 0.00 | 14.87 |
| 2221 | 0.0080 | 0.00 | 0.00 | 14.86 |
| 2222 | 0.0099 | 0.00 | 0.00 | 14.86 |
| 2223 | 0.0080 | 0.00 | 0.00 | 14.85 |
| 2224 | 0.0077 | 0.00 | 0.00 | 14.84 |
| 2225 | 0.0061 | 0.00 | 0.00 | 14.83 |
| 2226 | 0.0071 | 0.00 | 0.00 | 14.83 |
| 2227 | 0.0064 | 0.00 | 0.00 | 14.82 |
| 2228 | 0.0071 | 0.00 | 0.00 | 14.81 |
| 2229 | 0.0067 | 0.00 | 0.00 | 14.80 |
| 2230 | 0.0029 | 0.00 | 0.00 | 14.80 |
| 2231 | 0.0071 | 0.00 | 0.00 | 14.79 |
| 2232 | 0.0048 | 0.00 | 0.00 | 14.79 |
| 2233 | 0.0054 | 0.00 | 0.00 | 14.78 |
| 2234 | 0.0048 | 0.00 | 0.00 | 14.78 |
| 2235 | 0.0067 | 0.00 | 0.00 | 14.77 |
| 2236 | 0.0103 | 0.00 | 0.00 | 14.77 |
| 2237 | 0.0064 | 0.00 | 0.00 | 14.76 |
| 2238 | 0.0077 | 0.00 | 0.00 | 14.75 |
| 2239 | 0.0064 | 0.00 | 0.00 | 14.74 |
| 2240 | 0.0077 | 0.00 | 0.00 | 14.74 |
| 2241 | 0.0054 | 0.00 | 0.00 | 14.73 |
| 2242 | 0.0074 | 0.00 | 0.00 | 14.72 |
| 2243 | 0.0080 | 0.00 | 0.00 | 14.71 |
| 2244 | 0.0071 | 0.00 | 0.00 | 14.71 |

| | | | | |
|---|---|---|---|---|
| 2245 | 0.0087 | 0.00 | 0.00 | 14.70 |
| 2246 | 0.0083 | 0.00 | 0.00 | 14.69 |
| 2247 | 0.0077 | 0.00 | 0.00 | 14.68 |
| 2248 | 0.0051 | 0.00 | 0.00 | 14.68 |
| 2249 | 0.0077 | 0.00 | 0.00 | 14.67 |
| 2250 | 0.0061 | 0.00 | 0.00 | 14.66 |
| 2251 | 0.0064 | 0.00 | 0.00 | 14.66 |
| 2252 | 0.0083 | 0.00 | 0.00 | 14.65 |
| 2253 | 0.0067 | 0.00 | 0.00 | 14.64 |
| 2254 | 0.0099 | 0.00 | 0.00 | 14.63 |
| 2255 | 0.0074 | 0.00 | 0.00 | 14.62 |
| 2256 | 0.0083 | 0.00 | 0.00 | 14.62 |
| 2257 | 0.0080 | 0.00 | 0.00 | 14.61 |
| 2258 | 0.0067 | 0.00 | 0.00 | 14.60 |
| 2259 | 0.0077 | 0.00 | 0.00 | 14.59 |
| 2260 | 0.0067 | 0.00 | 0.00 | 14.59 |
| 2261 | 0.0074 | 0.00 | 0.00 | 14.58 |
| 2262 | 0.0061 | 0.00 | 0.00 | 14.57 |
| 2263 | 0.0061 | 0.00 | 0.00 | 14.57 |
| 2264 | 0.0061 | 0.00 | 0.00 | 14.56 |
| 2265 | 0.0045 | 0.00 | 0.00 | 14.55 |
| 2266 | 0.0054 | 0.00 | 0.00 | 14.55 |
| 2267 | 0.0054 | 0.00 | 0.00 | 14.54 |
| 2268 | 0.0029 | 0.00 | 0.00 | 14.54 |
| 2269 | 0.0054 | 0.00 | 0.00 | 14.54 |
| 2270 | 0.0077 | 0.00 | 0.00 | 14.53 |
| 2271 | 0.0099 | 0.00 | 0.00 | 14.52 |
| 2272 | 0.0048 | 0.00 | 0.00 | 14.51 |
| 2273 | 0.0048 | 0.00 | 0.00 | 14.51 |
| 2274 | 0.0058 | 0.00 | 0.00 | 14.50 |
| 2275 | 0.0038 | 0.00 | 0.00 | 14.50 |
| 2276 | 0.0087 | 0.00 | 0.00 | 14.49 |
| 2277 | 0.0067 | 0.00 | 0.00 | 14.48 |
| 2278 | 0.0080 | 0.00 | 0.00 | 14.48 |
| 2279 | 0.0074 | 0.00 | 0.00 | 14.47 |
| 2280 | 0.0045 | 0.00 | 0.00 | 14.46 |
| 2281 | 0.0077 | 0.00 | 0.00 | 14.46 |
| 2282 | 0.0048 | 0.00 | 0.00 | 14.45 |
| 2283 | 0.0067 | 0.00 | 0.00 | 14.45 |
| 2284 | 0.0080 | 0.00 | 0.00 | 14.44 |
| 2285 | 0.0083 | 0.00 | 0.00 | 14.43 |
| 2286 | 0.0048 | 0.00 | 0.00 | 14.42 |
| 2287 | 0.0080 | 0.00 | 0.00 | 14.42 |
| 2288 | 0.0074 | 0.00 | 0.00 | 14.41 |

| | | | | |
|---|---|---|---|---|
| 2289 | 0.0080 | 0.00 | 0.00 | 14.40 |
| 2290 | 0.0061 | 0.00 | 0.00 | 14.39 |
| 2291 | 0.0074 | 0.00 | 0.00 | 14.39 |
| 2292 | 0.0042 | 0.00 | 0.00 | 14.38 |
| 2293 | 0.0054 | 0.00 | 0.00 | 14.38 |
| 2294 | 0.0064 | 0.00 | 0.00 | 14.37 |
| 2295 | 0.0083 | 0.00 | 0.00 | 14.37 |
| 2296 | 0.0048 | 0.00 | 0.00 | 14.36 |
| 2297 | 0.0080 | 0.00 | 0.00 | 14.35 |
| 2298 | 0.0051 | 0.00 | 0.00 | 14.34 |
| 2299 | 0.0071 | 0.00 | 0.00 | 14.34 |
| 2300 | 0.0064 | 0.00 | 0.00 | 14.33 |
| 2301 | 0.0087 | 0.00 | 0.00 | 14.33 |
| 2302 | 0.0090 | 0.00 | 0.00 | 14.32 |
| 2303 | 0.0080 | 0.00 | 0.00 | 14.31 |
| 2304 | 0.0067 | 0.00 | 0.00 | 14.30 |
| 2305 | 0.0064 | 0.00 | 0.00 | 14.29 |
| 2306 | 0.0074 | 0.00 | 0.00 | 14.29 |
| 2307 | 0.0054 | 0.00 | 0.00 | 14.28 |
| 2308 | 0.0058 | 0.00 | 0.00 | 14.27 |
| 2309 | 0.0058 | 0.00 | 0.00 | 14.27 |
| 2310 | 0.0058 | 0.00 | 0.00 | 14.26 |
| 2311 | 0.0061 | 0.00 | 0.00 | 14.26 |
| 2312 | 0.0087 | 0.00 | 0.00 | 14.25 |
| 2313 | 0.0080 | 0.00 | 0.00 | 14.24 |
| 2314 | 0.0051 | 0.00 | 0.00 | 14.23 |
| 2315 | 0.0064 | 0.00 | 0.00 | 14.23 |
| 2316 | 0.0051 | 0.00 | 0.00 | 14.22 |
| 2317 | 0.0077 | 0.00 | 0.00 | 14.22 |
| 2318 | 0.0042 | 0.00 | 0.00 | 14.21 |
| 2319 | 0.0074 | 0.00 | 0.00 | 14.21 |
| 2320 | 0.0067 | 0.00 | 0.00 | 14.20 |
| 2321 | 0.0093 | 0.00 | 0.00 | 14.19 |
| 2322 | 0.0083 | 0.00 | 0.00 | 14.18 |
| 2323 | 0.0071 | 0.00 | 0.00 | 14.17 |
| 2324 | 0.0083 | 0.00 | 0.00 | 14.17 |
| 2325 | 0.0061 | 0.00 | 0.00 | 14.16 |
| 2326 | 0.0042 | 0.00 | 0.00 | 14.15 |
| 2327 | 0.0083 | 0.00 | 0.00 | 14.15 |
| 2328 | 0.0071 | 0.00 | 0.00 | 14.14 |
| 2329 | 0.0074 | 0.00 | 0.00 | 14.13 |
| 2330 | 0.0067 | 0.00 | 0.00 | 14.13 |
| 2331 | 0.0083 | 0.00 | 0.00 | 14.12 |
| 2332 | 0.0157 | 0.00 | 0.01 | 14.11 |

| | | | | |
|---|---|---|---|---|
| 2333 | 0.0074 | 0.00 | 0.00 | 14.09 |
| 2334 | 0.0080 | 0.00 | 0.00 | 14.09 |
| 2335 | 0.0080 | 0.00 | 0.00 | 14.08 |
| 2336 | 0.0090 | 0.00 | 0.00 | 14.07 |
| 2337 | 0.0067 | 0.00 | 0.00 | 14.06 |
| 2338 | 0.0080 | 0.00 | 0.00 | 14.06 |
| 2339 | 0.0077 | 0.00 | 0.00 | 14.05 |
| 2340 | 0.0074 | 0.00 | 0.00 | 14.04 |
| 2341 | 0.0058 | 0.00 | 0.00 | 14.03 |
| 2342 | 0.0074 | 0.00 | 0.00 | 14.03 |
| 2343 | 0.0087 | 0.00 | 0.00 | 14.02 |
| 2344 | 0.0071 | 0.00 | 0.00 | 14.01 |
| 2345 | 0.0048 | 0.00 | 0.00 | 14.00 |
| 2346 | 0.0071 | 0.00 | 0.00 | 14.00 |
| 2347 | 0.0035 | 0.00 | 0.00 | 13.99 |
| 2348 | 0.0064 | 0.00 | 0.00 | 13.99 |
| 2349 | 0.0099 | 0.00 | 0.00 | 13.98 |
| 2350 | 0.0077 | 0.00 | 0.00 | 13.97 |
| 2351 | 0.0067 | 0.00 | 0.00 | 13.96 |
| 2352 | 0.0048 | 0.00 | 0.00 | 13.96 |
| 2353 | 0.0051 | 0.00 | 0.00 | 13.95 |
| 2354 | 0.0067 | 0.00 | 0.00 | 13.95 |
| 2355 | 0.0054 | 0.00 | 0.00 | 13.94 |
| 2356 | 0.0064 | 0.00 | 0.00 | 13.93 |
| 2357 | 0.0071 | 0.00 | 0.00 | 13.93 |
| 2358 | 0.0061 | 0.00 | 0.00 | 13.92 |
| 2359 | 0.0096 | 0.00 | 0.00 | 13.92 |
| 2360 | 0.0051 | 0.00 | 0.00 | 13.91 |
| 2361 | 0.0058 | 0.00 | 0.00 | 13.90 |
| 2362 | 0.0045 | 0.00 | 0.00 | 13.89 |
| 2363 | 0.0042 | 0.00 | 0.00 | 13.89 |
| 2364 | 0.0058 | 0.00 | 0.00 | 13.89 |
| 2365 | 0.0067 | 0.00 | 0.00 | 13.88 |
| 2366 | 0.0074 | 0.00 | 0.00 | 13.87 |
| 2367 | 0.0090 | 0.00 | 0.00 | 13.87 |
| 2368 | 0.0074 | 0.00 | 0.00 | 13.86 |
| 2369 | 0.0067 | 0.00 | 0.00 | 13.85 |
| 2370 | 0.0058 | 0.00 | 0.00 | 13.84 |
| 2371 | 0.0061 | 0.00 | 0.00 | 13.84 |
| 2372 | 0.0061 | 0.00 | 0.00 | 13.83 |
| 2373 | 0.0077 | 0.00 | 0.00 | 13.83 |
| 2374 | 0.0051 | 0.00 | 0.00 | 13.82 |
| 2375 | 0.0087 | 0.00 | 0.00 | 13.81 |
| 2376 | 0.0064 | 0.00 | 0.00 | 13.80 |

| | | | | |
|---|---|---|---|---|
| 2377 | 0.0064 | 0.00 | 0.00 | 13.80 |
| 2378 | 0.0067 | 0.00 | 0.00 | 13.79 |
| 2379 | 0.0071 | 0.00 | 0.00 | 13.78 |
| 2380 | 0.0080 | 0.00 | 0.00 | 13.78 |
| 2381 | 0.0058 | 0.00 | 0.00 | 13.77 |
| 2382 | 0.0071 | 0.00 | 0.00 | 13.76 |
| 2383 | 0.0067 | 0.00 | 0.00 | 13.76 |
| 2384 | 0.0071 | 0.00 | 0.00 | 13.75 |
| 2385 | 0.0051 | 0.00 | 0.00 | 13.74 |
| 2386 | 0.0067 | 0.00 | 0.00 | 13.74 |
| 2387 | 0.0058 | 0.00 | 0.00 | 13.73 |
| 2388 | 0.0064 | 0.00 | 0.00 | 13.73 |
| 2389 | 0.0071 | 0.00 | 0.00 | 13.72 |
| 2390 | 0.0035 | 0.00 | 0.00 | 13.71 |
| 2391 | 0.0054 | 0.00 | 0.00 | 13.71 |
| 2392 | 0.0093 | 0.00 | 0.00 | 13.70 |
| 2393 | 0.0045 | 0.00 | 0.00 | 13.69 |
| 2394 | 0.0077 | 0.00 | 0.00 | 13.69 |
| 2395 | 0.0067 | 0.00 | 0.00 | 13.68 |
| 2396 | 0.0112 | 0.00 | 0.00 | 13.67 |
| 2397 | 0.0054 | 0.00 | 0.00 | 13.66 |
| 2398 | 0.0071 | 0.00 | 0.00 | 13.66 |
| 2399 | 0.0048 | 0.00 | 0.00 | 13.65 |
| 2400 | 0.0071 | 0.00 | 0.00 | 13.65 |
| 2401 | 0.0064 | 0.00 | 0.00 | 13.64 |
| 2402 | 0.0071 | 0.00 | 0.00 | 13.63 |
| 2403 | 0.0093 | 0.00 | 0.00 | 13.63 |
| 2404 | 0.0077 | 0.00 | 0.00 | 13.62 |
| 2405 | 0.0074 | 0.00 | 0.00 | 13.61 |
| 2406 | 0.0064 | 0.00 | 0.00 | 13.60 |
| 2407 | 0.0077 | 0.00 | 0.00 | 13.59 |
| 2408 | 0.0077 | 0.00 | 0.00 | 13.59 |
| 2409 | 0.0071 | 0.00 | 0.00 | 13.58 |
| 2410 | 0.0083 | 0.00 | 0.00 | 13.57 |
| 2411 | 0.0038 | 0.00 | 0.00 | 13.56 |
| 2412 | 0.0051 | 0.00 | 0.00 | 13.56 |
| 2413 | 0.0064 | 0.00 | 0.00 | 13.55 |
| 2414 | 0.0071 | 0.00 | 0.00 | 13.55 |
| 2415 | 0.0061 | 0.00 | 0.00 | 13.54 |
| 2416 | 0.0083 | 0.00 | 0.00 | 13.54 |
| 2417 | 0.0080 | 0.00 | 0.00 | 13.53 |
| 2418 | 0.0074 | 0.00 | 0.00 | 13.52 |
| 2419 | 0.0090 | 0.00 | 0.00 | 13.51 |
| 2420 | 0.0119 | 0.00 | 0.00 | 13.50 |

| | | | | |
|---|---|---|---|---|
| 2421 | 0.0051 | 0.00 | 0.00 | 13.49 |
| 2422 | 0.0080 | 0.00 | 0.00 | 13.49 |
| 2423 | 0.0054 | 0.00 | 0.00 | 13.48 |
| 2424 | 0.0080 | 0.00 | 0.00 | 13.47 |
| 2425 | 0.0071 | 0.00 | 0.00 | 13.46 |
| 2426 | 0.0058 | 0.00 | 0.00 | 13.46 |
| 2427 | 0.0074 | 0.00 | 0.00 | 13.45 |
| 2428 | 0.0071 | 0.00 | 0.00 | 13.44 |
| 2429 | 0.0071 | 0.00 | 0.00 | 13.44 |
| 2430 | 0.0067 | 0.00 | 0.00 | 13.43 |
| 2431 | 0.0074 | 0.00 | 0.00 | 13.42 |
| 2432 | 0.0080 | 0.00 | 0.00 | 13.42 |
| 2433 | 0.0064 | 0.00 | 0.00 | 13.41 |
| 2434 | 0.0061 | 0.00 | 0.00 | 13.40 |
| 2435 | 0.0074 | 0.00 | 0.00 | 13.40 |
| 2436 | 0.0064 | 0.00 | 0.00 | 13.39 |
| 2437 | 0.0106 | 0.00 | 0.00 | 13.38 |
| 2438 | 0.0109 | 0.00 | 0.00 | 13.37 |
| 2439 | 0.0061 | 0.00 | 0.00 | 13.36 |
| 2440 | 0.0051 | 0.00 | 0.00 | 13.35 |
| 2441 | 0.0061 | 0.00 | 0.00 | 13.35 |
| 2442 | 0.0054 | 0.00 | 0.00 | 13.34 |
| 2443 | 0.0067 | 0.00 | 0.00 | 13.34 |
| 2444 | 0.0096 | 0.00 | 0.00 | 13.33 |
| 2445 | 0.0064 | 0.00 | 0.00 | 13.32 |
| 2446 | 0.0074 | 0.00 | 0.00 | 13.31 |
| 2447 | 0.0071 | 0.00 | 0.00 | 13.31 |
| 2448 | 0.0064 | 0.00 | 0.00 | 13.30 |
| 2449 | 0.0051 | 0.00 | 0.00 | 13.29 |
| 2450 | 0.0074 | 0.00 | 0.00 | 13.29 |
| 2451 | 0.0077 | 0.00 | 0.00 | 13.28 |
| 2452 | 0.0071 | 0.00 | 0.00 | 13.27 |
| 2453 | 0.0090 | 0.00 | 0.00 | 13.27 |
| 2454 | 0.0058 | 0.00 | 0.00 | 13.26 |
| 2455 | 0.0083 | 0.00 | 0.00 | 13.25 |
| 2456 | 0.0064 | 0.00 | 0.00 | 13.24 |
| 2457 | 0.0048 | 0.00 | 0.00 | 13.24 |
| 2458 | 0.0144 | 0.00 | 0.01 | 13.23 |
| 2459 | 0.0067 | 0.00 | 0.00 | 13.22 |
| 2460 | 0.0080 | 0.00 | 0.00 | 13.21 |
| 2461 | 0.0058 | 0.00 | 0.00 | 13.20 |
| 2462 | 0.0071 | 0.00 | 0.00 | 13.20 |
| 2463 | 0.0042 | 0.00 | 0.00 | 13.19 |
| 2464 | 0.0045 | 0.00 | 0.00 | 13.19 |

| | | | | |
|---|---|---|---|---|
| 2465 | 0.0087 | 0.00 | 0.00 | 13.18 |
| 2466 | 0.0045 | 0.00 | 0.00 | 13.17 |
| 2467 | 0.0067 | 0.00 | 0.00 | 13.17 |
| 2468 | 0.0051 | 0.00 | 0.00 | 13.16 |
| 2469 | 0.0093 | 0.00 | 0.00 | 13.16 |
| 2470 | 0.0051 | 0.00 | 0.00 | 13.15 |
| 2471 | 0.0035 | 0.00 | 0.00 | 13.14 |
| 2472 | 0.0058 | 0.00 | 0.00 | 13.14 |
| 2473 | 0.0087 | 0.00 | 0.00 | 13.13 |
| 2474 | 0.0058 | 0.00 | 0.00 | 13.12 |
| 2475 | 0.0064 | 0.00 | 0.00 | 13.12 |
| 2476 | 0.0051 | 0.00 | 0.00 | 13.11 |
| 2477 | 0.0087 | 0.00 | 0.00 | 13.11 |
| 2478 | 0.0106 | 0.00 | 0.00 | 13.10 |
| 2479 | 0.0071 | 0.00 | 0.00 | 13.09 |
| 2480 | 0.0058 | 0.00 | 0.00 | 13.08 |
| 2481 | 0.0058 | 0.00 | 0.00 | 13.07 |
| 2482 | 0.0077 | 0.00 | 0.00 | 13.07 |
| 2483 | 0.0048 | 0.00 | 0.00 | 13.06 |
| 2484 | 0.0067 | 0.00 | 0.00 | 13.06 |
| 2485 | 0.0051 | 0.00 | 0.00 | 13.05 |
| 2486 | 0.0067 | 0.00 | 0.00 | 13.04 |
| 2487 | 0.0042 | 0.00 | 0.00 | 13.04 |
| 2488 | 0.0048 | 0.00 | 0.00 | 13.03 |
| 2489 | 0.0071 | 0.00 | 0.00 | 13.03 |
| 2490 | 0.0058 | 0.00 | 0.00 | 13.02 |
| 2491 | 0.0048 | 0.00 | 0.00 | 13.02 |
| 2492 | 0.0045 | 0.00 | 0.00 | 13.01 |
| 2493 | 0.0048 | 0.00 | 0.00 | 13.01 |
| 2494 | 0.0048 | 0.00 | 0.00 | 13.00 |
| 2495 | 0.0038 | 0.00 | 0.00 | 13.00 |
| 2496 | 0.0067 | 0.00 | 0.00 | 12.99 |
| 2497 | 0.0058 | 0.00 | 0.00 | 12.99 |
| 2498 | 0.0061 | 0.00 | 0.00 | 12.98 |
| 2499 | 0.0038 | 0.00 | 0.00 | 12.97 |
| 2500 | 0.0077 | 0.00 | 0.00 | 12.97 |
| 2501 | 0.0048 | 0.00 | 0.00 | 12.96 |
| 2502 | 0.0032 | 0.00 | 0.00 | 12.96 |
| 2503 | 0.0071 | 0.00 | 0.00 | 12.96 |
| 2504 | 0.0051 | 0.00 | 0.00 | 12.95 |
| 2505 | 0.0071 | 0.00 | 0.00 | 12.94 |
| 2506 | 0.0074 | 0.00 | 0.00 | 12.94 |
| 2507 | 0.0071 | 0.00 | 0.00 | 12.93 |
| 2508 | 0.0064 | 0.00 | 0.00 | 12.92 |

| | | | | |
|---|---|---|---|---|
| 2509 | 0.0071 | 0.00 | 0.00 | 12.92 |
| 2510 | 0.0067 | 0.00 | 0.00 | 12.91 |
| 2511 | 0.0048 | 0.00 | 0.00 | 12.90 |
| 2512 | 0.0051 | 0.00 | 0.00 | 12.90 |
| 2513 | 0.0067 | 0.00 | 0.00 | 12.89 |
| 2514 | 0.0045 | 0.00 | 0.00 | 12.88 |
| 2515 | 0.0048 | 0.00 | 0.00 | 12.88 |
| 2516 | 0.0026 | 0.00 | 0.00 | 12.88 |
| 2517 | 0.0087 | 0.00 | 0.00 | 12.87 |
| 2518 | 0.0054 | 0.00 | 0.00 | 12.86 |
| 2519 | 0.0058 | 0.00 | 0.00 | 12.86 |
| 2520 | 0.0071 | 0.00 | 0.00 | 12.85 |
| 2521 | 0.0051 | 0.00 | 0.00 | 12.85 |
| 2522 | 0.0042 | 0.00 | 0.00 | 12.84 |
| 2523 | 0.0054 | 0.00 | 0.00 | 12.84 |
| 2524 | 0.0071 | 0.00 | 0.00 | 12.83 |
| 2525 | 0.0051 | 0.00 | 0.00 | 12.82 |
| 2526 | 0.0048 | 0.00 | 0.00 | 12.82 |
| 2527 | 0.0051 | 0.00 | 0.00 | 12.81 |
| 2528 | 0.0042 | 0.00 | 0.00 | 12.81 |
| 2529 | 0.0090 | 0.00 | 0.00 | 12.80 |
| 2530 | 0.0087 | 0.00 | 0.00 | 12.80 |
| 2531 | 0.0061 | 0.00 | 0.00 | 12.79 |
| 2532 | 0.0074 | 0.00 | 0.00 | 12.78 |
| 2533 | 0.0080 | 0.00 | 0.00 | 12.77 |
| 2534 | 0.0048 | 0.00 | 0.00 | 12.77 |
| 2535 | 0.0064 | 0.00 | 0.00 | 12.76 |
| 2536 | 0.0042 | 0.00 | 0.00 | 12.75 |
| 2537 | 0.0074 | 0.00 | 0.00 | 12.75 |
| 2538 | 0.0074 | 0.00 | 0.00 | 12.74 |
| 2539 | 0.0054 | 0.00 | 0.00 | 12.74 |
| 2540 | 0.0093 | 0.00 | 0.00 | 12.73 |
| 2541 | 0.0045 | 0.00 | 0.00 | 12.72 |
| 2542 | 0.0061 | 0.00 | 0.00 | 12.72 |
| 2543 | 0.0061 | 0.00 | 0.00 | 12.71 |
| 2544 | 0.0077 | 0.00 | 0.00 | 12.70 |
| 2545 | 0.0061 | 0.00 | 0.00 | 12.70 |
| 2546 | 0.0061 | 0.00 | 0.00 | 12.69 |
| 2547 | 0.0058 | 0.00 | 0.00 | 12.68 |
| 2548 | 0.0045 | 0.00 | 0.00 | 12.68 |
| 2549 | 0.0067 | 0.00 | 0.00 | 12.67 |
| 2550 | 0.0067 | 0.00 | 0.00 | 12.67 |
| 2551 | 0.0061 | 0.00 | 0.00 | 12.66 |
| 2552 | 0.0067 | 0.00 | 0.00 | 12.65 |

| | | | | |
|---|---|---|---|---|
| 2553 | 0.0071 | 0.00 | 0.00 | 12.65 |
| 2554 | 0.0026 | 0.00 | 0.00 | 12.64 |
| 2555 | 0.0074 | 0.00 | 0.00 | 12.64 |
| 2556 | 0.0074 | 0.00 | 0.00 | 12.63 |
| 2557 | 0.0077 | 0.00 | 0.00 | 12.62 |
| 2558 | 0.0045 | 0.00 | 0.00 | 12.62 |
| 2559 | 0.0074 | 0.00 | 0.00 | 12.61 |
| 2560 | 0.0045 | 0.00 | 0.00 | 12.60 |
| 2561 | 0.0058 | 0.00 | 0.00 | 12.60 |
| 2562 | 0.0067 | 0.00 | 0.00 | 12.59 |
| 2563 | 0.0064 | 0.00 | 0.00 | 12.59 |
| 2564 | 0.0054 | 0.00 | 0.00 | 12.58 |
| 2565 | 0.0048 | 0.00 | 0.00 | 12.58 |
| 2566 | 0.0054 | 0.00 | 0.00 | 12.57 |
| 2567 | 0.0074 | 0.00 | 0.00 | 12.56 |
| 2568 | 0.0061 | 0.00 | 0.00 | 12.56 |
| 2569 | 0.0067 | 0.00 | 0.00 | 12.55 |
| 2570 | 0.0051 | 0.00 | 0.00 | 12.54 |
| 2571 | 0.0045 | 0.00 | 0.00 | 12.54 |
| 2572 | 0.0045 | 0.00 | 0.00 | 12.53 |
| 2573 | 0.0054 | 0.00 | 0.00 | 12.53 |
| 2574 | 0.0061 | 0.00 | 0.00 | 12.53 |
| 2575 | 0.0083 | 0.00 | 0.00 | 12.52 |
| 2576 | 0.0051 | 0.00 | 0.00 | 12.51 |
| 2577 | 0.0058 | 0.00 | 0.00 | 12.51 |
| 2578 | 0.0083 | 0.00 | 0.00 | 12.50 |
| 2579 | 0.0064 | 0.00 | 0.00 | 12.49 |
| 2580 | 0.0064 | 0.00 | 0.00 | 12.48 |
| 2581 | 0.0067 | 0.00 | 0.00 | 12.48 |
| 2582 | 0.0042 | 0.00 | 0.00 | 12.47 |
| 2583 | 0.0058 | 0.00 | 0.00 | 12.47 |
| 2584 | 0.0071 | 0.00 | 0.00 | 12.46 |
| 2585 | 0.0077 | 0.00 | 0.00 | 12.45 |
| 2586 | 0.0054 | 0.00 | 0.00 | 12.45 |
| 2587 | 0.0061 | 0.00 | 0.00 | 12.44 |
| 2588 | 0.0080 | 0.00 | 0.00 | 12.44 |
| 2589 | 0.0071 | 0.00 | 0.00 | 12.43 |
| 2590 | 0.0058 | 0.00 | 0.00 | 12.42 |
| 2591 | 0.0074 | 0.00 | 0.00 | 12.41 |
| 2592 | 0.0054 | 0.00 | 0.00 | 12.41 |
| 2593 | 0.0064 | 0.00 | 0.00 | 12.40 |
| 2594 | 0.0071 | 0.00 | 0.00 | 12.40 |
| 2595 | 0.0048 | 0.00 | 0.00 | 12.39 |
| 2596 | 0.0045 | 0.00 | 0.00 | 12.38 |

| | | | | |
|---|---|---|---|---|
| 2597 | 0.0061 | 0.00 | 0.00 | 12.38 |
| 2598 | 0.0061 | 0.00 | 0.00 | 12.37 |
| 2599 | 0.0035 | 0.00 | 0.00 | 12.37 |
| 2600 | 0.0058 | 0.00 | 0.00 | 12.36 |
| 2601 | 0.0045 | 0.00 | 0.00 | 12.36 |
| 2602 | 0.0058 | 0.00 | 0.00 | 12.35 |
| 2603 | 0.0064 | 0.00 | 0.00 | 12.35 |
| 2604 | 0.0061 | 0.00 | 0.00 | 12.34 |
| 2605 | 0.0071 | 0.00 | 0.00 | 12.33 |
| 2606 | 0.0064 | 0.00 | 0.00 | 12.33 |
| 2607 | 0.0045 | 0.00 | 0.00 | 12.32 |
| 2608 | 0.0064 | 0.00 | 0.00 | 12.32 |
| 2609 | 0.0064 | 0.00 | 0.00 | 12.31 |
| 2610 | 0.0048 | 0.00 | 0.00 | 12.30 |
| 2611 | 0.0067 | 0.00 | 0.00 | 12.30 |
| 2612 | 0.0054 | 0.00 | 0.00 | 12.29 |
| 2613 | 0.0058 | 0.00 | 0.00 | 12.29 |
| 2614 | 0.0045 | 0.00 | 0.00 | 12.28 |
| 2615 | 0.0038 | 0.00 | 0.00 | 12.28 |
| 2616 | 0.0029 | 0.00 | 0.00 | 12.27 |
| 2617 | 0.0045 | 0.00 | 0.00 | 12.27 |
| 2618 | 0.0045 | 0.00 | 0.00 | 12.27 |
| 2619 | 0.0067 | 0.00 | 0.00 | 12.26 |
| 2620 | 0.0077 | 0.00 | 0.00 | 12.25 |
| 2621 | 0.0038 | 0.00 | 0.00 | 12.25 |
| 2622 | 0.0087 | 0.00 | 0.00 | 12.24 |
| 2623 | 0.0054 | 0.00 | 0.00 | 12.23 |
| 2624 | 0.0077 | 0.00 | 0.00 | 12.23 |
| 2625 | 0.0058 | 0.00 | 0.00 | 12.22 |
| 2626 | 0.0058 | 0.00 | 0.00 | 12.22 |
| 2627 | 0.0058 | 0.00 | 0.00 | 12.21 |
| 2628 | 0.0061 | 0.00 | 0.00 | 12.20 |
| 2629 | 0.0045 | 0.00 | 0.00 | 12.20 |
| 2630 | 0.0035 | 0.00 | 0.00 | 12.19 |
| 2631 | 0.0058 | 0.00 | 0.00 | 12.19 |
| 2632 | 0.0038 | 0.00 | 0.00 | 12.18 |
| 2633 | 0.0067 | 0.00 | 0.00 | 12.18 |
| 2634 | 0.0051 | 0.00 | 0.00 | 12.17 |
| 2635 | 0.0054 | 0.00 | 0.00 | 12.17 |
| 2636 | 0.0038 | 0.00 | 0.00 | 12.16 |
| 2637 | 0.0026 | 0.00 | 0.00 | 12.16 |
| 2638 | 0.0035 | 0.00 | 0.00 | 12.16 |
| 2639 | 0.0054 | 0.00 | 0.00 | 12.15 |
| 2640 | 0.0054 | 0.00 | 0.00 | 12.15 |

| | | | | |
|---|---|---|---|---|
| 2641 | 0.0032 | 0.00 | 0.00 | 12.14 |
| 2642 | 0.0074 | 0.00 | 0.00 | 12.14 |
| 2643 | 0.0080 | 0.00 | 0.00 | 12.13 |
| 2644 | 0.0026 | 0.00 | 0.00 | 12.12 |
| 2645 | 0.0042 | 0.00 | 0.00 | 12.12 |
| 2646 | 0.0038 | 0.00 | 0.00 | 12.12 |
| 2647 | 0.0035 | 0.00 | 0.00 | 12.11 |
| 2648 | 0.0042 | 0.00 | 0.00 | 12.11 |
| 2649 | 0.0064 | 0.00 | 0.00 | 12.11 |
| 2650 | 0.0016 | 0.00 | 0.00 | 12.10 |
| 2651 | 0.0054 | 0.00 | 0.00 | 12.10 |
| 2652 | 0.0083 | 0.00 | 0.00 | 12.09 |
| 2653 | 0.0074 | 0.00 | 0.00 | 12.08 |
| 2654 | 0.0045 | 0.00 | 0.00 | 12.08 |
| 2655 | 0.0042 | 0.00 | 0.00 | 12.07 |
| 2656 | 0.0054 | 0.00 | 0.00 | 12.07 |
| 2657 | 0.0051 | 0.00 | 0.00 | 12.06 |
| 2658 | 0.0051 | 0.00 | 0.00 | 12.06 |
| 2659 | 0.0061 | 0.00 | 0.00 | 12.05 |
| 2660 | 0.0074 | 0.00 | 0.00 | 12.05 |
| 2661 | 0.0051 | 0.00 | 0.00 | 12.04 |
| 2662 | 0.0061 | 0.00 | 0.00 | 12.03 |
| 2663 | 0.0061 | 0.00 | 0.00 | 12.03 |
| 2664 | 0.0064 | 0.00 | 0.00 | 12.02 |
| 2665 | 0.0054 | 0.00 | 0.00 | 12.01 |
| 2666 | 0.0042 | 0.00 | 0.00 | 12.01 |
| 2667 | 0.0061 | 0.00 | 0.00 | 12.00 |
| 2668 | 0.0058 | 0.00 | 0.00 | 12.00 |
| 2669 | 0.0061 | 0.00 | 0.00 | 11.99 |
| 2670 | 0.0051 | 0.00 | 0.00 | 11.99 |
| 2671 | 0.0051 | 0.00 | 0.00 | 11.98 |
| 2672 | 0.0042 | 0.00 | 0.00 | 11.98 |
| 2673 | 0.0042 | 0.00 | 0.00 | 11.97 |
| 2674 | 0.0048 | 0.00 | 0.00 | 11.97 |
| 2675 | 0.0051 | 0.00 | 0.00 | 11.96 |
| 2676 | 0.0061 | 0.00 | 0.00 | 11.96 |
| 2677 | 0.0051 | 0.00 | 0.00 | 11.95 |
| 2678 | 0.0058 | 0.00 | 0.00 | 11.95 |
| 2679 | 0.0061 | 0.00 | 0.00 | 11.94 |
| 2680 | 0.0061 | 0.00 | 0.00 | 11.94 |
| 2681 | 0.0064 | 0.00 | 0.00 | 11.93 |
| 2682 | 0.0061 | 0.00 | 0.00 | 11.92 |
| 2683 | 0.0045 | 0.00 | 0.00 | 11.92 |
| 2684 | 0.0051 | 0.00 | 0.00 | 11.91 |

| | | | | |
|---|---|---|---|---|
| 2685 | 0.0080 | 0.00 | 0.00 | 11.91 |
| 2686 | 0.0058 | 0.00 | 0.00 | 11.90 |
| 2687 | 0.0051 | 0.00 | 0.00 | 11.89 |
| 2688 | 0.0058 | 0.00 | 0.00 | 11.89 |
| 2689 | 0.0080 | 0.00 | 0.00 | 11.88 |
| 2690 | 0.0058 | 0.00 | 0.00 | 11.87 |
| 2691 | 0.0032 | 0.00 | 0.00 | 11.87 |
| 2692 | 0.0061 | 0.00 | 0.00 | 11.87 |
| 2693 | 0.0067 | 0.00 | 0.00 | 11.86 |
| 2694 | 0.0042 | 0.00 | 0.00 | 11.85 |
| 2695 | 0.0071 | 0.00 | 0.00 | 11.85 |
| 2696 | 0.0067 | 0.00 | 0.00 | 11.84 |
| 2697 | 0.0054 | 0.00 | 0.00 | 11.83 |
| 2698 | 0.0048 | 0.00 | 0.00 | 11.83 |
| 2699 | 0.0064 | 0.00 | 0.00 | 11.82 |
| 2700 | 0.0058 | 0.00 | 0.00 | 11.82 |
| 2701 | 0.0029 | 0.00 | 0.00 | 11.81 |
| 2702 | 0.0067 | 0.00 | 0.00 | 11.81 |
| 2703 | 0.0051 | 0.00 | 0.00 | 11.80 |
| 2704 | 0.0048 | 0.00 | 0.00 | 11.80 |
| 2705 | 0.0051 | 0.00 | 0.00 | 11.79 |
| 2706 | 0.0051 | 0.00 | 0.00 | 11.79 |
| 2707 | 0.0074 | 0.00 | 0.00 | 11.78 |
| 2708 | 0.0029 | 0.00 | 0.00 | 11.78 |
| 2709 | 0.0054 | 0.00 | 0.00 | 11.77 |
| 2710 | 0.0058 | 0.00 | 0.00 | 11.77 |
| 2711 | 0.0054 | 0.00 | 0.00 | 11.76 |
| 2712 | 0.0061 | 0.00 | 0.00 | 11.76 |
| 2713 | 0.0032 | 0.00 | 0.00 | 11.75 |
| 2714 | 0.0054 | 0.00 | 0.00 | 11.75 |
| 2715 | 0.0054 | 0.00 | 0.00 | 11.74 |
| 2716 | 0.0048 | 0.00 | 0.00 | 11.74 |
| 2717 | 0.0048 | 0.00 | 0.00 | 11.73 |
| 2718 | 0.0051 | 0.00 | 0.00 | 11.73 |
| 2719 | 0.0035 | 0.00 | 0.00 | 11.72 |
| 2720 | 0.0080 | 0.00 | 0.00 | 11.72 |
| 2721 | 0.0061 | 0.00 | 0.00 | 11.71 |
| 2722 | 0.0071 | 0.00 | 0.00 | 11.70 |
| 2723 | 0.0045 | 0.00 | 0.00 | 11.70 |
| 2724 | 0.0048 | 0.00 | 0.00 | 11.69 |
| 2725 | 0.0080 | 0.00 | 0.00 | 11.69 |
| 2726 | 0.0045 | 0.00 | 0.00 | 11.68 |
| 2727 | 0.0074 | 0.00 | 0.00 | 11.67 |
| 2728 | 0.0074 | 0.00 | 0.00 | 11.67 |

| | | | | |
|---|---|---|---|---|
| 2729 | 0.0038 | 0.00 | 0.00 | 11.66 |
| 2730 | 0.0090 | 0.00 | 0.00 | 11.66 |
| 2731 | 0.0077 | 0.00 | 0.00 | 11.65 |
| 2732 | 0.0099 | 0.00 | 0.00 | 11.64 |
| 2733 | 0.0071 | 0.00 | 0.00 | 11.63 |
| 2734 | 0.0071 | 0.00 | 0.00 | 11.62 |
| 2735 | 0.0067 | 0.00 | 0.00 | 11.61 |
| 2736 | 0.0042 | 0.00 | 0.00 | 11.61 |
| 2737 | 0.0064 | 0.00 | 0.00 | 11.60 |
| 2738 | 0.0035 | 0.00 | 0.00 | 11.60 |
| 2739 | 0.0054 | 0.00 | 0.00 | 11.59 |
| 2740 | 0.0054 | 0.00 | 0.00 | 11.59 |
| 2741 | 0.0122 | 0.00 | 0.01 | 11.58 |
| 2742 | 0.0074 | 0.00 | 0.00 | 11.57 |
| 2743 | 0.0035 | 0.00 | 0.00 | 11.56 |
| 2744 | 0.0064 | 0.00 | 0.00 | 11.56 |
| 2745 | 0.0042 | 0.00 | 0.00 | 11.55 |
| 2746 | 0.0045 | 0.00 | 0.00 | 11.55 |
| 2747 | 0.0061 | 0.00 | 0.00 | 11.54 |
| 2748 | 0.0042 | 0.00 | 0.00 | 11.54 |
| 2749 | 0.0032 | 0.00 | 0.00 | 11.53 |
| 2750 | 0.0029 | 0.00 | 0.00 | 11.53 |
| 2751 | 0.0067 | 0.00 | 0.00 | 11.53 |
| 2752 | 0.0045 | 0.00 | 0.00 | 11.52 |
| 2753 | 0.0042 | 0.00 | 0.00 | 11.52 |
| 2754 | 0.0042 | 0.00 | 0.00 | 11.51 |
| 2755 | 0.0051 | 0.00 | 0.00 | 11.51 |
| 2756 | 0.0058 | 0.00 | 0.00 | 11.50 |
| 2757 | 0.0061 | 0.00 | 0.00 | 11.50 |
| 2758 | 0.0032 | 0.00 | 0.00 | 11.49 |
| 2759 | 0.0042 | 0.00 | 0.00 | 11.49 |
| 2760 | 0.0048 | 0.00 | 0.00 | 11.48 |
| 2761 | 0.0026 | 0.00 | 0.00 | 11.48 |
| 2762 | 0.0048 | 0.00 | 0.00 | 11.48 |
| 2763 | 0.0042 | 0.00 | 0.00 | 11.47 |
| 2764 | 0.0035 | 0.00 | 0.00 | 11.47 |
| 2765 | 0.0054 | 0.00 | 0.00 | 11.46 |
| 2766 | 0.0035 | 0.00 | 0.00 | 11.46 |
| 2767 | 0.0054 | 0.00 | 0.00 | 11.46 |
| 2768 | 0.0042 | 0.00 | 0.00 | 11.45 |
| 2769 | 0.0032 | 0.00 | 0.00 | 11.45 |
| 2770 | 0.0051 | 0.00 | 0.00 | 11.44 |
| 2771 | 0.0038 | 0.00 | 0.00 | 11.44 |
| 2772 | 0.0032 | 0.00 | 0.00 | 11.43 |

| | | | | |
|---|---|---|---|---|
| 2773 | 0.0071 | 0.00 | 0.00 | 11.43 |
| 2774 | 0.0038 | 0.00 | 0.00 | 11.42 |
| 2775 | 0.0048 | 0.00 | 0.00 | 11.42 |
| 2776 | 0.0029 | 0.00 | 0.00 | 11.42 |
| 2777 | 0.0038 | 0.00 | 0.00 | 11.41 |
| 2778 | 0.0051 | 0.00 | 0.00 | 11.41 |
| 2779 | 0.0061 | 0.00 | 0.00 | 11.40 |
| 2780 | 0.0035 | 0.00 | 0.00 | 11.40 |
| 2781 | 0.0067 | 0.00 | 0.00 | 11.39 |
| 2782 | 0.0042 | 0.00 | 0.00 | 11.39 |
| 2783 | 0.0042 | 0.00 | 0.00 | 11.38 |
| 2784 | 0.0022 | 0.00 | 0.00 | 11.38 |
| 2785 | 0.0058 | 0.00 | 0.00 | 11.38 |
| 2786 | 0.0035 | 0.00 | 0.00 | 11.37 |
| 2787 | 0.0038 | 0.00 | 0.00 | 11.37 |
| 2788 | 0.0019 | 0.00 | 0.00 | 11.36 |
| 2789 | 0.0045 | 0.00 | 0.00 | 11.36 |
| 2790 | 0.0029 | 0.00 | 0.00 | 11.36 |
| 2791 | 0.0067 | 0.00 | 0.00 | 11.35 |
| 2792 | 0.0048 | 0.00 | 0.00 | 11.35 |
| 2793 | 0.0026 | 0.00 | 0.00 | 11.34 |
| 2794 | 0.0048 | 0.00 | 0.00 | 11.34 |
| 2795 | 0.0061 | 0.00 | 0.00 | 11.33 |
| 2796 | 0.0029 | 0.00 | 0.00 | 11.33 |
| 2797 | 0.0048 | 0.00 | 0.00 | 11.33 |
| 2798 | 0.0051 | 0.00 | 0.00 | 11.32 |
| 2799 | 0.0048 | 0.00 | 0.00 | 11.32 |
| 2800 | 0.0045 | 0.00 | 0.00 | 11.31 |
| 2801 | 0.0038 | 0.00 | 0.00 | 11.31 |
| 2802 | 0.0038 | 0.00 | 0.00 | 11.30 |
| 2803 | 0.0064 | 0.00 | 0.00 | 11.30 |
| 2804 | 0.0035 | 0.00 | 0.00 | 11.29 |
| 2805 | 0.0038 | 0.00 | 0.00 | 11.29 |
| 2806 | 0.0061 | 0.00 | 0.00 | 11.29 |
| 2807 | 0.0029 | 0.00 | 0.00 | 11.28 |
| 2808 | 0.0064 | 0.00 | 0.00 | 11.28 |
| 2809 | 0.0051 | 0.00 | 0.00 | 11.27 |
| 2810 | 0.0048 | 0.00 | 0.00 | 11.26 |
| 2811 | 0.0029 | 0.00 | 0.00 | 11.26 |
| 2812 | 0.0051 | 0.00 | 0.00 | 11.26 |
| 2813 | 0.0051 | 0.00 | 0.00 | 11.25 |
| 2814 | 0.0064 | 0.00 | 0.00 | 11.25 |
| 2815 | 0.0064 | 0.00 | 0.00 | 11.24 |
| 2816 | 0.0080 | 0.00 | 0.00 | 11.23 |

| | | | | |
|---|---|---|---|---|
| 2817 | 0.0029 | 0.00 | 0.00 | 11.23 |
| 2818 | 0.0029 | 0.00 | 0.00 | 11.22 |
| 2819 | 0.0067 | 0.00 | 0.00 | 11.22 |
| 2820 | 0.0042 | 0.00 | 0.00 | 11.21 |
| 2821 | 0.0038 | 0.00 | 0.00 | 11.21 |
| 2822 | 0.0061 | 0.00 | 0.00 | 11.21 |
| 2823 | 0.0048 | 0.00 | 0.00 | 11.20 |
| 2824 | 0.0038 | 0.00 | 0.00 | 11.19 |
| 2825 | 0.0042 | 0.00 | 0.00 | 11.19 |
| 2826 | 0.0029 | 0.00 | 0.00 | 11.19 |
| 2827 | 0.0038 | 0.00 | 0.00 | 11.18 |
| 2828 | 0.0035 | 0.00 | 0.00 | 11.18 |
| 2829 | 0.0038 | 0.00 | 0.00 | 11.18 |
| 2830 | 0.0045 | 0.00 | 0.00 | 11.17 |
| 2831 | 0.0035 | 0.00 | 0.00 | 11.17 |
| 2832 | 0.0051 | 0.00 | 0.00 | 11.16 |
| 2833 | 0.0042 | 0.00 | 0.00 | 11.16 |
| 2834 | 0.0042 | 0.00 | 0.00 | 11.16 |
| 2835 | 0.0051 | 0.00 | 0.00 | 11.15 |
| 2836 | 0.0045 | 0.00 | 0.00 | 11.15 |
| 2837 | 0.0035 | 0.00 | 0.00 | 11.14 |
| 2838 | 0.0038 | 0.00 | 0.00 | 11.14 |
| 2839 | 0.0032 | 0.00 | 0.00 | 11.13 |
| 2840 | 0.0038 | 0.00 | 0.00 | 11.13 |
| 2841 | 0.0045 | 0.00 | 0.00 | 11.13 |
| 2842 | 0.0038 | 0.00 | 0.00 | 11.12 |
| 2843 | 0.0029 | 0.00 | 0.00 | 11.12 |
| 2844 | 0.0064 | 0.00 | 0.00 | 11.12 |
| 2845 | 0.0048 | 0.00 | 0.00 | 11.11 |
| 2846 | 0.0035 | 0.00 | 0.00 | 11.10 |
| 2847 | 0.0051 | 0.00 | 0.00 | 11.10 |
| 2848 | 0.0032 | 0.00 | 0.00 | 11.10 |
| 2849 | 0.0042 | 0.00 | 0.00 | 11.09 |
| 2850 | 0.0051 | 0.00 | 0.00 | 11.09 |
| 2851 | 0.0026 | 0.00 | 0.00 | 11.08 |
| 2852 | 0.0029 | 0.00 | 0.00 | 11.08 |
| 2853 | 0.0054 | 0.00 | 0.00 | 11.08 |
| 2854 | 0.0045 | 0.00 | 0.00 | 11.07 |
| 2855 | 0.0026 | 0.00 | 0.00 | 11.07 |
| 2856 | 0.0026 | 0.00 | 0.00 | 11.07 |
| 2857 | 0.0051 | 0.00 | 0.00 | 11.06 |
| 2858 | 0.0026 | 0.00 | 0.00 | 11.06 |
| 2859 | 0.0042 | 0.00 | 0.00 | 11.06 |
| 2860 | 0.0051 | 0.00 | 0.00 | 11.05 |

| | | | | |
|---|---|---|---|---|
| 2861 | 0.0026 | 0.00 | 0.00 | 11.05 |
| 2862 | 0.0035 | 0.00 | 0.00 | 11.04 |
| 2863 | 0.0035 | 0.00 | 0.00 | 11.04 |
| 2864 | 0.0058 | 0.00 | 0.00 | 11.04 |
| 2865 | 0.0029 | 0.00 | 0.00 | 11.03 |
| 2866 | 0.0042 | 0.00 | 0.00 | 11.03 |
| 2867 | 0.0032 | 0.00 | 0.00 | 11.02 |
| 2868 | 0.0054 | 0.00 | 0.00 | 11.02 |
| 2869 | 0.0029 | 0.00 | 0.00 | 11.01 |
| 2870 | 0.0054 | 0.00 | 0.00 | 11.01 |
| 2871 | 0.0038 | 0.00 | 0.00 | 11.01 |
| 2872 | 0.0051 | 0.00 | 0.00 | 11.00 |
| 2873 | 0.0045 | 0.00 | 0.00 | 11.00 |
| 2874 | 0.0032 | 0.00 | 0.00 | 10.99 |
| 2875 | 0.0035 | 0.00 | 0.00 | 10.99 |
| 2876 | 0.0032 | 0.00 | 0.00 | 10.99 |
| 2877 | 0.0038 | 0.00 | 0.00 | 10.98 |
| 2878 | 0.0042 | 0.00 | 0.00 | 10.98 |
| 2879 | 0.0054 | 0.00 | 0.00 | 10.98 |
| 2880 | 0.0045 | 0.00 | 0.00 | 10.97 |
| 2881 | 0.0026 | 0.00 | 0.00 | 10.97 |
| 2882 | 0.0054 | 0.00 | 0.00 | 10.96 |
| 2883 | 0.0067 | 0.00 | 0.00 | 10.96 |
| 2884 | 0.0061 | 0.00 | 0.00 | 10.95 |
| 2885 | 0.0045 | 0.00 | 0.00 | 10.94 |
| 2886 | 0.0064 | 0.00 | 0.00 | 10.94 |
| 2887 | 0.0038 | 0.00 | 0.00 | 10.93 |
| 2888 | 0.0029 | 0.00 | 0.00 | 10.93 |
| 2889 | 0.0038 | 0.00 | 0.00 | 10.93 |
| 2890 | 0.0064 | 0.00 | 0.00 | 10.92 |
| 2891 | 0.0038 | 0.00 | 0.00 | 10.92 |
| 2892 | 0.0045 | 0.00 | 0.00 | 10.91 |
| 2893 | 0.0051 | 0.00 | 0.00 | 10.91 |
| 2894 | 0.0035 | 0.00 | 0.00 | 10.90 |
| 2895 | 0.0032 | 0.00 | 0.00 | 10.90 |
| 2896 | 0.0038 | 0.00 | 0.00 | 10.90 |
| 2897 | 0.0048 | 0.00 | 0.00 | 10.89 |
| 2898 | 0.0051 | 0.00 | 0.00 | 10.89 |
| 2899 | 0.0038 | 0.00 | 0.00 | 10.88 |
| 2900 | 0.0038 | 0.00 | 0.00 | 10.88 |
| 2901 | 0.0038 | 0.00 | 0.00 | 10.87 |
| 2902 | 0.0048 | 0.00 | 0.00 | 10.87 |
| 2903 | 0.0029 | 0.00 | 0.00 | 10.87 |
| 2904 | 0.0038 | 0.00 | 0.00 | 10.86 |

| | | | | |
|---|---|---|---|---|
| 2905 | 0.0045 | 0.00 | 0.00 | 10.86 |
| 2906 | 0.0035 | 0.00 | 0.00 | 10.85 |
| 2907 | 0.0048 | 0.00 | 0.00 | 10.85 |
| 2908 | 0.0051 | 0.00 | 0.00 | 10.85 |
| 2909 | 0.0035 | 0.00 | 0.00 | 10.84 |
| 2910 | 0.0032 | 0.00 | 0.00 | 10.84 |
| 2911 | 0.0038 | 0.00 | 0.00 | 10.83 |
| 2912 | 0.0029 | 0.00 | 0.00 | 10.83 |
| 2913 | 0.0038 | 0.00 | 0.00 | 10.83 |
| 2914 | 0.0038 | 0.00 | 0.00 | 10.82 |
| 2915 | 0.0029 | 0.00 | 0.00 | 10.82 |
| 2916 | 0.0058 | 0.00 | 0.00 | 10.82 |
| 2917 | 0.0054 | 0.00 | 0.00 | 10.81 |
| 2918 | 0.0022 | 0.00 | 0.00 | 10.81 |
| 2919 | 0.0051 | 0.00 | 0.00 | 10.80 |
| 2920 | 0.0042 | 0.00 | 0.00 | 10.80 |
| 2921 | 0.0048 | 0.00 | 0.00 | 10.79 |
| 2922 | 0.0029 | 0.00 | 0.00 | 10.79 |
| 2923 | 0.0054 | 0.00 | 0.00 | 10.79 |
| 2924 | 0.0042 | 0.00 | 0.00 | 10.78 |
| 2925 | 0.0051 | 0.00 | 0.00 | 10.78 |
| 2926 | 0.0038 | 0.00 | 0.00 | 10.77 |
| 2927 | 0.0029 | 0.00 | 0.00 | 10.77 |
| 2928 | 0.0035 | 0.00 | 0.00 | 10.77 |
| 2929 | 0.0054 | 0.00 | 0.00 | 10.76 |
| 2930 | 0.0054 | 0.00 | 0.00 | 10.76 |
| 2931 | 0.0032 | 0.00 | 0.00 | 10.75 |
| 2932 | 0.0051 | 0.00 | 0.00 | 10.75 |
| 2933 | 0.0026 | 0.00 | 0.00 | 10.74 |
| 2934 | 0.0029 | 0.00 | 0.00 | 10.74 |
| 2935 | 0.0051 | 0.00 | 0.00 | 10.74 |
| 2936 | 0.0051 | 0.00 | 0.00 | 10.73 |
| 2937 | 0.0077 | 0.00 | 0.00 | 10.73 |
| 2938 | 0.0042 | 0.00 | 0.00 | 10.72 |
| 2939 | 0.0093 | 0.00 | 0.00 | 10.72 |
| 2940 | 0.0045 | 0.00 | 0.00 | 10.71 |
| 2941 | 0.0054 | 0.00 | 0.00 | 10.70 |
| 2942 | 0.0054 | 0.00 | 0.00 | 10.70 |
| 2943 | 0.0051 | 0.00 | 0.00 | 10.69 |
| 2944 | 0.0048 | 0.00 | 0.00 | 10.69 |
| 2945 | 0.0045 | 0.00 | 0.00 | 10.68 |
| 2946 | 0.0048 | 0.00 | 0.00 | 10.68 |
| 2947 | 0.0054 | 0.00 | 0.00 | 10.67 |
| 2948 | 0.0045 | 0.00 | 0.00 | 10.67 |

| | | | | |
|---|---|---|---|---|
| 2949 | 0.0045 | 0.00 | 0.00 | 10.66 |
| 2950 | 0.0038 | 0.00 | 0.00 | 10.66 |
| 2951 | 0.0032 | 0.00 | 0.00 | 10.65 |
| 2952 | 0.0029 | 0.00 | 0.00 | 10.65 |
| 2953 | 0.0045 | 0.00 | 0.00 | 10.65 |
| 2954 | 0.0026 | 0.00 | 0.00 | 10.64 |
| 2955 | 0.0029 | 0.00 | 0.00 | 10.64 |
| 2956 | 0.0029 | 0.00 | 0.00 | 10.64 |
| 2957 | 0.0035 | 0.00 | 0.00 | 10.63 |
| 2958 | 0.0038 | 0.00 | 0.00 | 10.63 |
| 2959 | 0.0048 | 0.00 | 0.00 | 10.63 |
| 2960 | 0.0019 | 0.00 | 0.00 | 10.62 |
| 2961 | 0.0022 | 0.00 | 0.00 | 10.62 |
| 2962 | 0.0061 | 0.00 | 0.00 | 10.62 |
| 2963 | 0.0042 | 0.00 | 0.00 | 10.61 |
| 2964 | 0.0042 | 0.00 | 0.00 | 10.61 |
| 2965 | 0.0038 | 0.00 | 0.00 | 10.60 |
| 2966 | 0.0042 | 0.00 | 0.00 | 10.60 |
| 2967 | 0.0026 | 0.00 | 0.00 | 10.60 |
| 2968 | 0.0029 | 0.00 | 0.00 | 10.59 |
| 2969 | 0.0019 | 0.00 | 0.00 | 10.59 |
| 2970 | 0.0058 | 0.00 | 0.00 | 10.59 |
| 2971 | 0.0051 | 0.00 | 0.00 | 10.58 |
| 2972 | 0.0038 | 0.00 | 0.00 | 10.58 |
| 2973 | 0.0045 | 0.00 | 0.00 | 10.57 |
| 2974 | 0.0061 | 0.00 | 0.00 | 10.57 |
| 2975 | 0.0045 | 0.00 | 0.00 | 10.56 |
| 2976 | 0.0042 | 0.00 | 0.00 | 10.56 |
| 2977 | 0.0048 | 0.00 | 0.00 | 10.55 |
| 2978 | 0.0042 | 0.00 | 0.00 | 10.55 |
| 2979 | 0.0045 | 0.00 | 0.00 | 10.54 |
| 2980 | 0.0058 | 0.00 | 0.00 | 10.54 |
| 2981 | 0.0029 | 0.00 | 0.00 | 10.53 |
| 2982 | 0.0045 | 0.00 | 0.00 | 10.53 |
| 2983 | 0.0042 | 0.00 | 0.00 | 10.53 |
| 2984 | 0.0035 | 0.00 | 0.00 | 10.52 |
| 2985 | 0.0048 | 0.00 | 0.00 | 10.52 |
| 2986 | 0.0038 | 0.00 | 0.00 | 10.51 |
| 2987 | 0.0029 | 0.00 | 0.00 | 10.51 |
| 2988 | 0.0045 | 0.00 | 0.00 | 10.51 |
| 2989 | 0.0032 | 0.00 | 0.00 | 10.50 |
| 2990 | 0.0048 | 0.00 | 0.00 | 10.50 |
| 2991 | 0.0045 | 0.00 | 0.00 | 10.50 |
| 2992 | 0.0032 | 0.00 | 0.00 | 10.49 |

| | | | | |
|---|---|---|---|---|
| 2993 | 0.0029 | 0.00 | 0.00 | 10.49 |
| 2994 | 0.0048 | 0.00 | 0.00 | 10.48 |
| 2995 | 0.0022 | 0.00 | 0.00 | 10.48 |
| 2996 | 0.0022 | 0.00 | 0.00 | 10.48 |
| 2997 | 0.0035 | 0.00 | 0.00 | 10.48 |
| 2998 | 0.0019 | 0.00 | 0.00 | 10.47 |
| 2999 | 0.0054 | 0.00 | 0.00 | 10.47 |
| 3000 | 0.0048 | 0.00 | 0.00 | 10.46 |
| 3001 | 0.0035 | 0.00 | 0.00 | 10.46 |
| 3002 | 0.0042 | 0.00 | 0.00 | 10.46 |
| 3003 | 0.0051 | 0.00 | 0.00 | 10.45 |
| 3004 | 0.0045 | 0.00 | 0.00 | 10.45 |
| 3005 | 0.0042 | 0.00 | 0.00 | 10.44 |
| 3006 | 0.0042 | 0.00 | 0.00 | 10.44 |
| 3007 | 0.0035 | 0.00 | 0.00 | 10.43 |
| 3008 | 0.0035 | 0.00 | 0.00 | 10.43 |
| 3009 | 0.0038 | 0.00 | 0.00 | 10.43 |
| 3010 | 0.0035 | 0.00 | 0.00 | 10.42 |
| 3011 | 0.0045 | 0.00 | 0.00 | 10.42 |
| 3012 | 0.0054 | 0.00 | 0.00 | 10.42 |
| 3013 | 0.0048 | 0.00 | 0.00 | 10.41 |
| 3014 | 0.0038 | 0.00 | 0.00 | 10.41 |
| 3015 | 0.0048 | 0.00 | 0.00 | 10.40 |
| 3016 | 0.0029 | 0.00 | 0.00 | 10.40 |
| 3017 | 0.0035 | 0.00 | 0.00 | 10.39 |
| 3018 | 0.0045 | 0.00 | 0.00 | 10.39 |
| 3019 | 0.0045 | 0.00 | 0.00 | 10.39 |
| 3020 | 0.0038 | 0.00 | 0.00 | 10.38 |
| 3021 | 0.0051 | 0.00 | 0.00 | 10.38 |
| 3022 | 0.0022 | 0.00 | 0.00 | 10.37 |
| 3023 | 0.0045 | 0.00 | 0.00 | 10.37 |
| 3024 | 0.0029 | 0.00 | 0.00 | 10.37 |
| 3025 | 0.0035 | 0.00 | 0.00 | 10.36 |
| 3026 | 0.0035 | 0.00 | 0.00 | 10.36 |
| 3027 | 0.0054 | 0.00 | 0.00 | 10.36 |
| 3028 | 0.0061 | 0.00 | 0.00 | 10.35 |
| 3029 | 0.0038 | 0.00 | 0.00 | 10.34 |
| 3030 | 0.0042 | 0.00 | 0.00 | 10.34 |
| 3031 | 0.0038 | 0.00 | 0.00 | 10.34 |
| 3032 | 0.0080 | 0.00 | 0.00 | 10.33 |
| 3033 | 0.0032 | 0.00 | 0.00 | 10.32 |
| 3034 | 0.0051 | 0.00 | 0.00 | 10.32 |
| 3035 | 0.0048 | 0.00 | 0.00 | 10.32 |
| 3036 | 0.0058 | 0.00 | 0.00 | 10.31 |

| | | | | |
|---|---|---|---|---|
| 3037 | 0.0064 | 0.00 | 0.00 | 10.31 |
| 3038 | 0.0013 | 0.00 | 0.00 | 10.30 |
| 3039 | 0.0022 | 0.00 | 0.00 | 10.30 |
| 3040 | 0.0045 | 0.00 | 0.00 | 10.30 |
| 3041 | 0.0035 | 0.00 | 0.00 | 10.29 |
| 3042 | 0.0035 | 0.00 | 0.00 | 10.29 |
| 3043 | 0.0022 | 0.00 | 0.00 | 10.28 |
| 3044 | 0.0019 | 0.00 | 0.00 | 10.28 |
| 3045 | 0.0038 | 0.00 | 0.00 | 10.28 |
| 3046 | 0.0029 | 0.00 | 0.00 | 10.28 |
| 3047 | 0.0048 | 0.00 | 0.00 | 10.27 |
| 3048 | 0.0058 | 0.00 | 0.00 | 10.27 |
| 3049 | 0.0029 | 0.00 | 0.00 | 10.26 |
| 3050 | 0.0048 | 0.00 | 0.00 | 10.26 |
| 3051 | 0.0042 | 0.00 | 0.00 | 10.25 |
| 3052 | 0.0042 | 0.00 | 0.00 | 10.25 |
| 3053 | 0.0038 | 0.00 | 0.00 | 10.25 |
| 3054 | 0.0058 | 0.00 | 0.00 | 10.24 |
| 3055 | 0.0045 | 0.00 | 0.00 | 10.24 |
| 3056 | 0.0035 | 0.00 | 0.00 | 10.23 |
| 3057 | 0.0026 | 0.00 | 0.00 | 10.23 |
| 3058 | 0.0035 | 0.00 | 0.00 | 10.23 |
| 3059 | 0.0035 | 0.00 | 0.00 | 10.22 |
| 3060 | 0.0054 | 0.00 | 0.00 | 10.22 |
| 3061 | 0.0061 | 0.00 | 0.00 | 10.21 |
| 3062 | 0.0026 | 0.00 | 0.00 | 10.21 |
| 3063 | 0.0045 | 0.00 | 0.00 | 10.20 |
| 3064 | 0.0048 | 0.00 | 0.00 | 10.20 |
| 3065 | 0.0045 | 0.00 | 0.00 | 10.20 |
| 3066 | 0.0032 | 0.00 | 0.00 | 10.19 |
| 3067 | 0.0032 | 0.00 | 0.00 | 10.19 |
| 3068 | 0.0051 | 0.00 | 0.00 | 10.18 |
| 3069 | 0.0026 | 0.00 | 0.00 | 10.18 |
| 3070 | 0.0045 | 0.00 | 0.00 | 10.18 |
| 3071 | 0.0064 | 0.00 | 0.00 | 10.17 |
| 3072 | 0.0026 | 0.00 | 0.00 | 10.17 |
| 3073 | 0.0045 | 0.00 | 0.00 | 10.16 |
| 3074 | 0.0048 | 0.00 | 0.00 | 10.16 |
| 3075 | 0.0038 | 0.00 | 0.00 | 10.15 |
| 3076 | 0.0022 | 0.00 | 0.00 | 10.15 |
| 3077 | 0.0026 | 0.00 | 0.00 | 10.15 |
| 3078 | 0.0048 | 0.00 | 0.00 | 10.15 |
| 3079 | 0.0029 | 0.00 | 0.00 | 10.14 |
| 3080 | 0.0038 | 0.00 | 0.00 | 10.14 |

| | | | | |
|---|---|---|---|---|
| 3081 | 0.0054 | 0.00 | 0.00 | 10.13 |
| 3082 | 0.0045 | 0.00 | 0.00 | 10.13 |
| 3083 | 0.0045 | 0.00 | 0.00 | 10.12 |
| 3084 | 0.0042 | 0.00 | 0.00 | 10.12 |
| 3085 | 0.0045 | 0.00 | 0.00 | 10.12 |
| 3086 | 0.0038 | 0.00 | 0.00 | 10.11 |
| 3087 | 0.0051 | 0.00 | 0.00 | 10.11 |
| 3088 | 0.0026 | 0.00 | 0.00 | 10.10 |
| 3089 | 0.0038 | 0.00 | 0.00 | 10.10 |
| 3090 | 0.0045 | 0.00 | 0.00 | 10.10 |
| 3091 | 0.0032 | 0.00 | 0.00 | 10.09 |
| 3092 | 0.0035 | 0.00 | 0.00 | 10.09 |
| 3093 | 0.0051 | 0.00 | 0.00 | 10.08 |
| 3094 | 0.0026 | 0.00 | 0.00 | 10.08 |
| 3095 | 0.0022 | 0.00 | 0.00 | 10.08 |
| 3096 | 0.0061 | 0.00 | 0.00 | 10.07 |
| 3097 | 0.0038 | 0.00 | 0.00 | 10.07 |
| 3098 | 0.0035 | 0.00 | 0.00 | 10.06 |
| 3099 | 0.0038 | 0.00 | 0.00 | 10.06 |
| 3100 | 0.0042 | 0.00 | 0.00 | 10.06 |
| 3101 | 0.0051 | 0.00 | 0.00 | 10.05 |
| 3102 | 0.0035 | 0.00 | 0.00 | 10.05 |
| 3103 | 0.0051 | 0.00 | 0.00 | 10.04 |
| 3104 | 0.0038 | 0.00 | 0.00 | 10.04 |
| 3105 | 0.0022 | 0.00 | 0.00 | 10.04 |
| 3106 | 0.0058 | 0.00 | 0.00 | 10.03 |
| 3107 | 0.0048 | 0.00 | 0.00 | 10.03 |
| 3108 | 0.0038 | 0.00 | 0.00 | 10.02 |
| 3109 | 0.0058 | 0.00 | 0.00 | 10.02 |
| 3110 | 0.0038 | 0.00 | 0.00 | 10.01 |
| 3111 | 0.0035 | 0.00 | 0.00 | 10.01 |
| 3112 | 0.0038 | 0.00 | 0.00 | 10.01 |
| 3113 | 0.0058 | 0.00 | 0.00 | 10.00 |
| 3114 | 0.0019 | 0.00 | 0.00 | 10.00 |
| 3115 | 0.0048 | 0.00 | 0.00 | 9.99 |
| 3116 | 0.0038 | 0.00 | 0.00 | 9.99 |
| 3117 | 0.0048 | 0.00 | 0.00 | 9.99 |
| 3118 | 0.0045 | 0.00 | 0.00 | 9.98 |
| 3119 | 0.0035 | 0.00 | 0.00 | 9.98 |
| 3120 | 0.0038 | 0.00 | 0.00 | 9.97 |
| 3121 | 0.0042 | 0.00 | 0.00 | 9.97 |
| 3122 | 0.0048 | 0.00 | 0.00 | 9.96 |
| 3123 | 0.0061 | 0.00 | 0.00 | 9.96 |
| 3124 | 0.0042 | 0.00 | 0.00 | 9.95 |

| | | | | |
|---|---|---|---|---|
| 3125 | 0.0048 | 0.00 | 0.00 | 9.95 |
| 3126 | 0.0026 | 0.00 | 0.00 | 9.94 |
| 3127 | 0.0026 | 0.00 | 0.00 | 9.94 |
| 3128 | 0.0045 | 0.00 | 0.00 | 9.94 |
| 3129 | 0.0054 | 0.00 | 0.00 | 9.93 |
| 3130 | 0.0035 | 0.00 | 0.00 | 9.93 |
| 3131 | 0.0051 | 0.00 | 0.00 | 9.93 |
| 3132 | 0.0026 | 0.00 | 0.00 | 9.92 |
| 3133 | 0.0042 | 0.00 | 0.00 | 9.92 |
| 3134 | 0.0054 | 0.00 | 0.00 | 9.91 |
| 3135 | 0.0042 | 0.00 | 0.00 | 9.91 |
| 3136 | 0.0045 | 0.00 | 0.00 | 9.90 |
| 3137 | 0.0038 | 0.00 | 0.00 | 9.90 |
| 3138 | 0.0042 | 0.00 | 0.00 | 9.90 |
| 3139 | 0.0051 | 0.00 | 0.00 | 9.89 |
| 3140 | 0.0045 | 0.00 | 0.00 | 9.89 |
| 3141 | 0.0061 | 0.00 | 0.00 | 9.88 |
| 3142 | 0.0029 | 0.00 | 0.00 | 9.88 |
| 3143 | 0.0029 | 0.00 | 0.00 | 9.87 |
| 3144 | 0.0032 | 0.00 | 0.00 | 9.87 |
| 3145 | 0.0045 | 0.00 | 0.00 | 9.87 |
| 3146 | 0.0029 | 0.00 | 0.00 | 9.86 |
| 3147 | 0.0029 | 0.00 | 0.00 | 9.86 |
| 3148 | 0.0022 | 0.00 | 0.00 | 9.86 |
| 3149 | 0.0029 | 0.00 | 0.00 | 9.85 |
| 3150 | 0.0032 | 0.00 | 0.00 | 9.85 |
| 3151 | 0.0035 | 0.00 | 0.00 | 9.85 |
| 3152 | 0.0051 | 0.00 | 0.00 | 9.85 |
| 3153 | 0.0042 | 0.00 | 0.00 | 9.84 |
| 3154 | 0.0042 | 0.00 | 0.00 | 9.84 |
| 3155 | 0.0064 | 0.00 | 0.00 | 9.83 |
| 3156 | 0.0035 | 0.00 | 0.00 | 9.83 |
| 3157 | 0.0032 | 0.00 | 0.00 | 9.82 |
| 3158 | 0.0022 | 0.00 | 0.00 | 9.82 |
| 3159 | 0.0029 | 0.00 | 0.00 | 9.82 |
| 3160 | 0.0035 | 0.00 | 0.00 | 9.81 |
| 3161 | 0.0029 | 0.00 | 0.00 | 9.81 |
| 3162 | 0.0051 | 0.00 | 0.00 | 9.81 |
| 3163 | 0.0026 | 0.00 | 0.00 | 9.80 |
| 3164 | 0.0035 | 0.00 | 0.00 | 9.80 |
| 3165 | 0.0038 | 0.00 | 0.00 | 9.80 |
| 3166 | 0.0035 | 0.00 | 0.00 | 9.79 |
| 3167 | 0.0032 | 0.00 | 0.00 | 9.79 |
| 3168 | 0.0051 | 0.00 | 0.00 | 9.79 |

| | | | | |
|---|---|---|---|---|
| 3169 | 0.0061 | 0.00 | 0.00 | 9.78 |
| 3170 | 0.0061 | 0.00 | 0.00 | 9.77 |
| 3171 | 0.0019 | 0.00 | 0.00 | 9.77 |
| 3172 | 0.0058 | 0.00 | 0.00 | 9.77 |
| 3173 | 0.0045 | 0.00 | 0.00 | 9.76 |
| 3174 | 0.0064 | 0.00 | 0.00 | 9.76 |
| 3175 | 0.0035 | 0.00 | 0.00 | 9.75 |
| 3176 | 0.0045 | 0.00 | 0.00 | 9.75 |
| 3177 | 0.0032 | 0.00 | 0.00 | 9.74 |
| 3178 | 0.0038 | 0.00 | 0.00 | 9.74 |
| 3179 | 0.0016 | 0.00 | 0.00 | 9.73 |
| 3180 | 0.0038 | 0.00 | 0.00 | 9.73 |
| 3181 | 0.0051 | 0.00 | 0.00 | 9.73 |
| 3182 | 0.0035 | 0.00 | 0.00 | 9.72 |
| 3183 | 0.0038 | 0.00 | 0.00 | 9.72 |
| 3184 | 0.0035 | 0.00 | 0.00 | 9.72 |
| 3185 | 0.0035 | 0.00 | 0.00 | 9.71 |
| 3186 | 0.0048 | 0.00 | 0.00 | 9.71 |
| 3187 | 0.0029 | 0.00 | 0.00 | 9.70 |
| 3188 | 0.0048 | 0.00 | 0.00 | 9.70 |
| 3189 | 0.0042 | 0.00 | 0.00 | 9.70 |
| 3190 | 0.0019 | 0.00 | 0.00 | 9.69 |
| 3191 | 0.0035 | 0.00 | 0.00 | 9.69 |
| 3192 | 0.0032 | 0.00 | 0.00 | 9.69 |
| 3193 | 0.0029 | 0.00 | 0.00 | 9.68 |
| 3194 | 0.0022 | 0.00 | 0.00 | 9.68 |
| 3195 | 0.0067 | 0.00 | 0.00 | 9.68 |
| 3196 | 0.0042 | 0.00 | 0.00 | 9.67 |
| 3197 | 0.0026 | 0.00 | 0.00 | 9.67 |
| 3198 | 0.0045 | 0.00 | 0.00 | 9.67 |
| 3199 | 0.0029 | 0.00 | 0.00 | 9.66 |
| 3200 | 0.0051 | 0.00 | 0.00 | 9.66 |
| 3201 | 0.0035 | 0.00 | 0.00 | 9.65 |
| 3202 | 0.0019 | 0.00 | 0.00 | 9.65 |
| 3203 | 0.0058 | 0.00 | 0.00 | 9.65 |
| 3204 | 0.0029 | 0.00 | 0.00 | 9.64 |
| 3205 | 0.0026 | 0.00 | 0.00 | 9.64 |
| 3206 | 0.0029 | 0.00 | 0.00 | 9.64 |
| 3207 | 0.0048 | 0.00 | 0.00 | 9.63 |
| 3208 | 0.0058 | 0.00 | 0.00 | 9.63 |
| 3209 | 0.0038 | 0.00 | 0.00 | 9.62 |
| 3210 | 0.0022 | 0.00 | 0.00 | 9.62 |
| 3211 | 0.0038 | 0.00 | 0.00 | 9.62 |
| 3212 | 0.0042 | 0.00 | 0.00 | 9.61 |

| | | | | |
|---|---|---|---|---|
| 3213 | 0.0058 | 0.00 | 0.00 | 9.61 |
| 3214 | 0.0029 | 0.00 | 0.00 | 9.60 |
| 3215 | 0.0026 | 0.00 | 0.00 | 9.60 |
| 3216 | 0.0038 | 0.00 | 0.00 | 9.60 |
| 3217 | 0.0042 | 0.00 | 0.00 | 9.59 |
| 3218 | 0.0038 | 0.00 | 0.00 | 9.59 |
| 3219 | 0.0026 | 0.00 | 0.00 | 9.59 |
| 3220 | 0.0029 | 0.00 | 0.00 | 9.58 |
| 3221 | 0.0026 | 0.00 | 0.00 | 9.58 |
| 3222 | 0.0029 | 0.00 | 0.00 | 9.58 |
| 3223 | 0.0026 | 0.00 | 0.00 | 9.57 |
| 3224 | 0.0054 | 0.00 | 0.00 | 9.57 |
| 3225 | 0.0054 | 0.00 | 0.00 | 9.57 |
| 3226 | 0.0026 | 0.00 | 0.00 | 9.56 |
| 3227 | 0.0042 | 0.00 | 0.00 | 9.56 |
| 3228 | 0.0042 | 0.00 | 0.00 | 9.55 |
| 3229 | 0.0032 | 0.00 | 0.00 | 9.55 |
| 3230 | 0.0058 | 0.00 | 0.00 | 9.55 |
| 3231 | 0.0051 | 0.00 | 0.00 | 9.54 |
| 3232 | 0.0035 | 0.00 | 0.00 | 9.54 |
| 3233 | 0.0026 | 0.00 | 0.00 | 9.53 |
| 3234 | 0.0045 | 0.00 | 0.00 | 9.53 |
| 3235 | 0.0038 | 0.00 | 0.00 | 9.53 |
| 3236 | 0.0035 | 0.00 | 0.00 | 9.52 |
| 3237 | 0.0035 | 0.00 | 0.00 | 9.52 |
| 3238 | 0.0051 | 0.00 | 0.00 | 9.51 |
| 3239 | 0.0032 | 0.00 | 0.00 | 9.51 |
| 3240 | 0.0045 | 0.00 | 0.00 | 9.51 |
| 3241 | 0.0029 | 0.00 | 0.00 | 9.50 |
| 3242 | 0.0035 | 0.00 | 0.00 | 9.50 |
| 3243 | 0.0026 | 0.00 | 0.00 | 9.50 |
| 3244 | 0.0054 | 0.00 | 0.00 | 9.49 |
| 3245 | 0.0032 | 0.00 | 0.00 | 9.49 |
| 3246 | 0.0045 | 0.00 | 0.00 | 9.48 |
| 3247 | 0.0042 | 0.00 | 0.00 | 9.48 |
| 3248 | 0.0042 | 0.00 | 0.00 | 9.48 |
| 3249 | 0.0029 | 0.00 | 0.00 | 9.47 |
| 3250 | 0.0045 | 0.00 | 0.00 | 9.47 |
| 3251 | 0.0045 | 0.00 | 0.00 | 9.46 |
| 3252 | 0.0058 | 0.00 | 0.00 | 9.46 |
| 3253 | 0.0061 | 0.00 | 0.00 | 9.45 |
| 3254 | 0.0026 | 0.00 | 0.00 | 9.45 |
| 3255 | 0.0045 | 0.00 | 0.00 | 9.45 |
| 3256 | 0.0054 | 0.00 | 0.00 | 9.44 |

| | | | | |
|---|---|---|---|---|
| 3257 | 0.0038 | 0.00 | 0.00 | 9.44 |
| 3258 | 0.0026 | 0.00 | 0.00 | 9.43 |
| 3259 | 0.0035 | 0.00 | 0.00 | 9.43 |
| 3260 | 0.0038 | 0.00 | 0.00 | 9.43 |
| 3261 | 0.0016 | 0.00 | 0.00 | 9.42 |
| 3262 | 0.0042 | 0.00 | 0.00 | 9.42 |
| 3263 | 0.0022 | 0.00 | 0.00 | 9.42 |
| 3264 | 0.0038 | 0.00 | 0.00 | 9.41 |
| 3265 | 0.0032 | 0.00 | 0.00 | 9.41 |
| 3266 | 0.0035 | 0.00 | 0.00 | 9.41 |
| 3267 | 0.0029 | 0.00 | 0.00 | 9.40 |
| 3268 | 0.0022 | 0.00 | 0.00 | 9.40 |
| 3269 | 0.0022 | 0.00 | 0.00 | 9.40 |
| 3270 | 0.0058 | 0.00 | 0.00 | 9.40 |
| 3271 | 0.0016 | 0.00 | 0.00 | 9.39 |
| 3272 | 0.0061 | 0.00 | 0.00 | 9.39 |
| 3273 | 0.0054 | 0.00 | 0.00 | 9.38 |
| 3274 | 0.0026 | 0.00 | 0.00 | 9.38 |
| 3275 | 0.0029 | 0.00 | 0.00 | 9.37 |
| 3276 | 0.0032 | 0.00 | 0.00 | 9.37 |
| 3277 | 0.0035 | 0.00 | 0.00 | 9.37 |
| 3278 | 0.0038 | 0.00 | 0.00 | 9.36 |
| 3279 | 0.0042 | 0.00 | 0.00 | 9.36 |
| 3280 | 0.0042 | 0.00 | 0.00 | 9.36 |
| 3281 | 0.0029 | 0.00 | 0.00 | 9.35 |
| 3282 | 0.0051 | 0.00 | 0.00 | 9.35 |
| 3283 | 0.0029 | 0.00 | 0.00 | 9.34 |
| 3284 | 0.0035 | 0.00 | 0.00 | 9.34 |
| 3285 | 0.0029 | 0.00 | 0.00 | 9.34 |
| 3286 | 0.0032 | 0.00 | 0.00 | 9.33 |
| 3287 | 0.0026 | 0.00 | 0.00 | 9.33 |
| 3288 | 0.0042 | 0.00 | 0.00 | 9.33 |
| 3289 | 0.0035 | 0.00 | 0.00 | 9.33 |
| 3290 | 0.0029 | 0.00 | 0.00 | 9.32 |
| 3291 | 0.0032 | 0.00 | 0.00 | 9.32 |
| 3292 | 0.0029 | 0.00 | 0.00 | 9.32 |
| 3293 | 0.0035 | 0.00 | 0.00 | 9.31 |
| 3294 | 0.0035 | 0.00 | 0.00 | 9.31 |
| 3295 | 0.0035 | 0.00 | 0.00 | 9.31 |
| 3296 | 0.0032 | 0.00 | 0.00 | 9.30 |
| 3297 | 0.0042 | 0.00 | 0.00 | 9.30 |
| 3298 | 0.0038 | 0.00 | 0.00 | 9.29 |
| 3299 | 0.0045 | 0.00 | 0.00 | 9.29 |
| 3300 | 0.0042 | 0.00 | 0.00 | 9.29 |

| | | | | |
|---|---|---|---|---|
| 3301 | 0.0048 | 0.00 | 0.00 | 9.28 |
| 3302 | 0.0029 | 0.00 | 0.00 | 9.28 |
| 3303 | 0.0035 | 0.00 | 0.00 | 9.27 |
| 3304 | 0.0045 | 0.00 | 0.00 | 9.27 |
| 3305 | 0.0038 | 0.00 | 0.00 | 9.27 |
| 3306 | 0.0058 | 0.00 | 0.00 | 9.26 |
| 3307 | 0.0048 | 0.00 | 0.00 | 9.26 |
| 3308 | 0.0045 | 0.00 | 0.00 | 9.25 |
| 3309 | 0.0061 | 0.00 | 0.00 | 9.25 |
| 3310 | 0.0022 | 0.00 | 0.00 | 9.24 |
| 3311 | 0.0032 | 0.00 | 0.00 | 9.24 |
| 3312 | 0.0042 | 0.00 | 0.00 | 9.24 |
| 3313 | 0.0038 | 0.00 | 0.00 | 9.23 |
| 3314 | 0.0022 | 0.00 | 0.00 | 9.23 |
| 3315 | 0.0035 | 0.00 | 0.00 | 9.23 |
| 3316 | 0.0032 | 0.00 | 0.00 | 9.22 |
| 3317 | 0.0038 | 0.00 | 0.00 | 9.22 |
| 3318 | 0.0022 | 0.00 | 0.00 | 9.22 |
| 3319 | 0.0054 | 0.00 | 0.00 | 9.21 |
| 3320 | 0.0016 | 0.00 | 0.00 | 9.21 |
| 3321 | 0.0054 | 0.00 | 0.00 | 9.21 |
| 3322 | 0.0029 | 0.00 | 0.00 | 9.20 |
| 3323 | 0.0051 | 0.00 | 0.00 | 9.20 |
| 3324 | 0.0026 | 0.00 | 0.00 | 9.19 |
| 3325 | 0.0032 | 0.00 | 0.00 | 9.19 |
| 3326 | 0.0035 | 0.00 | 0.00 | 9.19 |
| 3327 | 0.0045 | 0.00 | 0.00 | 9.18 |
| 3328 | 0.0045 | 0.00 | 0.00 | 9.18 |
| 3329 | 0.0042 | 0.00 | 0.00 | 9.17 |
| 3330 | 0.0048 | 0.00 | 0.00 | 9.17 |
| 3331 | 0.0029 | 0.00 | 0.00 | 9.17 |
| 3332 | 0.0029 | 0.00 | 0.00 | 9.16 |
| 3333 | 0.0022 | 0.00 | 0.00 | 9.16 |
| 3334 | 0.0026 | 0.00 | 0.00 | 9.16 |
| 3335 | 0.0026 | 0.00 | 0.00 | 9.15 |
| 3336 | 0.0038 | 0.00 | 0.00 | 9.15 |
| 3337 | 0.0054 | 0.00 | 0.00 | 9.15 |
| 3338 | 0.0048 | 0.00 | 0.00 | 9.14 |
| 3339 | 0.0029 | 0.00 | 0.00 | 9.14 |
| 3340 | 0.0032 | 0.00 | 0.00 | 9.13 |
| 3341 | 0.0045 | 0.00 | 0.00 | 9.13 |
| 3342 | 0.0035 | 0.00 | 0.00 | 9.13 |
| 3343 | 0.0035 | 0.00 | 0.00 | 9.12 |
| 3344 | 0.0035 | 0.00 | 0.00 | 9.12 |

| | | | | |
|---|---|---|---|---|
| 3345 | 0.0045 | 0.00 | 0.00 | 9.12 |
| 3346 | 0.0026 | 0.00 | 0.00 | 9.11 |
| 3347 | 0.0048 | 0.00 | 0.00 | 9.11 |
| 3348 | 0.0016 | 0.00 | 0.00 | 9.10 |
| 3349 | 0.0051 | 0.00 | 0.00 | 9.10 |
| 3350 | 0.0035 | 0.00 | 0.00 | 9.10 |
| 3351 | 0.0035 | 0.00 | 0.00 | 9.09 |
| 3352 | 0.0026 | 0.00 | 0.00 | 9.09 |
| 3353 | 0.0032 | 0.00 | 0.00 | 9.09 |
| 3354 | 0.0038 | 0.00 | 0.00 | 9.09 |
| 3355 | 0.0077 | 0.00 | 0.00 | 9.08 |
| 3356 | 0.0032 | 0.00 | 0.00 | 9.07 |
| 3357 | 0.0045 | 0.00 | 0.00 | 9.07 |
| 3358 | 0.0038 | 0.00 | 0.00 | 9.07 |
| 3359 | 0.0026 | 0.00 | 0.00 | 9.06 |
| 3360 | 0.0042 | 0.00 | 0.00 | 9.06 |
| 3361 | 0.0032 | 0.00 | 0.00 | 9.06 |
| 3362 | 0.0026 | 0.00 | 0.00 | 9.05 |
| 3363 | 0.0016 | 0.00 | 0.00 | 9.05 |
| 3364 | 0.0035 | 0.00 | 0.00 | 9.05 |
| 3365 | 0.0035 | 0.00 | 0.00 | 9.04 |
| 3366 | 0.0029 | 0.00 | 0.00 | 9.04 |
| 3367 | 0.0038 | 0.00 | 0.00 | 9.04 |
| 3368 | 0.0029 | 0.00 | 0.00 | 9.03 |
| 3369 | 0.0029 | 0.00 | 0.00 | 9.03 |
| 3370 | 0.0032 | 0.00 | 0.00 | 9.03 |
| 3371 | 0.0026 | 0.00 | 0.00 | 9.03 |
| 3372 | 0.0038 | 0.00 | 0.00 | 9.02 |
| 3373 | 0.0048 | 0.00 | 0.00 | 9.02 |
| 3374 | 0.0051 | 0.00 | 0.00 | 9.01 |
| 3375 | 0.0035 | 0.00 | 0.00 | 9.01 |
| 3376 | 0.0048 | 0.00 | 0.00 | 9.01 |
| 3377 | 0.0038 | 0.00 | 0.00 | 9.00 |
| 3378 | 0.0054 | 0.00 | 0.00 | 9.00 |
| 3379 | 0.0019 | 0.00 | 0.00 | 8.99 |
| 3380 | 0.0042 | 0.00 | 0.00 | 8.99 |
| 3381 | 0.0022 | 0.00 | 0.00 | 8.99 |
| 3382 | 0.0042 | 0.00 | 0.00 | 8.98 |
| 3383 | 0.0038 | 0.00 | 0.00 | 8.98 |
| 3384 | 0.0026 | 0.00 | 0.00 | 8.98 |
| 3385 | 0.0022 | 0.00 | 0.00 | 8.97 |
| 3386 | 0.0042 | 0.00 | 0.00 | 8.97 |
| 3387 | 0.0032 | 0.00 | 0.00 | 8.97 |
| 3388 | 0.0035 | 0.00 | 0.00 | 8.96 |

| | | | | |
|---|---|---|---|---|
| 3389 | 0.0035 | 0.00 | 0.00 | 8.96 |
| 3390 | 0.0032 | 0.00 | 0.00 | 8.96 |
| 3391 | 0.0054 | 0.00 | 0.00 | 8.95 |
| 3392 | 0.0029 | 0.00 | 0.00 | 8.95 |
| 3393 | 0.0035 | 0.00 | 0.00 | 8.94 |
| 3394 | 0.0029 | 0.00 | 0.00 | 8.94 |
| 3395 | 0.0016 | 0.00 | 0.00 | 8.94 |
| 3396 | 0.0032 | 0.00 | 0.00 | 8.94 |
| 3397 | 0.0029 | 0.00 | 0.00 | 8.93 |
| 3398 | 0.0026 | 0.00 | 0.00 | 8.93 |
| 3399 | 0.0013 | 0.00 | 0.00 | 8.93 |
| 3400 | 0.0038 | 0.00 | 0.00 | 8.93 |
| 3401 | 0.0022 | 0.00 | 0.00 | 8.92 |
| 3402 | 0.0051 | 0.00 | 0.00 | 8.92 |
| 3403 | 0.0061 | 0.00 | 0.00 | 8.92 |
| 3404 | 0.0016 | 0.00 | 0.00 | 8.91 |
| 3405 | 0.0038 | 0.00 | 0.00 | 8.91 |
| 3406 | 0.0035 | 0.00 | 0.00 | 8.90 |
| 3407 | 0.0048 | 0.00 | 0.00 | 8.90 |
| 3408 | 0.0035 | 0.00 | 0.00 | 8.90 |
| 3409 | 0.0035 | 0.00 | 0.00 | 8.89 |
| 3410 | 0.0026 | 0.00 | 0.00 | 8.89 |
| 3411 | 0.0019 | 0.00 | 0.00 | 8.89 |
| 3412 | 0.0026 | 0.00 | 0.00 | 8.88 |
| 3413 | 0.0058 | 0.00 | 0.00 | 8.88 |
| 3414 | 0.0016 | 0.00 | 0.00 | 8.88 |
| 3415 | 0.0048 | 0.00 | 0.00 | 8.87 |
| 3416 | 0.0019 | 0.00 | 0.00 | 8.87 |
| 3417 | 0.0019 | 0.00 | 0.00 | 8.87 |
| 3418 | 0.0045 | 0.00 | 0.00 | 8.87 |
| 3419 | 0.0048 | 0.00 | 0.00 | 8.86 |
| 3420 | 0.0032 | 0.00 | 0.00 | 8.86 |
| 3421 | 0.0038 | 0.00 | 0.00 | 8.85 |
| 3422 | 0.0032 | 0.00 | 0.00 | 8.85 |
| 3423 | 0.0026 | 0.00 | 0.00 | 8.85 |
| 3424 | 0.0035 | 0.00 | 0.00 | 8.84 |
| 3425 | 0.0029 | 0.00 | 0.00 | 8.84 |
| 3426 | 0.0026 | 0.00 | 0.00 | 8.84 |
| 3427 | 0.0042 | 0.00 | 0.00 | 8.83 |
| 3428 | 0.0045 | 0.00 | 0.00 | 8.83 |
| 3429 | 0.0054 | 0.00 | 0.00 | 8.83 |
| 3430 | 0.0042 | 0.00 | 0.00 | 8.82 |
| 3431 | 0.0032 | 0.00 | 0.00 | 8.82 |
| 3432 | 0.0038 | 0.00 | 0.00 | 8.81 |

| | | | | |
|---|---|---|---|---|
| 3433 | 0.0042 | 0.00 | 0.00 | 8.81 |
| 3434 | 0.0035 | 0.00 | 0.00 | 8.80 |
| 3435 | 0.0026 | 0.00 | 0.00 | 8.80 |
| 3436 | 0.0032 | 0.00 | 0.00 | 8.80 |
| 3437 | 0.0019 | 0.00 | 0.00 | 8.80 |
| 3438 | 0.0026 | 0.00 | 0.00 | 8.79 |
| 3439 | 0.0038 | 0.00 | 0.00 | 8.79 |
| 3440 | 0.0038 | 0.00 | 0.00 | 8.79 |
| 3441 | 0.0016 | 0.00 | 0.00 | 8.78 |
| 3442 | 0.0032 | 0.00 | 0.00 | 8.78 |
| 3443 | 0.0032 | 0.00 | 0.00 | 8.78 |
| 3444 | 0.0042 | 0.00 | 0.00 | 8.78 |
| 3445 | 0.0029 | 0.00 | 0.00 | 8.77 |
| 3446 | 0.0016 | 0.00 | 0.00 | 8.77 |
| 3447 | 0.0026 | 0.00 | 0.00 | 8.77 |
| 3448 | 0.0048 | 0.00 | 0.00 | 8.76 |
| 3449 | 0.0032 | 0.00 | 0.00 | 8.76 |
| 3450 | 0.0048 | 0.00 | 0.00 | 8.76 |
| 3451 | 0.0029 | 0.00 | 0.00 | 8.75 |
| 3452 | 0.0045 | 0.00 | 0.00 | 8.75 |
| 3453 | 0.0026 | 0.00 | 0.00 | 8.74 |
| 3454 | 0.0032 | 0.00 | 0.00 | 8.74 |
| 3455 | 0.0045 | 0.00 | 0.00 | 8.74 |
| 3456 | 0.0022 | 0.00 | 0.00 | 8.73 |
| 3457 | 0.0013 | 0.00 | 0.00 | 8.73 |
| 3458 | 0.0016 | 0.00 | 0.00 | 8.73 |
| 3459 | 0.0035 | 0.00 | 0.00 | 8.73 |
| 3460 | 0.0051 | 0.00 | 0.00 | 8.72 |
| 3461 | 0.0035 | 0.00 | 0.00 | 8.72 |
| 3462 | 0.0035 | 0.00 | 0.00 | 8.72 |
| 3463 | 0.0042 | 0.00 | 0.00 | 8.71 |
| 3464 | 0.0013 | 0.00 | 0.00 | 8.71 |
| 3465 | 0.0022 | 0.00 | 0.00 | 8.71 |
| 3466 | 0.0045 | 0.00 | 0.00 | 8.70 |
| 3467 | 0.0022 | 0.00 | 0.00 | 8.70 |
| 3468 | 0.0035 | 0.00 | 0.00 | 8.70 |
| 3469 | 0.0019 | 0.00 | 0.00 | 8.69 |
| 3470 | 0.0038 | 0.00 | 0.00 | 8.69 |
| 3471 | 0.0026 | 0.00 | 0.00 | 8.69 |
| 3472 | 0.0038 | 0.00 | 0.00 | 8.69 |
| 3473 | 0.0035 | 0.00 | 0.00 | 8.68 |
| 3474 | 0.0032 | 0.00 | 0.00 | 8.68 |
| 3475 | 0.0032 | 0.00 | 0.00 | 8.68 |
| 3476 | 0.0026 | 0.00 | 0.00 | 8.67 |

| | | | | |
|---|---|---|---|---|
| 3477 | 0.0026 | 0.00 | 0.00 | 8.67 |
| 3478 | 0.0042 | 0.00 | 0.00 | 8.67 |
| 3479 | 0.0032 | 0.00 | 0.00 | 8.66 |
| 3480 | 0.0022 | 0.00 | 0.00 | 8.66 |
| 3481 | 0.0022 | 0.00 | 0.00 | 8.66 |
| 3482 | 0.0026 | 0.00 | 0.00 | 8.66 |
| 3483 | 0.0042 | 0.00 | 0.00 | 8.65 |
| 3484 | 0.0026 | 0.00 | 0.00 | 8.65 |
| 3485 | 0.0026 | 0.00 | 0.00 | 8.65 |
| 3486 | 0.0016 | 0.00 | 0.00 | 8.64 |
| 3487 | 0.0022 | 0.00 | 0.00 | 8.64 |
| 3488 | 0.0035 | 0.00 | 0.00 | 8.64 |
| 3489 | 0.0032 | 0.00 | 0.00 | 8.64 |
| 3490 | 0.0042 | 0.00 | 0.00 | 8.63 |
| 3491 | 0.0029 | 0.00 | 0.00 | 8.63 |
| 3492 | 0.0035 | 0.00 | 0.00 | 8.63 |
| 3493 | 0.0022 | 0.00 | 0.00 | 8.62 |
| 3494 | 0.0029 | 0.00 | 0.00 | 8.62 |
| 3495 | 0.0038 | 0.00 | 0.00 | 8.62 |
| 3496 | 0.0038 | 0.00 | 0.00 | 8.61 |
| 3497 | 0.0016 | 0.00 | 0.00 | 8.61 |
| 3498 | 0.0016 | 0.00 | 0.00 | 8.61 |
| 3499 | 0.0013 | 0.00 | 0.00 | 8.61 |
| 3500 | 0.0042 | 0.00 | 0.00 | 8.61 |
| 3501 | 0.0035 | 0.00 | 0.00 | 8.60 |
| 3502 | 0.0035 | 0.00 | 0.00 | 8.60 |
| 3503 | 0.0026 | 0.00 | 0.00 | 8.59 |
| 3504 | 0.0038 | 0.00 | 0.00 | 8.59 |
| 3505 | 0.0042 | 0.00 | 0.00 | 8.59 |
| 3506 | 0.0032 | 0.00 | 0.00 | 8.58 |
| 3507 | 0.0045 | 0.00 | 0.00 | 8.58 |
| 3508 | 0.0035 | 0.00 | 0.00 | 8.58 |
| 3509 | 0.0022 | 0.00 | 0.00 | 8.57 |
| 3510 | 0.0013 | 0.00 | 0.00 | 8.57 |
| 3511 | 0.0042 | 0.00 | 0.00 | 8.57 |
| 3512 | 0.0019 | 0.00 | 0.00 | 8.56 |
| 3513 | 0.0038 | 0.00 | 0.00 | 8.56 |
| 3514 | 0.0029 | 0.00 | 0.00 | 8.56 |
| 3515 | 0.0038 | 0.00 | 0.00 | 8.56 |
| 3516 | 0.0029 | 0.00 | 0.00 | 8.55 |
| 3517 | 0.0045 | 0.00 | 0.00 | 8.55 |
| 3518 | 0.0029 | 0.00 | 0.00 | 8.54 |
| 3519 | 0.0071 | 0.00 | 0.00 | 8.54 |
| 3520 | 0.0054 | 0.00 | 0.00 | 8.53 |

| | | | | |
|---|---|---|---|---|
| 3521 | 0.0032 | 0.00 | 0.00 | 8.53 |
| 3522 | 0.0042 | 0.00 | 0.00 | 8.53 |
| 3523 | 0.0032 | 0.00 | 0.00 | 8.52 |
| 3524 | 0.0022 | 0.00 | 0.00 | 8.52 |
| 3525 | 0.0035 | 0.00 | 0.00 | 8.52 |
| 3526 | 0.0022 | 0.00 | 0.00 | 8.51 |
| 3527 | 0.0032 | 0.00 | 0.00 | 8.51 |
| 3528 | 0.0026 | 0.00 | 0.00 | 8.51 |
| 3529 | 0.0032 | 0.00 | 0.00 | 8.50 |
| 3530 | 0.0032 | 0.00 | 0.00 | 8.50 |
| 3531 | 0.0022 | 0.00 | 0.00 | 8.50 |
| 3532 | 0.0022 | 0.00 | 0.00 | 8.50 |
| 3533 | 0.0022 | 0.00 | 0.00 | 8.49 |
| 3534 | 0.0029 | 0.00 | 0.00 | 8.49 |
| 3535 | 0.0032 | 0.00 | 0.00 | 8.49 |
| 3536 | 0.0026 | 0.00 | 0.00 | 8.49 |
| 3537 | 0.0026 | 0.00 | 0.00 | 8.48 |
| 3538 | 0.0022 | 0.00 | 0.00 | 8.48 |
| 3539 | 0.0032 | 0.00 | 0.00 | 8.48 |
| 3540 | 0.0038 | 0.00 | 0.00 | 8.48 |
| 3541 | 0.0035 | 0.00 | 0.00 | 8.47 |
| 3542 | 0.0042 | 0.00 | 0.00 | 8.47 |
| 3543 | 0.0054 | 0.00 | 0.00 | 8.46 |
| 3544 | 0.0035 | 0.00 | 0.00 | 8.46 |
| 3545 | 0.0016 | 0.00 | 0.00 | 8.45 |
| 3546 | 0.0032 | 0.00 | 0.00 | 8.45 |
| 3547 | 0.0032 | 0.00 | 0.00 | 8.45 |
| 3548 | 0.0003 | 0.00 | 0.00 | 8.45 |
| 3549 | 0.0016 | 0.00 | 0.00 | 8.45 |
| 3550 | 0.0045 | 0.00 | 0.00 | 8.44 |
| 3551 | 0.0026 | 0.00 | 0.00 | 8.44 |
| 3552 | 0.0010 | 0.00 | 0.00 | 8.44 |
| 3553 | 0.0022 | 0.00 | 0.00 | 8.44 |
| 3554 | 0.0022 | 0.00 | 0.00 | 8.43 |
| 3555 | 0.0032 | 0.00 | 0.00 | 8.43 |
| 3556 | 0.0026 | 0.00 | 0.00 | 8.43 |
| 3557 | 0.0048 | 0.00 | 0.00 | 8.43 |
| 3558 | 0.0026 | 0.00 | 0.00 | 8.42 |
| 3559 | 0.0035 | 0.00 | 0.00 | 8.42 |
| 3560 | 0.0035 | 0.00 | 0.00 | 8.42 |
| 3561 | 0.0029 | 0.00 | 0.00 | 8.41 |
| 3562 | 0.0051 | 0.00 | 0.00 | 8.41 |
| 3563 | 0.0019 | 0.00 | 0.00 | 8.40 |
| 3564 | 0.0038 | 0.00 | 0.00 | 8.40 |

| | | | | |
|---|---|---|---|---|
| 3565 | 0.0016 | 0.00 | 0.00 | 8.40 |
| 3566 | 0.0032 | 0.00 | 0.00 | 8.40 |
| 3567 | 0.0026 | 0.00 | 0.00 | 8.39 |
| 3568 | 0.0048 | 0.00 | 0.00 | 8.39 |
| 3569 | 0.0019 | 0.00 | 0.00 | 8.39 |
| 3570 | 0.0032 | 0.00 | 0.00 | 8.38 |
| 3571 | 0.0035 | 0.00 | 0.00 | 8.38 |
| 3572 | 0.0038 | 0.00 | 0.00 | 8.38 |
| 3573 | 0.0048 | 0.00 | 0.00 | 8.37 |
| 3574 | 0.0029 | 0.00 | 0.00 | 8.37 |
| 3575 | 0.0029 | 0.00 | 0.00 | 8.37 |
| 3576 | 0.0019 | 0.00 | 0.00 | 8.36 |
| 3577 | 0.0045 | 0.00 | 0.00 | 8.36 |
| 3578 | 0.0029 | 0.00 | 0.00 | 8.36 |
| 3579 | 0.0035 | 0.00 | 0.00 | 8.35 |
| 3580 | 0.0010 | 0.00 | 0.00 | 8.35 |
| 3581 | 0.0038 | 0.00 | 0.00 | 8.35 |
| 3582 | 0.0042 | 0.00 | 0.00 | 8.35 |
| 3583 | 0.0042 | 0.00 | 0.00 | 8.34 |
| 3584 | 0.0032 | 0.00 | 0.00 | 8.34 |
| 3585 | 0.0026 | 0.00 | 0.00 | 8.33 |
| 3586 | 0.0045 | 0.00 | 0.00 | 8.33 |
| 3587 | 0.0022 | 0.00 | 0.00 | 8.33 |
| 3588 | 0.0029 | 0.00 | 0.00 | 8.32 |
| 3589 | 0.0022 | 0.00 | 0.00 | 8.32 |
| 3590 | 0.0032 | 0.00 | 0.00 | 8.32 |
| 3591 | 0.0019 | 0.00 | 0.00 | 8.32 |
| 3592 | 0.0022 | 0.00 | 0.00 | 8.31 |
| 3593 | 0.0042 | 0.00 | 0.00 | 8.31 |
| 3594 | 0.0038 | 0.00 | 0.00 | 8.31 |
| 3595 | 0.0026 | 0.00 | 0.00 | 8.30 |
| 3596 | 0.0022 | 0.00 | 0.00 | 8.30 |
| 3597 | 0.0019 | 0.00 | 0.00 | 8.30 |
| 3598 | 0.0019 | 0.00 | 0.00 | 8.30 |
| 3599 | 0.0026 | 0.00 | 0.00 | 8.30 |
| 3600 | 0.0042 | 0.00 | 0.00 | 8.29 |
| 3601 | 0.0032 | 0.00 | 0.00 | 8.29 |
| 3602 | 0.0013 | 0.00 | 0.00 | 8.29 |
| 3603 | 0.0026 | 0.00 | 0.00 | 8.28 |
| 3604 | 0.0026 | 0.00 | 0.00 | 8.28 |
| 3605 | 0.0016 | 0.00 | 0.00 | 8.28 |
| 3606 | 0.0042 | 0.00 | 0.00 | 8.28 |
| 3607 | 0.0038 | 0.00 | 0.00 | 8.27 |
| 3608 | 0.0026 | 0.00 | 0.00 | 8.27 |

| | | | | |
|---|---|---|---|---|
| 3609 | 0.0045 | 0.00 | 0.00 | 8.27 |
| 3610 | 0.0029 | 0.00 | 0.00 | 8.26 |
| 3611 | 0.0019 | 0.00 | 0.00 | 8.26 |
| 3612 | 0.0038 | 0.00 | 0.00 | 8.26 |
| 3613 | 0.0032 | 0.00 | 0.00 | 8.25 |
| 3614 | 0.0022 | 0.00 | 0.00 | 8.25 |
| 3615 | 0.0045 | 0.00 | 0.00 | 8.25 |
| 3616 | 0.0032 | 0.00 | 0.00 | 8.24 |
| 3617 | 0.0022 | 0.00 | 0.00 | 8.24 |
| 3618 | 0.0013 | 0.00 | 0.00 | 8.24 |
| 3619 | 0.0045 | 0.00 | 0.00 | 8.24 |
| 3620 | 0.0019 | 0.00 | 0.00 | 8.23 |
| 3621 | 0.0048 | 0.00 | 0.00 | 8.23 |
| 3622 | 0.0016 | 0.00 | 0.00 | 8.23 |
| 3623 | 0.0048 | 0.00 | 0.00 | 8.22 |
| 3624 | 0.0013 | 0.00 | 0.00 | 8.22 |
| 3625 | 0.0022 | 0.00 | 0.00 | 8.22 |
| 3626 | 0.0045 | 0.00 | 0.00 | 8.22 |
| 3627 | 0.0029 | 0.00 | 0.00 | 8.21 |
| 3628 | 0.0029 | 0.00 | 0.00 | 8.21 |
| 3629 | 0.0029 | 0.00 | 0.00 | 8.21 |
| 3630 | 0.0038 | 0.00 | 0.00 | 8.20 |
| 3631 | 0.0054 | 0.00 | 0.00 | 8.20 |
| 3632 | 0.0042 | 0.00 | 0.00 | 8.19 |
| 3633 | 0.0038 | 0.00 | 0.00 | 8.19 |
| 3634 | 0.0048 | 0.00 | 0.00 | 8.19 |
| 3635 | 0.0067 | 0.00 | 0.00 | 8.18 |
| 3636 | 0.0038 | 0.00 | 0.00 | 8.17 |
| 3637 | 0.0029 | 0.00 | 0.00 | 8.17 |
| 3638 | 0.0026 | 0.00 | 0.00 | 8.17 |
| 3639 | 0.0026 | 0.00 | 0.00 | 8.16 |
| 3640 | 0.0035 | 0.00 | 0.00 | 8.16 |
| 3641 | 0.0019 | 0.00 | 0.00 | 8.16 |
| 3642 | 0.0029 | 0.00 | 0.00 | 8.16 |
| 3643 | 0.0016 | 0.00 | 0.00 | 8.15 |
| 3644 | 0.0029 | 0.00 | 0.00 | 8.15 |
| 3645 | 0.0019 | 0.00 | 0.00 | 8.15 |
| 3646 | 0.0010 | 0.00 | 0.00 | 8.15 |
| 3647 | 0.0010 | 0.00 | 0.00 | 8.15 |
| 3648 | 0.0029 | 0.00 | 0.00 | 8.15 |
| 3649 | 0.0032 | 0.00 | 0.00 | 8.14 |
| 3650 | 0.0016 | 0.00 | 0.00 | 8.14 |
| 3651 | 0.0032 | 0.00 | 0.00 | 8.14 |
| 3652 | 0.0029 | 0.00 | 0.00 | 8.13 |

| | | | | |
|---|---|---|---|---|
| 3653 | 0.0006 | 0.00 | 0.00 | 8.13 |
| 3654 | 0.0032 | 0.00 | 0.00 | 8.13 |
| 3655 | 0.0022 | 0.00 | 0.00 | 8.13 |
| 3656 | 0.0016 | 0.00 | 0.00 | 8.13 |
| 3657 | 0.0029 | 0.00 | 0.00 | 8.12 |
| 3658 | 0.0013 | 0.00 | 0.00 | 8.12 |
| 3659 | 0.0042 | 0.00 | 0.00 | 8.12 |
| 3660 | 0.0032 | 0.00 | 0.00 | 8.12 |
| 3661 | 0.0026 | 0.00 | 0.00 | 8.11 |
| 3662 | 0.0022 | 0.00 | 0.00 | 8.11 |
| 3663 | 0.0042 | 0.00 | 0.00 | 8.11 |
| 3664 | 0.0016 | 0.00 | 0.00 | 8.10 |
| 3665 | 0.0019 | 0.00 | 0.00 | 8.10 |
| 3666 | 0.0029 | 0.00 | 0.00 | 8.10 |
| 3667 | 0.0038 | 0.00 | 0.00 | 8.10 |
| 3668 | 0.0029 | 0.00 | 0.00 | 8.09 |
| 3669 | 0.0038 | 0.00 | 0.00 | 8.09 |
| 3670 | 0.0019 | 0.00 | 0.00 | 8.09 |
| 3671 | 0.0035 | 0.00 | 0.00 | 8.08 |
| 3672 | 0.0058 | 0.00 | 0.00 | 8.08 |
| 3673 | 0.0013 | 0.00 | 0.00 | 8.08 |
| 3674 | 0.0026 | 0.00 | 0.00 | 8.07 |
| 3675 | 0.0035 | 0.00 | 0.00 | 8.07 |
| 3676 | 0.0029 | 0.00 | 0.00 | 8.07 |
| 3677 | 0.0051 | 0.00 | 0.00 | 8.06 |
| 3678 | 0.0029 | 0.00 | 0.00 | 8.06 |
| 3679 | 0.0026 | 0.00 | 0.00 | 8.06 |
| 3680 | 0.0016 | 0.00 | 0.00 | 8.05 |
| 3681 | 0.0016 | 0.00 | 0.00 | 8.05 |
| 3682 | 0.0022 | 0.00 | 0.00 | 8.05 |
| 3683 | 0.0016 | 0.00 | 0.00 | 8.05 |
| 3684 | 0.0038 | 0.00 | 0.00 | 8.05 |
| 3685 | 0.0029 | 0.00 | 0.00 | 8.04 |
| 3686 | 0.0010 | 0.00 | 0.00 | 8.04 |
| 3687 | 0.0035 | 0.00 | 0.00 | 8.04 |
| 3688 | 0.0010 | 0.00 | 0.00 | 8.04 |
| 3689 | 0.0029 | 0.00 | 0.00 | 8.03 |
| 3690 | 0.0022 | 0.00 | 0.00 | 8.03 |
| 3691 | 0.0022 | 0.00 | 0.00 | 8.03 |
| 3692 | 0.0006 | 0.00 | 0.00 | 8.03 |
| 3693 | 0.0019 | 0.00 | 0.00 | 8.03 |
| 3694 | 0.0026 | 0.00 | 0.00 | 8.03 |
| 3695 | 0.0013 | 0.00 | 0.00 | 8.02 |
| 3696 | 0.0038 | 0.00 | 0.00 | 8.02 |

| | | | | |
|---|---|---|---|---|
| 3697 | 0.0026 | 0.00 | 0.00 | 8.02 |
| 3698 | 0.0022 | 0.00 | 0.00 | 8.01 |
| 3699 | 0.0022 | 0.00 | 0.00 | 8.01 |
| 3700 | 0.0019 | 0.00 | 0.00 | 8.01 |
| 3701 | 0.0029 | 0.00 | 0.00 | 8.01 |
| 3702 | 0.0026 | 0.00 | 0.00 | 8.01 |
| 3703 | 0.0019 | 0.00 | 0.00 | 8.00 |
| 3704 | 0.0029 | 0.00 | 0.00 | 8.00 |
| 3705 | 0.0035 | 0.00 | 0.00 | 8.00 |
| 3706 | 0.0054 | 0.00 | 0.00 | 7.99 |
| 3707 | 0.0032 | 0.00 | 0.00 | 7.99 |
| 3708 | 0.0022 | 0.00 | 0.00 | 7.99 |
| 3709 | 0.0032 | 0.00 | 0.00 | 7.98 |
| 3710 | 0.0032 | 0.00 | 0.00 | 7.98 |
| 3711 | 0.0022 | 0.00 | 0.00 | 7.98 |
| 3712 | 0.0042 | 0.00 | 0.00 | 7.98 |
| 3713 | 0.0032 | 0.00 | 0.00 | 7.97 |
| 3714 | 0.0051 | 0.00 | 0.00 | 7.97 |
| 3715 | 0.0029 | 0.00 | 0.00 | 7.96 |
| 3716 | 0.0064 | 0.00 | 0.00 | 7.96 |
| 3717 | 0.0019 | 0.00 | 0.00 | 7.95 |
| 3718 | 0.0029 | 0.00 | 0.00 | 7.95 |
| 3719 | 0.0022 | 0.00 | 0.00 | 7.95 |
| 3720 | 0.0022 | 0.00 | 0.00 | 7.95 |
| 3721 | 0.0032 | 0.00 | 0.00 | 7.94 |
| 3722 | 0.0013 | 0.00 | 0.00 | 7.94 |
| 3723 | 0.0029 | 0.00 | 0.00 | 7.94 |
| 3724 | 0.0013 | 0.00 | 0.00 | 7.94 |
| 3725 | 0.0029 | 0.00 | 0.00 | 7.94 |
| 3726 | 0.0032 | 0.00 | 0.00 | 7.93 |
| 3727 | 0.0026 | 0.00 | 0.00 | 7.93 |
| 3728 | 0.0016 | 0.00 | 0.00 | 7.93 |
| 3729 | 0.0016 | 0.00 | 0.00 | 7.93 |
| 3730 | 0.0026 | 0.00 | 0.00 | 7.92 |
| 3731 | 0.0029 | 0.00 | 0.00 | 7.92 |
| 3732 | 0.0029 | 0.00 | 0.00 | 7.92 |
| 3733 | 0.0013 | 0.00 | 0.00 | 7.92 |
| 3734 | 0.0019 | 0.00 | 0.00 | 7.91 |
| 3735 | 0.0032 | 0.00 | 0.00 | 7.91 |
| 3736 | 0.0016 | 0.00 | 0.00 | 7.91 |
| 3737 | 0.0022 | 0.00 | 0.00 | 7.91 |
| 3738 | 0.0016 | 0.00 | 0.00 | 7.90 |
| 3739 | 0.0045 | 0.00 | 0.00 | 7.90 |
| 3740 | 0.0026 | 0.00 | 0.00 | 7.90 |

| | | | | |
|---|---|---|---|---|
| 3741 | 0.0035 | 0.00 | 0.00 | 7.90 |
| 3742 | 0.0016 | 0.00 | 0.00 | 7.89 |
| 3743 | 0.0013 | 0.00 | 0.00 | 7.89 |
| 3744 | 0.0019 | 0.00 | 0.00 | 7.89 |
| 3745 | 0.0042 | 0.00 | 0.00 | 7.89 |
| 3746 | 0.0016 | 0.00 | 0.00 | 7.88 |
| 3747 | 0.0029 | 0.00 | 0.00 | 7.88 |
| 3748 | 0.0022 | 0.00 | 0.00 | 7.88 |
| 3749 | 0.0019 | 0.00 | 0.00 | 7.88 |
| 3750 | 0.0026 | 0.00 | 0.00 | 7.88 |
| 3751 | 0.0026 | 0.00 | 0.00 | 7.87 |
| 3752 | 0.0035 | 0.00 | 0.00 | 7.87 |
| 3753 | 0.0038 | 0.00 | 0.00 | 7.87 |
| 3754 | 0.0029 | 0.00 | 0.00 | 7.86 |
| 3755 | 0.0032 | 0.00 | 0.00 | 7.86 |
| 3756 | 0.0022 | 0.00 | 0.00 | 7.86 |
| 3757 | 0.0006 | 0.00 | 0.00 | 7.85 |
| 3758 | 0.0019 | 0.00 | 0.00 | 7.85 |
| 3759 | 0.0032 | 0.00 | 0.00 | 7.85 |
| 3760 | 0.0019 | 0.00 | 0.00 | 7.85 |
| 3761 | 0.0038 | 0.00 | 0.00 | 7.85 |
| 3762 | 0.0013 | 0.00 | 0.00 | 7.84 |
| 3763 | 0.0016 | 0.00 | 0.00 | 7.84 |
| 3764 | 0.0022 | 0.00 | 0.00 | 7.84 |
| 3765 | 0.0016 | 0.00 | 0.00 | 7.84 |
| 3766 | 0.0022 | 0.00 | 0.00 | 7.84 |
| 3767 | 0.0016 | 0.00 | 0.00 | 7.83 |
| 3768 | 0.0022 | 0.00 | 0.00 | 7.83 |
| 3769 | 0.0013 | 0.00 | 0.00 | 7.83 |
| 3770 | 0.0013 | 0.00 | 0.00 | 7.83 |
| 3771 | 0.0019 | 0.00 | 0.00 | 7.83 |
| 3772 | 0.0029 | 0.00 | 0.00 | 7.83 |
| 3773 | 0.0038 | 0.00 | 0.00 | 7.82 |
| 3774 | 0.0026 | 0.00 | 0.00 | 7.82 |
| 3775 | 0.0013 | 0.00 | 0.00 | 7.82 |
| 3776 | 0.0032 | 0.00 | 0.00 | 7.81 |
| 3777 | 0.0026 | 0.00 | 0.00 | 7.81 |
| 3778 | 0.0035 | 0.00 | 0.00 | 7.81 |
| 3779 | 0.0051 | 0.00 | 0.00 | 7.81 |
| 3780 | 0.0022 | 0.00 | 0.00 | 7.80 |
| 3781 | 0.0029 | 0.00 | 0.00 | 7.80 |
| 3782 | 0.0029 | 0.00 | 0.00 | 7.80 |
| 3783 | 0.0029 | 0.00 | 0.00 | 7.79 |
| 3784 | 0.0022 | 0.00 | 0.00 | 7.79 |

| | | | | |
|---|---|---|---|---|
| 3785 | 0.0016 | 0.00 | 0.00 | 7.79 |
| 3786 | 0.0035 | 0.00 | 0.00 | 7.79 |
| 3787 | 0.0026 | 0.00 | 0.00 | 7.78 |
| 3788 | 0.0010 | 0.00 | 0.00 | 7.78 |
| 3789 | 0.0010 | 0.00 | 0.00 | 7.78 |
| 3790 | 0.0010 | 0.00 | 0.00 | 7.78 |
| 3791 | 0.0051 | 0.00 | 0.00 | 7.78 |
| 3792 | 0.0029 | 0.00 | 0.00 | 7.77 |
| 3793 | 0.0045 | 0.00 | 0.00 | 7.77 |
| 3794 | 0.0035 | 0.00 | 0.00 | 7.76 |
| 3795 | 0.0022 | 0.00 | 0.00 | 7.76 |
| 3796 | 0.0029 | 0.00 | 0.00 | 7.76 |
| 3797 | 0.0022 | 0.00 | 0.00 | 7.76 |
| 3798 | 0.0032 | 0.00 | 0.00 | 7.75 |
| 3799 | 0.0019 | 0.00 | 0.00 | 7.75 |
| 3800 | 0.0045 | 0.00 | 0.00 | 7.75 |
| 3801 | 0.0006 | 0.00 | 0.00 | 7.74 |
| 3802 | 0.0022 | 0.00 | 0.00 | 7.74 |
| 3803 | 0.0022 | 0.00 | 0.00 | 7.74 |
| 3804 | 0.0013 | 0.00 | 0.00 | 7.74 |
| 3805 | 0.0035 | 0.00 | 0.00 | 7.74 |
| 3806 | 0.0029 | 0.00 | 0.00 | 7.73 |
| 3807 | 0.0019 | 0.00 | 0.00 | 7.73 |
| 3808 | 0.0026 | 0.00 | 0.00 | 7.73 |
| 3809 | 0.0006 | 0.00 | 0.00 | 7.73 |
| 3810 | 0.0013 | 0.00 | 0.00 | 7.73 |
| 3811 | 0.0035 | 0.00 | 0.00 | 7.72 |
| 3812 | 0.0022 | 0.00 | 0.00 | 7.72 |
| 3813 | 0.0016 | 0.00 | 0.00 | 7.72 |
| 3814 | 0.0029 | 0.00 | 0.00 | 7.72 |
| 3815 | 0.0038 | 0.00 | 0.00 | 7.71 |
| 3816 | 0.0029 | 0.00 | 0.00 | 7.71 |
| 3817 | 0.0013 | 0.00 | 0.00 | 7.71 |
| 3818 | 0.0029 | 0.00 | 0.00 | 7.71 |
| 3819 | 0.0022 | 0.00 | 0.00 | 7.70 |
| 3820 | 0.0010 | 0.00 | 0.00 | 7.70 |
| 3821 | 0.0022 | 0.00 | 0.00 | 7.70 |
| 3822 | 0.0010 | 0.00 | 0.00 | 7.70 |
| 3823 | 0.0016 | 0.00 | 0.00 | 7.70 |
| 3824 | 0.0022 | 0.00 | 0.00 | 7.70 |
| 3825 | 0.0022 | 0.00 | 0.00 | 7.69 |
| 3826 | 0.0029 | 0.00 | 0.00 | 7.69 |
| 3827 | 0.0032 | 0.00 | 0.00 | 7.69 |
| 3828 | 0.0026 | 0.00 | 0.00 | 7.68 |

| | | | | |
|---|---|---|---|---|
| 3829 | 0.0051 | 0.00 | 0.00 | 7.68 |
| 3830 | 0.0019 | 0.00 | 0.00 | 7.68 |
| 3831 | 0.0038 | 0.00 | 0.00 | 7.68 |
| 3832 | 0.0013 | 0.00 | 0.00 | 7.67 |
| 3833 | 0.0022 | 0.00 | 0.00 | 7.67 |
| 3834 | 0.0006 | 0.00 | 0.00 | 7.67 |
| 3835 | 0.0019 | 0.00 | 0.00 | 7.67 |
| 3836 | 0.0013 | 0.00 | 0.00 | 7.67 |
| 3837 | 0.0019 | 0.00 | 0.00 | 7.66 |
| 3838 | 0.0022 | 0.00 | 0.00 | 7.66 |
| 3839 | 0.0026 | 0.00 | 0.00 | 7.66 |
| 3840 | 0.0035 | 0.00 | 0.00 | 7.66 |
| 3841 | 0.0019 | 0.00 | 0.00 | 7.65 |
| 3842 | 0.0032 | 0.00 | 0.00 | 7.65 |
| 3843 | 0.0016 | 0.00 | 0.00 | 7.65 |
| 3844 | 0.0042 | 0.00 | 0.00 | 7.65 |
| 3845 | 0.0026 | 0.00 | 0.00 | 7.64 |
| 3846 | 0.0019 | 0.00 | 0.00 | 7.64 |
| 3847 | 0.0035 | 0.00 | 0.00 | 7.64 |
| 3848 | 0.0010 | 0.00 | 0.00 | 7.63 |
| 3849 | 0.0016 | 0.00 | 0.00 | 7.63 |
| 3850 | 0.0019 | 0.00 | 0.00 | 7.63 |
| 3851 | 0.0016 | 0.00 | 0.00 | 7.63 |
| 3852 | 0.0026 | 0.00 | 0.00 | 7.63 |
| 3853 | 0.0026 | 0.00 | 0.00 | 7.63 |
| 3854 | 0.0022 | 0.00 | 0.00 | 7.62 |
| 3855 | 0.0010 | 0.00 | 0.00 | 7.62 |
| 3856 | 0.0026 | 0.00 | 0.00 | 7.62 |
| 3857 | 0.0022 | 0.00 | 0.00 | 7.62 |
| 3858 | 0.0010 | 0.00 | 0.00 | 7.62 |
| 3859 | 0.0019 | 0.00 | 0.00 | 7.61 |
| 3860 | 0.0029 | 0.00 | 0.00 | 7.61 |
| 3861 | 0.0022 | 0.00 | 0.00 | 7.61 |
| 3862 | 0.0029 | 0.00 | 0.00 | 7.61 |
| 3863 | 0.0032 | 0.00 | 0.00 | 7.60 |
| 3864 | 0.0022 | 0.00 | 0.00 | 7.60 |
| 3865 | 0.0029 | 0.00 | 0.00 | 7.60 |
| 3866 | 0.0026 | 0.00 | 0.00 | 7.60 |
| 3867 | 0.0029 | 0.00 | 0.00 | 7.59 |
| 3868 | 0.0029 | 0.00 | 0.00 | 7.59 |
| 3869 | 0.0016 | 0.00 | 0.00 | 7.59 |
| 3870 | 0.0026 | 0.00 | 0.00 | 7.59 |
| 3871 | 0.0029 | 0.00 | 0.00 | 7.58 |
| 3872 | 0.0013 | 0.00 | 0.00 | 7.58 |

| | | | | |
|---|---|---|---|---|
| 3873 | 0.0029 | 0.00 | 0.00 | 7.58 |
| 3874 | 0.0013 | 0.00 | 0.00 | 7.58 |
| 3875 | 0.0016 | 0.00 | 0.00 | 7.58 |
| 3876 | 0.0029 | 0.00 | 0.00 | 7.57 |
| 3877 | 0.0016 | 0.00 | 0.00 | 7.57 |
| 3878 | 0.0029 | 0.00 | 0.00 | 7.57 |
| 3879 | 0.0022 | 0.00 | 0.00 | 7.57 |
| 3880 | 0.0035 | 0.00 | 0.00 | 7.56 |
| 3881 | 0.0019 | 0.00 | 0.00 | 7.56 |
| 3882 | 0.0026 | 0.00 | 0.00 | 7.56 |
| 3883 | 0.0016 | 0.00 | 0.00 | 7.56 |
| 3884 | 0.0029 | 0.00 | 0.00 | 7.55 |
| 3885 | 0.0022 | 0.00 | 0.00 | 7.55 |
| 3886 | 0.0022 | 0.00 | 0.00 | 7.55 |
| 3887 | 0.0013 | 0.00 | 0.00 | 7.55 |
| 3888 | 0.0029 | 0.00 | 0.00 | 7.55 |
| 3889 | 0.0029 | 0.00 | 0.00 | 7.54 |
| 3890 | 0.0013 | 0.00 | 0.00 | 7.54 |
| 3891 | 0.0029 | 0.00 | 0.00 | 7.54 |
| 3892 | 0.0019 | 0.00 | 0.00 | 7.54 |
| 3893 | 0.0042 | 0.00 | 0.00 | 7.53 |
| 3894 | 0.0019 | 0.00 | 0.00 | 7.53 |
| 3895 | 0.0022 | 0.00 | 0.00 | 7.53 |
| 3896 | 0.0019 | 0.00 | 0.00 | 7.53 |
| 3897 | 0.0010 | 0.00 | 0.00 | 7.52 |
| 3898 | 0.0016 | 0.00 | 0.00 | 7.52 |
| 3899 | 0.0026 | 0.00 | 0.00 | 7.52 |
| 3900 | 0.0016 | 0.00 | 0.00 | 7.52 |
| 3901 | 0.0016 | 0.00 | 0.00 | 7.52 |
| 3902 | 0.0026 | 0.00 | 0.00 | 7.52 |
| 3903 | 0.0022 | 0.00 | 0.00 | 7.51 |
| 3904 | 0.0038 | 0.00 | 0.00 | 7.51 |
| 3905 | 0.0016 | 0.00 | 0.00 | 7.51 |
| 3906 | 0.0029 | 0.00 | 0.00 | 7.51 |
| 3907 | 0.0016 | 0.00 | 0.00 | 7.50 |
| 3908 | 0.0016 | 0.00 | 0.00 | 7.50 |
| 3909 | 0.0035 | 0.00 | 0.00 | 7.50 |
| 3910 | 0.0013 | 0.00 | 0.00 | 7.50 |
| 3911 | 0.0029 | 0.00 | 0.00 | 7.49 |
| 3912 | 0.0042 | 0.00 | 0.00 | 7.49 |
| 3913 | 0.0022 | 0.00 | 0.00 | 7.49 |
| 3914 | 0.0029 | 0.00 | 0.00 | 7.48 |
| 3915 | 0.0026 | 0.00 | 0.00 | 7.48 |
| 3916 | 0.0019 | 0.00 | 0.00 | 7.48 |

| | | | | |
|---|---|---|---|---|
| 3917 | 0.0016 | 0.00 | 0.00 | 7.48 |
| 3918 | 0.0038 | 0.00 | 0.00 | 7.48 |
| 3919 | 0.0016 | 0.00 | 0.00 | 7.47 |
| 3920 | 0.0022 | 0.00 | 0.00 | 7.47 |
| 3921 | 0.0022 | 0.00 | 0.00 | 7.47 |
| 3922 | 0.0019 | 0.00 | 0.00 | 7.47 |
| 3923 | 0.0013 | 0.00 | 0.00 | 7.46 |
| 3924 | 0.0029 | 0.00 | 0.00 | 7.46 |
| 3925 | 0.0035 | 0.00 | 0.00 | 7.46 |
| 3926 | 0.0022 | 0.00 | 0.00 | 7.46 |
| 3927 | 0.0032 | 0.00 | 0.00 | 7.45 |
| 3928 | 0.0016 | 0.00 | 0.00 | 7.45 |
| 3929 | 0.0035 | 0.00 | 0.00 | 7.45 |
| 3930 | 0.0026 | 0.00 | 0.00 | 7.45 |
| 3931 | 0.0038 | 0.00 | 0.00 | 7.44 |
| 3932 | 0.0013 | 0.00 | 0.00 | 7.44 |
| 3933 | 0.0032 | 0.00 | 0.00 | 7.44 |
| 3934 | 0.0019 | 0.00 | 0.00 | 7.43 |
| 3935 | 0.0019 | 0.00 | 0.00 | 7.43 |
| 3936 | 0.0022 | 0.00 | 0.00 | 7.43 |
| 3937 | 0.0038 | 0.00 | 0.00 | 7.43 |
| 3938 | 0.0022 | 0.00 | 0.00 | 7.43 |
| 3939 | 0.0016 | 0.00 | 0.00 | 7.42 |
| 3940 | 0.0029 | 0.00 | 0.00 | 7.42 |
| 3941 | 0.0022 | 0.00 | 0.00 | 7.42 |
| 3942 | 0.0032 | 0.00 | 0.00 | 7.42 |
| 3943 | 0.0022 | 0.00 | 0.00 | 7.41 |
| 3944 | 0.0013 | 0.00 | 0.00 | 7.41 |
| 3945 | 0.0026 | 0.00 | 0.00 | 7.41 |
| 3946 | 0.0029 | 0.00 | 0.00 | 7.41 |
| 3947 | 0.0022 | 0.00 | 0.00 | 7.40 |
| 3948 | 0.0022 | 0.00 | 0.00 | 7.40 |
| 3949 | 0.0032 | 0.00 | 0.00 | 7.40 |
| 3950 | 0.0013 | 0.00 | 0.00 | 7.40 |
| 3951 | 0.0026 | 0.00 | 0.00 | 7.39 |
| 3952 | 0.0006 | 0.00 | 0.00 | 7.39 |
| 3953 | 0.0045 | 0.00 | 0.00 | 7.39 |
| 3954 | 0.0016 | 0.00 | 0.00 | 7.39 |
| 3955 | 0.0022 | 0.00 | 0.00 | 7.39 |
| 3956 | 0.0026 | 0.00 | 0.00 | 7.38 |
| 3957 | 0.0016 | 0.00 | 0.00 | 7.38 |
| 3958 | 0.0029 | 0.00 | 0.00 | 7.38 |
| 3959 | 0.0032 | 0.00 | 0.00 | 7.38 |
| 3960 | 0.0003 | 0.00 | 0.00 | 7.37 |

| | | | | |
|---|---|---|---|---|
| 3961 | 0.0022 | 0.00 | 0.00 | 7.37 |
| 3962 | 0.0029 | 0.00 | 0.00 | 7.37 |
| 3963 | 0.0029 | 0.00 | 0.00 | 7.37 |
| 3964 | 0.0019 | 0.00 | 0.00 | 7.36 |
| 3965 | 0.0032 | 0.00 | 0.00 | 7.36 |
| 3966 | 0.0016 | 0.00 | 0.00 | 7.36 |
| 3967 | 0.0010 | 0.00 | 0.00 | 7.36 |
| 3968 | 0.0019 | 0.00 | 0.00 | 7.36 |
| 3969 | 0.0016 | 0.00 | 0.00 | 7.36 |
| 3970 | 0.0013 | 0.00 | 0.00 | 7.35 |
| 3971 | 0.0010 | 0.00 | 0.00 | 7.35 |
| 3972 | 0.0013 | 0.00 | 0.00 | 7.35 |
| 3973 | 0.0013 | 0.00 | 0.00 | 7.35 |
| 3974 | 0.0038 | 0.00 | 0.00 | 7.35 |
| 3975 | 0.0016 | 0.00 | 0.00 | 7.34 |
| 3976 | 0.0022 | 0.00 | 0.00 | 7.34 |
| 3977 | 0.0026 | 0.00 | 0.00 | 7.34 |
| 3978 | 0.0029 | 0.00 | 0.00 | 7.34 |
| 3979 | 0.0029 | 0.00 | 0.00 | 7.34 |
| 3980 | 0.0035 | 0.00 | 0.00 | 7.33 |
| 3981 | 0.0026 | 0.00 | 0.00 | 7.33 |
| 3982 | 0.0035 | 0.00 | 0.00 | 7.33 |
| 3983 | 0.0019 | 0.00 | 0.00 | 7.32 |
| 3984 | 0.0022 | 0.00 | 0.00 | 7.32 |
| 3985 | 0.0029 | 0.00 | 0.00 | 7.32 |
| 3986 | 0.0022 | 0.00 | 0.00 | 7.32 |
| 3988 | 0.0029 | 0.00 | 0.00 | 7.31 |
| 3989 | 0.0019 | 0.00 | 0.00 | 7.31 |
| 3990 | 0.0022 | 0.00 | 0.00 | 7.31 |
| 3991 | 0.0006 | 0.00 | 0.00 | 7.31 |
| 3992 | 0.0016 | 0.00 | 0.00 | 7.31 |
| 3993 | 0.0016 | 0.00 | 0.00 | 7.30 |
| 3994 | 0.0019 | 0.00 | 0.00 | 7.30 |
| 3995 | 0.0019 | 0.00 | 0.00 | 7.30 |
| 3996 | 0.0013 | 0.00 | 0.00 | 7.30 |
| 3997 | 0.0022 | 0.00 | 0.00 | 7.30 |
| 3998 | 0.0022 | 0.00 | 0.00 | 7.30 |
| 3999 | 0.0042 | 0.00 | 0.00 | 7.29 |
| 4000 | 0.0029 | 0.00 | 0.00 | 7.29 |
| 4001 | 0.0026 | 0.00 | 0.00 | 7.29 |
| 4002 | 0.0032 | 0.00 | 0.00 | 7.28 |
| 4003 | 0.0013 | 0.00 | 0.00 | 7.28 |
| 4004 | 0.0010 | 0.00 | 0.00 | 7.28 |
| 4005 | 0.0029 | 0.00 | 0.00 | 7.28 |

| | | | | |
|---|---|---|---|---|
| 4006 | 0.0016 | 0.00 | 0.00 | 7.28 |
| 4007 | 0.0019 | 0.00 | 0.00 | 7.27 |
| 4008 | 0.0019 | 0.00 | 0.00 | 7.27 |
| 4009 | 0.0019 | 0.00 | 0.00 | 7.27 |
| 4010 | 0.0022 | 0.00 | 0.00 | 7.27 |
| 4011 | 0.0019 | 0.00 | 0.00 | 7.27 |
| 4012 | 0.0019 | 0.00 | 0.00 | 7.26 |
| 4013 | 0.0013 | 0.00 | 0.00 | 7.26 |
| 4014 | 0.0026 | 0.00 | 0.00 | 7.26 |
| 4015 | 0.0032 | 0.00 | 0.00 | 7.26 |
| 4016 | 0.0013 | 0.00 | 0.00 | 7.25 |
| 4017 | 0.0010 | 0.00 | 0.00 | 7.25 |
| 4018 | 0.0029 | 0.00 | 0.00 | 7.25 |
| 4019 | 0.0006 | 0.00 | 0.00 | 7.25 |
| 4020 | 0.0032 | 0.00 | 0.00 | 7.25 |
| 4021 | 0.0022 | 0.00 | 0.00 | 7.25 |
| 4022 | 0.0019 | 0.00 | 0.00 | 7.24 |
| 4023 | 0.0016 | 0.00 | 0.00 | 7.24 |
| 4024 | 0.0016 | 0.00 | 0.00 | 7.24 |
| 4025 | 0.0016 | 0.00 | 0.00 | 7.24 |
| 4026 | 0.0022 | 0.00 | 0.00 | 7.24 |
| 4027 | 0.0029 | 0.00 | 0.00 | 7.23 |
| 4028 | 0.0013 | 0.00 | 0.00 | 7.23 |
| 4029 | 0.0022 | 0.00 | 0.00 | 7.23 |
| 4030 | 0.0013 | 0.00 | 0.00 | 7.23 |
| 4031 | 0.0019 | 0.00 | 0.00 | 7.23 |
| 4032 | 0.0016 | 0.00 | 0.00 | 7.23 |
| 4033 | 0.0022 | 0.00 | 0.00 | 7.22 |
| 4034 | 0.0016 | 0.00 | 0.00 | 7.22 |
| 4035 | 0.0022 | 0.00 | 0.00 | 7.22 |
| 4036 | 0.0022 | 0.00 | 0.00 | 7.22 |
| 4037 | 0.0022 | 0.00 | 0.00 | 7.22 |
| 4038 | 0.0016 | 0.00 | 0.00 | 7.21 |
| 4039 | 0.0035 | 0.00 | 0.00 | 7.21 |
| 4040 | 0.0022 | 0.00 | 0.00 | 7.21 |
| 4041 | 0.0029 | 0.00 | 0.00 | 7.21 |
| 4042 | 0.0016 | 0.00 | 0.00 | 7.20 |
| 4043 | 0.0022 | 0.00 | 0.00 | 7.20 |
| 4044 | 0.0032 | 0.00 | 0.00 | 7.20 |
| 4045 | 0.0026 | 0.00 | 0.00 | 7.20 |
| 4046 | 0.0016 | 0.00 | 0.00 | 7.19 |
| 4047 | 0.0022 | 0.00 | 0.00 | 7.19 |
| 4048 | 0.0019 | 0.00 | 0.00 | 7.19 |
| 4049 | 0.0026 | 0.00 | 0.00 | 7.19 |

| | | | | |
|---|---|---|---|---|
| 4050 | 0.0019 | 0.00 | 0.00 | 7.18 |
| 4051 | 0.0019 | 0.00 | 0.00 | 7.18 |
| 4052 | 0.0022 | 0.00 | 0.00 | 7.18 |
| 4053 | 0.0019 | 0.00 | 0.00 | 7.18 |
| 4054 | 0.0022 | 0.00 | 0.00 | 7.18 |
| 4055 | 0.0032 | 0.00 | 0.00 | 7.17 |
| 4056 | 0.0035 | 0.00 | 0.00 | 7.17 |
| 4057 | 0.0032 | 0.00 | 0.00 | 7.17 |
| 4058 | 0.0026 | 0.00 | 0.00 | 7.16 |
| 4059 | 0.0016 | 0.00 | 0.00 | 7.16 |
| 4060 | 0.0022 | 0.00 | 0.00 | 7.16 |
| 4061 | 0.0022 | 0.00 | 0.00 | 7.16 |
| 4062 | 0.0026 | 0.00 | 0.00 | 7.16 |
| 4063 | 0.0038 | 0.00 | 0.00 | 7.15 |
| 4064 | 0.0019 | 0.00 | 0.00 | 7.15 |
| 4065 | 0.0029 | 0.00 | 0.00 | 7.15 |
| 4066 | 0.0016 | 0.00 | 0.00 | 7.14 |
| 4067 | 0.0032 | 0.00 | 0.00 | 7.14 |
| 4068 | 0.0032 | 0.00 | 0.00 | 7.14 |
| 4069 | 0.0026 | 0.00 | 0.00 | 7.14 |
| 4070 | 0.0035 | 0.00 | 0.00 | 7.13 |
| 4071 | 0.0019 | 0.00 | 0.00 | 7.13 |
| 4072 | 0.0038 | 0.00 | 0.00 | 7.13 |
| 4073 | 0.0032 | 0.00 | 0.00 | 7.12 |
| 4074 | 0.0026 | 0.00 | 0.00 | 7.12 |
| 4075 | 0.0022 | 0.00 | 0.00 | 7.12 |
| 4076 | 0.0016 | 0.00 | 0.00 | 7.12 |
| 4077 | 0.0026 | 0.00 | 0.00 | 7.12 |
| 4078 | 0.0038 | 0.00 | 0.00 | 7.11 |
| 4079 | 0.0013 | 0.00 | 0.00 | 7.11 |
| 4080 | 0.0016 | 0.00 | 0.00 | 7.11 |
| 4081 | 0.0016 | 0.00 | 0.00 | 7.11 |
| 4082 | 0.0026 | 0.00 | 0.00 | 7.10 |
| 4083 | 0.0022 | 0.00 | 0.00 | 7.10 |
| 4084 | 0.0026 | 0.00 | 0.00 | 7.10 |
| 4085 | 0.0022 | 0.00 | 0.00 | 7.10 |
| 4086 | 0.0038 | 0.00 | 0.00 | 7.09 |
| 4087 | 0.0022 | 0.00 | 0.00 | 7.09 |
| 4088 | 0.0029 | 0.00 | 0.00 | 7.09 |
| 4089 | 0.0026 | 0.00 | 0.00 | 7.09 |
| 4090 | 0.0003 | 0.00 | 0.00 | 7.08 |
| 4091 | 0.0022 | 0.00 | 0.00 | 7.08 |
| 4092 | 0.0016 | 0.00 | 0.00 | 7.08 |
| 4093 | 0.0019 | 0.00 | 0.00 | 7.08 |

| | | | | |
|---|---|---|---|---|
| 4094 | 0.0013 | 0.00 | 0.00 | 7.08 |
| 4095 | 0.0022 | 0.00 | 0.00 | 7.08 |
| 4096 | 0.0016 | 0.00 | 0.00 | 7.07 |
| 4097 | 0.0022 | 0.00 | 0.00 | 7.07 |
| 4098 | 0.0038 | 0.00 | 0.00 | 7.07 |
| 4099 | 0.0016 | 0.00 | 0.00 | 7.07 |
| 4100 | 0.0010 | 0.00 | 0.00 | 7.06 |
| 4101 | 0.0006 | 0.00 | 0.00 | 7.06 |
| 4102 | 0.0016 | 0.00 | 0.00 | 7.06 |
| 4103 | 0.0022 | 0.00 | 0.00 | 7.06 |
| 4104 | 0.0045 | 0.00 | 0.00 | 7.06 |
| 4105 | 0.0022 | 0.00 | 0.00 | 7.05 |
| 4106 | 0.0038 | 0.00 | 0.00 | 7.05 |
| 4107 | 0.0032 | 0.00 | 0.00 | 7.05 |
| 4108 | 0.0019 | 0.00 | 0.00 | 7.04 |
| 4109 | 0.0019 | 0.00 | 0.00 | 7.04 |
| 4110 | 0.0061 | 0.00 | 0.00 | 7.04 |
| 4111 | 0.0013 | 0.00 | 0.00 | 7.03 |
| 4112 | 0.0022 | 0.00 | 0.00 | 7.03 |
| 4113 | 0.0026 | 0.00 | 0.00 | 7.03 |
| 4114 | 0.0038 | 0.00 | 0.00 | 7.03 |
| 4115 | 0.0035 | 0.00 | 0.00 | 7.02 |
| 4116 | 0.0029 | 0.00 | 0.00 | 7.02 |
| 4117 | 0.0029 | 0.00 | 0.00 | 7.02 |
| 4118 | 0.0010 | 0.00 | 0.00 | 7.02 |
| 4119 | 0.0016 | 0.00 | 0.00 | 7.01 |
| 4120 | 0.0051 | 0.00 | 0.00 | 7.01 |
| 4121 | 0.0038 | 0.00 | 0.00 | 7.01 |
| 4122 | 0.0035 | 0.00 | 0.00 | 7.00 |
| 4123 | 0.0016 | 0.00 | 0.00 | 7.00 |
| 4124 | 0.0026 | 0.00 | 0.00 | 7.00 |
| 4125 | 0.0032 | 0.00 | 0.00 | 7.00 |
| 4126 | 0.0026 | 0.00 | 0.00 | 6.99 |
| 4127 | 0.0032 | 0.00 | 0.00 | 6.99 |
| 4128 | 0.0006 | 0.00 | 0.00 | 6.99 |
| 4129 | 0.0026 | 0.00 | 0.00 | 6.99 |
| 4130 | 0.0051 | 0.00 | 0.00 | 6.98 |
| 4131 | 0.0013 | 0.00 | 0.00 | 6.98 |
| 4132 | 0.0038 | 0.00 | 0.00 | 6.98 |
| 4133 | 0.0035 | 0.00 | 0.00 | 6.97 |
| 4134 | 0.0022 | 0.00 | 0.00 | 6.97 |
| 4135 | 0.0032 | 0.00 | 0.00 | 6.97 |
| 4136 | 0.0026 | 0.00 | 0.00 | 6.97 |
| 4137 | 0.0029 | 0.00 | 0.00 | 6.96 |

| | | | | |
|---|---|---|---|---|
| 4138 | 0.0003 | 0.00 | 0.00 | 6.96 |
| 4139 | 0.0022 | 0.00 | 0.00 | 6.96 |
| 4140 | 0.0019 | 0.00 | 0.00 | 6.96 |
| 4141 | 0.0035 | 0.00 | 0.00 | 6.96 |
| 4142 | 0.0010 | 0.00 | 0.00 | 6.95 |
| 4143 | 0.0032 | 0.00 | 0.00 | 6.95 |
| 4144 | 0.0029 | 0.00 | 0.00 | 6.95 |
| 4145 | 0.0010 | 0.00 | 0.00 | 6.94 |
| 4146 | 0.0019 | 0.00 | 0.00 | 6.94 |
| 4147 | 0.0029 | 0.00 | 0.00 | 6.94 |
| 4148 | 0.0019 | 0.00 | 0.00 | 6.94 |
| 4149 | 0.0022 | 0.00 | 0.00 | 6.94 |
| 4150 | 0.0019 | 0.00 | 0.00 | 6.93 |
| 4151 | 0.0010 | 0.00 | 0.00 | 6.93 |
| 4152 | 0.0022 | 0.00 | 0.00 | 6.93 |
| 4153 | 0.0022 | 0.00 | 0.00 | 6.93 |
| 4154 | 0.0010 | 0.00 | 0.00 | 6.93 |
| 4155 | 0.0022 | 0.00 | 0.00 | 6.93 |
| 4156 | 0.0016 | 0.00 | 0.00 | 6.92 |
| 4157 | 0.0016 | 0.00 | 0.00 | 6.92 |
| 4158 | 0.0013 | 0.00 | 0.00 | 6.92 |
| 4159 | 0.0048 | 0.00 | 0.00 | 6.92 |
| 4160 | 0.0019 | 0.00 | 0.00 | 6.91 |
| 4161 | 0.0022 | 0.00 | 0.00 | 6.91 |
| 4162 | 0.0045 | 0.00 | 0.00 | 6.91 |
| 4163 | 0.0022 | 0.00 | 0.00 | 6.91 |
| 4164 | 0.0026 | 0.00 | 0.00 | 6.90 |
| 4165 | 0.0029 | 0.00 | 0.00 | 6.90 |
| 4166 | 0.0029 | 0.00 | 0.00 | 6.90 |
| 4167 | 0.0013 | 0.00 | 0.00 | 6.90 |
| 4168 | 0.0019 | 0.00 | 0.00 | 6.89 |
| 4169 | 0.0026 | 0.00 | 0.00 | 6.89 |
| 4170 | 0.0016 | 0.00 | 0.00 | 6.89 |
| 4171 | 0.0029 | 0.00 | 0.00 | 6.89 |
| 4172 | 0.0026 | 0.00 | 0.00 | 6.89 |
| 4173 | 0.0022 | 0.00 | 0.00 | 6.88 |
| 4174 | 0.0019 | 0.00 | 0.00 | 6.88 |
| 4175 | 0.0042 | 0.00 | 0.00 | 6.88 |
| 4176 | 0.0019 | 0.00 | 0.00 | 6.87 |
| 4177 | 0.0016 | 0.00 | 0.00 | 6.87 |
| 4178 | 0.0022 | 0.00 | 0.00 | 6.87 |
| 4179 | 0.0026 | 0.00 | 0.00 | 6.87 |
| 4180 | 0.0019 | 0.00 | 0.00 | 6.87 |
| 4181 | 0.0019 | 0.00 | 0.00 | 6.86 |

| | | | | |
|---|---|---|---|---|
| 4182 | 0.0032 | 0.00 | 0.00 | 6.86 |
| 4183 | 0.0032 | 0.00 | 0.00 | 6.86 |
| 4184 | 0.0035 | 0.00 | 0.00 | 6.86 |
| 4185 | 0.0016 | 0.00 | 0.00 | 6.85 |
| 4186 | 0.0019 | 0.00 | 0.00 | 6.85 |
| 4187 | 0.0029 | 0.00 | 0.00 | 6.85 |
| 4188 | 0.0032 | 0.00 | 0.00 | 6.85 |
| 4189 | 0.0077 | 0.00 | 0.01 | 6.84 |
| 4190 | 0.0061 | 0.00 | 0.00 | 6.83 |
| 4191 | 0.0048 | 0.00 | 0.00 | 6.83 |
| 4192 | 0.0026 | 0.00 | 0.00 | 6.82 |
| 4193 | 0.0032 | 0.00 | 0.00 | 6.82 |
| 4194 | 0.0026 | 0.00 | 0.00 | 6.82 |
| 4195 | 0.0022 | 0.00 | 0.00 | 6.82 |
| 4196 | 0.0016 | 0.00 | 0.00 | 6.81 |
| 4197 | 0.0010 | 0.00 | 0.00 | 6.81 |
| 4198 | 0.0022 | 0.00 | 0.00 | 6.81 |
| 4199 | 0.0019 | 0.00 | 0.00 | 6.81 |
| 4200 | 0.0029 | 0.00 | 0.00 | 6.81 |
| 4201 | 0.0003 | 0.00 | 0.00 | 6.80 |
| 4202 | 0.0022 | 0.00 | 0.00 | 6.80 |
| 4203 | 0.0013 | 0.00 | 0.00 | 6.80 |
| 4204 | 0.0016 | 0.00 | 0.00 | 6.80 |
| 4205 | 0.0019 | 0.00 | 0.00 | 6.80 |
| 4206 | 0.0042 | 0.00 | 0.00 | 6.80 |
| 4207 | 0.0035 | 0.00 | 0.00 | 6.79 |
| 4208 | 0.0019 | 0.00 | 0.00 | 6.79 |
| 4209 | 0.0019 | 0.00 | 0.00 | 6.79 |
| 4210 | 0.0038 | 0.00 | 0.00 | 6.78 |
| 4211 | 0.0016 | 0.00 | 0.00 | 6.78 |
| 4212 | 0.0016 | 0.00 | 0.00 | 6.78 |
| 4213 | 0.0019 | 0.00 | 0.00 | 6.78 |
| 4214 | 0.0022 | 0.00 | 0.00 | 6.78 |
| 4215 | 0.0006 | 0.00 | 0.00 | 6.77 |
| 4216 | 0.0013 | 0.00 | 0.00 | 6.77 |
| 4217 | 0.0029 | 0.00 | 0.00 | 6.77 |
| 4218 | 0.0019 | 0.00 | 0.00 | 6.77 |
| 4219 | 0.0026 | 0.00 | 0.00 | 6.77 |
| 4220 | 0.0019 | 0.00 | 0.00 | 6.76 |
| 4221 | 0.0029 | 0.00 | 0.00 | 6.76 |
| 4222 | 0.0022 | 0.00 | 0.00 | 6.76 |
| 4223 | 0.0022 | 0.00 | 0.00 | 6.76 |
| 4224 | 0.0016 | 0.00 | 0.00 | 6.76 |
| 4225 | 0.0022 | 0.00 | 0.00 | 6.75 |

| | | | | |
|---|---|---|---|---|
| 4226 | 0.0026 | 0.00 | 0.00 | 6.75 |
| 4227 | 0.0022 | 0.00 | 0.00 | 6.75 |
| 4228 | 0.0019 | 0.00 | 0.00 | 6.75 |
| 4229 | 0.0013 | 0.00 | 0.00 | 6.74 |
| 4230 | 0.0032 | 0.00 | 0.00 | 6.74 |
| 4231 | 0.0019 | 0.00 | 0.00 | 6.74 |
| 4232 | 0.0016 | 0.00 | 0.00 | 6.74 |
| 4233 | 0.0010 | 0.00 | 0.00 | 6.74 |
| 4234 | 0.0013 | 0.00 | 0.00 | 6.74 |
| 4235 | 0.0022 | 0.00 | 0.00 | 6.73 |
| 4236 | 0.0016 | 0.00 | 0.00 | 6.73 |
| 4237 | 0.0019 | 0.00 | 0.00 | 6.73 |
| 4238 | 0.0022 | 0.00 | 0.00 | 6.73 |
| 4239 | 0.0019 | 0.00 | 0.00 | 6.73 |
| 4240 | 0.0022 | 0.00 | 0.00 | 6.72 |
| 4241 | 0.0026 | 0.00 | 0.00 | 6.72 |
| 4242 | 0.0010 | 0.00 | 0.00 | 6.72 |
| 4243 | 0.0022 | 0.00 | 0.00 | 6.72 |
| 4244 | 0.0016 | 0.00 | 0.00 | 6.72 |
| 4245 | 0.0026 | 0.00 | 0.00 | 6.71 |
| 4246 | 0.0029 | 0.00 | 0.00 | 6.71 |
| 4247 | 0.0032 | 0.00 | 0.00 | 6.71 |
| 4248 | 0.0022 | 0.00 | 0.00 | 6.71 |
| 4249 | 0.0032 | 0.00 | 0.00 | 6.70 |
| 4250 | 0.0029 | 0.00 | 0.00 | 6.70 |
| 4251 | 0.0026 | 0.00 | 0.00 | 6.70 |
| 4252 | 0.0016 | 0.00 | 0.00 | 6.70 |
| 4253 | 0.0026 | 0.00 | 0.00 | 6.69 |
| 4254 | 0.0010 | 0.00 | 0.00 | 6.69 |
| 4255 | 0.0019 | 0.00 | 0.00 | 6.69 |
| 4256 | 0.0016 | 0.00 | 0.00 | 6.69 |
| 4257 | 0.0019 | 0.00 | 0.00 | 6.69 |
| 4258 | 0.0016 | 0.00 | 0.00 | 6.68 |
| 4259 | 0.0013 | 0.00 | 0.00 | 6.68 |
| 4260 | 0.0016 | 0.00 | 0.00 | 6.68 |
| 4261 | 0.0029 | 0.00 | 0.00 | 6.68 |
| 4262 | 0.0010 | 0.00 | 0.00 | 6.68 |
| 4263 | 0.0006 | 0.00 | 0.00 | 6.68 |
| 4264 | 0.0003 | 0.00 | 0.00 | 6.68 |
| 4265 | 0.0019 | 0.00 | 0.00 | 6.68 |
| 4266 | 0.0013 | 0.00 | 0.00 | 6.67 |
| 4267 | 0.0026 | 0.00 | 0.00 | 6.67 |
| 4268 | 0.0010 | 0.00 | 0.00 | 6.67 |
| 4269 | 0.0019 | 0.00 | 0.00 | 6.67 |

| | | | | |
|---|---|---|---|---|
| 4270 | 0.0029 | 0.00 | 0.00 | 6.67 |
| 4271 | 0.0026 | 0.00 | 0.00 | 6.66 |
| 4272 | 0.0022 | 0.00 | 0.00 | 6.66 |
| 4273 | 0.0016 | 0.00 | 0.00 | 6.66 |
| 4274 | 0.0010 | 0.00 | 0.00 | 6.66 |
| 4275 | 0.0022 | 0.00 | 0.00 | 6.66 |
| 4276 | 0.0013 | 0.00 | 0.00 | 6.65 |
| 4277 | 0.0016 | 0.00 | 0.00 | 6.65 |
| 4278 | 0.0026 | 0.00 | 0.00 | 6.65 |
| 4279 | 0.0029 | 0.00 | 0.00 | 6.65 |
| 4280 | 0.0016 | 0.00 | 0.00 | 6.65 |
| 4281 | 0.0019 | 0.00 | 0.00 | 6.64 |
| 4282 | 0.0022 | 0.00 | 0.00 | 6.64 |
| 4283 | 0.0010 | 0.00 | 0.00 | 6.64 |
| 4284 | 0.0016 | 0.00 | 0.00 | 6.64 |
| 4285 | 0.0022 | 0.00 | 0.00 | 6.64 |
| 4286 | 0.0010 | 0.00 | 0.00 | 6.64 |
| 4287 | 0.0019 | 0.00 | 0.00 | 6.63 |
| 4288 | 0.0013 | 0.00 | 0.00 | 6.63 |
| 4289 | 0.0019 | 0.00 | 0.00 | 6.63 |
| 4290 | 0.0029 | 0.00 | 0.00 | 6.63 |
| 4291 | 0.0019 | 0.00 | 0.00 | 6.63 |
| 4292 | 0.0003 | 0.00 | 0.00 | 6.62 |
| 4293 | 0.0019 | 0.00 | 0.00 | 6.62 |
| 4294 | 0.0006 | 0.00 | 0.00 | 6.62 |
| 4295 | 0.0010 | 0.00 | 0.00 | 6.62 |
| 4296 | 0.0022 | 0.00 | 0.00 | 6.62 |
| 4297 | 0.0016 | 0.00 | 0.00 | 6.62 |
| 4298 | 0.0013 | 0.00 | 0.00 | 6.62 |
| 4299 | 0.0019 | 0.00 | 0.00 | 6.62 |
| 4300 | 0.0013 | 0.00 | 0.00 | 6.61 |
| 4301 | 0.0010 | 0.00 | 0.00 | 6.61 |
| 4302 | 0.0016 | 0.00 | 0.00 | 6.61 |
| 4303 | 0.0029 | 0.00 | 0.00 | 6.61 |
| 4304 | 0.0019 | 0.00 | 0.00 | 6.61 |
| 4305 | 0.0019 | 0.00 | 0.00 | 6.60 |
| 4306 | 0.0029 | 0.00 | 0.00 | 6.60 |
| 4307 | 0.0022 | 0.00 | 0.00 | 6.60 |
| 4308 | 0.0010 | 0.00 | 0.00 | 6.60 |
| 4309 | 0.0019 | 0.00 | 0.00 | 6.60 |
| 4310 | 0.0035 | 0.00 | 0.00 | 6.59 |
| 4311 | 0.0016 | 0.00 | 0.00 | 6.59 |
| 4312 | 0.0016 | 0.00 | 0.00 | 6.59 |
| 4313 | 0.0026 | 0.00 | 0.00 | 6.59 |

| | | | | |
|---|---|---|---|---|
| 4314 | 0.0006 | 0.00 | 0.00 | 6.59 |
| 4315 | 0.0006 | 0.00 | 0.00 | 6.58 |
| 4316 | 0.0022 | 0.00 | 0.00 | 6.58 |
| 4317 | 0.0022 | 0.00 | 0.00 | 6.58 |
| 4318 | 0.0026 | 0.00 | 0.00 | 6.58 |
| 4319 | 0.0016 | 0.00 | 0.00 | 6.58 |
| 4320 | 0.0029 | 0.00 | 0.00 | 6.58 |
| 4321 | 0.0019 | 0.00 | 0.00 | 6.57 |
| 4322 | 0.0006 | 0.00 | 0.00 | 6.57 |
| 4323 | 0.0022 | 0.00 | 0.00 | 6.57 |
| 4324 | 0.0013 | 0.00 | 0.00 | 6.57 |
| 4325 | 0.0019 | 0.00 | 0.00 | 6.57 |
| 4326 | 0.0016 | 0.00 | 0.00 | 6.56 |
| 4327 | 0.0019 | 0.00 | 0.00 | 6.56 |
| 4328 | 0.0010 | 0.00 | 0.00 | 6.56 |
| 4329 | 0.0010 | 0.00 | 0.00 | 6.56 |
| 4330 | 0.0022 | 0.00 | 0.00 | 6.56 |
| 4331 | 0.0019 | 0.00 | 0.00 | 6.56 |
| 4332 | 0.0016 | 0.00 | 0.00 | 6.56 |
| 4333 | 0.0022 | 0.00 | 0.00 | 6.55 |
| 4334 | 0.0013 | 0.00 | 0.00 | 6.55 |
| 4335 | 0.0022 | 0.00 | 0.00 | 6.55 |
| 4336 | 0.0029 | 0.00 | 0.00 | 6.55 |
| 4337 | 0.0016 | 0.00 | 0.00 | 6.54 |
| 4338 | 0.0016 | 0.00 | 0.00 | 6.54 |
| 4339 | 0.0006 | 0.00 | 0.00 | 6.54 |
| 4340 | 0.0019 | 0.00 | 0.00 | 6.54 |
| 4341 | 0.0022 | 0.00 | 0.00 | 6.54 |
| 4342 | 0.0029 | 0.00 | 0.00 | 6.54 |
| 4343 | 0.0010 | 0.00 | 0.00 | 6.53 |
| 4344 | 0.0016 | 0.00 | 0.00 | 6.53 |
| 4345 | 0.0019 | 0.00 | 0.00 | 6.53 |
| 4346 | 0.0003 | 0.00 | 0.00 | 6.53 |
| 4347 | 0.0026 | 0.00 | 0.00 | 6.53 |
| 4348 | 0.0022 | 0.00 | 0.00 | 6.53 |
| 4349 | 0.0026 | 0.00 | 0.00 | 6.52 |
| 4350 | 0.0019 | 0.00 | 0.00 | 6.52 |
| 4351 | 0.0026 | 0.00 | 0.00 | 6.52 |
| 4352 | 0.0016 | 0.00 | 0.00 | 6.52 |
| 4353 | 0.0026 | 0.00 | 0.00 | 6.52 |
| 4354 | 0.0022 | 0.00 | 0.00 | 6.51 |
| 4355 | 0.0019 | 0.00 | 0.00 | 6.51 |
| 4356 | 0.0013 | 0.00 | 0.00 | 6.51 |
| 4357 | 0.0016 | 0.00 | 0.00 | 6.51 |

| | | | | |
|---|---|---|---|---|
| 4358 | 0.0013 | 0.00 | 0.00 | 6.51 |
| 4359 | 0.0026 | 0.00 | 0.00 | 6.50 |
| 4360 | 0.0010 | 0.00 | 0.00 | 6.50 |
| 4361 | 0.0022 | 0.00 | 0.00 | 6.50 |
| 4362 | 0.0022 | 0.00 | 0.00 | 6.50 |
| 4363 | 0.0019 | 0.00 | 0.00 | 6.50 |
| 4364 | 0.0013 | 0.00 | 0.00 | 6.49 |
| 4365 | 0.0013 | 0.00 | 0.00 | 6.49 |
| 4366 | 0.0019 | 0.00 | 0.00 | 6.49 |
| 4367 | 0.0006 | 0.00 | 0.00 | 6.49 |
| 4368 | 0.0022 | 0.00 | 0.00 | 6.49 |
| 4369 | 0.0006 | 0.00 | 0.00 | 6.49 |
| 4370 | 0.0013 | 0.00 | 0.00 | 6.49 |
| 4371 | 0.0026 | 0.00 | 0.00 | 6.49 |
| 4372 | 0.0029 | 0.00 | 0.00 | 6.48 |
| 4373 | 0.0013 | 0.00 | 0.00 | 6.48 |
| 4374 | 0.0016 | 0.00 | 0.00 | 6.48 |
| 4375 | 0.0013 | 0.00 | 0.00 | 6.48 |
| 4376 | 0.0019 | 0.00 | 0.00 | 6.48 |
| 4377 | 0.0016 | 0.00 | 0.00 | 6.47 |
| 4378 | 0.0016 | 0.00 | 0.00 | 6.47 |
| 4379 | 0.0026 | 0.00 | 0.00 | 6.47 |
| 4380 | 0.0022 | 0.00 | 0.00 | 6.47 |
| 4381 | 0.0016 | 0.00 | 0.00 | 6.47 |
| 4382 | 0.0016 | 0.00 | 0.00 | 6.46 |
| 4383 | 0.0016 | 0.00 | 0.00 | 6.46 |
| 4384 | 0.0026 | 0.00 | 0.00 | 6.46 |
| 4385 | 0.0006 | 0.00 | 0.00 | 6.46 |
| 4386 | 0.0019 | 0.00 | 0.00 | 6.46 |
| 4387 | 0.0013 | 0.00 | 0.00 | 6.46 |
| 4388 | 0.0010 | 0.00 | 0.00 | 6.45 |
| 4389 | 0.0010 | 0.00 | 0.00 | 6.45 |
| 4390 | 0.0026 | 0.00 | 0.00 | 6.45 |
| 4391 | 0.0019 | 0.00 | 0.00 | 6.45 |
| 4392 | 0.0026 | 0.00 | 0.00 | 6.45 |
| 4393 | 0.0026 | 0.00 | 0.00 | 6.45 |
| 4394 | 0.0019 | 0.00 | 0.00 | 6.44 |
| 4395 | 0.0013 | 0.00 | 0.00 | 6.44 |
| 4396 | 0.0016 | 0.00 | 0.00 | 6.44 |
| 4397 | 0.0006 | 0.00 | 0.00 | 6.44 |
| 4398 | 0.0006 | 0.00 | 0.00 | 6.44 |
| 4399 | 0.0026 | 0.00 | 0.00 | 6.44 |
| 4400 | 0.0013 | 0.00 | 0.00 | 6.43 |
| 4401 | 0.0019 | 0.00 | 0.00 | 6.43 |

| | | | | |
|---|---|---|---|---|
| 4402 | 0.0016 | 0.00 | 0.00 | 6.43 |
| 4403 | 0.0003 | 0.00 | 0.00 | 6.43 |
| 4404 | 0.0006 | 0.00 | 0.00 | 6.43 |
| 4405 | 0.0022 | 0.00 | 0.00 | 6.43 |
| 4406 | 0.0019 | 0.00 | 0.00 | 6.43 |
| 4407 | 0.0013 | 0.00 | 0.00 | 6.42 |
| 4408 | 0.0022 | 0.00 | 0.00 | 6.42 |
| 4409 | 0.0006 | 0.00 | 0.00 | 6.42 |
| 4410 | 0.0022 | 0.00 | 0.00 | 6.42 |
| 4411 | 0.0019 | 0.00 | 0.00 | 6.42 |
| 4412 | 0.0016 | 0.00 | 0.00 | 6.42 |
| 4413 | 0.0029 | 0.00 | 0.00 | 6.41 |
| 4414 | 0.0016 | 0.00 | 0.00 | 6.41 |
| 4415 | 0.0022 | 0.00 | 0.00 | 6.41 |
| 4416 | 0.0022 | 0.00 | 0.00 | 6.41 |
| 4417 | 0.0013 | 0.00 | 0.00 | 6.41 |
| 4418 | 0.0022 | 0.00 | 0.00 | 6.40 |
| 4419 | 0.0032 | 0.00 | 0.00 | 6.40 |
| 4420 | 0.0022 | 0.00 | 0.00 | 6.40 |
| 4421 | 0.0010 | 0.00 | 0.00 | 6.40 |
| 4422 | 0.0016 | 0.00 | 0.00 | 6.40 |
| 4423 | 0.0016 | 0.00 | 0.00 | 6.39 |
| 4424 | 0.0019 | 0.00 | 0.00 | 6.39 |
| 4425 | 0.0016 | 0.00 | 0.00 | 6.39 |
| 4426 | 0.0026 | 0.00 | 0.00 | 6.39 |
| 4427 | 0.0019 | 0.00 | 0.00 | 6.39 |
| 4428 | 0.0010 | 0.00 | 0.00 | 6.38 |
| 4429 | 0.0010 | 0.00 | 0.00 | 6.38 |
| 4430 | 0.0029 | 0.00 | 0.00 | 6.38 |
| 4431 | 0.0006 | 0.00 | 0.00 | 6.38 |
| 4432 | 0.0006 | 0.00 | 0.00 | 6.38 |
| 4433 | 0.0016 | 0.00 | 0.00 | 6.38 |
| 4434 | 0.0013 | 0.00 | 0.00 | 6.38 |
| 4435 | 0.0013 | 0.00 | 0.00 | 6.38 |
| 4436 | 0.0006 | 0.00 | 0.00 | 6.37 |
| 4437 | 0.0019 | 0.00 | 0.00 | 6.37 |
| 4438 | 0.0010 | 0.00 | 0.00 | 6.37 |
| 4439 | 0.0019 | 0.00 | 0.00 | 6.37 |
| 4440 | 0.0006 | 0.00 | 0.00 | 6.37 |
| 4441 | 0.0029 | 0.00 | 0.00 | 6.37 |
| 4442 | 0.0022 | 0.00 | 0.00 | 6.37 |
| 4443 | 0.0013 | 0.00 | 0.00 | 6.36 |
| 4444 | 0.0022 | 0.00 | 0.00 | 6.36 |
| 4445 | 0.0016 | 0.00 | 0.00 | 6.36 |

| | | | | |
|---|---|---|---|---|
| 4446 | 0.0022 | 0.00 | 0.00 | 6.36 |
| 4447 | 0.0013 | 0.00 | 0.00 | 6.36 |
| 4448 | 0.0019 | 0.00 | 0.00 | 6.35 |
| 4449 | 0.0029 | 0.00 | 0.00 | 6.35 |
| 4450 | 0.0019 | 0.00 | 0.00 | 6.35 |
| 4451 | 0.0006 | 0.00 | 0.00 | 6.35 |
| 4452 | 0.0019 | 0.00 | 0.00 | 6.35 |
| 4453 | 0.0010 | 0.00 | 0.00 | 6.35 |
| 4454 | 0.0013 | 0.00 | 0.00 | 6.34 |
| 4455 | 0.0003 | 0.00 | 0.00 | 6.34 |
| 4456 | 0.0016 | 0.00 | 0.00 | 6.34 |
| 4457 | 0.0016 | 0.00 | 0.00 | 6.34 |
| 4458 | 0.0022 | 0.00 | 0.00 | 6.34 |
| 4459 | 0.0006 | 0.00 | 0.00 | 6.34 |
| 4460 | 0.0019 | 0.00 | 0.00 | 6.34 |
| 4461 | 0.0006 | 0.00 | 0.00 | 6.33 |
| 4462 | 0.0019 | 0.00 | 0.00 | 6.33 |
| 4463 | 0.0019 | 0.00 | 0.00 | 6.33 |
| 4464 | 0.0019 | 0.00 | 0.00 | 6.33 |
| 4465 | 0.0019 | 0.00 | 0.00 | 6.33 |
| 4466 | 0.0016 | 0.00 | 0.00 | 6.33 |
| 4467 | 0.0016 | 0.00 | 0.00 | 6.32 |
| 4468 | 0.0019 | 0.00 | 0.00 | 6.32 |
| 4469 | 0.0003 | 0.00 | 0.00 | 6.32 |
| 4470 | 0.0016 | 0.00 | 0.00 | 6.32 |
| 4471 | 0.0006 | 0.00 | 0.00 | 6.32 |
| 4472 | 0.0016 | 0.00 | 0.00 | 6.32 |
| 4473 | 0.0026 | 0.00 | 0.00 | 6.32 |
| 4474 | 0.0022 | 0.00 | 0.00 | 6.31 |
| 4475 | 0.0029 | 0.00 | 0.00 | 6.31 |
| 4476 | 0.0013 | 0.00 | 0.00 | 6.31 |
| 4477 | 0.0019 | 0.00 | 0.00 | 6.31 |
| 4478 | 0.0013 | 0.00 | 0.00 | 6.31 |
| 4479 | 0.0026 | 0.00 | 0.00 | 6.30 |
| 4480 | 0.0013 | 0.00 | 0.00 | 6.30 |
| 4481 | 0.0016 | 0.00 | 0.00 | 6.30 |
| 4482 | 0.0032 | 0.00 | 0.00 | 6.30 |
| 4483 | 0.0013 | 0.00 | 0.00 | 6.30 |
| 4484 | 0.0016 | 0.00 | 0.00 | 6.29 |
| 4485 | 0.0026 | 0.00 | 0.00 | 6.29 |
| 4486 | 0.0016 | 0.00 | 0.00 | 6.29 |
| 4487 | 0.0010 | 0.00 | 0.00 | 6.29 |
| 4488 | 0.0016 | 0.00 | 0.00 | 6.29 |
| 4489 | 0.0022 | 0.00 | 0.00 | 6.29 |

| | | | | |
|---|---|---|---|---|
| 4490 | 0.0032 | 0.00 | 0.00 | 6.28 |
| 4491 | 0.0013 | 0.00 | 0.00 | 6.28 |
| 4492 | 0.0013 | 0.00 | 0.00 | 6.28 |
| 4493 | 0.0016 | 0.00 | 0.00 | 6.28 |
| 4494 | 0.0026 | 0.00 | 0.00 | 6.28 |
| 4495 | 0.0013 | 0.00 | 0.00 | 6.27 |
| 4496 | 0.0016 | 0.00 | 0.00 | 6.27 |
| 4497 | 0.0003 | 0.00 | 0.00 | 6.27 |
| 4498 | 0.0026 | 0.00 | 0.00 | 6.27 |
| 4499 | 0.0016 | 0.00 | 0.00 | 6.27 |
| 4500 | 0.0016 | 0.00 | 0.00 | 6.27 |
| 4501 | 0.0013 | 0.00 | 0.00 | 6.27 |
| 4502 | 0.0013 | 0.00 | 0.00 | 6.26 |
| 4503 | 0.0016 | 0.00 | 0.00 | 6.26 |
| 4504 | 0.0006 | 0.00 | 0.00 | 6.26 |
| 4505 | 0.0026 | 0.00 | 0.00 | 6.26 |
| 4506 | 0.0019 | 0.00 | 0.00 | 6.26 |
| 4507 | 0.0016 | 0.00 | 0.00 | 6.26 |
| 4508 | 0.0026 | 0.00 | 0.00 | 6.25 |
| 4509 | 0.0019 | 0.00 | 0.00 | 6.25 |
| 4510 | 0.0038 | 0.00 | 0.00 | 6.25 |
| 4511 | 0.0022 | 0.00 | 0.00 | 6.25 |
| 4512 | 0.0026 | 0.00 | 0.00 | 6.24 |
| 4513 | 0.0022 | 0.00 | 0.00 | 6.24 |
| 4514 | 0.0022 | 0.00 | 0.00 | 6.24 |
| 4515 | 0.0035 | 0.00 | 0.00 | 6.24 |
| 4516 | 0.0029 | 0.00 | 0.00 | 6.23 |
| 4517 | 0.0016 | 0.00 | 0.00 | 6.23 |
| 4518 | 0.0022 | 0.00 | 0.00 | 6.23 |
| 4519 | 0.0013 | 0.00 | 0.00 | 6.23 |
| 4520 | 0.0006 | 0.00 | 0.00 | 6.23 |
| 4521 | 0.0029 | 0.00 | 0.00 | 6.22 |
| 4522 | 0.0013 | 0.00 | 0.00 | 6.22 |
| 4523 | 0.0019 | 0.00 | 0.00 | 6.22 |
| 4524 | 0.0019 | 0.00 | 0.00 | 6.22 |
| 4525 | 0.0026 | 0.00 | 0.00 | 6.22 |
| 4526 | 0.0013 | 0.00 | 0.00 | 6.21 |
| 4527 | 0.0013 | 0.00 | 0.00 | 6.21 |
| 4528 | 0.0006 | 0.00 | 0.00 | 6.21 |
| 4529 | 0.0010 | 0.00 | 0.00 | 6.21 |
| 4530 | 0.0010 | 0.00 | 0.00 | 6.21 |
| 4531 | 0.0010 | 0.00 | 0.00 | 6.21 |
| 4532 | 0.0019 | 0.00 | 0.00 | 6.21 |
| 4533 | 0.0019 | 0.00 | 0.00 | 6.21 |

| | | | | |
|---|---|---|---|---|
| 4534 | 0.0029 | 0.00 | 0.00 | 6.20 |
| 4535 | 0.0013 | 0.00 | 0.00 | 6.20 |
| 4536 | 0.0019 | 0.00 | 0.00 | 6.20 |
| 4537 | 0.0019 | 0.00 | 0.00 | 6.20 |
| 4538 | 0.0013 | 0.00 | 0.00 | 6.20 |
| 4539 | 0.0010 | 0.00 | 0.00 | 6.19 |
| 4540 | 0.0013 | 0.00 | 0.00 | 6.19 |
| 4541 | 0.0038 | 0.00 | 0.00 | 6.19 |
| 4542 | 0.0029 | 0.00 | 0.00 | 6.19 |
| 4543 | 0.0026 | 0.00 | 0.00 | 6.19 |
| 4544 | 0.0006 | 0.00 | 0.00 | 6.18 |
| 4545 | 0.0026 | 0.00 | 0.00 | 6.18 |
| 4546 | 0.0026 | 0.00 | 0.00 | 6.18 |
| 4547 | 0.0013 | 0.00 | 0.00 | 6.18 |
| 4548 | 0.0026 | 0.00 | 0.00 | 6.18 |
| 4549 | 0.0010 | 0.00 | 0.00 | 6.17 |
| 4550 | 0.0019 | 0.00 | 0.00 | 6.17 |
| 4551 | 0.0022 | 0.00 | 0.00 | 6.17 |
| 4552 | 0.0022 | 0.00 | 0.00 | 6.17 |
| 4553 | 0.0016 | 0.00 | 0.00 | 6.17 |
| 4554 | 0.0019 | 0.00 | 0.00 | 6.16 |
| 4555 | 0.0019 | 0.00 | 0.00 | 6.16 |
| 4556 | 0.0026 | 0.00 | 0.00 | 6.16 |
| 4557 | 0.0010 | 0.00 | 0.00 | 6.16 |
| 4558 | 0.0035 | 0.00 | 0.00 | 6.16 |
| 4559 | 0.0035 | 0.00 | 0.00 | 6.15 |
| 4560 | 0.0029 | 0.00 | 0.00 | 6.15 |
| 4561 | 0.0026 | 0.00 | 0.00 | 6.15 |
| 4562 | 0.0010 | 0.00 | 0.00 | 6.14 |
| 4563 | 0.0013 | 0.00 | 0.00 | 6.14 |
| 4564 | 0.0010 | 0.00 | 0.00 | 6.14 |
| 4565 | 0.0013 | 0.00 | 0.00 | 6.14 |
| 4566 | 0.0016 | 0.00 | 0.00 | 6.14 |
| 4567 | 0.0010 | 0.00 | 0.00 | 6.14 |
| 4568 | 0.0019 | 0.00 | 0.00 | 6.14 |
| 4569 | 0.0016 | 0.00 | 0.00 | 6.14 |
| 4570 | 0.0026 | 0.00 | 0.00 | 6.13 |
| 4571 | 0.0013 | 0.00 | 0.00 | 6.13 |
| 4572 | 0.0019 | 0.00 | 0.00 | 6.13 |
| 4573 | 0.0010 | 0.00 | 0.00 | 6.13 |
| 4574 | 0.0032 | 0.00 | 0.00 | 6.13 |
| 4575 | 0.0026 | 0.00 | 0.00 | 6.12 |
| 4576 | 0.0010 | 0.00 | 0.00 | 6.12 |
| 4577 | 0.0026 | 0.00 | 0.00 | 6.12 |

| | | | | |
|---|---|---|---|---|
| 4578 | 0.0029 | 0.00 | 0.00 | 6.12 |
| 4579 | 0.0022 | 0.00 | 0.00 | 6.12 |
| 4580 | 0.0003 | 0.00 | 0.00 | 6.11 |
| 4581 | 0.0006 | 0.00 | 0.00 | 6.11 |
| 4582 | 0.0022 | 0.00 | 0.00 | 6.11 |
| 4583 | 0.0026 | 0.00 | 0.00 | 6.11 |
| 4584 | 0.0003 | 0.00 | 0.00 | 6.11 |
| 4585 | 0.0019 | 0.00 | 0.00 | 6.11 |
| 4586 | 0.0029 | 0.00 | 0.00 | 6.11 |
| 4587 | 0.0022 | 0.00 | 0.00 | 6.10 |
| 4588 | 0.0038 | 0.00 | 0.00 | 6.10 |
| 4589 | 0.0019 | 0.00 | 0.00 | 6.10 |
| 4590 | 0.0019 | 0.00 | 0.00 | 6.09 |
| 4591 | 0.0013 | 0.00 | 0.00 | 6.09 |
| 4592 | 0.0013 | 0.00 | 0.00 | 6.09 |
| 4593 | 0.0022 | 0.00 | 0.00 | 6.09 |
| 4594 | 0.0016 | 0.00 | 0.00 | 6.09 |
| 4595 | 0.0022 | 0.00 | 0.00 | 6.09 |
| 4596 | 0.0022 | 0.00 | 0.00 | 6.08 |
| 4597 | 0.0038 | 0.00 | 0.00 | 6.08 |
| 4598 | 0.0013 | 0.00 | 0.00 | 6.08 |
| 4599 | 0.0026 | 0.00 | 0.00 | 6.08 |
| 4600 | 0.0022 | 0.00 | 0.00 | 6.07 |
| 4601 | 0.0029 | 0.00 | 0.00 | 6.07 |
| 4602 | 0.0029 | 0.00 | 0.00 | 6.07 |
| 4603 | 0.0013 | 0.00 | 0.00 | 6.07 |
| 4604 | 0.0016 | 0.00 | 0.00 | 6.06 |
| 4605 | 0.0019 | 0.00 | 0.00 | 6.06 |
| 4606 | 0.0019 | 0.00 | 0.00 | 6.06 |
| 4607 | 0.0032 | 0.00 | 0.00 | 6.06 |
| 4608 | 0.0013 | 0.00 | 0.00 | 6.06 |
| 4609 | 0.0035 | 0.00 | 0.00 | 6.05 |
| 4610 | 0.0019 | 0.00 | 0.00 | 6.05 |
| 4611 | 0.0026 | 0.00 | 0.00 | 6.05 |
| 4612 | 0.0016 | 0.00 | 0.00 | 6.05 |
| 4613 | 0.0016 | 0.00 | 0.00 | 6.04 |
| 4614 | 0.0022 | 0.00 | 0.00 | 6.04 |
| 4615 | 0.0013 | 0.00 | 0.00 | 6.04 |
| 4616 | 0.0013 | 0.00 | 0.00 | 6.04 |
| 4617 | 0.0010 | 0.00 | 0.00 | 6.04 |
| 4618 | 0.0026 | 0.00 | 0.00 | 6.04 |
| 4619 | 0.0006 | 0.00 | 0.00 | 6.03 |
| 4620 | 0.0013 | 0.00 | 0.00 | 6.03 |
| 4621 | 0.0032 | 0.00 | 0.00 | 6.03 |

| | | | | |
|---|---|---|---|---|
| 4622 | 0.0010 | 0.00 | 0.00 | 6.03 |
| 4623 | 0.0019 | 0.00 | 0.00 | 6.03 |
| 4624 | 0.0019 | 0.00 | 0.00 | 6.03 |
| 4625 | 0.0019 | 0.00 | 0.00 | 6.03 |
| 4626 | 0.0019 | 0.00 | 0.00 | 6.02 |
| 4627 | 0.0022 | 0.00 | 0.00 | 6.02 |
| 4628 | 0.0026 | 0.00 | 0.00 | 6.02 |
| 4629 | 0.0016 | 0.00 | 0.00 | 6.02 |
| 4630 | 0.0019 | 0.00 | 0.00 | 6.01 |
| 4631 | 0.0029 | 0.00 | 0.00 | 6.01 |
| 4632 | 0.0019 | 0.00 | 0.00 | 6.01 |
| 4633 | 0.0016 | 0.00 | 0.00 | 6.01 |
| 4634 | 0.0019 | 0.00 | 0.00 | 6.01 |
| 4635 | 0.0032 | 0.00 | 0.00 | 6.00 |
| 4636 | 0.0032 | 0.00 | 0.00 | 6.00 |
| 4637 | 0.0010 | 0.00 | 0.00 | 6.00 |
| 4638 | 0.0038 | 0.00 | 0.00 | 6.00 |
| 4639 | 0.0026 | 0.00 | 0.00 | 5.99 |
| 4640 | 0.0026 | 0.00 | 0.00 | 5.99 |
| 4641 | 0.0022 | 0.00 | 0.00 | 5.99 |
| 4642 | 0.0019 | 0.00 | 0.00 | 5.99 |
| 4643 | 0.0019 | 0.00 | 0.00 | 5.98 |
| 4644 | 0.0013 | 0.00 | 0.00 | 5.98 |
| 4645 | 0.0029 | 0.00 | 0.00 | 5.98 |
| 4646 | 0.0013 | 0.00 | 0.00 | 5.98 |
| 4647 | 0.0016 | 0.00 | 0.00 | 5.98 |
| 4648 | 0.0016 | 0.00 | 0.00 | 5.98 |
| 4649 | 0.0016 | 0.00 | 0.00 | 5.97 |
| 4650 | 0.0013 | 0.00 | 0.00 | 5.97 |
| 4651 | 0.0003 | 0.00 | 0.00 | 5.97 |
| 4652 | 0.0022 | 0.00 | 0.00 | 5.97 |
| 4653 | 0.0022 | 0.00 | 0.00 | 5.97 |
| 4654 | 0.0026 | 0.00 | 0.00 | 5.97 |
| 4655 | 0.0019 | 0.00 | 0.00 | 5.96 |
| 4656 | 0.0042 | 0.00 | 0.00 | 5.96 |
| 4657 | 0.0016 | 0.00 | 0.00 | 5.96 |
| 4658 | 0.0022 | 0.00 | 0.00 | 5.96 |
| 4659 | 0.0016 | 0.00 | 0.00 | 5.95 |
| 4660 | 0.0029 | 0.00 | 0.00 | 5.95 |
| 4661 | 0.0013 | 0.00 | 0.00 | 5.95 |
| 4662 | 0.0013 | 0.00 | 0.00 | 5.95 |
| 4663 | 0.0032 | 0.00 | 0.00 | 5.95 |
| 4664 | 0.0022 | 0.00 | 0.00 | 5.94 |
| 4665 | 0.0029 | 0.00 | 0.00 | 5.94 |

| | | | | |
|---|---|---|---|---|
| 4666 | 0.0006 | 0.00 | 0.00 | 5.94 |
| 4667 | 0.0019 | 0.00 | 0.00 | 5.94 |
| 4668 | 0.0022 | 0.00 | 0.00 | 5.94 |
| 4669 | 0.0013 | 0.00 | 0.00 | 5.93 |
| 4670 | 0.0013 | 0.00 | 0.00 | 5.93 |
| 4671 | 0.0026 | 0.00 | 0.00 | 5.93 |
| 4672 | 0.0038 | 0.00 | 0.00 | 5.93 |
| 4673 | 0.0016 | 0.00 | 0.00 | 5.92 |
| 4674 | 0.0019 | 0.00 | 0.00 | 5.92 |
| 4675 | 0.0013 | 0.00 | 0.00 | 5.92 |
| 4676 | 0.0026 | 0.00 | 0.00 | 5.92 |
| 4677 | 0.0016 | 0.00 | 0.00 | 5.92 |
| 4678 | 0.0019 | 0.00 | 0.00 | 5.92 |
| 4679 | 0.0019 | 0.00 | 0.00 | 5.91 |
| 4680 | 0.0013 | 0.00 | 0.00 | 5.91 |
| 4681 | 0.0013 | 0.00 | 0.00 | 5.91 |
| 4682 | 0.0010 | 0.00 | 0.00 | 5.91 |
| 4683 | 0.0026 | 0.00 | 0.00 | 5.91 |
| 4684 | 0.0022 | 0.00 | 0.00 | 5.91 |
| 4685 | 0.0006 | 0.00 | 0.00 | 5.90 |
| 4686 | 0.0019 | 0.00 | 0.00 | 5.90 |
| 4687 | 0.0013 | 0.00 | 0.00 | 5.90 |
| 4688 | 0.0013 | 0.00 | 0.00 | 5.90 |
| 4689 | 0.0016 | 0.00 | 0.00 | 5.90 |
| 4690 | 0.0013 | 0.00 | 0.00 | 5.90 |
| 4691 | 0.0032 | 0.00 | 0.00 | 5.89 |
| 4692 | 0.0022 | 0.00 | 0.00 | 5.89 |
| 4693 | 0.0029 | 0.00 | 0.00 | 5.89 |
| 4694 | 0.0013 | 0.00 | 0.00 | 5.89 |
| 4695 | 0.0019 | 0.00 | 0.00 | 5.89 |
| 4696 | 0.0022 | 0.00 | 0.00 | 5.88 |
| 4697 | 0.0022 | 0.00 | 0.00 | 5.88 |
| 4698 | 0.0026 | 0.00 | 0.00 | 5.88 |
| 4699 | 0.0026 | 0.00 | 0.00 | 5.88 |
| 4700 | 0.0019 | 0.00 | 0.00 | 5.87 |
| 4701 | 0.0032 | 0.00 | 0.00 | 5.87 |
| 4702 | 0.0026 | 0.00 | 0.00 | 5.87 |
| 4703 | 0.0035 | 0.00 | 0.00 | 5.87 |
| 4704 | 0.0032 | 0.00 | 0.00 | 5.86 |
| 4705 | 0.0019 | 0.00 | 0.00 | 5.86 |
| 4706 | 0.0022 | 0.00 | 0.00 | 5.86 |
| 4707 | 0.0016 | 0.00 | 0.00 | 5.86 |
| 4708 | 0.0016 | 0.00 | 0.00 | 5.85 |
| 4709 | 0.0016 | 0.00 | 0.00 | 5.85 |

| | | | | |
|---|---|---|---|---|
| 4710 | 0.0013 | 0.00 | 0.00 | 5.85 |
| 4711 | 0.0026 | 0.00 | 0.00 | 5.85 |
| 4712 | 0.0022 | 0.00 | 0.00 | 5.85 |
| 4713 | 0.0026 | 0.00 | 0.00 | 5.84 |
| 4714 | 0.0003 | 0.00 | 0.00 | 5.84 |
| 4715 | 0.0013 | 0.00 | 0.00 | 5.84 |
| 4716 | 0.0013 | 0.00 | 0.00 | 5.84 |
| 4717 | 0.0013 | 0.00 | 0.00 | 5.84 |
| 4718 | 0.0019 | 0.00 | 0.00 | 5.84 |
| 4719 | 0.0022 | 0.00 | 0.00 | 5.84 |
| 4720 | 0.0022 | 0.00 | 0.00 | 5.83 |
| 4721 | 0.0010 | 0.00 | 0.00 | 5.83 |
| 4722 | 0.0016 | 0.00 | 0.00 | 5.83 |
| 4723 | 0.0029 | 0.00 | 0.00 | 5.83 |
| 4724 | 0.0010 | 0.00 | 0.00 | 5.83 |
| 4725 | 0.0019 | 0.00 | 0.00 | 5.82 |
| 4726 | 0.0029 | 0.00 | 0.00 | 5.82 |
| 4727 | 0.0022 | 0.00 | 0.00 | 5.82 |
| 4728 | 0.0019 | 0.00 | 0.00 | 5.82 |
| 4729 | 0.0016 | 0.00 | 0.00 | 5.82 |
| 4730 | 0.0019 | 0.00 | 0.00 | 5.81 |
| 4731 | 0.0032 | 0.00 | 0.00 | 5.81 |
| 4732 | 0.0016 | 0.00 | 0.00 | 5.81 |
| 4733 | 0.0013 | 0.00 | 0.00 | 5.81 |
| 4734 | 0.0016 | 0.00 | 0.00 | 5.81 |
| 4735 | 0.0022 | 0.00 | 0.00 | 5.80 |
| 4736 | 0.0019 | 0.00 | 0.00 | 5.80 |
| 4737 | 0.0022 | 0.00 | 0.00 | 5.80 |
| 4738 | 0.0019 | 0.00 | 0.00 | 5.80 |
| 4739 | 0.0026 | 0.00 | 0.00 | 5.80 |
| 4740 | 0.0010 | 0.00 | 0.00 | 5.79 |
| 4741 | 0.0026 | 0.00 | 0.00 | 5.79 |
| 4742 | 0.0006 | 0.00 | 0.00 | 5.79 |
| 4743 | 0.0010 | 0.00 | 0.00 | 5.79 |
| 4744 | 0.0019 | 0.00 | 0.00 | 5.79 |
| 4745 | 0.0010 | 0.00 | 0.00 | 5.79 |
| 4746 | 0.0019 | 0.00 | 0.00 | 5.79 |
| 4747 | 0.0013 | 0.00 | 0.00 | 5.78 |
| 4748 | 0.0013 | 0.00 | 0.00 | 5.78 |
| 4749 | 0.0026 | 0.00 | 0.00 | 5.78 |
| 4750 | 0.0032 | 0.00 | 0.00 | 5.78 |
| 4751 | 0.0013 | 0.00 | 0.00 | 5.78 |
| 4752 | 0.0016 | 0.00 | 0.00 | 5.77 |
| 4753 | 0.0026 | 0.00 | 0.00 | 5.77 |

| | | | | |
|---|---|---|---|---|
| 4754 | 0.0013 | 0.00 | 0.00 | 5.77 |
| 4755 | 0.0026 | 0.00 | 0.00 | 5.77 |
| 4756 | 0.0013 | 0.00 | 0.00 | 5.77 |
| 4757 | 0.0013 | 0.00 | 0.00 | 5.76 |
| 4758 | 0.0013 | 0.00 | 0.00 | 5.76 |
| 4759 | 0.0022 | 0.00 | 0.00 | 5.76 |
| 4760 | 0.0016 | 0.00 | 0.00 | 5.76 |
| 4761 | 0.0029 | 0.00 | 0.00 | 5.76 |
| 4762 | 0.0026 | 0.00 | 0.00 | 5.76 |
| 4763 | 0.0026 | 0.00 | 0.00 | 5.75 |
| 4764 | 0.0016 | 0.00 | 0.00 | 5.75 |
| 4765 | 0.0032 | 0.00 | 0.00 | 5.75 |
| 4766 | 0.0016 | 0.00 | 0.00 | 5.75 |
| 4767 | 0.0013 | 0.00 | 0.00 | 5.74 |
| 4768 | 0.0022 | 0.00 | 0.00 | 5.74 |
| 4769 | 0.0035 | 0.00 | 0.00 | 5.74 |
| 4770 | 0.0013 | 0.00 | 0.00 | 5.74 |
| 4771 | 0.0013 | 0.00 | 0.00 | 5.74 |
| 4772 | 0.0016 | 0.00 | 0.00 | 5.73 |
| 4773 | 0.0013 | 0.00 | 0.00 | 5.73 |
| 4774 | 0.0016 | 0.00 | 0.00 | 5.73 |
| 4775 | 0.0022 | 0.00 | 0.00 | 5.73 |
| 4776 | 0.0029 | 0.00 | 0.00 | 5.73 |
| 4777 | 0.0016 | 0.00 | 0.00 | 5.72 |
| 4778 | 0.0006 | 0.00 | 0.00 | 5.72 |
| 4779 | 0.0010 | 0.00 | 0.00 | 5.72 |
| 4780 | 0.0013 | 0.00 | 0.00 | 5.72 |
| 4781 | 0.0016 | 0.00 | 0.00 | 5.72 |
| 4782 | 0.0022 | 0.00 | 0.00 | 5.72 |
| 4783 | 0.0013 | 0.00 | 0.00 | 5.72 |
| 4784 | 0.0022 | 0.00 | 0.00 | 5.72 |
| 4785 | 0.0016 | 0.00 | 0.00 | 5.71 |
| 4786 | 0.0019 | 0.00 | 0.00 | 5.71 |
| 4787 | 0.0016 | 0.00 | 0.00 | 5.71 |
| 4788 | 0.0022 | 0.00 | 0.00 | 5.71 |
| 4789 | 0.0019 | 0.00 | 0.00 | 5.71 |
| 4790 | 0.0010 | 0.00 | 0.00 | 5.70 |
| 4791 | 0.0019 | 0.00 | 0.00 | 5.70 |
| 4792 | 0.0016 | 0.00 | 0.00 | 5.70 |
| 4793 | 0.0032 | 0.00 | 0.00 | 5.70 |
| 4794 | 0.0029 | 0.00 | 0.00 | 5.70 |
| 4795 | 0.0016 | 0.00 | 0.00 | 5.69 |
| 4796 | 0.0006 | 0.00 | 0.00 | 5.69 |
| 4797 | 0.0006 | 0.00 | 0.00 | 5.69 |

| | | | | |
|---|---|---|---|---|
| 4798 | 0.0013 | 0.00 | 0.00 | 5.69 |
| 4799 | 0.0019 | 0.00 | 0.00 | 5.69 |
| 4800 | 0.0013 | 0.00 | 0.00 | 5.69 |
| 4801 | 0.0022 | 0.00 | 0.00 | 5.69 |
| 4802 | 0.0016 | 0.00 | 0.00 | 5.68 |
| 4803 | 0.0038 | 0.00 | 0.00 | 5.68 |
| 4804 | 0.0010 | 0.00 | 0.00 | 5.68 |
| 4805 | 0.0010 | 0.00 | 0.00 | 5.68 |
| 4806 | 0.0022 | 0.00 | 0.00 | 5.68 |
| 4807 | 0.0029 | 0.00 | 0.00 | 5.67 |
| 4808 | 0.0016 | 0.00 | 0.00 | 5.67 |
| 4809 | 0.0042 | 0.00 | 0.00 | 5.67 |
| 4810 | 0.0013 | 0.00 | 0.00 | 5.67 |
| 4811 | 0.0013 | 0.00 | 0.00 | 5.66 |
| 4812 | 0.0022 | 0.00 | 0.00 | 5.66 |
| 4813 | 0.0016 | 0.00 | 0.00 | 5.66 |
| 4814 | 0.0013 | 0.00 | 0.00 | 5.66 |
| 4815 | 0.0016 | 0.00 | 0.00 | 5.66 |
| 4816 | 0.0006 | 0.00 | 0.00 | 5.66 |
| 4817 | 0.0013 | 0.00 | 0.00 | 5.66 |
| 4818 | 0.0019 | 0.00 | 0.00 | 5.65 |
| 4819 | 0.0013 | 0.00 | 0.00 | 5.65 |
| 4820 | 0.0016 | 0.00 | 0.00 | 5.65 |
| 4821 | 0.0026 | 0.00 | 0.00 | 5.65 |
| 4822 | 0.0022 | 0.00 | 0.00 | 5.65 |
| 4823 | 0.0010 | 0.00 | 0.00 | 5.64 |
| 4824 | 0.0019 | 0.00 | 0.00 | 5.64 |
| 4825 | 0.0013 | 0.00 | 0.00 | 5.64 |
| 4826 | 0.0006 | 0.00 | 0.00 | 5.64 |
| 4827 | 0.0013 | 0.00 | 0.00 | 5.64 |
| 4828 | 0.0013 | 0.00 | 0.00 | 5.64 |
| 4829 | 0.0029 | 0.00 | 0.00 | 5.64 |
| 4830 | 0.0013 | 0.00 | 0.00 | 5.63 |
| 4831 | 0.0010 | 0.00 | 0.00 | 5.63 |
| 4832 | 0.0010 | 0.00 | 0.00 | 5.63 |
| 4833 | 0.0016 | 0.00 | 0.00 | 5.63 |
| 4834 | 0.0013 | 0.00 | 0.00 | 5.63 |
| 4835 | 0.0016 | 0.00 | 0.00 | 5.63 |
| 4836 | 0.0026 | 0.00 | 0.00 | 5.63 |
| 4837 | 0.0019 | 0.00 | 0.00 | 5.62 |
| 4838 | 0.0022 | 0.00 | 0.00 | 5.62 |
| 4839 | 0.0016 | 0.00 | 0.00 | 5.62 |
| 4840 | 0.0026 | 0.00 | 0.00 | 5.62 |
| 4841 | 0.0016 | 0.00 | 0.00 | 5.62 |

| | | | | |
|---|---|---|---|---|
| 4842 | 0.0022 | 0.00 | 0.00 | 5.61 |
| 4843 | 0.0016 | 0.00 | 0.00 | 5.61 |
| 4844 | 0.0016 | 0.00 | 0.00 | 5.61 |
| 4845 | 0.0019 | 0.00 | 0.00 | 5.61 |
| 4846 | 0.0035 | 0.00 | 0.00 | 5.61 |
| 4847 | 0.0016 | 0.00 | 0.00 | 5.60 |
| 4848 | 0.0013 | 0.00 | 0.00 | 5.60 |
| 4849 | 0.0029 | 0.00 | 0.00 | 5.60 |
| 4850 | 0.0022 | 0.00 | 0.00 | 5.60 |
| 4851 | 0.0026 | 0.00 | 0.00 | 5.59 |
| 4852 | 0.0010 | 0.00 | 0.00 | 5.59 |
| 4853 | 0.0035 | 0.00 | 0.00 | 5.59 |
| 4854 | 0.0010 | 0.00 | 0.00 | 5.59 |
| 4855 | 0.0006 | 0.00 | 0.00 | 5.59 |
| 4856 | 0.0013 | 0.00 | 0.00 | 5.59 |
| 4857 | 0.0016 | 0.00 | 0.00 | 5.58 |
| 4858 | 0.0022 | 0.00 | 0.00 | 5.58 |
| 4859 | 0.0013 | 0.00 | 0.00 | 5.58 |
| 4860 | 0.0010 | 0.00 | 0.00 | 5.58 |
| 4861 | 0.0026 | 0.00 | 0.00 | 5.58 |
| 4862 | 0.0022 | 0.00 | 0.00 | 5.58 |
| 4863 | 0.0010 | 0.00 | 0.00 | 5.57 |
| 4864 | 0.0013 | 0.00 | 0.00 | 5.57 |
| 4865 | 0.0013 | 0.00 | 0.00 | 5.57 |
| 4867 | 0.0010 | 0.00 | 0.00 | 5.57 |
| 4868 | 0.0010 | 0.00 | 0.00 | 5.57 |
| 4869 | 0.0010 | 0.00 | 0.00 | 5.57 |
| 4870 | 0.0019 | 0.00 | 0.00 | 5.57 |
| 4871 | 0.0016 | 0.00 | 0.00 | 5.57 |
| 4872 | 0.0026 | 0.00 | 0.00 | 5.56 |
| 4873 | 0.0029 | 0.00 | 0.00 | 5.56 |
| 4874 | 0.0013 | 0.00 | 0.00 | 5.56 |
| 4875 | 0.0010 | 0.00 | 0.00 | 5.56 |
| 4876 | 0.0016 | 0.00 | 0.00 | 5.56 |
| 4877 | 0.0016 | 0.00 | 0.00 | 5.55 |
| 4878 | 0.0010 | 0.00 | 0.00 | 5.55 |
| 4879 | 0.0006 | 0.00 | 0.00 | 5.55 |
| 4880 | 0.0019 | 0.00 | 0.00 | 5.55 |
| 4881 | 0.0016 | 0.00 | 0.00 | 5.55 |
| 4882 | 0.0013 | 0.00 | 0.00 | 5.55 |
| 4883 | 0.0022 | 0.00 | 0.00 | 5.55 |
| 4884 | 0.0042 | 0.00 | 0.00 | 5.54 |
| 4885 | 0.0045 | 0.00 | 0.00 | 5.54 |
| 4886 | 0.0022 | 0.00 | 0.00 | 5.54 |

| | | | | |
|---|---|---|---|---|
| 4887 | 0.0019 | 0.00 | 0.00 | 5.53 |
| 4888 | 0.0026 | 0.00 | 0.00 | 5.53 |
| 4889 | 0.0045 | 0.00 | 0.00 | 5.53 |
| 4890 | 0.0016 | 0.00 | 0.00 | 5.52 |
| 4891 | 0.0026 | 0.00 | 0.00 | 5.52 |
| 4892 | 0.0019 | 0.00 | 0.00 | 5.52 |
| 4893 | 0.0016 | 0.00 | 0.00 | 5.52 |
| 4894 | 0.0019 | 0.00 | 0.00 | 5.52 |
| 4895 | 0.0016 | 0.00 | 0.00 | 5.52 |
| 4897 | 0.0029 | 0.00 | 0.00 | 5.51 |
| 4898 | 0.0016 | 0.00 | 0.00 | 5.51 |
| 4899 | 0.0006 | 0.00 | 0.00 | 5.51 |
| 4900 | 0.0016 | 0.00 | 0.00 | 5.51 |
| 4901 | 0.0010 | 0.00 | 0.00 | 5.51 |
| 4902 | 0.0026 | 0.00 | 0.00 | 5.51 |
| 4903 | 0.0019 | 0.00 | 0.00 | 5.50 |
| 4904 | 0.0019 | 0.00 | 0.00 | 5.50 |
| 4905 | 0.0032 | 0.00 | 0.00 | 5.50 |
| 4906 | 0.0016 | 0.00 | 0.00 | 5.50 |
| 4907 | 0.0013 | 0.00 | 0.00 | 5.49 |
| 4908 | 0.0026 | 0.00 | 0.00 | 5.49 |
| 4909 | 0.0016 | 0.00 | 0.00 | 5.49 |
| 4910 | 0.0029 | 0.00 | 0.00 | 5.49 |
| 4911 | 0.0026 | 0.00 | 0.00 | 5.49 |
| 4912 | 0.0013 | 0.00 | 0.00 | 5.48 |
| 4913 | 0.0029 | 0.00 | 0.00 | 5.48 |
| 4914 | 0.0010 | 0.00 | 0.00 | 5.48 |
| 4915 | 0.0006 | 0.00 | 0.00 | 5.48 |
| 4916 | 0.0013 | 0.00 | 0.00 | 5.48 |
| 4917 | 0.0010 | 0.00 | 0.00 | 5.48 |
| 4918 | 0.0013 | 0.00 | 0.00 | 5.48 |
| 4919 | 0.0022 | 0.00 | 0.00 | 5.47 |
| 4920 | 0.0019 | 0.00 | 0.00 | 5.47 |
| 4921 | 0.0013 | 0.00 | 0.00 | 5.47 |
| 4922 | 0.0019 | 0.00 | 0.00 | 5.47 |
| 4923 | 0.0010 | 0.00 | 0.00 | 5.47 |
| 4924 | 0.0029 | 0.00 | 0.00 | 5.47 |
| 4925 | 0.0006 | 0.00 | 0.00 | 5.46 |
| 4926 | 0.0013 | 0.00 | 0.00 | 5.46 |
| 4927 | 0.0016 | 0.00 | 0.00 | 5.46 |
| 4928 | 0.0013 | 0.00 | 0.00 | 5.46 |
| 4929 | 0.0019 | 0.00 | 0.00 | 5.46 |
| 4930 | 0.0006 | 0.00 | 0.00 | 5.46 |
| 4931 | 0.0035 | 0.00 | 0.00 | 5.46 |

| | | | | |
|---|---|---|---|---|
| 4932 | 0.0013 | 0.00 | 0.00 | 5.45 |
| 4933 | 0.0026 | 0.00 | 0.00 | 5.45 |
| 4934 | 0.0010 | 0.00 | 0.00 | 5.45 |
| 4935 | 0.0013 | 0.00 | 0.00 | 5.45 |
| 4936 | 0.0016 | 0.00 | 0.00 | 5.45 |
| 4937 | 0.0032 | 0.00 | 0.00 | 5.44 |
| 4938 | 0.0006 | 0.00 | 0.00 | 5.44 |
| 4939 | 0.0016 | 0.00 | 0.00 | 5.44 |
| 4940 | 0.0022 | 0.00 | 0.00 | 5.44 |
| 4941 | 0.0016 | 0.00 | 0.00 | 5.44 |
| 4942 | 0.0022 | 0.00 | 0.00 | 5.44 |
| 4943 | 0.0019 | 0.00 | 0.00 | 5.43 |
| 4944 | 0.0006 | 0.00 | 0.00 | 5.43 |
| 4945 | 0.0010 | 0.00 | 0.00 | 5.43 |
| 4946 | 0.0016 | 0.00 | 0.00 | 5.43 |
| 4947 | 0.0016 | 0.00 | 0.00 | 5.43 |
| 4948 | 0.0013 | 0.00 | 0.00 | 5.43 |
| 4949 | 0.0019 | 0.00 | 0.00 | 5.43 |
| 4950 | 0.0019 | 0.00 | 0.00 | 5.42 |
| 4951 | 0.0019 | 0.00 | 0.00 | 5.42 |
| 4952 | 0.0013 | 0.00 | 0.00 | 5.42 |
| 4953 | 0.0022 | 0.00 | 0.00 | 5.42 |
| 4954 | 0.0022 | 0.00 | 0.00 | 5.42 |
| 4955 | 0.0016 | 0.00 | 0.00 | 5.41 |
| 4956 | 0.0013 | 0.00 | 0.00 | 5.41 |
| 4957 | 0.0013 | 0.00 | 0.00 | 5.41 |
| 4958 | 0.0006 | 0.00 | 0.00 | 5.41 |
| 4959 | 0.0016 | 0.00 | 0.00 | 5.41 |
| 4960 | 0.0016 | 0.00 | 0.00 | 5.41 |
| 4961 | 0.0010 | 0.00 | 0.00 | 5.41 |
| 4962 | 0.0016 | 0.00 | 0.00 | 5.40 |
| 4963 | 0.0013 | 0.00 | 0.00 | 5.40 |
| 4964 | 0.0013 | 0.00 | 0.00 | 5.40 |
| 4965 | 0.0022 | 0.00 | 0.00 | 5.40 |
| 4966 | 0.0019 | 0.00 | 0.00 | 5.40 |
| 4967 | 0.0022 | 0.00 | 0.00 | 5.40 |
| 4968 | 0.0026 | 0.00 | 0.00 | 5.39 |
| 4969 | 0.0016 | 0.00 | 0.00 | 5.39 |
| 4970 | 0.0019 | 0.00 | 0.00 | 5.39 |
| 4971 | 0.0035 | 0.00 | 0.00 | 5.39 |
| 4972 | 0.0006 | 0.00 | 0.00 | 5.38 |
| 4973 | 0.0013 | 0.00 | 0.00 | 5.38 |
| 4974 | 0.0029 | 0.00 | 0.00 | 5.38 |
| 4975 | 0.0013 | 0.00 | 0.00 | 5.38 |

| | | | | |
|---|---|---|---|---|
| 4976 | 0.0026 | 0.00 | 0.00 | 5.38 |
| 4977 | 0.0019 | 0.00 | 0.00 | 5.38 |
| 4978 | 0.0003 | 0.00 | 0.00 | 5.37 |
| 4979 | 0.0013 | 0.00 | 0.00 | 5.37 |
| 4980 | 0.0006 | 0.00 | 0.00 | 5.37 |
| 4981 | 0.0013 | 0.00 | 0.00 | 5.37 |
| 4982 | 0.0010 | 0.00 | 0.00 | 5.37 |
| 4983 | 0.0016 | 0.00 | 0.00 | 5.37 |
| 4984 | 0.0022 | 0.00 | 0.00 | 5.37 |
| 4985 | 0.0019 | 0.00 | 0.00 | 5.37 |
| 4986 | 0.0003 | 0.00 | 0.00 | 5.36 |
| 4987 | 0.0016 | 0.00 | 0.00 | 5.36 |
| 4988 | 0.0003 | 0.00 | 0.00 | 5.36 |
| 4989 | 0.0022 | 0.00 | 0.00 | 5.36 |
| 4990 | 0.0006 | 0.00 | 0.00 | 5.36 |
| 4991 | 0.0003 | 0.00 | 0.00 | 5.36 |
| 4992 | 0.0006 | 0.00 | 0.00 | 5.36 |
| 4993 | 0.0006 | 0.00 | 0.00 | 5.36 |
| 4994 | 0.0022 | 0.00 | 0.00 | 5.36 |
| 4995 | 0.0013 | 0.00 | 0.00 | 5.35 |
| 4996 | 0.0010 | 0.00 | 0.00 | 5.35 |
| 4997 | 0.0022 | 0.00 | 0.00 | 5.35 |
| 4998 | 0.0032 | 0.00 | 0.00 | 5.35 |
| 4999 | 0.0010 | 0.00 | 0.00 | 5.35 |
| 5000 | 0.0016 | 0.00 | 0.00 | 5.35 |
| 5001 | 0.0022 | 0.00 | 0.00 | 5.34 |
| 5002 | 0.0006 | 0.00 | 0.00 | 5.34 |
| 5003 | 0.0016 | 0.00 | 0.00 | 5.34 |
| 5004 | 0.0016 | 0.00 | 0.00 | 5.34 |
| 5005 | 0.0013 | 0.00 | 0.00 | 5.34 |
| 5006 | 0.0019 | 0.00 | 0.00 | 5.34 |
| 5007 | 0.0022 | 0.00 | 0.00 | 5.33 |
| 5008 | 0.0006 | 0.00 | 0.00 | 5.33 |
| 5009 | 0.0016 | 0.00 | 0.00 | 5.33 |
| 5010 | 0.0010 | 0.00 | 0.00 | 5.33 |
| 5011 | 0.0026 | 0.00 | 0.00 | 5.33 |
| 5012 | 0.0010 | 0.00 | 0.00 | 5.33 |
| 5013 | 0.0016 | 0.00 | 0.00 | 5.33 |
| 5014 | 0.0013 | 0.00 | 0.00 | 5.32 |
| 5015 | 0.0029 | 0.00 | 0.00 | 5.32 |
| 5016 | 0.0010 | 0.00 | 0.00 | 5.32 |
| 5017 | 0.0006 | 0.00 | 0.00 | 5.32 |
| 5018 | 0.0013 | 0.00 | 0.00 | 5.32 |
| 5019 | 0.0010 | 0.00 | 0.00 | 5.32 |

| | | | | |
|---|---|---|---|---|
| 5020 | 0.0016 | 0.00 | 0.00 | 5.32 |
| 5021 | 0.0010 | 0.00 | 0.00 | 5.31 |
| 5022 | 0.0013 | 0.00 | 0.00 | 5.31 |
| 5023 | 0.0019 | 0.00 | 0.00 | 5.31 |
| 5024 | 0.0016 | 0.00 | 0.00 | 5.31 |
| 5025 | 0.0010 | 0.00 | 0.00 | 5.31 |
| 5026 | 0.0010 | 0.00 | 0.00 | 5.31 |
| 5027 | 0.0022 | 0.00 | 0.00 | 5.31 |
| 5028 | 0.0013 | 0.00 | 0.00 | 5.30 |
| 5029 | 0.0003 | 0.00 | 0.00 | 5.30 |
| 5030 | 0.0003 | 0.00 | 0.00 | 5.30 |
| 5031 | 0.0019 | 0.00 | 0.00 | 5.30 |
| 5032 | 0.0019 | 0.00 | 0.00 | 5.30 |
| 5033 | 0.0016 | 0.00 | 0.00 | 5.30 |
| 5034 | 0.0010 | 0.00 | 0.00 | 5.30 |
| 5035 | 0.0013 | 0.00 | 0.00 | 5.30 |
| 5036 | 0.0010 | 0.00 | 0.00 | 5.30 |
| 5037 | 0.0006 | 0.00 | 0.00 | 5.29 |
| 5038 | 0.0019 | 0.00 | 0.00 | 5.29 |
| 5039 | 0.0022 | 0.00 | 0.00 | 5.29 |
| 5040 | 0.0006 | 0.00 | 0.00 | 5.29 |
| 5041 | 0.0022 | 0.00 | 0.00 | 5.29 |
| 5042 | 0.0035 | 0.00 | 0.00 | 5.29 |
| 5043 | 0.0019 | 0.00 | 0.00 | 5.28 |
| 5044 | 0.0013 | 0.00 | 0.00 | 5.28 |
| 5045 | 0.0019 | 0.00 | 0.00 | 5.28 |
| 5046 | 0.0029 | 0.00 | 0.00 | 5.28 |
| 5047 | 0.0019 | 0.00 | 0.00 | 5.28 |
| 5048 | 0.0010 | 0.00 | 0.00 | 5.27 |
| 5049 | 0.0019 | 0.00 | 0.00 | 5.27 |
| 5050 | 0.0016 | 0.00 | 0.00 | 5.27 |
| 5051 | 0.0016 | 0.00 | 0.00 | 5.27 |
| 5052 | 0.0013 | 0.00 | 0.00 | 5.27 |
| 5053 | 0.0010 | 0.00 | 0.00 | 5.27 |
| 5054 | 0.0006 | 0.00 | 0.00 | 5.26 |
| 5055 | 0.0003 | 0.00 | 0.00 | 5.26 |
| 5056 | 0.0006 | 0.00 | 0.00 | 5.26 |
| 5057 | 0.0016 | 0.00 | 0.00 | 5.26 |
| 5058 | 0.0010 | 0.00 | 0.00 | 5.26 |
| 5059 | 0.0019 | 0.00 | 0.00 | 5.26 |
| 5060 | 0.0006 | 0.00 | 0.00 | 5.26 |
| 5061 | 0.0026 | 0.00 | 0.00 | 5.26 |
| 5062 | 0.0010 | 0.00 | 0.00 | 5.26 |
| 5063 | 0.0029 | 0.00 | 0.00 | 5.25 |

| | | | | |
|---|---|---|---|---|
| 5064 | 0.0013 | 0.00 | 0.00 | 5.25 |
| 5065 | 0.0010 | 0.00 | 0.00 | 5.25 |
| 5066 | 0.0013 | 0.00 | 0.00 | 5.25 |
| 5067 | 0.0035 | 0.00 | 0.00 | 5.25 |
| 5068 | 0.0013 | 0.00 | 0.00 | 5.24 |
| 5069 | 0.0003 | 0.00 | 0.00 | 5.24 |
| 5070 | 0.0013 | 0.00 | 0.00 | 5.24 |
| 5071 | 0.0016 | 0.00 | 0.00 | 5.24 |
| 5072 | 0.0019 | 0.00 | 0.00 | 5.24 |
| 5073 | 0.0013 | 0.00 | 0.00 | 5.24 |
| 5074 | 0.0013 | 0.00 | 0.00 | 5.24 |
| 5075 | 0.0013 | 0.00 | 0.00 | 5.24 |
| 5076 | 0.0010 | 0.00 | 0.00 | 5.23 |
| 5077 | 0.0013 | 0.00 | 0.00 | 5.23 |
| 5078 | 0.0016 | 0.00 | 0.00 | 5.23 |
| 5079 | 0.0013 | 0.00 | 0.00 | 5.23 |
| 5080 | 0.0029 | 0.00 | 0.00 | 5.23 |
| 5081 | 0.0019 | 0.00 | 0.00 | 5.23 |
| 5082 | 0.0016 | 0.00 | 0.00 | 5.22 |
| 5083 | 0.0010 | 0.00 | 0.00 | 5.22 |
| 5084 | 0.0022 | 0.00 | 0.00 | 5.22 |
| 5085 | 0.0016 | 0.00 | 0.00 | 5.22 |
| 5086 | 0.0010 | 0.00 | 0.00 | 5.22 |
| 5087 | 0.0019 | 0.00 | 0.00 | 5.22 |
| 5088 | 0.0013 | 0.00 | 0.00 | 5.22 |
| 5089 | 0.0016 | 0.00 | 0.00 | 5.21 |
| 5090 | 0.0013 | 0.00 | 0.00 | 5.21 |
| 5091 | 0.0013 | 0.00 | 0.00 | 5.21 |
| 5092 | 0.0022 | 0.00 | 0.00 | 5.21 |
| 5093 | 0.0016 | 0.00 | 0.00 | 5.21 |
| 5094 | 0.0019 | 0.00 | 0.00 | 5.21 |
| 5095 | 0.0016 | 0.00 | 0.00 | 5.20 |
| 5096 | 0.0016 | 0.00 | 0.00 | 5.20 |
| 5097 | 0.0019 | 0.00 | 0.00 | 5.20 |
| 5098 | 0.0006 | 0.00 | 0.00 | 5.20 |
| 5099 | 0.0016 | 0.00 | 0.00 | 5.20 |
| 5100 | 0.0013 | 0.00 | 0.00 | 5.20 |
| 5101 | 0.0013 | 0.00 | 0.00 | 5.20 |
| 5102 | 0.0022 | 0.00 | 0.00 | 5.19 |
| 5103 | 0.0026 | 0.00 | 0.00 | 5.19 |
| 5104 | 0.0022 | 0.00 | 0.00 | 5.19 |
| 5105 | 0.0019 | 0.00 | 0.00 | 5.19 |
| 5106 | 0.0026 | 0.00 | 0.00 | 5.18 |
| 5107 | 0.0019 | 0.00 | 0.00 | 5.18 |

| | | | | |
|---|---|---|---|---|
| 5108 | 0.0006 | 0.00 | 0.00 | 5.18 |
| 5109 | 0.0010 | 0.00 | 0.00 | 5.18 |
| 5110 | 0.0016 | 0.00 | 0.00 | 5.18 |
| 5111 | 0.0016 | 0.00 | 0.00 | 5.18 |
| 5112 | 0.0016 | 0.00 | 0.00 | 5.18 |
| 5113 | 0.0006 | 0.00 | 0.00 | 5.17 |
| 5114 | 0.0029 | 0.00 | 0.00 | 5.17 |
| 5115 | 0.0026 | 0.00 | 0.00 | 5.17 |
| 5116 | 0.0026 | 0.00 | 0.00 | 5.17 |
| 5117 | 0.0026 | 0.00 | 0.00 | 5.17 |
| 5118 | 0.0019 | 0.00 | 0.00 | 5.16 |
| 5119 | 0.0013 | 0.00 | 0.00 | 5.16 |
| 5120 | 0.0013 | 0.00 | 0.00 | 5.16 |
| 5121 | 0.0029 | 0.00 | 0.00 | 5.16 |
| 5122 | 0.0016 | 0.00 | 0.00 | 5.16 |
| 5123 | 0.0022 | 0.00 | 0.00 | 5.15 |
| 5124 | 0.0010 | 0.00 | 0.00 | 5.15 |
| 5125 | 0.0026 | 0.00 | 0.00 | 5.15 |
| 5126 | 0.0010 | 0.00 | 0.00 | 5.15 |
| 5127 | 0.0010 | 0.00 | 0.00 | 5.15 |
| 5128 | 0.0016 | 0.00 | 0.00 | 5.15 |
| 5129 | 0.0010 | 0.00 | 0.00 | 5.14 |
| 5130 | 0.0019 | 0.00 | 0.00 | 5.14 |
| 5131 | 0.0029 | 0.00 | 0.00 | 5.14 |
| 5132 | 0.0013 | 0.00 | 0.00 | 5.14 |
| 5133 | 0.0010 | 0.00 | 0.00 | 5.14 |
| 5134 | 0.0026 | 0.00 | 0.00 | 5.14 |
| 5135 | 0.0019 | 0.00 | 0.00 | 5.13 |
| 5136 | 0.0019 | 0.00 | 0.00 | 5.13 |
| 5137 | 0.0016 | 0.00 | 0.00 | 5.13 |
| 5138 | 0.0006 | 0.00 | 0.00 | 5.13 |
| 5139 | 0.0029 | 0.00 | 0.00 | 5.13 |
| 5140 | 0.0029 | 0.00 | 0.00 | 5.13 |
| 5141 | 0.0016 | 0.00 | 0.00 | 5.12 |
| 5142 | 0.0016 | 0.00 | 0.00 | 5.12 |
| 5143 | 0.0013 | 0.00 | 0.00 | 5.12 |
| 5144 | 0.0013 | 0.00 | 0.00 | 5.12 |
| 5145 | 0.0026 | 0.00 | 0.00 | 5.12 |
| 5146 | 0.0022 | 0.00 | 0.00 | 5.11 |
| 5147 | 0.0022 | 0.00 | 0.00 | 5.11 |
| 5148 | 0.0016 | 0.00 | 0.00 | 5.11 |
| 5149 | 0.0016 | 0.00 | 0.00 | 5.11 |
| 5150 | 0.0016 | 0.00 | 0.00 | 5.11 |
| 5151 | 0.0026 | 0.00 | 0.00 | 5.10 |

| | | | | |
|---|---|---|---|---|
| 5152 | 0.0032 | 0.00 | 0.00 | 5.10 |
| 5153 | 0.0006 | 0.00 | 0.00 | 5.10 |
| 5154 | 0.0026 | 0.00 | 0.00 | 5.10 |
| 5155 | 0.0029 | 0.00 | 0.00 | 5.10 |
| 5156 | 0.0029 | 0.00 | 0.00 | 5.09 |
| 5157 | 0.0010 | 0.00 | 0.00 | 5.09 |
| 5158 | 0.0006 | 0.00 | 0.00 | 5.09 |
| 5159 | 0.0010 | 0.00 | 0.00 | 5.09 |
| 5160 | 0.0029 | 0.00 | 0.00 | 5.09 |
| 5161 | 0.0016 | 0.00 | 0.00 | 5.08 |
| 5162 | 0.0019 | 0.00 | 0.00 | 5.08 |
| 5163 | 0.0026 | 0.00 | 0.00 | 5.08 |
| 5164 | 0.0022 | 0.00 | 0.00 | 5.08 |
| 5165 | 0.0029 | 0.00 | 0.00 | 5.08 |
| 5166 | 0.0026 | 0.00 | 0.00 | 5.07 |
| 5167 | 0.0032 | 0.00 | 0.00 | 5.07 |
| 5168 | 0.0026 | 0.00 | 0.00 | 5.07 |
| 5169 | 0.0019 | 0.00 | 0.00 | 5.06 |
| 5170 | 0.0016 | 0.00 | 0.00 | 5.06 |
| 5171 | 0.0016 | 0.00 | 0.00 | 5.06 |
| 5172 | 0.0016 | 0.00 | 0.00 | 5.06 |
| 5173 | 0.0026 | 0.00 | 0.00 | 5.06 |
| 5174 | 0.0013 | 0.00 | 0.00 | 5.06 |
| 5175 | 0.0022 | 0.00 | 0.00 | 5.05 |
| 5176 | 0.0013 | 0.00 | 0.00 | 5.05 |
| 5177 | 0.0054 | 0.00 | 0.00 | 5.05 |
| 5178 | 0.0038 | 0.00 | 0.00 | 5.05 |
| 5179 | 0.0032 | 0.00 | 0.00 | 5.04 |
| 5180 | 0.0016 | 0.00 | 0.00 | 5.04 |
| 5181 | 0.0016 | 0.00 | 0.00 | 5.04 |
| 5182 | 0.0013 | 0.00 | 0.00 | 5.03 |
| 5183 | 0.0013 | 0.00 | 0.00 | 5.03 |
| 5184 | 0.0013 | 0.00 | 0.00 | 5.03 |
| 5185 | 0.0013 | 0.00 | 0.00 | 5.03 |
| 5186 | 0.0010 | 0.00 | 0.00 | 5.03 |
| 5187 | 0.0016 | 0.00 | 0.00 | 5.03 |
| 5188 | 0.0016 | 0.00 | 0.00 | 5.03 |
| 5189 | 0.0010 | 0.00 | 0.00 | 5.03 |
| 5190 | 0.0022 | 0.00 | 0.00 | 5.02 |
| 5191 | 0.0010 | 0.00 | 0.00 | 5.02 |
| 5192 | 0.0022 | 0.00 | 0.00 | 5.02 |
| 5193 | 0.0006 | 0.00 | 0.00 | 5.02 |
| 5194 | 0.0013 | 0.00 | 0.00 | 5.02 |
| 5195 | 0.0010 | 0.00 | 0.00 | 5.02 |

| | | | | |
|---|---|---|---|---|
| 5196 | 0.0013 | 0.00 | 0.00 | 5.02 |
| 5197 | 0.0022 | 0.00 | 0.00 | 5.02 |
| 5198 | 0.0013 | 0.00 | 0.00 | 5.01 |
| 5199 | 0.0010 | 0.00 | 0.00 | 5.01 |
| 5200 | 0.0006 | 0.00 | 0.00 | 5.01 |
| 5201 | 0.0010 | 0.00 | 0.00 | 5.01 |
| 5202 | 0.0013 | 0.00 | 0.00 | 5.01 |
| 5203 | 0.0010 | 0.00 | 0.00 | 5.01 |
| 5204 | 0.0016 | 0.00 | 0.00 | 5.01 |
| 5205 | 0.0016 | 0.00 | 0.00 | 5.01 |
| 5206 | 0.0026 | 0.00 | 0.00 | 5.00 |
| 5207 | 0.0016 | 0.00 | 0.00 | 5.00 |
| 5208 | 0.0010 | 0.00 | 0.00 | 5.00 |
| 5209 | 0.0029 | 0.00 | 0.00 | 5.00 |
| 5210 | 0.0013 | 0.00 | 0.00 | 5.00 |
| 5211 | 0.0013 | 0.00 | 0.00 | 4.99 |
| 5212 | 0.0013 | 0.00 | 0.00 | 4.99 |
| 5213 | 0.0013 | 0.00 | 0.00 | 4.99 |
| 5214 | 0.0026 | 0.00 | 0.00 | 4.99 |
| 5215 | 0.0016 | 0.00 | 0.00 | 4.99 |
| 5216 | 0.0016 | 0.00 | 0.00 | 4.99 |
| 5217 | 0.0019 | 0.00 | 0.00 | 4.98 |
| 5218 | 0.0016 | 0.00 | 0.00 | 4.98 |
| 5219 | 0.0003 | 0.00 | 0.00 | 4.98 |
| 5220 | 0.0010 | 0.00 | 0.00 | 4.98 |
| 5221 | 0.0019 | 0.00 | 0.00 | 4.98 |
| 5222 | 0.0013 | 0.00 | 0.00 | 4.98 |
| 5223 | 0.0006 | 0.00 | 0.00 | 4.98 |
| 5224 | 0.0016 | 0.00 | 0.00 | 4.98 |
| 5225 | 0.0013 | 0.00 | 0.00 | 4.97 |
| 5226 | 0.0026 | 0.00 | 0.00 | 4.97 |
| 5227 | 0.0006 | 0.00 | 0.00 | 4.97 |
| 5228 | 0.0013 | 0.00 | 0.00 | 4.97 |
| 5229 | 0.0013 | 0.00 | 0.00 | 4.97 |
| 5230 | 0.0003 | 0.00 | 0.00 | 4.97 |
| 5231 | 0.0016 | 0.00 | 0.00 | 4.97 |
| 5232 | 0.0013 | 0.00 | 0.00 | 4.97 |
| 5233 | 0.0003 | 0.00 | 0.00 | 4.96 |
| 5234 | 0.0010 | 0.00 | 0.00 | 4.96 |
| 5235 | 0.0026 | 0.00 | 0.00 | 4.96 |
| 5236 | 0.0010 | 0.00 | 0.00 | 4.96 |
| 5237 | 0.0010 | 0.00 | 0.00 | 4.96 |
| 5238 | 0.0016 | 0.00 | 0.00 | 4.96 |
| 5239 | 0.0035 | 0.00 | 0.00 | 4.96 |

| | | | | |
|---|---|---|---|---|
| 5240 | 0.0003 | 0.00 | 0.00 | 4.95 |
| 5241 | 0.0026 | 0.00 | 0.00 | 4.95 |
| 5242 | 0.0013 | 0.00 | 0.00 | 4.95 |
| 5243 | 0.0016 | 0.00 | 0.00 | 4.95 |
| 5244 | 0.0010 | 0.00 | 0.00 | 4.95 |
| 5245 | 0.0010 | 0.00 | 0.00 | 4.95 |
| 5246 | 0.0022 | 0.00 | 0.00 | 4.95 |
| 5247 | 0.0006 | 0.00 | 0.00 | 4.94 |
| 5248 | 0.0013 | 0.00 | 0.00 | 4.94 |
| 5249 | 0.0010 | 0.00 | 0.00 | 4.94 |
| 5250 | 0.0003 | 0.00 | 0.00 | 4.94 |
| 5251 | 0.0013 | 0.00 | 0.00 | 4.94 |
| 5252 | 0.0032 | 0.00 | 0.00 | 4.94 |
| 5253 | 0.0022 | 0.00 | 0.00 | 4.94 |
| 5254 | 0.0013 | 0.00 | 0.00 | 4.93 |
| 5255 | 0.0010 | 0.00 | 0.00 | 4.93 |
| 5256 | 0.0006 | 0.00 | 0.00 | 4.93 |
| 5257 | 0.0026 | 0.00 | 0.00 | 4.93 |
| 5258 | 0.0022 | 0.00 | 0.00 | 4.93 |
| 5259 | 0.0019 | 0.00 | 0.00 | 4.93 |
| 5260 | 0.0013 | 0.00 | 0.00 | 4.92 |
| 5261 | 0.0006 | 0.00 | 0.00 | 4.92 |
| 5262 | 0.0013 | 0.00 | 0.00 | 4.92 |
| 5263 | 0.0019 | 0.00 | 0.00 | 4.92 |
| 5264 | 0.0019 | 0.00 | 0.00 | 4.92 |
| 5265 | 0.0010 | 0.00 | 0.00 | 4.92 |
| 5266 | 0.0029 | 0.00 | 0.00 | 4.92 |
| 5267 | 0.0016 | 0.00 | 0.00 | 4.91 |
| 5268 | 0.0016 | 0.00 | 0.00 | 4.91 |
| 5269 | 0.0022 | 0.00 | 0.00 | 4.91 |
| 5270 | 0.0003 | 0.00 | 0.00 | 4.91 |
| 5271 | 0.0026 | 0.00 | 0.00 | 4.91 |
| 5272 | 0.0035 | 0.00 | 0.00 | 4.90 |
| 5273 | 0.0010 | 0.00 | 0.00 | 4.90 |
| 5274 | 0.0026 | 0.00 | 0.00 | 4.90 |
| 5275 | 0.0019 | 0.00 | 0.00 | 4.90 |
| 5276 | 0.0013 | 0.00 | 0.00 | 4.90 |
| 5277 | 0.0026 | 0.00 | 0.00 | 4.89 |
| 5278 | 0.0016 | 0.00 | 0.00 | 4.89 |
| 5279 | 0.0019 | 0.00 | 0.00 | 4.89 |
| 5280 | 0.0029 | 0.00 | 0.00 | 4.89 |
| 5281 | 0.0029 | 0.00 | 0.00 | 4.89 |
| 5282 | 0.0022 | 0.00 | 0.00 | 4.88 |
| 5283 | 0.0003 | 0.00 | 0.00 | 4.88 |

| | | | | |
|---|---|---|---|---|
| 5284 | 0.0013 | 0.00 | 0.00 | 4.88 |
| 5285 | 0.0006 | 0.00 | 0.00 | 4.88 |
| 5286 | 0.0010 | 0.00 | 0.00 | 4.88 |
| 5287 | 0.0019 | 0.00 | 0.00 | 4.88 |
| 5288 | 0.0026 | 0.00 | 0.00 | 4.88 |
| 5289 | 0.0016 | 0.00 | 0.00 | 4.87 |
| 5290 | 0.0026 | 0.00 | 0.00 | 4.87 |
| 5291 | 0.0019 | 0.00 | 0.00 | 4.87 |
| 5292 | 0.0006 | 0.00 | 0.00 | 4.87 |
| 5293 | 0.0013 | 0.00 | 0.00 | 4.87 |
| 5294 | 0.0019 | 0.00 | 0.00 | 4.86 |
| 5295 | 0.0013 | 0.00 | 0.00 | 4.86 |
| 5296 | 0.0016 | 0.00 | 0.00 | 4.86 |
| 5297 | 0.0013 | 0.00 | 0.00 | 4.86 |
| 5298 | 0.0029 | 0.00 | 0.00 | 4.86 |
| 5299 | 0.0042 | 0.00 | 0.00 | 4.86 |
| 5300 | 0.0042 | 0.00 | 0.00 | 4.85 |
| 5301 | 0.0022 | 0.00 | 0.00 | 4.85 |
| 5302 | 0.0013 | 0.00 | 0.00 | 4.84 |
| 5303 | 0.0013 | 0.00 | 0.00 | 4.84 |
| 5304 | 0.0016 | 0.00 | 0.00 | 4.84 |
| 5305 | 0.0022 | 0.00 | 0.00 | 4.84 |
| 5306 | 0.0029 | 0.00 | 0.00 | 4.84 |
| 5307 | 0.0010 | 0.00 | 0.00 | 4.84 |
| 5308 | 0.0010 | 0.00 | 0.00 | 4.83 |
| 5309 | 0.0022 | 0.00 | 0.00 | 4.83 |
| 5310 | 0.0016 | 0.00 | 0.00 | 4.83 |
| 5311 | 0.0010 | 0.00 | 0.00 | 4.83 |
| 5312 | 0.0016 | 0.00 | 0.00 | 4.83 |
| 5313 | 0.0013 | 0.00 | 0.00 | 4.83 |
| 5314 | 0.0013 | 0.00 | 0.00 | 4.83 |
| 5315 | 0.0022 | 0.00 | 0.00 | 4.82 |
| 5316 | 0.0022 | 0.00 | 0.00 | 4.82 |
| 5317 | 0.0032 | 0.00 | 0.00 | 4.82 |
| 5318 | 0.0010 | 0.00 | 0.00 | 4.82 |
| 5319 | 0.0019 | 0.00 | 0.00 | 4.82 |
| 5320 | 0.0006 | 0.00 | 0.00 | 4.81 |
| 5321 | 0.0019 | 0.00 | 0.00 | 4.81 |
| 5322 | 0.0016 | 0.00 | 0.00 | 4.81 |
| 5323 | 0.0019 | 0.00 | 0.00 | 4.81 |
| 5324 | 0.0019 | 0.00 | 0.00 | 4.81 |
| 5325 | 0.0029 | 0.00 | 0.00 | 4.81 |
| 5326 | 0.0010 | 0.00 | 0.00 | 4.80 |
| 5327 | 0.0016 | 0.00 | 0.00 | 4.80 |

| | | | | |
|---|---|---|---|---|
| 5328 | 0.0006 | 0.00 | 0.00 | 4.80 |
| 5329 | 0.0013 | 0.00 | 0.00 | 4.80 |
| 5330 | 0.0016 | 0.00 | 0.00 | 4.80 |
| 5331 | 0.0019 | 0.00 | 0.00 | 4.80 |
| 5332 | 0.0022 | 0.00 | 0.00 | 4.80 |
| 5333 | 0.0010 | 0.00 | 0.00 | 4.79 |
| 5334 | 0.0016 | 0.00 | 0.00 | 4.79 |
| 5335 | 0.0022 | 0.00 | 0.00 | 4.79 |
| 5336 | 0.0010 | 0.00 | 0.00 | 4.79 |
| 5337 | 0.0022 | 0.00 | 0.00 | 4.79 |
| 5338 | 0.0026 | 0.00 | 0.00 | 4.78 |
| 5339 | 0.0016 | 0.00 | 0.00 | 4.78 |
| 5340 | 0.0022 | 0.00 | 0.00 | 4.78 |
| 5341 | 0.0003 | 0.00 | 0.00 | 4.78 |
| 5342 | 0.0016 | 0.00 | 0.00 | 4.78 |
| 5343 | 0.0006 | 0.00 | 0.00 | 4.78 |
| 5344 | 0.0013 | 0.00 | 0.00 | 4.78 |
| 5345 | 0.0016 | 0.00 | 0.00 | 4.77 |
| 5346 | 0.0038 | 0.00 | 0.00 | 4.77 |
| 5347 | 0.0003 | 0.00 | 0.00 | 4.77 |
| 5348 | 0.0013 | 0.00 | 0.00 | 4.77 |
| 5349 | 0.0026 | 0.00 | 0.00 | 4.77 |
| 5350 | 0.0026 | 0.00 | 0.00 | 4.77 |
| 5351 | 0.0003 | 0.00 | 0.00 | 4.76 |
| 5352 | 0.0022 | 0.00 | 0.00 | 4.76 |
| 5353 | 0.0013 | 0.00 | 0.00 | 4.76 |
| 5354 | 0.0006 | 0.00 | 0.00 | 4.76 |
| 5355 | 0.0003 | 0.00 | 0.00 | 4.76 |
| 5356 | 0.0013 | 0.00 | 0.00 | 4.76 |
| 5357 | 0.0013 | 0.00 | 0.00 | 4.76 |
| 5358 | 0.0013 | 0.00 | 0.00 | 4.76 |
| 5359 | 0.0013 | 0.00 | 0.00 | 4.75 |
| 5360 | 0.0016 | 0.00 | 0.00 | 4.75 |
| 5361 | 0.0019 | 0.00 | 0.00 | 4.75 |
| 5362 | 0.0022 | 0.00 | 0.00 | 4.75 |
| 5363 | 0.0022 | 0.00 | 0.00 | 4.75 |
| 5364 | 0.0013 | 0.00 | 0.00 | 4.74 |
| 5365 | 0.0019 | 0.00 | 0.00 | 4.74 |
| 5366 | 0.0013 | 0.00 | 0.00 | 4.74 |
| 5367 | 0.0013 | 0.00 | 0.00 | 4.74 |
| 5368 | 0.0016 | 0.00 | 0.00 | 4.74 |
| 5369 | 0.0019 | 0.00 | 0.00 | 4.74 |
| 5370 | 0.0029 | 0.00 | 0.00 | 4.74 |
| 5371 | 0.0019 | 0.00 | 0.00 | 4.73 |

| | | | | |
|---|---|---|---|---|
| 5372 | 0.0032 | 0.00 | 0.00 | 4.73 |
| 5373 | 0.0013 | 0.00 | 0.00 | 4.73 |
| 5374 | 0.0016 | 0.00 | 0.00 | 4.73 |
| 5375 | 0.0016 | 0.00 | 0.00 | 4.72 |
| 5376 | 0.0019 | 0.00 | 0.00 | 4.72 |
| 5377 | 0.0022 | 0.00 | 0.00 | 4.72 |
| 5378 | 0.0016 | 0.00 | 0.00 | 4.72 |
| 5379 | 0.0016 | 0.00 | 0.00 | 4.72 |
| 5380 | 0.0019 | 0.00 | 0.00 | 4.72 |
| 5381 | 0.0010 | 0.00 | 0.00 | 4.71 |
| 5382 | 0.0013 | 0.00 | 0.00 | 4.71 |
| 5383 | 0.0013 | 0.00 | 0.00 | 4.71 |
| 5384 | 0.0013 | 0.00 | 0.00 | 4.71 |
| 5385 | 0.0010 | 0.00 | 0.00 | 4.71 |
| 5386 | 0.0016 | 0.00 | 0.00 | 4.71 |
| 5387 | 0.0026 | 0.00 | 0.00 | 4.71 |
| 5388 | 0.0022 | 0.00 | 0.00 | 4.70 |
| 5389 | 0.0016 | 0.00 | 0.00 | 4.70 |
| 5390 | 0.0026 | 0.00 | 0.00 | 4.70 |
| 5391 | 0.0019 | 0.00 | 0.00 | 4.70 |
| 5392 | 0.0022 | 0.00 | 0.00 | 4.70 |
| 5393 | 0.0016 | 0.00 | 0.00 | 4.69 |
| 5394 | 0.0016 | 0.00 | 0.00 | 4.69 |
| 5395 | 0.0022 | 0.00 | 0.00 | 4.69 |
| 5396 | 0.0013 | 0.00 | 0.00 | 4.69 |
| 5397 | 0.0006 | 0.00 | 0.00 | 4.69 |
| 5398 | 0.0016 | 0.00 | 0.00 | 4.69 |
| 5399 | 0.0016 | 0.00 | 0.00 | 4.68 |
| 5400 | 0.0003 | 0.00 | 0.00 | 4.68 |
| 5401 | 0.0022 | 0.00 | 0.00 | 4.68 |
| 5402 | 0.0013 | 0.00 | 0.00 | 4.68 |
| 5403 | 0.0029 | 0.00 | 0.00 | 4.68 |
| 5404 | 0.0006 | 0.00 | 0.00 | 4.68 |
| 5405 | 0.0016 | 0.00 | 0.00 | 4.68 |
| 5406 | 0.0013 | 0.00 | 0.00 | 4.67 |
| 5407 | 0.0016 | 0.00 | 0.00 | 4.67 |
| 5408 | 0.0016 | 0.00 | 0.00 | 4.67 |
| 5409 | 0.0019 | 0.00 | 0.00 | 4.67 |
| 5410 | 0.0038 | 0.00 | 0.00 | 4.67 |
| 5412 | 0.0019 | 0.00 | 0.00 | 4.66 |
| 5413 | 0.0010 | 0.00 | 0.00 | 4.66 |
| 5414 | 0.0026 | 0.00 | 0.00 | 4.66 |
| 5415 | 0.0016 | 0.00 | 0.00 | 4.66 |
| 5416 | 0.0022 | 0.00 | 0.00 | 4.66 |

| | | | | |
|---|---|---|---|---|
| 5417 | 0.0026 | 0.00 | 0.00 | 4.65 |
| 5418 | 0.0016 | 0.00 | 0.00 | 4.65 |
| 5419 | 0.0016 | 0.00 | 0.00 | 4.65 |
| 5420 | 0.0016 | 0.00 | 0.00 | 4.65 |
| 5421 | 0.0022 | 0.00 | 0.00 | 4.65 |
| 5422 | 0.0006 | 0.00 | 0.00 | 4.64 |
| 5423 | 0.0013 | 0.00 | 0.00 | 4.64 |
| 5424 | 0.0022 | 0.00 | 0.00 | 4.64 |
| 5425 | 0.0010 | 0.00 | 0.00 | 4.64 |
| 5426 | 0.0006 | 0.00 | 0.00 | 4.64 |
| 5427 | 0.0013 | 0.00 | 0.00 | 4.64 |
| 5428 | 0.0029 | 0.00 | 0.00 | 4.64 |
| 5429 | 0.0013 | 0.00 | 0.00 | 4.63 |
| 5430 | 0.0006 | 0.00 | 0.00 | 4.63 |
| 5431 | 0.0010 | 0.00 | 0.00 | 4.63 |
| 5432 | 0.0022 | 0.00 | 0.00 | 4.63 |
| 5433 | 0.0013 | 0.00 | 0.00 | 4.63 |
| 5434 | 0.0035 | 0.00 | 0.00 | 4.63 |
| 5435 | 0.0010 | 0.00 | 0.00 | 4.62 |
| 5436 | 0.0022 | 0.00 | 0.00 | 4.62 |
| 5437 | 0.0006 | 0.00 | 0.00 | 4.62 |
| 5438 | 0.0032 | 0.00 | 0.00 | 4.62 |
| 5439 | 0.0006 | 0.00 | 0.00 | 4.62 |
| 5440 | 0.0019 | 0.00 | 0.00 | 4.62 |
| 5441 | 0.0003 | 0.00 | 0.00 | 4.61 |
| 5442 | 0.0016 | 0.00 | 0.00 | 4.61 |
| 5443 | 0.0016 | 0.00 | 0.00 | 4.61 |
| 5444 | 0.0006 | 0.00 | 0.00 | 4.61 |
| 5445 | 0.0016 | 0.00 | 0.00 | 4.61 |
| 5446 | 0.0016 | 0.00 | 0.00 | 4.61 |
| 5447 | 0.0019 | 0.00 | 0.00 | 4.61 |
| 5448 | 0.0016 | 0.00 | 0.00 | 4.61 |
| 5449 | 0.0006 | 0.00 | 0.00 | 4.60 |
| 5450 | 0.0013 | 0.00 | 0.00 | 4.60 |
| 5451 | 0.0006 | 0.00 | 0.00 | 4.60 |
| 5452 | 0.0013 | 0.00 | 0.00 | 4.60 |
| 5453 | 0.0019 | 0.00 | 0.00 | 4.60 |
| 5454 | 0.0006 | 0.00 | 0.00 | 4.60 |
| 5455 | 0.0026 | 0.00 | 0.00 | 4.60 |
| 5456 | 0.0013 | 0.00 | 0.00 | 4.59 |
| 5457 | 0.0019 | 0.00 | 0.00 | 4.59 |
| 5458 | 0.0019 | 0.00 | 0.00 | 4.59 |
| 5459 | 0.0010 | 0.00 | 0.00 | 4.59 |
| 5460 | 0.0006 | 0.00 | 0.00 | 4.59 |

| | | | | |
|---|---|---|---|---|
| 5461 | 0.0016 | 0.00 | 0.00 | 4.59 |
| 5462 | 0.0019 | 0.00 | 0.00 | 4.59 |
| 5463 | 0.0016 | 0.00 | 0.00 | 4.58 |
| 5464 | 0.0019 | 0.00 | 0.00 | 4.58 |
| 5465 | 0.0010 | 0.00 | 0.00 | 4.58 |
| 5466 | 0.0019 | 0.00 | 0.00 | 4.58 |
| 5467 | 0.0006 | 0.00 | 0.00 | 4.58 |
| 5468 | 0.0016 | 0.00 | 0.00 | 4.58 |
| 5469 | 0.0019 | 0.00 | 0.00 | 4.58 |
| 5470 | 0.0013 | 0.00 | 0.00 | 4.57 |
| 5471 | 0.0003 | 0.00 | 0.00 | 4.57 |
| 5472 | 0.0003 | 0.00 | 0.00 | 4.57 |
| 5473 | 0.0019 | 0.00 | 0.00 | 4.57 |
| 5474 | 0.0019 | 0.00 | 0.00 | 4.57 |
| 5475 | 0.0006 | 0.00 | 0.00 | 4.57 |
| 5476 | 0.0016 | 0.00 | 0.00 | 4.57 |
| 5477 | 0.0006 | 0.00 | 0.00 | 4.57 |
| 5478 | 0.0016 | 0.00 | 0.00 | 4.57 |
| 5479 | 0.0026 | 0.00 | 0.00 | 4.56 |
| 5480 | 0.0010 | 0.00 | 0.00 | 4.56 |
| 5481 | 0.0019 | 0.00 | 0.00 | 4.56 |
| 5482 | 0.0029 | 0.00 | 0.00 | 4.56 |
| 5483 | 0.0019 | 0.00 | 0.00 | 4.56 |
| 5484 | 0.0019 | 0.00 | 0.00 | 4.55 |
| 5485 | 0.0013 | 0.00 | 0.00 | 4.55 |
| 5486 | 0.0010 | 0.00 | 0.00 | 4.55 |
| 5487 | 0.0019 | 0.00 | 0.00 | 4.55 |
| 5488 | 0.0022 | 0.00 | 0.00 | 4.55 |
| 5489 | 0.0026 | 0.00 | 0.00 | 4.55 |
| 5490 | 0.0006 | 0.00 | 0.00 | 4.54 |
| 5491 | 0.0019 | 0.00 | 0.00 | 4.54 |
| 5492 | 0.0013 | 0.00 | 0.00 | 4.54 |
| 5493 | 0.0016 | 0.00 | 0.00 | 4.54 |
| 5494 | 0.0013 | 0.00 | 0.00 | 4.54 |
| 5495 | 0.0022 | 0.00 | 0.00 | 4.54 |
| 5496 | 0.0013 | 0.00 | 0.00 | 4.53 |
| 5497 | 0.0032 | 0.00 | 0.00 | 4.53 |
| 5498 | 0.0010 | 0.00 | 0.00 | 4.53 |
| 5499 | 0.0019 | 0.00 | 0.00 | 4.53 |
| 5500 | 0.0013 | 0.00 | 0.00 | 4.53 |
| 5501 | 0.0016 | 0.00 | 0.00 | 4.53 |
| 5502 | 0.0019 | 0.00 | 0.00 | 4.52 |
| 5503 | 0.0003 | 0.00 | 0.00 | 4.52 |
| 5504 | 0.0006 | 0.00 | 0.00 | 4.52 |

| | | | | |
|---|---|---|---|---|
| 5505 | 0.0006 | 0.00 | 0.00 | 4.52 |
| 5506 | 0.0013 | 0.00 | 0.00 | 4.52 |
| 5507 | 0.0010 | 0.00 | 0.00 | 4.52 |
| 5508 | 0.0022 | 0.00 | 0.00 | 4.52 |
| 5509 | 0.0003 | 0.00 | 0.00 | 4.52 |
| 5510 | 0.0010 | 0.00 | 0.00 | 4.52 |
| 5511 | 0.0019 | 0.00 | 0.00 | 4.51 |
| 5512 | 0.0013 | 0.00 | 0.00 | 4.51 |
| 5513 | 0.0006 | 0.00 | 0.00 | 4.51 |
| 5514 | 0.0006 | 0.00 | 0.00 | 4.51 |
| 5515 | 0.0019 | 0.00 | 0.00 | 4.51 |
| 5516 | 0.0016 | 0.00 | 0.00 | 4.51 |
| 5517 | 0.0010 | 0.00 | 0.00 | 4.51 |
| 5518 | 0.0013 | 0.00 | 0.00 | 4.51 |
| 5519 | 0.0019 | 0.00 | 0.00 | 4.50 |
| 5520 | 0.0003 | 0.00 | 0.00 | 4.50 |
| 5521 | 0.0010 | 0.00 | 0.00 | 4.50 |
| 5522 | 0.0016 | 0.00 | 0.00 | 4.50 |
| 5523 | 0.0006 | 0.00 | 0.00 | 4.50 |
| 5524 | 0.0019 | 0.00 | 0.00 | 4.50 |
| 5525 | 0.0022 | 0.00 | 0.00 | 4.50 |
| 5526 | 0.0029 | 0.00 | 0.00 | 4.49 |
| 5527 | 0.0006 | 0.00 | 0.00 | 4.49 |
| 5528 | 0.0029 | 0.00 | 0.00 | 4.49 |
| 5529 | 0.0010 | 0.00 | 0.00 | 4.49 |
| 5530 | 0.0016 | 0.00 | 0.00 | 4.49 |
| 5531 | 0.0013 | 0.00 | 0.00 | 4.49 |
| 5532 | 0.0010 | 0.00 | 0.00 | 4.48 |
| 5533 | 0.0016 | 0.00 | 0.00 | 4.48 |
| 5534 | 0.0019 | 0.00 | 0.00 | 4.48 |
| 5535 | 0.0003 | 0.00 | 0.00 | 4.48 |
| 5536 | 0.0016 | 0.00 | 0.00 | 4.48 |
| 5537 | 0.0013 | 0.00 | 0.00 | 4.48 |
| 5538 | 0.0010 | 0.00 | 0.00 | 4.48 |
| 5539 | 0.0003 | 0.00 | 0.00 | 4.48 |
| 5540 | 0.0006 | 0.00 | 0.00 | 4.48 |
| 5541 | 0.0016 | 0.00 | 0.00 | 4.47 |
| 5542 | 0.0013 | 0.00 | 0.00 | 4.47 |
| 5543 | 0.0019 | 0.00 | 0.00 | 4.47 |
| 5544 | 0.0013 | 0.00 | 0.00 | 4.47 |
| 5545 | 0.0026 | 0.00 | 0.00 | 4.47 |
| 5546 | 0.0013 | 0.00 | 0.00 | 4.47 |
| 5547 | 0.0019 | 0.00 | 0.00 | 4.46 |
| 5548 | 0.0006 | 0.00 | 0.00 | 4.46 |

| | | | | |
|---|---|---|---|---|
| 5549 | 0.0013 | 0.00 | 0.00 | 4.46 |
| 5550 | 0.0010 | 0.00 | 0.00 | 4.46 |
| 5551 | 0.0010 | 0.00 | 0.00 | 4.46 |
| 5552 | 0.0019 | 0.00 | 0.00 | 4.46 |
| 5553 | 0.0022 | 0.00 | 0.00 | 4.46 |
| 5554 | 0.0006 | 0.00 | 0.00 | 4.45 |
| 5555 | 0.0019 | 0.00 | 0.00 | 4.45 |
| 5556 | 0.0016 | 0.00 | 0.00 | 4.45 |
| 5557 | 0.0026 | 0.00 | 0.00 | 4.45 |
| 5558 | 0.0026 | 0.00 | 0.00 | 4.45 |
| 5559 | 0.0006 | 0.00 | 0.00 | 4.45 |
| 5560 | 0.0010 | 0.00 | 0.00 | 4.44 |
| 5561 | 0.0013 | 0.00 | 0.00 | 4.44 |
| 5562 | 0.0016 | 0.00 | 0.00 | 4.44 |
| 5563 | 0.0006 | 0.00 | 0.00 | 4.44 |
| 5564 | 0.0022 | 0.00 | 0.00 | 4.44 |
| 5565 | 0.0013 | 0.00 | 0.00 | 4.44 |
| 5566 | 0.0006 | 0.00 | 0.00 | 4.44 |
| 5567 | 0.0013 | 0.00 | 0.00 | 4.44 |
| 5568 | 0.0013 | 0.00 | 0.00 | 4.43 |
| 5569 | 0.0013 | 0.00 | 0.00 | 4.43 |
| 5570 | 0.0013 | 0.00 | 0.00 | 4.43 |
| 5571 | 0.0010 | 0.00 | 0.00 | 4.43 |
| 5572 | 0.0022 | 0.00 | 0.00 | 4.43 |
| 5573 | 0.0016 | 0.00 | 0.00 | 4.43 |
| 5574 | 0.0013 | 0.00 | 0.00 | 4.43 |
| 5575 | 0.0019 | 0.00 | 0.00 | 4.42 |
| 5576 | 0.0010 | 0.00 | 0.00 | 4.42 |
| 5577 | 0.0013 | 0.00 | 0.00 | 4.42 |
| 5578 | 0.0032 | 0.00 | 0.00 | 4.42 |
| 5579 | 0.0016 | 0.00 | 0.00 | 4.42 |
| 5580 | 0.0013 | 0.00 | 0.00 | 4.42 |
| 5581 | 0.0019 | 0.00 | 0.00 | 4.41 |
| 5582 | 0.0006 | 0.00 | 0.00 | 4.41 |
| 5583 | 0.0042 | 0.00 | 0.00 | 4.41 |
| 5584 | 0.0026 | 0.00 | 0.00 | 4.41 |
| 5585 | 0.0006 | 0.00 | 0.00 | 4.41 |
| 5586 | 0.0019 | 0.00 | 0.00 | 4.40 |
| 5587 | 0.0013 | 0.00 | 0.00 | 4.40 |
| 5588 | 0.0019 | 0.00 | 0.00 | 4.40 |
| 5589 | 0.0022 | 0.00 | 0.00 | 4.40 |
| 5590 | 0.0016 | 0.00 | 0.00 | 4.40 |
| 5591 | 0.0013 | 0.00 | 0.00 | 4.40 |
| 5592 | 0.0032 | 0.00 | 0.00 | 4.39 |

| | | | | |
|---|---|---|---|---|
| 5593 | 0.0016 | 0.00 | 0.00 | 4.39 |
| 5594 | 0.0013 | 0.00 | 0.00 | 4.39 |
| 5595 | 0.0006 | 0.00 | 0.00 | 4.39 |
| 5596 | 0.0022 | 0.00 | 0.00 | 4.39 |
| 5597 | 0.0026 | 0.00 | 0.00 | 4.39 |
| 5598 | 0.0019 | 0.00 | 0.00 | 4.38 |
| 5599 | 0.0026 | 0.00 | 0.00 | 4.38 |
| 5600 | 0.0016 | 0.00 | 0.00 | 4.38 |
| 5601 | 0.0013 | 0.00 | 0.00 | 4.38 |
| 5602 | 0.0013 | 0.00 | 0.00 | 4.38 |
| 5603 | 0.0019 | 0.00 | 0.00 | 4.37 |
| 5604 | 0.0029 | 0.00 | 0.00 | 4.37 |
| 5605 | 0.0035 | 0.00 | 0.00 | 4.37 |
| 5606 | 0.0016 | 0.00 | 0.00 | 4.37 |
| 5607 | 0.0016 | 0.00 | 0.00 | 4.36 |
| 5608 | 0.0022 | 0.00 | 0.00 | 4.36 |
| 5609 | 0.0019 | 0.00 | 0.00 | 4.36 |
| 5610 | 0.0035 | 0.00 | 0.00 | 4.36 |
| 5611 | 0.0010 | 0.00 | 0.00 | 4.35 |
| 5612 | 0.0006 | 0.00 | 0.00 | 4.35 |
| 5613 | 0.0022 | 0.00 | 0.00 | 4.35 |
| 5614 | 0.0016 | 0.00 | 0.00 | 4.35 |
| 5615 | 0.0013 | 0.00 | 0.00 | 4.35 |
| 5616 | 0.0022 | 0.00 | 0.00 | 4.35 |
| 5617 | 0.0019 | 0.00 | 0.00 | 4.35 |
| 5618 | 0.0019 | 0.00 | 0.00 | 4.34 |
| 5619 | 0.0022 | 0.00 | 0.00 | 4.34 |
| 5620 | 0.0013 | 0.00 | 0.00 | 4.34 |
| 5621 | 0.0029 | 0.00 | 0.00 | 4.34 |
| 5622 | 0.0006 | 0.00 | 0.00 | 4.34 |
| 5623 | 0.0022 | 0.00 | 0.00 | 4.33 |
| 5624 | 0.0016 | 0.00 | 0.00 | 4.33 |
| 5625 | 0.0026 | 0.00 | 0.00 | 4.33 |
| 5626 | 0.0013 | 0.00 | 0.00 | 4.33 |
| 5627 | 0.0022 | 0.00 | 0.00 | 4.33 |
| 5628 | 0.0013 | 0.00 | 0.00 | 4.33 |
| 5629 | 0.0019 | 0.00 | 0.00 | 4.32 |
| 5630 | 0.0006 | 0.00 | 0.00 | 4.32 |
| 5631 | 0.0006 | 0.00 | 0.00 | 4.32 |
| 5632 | 0.0006 | 0.00 | 0.00 | 4.32 |
| 5633 | 0.0003 | 0.00 | 0.00 | 4.32 |
| 5634 | 0.0016 | 0.00 | 0.00 | 4.32 |
| 5635 | 0.0022 | 0.00 | 0.00 | 4.32 |
| 5636 | 0.0019 | 0.00 | 0.00 | 4.32 |

| | | | | |
|---|---|---|---|---|
| 5637 | 0.0003 | 0.00 | 0.00 | 4.31 |
| 5638 | 0.0013 | 0.00 | 0.00 | 4.31 |
| 5639 | 0.0016 | 0.00 | 0.00 | 4.31 |
| 5640 | 0.0019 | 0.00 | 0.00 | 4.31 |
| 5641 | 0.0013 | 0.00 | 0.00 | 4.31 |
| 5642 | 0.0016 | 0.00 | 0.00 | 4.31 |
| 5643 | 0.0013 | 0.00 | 0.00 | 4.31 |
| 5644 | 0.0032 | 0.00 | 0.00 | 4.30 |
| 5645 | 0.0019 | 0.00 | 0.00 | 4.30 |
| 5646 | 0.0032 | 0.00 | 0.00 | 4.30 |
| 5647 | 0.0013 | 0.00 | 0.00 | 4.30 |
| 5648 | 0.0026 | 0.00 | 0.00 | 4.29 |
| 5649 | 0.0016 | 0.00 | 0.00 | 4.29 |
| 5650 | 0.0013 | 0.00 | 0.00 | 4.29 |
| 5651 | 0.0013 | 0.00 | 0.00 | 4.29 |
| 5652 | 0.0016 | 0.00 | 0.00 | 4.29 |
| 5653 | 0.0026 | 0.00 | 0.00 | 4.29 |
| 5654 | 0.0019 | 0.00 | 0.00 | 4.28 |
| 5655 | 0.0019 | 0.00 | 0.00 | 4.28 |
| 5656 | 0.0016 | 0.00 | 0.00 | 4.28 |
| 5657 | 0.0019 | 0.00 | 0.00 | 4.28 |
| 5658 | 0.0022 | 0.00 | 0.00 | 4.28 |
| 5659 | 0.0006 | 0.00 | 0.00 | 4.27 |
| 5660 | 0.0019 | 0.00 | 0.00 | 4.27 |
| 5661 | 0.0010 | 0.00 | 0.00 | 4.27 |
| 5662 | 0.0010 | 0.00 | 0.00 | 4.27 |
| 5663 | 0.0003 | 0.00 | 0.00 | 4.27 |
| 5664 | 0.0019 | 0.00 | 0.00 | 4.27 |
| 5665 | 0.0006 | 0.00 | 0.00 | 4.27 |
| 5666 | 0.0006 | 0.00 | 0.00 | 4.27 |
| 5667 | 0.0006 | 0.00 | 0.00 | 4.27 |
| 5668 | 0.0022 | 0.00 | 0.00 | 4.27 |
| 5669 | 0.0013 | 0.00 | 0.00 | 4.26 |
| 5670 | 0.0019 | 0.00 | 0.00 | 4.26 |
| 5671 | 0.0010 | 0.00 | 0.00 | 4.26 |
| 5672 | 0.0022 | 0.00 | 0.00 | 4.26 |
| 5673 | 0.0022 | 0.00 | 0.00 | 4.26 |
| 5674 | 0.0016 | 0.00 | 0.00 | 4.25 |
| 5675 | 0.0019 | 0.00 | 0.00 | 4.25 |
| 5677 | 0.0010 | 0.00 | 0.00 | 4.25 |
| 5678 | 0.0010 | 0.00 | 0.00 | 4.25 |
| 5679 | 0.0016 | 0.00 | 0.00 | 4.25 |
| 5680 | 0.0013 | 0.00 | 0.00 | 4.25 |
| 5681 | 0.0013 | 0.00 | 0.00 | 4.25 |

| | | | | |
|---|---|---|---|---|
| 5682 | 0.0013 | 0.00 | 0.00 | 4.25 |
| 5683 | 0.0006 | 0.00 | 0.00 | 4.24 |
| 5684 | 0.0016 | 0.00 | 0.00 | 4.24 |
| 5685 | 0.0013 | 0.00 | 0.00 | 4.24 |
| 5686 | 0.0010 | 0.00 | 0.00 | 4.24 |
| 5687 | 0.0010 | 0.00 | 0.00 | 4.24 |
| 5688 | 0.0016 | 0.00 | 0.00 | 4.24 |
| 5689 | 0.0003 | 0.00 | 0.00 | 4.24 |
| 5690 | 0.0006 | 0.00 | 0.00 | 4.24 |
| 5691 | 0.0013 | 0.00 | 0.00 | 4.24 |
| 5692 | 0.0019 | 0.00 | 0.00 | 4.23 |
| 5693 | 0.0010 | 0.00 | 0.00 | 4.23 |
| 5694 | 0.0006 | 0.00 | 0.00 | 4.23 |
| 5695 | 0.0016 | 0.00 | 0.00 | 4.23 |
| 5696 | 0.0010 | 0.00 | 0.00 | 4.23 |
| 5697 | 0.0006 | 0.00 | 0.00 | 4.23 |
| 5698 | 0.0016 | 0.00 | 0.00 | 4.23 |
| 5699 | 0.0006 | 0.00 | 0.00 | 4.23 |
| 5700 | 0.0010 | 0.00 | 0.00 | 4.23 |
| 5701 | 0.0016 | 0.00 | 0.00 | 4.22 |
| 5702 | 0.0029 | 0.00 | 0.00 | 4.22 |
| 5703 | 0.0016 | 0.00 | 0.00 | 4.22 |
| 5704 | 0.0010 | 0.00 | 0.00 | 4.22 |
| 5706 | 0.0013 | 0.00 | 0.00 | 4.22 |
| 5707 | 0.0006 | 0.00 | 0.00 | 4.22 |
| 5708 | 0.0006 | 0.00 | 0.00 | 4.22 |
| 5709 | 0.0003 | 0.00 | 0.00 | 4.22 |
| 5710 | 0.0010 | 0.00 | 0.00 | 4.21 |
| 5711 | 0.0013 | 0.00 | 0.00 | 4.21 |
| 5712 | 0.0013 | 0.00 | 0.00 | 4.21 |
| 5713 | 0.0013 | 0.00 | 0.00 | 4.21 |
| 5714 | 0.0016 | 0.00 | 0.00 | 4.21 |
| 5715 | 0.0010 | 0.00 | 0.00 | 4.21 |
| 5716 | 0.0022 | 0.00 | 0.00 | 4.21 |
| 5717 | 0.0006 | 0.00 | 0.00 | 4.21 |
| 5718 | 0.0013 | 0.00 | 0.00 | 4.20 |
| 5719 | 0.0003 | 0.00 | 0.00 | 4.20 |
| 5720 | 0.0013 | 0.00 | 0.00 | 4.20 |
| 5721 | 0.0013 | 0.00 | 0.00 | 4.20 |
| 5722 | 0.0010 | 0.00 | 0.00 | 4.20 |
| 5723 | 0.0016 | 0.00 | 0.00 | 4.20 |
| 5724 | 0.0013 | 0.00 | 0.00 | 4.20 |
| 5725 | 0.0006 | 0.00 | 0.00 | 4.20 |
| 5726 | 0.0006 | 0.00 | 0.00 | 4.20 |

| | | | | |
|---|---|---|---|---|
| 5727 | 0.0013 | 0.00 | 0.00 | 4.20 |
| 5728 | 0.0016 | 0.00 | 0.00 | 4.19 |
| 5729 | 0.0016 | 0.00 | 0.00 | 4.19 |
| 5730 | 0.0029 | 0.00 | 0.00 | 4.19 |
| 5731 | 0.0003 | 0.00 | 0.00 | 4.19 |
| 5732 | 0.0016 | 0.00 | 0.00 | 4.19 |
| 5733 | 0.0016 | 0.00 | 0.00 | 4.19 |
| 5734 | 0.0013 | 0.00 | 0.00 | 4.18 |
| 5735 | 0.0019 | 0.00 | 0.00 | 4.18 |
| 5736 | 0.0026 | 0.00 | 0.00 | 4.18 |
| 5737 | 0.0010 | 0.00 | 0.00 | 4.18 |
| 5738 | 0.0010 | 0.00 | 0.00 | 4.18 |
| 5739 | 0.0003 | 0.00 | 0.00 | 4.18 |
| 5740 | 0.0026 | 0.00 | 0.00 | 4.18 |
| 5741 | 0.0013 | 0.00 | 0.00 | 4.17 |
| 5742 | 0.0006 | 0.00 | 0.00 | 4.17 |
| 5743 | 0.0006 | 0.00 | 0.00 | 4.17 |
| 5744 | 0.0010 | 0.00 | 0.00 | 4.17 |
| 5745 | 0.0010 | 0.00 | 0.00 | 4.17 |
| 5746 | 0.0022 | 0.00 | 0.00 | 4.17 |
| 5747 | 0.0019 | 0.00 | 0.00 | 4.17 |
| 5748 | 0.0010 | 0.00 | 0.00 | 4.17 |
| 5749 | 0.0016 | 0.00 | 0.00 | 4.16 |
| 5750 | 0.0003 | 0.00 | 0.00 | 4.16 |
| 5751 | 0.0029 | 0.00 | 0.00 | 4.16 |
| 5752 | 0.0035 | 0.00 | 0.00 | 4.16 |
| 5753 | 0.0006 | 0.00 | 0.00 | 4.16 |
| 5755 | 0.0010 | 0.00 | 0.00 | 4.16 |
| 5756 | 0.0006 | 0.00 | 0.00 | 4.15 |
| 5757 | 0.0006 | 0.00 | 0.00 | 4.15 |
| 5758 | 0.0006 | 0.00 | 0.00 | 4.15 |
| 5759 | 0.0019 | 0.00 | 0.00 | 4.15 |
| 5760 | 0.0013 | 0.00 | 0.00 | 4.15 |
| 5761 | 0.0013 | 0.00 | 0.00 | 4.15 |
| 5762 | 0.0016 | 0.00 | 0.00 | 4.15 |
| 5763 | 0.0010 | 0.00 | 0.00 | 4.15 |
| 5764 | 0.0019 | 0.00 | 0.00 | 4.15 |
| 5765 | 0.0022 | 0.00 | 0.00 | 4.14 |
| 5766 | 0.0013 | 0.00 | 0.00 | 4.14 |
| 5767 | 0.0013 | 0.00 | 0.00 | 4.14 |
| 5768 | 0.0013 | 0.00 | 0.00 | 4.14 |
| 5769 | 0.0010 | 0.00 | 0.00 | 4.14 |
| 5770 | 0.0010 | 0.00 | 0.00 | 4.14 |
| 5771 | 0.0006 | 0.00 | 0.00 | 4.14 |

| | | | | |
|---|---|---|---|---|
| 5772 | 0.0016 | 0.00 | 0.00 | 4.13 |
| 5773 | 0.0010 | 0.00 | 0.00 | 4.13 |
| 5774 | 0.0019 | 0.00 | 0.00 | 4.13 |
| 5775 | 0.0003 | 0.00 | 0.00 | 4.13 |
| 5776 | 0.0010 | 0.00 | 0.00 | 4.13 |
| 5777 | 0.0019 | 0.00 | 0.00 | 4.13 |
| 5778 | 0.0022 | 0.00 | 0.00 | 4.13 |
| 5779 | 0.0019 | 0.00 | 0.00 | 4.12 |
| 5780 | 0.0019 | 0.00 | 0.00 | 4.12 |
| 5781 | 0.0019 | 0.00 | 0.00 | 4.12 |
| 5782 | 0.0013 | 0.00 | 0.00 | 4.12 |
| 5783 | 0.0019 | 0.00 | 0.00 | 4.12 |
| 5784 | 0.0006 | 0.00 | 0.00 | 4.12 |
| 5785 | 0.0013 | 0.00 | 0.00 | 4.12 |
| 5786 | 0.0010 | 0.00 | 0.00 | 4.11 |
| 5787 | 0.0019 | 0.00 | 0.00 | 4.11 |
| 5788 | 0.0006 | 0.00 | 0.00 | 4.11 |
| 5789 | 0.0019 | 0.00 | 0.00 | 4.11 |
| 5790 | 0.0003 | 0.00 | 0.00 | 4.11 |
| 5791 | 0.0006 | 0.00 | 0.00 | 4.11 |
| 5792 | 0.0013 | 0.00 | 0.00 | 4.11 |
| 5793 | 0.0016 | 0.00 | 0.00 | 4.11 |
| 5794 | 0.0010 | 0.00 | 0.00 | 4.10 |
| 5795 | 0.0010 | 0.00 | 0.00 | 4.10 |
| 5796 | 0.0006 | 0.00 | 0.00 | 4.10 |
| 5797 | 0.0022 | 0.00 | 0.00 | 4.10 |
| 5798 | 0.0019 | 0.00 | 0.00 | 4.10 |
| 5799 | 0.0016 | 0.00 | 0.00 | 4.10 |
| 5800 | 0.0013 | 0.00 | 0.00 | 4.10 |
| 5801 | 0.0010 | 0.00 | 0.00 | 4.09 |
| 5802 | 0.0010 | 0.00 | 0.00 | 4.09 |
| 5803 | 0.0019 | 0.00 | 0.00 | 4.09 |
| 5804 | 0.0013 | 0.00 | 0.00 | 4.09 |
| 5805 | 0.0003 | 0.00 | 0.00 | 4.09 |
| 5806 | 0.0019 | 0.00 | 0.00 | 4.09 |
| 5807 | 0.0016 | 0.00 | 0.00 | 4.09 |
| 5808 | 0.0013 | 0.00 | 0.00 | 4.09 |
| 5809 | 0.0010 | 0.00 | 0.00 | 4.08 |
| 5810 | 0.0010 | 0.00 | 0.00 | 4.08 |
| 5811 | 0.0010 | 0.00 | 0.00 | 4.08 |
| 5812 | 0.0016 | 0.00 | 0.00 | 4.08 |
| 5813 | 0.0010 | 0.00 | 0.00 | 4.08 |
| 5814 | 0.0016 | 0.00 | 0.00 | 4.08 |
| 5815 | 0.0003 | 0.00 | 0.00 | 4.08 |

| | | | | |
|---|---|---|---|---|
| 5816 | 0.0006 | 0.00 | 0.00 | 4.08 |
| 5817 | 0.0003 | 0.00 | 0.00 | 4.08 |
| 5818 | 0.0022 | 0.00 | 0.00 | 4.08 |
| 5819 | 0.0010 | 0.00 | 0.00 | 4.07 |
| 5820 | 0.0006 | 0.00 | 0.00 | 4.07 |
| 5821 | 0.0003 | 0.00 | 0.00 | 4.07 |
| 5822 | 0.0016 | 0.00 | 0.00 | 4.07 |
| 5823 | 0.0016 | 0.00 | 0.00 | 4.07 |
| 5824 | 0.0003 | 0.00 | 0.00 | 4.07 |
| 5825 | 0.0019 | 0.00 | 0.00 | 4.07 |
| 5826 | 0.0006 | 0.00 | 0.00 | 4.07 |
| 5827 | 0.0003 | 0.00 | 0.00 | 4.07 |
| 5829 | 0.0013 | 0.00 | 0.00 | 4.07 |
| 5830 | 0.0016 | 0.00 | 0.00 | 4.06 |
| 5831 | 0.0013 | 0.00 | 0.00 | 4.06 |
| 5832 | 0.0013 | 0.00 | 0.00 | 4.06 |
| 5833 | 0.0003 | 0.00 | 0.00 | 4.06 |
| 5835 | 0.0016 | 0.00 | 0.00 | 4.06 |
| 5836 | 0.0013 | 0.00 | 0.00 | 4.06 |
| 5837 | 0.0010 | 0.00 | 0.00 | 4.06 |
| 5838 | 0.0013 | 0.00 | 0.00 | 4.06 |
| 5839 | 0.0010 | 0.00 | 0.00 | 4.05 |
| 5840 | 0.0010 | 0.00 | 0.00 | 4.05 |
| 5841 | 0.0019 | 0.00 | 0.00 | 4.05 |
| 5842 | 0.0016 | 0.00 | 0.00 | 4.05 |
| 5843 | 0.0026 | 0.00 | 0.00 | 4.05 |
| 5844 | 0.0016 | 0.00 | 0.00 | 4.05 |
| 5845 | 0.0003 | 0.00 | 0.00 | 4.05 |
| 5846 | 0.0013 | 0.00 | 0.00 | 4.04 |
| 5847 | 0.0019 | 0.00 | 0.00 | 4.04 |
| 5848 | 0.0006 | 0.00 | 0.00 | 4.04 |
| 5849 | 0.0006 | 0.00 | 0.00 | 4.04 |
| 5850 | 0.0029 | 0.00 | 0.00 | 4.04 |
| 5851 | 0.0003 | 0.00 | 0.00 | 4.04 |
| 5852 | 0.0032 | 0.00 | 0.00 | 4.04 |
| 5853 | 0.0010 | 0.00 | 0.00 | 4.03 |
| 5854 | 0.0019 | 0.00 | 0.00 | 4.03 |
| 5855 | 0.0019 | 0.00 | 0.00 | 4.03 |
| 5856 | 0.0003 | 0.00 | 0.00 | 4.03 |
| 5857 | 0.0016 | 0.00 | 0.00 | 4.03 |
| 5858 | 0.0013 | 0.00 | 0.00 | 4.03 |
| 5859 | 0.0010 | 0.00 | 0.00 | 4.03 |
| 5860 | 0.0019 | 0.00 | 0.00 | 4.03 |
| 5861 | 0.0016 | 0.00 | 0.00 | 4.02 |

| | | | | |
|---|---|---|---|---|
| 5862 | 0.0010 | 0.00 | 0.00 | 4.02 |
| 5863 | 0.0016 | 0.00 | 0.00 | 4.02 |
| 5864 | 0.0026 | 0.00 | 0.00 | 4.02 |
| 5865 | 0.0003 | 0.00 | 0.00 | 4.02 |
| 5866 | 0.0010 | 0.00 | 0.00 | 4.02 |
| 5867 | 0.0003 | 0.00 | 0.00 | 4.02 |
| 5868 | 0.0022 | 0.00 | 0.00 | 4.01 |
| 5869 | 0.0019 | 0.00 | 0.00 | 4.01 |
| 5870 | 0.0016 | 0.00 | 0.00 | 4.01 |
| 5871 | 0.0013 | 0.00 | 0.00 | 4.01 |
| 5872 | 0.0013 | 0.00 | 0.00 | 4.01 |
| 5873 | 0.0022 | 0.00 | 0.00 | 4.01 |
| 5874 | 0.0022 | 0.00 | 0.00 | 4.00 |
| 5875 | 0.0019 | 0.00 | 0.00 | 4.00 |
| 5876 | 0.0010 | 0.00 | 0.00 | 4.00 |
| 5877 | 0.0022 | 0.00 | 0.00 | 4.00 |
| 5878 | 0.0006 | 0.00 | 0.00 | 4.00 |
| 5880 | 0.0013 | 0.00 | 0.00 | 4.00 |
| 5881 | 0.0022 | 0.00 | 0.00 | 3.99 |
| 5882 | 0.0006 | 0.00 | 0.00 | 3.99 |
| 5883 | 0.0016 | 0.00 | 0.00 | 3.99 |
| 5884 | 0.0029 | 0.00 | 0.00 | 3.99 |
| 5885 | 0.0003 | 0.00 | 0.00 | 3.99 |
| 5886 | 0.0022 | 0.00 | 0.00 | 3.99 |
| 5887 | 0.0022 | 0.00 | 0.00 | 3.98 |
| 5888 | 0.0003 | 0.00 | 0.00 | 3.98 |
| 5889 | 0.0022 | 0.00 | 0.00 | 3.98 |
| 5890 | 0.0003 | 0.00 | 0.00 | 3.98 |
| 5891 | 0.0013 | 0.00 | 0.00 | 3.98 |
| 5892 | 0.0016 | 0.00 | 0.00 | 3.98 |
| 5893 | 0.0032 | 0.00 | 0.00 | 3.98 |
| 5894 | 0.0013 | 0.00 | 0.00 | 3.97 |
| 5895 | 0.0026 | 0.00 | 0.00 | 3.97 |
| 5896 | 0.0019 | 0.00 | 0.00 | 3.97 |
| 5897 | 0.0013 | 0.00 | 0.00 | 3.97 |
| 5898 | 0.0010 | 0.00 | 0.00 | 3.97 |
| 5899 | 0.0026 | 0.00 | 0.00 | 3.97 |
| 5900 | 0.0016 | 0.00 | 0.00 | 3.96 |
| 5901 | 0.0016 | 0.00 | 0.00 | 3.96 |
| 5902 | 0.0016 | 0.00 | 0.00 | 3.96 |
| 5903 | 0.0010 | 0.00 | 0.00 | 3.96 |
| 5904 | 0.0013 | 0.00 | 0.00 | 3.96 |
| 5905 | 0.0016 | 0.00 | 0.00 | 3.96 |
| 5906 | 0.0006 | 0.00 | 0.00 | 3.95 |

| | | | | |
|---|---|---|---|---|
| 5907 | 0.0006 | 0.00 | 0.00 | 3.95 |
| 5908 | 0.0013 | 0.00 | 0.00 | 3.95 |
| 5909 | 0.0016 | 0.00 | 0.00 | 3.95 |
| 5910 | 0.0010 | 0.00 | 0.00 | 3.95 |
| 5911 | 0.0026 | 0.00 | 0.00 | 3.95 |
| 5912 | 0.0016 | 0.00 | 0.00 | 3.95 |
| 5913 | 0.0019 | 0.00 | 0.00 | 3.95 |
| 5914 | 0.0003 | 0.00 | 0.00 | 3.94 |
| 5915 | 0.0006 | 0.00 | 0.00 | 3.94 |
| 5916 | 0.0010 | 0.00 | 0.00 | 3.94 |
| 5917 | 0.0019 | 0.00 | 0.00 | 3.94 |
| 5918 | 0.0013 | 0.00 | 0.00 | 3.94 |
| 5919 | 0.0006 | 0.00 | 0.00 | 3.94 |
| 5920 | 0.0003 | 0.00 | 0.00 | 3.94 |
| 5921 | 0.0003 | 0.00 | 0.00 | 3.94 |
| 5922 | 0.0016 | 0.00 | 0.00 | 3.94 |
| 5923 | 0.0016 | 0.00 | 0.00 | 3.94 |
| 5924 | 0.0016 | 0.00 | 0.00 | 3.93 |
| 5925 | 0.0013 | 0.00 | 0.00 | 3.93 |
| 5926 | 0.0013 | 0.00 | 0.00 | 3.93 |
| 5927 | 0.0006 | 0.00 | 0.00 | 3.93 |
| 5928 | 0.0006 | 0.00 | 0.00 | 3.93 |
| 5929 | 0.0019 | 0.00 | 0.00 | 3.93 |
| 5930 | 0.0010 | 0.00 | 0.00 | 3.93 |
| 5931 | 0.0019 | 0.00 | 0.00 | 3.93 |
| 5932 | 0.0006 | 0.00 | 0.00 | 3.92 |
| 5933 | 0.0003 | 0.00 | 0.00 | 3.92 |
| 5934 | 0.0013 | 0.00 | 0.00 | 3.92 |
| 5935 | 0.0006 | 0.00 | 0.00 | 3.92 |
| 5936 | 0.0010 | 0.00 | 0.00 | 3.92 |
| 5937 | 0.0013 | 0.00 | 0.00 | 3.92 |
| 5938 | 0.0003 | 0.00 | 0.00 | 3.92 |
| 5939 | 0.0010 | 0.00 | 0.00 | 3.92 |
| 5940 | 0.0003 | 0.00 | 0.00 | 3.92 |
| 5941 | 0.0013 | 0.00 | 0.00 | 3.92 |
| 5942 | 0.0010 | 0.00 | 0.00 | 3.92 |
| 5943 | 0.0026 | 0.00 | 0.00 | 3.91 |
| 5944 | 0.0022 | 0.00 | 0.00 | 3.91 |
| 5945 | 0.0013 | 0.00 | 0.00 | 3.91 |
| 5946 | 0.0016 | 0.00 | 0.00 | 3.91 |
| 5947 | 0.0006 | 0.00 | 0.00 | 3.91 |
| 5948 | 0.0010 | 0.00 | 0.00 | 3.91 |
| 5949 | 0.0013 | 0.00 | 0.00 | 3.91 |
| 5950 | 0.0019 | 0.00 | 0.00 | 3.90 |

| | | | | |
|---|---|---|---|---|
| 5951 | 0.0006 | 0.00 | 0.00 | 3.90 |
| 5952 | 0.0003 | 0.00 | 0.00 | 3.90 |
| 5953 | 0.0003 | 0.00 | 0.00 | 3.90 |
| 5954 | 0.0016 | 0.00 | 0.00 | 3.90 |
| 5955 | 0.0010 | 0.00 | 0.00 | 3.90 |
| 5956 | 0.0010 | 0.00 | 0.00 | 3.90 |
| 5957 | 0.0016 | 0.00 | 0.00 | 3.90 |
| 5958 | 0.0006 | 0.00 | 0.00 | 3.90 |
| 5959 | 0.0013 | 0.00 | 0.00 | 3.89 |
| 5960 | 0.0013 | 0.00 | 0.00 | 3.89 |
| 5961 | 0.0006 | 0.00 | 0.00 | 3.89 |
| 5962 | 0.0010 | 0.00 | 0.00 | 3.89 |
| 5963 | 0.0013 | 0.00 | 0.00 | 3.89 |
| 5964 | 0.0013 | 0.00 | 0.00 | 3.89 |
| 5965 | 0.0003 | 0.00 | 0.00 | 3.89 |
| 5966 | 0.0022 | 0.00 | 0.00 | 3.89 |
| 5967 | 0.0013 | 0.00 | 0.00 | 3.89 |
| 5968 | 0.0006 | 0.00 | 0.00 | 3.88 |
| 5969 | 0.0003 | 0.00 | 0.00 | 3.88 |
| 5970 | 0.0013 | 0.00 | 0.00 | 3.88 |
| 5971 | 0.0019 | 0.00 | 0.00 | 3.88 |
| 5972 | 0.0022 | 0.00 | 0.00 | 3.88 |
| 5973 | 0.0010 | 0.00 | 0.00 | 3.88 |
| 5974 | 0.0006 | 0.00 | 0.00 | 3.88 |
| 5975 | 0.0016 | 0.00 | 0.00 | 3.88 |
| 5976 | 0.0019 | 0.00 | 0.00 | 3.87 |
| 5977 | 0.0022 | 0.00 | 0.00 | 3.87 |
| 5978 | 0.0016 | 0.00 | 0.00 | 3.87 |
| 5979 | 0.0010 | 0.00 | 0.00 | 3.87 |
| 5980 | 0.0010 | 0.00 | 0.00 | 3.87 |
| 5981 | 0.0013 | 0.00 | 0.00 | 3.87 |
| 5982 | 0.0016 | 0.00 | 0.00 | 3.87 |
| 5983 | 0.0010 | 0.00 | 0.00 | 3.86 |
| 5984 | 0.0010 | 0.00 | 0.00 | 3.86 |
| 5985 | 0.0006 | 0.00 | 0.00 | 3.86 |
| 5986 | 0.0003 | 0.00 | 0.00 | 3.86 |
| 5987 | 0.0013 | 0.00 | 0.00 | 3.86 |
| 5988 | 0.0006 | 0.00 | 0.00 | 3.86 |
| 5989 | 0.0003 | 0.00 | 0.00 | 3.86 |
| 5990 | 0.0013 | 0.00 | 0.00 | 3.86 |
| 5991 | 0.0003 | 0.00 | 0.00 | 3.86 |
| 5992 | 0.0006 | 0.00 | 0.00 | 3.86 |
| 5993 | 0.0013 | 0.00 | 0.00 | 3.86 |
| 5994 | 0.0026 | 0.00 | 0.00 | 3.86 |

| | | | | |
|---|---|---|---|---|
| 5995 | 0.0010 | 0.00 | 0.00 | 3.85 |
| 5996 | 0.0010 | 0.00 | 0.00 | 3.85 |
| 5997 | 0.0016 | 0.00 | 0.00 | 3.85 |
| 5998 | 0.0016 | 0.00 | 0.00 | 3.85 |
| 5999 | 0.0010 | 0.00 | 0.00 | 3.85 |
| 6000 | 0.0010 | 0.00 | 0.00 | 3.85 |
| 6001 | 0.0019 | 0.00 | 0.00 | 3.85 |
| 6002 | 0.0003 | 0.00 | 0.00 | 3.84 |
| 6003 | 0.0006 | 0.00 | 0.00 | 3.84 |
| 6004 | 0.0006 | 0.00 | 0.00 | 3.84 |
| 6005 | 0.0013 | 0.00 | 0.00 | 3.84 |
| 6006 | 0.0019 | 0.00 | 0.00 | 3.84 |
| 6007 | 0.0010 | 0.00 | 0.00 | 3.84 |
| 6008 | 0.0010 | 0.00 | 0.00 | 3.84 |
| 6009 | 0.0016 | 0.00 | 0.00 | 3.84 |
| 6010 | 0.0010 | 0.00 | 0.00 | 3.84 |
| 6011 | 0.0016 | 0.00 | 0.00 | 3.83 |
| 6012 | 0.0026 | 0.00 | 0.00 | 3.83 |
| 6013 | 0.0013 | 0.00 | 0.00 | 3.83 |
| 6014 | 0.0016 | 0.00 | 0.00 | 3.83 |
| 6015 | 0.0013 | 0.00 | 0.00 | 3.83 |
| 6016 | 0.0006 | 0.00 | 0.00 | 3.83 |
| 6017 | 0.0013 | 0.00 | 0.00 | 3.83 |
| 6018 | 0.0010 | 0.00 | 0.00 | 3.82 |
| 6019 | 0.0019 | 0.00 | 0.00 | 3.82 |
| 6020 | 0.0010 | 0.00 | 0.00 | 3.82 |
| 6021 | 0.0016 | 0.00 | 0.00 | 3.82 |
| 6022 | 0.0013 | 0.00 | 0.00 | 3.82 |
| 6023 | 0.0010 | 0.00 | 0.00 | 3.82 |
| 6024 | 0.0016 | 0.00 | 0.00 | 3.82 |
| 6025 | 0.0022 | 0.00 | 0.00 | 3.82 |
| 6026 | 0.0019 | 0.00 | 0.00 | 3.81 |
| 6027 | 0.0013 | 0.00 | 0.00 | 3.81 |
| 6028 | 0.0010 | 0.00 | 0.00 | 3.81 |
| 6029 | 0.0010 | 0.00 | 0.00 | 3.81 |
| 6030 | 0.0010 | 0.00 | 0.00 | 3.81 |
| 6031 | 0.0022 | 0.00 | 0.00 | 3.81 |
| 6032 | 0.0022 | 0.00 | 0.00 | 3.80 |
| 6033 | 0.0016 | 0.00 | 0.00 | 3.80 |
| 6034 | 0.0003 | 0.00 | 0.00 | 3.80 |
| 6035 | 0.0016 | 0.00 | 0.00 | 3.80 |
| 6036 | 0.0010 | 0.00 | 0.00 | 3.80 |
| 6037 | 0.0013 | 0.00 | 0.00 | 3.80 |
| 6038 | 0.0022 | 0.00 | 0.00 | 3.80 |

| | | | | |
|---|---|---|---|---|
| 6039 | 0.0026 | 0.00 | 0.00 | 3.79 |
| 6040 | 0.0010 | 0.00 | 0.00 | 3.79 |
| 6041 | 0.0010 | 0.00 | 0.00 | 3.79 |
| 6042 | 0.0013 | 0.00 | 0.00 | 3.79 |
| 6043 | 0.0003 | 0.00 | 0.00 | 3.79 |
| 6045 | 0.0016 | 0.00 | 0.00 | 3.79 |
| 6046 | 0.0010 | 0.00 | 0.00 | 3.79 |
| 6047 | 0.0010 | 0.00 | 0.00 | 3.79 |
| 6048 | 0.0013 | 0.00 | 0.00 | 3.78 |
| 6049 | 0.0006 | 0.00 | 0.00 | 3.78 |
| 6050 | 0.0006 | 0.00 | 0.00 | 3.78 |
| 6051 | 0.0006 | 0.00 | 0.00 | 3.78 |
| 6052 | 0.0003 | 0.00 | 0.00 | 3.78 |
| 6053 | 0.0006 | 0.00 | 0.00 | 3.78 |
| 6054 | 0.0006 | 0.00 | 0.00 | 3.78 |
| 6055 | 0.0006 | 0.00 | 0.00 | 3.78 |
| 6056 | 0.0006 | 0.00 | 0.00 | 3.78 |
| 6057 | 0.0019 | 0.00 | 0.00 | 3.78 |
| 6058 | 0.0006 | 0.00 | 0.00 | 3.78 |
| 6059 | 0.0006 | 0.00 | 0.00 | 3.78 |
| 6060 | 0.0016 | 0.00 | 0.00 | 3.78 |
| 6061 | 0.0010 | 0.00 | 0.00 | 3.77 |
| 6062 | 0.0006 | 0.00 | 0.00 | 3.77 |
| 6063 | 0.0016 | 0.00 | 0.00 | 3.77 |
| 6064 | 0.0010 | 0.00 | 0.00 | 3.77 |
| 6065 | 0.0013 | 0.00 | 0.00 | 3.77 |
| 6066 | 0.0016 | 0.00 | 0.00 | 3.77 |
| 6067 | 0.0010 | 0.00 | 0.00 | 3.77 |
| 6068 | 0.0006 | 0.00 | 0.00 | 3.77 |
| 6069 | 0.0010 | 0.00 | 0.00 | 3.77 |
| 6070 | 0.0006 | 0.00 | 0.00 | 3.76 |
| 6071 | 0.0010 | 0.00 | 0.00 | 3.76 |
| 6072 | 0.0010 | 0.00 | 0.00 | 3.76 |
| 6073 | 0.0010 | 0.00 | 0.00 | 3.76 |
| 6074 | 0.0006 | 0.00 | 0.00 | 3.76 |
| 6076 | 0.0016 | 0.00 | 0.00 | 3.76 |
| 6077 | 0.0006 | 0.00 | 0.00 | 3.76 |
| 6078 | 0.0010 | 0.00 | 0.00 | 3.76 |
| 6079 | 0.0013 | 0.00 | 0.00 | 3.76 |
| 6080 | 0.0026 | 0.00 | 0.00 | 3.76 |
| 6081 | 0.0013 | 0.00 | 0.00 | 3.75 |
| 6082 | 0.0006 | 0.00 | 0.00 | 3.75 |
| 6083 | 0.0010 | 0.00 | 0.00 | 3.75 |
| 6084 | 0.0013 | 0.00 | 0.00 | 3.75 |

| | | | | |
|---|---|---|---|---|
| 6085 | 0.0006 | 0.00 | 0.00 | 3.75 |
| 6086 | 0.0019 | 0.00 | 0.00 | 3.75 |
| 6088 | 0.0006 | 0.00 | 0.00 | 3.75 |
| 6089 | 0.0003 | 0.00 | 0.00 | 3.75 |
| 6090 | 0.0010 | 0.00 | 0.00 | 3.75 |
| 6091 | 0.0013 | 0.00 | 0.00 | 3.74 |
| 6093 | 0.0013 | 0.00 | 0.00 | 3.74 |
| 6094 | 0.0010 | 0.00 | 0.00 | 3.74 |
| 6095 | 0.0016 | 0.00 | 0.00 | 3.74 |
| 6096 | 0.0010 | 0.00 | 0.00 | 3.74 |
| 6097 | 0.0006 | 0.00 | 0.00 | 3.74 |
| 6098 | 0.0010 | 0.00 | 0.00 | 3.74 |
| 6099 | 0.0019 | 0.00 | 0.00 | 3.74 |
| 6100 | 0.0006 | 0.00 | 0.00 | 3.73 |
| 6101 | 0.0013 | 0.00 | 0.00 | 3.73 |
| 6102 | 0.0006 | 0.00 | 0.00 | 3.73 |
| 6103 | 0.0010 | 0.00 | 0.00 | 3.73 |
| 6104 | 0.0010 | 0.00 | 0.00 | 3.73 |
| 6105 | 0.0006 | 0.00 | 0.00 | 3.73 |
| 6107 | 0.0013 | 0.00 | 0.00 | 3.73 |
| 6108 | 0.0013 | 0.00 | 0.00 | 3.73 |
| 6109 | 0.0006 | 0.00 | 0.00 | 3.73 |
| 6110 | 0.0006 | 0.00 | 0.00 | 3.73 |
| 6111 | 0.0016 | 0.00 | 0.00 | 3.73 |
| 6113 | 0.0006 | 0.00 | 0.00 | 3.72 |
| 6114 | 0.0013 | 0.00 | 0.00 | 3.72 |
| 6115 | 0.0013 | 0.00 | 0.00 | 3.72 |
| 6116 | 0.0013 | 0.00 | 0.00 | 3.72 |
| 6117 | 0.0010 | 0.00 | 0.00 | 3.72 |
| 6118 | 0.0010 | 0.00 | 0.00 | 3.72 |
| 6119 | 0.0003 | 0.00 | 0.00 | 3.72 |
| 6120 | 0.0003 | 0.00 | 0.00 | 3.72 |
| 6121 | 0.0006 | 0.00 | 0.00 | 3.72 |
| 6122 | 0.0013 | 0.00 | 0.00 | 3.72 |
| 6123 | 0.0013 | 0.00 | 0.00 | 3.72 |
| 6124 | 0.0010 | 0.00 | 0.00 | 3.71 |
| 6125 | 0.0010 | 0.00 | 0.00 | 3.71 |
| 6126 | 0.0010 | 0.00 | 0.00 | 3.71 |
| 6127 | 0.0019 | 0.00 | 0.00 | 3.71 |
| 6128 | 0.0010 | 0.00 | 0.00 | 3.71 |
| 6129 | 0.0003 | 0.00 | 0.00 | 3.71 |
| 6130 | 0.0022 | 0.00 | 0.00 | 3.71 |
| 6131 | 0.0022 | 0.00 | 0.00 | 3.71 |
| 6132 | 0.0022 | 0.00 | 0.00 | 3.70 |

| | | | | |
|---|---|---|---|---|
| 6133 | 0.0010 | 0.00 | 0.00 | 3.70 |
| 6134 | 0.0019 | 0.00 | 0.00 | 3.70 |
| 6135 | 0.0010 | 0.00 | 0.00 | 3.70 |
| 6136 | 0.0022 | 0.00 | 0.00 | 3.70 |
| 6137 | 0.0010 | 0.00 | 0.00 | 3.69 |
| 6138 | 0.0019 | 0.00 | 0.00 | 3.69 |
| 6139 | 0.0016 | 0.00 | 0.00 | 3.69 |
| 6140 | 0.0029 | 0.00 | 0.00 | 3.69 |
| 6141 | 0.0013 | 0.00 | 0.00 | 3.69 |
| 6142 | 0.0013 | 0.00 | 0.00 | 3.69 |
| 6143 | 0.0010 | 0.00 | 0.00 | 3.68 |
| 6144 | 0.0006 | 0.00 | 0.00 | 3.68 |
| 6145 | 0.0013 | 0.00 | 0.00 | 3.68 |
| 6146 | 0.0013 | 0.00 | 0.00 | 3.68 |
| 6147 | 0.0019 | 0.00 | 0.00 | 3.68 |
| 6148 | 0.0013 | 0.00 | 0.00 | 3.68 |
| 6149 | 0.0029 | 0.00 | 0.00 | 3.68 |
| 6150 | 0.0016 | 0.00 | 0.00 | 3.67 |
| 6151 | 0.0019 | 0.00 | 0.00 | 3.67 |
| 6152 | 0.0010 | 0.00 | 0.00 | 3.67 |
| 6153 | 0.0022 | 0.00 | 0.00 | 3.67 |
| 6154 | 0.0019 | 0.00 | 0.00 | 3.67 |
| 6155 | 0.0022 | 0.00 | 0.00 | 3.67 |
| 6156 | 0.0019 | 0.00 | 0.00 | 3.66 |
| 6157 | 0.0016 | 0.00 | 0.00 | 3.66 |
| 6158 | 0.0010 | 0.00 | 0.00 | 3.66 |
| 6159 | 0.0006 | 0.00 | 0.00 | 3.66 |
| 6160 | 0.0006 | 0.00 | 0.00 | 3.66 |
| 6161 | 0.0006 | 0.00 | 0.00 | 3.66 |
| 6162 | 0.0010 | 0.00 | 0.00 | 3.66 |
| 6163 | 0.0003 | 0.00 | 0.00 | 3.66 |
| 6165 | 0.0013 | 0.00 | 0.00 | 3.66 |
| 6166 | 0.0022 | 0.00 | 0.00 | 3.65 |
| 6167 | 0.0016 | 0.00 | 0.00 | 3.65 |
| 6168 | 0.0010 | 0.00 | 0.00 | 3.65 |
| 6169 | 0.0006 | 0.00 | 0.00 | 3.65 |
| 6170 | 0.0013 | 0.00 | 0.00 | 3.65 |
| 6171 | 0.0010 | 0.00 | 0.00 | 3.65 |
| 6172 | 0.0013 | 0.00 | 0.00 | 3.65 |
| 6173 | 0.0013 | 0.00 | 0.00 | 3.65 |
| 6174 | 0.0006 | 0.00 | 0.00 | 3.64 |
| 6175 | 0.0003 | 0.00 | 0.00 | 3.64 |
| 6176 | 0.0010 | 0.00 | 0.00 | 3.64 |
| 6177 | 0.0006 | 0.00 | 0.00 | 3.64 |

| | | | | |
|---|---|---|---|---|
| 6178 | 0.0003 | 0.00 | 0.00 | 3.64 |
| 6179 | 0.0006 | 0.00 | 0.00 | 3.64 |
| 6180 | 0.0013 | 0.00 | 0.00 | 3.64 |
| 6181 | 0.0016 | 0.00 | 0.00 | 3.64 |
| 6182 | 0.0006 | 0.00 | 0.00 | 3.64 |
| 6183 | 0.0016 | 0.00 | 0.00 | 3.64 |
| 6184 | 0.0006 | 0.00 | 0.00 | 3.64 |
| 6185 | 0.0013 | 0.00 | 0.00 | 3.64 |
| 6186 | 0.0013 | 0.00 | 0.00 | 3.63 |
| 6187 | 0.0013 | 0.00 | 0.00 | 3.63 |
| 6188 | 0.0016 | 0.00 | 0.00 | 3.63 |
| 6189 | 0.0006 | 0.00 | 0.00 | 3.63 |
| 6190 | 0.0010 | 0.00 | 0.00 | 3.63 |
| 6191 | 0.0013 | 0.00 | 0.00 | 3.63 |
| 6192 | 0.0016 | 0.00 | 0.00 | 3.63 |
| 6193 | 0.0006 | 0.00 | 0.00 | 3.63 |
| 6194 | 0.0016 | 0.00 | 0.00 | 3.62 |
| 6195 | 0.0010 | 0.00 | 0.00 | 3.62 |
| 6196 | 0.0013 | 0.00 | 0.00 | 3.62 |
| 6197 | 0.0010 | 0.00 | 0.00 | 3.62 |
| 6198 | 0.0006 | 0.00 | 0.00 | 3.62 |
| 6200 | 0.0022 | 0.00 | 0.00 | 3.62 |
| 6201 | 0.0019 | 0.00 | 0.00 | 3.62 |
| 6202 | 0.0010 | 0.00 | 0.00 | 3.62 |
| 6203 | 0.0010 | 0.00 | 0.00 | 3.61 |
| 6204 | 0.0006 | 0.00 | 0.00 | 3.61 |
| 6206 | 0.0003 | 0.00 | 0.00 | 3.61 |
| 6207 | 0.0003 | 0.00 | 0.00 | 3.61 |
| 6208 | 0.0016 | 0.00 | 0.00 | 3.61 |
| 6209 | 0.0006 | 0.00 | 0.00 | 3.61 |
| 6210 | 0.0006 | 0.00 | 0.00 | 3.61 |
| 6211 | 0.0026 | 0.00 | 0.00 | 3.61 |
| 6212 | 0.0006 | 0.00 | 0.00 | 3.61 |
| 6213 | 0.0010 | 0.00 | 0.00 | 3.61 |
| 6214 | 0.0010 | 0.00 | 0.00 | 3.60 |
| 6216 | 0.0006 | 0.00 | 0.00 | 3.60 |
| 6217 | 0.0016 | 0.00 | 0.00 | 3.60 |
| 6218 | 0.0019 | 0.00 | 0.00 | 3.60 |
| 6219 | 0.0016 | 0.00 | 0.00 | 3.60 |
| 6220 | 0.0010 | 0.00 | 0.00 | 3.60 |
| 6221 | 0.0010 | 0.00 | 0.00 | 3.60 |
| 6223 | 0.0013 | 0.00 | 0.00 | 3.60 |
| 6224 | 0.0010 | 0.00 | 0.00 | 3.59 |
| 6225 | 0.0006 | 0.00 | 0.00 | 3.59 |

| | | | | |
|---|---|---|---|---|
| 6226 | 0.0013 | 0.00 | 0.00 | 3.59 |
| 6227 | 0.0006 | 0.00 | 0.00 | 3.59 |
| 6228 | 0.0003 | 0.00 | 0.00 | 3.59 |
| 6229 | 0.0019 | 0.00 | 0.00 | 3.59 |
| 6230 | 0.0016 | 0.00 | 0.00 | 3.59 |
| 6231 | 0.0010 | 0.00 | 0.00 | 3.59 |
| 6232 | 0.0006 | 0.00 | 0.00 | 3.59 |
| 6233 | 0.0013 | 0.00 | 0.00 | 3.59 |
| 6234 | 0.0010 | 0.00 | 0.00 | 3.58 |
| 6235 | 0.0006 | 0.00 | 0.00 | 3.58 |
| 6236 | 0.0003 | 0.00 | 0.00 | 3.58 |
| 6237 | 0.0006 | 0.00 | 0.00 | 3.58 |
| 6238 | 0.0013 | 0.00 | 0.00 | 3.58 |
| 6239 | 0.0016 | 0.00 | 0.00 | 3.58 |
| 6240 | 0.0013 | 0.00 | 0.00 | 3.58 |
| 6241 | 0.0013 | 0.00 | 0.00 | 3.58 |
| 6242 | 0.0003 | 0.00 | 0.00 | 3.58 |
| 6243 | 0.0013 | 0.00 | 0.00 | 3.58 |
| 6244 | 0.0013 | 0.00 | 0.00 | 3.58 |
| 6245 | 0.0016 | 0.00 | 0.00 | 3.57 |
| 6246 | 0.0013 | 0.00 | 0.00 | 3.57 |
| 6247 | 0.0006 | 0.00 | 0.00 | 3.57 |
| 6248 | 0.0016 | 0.00 | 0.00 | 3.57 |
| 6249 | 0.0026 | 0.00 | 0.00 | 3.57 |
| 6250 | 0.0016 | 0.00 | 0.00 | 3.57 |
| 6251 | 0.0003 | 0.00 | 0.00 | 3.56 |
| 6252 | 0.0006 | 0.00 | 0.00 | 3.56 |
| 6254 | 0.0003 | 0.00 | 0.00 | 3.56 |
| 6255 | 0.0006 | 0.00 | 0.00 | 3.56 |
| 6256 | 0.0006 | 0.00 | 0.00 | 3.56 |
| 6257 | 0.0003 | 0.00 | 0.00 | 3.56 |
| 6258 | 0.0016 | 0.00 | 0.00 | 3.56 |
| 6259 | 0.0003 | 0.00 | 0.00 | 3.56 |
| 6260 | 0.0016 | 0.00 | 0.00 | 3.56 |
| 6261 | 0.0013 | 0.00 | 0.00 | 3.56 |
| 6262 | 0.0019 | 0.00 | 0.00 | 3.56 |
| 6264 | 0.0003 | 0.00 | 0.00 | 3.55 |
| 6265 | 0.0010 | 0.00 | 0.00 | 3.55 |
| 6266 | 0.0013 | 0.00 | 0.00 | 3.55 |
| 6267 | 0.0003 | 0.00 | 0.00 | 3.55 |
| 6268 | 0.0006 | 0.00 | 0.00 | 3.55 |
| 6270 | 0.0010 | 0.00 | 0.00 | 3.55 |
| 6271 | 0.0013 | 0.00 | 0.00 | 3.55 |
| 6272 | 0.0010 | 0.00 | 0.00 | 3.55 |

| | | | | |
|---|---|---|---|---|
| 6273 | 0.0016 | 0.00 | 0.00 | 3.55 |
| 6275 | 0.0010 | 0.00 | 0.00 | 3.55 |
| 6276 | 0.0010 | 0.00 | 0.00 | 3.55 |
| 6277 | 0.0013 | 0.00 | 0.00 | 3.54 |
| 6278 | 0.0010 | 0.00 | 0.00 | 3.54 |
| 6279 | 0.0013 | 0.00 | 0.00 | 3.54 |
| 6280 | 0.0016 | 0.00 | 0.00 | 3.54 |
| 6281 | 0.0003 | 0.00 | 0.00 | 3.54 |
| 6282 | 0.0019 | 0.00 | 0.00 | 3.54 |
| 6283 | 0.0003 | 0.00 | 0.00 | 3.54 |
| 6284 | 0.0016 | 0.00 | 0.00 | 3.54 |
| 6286 | 0.0010 | 0.00 | 0.00 | 3.54 |
| 6287 | 0.0006 | 0.00 | 0.00 | 3.53 |
| 6288 | 0.0013 | 0.00 | 0.00 | 3.53 |
| 6289 | 0.0013 | 0.00 | 0.00 | 3.53 |
| 6290 | 0.0013 | 0.00 | 0.00 | 3.53 |
| 6291 | 0.0019 | 0.00 | 0.00 | 3.53 |
| 6292 | 0.0016 | 0.00 | 0.00 | 3.53 |
| 6293 | 0.0013 | 0.00 | 0.00 | 3.53 |
| 6294 | 0.0010 | 0.00 | 0.00 | 3.53 |
| 6295 | 0.0010 | 0.00 | 0.00 | 3.52 |
| 6296 | 0.0013 | 0.00 | 0.00 | 3.52 |
| 6297 | 0.0010 | 0.00 | 0.00 | 3.52 |
| 6298 | 0.0010 | 0.00 | 0.00 | 3.52 |
| 6299 | 0.0003 | 0.00 | 0.00 | 3.52 |
| 6300 | 0.0016 | 0.00 | 0.00 | 3.52 |
| 6301 | 0.0016 | 0.00 | 0.00 | 3.52 |
| 6302 | 0.0016 | 0.00 | 0.00 | 3.52 |
| 6303 | 0.0003 | 0.00 | 0.00 | 3.51 |
| 6304 | 0.0019 | 0.00 | 0.00 | 3.51 |
| 6305 | 0.0010 | 0.00 | 0.00 | 3.51 |
| 6306 | 0.0010 | 0.00 | 0.00 | 3.51 |
| 6307 | 0.0006 | 0.00 | 0.00 | 3.51 |
| 6308 | 0.0006 | 0.00 | 0.00 | 3.51 |
| 6309 | 0.0006 | 0.00 | 0.00 | 3.51 |
| 6310 | 0.0013 | 0.00 | 0.00 | 3.51 |
| 6311 | 0.0013 | 0.00 | 0.00 | 3.51 |
| 6312 | 0.0016 | 0.00 | 0.00 | 3.51 |
| 6313 | 0.0010 | 0.00 | 0.00 | 3.50 |
| 6314 | 0.0006 | 0.00 | 0.00 | 3.50 |
| 6315 | 0.0010 | 0.00 | 0.00 | 3.50 |
| 6316 | 0.0016 | 0.00 | 0.00 | 3.50 |
| 6317 | 0.0003 | 0.00 | 0.00 | 3.50 |
| 6318 | 0.0006 | 0.00 | 0.00 | 3.50 |

| | | | | |
|---|---|---|---|---|
| 6319 | 0.0013 | 0.00 | 0.00 | 3.50 |
| 6320 | 0.0016 | 0.00 | 0.00 | 3.50 |
| 6321 | 0.0003 | 0.00 | 0.00 | 3.50 |
| 6322 | 0.0016 | 0.00 | 0.00 | 3.50 |
| 6323 | 0.0019 | 0.00 | 0.00 | 3.49 |
| 6324 | 0.0010 | 0.00 | 0.00 | 3.49 |
| 6325 | 0.0003 | 0.00 | 0.00 | 3.49 |
| 6326 | 0.0010 | 0.00 | 0.00 | 3.49 |
| 6327 | 0.0006 | 0.00 | 0.00 | 3.49 |
| 6328 | 0.0006 | 0.00 | 0.00 | 3.49 |
| 6329 | 0.0006 | 0.00 | 0.00 | 3.49 |
| 6330 | 0.0006 | 0.00 | 0.00 | 3.49 |
| 6331 | 0.0010 | 0.00 | 0.00 | 3.49 |
| 6332 | 0.0010 | 0.00 | 0.00 | 3.49 |
| 6333 | 0.0010 | 0.00 | 0.00 | 3.49 |
| 6334 | 0.0019 | 0.00 | 0.00 | 3.48 |
| 6335 | 0.0013 | 0.00 | 0.00 | 3.48 |
| 6336 | 0.0006 | 0.00 | 0.00 | 3.48 |
| 6337 | 0.0010 | 0.00 | 0.00 | 3.48 |
| 6338 | 0.0003 | 0.00 | 0.00 | 3.48 |
| 6339 | 0.0013 | 0.00 | 0.00 | 3.48 |
| 6340 | 0.0010 | 0.00 | 0.00 | 3.48 |
| 6341 | 0.0003 | 0.00 | 0.00 | 3.48 |
| 6342 | 0.0013 | 0.00 | 0.00 | 3.48 |
| 6343 | 0.0006 | 0.00 | 0.00 | 3.48 |
| 6344 | 0.0013 | 0.00 | 0.00 | 3.48 |
| 6345 | 0.0010 | 0.00 | 0.00 | 3.47 |
| 6346 | 0.0006 | 0.00 | 0.00 | 3.47 |
| 6347 | 0.0006 | 0.00 | 0.00 | 3.47 |
| 6348 | 0.0003 | 0.00 | 0.00 | 3.47 |
| 6349 | 0.0010 | 0.00 | 0.00 | 3.47 |
| 6351 | 0.0006 | 0.00 | 0.00 | 3.47 |
| 6352 | 0.0010 | 0.00 | 0.00 | 3.47 |
| 6353 | 0.0006 | 0.00 | 0.00 | 3.47 |
| 6354 | 0.0006 | 0.00 | 0.00 | 3.47 |
| 6355 | 0.0019 | 0.00 | 0.00 | 3.47 |
| 6356 | 0.0006 | 0.00 | 0.00 | 3.47 |
| 6357 | 0.0019 | 0.00 | 0.00 | 3.47 |
| 6358 | 0.0016 | 0.00 | 0.00 | 3.46 |
| 6359 | 0.0006 | 0.00 | 0.00 | 3.46 |
| 6360 | 0.0003 | 0.00 | 0.00 | 3.46 |
| 6361 | 0.0006 | 0.00 | 0.00 | 3.46 |
| 6362 | 0.0016 | 0.00 | 0.00 | 3.46 |
| 6363 | 0.0006 | 0.00 | 0.00 | 3.46 |

| | | | | |
|---|---|---|---|---|
| 6364 | 0.0010 | 0.00 | 0.00 | 3.46 |
| 6365 | 0.0006 | 0.00 | 0.00 | 3.46 |
| 6366 | 0.0006 | 0.00 | 0.00 | 3.46 |
| 6367 | 0.0003 | 0.00 | 0.00 | 3.46 |
| 6368 | 0.0022 | 0.00 | 0.00 | 3.46 |
| 6369 | 0.0006 | 0.00 | 0.00 | 3.45 |
| 6370 | 0.0013 | 0.00 | 0.00 | 3.45 |
| 6371 | 0.0010 | 0.00 | 0.00 | 3.45 |
| 6372 | 0.0026 | 0.00 | 0.00 | 3.45 |
| 6373 | 0.0016 | 0.00 | 0.00 | 3.45 |
| 6374 | 0.0006 | 0.00 | 0.00 | 3.45 |
| 6375 | 0.0010 | 0.00 | 0.00 | 3.45 |
| 6376 | 0.0013 | 0.00 | 0.00 | 3.44 |
| 6377 | 0.0010 | 0.00 | 0.00 | 3.44 |
| 6378 | 0.0016 | 0.00 | 0.00 | 3.44 |
| 6379 | 0.0003 | 0.00 | 0.00 | 3.44 |
| 6380 | 0.0003 | 0.00 | 0.00 | 3.44 |
| 6381 | 0.0003 | 0.00 | 0.00 | 3.44 |
| 6382 | 0.0003 | 0.00 | 0.00 | 3.44 |
| 6383 | 0.0013 | 0.00 | 0.00 | 3.44 |
| 6384 | 0.0006 | 0.00 | 0.00 | 3.44 |
| 6385 | 0.0006 | 0.00 | 0.00 | 3.44 |
| 6386 | 0.0010 | 0.00 | 0.00 | 3.44 |
| 6387 | 0.0019 | 0.00 | 0.00 | 3.44 |
| 6388 | 0.0010 | 0.00 | 0.00 | 3.43 |
| 6389 | 0.0016 | 0.00 | 0.00 | 3.43 |
| 6390 | 0.0010 | 0.00 | 0.00 | 3.43 |
| 6391 | 0.0010 | 0.00 | 0.00 | 3.43 |
| 6392 | 0.0010 | 0.00 | 0.00 | 3.43 |
| 6393 | 0.0016 | 0.00 | 0.00 | 3.43 |
| 6394 | 0.0003 | 0.00 | 0.00 | 3.43 |
| 6395 | 0.0006 | 0.00 | 0.00 | 3.43 |
| 6397 | 0.0010 | 0.00 | 0.00 | 3.43 |
| 6398 | 0.0016 | 0.00 | 0.00 | 3.42 |
| 6399 | 0.0010 | 0.00 | 0.00 | 3.42 |
| 6400 | 0.0013 | 0.00 | 0.00 | 3.42 |
| 6401 | 0.0003 | 0.00 | 0.00 | 3.42 |
| 6402 | 0.0013 | 0.00 | 0.00 | 3.42 |
| 6403 | 0.0010 | 0.00 | 0.00 | 3.42 |
| 6404 | 0.0010 | 0.00 | 0.00 | 3.42 |
| 6405 | 0.0016 | 0.00 | 0.00 | 3.42 |
| 6406 | 0.0019 | 0.00 | 0.00 | 3.42 |
| 6407 | 0.0006 | 0.00 | 0.00 | 3.41 |
| 6408 | 0.0003 | 0.00 | 0.00 | 3.41 |

| | | | | |
|---|---|---|---|---|
| 6409 | 0.0006 | 0.00 | 0.00 | 3.41 |
| 6410 | 0.0010 | 0.00 | 0.00 | 3.41 |
| 6411 | 0.0013 | 0.00 | 0.00 | 3.41 |
| 6412 | 0.0006 | 0.00 | 0.00 | 3.41 |
| 6413 | 0.0006 | 0.00 | 0.00 | 3.41 |
| 6415 | 0.0013 | 0.00 | 0.00 | 3.41 |
| 6416 | 0.0013 | 0.00 | 0.00 | 3.41 |
| 6417 | 0.0006 | 0.00 | 0.00 | 3.41 |
| 6418 | 0.0006 | 0.00 | 0.00 | 3.41 |
| 6419 | 0.0016 | 0.00 | 0.00 | 3.40 |
| 6420 | 0.0010 | 0.00 | 0.00 | 3.40 |
| 6421 | 0.0010 | 0.00 | 0.00 | 3.40 |
| 6422 | 0.0003 | 0.00 | 0.00 | 3.40 |
| 6423 | 0.0006 | 0.00 | 0.00 | 3.40 |
| 6424 | 0.0003 | 0.00 | 0.00 | 3.40 |
| 6425 | 0.0006 | 0.00 | 0.00 | 3.40 |
| 6426 | 0.0010 | 0.00 | 0.00 | 3.40 |
| 6427 | 0.0006 | 0.00 | 0.00 | 3.40 |
| 6428 | 0.0019 | 0.00 | 0.00 | 3.40 |
| 6429 | 0.0022 | 0.00 | 0.00 | 3.40 |
| 6430 | 0.0006 | 0.00 | 0.00 | 3.39 |
| 6431 | 0.0006 | 0.00 | 0.00 | 3.39 |
| 6432 | 0.0006 | 0.00 | 0.00 | 3.39 |
| 6433 | 0.0003 | 0.00 | 0.00 | 3.39 |
| 6435 | 0.0006 | 0.00 | 0.00 | 3.39 |
| 6436 | 0.0013 | 0.00 | 0.00 | 3.39 |
| 6437 | 0.0006 | 0.00 | 0.00 | 3.39 |
| 6438 | 0.0019 | 0.00 | 0.00 | 3.39 |
| 6439 | 0.0010 | 0.00 | 0.00 | 3.39 |
| 6440 | 0.0010 | 0.00 | 0.00 | 3.39 |
| 6441 | 0.0003 | 0.00 | 0.00 | 3.38 |
| 6442 | 0.0016 | 0.00 | 0.00 | 3.38 |
| 6443 | 0.0006 | 0.00 | 0.00 | 3.38 |
| 6444 | 0.0016 | 0.00 | 0.00 | 3.38 |
| 6445 | 0.0019 | 0.00 | 0.00 | 3.38 |
| 6446 | 0.0016 | 0.00 | 0.00 | 3.38 |
| 6447 | 0.0013 | 0.00 | 0.00 | 3.38 |
| 6448 | 0.0019 | 0.00 | 0.00 | 3.38 |
| 6449 | 0.0010 | 0.00 | 0.00 | 3.37 |
| 6450 | 0.0019 | 0.00 | 0.00 | 3.37 |
| 6451 | 0.0003 | 0.00 | 0.00 | 3.37 |
| 6452 | 0.0013 | 0.00 | 0.00 | 3.37 |
| 6453 | 0.0003 | 0.00 | 0.00 | 3.37 |
| 6454 | 0.0010 | 0.00 | 0.00 | 3.37 |

| | | | | |
|---|---|---|---|---|
| 6455 | 0.0003 | 0.00 | 0.00 | 3.37 |
| 6456 | 0.0019 | 0.00 | 0.00 | 3.37 |
| 6458 | 0.0013 | 0.00 | 0.00 | 3.37 |
| 6459 | 0.0006 | 0.00 | 0.00 | 3.36 |
| 6460 | 0.0006 | 0.00 | 0.00 | 3.36 |
| 6461 | 0.0003 | 0.00 | 0.00 | 3.36 |
| 6462 | 0.0010 | 0.00 | 0.00 | 3.36 |
| 6463 | 0.0010 | 0.00 | 0.00 | 3.36 |
| 6464 | 0.0006 | 0.00 | 0.00 | 3.36 |
| 6465 | 0.0019 | 0.00 | 0.00 | 3.36 |
| 6466 | 0.0010 | 0.00 | 0.00 | 3.36 |
| 6467 | 0.0016 | 0.00 | 0.00 | 3.36 |
| 6468 | 0.0013 | 0.00 | 0.00 | 3.36 |
| 6469 | 0.0016 | 0.00 | 0.00 | 3.35 |
| 6470 | 0.0006 | 0.00 | 0.00 | 3.35 |
| 6471 | 0.0003 | 0.00 | 0.00 | 3.35 |
| 6472 | 0.0010 | 0.00 | 0.00 | 3.35 |
| 6473 | 0.0006 | 0.00 | 0.00 | 3.35 |
| 6474 | 0.0003 | 0.00 | 0.00 | 3.35 |
| 6475 | 0.0019 | 0.00 | 0.00 | 3.35 |
| 6478 | 0.0006 | 0.00 | 0.00 | 3.35 |
| 6479 | 0.0013 | 0.00 | 0.00 | 3.35 |
| 6480 | 0.0003 | 0.00 | 0.00 | 3.35 |
| 6481 | 0.0003 | 0.00 | 0.00 | 3.35 |
| 6482 | 0.0006 | 0.00 | 0.00 | 3.35 |
| 6483 | 0.0010 | 0.00 | 0.00 | 3.35 |
| 6484 | 0.0010 | 0.00 | 0.00 | 3.34 |
| 6485 | 0.0010 | 0.00 | 0.00 | 3.34 |
| 6486 | 0.0013 | 0.00 | 0.00 | 3.34 |
| 6487 | 0.0006 | 0.00 | 0.00 | 3.34 |
| 6488 | 0.0003 | 0.00 | 0.00 | 3.34 |
| 6489 | 0.0019 | 0.00 | 0.00 | 3.34 |
| 6490 | 0.0006 | 0.00 | 0.00 | 3.34 |
| 6491 | 0.0006 | 0.00 | 0.00 | 3.34 |
| 6492 | 0.0006 | 0.00 | 0.00 | 3.34 |
| 6493 | 0.0006 | 0.00 | 0.00 | 3.34 |
| 6494 | 0.0016 | 0.00 | 0.00 | 3.34 |
| 6495 | 0.0016 | 0.00 | 0.00 | 3.33 |
| 6496 | 0.0010 | 0.00 | 0.00 | 3.33 |
| 6497 | 0.0010 | 0.00 | 0.00 | 3.33 |
| 6498 | 0.0006 | 0.00 | 0.00 | 3.33 |
| 6499 | 0.0019 | 0.00 | 0.00 | 3.33 |
| 6500 | 0.0013 | 0.00 | 0.00 | 3.33 |
| 6501 | 0.0010 | 0.00 | 0.00 | 3.33 |

| | | | | |
|---|---|---|---|---|
| 6502 | 0.0003 | 0.00 | 0.00 | 3.33 |
| 6503 | 0.0013 | 0.00 | 0.00 | 3.33 |
| 6505 | 0.0003 | 0.00 | 0.00 | 3.32 |
| 6506 | 0.0003 | 0.00 | 0.00 | 3.32 |
| 6507 | 0.0006 | 0.00 | 0.00 | 3.32 |
| 6508 | 0.0003 | 0.00 | 0.00 | 3.32 |
| 6509 | 0.0010 | 0.00 | 0.00 | 3.32 |
| 6510 | 0.0016 | 0.00 | 0.00 | 3.32 |
| 6511 | 0.0010 | 0.00 | 0.00 | 3.32 |
| 6512 | 0.0013 | 0.00 | 0.00 | 3.32 |
| 6513 | 0.0003 | 0.00 | 0.00 | 3.32 |
| 6514 | 0.0006 | 0.00 | 0.00 | 3.32 |
| 6515 | 0.0003 | 0.00 | 0.00 | 3.32 |
| 6518 | 0.0010 | 0.00 | 0.00 | 3.32 |
| 6519 | 0.0010 | 0.00 | 0.00 | 3.32 |
| 6520 | 0.0003 | 0.00 | 0.00 | 3.31 |
| 6521 | 0.0003 | 0.00 | 0.00 | 3.31 |
| 6522 | 0.0016 | 0.00 | 0.00 | 3.31 |
| 6523 | 0.0006 | 0.00 | 0.00 | 3.31 |
| 6524 | 0.0006 | 0.00 | 0.00 | 3.31 |
| 6525 | 0.0013 | 0.00 | 0.00 | 3.31 |
| 6529 | 0.0013 | 0.00 | 0.00 | 3.31 |
| 6531 | 0.0013 | 0.00 | 0.00 | 3.31 |
| 6532 | 0.0003 | 0.00 | 0.00 | 3.31 |
| 6533 | 0.0013 | 0.00 | 0.00 | 3.31 |
| 6534 | 0.0010 | 0.00 | 0.00 | 3.31 |
| 6535 | 0.0006 | 0.00 | 0.00 | 3.30 |
| 6536 | 0.0010 | 0.00 | 0.00 | 3.30 |
| 6537 | 0.0010 | 0.00 | 0.00 | 3.30 |
| 6538 | 0.0003 | 0.00 | 0.00 | 3.30 |
| 6539 | 0.0013 | 0.00 | 0.00 | 3.30 |
| 6540 | 0.0010 | 0.00 | 0.00 | 3.30 |
| 6541 | 0.0003 | 0.00 | 0.00 | 3.30 |
| 6542 | 0.0010 | 0.00 | 0.00 | 3.30 |
| 6543 | 0.0006 | 0.00 | 0.00 | 3.30 |
| 6544 | 0.0006 | 0.00 | 0.00 | 3.30 |
| 6545 | 0.0010 | 0.00 | 0.00 | 3.30 |
| 6546 | 0.0013 | 0.00 | 0.00 | 3.30 |
| 6547 | 0.0022 | 0.00 | 0.00 | 3.29 |
| 6548 | 0.0003 | 0.00 | 0.00 | 3.29 |
| 6549 | 0.0006 | 0.00 | 0.00 | 3.29 |
| 6550 | 0.0016 | 0.00 | 0.00 | 3.29 |
| 6551 | 0.0006 | 0.00 | 0.00 | 3.29 |
| 6552 | 0.0010 | 0.00 | 0.00 | 3.29 |

| | | | | |
|---|---|---|---|---|
| 6553 | 0.0006 | 0.00 | 0.00 | 3.29 |
| 6554 | 0.0010 | 0.00 | 0.00 | 3.29 |
| 6555 | 0.0006 | 0.00 | 0.00 | 3.29 |
| 6556 | 0.0010 | 0.00 | 0.00 | 3.29 |
| 6558 | 0.0010 | 0.00 | 0.00 | 3.28 |
| 6559 | 0.0010 | 0.00 | 0.00 | 3.28 |
| 6560 | 0.0010 | 0.00 | 0.00 | 3.28 |
| 6561 | 0.0010 | 0.00 | 0.00 | 3.28 |
| 6562 | 0.0003 | 0.00 | 0.00 | 3.28 |
| 6563 | 0.0019 | 0.00 | 0.00 | 3.28 |
| 6564 | 0.0013 | 0.00 | 0.00 | 3.28 |
| 6565 | 0.0013 | 0.00 | 0.00 | 3.28 |
| 6566 | 0.0013 | 0.00 | 0.00 | 3.28 |
| 6567 | 0.0010 | 0.00 | 0.00 | 3.28 |
| 6568 | 0.0013 | 0.00 | 0.00 | 3.27 |
| 6569 | 0.0006 | 0.00 | 0.00 | 3.27 |
| 6570 | 0.0013 | 0.00 | 0.00 | 3.27 |
| 6571 | 0.0010 | 0.00 | 0.00 | 3.27 |
| 6572 | 0.0019 | 0.00 | 0.00 | 3.27 |
| 6573 | 0.0006 | 0.00 | 0.00 | 3.27 |
| 6574 | 0.0010 | 0.00 | 0.00 | 3.27 |
| 6575 | 0.0016 | 0.00 | 0.00 | 3.27 |
| 6576 | 0.0013 | 0.00 | 0.00 | 3.26 |
| 6577 | 0.0006 | 0.00 | 0.00 | 3.26 |
| 6578 | 0.0013 | 0.00 | 0.00 | 3.26 |
| 6579 | 0.0003 | 0.00 | 0.00 | 3.26 |
| 6580 | 0.0003 | 0.00 | 0.00 | 3.26 |
| 6581 | 0.0022 | 0.00 | 0.00 | 3.26 |
| 6582 | 0.0006 | 0.00 | 0.00 | 3.26 |
| 6583 | 0.0006 | 0.00 | 0.00 | 3.26 |
| 6584 | 0.0003 | 0.00 | 0.00 | 3.26 |
| 6585 | 0.0006 | 0.00 | 0.00 | 3.26 |
| 6586 | 0.0010 | 0.00 | 0.00 | 3.26 |
| 6587 | 0.0006 | 0.00 | 0.00 | 3.26 |
| 6588 | 0.0013 | 0.00 | 0.00 | 3.25 |
| 6589 | 0.0010 | 0.00 | 0.00 | 3.25 |
| 6590 | 0.0010 | 0.00 | 0.00 | 3.25 |
| 6591 | 0.0006 | 0.00 | 0.00 | 3.25 |
| 6593 | 0.0010 | 0.00 | 0.00 | 3.25 |
| 6594 | 0.0013 | 0.00 | 0.00 | 3.25 |
| 6595 | 0.0016 | 0.00 | 0.00 | 3.25 |
| 6596 | 0.0003 | 0.00 | 0.00 | 3.25 |
| 6597 | 0.0010 | 0.00 | 0.00 | 3.25 |
| 6598 | 0.0010 | 0.00 | 0.00 | 3.25 |

| | | | | |
|---|---|---|---|---|
| 6599 | 0.0010 | 0.00 | 0.00 | 3.24 |
| 6600 | 0.0006 | 0.00 | 0.00 | 3.24 |
| 6601 | 0.0013 | 0.00 | 0.00 | 3.24 |
| 6602 | 0.0016 | 0.00 | 0.00 | 3.24 |
| 6604 | 0.0003 | 0.00 | 0.00 | 3.24 |
| 6605 | 0.0013 | 0.00 | 0.00 | 3.24 |
| 6606 | 0.0006 | 0.00 | 0.00 | 3.24 |
| 6607 | 0.0006 | 0.00 | 0.00 | 3.24 |
| 6608 | 0.0006 | 0.00 | 0.00 | 3.24 |
| 6609 | 0.0016 | 0.00 | 0.00 | 3.24 |
| 6610 | 0.0006 | 0.00 | 0.00 | 3.24 |
| 6611 | 0.0006 | 0.00 | 0.00 | 3.23 |
| 6612 | 0.0016 | 0.00 | 0.00 | 3.23 |
| 6613 | 0.0010 | 0.00 | 0.00 | 3.23 |
| 6614 | 0.0013 | 0.00 | 0.00 | 3.23 |
| 6615 | 0.0003 | 0.00 | 0.00 | 3.23 |
| 6616 | 0.0016 | 0.00 | 0.00 | 3.23 |
| 6617 | 0.0010 | 0.00 | 0.00 | 3.23 |
| 6618 | 0.0010 | 0.00 | 0.00 | 3.23 |
| 6619 | 0.0010 | 0.00 | 0.00 | 3.23 |
| 6620 | 0.0003 | 0.00 | 0.00 | 3.23 |
| 6621 | 0.0003 | 0.00 | 0.00 | 3.23 |
| 6622 | 0.0006 | 0.00 | 0.00 | 3.22 |
| 6623 | 0.0006 | 0.00 | 0.00 | 3.22 |
| 6624 | 0.0010 | 0.00 | 0.00 | 3.22 |
| 6625 | 0.0003 | 0.00 | 0.00 | 3.22 |
| 6626 | 0.0003 | 0.00 | 0.00 | 3.22 |
| 6627 | 0.0010 | 0.00 | 0.00 | 3.22 |
| 6628 | 0.0006 | 0.00 | 0.00 | 3.22 |
| 6629 | 0.0006 | 0.00 | 0.00 | 3.22 |
| 6630 | 0.0016 | 0.00 | 0.00 | 3.22 |
| 6631 | 0.0010 | 0.00 | 0.00 | 3.22 |
| 6632 | 0.0010 | 0.00 | 0.00 | 3.22 |
| 6633 | 0.0010 | 0.00 | 0.00 | 3.22 |
| 6634 | 0.0016 | 0.00 | 0.00 | 3.22 |
| 6635 | 0.0006 | 0.00 | 0.00 | 3.21 |
| 6636 | 0.0006 | 0.00 | 0.00 | 3.21 |
| 6637 | 0.0006 | 0.00 | 0.00 | 3.21 |
| 6638 | 0.0003 | 0.00 | 0.00 | 3.21 |
| 6639 | 0.0010 | 0.00 | 0.00 | 3.21 |
| 6640 | 0.0003 | 0.00 | 0.00 | 3.21 |
| 6641 | 0.0010 | 0.00 | 0.00 | 3.21 |
| 6642 | 0.0013 | 0.00 | 0.00 | 3.21 |
| 6643 | 0.0010 | 0.00 | 0.00 | 3.21 |

| | | | | |
|---|---|---|---|---|
| 6644 | 0.0006 | 0.00 | 0.00 | 3.21 |
| 6645 | 0.0019 | 0.00 | 0.00 | 3.21 |
| 6646 | 0.0006 | 0.00 | 0.00 | 3.20 |
| 6647 | 0.0013 | 0.00 | 0.00 | 3.20 |
| 6649 | 0.0010 | 0.00 | 0.00 | 3.20 |
| 6650 | 0.0010 | 0.00 | 0.00 | 3.20 |
| 6651 | 0.0006 | 0.00 | 0.00 | 3.20 |
| 6652 | 0.0013 | 0.00 | 0.00 | 3.20 |
| 6653 | 0.0010 | 0.00 | 0.00 | 3.20 |
| 6654 | 0.0010 | 0.00 | 0.00 | 3.20 |
| 6655 | 0.0003 | 0.00 | 0.00 | 3.20 |
| 6656 | 0.0006 | 0.00 | 0.00 | 3.20 |
| 6657 | 0.0013 | 0.00 | 0.00 | 3.20 |
| 6658 | 0.0013 | 0.00 | 0.00 | 3.19 |
| 6659 | 0.0003 | 0.00 | 0.00 | 3.19 |
| 6660 | 0.0010 | 0.00 | 0.00 | 3.19 |
| 6661 | 0.0010 | 0.00 | 0.00 | 3.19 |
| 6662 | 0.0003 | 0.00 | 0.00 | 3.19 |
| 6663 | 0.0010 | 0.00 | 0.00 | 3.19 |
| 6664 | 0.0006 | 0.00 | 0.00 | 3.19 |
| 6665 | 0.0013 | 0.00 | 0.00 | 3.19 |
| 6666 | 0.0010 | 0.00 | 0.00 | 3.19 |
| 6667 | 0.0013 | 0.00 | 0.00 | 3.19 |
| 6668 | 0.0013 | 0.00 | 0.00 | 3.19 |
| 6669 | 0.0013 | 0.00 | 0.00 | 3.18 |
| 6670 | 0.0013 | 0.00 | 0.00 | 3.18 |
| 6671 | 0.0016 | 0.00 | 0.00 | 3.18 |
| 6673 | 0.0010 | 0.00 | 0.00 | 3.18 |
| 6674 | 0.0006 | 0.00 | 0.00 | 3.18 |
| 6675 | 0.0010 | 0.00 | 0.00 | 3.18 |
| 6676 | 0.0006 | 0.00 | 0.00 | 3.18 |
| 6677 | 0.0013 | 0.00 | 0.00 | 3.18 |
| 6678 | 0.0003 | 0.00 | 0.00 | 3.18 |
| 6679 | 0.0010 | 0.00 | 0.00 | 3.18 |
| 6681 | 0.0010 | 0.00 | 0.00 | 3.17 |
| 6683 | 0.0006 | 0.00 | 0.00 | 3.17 |
| 6684 | 0.0010 | 0.00 | 0.00 | 3.17 |
| 6685 | 0.0029 | 0.00 | 0.00 | 3.17 |
| 6686 | 0.0013 | 0.00 | 0.00 | 3.17 |
| 6687 | 0.0016 | 0.00 | 0.00 | 3.17 |
| 6688 | 0.0006 | 0.00 | 0.00 | 3.17 |
| 6689 | 0.0019 | 0.00 | 0.00 | 3.17 |
| 6690 | 0.0003 | 0.00 | 0.00 | 3.16 |
| 6691 | 0.0006 | 0.00 | 0.00 | 3.16 |

| | | | | |
|---|---|---|---|---|
| 6692 | 0.0016 | 0.00 | 0.00 | 3.16 |
| 6693 | 0.0013 | 0.00 | 0.00 | 3.16 |
| 6694 | 0.0006 | 0.00 | 0.00 | 3.16 |
| 6695 | 0.0003 | 0.00 | 0.00 | 3.16 |
| 6696 | 0.0019 | 0.00 | 0.00 | 3.16 |
| 6697 | 0.0003 | 0.00 | 0.00 | 3.16 |
| 6699 | 0.0016 | 0.00 | 0.00 | 3.16 |
| 6700 | 0.0016 | 0.00 | 0.00 | 3.15 |
| 6701 | 0.0006 | 0.00 | 0.00 | 3.15 |
| 6702 | 0.0013 | 0.00 | 0.00 | 3.15 |
| 6703 | 0.0010 | 0.00 | 0.00 | 3.15 |
| 6704 | 0.0003 | 0.00 | 0.00 | 3.15 |
| 6706 | 0.0006 | 0.00 | 0.00 | 3.15 |
| 6707 | 0.0006 | 0.00 | 0.00 | 3.15 |
| 6708 | 0.0006 | 0.00 | 0.00 | 3.15 |
| 6709 | 0.0010 | 0.00 | 0.00 | 3.15 |
| 6711 | 0.0013 | 0.00 | 0.00 | 3.15 |
| 6712 | 0.0010 | 0.00 | 0.00 | 3.15 |
| 6713 | 0.0006 | 0.00 | 0.00 | 3.14 |
| 6714 | 0.0006 | 0.00 | 0.00 | 3.14 |
| 6715 | 0.0013 | 0.00 | 0.00 | 3.14 |
| 6716 | 0.0010 | 0.00 | 0.00 | 3.14 |
| 6717 | 0.0003 | 0.00 | 0.00 | 3.14 |
| 6719 | 0.0003 | 0.00 | 0.00 | 3.14 |
| 6720 | 0.0010 | 0.00 | 0.00 | 3.14 |
| 6721 | 0.0006 | 0.00 | 0.00 | 3.14 |
| 6722 | 0.0016 | 0.00 | 0.00 | 3.14 |
| 6723 | 0.0006 | 0.00 | 0.00 | 3.14 |
| 6724 | 0.0003 | 0.00 | 0.00 | 3.14 |
| 6725 | 0.0010 | 0.00 | 0.00 | 3.14 |
| 6726 | 0.0013 | 0.00 | 0.00 | 3.14 |
| 6727 | 0.0013 | 0.00 | 0.00 | 3.13 |
| 6728 | 0.0006 | 0.00 | 0.00 | 3.13 |
| 6729 | 0.0019 | 0.00 | 0.00 | 3.13 |
| 6730 | 0.0006 | 0.00 | 0.00 | 3.13 |
| 6731 | 0.0006 | 0.00 | 0.00 | 3.13 |
| 6732 | 0.0016 | 0.00 | 0.00 | 3.13 |
| 6733 | 0.0006 | 0.00 | 0.00 | 3.13 |
| 6734 | 0.0006 | 0.00 | 0.00 | 3.13 |
| 6735 | 0.0016 | 0.00 | 0.00 | 3.13 |
| 6737 | 0.0003 | 0.00 | 0.00 | 3.12 |
| 6738 | 0.0003 | 0.00 | 0.00 | 3.12 |
| 6739 | 0.0019 | 0.00 | 0.00 | 3.12 |
| 6740 | 0.0019 | 0.00 | 0.00 | 3.12 |

| | | | | |
|---|---|---|---|---|
| 6741 | 0.0013 | 0.00 | 0.00 | 3.12 |
| 6742 | 0.0006 | 0.00 | 0.00 | 3.12 |
| 6743 | 0.0006 | 0.00 | 0.00 | 3.12 |
| 6744 | 0.0013 | 0.00 | 0.00 | 3.12 |
| 6745 | 0.0010 | 0.00 | 0.00 | 3.12 |
| 6746 | 0.0006 | 0.00 | 0.00 | 3.12 |
| 6747 | 0.0010 | 0.00 | 0.00 | 3.11 |
| 6748 | 0.0006 | 0.00 | 0.00 | 3.11 |
| 6749 | 0.0016 | 0.00 | 0.00 | 3.11 |
| 6750 | 0.0013 | 0.00 | 0.00 | 3.11 |
| 6753 | 0.0010 | 0.00 | 0.00 | 3.11 |
| 6754 | 0.0006 | 0.00 | 0.00 | 3.11 |
| 6755 | 0.0006 | 0.00 | 0.00 | 3.11 |
| 6756 | 0.0003 | 0.00 | 0.00 | 3.11 |
| 6757 | 0.0006 | 0.00 | 0.00 | 3.11 |
| 6758 | 0.0006 | 0.00 | 0.00 | 3.11 |
| 6760 | 0.0010 | 0.00 | 0.00 | 3.11 |
| 6761 | 0.0010 | 0.00 | 0.00 | 3.11 |
| 6763 | 0.0013 | 0.00 | 0.00 | 3.10 |
| 6764 | 0.0010 | 0.00 | 0.00 | 3.10 |
| 6765 | 0.0010 | 0.00 | 0.00 | 3.10 |
| 6766 | 0.0010 | 0.00 | 0.00 | 3.10 |
| 6767 | 0.0013 | 0.00 | 0.00 | 3.10 |
| 6768 | 0.0006 | 0.00 | 0.00 | 3.10 |
| 6769 | 0.0013 | 0.00 | 0.00 | 3.10 |
| 6770 | 0.0003 | 0.00 | 0.00 | 3.10 |
| 6771 | 0.0006 | 0.00 | 0.00 | 3.10 |
| 6772 | 0.0010 | 0.00 | 0.00 | 3.10 |
| 6773 | 0.0010 | 0.00 | 0.00 | 3.09 |
| 6774 | 0.0006 | 0.00 | 0.00 | 3.09 |
| 6775 | 0.0003 | 0.00 | 0.00 | 3.09 |
| 6776 | 0.0010 | 0.00 | 0.00 | 3.09 |
| 6777 | 0.0019 | 0.00 | 0.00 | 3.09 |
| 6778 | 0.0016 | 0.00 | 0.00 | 3.09 |
| 6779 | 0.0006 | 0.00 | 0.00 | 3.09 |
| 6780 | 0.0019 | 0.00 | 0.00 | 3.09 |
| 6781 | 0.0013 | 0.00 | 0.00 | 3.09 |
| 6782 | 0.0003 | 0.00 | 0.00 | 3.08 |
| 6783 | 0.0003 | 0.00 | 0.00 | 3.08 |
| 6784 | 0.0016 | 0.00 | 0.00 | 3.08 |
| 6785 | 0.0016 | 0.00 | 0.00 | 3.08 |
| 6786 | 0.0016 | 0.00 | 0.00 | 3.08 |
| 6787 | 0.0013 | 0.00 | 0.00 | 3.08 |
| 6788 | 0.0003 | 0.00 | 0.00 | 3.08 |

| | | | | |
|---|---|---|---|---|
| 6789 | 0.0010 | 0.00 | 0.00 | 3.08 |
| 6790 | 0.0010 | 0.00 | 0.00 | 3.08 |
| 6792 | 0.0010 | 0.00 | 0.00 | 3.08 |
| 6793 | 0.0013 | 0.00 | 0.00 | 3.07 |
| 6794 | 0.0019 | 0.00 | 0.00 | 3.07 |
| 6795 | 0.0006 | 0.00 | 0.00 | 3.07 |
| 6796 | 0.0010 | 0.00 | 0.00 | 3.07 |
| 6797 | 0.0016 | 0.00 | 0.00 | 3.07 |
| 6798 | 0.0010 | 0.00 | 0.00 | 3.07 |
| 6799 | 0.0010 | 0.00 | 0.00 | 3.07 |
| 6800 | 0.0013 | 0.00 | 0.00 | 3.07 |
| 6801 | 0.0006 | 0.00 | 0.00 | 3.07 |
| 6802 | 0.0013 | 0.00 | 0.00 | 3.06 |
| 6803 | 0.0003 | 0.00 | 0.00 | 3.06 |
| 6804 | 0.0003 | 0.00 | 0.00 | 3.06 |
| 6805 | 0.0006 | 0.00 | 0.00 | 3.06 |
| 6806 | 0.0006 | 0.00 | 0.00 | 3.06 |
| 6807 | 0.0016 | 0.00 | 0.00 | 3.06 |
| 6808 | 0.0006 | 0.00 | 0.00 | 3.06 |
| 6809 | 0.0013 | 0.00 | 0.00 | 3.06 |
| 6810 | 0.0022 | 0.00 | 0.00 | 3.06 |
| 6811 | 0.0010 | 0.00 | 0.00 | 3.06 |
| 6812 | 0.0003 | 0.00 | 0.00 | 3.05 |
| 6813 | 0.0010 | 0.00 | 0.00 | 3.05 |
| 6815 | 0.0003 | 0.00 | 0.00 | 3.05 |
| 6817 | 0.0013 | 0.00 | 0.00 | 3.05 |
| 6819 | 0.0022 | 0.00 | 0.00 | 3.05 |
| 6820 | 0.0022 | 0.00 | 0.00 | 3.05 |
| 6821 | 0.0006 | 0.00 | 0.00 | 3.05 |
| 6822 | 0.0003 | 0.00 | 0.00 | 3.05 |
| 6823 | 0.0006 | 0.00 | 0.00 | 3.05 |
| 6824 | 0.0016 | 0.00 | 0.00 | 3.05 |
| 6826 | 0.0013 | 0.00 | 0.00 | 3.04 |
| 6827 | 0.0013 | 0.00 | 0.00 | 3.04 |
| 6828 | 0.0006 | 0.00 | 0.00 | 3.04 |
| 6829 | 0.0019 | 0.00 | 0.00 | 3.04 |
| 6830 | 0.0010 | 0.00 | 0.00 | 3.04 |
| 6831 | 0.0013 | 0.00 | 0.00 | 3.04 |
| 6832 | 0.0006 | 0.00 | 0.00 | 3.04 |
| 6833 | 0.0010 | 0.00 | 0.00 | 3.04 |
| 6834 | 0.0003 | 0.00 | 0.00 | 3.03 |
| 6835 | 0.0010 | 0.00 | 0.00 | 3.03 |
| 6836 | 0.0006 | 0.00 | 0.00 | 3.03 |
| 6837 | 0.0006 | 0.00 | 0.00 | 3.03 |

| | | | | |
|---|---|---|---|---|
| 6838 | 0.0006 | 0.00 | 0.00 | 3.03 |
| 6839 | 0.0003 | 0.00 | 0.00 | 3.03 |
| 6840 | 0.0010 | 0.00 | 0.00 | 3.03 |
| 6841 | 0.0006 | 0.00 | 0.00 | 3.03 |
| 6842 | 0.0013 | 0.00 | 0.00 | 3.03 |
| 6843 | 0.0003 | 0.00 | 0.00 | 3.03 |
| 6844 | 0.0006 | 0.00 | 0.00 | 3.03 |
| 6845 | 0.0016 | 0.00 | 0.00 | 3.03 |
| 6846 | 0.0010 | 0.00 | 0.00 | 3.03 |
| 6847 | 0.0010 | 0.00 | 0.00 | 3.03 |
| 6848 | 0.0003 | 0.00 | 0.00 | 3.02 |
| 6849 | 0.0013 | 0.00 | 0.00 | 3.02 |
| 6850 | 0.0010 | 0.00 | 0.00 | 3.02 |
| 6851 | 0.0003 | 0.00 | 0.00 | 3.02 |
| 6852 | 0.0006 | 0.00 | 0.00 | 3.02 |
| 6853 | 0.0010 | 0.00 | 0.00 | 3.02 |
| 6854 | 0.0010 | 0.00 | 0.00 | 3.02 |
| 6855 | 0.0019 | 0.00 | 0.00 | 3.02 |
| 6856 | 0.0006 | 0.00 | 0.00 | 3.02 |
| 6857 | 0.0013 | 0.00 | 0.00 | 3.02 |
| 6858 | 0.0006 | 0.00 | 0.00 | 3.01 |
| 6859 | 0.0019 | 0.00 | 0.00 | 3.01 |
| 6860 | 0.0010 | 0.00 | 0.00 | 3.01 |
| 6861 | 0.0016 | 0.00 | 0.00 | 3.01 |
| 6862 | 0.0003 | 0.00 | 0.00 | 3.01 |
| 6863 | 0.0010 | 0.00 | 0.00 | 3.01 |
| 6864 | 0.0010 | 0.00 | 0.00 | 3.01 |
| 6865 | 0.0003 | 0.00 | 0.00 | 3.01 |
| 6866 | 0.0006 | 0.00 | 0.00 | 3.01 |
| 6867 | 0.0003 | 0.00 | 0.00 | 3.01 |
| 6868 | 0.0006 | 0.00 | 0.00 | 3.01 |
| 6869 | 0.0006 | 0.00 | 0.00 | 3.01 |
| 6870 | 0.0010 | 0.00 | 0.00 | 3.00 |
| 6871 | 0.0013 | 0.00 | 0.00 | 3.00 |
| 6872 | 0.0003 | 0.00 | 0.00 | 3.00 |
| 6873 | 0.0006 | 0.00 | 0.00 | 3.00 |
| 6874 | 0.0016 | 0.00 | 0.00 | 3.00 |
| 6875 | 0.0013 | 0.00 | 0.00 | 3.00 |
| 6876 | 0.0003 | 0.00 | 0.00 | 3.00 |
| 6877 | 0.0010 | 0.00 | 0.00 | 3.00 |
| 6879 | 0.0013 | 0.00 | 0.00 | 3.00 |
| 6880 | 0.0006 | 0.00 | 0.00 | 3.00 |
| 6881 | 0.0010 | 0.00 | 0.00 | 3.00 |
| 6882 | 0.0013 | 0.00 | 0.00 | 2.99 |

| | | | | |
|---|---|---|---|---|
| 6883 | 0.0006 | 0.00 | 0.00 | 2.99 |
| 6884 | 0.0006 | 0.00 | 0.00 | 2.99 |
| 6885 | 0.0003 | 0.00 | 0.00 | 2.99 |
| 6886 | 0.0013 | 0.00 | 0.00 | 2.99 |
| 6887 | 0.0016 | 0.00 | 0.00 | 2.99 |
| 6888 | 0.0016 | 0.00 | 0.00 | 2.99 |
| 6889 | 0.0006 | 0.00 | 0.00 | 2.99 |
| 6890 | 0.0003 | 0.00 | 0.00 | 2.99 |
| 6891 | 0.0003 | 0.00 | 0.00 | 2.99 |
| 6892 | 0.0010 | 0.00 | 0.00 | 2.99 |
| 6893 | 0.0013 | 0.00 | 0.00 | 2.98 |
| 6894 | 0.0003 | 0.00 | 0.00 | 2.98 |
| 6895 | 0.0010 | 0.00 | 0.00 | 2.98 |
| 6896 | 0.0013 | 0.00 | 0.00 | 2.98 |
| 6898 | 0.0006 | 0.00 | 0.00 | 2.98 |
| 6899 | 0.0003 | 0.00 | 0.00 | 2.98 |
| 6900 | 0.0010 | 0.00 | 0.00 | 2.98 |
| 6901 | 0.0006 | 0.00 | 0.00 | 2.98 |
| 6902 | 0.0006 | 0.00 | 0.00 | 2.98 |
| 6904 | 0.0013 | 0.00 | 0.00 | 2.98 |
| 6905 | 0.0013 | 0.00 | 0.00 | 2.98 |
| 6906 | 0.0003 | 0.00 | 0.00 | 2.98 |
| 6907 | 0.0003 | 0.00 | 0.00 | 2.98 |
| 6908 | 0.0013 | 0.00 | 0.00 | 2.97 |
| 6909 | 0.0013 | 0.00 | 0.00 | 2.97 |
| 6910 | 0.0010 | 0.00 | 0.00 | 2.97 |
| 6911 | 0.0006 | 0.00 | 0.00 | 2.97 |
| 6912 | 0.0013 | 0.00 | 0.00 | 2.97 |
| 6913 | 0.0013 | 0.00 | 0.00 | 2.97 |
| 6914 | 0.0013 | 0.00 | 0.00 | 2.97 |
| 6915 | 0.0006 | 0.00 | 0.00 | 2.97 |
| 6916 | 0.0019 | 0.00 | 0.00 | 2.97 |
| 6917 | 0.0016 | 0.00 | 0.00 | 2.96 |
| 6920 | 0.0003 | 0.00 | 0.00 | 2.96 |
| 6921 | 0.0013 | 0.00 | 0.00 | 2.96 |
| 6922 | 0.0010 | 0.00 | 0.00 | 2.96 |
| 6923 | 0.0010 | 0.00 | 0.00 | 2.96 |
| 6924 | 0.0010 | 0.00 | 0.00 | 2.96 |
| 6925 | 0.0013 | 0.00 | 0.00 | 2.96 |
| 6926 | 0.0003 | 0.00 | 0.00 | 2.96 |
| 6927 | 0.0010 | 0.00 | 0.00 | 2.96 |
| 6928 | 0.0013 | 0.00 | 0.00 | 2.96 |
| 6929 | 0.0013 | 0.00 | 0.00 | 2.95 |
| 6930 | 0.0006 | 0.00 | 0.00 | 2.95 |

| | | | | |
|---|---|---|---|---|
| 6931 | 0.0003 | 0.00 | 0.00 | 2.95 |
| 6933 | 0.0013 | 0.00 | 0.00 | 2.95 |
| 6934 | 0.0006 | 0.00 | 0.00 | 2.95 |
| 6935 | 0.0006 | 0.00 | 0.00 | 2.95 |
| 6937 | 0.0013 | 0.00 | 0.00 | 2.95 |
| 6938 | 0.0013 | 0.00 | 0.00 | 2.95 |
| 6939 | 0.0013 | 0.00 | 0.00 | 2.95 |
| 6940 | 0.0003 | 0.00 | 0.00 | 2.95 |
| 6941 | 0.0010 | 0.00 | 0.00 | 2.95 |
| 6942 | 0.0003 | 0.00 | 0.00 | 2.94 |
| 6943 | 0.0010 | 0.00 | 0.00 | 2.94 |
| 6944 | 0.0013 | 0.00 | 0.00 | 2.94 |
| 6945 | 0.0016 | 0.00 | 0.00 | 2.94 |
| 6946 | 0.0003 | 0.00 | 0.00 | 2.94 |
| 6947 | 0.0022 | 0.00 | 0.00 | 2.94 |
| 6948 | 0.0006 | 0.00 | 0.00 | 2.94 |
| 6949 | 0.0010 | 0.00 | 0.00 | 2.94 |
| 6950 | 0.0013 | 0.00 | 0.00 | 2.94 |
| 6951 | 0.0013 | 0.00 | 0.00 | 2.93 |
| 6952 | 0.0006 | 0.00 | 0.00 | 2.93 |
| 6953 | 0.0016 | 0.00 | 0.00 | 2.93 |
| 6954 | 0.0010 | 0.00 | 0.00 | 2.93 |
| 6956 | 0.0006 | 0.00 | 0.00 | 2.93 |
| 6957 | 0.0003 | 0.00 | 0.00 | 2.93 |
| 6958 | 0.0019 | 0.00 | 0.00 | 2.93 |
| 6959 | 0.0003 | 0.00 | 0.00 | 2.93 |
| 6960 | 0.0010 | 0.00 | 0.00 | 2.93 |
| 6961 | 0.0006 | 0.00 | 0.00 | 2.93 |
| 6962 | 0.0013 | 0.00 | 0.00 | 2.93 |
| 6963 | 0.0006 | 0.00 | 0.00 | 2.92 |
| 6964 | 0.0010 | 0.00 | 0.00 | 2.92 |
| 6965 | 0.0010 | 0.00 | 0.00 | 2.92 |
| 6966 | 0.0003 | 0.00 | 0.00 | 2.92 |
| 6967 | 0.0022 | 0.00 | 0.00 | 2.92 |
| 6968 | 0.0013 | 0.00 | 0.00 | 2.92 |
| 6969 | 0.0013 | 0.00 | 0.00 | 2.92 |
| 6970 | 0.0019 | 0.00 | 0.00 | 2.92 |
| 6971 | 0.0016 | 0.00 | 0.00 | 2.91 |
| 6972 | 0.0010 | 0.00 | 0.00 | 2.91 |
| 6973 | 0.0016 | 0.00 | 0.00 | 2.91 |
| 6974 | 0.0013 | 0.00 | 0.00 | 2.91 |
| 6975 | 0.0022 | 0.00 | 0.00 | 2.91 |
| 6976 | 0.0019 | 0.00 | 0.00 | 2.91 |
| 6977 | 0.0013 | 0.00 | 0.00 | 2.90 |

| | | | | |
|---|---|---|---|---|
| 6978 | 0.0010 | 0.00 | 0.00 | 2.90 |
| 6979 | 0.0006 | 0.00 | 0.00 | 2.90 |
| 6980 | 0.0013 | 0.00 | 0.00 | 2.90 |
| 6981 | 0.0010 | 0.00 | 0.00 | 2.90 |
| 6982 | 0.0013 | 0.00 | 0.00 | 2.90 |
| 6983 | 0.0013 | 0.00 | 0.00 | 2.90 |
| 6984 | 0.0016 | 0.00 | 0.00 | 2.90 |
| 6985 | 0.0013 | 0.00 | 0.00 | 2.90 |
| 6986 | 0.0013 | 0.00 | 0.00 | 2.89 |
| 6987 | 0.0010 | 0.00 | 0.00 | 2.89 |
| 6988 | 0.0006 | 0.00 | 0.00 | 2.89 |
| 6989 | 0.0016 | 0.00 | 0.00 | 2.89 |
| 6991 | 0.0003 | 0.00 | 0.00 | 2.89 |
| 6992 | 0.0006 | 0.00 | 0.00 | 2.89 |
| 6993 | 0.0010 | 0.00 | 0.00 | 2.89 |
| 6994 | 0.0013 | 0.00 | 0.00 | 2.89 |
| 6995 | 0.0010 | 0.00 | 0.00 | 2.89 |
| 6996 | 0.0006 | 0.00 | 0.00 | 2.89 |
| 6997 | 0.0010 | 0.00 | 0.00 | 2.88 |
| 6998 | 0.0006 | 0.00 | 0.00 | 2.88 |
| 6999 | 0.0003 | 0.00 | 0.00 | 2.88 |
| 7000 | 0.0006 | 0.00 | 0.00 | 2.88 |
| 7001 | 0.0003 | 0.00 | 0.00 | 2.88 |
| 7002 | 0.0006 | 0.00 | 0.00 | 2.88 |
| 7003 | 0.0003 | 0.00 | 0.00 | 2.88 |
| 7004 | 0.0003 | 0.00 | 0.00 | 2.88 |
| 7005 | 0.0013 | 0.00 | 0.00 | 2.88 |
| 7006 | 0.0006 | 0.00 | 0.00 | 2.88 |
| 7007 | 0.0003 | 0.00 | 0.00 | 2.88 |
| 7008 | 0.0013 | 0.00 | 0.00 | 2.88 |
| 7009 | 0.0003 | 0.00 | 0.00 | 2.88 |
| 7010 | 0.0006 | 0.00 | 0.00 | 2.88 |
| 7011 | 0.0006 | 0.00 | 0.00 | 2.88 |
| 7013 | 0.0013 | 0.00 | 0.00 | 2.88 |
| 7014 | 0.0013 | 0.00 | 0.00 | 2.87 |
| 7015 | 0.0013 | 0.00 | 0.00 | 2.87 |
| 7016 | 0.0003 | 0.00 | 0.00 | 2.87 |
| 7017 | 0.0013 | 0.00 | 0.00 | 2.87 |
| 7018 | 0.0010 | 0.00 | 0.00 | 2.87 |
| 7020 | 0.0006 | 0.00 | 0.00 | 2.87 |
| 7021 | 0.0006 | 0.00 | 0.00 | 2.87 |
| 7022 | 0.0003 | 0.00 | 0.00 | 2.87 |
| 7023 | 0.0016 | 0.00 | 0.00 | 2.87 |
| 7024 | 0.0003 | 0.00 | 0.00 | 2.87 |

| | | | | |
|---|---|---|---|---|
| 7025 | 0.0010 | 0.00 | 0.00 | 2.87 |
| 7026 | 0.0006 | 0.00 | 0.00 | 2.86 |
| 7027 | 0.0019 | 0.00 | 0.00 | 2.86 |
| 7028 | 0.0010 | 0.00 | 0.00 | 2.86 |
| 7029 | 0.0010 | 0.00 | 0.00 | 2.86 |
| 7030 | 0.0010 | 0.00 | 0.00 | 2.86 |
| 7031 | 0.0010 | 0.00 | 0.00 | 2.86 |
| 7032 | 0.0006 | 0.00 | 0.00 | 2.86 |
| 7034 | 0.0003 | 0.00 | 0.00 | 2.86 |
| 7035 | 0.0010 | 0.00 | 0.00 | 2.86 |
| 7036 | 0.0003 | 0.00 | 0.00 | 2.86 |
| 7037 | 0.0006 | 0.00 | 0.00 | 2.86 |
| 7038 | 0.0006 | 0.00 | 0.00 | 2.86 |
| 7039 | 0.0013 | 0.00 | 0.00 | 2.85 |
| 7040 | 0.0010 | 0.00 | 0.00 | 2.85 |
| 7041 | 0.0003 | 0.00 | 0.00 | 2.85 |
| 7042 | 0.0016 | 0.00 | 0.00 | 2.85 |
| 7043 | 0.0022 | 0.00 | 0.00 | 2.85 |
| 7044 | 0.0010 | 0.00 | 0.00 | 2.85 |
| 7045 | 0.0006 | 0.00 | 0.00 | 2.85 |
| 7046 | 0.0003 | 0.00 | 0.00 | 2.85 |
| 7047 | 0.0016 | 0.00 | 0.00 | 2.85 |
| 7048 | 0.0010 | 0.00 | 0.00 | 2.84 |
| 7049 | 0.0010 | 0.00 | 0.00 | 2.84 |
| 7050 | 0.0006 | 0.00 | 0.00 | 2.84 |
| 7051 | 0.0010 | 0.00 | 0.00 | 2.84 |
| 7052 | 0.0016 | 0.00 | 0.00 | 2.84 |
| 7053 | 0.0010 | 0.00 | 0.00 | 2.84 |
| 7054 | 0.0003 | 0.00 | 0.00 | 2.84 |
| 7055 | 0.0013 | 0.00 | 0.00 | 2.84 |
| 7056 | 0.0003 | 0.00 | 0.00 | 2.84 |
| 7057 | 0.0003 | 0.00 | 0.00 | 2.84 |
| 7058 | 0.0010 | 0.00 | 0.00 | 2.84 |
| 7060 | 0.0006 | 0.00 | 0.00 | 2.84 |
| 7061 | 0.0010 | 0.00 | 0.00 | 2.83 |
| 7062 | 0.0003 | 0.00 | 0.00 | 2.83 |
| 7063 | 0.0010 | 0.00 | 0.00 | 2.83 |
| 7064 | 0.0016 | 0.00 | 0.00 | 2.83 |
| 7066 | 0.0010 | 0.00 | 0.00 | 2.83 |
| 7068 | 0.0006 | 0.00 | 0.00 | 2.83 |
| 7069 | 0.0006 | 0.00 | 0.00 | 2.83 |
| 7070 | 0.0010 | 0.00 | 0.00 | 2.83 |
| 7071 | 0.0016 | 0.00 | 0.00 | 2.83 |
| 7072 | 0.0003 | 0.00 | 0.00 | 2.83 |

| | | | | |
|---|---|---|---|---|
| 7073 | 0.0010 | 0.00 | 0.00 | 2.83 |
| 7074 | 0.0013 | 0.00 | 0.00 | 2.83 |
| 7075 | 0.0016 | 0.00 | 0.00 | 2.82 |
| 7076 | 0.0010 | 0.00 | 0.00 | 2.82 |
| 7077 | 0.0013 | 0.00 | 0.00 | 2.82 |
| 7078 | 0.0003 | 0.00 | 0.00 | 2.82 |
| 7079 | 0.0003 | 0.00 | 0.00 | 2.82 |
| 7080 | 0.0006 | 0.00 | 0.00 | 2.82 |
| 7081 | 0.0022 | 0.00 | 0.00 | 2.82 |
| 7082 | 0.0003 | 0.00 | 0.00 | 2.82 |
| 7083 | 0.0010 | 0.00 | 0.00 | 2.82 |
| 7084 | 0.0006 | 0.00 | 0.00 | 2.82 |
| 7085 | 0.0016 | 0.00 | 0.00 | 2.81 |
| 7086 | 0.0006 | 0.00 | 0.00 | 2.81 |
| 7088 | 0.0013 | 0.00 | 0.00 | 2.81 |
| 7089 | 0.0010 | 0.00 | 0.00 | 2.81 |
| 7090 | 0.0010 | 0.00 | 0.00 | 2.81 |
| 7091 | 0.0010 | 0.00 | 0.00 | 2.81 |
| 7092 | 0.0006 | 0.00 | 0.00 | 2.81 |
| 7093 | 0.0010 | 0.00 | 0.00 | 2.81 |
| 7094 | 0.0003 | 0.00 | 0.00 | 2.81 |
| 7095 | 0.0013 | 0.00 | 0.00 | 2.81 |
| 7096 | 0.0003 | 0.00 | 0.00 | 2.80 |
| 7097 | 0.0006 | 0.00 | 0.00 | 2.80 |
| 7098 | 0.0010 | 0.00 | 0.00 | 2.80 |
| 7099 | 0.0003 | 0.00 | 0.00 | 2.80 |
| 7100 | 0.0016 | 0.00 | 0.00 | 2.80 |
| 7101 | 0.0003 | 0.00 | 0.00 | 2.80 |
| 7102 | 0.0006 | 0.00 | 0.00 | 2.80 |
| 7103 | 0.0010 | 0.00 | 0.00 | 2.80 |
| 7104 | 0.0010 | 0.00 | 0.00 | 2.80 |
| 7105 | 0.0003 | 0.00 | 0.00 | 2.80 |
| 7106 | 0.0010 | 0.00 | 0.00 | 2.80 |
| 7107 | 0.0010 | 0.00 | 0.00 | 2.80 |
| 7108 | 0.0003 | 0.00 | 0.00 | 2.80 |
| 7109 | 0.0016 | 0.00 | 0.00 | 2.80 |
| 7110 | 0.0006 | 0.00 | 0.00 | 2.79 |
| 7111 | 0.0006 | 0.00 | 0.00 | 2.79 |
| 7112 | 0.0022 | 0.00 | 0.00 | 2.79 |
| 7114 | 0.0010 | 0.00 | 0.00 | 2.79 |
| 7115 | 0.0010 | 0.00 | 0.00 | 2.79 |
| 7116 | 0.0016 | 0.00 | 0.00 | 2.79 |
| 7117 | 0.0013 | 0.00 | 0.00 | 2.79 |
| 7118 | 0.0003 | 0.00 | 0.00 | 2.79 |

| | | | | |
|---|---|---|---|---|
| 7119 | 0.0010 | 0.00 | 0.00 | 2.79 |
| 7120 | 0.0013 | 0.00 | 0.00 | 2.78 |
| 7121 | 0.0006 | 0.00 | 0.00 | 2.78 |
| 7122 | 0.0019 | 0.00 | 0.00 | 2.78 |
| 7124 | 0.0006 | 0.00 | 0.00 | 2.78 |
| 7125 | 0.0010 | 0.00 | 0.00 | 2.78 |
| 7126 | 0.0019 | 0.00 | 0.00 | 2.78 |
| 7127 | 0.0010 | 0.00 | 0.00 | 2.78 |
| 7128 | 0.0006 | 0.00 | 0.00 | 2.78 |
| 7129 | 0.0013 | 0.00 | 0.00 | 2.78 |
| 7130 | 0.0016 | 0.00 | 0.00 | 2.77 |
| 7131 | 0.0019 | 0.00 | 0.00 | 2.77 |
| 7132 | 0.0006 | 0.00 | 0.00 | 2.77 |
| 7133 | 0.0010 | 0.00 | 0.00 | 2.77 |
| 7134 | 0.0010 | 0.00 | 0.00 | 2.77 |
| 7135 | 0.0010 | 0.00 | 0.00 | 2.77 |
| 7136 | 0.0016 | 0.00 | 0.00 | 2.77 |
| 7137 | 0.0013 | 0.00 | 0.00 | 2.77 |
| 7138 | 0.0006 | 0.00 | 0.00 | 2.76 |
| 7139 | 0.0010 | 0.00 | 0.00 | 2.76 |
| 7140 | 0.0006 | 0.00 | 0.00 | 2.76 |
| 7141 | 0.0013 | 0.00 | 0.00 | 2.76 |
| 7142 | 0.0010 | 0.00 | 0.00 | 2.76 |
| 7143 | 0.0019 | 0.00 | 0.00 | 2.76 |
| 7144 | 0.0019 | 0.00 | 0.00 | 2.76 |
| 7145 | 0.0010 | 0.00 | 0.00 | 2.76 |
| 7146 | 0.0013 | 0.00 | 0.00 | 2.75 |
| 7148 | 0.0016 | 0.00 | 0.00 | 2.75 |
| 7149 | 0.0010 | 0.00 | 0.00 | 2.75 |
| 7150 | 0.0006 | 0.00 | 0.00 | 2.75 |
| 7151 | 0.0006 | 0.00 | 0.00 | 2.75 |
| 7152 | 0.0016 | 0.00 | 0.00 | 2.75 |
| 7153 | 0.0010 | 0.00 | 0.00 | 2.75 |
| 7154 | 0.0006 | 0.00 | 0.00 | 2.75 |
| 7155 | 0.0006 | 0.00 | 0.00 | 2.75 |
| 7156 | 0.0003 | 0.00 | 0.00 | 2.75 |
| 7157 | 0.0013 | 0.00 | 0.00 | 2.75 |
| 7158 | 0.0022 | 0.00 | 0.00 | 2.74 |
| 7159 | 0.0016 | 0.00 | 0.00 | 2.74 |
| 7160 | 0.0013 | 0.00 | 0.00 | 2.74 |
| 7161 | 0.0010 | 0.00 | 0.00 | 2.74 |
| 7162 | 0.0013 | 0.00 | 0.00 | 2.74 |
| 7163 | 0.0010 | 0.00 | 0.00 | 2.74 |
| 7164 | 0.0010 | 0.00 | 0.00 | 2.74 |

| | | | | |
|---|---|---|---|---|
| 7165 | 0.0006 | 0.00 | 0.00 | 2.73 |
| 7166 | 0.0006 | 0.00 | 0.00 | 2.73 |
| 7167 | 0.0006 | 0.00 | 0.00 | 2.73 |
| 7168 | 0.0003 | 0.00 | 0.00 | 2.73 |
| 7169 | 0.0006 | 0.00 | 0.00 | 2.73 |
| 7170 | 0.0003 | 0.00 | 0.00 | 2.73 |
| 7171 | 0.0013 | 0.00 | 0.00 | 2.73 |
| 7172 | 0.0016 | 0.00 | 0.00 | 2.73 |
| 7173 | 0.0013 | 0.00 | 0.00 | 2.73 |
| 7174 | 0.0003 | 0.00 | 0.00 | 2.73 |
| 7175 | 0.0003 | 0.00 | 0.00 | 2.73 |
| 7176 | 0.0006 | 0.00 | 0.00 | 2.73 |
| 7177 | 0.0016 | 0.00 | 0.00 | 2.73 |
| 7178 | 0.0010 | 0.00 | 0.00 | 2.72 |
| 7179 | 0.0010 | 0.00 | 0.00 | 2.72 |
| 7180 | 0.0013 | 0.00 | 0.00 | 2.72 |
| 7181 | 0.0006 | 0.00 | 0.00 | 2.72 |
| 7182 | 0.0016 | 0.00 | 0.00 | 2.72 |
| 7183 | 0.0006 | 0.00 | 0.00 | 2.72 |
| 7184 | 0.0010 | 0.00 | 0.00 | 2.72 |
| 7185 | 0.0006 | 0.00 | 0.00 | 2.72 |
| 7186 | 0.0016 | 0.00 | 0.00 | 2.72 |
| 7187 | 0.0010 | 0.00 | 0.00 | 2.72 |
| 7188 | 0.0003 | 0.00 | 0.00 | 2.71 |
| 7189 | 0.0010 | 0.00 | 0.00 | 2.71 |
| 7190 | 0.0003 | 0.00 | 0.00 | 2.71 |
| 7191 | 0.0010 | 0.00 | 0.00 | 2.71 |
| 7192 | 0.0016 | 0.00 | 0.00 | 2.71 |
| 7193 | 0.0003 | 0.00 | 0.00 | 2.71 |
| 7194 | 0.0006 | 0.00 | 0.00 | 2.71 |
| 7195 | 0.0010 | 0.00 | 0.00 | 2.71 |
| 7196 | 0.0016 | 0.00 | 0.00 | 2.71 |
| 7197 | 0.0010 | 0.00 | 0.00 | 2.71 |
| 7198 | 0.0016 | 0.00 | 0.00 | 2.71 |
| 7199 | 0.0029 | 0.00 | 0.00 | 2.70 |
| 7200 | 0.0006 | 0.00 | 0.00 | 2.70 |
| 7201 | 0.0010 | 0.00 | 0.00 | 2.70 |
| 7202 | 0.0029 | 0.00 | 0.00 | 2.70 |
| 7203 | 0.0013 | 0.00 | 0.00 | 2.70 |
| 7204 | 0.0006 | 0.00 | 0.00 | 2.70 |
| 7205 | 0.0013 | 0.00 | 0.00 | 2.69 |
| 7206 | 0.0035 | 0.00 | 0.00 | 2.69 |
| 7207 | 0.0029 | 0.00 | 0.00 | 2.69 |
| 7208 | 0.0029 | 0.00 | 0.00 | 2.69 |

| | | | | |
|---|---|---|---|---|
| 7209 | 0.0019 | 0.00 | 0.00 | 2.68 |
| 7210 | 0.0010 | 0.00 | 0.00 | 2.68 |
| 7211 | 0.0035 | 0.00 | 0.00 | 2.68 |
| 7212 | 0.0016 | 0.00 | 0.00 | 2.68 |
| 7213 | 0.0010 | 0.00 | 0.00 | 2.68 |
| 7214 | 0.0016 | 0.00 | 0.00 | 2.68 |
| 7215 | 0.0016 | 0.00 | 0.00 | 2.67 |
| 7216 | 0.0019 | 0.00 | 0.00 | 2.67 |
| 7217 | 0.0016 | 0.00 | 0.00 | 2.67 |
| 7219 | 0.0006 | 0.00 | 0.00 | 2.67 |
| 7221 | 0.0006 | 0.00 | 0.00 | 2.67 |
| 7222 | 0.0013 | 0.00 | 0.00 | 2.67 |
| 7223 | 0.0003 | 0.00 | 0.00 | 2.67 |
| 7224 | 0.0003 | 0.00 | 0.00 | 2.67 |
| 7225 | 0.0010 | 0.00 | 0.00 | 2.67 |
| 7226 | 0.0022 | 0.00 | 0.00 | 2.66 |
| 7228 | 0.0013 | 0.00 | 0.00 | 2.66 |
| 7229 | 0.0016 | 0.00 | 0.00 | 2.66 |
| 7230 | 0.0010 | 0.00 | 0.00 | 2.66 |
| 7231 | 0.0006 | 0.00 | 0.00 | 2.66 |
| 7232 | 0.0016 | 0.00 | 0.00 | 2.66 |
| 7233 | 0.0006 | 0.00 | 0.00 | 2.66 |
| 7234 | 0.0019 | 0.00 | 0.00 | 2.66 |
| 7236 | 0.0022 | 0.00 | 0.00 | 2.65 |
| 7237 | 0.0019 | 0.00 | 0.00 | 2.65 |
| 7238 | 0.0003 | 0.00 | 0.00 | 2.65 |
| 7240 | 0.0016 | 0.00 | 0.00 | 2.65 |
| 7241 | 0.0003 | 0.00 | 0.00 | 2.65 |
| 7242 | 0.0016 | 0.00 | 0.00 | 2.65 |
| 7243 | 0.0029 | 0.00 | 0.00 | 2.65 |
| 7244 | 0.0006 | 0.00 | 0.00 | 2.64 |
| 7245 | 0.0010 | 0.00 | 0.00 | 2.64 |
| 7246 | 0.0010 | 0.00 | 0.00 | 2.64 |
| 7247 | 0.0013 | 0.00 | 0.00 | 2.64 |
| 7248 | 0.0010 | 0.00 | 0.00 | 2.64 |
| 7249 | 0.0013 | 0.00 | 0.00 | 2.64 |
| 7250 | 0.0013 | 0.00 | 0.00 | 2.64 |
| 7251 | 0.0006 | 0.00 | 0.00 | 2.64 |
| 7252 | 0.0006 | 0.00 | 0.00 | 2.63 |
| 7253 | 0.0003 | 0.00 | 0.00 | 2.63 |
| 7254 | 0.0006 | 0.00 | 0.00 | 2.63 |
| 7255 | 0.0003 | 0.00 | 0.00 | 2.63 |
| 7256 | 0.0003 | 0.00 | 0.00 | 2.63 |
| 7257 | 0.0016 | 0.00 | 0.00 | 2.63 |

| | | | | |
|---|---|---|---|---|
| 7258 | 0.0010 | 0.00 | 0.00 | 2.63 |
| 7259 | 0.0006 | 0.00 | 0.00 | 2.63 |
| 7260 | 0.0006 | 0.00 | 0.00 | 2.63 |
| 7261 | 0.0019 | 0.00 | 0.00 | 2.63 |
| 7262 | 0.0013 | 0.00 | 0.00 | 2.63 |
| 7263 | 0.0026 | 0.00 | 0.00 | 2.63 |
| 7264 | 0.0013 | 0.00 | 0.00 | 2.62 |
| 7265 | 0.0006 | 0.00 | 0.00 | 2.62 |
| 7266 | 0.0022 | 0.00 | 0.00 | 2.62 |
| 7267 | 0.0019 | 0.00 | 0.00 | 2.62 |
| 7268 | 0.0013 | 0.00 | 0.00 | 2.62 |
| 7269 | 0.0022 | 0.00 | 0.00 | 2.62 |
| 7270 | 0.0010 | 0.00 | 0.00 | 2.61 |
| 7271 | 0.0013 | 0.00 | 0.00 | 2.61 |
| 7272 | 0.0010 | 0.00 | 0.00 | 2.61 |
| 7274 | 0.0010 | 0.00 | 0.00 | 2.61 |
| 7275 | 0.0003 | 0.00 | 0.00 | 2.61 |
| 7276 | 0.0013 | 0.00 | 0.00 | 2.61 |
| 7278 | 0.0003 | 0.00 | 0.00 | 2.61 |
| 7279 | 0.0016 | 0.00 | 0.00 | 2.61 |
| 7280 | 0.0003 | 0.00 | 0.00 | 2.61 |
| 7281 | 0.0003 | 0.00 | 0.00 | 2.61 |
| 7282 | 0.0010 | 0.00 | 0.00 | 2.60 |
| 7283 | 0.0010 | 0.00 | 0.00 | 2.60 |
| 7284 | 0.0003 | 0.00 | 0.00 | 2.60 |
| 7285 | 0.0013 | 0.00 | 0.00 | 2.60 |
| 7286 | 0.0013 | 0.00 | 0.00 | 2.60 |
| 7287 | 0.0010 | 0.00 | 0.00 | 2.60 |
| 7288 | 0.0006 | 0.00 | 0.00 | 2.60 |
| 7289 | 0.0019 | 0.00 | 0.00 | 2.60 |
| 7290 | 0.0026 | 0.00 | 0.00 | 2.60 |
| 7291 | 0.0010 | 0.00 | 0.00 | 2.59 |
| 7292 | 0.0016 | 0.00 | 0.00 | 2.59 |
| 7293 | 0.0013 | 0.00 | 0.00 | 2.59 |
| 7294 | 0.0006 | 0.00 | 0.00 | 2.59 |
| 7295 | 0.0006 | 0.00 | 0.00 | 2.59 |
| 7296 | 0.0010 | 0.00 | 0.00 | 2.59 |
| 7297 | 0.0019 | 0.00 | 0.00 | 2.59 |
| 7298 | 0.0019 | 0.00 | 0.00 | 2.59 |
| 7299 | 0.0019 | 0.00 | 0.00 | 2.58 |
| 7300 | 0.0006 | 0.00 | 0.00 | 2.58 |
| 7301 | 0.0006 | 0.00 | 0.00 | 2.58 |
| 7302 | 0.0019 | 0.00 | 0.00 | 2.58 |
| 7303 | 0.0003 | 0.00 | 0.00 | 2.58 |

| | | | | |
|---|---|---|---|---|
| 7304 | 0.0006 | 0.00 | 0.00 | 2.58 |
| 7305 | 0.0003 | 0.00 | 0.00 | 2.58 |
| 7306 | 0.0022 | 0.00 | 0.00 | 2.58 |
| 7307 | 0.0010 | 0.00 | 0.00 | 2.58 |
| 7308 | 0.0016 | 0.00 | 0.00 | 2.57 |
| 7309 | 0.0019 | 0.00 | 0.00 | 2.57 |
| 7310 | 0.0003 | 0.00 | 0.00 | 2.57 |
| 7311 | 0.0003 | 0.00 | 0.00 | 2.57 |
| 7312 | 0.0022 | 0.00 | 0.00 | 2.57 |
| 7313 | 0.0016 | 0.00 | 0.00 | 2.57 |
| 7314 | 0.0006 | 0.00 | 0.00 | 2.57 |
| 7315 | 0.0010 | 0.00 | 0.00 | 2.57 |
| 7316 | 0.0019 | 0.00 | 0.00 | 2.56 |
| 7317 | 0.0022 | 0.00 | 0.00 | 2.56 |
| 7318 | 0.0016 | 0.00 | 0.00 | 2.56 |
| 7319 | 0.0010 | 0.00 | 0.00 | 2.56 |
| 7320 | 0.0006 | 0.00 | 0.00 | 2.56 |
| 7321 | 0.0003 | 0.00 | 0.00 | 2.56 |
| 7322 | 0.0003 | 0.00 | 0.00 | 2.56 |
| 7323 | 0.0010 | 0.00 | 0.00 | 2.56 |
| 7324 | 0.0016 | 0.00 | 0.00 | 2.56 |
| 7325 | 0.0026 | 0.00 | 0.00 | 2.55 |
| 7326 | 0.0006 | 0.00 | 0.00 | 2.55 |
| 7327 | 0.0010 | 0.00 | 0.00 | 2.55 |
| 7328 | 0.0013 | 0.00 | 0.00 | 2.55 |
| 7329 | 0.0010 | 0.00 | 0.00 | 2.55 |
| 7330 | 0.0010 | 0.00 | 0.00 | 2.55 |
| 7331 | 0.0006 | 0.00 | 0.00 | 2.55 |
| 7332 | 0.0013 | 0.00 | 0.00 | 2.55 |
| 7333 | 0.0003 | 0.00 | 0.00 | 2.54 |
| 7334 | 0.0010 | 0.00 | 0.00 | 2.54 |
| 7335 | 0.0013 | 0.00 | 0.00 | 2.54 |
| 7336 | 0.0016 | 0.00 | 0.00 | 2.54 |
| 7337 | 0.0019 | 0.00 | 0.00 | 2.54 |
| 7338 | 0.0010 | 0.00 | 0.00 | 2.54 |
| 7339 | 0.0010 | 0.00 | 0.00 | 2.54 |
| 7340 | 0.0006 | 0.00 | 0.00 | 2.54 |
| 7341 | 0.0006 | 0.00 | 0.00 | 2.54 |
| 7342 | 0.0013 | 0.00 | 0.00 | 2.54 |
| 7343 | 0.0016 | 0.00 | 0.00 | 2.53 |
| 7344 | 0.0003 | 0.00 | 0.00 | 2.53 |
| 7345 | 0.0013 | 0.00 | 0.00 | 2.53 |
| 7346 | 0.0006 | 0.00 | 0.00 | 2.53 |
| 7347 | 0.0006 | 0.00 | 0.00 | 2.53 |

| | | | | |
|---|---|---|---|---|
| 7348 | 0.0006 | 0.00 | 0.00 | 2.53 |
| 7350 | 0.0013 | 0.00 | 0.00 | 2.53 |
| 7351 | 0.0016 | 0.00 | 0.00 | 2.53 |
| 7352 | 0.0016 | 0.00 | 0.00 | 2.53 |
| 7353 | 0.0016 | 0.00 | 0.00 | 2.52 |
| 7354 | 0.0010 | 0.00 | 0.00 | 2.52 |
| 7356 | 0.0010 | 0.00 | 0.00 | 2.52 |
| 7357 | 0.0016 | 0.00 | 0.00 | 2.52 |
| 7358 | 0.0013 | 0.00 | 0.00 | 2.52 |
| 7359 | 0.0029 | 0.00 | 0.00 | 2.52 |
| 7360 | 0.0010 | 0.00 | 0.00 | 2.52 |
| 7361 | 0.0010 | 0.00 | 0.00 | 2.51 |
| 7362 | 0.0010 | 0.00 | 0.00 | 2.51 |
| 7363 | 0.0019 | 0.00 | 0.00 | 2.51 |
| 7364 | 0.0010 | 0.00 | 0.00 | 2.51 |
| 7365 | 0.0026 | 0.00 | 0.00 | 2.51 |
| 7366 | 0.0006 | 0.00 | 0.00 | 2.51 |
| 7367 | 0.0010 | 0.00 | 0.00 | 2.51 |
| 7369 | 0.0013 | 0.00 | 0.00 | 2.51 |
| 7370 | 0.0016 | 0.00 | 0.00 | 2.50 |
| 7371 | 0.0010 | 0.00 | 0.00 | 2.50 |
| 7372 | 0.0010 | 0.00 | 0.00 | 2.50 |
| 7373 | 0.0010 | 0.00 | 0.00 | 2.50 |
| 7374 | 0.0022 | 0.00 | 0.00 | 2.50 |
| 7375 | 0.0010 | 0.00 | 0.00 | 2.50 |
| 7376 | 0.0016 | 0.00 | 0.00 | 2.50 |
| 7377 | 0.0006 | 0.00 | 0.00 | 2.49 |
| 7378 | 0.0006 | 0.00 | 0.00 | 2.49 |
| 7379 | 0.0006 | 0.00 | 0.00 | 2.49 |
| 7380 | 0.0006 | 0.00 | 0.00 | 2.49 |
| 7381 | 0.0010 | 0.00 | 0.00 | 2.49 |
| 7382 | 0.0013 | 0.00 | 0.00 | 2.49 |
| 7383 | 0.0010 | 0.00 | 0.00 | 2.49 |
| 7384 | 0.0013 | 0.00 | 0.00 | 2.49 |
| 7385 | 0.0006 | 0.00 | 0.00 | 2.49 |
| 7386 | 0.0010 | 0.00 | 0.00 | 2.49 |
| 7387 | 0.0003 | 0.00 | 0.00 | 2.49 |
| 7388 | 0.0013 | 0.00 | 0.00 | 2.49 |
| 7389 | 0.0019 | 0.00 | 0.00 | 2.48 |
| 7390 | 0.0003 | 0.00 | 0.00 | 2.48 |
| 7391 | 0.0013 | 0.00 | 0.00 | 2.48 |
| 7392 | 0.0016 | 0.00 | 0.00 | 2.48 |
| 7393 | 0.0006 | 0.00 | 0.00 | 2.48 |
| 7394 | 0.0016 | 0.00 | 0.00 | 2.48 |

| | | | | |
|---|---|---|---|---|
| 7395 | 0.0016 | 0.00 | 0.00 | 2.48 |
| 7396 | 0.0003 | 0.00 | 0.00 | 2.48 |
| 7397 | 0.0010 | 0.00 | 0.00 | 2.48 |
| 7398 | 0.0010 | 0.00 | 0.00 | 2.47 |
| 7399 | 0.0010 | 0.00 | 0.00 | 2.47 |
| 7400 | 0.0010 | 0.00 | 0.00 | 2.47 |
| 7401 | 0.0016 | 0.00 | 0.00 | 2.47 |
| 7402 | 0.0010 | 0.00 | 0.00 | 2.47 |
| 7403 | 0.0006 | 0.00 | 0.00 | 2.47 |
| 7404 | 0.0010 | 0.00 | 0.00 | 2.47 |
| 7405 | 0.0003 | 0.00 | 0.00 | 2.47 |
| 7406 | 0.0010 | 0.00 | 0.00 | 2.47 |
| 7407 | 0.0019 | 0.00 | 0.00 | 2.47 |
| 7408 | 0.0006 | 0.00 | 0.00 | 2.46 |
| 7409 | 0.0006 | 0.00 | 0.00 | 2.46 |
| 7410 | 0.0003 | 0.00 | 0.00 | 2.46 |
| 7411 | 0.0016 | 0.00 | 0.00 | 2.46 |
| 7412 | 0.0003 | 0.00 | 0.00 | 2.46 |
| 7413 | 0.0006 | 0.00 | 0.00 | 2.46 |
| 7414 | 0.0035 | 0.00 | 0.00 | 2.46 |
| 7415 | 0.0010 | 0.00 | 0.00 | 2.46 |
| 7416 | 0.0016 | 0.00 | 0.00 | 2.46 |
| 7417 | 0.0013 | 0.00 | 0.00 | 2.45 |
| 7418 | 0.0022 | 0.00 | 0.00 | 2.45 |
| 7419 | 0.0010 | 0.00 | 0.00 | 2.45 |
| 7420 | 0.0016 | 0.00 | 0.00 | 2.45 |
| 7421 | 0.0013 | 0.00 | 0.00 | 2.45 |
| 7422 | 0.0013 | 0.00 | 0.00 | 2.45 |
| 7423 | 0.0006 | 0.00 | 0.00 | 2.45 |
| 7424 | 0.0010 | 0.00 | 0.00 | 2.44 |
| 7425 | 0.0003 | 0.00 | 0.00 | 2.44 |
| 7426 | 0.0013 | 0.00 | 0.00 | 2.44 |
| 7427 | 0.0026 | 0.00 | 0.00 | 2.44 |
| 7430 | 0.0006 | 0.00 | 0.00 | 2.44 |
| 7431 | 0.0013 | 0.00 | 0.00 | 2.44 |
| 7432 | 0.0010 | 0.00 | 0.00 | 2.44 |
| 7433 | 0.0006 | 0.00 | 0.00 | 2.44 |
| 7434 | 0.0006 | 0.00 | 0.00 | 2.44 |
| 7435 | 0.0010 | 0.00 | 0.00 | 2.44 |
| 7436 | 0.0003 | 0.00 | 0.00 | 2.43 |
| 7437 | 0.0010 | 0.00 | 0.00 | 2.43 |
| 7438 | 0.0010 | 0.00 | 0.00 | 2.43 |
| 7439 | 0.0006 | 0.00 | 0.00 | 2.43 |
| 7440 | 0.0006 | 0.00 | 0.00 | 2.43 |

| | | | | |
|---|---|---|---|---|
| 7441 | 0.0029 | 0.00 | 0.00 | 2.43 |
| 7442 | 0.0010 | 0.00 | 0.00 | 2.43 |
| 7443 | 0.0013 | 0.00 | 0.00 | 2.43 |
| 7444 | 0.0010 | 0.00 | 0.00 | 2.43 |
| 7445 | 0.0006 | 0.00 | 0.00 | 2.42 |
| 7446 | 0.0019 | 0.00 | 0.00 | 2.42 |
| 7447 | 0.0010 | 0.00 | 0.00 | 2.42 |
| 7448 | 0.0013 | 0.00 | 0.00 | 2.42 |
| 7449 | 0.0013 | 0.00 | 0.00 | 2.42 |
| 7450 | 0.0006 | 0.00 | 0.00 | 2.42 |
| 7451 | 0.0006 | 0.00 | 0.00 | 2.42 |
| 7452 | 0.0022 | 0.00 | 0.00 | 2.42 |
| 7453 | 0.0016 | 0.00 | 0.00 | 2.42 |
| 7454 | 0.0010 | 0.00 | 0.00 | 2.41 |
| 7455 | 0.0013 | 0.00 | 0.00 | 2.41 |
| 7456 | 0.0003 | 0.00 | 0.00 | 2.41 |
| 7457 | 0.0003 | 0.00 | 0.00 | 2.41 |
| 7458 | 0.0013 | 0.00 | 0.00 | 2.41 |
| 7459 | 0.0006 | 0.00 | 0.00 | 2.41 |
| 7460 | 0.0003 | 0.00 | 0.00 | 2.41 |
| 7461 | 0.0022 | 0.00 | 0.00 | 2.41 |
| 7462 | 0.0013 | 0.00 | 0.00 | 2.41 |
| 7463 | 0.0003 | 0.00 | 0.00 | 2.40 |
| 7464 | 0.0006 | 0.00 | 0.00 | 2.40 |
| 7465 | 0.0019 | 0.00 | 0.00 | 2.40 |
| 7466 | 0.0006 | 0.00 | 0.00 | 2.40 |
| 7467 | 0.0022 | 0.00 | 0.00 | 2.40 |
| 7468 | 0.0022 | 0.00 | 0.00 | 2.40 |
| 7469 | 0.0019 | 0.00 | 0.00 | 2.40 |
| 7470 | 0.0006 | 0.00 | 0.00 | 2.39 |
| 7471 | 0.0006 | 0.00 | 0.00 | 2.39 |
| 7472 | 0.0016 | 0.00 | 0.00 | 2.39 |
| 7473 | 0.0022 | 0.00 | 0.00 | 2.39 |
| 7474 | 0.0010 | 0.00 | 0.00 | 2.39 |
| 7475 | 0.0019 | 0.00 | 0.00 | 2.39 |
| 7476 | 0.0006 | 0.00 | 0.00 | 2.39 |
| 7477 | 0.0016 | 0.00 | 0.00 | 2.39 |
| 7478 | 0.0019 | 0.00 | 0.00 | 2.38 |
| 7479 | 0.0013 | 0.00 | 0.00 | 2.38 |
| 7480 | 0.0019 | 0.00 | 0.00 | 2.38 |
| 7481 | 0.0013 | 0.00 | 0.00 | 2.38 |
| 7482 | 0.0010 | 0.00 | 0.00 | 2.38 |
| 7483 | 0.0010 | 0.00 | 0.00 | 2.38 |
| 7484 | 0.0010 | 0.00 | 0.00 | 2.38 |

| | | | | |
|---|---|---|---|---|
| 7485 | 0.0003 | 0.00 | 0.00 | 2.38 |
| 7486 | 0.0016 | 0.00 | 0.00 | 2.38 |
| 7487 | 0.0010 | 0.00 | 0.00 | 2.37 |
| 7488 | 0.0026 | 0.00 | 0.00 | 2.37 |
| 7489 | 0.0013 | 0.00 | 0.00 | 2.37 |
| 7490 | 0.0013 | 0.00 | 0.00 | 2.37 |
| 7492 | 0.0003 | 0.00 | 0.00 | 2.37 |
| 7493 | 0.0019 | 0.00 | 0.00 | 2.37 |
| 7494 | 0.0006 | 0.00 | 0.00 | 2.37 |
| 7495 | 0.0016 | 0.00 | 0.00 | 2.36 |
| 7496 | 0.0006 | 0.00 | 0.00 | 2.36 |
| 7498 | 0.0006 | 0.00 | 0.00 | 2.36 |
| 7499 | 0.0013 | 0.00 | 0.00 | 2.36 |
| 7500 | 0.0003 | 0.00 | 0.00 | 2.36 |
| 7501 | 0.0006 | 0.00 | 0.00 | 2.36 |
| 7502 | 0.0016 | 0.00 | 0.00 | 2.36 |
| 7503 | 0.0010 | 0.00 | 0.00 | 2.36 |
| 7504 | 0.0006 | 0.00 | 0.00 | 2.36 |
| 7505 | 0.0016 | 0.00 | 0.00 | 2.36 |
| 7506 | 0.0006 | 0.00 | 0.00 | 2.35 |
| 7507 | 0.0035 | 0.00 | 0.00 | 2.35 |
| 7508 | 0.0026 | 0.00 | 0.00 | 2.35 |
| 7509 | 0.0016 | 0.00 | 0.00 | 2.35 |
| 7510 | 0.0010 | 0.00 | 0.00 | 2.35 |
| 7511 | 0.0022 | 0.00 | 0.00 | 2.35 |
| 7512 | 0.0006 | 0.00 | 0.00 | 2.34 |
| 7513 | 0.0013 | 0.00 | 0.00 | 2.34 |
| 7514 | 0.0006 | 0.00 | 0.00 | 2.34 |
| 7515 | 0.0022 | 0.00 | 0.00 | 2.34 |
| 7516 | 0.0010 | 0.00 | 0.00 | 2.34 |
| 7517 | 0.0013 | 0.00 | 0.00 | 2.34 |
| 7518 | 0.0019 | 0.00 | 0.00 | 2.34 |
| 7519 | 0.0010 | 0.00 | 0.00 | 2.33 |
| 7520 | 0.0013 | 0.00 | 0.00 | 2.33 |
| 7521 | 0.0010 | 0.00 | 0.00 | 2.33 |
| 7522 | 0.0010 | 0.00 | 0.00 | 2.33 |
| 7523 | 0.0003 | 0.00 | 0.00 | 2.33 |
| 7524 | 0.0003 | 0.00 | 0.00 | 2.33 |
| 7525 | 0.0010 | 0.00 | 0.00 | 2.33 |
| 7526 | 0.0006 | 0.00 | 0.00 | 2.33 |
| 7527 | 0.0006 | 0.00 | 0.00 | 2.33 |
| 7528 | 0.0016 | 0.00 | 0.00 | 2.33 |
| 7529 | 0.0022 | 0.00 | 0.00 | 2.33 |
| 7530 | 0.0013 | 0.00 | 0.00 | 2.32 |

| | | | | |
|---|---|---|---|---|
| 7531 | 0.0003 | 0.00 | 0.00 | 2.32 |
| 7532 | 0.0016 | 0.00 | 0.00 | 2.32 |
| 7533 | 0.0019 | 0.00 | 0.00 | 2.32 |
| 7534 | 0.0006 | 0.00 | 0.00 | 2.32 |
| 7535 | 0.0019 | 0.00 | 0.00 | 2.32 |
| 7536 | 0.0016 | 0.00 | 0.00 | 2.32 |
| 7537 | 0.0006 | 0.00 | 0.00 | 2.31 |
| 7538 | 0.0013 | 0.00 | 0.00 | 2.31 |
| 7539 | 0.0006 | 0.00 | 0.00 | 2.31 |
| 7540 | 0.0029 | 0.00 | 0.00 | 2.31 |
| 7541 | 0.0010 | 0.00 | 0.00 | 2.31 |
| 7542 | 0.0006 | 0.00 | 0.00 | 2.31 |
| 7543 | 0.0006 | 0.00 | 0.00 | 2.31 |
| 7544 | 0.0016 | 0.00 | 0.00 | 2.31 |
| 7545 | 0.0010 | 0.00 | 0.00 | 2.30 |
| 7546 | 0.0022 | 0.00 | 0.00 | 2.30 |
| 7547 | 0.0010 | 0.00 | 0.00 | 2.30 |
| 7548 | 0.0016 | 0.00 | 0.00 | 2.30 |
| 7549 | 0.0006 | 0.00 | 0.00 | 2.30 |
| 7550 | 0.0010 | 0.00 | 0.00 | 2.30 |
| 7551 | 0.0010 | 0.00 | 0.00 | 2.30 |
| 7552 | 0.0010 | 0.00 | 0.00 | 2.30 |
| 7553 | 0.0006 | 0.00 | 0.00 | 2.30 |
| 7554 | 0.0016 | 0.00 | 0.00 | 2.29 |
| 7555 | 0.0013 | 0.00 | 0.00 | 2.29 |
| 7556 | 0.0016 | 0.00 | 0.00 | 2.29 |
| 7557 | 0.0010 | 0.00 | 0.00 | 2.29 |
| 7558 | 0.0013 | 0.00 | 0.00 | 2.29 |
| 7559 | 0.0022 | 0.00 | 0.00 | 2.29 |
| 7560 | 0.0003 | 0.00 | 0.00 | 2.29 |
| 7561 | 0.0019 | 0.00 | 0.00 | 2.29 |
| 7562 | 0.0003 | 0.00 | 0.00 | 2.28 |
| 7563 | 0.0003 | 0.00 | 0.00 | 2.28 |
| 7564 | 0.0016 | 0.00 | 0.00 | 2.28 |
| 7565 | 0.0010 | 0.00 | 0.00 | 2.28 |
| 7566 | 0.0022 | 0.00 | 0.00 | 2.28 |
| 7567 | 0.0022 | 0.00 | 0.00 | 2.28 |
| 7568 | 0.0019 | 0.00 | 0.00 | 2.28 |
| 7569 | 0.0010 | 0.00 | 0.00 | 2.27 |
| 7570 | 0.0026 | 0.00 | 0.00 | 2.27 |
| 7571 | 0.0019 | 0.00 | 0.00 | 2.27 |
| 7572 | 0.0006 | 0.00 | 0.00 | 2.27 |
| 7573 | 0.0016 | 0.00 | 0.00 | 2.27 |
| 7574 | 0.0022 | 0.00 | 0.00 | 2.27 |

| | | | | |
|---|---|---|---|---|
| 7575 | 0.0013 | 0.00 | 0.00 | 2.26 |
| 7576 | 0.0010 | 0.00 | 0.00 | 2.26 |
| 7577 | 0.0019 | 0.00 | 0.00 | 2.26 |
| 7578 | 0.0029 | 0.00 | 0.00 | 2.26 |
| 7579 | 0.0006 | 0.00 | 0.00 | 2.26 |
| 7580 | 0.0006 | 0.00 | 0.00 | 2.26 |
| 7581 | 0.0010 | 0.00 | 0.00 | 2.26 |
| 7582 | 0.0013 | 0.00 | 0.00 | 2.25 |
| 7583 | 0.0010 | 0.00 | 0.00 | 2.25 |
| 7584 | 0.0016 | 0.00 | 0.00 | 2.25 |
| 7585 | 0.0019 | 0.00 | 0.00 | 2.25 |
| 7586 | 0.0003 | 0.00 | 0.00 | 2.25 |
| 7587 | 0.0006 | 0.00 | 0.00 | 2.25 |
| 7588 | 0.0026 | 0.00 | 0.00 | 2.25 |
| 7589 | 0.0013 | 0.00 | 0.00 | 2.25 |
| 7590 | 0.0016 | 0.00 | 0.00 | 2.24 |
| 7591 | 0.0010 | 0.00 | 0.00 | 2.24 |
| 7592 | 0.0010 | 0.00 | 0.00 | 2.24 |
| 7593 | 0.0019 | 0.00 | 0.00 | 2.24 |
| 7594 | 0.0016 | 0.00 | 0.00 | 2.24 |
| 7595 | 0.0016 | 0.00 | 0.00 | 2.24 |
| 7596 | 0.0013 | 0.00 | 0.00 | 2.24 |
| 7597 | 0.0029 | 0.00 | 0.00 | 2.23 |
| 7598 | 0.0013 | 0.00 | 0.00 | 2.23 |
| 7599 | 0.0026 | 0.00 | 0.00 | 2.23 |
| 7600 | 0.0013 | 0.00 | 0.00 | 2.23 |
| 7601 | 0.0003 | 0.00 | 0.00 | 2.23 |
| 7602 | 0.0013 | 0.00 | 0.00 | 2.23 |
| 7603 | 0.0016 | 0.00 | 0.00 | 2.22 |
| 7604 | 0.0013 | 0.00 | 0.00 | 2.22 |
| 7605 | 0.0003 | 0.00 | 0.00 | 2.22 |
| 7606 | 0.0006 | 0.00 | 0.00 | 2.22 |
| 7607 | 0.0019 | 0.00 | 0.00 | 2.22 |
| 7608 | 0.0019 | 0.00 | 0.00 | 2.22 |
| 7609 | 0.0026 | 0.00 | 0.00 | 2.22 |
| 7610 | 0.0010 | 0.00 | 0.00 | 2.21 |
| 7611 | 0.0003 | 0.00 | 0.00 | 2.21 |
| 7612 | 0.0019 | 0.00 | 0.00 | 2.21 |
| 7613 | 0.0016 | 0.00 | 0.00 | 2.21 |
| 7614 | 0.0010 | 0.00 | 0.00 | 2.21 |
| 7615 | 0.0019 | 0.00 | 0.00 | 2.21 |
| 7616 | 0.0013 | 0.00 | 0.00 | 2.21 |
| 7617 | 0.0006 | 0.00 | 0.00 | 2.21 |
| 7618 | 0.0010 | 0.00 | 0.00 | 2.20 |

| | | | | |
|---|---|---|---|---|
| 7619 | 0.0016 | 0.00 | 0.00 | 2.20 |
| 7620 | 0.0022 | 0.00 | 0.00 | 2.20 |
| 7621 | 0.0010 | 0.00 | 0.00 | 2.20 |
| 7622 | 0.0016 | 0.00 | 0.00 | 2.20 |
| 7623 | 0.0010 | 0.00 | 0.00 | 2.20 |
| 7624 | 0.0010 | 0.00 | 0.00 | 2.20 |
| 7625 | 0.0019 | 0.00 | 0.00 | 2.20 |
| 7626 | 0.0003 | 0.00 | 0.00 | 2.19 |
| 7627 | 0.0022 | 0.00 | 0.00 | 2.19 |
| 7628 | 0.0013 | 0.00 | 0.00 | 2.19 |
| 7629 | 0.0010 | 0.00 | 0.00 | 2.19 |
| 7630 | 0.0010 | 0.00 | 0.00 | 2.19 |
| 7631 | 0.0010 | 0.00 | 0.00 | 2.19 |
| 7632 | 0.0006 | 0.00 | 0.00 | 2.19 |
| 7633 | 0.0016 | 0.00 | 0.00 | 2.19 |
| 7634 | 0.0019 | 0.00 | 0.00 | 2.18 |
| 7635 | 0.0006 | 0.00 | 0.00 | 2.18 |
| 7636 | 0.0026 | 0.00 | 0.00 | 2.18 |
| 7637 | 0.0016 | 0.00 | 0.00 | 2.18 |
| 7638 | 0.0016 | 0.00 | 0.00 | 2.18 |
| 7639 | 0.0032 | 0.00 | 0.00 | 2.18 |
| 7640 | 0.0022 | 0.00 | 0.00 | 2.17 |
| 7641 | 0.0013 | 0.00 | 0.00 | 2.17 |
| 7642 | 0.0006 | 0.00 | 0.00 | 2.17 |
| 7643 | 0.0022 | 0.00 | 0.00 | 2.17 |
| 7644 | 0.0016 | 0.00 | 0.00 | 2.17 |
| 7645 | 0.0035 | 0.00 | 0.00 | 2.16 |
| 7646 | 0.0013 | 0.00 | 0.00 | 2.16 |
| 7647 | 0.0022 | 0.00 | 0.00 | 2.16 |
| 7648 | 0.0010 | 0.00 | 0.00 | 2.16 |
| 7649 | 0.0006 | 0.00 | 0.00 | 2.16 |
| 7650 | 0.0010 | 0.00 | 0.00 | 2.16 |
| 7651 | 0.0010 | 0.00 | 0.00 | 2.16 |
| 7652 | 0.0022 | 0.00 | 0.00 | 2.15 |
| 7653 | 0.0013 | 0.00 | 0.00 | 2.15 |
| 7654 | 0.0006 | 0.00 | 0.00 | 2.15 |
| 7655 | 0.0016 | 0.00 | 0.00 | 2.15 |
| 7656 | 0.0016 | 0.00 | 0.00 | 2.15 |
| 7657 | 0.0003 | 0.00 | 0.00 | 2.15 |
| 7658 | 0.0006 | 0.00 | 0.00 | 2.15 |
| 7659 | 0.0019 | 0.00 | 0.00 | 2.15 |
| 7660 | 0.0016 | 0.00 | 0.00 | 2.14 |
| 7661 | 0.0010 | 0.00 | 0.00 | 2.14 |
| 7662 | 0.0016 | 0.00 | 0.00 | 2.14 |

| | | | | |
|---|---|---|---|---|
| 7663 | 0.0006 | 0.00 | 0.00 | 2.14 |
| 7664 | 0.0035 | 0.00 | 0.00 | 2.14 |
| 7665 | 0.0016 | 0.00 | 0.00 | 2.14 |
| 7666 | 0.0029 | 0.00 | 0.00 | 2.13 |
| 7667 | 0.0019 | 0.00 | 0.00 | 2.13 |
| 7668 | 0.0010 | 0.00 | 0.00 | 2.13 |
| 7669 | 0.0016 | 0.00 | 0.00 | 2.13 |
| 7670 | 0.0019 | 0.00 | 0.00 | 2.13 |
| 7671 | 0.0016 | 0.00 | 0.00 | 2.12 |
| 7672 | 0.0016 | 0.00 | 0.00 | 2.12 |
| 7673 | 0.0019 | 0.00 | 0.00 | 2.12 |
| 7674 | 0.0016 | 0.00 | 0.00 | 2.12 |
| 7675 | 0.0010 | 0.00 | 0.00 | 2.12 |
| 7676 | 0.0003 | 0.00 | 0.00 | 2.12 |
| 7677 | 0.0029 | 0.00 | 0.00 | 2.12 |
| 7678 | 0.0035 | 0.00 | 0.00 | 2.11 |
| 7679 | 0.0013 | 0.00 | 0.00 | 2.11 |
| 7680 | 0.0016 | 0.00 | 0.00 | 2.11 |
| 7681 | 0.0022 | 0.00 | 0.00 | 2.11 |
| 7682 | 0.0006 | 0.00 | 0.00 | 2.11 |
| 7683 | 0.0010 | 0.00 | 0.00 | 2.10 |
| 7684 | 0.0016 | 0.00 | 0.00 | 2.10 |
| 7685 | 0.0013 | 0.00 | 0.00 | 2.10 |
| 7686 | 0.0010 | 0.00 | 0.00 | 2.10 |
| 7687 | 0.0022 | 0.00 | 0.00 | 2.10 |
| 7688 | 0.0006 | 0.00 | 0.00 | 2.10 |
| 7689 | 0.0022 | 0.00 | 0.00 | 2.10 |
| 7690 | 0.0010 | 0.00 | 0.00 | 2.09 |
| 7691 | 0.0019 | 0.00 | 0.00 | 2.09 |
| 7692 | 0.0013 | 0.00 | 0.00 | 2.09 |
| 7694 | 0.0016 | 0.00 | 0.00 | 2.09 |
| 7695 | 0.0022 | 0.00 | 0.00 | 2.09 |
| 7696 | 0.0003 | 0.00 | 0.00 | 2.09 |
| 7697 | 0.0016 | 0.00 | 0.00 | 2.09 |
| 7698 | 0.0010 | 0.00 | 0.00 | 2.08 |
| 7699 | 0.0019 | 0.00 | 0.00 | 2.08 |
| 7700 | 0.0019 | 0.00 | 0.00 | 2.08 |
| 7701 | 0.0022 | 0.00 | 0.00 | 2.08 |
| 7702 | 0.0026 | 0.00 | 0.00 | 2.08 |
| 7703 | 0.0013 | 0.00 | 0.00 | 2.08 |
| 7704 | 0.0013 | 0.00 | 0.00 | 2.07 |
| 7705 | 0.0019 | 0.00 | 0.00 | 2.07 |
| 7706 | 0.0006 | 0.00 | 0.00 | 2.07 |
| 7707 | 0.0016 | 0.00 | 0.00 | 2.07 |

| | | | | |
|---|---|---|---|---|
| 7708 | 0.0016 | 0.00 | 0.00 | 2.07 |
| 7709 | 0.0006 | 0.00 | 0.00 | 2.07 |
| 7710 | 0.0010 | 0.00 | 0.00 | 2.07 |
| 7711 | 0.0003 | 0.00 | 0.00 | 2.07 |
| 7712 | 0.0003 | 0.00 | 0.00 | 2.06 |
| 7713 | 0.0016 | 0.00 | 0.00 | 2.06 |
| 7714 | 0.0019 | 0.00 | 0.00 | 2.06 |
| 7715 | 0.0010 | 0.00 | 0.00 | 2.06 |
| 7716 | 0.0019 | 0.00 | 0.00 | 2.06 |
| 7717 | 0.0010 | 0.00 | 0.00 | 2.06 |
| 7718 | 0.0010 | 0.00 | 0.00 | 2.06 |
| 7719 | 0.0016 | 0.00 | 0.00 | 2.06 |
| 7720 | 0.0013 | 0.00 | 0.00 | 2.05 |
| 7721 | 0.0022 | 0.00 | 0.00 | 2.05 |
| 7722 | 0.0013 | 0.00 | 0.00 | 2.05 |
| 7723 | 0.0026 | 0.00 | 0.00 | 2.05 |
| 7724 | 0.0010 | 0.00 | 0.00 | 2.05 |
| 7725 | 0.0013 | 0.00 | 0.00 | 2.05 |
| 7726 | 0.0022 | 0.00 | 0.00 | 2.04 |
| 7727 | 0.0022 | 0.00 | 0.00 | 2.04 |
| 7728 | 0.0019 | 0.00 | 0.00 | 2.04 |
| 7729 | 0.0010 | 0.00 | 0.00 | 2.04 |
| 7730 | 0.0006 | 0.00 | 0.00 | 2.04 |
| 7731 | 0.0029 | 0.00 | 0.00 | 2.04 |
| 7732 | 0.0006 | 0.00 | 0.00 | 2.03 |
| 7733 | 0.0016 | 0.00 | 0.00 | 2.03 |
| 7734 | 0.0022 | 0.00 | 0.00 | 2.03 |
| 7735 | 0.0013 | 0.00 | 0.00 | 2.03 |
| 7736 | 0.0035 | 0.00 | 0.00 | 2.03 |
| 7737 | 0.0013 | 0.00 | 0.00 | 2.02 |
| 7738 | 0.0016 | 0.00 | 0.00 | 2.02 |
| 7739 | 0.0016 | 0.00 | 0.00 | 2.02 |
| 7740 | 0.0019 | 0.00 | 0.00 | 2.02 |
| 7741 | 0.0029 | 0.00 | 0.00 | 2.02 |
| 7742 | 0.0032 | 0.00 | 0.00 | 2.02 |
| 7743 | 0.0016 | 0.00 | 0.00 | 2.01 |
| 7744 | 0.0013 | 0.00 | 0.00 | 2.01 |
| 7745 | 0.0013 | 0.00 | 0.00 | 2.01 |
| 7746 | 0.0019 | 0.00 | 0.00 | 2.01 |
| 7747 | 0.0016 | 0.00 | 0.00 | 2.01 |
| 7748 | 0.0026 | 0.00 | 0.00 | 2.00 |
| 7749 | 0.0006 | 0.00 | 0.00 | 2.00 |
| 7750 | 0.0013 | 0.00 | 0.00 | 2.00 |
| 7751 | 0.0010 | 0.00 | 0.00 | 2.00 |

| | | | | |
|---|---|---|---|---|
| 7752 | 0.0006 | 0.00 | 0.00 | 2.00 |
| 7753 | 0.0013 | 0.00 | 0.00 | 2.00 |
| 7754 | 0.0016 | 0.00 | 0.00 | 2.00 |
| 7755 | 0.0032 | 0.00 | 0.00 | 2.00 |
| 7756 | 0.0026 | 0.00 | 0.00 | 1.99 |
| 7757 | 0.0010 | 0.00 | 0.00 | 1.99 |
| 7758 | 0.0029 | 0.00 | 0.00 | 1.99 |
| 7759 | 0.0016 | 0.00 | 0.00 | 1.99 |
| 7760 | 0.0016 | 0.00 | 0.00 | 1.98 |
| 7761 | 0.0006 | 0.00 | 0.00 | 1.98 |
| 7762 | 0.0019 | 0.00 | 0.00 | 1.98 |
| 7763 | 0.0019 | 0.00 | 0.00 | 1.98 |
| 7764 | 0.0026 | 0.00 | 0.00 | 1.98 |
| 7765 | 0.0003 | 0.00 | 0.00 | 1.98 |
| 7766 | 0.0019 | 0.00 | 0.00 | 1.98 |
| 7767 | 0.0019 | 0.00 | 0.00 | 1.97 |
| 7768 | 0.0016 | 0.00 | 0.00 | 1.97 |
| 7769 | 0.0013 | 0.00 | 0.00 | 1.97 |
| 7770 | 0.0016 | 0.00 | 0.00 | 1.97 |
| 7771 | 0.0010 | 0.00 | 0.00 | 1.97 |
| 7772 | 0.0022 | 0.00 | 0.00 | 1.97 |
| 7773 | 0.0010 | 0.00 | 0.00 | 1.96 |
| 7774 | 0.0013 | 0.00 | 0.00 | 1.96 |
| 7775 | 0.0022 | 0.00 | 0.00 | 1.96 |
| 7776 | 0.0019 | 0.00 | 0.00 | 1.96 |
| 7777 | 0.0019 | 0.00 | 0.00 | 1.96 |
| 7778 | 0.0010 | 0.00 | 0.00 | 1.96 |
| 7779 | 0.0022 | 0.00 | 0.00 | 1.95 |
| 7780 | 0.0016 | 0.00 | 0.00 | 1.95 |
| 7781 | 0.0022 | 0.00 | 0.00 | 1.95 |
| 7782 | 0.0013 | 0.00 | 0.00 | 1.95 |
| 7783 | 0.0010 | 0.00 | 0.00 | 1.95 |
| 7784 | 0.0016 | 0.00 | 0.00 | 1.95 |
| 7785 | 0.0022 | 0.00 | 0.00 | 1.94 |
| 7786 | 0.0016 | 0.00 | 0.00 | 1.94 |
| 7787 | 0.0026 | 0.00 | 0.00 | 1.94 |
| 7788 | 0.0026 | 0.00 | 0.00 | 1.94 |
| 7789 | 0.0032 | 0.00 | 0.00 | 1.94 |
| 7790 | 0.0016 | 0.00 | 0.00 | 1.93 |
| 7791 | 0.0019 | 0.00 | 0.00 | 1.93 |
| 7792 | 0.0019 | 0.00 | 0.00 | 1.93 |
| 7793 | 0.0022 | 0.00 | 0.00 | 1.93 |
| 7794 | 0.0026 | 0.00 | 0.00 | 1.92 |
| 7795 | 0.0013 | 0.00 | 0.00 | 1.92 |

| | | | | |
|---|---|---|---|---|
| 7796 | 0.0022 | 0.00 | 0.00 | 1.92 |
| 7797 | 0.0022 | 0.00 | 0.00 | 1.92 |
| 7798 | 0.0022 | 0.00 | 0.00 | 1.92 |
| 7799 | 0.0022 | 0.00 | 0.00 | 1.91 |
| 7800 | 0.0029 | 0.00 | 0.00 | 1.91 |
| 7801 | 0.0022 | 0.00 | 0.00 | 1.91 |
| 7802 | 0.0019 | 0.00 | 0.00 | 1.91 |
| 7803 | 0.0026 | 0.00 | 0.00 | 1.90 |
| 7804 | 0.0019 | 0.00 | 0.00 | 1.90 |
| 7805 | 0.0022 | 0.00 | 0.00 | 1.90 |
| 7806 | 0.0013 | 0.00 | 0.00 | 1.90 |
| 7807 | 0.0035 | 0.00 | 0.00 | 1.90 |
| 7808 | 0.0019 | 0.00 | 0.00 | 1.89 |
| 7809 | 0.0022 | 0.00 | 0.00 | 1.89 |
| 7810 | 0.0019 | 0.00 | 0.00 | 1.89 |
| 7811 | 0.0035 | 0.00 | 0.00 | 1.89 |
| 7812 | 0.0019 | 0.00 | 0.00 | 1.88 |
| 7813 | 0.0013 | 0.00 | 0.00 | 1.88 |
| 7814 | 0.0013 | 0.00 | 0.00 | 1.88 |
| 7815 | 0.0022 | 0.00 | 0.00 | 1.88 |
| 7816 | 0.0022 | 0.00 | 0.00 | 1.88 |
| 7817 | 0.0016 | 0.00 | 0.00 | 1.87 |
| 7818 | 0.0019 | 0.00 | 0.00 | 1.87 |
| 7819 | 0.0022 | 0.00 | 0.00 | 1.87 |
| 7820 | 0.0013 | 0.00 | 0.00 | 1.87 |
| 7821 | 0.0032 | 0.00 | 0.00 | 1.87 |
| 7822 | 0.0038 | 0.00 | 0.00 | 1.86 |
| 7823 | 0.0010 | 0.00 | 0.00 | 1.86 |
| 7824 | 0.0016 | 0.00 | 0.00 | 1.86 |
| 7825 | 0.0029 | 0.00 | 0.00 | 1.86 |
| 7826 | 0.0019 | 0.00 | 0.00 | 1.86 |
| 7827 | 0.0035 | 0.00 | 0.00 | 1.85 |
| 7828 | 0.0016 | 0.00 | 0.00 | 1.85 |
| 7829 | 0.0029 | 0.00 | 0.00 | 1.85 |
| 7830 | 0.0042 | 0.00 | 0.00 | 1.85 |
| 7831 | 0.0026 | 0.00 | 0.00 | 1.84 |
| 7832 | 0.0029 | 0.00 | 0.00 | 1.84 |
| 7833 | 0.0029 | 0.00 | 0.00 | 1.84 |
| 7834 | 0.0022 | 0.00 | 0.00 | 1.83 |
| 7835 | 0.0032 | 0.00 | 0.00 | 1.83 |
| 7836 | 0.0035 | 0.00 | 0.00 | 1.83 |
| 7837 | 0.0029 | 0.00 | 0.00 | 1.82 |
| 7838 | 0.0016 | 0.00 | 0.00 | 1.82 |
| 7839 | 0.0016 | 0.00 | 0.00 | 1.82 |

| | | | | |
|---|---|---|---|---|
| 7840 | 0.0035 | 0.00 | 0.00 | 1.82 |
| 7841 | 0.0022 | 0.00 | 0.00 | 1.81 |
| 7842 | 0.0048 | 0.00 | 0.00 | 1.81 |
| 7843 | 0.0026 | 0.00 | 0.00 | 1.81 |
| 7844 | 0.0019 | 0.00 | 0.00 | 1.80 |
| 7845 | 0.0016 | 0.00 | 0.00 | 1.80 |
| 7846 | 0.0029 | 0.00 | 0.00 | 1.80 |
| 7847 | 0.0013 | 0.00 | 0.00 | 1.80 |
| 7848 | 0.0019 | 0.00 | 0.00 | 1.80 |
| 7849 | 0.0032 | 0.00 | 0.00 | 1.79 |
| 7850 | 0.0026 | 0.00 | 0.00 | 1.79 |
| 7851 | 0.0045 | 0.00 | 0.00 | 1.79 |
| 7852 | 0.0032 | 0.00 | 0.00 | 1.78 |
| 7853 | 0.0054 | 0.00 | 0.00 | 1.78 |
| 7854 | 0.0026 | 0.00 | 0.00 | 1.78 |
| 7855 | 0.0032 | 0.00 | 0.00 | 1.77 |
| 7856 | 0.0022 | 0.00 | 0.00 | 1.77 |
| 7857 | 0.0026 | 0.00 | 0.00 | 1.77 |
| 7858 | 0.0026 | 0.00 | 0.00 | 1.77 |
| 7859 | 0.0022 | 0.00 | 0.00 | 1.76 |
| 7860 | 0.0035 | 0.00 | 0.00 | 1.76 |
| 7861 | 0.0026 | 0.00 | 0.00 | 1.76 |
| 7862 | 0.0022 | 0.00 | 0.00 | 1.75 |
| 7863 | 0.0019 | 0.00 | 0.00 | 1.75 |
| 7864 | 0.0029 | 0.00 | 0.00 | 1.75 |
| 7865 | 0.0042 | 0.00 | 0.00 | 1.75 |
| 7866 | 0.0029 | 0.00 | 0.00 | 1.74 |
| 7867 | 0.0032 | 0.00 | 0.00 | 1.74 |
| 7868 | 0.0032 | 0.00 | 0.00 | 1.74 |
| 7869 | 0.0029 | 0.00 | 0.00 | 1.73 |
| 7870 | 0.0032 | 0.00 | 0.00 | 1.73 |
| 7871 | 0.0022 | 0.00 | 0.00 | 1.73 |
| 7872 | 0.0042 | 0.00 | 0.00 | 1.73 |
| 7873 | 0.0026 | 0.00 | 0.00 | 1.72 |
| 7874 | 0.0035 | 0.00 | 0.00 | 1.72 |
| 7875 | 0.0051 | 0.00 | 0.00 | 1.72 |
| 7876 | 0.0026 | 0.00 | 0.00 | 1.71 |
| 7877 | 0.0035 | 0.00 | 0.00 | 1.71 |
| 7878 | 0.0035 | 0.00 | 0.00 | 1.70 |
| 7879 | 0.0042 | 0.00 | 0.00 | 1.70 |
| 7880 | 0.0067 | 0.00 | 0.00 | 1.70 |
| 7881 | 0.0035 | 0.00 | 0.00 | 1.69 |
| 7882 | 0.0035 | 0.00 | 0.00 | 1.69 |
| 7883 | 0.0032 | 0.00 | 0.00 | 1.68 |

| | | | | |
|---|---|---|---|---|
| 7884 | 0.0042 | 0.00 | 0.00 | 1.68 |
| 7885 | 0.0048 | 0.00 | 0.00 | 1.68 |
| 7886 | 0.0035 | 0.00 | 0.00 | 1.67 |
| 7887 | 0.0045 | 0.00 | 0.00 | 1.67 |
| 7888 | 0.0058 | 0.00 | 0.00 | 1.66 |
| 7889 | 0.0042 | 0.00 | 0.00 | 1.66 |
| 7890 | 0.0067 | 0.00 | 0.00 | 1.65 |
| 7891 | 0.0029 | 0.00 | 0.00 | 1.65 |
| 7892 | 0.0048 | 0.00 | 0.00 | 1.64 |
| 7893 | 0.0038 | 0.00 | 0.00 | 1.64 |
| 7894 | 0.0019 | 0.00 | 0.00 | 1.63 |
| 7895 | 0.0032 | 0.00 | 0.00 | 1.63 |
| 7896 | 0.0045 | 0.00 | 0.00 | 1.63 |
| 7897 | 0.0054 | 0.00 | 0.00 | 1.62 |
| 7898 | 0.0058 | 0.00 | 0.00 | 1.62 |
| 7899 | 0.0022 | 0.00 | 0.00 | 1.61 |
| 7900 | 0.0038 | 0.00 | 0.00 | 1.61 |
| 7901 | 0.0029 | 0.00 | 0.00 | 1.61 |
| 7902 | 0.0038 | 0.00 | 0.00 | 1.60 |
| 7903 | 0.0087 | 0.00 | 0.00 | 1.60 |
| 7904 | 0.0029 | 0.00 | 0.00 | 1.59 |
| 7905 | 0.0042 | 0.00 | 0.00 | 1.59 |
| 7906 | 0.0054 | 0.00 | 0.00 | 1.58 |
| 7907 | 0.0058 | 0.00 | 0.00 | 1.58 |
| 7908 | 0.0074 | 0.00 | 0.00 | 1.57 |
| 7909 | 0.0048 | 0.00 | 0.00 | 1.57 |
| 7910 | 0.0074 | 0.00 | 0.00 | 1.56 |
| 7911 | 0.0067 | 0.00 | 0.00 | 1.55 |
| 7912 | 0.0054 | 0.00 | 0.00 | 1.55 |
| 7913 | 0.0080 | 0.00 | 0.00 | 1.54 |
| 7914 | 0.0032 | 0.00 | 0.00 | 1.53 |
| 7915 | 0.0032 | 0.00 | 0.00 | 1.53 |
| 7916 | 0.0038 | 0.00 | 0.00 | 1.53 |
| 7917 | 0.0054 | 0.00 | 0.00 | 1.52 |
| 7918 | 0.0035 | 0.00 | 0.00 | 1.52 |
| 7919 | 0.0064 | 0.00 | 0.00 | 1.51 |
| 7920 | 0.0074 | 0.00 | 0.00 | 1.51 |
| 7921 | 0.0054 | 0.00 | 0.00 | 1.50 |
| 7922 | 0.0048 | 0.00 | 0.00 | 1.50 |
| 7923 | 0.0038 | 0.00 | 0.00 | 1.49 |
| 7924 | 0.0048 | 0.00 | 0.00 | 1.49 |
| 7925 | 0.0064 | 0.00 | 0.00 | 1.48 |
| 7926 | 0.0048 | 0.00 | 0.00 | 1.48 |
| 7927 | 0.0064 | 0.00 | 0.00 | 1.47 |

| | | | | |
|---|---|---|---|---|
| 7928 | 0.0077 | 0.00 | 0.00 | 1.46 |
| 7929 | 0.0080 | 0.00 | 0.00 | 1.46 |
| 7930 | 0.0067 | 0.00 | 0.00 | 1.45 |
| 7931 | 0.0074 | 0.00 | 0.00 | 1.44 |
| 7932 | 0.0048 | 0.00 | 0.00 | 1.43 |
| 7933 | 0.0096 | 0.00 | 0.00 | 1.43 |
| 7934 | 0.0045 | 0.00 | 0.00 | 1.42 |
| 7935 | 0.0064 | 0.00 | 0.00 | 1.42 |
| 7936 | 0.0090 | 0.00 | 0.00 | 1.41 |
| 7937 | 0.0090 | 0.00 | 0.00 | 1.40 |
| 7938 | 0.0099 | 0.00 | 0.00 | 1.39 |
| 7939 | 0.0064 | 0.00 | 0.00 | 1.38 |
| 7940 | 0.0099 | 0.00 | 0.00 | 1.38 |
| 7941 | 0.0115 | 0.00 | 0.00 | 1.37 |
| 7942 | 0.0122 | 0.00 | 0.00 | 1.35 |
| 7943 | 0.0099 | 0.00 | 0.00 | 1.34 |
| 7944 | 0.0141 | 0.00 | 0.00 | 1.33 |
| 7945 | 0.0131 | 0.00 | 0.00 | 1.32 |
| 7946 | 0.0087 | 0.00 | 0.00 | 1.30 |
| 7947 | 0.0135 | 0.00 | 0.00 | 1.30 |
| 7948 | 0.0115 | 0.00 | 0.00 | 1.28 |
| 7949 | 0.0096 | 0.00 | 0.00 | 1.27 |
| 7950 | 0.0115 | 0.00 | 0.00 | 1.26 |
| 7951 | 0.0138 | 0.00 | 0.00 | 1.25 |
| 7952 | 0.0147 | 0.00 | 0.00 | 1.24 |
| 7953 | 0.0106 | 0.00 | 0.00 | 1.22 |
| 7954 | 0.0115 | 0.00 | 0.00 | 1.21 |
| 7955 | 0.0103 | 0.00 | 0.00 | 1.20 |
| 7956 | 0.0147 | 0.00 | 0.00 | 1.19 |
| 7957 | 0.0144 | 0.00 | 0.00 | 1.17 |
| 7958 | 0.0115 | 0.00 | 0.00 | 1.16 |
| 7959 | 0.0135 | 0.00 | 0.00 | 1.15 |
| 7960 | 0.0128 | 0.00 | 0.00 | 1.13 |
| 7961 | 0.0189 | 0.00 | 0.00 | 1.12 |
| 7962 | 0.0186 | 0.00 | 0.00 | 1.10 |
| 7963 | 0.0202 | 0.00 | 0.00 | 1.08 |
| 7964 | 0.0202 | 0.00 | 0.00 | 1.06 |
| 7965 | 0.0202 | 0.00 | 0.00 | 1.04 |
| 7966 | 0.0186 | 0.00 | 0.00 | 1.02 |
| 7967 | 0.0215 | 0.00 | 0.00 | 1.00 |
| 7968 | 0.0228 | 0.00 | 0.00 | 0.98 |
| 7969 | 0.0212 | 0.00 | 0.00 | 0.96 |
| 7970 | 0.0176 | 0.00 | 0.00 | 0.94 |
| 7971 | 0.0234 | 0.00 | 0.00 | 0.92 |

| | | | | |
|---|---|---|---|---|
| 7972 | 0.0244 | 0.00 | 0.00 | 0.90 |
| 7973 | 0.0212 | 0.00 | 0.00 | 0.87 |
| 7974 | 0.0205 | 0.00 | 0.00 | 0.85 |
| 7975 | 0.0253 | 0.00 | 0.00 | 0.83 |
| 7976 | 0.0202 | 0.00 | 0.00 | 0.81 |
| 7977 | 0.0221 | 0.00 | 0.00 | 0.79 |
| 7978 | 0.0218 | 0.00 | 0.00 | 0.76 |
| 7979 | 0.0314 | 0.00 | 0.01 | 0.74 |
| 7980 | 0.0308 | 0.00 | 0.01 | 0.71 |
| 7981 | 0.0272 | 0.00 | 0.01 | 0.68 |
| 7982 | 0.0247 | 0.00 | 0.00 | 0.65 |
| 7983 | 0.0279 | 0.00 | 0.01 | 0.63 |
| 7984 | 0.0231 | 0.00 | 0.00 | 0.60 |
| 7985 | 0.0276 | 0.00 | 0.01 | 0.58 |
| 7986 | 0.0292 | 0.00 | 0.01 | 0.55 |
| 7987 | 0.0317 | 0.00 | 0.01 | 0.52 |
| 7988 | 0.0279 | 0.00 | 0.00 | 0.49 |
| 7989 | 0.0253 | 0.00 | 0.00 | 0.46 |
| 7990 | 0.0282 | 0.00 | 0.01 | 0.44 |
| 7991 | 0.0321 | 0.00 | 0.01 | 0.41 |
| 7992 | 0.0314 | 0.00 | 0.01 | 0.38 |
| 7993 | 0.0247 | 0.00 | 0.00 | 0.34 |
| 7994 | 0.0269 | 0.00 | 0.01 | 0.32 |
| 7995 | 0.0237 | 0.00 | 0.00 | 0.29 |
| 7996 | 0.0276 | 0.00 | 0.01 | 0.27 |
| 7997 | 0.0212 | 0.00 | 0.00 | 0.24 |
| 7998 | 0.0205 | 0.00 | 0.00 | 0.22 |
| 7999 | 0.0183 | 0.00 | 0.00 | 0.20 |
| 8000 | 0.0131 | 0.00 | 0.00 | 0.18 |
| 8001 | 0.0096 | 0.00 | 0.00 | 0.17 |
| 8002 | 0.0080 | 0.00 | 0.00 | 0.16 |
| 8003 | 0.0080 | 0.00 | 0.00 | 0.15 |
| 8004 | 0.0077 | 0.00 | 0.00 | 0.14 |
| 8005 | 0.0064 | 0.00 | 0.00 | 0.14 |
| 8006 | 0.0067 | 0.00 | 0.00 | 0.13 |
| 8007 | 0.0064 | 0.00 | 0.00 | 0.12 |
| 8008 | 0.0071 | 0.00 | 0.00 | 0.12 |
| 8009 | 0.0061 | 0.00 | 0.00 | 0.11 |
| 8010 | 0.0067 | 0.00 | 0.00 | 0.10 |
| 8011 | 0.0051 | 0.00 | 0.00 | 0.10 |
| 8012 | 0.0048 | 0.00 | 0.00 | 0.09 |
| 8013 | 0.0064 | 0.00 | 0.00 | 0.09 |
| 8014 | 0.0048 | 0.00 | 0.00 | 0.08 |
| 8015 | 0.0029 | 0.00 | 0.00 | 0.07 |

| | | | | |
|---|---|---|---|---|
| 8016 | 0.0042 | 0.00 | 0.00 | 0.07 |
| 8017 | 0.0029 | 0.00 | 0.00 | 0.07 |
| 8018 | 0.0026 | 0.00 | 0.00 | 0.06 |
| 8019 | 0.0038 | 0.00 | 0.00 | 0.06 |
| 8020 | 0.0029 | 0.00 | 0.00 | 0.06 |
| 8021 | 0.0032 | 0.00 | 0.00 | 0.06 |
| 8022 | 0.0026 | 0.00 | 0.00 | 0.05 |
| 8023 | 0.0022 | 0.00 | 0.00 | 0.05 |
| 8024 | 0.0016 | 0.00 | 0.00 | 0.05 |
| 8025 | 0.0016 | 0.00 | 0.00 | 0.05 |
| 8026 | 0.0019 | 0.00 | 0.00 | 0.04 |
| 8027 | 0.0013 | 0.00 | 0.00 | 0.04 |
| 8028 | 0.0013 | 0.00 | 0.00 | 0.04 |
| 8029 | 0.0013 | 0.00 | 0.00 | 0.04 |
| 8030 | 0.0013 | 0.00 | 0.00 | 0.04 |
| 8031 | 0.0016 | 0.00 | 0.00 | 0.04 |
| 8032 | 0.0013 | 0.00 | 0.00 | 0.04 |
| 8033 | 0.0013 | 0.00 | 0.00 | 0.03 |
| 8034 | 0.0013 | 0.00 | 0.00 | 0.03 |
| 8035 | 0.0016 | 0.00 | 0.00 | 0.03 |
| 8036 | 0.0013 | 0.00 | 0.00 | 0.03 |
| 8037 | 0.0013 | 0.00 | 0.00 | 0.03 |
| 8038 | 0.0006 | 0.00 | 0.00 | 0.03 |
| 8039 | 0.0010 | 0.00 | 0.00 | 0.03 |
| 8040 | 0.0013 | 0.00 | 0.00 | 0.03 |
| 8041 | 0.0010 | 0.00 | 0.00 | 0.02 |
| 8042 | 0.0010 | 0.00 | 0.00 | 0.02 |
| 8043 | 0.0006 | 0.00 | 0.00 | 0.02 |
| 8044 | 0.0010 | 0.00 | 0.00 | 0.02 |
| 8045 | 0.0013 | 0.00 | 0.00 | 0.02 |
| 8046 | 0.0010 | 0.00 | 0.00 | 0.02 |
| 8047 | 0.0013 | 0.00 | 0.00 | 0.02 |
| 8048 | 0.0006 | 0.00 | 0.00 | 0.02 |
| 8049 | 0.0006 | 0.00 | 0.00 | 0.02 |
| 8050 | 0.0010 | 0.00 | 0.00 | 0.02 |
| 8051 | 0.0006 | 0.00 | 0.00 | 0.02 |
| 8052 | 0.0003 | 0.00 | 0.00 | 0.01 |
| 8053 | 0.0003 | 0.00 | 0.00 | 0.01 |
| 8054 | 0.0003 | 0.00 | 0.00 | 0.01 |
| 8056 | 0.0006 | 0.00 | 0.00 | 0.01 |
| 8057 | 0.0003 | 0.00 | 0.00 | 0.01 |
| 8058 | 0.0003 | 0.00 | 0.00 | 0.01 |
| 8059 | 0.0003 | 0.00 | 0.00 | 0.01 |
| 8060 | 0.0003 | 0.00 | 0.00 | 0.01 |

| | | | | |
|---|---|---|---|---|
| 8061 | 0.0010 | 0.00 | 0.00 | 0.01 |
| 8062 | 0.0006 | 0.00 | 0.00 | 0.01 |
| 8063 | 0.0006 | 0.00 | 0.00 | 0.01 |
| 8065 | 0.0003 | 0.00 | 0.00 | 0.01 |
| 8067 | 0.0003 | 0.00 | 0.00 | 0.01 |
| 8068 | 0.0003 | 0.00 | 0.00 | 0.01 |
| 8069 | 0.0006 | 0.00 | 0.00 | 0.01 |
| 8070 | 0.0016 | 0.00 | 0.00 | 0.01 |
| 8071 | 0.0006 | 0.00 | 0.00 | 0.01 |
| 8072 | 0.0010 | 0.00 | 0.00 | 0.01 |
| 8073 | 0.0003 | 0.00 | 0.00 | 0.00 |
| 8074 | 0.0003 | 0.00 | 0.00 | 0.00 |
| 8075 | 0.0006 | 0.00 | 0.00 | 0.00 |
| 8076 | 0.0003 | 0.00 | 0.00 | 0.00 |
| 8077 | 0.0003 | 0.00 | 0.00 | 0.00 |
| 8078 | 0.0010 | 0.00 | 0.00 | 0.00 |
| 8079 | 0.0003 | 0.00 | 0.00 | 0.00 |
| 8080 | 0.0006 | 0.00 | 0.00 | 0.00 |
| 8081 | 0.0010 | 0.00 | 0.00 | 0.00 |

**Supplemental Table 3: Genomic coverage for non-reference sites for individual cells**

| depth | mean_count_var | median_count_var | se_count | cum_count_var |
|---|---|---|---|---|
| 2 | 38.3314 | 19 | 1.00 | 82.99 |
| 3 | 11.3105 | 6 | 0.29 | 44.66 |
| 4 | 6.6451 | 3 | 0.17 | 33.35 |
| 5 | 4.5918 | 2 | 0.12 | 26.70 |
| 6 | 3.4629 | 1 | 0.09 | 22.11 |
| 7 | 2.2919 | 1 | 0.06 | 18.65 |
| 8 | 1.8496 | 1 | 0.05 | 16.36 |
| 9 | 1.5554 | 0 | 0.05 | 14.51 |
| 10 | 1.3446 | 0 | 0.04 | 12.95 |
| 11 | 1.1302 | 0 | 0.04 | 11.61 |
| 12 | 0.8971 | 0 | 0.03 | 10.48 |
| 13 | 0.8161 | 0 | 0.03 | 9.58 |
| 14 | 0.6956 | 0 | 0.02 | 8.76 |
| 15 | 0.6490 | 0 | 0.02 | 8.07 |
| 16 | 0.5031 | 0 | 0.02 | 7.42 |
| 17 | 0.4471 | 0 | 0.02 | 6.92 |
| 18 | 0.4182 | 0 | 0.02 | 6.47 |
| 19 | 0.3893 | 0 | 0.02 | 6.05 |
| 20 | 0.3488 | 0 | 0.02 | 5.66 |
| 21 | 0.3047 | 0 | 0.01 | 5.31 |
| 22 | 0.2735 | 0 | 0.01 | 5.01 |
| 23 | 0.2578 | 0 | 0.01 | 4.74 |
| 24 | 0.2144 | 0 | 0.01 | 4.48 |
| 25 | 0.2170 | 0 | 0.01 | 4.26 |
| 26 | 0.1990 | 0 | 0.01 | 4.05 |
| 27 | 0.1803 | 0 | 0.01 | 3.85 |
| 28 | 0.1723 | 0 | 0.01 | 3.67 |
| 29 | 0.1671 | 0 | 0.01 | 3.49 |
| 30 | 0.1373 | 0 | 0.01 | 3.33 |
| 31 | 0.1250 | 0 | 0.01 | 3.19 |
| 32 | 0.1167 | 0 | 0.01 | 3.06 |
| 33 | 0.1119 | 0 | 0.01 | 2.95 |
| 34 | 0.1115 | 0 | 0.01 | 2.84 |
| 35 | 0.1054 | 0 | 0.01 | 2.72 |
| 36 | 0.1041 | 0 | 0.01 | 2.62 |
| 37 | 0.0871 | 0 | 0.01 | 2.52 |
| 38 | 0.0858 | 0 | 0.01 | 2.43 |
| 39 | 0.0726 | 0 | 0.01 | 2.34 |
| 40 | 0.0788 | 0 | 0.01 | 2.27 |
| 41 | 0.0681 | 0 | 0.01 | 2.19 |
| 42 | 0.0669 | 0 | 0.01 | 2.12 |
| 43 | 0.0659 | 0 | 0.01 | 2.06 |
| 44 | 0.0614 | 0 | 0.01 | 1.99 |

| | | | | |
|---|---|---|---|---|
| 45 | 0.0521 | 0 | 0.00 | 1.93 |
| 46 | 0.0511 | 0 | 0.00 | 1.88 |
| 47 | 0.0482 | 0 | 0.00 | 1.83 |
| 48 | 0.0485 | 0 | 0.00 | 1.78 |
| 49 | 0.0456 | 0 | 0.00 | 1.73 |
| 50 | 0.0476 | 0 | 0.00 | 1.68 |
| 51 | 0.0440 | 0 | 0.00 | 1.64 |
| 52 | 0.0421 | 0 | 0.00 | 1.59 |
| 53 | 0.0399 | 0 | 0.00 | 1.55 |
| 54 | 0.0392 | 0 | 0.00 | 1.51 |
| 55 | 0.0341 | 0 | 0.00 | 1.47 |
| 56 | 0.0328 | 0 | 0.00 | 1.44 |
| 57 | 0.0315 | 0 | 0.00 | 1.40 |
| 58 | 0.0331 | 0 | 0.00 | 1.37 |
| 59 | 0.0312 | 0 | 0.00 | 1.34 |
| 60 | 0.0238 | 0 | 0.00 | 1.31 |
| 61 | 0.0228 | 0 | 0.00 | 1.28 |
| 62 | 0.0302 | 0 | 0.00 | 1.26 |
| 63 | 0.0235 | 0 | 0.00 | 1.23 |
| 64 | 0.0222 | 0 | 0.00 | 1.21 |
| 65 | 0.0219 | 0 | 0.00 | 1.19 |
| 66 | 0.0174 | 0 | 0.00 | 1.16 |
| 67 | 0.0167 | 0 | 0.00 | 1.15 |
| 68 | 0.0225 | 0 | 0.00 | 1.13 |
| 69 | 0.0196 | 0 | 0.00 | 1.11 |
| 70 | 0.0164 | 0 | 0.00 | 1.09 |
| 71 | 0.0148 | 0 | 0.00 | 1.07 |
| 72 | 0.0151 | 0 | 0.00 | 1.06 |
| 73 | 0.0132 | 0 | 0.00 | 1.04 |
| 74 | 0.0190 | 0 | 0.00 | 1.03 |
| 75 | 0.0138 | 0 | 0.00 | 1.01 |
| 76 | 0.0119 | 0 | 0.00 | 0.99 |
| 77 | 0.0145 | 0 | 0.00 | 0.98 |
| 78 | 0.0119 | 0 | 0.00 | 0.97 |
| 79 | 0.0145 | 0 | 0.00 | 0.96 |
| 80 | 0.0125 | 0 | 0.00 | 0.94 |
| 81 | 0.0125 | 0 | 0.00 | 0.93 |
| 82 | 0.0116 | 0 | 0.00 | 0.92 |
| 83 | 0.0132 | 0 | 0.00 | 0.91 |
| 84 | 0.0090 | 0 | 0.00 | 0.89 |
| 85 | 0.0087 | 0 | 0.00 | 0.88 |
| 86 | 0.0119 | 0 | 0.00 | 0.87 |
| 87 | 0.0125 | 0 | 0.00 | 0.86 |
| 88 | 0.0090 | 0 | 0.00 | 0.85 |

| | | | | |
|---|---|---|---|---|
| 89 | 0.0100 | 0 | 0.00 | 0.84 |
| 90 | 0.0129 | 0 | 0.00 | 0.83 |
| 91 | 0.0103 | 0 | 0.00 | 0.82 |
| 92 | 0.0084 | 0 | 0.00 | 0.81 |
| 93 | 0.0096 | 0 | 0.00 | 0.80 |
| 94 | 0.0103 | 0 | 0.00 | 0.79 |
| 95 | 0.0071 | 0 | 0.00 | 0.78 |
| 96 | 0.0096 | 0 | 0.00 | 0.77 |
| 97 | 0.0080 | 0 | 0.00 | 0.76 |
| 98 | 0.0090 | 0 | 0.00 | 0.76 |
| 99 | 0.0080 | 0 | 0.00 | 0.75 |
| 100 | 0.0051 | 0 | 0.00 | 0.74 |
| 101 | 0.0077 | 0 | 0.00 | 0.73 |
| 102 | 0.0058 | 0 | 0.00 | 0.73 |
| 103 | 0.0103 | 0 | 0.00 | 0.72 |
| 104 | 0.0080 | 0 | 0.00 | 0.71 |
| 105 | 0.0042 | 0 | 0.00 | 0.70 |
| 106 | 0.0048 | 0 | 0.00 | 0.70 |
| 107 | 0.0048 | 0 | 0.00 | 0.69 |
| 108 | 0.0068 | 0 | 0.00 | 0.69 |
| 109 | 0.0048 | 0 | 0.00 | 0.68 |
| 110 | 0.0058 | 0 | 0.00 | 0.68 |
| 111 | 0.0061 | 0 | 0.00 | 0.67 |
| 112 | 0.0077 | 0 | 0.00 | 0.66 |
| 113 | 0.0058 | 0 | 0.00 | 0.66 |
| 114 | 0.0042 | 0 | 0.00 | 0.65 |
| 115 | 0.0032 | 0 | 0.00 | 0.65 |
| 116 | 0.0061 | 0 | 0.00 | 0.64 |
| 117 | 0.0039 | 0 | 0.00 | 0.64 |
| 118 | 0.0074 | 0 | 0.00 | 0.63 |
| 119 | 0.0055 | 0 | 0.00 | 0.63 |
| 120 | 0.0058 | 0 | 0.00 | 0.62 |
| 121 | 0.0023 | 0 | 0.00 | 0.61 |
| 122 | 0.0045 | 0 | 0.00 | 0.61 |
| 123 | 0.0035 | 0 | 0.00 | 0.61 |
| 124 | 0.0051 | 0 | 0.00 | 0.60 |
| 125 | 0.0032 | 0 | 0.00 | 0.60 |
| 126 | 0.0042 | 0 | 0.00 | 0.60 |
| 127 | 0.0048 | 0 | 0.00 | 0.59 |
| 128 | 0.0058 | 0 | 0.00 | 0.59 |
| 129 | 0.0032 | 0 | 0.00 | 0.58 |
| 130 | 0.0058 | 0 | 0.00 | 0.58 |
| 131 | 0.0042 | 0 | 0.00 | 0.57 |
| 132 | 0.0026 | 0 | 0.00 | 0.57 |

| | | | | |
|---|---|---|---|---|
| 133 | 0.0023 | 0 | 0.00 | 0.57 |
| 134 | 0.0039 | 0 | 0.00 | 0.56 |
| 135 | 0.0042 | 0 | 0.00 | 0.56 |
| 136 | 0.0048 | 0 | 0.00 | 0.55 |
| 137 | 0.0019 | 0 | 0.00 | 0.55 |
| 138 | 0.0045 | 0 | 0.00 | 0.55 |
| 139 | 0.0019 | 0 | 0.00 | 0.54 |
| 140 | 0.0023 | 0 | 0.00 | 0.54 |
| 141 | 0.0035 | 0 | 0.00 | 0.54 |
| 142 | 0.0023 | 0 | 0.00 | 0.54 |
| 143 | 0.0032 | 0 | 0.00 | 0.53 |
| 144 | 0.0035 | 0 | 0.00 | 0.53 |
| 145 | 0.0029 | 0 | 0.00 | 0.53 |
| 146 | 0.0029 | 0 | 0.00 | 0.52 |
| 147 | 0.0045 | 0 | 0.00 | 0.52 |
| 148 | 0.0039 | 0 | 0.00 | 0.52 |
| 149 | 0.0026 | 0 | 0.00 | 0.51 |
| 150 | 0.0029 | 0 | 0.00 | 0.51 |
| 151 | 0.0026 | 0 | 0.00 | 0.51 |
| 152 | 0.0013 | 0 | 0.00 | 0.50 |
| 153 | 0.0019 | 0 | 0.00 | 0.50 |
| 154 | 0.0023 | 0 | 0.00 | 0.50 |
| 155 | 0.0016 | 0 | 0.00 | 0.50 |
| 156 | 0.0032 | 0 | 0.00 | 0.50 |
| 157 | 0.0019 | 0 | 0.00 | 0.49 |
| 158 | 0.0035 | 0 | 0.00 | 0.49 |
| 159 | 0.0023 | 0 | 0.00 | 0.49 |
| 160 | 0.0026 | 0 | 0.00 | 0.49 |
| 161 | 0.0029 | 0 | 0.00 | 0.48 |
| 162 | 0.0029 | 0 | 0.00 | 0.48 |
| 163 | 0.0019 | 0 | 0.00 | 0.48 |
| 164 | 0.0010 | 0 | 0.00 | 0.48 |
| 165 | 0.0045 | 0 | 0.00 | 0.48 |
| 166 | 0.0029 | 0 | 0.00 | 0.47 |
| 167 | 0.0029 | 0 | 0.00 | 0.47 |
| 168 | 0.0023 | 0 | 0.00 | 0.47 |
| 169 | 0.0016 | 0 | 0.00 | 0.46 |
| 170 | 0.0019 | 0 | 0.00 | 0.46 |
| 171 | 0.0006 | 0 | 0.00 | 0.46 |
| 172 | 0.0016 | 0 | 0.00 | 0.46 |
| 173 | 0.0006 | 0 | 0.00 | 0.46 |
| 174 | 0.0016 | 0 | 0.00 | 0.46 |
| 175 | 0.0026 | 0 | 0.00 | 0.45 |
| 176 | 0.0016 | 0 | 0.00 | 0.45 |

| | | | | |
|---|---|---|---|---|
| 177 | 0.0023 | 0 | 0.00 | 0.45 |
| 178 | 0.0029 | 0 | 0.00 | 0.45 |
| 179 | 0.0010 | 0 | 0.00 | 0.45 |
| 180 | 0.0032 | 0 | 0.00 | 0.44 |
| 181 | 0.0026 | 0 | 0.00 | 0.44 |
| 182 | 0.0010 | 0 | 0.00 | 0.44 |
| 183 | 0.0006 | 0 | 0.00 | 0.44 |
| 184 | 0.0026 | 0 | 0.00 | 0.44 |
| 185 | 0.0010 | 0 | 0.00 | 0.43 |
| 186 | 0.0023 | 0 | 0.00 | 0.43 |
| 187 | 0.0010 | 0 | 0.00 | 0.43 |
| 188 | 0.0019 | 0 | 0.00 | 0.43 |
| 189 | 0.0019 | 0 | 0.00 | 0.43 |
| 190 | 0.0035 | 0 | 0.00 | 0.43 |
| 191 | 0.0003 | 0 | 0.00 | 0.42 |
| 192 | 0.0026 | 0 | 0.00 | 0.42 |
| 193 | 0.0003 | 0 | 0.00 | 0.42 |
| 194 | 0.0029 | 0 | 0.00 | 0.42 |
| 195 | 0.0026 | 0 | 0.00 | 0.42 |
| 196 | 0.0006 | 0 | 0.00 | 0.41 |
| 197 | 0.0006 | 0 | 0.00 | 0.41 |
| 198 | 0.0010 | 0 | 0.00 | 0.41 |
| 199 | 0.0016 | 0 | 0.00 | 0.41 |
| 200 | 0.0016 | 0 | 0.00 | 0.41 |
| 201 | 0.0035 | 0 | 0.00 | 0.41 |
| 202 | 0.0010 | 0 | 0.00 | 0.41 |
| 203 | 0.0010 | 0 | 0.00 | 0.40 |
| 204 | 0.0010 | 0 | 0.00 | 0.40 |
| 205 | 0.0023 | 0 | 0.00 | 0.40 |
| 206 | 0.0035 | 0 | 0.00 | 0.40 |
| 207 | 0.0013 | 0 | 0.00 | 0.40 |
| 208 | 0.0010 | 0 | 0.00 | 0.40 |
| 209 | 0.0016 | 0 | 0.00 | 0.39 |
| 210 | 0.0013 | 0 | 0.00 | 0.39 |
| 211 | 0.0006 | 0 | 0.00 | 0.39 |
| 212 | 0.0010 | 0 | 0.00 | 0.39 |
| 213 | 0.0010 | 0 | 0.00 | 0.39 |
| 214 | 0.0006 | 0 | 0.00 | 0.39 |
| 215 | 0.0019 | 0 | 0.00 | 0.39 |
| 217 | 0.0010 | 0 | 0.00 | 0.39 |
| 218 | 0.0013 | 0 | 0.00 | 0.39 |
| 219 | 0.0023 | 0 | 0.00 | 0.38 |
| 220 | 0.0010 | 0 | 0.00 | 0.38 |
| 221 | 0.0010 | 0 | 0.00 | 0.38 |

| | | | | |
|---|---|---|---|---|
| 223 | 0.0010 | 0 | 0.00 | 0.38 |
| 224 | 0.0006 | 0 | 0.00 | 0.38 |
| 225 | 0.0013 | 0 | 0.00 | 0.38 |
| 226 | 0.0006 | 0 | 0.00 | 0.38 |
| 227 | 0.0010 | 0 | 0.00 | 0.38 |
| 228 | 0.0013 | 0 | 0.00 | 0.38 |
| 229 | 0.0016 | 0 | 0.00 | 0.37 |
| 230 | 0.0026 | 0 | 0.00 | 0.37 |
| 231 | 0.0019 | 0 | 0.00 | 0.37 |
| 232 | 0.0016 | 0 | 0.00 | 0.37 |
| 233 | 0.0006 | 0 | 0.00 | 0.37 |
| 234 | 0.0013 | 0 | 0.00 | 0.37 |
| 235 | 0.0016 | 0 | 0.00 | 0.36 |
| 236 | 0.0029 | 0 | 0.00 | 0.36 |
| 237 | 0.0010 | 0 | 0.00 | 0.36 |
| 238 | 0.0029 | 0 | 0.00 | 0.36 |
| 239 | 0.0010 | 0 | 0.00 | 0.36 |
| 240 | 0.0010 | 0 | 0.00 | 0.36 |
| 241 | 0.0006 | 0 | 0.00 | 0.35 |
| 242 | 0.0016 | 0 | 0.00 | 0.35 |
| 244 | 0.0006 | 0 | 0.00 | 0.35 |
| 245 | 0.0006 | 0 | 0.00 | 0.35 |
| 246 | 0.0010 | 0 | 0.00 | 0.35 |
| 247 | 0.0013 | 0 | 0.00 | 0.35 |
| 248 | 0.0013 | 0 | 0.00 | 0.35 |
| 249 | 0.0003 | 0 | 0.00 | 0.35 |
| 250 | 0.0006 | 0 | 0.00 | 0.35 |
| 251 | 0.0016 | 0 | 0.00 | 0.35 |
| 252 | 0.0013 | 0 | 0.00 | 0.34 |
| 253 | 0.0003 | 0 | 0.00 | 0.34 |
| 254 | 0.0026 | 0 | 0.00 | 0.34 |
| 255 | 0.0010 | 0 | 0.00 | 0.34 |
| 256 | 0.0013 | 0 | 0.00 | 0.34 |
| 257 | 0.0003 | 0 | 0.00 | 0.34 |
| 258 | 0.0010 | 0 | 0.00 | 0.34 |
| 259 | 0.0006 | 0 | 0.00 | 0.34 |
| 260 | 0.0010 | 0 | 0.00 | 0.34 |
| 261 | 0.0010 | 0 | 0.00 | 0.34 |
| 262 | 0.0013 | 0 | 0.00 | 0.33 |
| 263 | 0.0013 | 0 | 0.00 | 0.33 |
| 264 | 0.0013 | 0 | 0.00 | 0.33 |
| 265 | 0.0013 | 0 | 0.00 | 0.33 |
| 266 | 0.0006 | 0 | 0.00 | 0.33 |
| 267 | 0.0003 | 0 | 0.00 | 0.33 |

| | | | | |
|---|---|---|---|---|
| 268 | 0.0006 | 0 | 0.00 | 0.33 |
| 269 | 0.0013 | 0 | 0.00 | 0.33 |
| 270 | 0.0013 | 0 | 0.00 | 0.33 |
| 271 | 0.0010 | 0 | 0.00 | 0.32 |
| 272 | 0.0010 | 0 | 0.00 | 0.32 |
| 273 | 0.0013 | 0 | 0.00 | 0.32 |
| 274 | 0.0013 | 0 | 0.00 | 0.32 |
| 275 | 0.0019 | 0 | 0.00 | 0.32 |
| 278 | 0.0006 | 0 | 0.00 | 0.32 |
| 279 | 0.0019 | 0 | 0.00 | 0.32 |
| 280 | 0.0013 | 0 | 0.00 | 0.32 |
| 281 | 0.0003 | 0 | 0.00 | 0.31 |
| 282 | 0.0010 | 0 | 0.00 | 0.31 |
| 283 | 0.0003 | 0 | 0.00 | 0.31 |
| 284 | 0.0006 | 0 | 0.00 | 0.31 |
| 285 | 0.0013 | 0 | 0.00 | 0.31 |
| 286 | 0.0003 | 0 | 0.00 | 0.31 |
| 287 | 0.0006 | 0 | 0.00 | 0.31 |
| 288 | 0.0003 | 0 | 0.00 | 0.31 |
| 289 | 0.0013 | 0 | 0.00 | 0.31 |
| 290 | 0.0016 | 0 | 0.00 | 0.31 |
| 291 | 0.0003 | 0 | 0.00 | 0.31 |
| 292 | 0.0010 | 0 | 0.00 | 0.31 |
| 293 | 0.0010 | 0 | 0.00 | 0.31 |
| 294 | 0.0019 | 0 | 0.00 | 0.30 |
| 295 | 0.0010 | 0 | 0.00 | 0.30 |
| 296 | 0.0003 | 0 | 0.00 | 0.30 |
| 297 | 0.0010 | 0 | 0.00 | 0.30 |
| 298 | 0.0006 | 0 | 0.00 | 0.30 |
| 299 | 0.0010 | 0 | 0.00 | 0.30 |
| 300 | 0.0003 | 0 | 0.00 | 0.30 |
| 301 | 0.0019 | 0 | 0.00 | 0.30 |
| 302 | 0.0006 | 0 | 0.00 | 0.30 |
| 303 | 0.0003 | 0 | 0.00 | 0.30 |
| 305 | 0.0003 | 0 | 0.00 | 0.30 |
| 306 | 0.0013 | 0 | 0.00 | 0.30 |
| 307 | 0.0010 | 0 | 0.00 | 0.29 |
| 308 | 0.0010 | 0 | 0.00 | 0.29 |
| 309 | 0.0010 | 0 | 0.00 | 0.29 |
| 310 | 0.0003 | 0 | 0.00 | 0.29 |
| 311 | 0.0006 | 0 | 0.00 | 0.29 |
| 312 | 0.0006 | 0 | 0.00 | 0.29 |
| 313 | 0.0006 | 0 | 0.00 | 0.29 |
| 314 | 0.0006 | 0 | 0.00 | 0.29 |

| | | | | |
|---|---|---|---|---|
| 315 | 0.0003 | 0 | 0.00 | 0.29 |
| 316 | 0.0016 | 0 | 0.00 | 0.29 |
| 317 | 0.0010 | 0 | 0.00 | 0.29 |
| 319 | 0.0013 | 0 | 0.00 | 0.29 |
| 320 | 0.0003 | 0 | 0.00 | 0.28 |
| 321 | 0.0006 | 0 | 0.00 | 0.28 |
| 323 | 0.0006 | 0 | 0.00 | 0.28 |
| 324 | 0.0006 | 0 | 0.00 | 0.28 |
| 325 | 0.0006 | 0 | 0.00 | 0.28 |
| 326 | 0.0003 | 0 | 0.00 | 0.28 |
| 328 | 0.0013 | 0 | 0.00 | 0.28 |
| 329 | 0.0010 | 0 | 0.00 | 0.28 |
| 330 | 0.0013 | 0 | 0.00 | 0.28 |
| 332 | 0.0013 | 0 | 0.00 | 0.28 |
| 333 | 0.0006 | 0 | 0.00 | 0.28 |
| 335 | 0.0010 | 0 | 0.00 | 0.28 |
| 336 | 0.0016 | 0 | 0.00 | 0.27 |
| 337 | 0.0003 | 0 | 0.00 | 0.27 |
| 338 | 0.0032 | 0 | 0.00 | 0.27 |
| 339 | 0.0019 | 0 | 0.00 | 0.27 |
| 340 | 0.0010 | 0 | 0.00 | 0.27 |
| 341 | 0.0003 | 0 | 0.00 | 0.27 |
| 342 | 0.0003 | 0 | 0.00 | 0.27 |
| 343 | 0.0003 | 0 | 0.00 | 0.27 |
| 344 | 0.0010 | 0 | 0.00 | 0.27 |
| 345 | 0.0003 | 0 | 0.00 | 0.26 |
| 346 | 0.0010 | 0 | 0.00 | 0.26 |
| 347 | 0.0003 | 0 | 0.00 | 0.26 |
| 348 | 0.0006 | 0 | 0.00 | 0.26 |
| 349 | 0.0003 | 0 | 0.00 | 0.26 |
| 350 | 0.0016 | 0 | 0.00 | 0.26 |
| 351 | 0.0013 | 0 | 0.00 | 0.26 |
| 353 | 0.0006 | 0 | 0.00 | 0.26 |
| 354 | 0.0013 | 0 | 0.00 | 0.26 |
| 355 | 0.0003 | 0 | 0.00 | 0.26 |
| 356 | 0.0006 | 0 | 0.00 | 0.26 |
| 357 | 0.0003 | 0 | 0.00 | 0.26 |
| 358 | 0.0013 | 0 | 0.00 | 0.26 |
| 359 | 0.0003 | 0 | 0.00 | 0.25 |
| 360 | 0.0003 | 0 | 0.00 | 0.25 |
| 361 | 0.0003 | 0 | 0.00 | 0.25 |
| 362 | 0.0010 | 0 | 0.00 | 0.25 |
| 363 | 0.0013 | 0 | 0.00 | 0.25 |
| 364 | 0.0006 | 0 | 0.00 | 0.25 |

| | | | | |
|---|---|---|---|---|
| 365 | 0.0003 | 0 | 0.00 | 0.25 |
| 366 | 0.0003 | 0 | 0.00 | 0.25 |
| 367 | 0.0003 | 0 | 0.00 | 0.25 |
| 368 | 0.0006 | 0 | 0.00 | 0.25 |
| 369 | 0.0010 | 0 | 0.00 | 0.25 |
| 370 | 0.0003 | 0 | 0.00 | 0.25 |
| 371 | 0.0013 | 0 | 0.00 | 0.25 |
| 372 | 0.0010 | 0 | 0.00 | 0.25 |
| 374 | 0.0023 | 0 | 0.00 | 0.25 |
| 375 | 0.0006 | 0 | 0.00 | 0.24 |
| 376 | 0.0003 | 0 | 0.00 | 0.24 |
| 378 | 0.0006 | 0 | 0.00 | 0.24 |
| 379 | 0.0003 | 0 | 0.00 | 0.24 |
| 380 | 0.0003 | 0 | 0.00 | 0.24 |
| 381 | 0.0006 | 0 | 0.00 | 0.24 |
| 383 | 0.0003 | 0 | 0.00 | 0.24 |
| 384 | 0.0006 | 0 | 0.00 | 0.24 |
| 385 | 0.0006 | 0 | 0.00 | 0.24 |
| 386 | 0.0003 | 0 | 0.00 | 0.24 |
| 387 | 0.0003 | 0 | 0.00 | 0.24 |
| 389 | 0.0006 | 0 | 0.00 | 0.24 |
| 391 | 0.0003 | 0 | 0.00 | 0.24 |
| 392 | 0.0003 | 0 | 0.00 | 0.24 |
| 393 | 0.0010 | 0 | 0.00 | 0.24 |
| 394 | 0.0006 | 0 | 0.00 | 0.24 |
| 395 | 0.0006 | 0 | 0.00 | 0.24 |
| 397 | 0.0003 | 0 | 0.00 | 0.23 |
| 398 | 0.0003 | 0 | 0.00 | 0.23 |
| 400 | 0.0013 | 0 | 0.00 | 0.23 |
| 401 | 0.0010 | 0 | 0.00 | 0.23 |
| 402 | 0.0013 | 0 | 0.00 | 0.23 |
| 403 | 0.0006 | 0 | 0.00 | 0.23 |
| 404 | 0.0006 | 0 | 0.00 | 0.23 |
| 405 | 0.0006 | 0 | 0.00 | 0.23 |
| 406 | 0.0006 | 0 | 0.00 | 0.23 |
| 408 | 0.0006 | 0 | 0.00 | 0.23 |
| 409 | 0.0006 | 0 | 0.00 | 0.23 |
| 410 | 0.0013 | 0 | 0.00 | 0.23 |
| 412 | 0.0006 | 0 | 0.00 | 0.23 |
| 413 | 0.0006 | 0 | 0.00 | 0.22 |
| 414 | 0.0003 | 0 | 0.00 | 0.22 |
| 415 | 0.0003 | 0 | 0.00 | 0.22 |
| 416 | 0.0003 | 0 | 0.00 | 0.22 |
| 417 | 0.0013 | 0 | 0.00 | 0.22 |

| | | | | |
|---|---|---|---|---|
| 418 | 0.0003 | 0 | 0.00 | 0.22 |
| 420 | 0.0010 | 0 | 0.00 | 0.22 |
| 423 | 0.0003 | 0 | 0.00 | 0.22 |
| 424 | 0.0006 | 0 | 0.00 | 0.22 |
| 425 | 0.0006 | 0 | 0.00 | 0.22 |
| 426 | 0.0003 | 0 | 0.00 | 0.22 |
| 427 | 0.0010 | 0 | 0.00 | 0.22 |
| 430 | 0.0003 | 0 | 0.00 | 0.22 |
| 432 | 0.0003 | 0 | 0.00 | 0.22 |
| 434 | 0.0003 | 0 | 0.00 | 0.22 |
| 436 | 0.0003 | 0 | 0.00 | 0.22 |
| 437 | 0.0006 | 0 | 0.00 | 0.22 |
| 438 | 0.0006 | 0 | 0.00 | 0.22 |
| 439 | 0.0006 | 0 | 0.00 | 0.22 |
| 441 | 0.0003 | 0 | 0.00 | 0.21 |
| 442 | 0.0010 | 0 | 0.00 | 0.21 |
| 444 | 0.0003 | 0 | 0.00 | 0.21 |
| 445 | 0.0006 | 0 | 0.00 | 0.21 |
| 447 | 0.0003 | 0 | 0.00 | 0.21 |
| 449 | 0.0010 | 0 | 0.00 | 0.21 |
| 450 | 0.0003 | 0 | 0.00 | 0.21 |
| 451 | 0.0003 | 0 | 0.00 | 0.21 |
| 454 | 0.0003 | 0 | 0.00 | 0.21 |
| 455 | 0.0003 | 0 | 0.00 | 0.21 |
| 456 | 0.0003 | 0 | 0.00 | 0.21 |
| 457 | 0.0003 | 0 | 0.00 | 0.21 |
| 459 | 0.0003 | 0 | 0.00 | 0.21 |
| 462 | 0.0003 | 0 | 0.00 | 0.21 |
| 463 | 0.0006 | 0 | 0.00 | 0.21 |
| 464 | 0.0006 | 0 | 0.00 | 0.21 |
| 465 | 0.0003 | 0 | 0.00 | 0.21 |
| 467 | 0.0003 | 0 | 0.00 | 0.21 |
| 471 | 0.0003 | 0 | 0.00 | 0.21 |
| 472 | 0.0006 | 0 | 0.00 | 0.21 |
| 473 | 0.0006 | 0 | 0.00 | 0.21 |
| 474 | 0.0006 | 0 | 0.00 | 0.20 |
| 475 | 0.0006 | 0 | 0.00 | 0.20 |
| 476 | 0.0003 | 0 | 0.00 | 0.20 |
| 477 | 0.0006 | 0 | 0.00 | 0.20 |
| 478 | 0.0003 | 0 | 0.00 | 0.20 |
| 480 | 0.0003 | 0 | 0.00 | 0.20 |
| 482 | 0.0006 | 0 | 0.00 | 0.20 |
| 484 | 0.0003 | 0 | 0.00 | 0.20 |
| 485 | 0.0003 | 0 | 0.00 | 0.20 |

| | | | | |
|---|---|---|---|---|
| 486 | 0.0006 | 0 | 0.00 | 0.20 |
| 487 | 0.0003 | 0 | 0.00 | 0.20 |
| 488 | 0.0006 | 0 | 0.00 | 0.20 |
| 489 | 0.0003 | 0 | 0.00 | 0.20 |
| 493 | 0.0003 | 0 | 0.00 | 0.20 |
| 494 | 0.0003 | 0 | 0.00 | 0.20 |
| 499 | 0.0006 | 0 | 0.00 | 0.20 |
| 501 | 0.0006 | 0 | 0.00 | 0.20 |
| 503 | 0.0006 | 0 | 0.00 | 0.20 |
| 504 | 0.0003 | 0 | 0.00 | 0.20 |
| 506 | 0.0003 | 0 | 0.00 | 0.20 |
| 507 | 0.0003 | 0 | 0.00 | 0.20 |
| 508 | 0.0003 | 0 | 0.00 | 0.20 |
| 510 | 0.0003 | 0 | 0.00 | 0.19 |
| 511 | 0.0003 | 0 | 0.00 | 0.19 |
| 512 | 0.0010 | 0 | 0.00 | 0.19 |
| 513 | 0.0003 | 0 | 0.00 | 0.19 |
| 515 | 0.0003 | 0 | 0.00 | 0.19 |
| 517 | 0.0003 | 0 | 0.00 | 0.19 |
| 520 | 0.0003 | 0 | 0.00 | 0.19 |
| 521 | 0.0003 | 0 | 0.00 | 0.19 |
| 522 | 0.0003 | 0 | 0.00 | 0.19 |
| 523 | 0.0010 | 0 | 0.00 | 0.19 |
| 525 | 0.0003 | 0 | 0.00 | 0.19 |
| 526 | 0.0003 | 0 | 0.00 | 0.19 |
| 527 | 0.0003 | 0 | 0.00 | 0.19 |
| 530 | 0.0006 | 0 | 0.00 | 0.19 |
| 531 | 0.0006 | 0 | 0.00 | 0.19 |
| 532 | 0.0006 | 0 | 0.00 | 0.19 |
| 535 | 0.0003 | 0 | 0.00 | 0.19 |
| 536 | 0.0006 | 0 | 0.00 | 0.19 |
| 537 | 0.0003 | 0 | 0.00 | 0.19 |
| 539 | 0.0010 | 0 | 0.00 | 0.19 |
| 540 | 0.0006 | 0 | 0.00 | 0.19 |
| 541 | 0.0006 | 0 | 0.00 | 0.18 |
| 542 | 0.0003 | 0 | 0.00 | 0.18 |
| 543 | 0.0003 | 0 | 0.00 | 0.18 |
| 544 | 0.0003 | 0 | 0.00 | 0.18 |
| 545 | 0.0003 | 0 | 0.00 | 0.18 |
| 547 | 0.0006 | 0 | 0.00 | 0.18 |
| 548 | 0.0006 | 0 | 0.00 | 0.18 |
| 549 | 0.0010 | 0 | 0.00 | 0.18 |
| 552 | 0.0003 | 0 | 0.00 | 0.18 |
| 555 | 0.0010 | 0 | 0.00 | 0.18 |

| | | | | |
|---|---|---|---|---|
| 556 | 0.0003 | 0 | 0.00 | 0.18 |
| 562 | 0.0003 | 0 | 0.00 | 0.18 |
| 563 | 0.0003 | 0 | 0.00 | 0.18 |
| 564 | 0.0003 | 0 | 0.00 | 0.18 |
| 566 | 0.0003 | 0 | 0.00 | 0.18 |
| 571 | 0.0010 | 0 | 0.00 | 0.18 |
| 575 | 0.0013 | 0 | 0.00 | 0.18 |
| 579 | 0.0003 | 0 | 0.00 | 0.18 |
| 580 | 0.0003 | 0 | 0.00 | 0.17 |
| 581 | 0.0006 | 0 | 0.00 | 0.17 |
| 582 | 0.0003 | 0 | 0.00 | 0.17 |
| 585 | 0.0006 | 0 | 0.00 | 0.17 |
| 586 | 0.0003 | 0 | 0.00 | 0.17 |
| 590 | 0.0010 | 0 | 0.00 | 0.17 |
| 593 | 0.0003 | 0 | 0.00 | 0.17 |
| 595 | 0.0010 | 0 | 0.00 | 0.17 |
| 596 | 0.0003 | 0 | 0.00 | 0.17 |
| 597 | 0.0003 | 0 | 0.00 | 0.17 |
| 598 | 0.0003 | 0 | 0.00 | 0.17 |
| 601 | 0.0013 | 0 | 0.00 | 0.17 |
| 602 | 0.0003 | 0 | 0.00 | 0.17 |
| 603 | 0.0006 | 0 | 0.00 | 0.17 |
| 605 | 0.0003 | 0 | 0.00 | 0.17 |
| 609 | 0.0003 | 0 | 0.00 | 0.17 |
| 610 | 0.0013 | 0 | 0.00 | 0.17 |
| 613 | 0.0003 | 0 | 0.00 | 0.17 |
| 615 | 0.0003 | 0 | 0.00 | 0.16 |
| 616 | 0.0003 | 0 | 0.00 | 0.16 |
| 618 | 0.0003 | 0 | 0.00 | 0.16 |
| 623 | 0.0003 | 0 | 0.00 | 0.16 |
| 624 | 0.0003 | 0 | 0.00 | 0.16 |
| 625 | 0.0003 | 0 | 0.00 | 0.16 |
| 627 | 0.0003 | 0 | 0.00 | 0.16 |
| 628 | 0.0003 | 0 | 0.00 | 0.16 |
| 629 | 0.0003 | 0 | 0.00 | 0.16 |
| 631 | 0.0006 | 0 | 0.00 | 0.16 |
| 632 | 0.0003 | 0 | 0.00 | 0.16 |
| 634 | 0.0003 | 0 | 0.00 | 0.16 |
| 636 | 0.0003 | 0 | 0.00 | 0.16 |
| 637 | 0.0010 | 0 | 0.00 | 0.16 |
| 640 | 0.0003 | 0 | 0.00 | 0.16 |
| 644 | 0.0006 | 0 | 0.00 | 0.16 |
| 646 | 0.0003 | 0 | 0.00 | 0.16 |
| 647 | 0.0003 | 0 | 0.00 | 0.16 |

| | | | | |
|---|---|---|---|---|
| 649 | 0.0006 | 0 | 0.00 | 0.16 |
| 653 | 0.0006 | 0 | 0.00 | 0.16 |
| 654 | 0.0006 | 0 | 0.00 | 0.16 |
| 655 | 0.0003 | 0 | 0.00 | 0.16 |
| 656 | 0.0003 | 0 | 0.00 | 0.16 |
| 657 | 0.0006 | 0 | 0.00 | 0.16 |
| 659 | 0.0003 | 0 | 0.00 | 0.15 |
| 661 | 0.0003 | 0 | 0.00 | 0.15 |
| 662 | 0.0003 | 0 | 0.00 | 0.15 |
| 664 | 0.0006 | 0 | 0.00 | 0.15 |
| 666 | 0.0003 | 0 | 0.00 | 0.15 |
| 667 | 0.0010 | 0 | 0.00 | 0.15 |
| 670 | 0.0006 | 0 | 0.00 | 0.15 |
| 671 | 0.0003 | 0 | 0.00 | 0.15 |
| 677 | 0.0003 | 0 | 0.00 | 0.15 |
| 678 | 0.0003 | 0 | 0.00 | 0.15 |
| 679 | 0.0006 | 0 | 0.00 | 0.15 |
| 682 | 0.0003 | 0 | 0.00 | 0.15 |
| 683 | 0.0003 | 0 | 0.00 | 0.15 |
| 689 | 0.0003 | 0 | 0.00 | 0.15 |
| 690 | 0.0013 | 0 | 0.00 | 0.15 |
| 691 | 0.0006 | 0 | 0.00 | 0.15 |
| 693 | 0.0003 | 0 | 0.00 | 0.15 |
| 694 | 0.0003 | 0 | 0.00 | 0.15 |
| 699 | 0.0003 | 0 | 0.00 | 0.15 |
| 704 | 0.0003 | 0 | 0.00 | 0.15 |
| 705 | 0.0003 | 0 | 0.00 | 0.15 |
| 706 | 0.0003 | 0 | 0.00 | 0.14 |
| 708 | 0.0003 | 0 | 0.00 | 0.14 |
| 712 | 0.0003 | 0 | 0.00 | 0.14 |
| 715 | 0.0003 | 0 | 0.00 | 0.14 |
| 719 | 0.0006 | 0 | 0.00 | 0.14 |
| 720 | 0.0003 | 0 | 0.00 | 0.14 |
| 725 | 0.0003 | 0 | 0.00 | 0.14 |
| 727 | 0.0003 | 0 | 0.00 | 0.14 |
| 731 | 0.0003 | 0 | 0.00 | 0.14 |
| 734 | 0.0006 | 0 | 0.00 | 0.14 |
| 741 | 0.0003 | 0 | 0.00 | 0.14 |
| 745 | 0.0006 | 0 | 0.00 | 0.14 |
| 746 | 0.0003 | 0 | 0.00 | 0.14 |
| 749 | 0.0003 | 0 | 0.00 | 0.14 |
| 750 | 0.0006 | 0 | 0.00 | 0.14 |
| 752 | 0.0006 | 0 | 0.00 | 0.14 |
| 754 | 0.0003 | 0 | 0.00 | 0.14 |

| | | | | |
|---|---|---|---|---|
| 755 | 0.0003 | 0 | 0.00 | 0.14 |
| 759 | 0.0003 | 0 | 0.00 | 0.14 |
| 760 | 0.0006 | 0 | 0.00 | 0.14 |
| 761 | 0.0003 | 0 | 0.00 | 0.14 |
| 762 | 0.0003 | 0 | 0.00 | 0.14 |
| 772 | 0.0003 | 0 | 0.00 | 0.14 |
| 773 | 0.0003 | 0 | 0.00 | 0.14 |
| 774 | 0.0003 | 0 | 0.00 | 0.14 |
| 776 | 0.0003 | 0 | 0.00 | 0.14 |
| 777 | 0.0003 | 0 | 0.00 | 0.13 |
| 778 | 0.0003 | 0 | 0.00 | 0.13 |
| 780 | 0.0003 | 0 | 0.00 | 0.13 |
| 782 | 0.0003 | 0 | 0.00 | 0.13 |
| 783 | 0.0003 | 0 | 0.00 | 0.13 |
| 785 | 0.0003 | 0 | 0.00 | 0.13 |
| 788 | 0.0003 | 0 | 0.00 | 0.13 |
| 789 | 0.0003 | 0 | 0.00 | 0.13 |
| 790 | 0.0003 | 0 | 0.00 | 0.13 |
| 791 | 0.0003 | 0 | 0.00 | 0.13 |
| 795 | 0.0003 | 0 | 0.00 | 0.13 |
| 796 | 0.0006 | 0 | 0.00 | 0.13 |
| 797 | 0.0003 | 0 | 0.00 | 0.13 |
| 799 | 0.0003 | 0 | 0.00 | 0.13 |
| 809 | 0.0003 | 0 | 0.00 | 0.13 |
| 811 | 0.0003 | 0 | 0.00 | 0.13 |
| 812 | 0.0003 | 0 | 0.00 | 0.13 |
| 816 | 0.0003 | 0 | 0.00 | 0.13 |
| 817 | 0.0003 | 0 | 0.00 | 0.13 |
| 819 | 0.0003 | 0 | 0.00 | 0.13 |
| 821 | 0.0003 | 0 | 0.00 | 0.13 |
| 823 | 0.0003 | 0 | 0.00 | 0.13 |
| 827 | 0.0003 | 0 | 0.00 | 0.13 |
| 828 | 0.0006 | 0 | 0.00 | 0.13 |
| 830 | 0.0003 | 0 | 0.00 | 0.13 |
| 831 | 0.0003 | 0 | 0.00 | 0.13 |
| 836 | 0.0003 | 0 | 0.00 | 0.13 |
| 837 | 0.0003 | 0 | 0.00 | 0.13 |
| 840 | 0.0003 | 0 | 0.00 | 0.13 |
| 846 | 0.0003 | 0 | 0.00 | 0.12 |
| 850 | 0.0003 | 0 | 0.00 | 0.12 |
| 854 | 0.0003 | 0 | 0.00 | 0.12 |
| 857 | 0.0003 | 0 | 0.00 | 0.12 |
| 860 | 0.0003 | 0 | 0.00 | 0.12 |
| 861 | 0.0006 | 0 | 0.00 | 0.12 |

| | | | | |
|---|---|---|---|---|
| 864 | 0.0003 | 0 | 0.00 | 0.12 |
| 868 | 0.0003 | 0 | 0.00 | 0.12 |
| 871 | 0.0003 | 0 | 0.00 | 0.12 |
| 872 | 0.0003 | 0 | 0.00 | 0.12 |
| 873 | 0.0006 | 0 | 0.00 | 0.12 |
| 877 | 0.0003 | 0 | 0.00 | 0.12 |
| 879 | 0.0003 | 0 | 0.00 | 0.12 |
| 883 | 0.0003 | 0 | 0.00 | 0.12 |
| 884 | 0.0003 | 0 | 0.00 | 0.12 |
| 894 | 0.0003 | 0 | 0.00 | 0.12 |
| 900 | 0.0003 | 0 | 0.00 | 0.12 |
| 902 | 0.0006 | 0 | 0.00 | 0.12 |
| 904 | 0.0003 | 0 | 0.00 | 0.12 |
| 906 | 0.0003 | 0 | 0.00 | 0.12 |
| 917 | 0.0006 | 0 | 0.00 | 0.12 |
| 925 | 0.0006 | 0 | 0.00 | 0.12 |
| 926 | 0.0003 | 0 | 0.00 | 0.12 |
| 929 | 0.0003 | 0 | 0.00 | 0.12 |
| 932 | 0.0003 | 0 | 0.00 | 0.12 |
| 935 | 0.0003 | 0 | 0.00 | 0.12 |
| 945 | 0.0003 | 0 | 0.00 | 0.11 |
| 950 | 0.0010 | 0 | 0.00 | 0.11 |
| 951 | 0.0003 | 0 | 0.00 | 0.11 |
| 953 | 0.0003 | 0 | 0.00 | 0.11 |
| 955 | 0.0003 | 0 | 0.00 | 0.11 |
| 956 | 0.0003 | 0 | 0.00 | 0.11 |
| 957 | 0.0006 | 0 | 0.00 | 0.11 |
| 960 | 0.0003 | 0 | 0.00 | 0.11 |
| 961 | 0.0003 | 0 | 0.00 | 0.11 |
| 964 | 0.0003 | 0 | 0.00 | 0.11 |
| 965 | 0.0006 | 0 | 0.00 | 0.11 |
| 968 | 0.0003 | 0 | 0.00 | 0.11 |
| 972 | 0.0003 | 0 | 0.00 | 0.11 |
| 973 | 0.0003 | 0 | 0.00 | 0.11 |
| 974 | 0.0003 | 0 | 0.00 | 0.11 |
| 975 | 0.0003 | 0 | 0.00 | 0.11 |
| 985 | 0.0003 | 0 | 0.00 | 0.11 |
| 987 | 0.0003 | 0 | 0.00 | 0.11 |
| 991 | 0.0003 | 0 | 0.00 | 0.11 |
| 993 | 0.0003 | 0 | 0.00 | 0.11 |
| 995 | 0.0003 | 0 | 0.00 | 0.11 |
| 996 | 0.0003 | 0 | 0.00 | 0.11 |
| 997 | 0.0006 | 0 | 0.00 | 0.11 |
| 998 | 0.0003 | 0 | 0.00 | 0.11 |

| | | | | |
|---|---|---|---|---|
| 1000 | 0.0003 | 0 | 0.00 | 0.11 |
| 1010 | 0.0003 | 0 | 0.00 | 0.11 |
| 1013 | 0.0003 | 0 | 0.00 | 0.10 |
| 1014 | 0.0003 | 0 | 0.00 | 0.10 |
| 1021 | 0.0003 | 0 | 0.00 | 0.10 |
| 1038 | 0.0003 | 0 | 0.00 | 0.10 |
| 1039 | 0.0003 | 0 | 0.00 | 0.10 |
| 1044 | 0.0006 | 0 | 0.00 | 0.10 |
| 1047 | 0.0003 | 0 | 0.00 | 0.10 |
| 1048 | 0.0003 | 0 | 0.00 | 0.10 |
| 1051 | 0.0003 | 0 | 0.00 | 0.10 |
| 1055 | 0.0003 | 0 | 0.00 | 0.10 |
| 1057 | 0.0003 | 0 | 0.00 | 0.10 |
| 1060 | 0.0003 | 0 | 0.00 | 0.10 |
| 1065 | 0.0006 | 0 | 0.00 | 0.10 |
| 1068 | 0.0003 | 0 | 0.00 | 0.10 |
| 1069 | 0.0003 | 0 | 0.00 | 0.10 |
| 1071 | 0.0006 | 0 | 0.00 | 0.10 |
| 1073 | 0.0003 | 0 | 0.00 | 0.10 |
| 1075 | 0.0006 | 0 | 0.00 | 0.10 |
| 1077 | 0.0003 | 0 | 0.00 | 0.10 |
| 1087 | 0.0003 | 0 | 0.00 | 0.10 |
| 1093 | 0.0003 | 0 | 0.00 | 0.10 |
| 1094 | 0.0006 | 0 | 0.00 | 0.10 |
| 1098 | 0.0003 | 0 | 0.00 | 0.10 |
| 1109 | 0.0006 | 0 | 0.00 | 0.10 |
| 1116 | 0.0003 | 0 | 0.00 | 0.10 |
| 1121 | 0.0003 | 0 | 0.00 | 0.09 |
| 1131 | 0.0003 | 0 | 0.00 | 0.09 |
| 1140 | 0.0003 | 0 | 0.00 | 0.09 |
| 1144 | 0.0006 | 0 | 0.00 | 0.09 |
| 1146 | 0.0003 | 0 | 0.00 | 0.09 |
| 1150 | 0.0003 | 0 | 0.00 | 0.09 |
| 1155 | 0.0003 | 0 | 0.00 | 0.09 |
| 1165 | 0.0003 | 0 | 0.00 | 0.09 |
| 1166 | 0.0003 | 0 | 0.00 | 0.09 |
| 1176 | 0.0006 | 0 | 0.00 | 0.09 |
| 1178 | 0.0003 | 0 | 0.00 | 0.09 |
| 1179 | 0.0003 | 0 | 0.00 | 0.09 |
| 1181 | 0.0003 | 0 | 0.00 | 0.09 |
| 1190 | 0.0003 | 0 | 0.00 | 0.09 |
| 1195 | 0.0003 | 0 | 0.00 | 0.09 |
| 1199 | 0.0003 | 0 | 0.00 | 0.09 |
| 1204 | 0.0003 | 0 | 0.00 | 0.09 |

| | | | | |
|---|---|---|---|---|
| 1205 | 0.0003 | 0 | 0.00 | 0.09 |
| 1215 | 0.0003 | 0 | 0.00 | 0.09 |
| 1230 | 0.0003 | 0 | 0.00 | 0.09 |
| 1231 | 0.0003 | 0 | 0.00 | 0.09 |
| 1236 | 0.0003 | 0 | 0.00 | 0.09 |
| 1237 | 0.0003 | 0 | 0.00 | 0.09 |
| 1238 | 0.0003 | 0 | 0.00 | 0.09 |
| 1240 | 0.0003 | 0 | 0.00 | 0.09 |
| 1244 | 0.0003 | 0 | 0.00 | 0.09 |
| 1253 | 0.0006 | 0 | 0.00 | 0.09 |
| 1257 | 0.0003 | 0 | 0.00 | 0.09 |
| 1264 | 0.0003 | 0 | 0.00 | 0.08 |
| 1267 | 0.0003 | 0 | 0.00 | 0.08 |
| 1271 | 0.0003 | 0 | 0.00 | 0.08 |
| 1276 | 0.0003 | 0 | 0.00 | 0.08 |
| 1279 | 0.0003 | 0 | 0.00 | 0.08 |
| 1310 | 0.0003 | 0 | 0.00 | 0.08 |
| 1320 | 0.0003 | 0 | 0.00 | 0.08 |
| 1322 | 0.0003 | 0 | 0.00 | 0.08 |
| 1324 | 0.0003 | 0 | 0.00 | 0.08 |
| 1326 | 0.0003 | 0 | 0.00 | 0.08 |
| 1328 | 0.0003 | 0 | 0.00 | 0.08 |
| 1329 | 0.0003 | 0 | 0.00 | 0.08 |
| 1330 | 0.0003 | 0 | 0.00 | 0.08 |
| 1337 | 0.0003 | 0 | 0.00 | 0.08 |
| 1339 | 0.0003 | 0 | 0.00 | 0.08 |
| 1349 | 0.0006 | 0 | 0.00 | 0.08 |
| 1350 | 0.0003 | 0 | 0.00 | 0.08 |
| 1351 | 0.0003 | 0 | 0.00 | 0.08 |
| 1360 | 0.0003 | 0 | 0.00 | 0.08 |
| 1362 | 0.0003 | 0 | 0.00 | 0.08 |
| 1363 | 0.0006 | 0 | 0.00 | 0.08 |
| 1372 | 0.0006 | 0 | 0.00 | 0.08 |
| 1375 | 0.0003 | 0 | 0.00 | 0.08 |
| 1379 | 0.0003 | 0 | 0.00 | 0.08 |
| 1381 | 0.0003 | 0 | 0.00 | 0.08 |
| 1383 | 0.0003 | 0 | 0.00 | 0.08 |
| 1391 | 0.0003 | 0 | 0.00 | 0.08 |
| 1392 | 0.0003 | 0 | 0.00 | 0.08 |
| 1394 | 0.0006 | 0 | 0.00 | 0.07 |
| 1400 | 0.0003 | 0 | 0.00 | 0.07 |
| 1401 | 0.0003 | 0 | 0.00 | 0.07 |
| 1408 | 0.0003 | 0 | 0.00 | 0.07 |
| 1415 | 0.0003 | 0 | 0.00 | 0.07 |

| | | | | |
|---|---|---|---|---|
| 1417 | 0.0003 | 0 | 0.00 | 0.07 |
| 1422 | 0.0003 | 0 | 0.00 | 0.07 |
| 1442 | 0.0003 | 0 | 0.00 | 0.07 |
| 1443 | 0.0003 | 0 | 0.00 | 0.07 |
| 1460 | 0.0006 | 0 | 0.00 | 0.07 |
| 1467 | 0.0003 | 0 | 0.00 | 0.07 |
| 1476 | 0.0003 | 0 | 0.00 | 0.07 |
| 1485 | 0.0003 | 0 | 0.00 | 0.07 |
| 1489 | 0.0003 | 0 | 0.00 | 0.07 |
| 1493 | 0.0003 | 0 | 0.00 | 0.07 |
| 1500 | 0.0010 | 0 | 0.00 | 0.07 |
| 1513 | 0.0003 | 0 | 0.00 | 0.07 |
| 1528 | 0.0003 | 0 | 0.00 | 0.07 |
| 1537 | 0.0003 | 0 | 0.00 | 0.07 |
| 1539 | 0.0003 | 0 | 0.00 | 0.07 |
| 1540 | 0.0003 | 0 | 0.00 | 0.07 |
| 1551 | 0.0003 | 0 | 0.00 | 0.07 |
| 1569 | 0.0003 | 0 | 0.00 | 0.07 |
| 1570 | 0.0003 | 0 | 0.00 | 0.07 |
| 1575 | 0.0003 | 0 | 0.00 | 0.07 |
| 1584 | 0.0003 | 0 | 0.00 | 0.07 |
| 1585 | 0.0003 | 0 | 0.00 | 0.07 |
| 1595 | 0.0003 | 0 | 0.00 | 0.06 |
| 1600 | 0.0006 | 0 | 0.00 | 0.06 |
| 1605 | 0.0006 | 0 | 0.00 | 0.06 |
| 1627 | 0.0003 | 0 | 0.00 | 0.06 |
| 1631 | 0.0003 | 0 | 0.00 | 0.06 |
| 1637 | 0.0003 | 0 | 0.00 | 0.06 |
| 1641 | 0.0003 | 0 | 0.00 | 0.06 |
| 1651 | 0.0003 | 0 | 0.00 | 0.06 |
| 1653 | 0.0003 | 0 | 0.00 | 0.06 |
| 1660 | 0.0003 | 0 | 0.00 | 0.06 |
| 1671 | 0.0003 | 0 | 0.00 | 0.06 |
| 1676 | 0.0003 | 0 | 0.00 | 0.06 |
| 1680 | 0.0003 | 0 | 0.00 | 0.06 |
| 1685 | 0.0010 | 0 | 0.00 | 0.06 |
| 1688 | 0.0003 | 0 | 0.00 | 0.06 |
| 1693 | 0.0003 | 0 | 0.00 | 0.06 |
| 1697 | 0.0003 | 0 | 0.00 | 0.06 |
| 1709 | 0.0003 | 0 | 0.00 | 0.06 |
| 1713 | 0.0003 | 0 | 0.00 | 0.06 |
| 1718 | 0.0006 | 0 | 0.00 | 0.06 |
| 1734 | 0.0003 | 0 | 0.00 | 0.06 |
| 1735 | 0.0003 | 0 | 0.00 | 0.06 |

| | | | | |
|---|---|---|---|---|
| 1762 | 0.0003 | 0 | 0.00 | 0.06 |
| 1767 | 0.0003 | 0 | 0.00 | 0.06 |
| 1768 | 0.0003 | 0 | 0.00 | 0.06 |
| 1782 | 0.0006 | 0 | 0.00 | 0.06 |
| 1791 | 0.0003 | 0 | 0.00 | 0.05 |
| 1792 | 0.0003 | 0 | 0.00 | 0.05 |
| 1801 | 0.0003 | 0 | 0.00 | 0.05 |
| 1813 | 0.0003 | 0 | 0.00 | 0.05 |
| 1844 | 0.0003 | 0 | 0.00 | 0.05 |
| 1848 | 0.0003 | 0 | 0.00 | 0.05 |
| 1866 | 0.0003 | 0 | 0.00 | 0.05 |
| 1877 | 0.0003 | 0 | 0.00 | 0.05 |
| 1888 | 0.0003 | 0 | 0.00 | 0.05 |
| 1894 | 0.0003 | 0 | 0.00 | 0.05 |
| 1899 | 0.0003 | 0 | 0.00 | 0.05 |
| 1903 | 0.0003 | 0 | 0.00 | 0.05 |
| 1905 | 0.0003 | 0 | 0.00 | 0.05 |
| 1911 | 0.0003 | 0 | 0.00 | 0.05 |
| 1914 | 0.0003 | 0 | 0.00 | 0.05 |
| 1916 | 0.0003 | 0 | 0.00 | 0.05 |
| 1923 | 0.0003 | 0 | 0.00 | 0.05 |
| 1940 | 0.0003 | 0 | 0.00 | 0.05 |
| 1943 | 0.0003 | 0 | 0.00 | 0.05 |
| 1958 | 0.0003 | 0 | 0.00 | 0.05 |
| 1961 | 0.0003 | 0 | 0.00 | 0.05 |
| 1968 | 0.0003 | 0 | 0.00 | 0.05 |
| 2007 | 0.0003 | 0 | 0.00 | 0.05 |
| 2023 | 0.0003 | 0 | 0.00 | 0.05 |
| 2026 | 0.0003 | 0 | 0.00 | 0.05 |
| 2048 | 0.0003 | 0 | 0.00 | 0.05 |
| 2063 | 0.0003 | 0 | 0.00 | 0.05 |
| 2070 | 0.0003 | 0 | 0.00 | 0.05 |
| 2073 | 0.0003 | 0 | 0.00 | 0.05 |
| 2090 | 0.0003 | 0 | 0.00 | 0.05 |
| 2108 | 0.0003 | 0 | 0.00 | 0.05 |
| 2113 | 0.0003 | 0 | 0.00 | 0.04 |
| 2118 | 0.0003 | 0 | 0.00 | 0.04 |
| 2122 | 0.0003 | 0 | 0.00 | 0.04 |
| 2130 | 0.0003 | 0 | 0.00 | 0.04 |
| 2148 | 0.0003 | 0 | 0.00 | 0.04 |
| 2166 | 0.0003 | 0 | 0.00 | 0.04 |
| 2169 | 0.0003 | 0 | 0.00 | 0.04 |
| 2172 | 0.0003 | 0 | 0.00 | 0.04 |
| 2173 | 0.0003 | 0 | 0.00 | 0.04 |

| | | | | |
|---|---|---|---|---|
| 2175 | 0.0003 | 0 | 0.00 | 0.04 |
| 2179 | 0.0003 | 0 | 0.00 | 0.04 |
| 2189 | 0.0003 | 0 | 0.00 | 0.04 |
| 2190 | 0.0003 | 0 | 0.00 | 0.04 |
| 2223 | 0.0003 | 0 | 0.00 | 0.04 |
| 2229 | 0.0003 | 0 | 0.00 | 0.04 |
| 2244 | 0.0003 | 0 | 0.00 | 0.04 |
| 2249 | 0.0003 | 0 | 0.00 | 0.04 |
| 2253 | 0.0003 | 0 | 0.00 | 0.04 |
| 2256 | 0.0003 | 0 | 0.00 | 0.04 |
| 2261 | 0.0006 | 0 | 0.00 | 0.04 |
| 2301 | 0.0003 | 0 | 0.00 | 0.04 |
| 2303 | 0.0003 | 0 | 0.00 | 0.04 |
| 2317 | 0.0003 | 0 | 0.00 | 0.04 |
| 2331 | 0.0003 | 0 | 0.00 | 0.04 |
| 2332 | 0.0003 | 0 | 0.00 | 0.04 |
| 2351 | 0.0003 | 0 | 0.00 | 0.04 |
| 2352 | 0.0003 | 0 | 0.00 | 0.04 |
| 2354 | 0.0003 | 0 | 0.00 | 0.04 |
| 2360 | 0.0003 | 0 | 0.00 | 0.04 |
| 2367 | 0.0003 | 0 | 0.00 | 0.04 |
| 2403 | 0.0003 | 0 | 0.00 | 0.03 |
| 2424 | 0.0003 | 0 | 0.00 | 0.03 |
| 2431 | 0.0003 | 0 | 0.00 | 0.03 |
| 2435 | 0.0003 | 0 | 0.00 | 0.03 |
| 2442 | 0.0003 | 0 | 0.00 | 0.03 |
| 2446 | 0.0003 | 0 | 0.00 | 0.03 |
| 2448 | 0.0003 | 0 | 0.00 | 0.03 |
| 2472 | 0.0003 | 0 | 0.00 | 0.03 |
| 2533 | 0.0003 | 0 | 0.00 | 0.03 |
| 2569 | 0.0003 | 0 | 0.00 | 0.03 |
| 2585 | 0.0003 | 0 | 0.00 | 0.03 |
| 2588 | 0.0003 | 0 | 0.00 | 0.03 |
| 2593 | 0.0003 | 0 | 0.00 | 0.03 |
| 2594 | 0.0003 | 0 | 0.00 | 0.03 |
| 2606 | 0.0003 | 0 | 0.00 | 0.03 |
| 2613 | 0.0003 | 0 | 0.00 | 0.03 |
| 2618 | 0.0003 | 0 | 0.00 | 0.03 |
| 2631 | 0.0003 | 0 | 0.00 | 0.03 |
| 2633 | 0.0003 | 0 | 0.00 | 0.03 |
| 2664 | 0.0003 | 0 | 0.00 | 0.03 |
| 2672 | 0.0003 | 0 | 0.00 | 0.03 |
| 2686 | 0.0006 | 0 | 0.00 | 0.03 |
| 2688 | 0.0003 | 0 | 0.00 | 0.03 |

| | | | | |
|---|---|---|---|---|
| 2716 | 0.0003 | 0 | 0.00 | 0.03 |
| 2720 | 0.0003 | 0 | 0.00 | 0.03 |
| 2722 | 0.0003 | 0 | 0.00 | 0.03 |
| 2730 | 0.0003 | 0 | 0.00 | 0.03 |
| 2746 | 0.0003 | 0 | 0.00 | 0.03 |
| 2773 | 0.0003 | 0 | 0.00 | 0.03 |
| 2779 | 0.0006 | 0 | 0.00 | 0.03 |
| 2792 | 0.0003 | 0 | 0.00 | 0.02 |
| 2809 | 0.0006 | 0 | 0.00 | 0.02 |
| 2814 | 0.0003 | 0 | 0.00 | 0.02 |
| 2836 | 0.0003 | 0 | 0.00 | 0.02 |
| 2878 | 0.0003 | 0 | 0.00 | 0.02 |
| 2908 | 0.0003 | 0 | 0.00 | 0.02 |
| 2971 | 0.0003 | 0 | 0.00 | 0.02 |
| 2990 | 0.0003 | 0 | 0.00 | 0.02 |
| 3057 | 0.0003 | 0 | 0.00 | 0.02 |
| 3064 | 0.0003 | 0 | 0.00 | 0.02 |
| 3113 | 0.0003 | 0 | 0.00 | 0.02 |
| 3123 | 0.0003 | 0 | 0.00 | 0.02 |
| 3151 | 0.0003 | 0 | 0.00 | 0.02 |
| 3153 | 0.0003 | 0 | 0.00 | 0.02 |
| 3165 | 0.0003 | 0 | 0.00 | 0.02 |
| 3238 | 0.0003 | 0 | 0.00 | 0.02 |
| 3247 | 0.0003 | 0 | 0.00 | 0.02 |
| 3276 | 0.0003 | 0 | 0.00 | 0.02 |
| 3280 | 0.0003 | 0 | 0.00 | 0.02 |
| 3290 | 0.0003 | 0 | 0.00 | 0.02 |
| 3328 | 0.0006 | 0 | 0.00 | 0.02 |
| 3337 | 0.0003 | 0 | 0.00 | 0.02 |
| 3377 | 0.0003 | 0 | 0.00 | 0.02 |
| 3448 | 0.0003 | 0 | 0.00 | 0.02 |
| 3449 | 0.0003 | 0 | 0.00 | 0.02 |
| 3501 | 0.0003 | 0 | 0.00 | 0.02 |
| 3511 | 0.0003 | 0 | 0.00 | 0.02 |
| 3540 | 0.0003 | 0 | 0.00 | 0.02 |
| 3566 | 0.0003 | 0 | 0.00 | 0.01 |
| 3607 | 0.0003 | 0 | 0.00 | 0.01 |
| 3619 | 0.0003 | 0 | 0.00 | 0.01 |
| 3623 | 0.0003 | 0 | 0.00 | 0.01 |
| 3689 | 0.0003 | 0 | 0.00 | 0.01 |
| 3724 | 0.0003 | 0 | 0.00 | 0.01 |
| 3778 | 0.0003 | 0 | 0.00 | 0.01 |
| 3881 | 0.0003 | 0 | 0.00 | 0.01 |
| 3902 | 0.0003 | 0 | 0.00 | 0.01 |

| | | | | |
|---|---|---|---|---|
| 3919 | 0.0003 | 0 | 0.00 | 0.01 |
| 4104 | 0.0003 | 0 | 0.00 | 0.01 |
| 4266 | 0.0003 | 0 | 0.00 | 0.01 |
| 4294 | 0.0003 | 0 | 0.00 | 0.01 |
| 4612 | 0.0003 | 0 | 0.00 | 0.01 |
| 4624 | 0.0003 | 0 | 0.00 | 0.01 |
| 4636 | 0.0003 | 0 | 0.00 | 0.01 |
| 4678 | 0.0003 | 0 | 0.00 | 0.01 |
| 4878 | 0.0003 | 0 | 0.00 | 0.01 |
| 5062 | 0.0003 | 0 | 0.00 | 0.01 |
| 5165 | 0.0003 | 0 | 0.00 | 0.01 |
| 5178 | 0.0003 | 0 | 0.00 | 0.01 |
| 5222 | 0.0003 | 0 | 0.00 | 0.01 |
| 5255 | 0.0003 | 0 | 0.00 | 0.01 |
| 5280 | 0.0003 | 0 | 0.00 | 0.01 |
| 5417 | 0.0003 | 0 | 0.00 | 0.01 |
| 5467 | 0.0003 | 0 | 0.00 | 0.01 |
| 5489 | 0.0003 | 0 | 0.00 | 0.01 |
| 5635 | 0.0003 | 0 | 0.00 | 0.01 |
| 5712 | 0.0003 | 0 | 0.00 | 0.01 |
| 5737 | 0.0003 | 0 | 0.00 | 0.01 |
| 5784 | 0.0003 | 0 | 0.00 | 0.01 |
| 5842 | 0.0003 | 0 | 0.00 | 0.00 |
| 6038 | 0.0003 | 0 | 0.00 | 0.00 |
| 6139 | 0.0003 | 0 | 0.00 | 0.00 |
| 6169 | 0.0003 | 0 | 0.00 | 0.00 |
| 6416 | 0.0003 | 0 | 0.00 | 0.00 |
| 6475 | 0.0003 | 0 | 0.00 | 0.00 |
| 6555 | 0.0003 | 0 | 0.00 | 0.00 |
| 6619 | 0.0003 | 0 | 0.00 | 0.00 |
| 6729 | 0.0003 | 0 | 0.00 | 0.00 |
| 6845 | 0.0003 | 0 | 0.00 | 0.00 |
| 7337 | 0.0003 | 0 | 0.00 | 0.00 |
| 7538 | 0.0003 | 0 | 0.00 | 0.00 |
| 7951 | 0.0003 | 0 | 0.00 | 0.00 |
| 8003 | 0.0003 | 0 | 0.00 | 0.00 |
| 8075 | 0.0003 | 0 | 0.00 | 0.00 |

**Supplemental Table 4: Percent of incorrect variant calls by depth calculated from Flex data**

| depth | mean_total_count | mean_perc_wrong |
|---|---|---|
| 2 | 94,938.09 | 0.03140169 |
| 3 | 69,177.17 | 0.02931339 |
| 4 | 55,006.38 | 0.02642018 |
| 5 | 46,149.11 | 0.01872583 |
| 6 | 40,364.98 | 0.01697670 |
| 7 | 36,153.04 | 0.01170630 |
| 8 | 32,713.77 | 0.01143594 |
| 9 | 30,021.79 | 0.01076566 |
| 10 | 28,064.19 | 0.00807667 |
| 11 | 26,263.30 | 0.00683810 |
| 12 | 24,451.74 | 0.00755145 |
| 13 | 23,283.17 | 0.00903013 |
| 14 | 22,048.68 | 0.00582028 |
| 15 | 21,090.11 | 0.00415463 |
| 16 | 20,098.55 | 0.00498335 |
| 17 | 19,496.00 | 0.00305278 |
| 18 | 18,487.87 | 0.00429449 |
| 19 | 17,863.51 | 0.00572546 |
| 20 | 17,091.87 | 0.00296816 |
| 21 | 16,480.36 | 0.00357076 |
| 22 | 15,992.94 | 0.00370538 |
| 23 | 15,567.68 | 0.00132888 |
| 24 | 14,982.47 | 0.00447906 |
| 25 | 14,529.98 | 0.00177145 |
| 26 | 14,316.17 | 0.00155853 |
| 27 | 13,845.68 | 0.00274359 |
| 28 | 13,220.28 | 0.00322707 |
| 29 | 12,961.43 | 0.00262358 |
| 30 | 12,684.04 | 0.00275081 |
| 31 | 12,428.34 | 0.00253660 |
| 32 | 12,070.45 | 0.00390209 |
| 33 | 11,648.96 | 0.00142724 |
| 34 | 11,429.04 | 0.00166911 |
| 35 | 11,087.36 | 0.00000000 |
| 36 | 10,535.45 | 0.00221955 |
| 37 | 10,369.74 | 0.00118801 |
| 38 | 10,428.00 | 0.00051887 |
| 39 | 9,932.02 | 0.00095320 |
| 40 | 9,852.09 | 0.00103382 |
| 41 | 9,468.98 | 0.00158097 |
| 42 | 9,283.70 | 0.00112263 |
| 43 | 9,142.23 | 0.00000000 |
| 44 | 9,133.28 | 0.00110909 |

| | | |
|---|---|---|
| 45 | 8,569.38 | 0.00160466 |
| 46 | 8,559.81 | 0.00081279 |
| 47 | 8,368.38 | 0.00234644 |
| 48 | 8,132.38 | 0.00074654 |
| 49 | 8,011.19 | 0.00031451 |
| 50 | 7,739.55 | 0.00128789 |
| 51 | 7,567.47 | 0.00273478 |
| 52 | 7,398.43 | 0.00040550 |
| 53 | 7,302.26 | 0.00048867 |
| 54 | 7,273.49 | 0.00063879 |
| 55 | 7,085.89 | 0.00000000 |
| 56 | 6,848.53 | 0.00000000 |
| 57 | 6,719.94 | 0.00026138 |
| 58 | 6,852.07 | 0.00074949 |
| 59 | 6,456.77 | 0.00263049 |
| 60 | 6,241.91 | 0.00000000 |
| 61 | 6,224.55 | 0.00076866 |
| 62 | 6,290.51 | 0.00000000 |
| 63 | 5,997.53 | 0.00116696 |
| 64 | 6,079.35 | 0.00000000 |
| 65 | 6,018.09 | 0.00000000 |
| 66 | 5,678.83 | 0.00071393 |
| 67 | 5,729.33 | 0.00000000 |
| 68 | 5,686.28 | 0.00045432 |
| 69 | 5,583.85 | 0.00000000 |
| 70 | 5,214.87 | 0.00000000 |
| 71 | 5,073.28 | 0.00035185 |
| 72 | 5,103.04 | 0.00057134 |
| 73 | 5,050.04 | 0.00071735 |
| 74 | 4,970.96 | 0.00000000 |
| 75 | 4,970.00 | 0.00000000 |
| 76 | 5,070.37 | 0.00000000 |
| 77 | 4,754.85 | 0.00036136 |
| 78 | 4,816.26 | 0.00061654 |
| 79 | 4,484.98 | 0.00000000 |
| 80 | 4,466.96 | 0.00000000 |
| 81 | 4,414.91 | 0.00000000 |
| 82 | 4,527.00 | 0.00125228 |
| 83 | 4,298.15 | 0.00031615 |
| 84 | 4,217.98 | 0.00077257 |
| 85 | 4,198.02 | 0.00000000 |
| 86 | 4,216.74 | 0.00000000 |
| 87 | 4,088.81 | 0.00000000 |
| 88 | 4,072.91 | 0.00000000 |

| | | |
|---|---|---|
| 89 | 4,017.63 | 0.00000000 |
| 90 | 3,790.26 | 0.00000000 |
| 91 | 3,920.09 | 0.00000000 |
| 92 | 3,933.54 | 0.00000000 |
| 93 | 3,747.70 | 0.00000000 |
| 94 | 3,694.68 | 0.00000000 |
| 95 | 3,652.43 | 0.00000000 |
| 96 | 3,735.50 | 0.00000000 |
| 97 | 3,624.32 | 0.00062856 |
| 98 | 3,535.06 | 0.00078919 |
| 99 | 3,455.55 | 0.00000000 |
| 100 | 3,363.47 | 0.00000000 |
| 101 | 3,345.87 | 0.00000000 |
| 102 | 3,373.57 | 0.00000000 |
| 103 | 3,302.72 | 0.00000000 |
| 104 | 3,319.87 | 0.00311785 |
| 105 | 3,146.39 | 0.00000000 |
| 106 | 3,135.34 | 0.00000000 |
| 107 | 3,155.00 | 0.00000000 |
| 108 | 3,015.26 | 0.00000000 |
| 109 | 3,118.67 | 0.00000000 |
| 110 | 3,071.83 | 0.00000000 |
| 111 | 3,157.07 | 0.00000000 |
| 112 | 3,024.98 | 0.00000000 |
| 113 | 3,012.43 | 0.00000000 |
| 114 | 2,945.83 | 0.00000000 |
| 115 | 2,992.87 | 0.00000000 |
| 116 | 2,889.98 | 0.00000000 |
| 117 | 2,853.43 | 0.00000000 |
| 118 | 2,789.13 | 0.00000000 |
| 119 | 2,713.22 | 0.00000000 |
| 120 | 2,870.87 | 0.00096534 |
| 121 | 2,876.98 | 0.00000000 |
| 122 | 2,660.24 | 0.00000000 |
| 123 | 2,666.89 | 0.00000000 |
| 124 | 2,675.83 | 0.00000000 |
| 125 | 2,670.20 | 0.00000000 |
| 126 | 2,605.85 | 0.00000000 |
| 127 | 2,583.83 | 0.00000000 |
| 128 | 2,526.58 | 0.00104477 |
| 129 | 2,532.91 | 0.00000000 |
| 130 | 2,606.16 | 0.00000000 |
| 131 | 2,368.83 | 0.00000000 |
| 132 | 2,404.40 | 0.00000000 |

| | | |
|---|---|---|
| 133 | 2,465.24 | 0.00000000 |
| 134 | 2,369.58 | 0.00000000 |
| 135 | 2,356.20 | 0.00000000 |
| 136 | 2,381.00 | 0.00000000 |
| 137 | 2,306.57 | 0.00041646 |
| 138 | 2,333.00 | 0.00000000 |
| 139 | 2,192.48 | 0.00000000 |
| 140 | 2,326.44 | 0.00000000 |
| 141 | 2,308.93 | 0.00260518 |
| 142 | 2,237.62 | 0.00000000 |
| 143 | 2,118.41 | 0.00000000 |
| 144 | 2,112.81 | 0.00000000 |
| 145 | 2,179.38 | 0.00000000 |
| 146 | 2,109.11 | 0.00000000 |
| 147 | 2,143.67 | 0.00000000 |
| 148 | 2,092.50 | 0.00000000 |
| 149 | 2,152.50 | 0.00000000 |
| 150 | 2,021.26 | 0.00067702 |
| 151 | 2,061.56 | 0.00000000 |
| 152 | 2,015.16 | 0.00000000 |
| 153 | 2,010.51 | 0.00000000 |
| 154 | 1,971.76 | 0.00000000 |
| 155 | 1,927.84 | 0.00000000 |
| 156 | 2,036.49 | 0.00000000 |
| 157 | 1,951.00 | 0.00000000 |
| 158 | 1,938.16 | 0.00000000 |
| 159 | 1,909.29 | 0.00000000 |
| 160 | 1,925.44 | 0.00000000 |
| 161 | 1,845.56 | 0.00105419 |
| 162 | 1,828.38 | 0.00000000 |
| 163 | 1,894.71 | 0.00000000 |
| 164 | 1,827.04 | 0.00000000 |
| 165 | 1,853.42 | 0.00000000 |
| 166 | 1,822.18 | 0.00000000 |
| 167 | 1,763.49 | 0.00000000 |
| 168 | 1,739.47 | 0.00000000 |
| 169 | 1,772.47 | 0.00000000 |
| 170 | 1,758.47 | 0.00000000 |
| 171 | 1,786.80 | 0.00000000 |
| 172 | 1,765.07 | 0.00000000 |
| 173 | 1,724.62 | 0.00000000 |
| 174 | 1,682.29 | 0.00000000 |
| 175 | 1,631.27 | 0.00000000 |
| 176 | 1,715.53 | 0.00000000 |

| | | |
|---|---|---|
| 177 | 1,701.75 | 0.00171268 |
| 178 | 1,700.85 | 0.00000000 |
| 179 | 1,595.83 | 0.00000000 |
| 180 | 1,611.44 | 0.00000000 |
| 181 | 1,642.07 | 0.00000000 |
| 182 | 1,578.29 | 0.00000000 |
| 183 | 1,624.56 | 0.00000000 |
| 184 | 1,659.27 | 0.00000000 |
| 185 | 1,580.56 | 0.00000000 |
| 186 | 1,602.76 | 0.00000000 |
| 187 | 1,526.16 | 0.00000000 |
| 188 | 1,583.91 | 0.00000000 |
| 189 | 1,573.28 | 0.00000000 |
| 190 | 1,507.98 | 0.00000000 |
| 191 | 1,472.31 | 0.00000000 |
| 192 | 1,468.85 | 0.00117129 |
| 193 | 1,500.62 | 0.00000000 |
| 194 | 1,520.84 | 0.00000000 |
| 195 | 1,494.41 | 0.00088623 |
| 196 | 1,443.20 | 0.00000000 |
| 197 | 1,425.70 | 0.00000000 |
| 198 | 1,423.50 | 0.00000000 |
| 199 | 1,355.67 | 0.00000000 |
| 200 | 1,431.89 | 0.00000000 |
| 201 | 1,462.53 | 0.00000000 |
| 202 | 1,397.71 | 0.00000000 |
| 203 | 1,437.98 | 0.00000000 |
| 204 | 1,378.05 | 0.00000000 |
| 205 | 1,352.98 | 0.00000000 |
| 206 | 1,390.73 | 0.00000000 |
| 207 | 1,388.56 | 0.00000000 |
| 208 | 1,340.76 | 0.00000000 |
| 209 | 1,329.70 | 0.00000000 |
| 210 | 1,331.44 | 0.00000000 |
| 211 | 1,350.29 | 0.00000000 |
| 212 | 1,325.67 | 0.00000000 |
| 213 | 1,303.29 | 0.00000000 |
| 214 | 1,337.29 | 0.00000000 |
| 215 | 1,242.04 | 0.00000000 |
| 216 | 1,316.27 | 0.00000000 |
| 217 | 1,261.36 | 0.00000000 |
| 218 | 1,338.36 | 0.00000000 |
| 219 | 1,258.57 | 0.00000000 |
| 220 | 1,272.30 | 0.00000000 |

| | | |
|---|---|---|
| 221 | 1,298.93 | 0.00000000 |
| 222 | 1,229.41 | 0.00000000 |
| 223 | 1,262.16 | 0.00000000 |
| 224 | 1,206.44 | 0.00000000 |
| 225 | 1,257.16 | 0.00000000 |
| 226 | 1,276.36 | 0.00000000 |
| 227 | 1,177.30 | 0.00000000 |
| 228 | 1,216.44 | 0.00166583 |
| 229 | 1,148.37 | 0.00000000 |
| 230 | 1,128.37 | 0.00000000 |
| 231 | 1,172.15 | 0.00000000 |
| 232 | 1,149.23 | 0.00000000 |
| 233 | 1,097.24 | 0.00000000 |
| 234 | 1,196.14 | 0.00000000 |
| 235 | 1,212.23 | 0.00000000 |
| 236 | 1,159.61 | 0.00000000 |
| 237 | 1,185.42 | 0.00000000 |
| 238 | 1,131.07 | 0.00000000 |
| 239 | 1,164.16 | 0.00000000 |
| 240 | 1,140.82 | 0.00000000 |
| 241 | 1,110.05 | 0.00000000 |
| 242 | 1,133.58 | 0.00000000 |
| 243 | 1,118.09 | 0.00000000 |
| 244 | 1,151.60 | 0.00000000 |
| 245 | 1,121.39 | 0.00000000 |
| 246 | 1,146.27 | 0.00000000 |
| 247 | 1,120.14 | 0.00000000 |
| 248 | 1,056.05 | 0.00000000 |
| 249 | 1,130.48 | 0.00000000 |
| 250 | 1,075.00 | 0.00000000 |
| 251 | 1,120.36 | 0.00089092 |
| 252 | 1,042.93 | 0.00000000 |
| 253 | 1,080.98 | 0.00000000 |
| 254 | 1,105.75 | 0.00000000 |
| 255 | 1,105.07 | 0.00000000 |
| 256 | 958.62 | 0.00644122 |
| 257 | 1,046.77 | 0.00161072 |
| 258 | 1,113.67 | 0.00000000 |
| 259 | 1,055.20 | 0.00000000 |
| 260 | 1,050.14 | 0.00000000 |
| 261 | 1,024.84 | 0.00000000 |
| 262 | 947.64 | 0.00000000 |
| 263 | 981.58 | 0.00000000 |
| 264 | 1,006.68 | 0.00000000 |

| | | |
|---|---|---|
| 265 | 960.98 | 0.00000000 |
| 266 | 969.49 | 0.00000000 |
| 267 | 971.42 | 0.00000000 |
| 268 | 983.02 | 0.00000000 |
| 269 | 1,040.38 | 0.00000000 |
| 270 | 1,047.25 | 0.00000000 |
| 271 | 942.30 | 0.00000000 |
| 272 | 958.52 | 0.00000000 |
| 273 | 1,042.86 | 0.00137325 |
| 274 | 904.11 | 0.00000000 |
| 275 | 936.55 | 0.00000000 |
| 276 | 931.46 | 0.00000000 |
| 277 | 960.02 | 0.00000000 |
| 278 | 930.00 | 0.00000000 |
| 279 | 995.29 | 0.00000000 |
| 280 | 938.43 | 0.00171397 |
| 281 | 946.09 | 0.00000000 |
| 282 | 945.18 | 0.00000000 |
| 283 | 929.20 | 0.00548987 |
| 284 | 880.42 | 0.00000000 |
| 285 | 921.29 | 0.00000000 |
| 286 | 917.47 | 0.00000000 |
| 287 | 920.62 | 0.00000000 |
| 288 | 909.50 | 0.00000000 |
| 289 | 888.09 | 0.00000000 |
| 290 | 947.32 | 0.00000000 |
| 291 | 876.43 | 0.00000000 |
| 292 | 903.52 | 0.00000000 |
| 293 | 911.60 | 0.00000000 |
| 294 | 913.77 | 0.00000000 |
| 295 | 885.98 | 0.00000000 |
| 296 | 937.75 | 0.00000000 |
| 297 | 917.73 | 0.00000000 |
| 298 | 863.84 | 0.00000000 |
| 299 | 861.73 | 0.00000000 |
| 300 | 907.89 | 0.00000000 |
| 301 | 901.05 | 0.00000000 |
| 302 | 876.61 | 0.00000000 |
| 303 | 847.66 | 0.00000000 |
| 304 | 816.82 | 0.00000000 |
| 305 | 847.05 | 0.00000000 |
| 306 | 825.27 | 0.00000000 |
| 307 | 823.09 | 0.00000000 |
| 308 | 825.62 | 0.00000000 |

| | | |
|---|---|---|
| 309 | 744.57 | 0.00000000 |
| 310 | 812.84 | 0.00000000 |
| 311 | 871.81 | 0.00000000 |
| 312 | 799.91 | 0.00000000 |
| 313 | 857.74 | 0.00000000 |
| 314 | 840.07 | 0.00000000 |
| 315 | 779.43 | 0.00000000 |
| 316 | 794.61 | 0.00000000 |
| 317 | 788.09 | 0.00000000 |
| 318 | 777.50 | 0.00000000 |
| 319 | 767.16 | 0.00000000 |
| 320 | 735.98 | 0.00000000 |
| 321 | 824.38 | 0.00000000 |
| 322 | 868.00 | 0.00000000 |
| 323 | 741.44 | 0.00000000 |
| 324 | 751.58 | 0.00000000 |
| 325 | 772.14 | 0.00000000 |
| 326 | 759.02 | 0.00000000 |
| 327 | 711.43 | 0.00183581 |
| 328 | 769.50 | 0.00000000 |
| 329 | 747.04 | 0.00000000 |
| 330 | 763.62 | 0.00279473 |
| 331 | 790.63 | 0.00000000 |
| 332 | 775.83 | 0.00000000 |
| 333 | 732.05 | 0.00000000 |
| 334 | 745.86 | 0.00000000 |
| 335 | 755.64 | 0.00000000 |
| 336 | 769.93 | 0.00000000 |
| 337 | 750.33 | 0.00000000 |
| 338 | 682.31 | 0.00000000 |
| 339 | 758.11 | 0.00000000 |
| 340 | 770.62 | 0.00000000 |
| 341 | 658.75 | 0.00000000 |
| 342 | 728.32 | 0.00000000 |
| 343 | 694.86 | 0.00000000 |
| 344 | 713.50 | 0.00000000 |
| 345 | 692.18 | 0.00000000 |
| 346 | 719.33 | 0.00000000 |
| 347 | 698.55 | 0.00000000 |
| 348 | 693.58 | 0.00000000 |
| 349 | 686.98 | 0.00000000 |
| 350 | 732.33 | 0.00000000 |
| 351 | 729.05 | 0.00000000 |
| 352 | 724.95 | 0.00000000 |

| | | |
|---|---|---|
| 353 | 686.23 | 0.00000000 |
| 354 | 690.20 | 0.00000000 |
| 355 | 685.00 | 0.00000000 |
| 356 | 677.09 | 0.00147282 |
| 357 | 663.40 | 0.00000000 |
| 358 | 736.44 | 0.00000000 |
| 359 | 697.02 | 0.00000000 |
| 360 | 637.70 | 0.00000000 |
| 361 | 662.44 | 0.00000000 |
| 362 | 675.98 | 0.00000000 |
| 363 | 636.80 | 0.00000000 |
| 364 | 657.73 | 0.00000000 |
| 365 | 681.22 | 0.00000000 |
| 366 | 623.77 | 0.00000000 |
| 367 | 662.84 | 0.00000000 |
| 368 | 652.43 | 0.00000000 |
| 369 | 600.80 | 0.00000000 |
| 370 | 670.50 | 0.00000000 |
| 371 | 637.14 | 0.00000000 |
| 372 | 688.02 | 0.00000000 |
| 373 | 619.36 | 0.00000000 |
| 374 | 679.62 | 0.00000000 |
| 375 | 622.53 | 0.00000000 |
| 376 | 710.40 | 0.00000000 |
| 377 | 675.69 | 0.00000000 |
| 378 | 636.35 | 0.00000000 |
| 379 | 634.98 | 0.00000000 |
| 380 | 647.84 | 0.00000000 |
| 381 | 672.49 | 0.00000000 |
| 382 | 610.74 | 0.00000000 |
| 383 | 649.10 | 0.00000000 |
| 384 | 646.21 | 0.00000000 |
| 385 | 606.12 | 0.00000000 |
| 386 | 585.43 | 0.00000000 |
| 387 | 645.09 | 0.00000000 |
| 388 | 626.14 | 0.00000000 |
| 389 | 669.49 | 0.00182399 |
| 390 | 589.80 | 0.00000000 |
| 391 | 572.07 | 0.00000000 |
| 392 | 609.09 | 0.00252245 |
| 393 | 570.95 | 0.00000000 |
| 394 | 613.80 | 0.00000000 |
| 395 | 566.95 | 0.00000000 |
| 396 | 592.49 | 0.00000000 |

| | | |
|---|---|---|
| 397 | 607.90 | 0.00000000 |
| 398 | 588.98 | 0.00000000 |
| 399 | 614.65 | 0.00000000 |
| 400 | 585.09 | 0.00000000 |
| 401 | 571.18 | 0.00000000 |
| 402 | 560.32 | 0.00000000 |
| 403 | 584.02 | 0.00000000 |
| 404 | 549.20 | 0.00000000 |
| 405 | 535.98 | 0.00000000 |
| 406 | 614.10 | 0.00000000 |
| 407 | 586.90 | 0.00000000 |
| 408 | 612.58 | 0.00000000 |
| 409 | 587.80 | 0.00000000 |
| 410 | 606.24 | 0.00000000 |
| 411 | 605.66 | 0.00000000 |
| 412 | 583.86 | 0.00000000 |
| 413 | 607.64 | 0.00000000 |
| 414 | 590.81 | 0.00000000 |
| 415 | 620.72 | 0.00000000 |
| 416 | 547.44 | 0.00000000 |
| 417 | 524.16 | 0.00000000 |
| 418 | 556.57 | 0.00000000 |
| 419 | 545.14 | 0.00000000 |
| 420 | 502.95 | 0.00270563 |
| 421 | 551.95 | 0.00000000 |
| 422 | 539.00 | 0.00000000 |
| 423 | 557.25 | 0.00000000 |
| 424 | 513.26 | 0.00000000 |
| 425 | 515.48 | 0.00000000 |
| 426 | 528.28 | 0.00000000 |
| 427 | 510.23 | 0.00000000 |
| 428 | 508.52 | 0.00000000 |
| 429 | 547.86 | 0.00000000 |
| 430 | 542.64 | 0.00000000 |
| 431 | 498.49 | 0.00000000 |
| 432 | 540.00 | 0.00000000 |
| 433 | 502.57 | 0.00000000 |
| 434 | 527.02 | 0.00000000 |
| 435 | 503.95 | 0.00000000 |
| 436 | 544.02 | 0.00141052 |
| 437 | 528.23 | 0.00000000 |
| 438 | 518.43 | 0.00000000 |
| 439 | 500.40 | 0.00000000 |
| 440 | 501.62 | 0.00000000 |

| | | |
|---|---|---|
| 441 | 538.14 | 0.00000000 |
| 442 | 523.02 | 0.00000000 |
| 443 | 555.07 | 0.00000000 |
| 444 | 591.73 | 0.00000000 |
| 445 | 538.56 | 0.00000000 |
| 446 | 556.93 | 0.00000000 |
| 447 | 510.16 | 0.00000000 |
| 448 | 476.16 | 0.00000000 |
| 449 | 452.23 | 0.00000000 |
| 450 | 492.58 | 0.00000000 |
| 451 | 515.02 | 0.00000000 |
| 452 | 493.76 | 0.00000000 |
| 453 | 499.43 | 0.00000000 |
| 454 | 544.05 | 0.00000000 |
| 455 | 465.56 | 0.01107420 |
| 456 | 498.29 | 0.00000000 |
| 457 | 528.44 | 0.00000000 |
| 458 | 507.02 | 0.00000000 |
| 459 | 508.95 | 0.00000000 |
| 460 | 503.53 | 0.00000000 |
| 461 | 477.63 | 0.00000000 |
| 462 | 533.60 | 0.00000000 |
| 463 | 470.83 | 0.00000000 |
| 464 | 525.34 | 0.00000000 |
| 465 | 487.60 | 0.00000000 |
| 466 | 483.35 | 0.00000000 |
| 467 | 515.86 | 0.00000000 |
| 468 | 458.22 | 0.00000000 |
| 469 | 504.70 | 0.00000000 |
| 470 | 483.52 | 0.00000000 |
| 471 | 468.02 | 0.00000000 |
| 472 | 481.21 | 0.00000000 |
| 473 | 425.09 | 0.00000000 |
| 474 | 463.76 | 0.00000000 |
| 475 | 468.11 | 0.00000000 |
| 476 | 476.51 | 0.00000000 |
| 477 | 449.53 | 0.00000000 |
| 478 | 485.26 | 0.00000000 |
| 479 | 431.35 | 0.00000000 |
| 480 | 435.86 | 0.00000000 |
| 481 | 474.61 | 0.00000000 |
| 482 | 427.22 | 0.00000000 |
| 483 | 433.23 | 0.00000000 |
| 484 | 439.02 | 0.00000000 |

| | | |
|---|---|---|
| 485 | 424.93 | 0.00000000 |
| 486 | 489.05 | 0.00000000 |
| 487 | 450.91 | 0.00000000 |
| 488 | 470.93 | 0.00000000 |
| 489 | 443.33 | 0.00000000 |
| 490 | 442.95 | 0.00000000 |
| 491 | 413.46 | 0.00000000 |
| 492 | 477.22 | 0.00000000 |
| 493 | 455.41 | 0.00000000 |
| 494 | 438.93 | 0.00000000 |
| 495 | 407.00 | 0.00000000 |
| 496 | 442.63 | 0.00000000 |
| 497 | 445.44 | 0.00000000 |
| 498 | 434.57 | 0.00000000 |
| 499 | 430.81 | 0.00000000 |
| 500 | 412.71 | 0.00000000 |
| 501 | 442.42 | 0.00000000 |
| 502 | 411.55 | 0.00000000 |
| 503 | 418.10 | 0.00000000 |
| 504 | 407.93 | 0.00000000 |
| 505 | 437.54 | 0.00268025 |
| 506 | 422.93 | 0.00000000 |
| 507 | 423.61 | 0.00000000 |
| 508 | 393.71 | 0.00000000 |
| 509 | 425.43 | 0.00216450 |
| 510 | 410.68 | 0.00000000 |
| 511 | 391.50 | 0.00000000 |
| 512 | 430.90 | 0.00000000 |
| 513 | 405.25 | 0.00000000 |
| 514 | 432.34 | 0.00000000 |
| 515 | 412.21 | 0.00000000 |
| 516 | 467.14 | 0.00000000 |
| 517 | 442.83 | 0.00000000 |
| 518 | 401.17 | 0.00000000 |
| 519 | 403.62 | 0.00000000 |
| 520 | 397.95 | 0.00000000 |
| 521 | 410.34 | 0.00000000 |
| 522 | 426.46 | 0.00000000 |
| 523 | 441.05 | 0.00000000 |
| 524 | 439.88 | 0.00000000 |
| 525 | 387.79 | 0.00000000 |
| 526 | 474.15 | 0.00000000 |
| 527 | 401.67 | 0.00000000 |
| 528 | 410.35 | 0.00000000 |

| | | |
|---|---|---|
| 529 | 364.74 | 0.00000000 |
| 530 | 367.30 | 0.00000000 |
| 531 | 385.24 | 0.00000000 |
| 532 | 389.95 | 0.00000000 |
| 533 | 371.40 | 0.00000000 |
| 534 | 369.67 | 0.00000000 |
| 535 | 410.78 | 0.00000000 |
| 536 | 369.48 | 0.00000000 |
| 537 | 367.90 | 0.00000000 |
| 538 | 383.40 | 0.00000000 |
| 539 | 346.38 | 0.00000000 |
| 540 | 386.66 | 0.00000000 |
| 541 | 473.90 | 0.00000000 |
| 542 | 388.74 | 0.00000000 |
| 543 | 413.20 | 0.00000000 |
| 544 | 377.12 | 0.00000000 |
| 545 | 357.19 | 0.00000000 |
| 546 | 363.23 | 0.00000000 |
| 547 | 348.00 | 0.00000000 |
| 548 | 374.14 | 0.00000000 |
| 549 | 375.44 | 0.00000000 |
| 550 | 354.14 | 0.00000000 |
| 551 | 343.51 | 0.00000000 |
| 552 | 378.88 | 0.00000000 |
| 553 | 422.73 | 0.00000000 |
| 554 | 382.66 | 0.00000000 |
| 555 | 369.05 | 0.00000000 |
| 556 | 375.61 | 0.00000000 |
| 557 | 389.08 | 0.00000000 |
| 558 | 372.25 | 0.00000000 |
| 559 | 374.95 | 0.00464576 |
| 560 | 377.10 | 0.00000000 |
| 561 | 344.07 | 0.00000000 |
| 562 | 371.85 | 0.00000000 |
| 563 | 387.17 | 0.00000000 |
| 564 | 399.10 | 0.00000000 |
| 565 | 380.58 | 0.00000000 |
| 566 | 343.40 | 0.00000000 |
| 567 | 364.95 | 0.00000000 |
| 568 | 301.09 | 0.00000000 |
| 569 | 326.38 | 0.00000000 |
| 570 | 353.95 | 0.00000000 |
| 571 | 351.60 | 0.00000000 |
| 572 | 351.69 | 0.00000000 |

| | | |
|---|---|---|
| 573 | 353.24 | 0.00000000 |
| 574 | 371.05 | 0.00000000 |
| 575 | 345.16 | 0.00000000 |
| 576 | 340.78 | 0.00000000 |
| 577 | 324.31 | 0.00000000 |
| 578 | 327.00 | 0.00000000 |
| 579 | 343.93 | 0.00000000 |
| 580 | 322.40 | 0.00000000 |
| 581 | 304.64 | 0.00000000 |
| 582 | 314.37 | 0.00000000 |
| 583 | 303.12 | 0.00000000 |
| 584 | 333.14 | 0.00000000 |
| 585 | 371.68 | 0.00000000 |
| 586 | 354.44 | 0.00000000 |
| 587 | 337.42 | 0.00000000 |
| 588 | 353.15 | 0.00000000 |
| 589 | 371.82 | 0.00000000 |
| 590 | 346.60 | 0.00000000 |
| 591 | 348.15 | 0.00000000 |
| 592 | 365.43 | 0.00000000 |
| 593 | 342.58 | 0.00000000 |
| 594 | 326.88 | 0.00442556 |
| 595 | 336.60 | 0.00000000 |
| 596 | 325.00 | 0.00000000 |
| 597 | 356.83 | 0.00000000 |
| 598 | 308.93 | 0.00000000 |
| 599 | 317.60 | 0.00000000 |
| 600 | 372.97 | 0.00000000 |
| 601 | 334.93 | 0.00000000 |
| 602 | 305.56 | 0.00000000 |
| 603 | 366.43 | 0.00000000 |
| 604 | 321.29 | 0.00000000 |
| 605 | 316.24 | 0.00000000 |
| 606 | 321.70 | 0.00000000 |
| 607 | 313.51 | 0.00000000 |
| 608 | 311.45 | 0.00000000 |
| 609 | 326.97 | 0.00000000 |
| 610 | 356.03 | 0.00000000 |
| 611 | 288.52 | 0.00000000 |
| 612 | 350.85 | 0.00000000 |
| 613 | 319.90 | 0.00000000 |
| 614 | 353.00 | 0.00000000 |
| 615 | 347.95 | 0.00000000 |
| 616 | 328.67 | 0.00000000 |

| | | |
|---|---|---|
| 617 | 380.65 | 0.00000000 |
| 618 | 324.58 | 0.00000000 |
| 619 | 347.69 | 0.00000000 |
| 620 | 371.68 | 0.00000000 |
| 621 | 320.84 | 0.00000000 |
| 622 | 372.00 | 0.00000000 |
| 623 | 347.13 | 0.00000000 |
| 624 | 342.77 | 0.00000000 |
| 625 | 310.18 | 0.00000000 |
| 626 | 338.18 | 0.00000000 |
| 627 | 296.93 | 0.00000000 |
| 628 | 317.30 | 0.00000000 |
| 629 | 314.25 | 0.00000000 |
| 630 | 334.25 | 0.00000000 |
| 631 | 303.95 | 0.00000000 |
| 632 | 325.72 | 0.00000000 |
| 633 | 309.69 | 0.00000000 |
| 634 | 282.69 | 0.00000000 |
| 635 | 355.40 | 0.00000000 |
| 636 | 362.19 | 0.00000000 |
| 637 | 323.64 | 0.00000000 |
| 638 | 308.63 | 0.00000000 |
| 639 | 344.33 | 0.00000000 |
| 640 | 290.69 | 0.00000000 |
| 641 | 276.93 | 0.00000000 |
| 642 | 309.18 | 0.00000000 |
| 643 | 344.20 | 0.00000000 |
| 644 | 332.62 | 0.00000000 |
| 645 | 363.59 | 0.00000000 |
| 646 | 302.93 | 0.00000000 |
| 647 | 345.87 | 0.00000000 |
| 648 | 330.62 | 0.00000000 |
| 649 | 296.03 | 0.00000000 |
| 650 | 317.39 | 0.00000000 |
| 651 | 268.86 | 0.00000000 |
| 652 | 312.08 | 0.00000000 |
| 653 | 321.38 | 0.00000000 |
| 654 | 278.58 | 0.00000000 |
| 655 | 278.70 | 0.00000000 |
| 656 | 288.13 | 0.00000000 |
| 657 | 310.58 | 0.00000000 |
| 658 | 290.16 | 0.00000000 |
| 659 | 304.77 | 0.00000000 |
| 660 | 301.00 | 0.00000000 |

| | | |
|---|---|---|
| 661 | 287.35 | 0.00000000 |
| 662 | 283.38 | 0.00000000 |
| 663 | 274.59 | 0.00000000 |
| 664 | 283.97 | 0.00000000 |
| 665 | 305.54 | 0.00000000 |
| 666 | 259.78 | 0.00000000 |
| 667 | 295.85 | 0.00000000 |
| 668 | 291.55 | 0.00000000 |
| 669 | 279.03 | 0.00000000 |
| 670 | 292.85 | 0.00000000 |
| 671 | 266.39 | 0.00000000 |
| 672 | 297.45 | 0.00000000 |
| 673 | 288.41 | 0.00000000 |
| 674 | 336.27 | 0.00000000 |
| 675 | 282.10 | 0.00000000 |
| 676 | 305.41 | 0.00000000 |
| 677 | 315.41 | 0.00000000 |
| 678 | 295.41 | 0.00000000 |
| 679 | 293.21 | 0.00000000 |
| 680 | 270.90 | 0.00000000 |
| 681 | 284.26 | 0.00000000 |
| 682 | 274.70 | 0.00000000 |
| 683 | 253.56 | 0.00000000 |
| 684 | 281.97 | 0.00000000 |
| 685 | 311.78 | 0.00000000 |
| 686 | 348.00 | 0.00000000 |
| 687 | 296.35 | 0.00000000 |
| 688 | 285.27 | 0.00000000 |
| 689 | 280.59 | 0.00000000 |
| 690 | 310.11 | 0.00000000 |
| 691 | 280.88 | 0.00000000 |
| 692 | 260.72 | 0.00000000 |
| 693 | 276.55 | 0.00000000 |
| 694 | 231.57 | 0.00000000 |
| 695 | 286.56 | 0.00000000 |
| 696 | 267.17 | 0.00000000 |
| 697 | 267.86 | 0.00000000 |
| 698 | 232.49 | 0.00000000 |
| 699 | 257.31 | 0.00000000 |
| 700 | 256.78 | 0.00000000 |
| 701 | 270.78 | 0.00000000 |
| 702 | 267.05 | 0.00000000 |
| 703 | 261.13 | 0.00000000 |
| 704 | 273.26 | 0.00000000 |

| | | |
|---|---|---|
| 705 | 271.15 | 0.00000000 |
| 706 | 249.05 | 0.00000000 |
| 707 | 263.11 | 0.00000000 |
| 708 | 243.38 | 0.00000000 |
| 709 | 264.37 | 0.00000000 |
| 710 | 259.65 | 0.00000000 |
| 711 | 270.27 | 0.00000000 |
| 712 | 268.82 | 0.00000000 |
| 713 | 220.34 | 0.00000000 |
| 714 | 232.40 | 0.00000000 |
| 715 | 202.05 | 0.00000000 |
| 716 | 267.31 | 0.00000000 |
| 717 | 267.93 | 0.00000000 |
| 718 | 243.85 | 0.00000000 |
| 719 | 258.76 | 0.00000000 |
| 720 | 251.53 | 0.00000000 |
| 721 | 246.08 | 0.00000000 |
| 722 | 265.68 | 0.00000000 |
| 723 | 223.71 | 0.00000000 |
| 724 | 227.03 | 0.00000000 |
| 725 | 262.41 | 0.00000000 |
| 726 | 259.61 | 0.00000000 |
| 727 | 241.92 | 0.00000000 |
| 728 | 286.43 | 0.00000000 |
| 729 | 273.21 | 0.00000000 |
| 730 | 245.76 | 0.00000000 |
| 731 | 255.10 | 0.00000000 |
| 732 | 235.19 | 0.00000000 |
| 733 | 254.47 | 0.00000000 |
| 734 | 253.63 | 0.00000000 |
| 735 | 245.83 | 0.00000000 |
| 736 | 237.89 | 0.00000000 |
| 737 | 254.41 | 0.00000000 |
| 738 | 212.55 | 0.00000000 |
| 739 | 234.72 | 0.00000000 |
| 740 | 218.11 | 0.00000000 |
| 741 | 239.82 | 0.00000000 |
| 742 | 228.13 | 0.00000000 |
| 743 | 215.54 | 0.00000000 |
| 744 | 269.92 | 0.00000000 |
| 745 | 217.49 | 0.00000000 |
| 746 | 228.78 | 0.00000000 |
| 747 | 285.79 | 0.00000000 |
| 748 | 247.30 | 0.00000000 |

| | | |
|---|---|---|
| 749 | 236.14 | 0.00000000 |
| 750 | 263.53 | 0.00000000 |
| 751 | 258.82 | 0.00000000 |
| 752 | 257.00 | 0.00000000 |
| 753 | 305.31 | 0.00000000 |
| 754 | 235.53 | 0.00000000 |
| 755 | 250.16 | 0.00000000 |
| 756 | 293.90 | 0.00000000 |
| 757 | 282.88 | 0.00000000 |
| 758 | 206.59 | 0.00000000 |
| 759 | 223.00 | 0.00000000 |
| 760 | 248.22 | 0.00000000 |
| 761 | 213.05 | 0.00000000 |
| 762 | 253.80 | 0.00000000 |
| 763 | 264.71 | 0.00000000 |
| 764 | 255.45 | 0.00000000 |
| 765 | 263.46 | 0.00000000 |
| 766 | 261.18 | 0.00000000 |
| 767 | 264.23 | 0.00000000 |
| 768 | 309.27 | 0.00000000 |
| 769 | 257.00 | 0.00000000 |
| 770 | 261.52 | 0.00000000 |
| 771 | 276.56 | 0.00000000 |
| 772 | 225.89 | 0.00000000 |
| 773 | 243.58 | 0.00000000 |
| 774 | 260.50 | 0.00000000 |
| 775 | 252.03 | 0.00000000 |
| 776 | 258.75 | 0.00000000 |
| 777 | 217.86 | 0.00000000 |
| 778 | 217.72 | 0.00000000 |
| 779 | 264.91 | 0.00000000 |
| 780 | 248.20 | 0.00000000 |
| 781 | 232.49 | 0.00000000 |
| 782 | 226.37 | 0.00000000 |
| 783 | 198.15 | 0.00000000 |
| 784 | 213.19 | 0.00000000 |
| 785 | 181.49 | 0.00000000 |
| 786 | 188.73 | 0.00000000 |
| 787 | 214.62 | 0.00000000 |
| 788 | 187.86 | 0.00000000 |
| 789 | 216.13 | 0.00000000 |
| 790 | 235.81 | 0.00000000 |
| 791 | 209.77 | 0.00000000 |
| 792 | 245.49 | 0.00000000 |

| | | |
|---|---|---|
| 793 | 253.66 | 0.00000000 |
| 794 | 211.10 | 0.00000000 |
| 795 | 253.97 | 0.00000000 |
| 796 | 240.89 | 0.00000000 |
| 797 | 216.50 | 0.00000000 |
| 798 | 222.79 | 0.00000000 |
| 799 | 217.37 | 0.00000000 |
| 800 | 215.19 | 0.00000000 |
| 801 | 263.06 | 0.00000000 |
| 802 | 256.72 | 0.00000000 |
| 803 | 249.42 | 0.00000000 |
| 804 | 224.00 | 0.00000000 |
| 805 | 225.97 | 0.00000000 |
| 806 | 245.28 | 0.00000000 |
| 807 | 225.00 | 0.00000000 |
| 808 | 238.17 | 0.00000000 |
| 809 | 263.91 | 0.00000000 |
| 810 | 198.28 | 0.00000000 |
| 811 | 239.23 | 0.00000000 |
| 812 | 213.16 | 0.00000000 |
| 813 | 223.38 | 0.00000000 |
| 814 | 218.11 | 0.00000000 |
| 815 | 223.31 | 0.00000000 |
| 816 | 227.53 | 0.00000000 |
| 817 | 227.51 | 0.00000000 |
| 818 | 220.46 | 0.00000000 |
| 819 | 219.95 | 0.00000000 |
| 820 | 225.53 | 0.00000000 |
| 821 | 216.17 | 0.00000000 |
| 822 | 236.28 | 0.00000000 |
| 823 | 251.09 | 0.00000000 |
| 824 | 245.19 | 0.00000000 |
| 825 | 250.61 | 0.00000000 |
| 826 | 228.97 | 0.00000000 |
| 827 | 217.11 | 0.00000000 |
| 828 | 239.48 | 0.00000000 |
| 829 | 244.44 | 0.00000000 |
| 830 | 200.76 | 0.00000000 |
| 831 | 182.15 | 0.00000000 |
| 832 | 202.00 | 0.00000000 |
| 833 | 219.42 | 0.00000000 |
| 834 | 214.69 | 0.00000000 |
| 835 | 247.97 | 0.00000000 |
| 836 | 202.55 | 0.00000000 |

| | | |
|---|---|---|
| 837 | 229.20 | 0.00000000 |
| 838 | 190.47 | 0.00000000 |
| 839 | 200.14 | 0.00000000 |
| 840 | 184.97 | 0.00000000 |
| 841 | 201.76 | 0.00000000 |
| 842 | 196.42 | 0.00000000 |
| 843 | 202.36 | 0.00000000 |
| 844 | 205.69 | 0.00000000 |
| 845 | 190.86 | 0.00000000 |
| 846 | 243.92 | 0.00000000 |
| 847 | 213.86 | 0.00000000 |
| 848 | 212.29 | 0.00000000 |
| 849 | 214.41 | 0.00000000 |
| 850 | 234.28 | 0.00000000 |
| 851 | 227.14 | 0.00000000 |
| 852 | 209.47 | 0.00000000 |
| 853 | 222.03 | 0.00000000 |
| 854 | 239.69 | 0.00000000 |
| 855 | 220.91 | 0.00000000 |
| 856 | 204.88 | 0.00000000 |
| 857 | 234.85 | 0.00000000 |
| 858 | 194.80 | 0.00000000 |
| 859 | 244.28 | 0.00000000 |
| 860 | 210.57 | 0.00000000 |
| 861 | 221.64 | 0.00000000 |
| 862 | 192.03 | 0.00000000 |
| 863 | 250.77 | 0.00000000 |
| 864 | 209.34 | 0.00000000 |
| 865 | 177.21 | 0.00000000 |
| 866 | 175.84 | 0.00000000 |
| 867 | 217.00 | 0.00000000 |
| 868 | 213.16 | 0.00000000 |
| 869 | 187.27 | 0.00000000 |
| 870 | 207.15 | 0.00000000 |
| 871 | 215.08 | 0.00000000 |
| 872 | 196.12 | 0.00000000 |
| 873 | 198.90 | 0.00000000 |
| 874 | 229.79 | 0.00000000 |
| 875 | 196.57 | 0.00000000 |
| 876 | 199.46 | 0.00000000 |
| 877 | 216.23 | 0.00000000 |
| 878 | 192.89 | 0.00000000 |
| 879 | 196.92 | 0.00000000 |
| 880 | 198.82 | 0.00000000 |

| | | |
|---|---|---|
| 881 | 214.38 | 0.00000000 |
| 882 | 173.37 | 0.00000000 |
| 883 | 179.64 | 0.00000000 |
| 884 | 182.92 | 0.00000000 |
| 885 | 205.97 | 0.00000000 |
| 886 | 170.14 | 0.00000000 |
| 887 | 217.58 | 0.00000000 |
| 888 | 213.55 | 0.00000000 |
| 889 | 207.40 | 0.00000000 |
| 890 | 204.21 | 0.00000000 |
| 891 | 189.72 | 0.00000000 |
| 892 | 208.06 | 0.00000000 |
| 893 | 210.65 | 0.00000000 |
| 894 | 188.30 | 0.00000000 |
| 895 | 228.16 | 0.00000000 |
| 896 | 206.66 | 0.00000000 |
| 897 | 181.63 | 0.00000000 |
| 898 | 218.15 | 0.00000000 |
| 899 | 211.82 | 0.00000000 |
| 900 | 220.45 | 0.00000000 |
| 901 | 215.72 | 0.00000000 |
| 902 | 206.13 | 0.00000000 |
| 903 | 177.30 | 0.00000000 |
| 904 | 209.15 | 0.00000000 |
| 905 | 173.24 | 0.00000000 |
| 906 | 186.78 | 0.00000000 |
| 907 | 225.44 | 0.00000000 |
| 908 | 184.45 | 0.00000000 |
| 909 | 178.61 | 0.00000000 |
| 910 | 203.09 | 0.00000000 |
| 911 | 214.67 | 0.00000000 |
| 912 | 209.80 | 0.00000000 |
| 913 | 194.79 | 0.00000000 |
| 914 | 205.77 | 0.00000000 |
| 915 | 197.50 | 0.00000000 |
| 916 | 208.25 | 0.00000000 |
| 917 | 195.53 | 0.00000000 |
| 918 | 164.95 | 0.00000000 |
| 919 | 178.00 | 0.00000000 |
| 920 | 203.15 | 0.00000000 |
| 921 | 194.48 | 0.00000000 |
| 922 | 184.06 | 0.00000000 |
| 923 | 170.86 | 0.00000000 |
| 924 | 179.06 | 0.00000000 |

| | | |
|---|---|---|
| 925 | 201.37 | 0.00000000 |
| 926 | 177.47 | 0.00000000 |
| 927 | 181.18 | 0.00000000 |
| 928 | 213.71 | 0.00000000 |
| 929 | 168.14 | 0.00000000 |
| 930 | 226.24 | 0.00000000 |
| 931 | 187.97 | 0.00000000 |
| 932 | 159.94 | 0.00000000 |
| 933 | 188.91 | 0.00000000 |
| 934 | 192.21 | 0.00000000 |
| 935 | 167.89 | 0.00000000 |
| 936 | 161.05 | 0.00000000 |
| 937 | 188.14 | 0.00000000 |
| 938 | 165.15 | 0.00000000 |
| 939 | 166.56 | 0.00000000 |
| 940 | 204.15 | 0.00000000 |
| 941 | 177.14 | 0.00000000 |
| 942 | 182.03 | 0.00000000 |
| 943 | 198.56 | 0.00000000 |
| 944 | 174.74 | 0.00000000 |
| 945 | 196.31 | 0.00000000 |
| 946 | 171.94 | 0.00000000 |
| 947 | 176.72 | 0.00000000 |
| 948 | 195.42 | 0.00000000 |
| 949 | 198.22 | 0.00000000 |
| 950 | 167.48 | 0.00000000 |
| 951 | 181.16 | 0.00000000 |
| 952 | 196.76 | 0.00000000 |
| 953 | 184.48 | 0.00000000 |
| 954 | 159.69 | 0.00000000 |
| 955 | 160.24 | 0.00000000 |
| 956 | 154.24 | 0.00000000 |
| 957 | 164.21 | 0.00000000 |
| 958 | 190.88 | 0.00000000 |
| 959 | 184.07 | 0.00000000 |
| 960 | 192.58 | 0.00000000 |
| 961 | 197.50 | 0.00000000 |
| 962 | 188.97 | 0.00000000 |
| 963 | 214.58 | 0.00000000 |
| 964 | 171.43 | 0.00000000 |
| 965 | 170.62 | 0.00000000 |
| 966 | 166.50 | 0.00000000 |
| 967 | 190.91 | 0.00000000 |
| 968 | 196.71 | 0.00000000 |

| | | |
|---|---|---|
| 969 | 166.34 | 0.00000000 |
| 970 | 208.53 | 0.00000000 |
| 971 | 162.63 | 0.00000000 |
| 972 | 160.21 | 0.00000000 |
| 973 | 167.42 | 0.00000000 |
| 974 | 172.82 | 0.00000000 |
| 975 | 178.35 | 0.00000000 |
| 976 | 164.16 | 0.00000000 |
| 977 | 158.91 | 0.00000000 |
| 978 | 180.03 | 0.00000000 |
| 979 | 175.00 | 0.00000000 |
| 980 | 214.21 | 0.00000000 |
| 981 | 190.07 | 0.00000000 |
| 982 | 167.72 | 0.00000000 |
| 983 | 188.34 | 0.00000000 |
| 984 | 179.87 | 0.00000000 |
| 985 | 168.84 | 0.00000000 |
| 986 | 206.35 | 0.00000000 |
| 987 | 214.46 | 0.00000000 |
| 988 | 214.41 | 0.00000000 |
| 989 | 150.65 | 0.00000000 |
| 990 | 144.81 | 0.00000000 |
| 991 | 161.09 | 0.00000000 |
| 992 | 169.19 | 0.00000000 |
| 993 | 137.34 | 0.00000000 |
| 994 | 190.43 | 0.00000000 |
| 995 | 180.60 | 0.00000000 |
| 996 | 160.03 | 0.00000000 |
| 997 | 207.83 | 0.00000000 |
| 998 | 175.82 | 0.00000000 |
| 999 | 172.58 | 0.00000000 |
| 1000 | 167.71 | 0.00000000 |
| 1001 | 172.74 | 0.00000000 |
| 1002 | 174.55 | 0.00000000 |
| 1003 | 178.73 | 0.00000000 |
| 1004 | 138.80 | 0.00000000 |
| 1005 | 164.03 | 0.00000000 |
| 1006 | 169.16 | 0.00000000 |
| 1007 | 168.63 | 0.00000000 |
| 1008 | 167.94 | 0.00000000 |
| 1009 | 167.09 | 0.00000000 |
| 1010 | 161.10 | 0.00000000 |
| 1011 | 176.94 | 0.00000000 |
| 1012 | 162.52 | 0.00000000 |

| | | |
|---|---|---|
| 1013 | 165.52 | 0.00000000 |
| 1014 | 185.60 | 0.00000000 |
| 1015 | 151.27 | 0.00000000 |
| 1016 | 172.06 | 0.00000000 |
| 1017 | 165.31 | 0.00000000 |
| 1018 | 151.23 | 0.00000000 |
| 1019 | 151.58 | 0.00000000 |
| 1020 | 165.15 | 0.00000000 |
| 1021 | 138.81 | 0.00000000 |
| 1022 | 164.25 | 0.00000000 |
| 1023 | 168.30 | 0.00000000 |
| 1024 | 167.33 | 0.00000000 |
| 1025 | 163.80 | 0.00000000 |
| 1026 | 179.52 | 0.00000000 |
| 1027 | 169.00 | 0.00000000 |
| 1028 | 161.62 | 0.00000000 |
| 1029 | 176.90 | 0.00000000 |
| 1030 | 166.45 | 0.00000000 |
| 1031 | 150.47 | 0.00000000 |
| 1032 | 158.70 | 0.00000000 |
| 1033 | 159.81 | 0.00000000 |
| 1034 | 164.28 | 0.00000000 |
| 1035 | 168.93 | 0.00000000 |
| 1036 | 165.17 | 0.00000000 |
| 1037 | 155.19 | 0.00000000 |
| 1038 | 169.75 | 0.00000000 |
| 1039 | 152.97 | 0.00000000 |
| 1040 | 183.47 | 0.00000000 |
| 1041 | 144.65 | 0.00000000 |
| 1042 | 168.87 | 0.00000000 |
| 1043 | 154.76 | 0.00000000 |
| 1044 | 142.94 | 0.00000000 |
| 1045 | 144.61 | 0.00000000 |
| 1046 | 167.07 | 0.00000000 |
| 1047 | 194.14 | 0.00000000 |
| 1048 | 127.18 | 0.00000000 |
| 1049 | 153.38 | 0.00000000 |
| 1050 | 171.10 | 0.00000000 |
| 1051 | 146.22 | 0.00000000 |
| 1052 | 158.90 | 0.00000000 |
| 1053 | 150.27 | 0.00000000 |
| 1054 | 159.76 | 0.00000000 |
| 1055 | 164.03 | 0.00000000 |
| 1056 | 165.28 | 0.00000000 |

| | | |
|---|---|---|
| 1057 | 165.58 | 0.00000000 |
| 1058 | 145.32 | 0.00000000 |
| 1059 | 151.28 | 0.00000000 |
| 1060 | 133.14 | 0.00000000 |
| 1061 | 142.06 | 0.00000000 |
| 1062 | 154.29 | 0.00000000 |
| 1063 | 160.84 | 0.00000000 |
| 1064 | 145.80 | 0.00000000 |
| 1065 | 146.16 | 0.00000000 |
| 1066 | 190.41 | 0.00000000 |
| 1067 | 146.40 | 0.00000000 |
| 1068 | 148.97 | 0.00000000 |
| 1069 | 171.03 | 0.00000000 |
| 1070 | 147.59 | 0.00000000 |
| 1071 | 158.03 | 0.00000000 |
| 1072 | 157.18 | 0.00000000 |
| 1073 | 152.09 | 0.00000000 |
| 1074 | 131.56 | 0.00000000 |
| 1075 | 176.87 | 0.00000000 |
| 1076 | 162.21 | 0.00000000 |
| 1077 | 175.53 | 0.00000000 |
| 1078 | 152.84 | 0.00000000 |
| 1079 | 148.23 | 0.00000000 |
| 1080 | 132.09 | 0.00000000 |
| 1081 | 118.97 | 0.00000000 |
| 1082 | 152.69 | 0.00000000 |
| 1083 | 153.07 | 0.00000000 |
| 1084 | 141.63 | 0.00000000 |
| 1085 | 139.42 | 0.00000000 |
| 1086 | 128.56 | 0.00000000 |
| 1087 | 129.03 | 0.00000000 |
| 1088 | 157.35 | 0.00000000 |
| 1089 | 135.53 | 0.00000000 |
| 1090 | 138.06 | 0.00000000 |
| 1091 | 157.39 | 0.00000000 |
| 1092 | 147.42 | 0.00000000 |
| 1093 | 144.34 | 0.00000000 |
| 1094 | 145.09 | 0.00000000 |
| 1095 | 169.97 | 0.00000000 |
| 1096 | 164.97 | 0.00000000 |
| 1097 | 161.42 | 0.00000000 |
| 1098 | 147.06 | 0.00000000 |
| 1099 | 184.47 | 0.00000000 |
| 1100 | 140.16 | 0.00000000 |

| | | |
|---|---|---|
| 1101 | 117.15 | 0.00000000 |
| 1102 | 138.41 | 0.00000000 |
| 1103 | 149.58 | 0.00000000 |
| 1104 | 181.71 | 0.00000000 |
| 1105 | 158.58 | 0.00000000 |
| 1106 | 185.13 | 0.00000000 |
| 1107 | 123.15 | 0.00000000 |
| 1108 | 135.00 | 0.00000000 |
| 1109 | 141.13 | 0.00000000 |
| 1110 | 137.25 | 0.00000000 |
| 1111 | 116.94 | 0.00000000 |
| 1112 | 138.41 | 0.00000000 |
| 1113 | 137.19 | 0.00000000 |
| 1114 | 147.61 | 0.00000000 |
| 1115 | 133.32 | 0.00000000 |
| 1116 | 137.41 | 0.00000000 |
| 1117 | 136.29 | 0.00000000 |
| 1118 | 145.72 | 0.00000000 |
| 1119 | 131.00 | 0.00000000 |
| 1120 | 150.72 | 0.00000000 |
| 1121 | 130.29 | 0.00000000 |
| 1122 | 136.39 | 0.00000000 |
| 1123 | 147.34 | 0.00000000 |
| 1124 | 139.29 | 0.00000000 |
| 1125 | 189.79 | 0.00000000 |
| 1126 | 155.03 | 0.00000000 |
| 1127 | 158.00 | 0.00000000 |
| 1128 | 134.31 | 0.00000000 |
| 1129 | 146.93 | 0.00000000 |
| 1130 | 143.52 | 0.00000000 |
| 1131 | 136.88 | 0.00000000 |
| 1132 | 164.69 | 0.00000000 |
| 1133 | 119.08 | 0.00000000 |
| 1134 | 114.21 | 0.00000000 |
| 1135 | 118.70 | 0.00000000 |
| 1136 | 126.79 | 0.00000000 |
| 1137 | 110.89 | 0.00000000 |
| 1138 | 120.21 | 0.00000000 |
| 1139 | 130.59 | 0.00000000 |
| 1140 | 109.55 | 0.00000000 |
| 1141 | 108.82 | 0.00000000 |
| 1142 | 112.39 | 0.00000000 |
| 1143 | 121.21 | 0.00000000 |
| 1144 | 153.53 | 0.00000000 |

| | | |
|---|---|---|
| 1145 | 107.03 | 0.00000000 |
| 1146 | 136.70 | 0.00000000 |
| 1147 | 141.43 | 0.00000000 |
| 1148 | 135.41 | 0.00000000 |
| 1149 | 118.18 | 0.00000000 |
| 1150 | 101.97 | 0.00000000 |
| 1151 | 116.25 | 0.00000000 |
| 1152 | 150.41 | 0.00000000 |
| 1153 | 135.83 | 0.00000000 |
| 1154 | 117.66 | 0.00000000 |
| 1155 | 119.21 | 0.00000000 |
| 1156 | 136.73 | 0.00000000 |
| 1157 | 140.58 | 0.00000000 |
| 1158 | 125.06 | 0.00000000 |
| 1159 | 109.37 | 0.00000000 |
| 1160 | 141.90 | 0.00000000 |
| 1161 | 134.65 | 0.00000000 |
| 1162 | 128.90 | 0.00000000 |
| 1163 | 114.58 | 0.00000000 |
| 1164 | 163.17 | 0.00000000 |
| 1165 | 149.90 | 0.00000000 |
| 1166 | 118.77 | 0.00000000 |
| 1167 | 125.61 | 0.00000000 |
| 1168 | 127.62 | 0.00000000 |
| 1169 | 106.41 | 0.00000000 |
| 1170 | 121.26 | 0.00000000 |
| 1171 | 123.88 | 0.00000000 |
| 1172 | 109.61 | 0.00000000 |
| 1173 | 128.44 | 0.00000000 |
| 1174 | 124.37 | 0.00000000 |
| 1175 | 132.11 | 0.00000000 |
| 1176 | 132.67 | 0.00000000 |
| 1177 | 152.77 | 0.00000000 |
| 1178 | 133.57 | 0.00000000 |
| 1179 | 147.26 | 0.00000000 |
| 1180 | 134.59 | 0.00000000 |
| 1181 | 142.97 | 0.00000000 |
| 1182 | 113.16 | 0.00000000 |
| 1183 | 103.83 | 0.00000000 |
| 1184 | 118.44 | 0.00000000 |
| 1185 | 102.72 | 0.00000000 |
| 1186 | 118.69 | 0.00000000 |
| 1187 | 119.32 | 0.00000000 |
| 1188 | 103.58 | 0.00000000 |

| | | |
|---|---|---|
| 1189 | 121.83 | 0.00000000 |
| 1190 | 132.38 | 0.00000000 |
| 1191 | 112.77 | 0.00000000 |
| 1192 | 117.16 | 0.00000000 |
| 1193 | 129.81 | 0.00000000 |
| 1194 | 124.32 | 0.00000000 |
| 1195 | 102.16 | 0.00000000 |
| 1196 | 128.60 | 0.00000000 |
| 1197 | 105.16 | 0.00000000 |
| 1198 | 123.47 | 0.00000000 |
| 1199 | 118.18 | 0.00000000 |
| 1200 | 144.86 | 0.00000000 |
| 1201 | 132.61 | 0.00000000 |
| 1202 | 132.03 | 0.00000000 |
| 1203 | 127.42 | 0.00000000 |
| 1204 | 120.58 | 0.00000000 |
| 1205 | 139.57 | 0.00000000 |
| 1206 | 126.15 | 0.00000000 |
| 1207 | 115.77 | 0.00000000 |
| 1208 | 137.32 | 0.00000000 |
| 1209 | 151.41 | 0.00000000 |
| 1210 | 133.04 | 0.00000000 |
| 1211 | 132.32 | 0.00000000 |
| 1212 | 127.57 | 0.00000000 |
| 1213 | 131.67 | 0.00000000 |
| 1214 | 118.42 | 0.00000000 |
| 1215 | 132.74 | 0.00000000 |
| 1216 | 116.96 | 0.00000000 |
| 1217 | 135.16 | 0.00000000 |
| 1218 | 121.97 | 0.00000000 |
| 1219 | 120.97 | 0.00000000 |
| 1220 | 114.83 | 0.00000000 |
| 1221 | 116.76 | 0.00000000 |
| 1222 | 110.23 | 0.00000000 |
| 1223 | 118.30 | 0.00000000 |
| 1224 | 125.86 | 0.00000000 |
| 1225 | 112.72 | 0.00000000 |
| 1226 | 128.56 | 0.00000000 |
| 1227 | 136.00 | 0.00000000 |
| 1228 | 121.50 | 0.00000000 |
| 1229 | 126.84 | 0.00000000 |
| 1230 | 131.68 | 0.00000000 |
| 1231 | 118.13 | 0.00000000 |
| 1232 | 109.48 | 0.00000000 |

| | | |
|---|---|---|
| 1233 | 125.72 | 0.00000000 |
| 1234 | 106.17 | 0.00000000 |
| 1235 | 138.08 | 0.00000000 |
| 1236 | 163.44 | 0.00000000 |
| 1237 | 135.19 | 0.00000000 |
| 1238 | 108.86 | 0.00000000 |
| 1239 | 96.20 | 0.00000000 |
| 1240 | 112.48 | 0.00000000 |
| 1241 | 136.61 | 0.00000000 |
| 1242 | 120.97 | 0.00000000 |
| 1243 | 116.48 | 0.00000000 |
| 1244 | 102.19 | 0.00000000 |
| 1245 | 155.79 | 0.00000000 |
| 1246 | 129.07 | 0.00000000 |
| 1247 | 130.96 | 0.00000000 |
| 1248 | 128.61 | 0.00000000 |
| 1249 | 134.52 | 0.00000000 |
| 1250 | 111.09 | 0.00000000 |
| 1251 | 106.60 | 0.00000000 |
| 1252 | 109.41 | 0.00000000 |
| 1253 | 117.10 | 0.00000000 |
| 1254 | 111.38 | 0.00000000 |
| 1255 | 125.66 | 0.00000000 |
| 1256 | 103.38 | 0.00000000 |
| 1257 | 116.75 | 0.00000000 |
| 1258 | 88.58 | 0.00000000 |
| 1259 | 94.30 | 0.00000000 |
| 1260 | 115.46 | 0.00000000 |
| 1261 | 102.94 | 0.00000000 |
| 1262 | 129.24 | 0.00000000 |
| 1263 | 117.62 | 0.00000000 |
| 1264 | 110.50 | 0.00000000 |
| 1265 | 116.10 | 0.00000000 |
| 1266 | 124.11 | 0.00000000 |
| 1267 | 103.69 | 0.00000000 |
| 1268 | 104.44 | 0.00000000 |
| 1269 | 126.67 | 0.00000000 |
| 1270 | 129.63 | 0.00000000 |
| 1271 | 132.43 | 0.00000000 |
| 1272 | 128.83 | 0.00000000 |
| 1273 | 102.33 | 0.00000000 |
| 1274 | 118.63 | 0.00000000 |
| 1275 | 138.17 | 0.00000000 |
| 1276 | 96.50 | 0.00000000 |

| | | |
|---|---|---|
| 1277 | 129.28 | 0.00000000 |
| 1278 | 137.52 | 0.00000000 |
| 1279 | 116.96 | 0.00000000 |
| 1280 | 110.89 | 0.00000000 |
| 1281 | 112.33 | 0.00000000 |
| 1282 | 142.76 | 0.00000000 |
| 1283 | 135.56 | 0.00000000 |
| 1284 | 100.97 | 0.00000000 |
| 1285 | 101.13 | 0.00000000 |
| 1286 | 101.78 | 0.00000000 |
| 1287 | 76.38 | 0.00000000 |
| 1288 | 115.19 | 0.00000000 |
| 1289 | 129.50 | 0.00000000 |
| 1290 | 122.93 | 0.00000000 |
| 1291 | 122.48 | 0.00000000 |
| 1292 | 117.89 | 0.00000000 |
| 1293 | 103.83 | 0.00000000 |
| 1294 | 108.94 | 0.00000000 |
| 1295 | 99.82 | 0.00000000 |
| 1296 | 129.34 | 0.00000000 |
| 1297 | 113.93 | 0.00000000 |
| 1298 | 136.15 | 0.00000000 |
| 1299 | 118.78 | 0.00000000 |
| 1300 | 108.87 | 0.00000000 |
| 1301 | 101.69 | 0.00000000 |
| 1302 | 107.59 | 0.00000000 |
| 1303 | 117.34 | 0.00000000 |
| 1304 | 114.77 | 0.00000000 |
| 1305 | 121.60 | 0.00000000 |
| 1306 | 120.19 | 0.00000000 |
| 1307 | 111.21 | 0.00000000 |
| 1308 | 122.32 | 0.00000000 |
| 1309 | 110.17 | 0.00000000 |
| 1310 | 128.47 | 0.00000000 |
| 1311 | 128.36 | 0.00000000 |
| 1312 | 130.58 | 0.00000000 |
| 1313 | 140.00 | 0.00000000 |
| 1314 | 111.42 | 0.00000000 |
| 1315 | 102.14 | 0.00000000 |
| 1316 | 112.25 | 0.00000000 |
| 1317 | 91.00 | 0.00000000 |
| 1318 | 99.62 | 0.00000000 |
| 1319 | 101.81 | 0.00000000 |
| 1320 | 94.00 | 0.00000000 |

| | | |
|---|---|---|
| 1321 | 114.68 | 0.00000000 |
| 1322 | 106.97 | 0.00000000 |
| 1323 | 105.33 | 0.00000000 |
| 1324 | 107.90 | 0.00000000 |
| 1325 | 99.31 | 0.00000000 |
| 1326 | 104.76 | 0.00000000 |
| 1327 | 111.32 | 0.00000000 |
| 1328 | 100.83 | 0.00000000 |
| 1329 | 83.93 | 0.00000000 |
| 1330 | 102.77 | 0.00000000 |
| 1331 | 104.21 | 0.00000000 |
| 1332 | 101.73 | 0.00000000 |
| 1333 | 108.43 | 0.00000000 |
| 1334 | 104.59 | 0.00000000 |
| 1335 | 104.43 | 0.00000000 |
| 1336 | 102.33 | 0.00000000 |
| 1337 | 121.42 | 0.00000000 |
| 1338 | 121.64 | 0.00000000 |
| 1339 | 91.50 | 0.00000000 |
| 1340 | 85.40 | 0.00000000 |
| 1341 | 93.29 | 0.00000000 |
| 1342 | 80.10 | 0.00000000 |
| 1343 | 93.94 | 0.00000000 |
| 1344 | 95.83 | 0.00000000 |
| 1345 | 93.42 | 0.00000000 |
| 1346 | 84.75 | 0.00000000 |
| 1347 | 86.28 | 0.00000000 |
| 1348 | 122.23 | 0.00000000 |
| 1349 | 107.90 | 0.00000000 |
| 1350 | 114.11 | 0.00000000 |
| 1351 | 102.61 | 0.00000000 |
| 1352 | 114.58 | 0.00000000 |
| 1353 | 95.00 | 0.00000000 |
| 1354 | 104.28 | 0.00000000 |
| 1355 | 91.38 | 0.00000000 |
| 1356 | 83.17 | 0.00000000 |
| 1357 | 97.89 | 0.00000000 |
| 1358 | 99.56 | 0.00000000 |
| 1359 | 112.12 | 0.00000000 |
| 1360 | 104.24 | 0.00000000 |
| 1361 | 120.75 | 0.00000000 |
| 1362 | 115.48 | 0.00000000 |
| 1363 | 128.11 | 0.00000000 |
| 1364 | 129.73 | 0.00000000 |

| | | |
|---|---|---|
| 1365 | 116.96 | 0.00000000 |
| 1366 | 93.88 | 0.00000000 |
| 1367 | 143.12 | 0.00000000 |
| 1368 | 107.69 | 0.00000000 |
| 1369 | 115.46 | 0.00000000 |
| 1370 | 116.00 | 0.00000000 |
| 1371 | 108.28 | 0.00000000 |
| 1372 | 111.28 | 0.00000000 |
| 1373 | 105.33 | 0.00000000 |
| 1374 | 75.77 | 0.00000000 |
| 1375 | 88.17 | 0.00000000 |
| 1376 | 94.50 | 0.00000000 |
| 1377 | 97.10 | 0.00000000 |
| 1378 | 93.36 | 0.00000000 |
| 1379 | 101.19 | 0.00000000 |
| 1380 | 95.59 | 0.00000000 |
| 1381 | 89.42 | 0.00000000 |
| 1382 | 93.21 | 0.00000000 |
| 1383 | 99.23 | 0.00000000 |
| 1384 | 95.70 | 0.00000000 |
| 1385 | 94.21 | 0.00000000 |
| 1386 | 81.06 | 0.00000000 |
| 1387 | 94.66 | 0.00000000 |
| 1388 | 93.36 | 0.00000000 |
| 1389 | 101.20 | 0.00000000 |
| 1390 | 90.79 | 0.00000000 |
| 1391 | 105.44 | 0.00000000 |
| 1392 | 85.28 | 0.00000000 |
| 1393 | 104.07 | 0.00000000 |
| 1394 | 95.31 | 0.00000000 |
| 1395 | 87.62 | 0.00000000 |
| 1396 | 98.85 | 0.00000000 |
| 1397 | 107.81 | 0.00000000 |
| 1398 | 117.96 | 0.00000000 |
| 1399 | 92.68 | 0.00000000 |
| 1400 | 130.44 | 0.00000000 |
| 1401 | 131.08 | 0.00000000 |
| 1402 | 106.69 | 0.00000000 |
| 1403 | 98.07 | 0.00000000 |
| 1404 | 89.87 | 0.00000000 |
| 1405 | 94.59 | 0.00000000 |
| 1406 | 102.86 | 0.00000000 |
| 1407 | 99.67 | 0.00000000 |
| 1408 | 99.63 | 0.00000000 |

| | | |
|---|---|---|
| 1409 | 86.10 | 0.00000000 |
| 1410 | 112.26 | 0.00000000 |
| 1411 | 125.54 | 0.00000000 |
| 1412 | 111.84 | 0.00000000 |
| 1413 | 101.12 | 0.00000000 |
| 1414 | 99.11 | 0.00000000 |
| 1415 | 98.96 | 0.00000000 |
| 1416 | 109.69 | 0.00000000 |
| 1417 | 108.54 | 0.00000000 |
| 1418 | 105.04 | 0.00000000 |
| 1419 | 122.52 | 0.00000000 |
| 1420 | 85.85 | 0.00000000 |
| 1421 | 98.61 | 0.00000000 |
| 1422 | 101.58 | 0.00000000 |
| 1423 | 94.54 | 0.00000000 |
| 1424 | 93.68 | 0.00000000 |
| 1425 | 79.07 | 0.00000000 |
| 1426 | 89.39 | 0.00000000 |
| 1427 | 100.90 | 0.00000000 |
| 1428 | 90.88 | 0.00000000 |
| 1429 | 71.69 | 0.00000000 |
| 1430 | 93.04 | 0.00000000 |
| 1431 | 75.80 | 0.00000000 |
| 1432 | 93.86 | 0.00000000 |
| 1433 | 92.86 | 0.00000000 |
| 1434 | 102.33 | 0.00000000 |
| 1435 | 119.71 | 0.00000000 |
| 1436 | 83.73 | 0.00000000 |
| 1437 | 76.29 | 0.00000000 |
| 1438 | 79.39 | 0.00000000 |
| 1439 | 94.56 | 0.00000000 |
| 1440 | 84.96 | 0.00000000 |
| 1441 | 89.90 | 0.00000000 |
| 1442 | 98.72 | 0.00000000 |
| 1443 | 96.68 | 0.00000000 |
| 1444 | 126.68 | 0.00000000 |
| 1445 | 86.29 | 0.00000000 |
| 1446 | 85.17 | 0.00000000 |
| 1447 | 102.23 | 0.00000000 |
| 1448 | 73.69 | 0.00000000 |
| 1449 | 90.19 | 0.00000000 |
| 1450 | 104.19 | 0.00000000 |
| 1451 | 103.37 | 0.00000000 |
| 1452 | 78.03 | 0.00000000 |

| | | |
|---|---|---|
| 1453 | 83.90 | 0.00000000 |
| 1454 | 87.24 | 0.00000000 |
| 1455 | 109.78 | 0.00000000 |
| 1456 | 110.23 | 0.00000000 |
| 1457 | 103.33 | 0.00000000 |
| 1458 | 93.24 | 0.00000000 |
| 1459 | 97.82 | 0.00000000 |
| 1460 | 95.61 | 0.00000000 |
| 1461 | 97.00 | 0.00000000 |
| 1462 | 111.04 | 0.00000000 |
| 1463 | 123.71 | 0.00000000 |
| 1464 | 109.82 | 0.00000000 |
| 1465 | 100.33 | 0.00000000 |
| 1466 | 92.04 | 0.00000000 |
| 1467 | 107.32 | 0.00000000 |
| 1468 | 94.39 | 0.00000000 |
| 1469 | 77.92 | 0.00000000 |
| 1470 | 88.52 | 0.00000000 |
| 1471 | 68.81 | 0.00000000 |
| 1472 | 82.83 | 0.00000000 |
| 1473 | 81.18 | 0.00000000 |
| 1474 | 92.61 | 0.00000000 |
| 1475 | 79.59 | 0.00000000 |
| 1476 | 78.43 | 0.00000000 |
| 1477 | 92.40 | 0.00000000 |
| 1478 | 89.52 | 0.00000000 |
| 1479 | 84.75 | 0.00000000 |
| 1480 | 100.19 | 0.00000000 |
| 1481 | 72.31 | 0.00000000 |
| 1482 | 97.42 | 0.00000000 |
| 1483 | 66.20 | 0.00000000 |
| 1484 | 95.29 | 0.00000000 |
| 1485 | 98.14 | 0.00000000 |
| 1486 | 82.21 | 0.00000000 |
| 1487 | 79.57 | 0.00000000 |
| 1488 | 82.70 | 0.00000000 |
| 1489 | 63.82 | 0.00000000 |
| 1490 | 103.52 | 0.00000000 |
| 1491 | 132.22 | 0.00000000 |
| 1492 | 91.35 | 0.00000000 |
| 1493 | 104.52 | 0.00000000 |
| 1494 | 85.19 | 0.00000000 |
| 1495 | 95.56 | 0.00000000 |
| 1496 | 68.14 | 0.00000000 |

| | | |
|---|---|---|
| 1497 | 86.28 | 0.00000000 |
| 1498 | 89.58 | 0.00000000 |
| 1499 | 79.18 | 0.00000000 |
| 1500 | 63.68 | 0.00000000 |
| 1501 | 78.80 | 0.00000000 |
| 1502 | 72.81 | 0.00000000 |
| 1503 | 88.04 | 0.00000000 |
| 1504 | 93.24 | 0.00000000 |
| 1505 | 80.96 | 0.00000000 |
| 1506 | 96.79 | 0.00000000 |
| 1507 | 90.44 | 0.00000000 |
| 1508 | 84.71 | 0.00000000 |
| 1509 | 84.96 | 0.00000000 |
| 1510 | 83.00 | 0.00000000 |
| 1511 | 74.93 | 0.00000000 |
| 1512 | 82.96 | 0.00000000 |
| 1513 | 80.73 | 0.00000000 |
| 1514 | 87.87 | 0.00000000 |
| 1515 | 91.88 | 0.00000000 |
| 1516 | 99.26 | 0.00000000 |
| 1517 | 93.67 | 0.00000000 |
| 1518 | 101.59 | 0.00000000 |
| 1519 | 104.46 | 0.00000000 |
| 1520 | 90.78 | 0.00000000 |
| 1521 | 80.21 | 0.00000000 |
| 1522 | 116.10 | 0.00000000 |
| 1523 | 104.61 | 0.00000000 |
| 1524 | 61.29 | 0.00000000 |
| 1525 | 97.27 | 0.00000000 |
| 1526 | 94.54 | 0.00000000 |
| 1527 | 88.71 | 0.00000000 |
| 1528 | 92.68 | 0.00000000 |
| 1529 | 89.42 | 0.00000000 |
| 1530 | 87.15 | 0.00000000 |
| 1531 | 85.82 | 0.00000000 |
| 1532 | 86.14 | 0.00000000 |
| 1533 | 102.28 | 0.00000000 |
| 1534 | 69.07 | 0.00000000 |
| 1535 | 81.88 | 0.00000000 |
| 1536 | 82.73 | 0.00000000 |
| 1537 | 83.60 | 0.00000000 |
| 1538 | 93.82 | 0.00000000 |
| 1539 | 92.12 | 0.00000000 |
| 1540 | 84.42 | 0.00000000 |

| | | |
|---|---|---|
| 1541 | 89.35 | 0.00000000 |
| 1542 | 90.75 | 0.00000000 |
| 1543 | 81.50 | 0.00000000 |
| 1544 | 80.72 | 0.00000000 |
| 1545 | 72.88 | 0.00000000 |
| 1546 | 89.19 | 0.00000000 |
| 1547 | 91.68 | 0.00000000 |
| 1548 | 64.35 | 0.00000000 |
| 1549 | 99.56 | 0.00000000 |
| 1550 | 100.18 | 0.00000000 |
| 1551 | 83.64 | 0.00000000 |
| 1552 | 96.76 | 0.00000000 |
| 1553 | 87.81 | 0.00000000 |
| 1554 | 84.96 | 0.00000000 |
| 1555 | 103.62 | 0.00000000 |
| 1556 | 92.15 | 0.00000000 |
| 1557 | 101.83 | 0.00000000 |
| 1558 | 78.96 | 0.00000000 |
| 1559 | 80.89 | 0.00000000 |
| 1560 | 75.00 | 0.00000000 |
| 1561 | 86.56 | 0.00000000 |
| 1562 | 82.27 | 0.00000000 |
| 1563 | 83.58 | 0.00000000 |
| 1564 | 79.38 | 0.00000000 |
| 1565 | 84.65 | 0.00000000 |
| 1566 | 77.88 | 0.00000000 |
| 1567 | 96.50 | 0.00000000 |
| 1568 | 82.88 | 0.00000000 |
| 1569 | 64.30 | 0.00000000 |
| 1570 | 76.59 | 0.00000000 |
| 1571 | 83.57 | 0.00000000 |
| 1572 | 84.08 | 0.00000000 |
| 1573 | 74.86 | 0.00000000 |
| 1574 | 73.43 | 0.00000000 |
| 1575 | 76.80 | 0.00000000 |
| 1576 | 75.11 | 0.00000000 |
| 1577 | 76.56 | 0.00000000 |
| 1578 | 83.67 | 0.00000000 |
| 1579 | 77.88 | 0.00000000 |
| 1580 | 72.75 | 0.00000000 |
| 1581 | 64.29 | 0.00000000 |
| 1582 | 59.97 | 0.00000000 |
| 1583 | 75.07 | 0.00000000 |
| 1584 | 90.57 | 0.00000000 |

| | | |
|---|---|---|
| 1585 | 93.25 | 0.00000000 |
| 1586 | 88.68 | 0.00000000 |
| 1587 | 83.04 | 0.00000000 |
| 1588 | 74.55 | 0.00000000 |
| 1589 | 71.24 | 0.00000000 |
| 1590 | 84.36 | 0.00000000 |
| 1591 | 85.42 | 0.00000000 |
| 1592 | 67.21 | 0.00000000 |
| 1593 | 87.77 | 0.00000000 |
| 1594 | 52.41 | 0.00000000 |
| 1595 | 79.43 | 0.00000000 |
| 1596 | 77.93 | 0.00000000 |
| 1597 | 74.45 | 0.00000000 |
| 1598 | 64.84 | 0.00000000 |
| 1599 | 92.15 | 0.00000000 |
| 1600 | 58.43 | 0.00000000 |
| 1601 | 65.88 | 0.00000000 |
| 1602 | 86.81 | 0.00000000 |
| 1603 | 101.00 | 0.00000000 |
| 1604 | 92.77 | 0.00000000 |
| 1605 | 66.50 | 0.00000000 |
| 1606 | 84.43 | 0.00000000 |
| 1607 | 62.22 | 0.00000000 |
| 1608 | 62.21 | 0.00000000 |
| 1609 | 84.93 | 0.00000000 |
| 1610 | 61.04 | 0.00000000 |
| 1611 | 83.68 | 0.00000000 |
| 1612 | 89.76 | 0.00000000 |
| 1613 | 61.59 | 0.00000000 |
| 1614 | 81.96 | 0.00000000 |
| 1615 | 58.66 | 0.00000000 |
| 1616 | 78.38 | 0.00000000 |
| 1617 | 85.19 | 0.00000000 |
| 1618 | 77.12 | 0.00000000 |
| 1619 | 73.00 | 0.00000000 |
| 1620 | 74.44 | 0.00000000 |
| 1621 | 52.77 | 0.00000000 |
| 1622 | 83.26 | 0.00000000 |
| 1623 | 71.08 | 0.00000000 |
| 1624 | 82.32 | 0.00000000 |
| 1625 | 70.67 | 0.00000000 |
| 1626 | 89.44 | 0.00000000 |
| 1627 | 74.93 | 0.00000000 |
| 1628 | 80.08 | 0.00000000 |

| | | |
|---|---|---|
| 1629 | 50.00 | 0.00000000 |
| 1630 | 101.80 | 0.00000000 |
| 1631 | 71.19 | 0.00000000 |
| 1632 | 74.96 | 0.00000000 |
| 1633 | 66.62 | 0.00000000 |
| 1634 | 65.75 | 0.00000000 |
| 1635 | 70.45 | 0.00000000 |
| 1636 | 69.52 | 0.00000000 |
| 1637 | 70.96 | 0.00000000 |
| 1638 | 72.25 | 0.00000000 |
| 1639 | 77.30 | 0.00000000 |
| 1640 | 79.71 | 0.00000000 |
| 1641 | 83.75 | 0.00000000 |
| 1642 | 74.56 | 0.00000000 |
| 1643 | 69.18 | 0.00000000 |
| 1644 | 71.00 | 0.00000000 |
| 1645 | 75.16 | 0.00000000 |
| 1646 | 61.16 | 0.00000000 |
| 1647 | 67.88 | 0.00000000 |
| 1648 | 97.00 | 0.00000000 |
| 1649 | 68.88 | 0.00000000 |
| 1650 | 75.77 | 0.00000000 |
| 1651 | 68.72 | 0.00000000 |
| 1652 | 82.56 | 0.00000000 |
| 1653 | 76.74 | 0.00000000 |
| 1654 | 73.35 | 0.00000000 |
| 1655 | 70.30 | 0.00000000 |
| 1656 | 85.36 | 0.00000000 |
| 1657 | 95.92 | 0.00000000 |
| 1658 | 59.38 | 0.00000000 |
| 1659 | 81.36 | 0.00000000 |
| 1660 | 58.23 | 0.00000000 |
| 1661 | 73.17 | 0.00000000 |
| 1662 | 87.14 | 0.00000000 |
| 1663 | 56.43 | 0.00000000 |
| 1664 | 55.69 | 0.00000000 |
| 1665 | 79.41 | 0.00000000 |
| 1666 | 85.54 | 0.00000000 |
| 1667 | 64.48 | 0.00000000 |
| 1668 | 70.83 | 0.00000000 |
| 1669 | 75.82 | 0.00000000 |
| 1670 | 78.40 | 0.00000000 |
| 1671 | 59.59 | 0.00000000 |
| 1672 | 86.95 | 0.00000000 |

| | | |
|---|---|---|
| 1673 | 89.64 | 0.00000000 |
| 1674 | 82.62 | 0.00000000 |
| 1675 | 73.23 | 0.00000000 |
| 1676 | 64.40 | 0.00000000 |
| 1677 | 71.65 | 0.00000000 |
| 1678 | 75.70 | 0.00000000 |
| 1679 | 88.64 | 0.00000000 |
| 1680 | 62.50 | 0.00000000 |
| 1681 | 53.85 | 0.00000000 |
| 1682 | 75.04 | 0.00000000 |
| 1683 | 88.26 | 0.00000000 |
| 1684 | 67.54 | 0.00000000 |
| 1685 | 70.04 | 0.00000000 |
| 1686 | 73.36 | 0.00000000 |
| 1687 | 61.93 | 0.00000000 |
| 1688 | 60.85 | 0.00000000 |
| 1689 | 74.04 | 0.00000000 |
| 1690 | 82.17 | 0.00000000 |
| 1691 | 72.88 | 0.00000000 |
| 1692 | 72.76 | 0.00000000 |
| 1693 | 106.24 | 0.00000000 |
| 1694 | 54.25 | 0.00000000 |
| 1695 | 73.79 | 0.00000000 |
| 1696 | 71.88 | 0.00000000 |
| 1697 | 57.07 | 0.00000000 |
| 1698 | 54.54 | 0.00000000 |
| 1699 | 64.63 | 0.00000000 |
| 1700 | 68.39 | 0.00000000 |
| 1701 | 75.29 | 0.00000000 |
| 1702 | 72.87 | 0.00000000 |
| 1703 | 60.17 | 0.00000000 |
| 1704 | 68.85 | 0.00000000 |
| 1705 | 52.27 | 0.00000000 |
| 1706 | 84.13 | 0.00000000 |
| 1707 | 75.96 | 0.00000000 |
| 1708 | 74.21 | 0.00000000 |
| 1709 | 66.25 | 0.00000000 |
| 1710 | 60.55 | 0.00000000 |
| 1711 | 87.13 | 0.00000000 |
| 1712 | 82.08 | 0.00000000 |
| 1713 | 83.48 | 0.00000000 |
| 1714 | 70.60 | 0.00000000 |
| 1715 | 76.00 | 0.00000000 |
| 1716 | 64.88 | 0.00000000 |

| | | |
|---|---|---|
| 1717 | 78.13 | 0.00000000 |
| 1718 | 87.78 | 0.00000000 |
| 1719 | 72.08 | 0.00000000 |
| 1720 | 66.77 | 0.00000000 |
| 1721 | 73.79 | 0.00000000 |
| 1722 | 69.24 | 0.00000000 |
| 1723 | 64.45 | 0.00000000 |
| 1724 | 87.16 | 0.00000000 |
| 1725 | 67.44 | 0.00000000 |
| 1726 | 89.85 | 0.00000000 |
| 1727 | 72.36 | 0.00000000 |
| 1728 | 86.13 | 0.00000000 |
| 1729 | 65.92 | 0.00000000 |
| 1730 | 52.92 | 0.00000000 |
| 1731 | 77.20 | 0.00000000 |
| 1732 | 67.28 | 0.00000000 |
| 1733 | 86.71 | 0.00000000 |
| 1734 | 62.19 | 0.00000000 |
| 1735 | 58.86 | 0.00000000 |
| 1736 | 63.15 | 0.00000000 |
| 1737 | 70.09 | 0.00000000 |
| 1738 | 62.12 | 0.00000000 |
| 1739 | 59.39 | 0.00000000 |
| 1740 | 46.04 | 0.00000000 |
| 1741 | 61.63 | 0.00000000 |
| 1742 | 61.70 | 0.00000000 |
| 1743 | 52.13 | 0.00000000 |
| 1744 | 73.26 | 0.00000000 |
| 1745 | 79.91 | 0.00000000 |
| 1746 | 76.92 | 0.00000000 |
| 1747 | 65.74 | 0.00000000 |
| 1748 | 76.70 | 0.00000000 |
| 1749 | 67.91 | 0.00000000 |
| 1750 | 53.62 | 0.00000000 |
| 1751 | 49.00 | 0.00000000 |
| 1752 | 54.77 | 0.00000000 |
| 1753 | 47.93 | 0.00000000 |
| 1754 | 56.37 | 0.00000000 |
| 1755 | 80.11 | 0.00000000 |
| 1756 | 58.85 | 0.00000000 |
| 1757 | 64.82 | 0.00000000 |
| 1758 | 51.18 | 0.00000000 |
| 1759 | 52.69 | 0.00000000 |
| 1760 | 47.67 | 0.00000000 |

| | | |
|---|---|---|
| 1761 | 63.55 | 0.00000000 |
| 1762 | 75.28 | 0.00000000 |
| 1763 | 73.60 | 0.00000000 |
| 1764 | 76.82 | 0.00000000 |
| 1765 | 70.58 | 0.00000000 |
| 1766 | 89.35 | 0.00000000 |
| 1767 | 68.00 | 0.00000000 |
| 1768 | 71.17 | 0.00000000 |
| 1769 | 72.08 | 0.00000000 |
| 1770 | 67.57 | 0.00000000 |
| 1771 | 65.59 | 0.00000000 |
| 1772 | 76.05 | 0.00000000 |
| 1773 | 63.59 | 0.00000000 |
| 1774 | 88.90 | 0.00000000 |
| 1775 | 86.62 | 0.00000000 |
| 1776 | 95.71 | 0.00000000 |
| 1777 | 86.23 | 0.00000000 |
| 1778 | 60.87 | 0.00000000 |
| 1779 | 77.64 | 0.00000000 |
| 1780 | 46.23 | 0.00000000 |
| 1781 | 62.65 | 0.00000000 |
| 1782 | 63.70 | 0.00000000 |
| 1783 | 65.84 | 0.00000000 |
| 1784 | 69.80 | 0.00000000 |
| 1785 | 63.43 | 0.00000000 |
| 1786 | 80.24 | 0.00000000 |
| 1787 | 48.80 | 0.00000000 |
| 1788 | 62.00 | 0.00000000 |
| 1789 | 59.92 | 0.00000000 |
| 1790 | 54.57 | 0.00000000 |
| 1791 | 54.68 | 0.00000000 |
| 1792 | 53.28 | 0.00000000 |
| 1793 | 57.88 | 0.00000000 |
| 1794 | 58.19 | 0.00000000 |
| 1795 | 49.35 | 0.00000000 |
| 1796 | 61.29 | 0.00000000 |
| 1797 | 55.41 | 0.00000000 |
| 1798 | 69.20 | 0.00000000 |
| 1799 | 47.68 | 0.00000000 |
| 1800 | 51.16 | 0.00000000 |
| 1801 | 59.15 | 0.00000000 |
| 1802 | 62.09 | 0.00000000 |
| 1803 | 64.36 | 0.00000000 |
| 1804 | 55.17 | 0.00000000 |

| | | |
|---|---|---|
| 1805 | 50.50 | 0.00000000 |
| 1806 | 62.66 | 0.00000000 |
| 1807 | 47.55 | 0.00000000 |
| 1808 | 64.42 | 0.00000000 |
| 1809 | 55.71 | 0.00000000 |
| 1810 | 77.27 | 0.00000000 |
| 1811 | 62.58 | 0.00000000 |
| 1812 | 51.96 | 0.00000000 |
| 1813 | 75.13 | 0.00000000 |
| 1814 | 65.96 | 0.00000000 |
| 1815 | 71.50 | 0.00000000 |
| 1816 | 73.08 | 0.00000000 |
| 1817 | 51.45 | 0.00000000 |
| 1818 | 54.39 | 0.00000000 |
| 1819 | 46.81 | 0.00000000 |
| 1820 | 64.56 | 0.00000000 |
| 1821 | 62.30 | 0.00000000 |
| 1822 | 58.42 | 0.00000000 |
| 1823 | 62.41 | 0.00000000 |
| 1824 | 53.20 | 0.00000000 |
| 1825 | 65.11 | 0.00000000 |
| 1826 | 62.27 | 0.00000000 |
| 1827 | 55.08 | 0.00000000 |
| 1828 | 53.54 | 0.00000000 |
| 1829 | 62.37 | 0.00000000 |
| 1830 | 62.00 | 0.00000000 |
| 1831 | 53.17 | 0.00000000 |
| 1832 | 62.33 | 0.00000000 |
| 1833 | 62.45 | 0.00000000 |
| 1834 | 39.96 | 0.00000000 |
| 1835 | 63.50 | 0.00000000 |
| 1836 | 69.04 | 0.00000000 |
| 1837 | 53.22 | 0.00000000 |
| 1838 | 49.00 | 0.00000000 |
| 1839 | 69.16 | 0.00000000 |
| 1840 | 63.16 | 0.00000000 |
| 1841 | 72.78 | 0.00000000 |
| 1842 | 61.83 | 0.00000000 |
| 1843 | 65.44 | 0.00000000 |
| 1844 | 41.79 | 0.00000000 |
| 1845 | 53.59 | 0.00000000 |
| 1846 | 46.78 | 0.00000000 |
| 1847 | 52.31 | 0.00000000 |
| 1848 | 48.48 | 0.00000000 |

| | | |
|---|---|---|
| 1849 | 61.52 | 0.00000000 |
| 1850 | 83.25 | 0.00000000 |
| 1851 | 54.92 | 0.00000000 |
| 1852 | 62.17 | 0.00000000 |
| 1853 | 59.25 | 0.00000000 |
| 1854 | 54.85 | 0.00000000 |
| 1855 | 52.50 | 0.00000000 |
| 1856 | 69.39 | 0.00000000 |
| 1857 | 65.29 | 0.00000000 |
| 1858 | 50.74 | 0.00000000 |
| 1859 | 58.72 | 0.00000000 |
| 1860 | 56.79 | 0.00000000 |
| 1861 | 53.12 | 0.00000000 |
| 1862 | 54.78 | 0.00000000 |
| 1863 | 68.09 | 0.00000000 |
| 1864 | 72.71 | 0.00000000 |
| 1865 | 54.68 | 0.00000000 |
| 1866 | 49.35 | 0.00000000 |
| 1867 | 45.56 | 0.00000000 |
| 1868 | 54.71 | 0.00000000 |
| 1869 | 56.48 | 0.00000000 |
| 1870 | 63.92 | 0.00000000 |
| 1871 | 78.09 | 0.00000000 |
| 1872 | 58.25 | 0.00000000 |
| 1873 | 50.46 | 0.00000000 |
| 1874 | 40.67 | 0.00000000 |
| 1875 | 57.63 | 0.00000000 |
| 1876 | 55.76 | 0.00000000 |
| 1877 | 65.58 | 0.00000000 |
| 1878 | 42.13 | 0.00000000 |
| 1879 | 38.61 | 0.00000000 |
| 1880 | 68.80 | 0.00000000 |
| 1881 | 53.44 | 0.00000000 |
| 1882 | 54.75 | 0.00000000 |
| 1883 | 58.62 | 0.00000000 |
| 1884 | 54.70 | 0.00000000 |
| 1885 | 69.52 | 0.00000000 |
| 1886 | 61.29 | 0.00000000 |
| 1887 | 76.40 | 0.00000000 |
| 1888 | 62.40 | 0.00000000 |
| 1889 | 71.05 | 0.00000000 |
| 1890 | 69.05 | 0.00000000 |
| 1891 | 72.26 | 0.00000000 |
| 1892 | 74.16 | 0.00000000 |

| | | |
|---|---|---|
| 1893 | 67.17 | 0.00000000 |
| 1894 | 85.23 | 0.00000000 |
| 1895 | 40.65 | 0.00000000 |
| 1896 | 52.50 | 0.00000000 |
| 1897 | 55.38 | 0.00000000 |
| 1898 | 54.41 | 0.00000000 |
| 1899 | 49.20 | 0.00000000 |
| 1900 | 59.79 | 0.00000000 |
| 1901 | 46.96 | 0.00000000 |
| 1902 | 65.91 | 0.00000000 |
| 1903 | 41.54 | 0.00000000 |
| 1904 | 57.67 | 0.00000000 |
| 1905 | 67.45 | 0.00000000 |
| 1906 | 55.00 | 0.00000000 |
| 1907 | 69.23 | 0.00000000 |
| 1908 | 54.52 | 0.00000000 |
| 1909 | 60.83 | 0.00000000 |
| 1910 | 65.13 | 0.00000000 |
| 1911 | 44.28 | 0.00000000 |
| 1912 | 49.04 | 0.00000000 |
| 1913 | 51.25 | 0.00000000 |
| 1914 | 73.22 | 0.00000000 |
| 1915 | 57.52 | 0.00000000 |
| 1916 | 56.88 | 0.00000000 |
| 1917 | 73.50 | 0.00000000 |
| 1918 | 51.00 | 0.00000000 |
| 1919 | 52.91 | 0.00000000 |
| 1920 | 46.27 | 0.00000000 |
| 1921 | 44.10 | 0.00000000 |
| 1922 | 50.36 | 0.00000000 |
| 1923 | 58.35 | 0.00000000 |
| 1924 | 53.80 | 0.00000000 |
| 1925 | 65.74 | 0.00000000 |
| 1926 | 54.74 | 0.00000000 |
| 1927 | 63.33 | 0.00000000 |
| 1928 | 59.41 | 0.00000000 |
| 1929 | 49.82 | 0.00000000 |
| 1930 | 58.14 | 0.00000000 |
| 1931 | 49.89 | 0.00000000 |
| 1932 | 65.65 | 0.00000000 |
| 1933 | 53.50 | 0.00000000 |
| 1934 | 56.28 | 0.00000000 |
| 1935 | 61.09 | 0.00000000 |
| 1936 | 58.24 | 0.00000000 |

| | | |
|---|---|---|
| 1937 | 50.37 | 0.00000000 |
| 1938 | 55.04 | 0.00000000 |
| 1939 | 51.83 | 0.00000000 |
| 1940 | 44.50 | 0.00000000 |
| 1941 | 54.23 | 0.00000000 |
| 1942 | 56.72 | 0.00000000 |
| 1943 | 41.11 | 0.00000000 |
| 1944 | 50.75 | 0.00000000 |
| 1945 | 74.46 | 0.00000000 |
| 1946 | 38.42 | 0.00000000 |
| 1947 | 53.76 | 0.00000000 |
| 1948 | 65.65 | 0.00000000 |
| 1949 | 49.23 | 0.00000000 |
| 1950 | 48.46 | 0.00000000 |
| 1951 | 57.68 | 0.00000000 |
| 1952 | 40.32 | 0.00000000 |
| 1953 | 46.24 | 0.00000000 |
| 1954 | 66.26 | 0.00000000 |
| 1955 | 61.00 | 0.00000000 |
| 1956 | 35.96 | 0.00000000 |
| 1957 | 49.57 | 0.00000000 |
| 1958 | 52.90 | 0.00000000 |
| 1959 | 53.61 | 0.00000000 |
| 1960 | 52.83 | 0.00000000 |
| 1961 | 60.57 | 0.00000000 |
| 1962 | 53.26 | 0.00000000 |
| 1963 | 55.13 | 0.00000000 |
| 1964 | 63.23 | 0.00000000 |
| 1965 | 50.35 | 0.00000000 |
| 1966 | 39.85 | 0.00000000 |
| 1967 | 48.33 | 0.00000000 |
| 1968 | 49.58 | 0.00000000 |
| 1969 | 69.33 | 0.00000000 |
| 1970 | 47.57 | 0.00000000 |
| 1971 | 46.13 | 0.00000000 |
| 1972 | 55.96 | 0.00000000 |
| 1973 | 41.14 | 0.00000000 |
| 1974 | 46.96 | 0.00000000 |
| 1975 | 56.62 | 0.00000000 |
| 1976 | 51.70 | 0.00000000 |
| 1977 | 44.15 | 0.00000000 |
| 1978 | 56.90 | 0.00000000 |
| 1979 | 49.96 | 0.00000000 |
| 1980 | 60.00 | 0.00000000 |

| | | |
|---|---|---|
| 1981 | 46.62 | 0.00000000 |
| 1982 | 50.88 | 0.00000000 |
| 1983 | 49.85 | 0.00000000 |
| 1984 | 50.64 | 0.00000000 |
| 1985 | 53.61 | 0.00000000 |
| 1986 | 47.32 | 0.00000000 |
| 1987 | 51.14 | 0.00000000 |
| 1988 | 38.55 | 0.00000000 |
| 1989 | 48.65 | 0.00000000 |
| 1990 | 57.13 | 0.00000000 |
| 1991 | 68.57 | 0.00000000 |
| 1992 | 53.04 | 0.00000000 |
| 1993 | 61.91 | 0.00000000 |
| 1994 | 60.73 | 0.00000000 |
| 1995 | 59.42 | 0.00000000 |
| 1996 | 67.09 | 0.00000000 |
| 1997 | 53.24 | 0.00000000 |
| 1998 | 54.92 | 0.00000000 |
| 1999 | 56.54 | 0.00000000 |
| 2000 | 35.19 | 0.00000000 |
| 2001 | 46.04 | 0.00000000 |
| 2002 | 57.91 | 0.00000000 |
| 2003 | 49.32 | 0.00000000 |
| 2004 | 51.59 | 0.00000000 |
| 2005 | 49.23 | 0.00000000 |
| 2006 | 36.36 | 0.00000000 |
| 2007 | 46.30 | 0.00000000 |
| 2008 | 56.33 | 0.00000000 |
| 2009 | 37.55 | 0.00000000 |
| 2010 | 61.33 | 0.00000000 |
| 2011 | 48.00 | 0.00000000 |
| 2012 | 39.13 | 0.00000000 |
| 2013 | 41.41 | 0.00000000 |
| 2014 | 47.07 | 0.00000000 |
| 2015 | 44.52 | 0.00000000 |
| 2016 | 51.84 | 0.00000000 |
| 2017 | 44.44 | 0.00000000 |
| 2018 | 64.00 | 0.00000000 |
| 2019 | 40.07 | 0.00000000 |
| 2020 | 48.68 | 0.00000000 |
| 2021 | 50.61 | 0.00000000 |
| 2022 | 55.29 | 0.00000000 |
| 2023 | 46.07 | 0.00000000 |
| 2024 | 62.19 | 0.00000000 |

| | | |
|---|---|---|
| 2025 | 39.96 | 0.00000000 |
| 2026 | 55.09 | 0.00000000 |
| 2027 | 51.20 | 0.00000000 |
| 2028 | 42.88 | 0.00000000 |
| 2029 | 48.77 | 0.00000000 |
| 2030 | 43.70 | 0.00000000 |
| 2031 | 50.45 | 0.00000000 |
| 2032 | 45.32 | 0.00000000 |
| 2033 | 47.38 | 0.00000000 |
| 2034 | 50.73 | 0.00000000 |
| 2035 | 61.58 | 0.00000000 |
| 2036 | 55.05 | 0.00000000 |
| 2037 | 48.73 | 0.00000000 |
| 2038 | 62.55 | 0.00000000 |
| 2039 | 31.44 | 0.00000000 |
| 2040 | 48.68 | 0.00000000 |
| 2041 | 43.23 | 0.00000000 |
| 2042 | 54.83 | 0.00000000 |
| 2043 | 54.71 | 0.00000000 |
| 2044 | 41.74 | 0.00000000 |
| 2045 | 49.74 | 0.00000000 |
| 2046 | 34.79 | 0.00000000 |
| 2047 | 36.96 | 0.00000000 |
| 2048 | 62.25 | 0.00000000 |
| 2049 | 50.32 | 0.00000000 |
| 2050 | 45.52 | 0.00000000 |
| 2051 | 58.00 | 0.00000000 |
| 2052 | 52.27 | 0.00000000 |
| 2053 | 57.26 | 0.00000000 |
| 2054 | 51.24 | 0.00000000 |
| 2055 | 38.17 | 0.00000000 |
| 2056 | 53.57 | 0.00000000 |
| 2057 | 48.44 | 0.00000000 |
| 2058 | 57.09 | 0.00000000 |
| 2059 | 36.24 | 0.00000000 |
| 2060 | 51.65 | 0.00000000 |
| 2061 | 59.10 | 0.00000000 |
| 2062 | 57.05 | 0.00000000 |
| 2063 | 50.77 | 0.00000000 |
| 2064 | 63.89 | 0.00000000 |
| 2065 | 50.48 | 0.00000000 |
| 2066 | 55.40 | 0.00000000 |
| 2067 | 51.50 | 0.00000000 |
| 2068 | 54.21 | 0.00000000 |

| | | |
|---|---|---|
| 2069 | 57.10 | 0.00000000 |
| 2070 | 45.29 | 0.00000000 |
| 2071 | 40.33 | 0.00000000 |
| 2072 | 39.74 | 0.00000000 |
| 2073 | 56.43 | 0.00000000 |
| 2074 | 58.83 | 0.00000000 |
| 2075 | 44.14 | 0.00000000 |
| 2076 | 58.11 | 0.00000000 |
| 2077 | 46.52 | 0.00000000 |
| 2078 | 43.65 | 0.00000000 |
| 2079 | 55.59 | 0.00000000 |
| 2080 | 51.42 | 0.00000000 |
| 2081 | 53.30 | 0.00000000 |
| 2082 | 51.19 | 0.00000000 |
| 2083 | 52.09 | 0.00000000 |
| 2084 | 39.64 | 0.00000000 |
| 2085 | 52.04 | 0.00000000 |
| 2086 | 40.10 | 0.00000000 |
| 2087 | 45.74 | 0.00000000 |
| 2088 | 52.36 | 0.00000000 |
| 2089 | 41.91 | 0.00000000 |
| 2090 | 59.05 | 0.00000000 |
| 2091 | 38.09 | 0.00000000 |
| 2092 | 44.68 | 0.00000000 |
| 2093 | 54.77 | 0.00000000 |
| 2094 | 53.71 | 0.00000000 |
| 2095 | 73.50 | 0.00000000 |
| 2096 | 43.29 | 0.00000000 |
| 2097 | 54.00 | 0.00000000 |
| 2098 | 53.55 | 0.00000000 |
| 2099 | 70.32 | 0.00000000 |
| 2100 | 41.65 | 0.00000000 |
| 2101 | 38.71 | 0.00000000 |
| 2102 | 58.45 | 0.00000000 |
| 2103 | 50.29 | 0.00000000 |
| 2104 | 69.00 | 0.00000000 |
| 2105 | 49.89 | 0.00000000 |
| 2106 | 60.14 | 0.00000000 |
| 2107 | 37.60 | 0.00000000 |
| 2108 | 45.45 | 0.00000000 |
| 2109 | 59.57 | 0.00000000 |
| 2110 | 51.67 | 0.00000000 |
| 2111 | 70.14 | 0.00000000 |
| 2112 | 63.05 | 0.00000000 |

| | | |
|---|---|---|
| 2113 | 58.14 | 0.00000000 |
| 2114 | 55.30 | 0.00000000 |
| 2115 | 68.14 | 0.00000000 |
| 2116 | 55.70 | 0.00000000 |
| 2117 | 61.61 | 0.00000000 |
| 2118 | 58.48 | 0.00000000 |
| 2119 | 58.05 | 0.00000000 |
| 2120 | 42.68 | 0.00000000 |
| 2121 | 66.05 | 0.00000000 |
| 2122 | 42.72 | 0.00000000 |
| 2123 | 57.91 | 0.00000000 |
| 2124 | 44.64 | 0.00000000 |
| 2125 | 47.92 | 0.00000000 |
| 2126 | 59.32 | 0.00000000 |
| 2127 | 39.91 | 0.00000000 |
| 2128 | 44.70 | 0.00000000 |
| 2129 | 46.39 | 0.00000000 |
| 2130 | 73.41 | 0.00000000 |
| 2131 | 40.62 | 0.00000000 |
| 2132 | 56.00 | 0.00000000 |
| 2133 | 52.73 | 0.00000000 |
| 2134 | 49.73 | 0.00000000 |
| 2135 | 55.00 | 0.00000000 |
| 2136 | 41.27 | 0.00000000 |
| 2137 | 43.55 | 0.00000000 |
| 2138 | 47.87 | 0.00000000 |
| 2139 | 37.33 | 0.00000000 |
| 2140 | 43.04 | 0.00000000 |
| 2141 | 46.09 | 0.00000000 |
| 2142 | 41.12 | 0.00000000 |
| 2143 | 44.00 | 0.00000000 |
| 2144 | 44.04 | 0.00000000 |
| 2145 | 31.57 | 0.00000000 |
| 2146 | 31.23 | 0.00000000 |
| 2147 | 40.92 | 0.00000000 |
| 2148 | 40.48 | 0.00000000 |
| 2149 | 41.13 | 0.00000000 |
| 2150 | 43.77 | 0.00000000 |
| 2151 | 34.88 | 0.00000000 |
| 2152 | 35.33 | 0.00000000 |
| 2153 | 33.95 | 0.00000000 |
| 2154 | 48.95 | 0.00000000 |
| 2155 | 53.25 | 0.00000000 |
| 2156 | 51.11 | 0.00000000 |

| | | |
|---|---|---|
| 2157 | 28.83 | 0.00000000 |
| 2158 | 33.70 | 0.00000000 |
| 2159 | 42.38 | 0.00000000 |
| 2160 | 47.05 | 0.00000000 |
| 2161 | 38.67 | 0.00000000 |
| 2162 | 41.21 | 0.00000000 |
| 2163 | 40.96 | 0.00000000 |
| 2164 | 44.86 | 0.00000000 |
| 2165 | 28.91 | 0.00000000 |
| 2166 | 32.00 | 0.00000000 |
| 2167 | 44.29 | 0.00000000 |
| 2168 | 41.53 | 0.00000000 |
| 2169 | 56.36 | 0.00000000 |
| 2170 | 38.68 | 0.00000000 |
| 2171 | 47.55 | 0.00000000 |
| 2172 | 49.87 | 0.00000000 |
| 2173 | 41.83 | 0.00000000 |
| 2174 | 44.33 | 0.00000000 |
| 2175 | 28.92 | 0.00000000 |
| 2176 | 36.08 | 0.00000000 |
| 2177 | 32.79 | 0.00000000 |
| 2178 | 46.96 | 0.00000000 |
| 2179 | 41.70 | 0.00000000 |
| 2180 | 38.18 | 0.00000000 |
| 2181 | 34.71 | 0.00000000 |
| 2182 | 39.41 | 0.00000000 |
| 2183 | 44.59 | 0.00000000 |
| 2184 | 54.09 | 0.00000000 |
| 2185 | 56.95 | 0.00000000 |
| 2186 | 51.81 | 0.00000000 |
| 2187 | 47.48 | 0.00000000 |
| 2188 | 50.00 | 0.00000000 |
| 2189 | 46.30 | 0.00000000 |
| 2190 | 42.57 | 0.00000000 |
| 2191 | 50.05 | 0.00000000 |
| 2192 | 41.89 | 0.00000000 |
| 2193 | 59.25 | 0.00000000 |
| 2194 | 68.94 | 0.00000000 |
| 2195 | 37.27 | 0.00000000 |
| 2196 | 64.74 | 0.00000000 |
| 2197 | 62.00 | 0.00000000 |
| 2198 | 36.53 | 0.00000000 |
| 2199 | 37.43 | 0.00000000 |
| 2200 | 60.33 | 0.00000000 |

| | | |
|---|---|---|
| 2201 | 60.81 | 0.00000000 |
| 2202 | 57.55 | 0.00000000 |
| 2203 | 37.09 | 0.00000000 |
| 2204 | 35.11 | 0.00000000 |
| 2205 | 30.54 | 0.00000000 |
| 2206 | 38.22 | 0.00000000 |
| 2207 | 45.00 | 0.00000000 |
| 2208 | 44.64 | 0.00000000 |
| 2209 | 61.00 | 0.00000000 |
| 2210 | 67.05 | 0.00000000 |
| 2211 | 22.46 | 0.00000000 |
| 2212 | 64.17 | 0.00000000 |
| 2213 | 36.95 | 0.00000000 |
| 2214 | 41.05 | 0.00000000 |
| 2215 | 47.17 | 0.00000000 |
| 2216 | 38.71 | 0.00000000 |
| 2217 | 45.48 | 0.00000000 |
| 2218 | 64.76 | 0.00000000 |
| 2219 | 86.53 | 0.00000000 |
| 2220 | 47.05 | 0.00000000 |
| 2221 | 54.95 | 0.00000000 |
| 2222 | 53.21 | 0.00000000 |
| 2223 | 34.82 | 0.00000000 |
| 2224 | 46.94 | 0.00000000 |
| 2225 | 45.50 | 0.00000000 |
| 2226 | 48.05 | 0.00000000 |
| 2227 | 70.73 | 0.00000000 |
| 2228 | 35.70 | 0.00000000 |
| 2229 | 48.50 | 0.00000000 |
| 2230 | 40.38 | 0.00000000 |
| 2231 | 43.00 | 0.00000000 |
| 2232 | 45.41 | 0.00000000 |
| 2233 | 45.29 | 0.00000000 |
| 2234 | 59.12 | 0.00000000 |
| 2235 | 39.21 | 0.00000000 |
| 2236 | 44.59 | 0.00000000 |
| 2237 | 46.84 | 0.00000000 |
| 2238 | 39.00 | 0.00000000 |
| 2239 | 44.10 | 0.00000000 |
| 2240 | 64.32 | 0.00000000 |
| 2241 | 38.23 | 0.00000000 |
| 2242 | 52.05 | 0.00000000 |
| 2243 | 38.90 | 0.00000000 |
| 2244 | 40.80 | 0.00000000 |

| | | |
|---|---|---|
| 2245 | 45.28 | 0.00000000 |
| 2246 | 41.22 | 0.00000000 |
| 2247 | 41.05 | 0.00000000 |
| 2248 | 34.76 | 0.00000000 |
| 2249 | 41.22 | 0.00000000 |
| 2250 | 68.38 | 0.00000000 |
| 2251 | 41.45 | 0.00000000 |
| 2252 | 46.59 | 0.00000000 |
| 2253 | 47.87 | 0.00000000 |
| 2254 | 45.70 | 0.00000000 |
| 2255 | 39.76 | 0.00000000 |
| 2256 | 40.05 | 0.00000000 |
| 2257 | 28.10 | 0.00000000 |
| 2258 | 60.85 | 0.00000000 |
| 2259 | 36.36 | 0.00000000 |
| 2260 | 34.67 | 0.00000000 |
| 2261 | 33.83 | 0.00000000 |
| 2262 | 35.16 | 0.00000000 |
| 2263 | 43.00 | 0.00000000 |
| 2264 | 45.65 | 0.00000000 |
| 2265 | 36.90 | 0.00000000 |
| 2266 | 42.11 | 0.00000000 |
| 2267 | 48.16 | 0.00000000 |
| 2268 | 41.23 | 0.00000000 |
| 2269 | 47.37 | 0.00000000 |
| 2270 | 41.35 | 0.00000000 |
| 2271 | 45.81 | 0.00000000 |
| 2272 | 48.43 | 0.00000000 |
| 2273 | 53.83 | 0.00000000 |
| 2274 | 42.10 | 0.00000000 |
| 2275 | 41.39 | 0.00000000 |
| 2276 | 29.95 | 0.00000000 |
| 2277 | 47.42 | 0.00000000 |
| 2278 | 31.57 | 0.00000000 |
| 2279 | 38.18 | 0.00000000 |
| 2280 | 39.38 | 0.00000000 |
| 2281 | 45.00 | 0.00000000 |
| 2282 | 51.50 | 0.00000000 |
| 2283 | 29.52 | 0.00000000 |
| 2284 | 48.95 | 0.00000000 |
| 2285 | 30.35 | 0.00000000 |
| 2286 | 40.35 | 0.00000000 |
| 2287 | 42.44 | 0.00000000 |
| 2288 | 40.57 | 0.00000000 |

| | | |
|---|---|---|
| 2289 | 32.00 | 0.00000000 |
| 2290 | 28.83 | 0.00000000 |
| 2291 | 38.37 | 0.00000000 |
| 2292 | 63.06 | 0.00000000 |
| 2293 | 48.47 | 0.00000000 |
| 2294 | 51.65 | 0.00000000 |
| 2295 | 53.44 | 0.00000000 |
| 2296 | 31.43 | 0.00000000 |
| 2297 | 32.16 | 0.00000000 |
| 2298 | 45.09 | 0.00000000 |
| 2299 | 34.10 | 0.00000000 |
| 2300 | 39.83 | 0.00000000 |
| 2301 | 35.05 | 0.00000000 |
| 2302 | 51.00 | 0.00000000 |
| 2303 | 34.50 | 0.00000000 |
| 2304 | 43.95 | 0.00000000 |
| 2305 | 49.74 | 0.00000000 |
| 2306 | 40.55 | 0.00000000 |
| 2307 | 53.29 | 0.00000000 |
| 2308 | 38.15 | 0.00000000 |
| 2309 | 49.91 | 0.00000000 |
| 2310 | 43.40 | 0.00000000 |
| 2311 | 40.56 | 0.00000000 |
| 2312 | 43.84 | 0.00000000 |
| 2313 | 43.58 | 0.00000000 |
| 2314 | 40.50 | 0.00000000 |
| 2315 | 34.70 | 0.00000000 |
| 2316 | 28.48 | 0.00000000 |
| 2317 | 42.09 | 0.00000000 |
| 2318 | 28.62 | 0.00000000 |
| 2319 | 42.41 | 0.00000000 |
| 2320 | 37.79 | 0.00000000 |
| 2321 | 47.91 | 0.00000000 |
| 2322 | 61.53 | 0.00000000 |
| 2323 | 35.83 | 0.00000000 |
| 2324 | 37.43 | 0.00000000 |
| 2325 | 49.30 | 0.00000000 |
| 2326 | 48.15 | 0.00000000 |
| 2327 | 44.89 | 0.00000000 |
| 2328 | 42.55 | 0.00000000 |
| 2329 | 37.95 | 0.00000000 |
| 2330 | 49.89 | 0.00000000 |
| 2331 | 44.65 | 0.00000000 |
| 2332 | 40.42 | 0.00000000 |

| | | |
|---|---|---|
| 2333 | 49.68 | 0.00000000 |
| 2334 | 43.67 | 0.00000000 |
| 2335 | 42.83 | 0.00000000 |
| 2336 | 36.74 | 0.00000000 |
| 2337 | 30.00 | 0.00000000 |
| 2338 | 37.32 | 0.00000000 |
| 2339 | 44.57 | 0.00000000 |
| 2340 | 27.27 | 0.00000000 |
| 2341 | 35.14 | 0.00000000 |
| 2342 | 26.65 | 0.00000000 |
| 2343 | 39.00 | 0.00000000 |
| 2344 | 33.18 | 0.00000000 |
| 2345 | 40.58 | 0.00000000 |
| 2346 | 37.14 | 0.00000000 |
| 2347 | 41.14 | 0.00000000 |
| 2348 | 38.10 | 0.00000000 |
| 2349 | 33.57 | 0.00000000 |
| 2350 | 36.25 | 0.00000000 |
| 2351 | 56.25 | 0.00000000 |
| 2352 | 49.16 | 0.00000000 |
| 2353 | 56.76 | 0.00000000 |
| 2354 | 36.58 | 0.00000000 |
| 2355 | 46.89 | 0.00000000 |
| 2356 | 43.41 | 0.00000000 |
| 2357 | 24.62 | 0.00000000 |
| 2358 | 21.14 | 0.00000000 |
| 2359 | 34.57 | 0.00000000 |
| 2360 | 30.06 | 0.00000000 |
| 2361 | 28.32 | 0.00000000 |
| 2362 | 19.92 | 0.00000000 |
| 2363 | 46.67 | 0.00000000 |
| 2364 | 46.71 | 0.00000000 |
| 2365 | 40.30 | 0.00000000 |
| 2366 | 29.17 | 0.00000000 |
| 2367 | 64.00 | 0.00000000 |
| 2368 | 35.65 | 0.00000000 |
| 2369 | 49.00 | 0.00000000 |
| 2370 | 31.17 | 0.00000000 |
| 2371 | 31.68 | 0.00000000 |
| 2372 | 31.35 | 0.00000000 |
| 2373 | 24.12 | 0.00000000 |
| 2374 | 29.26 | 0.00000000 |
| 2375 | 37.29 | 0.00000000 |
| 2376 | 45.16 | 0.00000000 |

| | | |
|---|---|---|
| 2377 | 36.90 | 0.00000000 |
| 2378 | 35.24 | 0.00000000 |
| 2379 | 42.95 | 0.00000000 |
| 2380 | 30.70 | 0.00000000 |
| 2381 | 31.96 | 0.00000000 |
| 2382 | 34.24 | 0.00000000 |
| 2383 | 37.27 | 0.00000000 |
| 2384 | 35.05 | 0.00000000 |
| 2385 | 32.04 | 0.00000000 |
| 2386 | 53.50 | 0.00000000 |
| 2387 | 34.96 | 0.00000000 |
| 2388 | 39.48 | 0.00000000 |
| 2389 | 43.86 | 0.00000000 |
| 2390 | 45.68 | 0.00000000 |
| 2391 | 24.58 | 0.00000000 |
| 2392 | 50.00 | 0.00000000 |
| 2393 | 40.88 | 0.00000000 |
| 2394 | 48.11 | 0.00000000 |
| 2395 | 46.75 | 0.00000000 |
| 2396 | 48.50 | 0.00000000 |
| 2397 | 24.57 | 0.00000000 |
| 2398 | 31.91 | 0.00000000 |
| 2399 | 37.43 | 0.00000000 |
| 2400 | 24.30 | 0.00000000 |
| 2401 | 24.57 | 0.00000000 |
| 2402 | 38.71 | 0.00000000 |
| 2403 | 27.90 | 0.00000000 |
| 2404 | 45.32 | 0.00000000 |
| 2405 | 44.11 | 0.00000000 |
| 2406 | 32.21 | 0.00000000 |
| 2407 | 42.23 | 0.00000000 |
| 2408 | 35.53 | 0.00000000 |
| 2409 | 39.59 | 0.00000000 |
| 2410 | 34.55 | 0.00000000 |
| 2411 | 29.22 | 0.00000000 |
| 2412 | 23.32 | 0.00000000 |
| 2413 | 50.80 | 0.00000000 |
| 2414 | 39.50 | 0.00000000 |
| 2415 | 41.24 | 0.00000000 |
| 2416 | 36.05 | 0.00000000 |
| 2417 | 26.29 | 0.00000000 |
| 2418 | 39.38 | 0.00000000 |
| 2419 | 48.21 | 0.00000000 |
| 2420 | 39.80 | 0.00000000 |

| | | |
|---|---|---|
| 2421 | 51.67 | 0.00000000 |
| 2422 | 29.78 | 0.00000000 |
| 2423 | 44.06 | 0.00000000 |
| 2424 | 34.14 | 0.00000000 |
| 2425 | 26.13 | 0.00000000 |
| 2426 | 26.79 | 0.00000000 |
| 2427 | 43.00 | 0.00000000 |
| 2428 | 48.41 | 0.00000000 |
| 2429 | 41.50 | 0.00000000 |
| 2430 | 48.56 | 0.00000000 |
| 2431 | 32.05 | 0.00000000 |
| 2432 | 34.78 | 0.00000000 |
| 2433 | 42.63 | 0.00000000 |
| 2434 | 27.39 | 0.00000000 |
| 2435 | 40.75 | 0.00000000 |
| 2436 | 33.88 | 0.00000000 |
| 2437 | 28.24 | 0.00000000 |
| 2438 | 35.15 | 0.00000000 |
| 2439 | 32.95 | 0.00000000 |
| 2440 | 24.50 | 0.00000000 |
| 2441 | 29.39 | 0.00000000 |
| 2442 | 45.71 | 0.00000000 |
| 2443 | 30.05 | 0.00000000 |
| 2444 | 38.79 | 0.00000000 |
| 2445 | 35.47 | 0.00000000 |
| 2446 | 33.79 | 0.00000000 |
| 2447 | 54.56 | 0.00000000 |
| 2448 | 35.45 | 0.00000000 |
| 2449 | 56.59 | 0.00000000 |
| 2450 | 31.45 | 0.00000000 |
| 2451 | 28.81 | 0.00000000 |
| 2452 | 29.95 | 0.00000000 |
| 2453 | 34.39 | 0.00000000 |
| 2454 | 39.65 | 0.00000000 |
| 2455 | 34.82 | 0.00000000 |
| 2456 | 49.38 | 0.00000000 |
| 2457 | 28.53 | 0.00000000 |
| 2458 | 26.29 | 0.00000000 |
| 2459 | 26.38 | 0.00000000 |
| 2460 | 30.05 | 0.00000000 |
| 2461 | 22.00 | 0.00000000 |
| 2462 | 28.95 | 0.00000000 |
| 2463 | 20.55 | 0.00000000 |
| 2464 | 15.09 | 0.00000000 |

| | | |
|---|---|---|
| 2465 | 47.30 | 0.00000000 |
| 2466 | 29.67 | 0.00000000 |
| 2467 | 39.00 | 0.00000000 |
| 2468 | 50.11 | 0.00000000 |
| 2469 | 33.53 | 0.00000000 |
| 2470 | 42.55 | 0.00000000 |
| 2471 | 32.63 | 0.00000000 |
| 2472 | 43.67 | 0.00000000 |
| 2473 | 38.00 | 0.00000000 |
| 2474 | 38.61 | 0.00000000 |
| 2475 | 42.29 | 0.00000000 |
| 2476 | 22.29 | 0.00000000 |
| 2477 | 56.11 | 0.00000000 |
| 2478 | 30.44 | 0.00000000 |
| 2479 | 45.71 | 0.00000000 |
| 2480 | 34.05 | 0.00000000 |
| 2481 | 22.94 | 0.00000000 |
| 2482 | 30.47 | 0.00000000 |
| 2483 | 23.47 | 0.00000000 |
| 2484 | 29.59 | 0.00000000 |
| 2485 | 33.05 | 0.00000000 |
| 2486 | 43.38 | 0.00000000 |
| 2487 | 37.53 | 0.00000000 |
| 2488 | 44.33 | 0.00000000 |
| 2489 | 32.56 | 0.00000000 |
| 2490 | 22.80 | 0.00000000 |
| 2491 | 50.29 | 0.00000000 |
| 2492 | 45.65 | 0.00000000 |
| 2493 | 23.35 | 0.00000000 |
| 2494 | 34.12 | 0.00000000 |
| 2495 | 31.35 | 0.00000000 |
| 2496 | 41.26 | 0.00000000 |
| 2497 | 46.21 | 0.00000000 |
| 2498 | 43.11 | 0.00000000 |
| 2499 | 45.84 | 0.00000000 |
| 2500 | 43.71 | 0.00000000 |
| 2501 | 34.65 | 0.00000000 |
| 2502 | 19.91 | 0.00000000 |
| 2503 | 39.25 | 0.00000000 |
| 2504 | 35.23 | 0.00000000 |
| 2505 | 27.52 | 0.00000000 |
| 2506 | 52.44 | 0.00000000 |
| 2507 | 37.09 | 0.00000000 |
| 2508 | 28.52 | 0.00000000 |

| | | |
|---|---|---|
| 2509 | 25.35 | 0.00000000 |
| 2510 | 32.27 | 0.00000000 |
| 2511 | 17.25 | 0.00000000 |
| 2512 | 32.12 | 0.00000000 |
| 2513 | 33.08 | 0.00000000 |
| 2514 | 25.36 | 0.00000000 |
| 2515 | 31.53 | 0.00000000 |
| 2516 | 26.43 | 0.00000000 |
| 2517 | 19.57 | 0.00000000 |
| 2518 | 39.68 | 0.00000000 |
| 2519 | 28.62 | 0.00000000 |
| 2520 | 28.36 | 0.00000000 |
| 2521 | 30.59 | 0.00000000 |
| 2522 | 30.48 | 0.00000000 |
| 2523 | 32.26 | 0.00000000 |
| 2524 | 42.26 | 0.00000000 |
| 2525 | 37.85 | 0.00000000 |
| 2526 | 40.79 | 0.00000000 |
| 2527 | 31.05 | 0.00000000 |
| 2528 | 28.59 | 0.00000000 |
| 2529 | 42.00 | 0.00000000 |
| 2530 | 43.05 | 0.00000000 |
| 2531 | 34.29 | 0.00000000 |
| 2532 | 43.25 | 0.00000000 |
| 2533 | 28.25 | 0.00000000 |
| 2534 | 41.65 | 0.00000000 |
| 2535 | 47.00 | 0.00000000 |
| 2536 | 47.58 | 0.00000000 |
| 2537 | 34.37 | 0.00000000 |
| 2538 | 43.68 | 0.00000000 |
| 2539 | 43.11 | 0.00000000 |
| 2540 | 30.05 | 0.00000000 |
| 2541 | 33.32 | 0.00000000 |
| 2542 | 54.00 | 0.00000000 |
| 2543 | 30.79 | 0.00000000 |
| 2544 | 30.22 | 0.00000000 |
| 2545 | 38.78 | 0.00000000 |
| 2546 | 19.52 | 0.00000000 |
| 2547 | 34.53 | 0.00000000 |
| 2548 | 33.50 | 0.00000000 |
| 2549 | 40.00 | 0.00000000 |
| 2550 | 34.28 | 0.00000000 |
| 2551 | 25.14 | 0.00000000 |
| 2552 | 35.00 | 0.00000000 |

| | | |
|---|---|---|
| 2553 | 22.55 | 0.00000000 |
| 2554 | 33.18 | 0.00000000 |
| 2555 | 42.00 | 0.00000000 |
| 2556 | 29.45 | 0.00000000 |
| 2557 | 26.50 | 0.00000000 |
| 2558 | 26.50 | 0.00000000 |
| 2559 | 31.91 | 0.00000000 |
| 2560 | 31.10 | 0.00000000 |
| 2561 | 55.16 | 0.00000000 |
| 2562 | 37.00 | 0.00000000 |
| 2563 | 34.42 | 0.00000000 |
| 2564 | 27.64 | 0.00000000 |
| 2565 | 30.86 | 0.00000000 |
| 2566 | 39.48 | 0.00000000 |
| 2567 | 31.90 | 0.00000000 |
| 2568 | 37.48 | 0.00000000 |
| 2569 | 29.64 | 0.00000000 |
| 2570 | 38.42 | 0.00000000 |
| 2571 | 25.11 | 0.00000000 |
| 2572 | 21.95 | 0.00000000 |
| 2573 | 25.08 | 0.00000000 |
| 2574 | 34.14 | 0.00000000 |
| 2575 | 35.91 | 0.00000000 |
| 2576 | 40.17 | 0.00000000 |
| 2577 | 26.65 | 0.00000000 |
| 2578 | 38.40 | 0.00000000 |
| 2579 | 33.55 | 0.00000000 |
| 2580 | 32.55 | 0.00000000 |
| 2581 | 37.20 | 0.00000000 |
| 2582 | 36.50 | 0.00000000 |
| 2583 | 46.95 | 0.00000000 |
| 2584 | 39.83 | 0.00000000 |
| 2585 | 38.95 | 0.00000000 |
| 2586 | 39.61 | 0.00000000 |
| 2587 | 49.81 | 0.00000000 |
| 2588 | 26.00 | 0.00000000 |
| 2589 | 34.18 | 0.00000000 |
| 2590 | 37.58 | 0.00000000 |
| 2591 | 17.15 | 0.00000000 |
| 2592 | 26.65 | 0.00000000 |
| 2593 | 30.00 | 0.00000000 |
| 2594 | 36.50 | 0.00000000 |
| 2595 | 34.41 | 0.00000000 |
| 2596 | 31.12 | 0.00000000 |

| | | |
|---|---|---|
| 2597 | 30.67 | 0.00000000 |
| 2598 | 28.06 | 0.00000000 |
| 2599 | 33.71 | 0.00000000 |
| 2600 | 34.86 | 0.00000000 |
| 2601 | 37.50 | 0.00000000 |
| 2602 | 28.14 | 0.00000000 |
| 2603 | 28.05 | 0.00000000 |
| 2604 | 34.05 | 0.00000000 |
| 2605 | 32.29 | 0.00000000 |
| 2606 | 38.89 | 0.00000000 |
| 2607 | 31.16 | 0.00000000 |
| 2608 | 31.58 | 0.00000000 |
| 2609 | 40.32 | 0.00000000 |
| 2610 | 30.47 | 0.00000000 |
| 2611 | 35.22 | 0.00000000 |
| 2612 | 40.18 | 0.00000000 |
| 2613 | 22.00 | 0.00000000 |
| 2614 | 27.06 | 0.00000000 |
| 2615 | 30.63 | 0.00000000 |
| 2616 | 30.48 | 0.00000000 |
| 2617 | 31.42 | 0.00000000 |
| 2618 | 27.65 | 0.00000000 |
| 2619 | 22.42 | 0.00000000 |
| 2620 | 33.47 | 0.00000000 |
| 2621 | 19.74 | 0.00000000 |
| 2622 | 27.72 | 0.00000000 |
| 2623 | 33.82 | 0.00000000 |
| 2624 | 44.90 | 0.00000000 |
| 2625 | 26.95 | 0.00000000 |
| 2626 | 20.00 | 0.00000000 |
| 2627 | 34.14 | 0.00000000 |
| 2628 | 23.95 | 0.00000000 |
| 2629 | 33.60 | 0.00000000 |
| 2630 | 32.79 | 0.00000000 |
| 2631 | 35.61 | 0.00000000 |
| 2632 | 43.10 | 0.00000000 |
| 2633 | 23.42 | 0.00000000 |
| 2634 | 20.16 | 0.00000000 |
| 2635 | 26.84 | 0.00000000 |
| 2636 | 39.90 | 0.00000000 |
| 2637 | 45.90 | 0.00000000 |
| 2638 | 31.67 | 0.00000000 |
| 2639 | 28.45 | 0.00000000 |
| 2640 | 37.48 | 0.00000000 |

| | | |
|---|---|---|
| 2641 | 24.95 | 0.00000000 |
| 2642 | 22.45 | 0.00000000 |
| 2643 | 28.35 | 0.00000000 |
| 2644 | 27.30 | 0.00000000 |
| 2645 | 32.29 | 0.00000000 |
| 2646 | 29.67 | 0.00000000 |
| 2647 | 29.76 | 0.00000000 |
| 2648 | 30.05 | 0.00000000 |
| 2649 | 39.52 | 0.00000000 |
| 2650 | 32.48 | 0.00000000 |
| 2651 | 26.23 | 0.00000000 |
| 2652 | 42.53 | 0.00000000 |
| 2653 | 33.23 | 0.00000000 |
| 2654 | 30.81 | 0.00000000 |
| 2655 | 39.82 | 0.00000000 |
| 2656 | 46.84 | 0.00000000 |
| 2657 | 32.45 | 0.00000000 |
| 2658 | 28.19 | 0.00000000 |
| 2659 | 39.78 | 0.00000000 |
| 2660 | 28.17 | 0.00000000 |
| 2661 | 39.39 | 0.00000000 |
| 2662 | 38.29 | 0.00000000 |
| 2663 | 55.13 | 0.00000000 |
| 2664 | 21.42 | 0.00000000 |
| 2665 | 33.87 | 0.00000000 |
| 2666 | 21.22 | 0.00000000 |
| 2667 | 33.63 | 0.00000000 |
| 2668 | 17.53 | 0.00000000 |
| 2669 | 26.47 | 0.00000000 |
| 2670 | 45.74 | 0.00000000 |
| 2671 | 31.00 | 0.00000000 |
| 2672 | 28.83 | 0.00000000 |
| 2673 | 26.95 | 0.00000000 |
| 2674 | 33.28 | 0.00000000 |
| 2675 | 22.67 | 0.00000000 |
| 2676 | 26.11 | 0.00000000 |
| 2677 | 20.85 | 0.00000000 |
| 2678 | 20.85 | 0.00000000 |
| 2679 | 25.78 | 0.00000000 |
| 2680 | 18.44 | 0.00000000 |
| 2681 | 37.47 | 0.00000000 |
| 2682 | 22.56 | 0.00000000 |
| 2683 | 29.00 | 0.00000000 |
| 2684 | 20.48 | 0.00000000 |

| | | |
|---|---|---|
| 2685 | 14.32 | 0.00000000 |
| 2686 | 32.26 | 0.00000000 |
| 2687 | 36.72 | 0.00000000 |
| 2688 | 31.00 | 0.00000000 |
| 2689 | 36.00 | 0.00000000 |
| 2690 | 24.48 | 0.00000000 |
| 2691 | 29.17 | 0.00000000 |
| 2692 | 39.90 | 0.00000000 |
| 2693 | 26.71 | 0.00000000 |
| 2694 | 23.95 | 0.00000000 |
| 2695 | 27.40 | 0.00000000 |
| 2696 | 30.52 | 0.00000000 |
| 2697 | 27.47 | 0.00000000 |
| 2698 | 44.81 | 0.00000000 |
| 2699 | 44.22 | 0.00000000 |
| 2700 | 52.12 | 0.00000000 |
| 2701 | 36.35 | 0.00000000 |
| 2702 | 33.76 | 0.00000000 |
| 2703 | 34.00 | 0.00000000 |
| 2704 | 20.33 | 0.00000000 |
| 2705 | 43.67 | 0.00000000 |
| 2706 | 22.47 | 0.00000000 |
| 2707 | 28.00 | 0.00000000 |
| 2708 | 18.47 | 0.00000000 |
| 2709 | 25.76 | 0.00000000 |
| 2710 | 39.89 | 0.00000000 |
| 2711 | 36.35 | 0.00000000 |
| 2712 | 14.82 | 0.00000000 |
| 2713 | 27.00 | 0.00000000 |
| 2714 | 28.95 | 0.00000000 |
| 2715 | 22.41 | 0.00000000 |
| 2716 | 22.29 | 0.00000000 |
| 2717 | 27.55 | 0.00000000 |
| 2718 | 24.75 | 0.00000000 |
| 2719 | 27.30 | 0.00000000 |
| 2720 | 26.71 | 0.00000000 |
| 2721 | 29.69 | 0.00000000 |
| 2722 | 18.70 | 0.00000000 |
| 2723 | 26.11 | 0.00000000 |
| 2724 | 27.11 | 0.00000000 |
| 2725 | 20.32 | 0.00000000 |
| 2726 | 51.10 | 0.00000000 |
| 2727 | 28.48 | 0.00000000 |
| 2728 | 31.71 | 0.00000000 |

| | | |
|---|---|---|
| 2729 | 28.95 | 0.00000000 |
| 2730 | 35.50 | 0.00000000 |
| 2731 | 28.95 | 0.00000000 |
| 2732 | 37.71 | 0.00000000 |
| 2733 | 28.95 | 0.00000000 |
| 2734 | 23.83 | 0.00000000 |
| 2735 | 22.13 | 0.00000000 |
| 2736 | 34.79 | 0.00000000 |
| 2737 | 33.32 | 0.00000000 |
| 2738 | 41.73 | 0.00000000 |
| 2739 | 26.19 | 0.00000000 |
| 2740 | 45.31 | 0.00000000 |
| 2741 | 20.53 | 0.00000000 |
| 2742 | 34.55 | 0.00000000 |
| 2743 | 39.56 | 0.00000000 |
| 2744 | 30.78 | 0.00000000 |
| 2745 | 37.05 | 0.00000000 |
| 2746 | 40.84 | 0.00000000 |
| 2747 | 23.05 | 0.00000000 |
| 2748 | 28.00 | 0.00000000 |
| 2749 | 37.94 | 0.00000000 |
| 2750 | 35.53 | 0.00000000 |
| 2751 | 34.31 | 0.00000000 |
| 2752 | 24.73 | 0.00000000 |
| 2753 | 32.24 | 0.00000000 |
| 2754 | 34.59 | 0.00000000 |
| 2755 | 52.25 | 0.00000000 |
| 2756 | 36.05 | 0.00000000 |
| 2757 | 30.43 | 0.00000000 |
| 2758 | 31.18 | 0.00000000 |
| 2759 | 25.42 | 0.00000000 |
| 2760 | 22.29 | 0.00000000 |
| 2761 | 25.74 | 0.00000000 |
| 2762 | 44.19 | 0.00000000 |
| 2763 | 21.44 | 0.00000000 |
| 2764 | 20.88 | 0.00000000 |
| 2765 | 22.11 | 0.00000000 |
| 2766 | 33.40 | 0.00000000 |
| 2767 | 25.67 | 0.00000000 |
| 2768 | 18.89 | 0.00000000 |
| 2769 | 20.06 | 0.00000000 |
| 2770 | 21.30 | 0.00000000 |
| 2771 | 34.75 | 0.00000000 |
| 2772 | 31.25 | 0.00000000 |

| | | |
|---|---|---|
| 2773 | 25.05 | 0.00000000 |
| 2774 | 27.80 | 0.00000000 |
| 2775 | 29.61 | 0.00000000 |
| 2776 | 19.72 | 0.00000000 |
| 2777 | 36.35 | 0.00000000 |
| 2778 | 28.22 | 0.00000000 |
| 2779 | 32.17 | 0.00000000 |
| 2780 | 26.55 | 0.00000000 |
| 2781 | 35.89 | 0.00000000 |
| 2782 | 55.29 | 0.00000000 |
| 2783 | 26.05 | 0.00000000 |
| 2784 | 35.00 | 0.00000000 |
| 2785 | 43.79 | 0.00000000 |
| 2786 | 22.82 | 0.00000000 |
| 2787 | 35.00 | 0.00000000 |
| 2788 | 37.67 | 0.00000000 |
| 2789 | 32.63 | 0.00000000 |
| 2790 | 23.74 | 0.00000000 |
| 2791 | 23.95 | 0.00000000 |
| 2792 | 24.79 | 0.00000000 |
| 2793 | 56.21 | 0.00000000 |
| 2794 | 22.63 | 0.00000000 |
| 2795 | 40.41 | 0.00000000 |
| 2796 | 25.22 | 0.00000000 |
| 2797 | 21.53 | 0.00000000 |
| 2798 | 31.56 | 0.00000000 |
| 2799 | 18.00 | 0.00000000 |
| 2800 | 20.65 | 0.00000000 |
| 2801 | 36.94 | 0.00000000 |
| 2802 | 29.78 | 0.00000000 |
| 2803 | 16.35 | 0.00000000 |
| 2804 | 44.41 | 0.00000000 |
| 2805 | 24.94 | 0.00000000 |
| 2806 | 34.53 | 0.00000000 |
| 2807 | 22.70 | 0.00000000 |
| 2808 | 27.33 | 0.00000000 |
| 2809 | 16.32 | 0.00000000 |
| 2810 | 24.94 | 0.00000000 |
| 2811 | 44.61 | 0.00000000 |
| 2812 | 23.11 | 0.00000000 |
| 2813 | 27.60 | 0.00000000 |
| 2814 | 27.60 | 0.00000000 |
| 2815 | 30.35 | 0.00000000 |
| 2816 | 32.29 | 0.00000000 |

| | | |
|---|---|---|
| 2817 | 33.00 | 0.00000000 |
| 2818 | 32.60 | 0.00000000 |
| 2819 | 26.24 | 0.00000000 |
| 2820 | 30.43 | 0.00000000 |
| 2821 | 30.45 | 0.00000000 |
| 2822 | 46.06 | 0.00000000 |
| 2823 | 41.33 | 0.00000000 |
| 2824 | 35.76 | 0.00000000 |
| 2825 | 40.06 | 0.00000000 |
| 2826 | 30.39 | 0.00000000 |
| 2827 | 22.28 | 0.00000000 |
| 2828 | 29.61 | 0.00000000 |
| 2829 | 29.84 | 0.00000000 |
| 2830 | 26.25 | 0.00000000 |
| 2831 | 26.94 | 0.00000000 |
| 2832 | 32.13 | 0.00000000 |
| 2833 | 36.37 | 0.00000000 |
| 2834 | 33.06 | 0.00000000 |
| 2835 | 35.82 | 0.00000000 |
| 2836 | 36.80 | 0.00000000 |
| 2837 | 38.13 | 0.00000000 |
| 2838 | 29.53 | 0.00000000 |
| 2839 | 32.39 | 0.00000000 |
| 2840 | 30.47 | 0.00000000 |
| 2841 | 31.47 | 0.00000000 |
| 2842 | 22.81 | 0.00000000 |
| 2843 | 46.24 | 0.00000000 |
| 2844 | 40.00 | 0.00000000 |
| 2845 | 24.55 | 0.00000000 |
| 2846 | 25.11 | 0.00000000 |
| 2847 | 30.94 | 0.00000000 |
| 2848 | 45.93 | 0.00000000 |
| 2849 | 15.47 | 0.00000000 |
| 2850 | 39.25 | 0.00000000 |
| 2851 | 35.94 | 0.00000000 |
| 2852 | 37.29 | 0.00000000 |
| 2853 | 33.90 | 0.00000000 |
| 2854 | 36.29 | 0.00000000 |
| 2855 | 27.29 | 0.00000000 |
| 2856 | 30.12 | 0.00000000 |
| 2857 | 27.88 | 0.00000000 |
| 2858 | 21.88 | 0.00000000 |
| 2859 | 25.95 | 0.00000000 |
| 2860 | 26.89 | 0.00000000 |

| | | |
|---|---|---|
| 2861 | 24.50 | 0.00000000 |
| 2862 | 39.56 | 0.00000000 |
| 2863 | 37.61 | 0.00000000 |
| 2864 | 25.06 | 0.00000000 |
| 2865 | 22.57 | 0.00000000 |
| 2866 | 24.05 | 0.00000000 |
| 2867 | 28.15 | 0.00000000 |
| 2868 | 26.95 | 0.00000000 |
| 2869 | 42.72 | 0.00000000 |
| 2870 | 26.21 | 0.00000000 |
| 2871 | 24.41 | 0.00000000 |
| 2872 | 23.39 | 0.00000000 |
| 2873 | 17.75 | 0.00000000 |
| 2874 | 46.43 | 0.00000000 |
| 2875 | 28.88 | 0.00000000 |
| 2876 | 42.63 | 0.00000000 |
| 2877 | 28.65 | 0.00000000 |
| 2878 | 35.24 | 0.00000000 |
| 2879 | 37.40 | 0.00000000 |
| 2880 | 21.13 | 0.00000000 |
| 2881 | 23.65 | 0.00000000 |
| 2882 | 23.56 | 0.00000000 |
| 2883 | 21.28 | 0.00000000 |
| 2884 | 42.40 | 0.00000000 |
| 2885 | 30.44 | 0.00000000 |
| 2886 | 31.69 | 0.00000000 |
| 2887 | 46.86 | 0.00000000 |
| 2888 | 46.69 | 0.00000000 |
| 2889 | 29.69 | 0.00000000 |
| 2890 | 20.82 | 0.00000000 |
| 2891 | 23.68 | 0.00000000 |
| 2892 | 28.89 | 0.00000000 |
| 2893 | 34.13 | 0.00000000 |
| 2894 | 20.22 | 0.00000000 |
| 2895 | 38.57 | 0.00000000 |
| 2896 | 21.28 | 0.00000000 |
| 2897 | 34.22 | 0.00000000 |
| 2898 | 34.00 | 0.00000000 |
| 2899 | 20.55 | 0.00000000 |
| 2900 | 18.12 | 0.00000000 |
| 2901 | 24.00 | 0.00000000 |
| 2902 | 25.35 | 0.00000000 |
| 2903 | 46.57 | 0.00000000 |
| 2904 | 18.15 | 0.00000000 |

| | | |
|---|---|---|
| 2905 | 35.29 | 0.00000000 |
| 2906 | 21.29 | 0.00000000 |
| 2907 | 28.19 | 0.00000000 |
| 2908 | 27.00 | 0.00000000 |
| 2909 | 20.80 | 0.00000000 |
| 2910 | 19.50 | 0.00000000 |
| 2911 | 38.67 | 0.00000000 |
| 2912 | 26.56 | 0.00000000 |
| 2913 | 30.17 | 0.00000000 |
| 2914 | 46.00 | 0.00000000 |
| 2915 | 28.13 | 0.00000000 |
| 2916 | 22.94 | 0.00000000 |
| 2917 | 23.00 | 0.00000000 |
| 2918 | 21.32 | 0.00000000 |
| 2919 | 18.06 | 0.00000000 |
| 2920 | 19.37 | 0.00000000 |
| 2921 | 25.44 | 0.00000000 |
| 2922 | 24.06 | 0.00000000 |
| 2923 | 20.44 | 0.00000000 |
| 2924 | 27.00 | 0.00000000 |
| 2925 | 35.75 | 0.00000000 |
| 2926 | 19.07 | 0.00000000 |
| 2927 | 31.44 | 0.00000000 |
| 2928 | 28.00 | 0.00000000 |
| 2929 | 30.35 | 0.00000000 |
| 2930 | 34.93 | 0.00000000 |
| 2931 | 27.93 | 0.00000000 |
| 2932 | 20.88 | 0.00000000 |
| 2933 | 31.27 | 0.00000000 |
| 2934 | 32.47 | 0.00000000 |
| 2935 | 42.73 | 0.00000000 |
| 2936 | 49.21 | 0.00000000 |
| 2937 | 30.24 | 0.00000000 |
| 2938 | 28.87 | 0.00000000 |
| 2939 | 35.13 | 0.00000000 |
| 2940 | 39.92 | 0.00000000 |
| 2941 | 34.62 | 0.00000000 |
| 2942 | 29.53 | 0.00000000 |
| 2943 | 17.29 | 0.00000000 |
| 2944 | 30.59 | 0.00000000 |
| 2945 | 30.40 | 0.00000000 |
| 2946 | 26.17 | 0.00000000 |
| 2947 | 30.22 | 0.00000000 |
| 2948 | 31.07 | 0.00000000 |

| | | |
|---|---|---|
| 2949 | 31.93 | 0.00000000 |
| 2950 | 42.93 | 0.00000000 |
| 2951 | 42.86 | 0.00000000 |
| 2952 | 19.89 | 0.00000000 |
| 2953 | 21.89 | 0.00000000 |
| 2954 | 14.95 | 0.00000000 |
| 2955 | 38.27 | 0.00000000 |
| 2956 | 18.56 | 0.00000000 |
| 2957 | 24.00 | 0.00000000 |
| 2958 | 38.39 | 0.00000000 |
| 2959 | 25.42 | 0.00000000 |
| 2960 | 23.42 | 0.00000000 |
| 2961 | 32.07 | 0.00000000 |
| 2962 | 23.50 | 0.00000000 |
| 2963 | 23.76 | 0.00000000 |
| 2964 | 34.53 | 0.00000000 |
| 2965 | 19.74 | 0.00000000 |
| 2966 | 27.44 | 0.00000000 |
| 2967 | 26.00 | 0.00000000 |
| 2968 | 14.27 | 0.00000000 |
| 2969 | 32.73 | 0.00000000 |
| 2970 | 31.28 | 0.00000000 |
| 2971 | 24.00 | 0.00000000 |
| 2972 | 21.69 | 0.00000000 |
| 2973 | 29.13 | 0.00000000 |
| 2974 | 35.40 | 0.00000000 |
| 2975 | 13.00 | 0.00000000 |
| 2976 | 25.72 | 0.00000000 |
| 2977 | 21.83 | 0.00000000 |
| 2978 | 28.00 | 0.00000000 |
| 2979 | 26.44 | 0.00000000 |
| 2980 | 30.41 | 0.00000000 |
| 2981 | 19.85 | 0.00000000 |
| 2982 | 25.18 | 0.00000000 |
| 2983 | 22.67 | 0.00000000 |
| 2984 | 27.75 | 0.00000000 |
| 2985 | 34.00 | 0.00000000 |
| 2986 | 31.31 | 0.00000000 |
| 2987 | 30.79 | 0.00000000 |
| 2988 | 29.71 | 0.00000000 |
| 2989 | 29.94 | 0.00000000 |
| 2990 | 22.88 | 0.00000000 |
| 2991 | 23.21 | 0.00000000 |
| 2992 | 18.65 | 0.00000000 |

| | | |
|---|---|---|
| 2993 | 20.11 | 0.00000000 |
| 2994 | 26.88 | 0.00000000 |
| 2995 | 21.19 | 0.00000000 |
| 2996 | 24.24 | 0.00000000 |
| 2997 | 35.74 | 0.00000000 |
| 2998 | 33.26 | 0.00000000 |
| 2999 | 30.71 | 0.00000000 |
| 3000 | 42.53 | 0.00000000 |
| 3001 | 39.93 | 0.00000000 |
| 3002 | 32.38 | 0.00000000 |
| 3003 | 41.24 | 0.00000000 |
| 3004 | 28.00 | 0.00000000 |
| 3005 | 20.79 | 0.00000000 |
| 3006 | 30.00 | 0.00000000 |
| 3007 | 27.06 | 0.00000000 |
| 3008 | 29.00 | 0.00000000 |
| 3009 | 39.06 | 0.00000000 |
| 3010 | 26.18 | 0.00000000 |
| 3011 | 25.58 | 0.00000000 |
| 3012 | 39.76 | 0.00000000 |
| 3013 | 21.21 | 0.00000000 |
| 3014 | 36.94 | 0.00000000 |
| 3015 | 40.73 | 0.00000000 |
| 3016 | 29.05 | 0.00000000 |
| 3017 | 26.69 | 0.00000000 |
| 3018 | 20.61 | 0.00000000 |
| 3019 | 20.27 | 0.00000000 |
| 3020 | 22.28 | 0.00000000 |
| 3021 | 30.53 | 0.00000000 |
| 3022 | 24.75 | 0.00000000 |
| 3023 | 29.15 | 0.00000000 |
| 3024 | 20.82 | 0.00000000 |
| 3025 | 21.25 | 0.00000000 |
| 3026 | 24.12 | 0.00000000 |
| 3027 | 26.93 | 0.00000000 |
| 3028 | 38.11 | 0.00000000 |
| 3029 | 16.84 | 0.00000000 |
| 3030 | 24.14 | 0.00000000 |
| 3031 | 28.26 | 0.00000000 |
| 3032 | 10.88 | 0.00000000 |
| 3033 | 31.39 | 0.00000000 |
| 3034 | 32.39 | 0.00000000 |
| 3035 | 27.44 | 0.00000000 |
| 3036 | 32.39 | 0.00000000 |

| | | |
|---|---|---|
| 3037 | 23.00 | 0.00000000 |
| 3038 | 18.41 | 0.00000000 |
| 3039 | 28.95 | 0.00000000 |
| 3040 | 31.35 | 0.00000000 |
| 3041 | 30.06 | 0.00000000 |
| 3042 | 25.35 | 0.00000000 |
| 3043 | 27.71 | 0.00000000 |
| 3044 | 23.53 | 0.00000000 |
| 3045 | 19.44 | 0.00000000 |
| 3046 | 36.78 | 0.00000000 |
| 3047 | 29.35 | 0.00000000 |
| 3048 | 25.05 | 0.00000000 |
| 3049 | 21.44 | 0.00000000 |
| 3050 | 27.07 | 0.00000000 |
| 3051 | 31.67 | 0.00000000 |
| 3052 | 30.00 | 0.00000000 |
| 3053 | 26.53 | 0.00000000 |
| 3054 | 16.37 | 0.00000000 |
| 3055 | 42.94 | 0.00000000 |
| 3056 | 14.61 | 0.00000000 |
| 3057 | 18.05 | 0.00000000 |
| 3058 | 31.72 | 0.00000000 |
| 3059 | 22.00 | 0.00000000 |
| 3060 | 45.79 | 0.00000000 |
| 3061 | 41.14 | 0.00000000 |
| 3062 | 21.27 | 0.00000000 |
| 3063 | 17.95 | 0.00000000 |
| 3064 | 19.41 | 0.00000000 |
| 3065 | 27.83 | 0.00000000 |
| 3066 | 29.38 | 0.00000000 |
| 3067 | 27.23 | 0.00000000 |
| 3068 | 14.13 | 0.00000000 |
| 3069 | 35.94 | 0.00000000 |
| 3070 | 32.06 | 0.00000000 |
| 3071 | 22.25 | 0.00000000 |
| 3072 | 23.13 | 0.00000000 |
| 3073 | 28.00 | 0.00000000 |
| 3074 | 26.73 | 0.00000000 |
| 3075 | 31.27 | 0.00000000 |
| 3076 | 36.83 | 0.00000000 |
| 3077 | 23.56 | 0.00000000 |
| 3078 | 30.54 | 0.00000000 |
| 3079 | 24.47 | 0.00000000 |
| 3080 | 19.08 | 0.00000000 |

| | | |
|---|---|---|
| 3081 | 24.19 | 0.00000000 |
| 3082 | 20.62 | 0.00000000 |
| 3083 | 14.60 | 0.00000000 |
| 3084 | 20.67 | 0.00000000 |
| 3085 | 22.81 | 0.00000000 |
| 3086 | 34.00 | 0.00000000 |
| 3087 | 23.69 | 0.00000000 |
| 3088 | 20.43 | 0.00000000 |
| 3089 | 26.88 | 0.00000000 |
| 3090 | 19.83 | 0.00000000 |
| 3091 | 11.47 | 0.00000000 |
| 3092 | 24.73 | 0.00000000 |
| 3093 | 35.71 | 0.00000000 |
| 3094 | 24.00 | 0.00000000 |
| 3095 | 22.75 | 0.00000000 |
| 3096 | 30.08 | 0.00000000 |
| 3097 | 24.44 | 0.00000000 |
| 3098 | 26.16 | 0.00000000 |
| 3099 | 25.79 | 0.00000000 |
| 3100 | 27.50 | 0.00000000 |
| 3101 | 16.47 | 0.00000000 |
| 3102 | 13.25 | 0.00000000 |
| 3103 | 30.00 | 0.00000000 |
| 3104 | 27.29 | 0.00000000 |
| 3105 | 24.19 | 0.00000000 |
| 3106 | 29.72 | 0.00000000 |
| 3107 | 15.82 | 0.00000000 |
| 3108 | 28.32 | 0.00000000 |
| 3109 | 39.44 | 0.00000000 |
| 3110 | 21.13 | 0.00000000 |
| 3111 | 21.63 | 0.00000000 |
| 3112 | 18.68 | 0.00000000 |
| 3113 | 23.47 | 0.00000000 |
| 3114 | 25.39 | 0.00000000 |
| 3115 | 29.05 | 0.00000000 |
| 3116 | 20.07 | 0.00000000 |
| 3117 | 27.81 | 0.00000000 |
| 3118 | 27.12 | 0.00000000 |
| 3119 | 15.17 | 0.00000000 |
| 3120 | 21.33 | 0.00000000 |
| 3121 | 20.94 | 0.00000000 |
| 3122 | 16.21 | 0.00000000 |
| 3123 | 27.18 | 0.00000000 |
| 3124 | 24.19 | 0.00000000 |

| | | |
|---|---|---|
| 3125 | 22.81 | 0.00000000 |
| 3126 | 22.84 | 0.00000000 |
| 3127 | 33.16 | 0.00000000 |
| 3128 | 30.24 | 0.00000000 |
| 3129 | 19.12 | 0.00000000 |
| 3130 | 17.47 | 0.00000000 |
| 3131 | 27.13 | 0.00000000 |
| 3132 | 7.80 | 0.00000000 |
| 3133 | 16.47 | 0.00000000 |
| 3134 | 27.20 | 0.00000000 |
| 3135 | 28.00 | 0.00000000 |
| 3136 | 20.19 | 0.00000000 |
| 3137 | 27.79 | 0.00000000 |
| 3138 | 32.50 | 0.00000000 |
| 3139 | 35.23 | 0.00000000 |
| 3140 | 14.72 | 0.00000000 |
| 3141 | 24.13 | 0.00000000 |
| 3142 | 31.94 | 0.00000000 |
| 3143 | 17.11 | 0.00000000 |
| 3144 | 21.41 | 0.00000000 |
| 3145 | 19.19 | 0.00000000 |
| 3146 | 13.35 | 0.00000000 |
| 3147 | 22.79 | 0.00000000 |
| 3148 | 26.56 | 0.00000000 |
| 3149 | 17.36 | 0.00000000 |
| 3150 | 20.11 | 0.00000000 |
| 3151 | 27.82 | 0.00000000 |
| 3152 | 22.75 | 0.00000000 |
| 3153 | 28.89 | 0.00000000 |
| 3154 | 24.41 | 0.00000000 |
| 3155 | 14.81 | 0.00000000 |
| 3156 | 26.50 | 0.00000000 |
| 3157 | 31.21 | 0.00000000 |
| 3158 | 28.07 | 0.00000000 |
| 3159 | 20.79 | 0.00000000 |
| 3160 | 23.94 | 0.00000000 |
| 3161 | 42.36 | 0.00000000 |
| 3162 | 29.35 | 0.00000000 |
| 3163 | 28.12 | 0.00000000 |
| 3164 | 19.00 | 0.00000000 |
| 3165 | 30.94 | 0.00000000 |
| 3166 | 39.20 | 0.00000000 |
| 3167 | 21.47 | 0.00000000 |
| 3168 | 16.29 | 0.00000000 |

| | | |
|---|---|---|
| 3169 | 12.26 | 0.00000000 |
| 3170 | 14.05 | 0.00000000 |
| 3171 | 16.42 | 0.00000000 |
| 3172 | 17.35 | 0.00000000 |
| 3173 | 22.12 | 0.00000000 |
| 3174 | 10.60 | 0.00000000 |
| 3175 | 15.32 | 0.00000000 |
| 3176 | 26.84 | 0.00000000 |
| 3177 | 35.47 | 0.00000000 |
| 3178 | 19.36 | 0.00000000 |
| 3179 | 26.84 | 0.00000000 |
| 3180 | 32.50 | 0.00000000 |
| 3181 | 25.94 | 0.00000000 |
| 3182 | 22.68 | 0.00000000 |
| 3183 | 31.79 | 0.00000000 |
| 3184 | 18.06 | 0.00000000 |
| 3185 | 22.89 | 0.00000000 |
| 3186 | 33.78 | 0.00000000 |
| 3187 | 19.60 | 0.00000000 |
| 3188 | 22.85 | 0.00000000 |
| 3189 | 14.40 | 0.00000000 |
| 3190 | 26.76 | 0.00000000 |
| 3191 | 26.00 | 0.00000000 |
| 3192 | 26.15 | 0.00000000 |
| 3193 | 19.50 | 0.00000000 |
| 3194 | 23.35 | 0.00000000 |
| 3195 | 22.12 | 0.00000000 |
| 3196 | 15.56 | 0.00000000 |
| 3197 | 11.63 | 0.00000000 |
| 3198 | 19.90 | 0.00000000 |
| 3199 | 18.90 | 0.00000000 |
| 3200 | 13.30 | 0.00000000 |
| 3201 | 16.67 | 0.00000000 |
| 3202 | 13.52 | 0.00000000 |
| 3203 | 19.27 | 0.00000000 |
| 3204 | 17.89 | 0.00000000 |
| 3205 | 16.20 | 0.00000000 |
| 3206 | 28.69 | 0.00000000 |
| 3207 | 34.82 | 0.00000000 |
| 3208 | 19.00 | 0.00000000 |
| 3209 | 20.41 | 0.00000000 |
| 3210 | 26.88 | 0.00000000 |
| 3211 | 15.83 | 0.00000000 |
| 3212 | 30.25 | 0.00000000 |

| | | |
|---|---|---|
| 3213 | 31.00 | 0.00000000 |
| 3214 | 36.47 | 0.00000000 |
| 3215 | 22.19 | 0.00000000 |
| 3216 | 12.19 | 0.00000000 |
| 3217 | 25.13 | 0.00000000 |
| 3218 | 15.05 | 0.00000000 |
| 3219 | 20.57 | 0.00000000 |
| 3220 | 12.93 | 0.00000000 |
| 3221 | 31.00 | 0.00000000 |
| 3222 | 17.31 | 0.00000000 |
| 3223 | 25.00 | 0.00000000 |
| 3224 | 27.41 | 0.00000000 |
| 3225 | 27.50 | 0.00000000 |
| 3226 | 28.06 | 0.00000000 |
| 3227 | 22.60 | 0.00000000 |
| 3228 | 16.29 | 0.00000000 |
| 3229 | 28.80 | 0.00000000 |
| 3230 | 24.71 | 0.00000000 |
| 3231 | 28.92 | 0.00000000 |
| 3232 | 23.38 | 0.00000000 |
| 3233 | 19.06 | 0.00000000 |
| 3234 | 28.80 | 0.00000000 |
| 3235 | 17.00 | 0.00000000 |
| 3236 | 16.05 | 0.00000000 |
| 3237 | 18.58 | 0.00000000 |
| 3238 | 23.71 | 0.00000000 |
| 3239 | 25.20 | 0.00000000 |
| 3240 | 25.14 | 0.00000000 |
| 3241 | 17.50 | 0.00000000 |
| 3242 | 14.56 | 0.00000000 |
| 3243 | 15.81 | 0.00000000 |
| 3244 | 13.25 | 0.00000000 |
| 3245 | 33.71 | 0.00000000 |
| 3246 | 16.19 | 0.00000000 |
| 3247 | 22.56 | 0.00000000 |
| 3248 | 26.94 | 0.00000000 |
| 3249 | 16.27 | 0.00000000 |
| 3250 | 20.93 | 0.00000000 |
| 3251 | 23.50 | 0.00000000 |
| 3252 | 20.13 | 0.00000000 |
| 3253 | 24.53 | 0.00000000 |
| 3254 | 18.53 | 0.00000000 |
| 3255 | 35.44 | 0.00000000 |
| 3256 | 24.06 | 0.00000000 |

| | | |
|---|---|---|
| 3257 | 26.88 | 0.00000000 |
| 3258 | 18.35 | 0.00000000 |
| 3259 | 35.80 | 0.00000000 |
| 3260 | 27.53 | 0.00000000 |
| 3261 | 20.67 | 0.00000000 |
| 3262 | 35.82 | 0.00000000 |
| 3263 | 16.79 | 0.00000000 |
| 3264 | 28.20 | 0.00000000 |
| 3265 | 21.83 | 0.00000000 |
| 3266 | 29.41 | 0.00000000 |
| 3267 | 13.00 | 0.00000000 |
| 3268 | 25.38 | 0.00000000 |
| 3269 | 26.27 | 0.00000000 |
| 3270 | 20.11 | 0.00000000 |
| 3271 | 30.73 | 0.00000000 |
| 3272 | 22.31 | 0.00000000 |
| 3273 | 18.08 | 0.00000000 |
| 3274 | 15.63 | 0.00000000 |
| 3275 | 23.53 | 0.00000000 |
| 3276 | 28.28 | 0.00000000 |
| 3277 | 25.88 | 0.00000000 |
| 3278 | 22.00 | 0.00000000 |
| 3279 | 23.47 | 0.00000000 |
| 3280 | 21.24 | 0.00000000 |
| 3281 | 34.73 | 0.00000000 |
| 3282 | 31.88 | 0.00000000 |
| 3283 | 34.87 | 0.00000000 |
| 3284 | 16.08 | 0.00000000 |
| 3285 | 28.00 | 0.00000000 |
| 3286 | 12.76 | 0.00000000 |
| 3287 | 20.41 | 0.00000000 |
| 3288 | 15.65 | 0.00000000 |
| 3289 | 18.35 | 0.00000000 |
| 3290 | 9.93 | 0.00000000 |
| 3291 | 31.05 | 0.00000000 |
| 3292 | 29.78 | 0.00000000 |
| 3293 | 18.41 | 0.00000000 |
| 3294 | 15.11 | 0.00000000 |
| 3295 | 29.27 | 0.00000000 |
| 3296 | 28.79 | 0.00000000 |
| 3297 | 13.35 | 0.00000000 |
| 3298 | 19.47 | 0.00000000 |
| 3299 | 21.47 | 0.00000000 |
| 3300 | 20.87 | 0.00000000 |

| | | |
|---|---|---|
| 3301 | 22.93 | 0.00000000 |
| 3302 | 17.50 | 0.00000000 |
| 3303 | 24.12 | 0.00000000 |
| 3304 | 19.42 | 0.00000000 |
| 3305 | 26.40 | 0.00000000 |
| 3306 | 21.88 | 0.00000000 |
| 3307 | 16.21 | 0.00000000 |
| 3308 | 23.71 | 0.00000000 |
| 3309 | 31.71 | 0.00000000 |
| 3310 | 24.00 | 0.00000000 |
| 3311 | 26.64 | 0.00000000 |
| 3312 | 15.69 | 0.00000000 |
| 3313 | 24.23 | 0.00000000 |
| 3314 | 14.81 | 0.00000000 |
| 3315 | 23.67 | 0.00000000 |
| 3316 | 35.13 | 0.00000000 |
| 3317 | 13.46 | 0.00000000 |
| 3318 | 18.47 | 0.00000000 |
| 3319 | 18.82 | 0.00000000 |
| 3320 | 21.60 | 0.00000000 |
| 3321 | 26.38 | 0.00000000 |
| 3322 | 20.53 | 0.00000000 |
| 3323 | 12.50 | 0.00000000 |
| 3324 | 25.06 | 0.00000000 |
| 3325 | 30.92 | 0.00000000 |
| 3326 | 16.24 | 0.00000000 |
| 3327 | 13.14 | 0.00000000 |
| 3328 | 31.07 | 0.00000000 |
| 3329 | 15.12 | 0.00000000 |
| 3330 | 14.47 | 0.00000000 |
| 3331 | 16.50 | 0.00000000 |
| 3332 | 19.65 | 0.00000000 |
| 3333 | 12.63 | 0.00000000 |
| 3334 | 27.12 | 0.00000000 |
| 3335 | 20.63 | 0.00000000 |
| 3336 | 22.83 | 0.00000000 |
| 3337 | 20.35 | 0.00000000 |
| 3338 | 20.19 | 0.00000000 |
| 3339 | 18.86 | 0.00000000 |
| 3340 | 27.12 | 0.00000000 |
| 3341 | 20.76 | 0.00000000 |
| 3342 | 28.15 | 0.00000000 |
| 3343 | 14.73 | 0.00000000 |
| 3344 | 17.32 | 0.00000000 |

| | | |
|---|---|---|
| 3345 | 21.13 | 0.00000000 |
| 3346 | 16.19 | 0.00000000 |
| 3347 | 9.26 | 0.00000000 |
| 3348 | 18.63 | 0.00000000 |
| 3349 | 13.31 | 0.00000000 |
| 3350 | 14.23 | 0.00000000 |
| 3351 | 21.40 | 0.00000000 |
| 3352 | 27.00 | 0.00000000 |
| 3353 | 20.81 | 0.00000000 |
| 3354 | 11.75 | 0.00000000 |
| 3355 | 25.88 | 0.00000000 |
| 3356 | 22.43 | 0.00000000 |
| 3357 | 14.86 | 0.00000000 |
| 3358 | 15.94 | 0.00000000 |
| 3359 | 22.53 | 0.00000000 |
| 3360 | 17.06 | 0.00000000 |
| 3361 | 23.25 | 0.00000000 |
| 3362 | 32.54 | 0.00000000 |
| 3363 | 29.46 | 0.00000000 |
| 3364 | 19.79 | 0.00000000 |
| 3365 | 15.47 | 0.00000000 |
| 3366 | 18.88 | 0.00000000 |
| 3367 | 15.93 | 0.00000000 |
| 3368 | 29.07 | 0.00000000 |
| 3369 | 14.30 | 0.00000000 |
| 3370 | 29.60 | 0.00000000 |
| 3371 | 25.81 | 0.00000000 |
| 3372 | 20.29 | 0.00000000 |
| 3373 | 21.82 | 0.00000000 |
| 3374 | 21.44 | 0.00000000 |
| 3375 | 21.69 | 0.00000000 |
| 3376 | 13.20 | 0.00000000 |
| 3377 | 10.16 | 0.00000000 |
| 3378 | 17.22 | 0.00000000 |
| 3379 | 8.25 | 0.00000000 |
| 3380 | 20.41 | 0.00000000 |
| 3381 | 15.47 | 0.00000000 |
| 3382 | 15.94 | 0.00000000 |
| 3383 | 15.93 | 0.00000000 |
| 3384 | 13.54 | 0.00000000 |
| 3385 | 16.06 | 0.00000000 |
| 3386 | 21.76 | 0.00000000 |
| 3387 | 22.94 | 0.00000000 |
| 3388 | 21.72 | 0.00000000 |

| | | |
|---|---|---|
| 3389 | 15.67 | 0.00000000 |
| 3390 | 26.64 | 0.00000000 |
| 3391 | 17.00 | 0.00000000 |
| 3392 | 10.62 | 0.00000000 |
| 3393 | 21.60 | 0.00000000 |
| 3394 | 15.56 | 0.00000000 |
| 3395 | 11.94 | 0.00000000 |
| 3396 | 18.79 | 0.00000000 |
| 3397 | 25.83 | 0.00000000 |
| 3398 | 24.29 | 0.00000000 |
| 3399 | 12.44 | 0.00000000 |
| 3400 | 27.44 | 0.00000000 |
| 3401 | 21.94 | 0.00000000 |
| 3402 | 29.53 | 0.00000000 |
| 3403 | 16.69 | 0.00000000 |
| 3404 | 19.00 | 0.00000000 |
| 3405 | 19.41 | 0.00000000 |
| 3406 | 23.82 | 0.00000000 |
| 3407 | 25.39 | 0.00000000 |
| 3408 | 9.68 | 0.00000000 |
| 3409 | 23.21 | 0.00000000 |
| 3410 | 22.80 | 0.00000000 |
| 3411 | 17.71 | 0.00000000 |
| 3412 | 28.45 | 0.00000000 |
| 3413 | 20.92 | 0.00000000 |
| 3414 | 16.85 | 0.00000000 |
| 3415 | 14.59 | 0.00000000 |
| 3416 | 10.20 | 0.00000000 |
| 3417 | 21.47 | 0.00000000 |
| 3418 | 11.82 | 0.00000000 |
| 3419 | 24.88 | 0.00000000 |
| 3420 | 15.87 | 0.00000000 |
| 3421 | 9.38 | 0.00000000 |
| 3422 | 14.86 | 0.00000000 |
| 3423 | 12.93 | 0.00000000 |
| 3424 | 24.60 | 0.00000000 |
| 3425 | 15.78 | 0.00000000 |
| 3426 | 33.14 | 0.00000000 |
| 3427 | 19.69 | 0.00000000 |
| 3428 | 12.93 | 0.00000000 |
| 3429 | 14.61 | 0.00000000 |
| 3430 | 18.38 | 0.00000000 |
| 3431 | 11.33 | 0.00000000 |
| 3432 | 26.88 | 0.00000000 |

| | | |
|---|---|---|
| 3433 | 21.21 | 0.00000000 |
| 3434 | 29.39 | 0.00000000 |
| 3435 | 22.12 | 0.00000000 |
| 3436 | 28.21 | 0.00000000 |
| 3437 | 23.47 | 0.00000000 |
| 3438 | 29.23 | 0.00000000 |
| 3439 | 14.00 | 0.00000000 |
| 3440 | 19.29 | 0.00000000 |
| 3441 | 26.33 | 0.00000000 |
| 3442 | 25.50 | 0.00000000 |
| 3443 | 21.06 | 0.00000000 |
| 3444 | 19.79 | 0.00000000 |
| 3445 | 22.44 | 0.00000000 |
| 3446 | 19.13 | 0.00000000 |
| 3447 | 27.46 | 0.00000000 |
| 3448 | 20.60 | 0.00000000 |
| 3449 | 11.06 | 0.00000000 |
| 3450 | 12.00 | 0.00000000 |
| 3451 | 18.88 | 0.00000000 |
| 3452 | 26.15 | 0.00000000 |
| 3453 | 22.94 | 0.00000000 |
| 3454 | 12.00 | 0.00000000 |
| 3455 | 22.87 | 0.00000000 |
| 3456 | 16.71 | 0.00000000 |
| 3457 | 10.26 | 0.00000000 |
| 3458 | 24.13 | 0.00000000 |
| 3459 | 13.42 | 0.00000000 |
| 3460 | 22.50 | 0.00000000 |
| 3461 | 15.95 | 0.00000000 |
| 3462 | 18.29 | 0.00000000 |
| 3463 | 10.16 | 0.00000000 |
| 3464 | 16.93 | 0.00000000 |
| 3465 | 30.05 | 0.00000000 |
| 3466 | 25.69 | 0.00000000 |
| 3467 | 10.35 | 0.00000000 |
| 3468 | 18.71 | 0.00000000 |
| 3469 | 23.13 | 0.00000000 |
| 3470 | 17.88 | 0.00000000 |
| 3471 | 14.53 | 0.00000000 |
| 3472 | 16.28 | 0.00000000 |
| 3473 | 26.44 | 0.00000000 |
| 3474 | 20.73 | 0.00000000 |
| 3475 | 22.53 | 0.00000000 |
| 3476 | 21.78 | 0.00000000 |

| | | |
|---|---|---|
| 3477 | 13.83 | 0.00000000 |
| 3478 | 13.13 | 0.00000000 |
| 3479 | 16.22 | 0.00000000 |
| 3480 | 14.44 | 0.00000000 |
| 3481 | 32.75 | 0.00000000 |
| 3482 | 20.33 | 0.00000000 |
| 3483 | 18.89 | 0.00000000 |
| 3484 | 18.00 | 0.00000000 |
| 3485 | 24.00 | 0.00000000 |
| 3486 | 18.44 | 0.00000000 |
| 3487 | 21.75 | 0.00000000 |
| 3488 | 15.36 | 0.00000000 |
| 3489 | 17.06 | 0.00000000 |
| 3490 | 12.44 | 0.00000000 |
| 3491 | 29.71 | 0.00000000 |
| 3492 | 11.25 | 0.00000000 |
| 3493 | 20.50 | 0.00000000 |
| 3494 | 23.53 | 0.00000000 |
| 3495 | 32.00 | 0.00000000 |
| 3496 | 13.35 | 0.00000000 |
| 3497 | 20.43 | 0.00000000 |
| 3498 | 17.52 | 0.00000000 |
| 3499 | 17.12 | 0.00000000 |
| 3500 | 25.25 | 0.00000000 |
| 3501 | 15.71 | 0.00000000 |
| 3502 | 12.05 | 0.00000000 |
| 3503 | 11.60 | 0.00000000 |
| 3504 | 26.69 | 0.00000000 |
| 3505 | 28.50 | 0.00000000 |
| 3506 | 15.29 | 0.00000000 |
| 3507 | 13.53 | 0.00000000 |
| 3508 | 10.88 | 0.00000000 |
| 3509 | 31.72 | 0.00000000 |
| 3510 | 19.33 | 0.00000000 |
| 3511 | 22.47 | 0.00000000 |
| 3512 | 14.42 | 0.00000000 |
| 3513 | 20.76 | 0.00000000 |
| 3514 | 16.13 | 0.00000000 |
| 3515 | 17.22 | 0.00000000 |
| 3516 | 21.82 | 0.00000000 |
| 3517 | 21.58 | 0.00000000 |
| 3518 | 15.81 | 0.00000000 |
| 3519 | 12.61 | 0.00000000 |
| 3520 | 22.94 | 0.00000000 |

| | | |
|---|---|---|
| 3521 | 17.88 | 0.00000000 |
| 3522 | 19.75 | 0.00000000 |
| 3523 | 23.29 | 0.00000000 |
| 3524 | 15.71 | 0.00000000 |
| 3525 | 16.88 | 0.00000000 |
| 3526 | 15.57 | 0.00000000 |
| 3527 | 34.29 | 0.00000000 |
| 3528 | 18.79 | 0.00000000 |
| 3529 | 14.13 | 0.00000000 |
| 3530 | 8.15 | 0.00000000 |
| 3531 | 28.38 | 0.00000000 |
| 3532 | 25.50 | 0.00000000 |
| 3533 | 11.80 | 0.00000000 |
| 3534 | 26.80 | 0.00000000 |
| 3535 | 9.87 | 0.00000000 |
| 3536 | 20.07 | 0.00000000 |
| 3537 | 14.00 | 0.00000000 |
| 3538 | 15.86 | 0.00000000 |
| 3539 | 26.33 | 0.00000000 |
| 3540 | 19.80 | 0.00000000 |
| 3541 | 13.31 | 0.00000000 |
| 3542 | 25.14 | 0.00000000 |
| 3543 | 25.44 | 0.00000000 |
| 3544 | 18.86 | 0.00000000 |
| 3545 | 25.25 | 0.00000000 |
| 3546 | 21.00 | 0.00000000 |
| 3547 | 13.57 | 0.00000000 |
| 3548 | 25.50 | 0.00000000 |
| 3549 | 29.46 | 0.00000000 |
| 3550 | 11.53 | 0.00000000 |
| 3551 | 14.94 | 0.00000000 |
| 3552 | 20.06 | 0.00000000 |
| 3553 | 17.07 | 0.00000000 |
| 3554 | 19.47 | 0.00000000 |
| 3555 | 29.92 | 0.00000000 |
| 3556 | 13.67 | 0.00000000 |
| 3557 | 19.08 | 0.00000000 |
| 3558 | 31.00 | 0.00000000 |
| 3559 | 30.08 | 0.00000000 |
| 3560 | 17.36 | 0.00000000 |
| 3561 | 15.53 | 0.00000000 |
| 3562 | 19.75 | 0.00000000 |
| 3563 | 17.20 | 0.00000000 |
| 3564 | 19.00 | 0.00000000 |

| | | |
|---|---|---|
| 3565 | 24.07 | 0.00000000 |
| 3566 | 24.77 | 0.00000000 |
| 3567 | 23.88 | 0.00000000 |
| 3568 | 23.62 | 0.00000000 |
| 3569 | 24.58 | 0.00000000 |
| 3570 | 29.85 | 0.00000000 |
| 3571 | 25.27 | 0.00000000 |
| 3572 | 25.83 | 0.00000000 |
| 3573 | 9.75 | 0.00000000 |
| 3574 | 22.18 | 0.00000000 |
| 3575 | 34.09 | 0.00000000 |
| 3576 | 14.07 | 0.00000000 |
| 3577 | 17.60 | 0.00000000 |
| 3578 | 25.83 | 0.00000000 |
| 3579 | 25.75 | 0.00000000 |
| 3580 | 26.57 | 0.00000000 |
| 3581 | 27.43 | 0.00000000 |
| 3582 | 20.42 | 0.00000000 |
| 3583 | 11.58 | 0.00000000 |
| 3584 | 23.40 | 0.00000000 |
| 3585 | 14.69 | 0.00000000 |
| 3586 | 33.54 | 0.00000000 |
| 3587 | 11.50 | 0.00000000 |
| 3588 | 22.00 | 0.00000000 |
| 3589 | 18.88 | 0.00000000 |
| 3590 | 24.58 | 0.00000000 |
| 3591 | 15.53 | 0.00000000 |
| 3592 | 15.53 | 0.00000000 |
| 3593 | 8.76 | 0.00000000 |
| 3594 | 27.07 | 0.00000000 |
| 3595 | 22.13 | 0.00000000 |
| 3596 | 16.87 | 0.00000000 |
| 3597 | 23.67 | 0.00000000 |
| 3598 | 18.47 | 0.00000000 |
| 3599 | 16.69 | 0.00000000 |
| 3600 | 16.18 | 0.00000000 |
| 3601 | 10.50 | 0.00000000 |
| 3602 | 26.93 | 0.00000000 |
| 3603 | 25.94 | 0.00000000 |
| 3604 | 20.57 | 0.00000000 |
| 3605 | 12.44 | 0.00000000 |
| 3606 | 19.89 | 0.00000000 |
| 3607 | 22.47 | 0.00000000 |
| 3608 | 24.24 | 0.00000000 |

| | | |
|---|---|---|
| 3609 | 20.47 | 0.00000000 |
| 3610 | 16.77 | 0.00000000 |
| 3611 | 14.13 | 0.00000000 |
| 3612 | 18.20 | 0.00000000 |
| 3613 | 31.14 | 0.00000000 |
| 3614 | 18.93 | 0.00000000 |
| 3615 | 18.67 | 0.00000000 |
| 3616 | 30.42 | 0.00000000 |
| 3617 | 22.07 | 0.00000000 |
| 3618 | 18.81 | 0.00000000 |
| 3619 | 17.82 | 0.00000000 |
| 3620 | 21.29 | 0.00000000 |
| 3621 | 15.39 | 0.00000000 |
| 3622 | 21.38 | 0.00000000 |
| 3623 | 29.28 | 0.00000000 |
| 3624 | 29.33 | 0.00000000 |
| 3625 | 15.39 | 0.00000000 |
| 3626 | 18.21 | 0.00000000 |
| 3627 | 23.79 | 0.00000000 |
| 3628 | 18.88 | 0.00000000 |
| 3629 | 15.71 | 0.00000000 |
| 3630 | 12.62 | 0.00000000 |
| 3631 | 16.94 | 0.00000000 |
| 3632 | 16.06 | 0.00000000 |
| 3633 | 17.94 | 0.00000000 |
| 3634 | 16.33 | 0.00000000 |
| 3635 | 18.47 | 0.00000000 |
| 3636 | 17.80 | 0.00000000 |
| 3637 | 6.62 | 0.00000000 |
| 3638 | 14.87 | 0.00000000 |
| 3639 | 10.93 | 0.00000000 |
| 3640 | 17.20 | 0.00000000 |
| 3641 | 29.53 | 0.00000000 |
| 3642 | 25.69 | 0.00000000 |
| 3643 | 18.81 | 0.00000000 |
| 3644 | 21.38 | 0.00000000 |
| 3645 | 20.36 | 0.00000000 |
| 3646 | 34.70 | 0.00000000 |
| 3647 | 18.17 | 0.00000000 |
| 3648 | 13.10 | 0.00000000 |
| 3649 | 23.50 | 0.00000000 |
| 3650 | 23.64 | 0.00000000 |
| 3651 | 39.20 | 0.00000000 |
| 3652 | 16.15 | 0.00000000 |

| | | |
|---|---|---|
| 3653 | 14.14 | 0.00000000 |
| 3654 | 21.50 | 0.00000000 |
| 3655 | 24.45 | 0.00000000 |
| 3656 | 22.82 | 0.00000000 |
| 3657 | 16.33 | 0.00000000 |
| 3658 | 20.77 | 0.00000000 |
| 3659 | 10.53 | 0.00000000 |
| 3660 | 16.40 | 0.00000000 |
| 3661 | 21.62 | 0.00000000 |
| 3662 | 14.87 | 0.00000000 |
| 3663 | 22.13 | 0.00000000 |
| 3664 | 19.69 | 0.00000000 |
| 3665 | 14.20 | 0.00000000 |
| 3666 | 18.00 | 0.00000000 |
| 3667 | 20.43 | 0.00000000 |
| 3668 | 19.21 | 0.00000000 |
| 3669 | 27.21 | 0.00000000 |
| 3670 | 17.53 | 0.00000000 |
| 3671 | 15.73 | 0.00000000 |
| 3672 | 16.40 | 0.00000000 |
| 3673 | 32.64 | 0.00000000 |
| 3674 | 28.75 | 0.00000000 |
| 3675 | 19.71 | 0.00000000 |
| 3676 | 24.77 | 0.00000000 |
| 3677 | 23.09 | 0.00000000 |
| 3678 | 35.46 | 0.00000000 |
| 3679 | 14.36 | 0.00000000 |
| 3680 | 11.00 | 0.00000000 |
| 3681 | 26.40 | 0.00000000 |
| 3682 | 12.92 | 0.00000000 |
| 3683 | 24.09 | 0.00000000 |
| 3684 | 29.38 | 0.00000000 |
| 3685 | 17.07 | 0.00000000 |
| 3686 | 31.93 | 0.00000000 |
| 3687 | 33.08 | 0.00000000 |
| 3688 | 20.33 | 0.00000000 |
| 3689 | 18.36 | 0.00000000 |
| 3690 | 20.38 | 0.00000000 |
| 3691 | 19.86 | 0.00000000 |
| 3692 | 14.47 | 0.00000000 |
| 3693 | 20.33 | 0.00000000 |
| 3694 | 19.71 | 0.00000000 |
| 3695 | 26.47 | 0.00000000 |
| 3696 | 6.71 | 0.00000000 |

| | | |
|---|---|---|
| 3697 | 17.05 | 0.00000000 |
| 3698 | 30.54 | 0.00000000 |
| 3699 | 19.00 | 0.00000000 |
| 3700 | 20.88 | 0.00000000 |
| 3701 | 16.80 | 0.00000000 |
| 3702 | 22.54 | 0.00000000 |
| 3703 | 21.07 | 0.00000000 |
| 3704 | 19.38 | 0.00000000 |
| 3705 | 25.59 | 0.00000000 |
| 3706 | 19.79 | 0.00000000 |
| 3707 | 18.81 | 0.00000000 |
| 3708 | 31.07 | 0.00000000 |
| 3709 | 18.86 | 0.00000000 |
| 3710 | 23.36 | 0.00000000 |
| 3711 | 9.85 | 0.00000000 |
| 3712 | 13.36 | 0.00000000 |
| 3713 | 16.31 | 0.00000000 |
| 3714 | 14.64 | 0.00000000 |
| 3715 | 20.80 | 0.00000000 |
| 3716 | 15.93 | 0.00000000 |
| 3717 | 21.36 | 0.00000000 |
| 3718 | 16.10 | 0.00000000 |
| 3719 | 8.25 | 0.00000000 |
| 3720 | 21.91 | 0.00000000 |
| 3721 | 27.27 | 0.00000000 |
| 3722 | 19.83 | 0.00000000 |
| 3723 | 17.57 | 0.00000000 |
| 3724 | 19.54 | 0.00000000 |
| 3725 | 17.46 | 0.00000000 |
| 3726 | 13.27 | 0.00000000 |
| 3727 | 15.91 | 0.00000000 |
| 3728 | 29.09 | 0.00000000 |
| 3729 | 8.58 | 0.00000000 |
| 3730 | 21.67 | 0.00000000 |
| 3731 | 21.17 | 0.00000000 |
| 3732 | 23.44 | 0.00000000 |
| 3733 | 23.88 | 0.00000000 |
| 3734 | 13.50 | 0.00000000 |
| 3735 | 17.27 | 0.00000000 |
| 3736 | 15.58 | 0.00000000 |
| 3737 | 25.36 | 0.00000000 |
| 3738 | 23.77 | 0.00000000 |
| 3739 | 30.20 | 0.00000000 |
| 3740 | 16.43 | 0.00000000 |

| | | |
|---|---|---|
| 3741 | 14.62 | 0.00000000 |
| 3742 | 20.91 | 0.00000000 |
| 3743 | 18.69 | 0.00000000 |
| 3744 | 15.00 | 0.00000000 |
| 3745 | 9.54 | 0.00000000 |
| 3746 | 21.00 | 0.00000000 |
| 3747 | 27.50 | 0.00000000 |
| 3748 | 28.21 | 0.00000000 |
| 3749 | 21.64 | 0.00000000 |
| 3750 | 17.58 | 0.00000000 |
| 3751 | 19.64 | 0.00000000 |
| 3752 | 26.58 | 0.00000000 |
| 3753 | 11.69 | 0.00000000 |
| 3754 | 10.09 | 0.00000000 |
| 3755 | 14.00 | 0.00000000 |
| 3756 | 22.83 | 0.00000000 |
| 3757 | 11.46 | 0.00000000 |
| 3758 | 17.63 | 0.00000000 |
| 3759 | 10.00 | 0.00000000 |
| 3760 | 21.50 | 0.00000000 |
| 3761 | 27.67 | 0.00000000 |
| 3762 | 18.56 | 0.00000000 |
| 3763 | 19.67 | 0.00000000 |
| 3764 | 28.33 | 0.00000000 |
| 3765 | 35.93 | 0.00000000 |
| 3766 | 21.92 | 0.00000000 |
| 3767 | 17.00 | 0.00000000 |
| 3768 | 12.64 | 0.00000000 |
| 3769 | 16.53 | 0.00000000 |
| 3770 | 16.70 | 0.00000000 |
| 3771 | 20.00 | 0.00000000 |
| 3772 | 8.57 | 0.00000000 |
| 3773 | 13.79 | 0.00000000 |
| 3774 | 11.21 | 0.00000000 |
| 3775 | 20.11 | 0.00000000 |
| 3776 | 11.50 | 0.00000000 |
| 3777 | 10.43 | 0.00000000 |
| 3778 | 13.79 | 0.00000000 |
| 3779 | 17.47 | 0.00000000 |
| 3780 | 8.36 | 0.00000000 |
| 3781 | 11.00 | 0.00000000 |
| 3782 | 14.31 | 0.00000000 |
| 3783 | 10.08 | 0.00000000 |
| 3784 | 11.81 | 0.00000000 |

| | | |
|---|---|---|
| 3785 | 27.42 | 0.00000000 |
| 3786 | 4.60 | 0.00000000 |
| 3787 | 9.50 | 0.00000000 |
| 3788 | 26.19 | 0.00000000 |
| 3789 | 17.45 | 0.00000000 |
| 3790 | 20.77 | 0.00000000 |
| 3791 | 18.81 | 0.00000000 |
| 3792 | 7.50 | 0.00000000 |
| 3793 | 17.73 | 0.00000000 |
| 3794 | 14.31 | 0.00000000 |
| 3795 | 5.83 | 0.00000000 |
| 3796 | 10.31 | 0.00000000 |
| 3797 | 18.80 | 0.00000000 |
| 3798 | 13.73 | 0.00000000 |
| 3799 | 5.93 | 0.00000000 |
| 3800 | 14.73 | 0.00000000 |
| 3801 | 15.73 | 0.00000000 |
| 3802 | 15.46 | 0.00000000 |
| 3803 | 11.00 | 0.00000000 |
| 3804 | 21.35 | 0.00000000 |
| 3805 | 14.13 | 0.00000000 |
| 3806 | 15.00 | 0.00000000 |
| 3807 | 27.57 | 0.00000000 |
| 3808 | 17.59 | 0.00000000 |
| 3809 | 15.11 | 0.00000000 |
| 3810 | 24.50 | 0.00000000 |
| 3811 | 21.59 | 0.00000000 |
| 3812 | 17.44 | 0.00000000 |
| 3813 | 19.00 | 0.00000000 |
| 3814 | 13.40 | 0.00000000 |
| 3815 | 17.50 | 0.00000000 |
| 3816 | 12.00 | 0.00000000 |
| 3817 | 20.80 | 0.00000000 |
| 3818 | 11.25 | 0.00000000 |
| 3819 | 13.25 | 0.00000000 |
| 3820 | 13.29 | 0.00000000 |
| 3821 | 26.50 | 0.00000000 |
| 3822 | 18.92 | 0.00000000 |
| 3823 | 15.27 | 0.00000000 |
| 3824 | 14.25 | 0.00000000 |
| 3825 | 16.60 | 0.00000000 |
| 3826 | 26.00 | 0.00000000 |
| 3827 | 16.38 | 0.00000000 |
| 3828 | 17.69 | 0.00000000 |

| | | |
|---|---|---|
| 3829 | 13.31 | 0.00000000 |
| 3830 | 23.25 | 0.00000000 |
| 3831 | 8.00 | 0.00000000 |
| 3832 | 22.18 | 0.00000000 |
| 3833 | 14.80 | 0.00000000 |
| 3834 | 12.35 | 0.00000000 |
| 3835 | 16.29 | 0.00000000 |
| 3836 | 18.25 | 0.00000000 |
| 3837 | 32.75 | 0.00000000 |
| 3838 | 12.50 | 0.00000000 |
| 3839 | 14.31 | 0.00000000 |
| 3840 | 20.33 | 0.00000000 |
| 3841 | 11.44 | 0.00000000 |
| 3842 | 18.50 | 0.00000000 |
| 3843 | 25.57 | 0.00000000 |
| 3844 | 26.90 | 0.00000000 |
| 3845 | 15.64 | 0.00000000 |
| 3846 | 20.85 | 0.00000000 |
| 3847 | 20.08 | 0.00000000 |
| 3848 | 35.60 | 0.00000000 |
| 3849 | 10.77 | 0.00000000 |
| 3850 | 13.63 | 0.00000000 |
| 3851 | 27.20 | 0.00000000 |
| 3852 | 15.00 | 0.00000000 |
| 3853 | 22.08 | 0.00000000 |
| 3854 | 15.22 | 0.00000000 |
| 3855 | 11.25 | 0.00000000 |
| 3856 | 6.21 | 0.00000000 |
| 3857 | 9.23 | 0.00000000 |
| 3858 | 14.07 | 0.00000000 |
| 3859 | 9.13 | 0.00000000 |
| 3860 | 12.59 | 0.00000000 |
| 3861 | 16.53 | 0.00000000 |
| 3862 | 16.14 | 0.00000000 |
| 3863 | 28.93 | 0.00000000 |
| 3864 | 11.00 | 0.00000000 |
| 3865 | 12.46 | 0.00000000 |
| 3866 | 18.25 | 0.00000000 |
| 3867 | 12.57 | 0.00000000 |
| 3868 | 21.93 | 0.00000000 |
| 3869 | 24.20 | 0.00000000 |
| 3870 | 17.73 | 0.00000000 |
| 3871 | 17.43 | 0.00000000 |
| 3872 | 23.42 | 0.00000000 |

| | | |
|---|---|---|
| 3873 | 22.86 | 0.00000000 |
| 3874 | 19.67 | 0.00000000 |
| 3875 | 24.21 | 0.00000000 |
| 3876 | 8.21 | 0.00000000 |
| 3877 | 15.33 | 0.00000000 |
| 3878 | 21.00 | 0.00000000 |
| 3879 | 15.69 | 0.00000000 |
| 3880 | 24.43 | 0.00000000 |
| 3881 | 17.00 | 0.00000000 |
| 3882 | 21.38 | 0.00000000 |
| 3883 | 22.71 | 0.00000000 |
| 3884 | 6.75 | 0.00000000 |
| 3885 | 11.33 | 0.00000000 |
| 3886 | 14.25 | 0.00000000 |
| 3887 | 24.82 | 0.00000000 |
| 3888 | 9.33 | 0.00000000 |
| 3889 | 6.86 | 0.00000000 |
| 3890 | 9.33 | 0.00000000 |
| 3891 | 16.15 | 0.00000000 |
| 3892 | 19.27 | 0.00000000 |
| 3893 | 18.64 | 0.00000000 |
| 3894 | 15.85 | 0.00000000 |
| 3895 | 19.08 | 0.00000000 |
| 3896 | 11.85 | 0.00000000 |
| 3897 | 13.40 | 0.00000000 |
| 3898 | 26.92 | 0.00000000 |
| 3899 | 21.25 | 0.00000000 |
| 3900 | 27.15 | 0.00000000 |
| 3901 | 23.94 | 0.00000000 |
| 3902 | 14.82 | 0.00000000 |
| 3903 | 12.55 | 0.00000000 |
| 3904 | 10.77 | 0.00000000 |
| 3905 | 9.80 | 0.00000000 |
| 3906 | 20.00 | 0.00000000 |
| 3907 | 19.82 | 0.00000000 |
| 3908 | 17.17 | 0.00000000 |
| 3909 | 36.50 | 0.00000000 |
| 3910 | 11.79 | 0.00000000 |
| 3911 | 15.18 | 0.00000000 |
| 3912 | 20.00 | 0.00000000 |
| 3913 | 27.50 | 0.00000000 |
| 3914 | 20.42 | 0.00000000 |
| 3915 | 22.86 | 0.00000000 |
| 3916 | 14.00 | 0.00000000 |

| | | |
|---|---|---|
| 3917 | 14.87 | 0.00000000 |
| 3918 | 19.33 | 0.00000000 |
| 3919 | 22.67 | 0.00000000 |
| 3920 | 15.09 | 0.00000000 |
| 3921 | 15.67 | 0.00000000 |
| 3922 | 16.85 | 0.00000000 |
| 3923 | 22.62 | 0.00000000 |
| 3924 | 25.62 | 0.00000000 |
| 3925 | 12.31 | 0.00000000 |
| 3926 | 19.00 | 0.00000000 |
| 3927 | 6.79 | 0.00000000 |
| 3928 | 27.33 | 0.00000000 |
| 3929 | 16.93 | 0.00000000 |
| 3930 | 21.00 | 0.00000000 |
| 3931 | 14.83 | 0.00000000 |
| 3932 | 11.50 | 0.00000000 |
| 3933 | 34.31 | 0.00000000 |
| 3934 | 16.71 | 0.00000000 |
| 3935 | 23.93 | 0.00000000 |
| 3936 | 21.43 | 0.00000000 |
| 3937 | 22.36 | 0.00000000 |
| 3938 | 17.55 | 0.00000000 |
| 3939 | 20.17 | 0.00000000 |
| 3940 | 10.00 | 0.00000000 |
| 3941 | 19.08 | 0.00000000 |
| 3942 | 21.64 | 0.00000000 |
| 3943 | 19.14 | 0.00000000 |
| 3944 | 19.46 | 0.00000000 |
| 3945 | 20.00 | 0.00000000 |
| 3946 | 12.11 | 0.00000000 |
| 3947 | 16.21 | 0.00000000 |
| 3948 | 18.75 | 0.00000000 |
| 3949 | 18.33 | 0.00000000 |
| 3950 | 19.62 | 0.00000000 |
| 3951 | 12.00 | 0.00000000 |
| 3952 | 20.17 | 0.00000000 |
| 3953 | 18.13 | 0.00000000 |
| 3954 | 12.69 | 0.00000000 |
| 3955 | 20.57 | 0.00000000 |
| 3956 | 10.80 | 0.00000000 |
| 3957 | 16.77 | 0.00000000 |
| 3958 | 25.50 | 0.00000000 |
| 3959 | 14.79 | 0.00000000 |
| 3960 | 8.54 | 0.00000000 |

| | | |
|---|---|---|
| 3961 | 22.75 | 0.00000000 |
| 3962 | 14.64 | 0.00000000 |
| 3963 | 15.08 | 0.00000000 |
| 3964 | 19.17 | 0.00000000 |
| 3965 | 17.50 | 0.00000000 |
| 3966 | 15.08 | 0.00000000 |
| 3967 | 19.69 | 0.00000000 |
| 3968 | 16.20 | 0.00000000 |
| 3969 | 18.62 | 0.00000000 |
| 3970 | 18.08 | 0.00000000 |
| 3971 | 18.00 | 0.00000000 |
| 3972 | 8.73 | 0.00000000 |
| 3973 | 18.07 | 0.00000000 |
| 3974 | 11.23 | 0.00000000 |
| 3975 | 12.17 | 0.00000000 |
| 3976 | 16.67 | 0.00000000 |
| 3977 | 21.23 | 0.00000000 |
| 3978 | 20.62 | 0.00000000 |
| 3979 | 15.56 | 0.00000000 |
| 3980 | 8.55 | 0.00000000 |
| 3981 | 22.55 | 0.00000000 |
| 3982 | 7.14 | 0.00000000 |
| 3983 | 4.00 | 0.00000000 |
| 3984 | 21.08 | 0.00000000 |
| 3985 | 28.85 | 0.00000000 |
| 3986 | 15.85 | 0.00000000 |
| 3987 | 14.33 | 0.00000000 |
| 3988 | 10.00 | 0.00000000 |
| 3989 | 22.86 | 0.00000000 |
| 3990 | 20.14 | 0.00000000 |
| 3991 | 17.18 | 0.00000000 |
| 3992 | 19.54 | 0.00000000 |
| 3993 | 27.17 | 0.00000000 |
| 3994 | 11.60 | 0.00000000 |
| 3995 | 22.62 | 0.00000000 |
| 3996 | 16.08 | 0.00000000 |
| 3997 | 15.00 | 0.00000000 |
| 3998 | 16.36 | 0.00000000 |
| 3999 | 18.46 | 0.00000000 |
| 4000 | 25.90 | 0.00000000 |
| 4001 | 28.07 | 0.00000000 |
| 4002 | 19.18 | 0.00000000 |
| 4003 | 13.00 | 0.00000000 |
| 4004 | 20.64 | 0.00000000 |

| | | |
|---|---|---|
| 4005 | 4.64 | 0.00000000 |
| 4006 | 17.55 | 0.00000000 |
| 4007 | 12.33 | 0.00000000 |
| 4008 | 12.93 | 0.00000000 |
| 4009 | 16.08 | 0.00000000 |
| 4010 | 15.75 | 0.00000000 |
| 4011 | 18.80 | 0.00000000 |
| 4012 | 15.54 | 0.00000000 |
| 4013 | 13.71 | 0.00000000 |
| 4014 | 16.45 | 0.00000000 |
| 4015 | 11.25 | 0.00000000 |
| 4016 | 26.54 | 0.00000000 |
| 4017 | 14.93 | 0.00000000 |
| 4018 | 15.92 | 0.00000000 |
| 4019 | 20.29 | 0.00000000 |
| 4020 | 14.08 | 0.00000000 |
| 4021 | 16.24 | 0.00000000 |
| 4022 | 22.79 | 0.00000000 |
| 4023 | 17.31 | 0.00000000 |
| 4024 | 14.00 | 0.00000000 |
| 4025 | 2.91 | 0.00000000 |
| 4026 | 11.07 | 0.00000000 |
| 4027 | 20.42 | 0.00000000 |
| 4028 | 17.92 | 0.00000000 |
| 4029 | 21.54 | 0.00000000 |
| 4030 | 18.69 | 0.00000000 |
| 4031 | 11.45 | 0.00000000 |
| 4032 | 10.13 | 0.00000000 |
| 4033 | 15.28 | 0.00000000 |
| 4034 | 5.79 | 0.00000000 |
| 4035 | 6.75 | 0.00000000 |
| 4036 | 15.62 | 0.00000000 |
| 4037 | 12.38 | 0.00000000 |
| 4038 | 13.08 | 0.00000000 |
| 4039 | 12.08 | 0.00000000 |
| 4040 | 9.71 | 0.00000000 |
| 4041 | 19.87 | 0.00000000 |
| 4042 | 8.56 | 0.00000000 |
| 4043 | 12.53 | 0.00000000 |
| 4044 | 26.00 | 0.00000000 |
| 4045 | 10.38 | 0.00000000 |
| 4046 | 15.88 | 0.00000000 |
| 4047 | 5.56 | 0.00000000 |
| 4048 | 13.38 | 0.00000000 |

| | | |
|---|---|---|
| 4049 | 10.73 | 0.00000000 |
| 4050 | 10.73 | 0.00000000 |
| 4051 | 21.94 | 0.00000000 |
| 4052 | 12.73 | 0.00000000 |
| 4053 | 13.33 | 0.00000000 |
| 4054 | 20.92 | 0.00000000 |
| 4055 | 7.43 | 0.00000000 |
| 4056 | 20.50 | 0.00000000 |
| 4057 | 10.60 | 0.00000000 |
| 4058 | 24.00 | 0.00000000 |
| 4059 | 3.60 | 0.00000000 |
| 4060 | 16.38 | 0.00000000 |
| 4061 | 9.21 | 0.00000000 |
| 4062 | 13.40 | 0.00000000 |
| 4063 | 8.00 | 0.00000000 |
| 4064 | 8.46 | 0.00000000 |
| 4065 | 7.00 | 0.00000000 |
| 4066 | 10.15 | 0.00000000 |
| 4067 | 15.67 | 0.00000000 |
| 4068 | 22.25 | 0.00000000 |
| 4069 | 18.42 | 0.00000000 |
| 4070 | 15.08 | 0.00000000 |
| 4071 | 22.27 | 0.00000000 |
| 4072 | 20.83 | 0.00000000 |
| 4073 | 20.93 | 0.00000000 |
| 4074 | 17.53 | 0.00000000 |
| 4075 | 13.09 | 0.00000000 |
| 4076 | 13.42 | 0.00000000 |
| 4077 | 7.08 | 0.00000000 |
| 4078 | 11.06 | 0.00000000 |
| 4079 | 16.25 | 0.00000000 |
| 4080 | 12.44 | 0.00000000 |
| 4081 | 12.36 | 0.00000000 |
| 4082 | 16.54 | 0.00000000 |
| 4083 | 16.24 | 0.00000000 |
| 4084 | 17.00 | 0.00000000 |
| 4085 | 14.69 | 0.00000000 |
| 4086 | 16.00 | 0.00000000 |
| 4087 | 14.17 | 0.00000000 |
| 4088 | 16.75 | 0.00000000 |
| 4089 | 15.78 | 0.00000000 |
| 4090 | 14.73 | 0.00000000 |
| 4091 | 24.40 | 0.00000000 |
| 4092 | 10.40 | 0.00000000 |

| | | |
|---|---|---|
| 4093 | 7.85 | 0.00000000 |
| 4094 | 7.50 | 0.00000000 |
| 4095 | 6.77 | 0.00000000 |
| 4096 | 13.91 | 0.00000000 |
| 4097 | 11.14 | 0.00000000 |
| 4098 | 15.46 | 0.00000000 |
| 4099 | 12.15 | 0.00000000 |
| 4100 | 13.73 | 0.00000000 |
| 4101 | 11.00 | 0.00000000 |
| 4102 | 12.86 | 0.00000000 |
| 4103 | 10.00 | 0.00000000 |
| 4104 | 13.80 | 0.00000000 |
| 4105 | 15.15 | 0.00000000 |
| 4106 | 13.08 | 0.00000000 |
| 4107 | 14.07 | 0.00000000 |
| 4108 | 17.79 | 0.00000000 |
| 4109 | 9.09 | 0.00000000 |
| 4110 | 9.79 | 0.00000000 |
| 4111 | 10.13 | 0.00000000 |
| 4112 | 5.69 | 0.00000000 |
| 4113 | 11.18 | 0.00000000 |
| 4114 | 17.36 | 0.00000000 |
| 4115 | 8.13 | 0.00000000 |
| 4116 | 22.55 | 0.00000000 |
| 4117 | 2.62 | 0.00000000 |
| 4118 | 8.77 | 0.00000000 |
| 4119 | 16.18 | 0.00000000 |
| 4120 | 20.00 | 0.00000000 |
| 4121 | 10.31 | 0.00000000 |
| 4122 | 11.58 | 0.00000000 |
| 4123 | 12.79 | 0.00000000 |
| 4124 | 29.27 | 0.00000000 |
| 4125 | 15.42 | 0.00000000 |
| 4126 | 19.31 | 0.00000000 |
| 4127 | 20.00 | 0.00000000 |
| 4128 | 15.18 | 0.00000000 |
| 4129 | 15.47 | 0.00000000 |
| 4130 | 13.08 | 0.00000000 |
| 4131 | 15.43 | 0.00000000 |
| 4132 | 15.64 | 0.00000000 |
| 4133 | 9.87 | 0.00000000 |
| 4134 | 21.21 | 0.00000000 |
| 4135 | 8.77 | 0.00000000 |
| 4136 | 9.92 | 0.00000000 |

| | | |
|---|---|---|
| 4137 | 16.85 | 0.00000000 |
| 4138 | 20.22 | 0.00000000 |
| 4139 | 17.91 | 0.00000000 |
| 4140 | 10.27 | 0.00000000 |
| 4141 | 12.21 | 0.00000000 |
| 4142 | 17.46 | 0.00000000 |
| 4143 | 15.83 | 0.00000000 |
| 4144 | 14.29 | 0.00000000 |
| 4145 | 16.08 | 0.00000000 |
| 4146 | 9.18 | 0.00000000 |
| 4147 | 11.38 | 0.00000000 |
| 4148 | 16.09 | 0.00000000 |
| 4149 | 22.17 | 0.00000000 |
| 4150 | 12.77 | 0.00000000 |
| 4151 | 18.83 | 0.00000000 |
| 4152 | 15.94 | 0.00000000 |
| 4153 | 18.00 | 0.00000000 |
| 4154 | 11.93 | 0.00000000 |
| 4155 | 12.89 | 0.00000000 |
| 4156 | 16.80 | 0.00000000 |
| 4157 | 12.00 | 0.00000000 |
| 4158 | 14.77 | 0.00000000 |
| 4159 | 18.62 | 0.00000000 |
| 4160 | 17.00 | 0.00000000 |
| 4161 | 17.23 | 0.00000000 |
| 4162 | 12.00 | 0.00000000 |
| 4163 | 12.46 | 0.00000000 |
| 4164 | 18.00 | 0.00000000 |
| 4165 | 16.00 | 0.00000000 |
| 4166 | 11.62 | 0.00000000 |
| 4167 | 8.08 | 0.00000000 |
| 4168 | 29.75 | 0.00000000 |
| 4169 | 15.18 | 0.00000000 |
| 4170 | 6.08 | 0.00000000 |
| 4171 | 11.40 | 0.00000000 |
| 4172 | 13.85 | 0.00000000 |
| 4173 | 18.80 | 0.00000000 |
| 4174 | 14.50 | 0.00000000 |
| 4175 | 6.92 | 0.00000000 |
| 4176 | 19.08 | 0.00000000 |
| 4177 | 6.93 | 0.00000000 |
| 4178 | 20.10 | 0.00000000 |
| 4179 | 15.08 | 0.00000000 |
| 4180 | 11.33 | 0.00000000 |

| | | |
|---|---|---|
| 4181 | 10.36 | 0.00000000 |
| 4182 | 13.50 | 0.00000000 |
| 4183 | 5.93 | 0.00000000 |
| 4184 | 10.42 | 0.00000000 |
| 4185 | 7.83 | 0.00000000 |
| 4186 | 18.86 | 0.00000000 |
| 4187 | 11.80 | 0.00000000 |
| 4188 | 2.67 | 0.00000000 |
| 4189 | 20.40 | 0.00000000 |
| 4190 | 22.00 | 0.00000000 |
| 4191 | 13.55 | 0.00000000 |
| 4192 | 4.23 | 0.00000000 |
| 4193 | 16.00 | 0.00000000 |
| 4194 | 18.55 | 0.00000000 |
| 4195 | 31.58 | 0.00000000 |
| 4196 | 22.38 | 0.00000000 |
| 4197 | 16.53 | 0.00000000 |
| 4198 | 13.20 | 0.00000000 |
| 4199 | 13.60 | 0.00000000 |
| 4200 | 15.80 | 0.00000000 |
| 4201 | 18.45 | 0.00000000 |
| 4202 | 16.71 | 0.00000000 |
| 4203 | 11.25 | 0.00000000 |
| 4204 | 8.00 | 0.00000000 |
| 4205 | 16.55 | 0.00000000 |
| 4206 | 21.90 | 0.00000000 |
| 4207 | 12.67 | 0.00000000 |
| 4208 | 13.09 | 0.00000000 |
| 4209 | 10.80 | 0.00000000 |
| 4210 | 34.18 | 0.00000000 |
| 4211 | 9.36 | 0.00000000 |
| 4212 | 12.92 | 0.00000000 |
| 4213 | 15.82 | 0.00000000 |
| 4214 | 17.25 | 0.00000000 |
| 4215 | 19.92 | 0.00000000 |
| 4216 | 14.75 | 0.00000000 |
| 4217 | 17.56 | 0.00000000 |
| 4218 | 5.86 | 0.00000000 |
| 4219 | 8.93 | 0.00000000 |
| 4220 | 10.92 | 0.00000000 |
| 4221 | 11.60 | 0.00000000 |
| 4222 | 18.33 | 0.00000000 |
| 4223 | 9.71 | 0.00000000 |
| 4224 | 17.14 | 0.00000000 |

| | | |
|---|---|---|
| 4225 | 9.00 | 0.00000000 |
| 4226 | 20.00 | 0.00000000 |
| 4227 | 10.44 | 0.00000000 |
| 4228 | 15.57 | 0.00000000 |
| 4229 | 13.81 | 0.00000000 |
| 4230 | 21.36 | 0.00000000 |
| 4231 | 19.13 | 0.00000000 |
| 4232 | 20.15 | 0.00000000 |
| 4233 | 11.00 | 0.00000000 |
| 4234 | 15.83 | 0.00000000 |
| 4235 | 14.75 | 0.00000000 |
| 4236 | 15.27 | 0.00000000 |
| 4237 | 21.00 | 0.00000000 |
| 4238 | 12.71 | 0.00000000 |
| 4239 | 25.07 | 0.00000000 |
| 4240 | 12.67 | 0.00000000 |
| 4241 | 10.20 | 0.00000000 |
| 4242 | 18.54 | 0.00000000 |
| 4243 | 7.67 | 0.00000000 |
| 4244 | 16.81 | 0.00000000 |
| 4245 | 15.93 | 0.00000000 |
| 4246 | 12.27 | 0.00000000 |
| 4247 | 13.77 | 0.00000000 |
| 4248 | 8.46 | 0.00000000 |
| 4249 | 17.90 | 0.00000000 |
| 4250 | 13.56 | 0.00000000 |
| 4251 | 18.14 | 0.00000000 |
| 4252 | 15.92 | 0.00000000 |
| 4253 | 13.36 | 0.00000000 |
| 4254 | 11.00 | 0.00000000 |
| 4255 | 17.00 | 0.00000000 |
| 4256 | 15.23 | 0.00000000 |
| 4257 | 12.00 | 0.00000000 |
| 4258 | 11.15 | 0.00000000 |
| 4259 | 9.82 | 0.00000000 |
| 4260 | 11.90 | 0.00000000 |
| 4261 | 16.40 | 0.00000000 |
| 4262 | 20.18 | 0.00000000 |
| 4263 | 17.82 | 0.00000000 |
| 4264 | 9.33 | 0.00000000 |
| 4265 | 20.44 | 0.00000000 |
| 4266 | 6.92 | 0.00000000 |
| 4267 | 8.44 | 0.00000000 |
| 4268 | 8.36 | 0.00000000 |

| | | |
|---|---|---|
| 4269 | 15.45 | 0.00000000 |
| 4270 | 17.86 | 0.00000000 |
| 4271 | 8.08 | 0.00000000 |
| 4272 | 12.36 | 0.00000000 |
| 4273 | 11.18 | 0.00000000 |
| 4274 | 18.50 | 0.00000000 |
| 4275 | 16.54 | 0.00000000 |
| 4276 | 14.40 | 0.00000000 |
| 4277 | 4.80 | 0.00000000 |
| 4278 | 11.08 | 0.00000000 |
| 4279 | 15.10 | 0.00000000 |
| 4280 | 19.64 | 0.00000000 |
| 4281 | 8.31 | 0.00000000 |
| 4282 | 14.36 | 0.00000000 |
| 4283 | 14.83 | 0.00000000 |
| 4284 | 20.73 | 0.00000000 |
| 4285 | 12.00 | 0.00000000 |
| 4286 | 29.18 | 0.00000000 |
| 4287 | 20.60 | 0.00000000 |
| 4288 | 21.33 | 0.00000000 |
| 4289 | 7.90 | 0.00000000 |
| 4290 | 11.58 | 0.00000000 |
| 4291 | 11.67 | 0.00000000 |
| 4292 | 18.25 | 0.00000000 |
| 4293 | 21.42 | 0.00000000 |
| 4294 | 7.83 | 0.00000000 |
| 4295 | 6.91 | 0.00000000 |
| 4296 | 13.08 | 0.00000000 |
| 4297 | 13.58 | 0.00000000 |
| 4298 | 11.69 | 0.00000000 |
| 4299 | 19.64 | 0.00000000 |
| 4300 | 11.14 | 0.00000000 |
| 4301 | 13.56 | 0.00000000 |
| 4302 | 14.91 | 0.00000000 |
| 4303 | 13.67 | 0.00000000 |
| 4304 | 11.00 | 0.00000000 |
| 4305 | 11.36 | 0.00000000 |
| 4306 | 10.80 | 0.00000000 |
| 4307 | 27.18 | 0.00000000 |
| 4308 | 29.42 | 0.00000000 |
| 4309 | 8.42 | 0.00000000 |
| 4310 | 7.22 | 0.00000000 |
| 4311 | 2.92 | 0.00000000 |
| 4312 | 16.50 | 0.00000000 |

| | | |
|---|---|---|
| 4313 | 16.13 | 0.00000000 |
| 4314 | 19.38 | 0.00000000 |
| 4315 | 8.13 | 0.00000000 |
| 4316 | 8.50 | 0.00000000 |
| 4317 | 14.18 | 0.00000000 |
| 4318 | 6.18 | 0.00000000 |
| 4319 | 4.07 | 0.00000000 |
| 4320 | 3.25 | 0.00000000 |
| 4321 | 8.92 | 0.00000000 |
| 4322 | 3.89 | 0.00000000 |
| 4323 | 10.31 | 0.00000000 |
| 4324 | 14.80 | 0.00000000 |
| 4325 | 14.36 | 0.00000000 |
| 4326 | 6.00 | 0.00000000 |
| 4327 | 13.20 | 0.00000000 |
| 4328 | 9.15 | 0.00000000 |
| 4329 | 9.79 | 0.00000000 |
| 4330 | 21.00 | 0.00000000 |
| 4331 | 15.47 | 0.00000000 |
| 4332 | 9.00 | 0.00000000 |
| 4333 | 30.08 | 0.00000000 |
| 4334 | 16.07 | 0.00000000 |
| 4335 | 21.92 | 0.00000000 |
| 4336 | 5.08 | 0.00000000 |
| 4337 | 11.07 | 0.00000000 |
| 4338 | 16.75 | 0.00000000 |
| 4339 | 9.54 | 0.00000000 |
| 4340 | 13.33 | 0.00000000 |
| 4341 | 13.57 | 0.00000000 |
| 4342 | 12.86 | 0.00000000 |
| 4343 | 24.43 | 0.00000000 |
| 4344 | 7.30 | 0.00000000 |
| 4345 | 22.86 | 0.00000000 |
| 4346 | 9.87 | 0.00000000 |
| 4347 | 18.29 | 0.00000000 |
| 4348 | 20.92 | 0.00000000 |
| 4349 | 4.00 | 0.00000000 |
| 4350 | 15.13 | 0.00000000 |
| 4351 | 25.36 | 0.00000000 |
| 4352 | 13.15 | 0.00000000 |
| 4353 | 20.08 | 0.00000000 |
| 4354 | 15.70 | 0.00000000 |
| 4355 | 10.88 | 0.00000000 |
| 4356 | 23.55 | 0.00000000 |

| | | |
|---|---|---|
| 4357 | 22.77 | 0.00000000 |
| 4358 | 18.09 | 0.00000000 |
| 4359 | 7.85 | 0.00000000 |
| 4360 | 20.18 | 0.00000000 |
| 4361 | 17.50 | 0.00000000 |
| 4362 | 7.50 | 0.00000000 |
| 4363 | 17.64 | 0.00000000 |
| 4364 | 3.69 | 0.00000000 |
| 4365 | 18.62 | 0.00000000 |
| 4366 | 27.45 | 0.00000000 |
| 4367 | 17.08 | 0.00000000 |
| 4368 | 15.63 | 0.00000000 |
| 4369 | 6.21 | 0.00000000 |
| 4370 | 9.06 | 0.00000000 |
| 4371 | 19.07 | 0.00000000 |
| 4372 | 8.36 | 0.00000000 |
| 4373 | 15.92 | 0.00000000 |
| 4374 | 20.31 | 0.00000000 |
| 4375 | 17.25 | 0.00000000 |
| 4376 | 6.20 | 0.00000000 |
| 4377 | 10.20 | 0.00000000 |
| 4378 | 6.08 | 0.00000000 |
| 4379 | 20.75 | 0.00000000 |
| 4380 | 12.93 | 0.00000000 |
| 4381 | 9.08 | 0.00000000 |
| 4382 | 6.82 | 0.00000000 |
| 4383 | 3.44 | 0.00000000 |
| 4384 | 11.15 | 0.00000000 |
| 4385 | 13.00 | 0.00000000 |
| 4386 | 15.71 | 0.00000000 |
| 4387 | 16.93 | 0.00000000 |
| 4388 | 15.07 | 0.00000000 |
| 4389 | 17.31 | 0.00000000 |
| 4390 | 8.27 | 0.00000000 |
| 4391 | 9.79 | 0.00000000 |
| 4392 | 10.20 | 0.00000000 |
| 4393 | 22.06 | 0.00000000 |
| 4394 | 16.36 | 0.00000000 |
| 4395 | 15.00 | 0.00000000 |
| 4396 | 17.00 | 0.00000000 |
| 4397 | 3.83 | 0.00000000 |
| 4398 | 11.36 | 0.00000000 |
| 4399 | 6.38 | 0.00000000 |
| 4400 | 25.64 | 0.00000000 |

| | | |
|---|---|---|
| 4401 | 9.82 | 0.00000000 |
| 4402 | 20.40 | 0.00000000 |
| 4403 | 15.23 | 0.00000000 |
| 4404 | 10.10 | 0.00000000 |
| 4405 | 12.43 | 0.00000000 |
| 4406 | 11.75 | 0.00000000 |
| 4407 | 16.00 | 0.00000000 |
| 4408 | 19.42 | 0.00000000 |
| 4409 | 15.90 | 0.00000000 |
| 4410 | 18.23 | 0.00000000 |
| 4411 | 11.91 | 0.00000000 |
| 4412 | 15.90 | 0.00000000 |
| 4413 | 12.27 | 0.00000000 |
| 4414 | 8.50 | 0.00000000 |
| 4415 | 13.27 | 0.00000000 |
| 4416 | 14.38 | 0.00000000 |
| 4417 | 8.38 | 0.00000000 |
| 4418 | 17.31 | 0.00000000 |
| 4419 | 7.07 | 0.00000000 |
| 4420 | 12.77 | 0.00000000 |
| 4421 | 17.58 | 0.00000000 |
| 4422 | 21.73 | 0.00000000 |
| 4423 | 9.81 | 0.00000000 |
| 4424 | 37.93 | 0.00000000 |
| 4425 | 17.42 | 0.00000000 |
| 4426 | 15.07 | 0.00000000 |
| 4427 | 2.91 | 0.00000000 |
| 4428 | 18.53 | 0.00000000 |
| 4429 | 14.82 | 0.00000000 |
| 4430 | 16.38 | 0.00000000 |
| 4431 | 8.80 | 0.00000000 |
| 4432 | 15.15 | 0.00000000 |
| 4433 | 17.60 | 0.00000000 |
| 4434 | 20.23 | 0.00000000 |
| 4435 | 16.57 | 0.00000000 |
| 4436 | 11.50 | 0.00000000 |
| 4437 | 5.86 | 0.00000000 |
| 4438 | 20.91 | 0.00000000 |
| 4439 | 16.92 | 0.00000000 |
| 4440 | 13.13 | 0.00000000 |
| 4441 | 11.45 | 0.00000000 |
| 4442 | 9.08 | 0.00000000 |
| 4443 | 8.36 | 0.00000000 |
| 4444 | 7.38 | 0.00000000 |

| | | |
|---|---|---|
| 4445 | 12.86 | 0.00000000 |
| 4446 | 30.08 | 0.00000000 |
| 4447 | 5.50 | 0.00000000 |
| 4448 | 10.85 | 0.00000000 |
| 4449 | 25.40 | 0.00000000 |
| 4450 | 9.82 | 0.00000000 |
| 4451 | 13.10 | 0.00000000 |
| 4452 | 18.58 | 0.00000000 |
| 4453 | 11.36 | 0.00000000 |
| 4454 | 7.18 | 0.00000000 |
| 4455 | 14.08 | 0.00000000 |
| 4456 | 12.60 | 0.00000000 |
| 4457 | 7.14 | 0.00000000 |
| 4458 | 4.23 | 0.00000000 |
| 4459 | 16.50 | 0.00000000 |
| 4460 | 11.45 | 0.00000000 |
| 4461 | 19.69 | 0.00000000 |
| 4462 | 5.92 | 0.00000000 |
| 4463 | 12.15 | 0.00000000 |
| 4464 | 7.67 | 0.00000000 |
| 4465 | 12.44 | 0.00000000 |
| 4466 | 11.17 | 0.00000000 |
| 4467 | 10.00 | 0.00000000 |
| 4468 | 8.85 | 0.00000000 |
| 4469 | 11.13 | 0.00000000 |
| 4470 | 16.94 | 0.00000000 |
| 4471 | 14.50 | 0.00000000 |
| 4472 | 13.54 | 0.00000000 |
| 4473 | 11.56 | 0.00000000 |
| 4474 | 8.85 | 0.00000000 |
| 4475 | 10.00 | 0.00000000 |
| 4476 | 5.00 | 0.00000000 |
| 4477 | 9.93 | 0.00000000 |
| 4478 | 10.85 | 0.00000000 |
| 4479 | 12.77 | 0.00000000 |
| 4480 | 10.54 | 0.00000000 |
| 4481 | 20.00 | 0.00000000 |
| 4482 | 8.86 | 0.00000000 |
| 4483 | 19.83 | 0.00000000 |
| 4484 | 24.15 | 0.00000000 |
| 4485 | 17.42 | 0.00000000 |
| 4486 | 14.36 | 0.00000000 |
| 4487 | 16.17 | 0.00000000 |
| 4488 | 12.08 | 0.00000000 |

| | | |
|---|---|---|
| 4489 | 9.58 | 0.00000000 |
| 4490 | 16.69 | 0.00000000 |
| 4491 | 11.64 | 0.00000000 |
| 4492 | 14.43 | 0.00000000 |
| 4493 | 12.46 | 0.00000000 |
| 4494 | 13.91 | 0.00000000 |
| 4495 | 13.92 | 0.00000000 |
| 4496 | 9.00 | 0.00000000 |
| 4497 | 3.50 | 0.00000000 |
| 4498 | 15.33 | 0.00000000 |
| 4499 | 14.10 | 0.00000000 |
| 4500 | 14.50 | 0.00000000 |
| 4501 | 11.92 | 0.00000000 |
| 4502 | 11.23 | 0.00000000 |
| 4503 | 10.50 | 0.00000000 |
| 4504 | 7.67 | 0.00000000 |
| 4505 | 11.00 | 0.00000000 |
| 4506 | 21.18 | 0.00000000 |
| 4507 | 11.67 | 0.00000000 |
| 4508 | 11.91 | 0.00000000 |
| 4509 | 16.92 | 0.00000000 |
| 4510 | 7.18 | 0.00000000 |
| 4511 | 28.18 | 0.00000000 |
| 4512 | 4.09 | 0.00000000 |
| 4513 | 12.33 | 0.00000000 |
| 4514 | 12.89 | 0.00000000 |
| 4515 | 10.21 | 0.00000000 |
| 4516 | 12.15 | 0.00000000 |
| 4517 | 16.92 | 0.00000000 |
| 4518 | 22.58 | 0.00000000 |
| 4519 | 10.75 | 0.00000000 |
| 4520 | 10.18 | 0.00000000 |
| 4521 | 13.63 | 0.00000000 |
| 4522 | 10.13 | 0.00000000 |
| 4523 | 15.00 | 0.00000000 |
| 4524 | 7.70 | 0.00000000 |
| 4525 | 14.22 | 0.00000000 |
| 4526 | 19.11 | 0.00000000 |
| 4527 | 20.67 | 0.00000000 |
| 4528 | 13.08 | 0.00000000 |
| 4529 | 7.67 | 0.00000000 |
| 4530 | 19.83 | 0.00000000 |
| 4531 | 16.00 | 0.00000000 |
| 4532 | 19.09 | 0.00000000 |

| | | |
|---|---|---|
| 4533 | 30.08 | 0.00000000 |
| 4534 | 14.56 | 0.00000000 |
| 4535 | 8.64 | 0.00000000 |
| 4536 | 6.15 | 0.00000000 |
| 4537 | 20.17 | 0.00000000 |
| 4538 | 10.15 | 0.00000000 |
| 4539 | 19.82 | 0.00000000 |
| 4540 | 11.85 | 0.00000000 |
| 4541 | 20.08 | 0.00000000 |
| 4542 | 12.69 | 0.00000000 |
| 4543 | 11.08 | 0.00000000 |
| 4544 | 5.30 | 0.00000000 |
| 4545 | 20.17 | 0.00000000 |
| 4546 | 9.58 | 0.00000000 |
| 4547 | 15.62 | 0.00000000 |
| 4548 | 14.00 | 0.00000000 |
| 4549 | 11.75 | 0.00000000 |
| 4550 | 12.08 | 0.00000000 |
| 4551 | 19.00 | 0.00000000 |
| 4552 | 25.93 | 0.00000000 |
| 4553 | 26.10 | 0.00000000 |
| 4554 | 8.69 | 0.00000000 |
| 4555 | 11.20 | 0.00000000 |
| 4556 | 8.23 | 0.00000000 |
| 4557 | 9.91 | 0.00000000 |
| 4558 | 10.20 | 0.00000000 |
| 4559 | 17.11 | 0.00000000 |
| 4560 | 12.73 | 0.00000000 |
| 4561 | 9.13 | 0.00000000 |
| 4562 | 12.43 | 0.00000000 |
| 4563 | 8.64 | 0.00000000 |
| 4564 | 7.17 | 0.00000000 |
| 4565 | 14.36 | 0.00000000 |
| 4566 | 25.25 | 0.00000000 |
| 4567 | 7.92 | 0.00000000 |
| 4568 | 12.36 | 0.00000000 |
| 4569 | 9.29 | 0.00000000 |
| 4570 | 9.08 | 0.00000000 |
| 4571 | 25.00 | 0.00000000 |
| 4572 | 15.18 | 0.00000000 |
| 4573 | 6.54 | 0.00000000 |
| 4574 | 11.00 | 0.00000000 |
| 4575 | 9.08 | 0.00000000 |
| 4576 | 11.42 | 0.00000000 |

| | | |
|---|---|---|
| 4577 | 6.17 | 0.00000000 |
| 4578 | 13.45 | 0.00000000 |
| 4579 | 13.90 | 0.00000000 |
| 4580 | 21.40 | 0.00000000 |
| 4581 | 21.10 | 0.00000000 |
| 4582 | 13.64 | 0.00000000 |
| 4583 | 9.69 | 0.00000000 |
| 4584 | 30.78 | 0.00000000 |
| 4585 | 20.70 | 0.00000000 |
| 4586 | 14.85 | 0.00000000 |
| 4587 | 19.30 | 0.00000000 |
| 4588 | 18.64 | 0.00000000 |
| 4589 | 12.50 | 0.00000000 |
| 4590 | 21.33 | 0.00000000 |
| 4591 | 13.80 | 0.00000000 |
| 4592 | 28.64 | 0.00000000 |
| 4593 | 8.00 | 0.00000000 |
| 4594 | 11.23 | 0.00000000 |
| 4595 | 17.50 | 0.00000000 |
| 4596 | 15.30 | 0.00000000 |
| 4597 | 22.20 | 0.00000000 |
| 4598 | 5.55 | 0.00000000 |
| 4599 | 3.08 | 0.00000000 |
| 4600 | 12.33 | 0.00000000 |
| 4601 | 6.60 | 0.00000000 |
| 4602 | 6.50 | 0.00000000 |
| 4603 | 9.64 | 0.00000000 |
| 4604 | 7.78 | 0.00000000 |
| 4605 | 14.64 | 0.00000000 |
| 4606 | 6.60 | 0.00000000 |
| 4607 | 7.00 | 0.00000000 |
| 4608 | 13.43 | 0.00000000 |
| 4609 | 9.27 | 0.00000000 |
| 4610 | 8.83 | 0.00000000 |
| 4611 | 6.80 | 0.00000000 |
| 4612 | 14.00 | 0.00000000 |
| 4613 | 24.18 | 0.00000000 |
| 4614 | 10.60 | 0.00000000 |
| 4615 | 16.67 | 0.00000000 |
| 4616 | 15.00 | 0.00000000 |
| 4617 | 10.40 | 0.00000000 |
| 4618 | 20.00 | 0.00000000 |
| 4619 | 6.18 | 0.00000000 |
| 4620 | 13.55 | 0.00000000 |

| | | |
|---|---|---|
| 4621 | 23.22 | 0.00000000 |
| 4622 | 11.00 | 0.00000000 |
| 4623 | 5.85 | 0.00000000 |
| 4624 | 3.33 | 0.00000000 |
| 4625 | 11.38 | 0.00000000 |
| 4626 | 13.40 | 0.00000000 |
| 4627 | 16.57 | 0.00000000 |
| 4628 | 15.27 | 0.00000000 |
| 4629 | 10.50 | 0.00000000 |
| 4630 | 5.00 | 0.00000000 |
| 4631 | 10.60 | 0.00000000 |
| 4632 | 11.45 | 0.00000000 |
| 4633 | 13.25 | 0.00000000 |
| 4634 | 20.90 | 0.00000000 |
| 4635 | 13.70 | 0.00000000 |
| 4636 | 10.36 | 0.00000000 |
| 4637 | 8.50 | 0.00000000 |
| 4638 | 7.45 | 0.00000000 |
| 4639 | 15.69 | 0.00000000 |
| 4640 | 11.92 | 0.00000000 |
| 4641 | 17.33 | 0.00000000 |
| 4642 | 11.20 | 0.00000000 |
| 4643 | 13.80 | 0.00000000 |
| 4644 | 9.46 | 0.00000000 |
| 4645 | 19.73 | 0.00000000 |
| 4646 | 8.58 | 0.00000000 |
| 4647 | 18.09 | 0.00000000 |
| 4648 | 22.10 | 0.00000000 |
| 4649 | 14.00 | 0.00000000 |
| 4650 | 7.00 | 0.00000000 |
| 4651 | 7.44 | 0.00000000 |
| 4652 | 12.08 | 0.00000000 |
| 4653 | 6.18 | 0.00000000 |
| 4654 | 15.00 | 0.00000000 |
| 4655 | 13.35 | 0.00000000 |
| 4656 | 11.53 | 0.00000000 |
| 4657 | 14.00 | 0.00000000 |
| 4658 | 13.57 | 0.00000000 |
| 4659 | 19.88 | 0.00000000 |
| 4660 | 7.56 | 0.00000000 |
| 4661 | 19.13 | 0.00000000 |
| 4662 | 6.27 | 0.00000000 |
| 4663 | 10.30 | 0.00000000 |
| 4664 | 5.33 | 0.00000000 |

| | | |
|---|---|---|
| 4665 | 32.30 | 0.00000000 |
| 4666 | 23.00 | 0.00000000 |
| 4667 | 9.57 | 0.00000000 |
| 4668 | 10.82 | 0.00000000 |
| 4669 | 14.45 | 0.00000000 |
| 4670 | 8.56 | 0.00000000 |
| 4671 | 23.38 | 0.00000000 |
| 4672 | 11.27 | 0.00000000 |
| 4673 | 7.15 | 0.00000000 |
| 4674 | 32.78 | 0.00000000 |
| 4675 | 14.27 | 0.00000000 |
| 4676 | 5.70 | 0.00000000 |
| 4677 | 16.18 | 0.00000000 |
| 4678 | 17.91 | 0.00000000 |
| 4679 | 9.00 | 0.00000000 |
| 4680 | 11.20 | 0.00000000 |
| 4681 | 16.33 | 0.00000000 |
| 4682 | 13.86 | 0.00000000 |
| 4683 | 21.11 | 0.00000000 |
| 4684 | 11.44 | 0.00000000 |
| 4685 | 9.70 | 0.00000000 |
| 4686 | 13.30 | 0.00000000 |
| 4687 | 4.25 | 0.00000000 |
| 4688 | 29.44 | 0.00000000 |
| 4689 | 26.78 | 0.00000000 |
| 4690 | 7.60 | 0.00000000 |
| 4691 | 14.70 | 0.00000000 |
| 4692 | 5.13 | 0.00000000 |
| 4693 | 7.75 | 0.00000000 |
| 4694 | 14.80 | 0.00000000 |
| 4695 | 7.38 | 0.00000000 |
| 4696 | 10.45 | 0.00000000 |
| 4697 | 16.55 | 0.00000000 |
| 4698 | 7.20 | 0.00000000 |
| 4699 | 15.44 | 0.00000000 |
| 4700 | 16.09 | 0.00000000 |
| 4701 | 8.73 | 0.00000000 |
| 4702 | 26.86 | 0.00000000 |
| 4703 | 11.50 | 0.00000000 |
| 4704 | 15.50 | 0.00000000 |
| 4705 | 4.14 | 0.00000000 |
| 4706 | 18.82 | 0.00000000 |
| 4707 | 7.40 | 0.00000000 |
| 4708 | 22.71 | 0.00000000 |

| | | |
|---|---|---|
| 4709 | 11.67 | 0.00000000 |
| 4710 | 3.22 | 0.00000000 |
| 4711 | 3.67 | 0.00000000 |
| 4712 | 8.73 | 0.00000000 |
| 4713 | 13.82 | 0.00000000 |
| 4714 | 18.25 | 0.00000000 |
| 4715 | 2.00 | 0.00000000 |
| 4716 | 7.82 | 0.00000000 |
| 4717 | 12.00 | 0.00000000 |
| 4718 | 20.93 | 0.00000000 |
| 4719 | 8.70 | 0.00000000 |
| 4720 | 3.18 | 0.00000000 |
| 4721 | 14.27 | 0.00000000 |
| 4722 | 9.82 | 0.00000000 |
| 4723 | 13.86 | 0.00000000 |
| 4724 | 8.83 | 0.00000000 |
| 4725 | 5.50 | 0.00000000 |
| 4726 | 15.23 | 0.00000000 |
| 4727 | 5.62 | 0.00000000 |
| 4728 | 8.22 | 0.00000000 |
| 4729 | 15.27 | 0.00000000 |
| 4730 | 5.89 | 0.00000000 |
| 4731 | 10.73 | 0.00000000 |
| 4732 | 4.27 | 0.00000000 |
| 4733 | 22.82 | 0.00000000 |
| 4734 | 5.60 | 0.00000000 |
| 4735 | 13.78 | 0.00000000 |
| 4736 | 9.89 | 0.00000000 |
| 4737 | 13.13 | 0.00000000 |
| 4738 | 2.91 | 0.00000000 |
| 4739 | 7.70 | 0.00000000 |
| 4740 | 10.67 | 0.00000000 |
| 4741 | 9.82 | 0.00000000 |
| 4742 | 2.73 | 0.00000000 |
| 4743 | 9.00 | 0.00000000 |
| 4744 | 21.27 | 0.00000000 |
| 4745 | 8.36 | 0.00000000 |
| 4746 | 15.92 | 0.00000000 |
| 4747 | 19.62 | 0.00000000 |
| 4748 | 15.07 | 0.00000000 |
| 4749 | 3.50 | 0.00000000 |
| 4750 | 8.36 | 0.00000000 |
| 4751 | 18.40 | 0.00000000 |
| 4752 | 14.67 | 0.00000000 |

| | | |
|---|---|---|
| 4753 | 8.00 | 0.00000000 |
| 4754 | 15.00 | 0.00000000 |
| 4755 | 4.33 | 0.00000000 |
| 4756 | 4.18 | 0.00000000 |
| 4757 | 17.67 | 0.00000000 |
| 4758 | 8.33 | 0.00000000 |
| 4759 | 13.22 | 0.00000000 |
| 4760 | 15.00 | 0.00000000 |
| 4761 | 14.08 | 0.00000000 |
| 4762 | 20.47 | 0.00000000 |
| 4763 | 8.75 | 0.00000000 |
| 4764 | 7.80 | 0.00000000 |
| 4765 | 23.67 | 0.00000000 |
| 4766 | 13.10 | 0.00000000 |
| 4767 | 13.50 | 0.00000000 |
| 4768 | 18.29 | 0.00000000 |
| 4769 | 14.78 | 0.00000000 |
| 4770 | 7.80 | 0.00000000 |
| 4771 | 7.00 | 0.00000000 |
| 4772 | 6.18 | 0.00000000 |
| 4773 | 5.50 | 0.00000000 |
| 4774 | 21.50 | 0.00000000 |
| 4775 | 13.30 | 0.00000000 |
| 4776 | 3.00 | 0.00000000 |
| 4777 | 4.91 | 0.00000000 |
| 4778 | 13.88 | 0.00000000 |
| 4779 | 12.89 | 0.00000000 |
| 4780 | 9.53 | 0.00000000 |
| 4781 | 10.38 | 0.00000000 |
| 4782 | 10.50 | 0.00000000 |
| 4783 | 8.33 | 0.00000000 |
| 4784 | 9.12 | 0.00000000 |
| 4785 | 12.14 | 0.00000000 |
| 4786 | 13.64 | 0.00000000 |
| 4787 | 7.10 | 0.00000000 |
| 4788 | 11.08 | 0.00000000 |
| 4789 | 10.92 | 0.00000000 |
| 4790 | 3.92 | 0.00000000 |
| 4791 | 11.57 | 0.00000000 |
| 4792 | 9.27 | 0.00000000 |
| 4793 | 12.62 | 0.00000000 |
| 4794 | 9.36 | 0.00000000 |
| 4795 | 13.29 | 0.00000000 |
| 4796 | 16.92 | 0.00000000 |

| | | |
|---|---|---|
| 4797 | 5.30 | 0.00000000 |
| 4798 | 11.29 | 0.00000000 |
| 4799 | 18.82 | 0.00000000 |
| 4800 | 10.83 | 0.00000000 |
| 4801 | 18.17 | 0.00000000 |
| 4802 | 12.25 | 0.00000000 |
| 4803 | 5.73 | 0.00000000 |
| 4804 | 1.92 | 0.00000000 |
| 4805 | 9.91 | 0.00000000 |
| 4806 | 13.60 | 0.00000000 |
| 4807 | 4.75 | 0.00000000 |
| 4808 | 9.86 | 0.00000000 |
| 4809 | 13.10 | 0.00000000 |
| 4810 | 3.67 | 0.00000000 |
| 4811 | 7.50 | 0.00000000 |
| 4812 | 14.62 | 0.00000000 |
| 4813 | 5.86 | 0.00000000 |
| 4814 | 13.81 | 0.00000000 |
| 4815 | 4.73 | 0.00000000 |
| 4816 | 12.58 | 0.00000000 |
| 4817 | 9.75 | 0.00000000 |
| 4818 | 26.40 | 0.00000000 |
| 4819 | 5.33 | 0.00000000 |
| 4820 | 14.58 | 0.00000000 |
| 4821 | 15.78 | 0.00000000 |
| 4822 | 16.82 | 0.00000000 |
| 4823 | 9.82 | 0.00000000 |
| 4824 | 2.78 | 0.00000000 |
| 4825 | 11.31 | 0.00000000 |
| 4826 | 16.08 | 0.00000000 |
| 4827 | 11.10 | 0.00000000 |
| 4828 | 12.64 | 0.00000000 |
| 4829 | 15.67 | 0.00000000 |
| 4830 | 16.67 | 0.00000000 |
| 4831 | 15.75 | 0.00000000 |
| 4832 | 12.08 | 0.00000000 |
| 4833 | 29.00 | 0.00000000 |
| 4834 | 8.91 | 0.00000000 |
| 4835 | 13.60 | 0.00000000 |
| 4836 | 5.82 | 0.00000000 |
| 4837 | 6.00 | 0.00000000 |
| 4838 | 15.07 | 0.00000000 |
| 4839 | 14.20 | 0.00000000 |
| 4840 | 10.14 | 0.00000000 |

| | | |
|---|---|---|
| 4841 | 5.67 | 0.00000000 |
| 4842 | 15.60 | 0.00000000 |
| 4843 | 8.81 | 0.00000000 |
| 4844 | 19.55 | 0.00000000 |
| 4845 | 5.90 | 0.00000000 |
| 4846 | 10.42 | 0.00000000 |
| 4847 | 11.50 | 0.00000000 |
| 4848 | 17.15 | 0.00000000 |
| 4849 | 14.08 | 0.00000000 |
| 4850 | 5.85 | 0.00000000 |
| 4851 | 7.31 | 0.00000000 |
| 4852 | 12.36 | 0.00000000 |
| 4853 | 10.00 | 0.00000000 |
| 4854 | 10.08 | 0.00000000 |
| 4855 | 10.69 | 0.00000000 |
| 4856 | 6.67 | 0.00000000 |
| 4857 | 25.31 | 0.00000000 |
| 4858 | 7.55 | 0.00000000 |
| 4859 | 17.50 | 0.00000000 |
| 4860 | 14.10 | 0.00000000 |
| 4861 | 6.30 | 0.00000000 |
| 4862 | 12.82 | 0.00000000 |
| 4863 | 11.50 | 0.00000000 |
| 4864 | 9.18 | 0.00000000 |
| 4865 | 7.44 | 0.00000000 |
| 4866 | 9.00 | 0.00000000 |
| 4867 | 6.42 | 0.00000000 |
| 4868 | 8.25 | 0.00000000 |
| 4869 | 27.90 | 0.00000000 |
| 4870 | 10.00 | 0.00000000 |
| 4871 | 16.60 | 0.00000000 |
| 4872 | 8.91 | 0.00000000 |
| 4873 | 8.45 | 0.00000000 |
| 4874 | 13.90 | 0.00000000 |
| 4875 | 12.54 | 0.00000000 |
| 4876 | 4.36 | 0.00000000 |
| 4877 | 10.14 | 0.00000000 |
| 4878 | 11.00 | 0.00000000 |
| 4879 | 25.93 | 0.00000000 |
| 4880 | 11.91 | 0.00000000 |
| 4881 | 8.73 | 0.00000000 |
| 4882 | 7.00 | 0.00000000 |
| 4883 | 9.64 | 0.00000000 |
| 4884 | 17.64 | 0.00000000 |

| | | |
|---|---|---|
| 4885 | 8.78 | 0.00000000 |
| 4886 | 23.89 | 0.00000000 |
| 4887 | 19.67 | 0.00000000 |
| 4888 | 16.38 | 0.00000000 |
| 4889 | 11.88 | 0.00000000 |
| 4890 | 15.33 | 0.00000000 |
| 4891 | 5.88 | 0.00000000 |
| 4892 | 15.63 | 0.00000000 |
| 4893 | 13.58 | 0.00000000 |
| 4894 | 9.09 | 0.00000000 |
| 4895 | 2.67 | 0.00000000 |
| 4896 | 13.80 | 0.00000000 |
| 4897 | 9.88 | 0.00000000 |
| 4898 | 5.78 | 0.00000000 |
| 4899 | 13.00 | 0.00000000 |
| 4900 | 2.62 | 0.00000000 |
| 4901 | 16.08 | 0.00000000 |
| 4902 | 15.56 | 0.00000000 |
| 4903 | 9.62 | 0.00000000 |
| 4904 | 4.92 | 0.00000000 |
| 4905 | 4.45 | 0.00000000 |
| 4906 | 7.42 | 0.00000000 |
| 4907 | 9.09 | 0.00000000 |
| 4908 | 6.18 | 0.00000000 |
| 4909 | 12.45 | 0.00000000 |
| 4910 | 6.92 | 0.00000000 |
| 4911 | 2.80 | 0.00000000 |
| 4912 | 6.00 | 0.00000000 |
| 4913 | 13.23 | 0.00000000 |
| 4914 | 9.44 | 0.00000000 |
| 4915 | 12.79 | 0.00000000 |
| 4916 | 11.91 | 0.00000000 |
| 4917 | 5.62 | 0.00000000 |
| 4918 | 12.20 | 0.00000000 |
| 4919 | 9.00 | 0.00000000 |
| 4920 | 23.64 | 0.00000000 |
| 4921 | 15.93 | 0.00000000 |
| 4922 | 14.10 | 0.00000000 |
| 4923 | 11.67 | 0.00000000 |
| 4924 | 8.46 | 0.00000000 |
| 4925 | 8.33 | 0.00000000 |
| 4926 | 16.50 | 0.00000000 |
| 4927 | 9.82 | 0.00000000 |
| 4928 | 5.43 | 0.00000000 |

| | | |
|---|---|---|
| 4929 | 7.00 | 0.00000000 |
| 4930 | 6.55 | 0.00000000 |
| 4931 | 10.69 | 0.00000000 |
| 4932 | 9.85 | 0.00000000 |
| 4933 | 9.73 | 0.00000000 |
| 4934 | 10.15 | 0.00000000 |
| 4935 | 6.70 | 0.00000000 |
| 4936 | 9.91 | 0.00000000 |
| 4937 | 17.40 | 0.00000000 |
| 4938 | 7.82 | 0.00000000 |
| 4939 | 15.33 | 0.00000000 |
| 4940 | 9.08 | 0.00000000 |
| 4941 | 8.13 | 0.00000000 |
| 4942 | 13.27 | 0.00000000 |
| 4943 | 21.13 | 0.00000000 |
| 4944 | 5.38 | 0.00000000 |
| 4945 | 5.50 | 0.00000000 |
| 4946 | 13.14 | 0.00000000 |
| 4947 | 10.50 | 0.00000000 |
| 4948 | 9.36 | 0.00000000 |
| 4949 | 14.40 | 0.00000000 |
| 4950 | 3.10 | 0.00000000 |
| 4951 | 7.18 | 0.00000000 |
| 4952 | 18.54 | 0.00000000 |
| 4953 | 3.00 | 0.00000000 |
| 4954 | 10.15 | 0.00000000 |
| 4955 | 19.85 | 0.00000000 |
| 4956 | 7.44 | 0.00000000 |
| 4957 | 10.42 | 0.00000000 |
| 4958 | 10.56 | 0.00000000 |
| 4959 | 8.73 | 0.00000000 |
| 4960 | 16.80 | 0.00000000 |
| 4961 | 4.20 | 0.00000000 |
| 4962 | 19.90 | 0.00000000 |
| 4963 | 11.55 | 0.00000000 |
| 4964 | 11.82 | 0.00000000 |
| 4965 | 14.67 | 0.00000000 |
| 4966 | 14.00 | 0.00000000 |
| 4967 | 21.50 | 0.00000000 |
| 4968 | 7.25 | 0.00000000 |
| 4969 | 21.00 | 0.00000000 |
| 4970 | 24.44 | 0.00000000 |
| 4971 | 6.89 | 0.00000000 |
| 4972 | 9.30 | 0.00000000 |

| | | |
|---|---|---|
| 4973 | 15.50 | 0.00000000 |
| 4974 | 10.36 | 0.00000000 |
| 4975 | 16.56 | 0.00000000 |
| 4976 | 10.30 | 0.00000000 |
| 4977 | 12.13 | 0.00000000 |
| 4978 | 10.40 | 0.00000000 |
| 4979 | 6.78 | 0.00000000 |
| 4980 | 6.44 | 0.00000000 |
| 4981 | 11.22 | 0.00000000 |
| 4982 | 11.91 | 0.00000000 |
| 4983 | 7.10 | 0.00000000 |
| 4984 | 14.67 | 0.00000000 |
| 4985 | 4.27 | 0.00000000 |
| 4986 | 3.56 | 0.00000000 |
| 4987 | 20.64 | 0.00000000 |
| 4988 | 10.56 | 0.00000000 |
| 4989 | 11.67 | 0.00000000 |
| 4990 | 8.62 | 0.00000000 |
| 4991 | 10.77 | 0.00000000 |
| 4992 | 2.58 | 0.00000000 |
| 4993 | 10.09 | 0.00000000 |
| 4994 | 13.27 | 0.00000000 |
| 4995 | 12.83 | 0.00000000 |
| 4996 | 10.09 | 0.00000000 |
| 4997 | 7.92 | 0.00000000 |
| 4998 | 7.58 | 0.00000000 |
| 4999 | 11.00 | 0.00000000 |
| 5000 | 18.15 | 0.00000000 |
| 5001 | 11.00 | 0.00000000 |
| 5002 | 11.75 | 0.00000000 |
| 5003 | 13.73 | 0.00000000 |
| 5004 | 11.80 | 0.00000000 |
| 5005 | 19.18 | 0.00000000 |
| 5006 | 8.36 | 0.00000000 |
| 5007 | 10.50 | 0.00000000 |
| 5008 | 17.17 | 0.00000000 |
| 5009 | 14.00 | 0.00000000 |
| 5010 | 14.43 | 0.00000000 |
| 5011 | 9.64 | 0.00000000 |
| 5012 | 6.10 | 0.00000000 |
| 5013 | 7.55 | 0.00000000 |
| 5014 | 11.33 | 0.00000000 |
| 5015 | 6.90 | 0.00000000 |
| 5016 | 14.33 | 0.00000000 |

| | | |
|---|---|---|
| 5017 | 6.25 | 0.00000000 |
| 5018 | 9.58 | 0.00000000 |
| 5019 | 4.00 | 0.00000000 |
| 5020 | 13.70 | 0.00000000 |
| 5021 | 6.75 | 0.00000000 |
| 5022 | 17.15 | 0.00000000 |
| 5023 | 6.86 | 0.00000000 |
| 5024 | 15.33 | 0.00000000 |
| 5025 | 17.60 | 0.00000000 |
| 5026 | 14.17 | 0.00000000 |
| 5027 | 17.58 | 0.00000000 |
| 5028 | 9.80 | 0.00000000 |
| 5029 | 15.54 | 0.00000000 |
| 5030 | 7.15 | 0.00000000 |
| 5031 | 7.43 | 0.00000000 |
| 5032 | 17.38 | 0.00000000 |
| 5033 | 8.40 | 0.00000000 |
| 5034 | 8.22 | 0.00000000 |
| 5035 | 17.00 | 0.00000000 |
| 5036 | 10.00 | 0.00000000 |
| 5037 | 14.14 | 0.00000000 |
| 5038 | 5.00 | 0.00000000 |
| 5039 | 14.55 | 0.00000000 |
| 5040 | 19.50 | 0.00000000 |
| 5041 | 13.85 | 0.00000000 |
| 5042 | 7.50 | 0.00000000 |
| 5043 | 19.56 | 0.00000000 |
| 5044 | 7.86 | 0.00000000 |
| 5045 | 4.00 | 0.00000000 |
| 5046 | 4.77 | 0.00000000 |
| 5047 | 13.00 | 0.00000000 |
| 5048 | 14.13 | 0.00000000 |
| 5049 | 6.70 | 0.00000000 |
| 5050 | 17.80 | 0.00000000 |
| 5051 | 2.75 | 0.00000000 |
| 5052 | 9.29 | 0.00000000 |
| 5053 | 12.23 | 0.00000000 |
| 5054 | 24.17 | 0.00000000 |
| 5055 | 20.55 | 0.00000000 |
| 5056 | 20.38 | 0.00000000 |
| 5057 | 6.89 | 0.00000000 |
| 5058 | 5.90 | 0.00000000 |
| 5059 | 9.00 | 0.00000000 |
| 5060 | 4.11 | 0.00000000 |

| | | |
|---|---|---|
| 5061 | 11.64 | 0.00000000 |
| 5062 | 9.55 | 0.00000000 |
| 5063 | 7.55 | 0.00000000 |
| 5064 | 20.89 | 0.00000000 |
| 5065 | 17.40 | 0.00000000 |
| 5066 | 9.38 | 0.00000000 |
| 5067 | 14.33 | 0.00000000 |
| 5068 | 5.89 | 0.00000000 |
| 5069 | 17.38 | 0.00000000 |
| 5070 | 17.57 | 0.00000000 |
| 5071 | 13.00 | 0.00000000 |
| 5072 | 11.44 | 0.00000000 |
| 5073 | 13.60 | 0.00000000 |
| 5074 | 11.89 | 0.00000000 |
| 5075 | 5.22 | 0.00000000 |
| 5076 | 2.20 | 0.00000000 |
| 5077 | 9.08 | 0.00000000 |
| 5078 | 10.11 | 0.00000000 |
| 5079 | 1.90 | 0.00000000 |
| 5080 | 8.55 | 0.00000000 |
| 5081 | 6.50 | 0.00000000 |
| 5082 | 5.80 | 0.00000000 |
| 5083 | 6.90 | 0.00000000 |
| 5084 | 20.50 | 0.00000000 |
| 5085 | 1.79 | 0.00000000 |
| 5086 | 9.25 | 0.00000000 |
| 5087 | 10.62 | 0.00000000 |
| 5088 | 6.42 | 0.00000000 |
| 5089 | 12.42 | 0.00000000 |
| 5090 | 11.90 | 0.00000000 |
| 5091 | 5.36 | 0.00000000 |
| 5092 | 11.75 | 0.00000000 |
| 5093 | 21.63 | 0.00000000 |
| 5094 | 5.73 | 0.00000000 |
| 5095 | 2.73 | 0.00000000 |
| 5096 | 3.00 | 0.00000000 |
| 5097 | 16.17 | 0.00000000 |
| 5098 | 12.18 | 0.00000000 |
| 5099 | 6.89 | 0.00000000 |
| 5100 | 6.77 | 0.00000000 |
| 5101 | 13.91 | 0.00000000 |
| 5102 | 12.67 | 0.00000000 |
| 5103 | 11.22 | 0.00000000 |
| 5104 | 13.90 | 0.00000000 |

| | | |
|---|---|---|
| 5105 | 7.00 | 0.00000000 |
| 5106 | 5.77 | 0.00000000 |
| 5107 | 8.44 | 0.00000000 |
| 5108 | 15.80 | 0.00000000 |
| 5109 | 15.69 | 0.00000000 |
| 5110 | 21.63 | 0.00000000 |
| 5111 | 6.67 | 0.00000000 |
| 5112 | 12.58 | 0.00000000 |
| 5113 | 5.60 | 0.00000000 |
| 5114 | 5.13 | 0.00000000 |
| 5115 | 8.33 | 0.00000000 |
| 5116 | 10.00 | 0.00000000 |
| 5117 | 18.33 | 0.00000000 |
| 5118 | 9.67 | 0.00000000 |
| 5119 | 19.45 | 0.00000000 |
| 5120 | 13.00 | 0.00000000 |
| 5121 | 17.73 | 0.00000000 |
| 5122 | 15.00 | 0.00000000 |
| 5123 | 7.00 | 0.00000000 |
| 5124 | 14.88 | 0.00000000 |
| 5125 | 20.54 | 0.00000000 |
| 5126 | 14.46 | 0.00000000 |
| 5127 | 12.79 | 0.00000000 |
| 5128 | 14.38 | 0.00000000 |
| 5129 | 14.17 | 0.00000000 |
| 5130 | 8.07 | 0.00000000 |
| 5131 | 12.83 | 0.00000000 |
| 5132 | 6.38 | 0.00000000 |
| 5133 | 5.67 | 0.00000000 |
| 5134 | 3.22 | 0.00000000 |
| 5135 | 9.89 | 0.00000000 |
| 5136 | 8.08 | 0.00000000 |
| 5137 | 5.54 | 0.00000000 |
| 5138 | 8.00 | 0.00000000 |
| 5139 | 9.91 | 0.00000000 |
| 5140 | 5.64 | 0.00000000 |
| 5141 | 17.55 | 0.00000000 |
| 5142 | 11.80 | 0.00000000 |
| 5143 | 15.73 | 0.00000000 |
| 5144 | 9.40 | 0.00000000 |
| 5145 | 7.64 | 0.00000000 |
| 5146 | 15.54 | 0.00000000 |
| 5147 | 7.44 | 0.00000000 |
| 5148 | 15.17 | 0.00000000 |

| | | |
|---|---|---|
| 5149 | 6.69 | 0.00000000 |
| 5150 | 10.83 | 0.00000000 |
| 5151 | 20.00 | 0.00000000 |
| 5152 | 14.18 | 0.00000000 |
| 5153 | 30.29 | 0.00000000 |
| 5154 | 5.33 | 0.00000000 |
| 5155 | 14.10 | 0.00000000 |
| 5156 | 5.08 | 0.00000000 |
| 5157 | 4.09 | 0.00000000 |
| 5158 | 6.70 | 0.00000000 |
| 5159 | 8.10 | 0.00000000 |
| 5160 | 10.00 | 0.00000000 |
| 5161 | 11.67 | 0.00000000 |
| 5162 | 8.64 | 0.00000000 |
| 5163 | 7.40 | 0.00000000 |
| 5164 | 8.27 | 0.00000000 |
| 5165 | 14.90 | 0.00000000 |
| 5166 | 34.27 | 0.00000000 |
| 5167 | 20.62 | 0.00000000 |
| 5168 | 14.90 | 0.00000000 |
| 5169 | 20.00 | 0.00000000 |
| 5170 | 15.33 | 0.00000000 |
| 5171 | 6.78 | 0.00000000 |
| 5172 | 3.58 | 0.00000000 |
| 5173 | 19.25 | 0.00000000 |
| 5174 | 10.75 | 0.00000000 |
| 5175 | 3.00 | 0.00000000 |
| 5176 | 5.17 | 0.00000000 |
| 5177 | 14.27 | 0.00000000 |
| 5178 | 32.00 | 0.00000000 |
| 5179 | 21.89 | 0.00000000 |
| 5180 | 5.64 | 0.00000000 |
| 5181 | 10.38 | 0.00000000 |
| 5182 | 9.38 | 0.00000000 |
| 5183 | 1.64 | 0.00000000 |
| 5184 | 4.33 | 0.00000000 |
| 5185 | 14.63 | 0.00000000 |
| 5186 | 8.73 | 0.00000000 |
| 5187 | 1.75 | 0.00000000 |
| 5188 | 9.44 | 0.00000000 |
| 5189 | 14.88 | 0.00000000 |
| 5190 | 1.91 | 0.00000000 |
| 5191 | 9.09 | 0.00000000 |
| 5192 | 10.30 | 0.00000000 |

| | | |
|---|---|---|
| 5193 | 14.56 | 0.00000000 |
| 5194 | 7.29 | 0.00000000 |
| 5195 | 10.10 | 0.00000000 |
| 5196 | 8.38 | 0.00000000 |
| 5197 | 14.89 | 0.00000000 |
| 5198 | 3.00 | 0.00000000 |
| 5199 | 5.55 | 0.00000000 |
| 5200 | 10.92 | 0.00000000 |
| 5201 | 13.14 | 0.00000000 |
| 5202 | 2.22 | 0.00000000 |
| 5203 | 12.22 | 0.00000000 |
| 5204 | 10.50 | 0.00000000 |
| 5205 | 12.45 | 0.00000000 |
| 5206 | 12.91 | 0.00000000 |
| 5207 | 13.50 | 0.00000000 |
| 5208 | 10.11 | 0.00000000 |
| 5209 | 11.73 | 0.00000000 |
| 5210 | 14.43 | 0.00000000 |
| 5211 | 4.13 | 0.00000000 |
| 5212 | 16.36 | 0.00000000 |
| 5213 | 6.20 | 0.00000000 |
| 5214 | 14.75 | 0.00000000 |
| 5215 | 15.38 | 0.00000000 |
| 5216 | 7.57 | 0.00000000 |
| 5217 | 14.10 | 0.00000000 |
| 5218 | 9.80 | 0.00000000 |
| 5219 | 13.88 | 0.00000000 |
| 5220 | 11.83 | 0.00000000 |
| 5221 | 10.62 | 0.00000000 |
| 5222 | 5.88 | 0.00000000 |
| 5223 | 5.46 | 0.00000000 |
| 5224 | 5.27 | 0.00000000 |
| 5225 | 7.30 | 0.00000000 |
| 5226 | 7.33 | 0.00000000 |
| 5227 | 8.62 | 0.00000000 |
| 5228 | 17.60 | 0.00000000 |
| 5229 | 9.70 | 0.00000000 |
| 5230 | 13.17 | 0.00000000 |
| 5231 | 5.89 | 0.00000000 |
| 5232 | 6.60 | 0.00000000 |
| 5233 | 17.83 | 0.00000000 |
| 5234 | 11.13 | 0.00000000 |
| 5235 | 8.33 | 0.00000000 |
| 5236 | 19.14 | 0.00000000 |

| | | |
|---|---|---|
| 5237 | 14.56 | 0.00000000 |
| 5238 | 8.14 | 0.00000000 |
| 5239 | 6.38 | 0.00000000 |
| 5240 | 6.10 | 0.00000000 |
| 5241 | 15.38 | 0.00000000 |
| 5242 | 9.78 | 0.00000000 |
| 5243 | 10.00 | 0.00000000 |
| 5244 | 11.38 | 0.00000000 |
| 5245 | 8.50 | 0.00000000 |
| 5246 | 3.63 | 0.00000000 |
| 5247 | 8.15 | 0.00000000 |
| 5248 | 8.50 | 0.00000000 |
| 5249 | 5.75 | 0.00000000 |
| 5250 | 13.33 | 0.00000000 |
| 5251 | 3.00 | 0.00000000 |
| 5252 | 23.50 | 0.00000000 |
| 5253 | 2.55 | 0.00000000 |
| 5254 | 11.40 | 0.00000000 |
| 5255 | 6.73 | 0.00000000 |
| 5256 | 12.93 | 0.00000000 |
| 5257 | 11.10 | 0.00000000 |
| 5258 | 26.70 | 0.00000000 |
| 5259 | 16.45 | 0.00000000 |
| 5260 | 8.50 | 0.00000000 |
| 5261 | 8.18 | 0.00000000 |
| 5262 | 18.11 | 0.00000000 |
| 5263 | 1.90 | 0.00000000 |
| 5264 | 16.38 | 0.00000000 |
| 5265 | 14.56 | 0.00000000 |
| 5266 | 13.78 | 0.00000000 |
| 5267 | 3.14 | 0.00000000 |
| 5268 | 7.63 | 0.00000000 |
| 5269 | 13.29 | 0.00000000 |
| 5270 | 10.75 | 0.00000000 |
| 5271 | 1.71 | 0.00000000 |
| 5272 | 3.00 | 0.00000000 |
| 5273 | 14.17 | 0.00000000 |
| 5274 | 2.60 | 0.00000000 |
| 5275 | 12.25 | 0.00000000 |
| 5276 | 6.88 | 0.00000000 |
| 5277 | 3.56 | 0.00000000 |
| 5278 | 22.63 | 0.00000000 |
| 5279 | 9.00 | 0.00000000 |
| 5280 | 8.44 | 0.00000000 |

| | | |
|---|---|---|
| 5281 | 11.00 | 0.00000000 |
| 5282 | 8.50 | 0.00000000 |
| 5283 | 3.43 | 0.00000000 |
| 5284 | 8.43 | 0.00000000 |
| 5285 | 4.38 | 0.00000000 |
| 5286 | 9.00 | 0.00000000 |
| 5287 | 8.82 | 0.00000000 |
| 5288 | 8.33 | 0.00000000 |
| 5289 | 17.57 | 0.00000000 |
| 5290 | 14.57 | 0.00000000 |
| 5291 | 14.40 | 0.00000000 |
| 5292 | 21.70 | 0.00000000 |
| 5293 | 6.33 | 0.00000000 |
| 5294 | 16.63 | 0.00000000 |
| 5295 | 10.00 | 0.00000000 |
| 5296 | 7.50 | 0.00000000 |
| 5297 | 11.33 | 0.00000000 |
| 5298 | 22.17 | 0.00000000 |
| 5299 | 3.56 | 0.00000000 |
| 5300 | 15.22 | 0.00000000 |
| 5301 | 11.64 | 0.00000000 |
| 5302 | 11.13 | 0.00000000 |
| 5303 | 3.89 | 0.00000000 |
| 5304 | 9.30 | 0.00000000 |
| 5305 | 7.91 | 0.00000000 |
| 5306 | 6.25 | 0.00000000 |
| 5307 | 11.50 | 0.00000000 |
| 5308 | 16.70 | 0.00000000 |
| 5309 | 18.11 | 0.00000000 |
| 5310 | 4.22 | 0.00000000 |
| 5311 | 9.09 | 0.00000000 |
| 5312 | 12.57 | 0.00000000 |
| 5313 | 2.71 | 0.00000000 |
| 5314 | 9.13 | 0.00000000 |
| 5315 | 11.33 | 0.00000000 |
| 5316 | 7.89 | 0.00000000 |
| 5317 | 16.78 | 0.00000000 |
| 5318 | 6.25 | 0.00000000 |
| 5319 | 6.50 | 0.00000000 |
| 5320 | 7.44 | 0.00000000 |
| 5321 | 20.17 | 0.00000000 |
| 5322 | 13.25 | 0.00000000 |
| 5323 | 8.30 | 0.00000000 |
| 5324 | 12.13 | 0.00000000 |

| | | |
|---|---|---|
| 5325 | 8.38 | 0.00000000 |
| 5326 | 8.18 | 0.00000000 |
| 5327 | 10.20 | 0.00000000 |
| 5328 | 13.40 | 0.00000000 |
| 5329 | 7.75 | 0.00000000 |
| 5330 | 9.80 | 0.00000000 |
| 5331 | 10.64 | 0.00000000 |
| 5332 | 17.86 | 0.00000000 |
| 5333 | 8.60 | 0.00000000 |
| 5334 | 5.64 | 0.00000000 |
| 5335 | 6.75 | 0.00000000 |
| 5336 | 6.40 | 0.00000000 |
| 5337 | 2.88 | 0.00000000 |
| 5338 | 9.82 | 0.00000000 |
| 5339 | 7.83 | 0.00000000 |
| 5340 | 18.00 | 0.00000000 |
| 5341 | 7.22 | 0.00000000 |
| 5342 | 14.89 | 0.00000000 |
| 5343 | 10.50 | 0.00000000 |
| 5344 | 2.30 | 0.00000000 |
| 5345 | 4.89 | 0.00000000 |
| 5346 | 18.17 | 0.00000000 |
| 5347 | 6.27 | 0.00000000 |
| 5348 | 2.75 | 0.00000000 |
| 5349 | 10.83 | 0.00000000 |
| 5350 | 5.82 | 0.00000000 |
| 5351 | 2.77 | 0.00000000 |
| 5352 | 3.89 | 0.00000000 |
| 5353 | 26.33 | 0.00000000 |
| 5354 | 14.50 | 0.00000000 |
| 5355 | 16.71 | 0.00000000 |
| 5356 | 6.33 | 0.00000000 |
| 5357 | 8.73 | 0.00000000 |
| 5358 | 9.43 | 0.00000000 |
| 5359 | 7.90 | 0.00000000 |
| 5360 | 8.69 | 0.00000000 |
| 5361 | 7.73 | 0.00000000 |
| 5362 | 15.82 | 0.00000000 |
| 5363 | 17.00 | 0.00000000 |
| 5364 | 13.69 | 0.00000000 |
| 5365 | 15.29 | 0.00000000 |
| 5366 | 4.11 | 0.00000000 |
| 5367 | 16.50 | 0.00000000 |
| 5368 | 9.00 | 0.00000000 |

| | | |
|---|---|---|
| 5369 | 15.75 | 0.00000000 |
| 5370 | 7.63 | 0.00000000 |
| 5371 | 7.56 | 0.00000000 |
| 5372 | 8.75 | 0.00000000 |
| 5373 | 10.90 | 0.00000000 |
| 5374 | 3.38 | 0.00000000 |
| 5375 | 17.42 | 0.00000000 |
| 5376 | 13.90 | 0.00000000 |
| 5377 | 7.60 | 0.00000000 |
| 5378 | 6.00 | 0.00000000 |
| 5379 | 7.57 | 0.00000000 |
| 5380 | 9.17 | 0.00000000 |
| 5381 | 26.25 | 0.00000000 |
| 5382 | 14.88 | 0.00000000 |
| 5383 | 16.50 | 0.00000000 |
| 5384 | 12.90 | 0.00000000 |
| 5385 | 12.67 | 0.00000000 |
| 5386 | 29.50 | 0.00000000 |
| 5387 | 6.88 | 0.00000000 |
| 5388 | 11.44 | 0.00000000 |
| 5389 | 5.92 | 0.00000000 |
| 5390 | 12.50 | 0.00000000 |
| 5391 | 17.22 | 0.00000000 |
| 5392 | 14.00 | 0.00000000 |
| 5393 | 9.18 | 0.00000000 |
| 5394 | 3.33 | 0.00000000 |
| 5395 | 15.80 | 0.00000000 |
| 5396 | 8.00 | 0.00000000 |
| 5397 | 6.14 | 0.00000000 |
| 5398 | 1.67 | 0.00000000 |
| 5399 | 16.83 | 0.00000000 |
| 5400 | 1.50 | 0.00000000 |
| 5401 | 2.57 | 0.00000000 |
| 5402 | 4.29 | 0.00000000 |
| 5403 | 13.70 | 0.00000000 |
| 5404 | 8.50 | 0.00000000 |
| 5405 | 3.11 | 0.00000000 |
| 5406 | 11.56 | 0.00000000 |
| 5407 | 11.33 | 0.00000000 |
| 5408 | 10.63 | 0.00000000 |
| 5409 | 12.55 | 0.00000000 |
| 5410 | 5.90 | 0.00000000 |
| 5411 | 14.11 | 0.00000000 |
| 5412 | 14.83 | 0.00000000 |

| | | |
|---|---|---|
| 5413 | 2.43 | 0.00000000 |
| 5414 | 1.82 | 0.00000000 |
| 5415 | 12.00 | 0.00000000 |
| 5416 | 7.69 | 0.00000000 |
| 5417 | 2.80 | 0.00000000 |
| 5418 | 21.33 | 0.00000000 |
| 5419 | 8.55 | 0.00000000 |
| 5420 | 8.80 | 0.00000000 |
| 5421 | 8.00 | 0.00000000 |
| 5422 | 4.00 | 0.00000000 |
| 5423 | 5.00 | 0.00000000 |
| 5424 | 13.22 | 0.00000000 |
| 5425 | 1.82 | 0.00000000 |
| 5426 | 8.25 | 0.00000000 |
| 5427 | 6.13 | 0.00000000 |
| 5428 | 7.85 | 0.00000000 |
| 5429 | 13.13 | 0.00000000 |
| 5430 | 2.60 | 0.00000000 |
| 5431 | 8.00 | 0.00000000 |
| 5432 | 6.80 | 0.00000000 |
| 5433 | 12.82 | 0.00000000 |
| 5434 | 5.70 | 0.00000000 |
| 5435 | 13.09 | 0.00000000 |
| 5436 | 5.80 | 0.00000000 |
| 5437 | 11.42 | 0.00000000 |
| 5438 | 2.64 | 0.00000000 |
| 5439 | 8.33 | 0.00000000 |
| 5440 | 5.92 | 0.00000000 |
| 5441 | 8.92 | 0.00000000 |
| 5442 | 13.00 | 0.00000000 |
| 5443 | 29.11 | 0.00000000 |
| 5444 | 13.30 | 0.00000000 |
| 5445 | 14.17 | 0.00000000 |
| 5446 | 2.89 | 0.00000000 |
| 5447 | 10.64 | 0.00000000 |
| 5448 | 2.11 | 0.00000000 |
| 5449 | 10.30 | 0.00000000 |
| 5450 | 13.20 | 0.00000000 |
| 5451 | 4.45 | 0.00000000 |
| 5452 | 14.92 | 0.00000000 |
| 5453 | 19.33 | 0.00000000 |
| 5454 | 2.11 | 0.00000000 |
| 5455 | 6.22 | 0.00000000 |
| 5456 | 6.13 | 0.00000000 |

| | | |
|---|---|---|
| 5457 | 3.55 | 0.00000000 |
| 5458 | 6.00 | 0.00000000 |
| 5459 | 16.70 | 0.00000000 |
| 5460 | 7.92 | 0.00000000 |
| 5461 | 7.82 | 0.00000000 |
| 5462 | 2.00 | 0.00000000 |
| 5463 | 6.75 | 0.00000000 |
| 5464 | 13.33 | 0.00000000 |
| 5465 | 3.27 | 0.00000000 |
| 5466 | 5.54 | 0.00000000 |
| 5467 | 11.89 | 0.00000000 |
| 5468 | 7.80 | 0.00000000 |
| 5469 | 10.40 | 0.00000000 |
| 5470 | 5.46 | 0.00000000 |
| 5471 | 12.20 | 0.00000000 |
| 5472 | 14.25 | 0.00000000 |
| 5473 | 10.44 | 0.00000000 |
| 5474 | 7.70 | 0.00000000 |
| 5475 | 9.91 | 0.00000000 |
| 5476 | 10.00 | 0.00000000 |
| 5477 | 12.64 | 0.00000000 |
| 5478 | 8.38 | 0.00000000 |
| 5479 | 11.46 | 0.00000000 |
| 5480 | 11.46 | 0.00000000 |
| 5481 | 17.63 | 0.00000000 |
| 5482 | 10.09 | 0.00000000 |
| 5483 | 21.91 | 0.00000000 |
| 5484 | 6.89 | 0.00000000 |
| 5485 | 5.83 | 0.00000000 |
| 5486 | 5.15 | 0.00000000 |
| 5487 | 8.38 | 0.00000000 |
| 5488 | 10.82 | 0.00000000 |
| 5489 | 13.42 | 0.00000000 |
| 5490 | 7.20 | 0.00000000 |
| 5491 | 17.55 | 0.00000000 |
| 5492 | 10.78 | 0.00000000 |
| 5493 | 5.64 | 0.00000000 |
| 5494 | 7.09 | 0.00000000 |
| 5495 | 10.10 | 0.00000000 |
| 5496 | 2.30 | 0.00000000 |
| 5497 | 3.31 | 0.00000000 |
| 5498 | 7.29 | 0.00000000 |
| 5499 | 18.50 | 0.00000000 |
| 5500 | 13.67 | 0.00000000 |

| | | |
|---|---|---|
| 5501 | 6.33 | 0.00000000 |
| 5502 | 5.18 | 0.00000000 |
| 5503 | 13.83 | 0.00000000 |
| 5504 | 12.58 | 0.00000000 |
| 5505 | 11.92 | 0.00000000 |
| 5506 | 6.82 | 0.00000000 |
| 5507 | 9.33 | 0.00000000 |
| 5508 | 13.00 | 0.00000000 |
| 5509 | 6.78 | 0.00000000 |
| 5510 | 13.60 | 0.00000000 |
| 5511 | 14.18 | 0.00000000 |
| 5512 | 7.69 | 0.00000000 |
| 5513 | 21.13 | 0.00000000 |
| 5514 | 14.11 | 0.00000000 |
| 5515 | 3.14 | 0.00000000 |
| 5516 | 18.29 | 0.00000000 |
| 5517 | 5.78 | 0.00000000 |
| 5518 | 13.63 | 0.00000000 |
| 5519 | 6.92 | 0.00000000 |
| 5520 | 2.90 | 0.00000000 |
| 5521 | 9.27 | 0.00000000 |
| 5522 | 11.40 | 0.00000000 |
| 5523 | 9.90 | 0.00000000 |
| 5524 | 5.46 | 0.00000000 |
| 5525 | 3.45 | 0.00000000 |
| 5526 | 9.10 | 0.00000000 |
| 5527 | 11.00 | 0.00000000 |
| 5528 | 15.63 | 0.00000000 |
| 5529 | 7.86 | 0.00000000 |
| 5530 | 13.57 | 0.00000000 |
| 5531 | 5.30 | 0.00000000 |
| 5532 | 10.88 | 0.00000000 |
| 5533 | 11.78 | 0.00000000 |
| 5534 | 7.56 | 0.00000000 |
| 5535 | 8.75 | 0.00000000 |
| 5536 | 22.30 | 0.00000000 |
| 5537 | 9.45 | 0.00000000 |
| 5538 | 15.00 | 0.00000000 |
| 5539 | 10.89 | 0.00000000 |
| 5540 | 7.00 | 0.00000000 |
| 5541 | 6.00 | 0.00000000 |
| 5542 | 3.44 | 0.00000000 |
| 5543 | 24.29 | 0.00000000 |
| 5544 | 23.43 | 0.00000000 |

| | | |
|---|---|---|
| 5545 | 7.00 | 0.00000000 |
| 5546 | 2.57 | 0.00000000 |
| 5547 | 1.22 | 0.00000000 |
| 5548 | 7.29 | 0.00000000 |
| 5549 | 6.11 | 0.00000000 |
| 5550 | 1.75 | 0.00000000 |
| 5551 | 1.82 | 0.00000000 |
| 5552 | 7.00 | 0.00000000 |
| 5553 | 6.13 | 0.00000000 |
| 5554 | 12.67 | 0.00000000 |
| 5555 | 2.11 | 0.00000000 |
| 5556 | 2.50 | 0.00000000 |
| 5557 | 4.08 | 0.00000000 |
| 5558 | 13.50 | 0.00000000 |
| 5559 | 1.80 | 0.00000000 |
| 5560 | 8.50 | 0.00000000 |
| 5561 | 4.90 | 0.00000000 |
| 5562 | 8.91 | 0.00000000 |
| 5563 | 11.71 | 0.00000000 |
| 5564 | 6.86 | 0.00000000 |
| 5565 | 15.45 | 0.00000000 |
| 5566 | 9.33 | 0.00000000 |
| 5567 | 5.64 | 0.00000000 |
| 5568 | 10.83 | 0.00000000 |
| 5569 | 5.00 | 0.00000000 |
| 5570 | 6.18 | 0.00000000 |
| 5571 | 9.67 | 0.00000000 |
| 5572 | 12.60 | 0.00000000 |
| 5573 | 15.30 | 0.00000000 |
| 5574 | 9.80 | 0.00000000 |
| 5575 | 9.60 | 0.00000000 |
| 5576 | 10.20 | 0.00000000 |
| 5577 | 2.91 | 0.00000000 |
| 5578 | 8.08 | 0.00000000 |
| 5579 | 17.69 | 0.00000000 |
| 5580 | 18.73 | 0.00000000 |
| 5581 | 11.38 | 0.00000000 |
| 5582 | 2.20 | 0.00000000 |
| 5583 | 17.44 | 0.00000000 |
| 5584 | 5.78 | 0.00000000 |
| 5585 | 10.00 | 0.00000000 |
| 5586 | 8.22 | 0.00000000 |
| 5587 | 16.67 | 0.00000000 |
| 5588 | 1.75 | 0.00000000 |

| | | |
|---|---|---|
| 5589 | 15.83 | 0.00000000 |
| 5590 | 2.43 | 0.00000000 |
| 5591 | 8.00 | 0.00000000 |
| 5592 | 18.90 | 0.00000000 |
| 5593 | 18.71 | 0.00000000 |
| 5594 | 12.44 | 0.00000000 |
| 5595 | 3.00 | 0.00000000 |
| 5596 | 11.22 | 0.00000000 |
| 5597 | 2.00 | 0.00000000 |
| 5598 | 12.25 | 0.00000000 |
| 5599 | 13.50 | 0.00000000 |
| 5600 | 7.88 | 0.00000000 |
| 5601 | 5.63 | 0.00000000 |
| 5602 | 4.00 | 0.00000000 |
| 5603 | 4.20 | 0.00000000 |
| 5604 | 10.11 | 0.00000000 |
| 5605 | 14.43 | 0.00000000 |
| 5606 | 10.14 | 0.00000000 |
| 5607 | 26.80 | 0.00000000 |
| 5608 | 10.20 | 0.00000000 |
| 5609 | 7.60 | 0.00000000 |
| 5610 | 2.25 | 0.00000000 |
| 5611 | 8.45 | 0.00000000 |
| 5612 | 13.89 | 0.00000000 |
| 5613 | 3.50 | 0.00000000 |
| 5614 | 16.00 | 0.00000000 |
| 5615 | 9.71 | 0.00000000 |
| 5616 | 7.14 | 0.00000000 |
| 5617 | 10.44 | 0.00000000 |
| 5618 | 6.55 | 0.00000000 |
| 5619 | 11.75 | 0.00000000 |
| 5620 | 11.80 | 0.00000000 |
| 5621 | 11.40 | 0.00000000 |
| 5622 | 22.50 | 0.00000000 |
| 5623 | 27.60 | 0.00000000 |
| 5624 | 13.10 | 0.00000000 |
| 5625 | 13.20 | 0.00000000 |
| 5626 | 9.50 | 0.00000000 |
| 5627 | 14.63 | 0.00000000 |
| 5628 | 9.89 | 0.00000000 |
| 5629 | 10.00 | 0.00000000 |
| 5630 | 19.63 | 0.00000000 |
| 5631 | 2.50 | 0.00000000 |
| 5632 | 13.20 | 0.00000000 |

| | | |
|---|---|---|
| 5633 | 13.00 | 0.00000000 |
| 5634 | 7.71 | 0.00000000 |
| 5635 | 7.11 | 0.00000000 |
| 5636 | 7.43 | 0.00000000 |
| 5637 | 9.29 | 0.00000000 |
| 5638 | 21.70 | 0.00000000 |
| 5639 | 3.17 | 0.00000000 |
| 5640 | 3.17 | 0.00000000 |
| 5641 | 5.00 | 0.00000000 |
| 5642 | 11.75 | 0.00000000 |
| 5643 | 8.83 | 0.00000000 |
| 5644 | 10.70 | 0.00000000 |
| 5645 | 14.00 | 0.00000000 |
| 5646 | 15.70 | 0.00000000 |
| 5647 | 2.25 | 0.00000000 |
| 5648 | 6.78 | 0.00000000 |
| 5649 | 4.40 | 0.00000000 |
| 5650 | 18.63 | 0.00000000 |
| 5651 | 19.00 | 0.00000000 |
| 5652 | 6.25 | 0.00000000 |
| 5653 | 6.89 | 0.00000000 |
| 5654 | 5.00 | 0.00000000 |
| 5655 | 2.80 | 0.00000000 |
| 5656 | 2.00 | 0.00000000 |
| 5657 | 15.88 | 0.00000000 |
| 5658 | 12.14 | 0.00000000 |
| 5659 | 8.86 | 0.00000000 |
| 5660 | 5.67 | 0.00000000 |
| 5661 | 6.50 | 0.00000000 |
| 5662 | 6.67 | 0.00000000 |
| 5663 | 5.60 | 0.00000000 |
| 5664 | 2.50 | 0.00000000 |
| 5665 | 5.73 | 0.00000000 |
| 5666 | 11.18 | 0.00000000 |
| 5667 | 10.64 | 0.00000000 |
| 5668 | 13.44 | 0.00000000 |
| 5669 | 4.86 | 0.00000000 |
| 5670 | 20.00 | 0.00000000 |
| 5671 | 5.82 | 0.00000000 |
| 5672 | 17.00 | 0.00000000 |
| 5673 | 11.30 | 0.00000000 |
| 5674 | 9.09 | 0.00000000 |
| 5675 | 6.56 | 0.00000000 |
| 5676 | 2.71 | 0.00000000 |

| | | |
|---|---|---|
| 5677 | 18.33 | 0.00000000 |
| 5678 | 6.40 | 0.00000000 |
| 5679 | 7.13 | 0.00000000 |
| 5680 | 9.33 | 0.00000000 |
| 5681 | 5.91 | 0.00000000 |
| 5682 | 4.75 | 0.00000000 |
| 5683 | 7.00 | 0.00000000 |
| 5684 | 5.60 | 0.00000000 |
| 5685 | 5.73 | 0.00000000 |
| 5686 | 8.17 | 0.00000000 |
| 5687 | 2.25 | 0.00000000 |
| 5688 | 3.50 | 0.00000000 |
| 5689 | 14.23 | 0.00000000 |
| 5690 | 10.11 | 0.00000000 |
| 5691 | 5.50 | 0.00000000 |
| 5692 | 7.22 | 0.00000000 |
| 5693 | 3.44 | 0.00000000 |
| 5694 | 25.67 | 0.00000000 |
| 5695 | 11.67 | 0.00000000 |
| 5696 | 8.75 | 0.00000000 |
| 5697 | 12.17 | 0.00000000 |
| 5698 | 10.33 | 0.00000000 |
| 5699 | 4.83 | 0.00000000 |
| 5700 | 21.82 | 0.00000000 |
| 5701 | 6.78 | 0.00000000 |
| 5702 | 17.56 | 0.00000000 |
| 5703 | 3.00 | 0.00000000 |
| 5704 | 8.91 | 0.00000000 |
| 5705 | 8.18 | 0.00000000 |
| 5706 | 11.20 | 0.00000000 |
| 5707 | 5.56 | 0.00000000 |
| 5708 | 8.17 | 0.00000000 |
| 5709 | 4.38 | 0.00000000 |
| 5710 | 11.23 | 0.00000000 |
| 5711 | 9.27 | 0.00000000 |
| 5712 | 5.23 | 0.00000000 |
| 5713 | 9.42 | 0.00000000 |
| 5714 | 8.57 | 0.00000000 |
| 5715 | 7.00 | 0.00000000 |
| 5716 | 3.33 | 0.00000000 |
| 5717 | 18.25 | 0.00000000 |
| 5718 | 14.27 | 0.00000000 |
| 5719 | 7.86 | 0.00000000 |
| 5720 | 22.00 | 0.00000000 |

| | | |
|---|---|---|
| 5721 | 9.11 | 0.00000000 |
| 5722 | 2.25 | 0.00000000 |
| 5723 | 2.91 | 0.00000000 |
| 5724 | 17.10 | 0.00000000 |
| 5725 | 7.11 | 0.00000000 |
| 5726 | 8.11 | 0.00000000 |
| 5727 | 6.89 | 0.00000000 |
| 5728 | 15.00 | 0.00000000 |
| 5729 | 20.38 | 0.00000000 |
| 5730 | 7.80 | 0.00000000 |
| 5731 | 6.73 | 0.00000000 |
| 5732 | 13.30 | 0.00000000 |
| 5733 | 8.45 | 0.00000000 |
| 5734 | 13.25 | 0.00000000 |
| 5735 | 14.55 | 0.00000000 |
| 5736 | 24.70 | 0.00000000 |
| 5737 | 8.00 | 0.00000000 |
| 5738 | 7.11 | 0.00000000 |
| 5739 | 5.43 | 0.00000000 |
| 5740 | 14.60 | 0.00000000 |
| 5741 | 6.00 | 0.00000000 |
| 5742 | 2.70 | 0.00000000 |
| 5743 | 8.33 | 0.00000000 |
| 5744 | 6.56 | 0.00000000 |
| 5745 | 6.22 | 0.00000000 |
| 5746 | 8.56 | 0.00000000 |
| 5747 | 23.38 | 0.00000000 |
| 5748 | 12.83 | 0.00000000 |
| 5749 | 7.13 | 0.00000000 |
| 5750 | 9.57 | 0.00000000 |
| 5751 | 7.75 | 0.00000000 |
| 5752 | 8.50 | 0.00000000 |
| 5753 | 14.14 | 0.00000000 |
| 5754 | 6.82 | 0.00000000 |
| 5755 | 13.29 | 0.00000000 |
| 5756 | 2.40 | 0.00000000 |
| 5757 | 11.91 | 0.00000000 |
| 5758 | 12.86 | 0.00000000 |
| 5759 | 7.22 | 0.00000000 |
| 5760 | 15.83 | 0.00000000 |
| 5761 | 19.00 | 0.00000000 |
| 5762 | 7.13 | 0.00000000 |
| 5763 | 9.60 | 0.00000000 |
| 5764 | 1.80 | 0.00000000 |

| | | |
|---|---|---|
| 5765 | 5.38 | 0.00000000 |
| 5766 | 12.82 | 0.00000000 |
| 5767 | 14.89 | 0.00000000 |
| 5768 | 2.45 | 0.00000000 |
| 5769 | 9.33 | 0.00000000 |
| 5770 | 5.80 | 0.00000000 |
| 5771 | 16.90 | 0.00000000 |
| 5772 | 7.09 | 0.00000000 |
| 5773 | 21.78 | 0.00000000 |
| 5774 | 7.00 | 0.00000000 |
| 5775 | 13.50 | 0.00000000 |
| 5776 | 15.20 | 0.00000000 |
| 5777 | 2.67 | 0.00000000 |
| 5778 | 11.27 | 0.00000000 |
| 5779 | 12.33 | 0.00000000 |
| 5780 | 11.63 | 0.00000000 |
| 5781 | 6.56 | 0.00000000 |
| 5782 | 10.00 | 0.00000000 |
| 5783 | 2.50 | 0.00000000 |
| 5784 | 7.70 | 0.00000000 |
| 5785 | 19.36 | 0.00000000 |
| 5786 | 2.86 | 0.00000000 |
| 5787 | 18.67 | 0.00000000 |
| 5788 | 3.00 | 0.00000000 |
| 5789 | 18.78 | 0.00000000 |
| 5790 | 6.25 | 0.00000000 |
| 5791 | 2.20 | 0.00000000 |
| 5792 | 2.45 | 0.00000000 |
| 5793 | 3.44 | 0.00000000 |
| 5794 | 9.50 | 0.00000000 |
| 5795 | 12.00 | 0.00000000 |
| 5796 | 6.25 | 0.00000000 |
| 5797 | 2.75 | 0.00000000 |
| 5798 | 19.40 | 0.00000000 |
| 5799 | 7.67 | 0.00000000 |
| 5800 | 14.10 | 0.00000000 |
| 5801 | 16.33 | 0.00000000 |
| 5802 | 8.71 | 0.00000000 |
| 5803 | 15.75 | 0.00000000 |
| 5804 | 16.45 | 0.00000000 |
| 5805 | 7.86 | 0.00000000 |
| 5806 | 6.30 | 0.00000000 |
| 5807 | 3.13 | 0.00000000 |
| 5808 | 31.00 | 0.00000000 |

| | | |
|---|---|---|
| 5809 | 8.14 | 0.00000000 |
| 5810 | 2.89 | 0.00000000 |
| 5811 | 17.25 | 0.00000000 |
| 5812 | 9.00 | 0.00000000 |
| 5813 | 10.80 | 0.00000000 |
| 5814 | 13.88 | 0.00000000 |
| 5815 | 7.86 | 0.00000000 |
| 5816 | 5.40 | 0.00000000 |
| 5817 | 15.30 | 0.00000000 |
| 5818 | 3.71 | 0.00000000 |
| 5819 | 6.38 | 0.00000000 |
| 5820 | 8.50 | 0.00000000 |
| 5821 | 3.25 | 0.00000000 |
| 5822 | 11.00 | 0.00000000 |
| 5823 | 7.80 | 0.00000000 |
| 5824 | 3.50 | 0.00000000 |
| 5825 | 20.00 | 0.00000000 |
| 5826 | 9.00 | 0.00000000 |
| 5827 | 4.58 | 0.00000000 |
| 5828 | 3.86 | 0.00000000 |
| 5829 | 7.33 | 0.00000000 |
| 5830 | 7.86 | 0.00000000 |
| 5831 | 9.44 | 0.00000000 |
| 5832 | 17.22 | 0.00000000 |
| 5833 | 14.50 | 0.00000000 |
| 5834 | 6.29 | 0.00000000 |
| 5835 | 11.13 | 0.00000000 |
| 5836 | 21.83 | 0.00000000 |
| 5837 | 2.33 | 0.00000000 |
| 5838 | 2.57 | 0.00000000 |
| 5839 | 14.29 | 0.00000000 |
| 5840 | 9.00 | 0.00000000 |
| 5841 | 7.57 | 0.00000000 |
| 5842 | 11.50 | 0.00000000 |
| 5843 | 16.00 | 0.00000000 |
| 5844 | 3.22 | 0.00000000 |
| 5845 | 12.00 | 0.00000000 |
| 5846 | 8.75 | 0.00000000 |
| 5847 | 12.38 | 0.00000000 |
| 5848 | 3.75 | 0.00000000 |
| 5849 | 9.63 | 0.00000000 |
| 5850 | 26.63 | 0.00000000 |
| 5851 | 8.57 | 0.00000000 |
| 5852 | 9.57 | 0.00000000 |

| | | |
|---|---|---|
| 5853 | 11.00 | 0.00000000 |
| 5854 | 11.22 | 0.00000000 |
| 5855 | 23.57 | 0.00000000 |
| 5856 | 9.00 | 0.00000000 |
| 5857 | 7.00 | 0.00000000 |
| 5858 | 17.29 | 0.00000000 |
| 5859 | 9.88 | 0.00000000 |
| 5860 | 1.57 | 0.00000000 |
| 5861 | 7.43 | 0.00000000 |
| 5862 | 1.00 | 0.00000000 |
| 5863 | 1.50 | 0.00000000 |
| 5864 | 3.43 | 0.00000000 |
| 5865 | 10.00 | 0.00000000 |
| 5866 | 6.83 | 0.00000000 |
| 5867 | 1.70 | 0.00000000 |
| 5868 | 2.43 | 0.00000000 |
| 5869 | 5.80 | 0.00000000 |
| 5870 | 4.14 | 0.00000000 |
| 5871 | 10.00 | 0.00000000 |
| 5872 | 11.00 | 0.00000000 |
| 5873 | 8.43 | 0.00000000 |
| 5874 | 14.00 | 0.00000000 |
| 5875 | 7.60 | 0.00000000 |
| 5876 | 5.67 | 0.00000000 |
| 5877 | 13.56 | 0.00000000 |
| 5878 | 6.78 | 0.00000000 |
| 5879 | 8.90 | 0.00000000 |
| 5880 | 13.56 | 0.00000000 |
| 5881 | 15.00 | 0.00000000 |
| 5882 | 8.82 | 0.00000000 |
| 5883 | 3.57 | 0.00000000 |
| 5884 | 9.17 | 0.00000000 |
| 5885 | 12.78 | 0.00000000 |
| 5886 | 5.20 | 0.00000000 |
| 5887 | 2.60 | 0.00000000 |
| 5888 | 6.78 | 0.00000000 |
| 5889 | 6.45 | 0.00000000 |
| 5890 | 1.82 | 0.00000000 |
| 5891 | 11.23 | 0.00000000 |
| 5892 | 7.70 | 0.00000000 |
| 5893 | 3.89 | 0.00000000 |
| 5894 | 5.91 | 0.00000000 |
| 5895 | 10.20 | 0.00000000 |
| 5896 | 1.70 | 0.00000000 |

| | | |
|---|---|---|
| 5897 | 5.64 | 0.00000000 |
| 5898 | 21.50 | 0.00000000 |
| 5899 | 13.23 | 0.00000000 |
| 5900 | 2.92 | 0.00000000 |
| 5901 | 9.57 | 0.00000000 |
| 5902 | 8.43 | 0.00000000 |
| 5903 | 4.38 | 0.00000000 |
| 5904 | 7.70 | 0.00000000 |
| 5905 | 19.56 | 0.00000000 |
| 5906 | 11.25 | 0.00000000 |
| 5907 | 3.23 | 0.00000000 |
| 5908 | 6.56 | 0.00000000 |
| 5909 | 7.00 | 0.00000000 |
| 5910 | 12.38 | 0.00000000 |
| 5911 | 19.38 | 0.00000000 |
| 5912 | 8.57 | 0.00000000 |
| 5913 | 16.40 | 0.00000000 |
| 5914 | 1.80 | 0.00000000 |
| 5915 | 11.29 | 0.00000000 |
| 5916 | 8.14 | 0.00000000 |
| 5917 | 1.80 | 0.00000000 |
| 5918 | 11.63 | 0.00000000 |
| 5919 | 1.83 | 0.00000000 |
| 5920 | 8.40 | 0.00000000 |
| 5921 | 18.40 | 0.00000000 |
| 5922 | 1.75 | 0.00000000 |
| 5923 | 3.00 | 0.00000000 |
| 5924 | 7.17 | 0.00000000 |
| 5925 | 4.20 | 0.00000000 |
| 5926 | 2.50 | 0.00000000 |
| 5927 | 14.00 | 0.00000000 |
| 5928 | 7.50 | 0.00000000 |
| 5929 | 10.38 | 0.00000000 |
| 5930 | 3.67 | 0.00000000 |
| 5931 | 5.29 | 0.00000000 |
| 5932 | 11.63 | 0.00000000 |
| 5933 | 11.89 | 0.00000000 |
| 5934 | 12.86 | 0.00000000 |
| 5935 | 5.86 | 0.00000000 |
| 5936 | 3.00 | 0.00000000 |
| 5937 | 10.20 | 0.00000000 |
| 5938 | 13.00 | 0.00000000 |
| 5939 | 4.89 | 0.00000000 |
| 5940 | 9.57 | 0.00000000 |

| | | |
|---|---|---|
| 5941 | 9.64 | 0.00000000 |
| 5942 | 6.14 | 0.00000000 |
| 5943 | 2.57 | 0.00000000 |
| 5944 | 7.38 | 0.00000000 |
| 5945 | 10.56 | 0.00000000 |
| 5946 | 4.88 | 0.00000000 |
| 5947 | 7.67 | 0.00000000 |
| 5948 | 5.78 | 0.00000000 |
| 5949 | 12.00 | 0.00000000 |
| 5950 | 2.20 | 0.00000000 |
| 5951 | 1.88 | 0.00000000 |
| 5952 | 9.33 | 0.00000000 |
| 5953 | 10.11 | 0.00000000 |
| 5954 | 25.80 | 0.00000000 |
| 5955 | 11.50 | 0.00000000 |
| 5956 | 5.58 | 0.00000000 |
| 5957 | 8.36 | 0.00000000 |
| 5958 | 7.00 | 0.00000000 |
| 5959 | 11.29 | 0.00000000 |
| 5960 | 6.29 | 0.00000000 |
| 5961 | 2.38 | 0.00000000 |
| 5962 | 5.40 | 0.00000000 |
| 5963 | 9.00 | 0.00000000 |
| 5964 | 8.80 | 0.00000000 |
| 5965 | 1.44 | 0.00000000 |
| 5966 | 3.44 | 0.00000000 |
| 5967 | 11.57 | 0.00000000 |
| 5968 | 5.78 | 0.00000000 |
| 5969 | 2.10 | 0.00000000 |
| 5970 | 12.00 | 0.00000000 |
| 5971 | 13.56 | 0.00000000 |
| 5972 | 11.67 | 0.00000000 |
| 5973 | 5.00 | 0.00000000 |
| 5974 | 2.29 | 0.00000000 |
| 5975 | 5.60 | 0.00000000 |
| 5976 | 7.60 | 0.00000000 |
| 5977 | 15.78 | 0.00000000 |
| 5978 | 10.90 | 0.00000000 |
| 5979 | 6.18 | 0.00000000 |
| 5980 | 15.13 | 0.00000000 |
| 5981 | 11.20 | 0.00000000 |
| 5982 | 13.11 | 0.00000000 |
| 5983 | 11.00 | 0.00000000 |
| 5984 | 6.89 | 0.00000000 |

| | | |
|---|---|---|
| 5985 | 11.00 | 0.00000000 |
| 5986 | 7.83 | 0.00000000 |
| 5987 | 8.50 | 0.00000000 |
| 5988 | 10.71 | 0.00000000 |
| 5989 | 7.75 | 0.00000000 |
| 5990 | 5.25 | 0.00000000 |
| 5991 | 5.00 | 0.00000000 |
| 5992 | 9.17 | 0.00000000 |
| 5993 | 17.43 | 0.00000000 |
| 5994 | 16.00 | 0.00000000 |
| 5995 | 2.20 | 0.00000000 |
| 5996 | 3.17 | 0.00000000 |
| 5997 | 2.13 | 0.00000000 |
| 5998 | 14.00 | 0.00000000 |
| 5999 | 1.60 | 0.00000000 |
| 6000 | 1.44 | 0.00000000 |
| 6001 | 1.50 | 0.00000000 |
| 6002 | 5.44 | 0.00000000 |
| 6003 | 12.13 | 0.00000000 |
| 6004 | 7.29 | 0.00000000 |
| 6005 | 10.75 | 0.00000000 |
| 6006 | 8.00 | 0.00000000 |
| 6007 | 2.71 | 0.00000000 |
| 6008 | 1.67 | 0.00000000 |
| 6009 | 5.44 | 0.00000000 |
| 6010 | 8.83 | 0.00000000 |
| 6011 | 4.57 | 0.00000000 |
| 6012 | 9.60 | 0.00000000 |
| 6013 | 9.50 | 0.00000000 |
| 6014 | 14.86 | 0.00000000 |
| 6015 | 1.00 | 0.00000000 |
| 6016 | 2.40 | 0.00000000 |
| 6017 | 5.13 | 0.00000000 |
| 6018 | 7.70 | 0.00000000 |
| 6019 | 3.00 | 0.00000000 |
| 6020 | 3.08 | 0.00000000 |
| 6021 | 6.33 | 0.00000000 |
| 6022 | 13.09 | 0.00000000 |
| 6023 | 6.33 | 0.00000000 |
| 6024 | 3.22 | 0.00000000 |
| 6025 | 17.00 | 0.00000000 |
| 6026 | 2.20 | 0.00000000 |
| 6027 | 20.64 | 0.00000000 |
| 6028 | 12.89 | 0.00000000 |

| | | |
|---|---|---|
| 6029 | 7.43 | 0.00000000 |
| 6030 | 10.44 | 0.00000000 |
| 6031 | 8.60 | 0.00000000 |
| 6032 | 15.00 | 0.00000000 |
| 6033 | 8.86 | 0.00000000 |
| 6034 | 6.88 | 0.00000000 |
| 6035 | 8.60 | 0.00000000 |
| 6036 | 18.67 | 0.00000000 |
| 6037 | 2.75 | 0.00000000 |
| 6038 | 13.44 | 0.00000000 |
| 6039 | 10.00 | 0.00000000 |
| 6040 | 1.71 | 0.00000000 |
| 6041 | 5.09 | 0.00000000 |
| 6042 | 11.17 | 0.00000000 |
| 6043 | 10.22 | 0.00000000 |
| 6044 | 5.63 | 0.00000000 |
| 6045 | 6.33 | 0.00000000 |
| 6046 | 2.38 | 0.00000000 |
| 6047 | 8.75 | 0.00000000 |
| 6048 | 7.22 | 0.00000000 |
| 6049 | 11.22 | 0.00000000 |
| 6050 | 8.40 | 0.00000000 |
| 6051 | 7.38 | 0.00000000 |
| 6052 | 7.00 | 0.00000000 |
| 6053 | 1.50 | 0.00000000 |
| 6054 | 12.00 | 0.00000000 |
| 6055 | 6.63 | 0.00000000 |
| 6056 | 17.14 | 0.00000000 |
| 6057 | 5.50 | 0.00000000 |
| 6058 | 9.80 | 0.00000000 |
| 6059 | 1.43 | 0.00000000 |
| 6060 | 3.17 | 0.00000000 |
| 6061 | 11.40 | 0.00000000 |
| 6062 | 8.27 | 0.00000000 |
| 6063 | 14.22 | 0.00000000 |
| 6064 | 5.56 | 0.00000000 |
| 6065 | 8.25 | 0.00000000 |
| 6066 | 8.25 | 0.00000000 |
| 6067 | 6.00 | 0.00000000 |
| 6068 | 6.71 | 0.00000000 |
| 6069 | 14.73 | 0.00000000 |
| 6070 | 11.67 | 0.00000000 |
| 6071 | 17.20 | 0.00000000 |
| 6072 | 8.67 | 0.00000000 |

| | | |
|---|---|---|
| 6073 | 5.40 | 0.00000000 |
| 6074 | 10.29 | 0.00000000 |
| 6075 | 2.73 | 0.00000000 |
| 6076 | 9.80 | 0.00000000 |
| 6077 | 10.60 | 0.00000000 |
| 6078 | 8.55 | 0.00000000 |
| 6079 | 13.57 | 0.00000000 |
| 6080 | 5.73 | 0.00000000 |
| 6081 | 9.00 | 0.00000000 |
| 6082 | 6.75 | 0.00000000 |
| 6083 | 4.50 | 0.00000000 |
| 6084 | 17.83 | 0.00000000 |
| 6085 | 16.11 | 0.00000000 |
| 6086 | 6.56 | 0.00000000 |
| 6087 | 12.25 | 0.00000000 |
| 6088 | 13.63 | 0.00000000 |
| 6089 | 24.43 | 0.00000000 |
| 6090 | 2.14 | 0.00000000 |
| 6091 | 9.33 | 0.00000000 |
| 6092 | 9.14 | 0.00000000 |
| 6093 | 6.56 | 0.00000000 |
| 6094 | 8.17 | 0.00000000 |
| 6095 | 2.50 | 0.00000000 |
| 6096 | 2.71 | 0.00000000 |
| 6097 | 7.00 | 0.00000000 |
| 6098 | 6.88 | 0.00000000 |
| 6099 | 7.14 | 0.00000000 |
| 6100 | 7.83 | 0.00000000 |
| 6101 | 13.29 | 0.00000000 |
| 6102 | 8.43 | 0.00000000 |
| 6103 | 13.44 | 0.00000000 |
| 6104 | 2.25 | 0.00000000 |
| 6105 | 7.57 | 0.00000000 |
| 6106 | 9.80 | 0.00000000 |
| 6107 | 20.25 | 0.00000000 |
| 6108 | 5.70 | 0.00000000 |
| 6109 | 6.40 | 0.00000000 |
| 6110 | 19.25 | 0.00000000 |
| 6111 | 15.00 | 0.00000000 |
| 6112 | 10.75 | 0.00000000 |
| 6113 | 1.67 | 0.00000000 |
| 6114 | 12.75 | 0.00000000 |
| 6115 | 1.71 | 0.00000000 |
| 6116 | 6.00 | 0.00000000 |

| | | |
|---|---|---|
| 6117 | 6.29 | 0.00000000 |
| 6118 | 5.63 | 0.00000000 |
| 6119 | 6.88 | 0.00000000 |
| 6120 | 12.88 | 0.00000000 |
| 6121 | 11.67 | 0.00000000 |
| 6122 | 1.60 | 0.00000000 |
| 6123 | 2.20 | 0.00000000 |
| 6124 | 8.80 | 0.00000000 |
| 6125 | 2.43 | 0.00000000 |
| 6126 | 9.67 | 0.00000000 |
| 6127 | 10.88 | 0.00000000 |
| 6128 | 14.43 | 0.00000000 |
| 6129 | 3.14 | 0.00000000 |
| 6130 | 2.50 | 0.00000000 |
| 6131 | 8.13 | 0.00000000 |
| 6132 | 16.75 | 0.00000000 |
| 6133 | 13.50 | 0.00000000 |
| 6134 | 6.86 | 0.00000000 |
| 6135 | 9.14 | 0.00000000 |
| 6136 | 8.00 | 0.00000000 |
| 6137 | 15.57 | 0.00000000 |
| 6138 | 10.86 | 0.00000000 |
| 6139 | 10.75 | 0.00000000 |
| 6140 | 19.75 | 0.00000000 |
| 6141 | 6.57 | 0.00000000 |
| 6142 | 8.60 | 0.00000000 |
| 6143 | 3.29 | 0.00000000 |
| 6144 | 1.57 | 0.00000000 |
| 6145 | 15.60 | 0.00000000 |
| 6146 | 7.50 | 0.00000000 |
| 6147 | 2.00 | 0.00000000 |
| 6148 | 3.50 | 0.00000000 |
| 6149 | 7.83 | 0.00000000 |
| 6150 | 2.83 | 0.00000000 |
| 6151 | 8.20 | 0.00000000 |
| 6152 | 4.75 | 0.00000000 |
| 6153 | 10.60 | 0.00000000 |
| 6154 | 9.50 | 0.00000000 |
| 6155 | 4.78 | 0.00000000 |
| 6156 | 9.20 | 0.00000000 |
| 6157 | 3.33 | 0.00000000 |
| 6158 | 7.83 | 0.00000000 |
| 6159 | 8.60 | 0.00000000 |
| 6160 | 9.50 | 0.00000000 |

| | | |
|---|---|---|
| 6161 | 13.00 | 0.00000000 |
| 6162 | 23.83 | 0.00000000 |
| 6163 | 10.00 | 0.00000000 |
| 6164 | 9.71 | 0.00000000 |
| 6165 | 16.11 | 0.00000000 |
| 6166 | 2.14 | 0.00000000 |
| 6167 | 9.33 | 0.00000000 |
| 6168 | 12.00 | 0.00000000 |
| 6169 | 13.00 | 0.00000000 |
| 6170 | 6.86 | 0.00000000 |
| 6171 | 2.00 | 0.00000000 |
| 6172 | 2.80 | 0.00000000 |
| 6173 | 7.57 | 0.00000000 |
| 6174 | 5.43 | 0.00000000 |
| 6175 | 2.00 | 0.00000000 |
| 6176 | 2.63 | 0.00000000 |
| 6177 | 3.60 | 0.00000000 |
| 6178 | 11.63 | 0.00000000 |
| 6179 | 7.25 | 0.00000000 |
| 6180 | 2.14 | 0.00000000 |
| 6181 | 8.71 | 0.00000000 |
| 6182 | 2.57 | 0.00000000 |
| 6183 | 13.00 | 0.00000000 |
| 6184 | 9.30 | 0.00000000 |
| 6185 | 2.14 | 0.00000000 |
| 6186 | 2.10 | 0.00000000 |
| 6187 | 7.22 | 0.00000000 |
| 6188 | 2.50 | 0.00000000 |
| 6189 | 7.22 | 0.00000000 |
| 6190 | 14.00 | 0.00000000 |
| 6191 | 14.22 | 0.00000000 |
| 6192 | 9.78 | 0.00000000 |
| 6193 | 3.75 | 0.00000000 |
| 6194 | 26.83 | 0.00000000 |
| 6195 | 13.14 | 0.00000000 |
| 6196 | 16.56 | 0.00000000 |
| 6197 | 11.50 | 0.00000000 |
| 6198 | 10.63 | 0.00000000 |
| 6199 | 6.50 | 0.00000000 |
| 6200 | 4.00 | 0.00000000 |
| 6201 | 11.75 | 0.00000000 |
| 6202 | 7.29 | 0.00000000 |
| 6203 | 8.78 | 0.00000000 |
| 6204 | 2.90 | 0.00000000 |

| | | |
|---|---|---|
| 6205 | 12.50 | 0.00000000 |
| 6206 | 11.36 | 0.00000000 |
| 6207 | 11.44 | 0.00000000 |
| 6208 | 1.57 | 0.00000000 |
| 6209 | 10.30 | 0.00000000 |
| 6210 | 2.63 | 0.00000000 |
| 6211 | 6.86 | 0.00000000 |
| 6212 | 1.90 | 0.00000000 |
| 6213 | 2.25 | 0.00000000 |
| 6214 | 3.86 | 0.00000000 |
| 6215 | 10.11 | 0.00000000 |
| 6216 | 3.00 | 0.00000000 |
| 6217 | 23.22 | 0.00000000 |
| 6218 | 8.91 | 0.00000000 |
| 6219 | 8.90 | 0.00000000 |
| 6220 | 7.88 | 0.00000000 |
| 6221 | 6.30 | 0.00000000 |
| 6222 | 10.56 | 0.00000000 |
| 6223 | 20.60 | 0.00000000 |
| 6224 | 1.83 | 0.00000000 |
| 6225 | 1.71 | 0.00000000 |
| 6226 | 8.00 | 0.00000000 |
| 6227 | 13.82 | 0.00000000 |
| 6228 | 1.56 | 0.00000000 |
| 6229 | 7.11 | 0.00000000 |
| 6230 | 10.88 | 0.00000000 |
| 6231 | 7.11 | 0.00000000 |
| 6232 | 2.92 | 0.00000000 |
| 6233 | 16.38 | 0.00000000 |
| 6234 | 16.90 | 0.00000000 |
| 6235 | 11.86 | 0.00000000 |
| 6236 | 8.43 | 0.00000000 |
| 6237 | 4.89 | 0.00000000 |
| 6238 | 8.38 | 0.00000000 |
| 6239 | 3.00 | 0.00000000 |
| 6240 | 1.56 | 0.00000000 |
| 6241 | 5.75 | 0.00000000 |
| 6242 | 20.75 | 0.00000000 |
| 6243 | 3.00 | 0.00000000 |
| 6244 | 5.56 | 0.00000000 |
| 6245 | 7.43 | 0.00000000 |
| 6246 | 12.00 | 0.00000000 |
| 6247 | 2.00 | 0.00000000 |
| 6248 | 11.75 | 0.00000000 |

| | | |
|---|---|---|
| 6249 | 2.29 | 0.00000000 |
| 6250 | 2.36 | 0.00000000 |
| 6251 | 8.50 | 0.00000000 |
| 6252 | 8.42 | 0.00000000 |
| 6253 | 8.58 | 0.00000000 |
| 6254 | 14.33 | 0.00000000 |
| 6255 | 11.00 | 0.00000000 |
| 6256 | 8.50 | 0.00000000 |
| 6257 | 7.71 | 0.00000000 |
| 6258 | 4.00 | 0.00000000 |
| 6259 | 7.00 | 0.00000000 |
| 6260 | 18.75 | 0.00000000 |
| 6261 | 11.89 | 0.00000000 |
| 6262 | 9.63 | 0.00000000 |
| 6263 | 4.75 | 0.00000000 |
| 6264 | 17.75 | 0.00000000 |
| 6265 | 13.11 | 0.00000000 |
| 6266 | 12.29 | 0.00000000 |
| 6267 | 14.22 | 0.00000000 |
| 6268 | 7.17 | 0.00000000 |
| 6269 | 4.20 | 0.00000000 |
| 6270 | 16.80 | 0.00000000 |
| 6271 | 2.13 | 0.00000000 |
| 6272 | 7.14 | 0.00000000 |
| 6273 | 11.78 | 0.00000000 |
| 6274 | 5.70 | 0.00000000 |
| 6275 | 10.11 | 0.00000000 |
| 6276 | 2.43 | 0.00000000 |
| 6277 | 6.08 | 0.00000000 |
| 6278 | 4.50 | 0.00000000 |
| 6279 | 2.78 | 0.00000000 |
| 6280 | 11.57 | 0.00000000 |
| 6281 | 13.30 | 0.00000000 |
| 6282 | 15.50 | 0.00000000 |
| 6283 | 20.67 | 0.00000000 |
| 6284 | 9.17 | 0.00000000 |
| 6285 | 17.50 | 0.00000000 |
| 6286 | 9.50 | 0.00000000 |
| 6287 | 7.50 | 0.00000000 |
| 6288 | 2.75 | 0.00000000 |
| 6289 | 20.50 | 0.00000000 |
| 6290 | 2.33 | 0.00000000 |
| 6291 | 3.17 | 0.00000000 |
| 6292 | 6.78 | 0.00000000 |

| | | |
|---|---|---|
| 6293 | 9.22 | 0.00000000 |
| 6294 | 12.38 | 0.00000000 |
| 6295 | 3.83 | 0.00000000 |
| 6296 | 10.43 | 0.00000000 |
| 6297 | 8.50 | 0.00000000 |
| 6298 | 2.29 | 0.00000000 |
| 6299 | 12.40 | 0.00000000 |
| 6300 | 5.78 | 0.00000000 |
| 6301 | 11.14 | 0.00000000 |
| 6302 | 2.29 | 0.00000000 |
| 6303 | 2.86 | 0.00000000 |
| 6304 | 3.25 | 0.00000000 |
| 6305 | 17.13 | 0.00000000 |
| 6306 | 11.43 | 0.00000000 |
| 6307 | 7.00 | 0.00000000 |
| 6308 | 1.17 | 0.00000000 |
| 6309 | 1.80 | 0.00000000 |
| 6310 | 4.25 | 0.00000000 |
| 6311 | 5.25 | 0.00000000 |
| 6312 | 14.09 | 0.00000000 |
| 6313 | 4.00 | 0.00000000 |
| 6314 | 5.00 | 0.00000000 |
| 6315 | 5.78 | 0.00000000 |
| 6316 | 9.20 | 0.00000000 |
| 6317 | 2.17 | 0.00000000 |
| 6318 | 2.33 | 0.00000000 |
| 6319 | 7.11 | 0.00000000 |
| 6320 | 4.73 | 0.00000000 |
| 6321 | 13.50 | 0.00000000 |
| 6322 | 2.27 | 0.00000000 |
| 6323 | 19.43 | 0.00000000 |
| 6324 | 8.00 | 0.00000000 |
| 6325 | 3.33 | 0.00000000 |
| 6326 | 2.56 | 0.00000000 |
| 6327 | 8.43 | 0.00000000 |
| 6328 | 16.20 | 0.00000000 |
| 6329 | 10.56 | 0.00000000 |
| 6330 | 13.10 | 0.00000000 |
| 6331 | 9.75 | 0.00000000 |
| 6332 | 12.86 | 0.00000000 |
| 6333 | 14.38 | 0.00000000 |
| 6334 | 2.00 | 0.00000000 |
| 6335 | 7.29 | 0.00000000 |
| 6336 | 10.30 | 0.00000000 |

| | | |
|---|---|---|
| 6337 | 6.88 | 0.00000000 |
| 6338 | 37.50 | 0.00000000 |
| 6339 | 3.80 | 0.00000000 |
| 6340 | 2.25 | 0.00000000 |
| 6341 | 15.11 | 0.00000000 |
| 6342 | 8.33 | 0.00000000 |
| 6343 | 15.50 | 0.00000000 |
| 6344 | 6.57 | 0.00000000 |
| 6345 | 1.33 | 0.00000000 |
| 6346 | 2.50 | 0.00000000 |
| 6347 | 7.67 | 0.00000000 |
| 6348 | 2.43 | 0.00000000 |
| 6349 | 7.00 | 0.00000000 |
| 6350 | 13.43 | 0.00000000 |
| 6351 | 10.75 | 0.00000000 |
| 6352 | 3.63 | 0.00000000 |
| 6353 | 2.43 | 0.00000000 |
| 6354 | 13.13 | 0.00000000 |
| 6355 | 6.00 | 0.00000000 |
| 6356 | 3.86 | 0.00000000 |
| 6357 | 10.67 | 0.00000000 |
| 6358 | 10.80 | 0.00000000 |
| 6359 | 14.89 | 0.00000000 |
| 6360 | 13.75 | 0.00000000 |
| 6361 | 15.25 | 0.00000000 |
| 6362 | 16.14 | 0.00000000 |
| 6363 | 3.17 | 0.00000000 |
| 6364 | 6.38 | 0.00000000 |
| 6365 | 6.09 | 0.00000000 |
| 6366 | 9.40 | 0.00000000 |
| 6367 | 5.38 | 0.00000000 |
| 6368 | 18.57 | 0.00000000 |
| 6369 | 1.67 | 0.00000000 |
| 6370 | 2.20 | 0.00000000 |
| 6371 | 6.56 | 0.00000000 |
| 6372 | 13.00 | 0.00000000 |
| 6373 | 2.60 | 0.00000000 |
| 6374 | 9.22 | 0.00000000 |
| 6375 | 7.75 | 0.00000000 |
| 6376 | 6.64 | 0.00000000 |
| 6377 | 16.80 | 0.00000000 |
| 6378 | 9.40 | 0.00000000 |
| 6379 | 11.86 | 0.00000000 |
| 6380 | 4.82 | 0.00000000 |

| | | |
|---|---|---|
| 6381 | 7.42 | 0.00000000 |
| 6382 | 1.43 | 0.00000000 |
| 6383 | 1.44 | 0.00000000 |
| 6384 | 9.10 | 0.00000000 |
| 6385 | 2.33 | 0.00000000 |
| 6386 | 10.90 | 0.00000000 |
| 6387 | 13.42 | 0.00000000 |
| 6388 | 7.71 | 0.00000000 |
| 6389 | 6.71 | 0.00000000 |
| 6390 | 1.86 | 0.00000000 |
| 6391 | 4.43 | 0.00000000 |
| 6392 | 11.50 | 0.00000000 |
| 6393 | 9.44 | 0.00000000 |
| 6394 | 5.91 | 0.00000000 |
| 6395 | 14.86 | 0.00000000 |
| 6396 | 11.90 | 0.00000000 |
| 6397 | 2.63 | 0.00000000 |
| 6398 | 14.63 | 0.00000000 |
| 6399 | 12.80 | 0.00000000 |
| 6400 | 14.00 | 0.00000000 |
| 6401 | 1.50 | 0.00000000 |
| 6402 | 10.00 | 0.00000000 |
| 6403 | 6.25 | 0.00000000 |
| 6404 | 6.40 | 0.00000000 |
| 6405 | 16.14 | 0.00000000 |
| 6406 | 1.57 | 0.00000000 |
| 6407 | 4.67 | 0.00000000 |
| 6408 | 6.70 | 0.00000000 |
| 6409 | 12.86 | 0.00000000 |
| 6410 | 3.80 | 0.00000000 |
| 6411 | 11.67 | 0.00000000 |
| 6412 | 5.11 | 0.00000000 |
| 6413 | 5.11 | 0.00000000 |
| 6414 | 2.71 | 0.00000000 |
| 6415 | 7.57 | 0.00000000 |
| 6416 | 10.40 | 0.00000000 |
| 6417 | 10.60 | 0.00000000 |
| 6418 | 2.38 | 0.00000000 |
| 6419 | 1.71 | 0.00000000 |
| 6420 | 3.57 | 0.00000000 |
| 6421 | 14.14 | 0.00000000 |
| 6422 | 23.43 | 0.00000000 |
| 6423 | 6.75 | 0.00000000 |
| 6424 | 10.60 | 0.00000000 |

| | | |
|---|---|---|
| 6425 | 8.90 | 0.00000000 |
| 6426 | 13.86 | 0.00000000 |
| 6427 | 3.33 | 0.00000000 |
| 6428 | 3.00 | 0.00000000 |
| 6429 | 20.83 | 0.00000000 |
| 6430 | 4.67 | 0.00000000 |
| 6431 | 14.00 | 0.00000000 |
| 6432 | 9.60 | 0.00000000 |
| 6433 | 2.38 | 0.00000000 |
| 6434 | 12.13 | 0.00000000 |
| 6435 | 30.89 | 0.00000000 |
| 6436 | 7.00 | 0.00000000 |
| 6437 | 2.57 | 0.00000000 |
| 6438 | 8.22 | 0.00000000 |
| 6439 | 6.57 | 0.00000000 |
| 6440 | 2.00 | 0.00000000 |
| 6441 | 7.60 | 0.00000000 |
| 6442 | 12.00 | 0.00000000 |
| 6443 | 19.50 | 0.00000000 |
| 6444 | 1.86 | 0.00000000 |
| 6445 | 7.29 | 0.00000000 |
| 6446 | 11.25 | 0.00000000 |
| 6447 | 8.17 | 0.00000000 |
| 6448 | 1.83 | 0.00000000 |
| 6449 | 2.83 | 0.00000000 |
| 6450 | 2.14 | 0.00000000 |
| 6451 | 20.17 | 0.00000000 |
| 6452 | 3.00 | 0.00000000 |
| 6453 | 8.60 | 0.00000000 |
| 6454 | 2.50 | 0.00000000 |
| 6455 | 14.67 | 0.00000000 |
| 6456 | 3.00 | 0.00000000 |
| 6457 | 11.75 | 0.00000000 |
| 6458 | 2.63 | 0.00000000 |
| 6459 | 1.44 | 0.00000000 |
| 6460 | 18.25 | 0.00000000 |
| 6461 | 22.33 | 0.00000000 |
| 6462 | 2.29 | 0.00000000 |
| 6463 | 4.30 | 0.00000000 |
| 6464 | 10.13 | 0.00000000 |
| 6465 | 3.25 | 0.00000000 |
| 6466 | 2.33 | 0.00000000 |
| 6467 | 2.83 | 0.00000000 |
| 6468 | 3.00 | 0.00000000 |

| | | |
|---|---|---|
| 6469 | 16.80 | 0.00000000 |
| 6470 | 16.25 | 0.00000000 |
| 6471 | 2.29 | 0.00000000 |
| 6472 | 4.17 | 0.00000000 |
| 6473 | 6.00 | 0.00000000 |
| 6474 | 7.63 | 0.00000000 |
| 6475 | 11.14 | 0.00000000 |
| 6476 | 8.00 | 0.00000000 |
| 6477 | 5.78 | 0.00000000 |
| 6478 | 17.00 | 0.00000000 |
| 6479 | 6.43 | 0.00000000 |
| 6480 | 1.00 | 0.00000000 |
| 6481 | 10.33 | 0.00000000 |
| 6482 | 10.25 | 0.00000000 |
| 6483 | 9.40 | 0.00000000 |
| 6484 | 1.75 | 0.00000000 |
| 6485 | 16.40 | 0.00000000 |
| 6486 | 5.50 | 0.00000000 |
| 6487 | 14.00 | 0.00000000 |
| 6488 | 11.43 | 0.00000000 |
| 6489 | 1.60 | 0.00000000 |
| 6490 | 8.25 | 0.00000000 |
| 6491 | 2.67 | 0.00000000 |
| 6492 | 11.00 | 0.00000000 |
| 6493 | 2.40 | 0.00000000 |
| 6494 | 16.13 | 0.00000000 |
| 6495 | 1.67 | 0.00000000 |
| 6496 | 18.14 | 0.00000000 |
| 6497 | 2.86 | 0.00000000 |
| 6498 | 2.00 | 0.00000000 |
| 6499 | 7.00 | 0.00000000 |
| 6500 | 10.50 | 0.00000000 |
| 6501 | 7.14 | 0.00000000 |
| 6502 | 1.33 | 0.00000000 |
| 6503 | 2.20 | 0.00000000 |
| 6504 | 7.00 | 0.00000000 |
| 6505 | 2.25 | 0.00000000 |
| 6506 | 4.60 | 0.00000000 |
| 6507 | 10.20 | 0.00000000 |
| 6508 | 8.67 | 0.00000000 |
| 6509 | 6.00 | 0.00000000 |
| 6510 | 2.75 | 0.00000000 |
| 6511 | 2.67 | 0.00000000 |
| 6512 | 5.71 | 0.00000000 |

| | | |
|---|---|---|
| 6513 | 12.44 | 0.00000000 |
| 6514 | 13.00 | 0.00000000 |
| 6515 | 8.17 | 0.00000000 |
| 6516 | 1.67 | 0.00000000 |
| 6517 | 8.60 | 0.00000000 |
| 6518 | 10.00 | 0.00000000 |
| 6519 | 11.71 | 0.00000000 |
| 6520 | 1.75 | 0.00000000 |
| 6521 | 1.86 | 0.00000000 |
| 6522 | 2.00 | 0.00000000 |
| 6523 | 5.50 | 0.00000000 |
| 6524 | 6.63 | 0.00000000 |
| 6525 | 6.29 | 0.00000000 |
| 6526 | 6.13 | 0.00000000 |
| 6527 | 3.14 | 0.00000000 |
| 6528 | 11.13 | 0.00000000 |
| 6529 | 15.50 | 0.00000000 |
| 6530 | 10.11 | 0.00000000 |
| 6531 | 11.25 | 0.00000000 |
| 6532 | 6.43 | 0.00000000 |
| 6533 | 2.20 | 0.00000000 |
| 6534 | 2.00 | 0.00000000 |
| 6535 | 10.25 | 0.00000000 |
| 6536 | 6.38 | 0.00000000 |
| 6537 | 4.20 | 0.00000000 |
| 6538 | 4.67 | 0.00000000 |
| 6539 | 1.83 | 0.00000000 |
| 6540 | 2.00 | 0.00000000 |
| 6541 | 1.83 | 0.00000000 |
| 6542 | 15.17 | 0.00000000 |
| 6543 | 10.20 | 0.00000000 |
| 6544 | 4.13 | 0.00000000 |
| 6545 | 3.50 | 0.00000000 |
| 6546 | 6.56 | 0.00000000 |
| 6547 | 3.44 | 0.00000000 |
| 6548 | 21.25 | 0.00000000 |
| 6549 | 16.00 | 0.00000000 |
| 6550 | 12.50 | 0.00000000 |
| 6551 | 8.00 | 0.00000000 |
| 6552 | 2.13 | 0.00000000 |
| 6553 | 8.71 | 0.00000000 |
| 6554 | 2.67 | 0.00000000 |
| 6555 | 15.75 | 0.00000000 |
| 6556 | 8.27 | 0.00000000 |

| | | |
|---|---|---|
| 6557 | 9.56 | 0.00000000 |
| 6558 | 9.89 | 0.00000000 |
| 6559 | 5.67 | 0.00000000 |
| 6560 | 7.33 | 0.00000000 |
| 6561 | 14.86 | 0.00000000 |
| 6562 | 16.00 | 0.00000000 |
| 6563 | 2.00 | 0.00000000 |
| 6564 | 2.86 | 0.00000000 |
| 6565 | 6.50 | 0.00000000 |
| 6566 | 11.71 | 0.00000000 |
| 6567 | 3.60 | 0.00000000 |
| 6568 | 6.75 | 0.00000000 |
| 6569 | 5.56 | 0.00000000 |
| 6570 | 2.50 | 0.00000000 |
| 6571 | 6.44 | 0.00000000 |
| 6572 | 2.67 | 0.00000000 |
| 6573 | 8.50 | 0.00000000 |
| 6574 | 11.33 | 0.00000000 |
| 6575 | 2.80 | 0.00000000 |
| 6576 | 11.57 | 0.00000000 |
| 6577 | 1.50 | 0.00000000 |
| 6578 | 9.83 | 0.00000000 |
| 6579 | 10.13 | 0.00000000 |
| 6580 | 11.71 | 0.00000000 |
| 6581 | 8.17 | 0.00000000 |
| 6582 | 4.00 | 0.00000000 |
| 6583 | 8.40 | 0.00000000 |
| 6584 | 6.67 | 0.00000000 |
| 6585 | 5.20 | 0.00000000 |
| 6586 | 6.50 | 0.00000000 |
| 6587 | 10.00 | 0.00000000 |
| 6588 | 4.50 | 0.00000000 |
| 6589 | 7.57 | 0.00000000 |
| 6590 | 6.57 | 0.00000000 |
| 6591 | 7.67 | 0.00000000 |
| 6592 | 2.33 | 0.00000000 |
| 6593 | 10.60 | 0.00000000 |
| 6594 | 5.11 | 0.00000000 |
| 6595 | 8.60 | 0.00000000 |
| 6596 | 2.75 | 0.00000000 |
| 6597 | 4.00 | 0.00000000 |
| 6598 | 17.20 | 0.00000000 |
| 6599 | 7.43 | 0.00000000 |
| 6600 | 6.00 | 0.00000000 |

| | | |
|---|---|---|
| 6601 | 8.50 | 0.00000000 |
| 6602 | 7.00 | 0.00000000 |
| 6603 | 6.75 | 0.00000000 |
| 6604 | 5.88 | 0.00000000 |
| 6605 | 10.00 | 0.00000000 |
| 6606 | 2.71 | 0.00000000 |
| 6607 | 5.88 | 0.00000000 |
| 6608 | 1.00 | 0.00000000 |
| 6609 | 7.71 | 0.00000000 |
| 6610 | 2.13 | 0.00000000 |
| 6611 | 4.71 | 0.00000000 |
| 6612 | 3.71 | 0.00000000 |
| 6613 | 14.09 | 0.00000000 |
| 6614 | 16.50 | 0.00000000 |
| 6615 | 12.13 | 0.00000000 |
| 6616 | 2.33 | 0.00000000 |
| 6617 | 7.13 | 0.00000000 |
| 6618 | 14.18 | 0.00000000 |
| 6619 | 3.33 | 0.00000000 |
| 6620 | 10.17 | 0.00000000 |
| 6621 | 8.86 | 0.00000000 |
| 6622 | 26.00 | 0.00000000 |
| 6623 | 7.00 | 0.00000000 |
| 6624 | 7.80 | 0.00000000 |
| 6625 | 2.71 | 0.00000000 |
| 6626 | 8.14 | 0.00000000 |
| 6627 | 20.17 | 0.00000000 |
| 6628 | 7.00 | 0.00000000 |
| 6629 | 3.20 | 0.00000000 |
| 6630 | 2.83 | 0.00000000 |
| 6631 | 6.78 | 0.00000000 |
| 6632 | 11.89 | 0.00000000 |
| 6633 | 3.67 | 0.00000000 |
| 6634 | 3.18 | 0.00000000 |
| 6635 | 36.43 | 0.00000000 |
| 6636 | 8.25 | 0.00000000 |
| 6637 | 9.14 | 0.00000000 |
| 6638 | 2.13 | 0.00000000 |
| 6639 | 15.14 | 0.00000000 |
| 6640 | 11.00 | 0.00000000 |
| 6641 | 10.22 | 0.00000000 |
| 6642 | 9.11 | 0.00000000 |
| 6643 | 5.88 | 0.00000000 |
| 6644 | 11.57 | 0.00000000 |

| | | |
|---|---|---|
| 6645 | 7.29 | 0.00000000 |
| 6646 | 7.38 | 0.00000000 |
| 6647 | 11.57 | 0.00000000 |
| 6648 | 14.83 | 0.00000000 |
| 6649 | 28.00 | 0.00000000 |
| 6650 | 9.89 | 0.00000000 |
| 6651 | 7.14 | 0.00000000 |
| 6652 | 26.00 | 0.00000000 |
| 6653 | 10.00 | 0.00000000 |
| 6654 | 3.20 | 0.00000000 |
| 6655 | 1.86 | 0.00000000 |
| 6656 | 3.00 | 0.00000000 |
| 6657 | 6.00 | 0.00000000 |
| 6658 | 7.33 | 0.00000000 |
| 6659 | 2.29 | 0.00000000 |
| 6660 | 12.67 | 0.00000000 |
| 6661 | 14.20 | 0.00000000 |
| 6662 | 6.14 | 0.00000000 |
| 6663 | 1.75 | 0.00000000 |
| 6664 | 2.60 | 0.00000000 |
| 6665 | 2.00 | 0.00000000 |
| 6666 | 2.11 | 0.00000000 |
| 6667 | 2.33 | 0.00000000 |
| 6668 | 9.60 | 0.00000000 |
| 6669 | 14.83 | 0.00000000 |
| 6670 | 8.43 | 0.00000000 |
| 6671 | 3.00 | 0.00000000 |
| 6672 | 8.70 | 0.00000000 |
| 6673 | 7.88 | 0.00000000 |
| 6674 | 7.57 | 0.00000000 |
| 6675 | 8.63 | 0.00000000 |
| 6676 | 23.00 | 0.00000000 |
| 6677 | 8.60 | 0.00000000 |
| 6678 | 17.63 | 0.00000000 |
| 6679 | 13.00 | 0.00000000 |
| 6680 | 5.11 | 0.00000000 |
| 6681 | 7.91 | 0.00000000 |
| 6682 | 2.00 | 0.00000000 |
| 6683 | 2.29 | 0.00000000 |
| 6684 | 6.63 | 0.00000000 |
| 6685 | 6.13 | 0.00000000 |
| 6686 | 5.56 | 0.00000000 |
| 6687 | 7.71 | 0.00000000 |
| 6688 | 10.40 | 0.00000000 |

| | | |
|---|---|---|
| 6689 | 18.75 | 0.00000000 |
| 6690 | 3.33 | 0.00000000 |
| 6691 | 4.91 | 0.00000000 |
| 6692 | 5.13 | 0.00000000 |
| 6693 | 2.00 | 0.00000000 |
| 6694 | 1.67 | 0.00000000 |
| 6695 | 1.56 | 0.00000000 |
| 6696 | 2.14 | 0.00000000 |
| 6697 | 11.25 | 0.00000000 |
| 6698 | 2.33 | 0.00000000 |
| 6699 | 4.73 | 0.00000000 |
| 6700 | 9.17 | 0.00000000 |
| 6701 | 12.43 | 0.00000000 |
| 6702 | 5.80 | 0.00000000 |
| 6703 | 1.90 | 0.00000000 |
| 6704 | 1.88 | 0.00000000 |
| 6705 | 3.27 | 0.00000000 |
| 6706 | 13.58 | 0.00000000 |
| 6707 | 8.55 | 0.00000000 |
| 6708 | 7.55 | 0.00000000 |
| 6709 | 11.00 | 0.00000000 |
| 6710 | 14.18 | 0.00000000 |
| 6711 | 9.67 | 0.00000000 |
| 6712 | 6.70 | 0.00000000 |
| 6713 | 7.33 | 0.00000000 |
| 6714 | 5.78 | 0.00000000 |
| 6715 | 6.43 | 0.00000000 |
| 6716 | 12.25 | 0.00000000 |
| 6717 | 4.43 | 0.00000000 |
| 6718 | 15.11 | 0.00000000 |
| 6719 | 8.14 | 0.00000000 |
| 6720 | 6.38 | 0.00000000 |
| 6721 | 13.60 | 0.00000000 |
| 6722 | 8.20 | 0.00000000 |
| 6723 | 6.91 | 0.00000000 |
| 6724 | 9.50 | 0.00000000 |
| 6725 | 6.22 | 0.00000000 |
| 6726 | 4.75 | 0.00000000 |
| 6727 | 5.78 | 0.00000000 |
| 6728 | 17.00 | 0.00000000 |
| 6729 | 18.30 | 0.00000000 |
| 6730 | 2.88 | 0.00000000 |
| 6731 | 7.17 | 0.00000000 |
| 6732 | 9.89 | 0.00000000 |

| | | |
|---|---|---|
| 6733 | 2.40 | 0.00000000 |
| 6734 | 3.43 | 0.00000000 |
| 6735 | 6.78 | 0.00000000 |
| 6736 | 15.88 | 0.00000000 |
| 6737 | 2.13 | 0.00000000 |
| 6738 | 6.29 | 0.00000000 |
| 6739 | 11.80 | 0.00000000 |
| 6740 | 8.43 | 0.00000000 |
| 6741 | 5.22 | 0.00000000 |
| 6742 | 7.50 | 0.00000000 |
| 6743 | 13.33 | 0.00000000 |
| 6744 | 2.60 | 0.00000000 |
| 6745 | 7.00 | 0.00000000 |
| 6746 | 9.00 | 0.00000000 |
| 6747 | 8.50 | 0.00000000 |
| 6748 | 7.13 | 0.00000000 |
| 6749 | 5.50 | 0.00000000 |
| 6750 | 2.44 | 0.00000000 |
| 6751 | 9.10 | 0.00000000 |
| 6752 | 4.20 | 0.00000000 |
| 6753 | 6.69 | 0.00000000 |
| 6754 | 8.30 | 0.00000000 |
| 6755 | 5.30 | 0.00000000 |
| 6756 | 2.42 | 0.00000000 |
| 6757 | 9.82 | 0.00000000 |
| 6758 | 4.14 | 0.00000000 |
| 6759 | 12.40 | 0.00000000 |
| 6760 | 9.40 | 0.00000000 |
| 6761 | 7.80 | 0.00000000 |
| 6762 | 15.45 | 0.00000000 |
| 6763 | 22.91 | 0.00000000 |
| 6764 | 6.60 | 0.00000000 |
| 6765 | 13.57 | 0.00000000 |
| 6766 | 9.22 | 0.00000000 |
| 6767 | 11.29 | 0.00000000 |
| 6768 | 2.17 | 0.00000000 |
| 6769 | 9.00 | 0.00000000 |
| 6770 | 7.30 | 0.00000000 |
| 6771 | 7.18 | 0.00000000 |
| 6772 | 9.82 | 0.00000000 |
| 6773 | 6.86 | 0.00000000 |
| 6774 | 7.91 | 0.00000000 |
| 6775 | 11.33 | 0.00000000 |
| 6776 | 3.25 | 0.00000000 |

| | | |
|---|---|---|
| 6777 | 3.86 | 0.00000000 |
| 6778 | 12.56 | 0.00000000 |
| 6779 | 15.40 | 0.00000000 |
| 6780 | 2.71 | 0.00000000 |
| 6781 | 6.00 | 0.00000000 |
| 6782 | 15.67 | 0.00000000 |
| 6783 | 6.20 | 0.00000000 |
| 6784 | 2.00 | 0.00000000 |
| 6785 | 14.17 | 0.00000000 |
| 6786 | 26.67 | 0.00000000 |
| 6787 | 1.75 | 0.00000000 |
| 6788 | 1.14 | 0.00000000 |
| 6789 | 1.50 | 0.00000000 |
| 6790 | 2.33 | 0.00000000 |
| 6791 | 6.67 | 0.00000000 |
| 6792 | 10.75 | 0.00000000 |
| 6793 | 3.50 | 0.00000000 |
| 6794 | 2.80 | 0.00000000 |
| 6795 | 7.75 | 0.00000000 |
| 6796 | 12.67 | 0.00000000 |
| 6797 | 6.25 | 0.00000000 |
| 6798 | 2.00 | 0.00000000 |
| 6799 | 2.25 | 0.00000000 |
| 6800 | 10.00 | 0.00000000 |
| 6801 | 5.78 | 0.00000000 |
| 6802 | 1.50 | 0.00000000 |
| 6803 | 3.00 | 0.00000000 |
| 6804 | 3.50 | 0.00000000 |
| 6805 | 8.00 | 0.00000000 |
| 6806 | 13.17 | 0.00000000 |
| 6807 | 14.50 | 0.00000000 |
| 6808 | 11.83 | 0.00000000 |
| 6809 | 14.00 | 0.00000000 |
| 6810 | 5.33 | 0.00000000 |
| 6811 | 15.67 | 0.00000000 |
| 6812 | 6.00 | 0.00000000 |
| 6813 | 11.20 | 0.00000000 |
| 6814 | 7.86 | 0.00000000 |
| 6815 | 3.00 | 0.00000000 |
| 6816 | 8.17 | 0.00000000 |
| 6817 | 4.50 | 0.00000000 |
| 6818 | 10.88 | 0.00000000 |
| 6819 | 9.57 | 0.00000000 |
| 6820 | 2.67 | 0.00000000 |

| | | |
|---|---|---|
| 6821 | 14.17 | 0.00000000 |
| 6822 | 9.00 | 0.00000000 |
| 6823 | 10.63 | 0.00000000 |
| 6824 | 8.00 | 0.00000000 |
| 6825 | 19.33 | 0.00000000 |
| 6826 | 1.60 | 0.00000000 |
| 6827 | 1.43 | 0.00000000 |
| 6828 | 11.38 | 0.00000000 |
| 6829 | 2.00 | 0.00000000 |
| 6830 | 2.29 | 0.00000000 |
| 6831 | 13.80 | 0.00000000 |
| 6832 | 1.43 | 0.00000000 |
| 6833 | 6.33 | 0.00000000 |
| 6834 | 2.57 | 0.00000000 |
| 6835 | 7.57 | 0.00000000 |
| 6836 | 2.30 | 0.00000000 |
| 6837 | 11.38 | 0.00000000 |
| 6838 | 1.43 | 0.00000000 |
| 6839 | 19.83 | 0.00000000 |
| 6840 | 2.60 | 0.00000000 |
| 6841 | 7.67 | 0.00000000 |
| 6842 | 11.86 | 0.00000000 |
| 6843 | 9.50 | 0.00000000 |
| 6844 | 7.57 | 0.00000000 |
| 6845 | 2.00 | 0.00000000 |
| 6846 | 5.78 | 0.00000000 |
| 6847 | 12.86 | 0.00000000 |
| 6848 | 1.20 | 0.00000000 |
| 6849 | 2.00 | 0.00000000 |
| 6850 | 3.50 | 0.00000000 |
| 6851 | 6.43 | 0.00000000 |
| 6852 | 5.89 | 0.00000000 |
| 6853 | 6.20 | 0.00000000 |
| 6854 | 12.50 | 0.00000000 |
| 6855 | 8.73 | 0.00000000 |
| 6856 | 7.00 | 0.00000000 |
| 6857 | 7.55 | 0.00000000 |
| 6858 | 10.11 | 0.00000000 |
| 6859 | 6.33 | 0.00000000 |
| 6860 | 13.83 | 0.00000000 |
| 6861 | 2.70 | 0.00000000 |
| 6862 | 18.88 | 0.00000000 |
| 6863 | 21.25 | 0.00000000 |
| 6864 | 3.67 | 0.00000000 |

| | | |
|---|---|---|
| 6865 | 12.00 | 0.00000000 |
| 6866 | 20.00 | 0.00000000 |
| 6867 | 5.00 | 0.00000000 |
| 6868 | 2.33 | 0.00000000 |
| 6869 | 2.67 | 0.00000000 |
| 6870 | 10.56 | 0.00000000 |
| 6871 | 12.17 | 0.00000000 |
| 6872 | 15.20 | 0.00000000 |
| 6873 | 1.38 | 0.00000000 |
| 6874 | 1.33 | 0.00000000 |
| 6875 | 10.00 | 0.00000000 |
| 6876 | 2.83 | 0.00000000 |
| 6877 | 6.83 | 0.00000000 |
| 6878 | 7.00 | 0.00000000 |
| 6879 | 2.17 | 0.00000000 |
| 6880 | 7.71 | 0.00000000 |
| 6881 | 8.57 | 0.00000000 |
| 6882 | 9.18 | 0.00000000 |
| 6883 | 4.00 | 0.00000000 |
| 6884 | 7.14 | 0.00000000 |
| 6885 | 1.90 | 0.00000000 |
| 6886 | 11.75 | 0.00000000 |
| 6887 | 8.00 | 0.00000000 |
| 6888 | 8.80 | 0.00000000 |
| 6889 | 9.30 | 0.00000000 |
| 6890 | 9.50 | 0.00000000 |
| 6891 | 6.67 | 0.00000000 |
| 6892 | 5.08 | 0.00000000 |
| 6893 | 10.25 | 0.00000000 |
| 6894 | 11.25 | 0.00000000 |
| 6895 | 1.60 | 0.00000000 |
| 6896 | 10.63 | 0.00000000 |
| 6897 | 9.56 | 0.00000000 |
| 6898 | 6.86 | 0.00000000 |
| 6899 | 7.67 | 0.00000000 |
| 6900 | 7.29 | 0.00000000 |
| 6901 | 6.14 | 0.00000000 |
| 6902 | 9.40 | 0.00000000 |
| 6903 | 9.33 | 0.00000000 |
| 6904 | 14.83 | 0.00000000 |
| 6905 | 5.63 | 0.00000000 |
| 6906 | 7.43 | 0.00000000 |
| 6907 | 2.14 | 0.00000000 |
| 6908 | 6.13 | 0.00000000 |

| | | |
|---|---|---|
| 6909 | 1.43 | 0.00000000 |
| 6910 | 3.29 | 0.00000000 |
| 6911 | 3.20 | 0.00000000 |
| 6912 | 1.67 | 0.00000000 |
| 6913 | 2.50 | 0.00000000 |
| 6914 | 7.00 | 0.00000000 |
| 6915 | 5.55 | 0.00000000 |
| 6916 | 8.18 | 0.00000000 |
| 6917 | 6.30 | 0.00000000 |
| 6918 | 9.27 | 0.00000000 |
| 6919 | 15.50 | 0.00000000 |
| 6920 | 13.86 | 0.00000000 |
| 6921 | 14.00 | 0.00000000 |
| 6922 | 13.17 | 0.00000000 |
| 6923 | 3.20 | 0.00000000 |
| 6924 | 4.73 | 0.00000000 |
| 6925 | 5.00 | 0.00000000 |
| 6926 | 10.13 | 0.00000000 |
| 6927 | 13.71 | 0.00000000 |
| 6928 | 4.43 | 0.00000000 |
| 6929 | 15.57 | 0.00000000 |
| 6930 | 10.22 | 0.00000000 |
| 6931 | 7.13 | 0.00000000 |
| 6932 | 7.86 | 0.00000000 |
| 6933 | 2.00 | 0.00000000 |
| 6934 | 7.88 | 0.00000000 |
| 6935 | 12.50 | 0.00000000 |
| 6936 | 15.17 | 0.00000000 |
| 6937 | 17.44 | 0.00000000 |
| 6938 | 10.38 | 0.00000000 |
| 6939 | 9.75 | 0.00000000 |
| 6940 | 1.50 | 0.00000000 |
| 6941 | 1.17 | 0.00000000 |
| 6942 | 2.50 | 0.00000000 |
| 6943 | 6.14 | 0.00000000 |
| 6944 | 1.75 | 0.00000000 |
| 6945 | 7.13 | 0.00000000 |
| 6946 | 8.55 | 0.00000000 |
| 6947 | 2.25 | 0.00000000 |
| 6948 | 10.75 | 0.00000000 |
| 6949 | 10.88 | 0.00000000 |
| 6950 | 5.18 | 0.00000000 |
| 6951 | 2.00 | 0.00000000 |
| 6952 | 2.33 | 0.00000000 |

| | | |
|---|---|---|
| 6953 | 6.67 | 0.00000000 |
| 6954 | 23.60 | 0.00000000 |
| 6955 | 9.20 | 0.00000000 |
| 6956 | 10.13 | 0.00000000 |
| 6957 | 8.00 | 0.00000000 |
| 6958 | 4.50 | 0.00000000 |
| 6959 | 2.57 | 0.00000000 |
| 6960 | 19.00 | 0.00000000 |
| 6961 | 2.00 | 0.00000000 |
| 6962 | 3.33 | 0.00000000 |
| 6963 | 14.20 | 0.00000000 |
| 6964 | 4.67 | 0.00000000 |
| 6965 | 1.20 | 0.00000000 |
| 6966 | 1.43 | 0.00000000 |
| 6967 | 1.25 | 0.00000000 |
| 6968 | 2.29 | 0.00000000 |
| 6969 | 2.86 | 0.00000000 |
| 6970 | 9.88 | 0.00000000 |
| 6971 | 2.63 | 0.00000000 |
| 6972 | 11.14 | 0.00000000 |
| 6973 | 7.00 | 0.00000000 |
| 6974 | 4.33 | 0.00000000 |
| 6975 | 11.75 | 0.00000000 |
| 6976 | 10.29 | 0.00000000 |
| 6977 | 11.00 | 0.00000000 |
| 6978 | 11.80 | 0.00000000 |
| 6979 | 1.71 | 0.00000000 |
| 6980 | 3.50 | 0.00000000 |
| 6981 | 13.75 | 0.00000000 |
| 6982 | 13.71 | 0.00000000 |
| 6983 | 6.43 | 0.00000000 |
| 6984 | 7.67 | 0.00000000 |
| 6985 | 9.29 | 0.00000000 |
| 6986 | 11.25 | 0.00000000 |
| 6987 | 16.60 | 0.00000000 |
| 6988 | 6.38 | 0.00000000 |
| 6989 | 7.67 | 0.00000000 |
| 6990 | 1.86 | 0.00000000 |
| 6991 | 3.75 | 0.00000000 |
| 6992 | 5.78 | 0.00000000 |
| 6993 | 2.20 | 0.00000000 |
| 6994 | 9.75 | 0.00000000 |
| 6995 | 2.63 | 0.00000000 |
| 6996 | 1.50 | 0.00000000 |

| | | |
|---|---|---|
| 6997 | 17.33 | 0.00000000 |
| 6998 | 5.88 | 0.00000000 |
| 6999 | 13.33 | 0.00000000 |
| 7000 | 9.17 | 0.00000000 |
| 7001 | 7.83 | 0.00000000 |
| 7002 | 3.00 | 0.00000000 |
| 7003 | 10.00 | 0.00000000 |
| 7004 | 2.44 | 0.00000000 |
| 7005 | 8.75 | 0.00000000 |
| 7006 | 11.57 | 0.00000000 |
| 7007 | 16.00 | 0.00000000 |
| 7008 | 9.50 | 0.00000000 |
| 7009 | 16.20 | 0.00000000 |
| 7010 | 11.50 | 0.00000000 |
| 7011 | 2.50 | 0.00000000 |
| 7012 | 15.50 | 0.00000000 |
| 7013 | 8.00 | 0.00000000 |
| 7014 | 17.20 | 0.00000000 |
| 7015 | 16.00 | 0.00000000 |
| 7016 | 14.17 | 0.00000000 |
| 7017 | 1.80 | 0.00000000 |
| 7018 | 7.67 | 0.00000000 |
| 7019 | 36.00 | 0.00000000 |
| 7020 | 1.50 | 0.00000000 |
| 7021 | 12.00 | 0.00000000 |
| 7022 | 2.00 | 0.00000000 |
| 7023 | 22.57 | 0.00000000 |
| 7024 | 17.33 | 0.00000000 |
| 7025 | 1.00 | 0.00000000 |
| 7026 | 9.25 | 0.00000000 |
| 7027 | 1.25 | 0.00000000 |
| 7028 | 1.38 | 0.00000000 |
| 7029 | 3.75 | 0.00000000 |
| 7030 | 1.00 | 0.00000000 |
| 7031 | 1.83 | 0.00000000 |
| 7032 | 10.00 | 0.00000000 |
| 7033 | 2.25 | 0.00000000 |
| 7034 | 2.17 | 0.00000000 |
| 7035 | 19.40 | 0.00000000 |
| 7036 | 21.50 | 0.00000000 |
| 7037 | 1.60 | 0.00000000 |
| 7038 | 7.80 | 0.00000000 |
| 7039 | 8.40 | 0.00000000 |
| 7040 | 4.33 | 0.00000000 |

| | | |
|---|---|---|
| 7041 | 6.63 | 0.00000000 |
| 7042 | 8.14 | 0.00000000 |
| 7043 | 19.83 | 0.00000000 |
| 7044 | 2.67 | 0.00000000 |
| 7045 | 2.00 | 0.00000000 |
| 7046 | 11.00 | 0.00000000 |
| 7047 | 16.00 | 0.00000000 |
| 7048 | 9.80 | 0.00000000 |
| 7049 | 1.20 | 0.00000000 |
| 7050 | 1.00 | 0.00000000 |
| 7051 | 1.67 | 0.00000000 |
| 7052 | 9.00 | 0.00000000 |
| 7053 | 6.71 | 0.00000000 |
| 7054 | 2.00 | 0.00000000 |
| 7055 | 7.00 | 0.00000000 |
| 7056 | 9.80 | 0.00000000 |
| 7057 | 6.20 | 0.00000000 |
| 7058 | 1.33 | 0.00000000 |
| 7059 | 8.80 | 0.00000000 |
| 7060 | 2.00 | 0.00000000 |
| 7061 | 12.00 | 0.00000000 |
| 7062 | 1.67 | 0.00000000 |
| 7063 | 2.20 | 0.00000000 |
| 7064 | 2.00 | 0.00000000 |
| 7065 | 9.00 | 0.00000000 |
| 7066 | 7.17 | 0.00000000 |
| 7067 | 3.75 | 0.00000000 |
| 7068 | 1.88 | 0.00000000 |
| 7069 | 8.67 | 0.00000000 |
| 7070 | 17.29 | 0.00000000 |
| 7071 | 2.22 | 0.00000000 |
| 7072 | 10.00 | 0.00000000 |
| 7073 | 13.50 | 0.00000000 |
| 7074 | 4.33 | 0.00000000 |
| 7075 | 6.17 | 0.00000000 |
| 7076 | 13.00 | 0.00000000 |
| 7077 | 1.00 | 0.00000000 |
| 7078 | 3.71 | 0.00000000 |
| 7079 | 6.00 | 0.00000000 |
| 7080 | 27.67 | 0.00000000 |
| 7081 | 6.86 | 0.00000000 |
| 7082 | 6.71 | 0.00000000 |
| 7083 | 1.00 | 0.00000000 |
| 7084 | 2.20 | 0.00000000 |

| | | |
|---|---|---|
| 7085 | 7.71 | 0.00000000 |
| 7086 | 9.00 | 0.00000000 |
| 7087 | 1.33 | 0.00000000 |
| 7088 | 1.60 | 0.00000000 |
| 7089 | 9.00 | 0.00000000 |
| 7090 | 3.67 | 0.00000000 |
| 7091 | 9.20 | 0.00000000 |
| 7092 | 6.75 | 0.00000000 |
| 7093 | 2.25 | 0.00000000 |
| 7094 | 10.60 | 0.00000000 |
| 7095 | 10.14 | 0.00000000 |
| 7096 | 2.75 | 0.00000000 |
| 7097 | 17.60 | 0.00000000 |
| 7098 | 2.50 | 0.00000000 |
| 7099 | 1.86 | 0.00000000 |
| 7100 | 2.83 | 0.00000000 |
| 7101 | 1.80 | 0.00000000 |
| 7102 | 2.86 | 0.00000000 |
| 7103 | 15.33 | 0.00000000 |
| 7104 | 14.00 | 0.00000000 |
| 7105 | 3.33 | 0.00000000 |
| 7106 | 15.00 | 0.00000000 |
| 7107 | 12.67 | 0.00000000 |
| 7108 | 12.89 | 0.00000000 |
| 7109 | 12.00 | 0.00000000 |
| 7110 | 2.80 | 0.00000000 |
| 7111 | 18.75 | 0.00000000 |
| 7112 | 1.67 | 0.00000000 |
| 7113 | 3.25 | 0.00000000 |
| 7114 | 16.80 | 0.00000000 |
| 7115 | 2.00 | 0.00000000 |
| 7116 | 1.00 | 0.00000000 |
| 7117 | 1.57 | 0.00000000 |
| 7118 | 8.17 | 0.00000000 |
| 7119 | 17.00 | 0.00000000 |
| 7120 | 1.43 | 0.00000000 |
| 7121 | 1.00 | 0.00000000 |
| 7122 | 1.43 | 0.00000000 |
| 7123 | 11.75 | 0.00000000 |
| 7124 | 2.29 | 0.00000000 |
| 7125 | 3.29 | 0.00000000 |
| 7126 | 6.83 | 0.00000000 |
| 7127 | 2.00 | 0.00000000 |
| 7128 | 2.20 | 0.00000000 |

| | | |
|---|---|---|
| 7129 | 6.00 | 0.00000000 |
| 7130 | 7.67 | 0.00000000 |
| 7131 | 7.67 | 0.00000000 |
| 7132 | 3.00 | 0.00000000 |
| 7133 | 11.57 | 0.00000000 |
| 7134 | 7.71 | 0.00000000 |
| 7135 | 8.80 | 0.00000000 |
| 7136 | 2.60 | 0.00000000 |
| 7137 | 2.00 | 0.00000000 |
| 7138 | 9.00 | 0.00000000 |
| 7139 | 9.30 | 0.00000000 |
| 7140 | 14.50 | 0.00000000 |
| 7141 | 1.67 | 0.00000000 |
| 7142 | 15.00 | 0.00000000 |
| 7143 | 12.67 | 0.00000000 |
| 7144 | 1.00 | 0.00000000 |
| 7145 | 9.33 | 0.00000000 |
| 7146 | 1.67 | 0.00000000 |
| 7147 | 2.60 | 0.00000000 |
| 7148 | 11.71 | 0.00000000 |
| 7149 | 3.29 | 0.00000000 |
| 7150 | 5.43 | 0.00000000 |
| 7151 | 2.50 | 0.00000000 |
| 7152 | 9.00 | 0.00000000 |
| 7153 | 7.25 | 0.00000000 |
| 7154 | 23.33 | 0.00000000 |
| 7155 | 8.80 | 0.00000000 |
| 7156 | 2.00 | 0.00000000 |
| 7157 | 5.00 | 0.00000000 |
| 7158 | 2.20 | 0.00000000 |
| 7159 | 7.83 | 0.00000000 |
| 7160 | 17.00 | 0.00000000 |
| 7161 | 9.00 | 0.00000000 |
| 7162 | 9.20 | 0.00000000 |
| 7163 | 3.00 | 0.00000000 |
| 7164 | 11.00 | 0.00000000 |
| 7165 | 1.20 | 0.00000000 |
| 7166 | 7.43 | 0.00000000 |
| 7167 | 2.00 | 0.00000000 |
| 7168 | 10.00 | 0.00000000 |
| 7169 | 2.38 | 0.00000000 |
| 7170 | 2.83 | 0.00000000 |
| 7171 | 2.00 | 0.00000000 |
| 7172 | 16.40 | 0.00000000 |

| | | |
|---|---|---|
| 7173 | 2.71 | 0.00000000 |
| 7174 | 2.83 | 0.00000000 |
| 7175 | 12.89 | 0.00000000 |
| 7176 | 5.00 | 0.00000000 |
| 7177 | 2.33 | 0.00000000 |
| 7178 | 9.50 | 0.00000000 |
| 7179 | 11.00 | 0.00000000 |
| 7180 | 9.38 | 0.00000000 |
| 7181 | 15.50 | 0.00000000 |
| 7182 | 1.00 | 0.00000000 |
| 7183 | 2.33 | 0.00000000 |
| 7184 | 13.33 | 0.00000000 |
| 7185 | 17.67 | 0.00000000 |
| 7186 | 8.60 | 0.00000000 |
| 7187 | 8.80 | 0.00000000 |
| 7188 | 2.25 | 0.00000000 |
| 7189 | 11.25 | 0.00000000 |
| 7190 | 7.80 | 0.00000000 |
| 7191 | 2.40 | 0.00000000 |
| 7192 | 7.40 | 0.00000000 |
| 7193 | 3.17 | 0.00000000 |
| 7194 | 18.20 | 0.00000000 |
| 7195 | 22.80 | 0.00000000 |
| 7196 | 2.20 | 0.00000000 |
| 7197 | 2.14 | 0.00000000 |
| 7198 | 8.43 | 0.00000000 |
| 7199 | 2.40 | 0.00000000 |
| 7200 | 9.80 | 0.00000000 |
| 7201 | 10.00 | 0.00000000 |
| 7202 | 6.00 | 0.00000000 |
| 7203 | 9.75 | 0.00000000 |
| 7204 | 10.40 | 0.00000000 |
| 7205 | 1.50 | 0.00000000 |
| 7206 | 15.33 | 0.00000000 |
| 7207 | 6.71 | 0.00000000 |
| 7208 | 1.71 | 0.00000000 |
| 7209 | 3.14 | 0.00000000 |
| 7210 | 6.57 | 0.00000000 |
| 7211 | 2.00 | 0.00000000 |
| 7212 | 2.25 | 0.00000000 |
| 7213 | 9.29 | 0.00000000 |
| 7214 | 1.44 | 0.00000000 |
| 7215 | 4.25 | 0.00000000 |
| 7216 | 8.25 | 0.00000000 |

| | | |
|---|---|---|
| 7217 | 5.78 | 0.00000000 |
| 7218 | 4.80 | 0.00000000 |
| 7219 | 9.88 | 0.00000000 |
| 7220 | 6.50 | 0.00000000 |
| 7221 | 15.67 | 0.00000000 |
| 7222 | 2.00 | 0.00000000 |
| 7223 | 2.57 | 0.00000000 |
| 7224 | 10.50 | 0.00000000 |
| 7225 | 14.00 | 0.00000000 |
| 7226 | 9.78 | 0.00000000 |
| 7227 | 6.83 | 0.00000000 |
| 7228 | 17.67 | 0.00000000 |
| 7229 | 13.83 | 0.00000000 |
| 7230 | 16.00 | 0.00000000 |
| 7231 | 16.00 | 0.00000000 |
| 7232 | 10.75 | 0.00000000 |
| 7233 | 1.20 | 0.00000000 |
| 7234 | 2.67 | 0.00000000 |
| 7235 | 2.60 | 0.00000000 |
| 7236 | 20.25 | 0.00000000 |
| 7237 | 1.70 | 0.00000000 |
| 7238 | 12.83 | 0.00000000 |
| 7239 | 8.80 | 0.00000000 |
| 7240 | 11.50 | 0.00000000 |
| 7241 | 1.20 | 0.00000000 |
| 7242 | 2.75 | 0.00000000 |
| 7243 | 6.86 | 0.00000000 |
| 7244 | 11.75 | 0.00000000 |
| 7245 | 14.33 | 0.00000000 |
| 7246 | 1.57 | 0.00000000 |
| 7247 | 7.40 | 0.00000000 |
| 7248 | 2.50 | 0.00000000 |
| 7249 | 2.00 | 0.00000000 |
| 7250 | 1.60 | 0.00000000 |
| 7251 | 3.00 | 0.00000000 |
| 7252 | 8.60 | 0.00000000 |
| 7253 | 2.57 | 0.00000000 |
| 7254 | 6.63 | 0.00000000 |
| 7255 | 3.00 | 0.00000000 |
| 7256 | 8.00 | 0.00000000 |
| 7257 | 2.17 | 0.00000000 |
| 7258 | 5.67 | 0.00000000 |
| 7259 | 7.43 | 0.00000000 |
| 7260 | 12.43 | 0.00000000 |

| | | |
|---|---|---|
| 7261 | 9.17 | 0.00000000 |
| 7262 | 9.17 | 0.00000000 |
| 7263 | 9.40 | 0.00000000 |
| 7264 | 11.00 | 0.00000000 |
| 7265 | 13.67 | 0.00000000 |
| 7266 | 8.33 | 0.00000000 |
| 7267 | 6.67 | 0.00000000 |
| 7268 | 1.00 | 0.00000000 |
| 7269 | 1.00 | 0.00000000 |
| 7270 | 1.25 | 0.00000000 |
| 7271 | 1.11 | 0.00000000 |
| 7272 | 1.40 | 0.00000000 |
| 7273 | 11.75 | 0.00000000 |
| 7274 | 3.22 | 0.00000000 |
| 7275 | 9.33 | 0.00000000 |
| 7276 | 2.45 | 0.00000000 |
| 7277 | 6.57 | 0.00000000 |
| 7278 | 5.78 | 0.00000000 |
| 7279 | 6.50 | 0.00000000 |
| 7280 | 5.75 | 0.00000000 |
| 7281 | 2.25 | 0.00000000 |
| 7282 | 14.57 | 0.00000000 |
| 7283 | 5.50 | 0.00000000 |
| 7284 | 1.17 | 0.00000000 |
| 7285 | 2.25 | 0.00000000 |
| 7286 | 4.25 | 0.00000000 |
| 7287 | 9.40 | 0.00000000 |
| 7288 | 15.86 | 0.00000000 |
| 7289 | 9.00 | 0.00000000 |
| 7290 | 3.75 | 0.00000000 |
| 7291 | 12.57 | 0.00000000 |
| 7292 | 8.40 | 0.00000000 |
| 7293 | 2.50 | 0.00000000 |
| 7294 | 1.60 | 0.00000000 |
| 7295 | 1.50 | 0.00000000 |
| 7296 | 2.25 | 0.00000000 |
| 7297 | 9.60 | 0.00000000 |
| 7298 | 8.17 | 0.00000000 |
| 7299 | 7.83 | 0.00000000 |
| 7300 | 2.00 | 0.00000000 |
| 7301 | 9.20 | 0.00000000 |
| 7302 | 16.38 | 0.00000000 |
| 7303 | 11.25 | 0.00000000 |
| 7304 | 9.80 | 0.00000000 |

| | | |
|---|---|---|
| 7305 | 10.25 | 0.00000000 |
| 7306 | 1.60 | 0.00000000 |
| 7307 | 1.50 | 0.00000000 |
| 7308 | 2.25 | 0.00000000 |
| 7309 | 2.71 | 0.00000000 |
| 7310 | 12.14 | 0.00000000 |
| 7311 | 5.88 | 0.00000000 |
| 7312 | 6.50 | 0.00000000 |
| 7313 | 9.00 | 0.00000000 |
| 7314 | 1.20 | 0.00000000 |
| 7315 | 6.86 | 0.00000000 |
| 7316 | 2.00 | 0.00000000 |
| 7317 | 11.00 | 0.00000000 |
| 7318 | 1.00 | 0.00000000 |
| 7319 | 1.80 | 0.00000000 |
| 7320 | 6.71 | 0.00000000 |
| 7321 | 3.80 | 0.00000000 |
| 7322 | 6.88 | 0.00000000 |
| 7323 | 6.75 | 0.00000000 |
| 7324 | 8.40 | 0.00000000 |
| 7325 | 1.50 | 0.00000000 |
| 7326 | 7.88 | 0.00000000 |
| 7327 | 9.60 | 0.00000000 |
| 7328 | 9.22 | 0.00000000 |
| 7329 | 2.00 | 0.00000000 |
| 7330 | 7.14 | 0.00000000 |
| 7331 | 2.86 | 0.00000000 |
| 7332 | 2.50 | 0.00000000 |
| 7333 | 7.29 | 0.00000000 |
| 7334 | 16.00 | 0.00000000 |
| 7335 | 14.00 | 0.00000000 |
| 7336 | 1.50 | 0.00000000 |
| 7337 | 2.00 | 0.00000000 |
| 7338 | 2.80 | 0.00000000 |
| 7339 | 15.40 | 0.00000000 |
| 7340 | 3.83 | 0.00000000 |
| 7341 | 3.14 | 0.00000000 |
| 7342 | 16.00 | 0.00000000 |
| 7343 | 10.75 | 0.00000000 |
| 7344 | 2.60 | 0.00000000 |
| 7345 | 7.13 | 0.00000000 |
| 7346 | 17.60 | 0.00000000 |
| 7347 | 25.20 | 0.00000000 |
| 7348 | 10.86 | 0.00000000 |

| | | |
|---|---|---|
| 7349 | 8.20 | 0.00000000 |
| 7350 | 2.20 | 0.00000000 |
| 7351 | 1.86 | 0.00000000 |
| 7352 | 7.92 | 0.00000000 |
| 7353 | 9.67 | 0.00000000 |
| 7354 | 14.56 | 0.00000000 |
| 7355 | 13.43 | 0.00000000 |
| 7356 | 18.67 | 0.00000000 |
| 7357 | 1.50 | 0.00000000 |
| 7358 | 14.00 | 0.00000000 |
| 7359 | 1.00 | 0.00000000 |
| 7360 | 2.40 | 0.00000000 |
| 7361 | 8.60 | 0.00000000 |
| 7362 | 1.14 | 0.00000000 |
| 7363 | 2.00 | 0.00000000 |
| 7364 | 2.71 | 0.00000000 |
| 7365 | 6.57 | 0.00000000 |
| 7366 | 10.38 | 0.00000000 |
| 7367 | 7.83 | 0.00000000 |
| 7368 | 6.83 | 0.00000000 |
| 7369 | 2.25 | 0.00000000 |
| 7370 | 7.50 | 0.00000000 |
| 7371 | 10.00 | 0.00000000 |
| 7372 | 2.20 | 0.00000000 |
| 7373 | 2.29 | 0.00000000 |
| 7374 | 1.83 | 0.00000000 |
| 7375 | 3.00 | 0.00000000 |
| 7376 | 6.86 | 0.00000000 |
| 7377 | 8.00 | 0.00000000 |
| 7378 | 10.00 | 0.00000000 |
| 7379 | 10.38 | 0.00000000 |
| 7380 | 13.50 | 0.00000000 |
| 7381 | 1.83 | 0.00000000 |
| 7382 | 10.43 | 0.00000000 |
| 7383 | 5.44 | 0.00000000 |
| 7384 | 1.25 | 0.00000000 |
| 7385 | 7.86 | 0.00000000 |
| 7386 | 18.50 | 0.00000000 |
| 7387 | 2.14 | 0.00000000 |
| 7388 | 2.83 | 0.00000000 |
| 7389 | 3.00 | 0.00000000 |
| 7390 | 36.50 | 0.00000000 |
| 7391 | 12.50 | 0.00000000 |
| 7392 | 6.33 | 0.00000000 |

| | | |
|---|---|---|
| 7393 | 9.00 | 0.00000000 |
| 7394 | 1.50 | 0.00000000 |
| 7395 | 2.00 | 0.00000000 |
| 7396 | 9.60 | 0.00000000 |
| 7397 | 11.33 | 0.00000000 |
| 7398 | 1.40 | 0.00000000 |
| 7399 | 1.80 | 0.00000000 |
| 7400 | 13.00 | 0.00000000 |
| 7401 | 1.20 | 0.00000000 |
| 7402 | 27.00 | 0.00000000 |
| 7403 | 4.00 | 0.00000000 |
| 7404 | 8.80 | 0.00000000 |
| 7405 | 2.40 | 0.00000000 |
| 7406 | 2.00 | 0.00000000 |
| 7407 | 5.00 | 0.00000000 |
| 7408 | 19.00 | 0.00000000 |
| 7409 | 9.25 | 0.00000000 |
| 7410 | 3.00 | 0.00000000 |
| 7411 | 2.00 | 0.00000000 |
| 7412 | 1.75 | 0.00000000 |
| 7413 | 12.67 | 0.00000000 |
| 7414 | 3.00 | 0.00000000 |
| 7415 | 13.60 | 0.00000000 |
| 7416 | 2.33 | 0.00000000 |
| 7417 | 11.50 | 0.00000000 |
| 7418 | 1.00 | 0.00000000 |
| 7419 | 1.33 | 0.00000000 |
| 7420 | 2.00 | 0.00000000 |
| 7421 | 2.40 | 0.00000000 |
| 7422 | 1.25 | 0.00000000 |
| 7423 | 8.80 | 0.00000000 |
| 7424 | 1.50 | 0.00000000 |
| 7425 | 1.50 | 0.00000000 |
| 7426 | 3.00 | 0.00000000 |
| 7427 | 12.43 | 0.00000000 |
| 7428 | 1.40 | 0.00000000 |
| 7429 | 1.83 | 0.00000000 |
| 7430 | 2.00 | 0.00000000 |
| 7431 | 3.75 | 0.00000000 |
| 7432 | 12.25 | 0.00000000 |
| 7433 | 1.29 | 0.00000000 |
| 7434 | 7.13 | 0.00000000 |
| 7435 | 6.86 | 0.00000000 |
| 7436 | 15.20 | 0.00000000 |

| | | |
|---|---|---|
| 7437 | 3.20 | 0.00000000 |
| 7438 | 6.25 | 0.00000000 |
| 7439 | 13.60 | 0.00000000 |
| 7440 | 1.40 | 0.00000000 |
| 7441 | 1.50 | 0.00000000 |
| 7442 | 1.50 | 0.00000000 |
| 7443 | 2.60 | 0.00000000 |
| 7444 | 3.00 | 0.00000000 |
| 7445 | 5.71 | 0.00000000 |
| 7446 | 3.00 | 0.00000000 |
| 7447 | 2.75 | 0.00000000 |
| 7448 | 6.00 | 0.00000000 |
| 7449 | 9.50 | 0.00000000 |
| 7450 | 4.67 | 0.00000000 |
| 7451 | 18.83 | 0.00000000 |
| 7452 | 1.75 | 0.00000000 |
| 7453 | 2.43 | 0.00000000 |
| 7454 | 8.33 | 0.00000000 |
| 7455 | 23.80 | 0.00000000 |
| 7456 | 16.83 | 0.00000000 |
| 7457 | 10.63 | 0.00000000 |
| 7458 | 13.83 | 0.00000000 |
| 7459 | 1.50 | 0.00000000 |
| 7460 | 1.20 | 0.00000000 |
| 7461 | 3.00 | 0.00000000 |
| 7462 | 7.83 | 0.00000000 |
| 7463 | 1.83 | 0.00000000 |
| 7464 | 4.33 | 0.00000000 |
| 7465 | 10.50 | 0.00000000 |
| 7466 | 11.70 | 0.00000000 |
| 7467 | 14.00 | 0.00000000 |
| 7468 | 1.67 | 0.00000000 |
| 7469 | 6.86 | 0.00000000 |
| 7470 | 13.33 | 0.00000000 |
| 7471 | 2.00 | 0.00000000 |
| 7472 | 16.20 | 0.00000000 |
| 7473 | 6.67 | 0.00000000 |
| 7474 | 16.00 | 0.00000000 |
| 7475 | 1.00 | 0.00000000 |
| 7476 | 1.25 | 0.00000000 |
| 7477 | 1.00 | 0.00000000 |
| 7478 | 2.75 | 0.00000000 |
| 7479 | 1.25 | 0.00000000 |
| 7480 | 3.20 | 0.00000000 |

| | | |
|---|---|---|
| 7481 | 2.33 | 0.00000000 |
| 7482 | 9.89 | 0.00000000 |
| 7483 | 9.60 | 0.00000000 |
| 7484 | 5.38 | 0.00000000 |
| 7485 | 1.80 | 0.00000000 |
| 7486 | 18.33 | 0.00000000 |
| 7487 | 7.25 | 0.00000000 |
| 7488 | 3.43 | 0.00000000 |
| 7489 | 10.29 | 0.00000000 |
| 7490 | 2.22 | 0.00000000 |
| 7491 | 2.00 | 0.00000000 |
| 7492 | 6.44 | 0.00000000 |
| 7493 | 2.57 | 0.00000000 |
| 7494 | 11.00 | 0.00000000 |
| 7495 | 6.92 | 0.00000000 |
| 7496 | 1.20 | 0.00000000 |
| 7497 | 2.13 | 0.00000000 |
| 7498 | 10.00 | 0.00000000 |
| 7499 | 5.10 | 0.00000000 |
| 7500 | 14.40 | 0.00000000 |
| 7501 | 10.00 | 0.00000000 |
| 7502 | 2.57 | 0.00000000 |
| 7503 | 4.63 | 0.00000000 |
| 7504 | 7.57 | 0.00000000 |
| 7505 | 2.57 | 0.00000000 |
| 7506 | 13.33 | 0.00000000 |
| 7507 | 8.20 | 0.00000000 |
| 7508 | 1.29 | 0.00000000 |
| 7509 | 1.88 | 0.00000000 |
| 7510 | 12.00 | 0.00000000 |
| 7511 | 3.40 | 0.00000000 |
| 7512 | 8.00 | 0.00000000 |
| 7513 | 8.50 | 0.00000000 |
| 7514 | 1.75 | 0.00000000 |
| 7515 | 4.70 | 0.00000000 |
| 7516 | 3.20 | 0.00000000 |
| 7517 | 1.83 | 0.00000000 |
| 7518 | 10.67 | 0.00000000 |
| 7519 | 1.71 | 0.00000000 |
| 7520 | 12.67 | 0.00000000 |
| 7521 | 7.67 | 0.00000000 |
| 7522 | 6.88 | 0.00000000 |
| 7523 | 6.25 | 0.00000000 |
| 7524 | 11.00 | 0.00000000 |

| | | |
|---|---|---|
| 7525 | 7.14 | 0.00000000 |
| 7526 | 7.00 | 0.00000000 |
| 7527 | 15.88 | 0.00000000 |
| 7528 | 9.00 | 0.00000000 |
| 7529 | 9.63 | 0.00000000 |
| 7530 | 11.71 | 0.00000000 |
| 7531 | 11.00 | 0.00000000 |
| 7532 | 1.33 | 0.00000000 |
| 7533 | 12.50 | 0.00000000 |
| 7534 | 10.50 | 0.00000000 |
| 7535 | 19.33 | 0.00000000 |
| 7536 | 1.00 | 0.00000000 |
| 7537 | 2.00 | 0.00000000 |
| 7538 | 2.50 | 0.00000000 |
| 7539 | 1.50 | 0.00000000 |
| 7540 | 7.83 | 0.00000000 |
| 7541 | 1.50 | 0.00000000 |
| 7542 | 2.67 | 0.00000000 |
| 7543 | 16.00 | 0.00000000 |
| 7544 | 1.83 | 0.00000000 |
| 7545 | 6.33 | 0.00000000 |
| 7546 | 1.75 | 0.00000000 |
| 7547 | 3.44 | 0.00000000 |
| 7548 | 17.83 | 0.00000000 |
| 7549 | 1.33 | 0.00000000 |
| 7550 | 7.80 | 0.00000000 |
| 7551 | 2.67 | 0.00000000 |
| 7552 | 2.83 | 0.00000000 |
| 7553 | 7.83 | 0.00000000 |
| 7554 | 8.80 | 0.00000000 |
| 7555 | 7.57 | 0.00000000 |
| 7556 | 7.78 | 0.00000000 |
| 7557 | 9.60 | 0.00000000 |
| 7558 | 1.40 | 0.00000000 |
| 7559 | 2.00 | 0.00000000 |
| 7560 | 6.86 | 0.00000000 |
| 7561 | 8.67 | 0.00000000 |
| 7562 | 2.29 | 0.00000000 |
| 7563 | 7.57 | 0.00000000 |
| 7564 | 20.00 | 0.00000000 |
| 7565 | 10.13 | 0.00000000 |
| 7566 | 2.83 | 0.00000000 |
| 7567 | 9.50 | 0.00000000 |
| 7568 | 7.40 | 0.00000000 |

| | | |
|---|---|---|
| 7569 | 3.50 | 0.00000000 |
| 7570 | 8.33 | 0.00000000 |
| 7571 | 3.29 | 0.00000000 |
| 7572 | 9.60 | 0.00000000 |
| 7573 | 15.40 | 0.00000000 |
| 7574 | 1.70 | 0.00000000 |
| 7575 | 2.71 | 0.00000000 |
| 7576 | 6.40 | 0.00000000 |
| 7577 | 23.40 | 0.00000000 |
| 7578 | 7.29 | 0.00000000 |
| 7579 | 1.80 | 0.00000000 |
| 7580 | 8.67 | 0.00000000 |
| 7581 | 2.29 | 0.00000000 |
| 7582 | 1.67 | 0.00000000 |
| 7583 | 2.00 | 0.00000000 |
| 7584 | 13.00 | 0.00000000 |
| 7585 | 11.57 | 0.00000000 |
| 7586 | 13.43 | 0.00000000 |
| 7587 | 2.50 | 0.00000000 |
| 7588 | 6.63 | 0.00000000 |
| 7589 | 3.40 | 0.00000000 |
| 7590 | 12.13 | 0.00000000 |
| 7591 | 7.00 | 0.00000000 |
| 7592 | 2.20 | 0.00000000 |
| 7593 | 13.33 | 0.00000000 |
| 7594 | 9.86 | 0.00000000 |
| 7595 | 2.00 | 0.00000000 |
| 7596 | 1.71 | 0.00000000 |
| 7597 | 1.60 | 0.00000000 |
| 7598 | 1.25 | 0.00000000 |
| 7599 | 1.71 | 0.00000000 |
| 7600 | 7.83 | 0.00000000 |
| 7601 | 10.86 | 0.00000000 |
| 7602 | 2.33 | 0.00000000 |
| 7603 | 6.00 | 0.00000000 |
| 7604 | 1.14 | 0.00000000 |
| 7605 | 12.00 | 0.00000000 |
| 7606 | 7.00 | 0.00000000 |
| 7607 | 2.00 | 0.00000000 |
| 7608 | 6.50 | 0.00000000 |
| 7609 | 8.17 | 0.00000000 |
| 7610 | 8.00 | 0.00000000 |
| 7611 | 3.25 | 0.00000000 |
| 7612 | 6.67 | 0.00000000 |

| | | |
|---|---|---|
| 7613 | 2.17 | 0.00000000 |
| 7614 | 1.00 | 0.00000000 |
| 7615 | 17.71 | 0.00000000 |
| 7616 | 1.33 | 0.00000000 |
| 7617 | 3.00 | 0.00000000 |
| 7618 | 8.40 | 0.00000000 |
| 7619 | 2.20 | 0.00000000 |
| 7620 | 4.00 | 0.00000000 |
| 7621 | 12.00 | 0.00000000 |
| 7622 | 16.20 | 0.00000000 |
| 7623 | 8.00 | 0.00000000 |
| 7624 | 4.33 | 0.00000000 |
| 7625 | 7.83 | 0.00000000 |
| 7626 | 11.88 | 0.00000000 |
| 7627 | 1.33 | 0.00000000 |
| 7628 | 1.33 | 0.00000000 |
| 7629 | 14.89 | 0.00000000 |
| 7630 | 1.33 | 0.00000000 |
| 7631 | 7.60 | 0.00000000 |
| 7632 | 4.00 | 0.00000000 |
| 7633 | 1.43 | 0.00000000 |
| 7634 | 3.71 | 0.00000000 |
| 7635 | 8.20 | 0.00000000 |
| 7636 | 2.29 | 0.00000000 |
| 7637 | 9.20 | 0.00000000 |
| 7638 | 1.60 | 0.00000000 |
| 7639 | 2.33 | 0.00000000 |
| 7640 | 2.20 | 0.00000000 |
| 7641 | 29.00 | 0.00000000 |
| 7642 | 1.50 | 0.00000000 |
| 7643 | 8.60 | 0.00000000 |
| 7644 | 11.67 | 0.00000000 |
| 7645 | 6.50 | 0.00000000 |
| 7646 | 7.67 | 0.00000000 |
| 7647 | 1.83 | 0.00000000 |
| 7648 | 3.80 | 0.00000000 |
| 7649 | 4.25 | 0.00000000 |
| 7650 | 14.63 | 0.00000000 |
| 7651 | 17.00 | 0.00000000 |
| 7652 | 1.60 | 0.00000000 |
| 7653 | 7.44 | 0.00000000 |
| 7654 | 7.67 | 0.00000000 |
| 7655 | 16.60 | 0.00000000 |
| 7656 | 10.25 | 0.00000000 |

| | | |
|---|---|---|
| 7657 | 8.43 | 0.00000000 |
| 7658 | 11.29 | 0.00000000 |
| 7659 | 13.20 | 0.00000000 |
| 7660 | 1.00 | 0.00000000 |
| 7661 | 15.33 | 0.00000000 |
| 7662 | 1.67 | 0.00000000 |
| 7663 | 10.00 | 0.00000000 |
| 7664 | 21.00 | 0.00000000 |
| 7665 | 2.80 | 0.00000000 |
| 7666 | 12.80 | 0.00000000 |
| 7667 | 15.00 | 0.00000000 |
| 7668 | 14.60 | 0.00000000 |
| 7669 | 3.33 | 0.00000000 |
| 7670 | 38.67 | 0.00000000 |
| 7671 | 6.83 | 0.00000000 |
| 7672 | 22.00 | 0.00000000 |
| 7673 | 2.25 | 0.00000000 |
| 7674 | 1.80 | 0.00000000 |
| 7675 | 9.00 | 0.00000000 |
| 7676 | 3.67 | 0.00000000 |
| 7677 | 1.67 | 0.00000000 |
| 7678 | 7.80 | 0.00000000 |
| 7679 | 9.83 | 0.00000000 |
| 7680 | 20.25 | 0.00000000 |
| 7681 | 1.67 | 0.00000000 |
| 7682 | 12.50 | 0.00000000 |
| 7683 | 1.50 | 0.00000000 |
| 7684 | 1.29 | 0.00000000 |
| 7685 | 1.83 | 0.00000000 |
| 7686 | 1.25 | 0.00000000 |
| 7687 | 1.75 | 0.00000000 |
| 7688 | 9.20 | 0.00000000 |
| 7689 | 3.00 | 0.00000000 |
| 7690 | 8.17 | 0.00000000 |
| 7691 | 7.40 | 0.00000000 |
| 7692 | 9.80 | 0.00000000 |
| 7693 | 7.67 | 0.00000000 |
| 7694 | 14.00 | 0.00000000 |
| 7695 | 1.00 | 0.00000000 |
| 7696 | 7.83 | 0.00000000 |
| 7697 | 11.33 | 0.00000000 |
| 7698 | 22.50 | 0.00000000 |
| 7699 | 1.75 | 0.00000000 |
| 7700 | 2.67 | 0.00000000 |

| | | |
|---|---|---|
| 7701 | 1.33 | 0.00000000 |
| 7702 | 8.17 | 0.00000000 |
| 7703 | 1.14 | 0.00000000 |
| 7704 | 14.60 | 0.00000000 |
| 7705 | 1.50 | 0.00000000 |
| 7706 | 18.80 | 0.00000000 |
| 7707 | 8.29 | 0.00000000 |
| 7708 | 3.33 | 0.00000000 |
| 7709 | 4.27 | 0.00000000 |
| 7710 | 10.25 | 0.00000000 |
| 7711 | 1.60 | 0.00000000 |
| 7712 | 1.40 | 0.00000000 |
| 7713 | 4.75 | 0.00000000 |
| 7714 | 15.40 | 0.00000000 |
| 7715 | 2.80 | 0.00000000 |
| 7716 | 2.83 | 0.00000000 |
| 7717 | 14.00 | 0.00000000 |
| 7718 | 3.75 | 0.00000000 |
| 7719 | 13.50 | 0.00000000 |
| 7720 | 14.00 | 0.00000000 |
| 7721 | 2.40 | 0.00000000 |
| 7722 | 5.13 | 0.00000000 |
| 7723 | 6.17 | 0.00000000 |
| 7724 | 3.00 | 0.00000000 |
| 7725 | 1.00 | 0.00000000 |
| 7726 | 1.00 | 0.00000000 |
| 7727 | 20.50 | 0.00000000 |
| 7728 | 1.60 | 0.00000000 |
| 7729 | 14.33 | 0.00000000 |
| 7730 | 2.50 | 0.00000000 |
| 7731 | 1.80 | 0.00000000 |
| 7732 | 7.14 | 0.00000000 |
| 7733 | 2.40 | 0.00000000 |
| 7734 | 2.00 | 0.00000000 |
| 7735 | 10.25 | 0.00000000 |
| 7736 | 10.80 | 0.00000000 |
| 7737 | 2.33 | 0.00000000 |
| 7738 | 2.50 | 0.00000000 |
| 7739 | 9.40 | 0.00000000 |
| 7740 | 10.25 | 0.00000000 |
| 7741 | 14.00 | 0.00000000 |
| 7742 | 1.83 | 0.00000000 |
| 7743 | 9.50 | 0.00000000 |
| 7744 | 7.57 | 0.00000000 |

| | | |
|---|---|---|
| 7745 | 6.00 | 0.00000000 |
| 7746 | 2.33 | 0.00000000 |
| 7747 | 1.67 | 0.00000000 |
| 7748 | 10.60 | 0.00000000 |
| 7749 | 5.20 | 0.00000000 |
| 7750 | 2.13 | 0.00000000 |
| 7751 | 3.83 | 0.00000000 |
| 7752 | 10.11 | 0.00000000 |
| 7753 | 12.00 | 0.00000000 |
| 7754 | 5.71 | 0.00000000 |
| 7755 | 10.86 | 0.00000000 |
| 7756 | 2.80 | 0.00000000 |
| 7757 | 14.83 | 0.00000000 |
| 7758 | 5.88 | 0.00000000 |
| 7759 | 6.29 | 0.00000000 |
| 7760 | 14.80 | 0.00000000 |
| 7761 | 8.00 | 0.00000000 |
| 7762 | 4.00 | 0.00000000 |
| 7763 | 11.00 | 0.00000000 |
| 7764 | 2.50 | 0.00000000 |
| 7765 | 4.20 | 0.00000000 |
| 7766 | 7.00 | 0.00000000 |
| 7767 | 5.00 | 0.00000000 |
| 7768 | 1.60 | 0.00000000 |
| 7769 | 6.63 | 0.00000000 |
| 7770 | 6.33 | 0.00000000 |
| 7771 | 3.86 | 0.00000000 |
| 7772 | 11.25 | 0.00000000 |
| 7773 | 2.38 | 0.00000000 |
| 7774 | 7.50 | 0.00000000 |
| 7775 | 2.40 | 0.00000000 |
| 7776 | 12.50 | 0.00000000 |
| 7777 | 9.20 | 0.00000000 |
| 7778 | 19.50 | 0.00000000 |
| 7779 | 8.83 | 0.00000000 |
| 7780 | 8.50 | 0.00000000 |
| 7781 | 17.40 | 0.00000000 |
| 7782 | 15.50 | 0.00000000 |
| 7783 | 7.33 | 0.00000000 |
| 7784 | 2.00 | 0.00000000 |
| 7785 | 1.00 | 0.00000000 |
| 7786 | 3.00 | 0.00000000 |
| 7787 | 7.43 | 0.00000000 |
| 7788 | 2.60 | 0.00000000 |

| | | |
|---|---|---|
| 7789 | 13.17 | 0.00000000 |
| 7790 | 7.00 | 0.00000000 |
| 7791 | 13.17 | 0.00000000 |
| 7792 | 24.50 | 0.00000000 |
| 7793 | 10.00 | 0.00000000 |
| 7794 | 1.67 | 0.00000000 |
| 7795 | 1.75 | 0.00000000 |
| 7796 | 3.75 | 0.00000000 |
| 7797 | 2.43 | 0.00000000 |
| 7798 | 9.11 | 0.00000000 |
| 7799 | 16.67 | 0.00000000 |
| 7800 | 4.25 | 0.00000000 |
| 7801 | 22.80 | 0.00000000 |
| 7802 | 10.50 | 0.00000000 |
| 7803 | 13.67 | 0.00000000 |
| 7804 | 6.43 | 0.00000000 |
| 7805 | 4.00 | 0.00000000 |
| 7806 | 2.75 | 0.00000000 |
| 7807 | 13.33 | 0.00000000 |
| 7808 | 3.00 | 0.00000000 |
| 7809 | 17.00 | 0.00000000 |
| 7810 | 10.60 | 0.00000000 |
| 7811 | 1.67 | 0.00000000 |
| 7812 | 2.00 | 0.00000000 |
| 7813 | 10.25 | 0.00000000 |
| 7814 | 1.25 | 0.00000000 |
| 7815 | 1.71 | 0.00000000 |
| 7816 | 10.50 | 0.00000000 |
| 7817 | 3.00 | 0.00000000 |
| 7818 | 7.00 | 0.00000000 |
| 7819 | 7.33 | 0.00000000 |
| 7820 | 10.75 | 0.00000000 |
| 7821 | 2.20 | 0.00000000 |
| 7822 | 1.50 | 0.00000000 |
| 7823 | 2.00 | 0.00000000 |
| 7824 | 6.86 | 0.00000000 |
| 7825 | 8.50 | 0.00000000 |
| 7826 | 1.33 | 0.00000000 |
| 7827 | 2.17 | 0.00000000 |
| 7828 | 6.86 | 0.00000000 |
| 7829 | 1.25 | 0.00000000 |
| 7830 | 2.00 | 0.00000000 |
| 7831 | 6.43 | 0.00000000 |
| 7832 | 36.67 | 0.00000000 |

| | | |
|---|---|---|
| 7833 | 2.00 | 0.00000000 |
| 7834 | 1.25 | 0.00000000 |
| 7835 | 1.00 | 0.00000000 |
| 7836 | 1.71 | 0.00000000 |
| 7837 | 2.89 | 0.00000000 |
| 7838 | 8.50 | 0.00000000 |
| 7839 | 5.38 | 0.00000000 |
| 7840 | 3.20 | 0.00000000 |
| 7841 | 9.63 | 0.00000000 |
| 7842 | 2.67 | 0.00000000 |
| 7843 | 9.50 | 0.00000000 |
| 7844 | 5.50 | 0.00000000 |
| 7845 | 7.83 | 0.00000000 |
| 7846 | 10.20 | 0.00000000 |
| 7847 | 1.50 | 0.00000000 |
| 7848 | 2.50 | 0.00000000 |
| 7849 | 8.00 | 0.00000000 |
| 7850 | 2.29 | 0.00000000 |
| 7851 | 12.00 | 0.00000000 |
| 7852 | 2.00 | 0.00000000 |
| 7853 | 11.14 | 0.00000000 |
| 7854 | 7.57 | 0.00000000 |
| 7855 | 6.75 | 0.00000000 |
| 7856 | 1.33 | 0.00000000 |
| 7857 | 2.00 | 0.00000000 |
| 7858 | 2.33 | 0.00000000 |
| 7859 | 9.33 | 0.00000000 |
| 7860 | 2.00 | 0.00000000 |
| 7861 | 7.71 | 0.00000000 |
| 7862 | 1.83 | 0.00000000 |
| 7863 | 2.60 | 0.00000000 |
| 7864 | 8.71 | 0.00000000 |
| 7865 | 11.29 | 0.00000000 |
| 7866 | 7.43 | 0.00000000 |
| 7867 | 14.60 | 0.00000000 |
| 7868 | 13.17 | 0.00000000 |
| 7869 | 2.29 | 0.00000000 |
| 7870 | 16.40 | 0.00000000 |
| 7871 | 15.60 | 0.00000000 |
| 7872 | 1.20 | 0.00000000 |
| 7873 | 2.71 | 0.00000000 |
| 7874 | 1.00 | 0.00000000 |
| 7875 | 1.25 | 0.00000000 |
| 7876 | 1.33 | 0.00000000 |

| | | |
|---|---|---|
| 7877 | 2.29 | 0.00000000 |
| 7878 | 8.43 | 0.00000000 |
| 7879 | 1.75 | 0.00000000 |
| 7880 | 5.89 | 0.00000000 |
| 7881 | 10.88 | 0.00000000 |
| 7882 | 20.00 | 0.00000000 |
| 7883 | 12.00 | 0.00000000 |
| 7884 | 1.25 | 0.00000000 |
| 7885 | 1.60 | 0.00000000 |
| 7886 | 1.83 | 0.00000000 |
| 7887 | 3.00 | 0.00000000 |
| 7888 | 2.67 | 0.00000000 |
| 7889 | 1.57 | 0.00000000 |
| 7890 | 1.75 | 0.00000000 |
| 7891 | 7.43 | 0.00000000 |
| 7892 | 6.14 | 0.00000000 |
| 7893 | 7.10 | 0.00000000 |
| 7894 | 9.78 | 0.00000000 |
| 7895 | 3.00 | 0.00000000 |
| 7896 | 11.71 | 0.00000000 |
| 7897 | 7.00 | 0.00000000 |
| 7898 | 2.00 | 0.00000000 |
| 7899 | 1.82 | 0.00000000 |
| 7900 | 6.75 | 0.00000000 |
| 7901 | 2.88 | 0.00000000 |
| 7902 | 5.82 | 0.00000000 |
| 7903 | 1.71 | 0.00000000 |
| 7904 | 1.29 | 0.00000000 |
| 7905 | 3.56 | 0.00000000 |
| 7906 | 7.00 | 0.00000000 |
| 7907 | 12.00 | 0.00000000 |
| 7908 | 3.22 | 0.00000000 |
| 7909 | 12.13 | 0.00000000 |
| 7910 | 11.00 | 0.00000000 |
| 7911 | 3.00 | 0.00000000 |
| 7912 | 7.14 | 0.00000000 |
| 7913 | 5.14 | 0.00000000 |
| 7914 | 2.56 | 0.00000000 |
| 7915 | 2.00 | 0.00000000 |
| 7916 | 1.92 | 0.00000000 |
| 7917 | 6.73 | 0.00000000 |
| 7918 | 2.14 | 0.00000000 |
| 7919 | 7.00 | 0.00000000 |
| 7920 | 4.90 | 0.00000000 |

| | | |
|---|---|---|
| 7921 | 10.00 | 0.00000000 |
| 7922 | 2.38 | 0.00000000 |
| 7923 | 6.10 | 0.00000000 |
| 7924 | 2.22 | 0.00000000 |
| 7925 | 2.00 | 0.00000000 |
| 7926 | 10.20 | 0.00000000 |
| 7927 | 2.17 | 0.00000000 |
| 7928 | 5.57 | 0.00000000 |
| 7929 | 6.33 | 0.00000000 |
| 7930 | 1.80 | 0.00000000 |
| 7931 | 2.69 | 0.00000000 |
| 7932 | 2.15 | 0.00000000 |
| 7933 | 8.00 | 0.00000000 |
| 7934 | 8.82 | 0.00000000 |
| 7935 | 2.33 | 0.00000000 |
| 7936 | 9.69 | 0.00000000 |
| 7937 | 2.69 | 0.00000000 |
| 7938 | 2.77 | 0.00000000 |
| 7939 | 3.36 | 0.00000000 |
| 7940 | 4.00 | 0.00000000 |
| 7941 | 3.27 | 0.00000000 |
| 7942 | 4.29 | 0.00000000 |
| 7943 | 4.17 | 0.00000000 |
| 7944 | 3.50 | 0.00000000 |
| 7945 | 4.07 | 0.00000000 |
| 7946 | 3.50 | 0.00000000 |
| 7947 | 4.44 | 0.00000000 |
| 7948 | 3.24 | 0.00000000 |
| 7949 | 4.26 | 0.00000000 |
| 7950 | 11.86 | 0.00000000 |
| 7951 | 4.94 | 0.00000000 |
| 7952 | 4.00 | 0.00000000 |
| 7953 | 7.11 | 0.00000000 |
| 7954 | 8.93 | 0.00000000 |
| 7955 | 6.00 | 0.00000000 |
| 7956 | 7.29 | 0.00000000 |
| 7957 | 6.18 | 0.00000000 |
| 7958 | 10.44 | 0.00000000 |
| 7959 | 8.44 | 0.00000000 |
| 7960 | 8.38 | 0.00000000 |
| 7961 | 5.58 | 0.00000000 |
| 7962 | 5.59 | 0.00000000 |
| 7963 | 7.79 | 0.00000000 |
| 7964 | 8.24 | 0.00000000 |

| | | |
|---|---|---|
| 7965 | 5.45 | 0.00000000 |
| 7966 | 5.52 | 0.00000000 |
| 7967 | 6.86 | 0.00000000 |
| 7968 | 8.17 | 0.00000000 |
| 7969 | 7.82 | 0.00000000 |
| 7970 | 8.46 | 0.00000000 |
| 7971 | 9.54 | 0.00000000 |
| 7972 | 8.82 | 0.00000000 |
| 7973 | 9.04 | 0.00000000 |
| 7974 | 9.79 | 0.00000000 |
| 7975 | 8.11 | 0.00000000 |
| 7976 | 8.13 | 0.00000000 |
| 7977 | 10.70 | 0.00000000 |
| 7978 | 9.16 | 0.00000000 |
| 7979 | 9.64 | 0.00000000 |
| 7980 | 9.89 | 0.00000000 |
| 7981 | 10.45 | 0.00000000 |
| 7982 | 14.00 | 0.00000000 |
| 7983 | 13.42 | 0.00000000 |
| 7984 | 9.44 | 0.00000000 |
| 7985 | 14.21 | 0.00000000 |
| 7986 | 17.92 | 0.00000000 |
| 7987 | 16.44 | 0.00000000 |
| 7988 | 14.42 | 0.00000000 |
| 7989 | 16.14 | 0.00000000 |
| 7990 | 17.90 | 0.00000000 |
| 7991 | 20.56 | 0.00000000 |
| 7992 | 17.48 | 0.00000000 |
| 7993 | 27.26 | 0.00000000 |
| 7994 | 22.06 | 0.00000000 |
| 7995 | 25.53 | 0.00000000 |
| 7996 | 35.74 | 0.00000000 |
| 7997 | 38.00 | 0.00000000 |
| 7998 | 50.34 | 0.00000000 |
| 7999 | 74.28 | 0.00000000 |
| 8000 | 147.81 | 0.00000000 |
| 8001 | 205.98 | 0.00000000 |
| 8002 | 487.67 | 0.00000000 |
| 8003 | 593.98 | 0.00000000 |
| 8004 | 605.08 | 0.00000000 |
| 8005 | 502.83 | 0.00000000 |
| 8006 | 431.29 | 0.00000000 |
| 8007 | 374.91 | 0.00000000 |
| 8008 | 339.91 | 0.00000000 |

| | | |
|---|---|---|
| 8009 | 269.65 | 0.00000000 |
| 8010 | 219.39 | 0.00000000 |
| 8011 | 158.96 | 0.00000000 |
| 8012 | 117.19 | 0.00000000 |
| 8013 | 86.18 | 0.00000000 |
| 8014 | 32.00 | 0.00000000 |
| 8015 | 34.75 | 0.00000000 |
| 8016 | 22.50 | 0.00000000 |
| 8017 | 14.50 | 0.00000000 |
| 8018 | 31.50 | 0.00000000 |
